# An RGS5-positive fibroblast to WNT-like epithelial TGFB axis in colorectal liver metastasis, replicated in an independent paired cohort

**DOI:** 10.64898/2026.09.06.749652

**Authors:** Yingyu Yu, Feifei Ma, Yige Yu

## Abstract

**Background:** Colorectal cancer liver metastasis is accompanied by coordinated stromal remodelling, but which fibroblast and malignant epithelial states are coupled, and whether any such coupling holds outside the dataset it was found in, is rarely tested.

**Methods:** Paired primary tumours and liver metastases from six patients were analysed by single-cell RNA sequencing with copy-number-based identification of malignant cells, trajectory inference, cell-cell communication and ligand-target modelling, and spatial transcriptomics of three metastatic sections. Protein staining was taken from an independent tissue resource. Every step was then repeated in a second, independent paired cohort of five patients and two further metastatic sections, with cell identities fixed by transfer from the discovery data rather than refitted.

**Results:** An RGS5-positive fibroblast/pericyte-like state (F2) and a WNT-like malignant epithelial state (C2) were both enriched in metastases and occupied late trajectory positions. Communication analysis nominated a directional F2-to-C2 programme with transforming growth factor beta as the highest-ranked pathway and its receptor *TGFBR2* enriched in C2. Spatial data placed the two states in the same neighbourhoods. Hub genes of the programme were *TPT1*, *SLC1A5*, *SOX4* and *TSC22D1*, and protein staining was higher in tumour than normal colon for the three with available data. In the independent cohort the same states reappeared: F2 was enriched in metastases (Ro/E 1.22 versus 0.52; 32% versus 14% of fibroblasts), C2-like cells were the most metastasis-enriched epithelial state (1.10 versus 0.86), 92% of the cells that cohort’s own authors called MCAM-positive fibroblasts mapped to F2, and F2 was again the strongest TGFB sender to C2-like epithelium at both sites.

**Conclusions:** A directional TGFB axis from an RGS5-positive fibroblast state to a WNT-like epithelial state is a reproducible feature of colorectal liver metastasis rather than a property of one dataset. Its spatial expression varies between sections, which constrains how tightly the two states can be said to be co-localised.

## Background

Colorectal cancer (CRC) remains a leading cause of cancer death, and metastatic disease accounts for most CRC-related mortality, with the liver as the dominant site of spread ^1,2^. Although systemic therapy, molecular stratification and local treatment have improved management of colorectal liver metastasis, outcomes remain highly variable ^3,4^. This variability reflects more than tumour-intrinsic genomic alterations; adaptive malignant-cell states and tissue-specific microenvironmental programmes contribute to colonization, resistance and recurrence ^5,6^.

Bulk transcriptomic classifications, including the consensus molecular subtypes, capture major epithelial WNT/*MYC* and mesenchymal stromal axes but average across coexisting cell states and cannot resolve which malignant epithelial populations are enriched during metastasis ^7^. WNT-high and intestinal stem-like programmes have been linked to colorectal cancer stemness, plasticity and relapse ^8,9^. Cancer-associated fibroblasts remodel extracellular matrix, regulate paracrine signalling and shape immune contexture ^10,11^, while fibroblast diversity is organized across tissues and disease states ^12^. Single-cell and spatial transcriptomics can resolve these compartments and their shared architecture ^13–17^. Recent colorectal liver-metastasis studies have reported myofibroblastic fibroblast– tumour crosstalk, *SEMA3C*–*NRP2* signalling, desmoplastic zonation and other spatial stromal–tumour programmes ^18–22^.

Building on this work, we asked a defined question: whether an independently derived RGS5-positive, pericyte-like fibroblast state and a WNT-like malignant epithelial state are linked across matched primary–metastatic origin, inferred trajectory, ligand–target modelling and spatial position, and whether that link is reflected in the immune contexture of colorectal tumours. We addressed it by integrating paired single-cell data, copy-number-based malignant-cell identification, trajectory inference, intercellular communication, spatial deconvolution, protein-level staining from an independent tissue resource and two bulk-expression cohorts. Each layer is assessed on its own terms and associations are reported as associations; the analyses define a testable niche model rather than infer causality from observational transcriptomic data.

## Methods

### Data sources

Bulk RNA-seq profiles and matched clinical data were obtained from The Cancer Genome Atlas (TCGA; https://portal.gdc.cancer.gov/), comprising 449 colon and 164 rectal adenocarcinomas; 593 cases with recurrence-free survival data entered the survival analysis, and tumour plus normal samples entered diagnostic analyses. The external cohort GSE39582 (19 normal and 566 colorectal cancer samples; 519 with eligible recurrence-free survival data) was downloaded from the Gene Expression Omnibus (GEO; https://www.ncbi.nlm.nih.gov/geo/). Single-cell RNA-seq data were obtained from GSE178318 (six paired primary colorectal cancers and matched liver metastases). Spatial transcriptomic data were obtained from GSE217414 (four liver-metastasis sections, of which three met the pre-specified technical criteria for downstream niche analysis).

### Single-cell processing and malignant-cell identification

Single-cell data were analysed in Seurat v5.5.0 ^23^. Raw counts were re-used whenever extracted subpopulations were reprocessed. Counts were log-normalized, 2,000 highly variable genes were selected, data were scaled, and principal-component analysis used 50 dimensions; Harmony (sample as batch variable) corrected batch effects, followed by shared-nearest-neighbour graph construction and Louvain clustering. Major lineages were annotated with canonical markers. Epithelial cells were evaluated with CopyKAT using high-confidence T/NK and B/plasma cells as diploid references, and cells with broad aneuploid profiles were classified as malignant epithelial cells.

Malignant epithelial cells were re-normalized from raw counts, re-clustered (50 principal components, Harmony) and annotated into seven subclusters; fibroblasts were reprocessed identically. In each compartment, a coarse cluster showing lineage-inconsistent identity was removed before final reclustering using an explicit marker rule: its top markers were dominated by T-cell genes (*CD3D*, *CD3E*, *CD2*, *TRAC* and *IL7R*), with low epithelial (*EPCAM* and keratins) or fibroblast (*COL1A1*, *DCN* and *PDGFRB*) marker expression. DoubletFinder had been applied upstream, but doublet filtering does not remove ambient-RNA background or lineage cells co-embedded during coarse clustering. Discriminating markers across the retained subclusters are shown in Additional file 1: Figure S1. Differential expression used the Wilcoxon rank-sum test (Seurat FindMarkers), requiring adjusted P < 0.05, positive average log2 fold-change and detectable expression in a sufficient fraction of target-cluster cells. Full marker lists are provided in Additional file 3: Table S1.

### Enrichment, distribution and trajectory analysis

Tissue distribution across primary and metastatic groups was quantified by the ratio of observed to expected cell numbers (Ro/E), with the expected fraction estimated from the overall cross-group distribution; Ro/E > 1 denotes enrichment and Ro/E < 1 depletion. Gene Ontology biological-process (GO-BP) enrichment of cluster markers used clusterProfiler v4.18.4 with org.Hs.eg.db v3.22.0 (Benjamini–Hochberg-adjusted P < 0.05; full results in Additional file 3: Table S2). Differentiation potential was estimated with CytoTRACE2 v1.1.0, and Monocle3 v1.4.27 inferred pseudotime with the trajectory root chosen in the CytoTRACE2-defined least-differentiated region; subclusters were ordered along pseudotime and binned into early, middle and late stages.

### Intercellular communication and NicheNet

Ligand–receptor communication was inferred with CellChat v2.1.2 using the default human database ^24^ and truncated-mean expression, analysed separately in primary and metastatic samples and compared between conditions, with attention to F2 *RGS5*-positive fibroblast–C2 WNT-like epithelial signalling. NicheNet (nichenetr v2.2.1.1) then tested whether sender-derived ligands could explain receiver transcriptional programmes, treating F2 fibroblasts as senders and C2 epithelial cells as receivers in metastatic samples. Expressed genes were detected in at least 10% of the respective population; ligand activity was ranked by corrected area under the precision–recall curve against the C2 gene set, and the top 15 ligands and their regulatory-potential links to C2 targets were retained. C2 target-programme and TGFB-linked target-programme activity per cell were scored with Seurat AddModuleScore using the full NicheNet C2 target set and the *TGFB1*-specific target set, respectively.

### Spatial transcriptomics

Spatial data were processed in Seurat (quality control, normalization, principal-component analysis and UMAP). Cell-type deconvolution used RCTD (spacexr v1.2.0) in full mode with the refined single-cell object as reference, and continuous weights quantified spatial abundance of malignant epithelial and fibroblast subclusters. F2-high and C2-high spots were defined from z-scored top-50 marker scores; niche relationships were assessed by comparing continuous F2 and C2 scores and by computing, for each spot, the distance to the nearest F2-high spot, compared between C2-high and non-C2-high spots using the Wilcoxon test. Neighbourhood F2 ligand activity was aggregated from nearby spots by inverse-distance weighting and tested against C2 target activity within C2-high regions by Spearman correlation. Because Visium spots contain mixed cells and RCTD estimates continuous weights, these analyses test niche-level association rather than single-cell contact.

### Survival and diagnostic modelling

Hub genes were scored per sample by single-sample gene-set enrichment analysis (ssGSEA; GSVA v2.4.9) against the whole-transcriptome background. For RFS, patients were split at the median score; Kaplan–Meier curves were compared by the log-rank test, and hazard ratios were estimated with Cox proportional-hazards models both univariably and multivariably with adjustment for tumour stage. For diagnosis, single-gene performance was assessed by logistic-regression ROC analysis with Youden-derived cut-offs; TCGA samples were split 7:3 into training and locked-threshold validation sets, and external validation used GSE39582. Performance was summarized by area under the curve (AUC) with confidence intervals, sensitivity, specificity and accuracy.

### Protein-level expression

Immunohistochemical images for the hub genes were retrieved from the Human Protein Atlas (https://www.proteinatlas.org/), taking one normal colon and one colorectal cancer core per gene. Staining intensity, stained fraction and subcellular location were used as annotated in the resource; no re-scoring was performed. Antibody, donor and scoring detail for each image are listed in Additional file 2. For TPT1 and SLC1A5 the normal and tumour images derive from different antibodies, so these comparisons are qualitative; for SOX4 the same antibody was used in both tissues. TSC22D1 has no colorectal staining data in the resource and could not be assessed.

### Independent-cohort replication

Single-cell and spatial data from an independent paired cohort (GSE225857) were downloaded from the Gene Expression Omnibus. Non-immune cells with the authors’ own annotation were split into fibroblast and malignant epithelial compartments, normalized, and integrated over patients with Harmony. Fibroblast identities were assigned by reference-based label transfer from the discovery fibroblast object using FindTransferAnchors and TransferData over 30 dimensions, with no reclustering, so the labels are fixed by the discovery data rather than fitted to the validation data. Epithelial cells were scored against the seven discovery signatures with AddModuleScore and assigned to the best-matching state after column-wise scaling. Enrichment by site was quantified by the ratio of observed to expected cell numbers. Communication was inferred with CellChat as in the discovery analysis, down-sampled to at most 1,500 cells per group and run separately for each site. For the two liver-metastasis sections, spots were scored with the same signatures; spots in the top quartile were called F2-high or C2-high; the neighbourhood F2 score is the mean F2 score of spots within 1.5 spot pitches, and both the neighbourhood score and the distance to the nearest F2-high spot were tested against 2,000 random relabellings.

### Statistical analysis and reproducibility

Analyses used R v4.5.3 with SeuratObject v5.4.0, Harmony v2.0.4, copyKAT v1.1.0, CytoTRACE2 v1.1.0, Monocle3 v1.4.27, clusterProfiler v4.18.4, GSVA v2.4.9, CellChat v2.1.2, nichenetr v2.2.1.1, spacexr v1.2.0, survival v3.8.6, survminer v0.5.2, pROC v1.19.0.1 and ggplot2. Two-group comparisons used the Wilcoxon rank-sum test and correlations the Spearman coefficient unless stated otherwise; multiple testing was controlled by the Benjamini–Hochberg method, with adjusted P < 0.05 considered significant. All figures were generated from the analysis objects with a single shared plotting configuration; analysis code is available as described under Data Availability Statement.

## Results

### A single-cell landscape of paired primary and metastatic CRC

Analysis of GSE178318 resolved T/NK, B, plasma, myeloid, mast, fibroblast, endothelial and epithelial compartments, supported by canonical markers, with both primary and metastatic samples contributing to all lineages (Figure 1A–C). Metastatic lesions contained a higher proportion of T/NK cells and relatively fewer epithelial, B, plasma and fibroblast cells than primary tumours (Figure 1D). Because epithelial cells represent the malignant compartment and fibroblasts shape stromal organization, both lineages were taken forward for subclustering.

**Figure 1.**
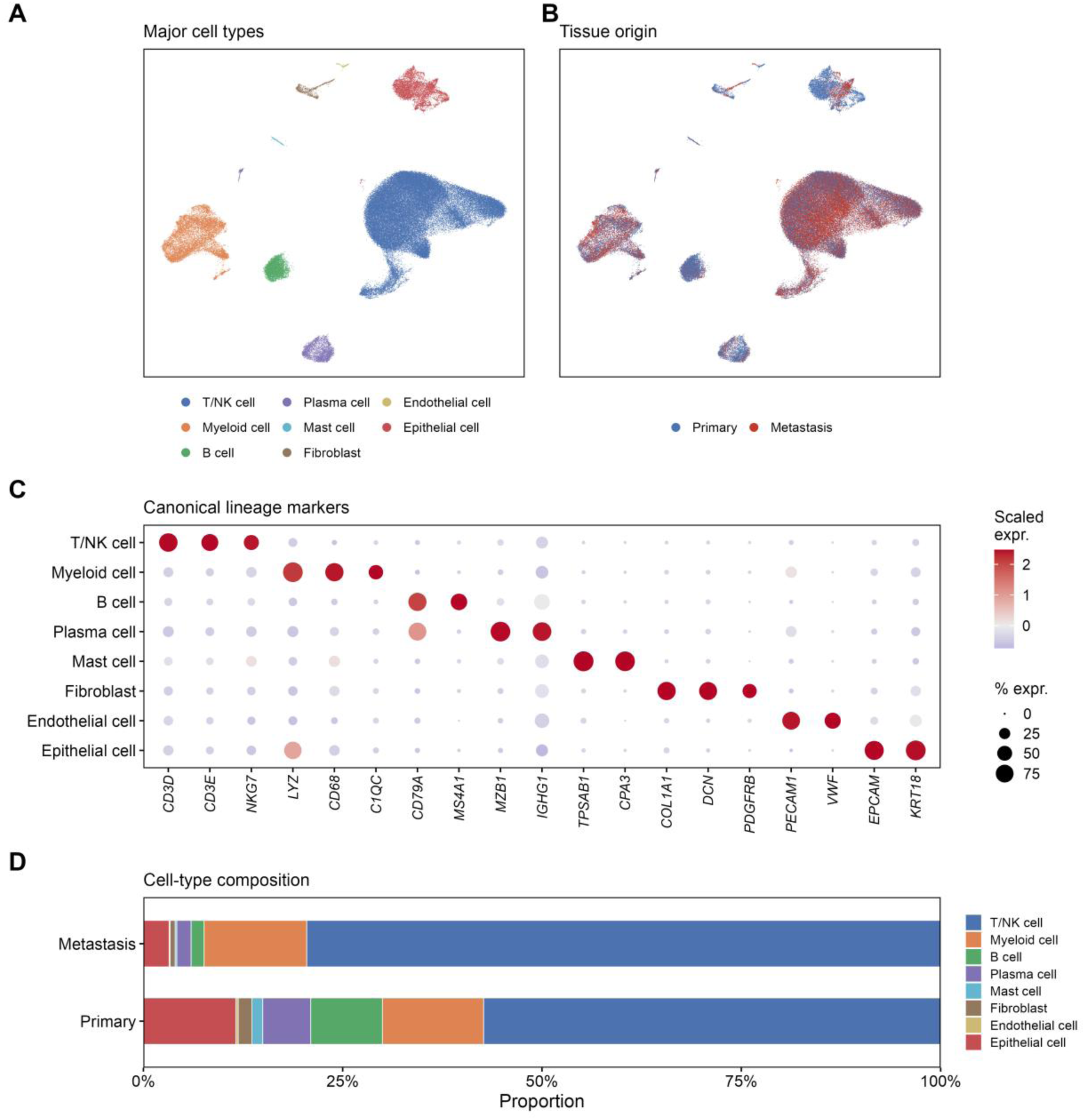
Single-cell landscape of GSE178318 CRC samples. (A) UMAP of major annotated cell types. (B) UMAP coloured by primary versus metastatic origin. (C) Dot plot of canonical lineage markers across cell types. (D) Cell-type composition in primary and metastatic samples.

### A metastasis-enriched C2 WNT malignant epithelial state occupies a late trajectory position

CopyKAT identified aneuploid malignant epithelial cells, which resolved into seven subclusters after removal of an immune-contaminated cluster: C0 Is, C1 Das, C2 WNT, C3 Mp, C4 T/r-h, C5 Am/CEA-like and C6 DNA-r p (Figure 2A–C, G). Composition and Ro/E analyses showed non-random enrichment across sites, with C2 WNT cells preferentially enriched in liver metastases (Figure 2D–F). Consistent with a WNT-like identity, C2 markers included WNT targets such as *NKD1* and *NOTUM*, and C2 was enriched for Wnt-signalling terms (Figure 2C, G). CytoTRACE2 and Monocle3 placed C2 WNT cells toward the late pseudotime stage, together with metastasis-derived cells, indicating that this metastasis-enriched state corresponds to a relatively advanced transcriptional position (Figure 2H–J).

**Figure 2.**
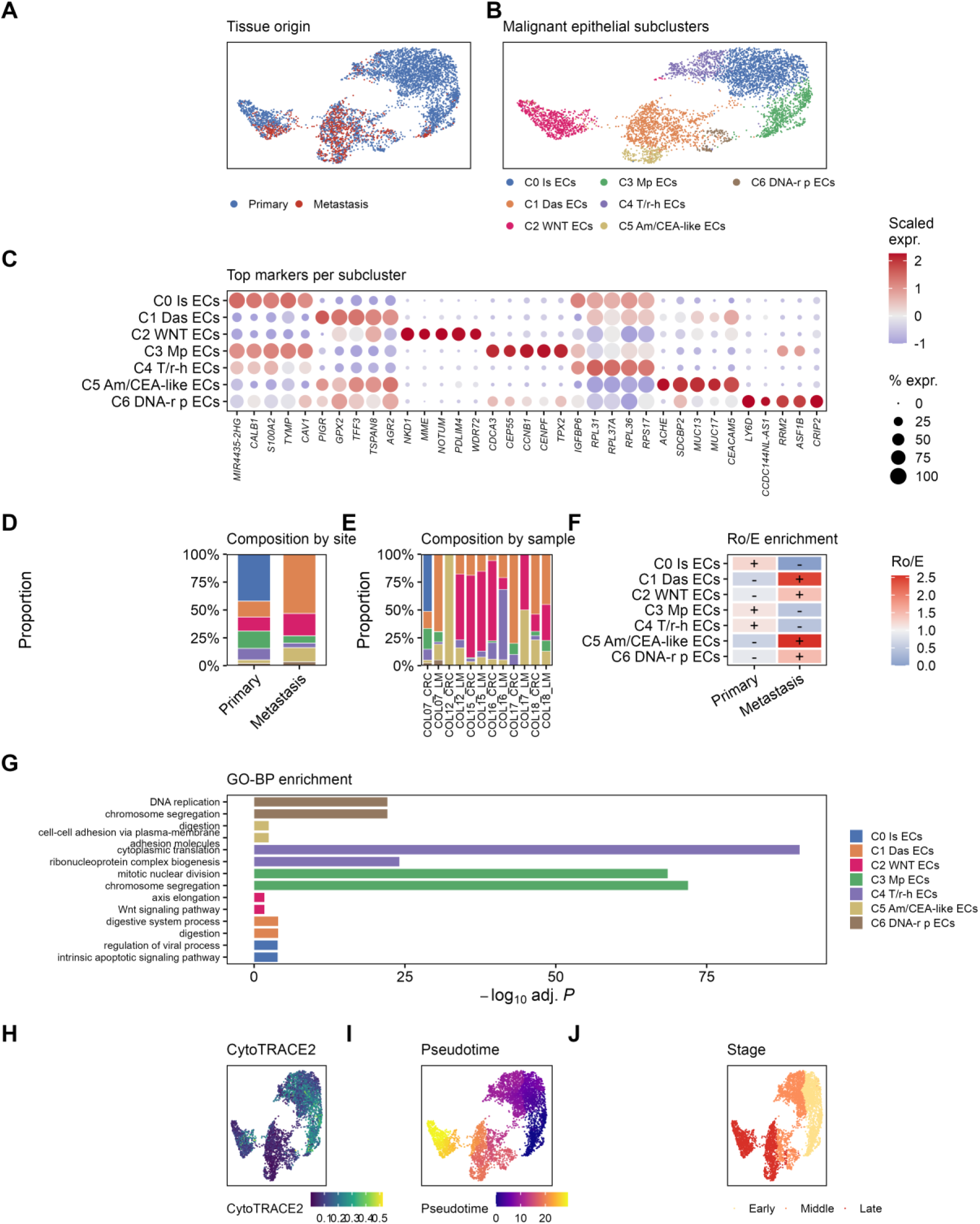
Malignant epithelial heterogeneity. (A) UMAP by origin. (B) UMAP of seven subclusters. (C) Marker dot plot. (D) Composition by site and (E) by sample. (F) Ro/E enrichment. (G) GO-BP enrichment. (H) CytoTRACE2 score. (I) Monocle3 pseudotime. (J) Pseudotime stage.

### Fibroblast reclustering identifies a metastasis-enriched F2 RGS5+ population

Fibroblasts reclustered after removal of a T-cell-contaminated cluster into five subtypes: F0 SFRP2+, F1 MYH11+, F2 *RGS5*+, F3 COL11A1+ mCAF and F4 ADAMDEC1+ (Figure 3A–C). F2 *RGS5*+ fibroblasts expressed pericyte/mural markers (*RGS5*, *NOTCH3*, *MCAM*) and were preferentially enriched in liver metastases by composition and Ro/E analysis (Figure 3D–F). GO-BP enrichment distinguished collagen/extracellular-matrix programmes in F0 and F3 from muscle-contraction, Notch-signalling and muscle-differentiation programmes in F2, supporting a contractile, pericyte-like identity (Figure 3G). CytoTRACE2 and Monocle3 placed F2 toward the late pseudotime stage, overlapping metastasis-derived fibroblasts (Figure 3H–J). Thus F2 *RGS5*+ fibroblasts represent both a metastasis-enriched and a relatively late stromal state.

**Figure 3.**
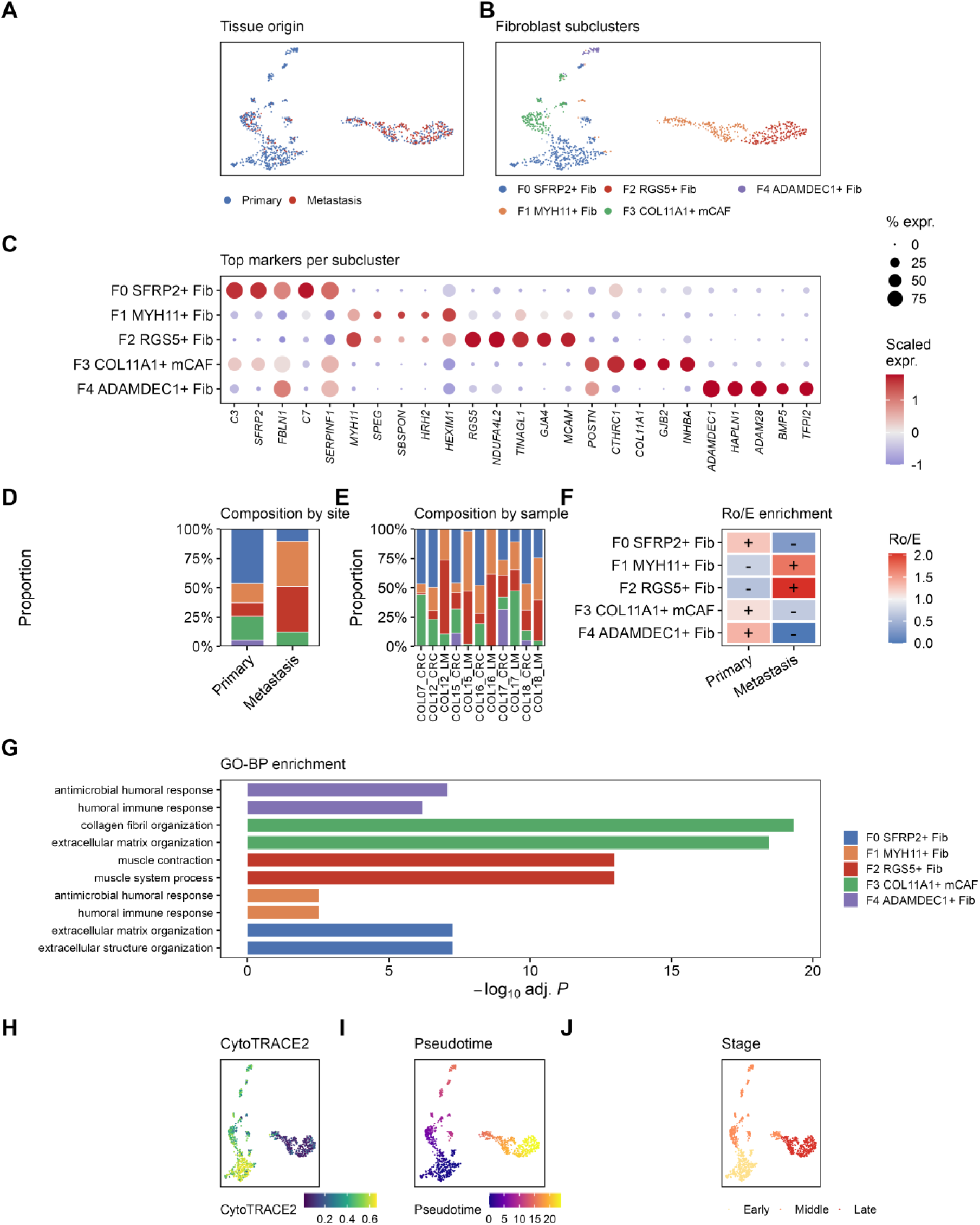
Fibroblast heterogeneity in colorectal liver metastasis. (A) UMAP by tissue origin. (B) UMAP of the five fibroblast subclusters (F0-F4). (C) Dot plot of top marker genes. (D) Subcluster composition by site and (E) by sample. (F) Ro/E enrichment across primary and metastatic lesions. (G) GO biological-process enrichment. (H) CytoTRACE2 score. (I) Monocle3 pseudotime. (J) Pseudotime stage.

### CellChat and NicheNet nominate an F2-to-C2 TGFB-linked programme

Given their parallel enrichment, communication between F2 and C2 was examined directly. CellChat identified multiple F2-to-C2 ligand–receptor pairs in primary and metastatic samples, dominated by extracellular-matrix and adhesion interactions (collagens with SDC1/SDC4 and integrins, laminins, fibronectin, *APP* and *THBS1*), several of which were retained or increased in metastasis (Figure 4A). NicheNet ranked *TGFB1* as the top predicted ligand explaining the C2 programme, followed by *ANG*, *TSPAN3*, *EFNA1*, *TIMP1*, *PROS1*, *SERPING1*, *CLDN3*, C1QB and *GDF11*, and linked these ligands to numerous C2 target genes (Figure 4B, C; Additional file 3: Table S5).

**Figure 4.**
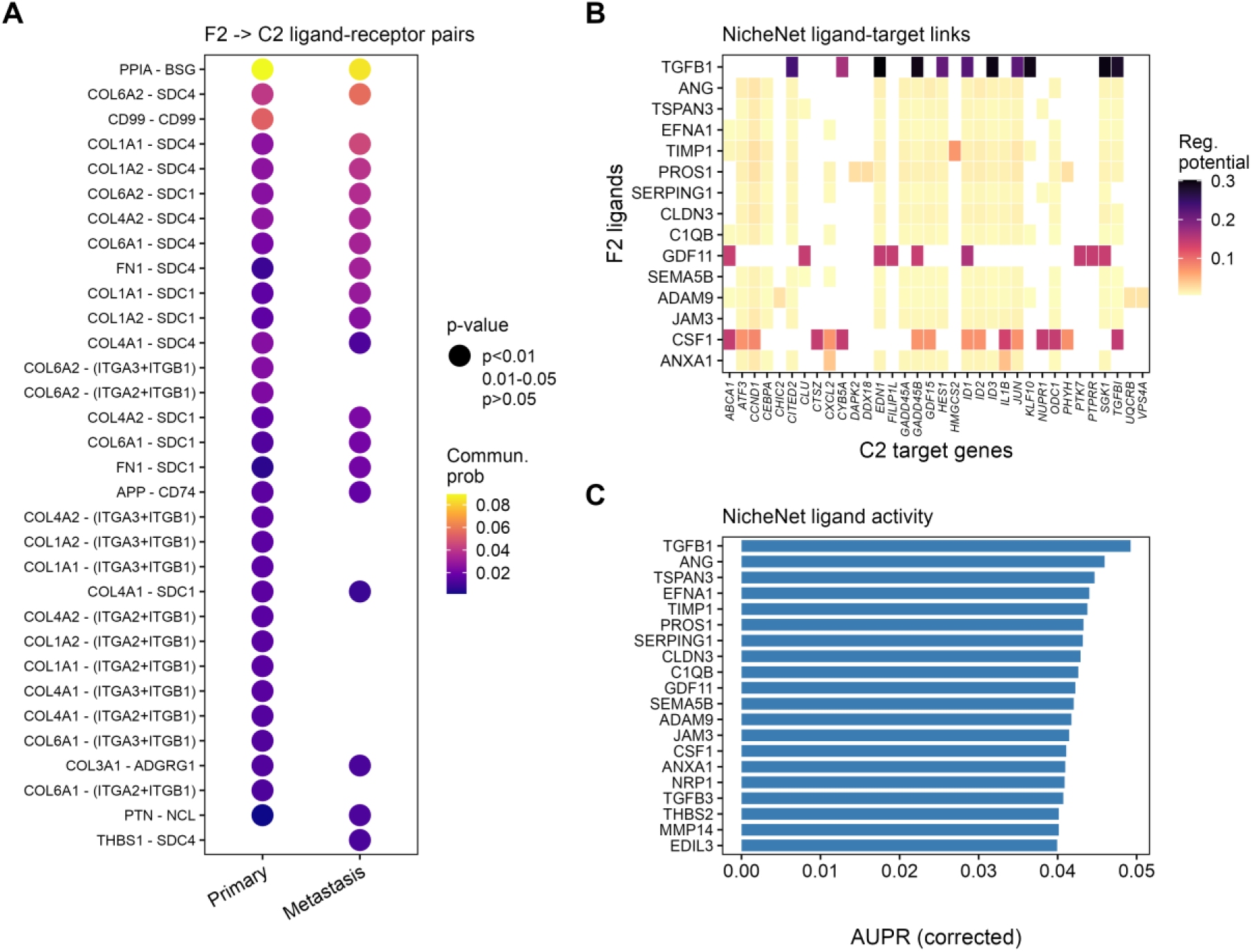
F2-to-C2 communication. (A) CellChat ligand–receptor bubble plot between F2 *RGS5*+ fibroblasts and C2 WNT epithelial cells in primary versus metastatic samples. (B) NicheNet ligand–target regulatory-potential heatmap. (C) NicheNet ligand-activity ranking (AUPR).

The TGFB axis was concordant across layers. Among fibroblasts, *TGFB1* was highest in F2 *RGS5*+ cells, whereas *TGFB3* was broadly distributed; among malignant epithelial cells, *TGFBR2* was enriched in C2 WNT cells, which also showed the highest C2 target-programme and TGFB-linked target-programme scores (Figure 5A–C). These data are consistent with F2 fibroblasts providing a TGFB-rich stromal environment aligned with the C2 epithelial programme, while remaining a hypothesis-generating rather than a causal inference.

**Figure 5.**
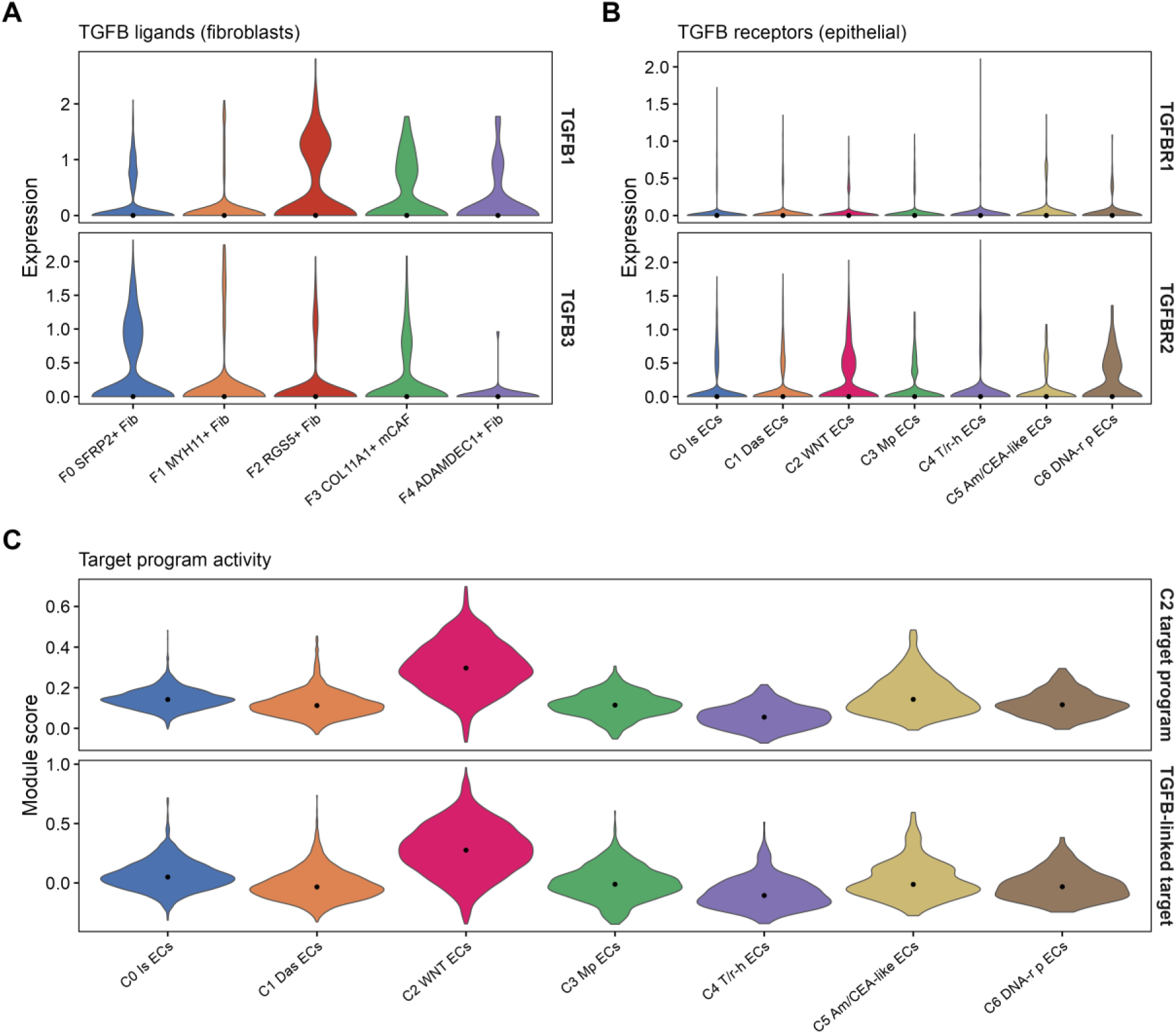
TGFB signalling axis. (A) *TGFB1* and *TGFB3* across fibroblast subclusters. (B) *TGFBR1* and *TGFBR2* across malignant epithelial subclusters. (C) C2 target-programme and TGFB-linked target-programme module scores across malignant epithelial subclusters.

### Spatial transcriptomics supports an F2–C2 niche across metastatic sections

Spatial data were available for four liver-metastasis sections from GSE217414. Sample 2 was excluded before downstream niche analysis because it had fewer tissue-covered spots and lower deconvolution confidence for the epithelial and fibroblast subclusters of interest; Samples 1, 3 and 4 were processed identically and reported in full. In the retained sections, deconvolution and z-scored F2 and C2 top-50 marker scores showed overlapping or adjacent high-score regions. Continuous F2 and C2 scores were positively but modestly correlated (Spearman rho 0.20–0.30; Figure 6A–F). C2-high spots were closer to F2-high regions than C2-low spots in each section (Wilcoxon P from 1.5×10⁻⁷ to 3.8×10⁻¹⁶; Figure 6G–I). Within C2-high regions, neighbourhood F2 ligand activity correlated positively with C2 target-programme activity (Spearman rho 0.13, 0.43 and 0.16 for Samples 1, 3 and 4, respectively; Figure 6J–L). These spot-level analyses support spatial association but do not establish cell–cell contact or ligand causality.

**Figure 6.**
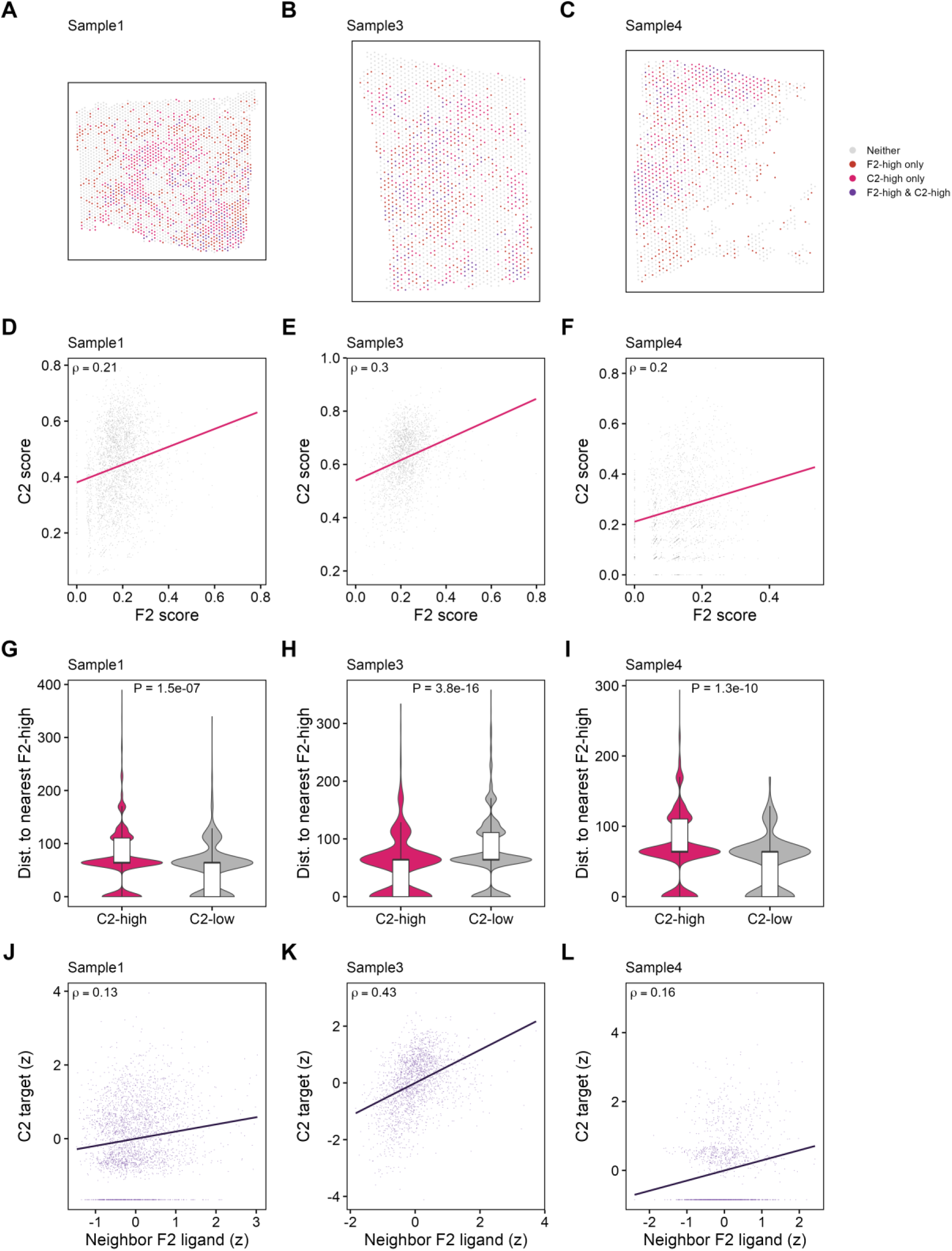
Spatial validation across three liver-metastasis sections (Sample 1, 3, 4). (A–C) Spatial maps of F2-high, C2-high and co-high regions. (D–F) Correlation of F2 and C2 signature scores. (G–I) Distance from C2-high versus C2-low spots to the nearest F2-high spot. (J–L) Neighbourhood F2 ligand activity versus C2 target activity within C2-high regions.

### A four-gene programme links the niche to recurrence and diagnosis

Intersecting NicheNet-predicted C2 targets, C2 markers and late-pseudotime up-regulated genes identified four overlapping hub genes, TPT1, SLC1A5, SOX4 and TSC22D1 (Figure 7A). In TCGA these genes were generally higher in tumours than normal tissue across stages, with SOX4 and SLC1A5 showing the strongest tumour-associated patterns (Figure 7B). A four-gene ssGSEA score stratified RFS, with high scores associated with worse RFS in TCGA (log-rank P = 0.049) and GSE39582 (log-rank P = 0.033; Figure 7C, D). In multivariable Cox models adjusting for tumour stage, the high-score group remained independently associated with RFS in GSE39582 (HR 1.54, P = 0.012) but was attenuated and non-significant in TCGA (HR 1.31, P = 0.17), indicating that the score’s prognostic value is partly linked to stage (Additional file 3: Table S3). For tumour-versus-normal discrimination, SOX4 and SLC1A5 achieved high single-gene AUCs (TCGA validation 0.966 and 0.924; GSE39582 0.971 and 0.926), whereas TPT1 and TSC22D1 were weaker (Figure 7E, F; Additional file 3: Table S4). At protein level, annotated staining in an independent tissue resource was higher in colorectal cancer than in normal colon for TPT1, SLC1A5 and SOX4, consistent with the transcriptional pattern; TSC22D1 has no colorectal staining data in that resource and could not be assessed (Figure 7C; antibody and donor details in Additional file 2). Together these analyses connect the F2–C2 niche programme to recurrence risk and tumour detection (Additional file 4).

**Figure 7.**
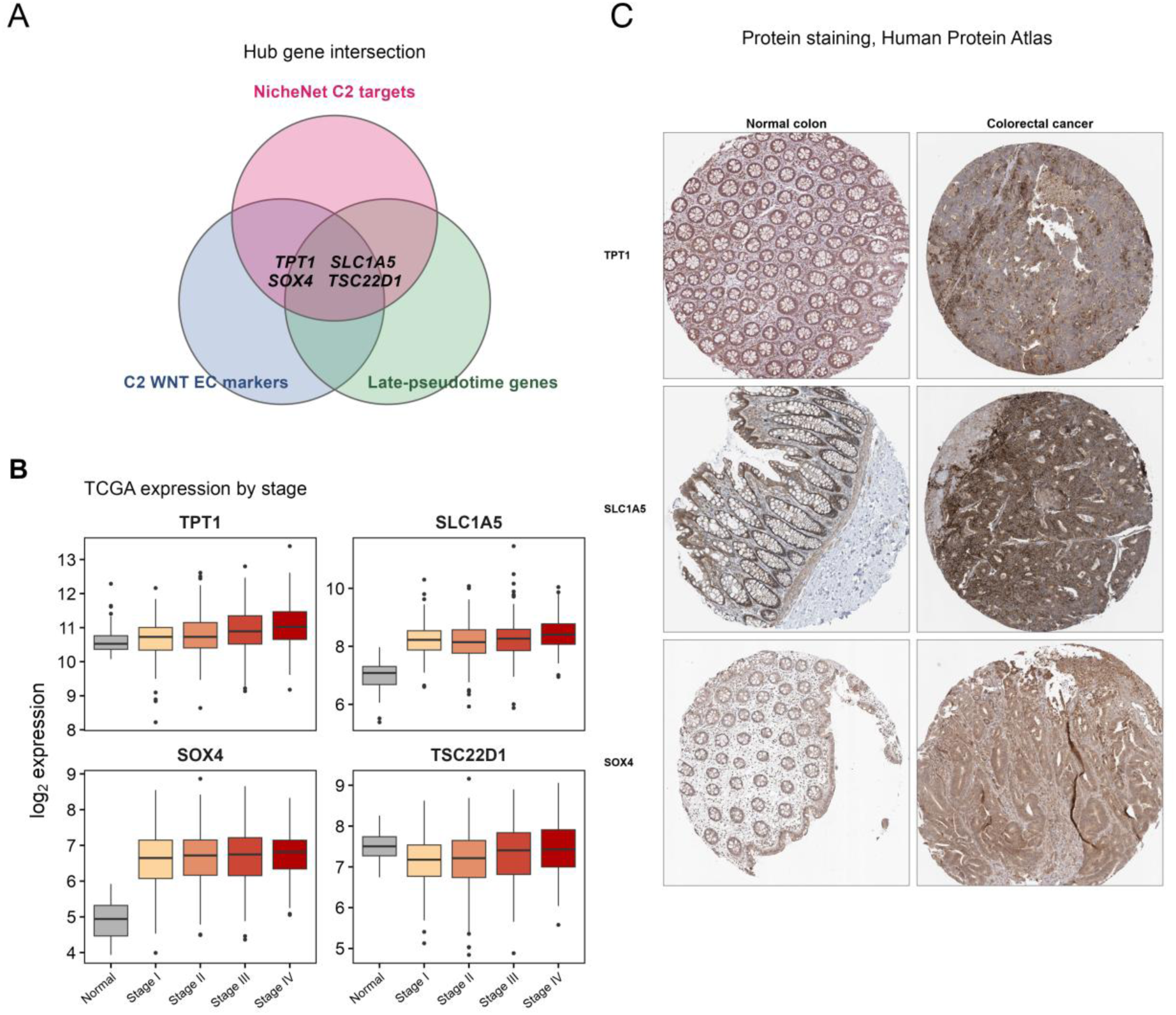
Hub genes of the F2-to-C2 programme. (A) Intersection of NicheNet-predicted C2 targets, C2 marker genes and late-pseudotime genes, yielding TPT1, SLC1A5, SOX4 and TSC22D1. (B) Expression of the four genes across tumour stages in The Cancer Genome Atlas. (C) Immunohistochemical staining in normal colon and colorectal cancer from the Human Protein Atlas; antibodies and staining categories are given in Additional file 2.

### The F2 and C2 states are recovered in an independent paired cohort

To test whether the two states are specific to the discovery data, we analysed an independent cohort of five patients with paired primary colorectal tumour and liver metastasis profiled by a different group (15,093 fibroblasts and 23,954 malignant epithelial cells). Fibroblast identities were assigned by reference-based label transfer from the discovery object, without reclustering. The F2 state was enriched in metastases (Ro/E 1.22 versus 0.52; 31.9% of fibroblasts in metastases versus 13.5% in primaries), and the increase held in 5 of five patients individually (Figure 8A-C). Transferred F2 cells expressed the markers that define the state in the discovery data, including *RGS5*, *NOTCH3*, *MCAM*, *PDGFRB* and *NDUFA4L2* (Figure 8D). The cohort’s own annotation supports the mapping: 92% of the cells the original authors called MCAM-positive fibroblasts were assigned to F2, an independently derived label (Figure 8E).

**Figure 8.**
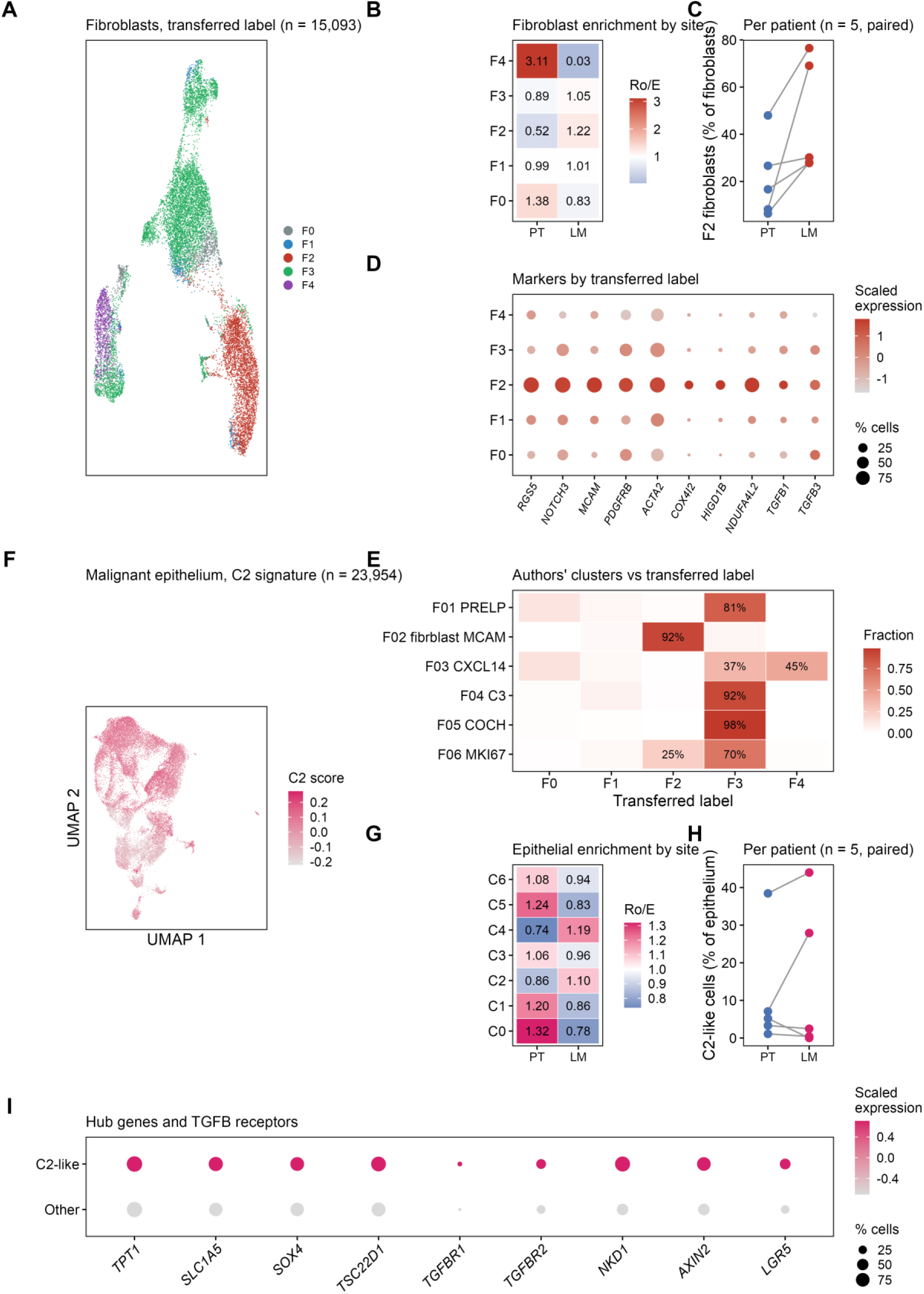
Replication of the F2 and C2 states in an independent paired cohort (GSE225857; five patients, paired primary tumour and liver metastasis). (A) Fibroblasts coloured by label transferred from the discovery object. (B) Ratio of observed to expected fibroblast numbers by site. (C) F2 fraction per patient, primary tumour (PT) versus liver metastasis (LM). (D) Marker expression by transferred label. (E) Concordance between the original authors’ fibroblast clusters and the transferred labels, row-normalised. (F) Malignant epithelium coloured by C2 signature score. (G) Ratio of observed to expected epithelial states by site. (H) C2-like fraction per patient. (I) Hub genes and TGFB receptors in C2-like versus remaining epithelium.

In the malignant epithelium, cells were scored against the seven discovery signatures and assigned to the best-matching state. C2-like cells were the most metastasis-enriched epithelial state (Ro/E 1.10 versus 0.86; 32.8% versus 25.8%; Figure 8F-H). Relative to the remaining epithelium, C2- like cells expressed higher *NKD1* (log2 fold change 1.70), *AXIN2* (1.10) and *LGR5* (0.95), confirming the WNT character of the state, together with the TGFB receptors *TGFBR2* (0.56) and *TGFBR1* (0.46) and three of the four hub genes, *TSC22D1* (0.66), *SLC1A5* (0.73) and *TPT1* (0.35); all adjusted P < 0.001. *SOX4* was higher but only marginally (0.10; Figure 8I).

### The F2-to-C2 TGFB axis and its tissue arrangement replicate

Communication analysis was repeated in the validation cohort with the transferred fibroblast labels. Among the five fibroblast states, F2 was the strongest TGFB sender to C2-like epithelium at both sites, in primary tumours and in metastases alike (Figure 9H), reproducing the direction nominated in the discovery data. The receptor complex was the same in both cohorts, with *TGFB1*, *TGFB2* and *TGFB3* signalling to ACVR1B and TGFBR2 (Figure 9I).

**Figure 9.**
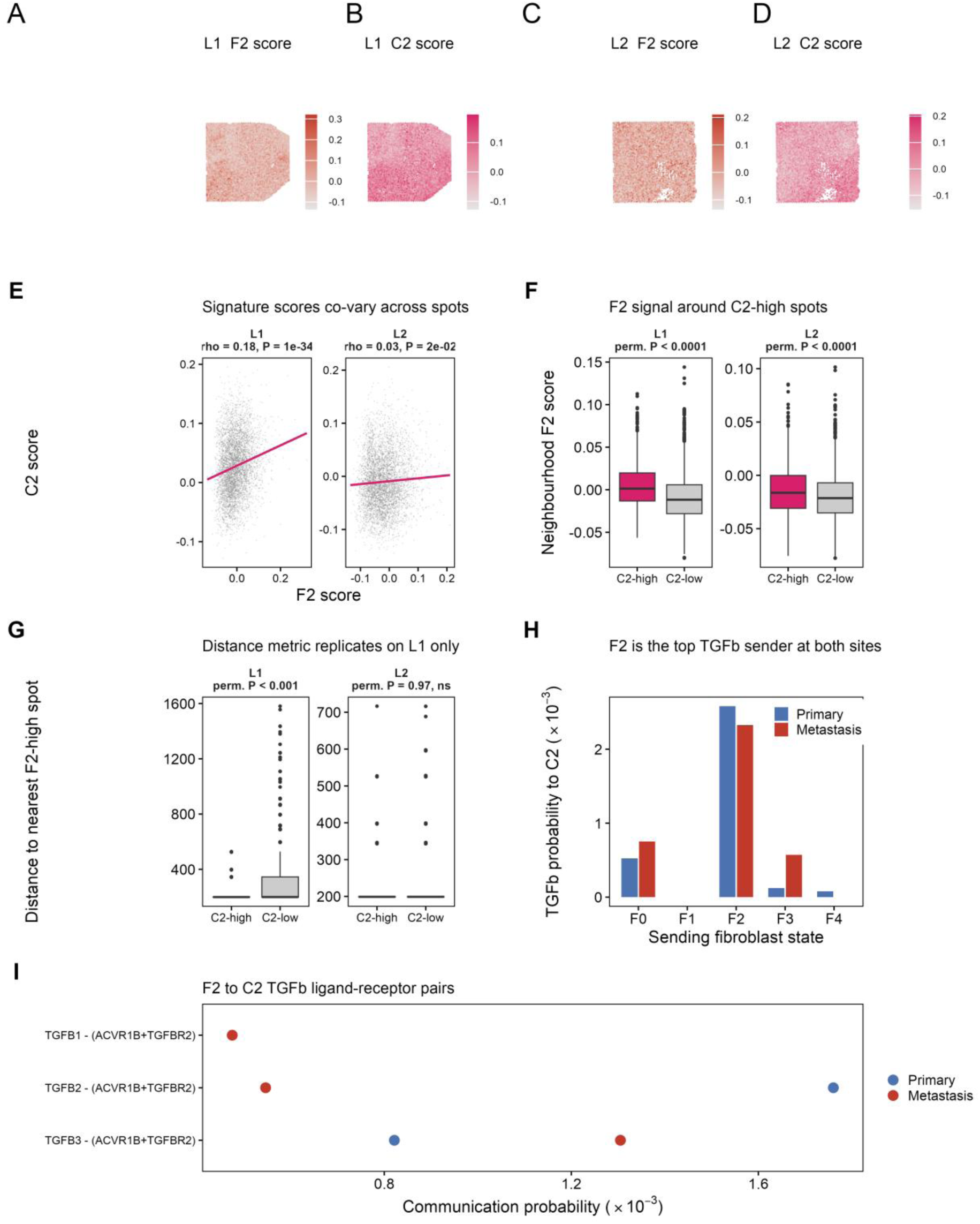
Tissue and communication layers of the validation cohort. (A-D) F2 and C2 signature scores across the two liver-metastasis sections L1 and L2. (E) Correlation of the two scores per section, with Spearman statistics. (F) Mean F2 score of the immediate neighbourhood around C2-high versus other spots; P values from 2,000 label permutations. (G) Distance to the nearest F2-high spot, which separates the groups on L1 but not on L2. (H) CellChat TGFB communication probability from each fibroblast state to C2-like epithelium, by site. (I) TGFB ligand-receptor pairs from F2 to C2-like epithelium.

Two liver-metastasis sections from the same cohort were scored spot by spot. F2 and C2 scores were positively correlated on both sections (Spearman rho 0.18, P = 1e-34 on L1; rho 0.03, P = 0.02 on L2; Figure 9A-E), and the mean F2 score of the immediate neighbourhood was higher around C2-high spots than around the rest on both sections (permutation P < 0.0001 for each; Figure 9F). The distance metric used in the discovery data behaved differently: C2-high spots lay closer to F2-high spots on L1 (permutation P < 0.001) but not on L2 (permutation P = 0.97), where F2-high spots were dense enough that almost every spot had one adjacent to it and the measure saturated (Figure 9G). The neighbourhood score, which does not saturate, is therefore the more informative of the two in this cohort. Taken together, the state definitions, their metastatic enrichment and the directional TGFB signal reproduce in an independent cohort, while the strength of the spatial arrangement varies between sections.

## Discussion

Integrating single-cell, trajectory, communication, spatial and bulk analyses, we define a metastasis-associated epithelial–stromal niche in colorectal liver metastasis. The central observation is the parallel enrichment of C2 WNT malignant epithelial cells and F2 *RGS5*+ fibroblast/pericyte-like cells in metastatic samples, both occupying relatively advanced transcriptional positions. This suggests that liver metastasis is accompanied not only by expansion of malignant epithelial states but also by coordinated stromal remodelling.

The enrichment of C2 WNT-like cells is consistent with the established role of WNT signalling in colorectal cancer stemness and progression ^25,26^. Recent studies have also connected colorectal liver-metastasis epithelial states to myofibroblastic fibroblasts, *SEMA3C*–*NRP2* signalling, desmoplastic architecture and *HGF*–*MET*–*MYC* interactions ^19–22^. The present analysis does not displace those axes. Its narrower contribution is the reproducible alignment of an *RGS5*-positive, mural-like fibroblast state with a WNT-like epithelial state across paired tissue origin, trajectory, communication and spatial layers. *RGS5* and *NOTCH3* expression supports a vascular-associated mural identity ^27^, so the term fibroblast/pericyte-like is retained rather than asserting a pure fibroblast lineage. CellChat and NicheNet provided complementary candidate interactions ^28,29^; their convergence on a TGFB-linked programme is coherent with stromal TGFB in colorectal metastasis ^30^, but it does not demonstrate that TGFB signalling is required for the C2 state.

Spatial transcriptomics adds an independent layer: F2-high and C2-high regions were coupled, C2-high spots lay closer to F2-high neighbourhoods, and neighbourhood F2 ligand activity tracked C2 target activity. These patterns reduce the likelihood that the relationship is an artefact of dissociation, but Visium spots are mixtures and RCTD yields continuous weights ^31^, so the evidence supports niche-level association rather than single-cell contact. The four-gene programme connected the niche to clinically relevant outcomes; importantly, multivariable analysis showed that its prognostic association is stage-independent in GSE39582 but not in TCGA, so the score is best regarded as a niche-linked correlate of recurrence risk that partly reflects tumour progression, not yet as a stage-independent biomarker. Previous colorectal cancer studies have independently linked SOX4 expression to tumour progression and SLC1A5 to tumour-cell growth ^32,33^, providing biological context for two components of the score. Protein staining in an independent tissue resource was higher in tumour than in normal colon for the three genes with available data, which supports the direction of the transcriptional finding; because two of the three comparisons use different antibodies in the two tissues, and because staining is annotated rather than quantified, this is corroborative rather than quantitative validation.

Several limitations constrain interpretation. All central conclusions rest on computational inference: CytoTRACE2 and Monocle3 estimate relative states rather than lineage history ^34,35^, and communication and spatial correlations nominate candidates rather than demonstrate causality. The F2 population has mixed fibroblast/pericyte features that require orthogonal histological markers. Spatial validation rests on three retained sections; one section was excluded for insufficient spot-level quality, and the small number of sections limits the precision of the spatial estimates. Protein-level support is limited to annotated single-core images from a public resource and does not substitute for quantified staining in a patient series. Functional testing should include fibroblast–epithelial co-culture, organoid models, ligand or receptor blockade and multiplex protein-level spatial validation. Within these limits, the cross-layer consistency defines a testable RGS5-positive fibroblast/pericyte-like–WNT epithelial programme associated with colorectal liver metastasis and recurrence. In the validation cohort the distance-based spatial measure separated C2-high from other spots on one of the two sections only; the neighbourhood score, which does not saturate when F2-high spots are dense, separated them on both. Spatial coupling should therefore be read as a neighbourhood-level association whose strength varies between sections.

## Conclusions

We delineate a stromal-epithelial niche in colorectal liver metastasis formed by RGS5-positive fibroblast/pericyte-like cells and WNT-like malignant epithelial cells, which are co-enriched in metastases and occupy late transcriptional states. Communication analyses nominate a directional TGFB axis from the fibroblast state to the epithelial state, and spatial data place the two states in shared neighbourhoods. The same states, the same metastatic enrichment and the same directional signal are recovered in an independent paired cohort whose cell identities were fixed by transfer rather than refitted, so the programme is a reproducible feature of the disease rather than a property of one dataset. The strength of the spatial arrangement varies between sections, and the axis remains a candidate for functional testing rather than an established mechanism.

## Declarations

### Ethics approval and consent to participate

Not applicable. This study analysed publicly available, de-identified datasets and collected no new human or animal samples, so no ethical approval or informed consent was required.

### Consent for publication

Not applicable.

### Availability of data and materials

The single-cell (GSE178318), spatial (GSE217414) validation (GSE39582) and independent replication (GSE225857) datasets are available from the Gene Expression Omnibus; colorectal cancer data from The Cancer Genome Atlas are available through the Genomic Data Commons. Processed analysis objects and figure-generation code are available from the corresponding author on reasonable request. Additional files 1 to 4 contain the supporting figures (Figures S1 to S3) and the supporting tables (Tables S1 to S6).

### Authors’ contributions

Y.Y. conceived the study, designed the analyses and drafted the manuscript. F.M. implemented the analysis pipelines, performed the statistical analysis and contributed to validation and investigation. Y.G.Y. curated the data, produced the figures, interpreted the results and revised the manuscript. Y.Y. and F.M. contributed equally. All authors read and approved the final manuscript.

## Acknowledgements

Not applicable.

## Abbreviations

AUC: area under the curve
AUPR: area under the precision–recall curve
CI: confidence interval
COAD: colon adenocarcinoma
CRC: colorectal cancer
GEO: Gene Expression Omnibus
GO-BP: Gene Ontology biological process
HR: hazard ratio
PCA: principal component analysis
RCTD: robust cell-type decomposition
READ: rectal adenocarcinoma
RFS: recurrence-free survival
ROC: receiver operating characteristic
Ro/E: ratio of observed to expected cell numbers
scRNA-seq: single-cell RNA sequencing
ssGSEA: single-sample gene-set enrichment analysis
TCGA: The Cancer Genome Atlas
UMAP: uniform manifold approximation and projection.

## Additional files

Additional file 1: Figure S1 Quality control. Dot plots of discriminating markers (T-cell *CD3D*, *CD3E*, *CD2*, *TRAC*, *IL7R*; epithelial *EPCAM*, *KRT8*, *KRT18*; fibroblast *COL1A1*, *DCN*, *PDGFRB*) across the retained malignant epithelial (A) and fibroblast (B) subclusters, supporting lineage identity and low T-cell-marker expression after removal of the contaminated clusters.

Additional file 2: Table S7 Immunohistochemistry metadata for the images shown in Figure 7C. Antibody, donor identifier, sex and age, annotated cell type, staining category, intensity, stained fraction and subcellular location for each of the six Human Protein Atlas images. For TPT1 and SLC1A5 the normal and tumour images derive from different antibodies, so those comparisons are qualitative; SOX4 uses the same antibody in both tissues.

Additional file 3: Tables S1–S6 Supplementary tables. Table S1, subcluster marker genes; Table S2, GO-BP enrichment; Table S3, univariable and stage-adjusted Cox models for recurrence-free survival; Table S4, single-gene diagnostic performance; Table S5, NicheNet ligand-activity ranking and ligand–target links; Table S6, gene sets used for the immune and therapeutic analyses.

Additional file 4: Figure S3 Recurrence-free survival by four-gene score. Kaplan-Meier curves for patients above and below the median four-gene single-sample enrichment score in (A) The Cancer Genome Atlas (n = 593) and (B) GSE39582 (n = 519), with log-rank P values.

## References

1. Bray F, Laversanne M, Sung H, et al. Global cancer statistics 2022: GLOBOCAN estimates of incidence and mortality worldwide for 36 cancers in 185 countries. CA Cancer J Clin. 2024;74(3):229–263.

2. Dekker E, Tanis PJ, Vleugels JLA, Kasi PM, Wallace MB. Colorectal cancer. Lancet. 2019;394(10207):1467–1480.

3. Van Cutsem E, Cervantes A, Adam R, et al. ESMO consensus guidelines for the management of patients with metastatic colorectal cancer. Ann Oncol. 2016;27(8):1386–1422.

4. Biller LH, Schrag D. Diagnosis and treatment of metastatic colorectal cancer: a review. JAMA. 2021;325(7):669–685.

5. Massagué J, Obenauf AC. Metastatic colonization by circulating tumour cells. Nature. 2016;529(7586):298–306.

6. Lambert AW, Pattabiraman DR, Weinberg RA. Emerging biological principles of metastasis. Cell. 2017;168(4):670–691.

7. Guinney J, Dienstmann R, Wang X, et al. The consensus molecular subtypes of colorectal cancer. Nat Med. 2015;21(11):1350–1356.

8. Merlos-Suárez A, Barriga FM, Jung P, et al. The intestinal stem cell signature identifies colorectal cancer stem cells and predicts disease relapse. Cell Stem Cell. 2011;8(5):511–524.

9. Vermeulen L, De Sousa E Melo F, van der Heijden M, et al. Wnt activity defines colon cancer stem cells and is regulated by the microenvironment. Nat Cell Biol. 2010;12(5):468–476.

10. Kalluri R. The biology and function of fibroblasts in cancer. Nat Rev Cancer. 2016;16(9):582–598.

11. Sahai E, Astsaturov I, Cukierman E, et al. A framework for advancing our understanding of cancer-associated fibroblasts. Nat Rev Cancer. 2020;20(3):174–186.

12. Buechler MB, Pradhan RN, Krishnamurty AT, et al. Cross-tissue organization of the fibroblast lineage. Nature. 2021;593(7860):575–579.

13. Joanito I, Wirapati P, Zhao N, et al. Single-cell and bulk transcriptome sequencing identifies two epithelial tumour cell states and refines the consensus molecular classification of colorectal cancer. Nat Genet. 2022;54(7):963–975.

14. Li H, Courtois ET, Sengupta D, et al. Reference component analysis of single-cell transcriptomes elucidates cellular heterogeneity in human colorectal tumours. Nat Genet. 2017;49(5):708–718.

15. Ståhl PL, Salmén F, Vickovic S, et al. Visualization and analysis of gene expression in tissue sections by spatial transcriptomics. Science. 2016;353(6294):78–82.

16. Pelka K, Hofree M, Chen JH, et al. Spatially organized multicellular immune hubs in human colorectal cancer. Cell. 2021;184(18):4734–4752.e20.

17. Xiao J, Yu X, Meng F, et al. Integrating spatial and single-cell transcriptomics reveals tumour heterogeneity and intercellular networks in colorectal cancer. Cell Death Dis. 2024;15(5):326.

18. Wang H, Semba T, Yonemura A, et al. Organ-specific cancer-associated fibroblast subtypes across the digestive system linked to cancer hallmarks. Cancer Sci. Published online June 1, 2026. doi:10.1111/cas.70435.

19. Zhan Y, Sun D, Gao J, et al. Single-cell transcriptomics reveals intratumor heterogeneity and the potential roles of cancer stem cells and myCAFs in colorectal cancer liver metastasis and recurrence. Cancer Lett. 2025;612:217452. doi:10.1016/j.canlet.2025.217452.

20. Zhang Y, Zuo A, Ba Y, et al. Cancer-associated fibroblast-derived *SEMA3C* facilitates colorectal cancer liver metastasis via *NRP2*-mediated MAPK activation. Proc Natl Acad Sci U S A. 2025;122(21):e2423077122. doi:10.1073/pnas.2423077122.

21. Andersson A, Escriva Conde M, Surova O, et al. Spatial transcriptome mapping of the desmoplastic growth pattern of colorectal liver metastases by in situ sequencing reveals a biologically relevant zonation of the desmoplastic rim. Clin Cancer Res. 2024;30(19):4517–4529. doi:10.1158/1078-0432.CCR-23-3461.

22. Chen J, Wang Z, Zhu B, Guan G. A spatially resolved single-cell landscape of colorectal cancer liver metastasis reveals a stromal–tumor glycolytic signaling interaction. Front Cell Dev Biol. 2025;13:1687485. doi:10.3389/fcell.2025.1687485.

23. Stuart T, Butler A, Hoffman P, et al. Comprehensive integration of single-cell data. Cell. 2019;177(7):1888–1902.e21.

24. Ramilowski JA, Goldberg T, Harshbarger J, et al. A draft network of ligand–receptor-mediated multicellular signalling in human. Nat Commun. 2015;6:7866.

25. Zhan T, Rindtorff N, Boutros M. Wnt signaling in cancer. Oncogene. 2017;36(11):1461–1473.

26. Wang F, Long J, Li L, et al. Single-cell and spatial transcriptome analysis reveals the cellular heterogeneity of liver metastatic colorectal cancer. Sci Adv. 2023;9(24):eadf5464.

27. Mitchell TS, Bradley J, Robinson GS, Shima DT, Ng YS. *RGS5* expression is a quantitative measure of pericyte coverage of blood vessels. Angiogenesis. 2008;11(2):141–151.

28. Jin S, Guerrero-Juarez CF, Zhang L, et al. Inference and analysis of cell-cell communication using CellChat. Nat Commun. 2021;12(1):1088.

29. Browaeys R, Saelens W, Saeys Y. NicheNet: modeling intercellular communication by linking ligands to target genes. Nat Methods. 2020;17(2):159–162.

30. Calon A, Espinet E, Palomo-Ponce S, et al. Dependency of colorectal cancer on a TGF-β-driven programme in stromal cells for metastasis initiation. Cancer Cell. 2012;22(5):571–584.

31. Cable DM, Murray E, Zou LS, et al. Robust decomposition of cell type mixtures in spatial transcriptomics. Nat Biotechnol. 2022;40(4):517–526.

32. Wang B, Li Y, Tan F, Xiao Z. Increased expression of *SOX4* is associated with colorectal cancer progression. Tumour Biol. 2016;37(7):9131–9137.

33. Huang F, Zhao Y, Zhao J, et al. Upregulated *SLC1A5* promotes cell growth and survival in colorectal cancer. Int J Clin Exp Pathol. 2014;7(9):6006–6014.

34. Cao J, Spielmann M, Qiu X, et al. The single-cell transcriptional landscape of mammalian organogenesis. Nature. 2019;566(7745):496–502.

35. Kang M, Gulati GS, Brown EL, et al. Improved reconstruction of single-cell developmental potential with CytoTRACE 2. Nat Methods. 2025;22(11):2258–2263.

