## Additional file for "An RGS5-positive fibroblast to WNT-like epithelial TGFB axis in colorectal liver metastasis, replicated in an independent paired cohort": Additional_file_2.docx

**Additional file 2: Table S7 Immunohistochemistry metadata for the images shown in Figure 7C**

Staining categories are those annotated in the Human Protein Atlas (https://www.proteinatlas.org/); no re-scoring was performed. For TPT1 and SLC1A5 the normal and tumour images derive from different antibodies, so those comparisons are qualitative; SOX4 uses the same antibody in both tissues. TSC22D1 has no colorectal staining data in the resource.

| **Gene** | **Tissue** | **Antibody** | **Donor ID** | **Sex, age** | **Annotated cell type** | **Staining** | **Intensity** | **Fraction** | **Location** |
| --- | --- | --- | --- | --- | --- | --- | --- | --- | --- |
| *TPT1* | Normal colon | HPA039437 | 1960 | Male, 84 years | Glandular cells | Medium | Moderate | 75%–25% | Cytoplasmic/membranous, nuclear |
| *TPT1* | Colorectal cancer | HPA073929 | 2947 | Male, 83 years | Tumor cells | High | Strong | >75% | Cytoplasmic/membranous |
| *SLC1A5* | Normal colon | HPA035239 | 1857 | Male, 47 years | Endothelial cells | Medium | Moderate | >75% | Cytoplasmic/membranous |
| *SLC1A5* | Colorectal cancer | HPA035240 | 2947 | - | Tumor cells | High | Strong | >75% | Cytoplasmic/membranous |
| *SOX4* | Normal colon | HPA029901 | 1993 | Male, 56 years | Glandular cells | Low | Weak | >75% | Cytoplasmic/membranous, nuclear |
| *SOX4* | Colorectal cancer | HPA029901 | 2947 | Male, 83 years | Tumor cells | Medium | Moderate | >75% | Cytoplasmic/membranous, nuclear |
