## Additional file for "An RGS5-positive fibroblast to WNT-like epithelial TGFB axis in colorectal liver metastasis, replicated in an independent paired cohort": Additional_file_3.pdf

### Table S1

Table S1. Differentially expressed marker genes for each malignant epithelial (C0-C6) and fibroblast (F0-F4) subcluster.

| compartment | subcluster | gene | avg_log2FC | pct.1 | pct.2 | p_val | p_val_adj |
|---|---|---|---|---|---|---|---|
| Malignant epithelial | C0 Is ECs | ITGA10 | 2.98 | 0.108 | 0.018 | 5.16e-47 | 1.74e-42 |
| Malignant epithelial | C0 Is ECs | CDH26 | 2.416 | 0.244 | 0.051 | 7.78e-96 | 2.62e-91 |
| Malignant epithelial | C0 Is ECs | CLCF1 | 2.095 | 0.588 | 0.233 | 5.509999999999999e-180 | 1.86e-175 |
| Malignant epithelial | C0 Is ECs | CP | 2.08 | 0.398 | 0.107 | 6.959999999999999e-138 | 2.35e-133 |
| Malignant epithelial | C0 Is ECs | SAA1 | 2.016 | 0.649 | 0.267 | 4.6e-176 | 1.55e-171 |
| Malignant epithelial | C0 Is ECs | SAA2 | 1.974 | 0.286 | 0.105 | 1.38e-66 | 4.66e-62 |
| Malignant epithelial | C0 Is ECs | C3 | 1.96 | 0.427 | 0.124 | 5.89e-139 | 1.99e-134 |
| Malignant epithelial | C0 Is ECs | MAP3K8 | 1.959 | 0.57 | 0.248 | 6.429999999999999e-152 | 2.17e-147 |
| Malignant epithelial | C0 Is ECs | ANGPTL4 | 1.934 | 0.305 | 0.11 | 3.46e-73 | 1.17e-68 |
| Malignant epithelial | C0 Is ECs | RHCG | 1.932 | 0.109 | 0.034 | 1.02e-27 | 3.45e-23 |
| Malignant epithelial | C0 Is ECs | CREG2 | 1.918 | 0.114 | 0.037 | 2.28e-28 | 7.689999999999999e-24 |
| Malignant epithelial | C0 Is ECs | NEURL3 | 1.914 | 0.106 | 0.029 | 4.93e-31 | 1.66e-26 |
| Malignant epithelial | C0 Is ECs | PAEP | 1.837 | 0.19 | 0.073 | 5.29e-38 | 1.78e-33 |
| Malignant epithelial | C0 Is ECs | IDO1 | 1.836 | 0.289 | 0.103 | 1.53e-68 | 5.140000000000001e-64 |
| Malignant epithelial | C0 Is ECs | TNFAIP2 | 1.816 | 0.365 | 0.13 | 6.08e-91 | 2.05e-86 |
| Malignant epithelial | C0 Is ECs | SNAI2 | 1.81 | 0.156 | 0.057 | 1.59e-33 | 5.36e-29 |
| Malignant epithelial | C0 Is ECs | GGT5 | 1.772 | 0.21 | 0.068 | 1.2e-52 | 4.06e-48 |
| Malignant epithelial | C0 Is ECs | CFL2 | 1.763 | 0.238 | 0.095 | 1.07e-47 | 3.61e-43 |
| Malignant epithelial | C0 Is ECs | LINC01503 | 1.76 | 0.195 | 0.063 | 2.51e-49 | 8.47e-45 |
| Malignant epithelial | C0 Is ECs | NR4A2 | 1.757 | 0.721 | 0.394 | 5.68e-183 | 1.91e-178 |
| Malignant epithelial | C0 Is ECs | PTGES | 1.704 | 0.447 | 0.187 | 1.09e-101 | 3.68e-97 |
| Malignant epithelial | C0 Is ECs | TM4SF18 | 1.679 | 0.469 | 0.145 | 4.849999999999999e-140 | 1.63e-135 |
| Malignant epithelial | C0 Is ECs | DLL3 | 1.659 | 0.104 | 0.041 | 4.03e-20 | 1.36e-15 |
| Malignant epithelial | C0 Is ECs | CALB1 | 1.658 | 0.868 | 0.363 | 1.39e-290 | 4.67e-286 |
| Malignant epithelial | C0 Is ECs | XDH | 1.658 | 0.157 | 0.051 | 3.36e-38 | 1.13e-33 |
| Malignant epithelial | C0 Is ECs | RHOV | 1.656 | 0.448 | 0.23 | 2.23e-75 | 7.51e-71 |
| Malignant epithelial | C0 Is ECs | SECTM1 | 1.647 | 0.177 | 0.058 | 2e-43 | 6.75e-39 |
| Malignant epithelial | C0 Is ECs | TSNARE1 | 1.614 | 0.214 | 0.093 | 3.34e-36 | 1.12e-31 |
| Malignant epithelial | C0 Is ECs | LYPD3 | 1.604 | 0.278 | 0.166 | 1.58e-26 | 5.33e-22 |
| Malignant epithelial | C0 Is ECs | RP11-142A22.4 | 1.577 | 0.112 | 0.038 | 1.98e-25 | 6.680000000000001e-21 |
| Malignant epithelial | C0 Is ECs | MIR4435-2HG | 1.575 | 0.933 | 0.504 | 3.39e-311 | 1.14e-306 |
| Malignant epithelial | C0 Is ECs | ADGRF1 | 1.573 | 0.196 | 0.065 | 5.4e-48 | 1.82e-43 |
| Malignant epithelial | C0 Is ECs | SERPINA5 | 1.572 | 0.116 | 0.038 | 2.61e-28 | 8.78e-24 |
| Malignant epithelial | C0 Is ECs | LINC01589 | 1.572 | 0.119 | 0.045 | 3.27e-24 | 1.1e-19 |
| Malignant epithelial | C0 Is ECs | FLRT3 | 1.548 | 0.119 | 0.041 | 5.64e-27 | 1.9e-22 |
| Malignant epithelial | C0 Is ECs | HAS2 | 1.534 | 0.212 | 0.083 | 2.26e-42 | 7.63e-38 |
| Malignant epithelial | C0 Is ECs | TRIB2 | 1.511 | 0.226 | 0.092 | 1.63e-42 | 5.48e-38 |
| Malignant epithelial | C0 Is ECs | HACD1 | 1.509 | 0.337 | 0.127 | 1.11e-75 | 3.72e-71 |
| Malignant epithelial | C0 Is ECs | MAPK12 | 1.508 | 0.13 | 0.055 | 1.07e-21 | 3.61e-17 |
| Malignant epithelial | C0 Is ECs | UPK3B | 1.489 | 0.158 | 0.063 | 1.41e-29 | 4.76e-25 |
| Malignant epithelial | C0 Is ECs | LIF | 1.481 | 0.467 | 0.222 | 4.77e-86 | 1.61e-81 |
| Malignant epithelial | C0 Is ECs | C9orf116 | 1.478 | 0.168 | 0.073 | 9.279999999999999e-28 | 3.129999999999999e-23 |
| Malignant epithelial | C0 Is ECs | ALDH1A3 | 1.476 | 0.27 | 0.096 | 6.39e-62 | 2.15e-57 |
| Malignant epithelial | C0 Is ECs | DIO3 | 1.451 | 0.389 | 0.188 | 1.45e-62 | 4.87e-58 |
| Malignant epithelial | C0 Is ECs | CALB2 | 1.425 | 0.499 | 0.176 | 3.3e-128 | 1.11e-123 |
| Malignant epithelial | C0 Is ECs | OBSCN | 1.425 | 0.104 | 0.042 | 1.53e-18 | 5.14e-14 |
| Malignant epithelial | C0 Is ECs | WNT10A | 1.419 | 0.419 | 0.188 | 1.12e-79 | 3.78e-75 |
| Malignant epithelial | C0 Is ECs | PLTP | 1.413 | 0.481 | 0.241 | 9.82e-87 | 3.31e-82 |
| Malignant epithelial | C0 Is ECs | ABCA7 | 1.404 | 0.348 | 0.161 | 2.13e-57 | 7.17e-53 |
| Malignant epithelial | C0 Is ECs | SOCS3 | 1.403 | 0.684 | 0.368 | 9.81e-140 | 3.3e-135 |
| Malignant epithelial | C0 Is ECs | CXCL1 | 1.392 | 0.79 | 0.471 | 1.52e-150 | 5.12e-146 |
| Malignant epithelial | C0 Is ECs | MT1A | 1.384 | 0.177 | 0.078 | 7.52e-28 | 2.53e-23 |
| Malignant epithelial | C0 Is ECs | C15orf48 | 1.382 | 0.974 | 0.776 | 1.5e-225 | 5.04e-221 |
| Malignant epithelial | C0 Is ECs | S100A2 | 1.372 | 0.946 | 0.516 | 1.54e-276 | 5.17e-272 |
| Malignant epithelial | C0 Is ECs | SLC9A7 | 1.371 | 0.167 | 0.072 | 4.39e-27 | 1.48e-22 |
| Malignant epithelial | C0 Is ECs | IQCD | 1.371 | 0.104 | 0.044 | 8.72e-18 | 2.94e-13 |
| Malignant epithelial | C0 Is ECs | SOX12 | 1.368 | 0.178 | 0.081 | 4.37e-27 | 1.47e-22 |
| Malignant epithelial | C0 Is ECs | MUC16 | 1.366 | 0.244 | 0.087 | 2.8e-54 | 9.420000000000001e-50 |
| Malignant epithelial | C0 Is ECs | WNT7B | 1.365 | 0.176 | 0.078 | 3.84e-28 | 1.29e-23 |

| compartment | subcluster | gene | avg_log2FC | pct.1 | pct.2 | p_val | p_val_adj |
|---|---|---|---|---|---|---|---|
| Malignant epithelial | C0 Is ECs | CXCL2 | 1.363 | 0.857 | 0.561 | 8.549999999999999e-187 | 2.88e-182 |
| Malignant epithelial | C0 Is ECs | TYMP | 1.362 | 0.914 | 0.582 | 2.47e-238 | 8.33e-234 |
| Malignant epithelial | C0 Is ECs | CMTM3 | 1.361 | 0.259 | 0.116 | 2.56e-42 | 8.63e-38 |
| Malignant epithelial | C0 Is ECs | CAV1 | 1.354 | 0.836 | 0.379 | 5.859999999999999e-229 | 1.97e-224 |
| Malignant epithelial | C0 Is ECs | NNMT | 1.342 | 0.141 | 0.06 | 5.189999999999999e-23 | 1.75e-18 |
| Malignant epithelial | C0 Is ECs | ID1 | 1.338 | 0.947 | 0.776 | 1.89e-160 | 6.36e-156 |
| Malignant epithelial | C0 Is ECs | PHLDA3 | 1.337 | 0.67 | 0.382 | 7.6e-124 | 2.56e-119 |
| Malignant epithelial | C0 Is ECs | TSPAN4 | 1.337 | 0.234 | 0.102 | 1.62e-38 | 5.47e-34 |
| Malignant epithelial | C0 Is ECs | LETM2 | 1.332 | 0.369 | 0.157 | 1.73e-70 | 5.83e-66 |
| Malignant epithelial | C0 Is ECs | TMEM132A | 1.328 | 0.182 | 0.092 | 2.33e-22 | 7.85e-18 |
| Malignant epithelial | C0 Is ECs | NFKBID | 1.326 | 0.174 | 0.085 | 8.5e-23 | 2.86e-18 |
| Malignant epithelial | C0 Is ECs | PPP1R1C | 1.322 | 0.243 | 0.108 | 1.89e-39 | 6.38e-35 |
| Malignant epithelial | C0 Is ECs | HMGA2 | 1.313 | 0.475 | 0.234 | 9.490000000000001e-79 | 3.2e-74 |
| Malignant epithelial | C0 Is ECs | GBP4 | 1.311 | 0.21 | 0.1 | 3.45e-29 | 1.16e-24 |
| Malignant epithelial | C0 Is ECs | DDIT4 | 1.3 | 0.957 | 0.739 | 6.66e-201 | 2.24e-196 |
| Malignant epithelial | C0 Is ECs | EMP3 | 1.293 | 0.695 | 0.379 | 8.619999999999999e-136 | 2.91e-131 |
| Malignant epithelial | C0 Is ECs | MYADM | 1.29 | 0.813 | 0.47 | 5.289999999999999e-178 | 1.78e-173 |
| Malignant epithelial | C0 Is ECs | CACNA1A | 1.29 | 0.264 | 0.115 | 5.63e-44 | 1.9e-39 |
| Malignant epithelial | C0 Is ECs | DCBLD1 | 1.288 | 0.388 | 0.183 | 9.81e-65 | 3.3e-60 |
| Malignant epithelial | C0 Is ECs | TPST1 | 1.286 | 0.151 | 0.067 | 4.039999999999999e-23 | 1.36e-18 |
| Malignant epithelial | C0 Is ECs | CTSC | 1.277 | 0.864 | 0.555 | 3.15e-187 | 1.06e-182 |
| Malignant epithelial | C0 Is ECs | CXCL8 | 1.277 | 0.858 | 0.599 | 7.19e-162 | 2.42e-157 |
| Malignant epithelial | C0 Is ECs | PRDM1 | 1.273 | 0.404 | 0.228 | 2.01e-49 | 6.78e-45 |
| Malignant epithelial | C0 Is ECs | SLC5A3 | 1.27 | 0.609 | 0.34 | 7.49e-100 | 2.52e-95 |
| Malignant epithelial | C0 Is ECs | PSD3 | 1.264 | 0.131 | 0.055 | 6.680000000000001e-22 | 2.25e-17 |
| Malignant epithelial | C0 Is ECs | SULF2 | 1.26 | 0.639 | 0.353 | 6.83e-116 | 2.3e-111 |
| Malignant epithelial | C0 Is ECs | LINC00460 | 1.258 | 0.331 | 0.178 | 7.63e-41 | 2.57e-36 |
| Malignant epithelial | C0 Is ECs | LINC00152 | 1.252 | 0.82 | 0.522 | 5.23e-172 | 1.76e-167 |
| Malignant epithelial | C0 Is ECs | ANPEP | 1.246 | 0.254 | 0.102 | 6.219999999999999e-48 | 2.1e-43 |
| Malignant epithelial | C0 Is ECs | ICAM2 | 1.246 | 0.238 | 0.117 | 1.97e-31 | 6.630000000000001e-27 |
| Malignant epithelial | C0 Is ECs | CXCL10 | 1.246 | 0.129 | 0.077 | 4.34e-10 | 1.46e-05 |
| Malignant epithelial | C0 Is ECs | CH17-360D5.2 | 1.243 | 0.201 | 0.09 | 6.67e-31 | 2.25e-26 |
| Malignant epithelial | C0 Is ECs | TNFSF12 | 1.242 | 0.11 | 0.044 | 7.03e-20 | 2.37e-15 |
| Malignant epithelial | C0 Is ECs | EHD2 | 1.237 | 0.112 | 0.048 | 7.02e-18 | 2.37e-13 |
| Malignant epithelial | C0 Is ECs | FKBP10 | 1.234 | 0.498 | 0.242 | 1.46e-86 | 4.94e-82 |
| Malignant epithelial | C0 Is ECs | EREG | 1.233 | 0.713 | 0.458 | 1.32e-109 | 4.44e-105 |
| Malignant epithelial | C0 Is ECs | CEBPD | 1.23 | 0.908 | 0.703 | 2.83e-185 | 9.53e-181 |
| Malignant epithelial | C0 Is ECs | PLAUR | 1.219 | 0.882 | 0.634 | 2.19e-179 | 7.39e-175 |
| Malignant epithelial | C0 Is ECs | TNFSF9 | 1.212 | 0.23 | 0.116 | 2.76e-29 | 9.28e-25 |
| Malignant epithelial | C0 Is ECs | SPON1 | 1.21 | 0.424 | 0.229 | 4.25e-58 | 1.43e-53 |
| Malignant epithelial | C0 Is ECs | TNNT1 | 1.207 | 0.761 | 0.373 | 1.89e-170 | 6.35e-166 |
| Malignant epithelial | C0 Is ECs | KCP | 1.204 | 0.229 | 0.119 | 4.87e-28 | 1.64e-23 |
| Malignant epithelial | C0 Is ECs | PLAU | 1.203 | 0.706 | 0.323 | 4.71e-155 | 1.59e-150 |
| Malignant epithelial | C0 Is ECs | CXCL3 | 1.2 | 0.867 | 0.628 | 4.34e-146 | 1.46e-141 |
| Malignant epithelial | C0 Is ECs | TFPI | 1.198 | 0.447 | 0.245 | 1.04e-60 | 3.51e-56 |
| Malignant epithelial | C0 Is ECs | RBM43 | 1.192 | 0.111 | 0.048 | 2.41e-17 | 8.12e-13 |
| Malignant epithelial | C0 Is ECs | EGFL7 | 1.191 | 0.183 | 0.084 | 6.12e-27 | 2.06e-22 |
| Malignant epithelial | C0 Is ECs | EID3 | 1.189 | 0.154 | 0.087 | 2.4e-14 | 8.08e-10 |
| Malignant epithelial | C0 Is ECs | RP11-11N9.4 | 1.174 | 0.324 | 0.149 | 1.41e-52 | 4.75e-48 |
| Malignant epithelial | C0 Is ECs | TGM2 | 1.171 | 0.69 | 0.374 | 3.25e-129 | 1.1e-124 |
| Malignant epithelial | C0 Is ECs | SNRK | 1.161 | 0.246 | 0.145 | 7.379999999999999e-23 | 2.49e-18 |
| Malignant epithelial | C0 Is ECs | LOXL1 | 1.158 | 0.132 | 0.066 | 3.28e-16 | 1.1e-11 |
| Malignant epithelial | C0 Is ECs | BIRC3 | 1.154 | 0.589 | 0.328 | 4.78e-84 | 1.61e-79 |
| Malignant epithelial | C0 Is ECs | CITED4 | 1.154 | 0.152 | 0.079 | 2.01e-17 | 6.76e-13 |
| Malignant epithelial | C0 Is ECs | LINC01234 | 1.151 | 0.121 | 0.05 | 2.8e-20 | 9.43e-16 |
| Malignant epithelial | C0 Is ECs | PCDHB14 | 1.15 | 0.133 | 0.059 | 4.13e-20 | 1.39e-15 |
| Malignant epithelial | C0 Is ECs | RAB27B | 1.15 | 0.167 | 0.085 | 5.54e-20 | 1.87e-15 |
| Malignant epithelial | C0 Is ECs | NR2F1-AS1 | 1.15 | 0.108 | 0.052 | 4.04e-14 | 1.36e-09 |
| Malignant epithelial | C0 Is ECs | SNX24 | 1.142 | 0.109 | 0.06 | 9.93e-11 | 3.35e-06 |
| Malignant epithelial | C0 Is ECs | CALD1 | 1.137 | 0.234 | 0.105 | 2.43e-35 | 8.18e-31 |
| Malignant epithelial | C0 Is ECs | MT1E | 1.136 | 0.71 | 0.367 | 5.31e-131 | 1.79e-126 |
| Malignant epithelial | C0 Is ECs | ITPKC | 1.133 | 0.42 | 0.251 | 3.78e-46 | 1.27e-41 |
| Malignant epithelial | C0 Is ECs | NMB | 1.131 | 0.512 | 0.364 | 2.8e-41 | 9.420000000000001e-37 |
| Malignant epithelial | C0 Is ECs | RASA1 | 1.131 | 0.352 | 0.195 | 1.29e-39 | 4.36e-35 |

| compartment | subcluster | gene | avg_log2FC | pct.1 | pct.2 | p_val | p_val_adj |
|---|---|---|---|---|---|---|---|
| Malignant epithelial | C0 Is ECs | IKBKE | 1.128 | 0.238 | 0.137 | 6.87e-23 | 2.32e-18 |
| Malignant epithelial | C0 Is ECs | HYAL3 | 1.128 | 0.106 | 0.061 | 1.69e-09 | 5.69e-05 |
| Malignant epithelial | C0 Is ECs | MESP1 | 1.127 | 0.33 | 0.171 | 1.48e-41 | 5.000000000000001e-37 |
| Malignant epithelial | C0 Is ECs | ELFN1-AS1 | 1.12 | 0.469 | 0.259 | 1.31e-60 | 4.43e-56 |
| Malignant epithelial | C0 Is ECs | PADI2 | 1.12 | 0.146 | 0.062 | 8.989999999999999e-24 | 3.03e-19 |
| Malignant epithelial | C0 Is ECs | SIK1 | 1.119 | 0.232 | 0.121 | 4.850000000000001e-27 | 1.64e-22 |
| Malignant epithelial | C0 Is ECs | HS3ST1 | 1.119 | 0.192 | 0.113 | 2.23e-16 | 7.5e-12 |
| Malignant epithelial | C0 Is ECs | CTD-2325P2.4 | 1.119 | 0.103 | 0.054 | 3.1e-11 | 1.05e-06 |
| Malignant epithelial | C0 Is ECs | ALDH3B1 | 1.118 | 0.335 | 0.191 | 4.2e-36 | 1.42e-31 |
| Malignant epithelial | C0 Is ECs | AGMO | 1.118 | 0.194 | 0.089 | 4.1e-28 | 1.38e-23 |
| Malignant epithelial | C0 Is ECs | MAP4K4 | 1.109 | 0.451 | 0.254 | 5.14e-57 | 1.73e-52 |
| Malignant epithelial | C0 Is ECs | PDP1 | 1.103 | 0.397 | 0.242 | 6.33e-39 | 2.13e-34 |
| Malignant epithelial | C0 Is ECs | DMKN | 1.101 | 0.89 | 0.554 | 6.470000000000001e-184 | 2.18e-179 |
| Malignant epithelial | C0 Is ECs | PIM3 | 1.096 | 0.757 | 0.57 | 1.86e-101 | 6.25e-97 |
| Malignant epithelial | C0 Is ECs | SERPINE2 | 1.096 | 0.425 | 0.283 | 1.65e-33 | 5.550000000000001e-29 |
| Malignant epithelial | C0 Is ECs | KIAA0895 | 1.096 | 0.104 | 0.053 | 1.64e-12 | 5.53e-08 |
| Malignant epithelial | C0 Is ECs | RASAL2 | 1.093 | 0.292 | 0.157 | 5.92e-34 | 1.99e-29 |
| Malignant epithelial | C0 Is ECs | AGAP2-AS1 | 1.092 | 0.196 | 0.104 | 2.12e-21 | 7.15e-17 |
| Malignant epithelial | C0 Is ECs | MARVELD1 | 1.091 | 0.245 | 0.131 | 4.380000000000001e-27 | 1.48e-22 |
| Malignant epithelial | C0 Is ECs | CSTB | 1.089 | 0.998 | 0.973 | 2.09e-215 | 7.04e-211 |
| Malignant epithelial | C0 Is ECs | AVPI1 | 1.089 | 0.671 | 0.51 | 4.23e-65 | 1.43e-60 |
| Malignant epithelial | C0 Is ECs | ODF3B | 1.082 | 0.442 | 0.246 | 1.46e-53 | 4.93e-49 |
| Malignant epithelial | C0 Is ECs | SYT17 | 1.08 | 0.111 | 0.057 | 2.47e-12 | 8.32e-08 |
| Malignant epithelial | C0 Is ECs | LY6E | 1.078 | 0.977 | 0.734 | 3.76e-217 | 1.27e-212 |
| Malignant epithelial | C0 Is ECs | GBP1 | 1.077 | 0.239 | 0.15 | 1.63e-17 | 5.48e-13 |
| Malignant epithelial | C0 Is ECs | ADCY7 | 1.076 | 0.151 | 0.082 | 3.31e-15 | 1.11e-10 |
| Malignant epithelial | C0 Is ECs | MIA | 1.069 | 0.781 | 0.474 | 7.000000000000001e-119 | 2.36e-114 |
| Malignant epithelial | C0 Is ECs | RIPK2 | 1.067 | 0.5 | 0.333 | 4.51e-49 | 1.52e-44 |
| Malignant epithelial | C0 Is ECs | ADA | 1.067 | 0.189 | 0.107 | 2.43e-17 | 8.19e-13 |
| Malignant epithelial | C0 Is ECs | WFDC2 | 1.062 | 0.604 | 0.384 | 8.499999999999999e-68 | 2.86e-63 |
| Malignant epithelial | C0 Is ECs | RP11-37B2.1 | 1.06 | 0.119 | 0.072 | 4.11e-09 | 0.000138 |
| Malignant epithelial | C0 Is ECs | NFKBIA | 1.055 | 0.979 | 0.835 | 8.44e-201 | 2.84e-196 |
| Malignant epithelial | C0 Is ECs | LYL1 | 1.055 | 0.111 | 0.056 | 3.89e-13 | 1.31e-08 |
| Malignant epithelial | C0 Is ECs | LINC01468 | 1.052 | 0.166 | 0.091 | 4.87e-16 | 1.64e-11 |
| Malignant epithelial | C0 Is ECs | HDAC9 | 1.049 | 0.133 | 0.059 | 2.04e-19 | 6.89e-15 |
| Malignant epithelial | C0 Is ECs | CTD-3252C9.4 | 1.042 | 0.737 | 0.47 | 1.24e-103 | 4.19e-99 |
| Malignant epithelial | C0 Is ECs | SLC35G2 | 1.042 | 0.117 | 0.063 | 2.77e-12 | 9.35e-08 |
| Malignant epithelial | C0 Is ECs | CMTM7 | 1.041 | 0.607 | 0.409 | 2.67e-69 | 9.01e-65 |
| Malignant epithelial | C0 Is ECs | IGFBP6 | 1.039 | 0.809 | 0.417 | 2.21e-172 | 7.44e-168 |
| Malignant epithelial | C0 Is ECs | TNNI3 | 1.037 | 0.301 | 0.166 | 9.180000000000001e-32 | 3.09e-27 |
| Malignant epithelial | C0 Is ECs | LGALS1 | 1.036 | 0.962 | 0.672 | 7.42e-224 | 2.5e-219 |
| Malignant epithelial | C0 Is ECs | NAT6 | 1.034 | 0.159 | 0.101 | 8.77e-11 | 2.96e-06 |
| Malignant epithelial | C0 Is ECs | TM4SF1-AS1 | 1.03 | 0.2 | 0.117 | 1.79e-17 | 6.01e-13 |
| Malignant epithelial | C0 Is ECs | SERPING1 | 1.028 | 0.204 | 0.105 | 2.06e-23 | 6.96e-19 |
| Malignant epithelial | C0 Is ECs | PDK4 | 1.025 | 0.28 | 0.173 | 4.75e-21 | 1.6e-16 |
| Malignant epithelial | C0 Is ECs | TFF2 | 1.022 | 0.282 | 0.129 | 3.56e-45 | 1.2e-40 |
| Malignant epithelial | C0 Is ECs | WFDC3 | 1.022 | 0.29 | 0.177 | 1.05e-24 | 3.52e-20 |
| Malignant epithelial | C0 Is ECs | KRT6B | 1.019 | 0.398 | 0.243 | 2.1e-35 | 7.08e-31 |
| Malignant epithelial | C0 Is ECs | SAV1 | 1.019 | 0.342 | 0.196 | 6.2e-34 | 2.09e-29 |
| Malignant epithelial | C0 Is ECs | LINC00346 | 1.017 | 0.185 | 0.097 | 2.52e-20 | 8.48e-16 |
| Malignant epithelial | C0 Is ECs | LINC00936 | 1.017 | 0.259 | 0.161 | 1.12e-19 | 3.76e-15 |
| Malignant epithelial | C0 Is ECs | SOCS6 | 1.016 | 0.435 | 0.254 | 3.52e-48 | 1.19e-43 |
| Malignant epithelial | C0 Is ECs | CAV2 | 1.011 | 0.74 | 0.426 | 1.01e-118 | 3.4e-114 |
| Malignant epithelial | C0 Is ECs | KLF2 | 1.011 | 0.562 | 0.341 | 1.89e-63 | 6.370000000000001e-59 |
| Malignant epithelial | C0 Is ECs | SLPI | 1.01 | 0.72 | 0.544 | 1.66e-62 | 5.58e-58 |
| Malignant epithelial | C0 Is ECs | PDE9A | 1.006 | 0.111 | 0.054 | 2.97e-14 | 1e-09 |
| Malignant epithelial | C0 Is ECs | C16orf74 | 1.004 | 0.326 | 0.181 | 1.17e-34 | 3.95e-30 |
| Malignant epithelial | C0 Is ECs | C1RL | 1.003 | 0.157 | 0.083 | 8.19e-17 | 2.76e-12 |
| Malignant epithelial | C0 Is ECs | PMP22 | 1.002 | 0.137 | 0.09 | 2.16e-08 | 0.000727 |
| Malignant epithelial | C0 Is ECs | ABL2 | 1.001 | 0.391 | 0.215 | 4.149999999999999e-46 | 1.4e-41 |
| Malignant epithelial | C0 Is ECs | SNX21 | 1.001 | 0.234 | 0.134 | 1.07e-22 | 3.62e-18 |
| Malignant epithelial | C0 Is ECs | TIPARP | 0.999 | 0.404 | 0.261 | 1.15e-33 | 3.88e-29 |
| Malignant epithelial | C0 Is ECs | NOTCH2 | 0.997 | 0.208 | 0.109 | 3.56e-23 | 1.2e-18 |
| Malignant epithelial | C0 Is ECs | ICAM1 | 0.996 | 0.188 | 0.12 | 2.22e-12 | 7.47e-08 |

| compartment | subcluster | gene | avg_log2FC | pct.1 | pct.2 | p_val | p_val_adj |
|---|---|---|---|---|---|---|---|
| Malignant epithelial | C0 Is ECs | PHLDA1 | 0.992 | 0.931 | 0.647 | 1.35e-178 | 4.54e-174 |
| Malignant epithelial | C0 Is ECs | S100A4 | 0.99 | 0.984 | 0.831 | 5.02e-226 | 1.69e-221 |
| Malignant epithelial | C0 Is ECs | WISP3 | 0.99 | 0.103 | 0.062 | 3.08e-08 | 0.00104 |
| Malignant epithelial | C0 Is ECs | IRS1 | 0.989 | 0.219 | 0.132 | 4.23e-17 | 1.43e-12 |
| Malignant epithelial | C0 Is ECs | MUC1 | 0.986 | 0.471 | 0.287 | 6.31e-48 | 2.13e-43 |
| Malignant epithelial | C0 Is ECs | MROH6 | 0.985 | 0.421 | 0.272 | 1.82e-36 | 6.12e-32 |
| Malignant epithelial | C0 Is ECs | SLCO3A1 | 0.979 | 0.238 | 0.143 | 4.62e-19 | 1.56e-14 |
| Malignant epithelial | C0 Is ECs | ANXA2R | 0.978 | 0.177 | 0.108 | 3.21e-13 | 1.08e-08 |
| Malignant epithelial | C0 Is ECs | OASL | 0.975 | 0.124 | 0.083 | 5.53e-07 | 0.0186 |
| Malignant epithelial | C0 Is ECs | RP11-489E7.4 | 0.974 | 0.208 | 0.14 | 1.41e-11 | 4.75e-07 |
| Malignant epithelial | C0 Is ECs | CYSRT1 | 0.969 | 0.253 | 0.13 | 1.93e-30 | 6.5e-26 |
| Malignant epithelial | C0 Is ECs | PRSS12 | 0.962 | 0.102 | 0.057 | 1.69e-09 | 5.7e-05 |
| Malignant epithelial | C0 Is ECs | SCRN1 | 0.96 | 0.181 | 0.107 | 8.22e-15 | 2.77e-10 |
| Malignant epithelial | C0 Is ECs | RP11-65I12.1 | 0.959 | 0.46 | 0.269 | 4.98e-49 | 1.68e-44 |
| Malignant epithelial | C0 Is ECs | RAP2B | 0.959 | 0.471 | 0.303 | 2.18e-42 | 7.33e-38 |
| Malignant epithelial | C0 Is ECs | FJX1 | 0.959 | 0.101 | 0.061 | 8.98e-08 | 0.00303 |
| Malignant epithelial | C0 Is ECs | SLC20A1 | 0.958 | 0.762 | 0.515 | 1.24e-94 | 4.19e-90 |
| Malignant epithelial | C0 Is ECs | ASPHD1 | 0.958 | 0.263 | 0.149 | 1.39e-24 | 4.69e-20 |
| Malignant epithelial | C0 Is ECs | APOL1 | 0.95 | 0.442 | 0.266 | 4.23e-46 | 1.43e-41 |
| Malignant epithelial | C0 Is ECs | DNAAF2 | 0.95 | 0.398 | 0.277 | 1.27e-25 | 4.28e-21 |
| Malignant epithelial | C0 Is ECs | ABHD14A | 0.95 | 0.284 | 0.177 | 3.39e-22 | 1.14e-17 |
| Malignant epithelial | C0 Is ECs | PPP1R15B | 0.948 | 0.427 | 0.279 | 3.66e-36 | 1.23e-31 |
| Malignant epithelial | C0 Is ECs | PER1 | 0.944 | 0.258 | 0.17 | 1.34e-16 | 4.51e-12 |
| Malignant epithelial | C0 Is ECs | PKP1 | 0.943 | 0.121 | 0.068 | 6.61e-11 | 2.23e-06 |
| Malignant epithelial | C0 Is ECs | CAPRIN2 | 0.941 | 0.13 | 0.083 | 2.7e-08 | 0.000911 |
| Malignant epithelial | C0 Is ECs | PLAC8 | 0.934 | 0.519 | 0.377 | 3.99e-34 | 1.35e-29 |
| Malignant epithelial | C0 Is ECs | FGD6 | 0.931 | 0.23 | 0.14 | 3.88e-18 | 1.31e-13 |
| Malignant epithelial | C0 Is ECs | CERCAM | 0.931 | 0.13 | 0.066 | 4.23e-15 | 1.43e-10 |
| Malignant epithelial | C0 Is ECs | FAM84B | 0.928 | 0.39 | 0.256 | 2.42e-31 | 8.150000000000001e-27 |
| Malignant epithelial | C0 Is ECs | KLF4 | 0.926 | 0.894 | 0.686 | 2.8e-122 | 9.42e-118 |
| Malignant epithelial | C0 Is ECs | SLC7A7 | 0.923 | 0.262 | 0.154 | 5.059999999999999e-23 | 1.7e-18 |
| Malignant epithelial | C0 Is ECs | AHNAK2 | 0.922 | 0.528 | 0.309 | 2.76e-60 | 9.29e-56 |
| Malignant epithelial | C0 Is ECs | SPRY1 | 0.922 | 0.346 | 0.242 | 3.22e-20 | 1.09e-15 |
| Malignant epithelial | C0 Is ECs | PYGL | 0.922 | 0.231 | 0.138 | 1.29e-18 | 4.34e-14 |
| Malignant epithelial | C0 Is ECs | MAML3 | 0.921 | 0.14 | 0.08 | 2.24e-12 | 7.54e-08 |
| Malignant epithelial | C0 Is ECs | PBX3 | 0.92 | 0.122 | 0.068 | 1.29e-11 | 4.35e-07 |
| Malignant epithelial | C0 Is ECs | RP11-467L13.7 | 0.919 | 0.283 | 0.204 | 9.38e-13 | 3.16e-08 |
| Malignant epithelial | C0 Is ECs | ANXA1 | 0.916 | 0.965 | 0.712 | 1.39e-172 | 4.68e-168 |
| Malignant epithelial | C0 Is ECs | TMCC3 | 0.915 | 0.188 | 0.115 | 2.03e-14 | 6.83e-10 |
| Malignant epithelial | C0 Is ECs | RARRES1 | 0.914 | 0.289 | 0.229 | 6.81e-09 | 0.000229 |
| Malignant epithelial | C0 Is ECs | SUSD3 | 0.911 | 0.138 | 0.08 | 1.37e-11 | 4.61e-07 |
| Malignant epithelial | C0 Is ECs | CASC8 | 0.91 | 0.226 | 0.128 | 6.490000000000001e-21 | 2.19e-16 |
| Malignant epithelial | C0 Is ECs | MIR4458HG | 0.909 | 0.596 | 0.397 | 1.97e-59 | 6.62e-55 |
| Malignant epithelial | C0 Is ECs | ANKRD28 | 0.909 | 0.287 | 0.183 | 3e-21 | 1.01e-16 |
| Malignant epithelial | C0 Is ECs | SNCG | 0.909 | 0.147 | 0.076 | 3.96e-16 | 1.33e-11 |
| Malignant epithelial | C0 Is ECs | WEE1 | 0.908 | 0.554 | 0.403 | 6.52e-40 | 2.2e-35 |
| Malignant epithelial | C0 Is ECs | BCAM | 0.905 | 0.365 | 0.206 | 5.63e-38 | 1.9e-33 |
| Malignant epithelial | C0 Is ECs | TMEM42 | 0.903 | 0.233 | 0.144 | 3.12e-17 | 1.05e-12 |
| Malignant epithelial | C0 Is ECs | RP11-2C24.9 | 0.901 | 0.182 | 0.108 | 1.79e-14 | 6.04e-10 |
| Malignant epithelial | C0 Is ECs | SCO2 | 0.898 | 0.52 | 0.368 | 3.36e-40 | 1.13e-35 |
| Malignant epithelial | C0 Is ECs | CCDC6 | 0.89 | 0.444 | 0.302 | 1.53e-33 | 5.17e-29 |
| Malignant epithelial | C0 Is ECs | RP11-796E2.4 | 0.888 | 0.125 | 0.077 | 4.52e-09 | 0.000152 |
| Malignant epithelial | C0 Is ECs | ITGB6 | 0.887 | 0.263 | 0.157 | 4.98e-22 | 1.68e-17 |
| Malignant epithelial | C0 Is ECs | ISYNA1 | 0.887 | 0.251 | 0.155 | 2.63e-18 | 8.87e-14 |
| Malignant epithelial | C0 Is ECs | FAM217B | 0.887 | 0.249 | 0.167 | 3.75e-15 | 1.27e-10 |
| Malignant epithelial | C0 Is ECs | IFITM1 | 0.886 | 0.765 | 0.538 | 5.02e-84 | 1.69e-79 |
| Malignant epithelial | C0 Is ECs | RIPK4 | 0.885 | 0.301 | 0.207 | 1.06e-17 | 3.57e-13 |
| Malignant epithelial | C0 Is ECs | EMP1 | 0.883 | 0.732 | 0.519 | 3.1e-77 | 1.05e-72 |
| Malignant epithelial | C0 Is ECs | C15orf52 | 0.882 | 0.214 | 0.136 | 1.88e-14 | 6.33e-10 |
| Malignant epithelial | C0 Is ECs | STIM1 | 0.882 | 0.183 | 0.128 | 7.64e-09 | 0.000257 |
| Malignant epithelial | C0 Is ECs | MBD5 | 0.881 | 0.124 | 0.078 | 3.77e-08 | 0.00127 |
| Malignant epithelial | C0 Is ECs | MT1F | 0.88 | 0.271 | 0.157 | 4.86e-25 | 1.64e-20 |
| Malignant epithelial | C0 Is ECs | PAX8 | 0.879 | 0.222 | 0.13 | 6.62e-19 | 2.23e-14 |
| Malignant epithelial | C0 Is ECs | ACE | 0.879 | 0.138 | 0.088 | 9.13e-09 | 0.000308 |

| compartment | subcluster | gene | avg_log2FC | pct.1 | pct.2 | p_val | p_val_adj |
|---|---|---|---|---|---|---|---|
| Malignant epithelial | C0 Is ECs | NR2F1 | 0.878 | 0.175 | 0.11 | 5.48e-12 | 1.85e-07 |
| Malignant epithelial | C0 Is ECs | KLC3 | 0.871 | 0.162 | 0.079 | 8.44e-20 | 2.85e-15 |
| Malignant epithelial | C0 Is ECs | SH3PXD2A-AS1 | 0.869 | 0.126 | 0.08 | 1.94e-08 | 0.000652 |
| Malignant epithelial | C0 Is ECs | FRMD6-AS1 | 0.868 | 0.106 | 0.058 | 2.25e-10 | 7.58e-06 |
| Malignant epithelial | C0 Is ECs | VMP1 | 0.867 | 0.95 | 0.806 | 2.34e-145 | 7.87e-141 |
| Malignant epithelial | C0 Is ECs | PLEKHF2 | 0.867 | 0.381 | 0.302 | 3e-15 | 1.01e-10 |
| Malignant epithelial | C0 Is ECs | ARL4C | 0.866 | 0.513 | 0.301 | 9.78e-56 | 3.29e-51 |
| Malignant epithelial | C0 Is ECs | NME4 | 0.861 | 0.614 | 0.423 | 8.83e-60 | 2.98e-55 |
| Malignant epithelial | C0 Is ECs | TWISTNB | 0.861 | 0.422 | 0.334 | 6.34e-17 | 2.14e-12 |
| Malignant epithelial | C0 Is ECs | ZBP1 | 0.857 | 0.2 | 0.121 | 3.15e-15 | 1.06e-10 |
| Malignant epithelial | C0 Is ECs | YPEL5 | 0.856 | 0.729 | 0.584 | 1.13e-58 | 3.8e-54 |
| Malignant epithelial | C0 Is ECs | DUSP1 | 0.855 | 0.979 | 0.849 | 8.339999999999999e-147 | 2.81e-142 |
| Malignant epithelial | C0 Is ECs | CCL20 | 0.855 | 0.607 | 0.495 | 2.23e-28 | 7.499999999999999e-24 |
| Malignant epithelial | C0 Is ECs | PITPNC1 | 0.855 | 0.322 | 0.211 | 9.859999999999999e-23 | 3.32e-18 |
| Malignant epithelial | C0 Is ECs | IL15RA | 0.855 | 0.195 | 0.124 | 1.91e-13 | 6.43e-09 |
| Malignant epithelial | C0 Is ECs | ARHGAP5-AS1 | 0.851 | 0.238 | 0.147 | 8.38e-18 | 2.82e-13 |
| Malignant epithelial | C0 Is ECs | DNAL4 | 0.851 | 0.227 | 0.147 | 1.03e-14 | 3.47e-10 |
| Malignant epithelial | C0 Is ECs | TNFRSF1A | 0.849 | 0.563 | 0.393 | 4.22e-48 | 1.42e-43 |
| Malignant epithelial | C0 Is ECs | SYNE4 | 0.848 | 0.162 | 0.101 | 1.99e-11 | 6.71e-07 |
| Malignant epithelial | C0 Is ECs | CSTA | 0.847 | 0.331 | 0.209 | 2.75e-25 | 9.28e-21 |
| Malignant epithelial | C0 Is ECs | ZNF217 | 0.847 | 0.326 | 0.207 | 1.06e-23 | 3.57e-19 |
| Malignant epithelial | C0 Is ECs | BBC3 | 0.845 | 0.662 | 0.492 | 1.33e-57 | 4.48e-53 |
| Malignant epithelial | C0 Is ECs | NFKB2 | 0.842 | 0.306 | 0.207 | 1.29e-18 | 4.36e-14 |
| Malignant epithelial | C0 Is ECs | RP11-983P16.4 | 0.842 | 0.117 | 0.07 | 6.78e-09 | 0.000229 |
| Malignant epithelial | C0 Is ECs | PCGF2 | 0.839 | 0.251 | 0.163 | 3.46e-16 | 1.17e-11 |
| Malignant epithelial | C0 Is ECs | NAB2 | 0.838 | 0.154 | 0.101 | 2.28e-09 | 7.69e-05 |
| Malignant epithelial | C0 Is ECs | IRF1 | 0.837 | 0.824 | 0.652 | 1.34e-77 | 4.530000000000001e-73 |
| Malignant epithelial | C0 Is ECs | CDC42EP4 | 0.833 | 0.442 | 0.326 | 5.94e-24 | 2e-19 |
| Malignant epithelial | C0 Is ECs | TMC8 | 0.833 | 0.253 | 0.173 | 8.47e-14 | 2.86e-09 |
| Malignant epithelial | C0 Is ECs | ALCAM | 0.832 | 0.391 | 0.241 | 1.63e-33 | 5.48e-29 |
| Malignant epithelial | C0 Is ECs | NFIC | 0.832 | 0.299 | 0.214 | 8.44e-15 | 2.84e-10 |
| Malignant epithelial | C0 Is ECs | KYNU | 0.832 | 0.211 | 0.133 | 2.59e-14 | 8.73e-10 |
| Malignant epithelial | C0 Is ECs | ATHL1 | 0.83 | 0.218 | 0.139 | 3.1e-14 | 1.04e-09 |
| Malignant epithelial | C0 Is ECs | LITAF | 0.828 | 0.705 | 0.549 | 2.05e-62 | 6.9e-58 |
| Malignant epithelial | C0 Is ECs | KCNK1 | 0.824 | 0.34 | 0.244 | 9.29e-18 | 3.13e-13 |
| Malignant epithelial | C0 Is ECs | APOL2 | 0.823 | 0.24 | 0.139 | 4e-21 | 1.35e-16 |
| Malignant epithelial | C0 Is ECs | LAMA3 | 0.82 | 0.555 | 0.344 | 3.25e-55 | 1.1e-50 |
| Malignant epithelial | C0 Is ECs | CCNL1 | 0.818 | 0.942 | 0.813 | 6.089999999999999e-104 | 2.05e-99 |
| Malignant epithelial | C0 Is ECs | RHOD | 0.818 | 0.562 | 0.36 | 6.67e-53 | 2.25e-48 |
| Malignant epithelial | C0 Is ECs | ANKRD37 | 0.818 | 0.384 | 0.284 | 1.15e-17 | 3.88e-13 |
| Malignant epithelial | C0 Is ECs | ITGB8 | 0.817 | 0.348 | 0.222 | 1.39e-25 | 4.680000000000001e-21 |
| Malignant epithelial | C0 Is ECs | PTPN14 | 0.815 | 0.144 | 0.084 | 8.64e-12 | 2.91e-07 |
| Malignant epithelial | C0 Is ECs | SOCS1 | 0.815 | 0.235 | 0.175 | 5.15e-09 | 0.000173 |
| Malignant epithelial | C0 Is ECs | GLI4 | 0.811 | 0.252 | 0.17 | 2.85e-14 | 9.6e-10 |
| Malignant epithelial | C0 Is ECs | ADAM19 | 0.811 | 0.22 | 0.153 | 4.09e-11 | 1.38e-06 |
| Malignant epithelial | C0 Is ECs | RP11-350N15.5 | 0.808 | 0.115 | 0.071 | 5.03e-08 | 0.0017 |
| Malignant epithelial | C0 Is ECs | INPP4B | 0.807 | 0.26 | 0.174 | 2.56e-15 | 8.63e-11 |
| Malignant epithelial | C0 Is ECs | PLIN2 | 0.805 | 0.635 | 0.485 | 2.3e-42 | 7.75e-38 |
| Malignant epithelial | C0 Is ECs | SEMA3A | 0.805 | 0.125 | 0.078 | 1.7e-08 | 0.000572 |
| Malignant epithelial | C0 Is ECs | KLK7 | 0.804 | 0.232 | 0.132 | 5.940000000000001e-21 | 2e-16 |
| Malignant epithelial | C0 Is ECs | SLC22A18AS | 0.803 | 0.196 | 0.133 | 1.17e-10 | 3.94e-06 |
| Malignant epithelial | C0 Is ECs | ELK3 | 0.802 | 0.19 | 0.113 | 2.17e-15 | 7.32e-11 |
| Malignant epithelial | C0 Is ECs | RUFY2 | 0.801 | 0.127 | 0.086 | 1.08e-06 | 0.0365 |
| Malignant epithelial | C0 Is ECs | ZNF414 | 0.8 | 0.357 | 0.254 | 9.26e-20 | 3.12e-15 |
| Malignant epithelial | C0 Is ECs | NFKBIZ | 0.798 | 0.584 | 0.408 | 3.97e-45 | 1.34e-40 |
| Malignant epithelial | C0 Is ECs | BCL2L11 | 0.794 | 0.298 | 0.205 | 1.85e-17 | 6.25e-13 |
| Malignant epithelial | C0 Is ECs | SEMA3C | 0.787 | 0.334 | 0.235 | 3.92e-18 | 1.32e-13 |
| Malignant epithelial | C0 Is ECs | DYRK4 | 0.785 | 0.258 | 0.182 | 1.8e-12 | 6.06e-08 |
| Malignant epithelial | C0 Is ECs | RIN1 | 0.783 | 0.182 | 0.119 | 5.66e-11 | 1.91e-06 |
| Malignant epithelial | C0 Is ECs | TMEM55A | 0.782 | 0.22 | 0.148 | 5.03e-12 | 1.69e-07 |
| Malignant epithelial | C0 Is ECs | PMAIP1 | 0.781 | 0.803 | 0.617 | 5.68e-73 | 1.91e-68 |
| Malignant epithelial | C0 Is ECs | UBTD1 | 0.781 | 0.218 | 0.15 | 2.58e-11 | 8.68e-07 |
| Malignant epithelial | C0 Is ECs | EHBP1 | 0.779 | 0.266 | 0.18 | 3.02e-15 | 1.02e-10 |
| Malignant epithelial | C0 Is ECs | FAM174A | 0.775 | 0.33 | 0.228 | 2.71e-18 | 9.14e-14 |

| compartment | subcluster | gene | avg_log2FC | pct.1 | pct.2 | p_val | p_val_adj |
|---|---|---|---|---|---|---|---|
| Malignant epithelial | C0 Is ECs | ZFP36 | 0.772 | 0.985 | 0.866 | 2.78e-120 | 9.38e-116 |
| Malignant epithelial | C0 Is ECs | B4GALT5 | 0.772 | 0.401 | 0.302 | 1.18e-19 | 3.97e-15 |
| Malignant epithelial | C0 Is ECs | IRF9 | 0.769 | 0.272 | 0.195 | 6.55e-13 | 2.21e-08 |
| Malignant epithelial | C0 Is ECs | AHR | 0.767 | 0.444 | 0.32 | 2.21e-26 | 7.46e-22 |
| Malignant epithelial | C0 Is ECs | JUNB | 0.765 | 0.995 | 0.919 | 3.19e-106 | 1.07e-101 |
| Malignant epithelial | C0 Is ECs | PNRC1 | 0.764 | 0.834 | 0.69 | 2.06e-68 | 6.93e-64 |
| Malignant epithelial | C0 Is ECs | NOL3 | 0.764 | 0.375 | 0.268 | 1.24e-20 | 4.18e-16 |
| Malignant epithelial | C0 Is ECs | MRPS6 | 0.763 | 0.765 | 0.657 | 4.69e-58 | 1.58e-53 |
| Malignant epithelial | C0 Is ECs | SRGAP2 | 0.763 | 0.114 | 0.074 | 4.97e-07 | 0.0167 |
| Malignant epithelial | C0 Is ECs | PKN1 | 0.762 | 0.213 | 0.141 | 4.22e-12 | 1.42e-07 |
| Malignant epithelial | C0 Is ECs | IFITM3 | 0.757 | 0.992 | 0.927 | 1.22e-158 | 4.12e-154 |
| Malignant epithelial | C0 Is ECs | COQ7 | 0.756 | 0.328 | 0.259 | 6.39e-10 | 2.15e-05 |
| Malignant epithelial | C0 Is ECs | PIAS3 | 0.756 | 0.126 | 0.08 | 3.05e-08 | 0.00103 |
| Malignant epithelial | C0 Is ECs | SSBP4 | 0.755 | 0.474 | 0.342 | 6.81e-31 | 2.3e-26 |
| Malignant epithelial | C0 Is ECs | CEBPB | 0.749 | 0.951 | 0.819 | 1.55e-115 | 5.23e-111 |
| Malignant epithelial | C0 Is ECs | CDA | 0.749 | 0.831 | 0.538 | 1.38e-99 | 4.64e-95 |
| Malignant epithelial | C0 Is ECs | SLC7A11 | 0.746 | 0.26 | 0.168 | 4.5e-17 | 1.52e-12 |
| Malignant epithelial | C0 Is ECs | EPB41L4A-AS1 | 0.745 | 0.74 | 0.61 | 1.08e-54 | 3.63e-50 |
| Malignant epithelial | C0 Is ECs | MYEOV | 0.743 | 0.428 | 0.29 | 1.13e-27 | 3.82e-23 |
| Malignant epithelial | C0 Is ECs | RP11-841O20.2 | 0.742 | 0.166 | 0.102 | 8.56e-12 | 2.88e-07 |
| Malignant epithelial | C0 Is ECs | C19orf66 | 0.741 | 0.42 | 0.313 | 3.77e-21 | 1.27e-16 |
| Malignant epithelial | C0 Is ECs | ITPRIPL2 | 0.739 | 0.298 | 0.203 | 2.41e-17 | 8.12e-13 |
| Malignant epithelial | C0 Is ECs | AXL | 0.737 | 0.315 | 0.18 | 1.57e-28 | 5.3e-24 |
| Malignant epithelial | C0 Is ECs | RNF223 | 0.736 | 0.191 | 0.141 | 3.18e-07 | 0.0107 |
| Malignant epithelial | C0 Is ECs | TNFAIP3 | 0.735 | 0.57 | 0.415 | 1.19e-38 | 4.02e-34 |
| Malignant epithelial | C0 Is ECs | STEAP2 | 0.734 | 0.386 | 0.249 | 7.88e-30 | 2.65e-25 |
| Malignant epithelial | C0 Is ECs | RNF207 | 0.734 | 0.122 | 0.076 | 2.26e-08 | 0.000763 |
| Malignant epithelial | C0 Is ECs | APOL6 | 0.73 | 0.304 | 0.221 | 1.27e-13 | 4.26e-09 |
| Malignant epithelial | C0 Is ECs | LTBR | 0.729 | 0.622 | 0.463 | 4.67e-45 | 1.57e-40 |
| Malignant epithelial | C0 Is ECs | IER3 | 0.728 | 0.963 | 0.869 | 1.53e-80 | 5.14e-76 |
| Malignant epithelial | C0 Is ECs | RP11-670E13.6 | 0.728 | 0.463 | 0.347 | 6.72e-23 | 2.26e-18 |
| Malignant epithelial | C0 Is ECs | RARRES3 | 0.727 | 0.769 | 0.567 | 3.489999999999999e-54 | 1.18e-49 |
| Malignant epithelial | C0 Is ECs | ARL5B | 0.723 | 0.328 | 0.238 | 5.69e-16 | 1.92e-11 |
| Malignant epithelial | C0 Is ECs | IFITM2 | 0.722 | 0.599 | 0.451 | 7.59e-41 | 2.56e-36 |
| Malignant epithelial | C0 Is ECs | TGIF1 | 0.721 | 0.681 | 0.554 | 1.06e-39 | 3.58e-35 |
| Malignant epithelial | C0 Is ECs | OSBPL10 | 0.718 | 0.263 | 0.182 | 3.77e-14 | 1.27e-09 |
| Malignant epithelial | C0 Is ECs | BTG1 | 0.716 | 0.937 | 0.866 | 4.850000000000001e-72 | 1.63e-67 |
| Malignant epithelial | C0 Is ECs | CYP2W1 | 0.716 | 0.416 | 0.323 | 1.15e-16 | 3.87e-12 |
| Malignant epithelial | C0 Is ECs | PRKCDBP | 0.715 | 0.801 | 0.48 | 2.11e-110 | 7.11e-106 |
| Malignant epithelial | C0 Is ECs | JMJD1C | 0.715 | 0.612 | 0.474 | 2.13e-37 | 7.17e-33 |
| Malignant epithelial | C0 Is ECs | MORN4 | 0.715 | 0.101 | 0.059 | 7.06e-08 | 0.00238 |
| Malignant epithelial | C0 Is ECs | BEST1 | 0.714 | 0.136 | 0.088 | 3.93e-08 | 0.00133 |
| Malignant epithelial | C0 Is ECs | TOB1 | 0.713 | 0.881 | 0.765 | 2.42e-77 | 8.160000000000001e-73 |
| Malignant epithelial | C0 Is ECs | HSBP1L1 | 0.71 | 0.561 | 0.437 | 1.97e-30 | 6.620000000000001e-26 |
| Malignant epithelial | C0 Is ECs | RRAS | 0.708 | 0.603 | 0.461 | 6.35e-40 | 2.14e-35 |
| Malignant epithelial | C0 Is ECs | F2RL1 | 0.708 | 0.427 | 0.321 | 8.430000000000001e-21 | 2.84e-16 |
| Malignant epithelial | C0 Is ECs | SLMAP | 0.708 | 0.416 | 0.309 | 1.35e-20 | 4.56e-16 |
| Malignant epithelial | C0 Is ECs | HIST3H2A | 0.705 | 0.592 | 0.433 | 1.45e-39 | 4.89e-35 |
| Malignant epithelial | C0 Is ECs | GBP2 | 0.704 | 0.318 | 0.253 | 2.04e-10 | 6.87e-06 |
| Malignant epithelial | C0 Is ECs | C1orf53 | 0.704 | 0.198 | 0.144 | 3.49e-08 | 0.00118 |
| Malignant epithelial | C0 Is ECs | ARF6 | 0.703 | 0.837 | 0.713 | 2.97e-72 | 1e-67 |
| Malignant epithelial | C0 Is ECs | RARG | 0.702 | 0.192 | 0.13 | 3.7e-10 | 1.25e-05 |
| Malignant epithelial | C0 Is ECs | TGFB1I1 | 0.701 | 0.276 | 0.194 | 8.61e-14 | 2.9e-09 |
| Malignant epithelial | C0 Is ECs | IER5L | 0.699 | 0.324 | 0.244 | 5.68e-13 | 1.91e-08 |
| Malignant epithelial | C0 Is ECs | WARS | 0.699 | 0.342 | 0.275 | 1.49e-10 | 5.01e-06 |
| Malignant epithelial | C0 Is ECs | BCAR3 | 0.699 | 0.157 | 0.103 | 4.06e-09 | 0.000137 |
| Malignant epithelial | C0 Is ECs | IL15 | 0.696 | 0.107 | 0.07 | 1.13e-06 | 0.038 |
| Malignant epithelial | C0 Is ECs | FABP6 | 0.695 | 0.529 | 0.352 | 1.96e-39 | 6.61e-35 |
| Malignant epithelial | C0 Is ECs | EGLN3 | 0.695 | 0.243 | 0.179 | 2.68e-10 | 9.03e-06 |
| Malignant epithelial | C0 Is ECs | GPX1 | 0.694 | 0.98 | 0.928 | 2.65e-154 | 8.919999999999999e-150 |
| Malignant epithelial | C0 Is ECs | PMEPA1 | 0.693 | 0.813 | 0.614 | 3.17e-79 | 1.07e-74 |
| Malignant epithelial | C0 Is ECs | CSRNP1 | 0.692 | 0.613 | 0.479 | 1.89e-34 | 6.349999999999999e-30 |
| Malignant epithelial | C0 Is ECs | P3H4 | 0.692 | 0.218 | 0.165 | 5.96e-08 | 0.00201 |
| Malignant epithelial | C0 Is ECs | FAXDC2 | 0.688 | 0.129 | 0.069 | 6e-13 | 2.02e-08 |

| compartment | subcluster | gene | avg_log2FC | pct.1 | pct.2 | p_val | p_val_adj |
|---|---|---|---|---|---|---|---|
| Malignant epithelial | C0 Is ECs | CTD-2020K17.4 | 0.687 | 0.252 | 0.172 | 1.61e-13 | 5.42e-09 |
| Malignant epithelial | C0 Is ECs | PTPN2 | 0.685 | 0.363 | 0.287 | 1.21e-12 | 4.09e-08 |
| Malignant epithelial | C0 Is ECs | SLC2A4RG | 0.683 | 0.428 | 0.296 | 2.05e-25 | 6.9e-21 |
| Malignant epithelial | C0 Is ECs | FTH1 | 0.682 | 1.0 | 1.0 | 7.230000000000001e-163 | 2.44e-158 |
| Malignant epithelial | C0 Is ECs | SYTL1 | 0.682 | 0.284 | 0.202 | 6.37e-14 | 2.15e-09 |
| Malignant epithelial | C0 Is ECs | BCL9L | 0.682 | 0.199 | 0.15 | 4.28e-07 | 0.0144 |
| Malignant epithelial | C0 Is ECs | C4orf3 | 0.679 | 0.92 | 0.843 | 5.609999999999999e-95 | 1.89e-90 |
| Malignant epithelial | C0 Is ECs | STMN3 | 0.679 | 0.526 | 0.362 | 1.05e-35 | 3.55e-31 |
| Malignant epithelial | C0 Is ECs | CASP8 | 0.677 | 0.351 | 0.26 | 7.81e-15 | 2.63e-10 |
| Malignant epithelial | C0 Is ECs | H2AFY2 | 0.676 | 0.288 | 0.197 | 4.04e-15 | 1.36e-10 |
| Malignant epithelial | C0 Is ECs | CRIP1 | 0.674 | 0.907 | 0.734 | 1.84e-90 | 6.19e-86 |
| Malignant epithelial | C0 Is ECs | PGLS | 0.674 | 0.842 | 0.744 | 8.91e-78 | 3e-73 |
| Malignant epithelial | C0 Is ECs | CFD | 0.669 | 0.444 | 0.325 | 1.59e-22 | 5.34e-18 |
| Malignant epithelial | C0 Is ECs | CCDC88B | 0.669 | 0.18 | 0.132 | 4.9e-07 | 0.0165 |
| Malignant epithelial | C0 Is ECs | ERC1 | 0.669 | 0.124 | 0.082 | 5.83e-07 | 0.0197 |
| Malignant epithelial | C0 Is ECs | MAML2 | 0.668 | 0.187 | 0.134 | 7.52e-08 | 0.00253 |
| Malignant epithelial | C0 Is ECs | KLK8 | 0.667 | 0.362 | 0.251 | 6.15e-21 | 2.07e-16 |
| Malignant epithelial | C0 Is ECs | BCAR1 | 0.667 | 0.41 | 0.316 | 1.79e-17 | 6.05e-13 |
| Malignant epithelial | C0 Is ECs | UNC13D | 0.665 | 0.332 | 0.221 | 8.29e-20 | 2.79e-15 |
| Malignant epithelial | C0 Is ECs | MIF | 0.664 | 0.99 | 0.928 | 1.16e-161 | 3.91e-157 |
| Malignant epithelial | C0 Is ECs | HYAL2 | 0.663 | 0.372 | 0.285 | 6.23e-16 | 2.1e-11 |
| Malignant epithelial | C0 Is ECs | ONECUT3 | 0.662 | 0.309 | 0.201 | 4.35e-20 | 1.47e-15 |
| Malignant epithelial | C0 Is ECs | GAPDH | 0.66 | 1.0 | 0.997 | 1.69e-199 | 5.700000000000001e-195 |
| Malignant epithelial | C0 Is ECs | MTAP | 0.66 | 0.494 | 0.347 | 2.15e-31 | 7.260000000000001e-27 |
| Malignant epithelial | C0 Is ECs | CHMP1B | 0.658 | 0.599 | 0.509 | 6.02e-22 | 2.03e-17 |
| Malignant epithelial | C0 Is ECs | RAC2 | 0.656 | 0.395 | 0.276 | 8.48e-22 | 2.86e-17 |
| Malignant epithelial | C0 Is ECs | PGM1 | 0.654 | 0.156 | 0.108 | 1.89e-07 | 0.00636 |
| Malignant epithelial | C0 Is ECs | SAT1 | 0.653 | 0.981 | 0.964 | 7.809999999999999e-61 | 2.63e-56 |
| Malignant epithelial | C0 Is ECs | U47924.27 | 0.653 | 0.151 | 0.104 | 4.63e-07 | 0.0156 |
| Malignant epithelial | C0 Is ECs | WDR24 | 0.65 | 0.438 | 0.317 | 1.49e-22 | 5.02e-18 |
| Malignant epithelial | C0 Is ECs | SLC16A3 | 0.648 | 0.479 | 0.335 | 1.69e-28 | 5.68e-24 |
| Malignant epithelial | C0 Is ECs | TRIM47 | 0.647 | 0.376 | 0.277 | 5.68e-17 | 1.91e-12 |
| Malignant epithelial | C0 Is ECs | TMEM41A | 0.647 | 0.422 | 0.336 | 6.75e-16 | 2.27e-11 |
| Malignant epithelial | C0 Is ECs | LCN2 | 0.646 | 0.91 | 0.679 | 1.71e-93 | 5.75e-89 |
| Malignant epithelial | C0 Is ECs | FAM3C | 0.646 | 0.708 | 0.564 | 1.02e-52 | 3.43e-48 |
| Malignant epithelial | C0 Is ECs | ERO1A | 0.646 | 0.634 | 0.51 | 2.68e-38 | 9.03e-34 |
| Malignant epithelial | C0 Is ECs | FOSL1 | 0.645 | 0.553 | 0.401 | 3.02e-33 | 1.02e-28 |
| Malignant epithelial | C0 Is ECs | RPL22L1 | 0.644 | 0.926 | 0.839 | 2.19e-87 | 7.39e-83 |
| Malignant epithelial | C0 Is ECs | KIFC3 | 0.644 | 0.167 | 0.113 | 1.75e-08 | 0.000591 |
| Malignant epithelial | C0 Is ECs | F3 | 0.642 | 0.494 | 0.319 | 9.22e-37 | 3.11e-32 |
| Malignant epithelial | C0 Is ECs | RALB | 0.642 | 0.469 | 0.352 | 4.27e-23 | 1.44e-18 |
| Malignant epithelial | C0 Is ECs | DTX2 | 0.641 | 0.407 | 0.314 | 1.48e-16 | 4.97e-12 |
| Malignant epithelial | C0 Is ECs | LTA4H | 0.639 | 0.46 | 0.384 | 1.73e-14 | 5.81e-10 |
| Malignant epithelial | C0 Is ECs | CLIP2 | 0.636 | 0.248 | 0.177 | 4.27e-11 | 1.44e-06 |
| Malignant epithelial | C0 Is ECs | C9orf16 | 0.635 | 0.961 | 0.912 | 1.42e-105 | 4.78e-101 |
| Malignant epithelial | C0 Is ECs | ZFP36L1 | 0.635 | 0.924 | 0.822 | 1.01e-82 | 3.39e-78 |
| Malignant epithelial | C0 Is ECs | BHLHE40 | 0.634 | 0.621 | 0.475 | 1.51e-37 | 5.09e-33 |
| Malignant epithelial | C0 Is ECs | RP11-54H7.4 | 0.634 | 0.48 | 0.306 | 1.29e-35 | 4.34e-31 |
| Malignant epithelial | C0 Is ECs | TACSTD2 | 0.633 | 0.818 | 0.632 | 2.21e-66 | 7.46e-62 |
| Malignant epithelial | C0 Is ECs | RGS10 | 0.632 | 0.561 | 0.422 | 3.37e-30 | 1.13e-25 |
| Malignant epithelial | C0 Is ECs | ATG12 | 0.631 | 0.661 | 0.552 | 2.37e-35 | 7.98e-31 |
| Malignant epithelial | C0 Is ECs | LEMD1 | 0.63 | 0.25 | 0.145 | 4.240000000000001e-21 | 1.43e-16 |
| Malignant epithelial | C0 Is ECs | CBX6 | 0.629 | 0.173 | 0.123 | 1.85e-07 | 0.00624 |
| Malignant epithelial | C0 Is ECs | NCOA3 | 0.626 | 0.256 | 0.18 | 2.38e-12 | 8.01e-08 |
| Malignant epithelial | C0 Is ECs | GAA | 0.626 | 0.206 | 0.146 | 5.17e-09 | 0.000174 |
| Malignant epithelial | C0 Is ECs | PRSS21 | 0.625 | 0.484 | 0.316 | 3.27e-31 | 1.1e-26 |
| Malignant epithelial | C0 Is ECs | CKLF | 0.624 | 0.695 | 0.554 | 2.18e-39 | 7.350000000000001e-35 |
| Malignant epithelial | C0 Is ECs | DENND3 | 0.623 | 0.22 | 0.157 | 1.35e-09 | 4.53e-05 |
| Malignant epithelial | C0 Is ECs | SFTA2 | 0.621 | 0.559 | 0.426 | 9.06e-28 | 3.05e-23 |
| Malignant epithelial | C0 Is ECs | AAED1 | 0.62 | 0.226 | 0.169 | 4.54e-08 | 0.00153 |
| Malignant epithelial | C0 Is ECs | TM4SF1 | 0.617 | 0.969 | 0.859 | 2.7e-104 | 9.11e-100 |
| Malignant epithelial | C0 Is ECs | CARD19 | 0.615 | 0.625 | 0.523 | 7.42e-30 | 2.5e-25 |
| Malignant epithelial | C0 Is ECs | ADSS | 0.614 | 0.448 | 0.354 | 5.14e-19 | 1.73e-14 |
| Malignant epithelial | C0 Is ECs | GCNT3 | 0.61 | 0.537 | 0.413 | 3.22e-25 | 1.08e-20 |

| compartment | subcluster | gene | avg_log2FC | pct.1 | pct.2 | p_val | p_val_adj |
|---|---|---|---|---|---|---|---|
| Malignant epithelial | C0 Is ECs | NTAN1 | 0.609 | 0.272 | 0.203 | 2.26e-10 | 7.63e-06 |
| Malignant epithelial | C0 Is ECs | TP53BP2 | 0.609 | 0.146 | 0.101 | 6.34e-07 | 0.0213 |
| Malignant epithelial | C0 Is ECs | OXSR1 | 0.607 | 0.293 | 0.222 | 1.51e-10 | 5.08e-06 |
| Malignant epithelial | C0 Is ECs | HES4 | 0.604 | 0.667 | 0.478 | 4.27e-52 | 1.44e-47 |
| Malignant epithelial | C0 Is ECs | CD163L1 | 0.604 | 0.246 | 0.152 | 3.92e-16 | 1.32e-11 |
| Malignant epithelial | C0 Is ECs | SERTAD2 | 0.602 | 0.278 | 0.216 | 9.41e-09 | 0.000317 |
| Malignant epithelial | C0 Is ECs | CFLAR | 0.6 | 0.633 | 0.513 | 1e-33 | 3.37e-29 |
| Malignant epithelial | C0 Is ECs | SSFA2 | 0.6 | 0.578 | 0.468 | 5.94e-26 | 2e-21 |
| Malignant epithelial | C0 Is ECs | AP3S1 | 0.599 | 0.799 | 0.711 | 6.1e-49 | 2.05e-44 |
| Malignant epithelial | C0 Is ECs | HAGHL | 0.597 | 0.245 | 0.18 | 3.28e-09 | 0.00011 |
| Malignant epithelial | C0 Is ECs | LAMB3 | 0.596 | 0.713 | 0.587 | 3.58e-40 | 1.21e-35 |
| Malignant epithelial | C0 Is ECs | DSC2 | 0.596 | 0.471 | 0.352 | 3.8e-23 | 1.28e-18 |
| Malignant epithelial | C0 Is ECs | UROS | 0.594 | 0.443 | 0.352 | 2.1e-16 | 7.09e-12 |
| Malignant epithelial | C0 Is ECs | SP140L | 0.594 | 0.233 | 0.159 | 3.74e-12 | 1.26e-07 |
| Malignant epithelial | C0 Is ECs | C19orf33 | 0.592 | 0.991 | 0.936 | 1.69e-142 | 5.69e-138 |
| Malignant epithelial | C0 Is ECs | EFNB2 | 0.591 | 0.599 | 0.457 | 2.02e-31 | 6.82e-27 |
| Malignant epithelial | C0 Is ECs | JUND | 0.589 | 0.957 | 0.816 | 1.18e-72 | 3.99e-68 |
| Malignant epithelial | C0 Is ECs | COL17A1 | 0.589 | 0.635 | 0.453 | 3.4e-43 | 1.15e-38 |
| Malignant epithelial | C0 Is ECs | KLF6 | 0.588 | 0.975 | 0.842 | 8.65e-116 | 2.91e-111 |
| Malignant epithelial | C0 Is ECs | RTP4 | 0.588 | 0.172 | 0.124 | 5.26e-07 | 0.0177 |
| Malignant epithelial | C0 Is ECs | PPP1R13L | 0.584 | 0.35 | 0.268 | 1.65e-12 | 5.57e-08 |
| Malignant epithelial | C0 Is ECs | HIST2H2BE | 0.583 | 0.319 | 0.239 | 1.48e-11 | 4.98e-07 |
| Malignant epithelial | C0 Is ECs | ATN1 | 0.582 | 0.281 | 0.207 | 2.2e-11 | 7.4e-07 |
| Malignant epithelial | C0 Is ECs | MRPS21 | 0.58 | 0.884 | 0.749 | 7.98e-71 | 2.69e-66 |
| Malignant epithelial | C0 Is ECs | MYOF | 0.578 | 0.311 | 0.215 | 2.33e-15 | 7.86e-11 |
| Malignant epithelial | C0 Is ECs | CH17-189H20.1 | 0.577 | 0.442 | 0.351 | 1.73e-15 | 5.84e-11 |
| Malignant epithelial | C0 Is ECs | CDC25B | 0.576 | 0.428 | 0.335 | 1.82e-15 | 6.13e-11 |
| Malignant epithelial | C0 Is ECs | MT2A | 0.575 | 0.857 | 0.644 | 2.7e-71 | 9.100000000000001e-67 |
| Malignant epithelial | C0 Is ECs | CHCHD10 | 0.574 | 0.898 | 0.763 | 1.07e-72 | 3.61e-68 |
| Malignant epithelial | C0 Is ECs | HCFC1R1 | 0.569 | 0.548 | 0.431 | 2.15e-24 | 7.25e-20 |
| Malignant epithelial | C0 Is ECs | EPHX1 | 0.567 | 0.26 | 0.205 | 5.3e-07 | 0.0179 |
| Malignant epithelial | C0 Is ECs | PFKP | 0.562 | 0.421 | 0.307 | 8.71e-21 | 2.93e-16 |
| Malignant epithelial | C0 Is ECs | NR2F6 | 0.56 | 0.444 | 0.354 | 2.96e-16 | 9.97e-12 |
| Malignant epithelial | C0 Is ECs | TRIP6 | 0.56 | 0.435 | 0.352 | 2.08e-14 | 7.01e-10 |
| Malignant epithelial | C0 Is ECs | SFR1 | 0.56 | 0.242 | 0.19 | 1.25e-06 | 0.042 |
| Malignant epithelial | C0 Is ECs | PML | 0.559 | 0.335 | 0.269 | 4.69e-09 | 0.000158 |
| Malignant epithelial | C0 Is ECs | MRPL40 | 0.556 | 0.584 | 0.489 | 6.64e-22 | 2.24e-17 |
| Malignant epithelial | C0 Is ECs | PTRHD1 | 0.556 | 0.582 | 0.488 | 9.51e-22 | 3.2e-17 |
| Malignant epithelial | C0 Is ECs | KLF10 | 0.555 | 0.684 | 0.567 | 1.79e-26 | 6.03e-22 |
| Malignant epithelial | C0 Is ECs | C12orf73 | 0.554 | 0.258 | 0.21 | 1.04e-06 | 0.0349 |
| Malignant epithelial | C0 Is ECs | RBMS1 | 0.554 | 0.222 | 0.171 | 1.27e-06 | 0.0428 |
| Malignant epithelial | C0 Is ECs | TMSB10 | 0.552 | 1.0 | 1.0 | 1.13e-171 | 3.81e-167 |
| Malignant epithelial | C0 Is ECs | CHD2 | 0.549 | 0.425 | 0.343 | 9.2e-14 | 3.1e-09 |
| Malignant epithelial | C0 Is ECs | SRGAP1 | 0.549 | 0.249 | 0.188 | 7.13e-09 | 0.00024 |
| Malignant epithelial | C0 Is ECs | NCK1 | 0.548 | 0.261 | 0.201 | 1.03e-08 | 0.000348 |
| Malignant epithelial | C0 Is ECs | PARP10 | 0.548 | 0.26 | 0.198 | 1.93e-08 | 0.00065 |
| Malignant epithelial | C0 Is ECs | IRS2 | 0.546 | 0.502 | 0.416 | 8.78e-17 | 2.96e-12 |
| Malignant epithelial | C0 Is ECs | BST2 | 0.545 | 0.897 | 0.678 | 5.3e-66 | 1.79e-61 |
| Malignant epithelial | C0 Is ECs | MAP4 | 0.544 | 0.425 | 0.339 | 1.42e-14 | 4.77e-10 |
| Malignant epithelial | C0 Is ECs | GPX4 | 0.542 | 0.972 | 0.927 | 8.210000000000001e-103 | 2.77e-98 |
| Malignant epithelial | C0 Is ECs | RNF135 | 0.542 | 0.302 | 0.233 | 1.13e-09 | 3.82e-05 |
| Malignant epithelial | C0 Is ECs | DYRK2 | 0.541 | 0.304 | 0.236 | 5.43e-10 | 1.83e-05 |
| Malignant epithelial | C0 Is ECs | LYST | 0.541 | 0.195 | 0.139 | 5.65e-08 | 0.0019 |
| Malignant epithelial | C0 Is ECs | LAMB2 | 0.538 | 0.181 | 0.127 | 5e-08 | 0.00169 |
| Malignant epithelial | C0 Is ECs | TICAM1 | 0.537 | 0.326 | 0.255 | 1.91e-10 | 6.43e-06 |
| Malignant epithelial | C0 Is ECs | TP53I3 | 0.536 | 0.385 | 0.313 | 2.01e-11 | 6.77e-07 |
| Malignant epithelial | C0 Is ECs | C6orf132 | 0.535 | 0.54 | 0.501 | 5.52e-10 | 1.86e-05 |
| Malignant epithelial | C0 Is ECs | ABI2 | 0.534 | 0.313 | 0.24 | 1.37e-10 | 4.6e-06 |
| Malignant epithelial | C0 Is ECs | IGF2BP3 | 0.53 | 0.26 | 0.196 | 7.89e-09 | 0.000266 |
| Malignant epithelial | C0 Is ECs | TMSB4X | 0.528 | 1.0 | 0.999 | 6.56e-159 | 2.21e-154 |
| Malignant epithelial | C0 Is ECs | BCL2L13 | 0.526 | 0.343 | 0.272 | 1.32e-10 | 4.45e-06 |
| Malignant epithelial | C0 Is ECs | GPR35 | 0.526 | 0.315 | 0.26 | 1.5e-07 | 0.00505 |
| Malignant epithelial | C0 Is ECs | ANAPC16 | 0.525 | 0.878 | 0.813 | 2.98e-64 | 1e-59 |
| Malignant epithelial | C0 Is ECs | GADD45B | 0.524 | 0.837 | 0.71 | 1.39e-38 | 4.69e-34 |

| compartment | subcluster | gene | avg_log2FC | pct.1 | pct.2 | p_val | p_val_adj |
|---|---|---|---|---|---|---|---|
| Malignant epithelial | C0 Is ECs | OAS2 | 0.524 | 0.19 | 0.136 | 1.22e-07 | 0.0041 |
| Malignant epithelial | C0 Is ECs | EIF4EBP1 | 0.523 | 0.771 | 0.664 | 1.19e-34 | 4e-30 |
| Malignant epithelial | C0 Is ECs | KAT6B | 0.523 | 0.226 | 0.177 | 1.3e-06 | 0.0437 |
| Malignant epithelial | C0 Is ECs | ZFAS1 | 0.512 | 0.995 | 0.978 | 2.21e-84 | 7.45e-80 |
| Malignant epithelial | C0 Is ECs | ARL4A | 0.511 | 0.852 | 0.743 | 1.25e-40 | 4.21e-36 |
| Malignant epithelial | C0 Is ECs | MT1X | 0.511 | 0.465 | 0.361 | 4.94e-19 | 1.67e-14 |
| Malignant epithelial | C0 Is ECs | FOSB | 0.51 | 0.967 | 0.818 | 1.6e-64 | 5.4e-60 |
| Malignant epithelial | C0 Is ECs | ANKLE2 | 0.51 | 0.394 | 0.321 | 1.21e-11 | 4.07e-07 |
| Malignant epithelial | C0 Is ECs | APOBEC3C | 0.508 | 0.301 | 0.228 | 5.71e-10 | 1.92e-05 |
| Malignant epithelial | C0 Is ECs | TNFRSF21 | 0.507 | 0.394 | 0.317 | 2.18e-11 | 7.34e-07 |
| Malignant epithelial | C0 Is ECs | BTBD2 | 0.506 | 0.243 | 0.182 | 3.12e-08 | 0.00105 |
| Malignant epithelial | C0 Is ECs | HSD3B7 | 0.501 | 0.269 | 0.207 | 2.43e-08 | 0.00082 |
| Malignant epithelial | C0 Is ECs | SUN1 | 0.499 | 0.64 | 0.51 | 2.26e-31 | 7.62e-27 |
| Malignant epithelial | C0 Is ECs | USF2 | 0.498 | 0.381 | 0.319 | 9.14e-10 | 3.08e-05 |
| Malignant epithelial | C0 Is ECs | SNHG8 | 0.496 | 0.9 | 0.816 | 1.73e-49 | 5.819999999999999e-45 |
| Malignant epithelial | C0 Is ECs | TRIB1 | 0.492 | 0.725 | 0.591 | 5.47e-37 | 1.84e-32 |
| Malignant epithelial | C0 Is ECs | P4HA1 | 0.492 | 0.335 | 0.263 | 7.02e-10 | 2.36e-05 |
| Malignant epithelial | C0 Is ECs | COMT | 0.491 | 0.826 | 0.729 | 2.6e-40 | 8.76e-36 |
| Malignant epithelial | C0 Is ECs | TM2D2 | 0.491 | 0.419 | 0.312 | 5.61e-17 | 1.89e-12 |
| Malignant epithelial | C0 Is ECs | RP11-792A8.4 | 0.491 | 0.272 | 0.22 | 1.24e-06 | 0.0417 |
| Malignant epithelial | C0 Is ECs | TNKS1BP1 | 0.489 | 0.248 | 0.187 | 1.58e-08 | 0.000531 |
| Malignant epithelial | C0 Is ECs | CMBL | 0.487 | 0.346 | 0.269 | 8.51e-11 | 2.87e-06 |
| Malignant epithelial | C0 Is ECs | TOB2 | 0.487 | 0.457 | 0.388 | 1.48e-10 | 4.99e-06 |
| Malignant epithelial | C0 Is ECs | DALRD3 | 0.486 | 0.289 | 0.238 | 7.37e-07 | 0.0248 |
| Malignant epithelial | C0 Is ECs | GPRC5A | 0.485 | 0.901 | 0.78 | 2.43e-54 | 8.19e-50 |
| Malignant epithelial | C0 Is ECs | GLIPR1 | 0.485 | 0.48 | 0.327 | 7.240000000000001e-29 | 2.44e-24 |
| Malignant epithelial | C0 Is ECs | MZT2A | 0.484 | 0.915 | 0.797 | 5.65e-75 | 1.9e-70 |
| Malignant epithelial | C0 Is ECs | ITGA3 | 0.484 | 0.476 | 0.359 | 1.06e-19 | 3.56e-15 |
| Malignant epithelial | C0 Is ECs | ERRFI1 | 0.484 | 0.537 | 0.451 | 3.02e-15 | 1.02e-10 |
| Malignant epithelial | C0 Is ECs | B3GNT5 | 0.478 | 0.411 | 0.338 | 4.05e-11 | 1.36e-06 |
| Malignant epithelial | C0 Is ECs | NMI | 0.476 | 0.333 | 0.278 | 4.31e-07 | 0.0145 |
| Malignant epithelial | C0 Is ECs | IFT43 | 0.475 | 0.336 | 0.277 | 3.61e-08 | 0.00122 |
| Malignant epithelial | C0 Is ECs | GSTO2 | 0.473 | 0.506 | 0.4 | 1.96e-17 | 6.61e-13 |
| Malignant epithelial | C0 Is ECs | TSC22D3 | 0.473 | 0.545 | 0.496 | 1.49e-08 | 0.000503 |
| Malignant epithelial | C0 Is ECs | THG1L | 0.472 | 0.275 | 0.22 | 4.04e-07 | 0.0136 |
| Malignant epithelial | C0 Is ECs | DPCD | 0.471 | 0.402 | 0.34 | 2.27e-09 | 7.65e-05 |
| Malignant epithelial | C0 Is ECs | DNTTIP1 | 0.47 | 0.698 | 0.612 | 1.67e-22 | 5.64e-18 |
| Malignant epithelial | C0 Is ECs | OXLD1 | 0.47 | 0.417 | 0.344 | 2.49e-11 | 8.38e-07 |
| Malignant epithelial | C0 Is ECs | TSPO | 0.469 | 0.968 | 0.924 | 2.03e-84 | 6.830000000000001e-80 |
| Malignant epithelial | C0 Is ECs | NGFRAP1 | 0.468 | 0.686 | 0.494 | 7.51e-40 | 2.53e-35 |
| Malignant epithelial | C0 Is ECs | ARHGAP5 | 0.467 | 0.47 | 0.386 | 5.62e-14 | 1.89e-09 |
| Malignant epithelial | C0 Is ECs | LYSMD2 | 0.467 | 0.286 | 0.214 | 4.98e-10 | 1.68e-05 |
| Malignant epithelial | C0 Is ECs | AC090498.1 | 0.466 | 0.897 | 0.834 | 3.149999999999999e-54 | 1.06e-49 |
| Malignant epithelial | C0 Is ECs | ISG15 | 0.466 | 0.918 | 0.816 | 5.92e-38 | 2e-33 |
| Malignant epithelial | C0 Is ECs | GLS | 0.466 | 0.416 | 0.343 | 1.67e-10 | 5.61e-06 |
| Malignant epithelial | C0 Is ECs | USE1 | 0.465 | 0.443 | 0.376 | 2.13e-11 | 7.19e-07 |
| Malignant epithelial | C0 Is ECs | RICTOR | 0.464 | 0.302 | 0.24 | 2.51e-08 | 0.000844 |
| Malignant epithelial | C0 Is ECs | SIPA1 | 0.461 | 0.322 | 0.254 | 1.75e-09 | 5.89e-05 |
| Malignant epithelial | C0 Is ECs | LDLR | 0.46 | 0.562 | 0.463 | 9.48e-18 | 3.19e-13 |
| Malignant epithelial | C0 Is ECs | BLVRB | 0.457 | 0.73 | 0.647 | 3.29e-26 | 1.11e-21 |
| Malignant epithelial | C0 Is ECs | RPL29 | 0.455 | 1.0 | 0.999 | 3.28e-171 | 1.11e-166 |
| Malignant epithelial | C0 Is ECs | XAF1 | 0.453 | 0.288 | 0.219 | 1e-08 | 0.000337 |
| Malignant epithelial | C0 Is ECs | LURAP1L | 0.449 | 0.307 | 0.252 | 9.23e-07 | 0.0311 |
| Malignant epithelial | C0 Is ECs | CASP4 | 0.447 | 0.394 | 0.337 | 3.18e-08 | 0.00107 |
| Malignant epithelial | C0 Is ECs | TMC5 | 0.446 | 0.56 | 0.496 | 4.08e-14 | 1.37e-09 |
| Malignant epithelial | C0 Is ECs | C12orf45 | 0.44 | 0.4 | 0.333 | 1.28e-09 | 4.32e-05 |
| Malignant epithelial | C0 Is ECs | NPC2 | 0.436 | 0.896 | 0.867 | 1.48e-27 | 4.98e-23 |
| Malignant epithelial | C0 Is ECs | ATP5G2 | 0.434 | 0.988 | 0.966 | 4.97e-97 | 1.67e-92 |
| Malignant epithelial | C0 Is ECs | EMG1 | 0.434 | 0.469 | 0.38 | 1.03e-13 | 3.48e-09 |
| Malignant epithelial | C0 Is ECs | STEAP1 | 0.433 | 0.578 | 0.472 | 9.52e-19 | 3.21e-14 |
| Malignant epithelial | C0 Is ECs | COTL1 | 0.432 | 0.558 | 0.473 | 1.73e-16 | 5.82e-12 |
| Malignant epithelial | C0 Is ECs | MAP3K13 | 0.431 | 0.599 | 0.531 | 2.96e-15 | 9.98e-11 |
| Malignant epithelial | C0 Is ECs | PLSCR1 | 0.43 | 0.704 | 0.619 | 2.77e-24 | 9.33e-20 |
| Malignant epithelial | C0 Is ECs | BIK | 0.428 | 0.698 | 0.572 | 2.87e-26 | 9.66e-22 |

| compartment | subcluster | gene | avg_log2FC | pct.1 | pct.2 | p_val | p_val_adj |
|---|---|---|---|---|---|---|---|
| Malignant epithelial | C0 Is ECs | MZT2B | 0.425 | 0.962 | 0.881 | 5.87e-72 | 1.98e-67 |
| Malignant epithelial | C0 Is ECs | HIST1H2BD | 0.425 | 0.344 | 0.272 | 4.56e-09 | 0.000154 |
| Malignant epithelial | C0 Is ECs | MMP7 | 0.423 | 0.52 | 0.367 | 8.21e-25 | 2.77e-20 |
| Malignant epithelial | C0 Is ECs | FSCN1 | 0.423 | 0.478 | 0.327 | 6.680000000000001e-22 | 2.25e-17 |
| Malignant epithelial | C0 Is ECs | SKIL | 0.423 | 0.375 | 0.322 | 9.32e-07 | 0.0314 |
| Malignant epithelial | C0 Is ECs | REEP3 | 0.422 | 0.602 | 0.48 | 8.87e-22 | 2.99e-17 |
| Malignant epithelial | C0 Is ECs | SOX9 | 0.422 | 0.671 | 0.607 | 1.48e-16 | 4.97e-12 |
| Malignant epithelial | C0 Is ECs | ID2 | 0.421 | 0.828 | 0.74 | 2.54e-25 | 8.57e-21 |
| Malignant epithelial | C0 Is ECs | FAM213A | 0.419 | 0.549 | 0.467 | 4.77e-16 | 1.61e-11 |
| Malignant epithelial | C0 Is ECs | RUNX1 | 0.418 | 0.504 | 0.427 | 2.28e-11 | 7.69e-07 |
| Malignant epithelial | C0 Is ECs | TPM2 | 0.414 | 0.908 | 0.7 | 2.25e-51 | 7.570000000000001e-47 |
| Malignant epithelial | C0 Is ECs | TCEAL4 | 0.411 | 0.546 | 0.465 | 6.73e-13 | 2.27e-08 |
| Malignant epithelial | C0 Is ECs | EFNA1 | 0.411 | 0.579 | 0.537 | 8.52e-07 | 0.0287 |
| Malignant epithelial | C0 Is ECs | CDV3 | 0.41 | 0.394 | 0.347 | 4.6e-07 | 0.0155 |
| Malignant epithelial | C0 Is ECs | CYBA | 0.406 | 0.986 | 0.966 | 5.41e-60 | 1.82e-55 |
| Malignant epithelial | C0 Is ECs | TMEM256 | 0.404 | 0.599 | 0.513 | 8.29e-18 | 2.79e-13 |
| Malignant epithelial | C0 Is ECs | PDXK | 0.403 | 0.667 | 0.593 | 4.61e-15 | 1.55e-10 |
| Malignant epithelial | C0 Is ECs | LSM2 | 0.402 | 0.734 | 0.639 | 9.55e-25 | 3.22e-20 |
| Malignant epithelial | C0 Is ECs | MRPL54 | 0.4 | 0.663 | 0.589 | 5.71e-16 | 1.93e-11 |
| Malignant epithelial | C0 Is ECs | C12orf57 | 0.399 | 0.735 | 0.634 | 1.52e-22 | 5.13e-18 |
| Malignant epithelial | C0 Is ECs | LIPH | 0.398 | 0.482 | 0.405 | 1.54e-11 | 5.18e-07 |
| Malignant epithelial | C0 Is ECs | IAH1 | 0.395 | 0.499 | 0.434 | 1.11e-10 | 3.73e-06 |
| Malignant epithelial | C0 Is ECs | SPAG9 | 0.394 | 0.465 | 0.394 | 2.61e-11 | 8.79e-07 |
| Malignant epithelial | C0 Is ECs | ZNF581 | 0.394 | 0.409 | 0.358 | 1.87e-07 | 0.00631 |
| Malignant epithelial | C0 Is ECs | MRPL15 | 0.393 | 0.625 | 0.543 | 9.31e-16 | 3.14e-11 |
| Malignant epithelial | C0 Is ECs | IER2 | 0.392 | 0.973 | 0.892 | 3.93e-29 | 1.32e-24 |
| Malignant epithelial | C0 Is ECs | UBE2L6 | 0.391 | 0.578 | 0.473 | 4.13e-16 | 1.39e-11 |
| Malignant epithelial | C0 Is ECs | SSSCA1 | 0.39 | 0.533 | 0.479 | 5.95e-10 | 2.01e-05 |
| Malignant epithelial | C0 Is ECs | NCOA7 | 0.39 | 0.557 | 0.493 | 4.24e-09 | 0.000143 |
| Malignant epithelial | C0 Is ECs | LDHA | 0.389 | 0.952 | 0.908 | 1.8e-42 | 6.079999999999999e-38 |
| Malignant epithelial | C0 Is ECs | WDR33 | 0.389 | 0.412 | 0.36 | 4.32e-07 | 0.0146 |
| Malignant epithelial | C0 Is ECs | COMTD1 | 0.386 | 0.634 | 0.572 | 2.33e-15 | 7.85e-11 |
| Malignant epithelial | C0 Is ECs | RPS24 | 0.384 | 0.999 | 0.997 | 1.18e-101 | 3.96e-97 |
| Malignant epithelial | C0 Is ECs | ELF3 | 0.384 | 0.996 | 0.928 | 2.58e-45 | 8.7e-41 |
| Malignant epithelial | C0 Is ECs | CD74 | 0.384 | 0.955 | 0.868 | 6.43e-43 | 2.17e-38 |
| Malignant epithelial | C0 Is ECs | TAPBP | 0.384 | 0.715 | 0.631 | 1.42e-20 | 4.78e-16 |
| Malignant epithelial | C0 Is ECs | CDCP1 | 0.384 | 0.397 | 0.327 | 3.54e-09 | 0.000119 |
| Malignant epithelial | C0 Is ECs | MX1 | 0.384 | 0.403 | 0.335 | 1.69e-08 | 0.000571 |
| Malignant epithelial | C0 Is ECs | PPP1R15A | 0.378 | 0.975 | 0.864 | 6.27e-48 | 2.11e-43 |
| Malignant epithelial | C0 Is ECs | IL18 | 0.378 | 0.461 | 0.357 | 5.77e-15 | 1.95e-10 |
| Malignant epithelial | C0 Is ECs | CRB3 | 0.377 | 0.512 | 0.467 | 1.11e-07 | 0.00374 |
| Malignant epithelial | C0 Is ECs | ARL6IP6 | 0.377 | 0.33 | 0.272 | 3.4e-07 | 0.0114 |
| Malignant epithelial | C0 Is ECs | ADAM9 | 0.376 | 0.44 | 0.337 | 1.63e-15 | 5.48e-11 |
| Malignant epithelial | C0 Is ECs | SEZ6L2 | 0.376 | 0.274 | 0.215 | 3.37e-07 | 0.0113 |
| Malignant epithelial | C0 Is ECs | C19orf53 | 0.373 | 0.943 | 0.892 | 1.3e-45 | 4.37e-41 |
| Malignant epithelial | C0 Is ECs | NOP10 | 0.373 | 0.915 | 0.847 | 3.4e-42 | 1.15e-37 |
| Malignant epithelial | C0 Is ECs | NET1 | 0.371 | 0.636 | 0.539 | 3.95e-19 | 1.33e-14 |
| Malignant epithelial | C0 Is ECs | LINC00116 | 0.371 | 0.544 | 0.479 | 4.42e-11 | 1.49e-06 |
| Malignant epithelial | C0 Is ECs | S100A6 | 0.37 | 1.0 | 1.0 | 1.06e-90 | 3.57e-86 |
| Malignant epithelial | C0 Is ECs | LAMC2 | 0.369 | 0.45 | 0.334 | 2.51e-18 | 8.46e-14 |
| Malignant epithelial | C0 Is ECs | TIMM9 | 0.368 | 0.52 | 0.447 | 4.69e-11 | 1.58e-06 |
| Malignant epithelial | C0 Is ECs | NDUFAF3 | 0.366 | 0.769 | 0.685 | 9.44e-22 | 3.18e-17 |
| Malignant epithelial | C0 Is ECs | SEC11A | 0.365 | 0.817 | 0.775 | 1.23e-29 | 4.13e-25 |
| Malignant epithelial | C0 Is ECs | AGPAT2 | 0.365 | 0.825 | 0.771 | 1.82e-21 | 6.14e-17 |
| Malignant epithelial | C0 Is ECs | NT5C | 0.363 | 0.645 | 0.594 | 1.76e-11 | 5.93e-07 |
| Malignant epithelial | C0 Is ECs | TRA2A | 0.363 | 0.626 | 0.572 | 1.4e-08 | 0.000473 |
| Malignant epithelial | C0 Is ECs | OXA1L | 0.363 | 0.488 | 0.44 | 6.44e-08 | 0.00217 |
| Malignant epithelial | C0 Is ECs | RPL36 | 0.36 | 1.0 | 1.0 | 1.75e-123 | 5.9e-119 |
| Malignant epithelial | C0 Is ECs | PTMA | 0.36 | 1.0 | 0.998 | 2.32e-82 | 7.8e-78 |
| Malignant epithelial | C0 Is ECs | AHNAK | 0.357 | 0.63 | 0.527 | 2.77e-19 | 9.32e-15 |
| Malignant epithelial | C0 Is ECs | NAP1L1 | 0.355 | 0.92 | 0.879 | 5.25e-39 | 1.77e-34 |
| Malignant epithelial | C0 Is ECs | RND3 | 0.355 | 0.721 | 0.589 | 1.68e-29 | 5.67e-25 |
| Malignant epithelial | C0 Is ECs | ARHGDIB | 0.354 | 0.527 | 0.453 | 9.27e-10 | 3.12e-05 |
| Malignant epithelial | C0 Is ECs | SERPINA1 | 0.351 | 0.543 | 0.371 | 1.77e-32 | 5.98e-28 |

| compartment | subcluster | gene | avg_log2FC | pct.1 | pct.2 | p_val | p_val_adj |
|---|---|---|---|---|---|---|---|
| Malignant epithelial | C0 Is ECs | MRPS16 | 0.351 | 0.733 | 0.646 | 3.44e-19 | 1.16e-14 |
| Malignant epithelial | C0 Is ECs | HLA-DMA | 0.35 | 0.546 | 0.44 | 3.47e-14 | 1.17e-09 |
| Malignant epithelial | C0 Is ECs | OAS1 | 0.35 | 0.51 | 0.457 | 4.53e-08 | 0.00153 |
| Malignant epithelial | C0 Is ECs | LINC00493 | 0.349 | 0.91 | 0.835 | 5.2e-38 | 1.75e-33 |
| Malignant epithelial | C0 Is ECs | EXOSC7 | 0.348 | 0.384 | 0.323 | 2.45e-07 | 0.00824 |
| Malignant epithelial | C0 Is ECs | RPS15 | 0.347 | 0.999 | 0.999 | 1.66e-130 | 5.6e-126 |
| Malignant epithelial | C0 Is ECs | SPATS2L | 0.347 | 0.697 | 0.623 | 1.27e-15 | 4.26e-11 |
| Malignant epithelial | C0 Is ECs | RNF213 | 0.347 | 0.523 | 0.453 | 1.63e-10 | 5.48e-06 |
| Malignant epithelial | C0 Is ECs | GADD45GIP1 | 0.346 | 0.869 | 0.79 | 4.02e-29 | 1.36e-24 |
| Malignant epithelial | C0 Is ECs | CISD1 | 0.345 | 0.578 | 0.504 | 2.49e-11 | 8.39e-07 |
| Malignant epithelial | C0 Is ECs | RHOG | 0.345 | 0.485 | 0.432 | 2.56e-08 | 0.000864 |
| Malignant epithelial | C0 Is ECs | ESRP1 | 0.345 | 0.457 | 0.401 | 1.84e-07 | 0.0062 |
| Malignant epithelial | C0 Is ECs | BIRC2 | 0.345 | 0.361 | 0.299 | 2.73e-07 | 0.00921 |
| Malignant epithelial | C0 Is ECs | DCBLD2 | 0.341 | 0.539 | 0.393 | 6.649999999999999e-23 | 2.24e-18 |
| Malignant epithelial | C0 Is ECs | GNAI2 | 0.339 | 0.483 | 0.388 | 5.36e-14 | 1.81e-09 |
| Malignant epithelial | C0 Is ECs | DUSP5 | 0.338 | 0.461 | 0.273 | 1.88e-42 | 6.339999999999999e-38 |
| Malignant epithelial | C0 Is ECs | CDKN2A | 0.335 | 0.828 | 0.73 | 2.74e-23 | 9.22e-19 |
| Malignant epithelial | C0 Is ECs | S100A13 | 0.334 | 0.831 | 0.746 | 6.43e-25 | 2.17e-20 |
| Malignant epithelial | C0 Is ECs | RPL36AL | 0.332 | 0.987 | 0.954 | 3.41e-60 | 1.15e-55 |
| Malignant epithelial | C0 Is ECs | IFI16 | 0.332 | 0.402 | 0.321 | 3.36e-10 | 1.13e-05 |
| Malignant epithelial | C0 Is ECs | ADAR | 0.331 | 0.44 | 0.377 | 4.4e-08 | 0.00148 |
| Malignant epithelial | C0 Is ECs | SNU13 | 0.33 | 0.911 | 0.842 | 1.14e-28 | 3.85e-24 |
| Malignant epithelial | C0 Is ECs | CTNNB1 | 0.33 | 0.68 | 0.619 | 2.05e-09 | 6.91e-05 |
| Malignant epithelial | C0 Is ECs | ATF3 | 0.329 | 0.891 | 0.736 | 1.48e-34 | 4.97e-30 |
| Malignant epithelial | C0 Is ECs | HOOK2 | 0.329 | 0.588 | 0.533 | 2.22e-10 | 7.48e-06 |
| Malignant epithelial | C0 Is ECs | STAT1 | 0.329 | 0.499 | 0.444 | 3.57e-07 | 0.012 |
| Malignant epithelial | C0 Is ECs | TOMM22 | 0.325 | 0.728 | 0.664 | 2.98e-18 | 1e-13 |
| Malignant epithelial | C0 Is ECs | UQCC2 | 0.325 | 0.595 | 0.534 | 1.04e-10 | 3.52e-06 |
| Malignant epithelial | C0 Is ECs | HIGD2A | 0.324 | 0.89 | 0.833 | 9.31e-30 | 3.14e-25 |
| Malignant epithelial | C0 Is ECs | C19orf70 | 0.323 | 0.819 | 0.745 | 1.4e-23 | 4.71e-19 |
| Malignant epithelial | C0 Is ECs | FAM162A | 0.323 | 0.669 | 0.614 | 4.05e-11 | 1.37e-06 |
| Malignant epithelial | C0 Is ECs | SNHG7 | 0.322 | 0.819 | 0.735 | 4.12e-24 | 1.39e-19 |
| Malignant epithelial | C0 Is ECs | RPL26L1 | 0.319 | 0.604 | 0.544 | 1.83e-11 | 6.16e-07 |
| Malignant epithelial | C0 Is ECs | RPL23 | 0.312 | 0.997 | 0.987 | 2.51e-76 | 8.46e-72 |
| Malignant epithelial | C0 Is ECs | ZDHHC3 | 0.311 | 0.549 | 0.488 | 7.39e-10 | 2.49e-05 |
| Malignant epithelial | C0 Is ECs | COX19 | 0.311 | 0.552 | 0.506 | 3.58e-08 | 0.00121 |
| Malignant epithelial | C0 Is ECs | CCDC85B | 0.31 | 0.786 | 0.699 | 2.71e-19 | 9.14e-15 |
| Malignant epithelial | C0 Is ECs | GTF3C6 | 0.31 | 0.68 | 0.628 | 3.25e-12 | 1.1e-07 |
| Malignant epithelial | C0 Is ECs | RRP7A | 0.31 | 0.546 | 0.49 | 4.73e-08 | 0.00159 |
| Malignant epithelial | C0 Is ECs | APOE | 0.308 | 0.553 | 0.471 | 8.76e-12 | 2.95e-07 |
| Malignant epithelial | C0 Is ECs | RNF149 | 0.308 | 0.531 | 0.475 | 1.79e-09 | 6.02e-05 |
| Malignant epithelial | C0 Is ECs | ZFAND1 | 0.307 | 0.598 | 0.547 | 2.81e-10 | 9.45e-06 |
| Malignant epithelial | C0 Is ECs | VAMP8 | 0.306 | 0.907 | 0.873 | 3.2e-25 | 1.08e-20 |
| Malignant epithelial | C0 Is ECs | CCDC59 | 0.302 | 0.579 | 0.539 | 5.44e-08 | 0.00183 |
| Malignant epithelial | C0 Is ECs | PYCARD | 0.3 | 0.824 | 0.735 | 5.28e-21 | 1.78e-16 |
| Malignant epithelial | C0 Is ECs | NDUFB1 | 0.296 | 0.914 | 0.852 | 2.33e-23 | 7.85e-19 |
| Malignant epithelial | C0 Is ECs | RPS29 | 0.295 | 1.0 | 0.999 | 6.53e-99 | 2.2e-94 |
| Malignant epithelial | C0 Is ECs | RPL38 | 0.294 | 0.997 | 0.997 | 4.78e-75 | 1.61e-70 |
| Malignant epithelial | C0 Is ECs | MYL9 | 0.294 | 0.3 | 0.217 | 3.7e-11 | 1.25e-06 |
| Malignant epithelial | C0 Is ECs | NT5E | 0.293 | 0.509 | 0.436 | 4.08e-09 | 0.000137 |
| Malignant epithelial | C0 Is ECs | IRF2BP2 | 0.293 | 0.632 | 0.596 | 5.4e-09 | 0.000182 |
| Malignant epithelial | C0 Is ECs | TIMP2 | 0.29 | 0.338 | 0.223 | 2.2e-18 | 7.4e-14 |
| Malignant epithelial | C0 Is ECs | FKBP8 | 0.289 | 0.8 | 0.769 | 4.26e-15 | 1.44e-10 |
| Malignant epithelial | C0 Is ECs | EIF1 | 0.288 | 0.999 | 0.991 | 5.470000000000001e-56 | 1.84e-51 |
| Malignant epithelial | C0 Is ECs | MGST1 | 0.287 | 0.781 | 0.685 | 2.19e-17 | 7.37e-13 |
| Malignant epithelial | C0 Is ECs | MALL | 0.287 | 0.536 | 0.435 | 1.34e-13 | 4.51e-09 |
| Malignant epithelial | C0 Is ECs | SRSF7 | 0.287 | 0.796 | 0.743 | 1.16e-08 | 0.000391 |
| Malignant epithelial | C0 Is ECs | SDC4 | 0.285 | 0.764 | 0.706 | 5.11e-13 | 1.72e-08 |
| Malignant epithelial | C0 Is ECs | YBX3 | 0.284 | 0.797 | 0.73 | 8.2e-16 | 2.76e-11 |
| Malignant epithelial | C0 Is ECs | CTSH | 0.281 | 0.537 | 0.47 | 4.64e-09 | 0.000156 |
| Malignant epithelial | C0 Is ECs | IFI27L2 | 0.278 | 0.728 | 0.649 | 4.79e-15 | 1.61e-10 |
| Malignant epithelial | C0 Is ECs | CLDN4 | 0.277 | 1.0 | 0.951 | 3.58e-46 | 1.21e-41 |
| Malignant epithelial | C0 Is ECs | PRDX6 | 0.276 | 0.846 | 0.798 | 7.98e-20 | 2.69e-15 |
| Malignant epithelial | C0 Is ECs | RPL18A | 0.275 | 1.0 | 0.996 | 4.37e-70 | 1.47e-65 |

| compartment | subcluster | gene | avg_log2FC | pct.1 | pct.2 | p_val | p_val_adj |
|---|---|---|---|---|---|---|---|
| Malignant epithelial | C0 Is ECs | RPL34 | 0.272 | 0.999 | 0.998 | 1.06e-72 | 3.58e-68 |
| Malignant epithelial | C0 Is ECs | CHCHD5 | 0.271 | 0.639 | 0.573 | 1.68e-11 | 5.68e-07 |
| Malignant epithelial | C0 Is ECs | TOMM20 | 0.265 | 0.808 | 0.76 | 1.68e-17 | 5.66e-13 |
| Malignant epithelial | C0 Is ECs | RPL41 | 0.263 | 1.0 | 1.0 | 1.03e-75 | 3.46e-71 |
| Malignant epithelial | C0 Is ECs | KRTCAP2 | 0.263 | 0.866 | 0.807 | 1.19e-15 | 4.01e-11 |
| Malignant epithelial | C0 Is ECs | HIST1H1C | 0.262 | 0.495 | 0.436 | 1.99e-07 | 0.0067 |
| Malignant epithelial | C0 Is ECs | RPL6 | 0.261 | 0.998 | 0.991 | 9.54e-57 | 3.21e-52 |
| Malignant epithelial | C0 Is ECs | RPL35 | 0.258 | 0.999 | 0.999 | 8.9e-58 | 3e-53 |
| Malignant epithelial | C0 Is ECs | TATDN1 | 0.258 | 0.62 | 0.575 | 1.36e-08 | 0.000457 |
| Malignant epithelial | C0 Is ECs | RPL4 | 0.254 | 0.984 | 0.969 | 3.57e-35 | 1.2e-30 |
| Malignant epithelial | C0 Is ECs | RPL37 | 0.253 | 0.999 | 0.999 | 1.6e-74 | 5.39e-70 |
| Malignant epithelial | C0 Is ECs | NBEAL1 | 0.253 | 0.912 | 0.876 | 2.77e-24 | 9.33e-20 |
| Malignant epithelial | C0 Is ECs | NFAT5 | 0.253 | 0.441 | 0.374 | 1.07e-08 | 0.000362 |
| Malignant epithelial | C0 Is ECs | FOS | 0.251 | 0.99 | 0.921 | 4.91e-24 | 1.65e-19 |
| Malignant epithelial | C0 Is ECs | TAF1D | 0.251 | 0.774 | 0.749 | 9.75e-10 | 3.29e-05 |
| Malignant epithelial | C0 Is ECs | NDUFS4 | 0.251 | 0.648 | 0.619 | 5.61e-08 | 0.00189 |
| Malignant epithelial | C0 Is ECs | NACA | 0.25 | 0.994 | 0.992 | 1.07e-60 | 3.6e-56 |
| Malignant epithelial | C1 Das ECs | MYBPC1 | 4.246 | 0.235 | 0.017 | 1.02e-156 | 3.42e-152 |
| Malignant epithelial | C1 Das ECs | OLFM4 | 4.107 | 0.101 | 0.017 | 1.53e-43 | 5.17e-39 |
| Malignant epithelial | C1 Das ECs | LEFTY1 | 4.028 | 0.466 | 0.055 | 3.89e-279 | 1.31e-274 |
| Malignant epithelial | C1 Das ECs | GABRP | 3.849 | 0.124 | 0.015 | 6.36e-66 | 2.14e-61 |
| Malignant epithelial | C1 Das ECs | PIGR | 3.598 | 0.667 | 0.096 | 0.0 | 0.0 |
| Malignant epithelial | C1 Das ECs | DMRT2 | 3.582 | 0.112 | 0.011 | 4.08e-64 | 1.38e-59 |
| Malignant epithelial | C1 Das ECs | AQP1 | 3.521 | 0.476 | 0.053 | 3.06e-289 | 1.03e-284 |
| Malignant epithelial | C1 Das ECs | MS4A8 | 3.505 | 0.176 | 0.016 | 3.49e-107 | 1.18e-102 |
| Malignant epithelial | C1 Das ECs | ARHGAP6 | 3.401 | 0.307 | 0.037 | 6.42e-171 | 2.16e-166 |
| Malignant epithelial | C1 Das ECs | CLDN2 | 3.398 | 0.143 | 0.014 | 1.18e-83 | 3.96e-79 |
| Malignant epithelial | C1 Das ECs | NOX1 | 3.151 | 0.444 | 0.055 | 4.82e-252 | 1.62e-247 |
| Malignant epithelial | C1 Das ECs | AC016735.2 | 3.137 | 0.185 | 0.025 | 1.4e-91 | 4.7e-87 |
| Malignant epithelial | C1 Das ECs | BLNK | 3.101 | 0.128 | 0.018 | 2.18e-60 | 7.340000000000001e-56 |
| Malignant epithelial | C1 Das ECs | FEZF1-AS1 | 3.09 | 0.115 | 0.012 | 8.45e-65 | 2.85e-60 |
| Malignant epithelial | C1 Das ECs | BCL2L14 | 3.029 | 0.277 | 0.045 | 3.74e-128 | 1.26e-123 |
| Malignant epithelial | C1 Das ECs | PLCB4 | 3.013 | 0.221 | 0.034 | 7.470000000000001e-103 | 2.52e-98 |
| Malignant epithelial | C1 Das ECs | IQGAP2 | 2.988 | 0.215 | 0.032 | 3.83e-101 | 1.29e-96 |
| Malignant epithelial | C1 Das ECs | MUC5B | 2.943 | 0.315 | 0.049 | 3.04e-149 | 1.02e-144 |
| Malignant epithelial | C1 Das ECs | SHD | 2.85 | 0.178 | 0.031 | 1.09e-74 | 3.66e-70 |
| Malignant epithelial | C1 Das ECs | ST6GALNAC1 | 2.814 | 0.218 | 0.037 | 1.14e-93 | 3.85e-89 |
| Malignant epithelial | C1 Das ECs | CFTR | 2.804 | 0.314 | 0.051 | 2.13e-144 | 7.189999999999999e-140 |
| Malignant epithelial | C1 Das ECs | PLEKHS1 | 2.791 | 0.183 | 0.032 | 1.62e-76 | 5.47e-72 |
| Malignant epithelial | C1 Das ECs | CTTNBP2 | 2.787 | 0.152 | 0.028 | 7.23e-60 | 2.43e-55 |
| Malignant epithelial | C1 Das ECs | PKDCC | 2.753 | 0.175 | 0.033 | 4.53e-69 | 1.53e-64 |
| Malignant epithelial | C1 Das ECs | DEGS2 | 2.725 | 0.385 | 0.062 | 6.21e-182 | 2.09e-177 |
| Malignant epithelial | C1 Das ECs | SSPN | 2.703 | 0.101 | 0.018 | 1.32e-40 | 4.43e-36 |
| Malignant epithelial | C1 Das ECs | PRLR | 2.697 | 0.113 | 0.019 | 2.99e-47 | 1.01e-42 |
| Malignant epithelial | C1 Das ECs | PPP1R1B | 2.658 | 0.701 | 0.168 | 2.93e-313 | 9.87e-309 |
| Malignant epithelial | C1 Das ECs | BCL11A | 2.63 | 0.2 | 0.039 | 3.14e-77 | 1.06e-72 |
| Malignant epithelial | C1 Das ECs | FAM3D | 2.62 | 0.641 | 0.128 | 5.83e-295 | 1.97e-290 |
| Malignant epithelial | C1 Das ECs | GPX2 | 2.582 | 0.957 | 0.368 | 0.0 | 0.0 |
| Malignant epithelial | C1 Das ECs | SLC5A1 | 2.549 | 0.158 | 0.036 | 3.63e-52 | 1.22e-47 |
| Malignant epithelial | C1 Das ECs | TBXAS1 | 2.534 | 0.221 | 0.043 | 5.06e-86 | 1.71e-81 |
| Malignant epithelial | C1 Das ECs | MLLT3 | 2.521 | 0.326 | 0.065 | 2.11e-130 | 7.129999999999999e-126 |
| Malignant epithelial | C1 Das ECs | C1orf21 | 2.502 | 0.649 | 0.155 | 4.02e-270 | 1.36e-265 |
| Malignant epithelial | C1 Das ECs | KIAA1324 | 2.492 | 0.107 | 0.021 | 2.26e-40 | 7.61e-36 |
| Malignant epithelial | C1 Das ECs | WNK2 | 2.485 | 0.171 | 0.035 | 2.98e-62 | 1e-57 |
| Malignant epithelial | C1 Das ECs | PBX1 | 2.475 | 0.193 | 0.043 | 5.21e-66 | 1.76e-61 |
| Malignant epithelial | C1 Das ECs | LINC00926 | 2.459 | 0.25 | 0.058 | 6.83e-85 | 2.3e-80 |
| Malignant epithelial | C1 Das ECs | PLA2G10 | 2.44 | 0.27 | 0.064 | 2.23e-90 | 7.51e-86 |
| Malignant epithelial | C1 Das ECs | RP11-150O12.3 | 2.432 | 0.123 | 0.027 | 8.860000000000001e-43 | 2.99e-38 |
| Malignant epithelial | C1 Das ECs | TFF3 | 2.383 | 0.975 | 0.528 | 0.0 | 0.0 |
| Malignant epithelial | C1 Das ECs | GJB1 | 2.373 | 0.413 | 0.105 | 4.849999999999999e-140 | 1.63e-135 |
| Malignant epithelial | C1 Das ECs | KIF12 | 2.364 | 0.187 | 0.048 | 2.17e-55 | 7.33e-51 |
| Malignant epithelial | C1 Das ECs | ST6GAL1 | 2.352 | 0.185 | 0.046 | 7.39e-57 | 2.49e-52 |
| Malignant epithelial | C1 Das ECs | LINC01207 | 2.339 | 0.306 | 0.053 | 1.2e-132 | 4.05e-128 |
| Malignant epithelial | C1 Das ECs | C10orf99 | 2.338 | 0.118 | 0.025 | 1.13e-41 | 3.82e-37 |

| compartment | subcluster | gene | avg_log2FC | pct.1 | pct.2 | p_val | p_val_adj |
|---|---|---|---|---|---|---|---|
| Malignant epithelial | C1 Das ECs | ANXA13 | 2.289 | 0.273 | 0.054 | 1.88e-105 | 6.33e-101 |
| Malignant epithelial | C1 Das ECs | SORBS2 | 2.287 | 0.156 | 0.036 | 5.68e-51 | 1.91e-46 |
| Malignant epithelial | C1 Das ECs | HNF1B | 2.28 | 0.115 | 0.026 | 1.33e-38 | 4.47e-34 |
| Malignant epithelial | C1 Das ECs | BTNL8 | 2.27 | 0.113 | 0.02 | 5.359999999999999e-45 | 1.81e-40 |
| Malignant epithelial | C1 Das ECs | PLCE1 | 2.269 | 0.102 | 0.024 | 3.86e-33 | 1.3e-28 |
| Malignant epithelial | C1 Das ECs | CREB3L1 | 2.232 | 0.237 | 0.053 | 2.02e-81 | 6.82e-77 |
| Malignant epithelial | C1 Das ECs | AC005550.3 | 2.217 | 0.158 | 0.036 | 2.93e-52 | 9.879999999999999e-48 |
| Malignant epithelial | C1 Das ECs | FBP1 | 2.208 | 0.42 | 0.11 | 6.82e-140 | 2.3e-135 |
| Malignant epithelial | C1 Das ECs | ARFGEF3 | 2.206 | 0.101 | 0.023 | 2.11e-32 | 7.12e-28 |
| Malignant epithelial | C1 Das ECs | BHLHE41 | 2.203 | 0.151 | 0.03 | 2.27e-55 | 7.66e-51 |
| Malignant epithelial | C1 Das ECs | SCIN | 2.201 | 0.141 | 0.025 | 1.37e-57 | 4.63e-53 |
| Malignant epithelial | C1 Das ECs | NDRG2 | 2.19 | 0.275 | 0.07 | 1.31e-84 | 4.42e-80 |
| Malignant epithelial | C1 Das ECs | PIP5K1B | 2.189 | 0.207 | 0.046 | 4.57e-70 | 1.54e-65 |
| Malignant epithelial | C1 Das ECs | C9orf152 | 2.183 | 0.313 | 0.087 | 1.63e-91 | 5.49e-87 |
| Malignant epithelial | C1 Das ECs | DEPTOR | 2.156 | 0.197 | 0.05 | 5.74e-60 | 1.94e-55 |
| Malignant epithelial | C1 Das ECs | REPS2 | 2.155 | 0.102 | 0.026 | 6.37e-30 | 2.15e-25 |
| Malignant epithelial | C1 Das ECs | CDX1 | 2.134 | 0.552 | 0.153 | 2.42e-183 | 8.15e-179 |
| Malignant epithelial | C1 Das ECs | MARCH3 | 2.133 | 0.287 | 0.069 | 1.87e-94 | 6.29e-90 |
| Malignant epithelial | C1 Das ECs | AGR3 | 2.13 | 0.274 | 0.059 | 3.13e-97 | 1.05e-92 |
| Malignant epithelial | C1 Das ECs | DDC | 2.126 | 0.213 | 0.059 | 3.66e-59 | 1.23e-54 |
| Malignant epithelial | C1 Das ECs | MLXIPL | 2.12 | 0.231 | 0.065 | 5.47e-63 | 1.84e-58 |
| Malignant epithelial | C1 Das ECs | PBLD | 2.106 | 0.109 | 0.032 | 1.8e-27 | 6.079999999999999e-23 |
| Malignant epithelial | C1 Das ECs | NFIA | 2.095 | 0.22 | 0.059 | 1.73e-62 | 5.81e-58 |
| Malignant epithelial | C1 Das ECs | MOB3B | 2.095 | 0.126 | 0.029 | 3.63e-40 | 1.22e-35 |
| Malignant epithelial | C1 Das ECs | SLCO4A1-AS1 | 2.085 | 0.244 | 0.092 | 6.62e-46 | 2.23e-41 |
| Malignant epithelial | C1 Das ECs | GPR160 | 2.083 | 0.54 | 0.143 | 8.93e-183 | 3.01e-178 |
| Malignant epithelial | C1 Das ECs | CCND2 | 2.067 | 0.232 | 0.063 | 2.81e-65 | 9.479999999999999e-61 |
| Malignant epithelial | C1 Das ECs | IGHM | 2.066 | 0.151 | 0.034 | 5.71e-50 | 1.92e-45 |
| Malignant epithelial | C1 Das ECs | SELENBP1 | 2.061 | 0.263 | 0.085 | 1.44e-62 | 4.84e-58 |
| Malignant epithelial | C1 Das ECs | SLC40A1 | 2.054 | 0.517 | 0.101 | 2.8e-220 | 9.44e-216 |
| Malignant epithelial | C1 Das ECs | GCNT1 | 2.032 | 0.148 | 0.036 | 6.28e-45 | 2.12e-40 |
| Malignant epithelial | C1 Das ECs | GGT6 | 2.023 | 0.123 | 0.035 | 2.7e-32 | 9.09e-28 |
| Malignant epithelial | C1 Das ECs | RP11-295M3.4 | 2.01 | 0.155 | 0.04 | 2.24e-44 | 7.53e-40 |
| Malignant epithelial | C1 Das ECs | LINC00675 | 2.005 | 0.219 | 0.051 | 9.52e-72 | 3.21e-67 |
| Malignant epithelial | C1 Das ECs | PROM1 | 2.001 | 0.132 | 0.034 | 1.7e-38 | 5.73e-34 |
| Malignant epithelial | C1 Das ECs | FOXA1 | 2.0 | 0.109 | 0.029 | 6.18e-30 | 2.08e-25 |
| Malignant epithelial | C1 Das ECs | TSPAN8 | 1.993 | 0.957 | 0.456 | 0.0 | 0.0 |
| Malignant epithelial | C1 Das ECs | IYD | 1.963 | 0.117 | 0.033 | 1.89e-30 | 6.36e-26 |
| Malignant epithelial | C1 Das ECs | SATB2 | 1.952 | 0.152 | 0.044 | 3.61e-38 | 1.22e-33 |
| Malignant epithelial | C1 Das ECs | CA12 | 1.95 | 0.101 | 0.027 | 5.249999999999999e-28 | 1.77e-23 |
| Malignant epithelial | C1 Das ECs | GPC4 | 1.947 | 0.185 | 0.046 | 4.460000000000001e-56 | 1.5e-51 |
| Malignant epithelial | C1 Das ECs | DNAJC12 | 1.929 | 0.137 | 0.038 | 1.86e-36 | 6.26e-32 |
| Malignant epithelial | C1 Das ECs | SMPD3 | 1.928 | 0.13 | 0.041 | 3.27e-30 | 1.1e-25 |
| Malignant epithelial | C1 Das ECs | SLC12A2 | 1.921 | 0.643 | 0.265 | 2.68e-158 | 9.02e-154 |
| Malignant epithelial | C1 Das ECs | ALDH1B1 | 1.92 | 0.281 | 0.103 | 7.710000000000001e-58 | 2.6e-53 |
| Malignant epithelial | C1 Das ECs | ZG16B | 1.91 | 0.39 | 0.107 | 1.49e-118 | 5.02e-114 |
| Malignant epithelial | C1 Das ECs | FGGY | 1.89 | 0.181 | 0.066 | 2.17e-34 | 7.3e-30 |
| Malignant epithelial | C1 Das ECs | CASP1 | 1.888 | 0.301 | 0.101 | 1.46e-68 | 4.930000000000001e-64 |
| Malignant epithelial | C1 Das ECs | PRR15L | 1.88 | 0.562 | 0.194 | 9.5e-151 | 3.2e-146 |
| Malignant epithelial | C1 Das ECs | AGR2 | 1.876 | 0.935 | 0.465 | 2.07e-320 | 6.99e-316 |
| Malignant epithelial | C1 Das ECs | AGFG2 | 1.864 | 0.174 | 0.051 | 1.25e-43 | 4.21e-39 |
| Malignant epithelial | C1 Das ECs | LGALS4 | 1.863 | 0.939 | 0.376 | 7.25e-313 | 2.44e-308 |
| Malignant epithelial | C1 Das ECs | LINC01133 | 1.848 | 0.28 | 0.07 | 1.04e-86 | 3.5e-82 |
| Malignant epithelial | C1 Das ECs | PRKACB | 1.847 | 0.135 | 0.042 | 4.42e-31 | 1.49e-26 |
| Malignant epithelial | C1 Das ECs | CCL15 | 1.845 | 0.299 | 0.093 | 1.14e-75 | 3.83e-71 |
| Malignant epithelial | C1 Das ECs | GIPC2 | 1.844 | 0.132 | 0.04 | 3.98e-32 | 1.34e-27 |
| Malignant epithelial | C1 Das ECs | ANG | 1.839 | 0.525 | 0.164 | 6.959999999999999e-146 | 2.34e-141 |
| Malignant epithelial | C1 Das ECs | ATP10B | 1.827 | 0.389 | 0.117 | 2.82e-105 | 9.5e-101 |
| Malignant epithelial | C1 Das ECs | MCF2L-AS1 | 1.819 | 0.108 | 0.037 | 4.11e-22 | 1.39e-17 |
| Malignant epithelial | C1 Das ECs | ACSM3 | 1.815 | 0.132 | 0.048 | 3.27e-25 | 1.1e-20 |
| Malignant epithelial | C1 Das ECs | SNHG18 | 1.809 | 0.316 | 0.105 | 4.17e-73 | 1.41e-68 |
| Malignant epithelial | C1 Das ECs | GUCY2C | 1.807 | 0.115 | 0.034 | 2.23e-28 | 7.52e-24 |
| Malignant epithelial | C1 Das ECs | SEMA4A | 1.806 | 0.112 | 0.039 | 3.68e-22 | 1.24e-17 |
| Malignant epithelial | C1 Das ECs | MIB2 | 1.798 | 0.369 | 0.136 | 2.78e-79 | 9.360000000000001e-75 |

| compartment | subcluster | gene | avg_log2FC | pct.1 | pct.2 | p_val | p_val_adj |
|---|---|---|---|---|---|---|---|
| Malignant epithelial | C1 Das ECs | LRIG1 | 1.794 | 0.122 | 0.044 | 3.03e-23 | 1.02e-18 |
| Malignant epithelial | C1 Das ECs | LGR4 | 1.775 | 0.167 | 0.057 | 3.51e-34 | 1.18e-29 |
| Malignant epithelial | C1 Das ECs | KCNE3 | 1.773 | 0.188 | 0.066 | 1.1e-37 | 3.69e-33 |
| Malignant epithelial | C1 Das ECs | A1CF | 1.76 | 0.162 | 0.057 | 1.36e-32 | 4.58e-28 |
| Malignant epithelial | C1 Das ECs | TCEA3 | 1.758 | 0.214 | 0.083 | 5.72e-38 | 1.93e-33 |
| Malignant epithelial | C1 Das ECs | DENND2A | 1.754 | 0.102 | 0.027 | 7.96e-28 | 2.68e-23 |
| Malignant epithelial | C1 Das ECs | FAM221A | 1.747 | 0.165 | 0.06 | 4.06e-32 | 1.37e-27 |
| Malignant epithelial | C1 Das ECs | LINC01315 | 1.743 | 0.236 | 0.087 | 2.49e-45 | 8.39e-41 |
| Malignant epithelial | C1 Das ECs | NOXO1 | 1.74 | 0.457 | 0.17 | 5.44e-102 | 1.83e-97 |
| Malignant epithelial | C1 Das ECs | RNASE4 | 1.739 | 0.185 | 0.054 | 6.550000000000001e-47 | 2.21e-42 |
| Malignant epithelial | C1 Das ECs | PCK1 | 1.732 | 0.231 | 0.066 | 5.65e-61 | 1.9e-56 |
| Malignant epithelial | C1 Das ECs | PIK3R1 | 1.726 | 0.207 | 0.075 | 1.85e-40 | 6.23e-36 |
| Malignant epithelial | C1 Das ECs | CTB-55O6.8 | 1.722 | 0.111 | 0.043 | 4.65e-19 | 1.57e-14 |
| Malignant epithelial | C1 Das ECs | FGFR4 | 1.718 | 0.252 | 0.091 | 2.85e-50 | 9.609999999999999e-46 |
| Malignant epithelial | C1 Das ECs | ETV4 | 1.711 | 0.402 | 0.156 | 2.12e-82 | 7.15e-78 |
| Malignant epithelial | C1 Das ECs | HEPH | 1.707 | 0.24 | 0.071 | 7.74e-61 | 2.61e-56 |
| Malignant epithelial | C1 Das ECs | TRIM2 | 1.701 | 0.523 | 0.209 | 3e-111 | 1.01e-106 |
| Malignant epithelial | C1 Das ECs | PID1 | 1.693 | 0.122 | 0.047 | 1.41e-20 | 4.74e-16 |
| Malignant epithelial | C1 Das ECs | UBD | 1.692 | 0.202 | 0.07 | 3.88e-41 | 1.31e-36 |
| Malignant epithelial | C1 Das ECs | NPDC1 | 1.69 | 0.709 | 0.34 | 2.99e-161 | 1.01e-156 |
| Malignant epithelial | C1 Das ECs | SLC44A4 | 1.688 | 0.56 | 0.203 | 3.54e-138 | 1.19e-133 |
| Malignant epithelial | C1 Das ECs | C6orf222 | 1.688 | 0.195 | 0.059 | 3.94e-47 | 1.33e-42 |
| Malignant epithelial | C1 Das ECs | ENTPD2 | 1.686 | 0.247 | 0.083 | 5.23e-54 | 1.76e-49 |
| Malignant epithelial | C1 Das ECs | RALGPS1 | 1.685 | 0.127 | 0.044 | 2.2e-25 | 7.42e-21 |
| Malignant epithelial | C1 Das ECs | ERN2 | 1.681 | 0.141 | 0.053 | 2.31e-25 | 7.8e-21 |
| Malignant epithelial | C1 Das ECs | ATP2A3 | 1.677 | 0.101 | 0.035 | 7.56e-20 | 2.55e-15 |
| Malignant epithelial | C1 Das ECs | CDH17 | 1.675 | 0.658 | 0.209 | 9.780000000000001e-192 | 3.3e-187 |
| Malignant epithelial | C1 Das ECs | MAOA | 1.672 | 0.239 | 0.075 | 3.34e-56 | 1.12e-51 |
| Malignant epithelial | C1 Das ECs | MIR194-2HG | 1.664 | 0.122 | 0.04 | 7.020000000000001e-26 | 2.36e-21 |
| Malignant epithelial | C1 Das ECs | SLC39A5 | 1.659 | 0.217 | 0.076 | 1.95e-43 | 6.58e-39 |
| Malignant epithelial | C1 Das ECs | ACE2 | 1.652 | 0.116 | 0.035 | 5.59e-28 | 1.88e-23 |
| Malignant epithelial | C1 Das ECs | POF1B | 1.65 | 0.302 | 0.11 | 1.57e-61 | 5.300000000000001e-57 |
| Malignant epithelial | C1 Das ECs | SIAE | 1.646 | 0.144 | 0.049 | 5.93e-30 | 2e-25 |
| Malignant epithelial | C1 Das ECs | ITGA1 | 1.646 | 0.106 | 0.037 | 9.070000000000001e-21 | 3.06e-16 |
| Malignant epithelial | C1 Das ECs | ANKS4B | 1.641 | 0.134 | 0.046 | 4.11e-27 | 1.38e-22 |
| Malignant epithelial | C1 Das ECs | GDA | 1.635 | 0.107 | 0.043 | 1.93e-17 | 6.51e-13 |
| Malignant epithelial | C1 Das ECs | RP11-1143G9.4 | 1.632 | 0.297 | 0.127 | 7.6e-45 | 2.56e-40 |
| Malignant epithelial | C1 Das ECs | TOX3 | 1.631 | 0.297 | 0.115 | 8.87e-55 | 2.99e-50 |
| Malignant epithelial | C1 Das ECs | ARID5B | 1.623 | 0.365 | 0.135 | 1.08e-74 | 3.64e-70 |
| Malignant epithelial | C1 Das ECs | KCNQ1 | 1.623 | 0.225 | 0.079 | 4.83e-45 | 1.63e-40 |
| Malignant epithelial | C1 Das ECs | APIP | 1.617 | 0.459 | 0.226 | 3.22e-69 | 1.08e-64 |
| Malignant epithelial | C1 Das ECs | SCG5 | 1.615 | 0.14 | 0.05 | 2.23e-26 | 7.500000000000001e-22 |
| Malignant epithelial | C1 Das ECs | MFHAS1 | 1.615 | 0.145 | 0.055 | 6.69e-26 | 2.25e-21 |
| Malignant epithelial | C1 Das ECs | MECOM | 1.614 | 0.193 | 0.065 | 1.57e-40 | 5.3e-36 |
| Malignant epithelial | C1 Das ECs | PPP1R9A | 1.607 | 0.108 | 0.038 | 1.06e-20 | 3.58e-16 |
| Malignant epithelial | C1 Das ECs | MID1 | 1.6 | 0.116 | 0.038 | 7.02e-25 | 2.37e-20 |
| Malignant epithelial | C1 Das ECs | ASS1 | 1.597 | 0.747 | 0.37 | 3.01e-163 | 1.01e-158 |
| Malignant epithelial | C1 Das ECs | BMP2 | 1.593 | 0.126 | 0.036 | 1.97e-31 | 6.62e-27 |
| Malignant epithelial | C1 Das ECs | RNF128 | 1.592 | 0.404 | 0.18 | 1.2e-64 | 4.03e-60 |
| Malignant epithelial | C1 Das ECs | RBPMS | 1.59 | 0.168 | 0.065 | 1.03e-28 | 3.46e-24 |
| Malignant epithelial | C1 Das ECs | SYBU | 1.587 | 0.159 | 0.054 | 1.57e-32 | 5.29e-28 |
| Malignant epithelial | C1 Das ECs | CDCA7 | 1.581 | 0.424 | 0.179 | 8.75e-76 | 2.95e-71 |
| Malignant epithelial | C1 Das ECs | RAPGEF5 | 1.578 | 0.112 | 0.044 | 9.71e-19 | 3.27e-14 |
| Malignant epithelial | C1 Das ECs | VDR | 1.577 | 0.168 | 0.064 | 4.36e-29 | 1.47e-24 |
| Malignant epithelial | C1 Das ECs | ACTL10 | 1.568 | 0.134 | 0.056 | 7.72e-20 | 2.6e-15 |
| Malignant epithelial | C1 Das ECs | SMIM24 | 1.561 | 0.332 | 0.125 | 2.15e-63 | 7.26e-59 |
| Malignant epithelial | C1 Das ECs | CRACR2B | 1.559 | 0.463 | 0.201 | 1.04e-81 | 3.49e-77 |
| Malignant epithelial | C1 Das ECs | RNASE1 | 1.558 | 0.514 | 0.167 | 2e-135 | 6.73e-131 |
| Malignant epithelial | C1 Das ECs | ARSE | 1.557 | 0.408 | 0.176 | 9.49e-70 | 3.2e-65 |
| Malignant epithelial | C1 Das ECs | MAP2K6 | 1.54 | 0.159 | 0.064 | 2.03e-25 | 6.840000000000001e-21 |
| Malignant epithelial | C1 Das ECs | NEDD9 | 1.538 | 0.278 | 0.098 | 9.85e-57 | 3.32e-52 |
| Malignant epithelial | C1 Das ECs | PRSS3 | 1.529 | 0.914 | 0.638 | 3.32e-229 | 1.12e-224 |
| Malignant epithelial | C1 Das ECs | MUC17 | 1.529 | 0.287 | 0.073 | 4.52e-85 | 1.52e-80 |
| Malignant epithelial | C1 Das ECs | NOTCH1 | 1.516 | 0.133 | 0.054 | 8.49e-21 | 2.86e-16 |

| compartment | subcluster | gene | avg_log2FC | pct.1 | pct.2 | p_val | p_val_adj |
|---|---|---|---|---|---|---|---|
| Malignant epithelial | C1 Das ECs | MUC3A | 1.502 | 0.543 | 0.221 | 3.54e-114 | 1.19e-109 |
| Malignant epithelial | C1 Das ECs | TNIK | 1.502 | 0.16 | 0.068 | 3.29e-23 | 1.11e-18 |
| Malignant epithelial | C1 Das ECs | MMEL1 | 1.501 | 0.174 | 0.061 | 4.16e-34 | 1.4e-29 |
| Malignant epithelial | C1 Das ECs | HNF4G | 1.498 | 0.127 | 0.048 | 2.5e-22 | 8.42e-18 |
| Malignant epithelial | C1 Das ECs | CEACAM5 | 1.496 | 0.819 | 0.256 | 2.39e-274 | 8.069999999999999e-270 |
| Malignant epithelial | C1 Das ECs | ZBTB18 | 1.496 | 0.117 | 0.046 | 7.63e-20 | 2.57e-15 |
| Malignant epithelial | C1 Das ECs | AKR7A3 | 1.495 | 0.284 | 0.113 | 2.35e-50 | 7.93e-46 |
| Malignant epithelial | C1 Das ECs | USP30-AS1 | 1.482 | 0.127 | 0.052 | 5.16e-19 | 1.74e-14 |
| Malignant epithelial | C1 Das ECs | TLE2 | 1.478 | 0.161 | 0.06 | 2.32e-28 | 7.83e-24 |
| Malignant epithelial | C1 Das ECs | CKMT1B | 1.476 | 0.205 | 0.093 | 2.3e-27 | 7.729999999999999e-23 |
| Malignant epithelial | C1 Das ECs | CTSO | 1.476 | 0.124 | 0.049 | 5.37e-20 | 1.81e-15 |
| Malignant epithelial | C1 Das ECs | TMEM246 | 1.475 | 0.16 | 0.06 | 1.02e-28 | 3.45e-24 |
| Malignant epithelial | C1 Das ECs | MYB | 1.472 | 0.31 | 0.123 | 1.98e-54 | 6.66e-50 |
| Malignant epithelial | C1 Das ECs | HSD11B2 | 1.469 | 0.298 | 0.12 | 3.07e-50 | 1.04e-45 |
| Malignant epithelial | C1 Das ECs | IGFBP2 | 1.466 | 0.476 | 0.161 | 1.01e-110 | 3.42e-106 |
| Malignant epithelial | C1 Das ECs | GALC | 1.466 | 0.135 | 0.052 | 2.05e-22 | 6.92e-18 |
| Malignant epithelial | C1 Das ECs | TNFRSF14 | 1.463 | 0.5 | 0.24 | 1.06e-81 | 3.57e-77 |
| Malignant epithelial | C1 Das ECs | RALGAPA2 | 1.46 | 0.214 | 0.083 | 3.85e-36 | 1.3e-31 |
| Malignant epithelial | C1 Das ECs | RAPGEFL1 | 1.45 | 0.211 | 0.093 | 2.23e-29 | 7.51e-25 |
| Malignant epithelial | C1 Das ECs | RAB15 | 1.429 | 0.107 | 0.042 | 1.49e-17 | 5.02e-13 |
| Malignant epithelial | C1 Das ECs | EPB41L4B | 1.424 | 0.168 | 0.075 | 5.92e-23 | 2e-18 |
| Malignant epithelial | C1 Das ECs | ACVR1B | 1.421 | 0.139 | 0.06 | 1.4e-19 | 4.73e-15 |
| Malignant epithelial | C1 Das ECs | TNFSF13B | 1.42 | 0.124 | 0.053 | 8.22e-18 | 2.77e-13 |
| Malignant epithelial | C1 Das ECs | SPDEF | 1.42 | 0.131 | 0.067 | 2.36e-13 | 7.96e-09 |
| Malignant epithelial | C1 Das ECs | STX19 | 1.414 | 0.109 | 0.046 | 2.77e-16 | 9.33e-12 |
| Malignant epithelial | C1 Das ECs | NPIPB15 | 1.411 | 0.115 | 0.052 | 1.6e-14 | 5.4e-10 |
| Malignant epithelial | C1 Das ECs | ACSS1 | 1.406 | 0.263 | 0.11 | 5.16e-41 | 1.74e-36 |
| Malignant epithelial | C1 Das ECs | CYP3A5 | 1.404 | 0.602 | 0.296 | 1.12e-100 | 3.78e-96 |
| Malignant epithelial | C1 Das ECs | CYP4F3 | 1.404 | 0.216 | 0.085 | 3.54e-36 | 1.19e-31 |
| Malignant epithelial | C1 Das ECs | JPH1 | 1.399 | 0.184 | 0.089 | 1.76e-21 | 5.94e-17 |
| Malignant epithelial | C1 Das ECs | GATA6 | 1.397 | 0.161 | 0.064 | 2.92e-26 | 9.84e-22 |
| Malignant epithelial | C1 Das ECs | GPA33 | 1.391 | 0.173 | 0.052 | 2.33e-41 | 7.85e-37 |
| Malignant epithelial | C1 Das ECs | CAPN5 | 1.39 | 0.244 | 0.094 | 2.6e-42 | 8.77e-38 |
| Malignant epithelial | C1 Das ECs | CYP27A1 | 1.383 | 0.139 | 0.06 | 2.19e-19 | 7.38e-15 |
| Malignant epithelial | C1 Das ECs | MYO7B | 1.375 | 0.13 | 0.051 | 4.65e-21 | 1.57e-16 |
| Malignant epithelial | C1 Das ECs | UTP14A | 1.359 | 0.264 | 0.12 | 3.15e-36 | 1.06e-31 |
| Malignant epithelial | C1 Das ECs | NHSL1 | 1.353 | 0.186 | 0.084 | 8.92e-24 | 3.01e-19 |
| Malignant epithelial | C1 Das ECs | PDE4C | 1.352 | 0.526 | 0.229 | 2.74e-92 | 9.23e-88 |
| Malignant epithelial | C1 Das ECs | CASK | 1.347 | 0.254 | 0.112 | 3.1e-36 | 1.05e-31 |
| Malignant epithelial | C1 Das ECs | KIAA0226L | 1.347 | 0.121 | 0.044 | 2.84e-22 | 9.57e-18 |
| Malignant epithelial | C1 Das ECs | LYZ | 1.345 | 0.908 | 0.737 | 2.65e-122 | 8.94e-118 |
| Malignant epithelial | C1 Das ECs | C9orf72 | 1.344 | 0.138 | 0.048 | 5.180000000000001e-27 | 1.75e-22 |
| Malignant epithelial | C1 Das ECs | BARX2 | 1.339 | 0.11 | 0.055 | 1.32e-11 | 4.45e-07 |
| Malignant epithelial | C1 Das ECs | PPFIBP2 | 1.336 | 0.115 | 0.045 | 7.95e-19 | 2.68e-14 |
| Malignant epithelial | C1 Das ECs | RBBP8NL | 1.335 | 0.134 | 0.052 | 6.96e-22 | 2.34e-17 |
| Malignant epithelial | C1 Das ECs | NEO1 | 1.334 | 0.125 | 0.053 | 8.99e-18 | 3.03e-13 |
| Malignant epithelial | C1 Das ECs | KLHL23 | 1.333 | 0.159 | 0.072 | 1.8e-20 | 6.06e-16 |
| Malignant epithelial | C1 Das ECs | TSPAN3 | 1.33 | 0.706 | 0.387 | 4.450000000000001e-124 | 1.5e-119 |
| Malignant epithelial | C1 Das ECs | GALNT12 | 1.328 | 0.175 | 0.083 | 3.23e-21 | 1.09e-16 |
| Malignant epithelial | C1 Das ECs | ARHGEF10L | 1.327 | 0.143 | 0.068 | 9.34e-17 | 3.15e-12 |
| Malignant epithelial | C1 Das ECs | QTRT1 | 1.317 | 0.557 | 0.322 | 4.38e-67 | 1.48e-62 |
| Malignant epithelial | C1 Das ECs | RHOBTB3 | 1.314 | 0.378 | 0.152 | 5.45e-65 | 1.84e-60 |
| Malignant epithelial | C1 Das ECs | PIGZ | 1.314 | 0.109 | 0.046 | 1.46e-15 | 4.91e-11 |
| Malignant epithelial | C1 Das ECs | KCTD14 | 1.313 | 0.131 | 0.062 | 3.16e-15 | 1.06e-10 |
| Malignant epithelial | C1 Das ECs | KCNK9 | 1.31 | 0.132 | 0.051 | 9.35e-22 | 3.15e-17 |
| Malignant epithelial | C1 Das ECs | CD24 | 1.306 | 0.915 | 0.367 | 1.63e-227 | 5.480000000000001e-223 |
| Malignant epithelial | C1 Das ECs | DIAPH2 | 1.306 | 0.272 | 0.136 | 4.83e-31 | 1.63e-26 |
| Malignant epithelial | C1 Das ECs | KALRN | 1.302 | 0.117 | 0.045 | 1.57e-19 | 5.29e-15 |
| Malignant epithelial | C1 Das ECs | CRYL1 | 1.301 | 0.299 | 0.144 | 2.71e-37 | 9.120000000000001e-33 |
| Malignant epithelial | C1 Das ECs | ETFDH | 1.299 | 0.179 | 0.078 | 9.52e-25 | 3.21e-20 |
| Malignant epithelial | C1 Das ECs | IL17RB | 1.297 | 0.122 | 0.056 | 6.7e-15 | 2.26e-10 |
| Malignant epithelial | C1 Das ECs | HOTTIP | 1.293 | 0.126 | 0.053 | 4.54e-18 | 1.53e-13 |
| Malignant epithelial | C1 Das ECs | CALML4 | 1.292 | 0.113 | 0.054 | 3.65e-13 | 1.23e-08 |
| Malignant epithelial | C1 Das ECs | SPINK1 | 1.289 | 0.594 | 0.183 | 1.26e-163 | 4.26e-159 |

| compartment | subcluster | gene | avg_log2FC | pct.1 | pct.2 | p_val | p_val_adj |
|---|---|---|---|---|---|---|---|
| Malignant epithelial | C1 Das ECs | LYN | 1.288 | 0.223 | 0.107 | 3.55e-27 | 1.2e-22 |
| Malignant epithelial | C1 Das ECs | PCLO | 1.286 | 0.167 | 0.075 | 6.49e-22 | 2.19e-17 |
| Malignant epithelial | C1 Das ECs | XK | 1.286 | 0.124 | 0.058 | 3.74e-15 | 1.26e-10 |
| Malignant epithelial | C1 Das ECs | HDHD3 | 1.285 | 0.609 | 0.355 | 2.77e-78 | 9.349999999999999e-74 |
| Malignant epithelial | C1 Das ECs | POC1B | 1.279 | 0.232 | 0.107 | 8.63e-30 | 2.91e-25 |
| Malignant epithelial | C1 Das ECs | FERMT1 | 1.278 | 0.691 | 0.422 | 1e-95 | 3.38e-91 |
| Malignant epithelial | C1 Das ECs | BDH1 | 1.276 | 0.232 | 0.114 | 1.01e-26 | 3.39e-22 |
| Malignant epithelial | C1 Das ECs | PRR15 | 1.275 | 0.451 | 0.23 | 2.8e-57 | 9.45e-53 |
| Malignant epithelial | C1 Das ECs | AUTS2 | 1.275 | 0.136 | 0.063 | 5.33e-17 | 1.8e-12 |
| Malignant epithelial | C1 Das ECs | TSPAN12 | 1.273 | 0.187 | 0.066 | 2.83e-35 | 9.520000000000001e-31 |
| Malignant epithelial | C1 Das ECs | SMAD6 | 1.273 | 0.14 | 0.068 | 8.82e-16 | 2.97e-11 |
| Malignant epithelial | C1 Das ECs | RSPH1 | 1.271 | 0.136 | 0.061 | 3.77e-18 | 1.27e-13 |
| Malignant epithelial | C1 Das ECs | HNF4A | 1.27 | 0.371 | 0.2 | 7.52e-39 | 2.53e-34 |
| Malignant epithelial | C1 Das ECs | SEMA4D | 1.269 | 0.168 | 0.081 | 1.08e-18 | 3.65e-14 |
| Malignant epithelial | C1 Das ECs | RP3-325F22.5 | 1.268 | 0.11 | 0.048 | 7.37e-15 | 2.48e-10 |
| Malignant epithelial | C1 Das ECs | TRMT10B | 1.265 | 0.141 | 0.067 | 3.47e-16 | 1.17e-11 |
| Malignant epithelial | C1 Das ECs | CHDH | 1.263 | 0.267 | 0.13 | 2.56e-31 | 8.630000000000001e-27 |
| Malignant epithelial | C1 Das ECs | HAUS7 | 1.256 | 0.154 | 0.081 | 1.53e-14 | 5.16e-10 |
| Malignant epithelial | C1 Das ECs | TMEM192 | 1.251 | 0.191 | 0.095 | 1.94e-20 | 6.54e-16 |
| Malignant epithelial | C1 Das ECs | ALDH6A1 | 1.251 | 0.117 | 0.055 | 5.96e-14 | 2.01e-09 |
| Malignant epithelial | C1 Das ECs | DUSP8 | 1.249 | 0.171 | 0.08 | 2.15e-20 | 7.25e-16 |
| Malignant epithelial | C1 Das ECs | SLC44A3 | 1.246 | 0.203 | 0.094 | 1.92e-25 | 6.46e-21 |
| Malignant epithelial | C1 Das ECs | CYP4F12 | 1.244 | 0.121 | 0.052 | 1.05e-16 | 3.54e-12 |
| Malignant epithelial | C1 Das ECs | CTSS | 1.243 | 0.688 | 0.368 | 1.1e-110 | 3.72e-106 |
| Malignant epithelial | C1 Das ECs | SYTL2 | 1.238 | 0.174 | 0.071 | 1.34e-26 | 4.500000000000001e-22 |
| Malignant epithelial | C1 Das ECs | HID1 | 1.237 | 0.11 | 0.058 | 2.2e-10 | 7.43e-06 |
| Malignant epithelial | C1 Das ECs | NUDT7 | 1.236 | 0.155 | 0.077 | 1.9e-16 | 6.39e-12 |
| Malignant epithelial | C1 Das ECs | PAQR8 | 1.235 | 0.114 | 0.055 | 7.56e-13 | 2.55e-08 |
| Malignant epithelial | C1 Das ECs | PRMT7 | 1.231 | 0.165 | 0.086 | 8.69e-16 | 2.93e-11 |
| Malignant epithelial | C1 Das ECs | CNKSR3 | 1.228 | 0.135 | 0.055 | 3.44e-21 | 1.16e-16 |
| Malignant epithelial | C1 Das ECs | STARD10 | 1.225 | 0.767 | 0.477 | 1.13e-110 | 3.82e-106 |
| Malignant epithelial | C1 Das ECs | SPAG1 | 1.224 | 0.217 | 0.1 | 1.8e-27 | 6.059999999999999e-23 |
| Malignant epithelial | C1 Das ECs | CLDN15 | 1.223 | 0.259 | 0.118 | 4.92e-34 | 1.66e-29 |
| Malignant epithelial | C1 Das ECs | HOXD-AS2 | 1.222 | 0.15 | 0.061 | 1.04e-22 | 3.52e-18 |
| Malignant epithelial | C1 Das ECs | SEMA4G | 1.221 | 0.148 | 0.067 | 1.01e-18 | 3.39e-14 |
| Malignant epithelial | C1 Das ECs | NANS | 1.22 | 0.733 | 0.45 | 5.45e-107 | 1.84e-102 |
| Malignant epithelial | C1 Das ECs | SCML1 | 1.216 | 0.278 | 0.145 | 1.29e-28 | 4.35e-24 |
| Malignant epithelial | C1 Das ECs | TPPP3 | 1.214 | 0.279 | 0.116 | 2.01e-43 | 6.77e-39 |
| Malignant epithelial | C1 Das ECs | MT1G | 1.212 | 0.183 | 0.084 | 2.73e-23 | 9.2e-19 |
| Malignant epithelial | C1 Das ECs | MINCR | 1.207 | 0.225 | 0.115 | 4.47e-23 | 1.51e-18 |
| Malignant epithelial | C1 Das ECs | LACTB2 | 1.206 | 0.507 | 0.293 | 4.96e-55 | 1.67e-50 |
| Malignant epithelial | C1 Das ECs | LIMA1 | 1.198 | 0.463 | 0.245 | 1.37e-55 | 4.6e-51 |
| Malignant epithelial | C1 Das ECs | PRKAB1 | 1.197 | 0.197 | 0.107 | 9.41e-18 | 3.17e-13 |
| Malignant epithelial | C1 Das ECs | TPRN | 1.191 | 0.184 | 0.092 | 1.44e-19 | 4.86e-15 |
| Malignant epithelial | C1 Das ECs | CXADR | 1.19 | 0.589 | 0.284 | 5.96e-95 | 2.01e-90 |
| Malignant epithelial | C1 Das ECs | CDC42EP5 | 1.189 | 0.277 | 0.131 | 2e-34 | 6.74e-30 |
| Malignant epithelial | C1 Das ECs | SPIRE2 | 1.189 | 0.27 | 0.13 | 2.4e-32 | 8.07e-28 |
| Malignant epithelial | C1 Das ECs | TTC39B | 1.188 | 0.155 | 0.074 | 1.6e-17 | 5.4e-13 |
| Malignant epithelial | C1 Das ECs | CCHCR1 | 1.182 | 0.176 | 0.09 | 9.15e-18 | 3.08e-13 |
| Malignant epithelial | C1 Das ECs | VILL | 1.181 | 0.2 | 0.09 | 1.35e-25 | 4.56e-21 |
| Malignant epithelial | C1 Das ECs | TMEM98 | 1.175 | 0.323 | 0.118 | 6.96e-60 | 2.35e-55 |
| Malignant epithelial | C1 Das ECs | PNPLA4 | 1.174 | 0.364 | 0.213 | 1.81e-31 | 6.09e-27 |
| Malignant epithelial | C1 Das ECs | DDAH1 | 1.172 | 0.215 | 0.101 | 3.62e-26 | 1.22e-21 |
| Malignant epithelial | C1 Das ECs | RNF43 | 1.171 | 0.485 | 0.247 | 6.02e-59 | 2.03e-54 |
| Malignant epithelial | C1 Das ECs | C6orf223 | 1.169 | 0.107 | 0.049 | 4.56e-13 | 1.54e-08 |
| Malignant epithelial | C1 Das ECs | SLC2A8 | 1.168 | 0.162 | 0.083 | 2.04e-16 | 6.88e-12 |
| Malignant epithelial | C1 Das ECs | IFT74 | 1.167 | 0.123 | 0.062 | 1.3e-12 | 4.39e-08 |
| Malignant epithelial | C1 Das ECs | PHGR1 | 1.166 | 0.957 | 0.451 | 4.869999999999999e-263 | 1.64e-258 |
| Malignant epithelial | C1 Das ECs | ENOX2 | 1.166 | 0.108 | 0.057 | 6.52e-10 | 2.2e-05 |
| Malignant epithelial | C1 Das ECs | COL27A1 | 1.165 | 0.141 | 0.062 | 1.26e-18 | 4.25e-14 |
| Malignant epithelial | C1 Das ECs | C2 | 1.163 | 0.312 | 0.157 | 5.96e-35 | 2.01e-30 |
| Malignant epithelial | C1 Das ECs | VSIG2 | 1.162 | 0.127 | 0.058 | 8.61e-16 | 2.9e-11 |
| Malignant epithelial | C1 Das ECs | RP11-452F19.3 | 1.161 | 0.151 | 0.076 | 2.1e-15 | 7.07e-11 |
| Malignant epithelial | C1 Das ECs | PLS1 | 1.157 | 0.351 | 0.165 | 1.17e-45 | 3.94e-41 |

| compartment | subcluster | gene | avg_log2FC | pct.1 | pct.2 | p_val | p_val_adj |
|---|---|---|---|---|---|---|---|
| Malignant epithelial | C1 Das ECs | SYNJ2 | 1.155 | 0.128 | 0.068 | 1.49e-11 | 5.03e-07 |
| Malignant epithelial | C1 Das ECs | SHROOM3 | 1.153 | 0.181 | 0.087 | 1.15e-20 | 3.86e-16 |
| Malignant epithelial | C1 Das ECs | SNTB1 | 1.151 | 0.136 | 0.079 | 5.1e-10 | 1.72e-05 |
| Malignant epithelial | C1 Das ECs | BAIAP2L2 | 1.15 | 0.234 | 0.116 | 1.81e-25 | 6.110000000000001e-21 |
| Malignant epithelial | C1 Das ECs | RP11-395B7.4 | 1.15 | 0.122 | 0.056 | 1.39e-14 | 4.67e-10 |
| Malignant epithelial | C1 Das ECs | FOXN3 | 1.149 | 0.134 | 0.061 | 1.75e-16 | 5.91e-12 |
| Malignant epithelial | C1 Das ECs | ZDHHC9 | 1.141 | 0.359 | 0.203 | 4.149999999999999e-34 | 1.4e-29 |
| Malignant epithelial | C1 Das ECs | PYGB | 1.139 | 0.589 | 0.35 | 3.89e-63 | 1.31e-58 |
| Malignant epithelial | C1 Das ECs | MEIS2 | 1.135 | 0.135 | 0.063 | 7.76e-16 | 2.61e-11 |
| Malignant epithelial | C1 Das ECs | MSI2 | 1.134 | 0.371 | 0.199 | 5.59e-38 | 1.88e-33 |
| Malignant epithelial | C1 Das ECs | PRADC1 | 1.133 | 0.183 | 0.094 | 6.24e-18 | 2.1e-13 |
| Malignant epithelial | C1 Das ECs | GMDS | 1.13 | 0.479 | 0.272 | 1.01e-49 | 3.4e-45 |
| Malignant epithelial | C1 Das ECs | RNF186 | 1.127 | 0.103 | 0.051 | 7.14e-11 | 2.41e-06 |
| Malignant epithelial | C1 Das ECs | MLXIP | 1.126 | 0.475 | 0.277 | 1.02e-46 | 3.43e-42 |
| Malignant epithelial | C1 Das ECs | PHF8 | 1.126 | 0.101 | 0.058 | 8.89e-08 | 0.003 |
| Malignant epithelial | C1 Das ECs | RMND1 | 1.125 | 0.162 | 0.089 | 1.01e-13 | 3.39e-09 |
| Malignant epithelial | C1 Das ECs | RP11-362F19.1 | 1.123 | 0.117 | 0.055 | 2e-13 | 6.73e-09 |
| Malignant epithelial | C1 Das ECs | CDX2 | 1.122 | 0.332 | 0.145 | 1.52e-46 | 5.12e-42 |
| Malignant epithelial | C1 Das ECs | ZCCHC7 | 1.117 | 0.394 | 0.228 | 2.9e-36 | 9.780000000000001e-32 |
| Malignant epithelial | C1 Das ECs | AC021218.2 | 1.116 | 0.164 | 0.083 | 1.41e-16 | 4.73e-12 |
| Malignant epithelial | C1 Das ECs | CES2 | 1.114 | 0.333 | 0.193 | 3.17e-28 | 1.07e-23 |
| Malignant epithelial | C1 Das ECs | CTGF | 1.109 | 0.321 | 0.144 | 1.25e-43 | 4.21e-39 |
| Malignant epithelial | C1 Das ECs | IFI44L | 1.108 | 0.283 | 0.155 | 2.56e-25 | 8.63e-21 |
| Malignant epithelial | C1 Das ECs | RP11-734K2.4 | 1.108 | 0.102 | 0.059 | 1.35e-07 | 0.00456 |
| Malignant epithelial | C1 Das ECs | FUT3 | 1.107 | 0.217 | 0.097 | 5.74e-28 | 1.93e-23 |
| Malignant epithelial | C1 Das ECs | CCNO | 1.105 | 0.128 | 0.067 | 4.77e-12 | 1.61e-07 |
| Malignant epithelial | C1 Das ECs | IGBP1 | 1.102 | 0.746 | 0.524 | 1.48e-89 | 5e-85 |
| Malignant epithelial | C1 Das ECs | ACSF3 | 1.096 | 0.208 | 0.122 | 3.22e-15 | 1.08e-10 |
| Malignant epithelial | C1 Das ECs | PPM1N | 1.094 | 0.178 | 0.089 | 3.36e-18 | 1.13e-13 |
| Malignant epithelial | C1 Das ECs | TMEM243 | 1.09 | 0.39 | 0.232 | 2.13e-33 | 7.160000000000001e-29 |
| Malignant epithelial | C1 Das ECs | C12orf75 | 1.089 | 0.809 | 0.51 | 3.25e-127 | 1.09e-122 |
| Malignant epithelial | C1 Das ECs | NEK6 | 1.089 | 0.291 | 0.156 | 4.9e-28 | 1.65e-23 |
| Malignant epithelial | C1 Das ECs | FOXA2 | 1.089 | 0.248 | 0.124 | 2.2e-26 | 7.41e-22 |
| Malignant epithelial | C1 Das ECs | OAF | 1.087 | 0.248 | 0.137 | 3.32e-21 | 1.12e-16 |
| Malignant epithelial | C1 Das ECs | RAB40B | 1.086 | 0.334 | 0.17 | 5.07e-37 | 1.71e-32 |
| Malignant epithelial | C1 Das ECs | PKP2 | 1.086 | 0.289 | 0.155 | 3.83e-28 | 1.29e-23 |
| Malignant epithelial | C1 Das ECs | TRIM14 | 1.086 | 0.191 | 0.098 | 1.03e-18 | 3.46e-14 |
| Malignant epithelial | C1 Das ECs | TACC1 | 1.084 | 0.321 | 0.168 | 2.41e-33 | 8.13e-29 |
| Malignant epithelial | C1 Das ECs | KIAA1161 | 1.083 | 0.105 | 0.048 | 3.34e-13 | 1.12e-08 |
| Malignant epithelial | C1 Das ECs | AC009403.2 | 1.082 | 0.202 | 0.109 | 5.88e-18 | 1.98e-13 |
| Malignant epithelial | C1 Das ECs | GADD45G | 1.08 | 0.346 | 0.198 | 2.32e-29 | 7.8e-25 |
| Malignant epithelial | C1 Das ECs | SH3BGRL2 | 1.074 | 0.24 | 0.115 | 1.92e-27 | 6.46e-23 |
| Malignant epithelial | C1 Das ECs | SMKR1 | 1.074 | 0.121 | 0.057 | 6.9e-14 | 2.33e-09 |
| Malignant epithelial | C1 Das ECs | TMEM187 | 1.072 | 0.15 | 0.081 | 1.98e-13 | 6.68e-09 |
| Malignant epithelial | C1 Das ECs | FAM134B | 1.071 | 0.166 | 0.062 | 8.05e-29 | 2.71e-24 |
| Malignant epithelial | C1 Das ECs | CORO2A | 1.071 | 0.21 | 0.107 | 2.58e-21 | 8.7e-17 |
| Malignant epithelial | C1 Das ECs | CKMT1A | 1.071 | 0.112 | 0.057 | 4.07e-11 | 1.37e-06 |
| Malignant epithelial | C1 Das ECs | RAB20 | 1.07 | 0.326 | 0.192 | 2.98e-25 | 1e-20 |
| Malignant epithelial | C1 Das ECs | FAM213B | 1.069 | 0.345 | 0.195 | 4.790000000000001e-31 | 1.61e-26 |
| Malignant epithelial | C1 Das ECs | MYO1A | 1.066 | 0.135 | 0.068 | 1.55e-13 | 5.24e-09 |
| Malignant epithelial | C1 Das ECs | BACE2 | 1.065 | 0.576 | 0.351 | 2.19e-61 | 7.39e-57 |
| Malignant epithelial | C1 Das ECs | FABP1 | 1.065 | 0.13 | 0.072 | 1.19e-10 | 4.02e-06 |
| Malignant epithelial | C1 Das ECs | C5orf45 | 1.063 | 0.195 | 0.118 | 6.84e-13 | 2.3e-08 |
| Malignant epithelial | C1 Das ECs | FAM3A | 1.058 | 0.42 | 0.254 | 8.22e-36 | 2.77e-31 |
| Malignant epithelial | C1 Das ECs | HLA-DMB | 1.054 | 0.236 | 0.132 | 3.48e-19 | 1.17e-14 |
| Malignant epithelial | C1 Das ECs | BCL2L15 | 1.051 | 0.277 | 0.147 | 1.15e-26 | 3.88e-22 |
| Malignant epithelial | C1 Das ECs | LINC01558 | 1.046 | 0.101 | 0.055 | 3.03e-08 | 0.00102 |
| Malignant epithelial | C1 Das ECs | KANK1 | 1.045 | 0.141 | 0.072 | 5.67e-14 | 1.91e-09 |
| Malignant epithelial | C1 Das ECs | ARHGEF9 | 1.044 | 0.168 | 0.094 | 2.41e-13 | 8.11e-09 |
| Malignant epithelial | C1 Das ECs | NFIB | 1.043 | 0.346 | 0.19 | 3.33e-32 | 1.12e-27 |
| Malignant epithelial | C1 Das ECs | TSPAN5 | 1.042 | 0.191 | 0.095 | 1.04e-19 | 3.5e-15 |
| Malignant epithelial | C1 Das ECs | IQSEC2 | 1.042 | 0.117 | 0.06 | 1.17e-11 | 3.94e-07 |
| Malignant epithelial | C1 Das ECs | RERE | 1.04 | 0.274 | 0.156 | 1.07e-22 | 3.6e-18 |
| Malignant epithelial | C1 Das ECs | CXorf40A | 1.035 | 0.138 | 0.075 | 1.05e-11 | 3.52e-07 |

| compartment | subcluster | gene | avg_log2FC | pct.1 | pct.2 | p_val | p_val_adj |
|---|---|---|---|---|---|---|---|
| Malignant epithelial | C1 Das ECs | ASCL2 | 1.034 | 0.646 | 0.306 | 4.8e-106 | 1.62e-101 |
| Malignant epithelial | C1 Das ECs | GIMAP2 | 1.034 | 0.203 | 0.107 | 3.83e-19 | 1.29e-14 |
| Malignant epithelial | C1 Das ECs | HIPK2 | 1.032 | 0.133 | 0.074 | 1.12e-10 | 3.78e-06 |
| Malignant epithelial | C1 Das ECs | FAM84A | 1.03 | 0.373 | 0.172 | 7.76e-49 | 2.62e-44 |
| Malignant epithelial | C1 Das ECs | CLDN3 | 1.029 | 0.99 | 0.889 | 4.2e-179 | 1.42e-174 |
| Malignant epithelial | C1 Das ECs | IFT172 | 1.025 | 0.503 | 0.279 | 3.81e-53 | 1.28e-48 |
| Malignant epithelial | C1 Das ECs | LRRC28 | 1.022 | 0.155 | 0.09 | 1.11e-11 | 3.75e-07 |
| Malignant epithelial | C1 Das ECs | CERS6 | 1.021 | 0.352 | 0.204 | 1.11e-29 | 3.75e-25 |
| Malignant epithelial | C1 Das ECs | ETV7 | 1.02 | 0.219 | 0.124 | 3.91e-17 | 1.32e-12 |
| Malignant epithelial | C1 Das ECs | TP53TG1 | 1.019 | 0.402 | 0.231 | 2.47e-36 | 8.340000000000001e-32 |
| Malignant epithelial | C1 Das ECs | REPIN1 | 1.018 | 0.552 | 0.318 | 3.17e-57 | 1.07e-52 |
| Malignant epithelial | C1 Das ECs | GEMIN8 | 1.018 | 0.177 | 0.1 | 1.32e-13 | 4.43e-09 |
| Malignant epithelial | C1 Das ECs | SLC27A5 | 1.018 | 0.13 | 0.072 | 1.14e-10 | 3.83e-06 |
| Malignant epithelial | C1 Das ECs | GSDMB | 1.017 | 0.202 | 0.108 | 1.12e-17 | 3.78e-13 |
| Malignant epithelial | C1 Das ECs | ZBED1 | 1.016 | 0.15 | 0.082 | 1.27e-12 | 4.29e-08 |
| Malignant epithelial | C1 Das ECs | RP13-188A5.1 | 1.015 | 0.116 | 0.068 | 3.19e-08 | 0.00107 |
| Malignant epithelial | C1 Das ECs | RP4-568C11.4 | 1.011 | 0.224 | 0.12 | 3.12e-20 | 1.05e-15 |
| Malignant epithelial | C1 Das ECs | KHK | 1.008 | 0.146 | 0.085 | 9.54e-11 | 3.21e-06 |
| Malignant epithelial | C1 Das ECs | MPST | 1.006 | 0.852 | 0.626 | 1.75e-109 | 5.88e-105 |
| Malignant epithelial | C1 Das ECs | ALDH2 | 1.004 | 0.469 | 0.241 | 1.32e-54 | 4.44e-50 |
| Malignant epithelial | C1 Das ECs | SMCO4 | 1.003 | 0.273 | 0.142 | 6.160000000000001e-27 | 2.07e-22 |
| Malignant epithelial | C1 Das ECs | TXLNG | 1.003 | 0.236 | 0.134 | 1.14e-18 | 3.83e-14 |
| Malignant epithelial | C1 Das ECs | SLC25A5 | 1.002 | 0.95 | 0.87 | 2.14e-130 | 7.2e-126 |
| Malignant epithelial | C1 Das ECs | APTR | 1.002 | 0.319 | 0.169 | 2.07e-31 | 6.970000000000001e-27 |
| Malignant epithelial | C1 Das ECs | C9orf85 | 1.001 | 0.244 | 0.144 | 5.79e-18 | 1.95e-13 |
| Malignant epithelial | C1 Das ECs | HMHA1 | 0.999 | 0.273 | 0.161 | 6.36e-20 | 2.14e-15 |
| Malignant epithelial | C1 Das ECs | SLC25A33 | 0.997 | 0.276 | 0.16 | 1.36e-21 | 4.58e-17 |
| Malignant epithelial | C1 Das ECs | TMEM2 | 0.997 | 0.244 | 0.138 | 3.68e-19 | 1.24e-14 |
| Malignant epithelial | C1 Das ECs | C9orf9 | 0.997 | 0.14 | 0.072 | 2.85e-13 | 9.61e-09 |
| Malignant epithelial | C1 Das ECs | NRIP1 | 0.997 | 0.152 | 0.084 | 2.45e-12 | 8.26e-08 |
| Malignant epithelial | C1 Das ECs | GK | 0.996 | 0.115 | 0.054 | 1.96e-13 | 6.62e-09 |
| Malignant epithelial | C1 Das ECs | BSPRY | 0.995 | 0.305 | 0.172 | 2.9e-26 | 9.76e-22 |
| Malignant epithelial | C1 Das ECs | MPC2 | 0.994 | 0.675 | 0.481 | 9.09e-64 | 3.06e-59 |
| Malignant epithelial | C1 Das ECs | TM4SF5 | 0.994 | 0.282 | 0.142 | 4.65e-30 | 1.57e-25 |
| Malignant epithelial | C1 Das ECs | STX17 | 0.994 | 0.188 | 0.107 | 1.99e-14 | 6.71e-10 |
| Malignant epithelial | C1 Das ECs | N4BP2L1 | 0.992 | 0.149 | 0.073 | 7.16e-16 | 2.41e-11 |
| Malignant epithelial | C1 Das ECs | RNASET2 | 0.991 | 0.856 | 0.614 | 2.67e-104 | 9e-100 |
| Malignant epithelial | C1 Das ECs | UGT8 | 0.991 | 0.106 | 0.062 | 2.04e-07 | 0.00688 |
| Malignant epithelial | C1 Das ECs | DAB2IP | 0.988 | 0.155 | 0.081 | 4.82e-14 | 1.63e-09 |
| Malignant epithelial | C1 Das ECs | SCAMP1-AS1 | 0.984 | 0.174 | 0.098 | 1.73e-13 | 5.82e-09 |
| Malignant epithelial | C1 Das ECs | RSPO3 | 0.982 | 0.317 | 0.169 | 5.190000000000001e-29 | 1.75e-24 |
| Malignant epithelial | C1 Das ECs | NXT2 | 0.982 | 0.113 | 0.071 | 1.17e-06 | 0.0394 |
| Malignant epithelial | C1 Das ECs | IDNK | 0.98 | 0.107 | 0.065 | 1e-06 | 0.0337 |
| Malignant epithelial | C1 Das ECs | AIFM1 | 0.978 | 0.223 | 0.128 | 7.41e-17 | 2.5e-12 |
| Malignant epithelial | C1 Das ECs | LINC00668 | 0.977 | 0.121 | 0.065 | 1.88e-10 | 6.34e-06 |
| Malignant epithelial | C1 Das ECs | TAB3 | 0.976 | 0.106 | 0.058 | 1.05e-08 | 0.000353 |
| Malignant epithelial | C1 Das ECs | WIPI1 | 0.973 | 0.162 | 0.079 | 1.12e-17 | 3.78e-13 |
| Malignant epithelial | C1 Das ECs | RAB11FIP1 | 0.972 | 0.75 | 0.524 | 2.68e-75 | 9.03e-71 |
| Malignant epithelial | C1 Das ECs | SAMD12 | 0.971 | 0.168 | 0.101 | 7.02e-11 | 2.37e-06 |
| Malignant epithelial | C1 Das ECs | ABCD1 | 0.971 | 0.119 | 0.069 | 1.91e-08 | 0.000642 |
| Malignant epithelial | C1 Das ECs | ACADS | 0.97 | 0.235 | 0.145 | 5.33e-15 | 1.8e-10 |
| Malignant epithelial | C1 Das ECs | ASNS | 0.965 | 0.204 | 0.121 | 5.06e-14 | 1.7e-09 |
| Malignant epithelial | C1 Das ECs | NOSTRIN | 0.965 | 0.168 | 0.093 | 2.27e-13 | 7.65e-09 |
| Malignant epithelial | C1 Das ECs | FBLIM1 | 0.965 | 0.157 | 0.089 | 4.02e-12 | 1.35e-07 |
| Malignant epithelial | C1 Das ECs | BRCC3 | 0.964 | 0.2 | 0.119 | 1.38e-13 | 4.63e-09 |
| Malignant epithelial | C1 Das ECs | HMGN5 | 0.964 | 0.155 | 0.086 | 2.44e-12 | 8.23e-08 |
| Malignant epithelial | C1 Das ECs | TSPAN13 | 0.963 | 0.542 | 0.306 | 1.83e-57 | 6.16e-53 |
| Malignant epithelial | C1 Das ECs | GOLM1 | 0.963 | 0.525 | 0.325 | 1.35e-44 | 4.54e-40 |
| Malignant epithelial | C1 Das ECs | AKAP1 | 0.958 | 0.486 | 0.317 | 5.5e-37 | 1.85e-32 |
| Malignant epithelial | C1 Das ECs | IL7 | 0.958 | 0.12 | 0.064 | 1.76e-10 | 5.91e-06 |
| Malignant epithelial | C1 Das ECs | FOXA3 | 0.957 | 0.206 | 0.1 | 2.45e-22 | 8.26e-18 |
| Malignant epithelial | C1 Das ECs | HSD17B10 | 0.956 | 0.728 | 0.537 | 1.02e-68 | 3.44e-64 |
| Malignant epithelial | C1 Das ECs | TMEM161B-AS1 | 0.956 | 0.107 | 0.056 | 1.01e-09 | 3.39e-05 |
| Malignant epithelial | C1 Das ECs | NOL9 | 0.956 | 0.108 | 0.065 | 5.4e-07 | 0.0182 |

| compartment | subcluster | gene | avg_log2FC | pct.1 | pct.2 | p_val | p_val_adj |
|---|---|---|---|---|---|---|---|
| Malignant epithelial | C1 Das ECs | ARID1B | 0.955 | 0.192 | 0.121 | 6.95e-11 | 2.34e-06 |
| Malignant epithelial | C1 Das ECs | CBWD3 | 0.952 | 0.187 | 0.107 | 3.86e-14 | 1.3e-09 |
| Malignant epithelial | C1 Das ECs | SMARCA2 | 0.95 | 0.164 | 0.097 | 2.04e-11 | 6.87e-07 |
| Malignant epithelial | C1 Das ECs | CNNM4 | 0.947 | 0.124 | 0.074 | 4.71e-08 | 0.00159 |
| Malignant epithelial | C1 Das ECs | SUCLG1 | 0.946 | 0.627 | 0.424 | 1.3e-54 | 4.38e-50 |
| Malignant epithelial | C1 Das ECs | PTGER4 | 0.945 | 0.129 | 0.065 | 4.88e-13 | 1.64e-08 |
| Malignant epithelial | C1 Das ECs | PEX7 | 0.943 | 0.205 | 0.121 | 1.65e-14 | 5.58e-10 |
| Malignant epithelial | C1 Das ECs | BTG2 | 0.942 | 0.8 | 0.514 | 3.53e-96 | 1.19e-91 |
| Malignant epithelial | C1 Das ECs | STXBP1 | 0.942 | 0.128 | 0.068 | 2.66e-11 | 8.96e-07 |
| Malignant epithelial | C1 Das ECs | IGHG3 | 0.938 | 0.655 | 0.451 | 6.71e-60 | 2.26e-55 |
| Malignant epithelial | C1 Das ECs | TPM1 | 0.936 | 0.79 | 0.546 | 4.930000000000001e-86 | 1.66e-81 |
| Malignant epithelial | C1 Das ECs | RP11-462G2.1 | 0.935 | 0.534 | 0.275 | 1.31e-60 | 4.4e-56 |
| Malignant epithelial | C1 Das ECs | BCAS1 | 0.935 | 0.135 | 0.064 | 1.49e-15 | 5.02e-11 |
| Malignant epithelial | C1 Das ECs | ACO1 | 0.934 | 0.181 | 0.094 | 3.6e-17 | 1.21e-12 |
| Malignant epithelial | C1 Das ECs | SH3RF1 | 0.931 | 0.181 | 0.106 | 8.4e-13 | 2.83e-08 |
| Malignant epithelial | C1 Das ECs | MED14 | 0.929 | 0.141 | 0.078 | 1.59e-11 | 5.37e-07 |
| Malignant epithelial | C1 Das ECs | PPARA | 0.927 | 0.134 | 0.075 | 1.65e-10 | 5.57e-06 |
| Malignant epithelial | C1 Das ECs | SPR | 0.926 | 0.458 | 0.285 | 6.65e-36 | 2.24e-31 |
| Malignant epithelial | C1 Das ECs | TMPRSS2 | 0.925 | 0.401 | 0.193 | 1.75e-51 | 5.91e-47 |
| Malignant epithelial | C1 Das ECs | ARHGEF40 | 0.924 | 0.112 | 0.052 | 2.58e-13 | 8.68e-09 |
| Malignant epithelial | C1 Das ECs | IRF2 | 0.923 | 0.26 | 0.155 | 1.38e-18 | 4.66e-14 |
| Malignant epithelial | C1 Das ECs | FKBP11 | 0.922 | 0.485 | 0.289 | 7.8e-43 | 2.63e-38 |
| Malignant epithelial | C1 Das ECs | PLD1 | 0.922 | 0.136 | 0.079 | 1.16e-09 | 3.9e-05 |
| Malignant epithelial | C1 Das ECs | GNAQ | 0.921 | 0.383 | 0.219 | 2.04e-32 | 6.89e-28 |
| Malignant epithelial | C1 Das ECs | RHBDL2 | 0.919 | 0.106 | 0.059 | 1.67e-08 | 0.000564 |
| Malignant epithelial | C1 Das ECs | FXYD3 | 0.918 | 0.967 | 0.842 | 1.53e-147 | 5.15e-143 |
| Malignant epithelial | C1 Das ECs | CXorf56 | 0.917 | 0.139 | 0.086 | 2.55e-08 | 0.00086 |
| Malignant epithelial | C1 Das ECs | PARD6A | 0.913 | 0.127 | 0.067 | 2.27e-11 | 7.64e-07 |
| Malignant epithelial | C1 Das ECs | TMEM144 | 0.91 | 0.107 | 0.056 | 9.12e-10 | 3.07e-05 |
| Malignant epithelial | C1 Das ECs | ELK4 | 0.909 | 0.188 | 0.107 | 8.83e-14 | 2.97e-09 |
| Malignant epithelial | C1 Das ECs | MANSC1 | 0.908 | 0.304 | 0.165 | 1.95e-26 | 6.58e-22 |
| Malignant epithelial | C1 Das ECs | GFPT1 | 0.907 | 0.357 | 0.193 | 6.33e-34 | 2.13e-29 |
| Malignant epithelial | C1 Das ECs | MORC4 | 0.905 | 0.114 | 0.069 | 3.93e-07 | 0.0133 |
| Malignant epithelial | C1 Das ECs | MUC13 | 0.904 | 0.589 | 0.232 | 1.38e-120 | 4.66e-116 |
| Malignant epithelial | C1 Das ECs | RPS6KA3 | 0.904 | 0.241 | 0.146 | 2.15e-16 | 7.23e-12 |
| Malignant epithelial | C1 Das ECs | TCEAL1 | 0.904 | 0.22 | 0.142 | 3.98e-12 | 1.34e-07 |
| Malignant epithelial | C1 Das ECs | SLC25A4 | 0.902 | 0.237 | 0.145 | 2.22e-15 | 7.5e-11 |
| Malignant epithelial | C1 Das ECs | ADCK3 | 0.899 | 0.23 | 0.13 | 6.39e-18 | 2.15e-13 |
| Malignant epithelial | C1 Das ECs | SGPP2 | 0.898 | 0.146 | 0.082 | 1.67e-11 | 5.61e-07 |
| Malignant epithelial | C1 Das ECs | PHOSPHO2 | 0.898 | 0.115 | 0.07 | 2.87e-07 | 0.00968 |
| Malignant epithelial | C1 Das ECs | PODXL2 | 0.897 | 0.207 | 0.115 | 3.2e-16 | 1.08e-11 |
| Malignant epithelial | C1 Das ECs | COBLL1 | 0.897 | 0.148 | 0.082 | 1.04e-11 | 3.52e-07 |
| Malignant epithelial | C1 Das ECs | ACOT2 | 0.897 | 0.136 | 0.083 | 1.7e-08 | 0.000572 |
| Malignant epithelial | C1 Das ECs | FOXP1 | 0.896 | 0.491 | 0.315 | 1.1e-34 | 3.7e-30 |
| Malignant epithelial | C1 Das ECs | FAM122A | 0.896 | 0.184 | 0.101 | 1.08e-14 | 3.65e-10 |
| Malignant epithelial | C1 Das ECs | TMEM38B | 0.889 | 0.273 | 0.16 | 3.87e-20 | 1.3e-15 |
| Malignant epithelial | C1 Das ECs | FER | 0.887 | 0.14 | 0.082 | 9.96e-10 | 3.36e-05 |
| Malignant epithelial | C1 Das ECs | LGMN | 0.885 | 0.412 | 0.25 | 3.31e-31 | 1.11e-26 |
| Malignant epithelial | C1 Das ECs | EBLN3 | 0.882 | 0.389 | 0.251 | 4.54e-24 | 1.53e-19 |
| Malignant epithelial | C1 Das ECs | FAM105A | 0.881 | 0.136 | 0.063 | 7.18e-16 | 2.42e-11 |
| Malignant epithelial | C1 Das ECs | RASSF7 | 0.88 | 0.554 | 0.37 | 4.45e-40 | 1.5e-35 |
| Malignant epithelial | C1 Das ECs | GLRX | 0.878 | 0.521 | 0.256 | 2.73e-67 | 9.2e-63 |
| Malignant epithelial | C1 Das ECs | CAPN8 | 0.876 | 0.207 | 0.108 | 1.14e-18 | 3.85e-14 |
| Malignant epithelial | C1 Das ECs | PTPRN2 | 0.875 | 0.173 | 0.102 | 1.3e-11 | 4.37e-07 |
| Malignant epithelial | C1 Das ECs | GTPBP6 | 0.874 | 0.292 | 0.192 | 4.54e-16 | 1.53e-11 |
| Malignant epithelial | C1 Das ECs | GDPD5 | 0.872 | 0.102 | 0.059 | 2.29e-07 | 0.00772 |
| Malignant epithelial | C1 Das ECs | ETS2 | 0.871 | 0.523 | 0.342 | 1.71e-38 | 5.75e-34 |
| Malignant epithelial | C1 Das ECs | ADAM15 | 0.871 | 0.309 | 0.201 | 3.54e-18 | 1.19e-13 |
| Malignant epithelial | C1 Das ECs | CLMN | 0.871 | 0.172 | 0.089 | 8.42e-16 | 2.84e-11 |
| Malignant epithelial | C1 Das ECs | ASL | 0.87 | 0.514 | 0.339 | 3.21e-36 | 1.08e-31 |
| Malignant epithelial | C1 Das ECs | SYNE2 | 0.87 | 0.378 | 0.244 | 1.96e-22 | 6.61e-18 |
| Malignant epithelial | C1 Das ECs | TMEM51 | 0.869 | 0.159 | 0.094 | 1.02e-10 | 3.42e-06 |
| Malignant epithelial | C1 Das ECs | TMEM141 | 0.868 | 0.89 | 0.777 | 3.75e-88 | 1.26e-83 |
| Malignant epithelial | C1 Das ECs | GALM | 0.867 | 0.216 | 0.111 | 1.03e-20 | 3.47e-16 |

| compartment | subcluster | gene | avg_log2FC | pct.1 | pct.2 | p_val | p_val_adj |
|---|---|---|---|---|---|---|---|
| Malignant epithelial | C1 Das ECs | FAM120A | 0.863 | 0.516 | 0.354 | 7.04e-36 | 2.37e-31 |
| Malignant epithelial | C1 Das ECs | CCDC34 | 0.863 | 0.471 | 0.296 | 4.17e-34 | 1.4e-29 |
| Malignant epithelial | C1 Das ECs | BAG1 | 0.862 | 0.733 | 0.548 | 1.29e-59 | 4.34e-55 |
| Malignant epithelial | C1 Das ECs | RNF113A | 0.862 | 0.349 | 0.223 | 1.52e-21 | 5.11e-17 |
| Malignant epithelial | C1 Das ECs | CBR3 | 0.862 | 0.199 | 0.11 | 1.26e-15 | 4.23e-11 |
| Malignant epithelial | C1 Das ECs | REEP6 | 0.861 | 0.218 | 0.147 | 2.03e-10 | 6.84e-06 |
| Malignant epithelial | C1 Das ECs | ENTPD5 | 0.861 | 0.1 | 0.056 | 5.94e-08 | 0.002 |
| Malignant epithelial | C1 Das ECs | RP11-111M22.3 | 0.861 | 0.138 | 0.087 | 1.04e-07 | 0.0035 |
| Malignant epithelial | C1 Das ECs | TSTD1 | 0.858 | 0.809 | 0.623 | 3.8e-76 | 1.28e-71 |
| Malignant epithelial | C1 Das ECs | ZNF703 | 0.857 | 0.696 | 0.456 | 8.67e-61 | 2.92e-56 |
| Malignant epithelial | C1 Das ECs | PDLIM1 | 0.856 | 0.806 | 0.563 | 5.71e-85 | 1.92e-80 |
| Malignant epithelial | C1 Das ECs | IL10RB | 0.856 | 0.356 | 0.219 | 7.52e-25 | 2.53e-20 |
| Malignant epithelial | C1 Das ECs | TMEM63A | 0.855 | 0.386 | 0.238 | 3.25e-26 | 1.1e-21 |
| Malignant epithelial | C1 Das ECs | APOL3 | 0.855 | 0.128 | 0.081 | 4.69e-07 | 0.0158 |
| Malignant epithelial | C1 Das ECs | SECISBP2 | 0.854 | 0.382 | 0.249 | 1.33e-22 | 4.48e-18 |
| Malignant epithelial | C1 Das ECs | IDH3G | 0.852 | 0.589 | 0.407 | 3.42e-41 | 1.15e-36 |
| Malignant epithelial | C1 Das ECs | SLC12A9 | 0.85 | 0.282 | 0.168 | 8.44e-20 | 2.84e-15 |
| Malignant epithelial | C1 Das ECs | TLE4 | 0.849 | 0.228 | 0.119 | 3.17e-21 | 1.07e-16 |
| Malignant epithelial | C1 Das ECs | SCPEP1 | 0.847 | 0.114 | 0.069 | 3.27e-07 | 0.011 |
| Malignant epithelial | C1 Das ECs | TRIM22 | 0.847 | 0.129 | 0.083 | 9.18e-07 | 0.0309 |
| Malignant epithelial | C1 Das ECs | CLINT1 | 0.846 | 0.345 | 0.218 | 2.38e-21 | 8.01e-17 |
| Malignant epithelial | C1 Das ECs | C5orf56 | 0.846 | 0.195 | 0.124 | 1.09e-10 | 3.67e-06 |
| Malignant epithelial | C1 Das ECs | MYO1F | 0.846 | 0.102 | 0.057 | 6.8e-08 | 0.00229 |
| Malignant epithelial | C1 Das ECs | CEACAM6 | 0.844 | 0.746 | 0.334 | 8.6e-138 | 2.9e-133 |
| Malignant epithelial | C1 Das ECs | CCDC22 | 0.844 | 0.174 | 0.11 | 1.15e-09 | 3.87e-05 |
| Malignant epithelial | C1 Das ECs | C1GALT1C1 | 0.843 | 0.289 | 0.188 | 2.97e-16 | 1e-11 |
| Malignant epithelial | C1 Das ECs | AKR1A1 | 0.842 | 0.494 | 0.337 | 2.22e-31 | 7.46e-27 |
| Malignant epithelial | C1 Das ECs | RGS3 | 0.841 | 0.148 | 0.083 | 2.96e-11 | 9.98e-07 |
| Malignant epithelial | C1 Das ECs | PLEKHA5 | 0.84 | 0.323 | 0.218 | 1.28e-16 | 4.31e-12 |
| Malignant epithelial | C1 Das ECs | CYP2S1 | 0.84 | 0.25 | 0.159 | 8.08e-15 | 2.72e-10 |
| Malignant epithelial | C1 Das ECs | TMEM168 | 0.84 | 0.119 | 0.07 | 5.04e-08 | 0.0017 |
| Malignant epithelial | C1 Das ECs | TRAK1 | 0.839 | 0.243 | 0.154 | 5.25e-14 | 1.77e-09 |
| Malignant epithelial | C1 Das ECs | RPS4X | 0.835 | 1.0 | 0.997 | 1.74e-215 | 5.85e-211 |
| Malignant epithelial | C1 Das ECs | APEX2 | 0.835 | 0.154 | 0.095 | 2.55e-09 | 8.59e-05 |
| Malignant epithelial | C1 Das ECs | TMEM106C | 0.834 | 0.467 | 0.294 | 4.13e-34 | 1.39e-29 |
| Malignant epithelial | C1 Das ECs | EXD3 | 0.834 | 0.188 | 0.115 | 1.04e-11 | 3.52e-07 |
| Malignant epithelial | C1 Das ECs | PIGH | 0.834 | 0.205 | 0.131 | 3.49e-11 | 1.18e-06 |
| Malignant epithelial | C1 Das ECs | DOLPP1 | 0.833 | 0.136 | 0.078 | 1e-09 | 3.38e-05 |
| Malignant epithelial | C1 Das ECs | DHRS4 | 0.832 | 0.254 | 0.162 | 1.72e-14 | 5.81e-10 |
| Malignant epithelial | C1 Das ECs | SUSD6 | 0.832 | 0.151 | 0.098 | 9.87e-08 | 0.00332 |
| Malignant epithelial | C1 Das ECs | TRIM31 | 0.831 | 0.515 | 0.312 | 7.67e-40 | 2.58e-35 |
| Malignant epithelial | C1 Das ECs | VEGFA | 0.831 | 0.494 | 0.311 | 1.13e-34 | 3.8199999999999996e-30 |
| Malignant epithelial | C1 Das ECs | MARVELD2 | 0.83 | 0.18 | 0.119 | 9.11e-09 | 0.000307 |
| Malignant epithelial | C1 Das ECs | SNHG21 | 0.83 | 0.132 | 0.082 | 1.38e-07 | 0.00465 |
| Malignant epithelial | C1 Das ECs | DNLZ | 0.83 | 0.1 | 0.058 | 3.95e-07 | 0.0133 |
| Malignant epithelial | C1 Das ECs | EXOC6 | 0.83 | 0.131 | 0.086 | 1.45e-06 | 0.0488 |
| Malignant epithelial | C1 Das ECs | HMGN3 | 0.829 | 0.452 | 0.297 | 2.63e-27 | 8.849999999999999e-23 |
| Malignant epithelial | C1 Das ECs | URM1 | 0.828 | 0.51 | 0.355 | 4.6e-32 | 1.55e-27 |
| Malignant epithelial | C1 Das ECs | RRBP1 | 0.826 | 0.826 | 0.599 | 3.05e-84 | 1.03e-79 |
| Malignant epithelial | C1 Das ECs | CARD16 | 0.822 | 0.282 | 0.174 | 5.56e-18 | 1.87e-13 |
| Malignant epithelial | C1 Das ECs | SSR4 | 0.82 | 0.973 | 0.908 | 9.19e-131 | 3.1e-126 |
| Malignant epithelial | C1 Das ECs | NR4A1 | 0.82 | 0.802 | 0.623 | 1.22e-62 | 4.11e-58 |
| Malignant epithelial | C1 Das ECs | NOXA1 | 0.819 | 0.153 | 0.095 | 7.25e-09 | 0.000244 |
| Malignant epithelial | C1 Das ECs | SDHAF3 | 0.818 | 0.395 | 0.274 | 8.44e-20 | 2.84e-15 |
| Malignant epithelial | C1 Das ECs | CDHR5 | 0.818 | 0.253 | 0.151 | 4.19e-17 | 1.41e-12 |
| Malignant epithelial | C1 Das ECs | SLC6A14 | 0.817 | 0.114 | 0.06 | 6.5e-10 | 2.19e-05 |
| Malignant epithelial | C1 Das ECs | RPP25L | 0.815 | 0.465 | 0.312 | 2.66e-28 | 8.95e-24 |
| Malignant epithelial | C1 Das ECs | ADCK2 | 0.815 | 0.174 | 0.109 | 2.78e-10 | 9.35e-06 |
| Malignant epithelial | C1 Das ECs | KBTBD3 | 0.814 | 0.142 | 0.091 | 1.06e-07 | 0.00358 |
| Malignant epithelial | C1 Das ECs | RFFL | 0.813 | 0.151 | 0.089 | 3.31e-10 | 1.12e-05 |
| Malignant epithelial | C1 Das ECs | ERVK3-1 | 0.813 | 0.193 | 0.124 | 3.49e-10 | 1.17e-05 |
| Malignant epithelial | C1 Das ECs | ABCB7 | 0.813 | 0.185 | 0.129 | 1.41e-07 | 0.00476 |
| Malignant epithelial | C1 Das ECs | NSMCE4A | 0.812 | 0.374 | 0.249 | 7.21e-21 | 2.43e-16 |
| Malignant epithelial | C1 Das ECs | PDCD2 | 0.811 | 0.496 | 0.326 | 2.47e-34 | 8.33e-30 |

| compartment | subcluster | gene | avg_log2FC | pct.1 | pct.2 | p_val | p_val_adj |
|---|---|---|---|---|---|---|---|
| Malignant epithelial | C1 Das ECs | MAGED2 | 0.811 | 0.393 | 0.253 | 3.24e-24 | 1.09e-19 |
| Malignant epithelial | C1 Das ECs | S100P | 0.808 | 0.841 | 0.474 | 2.83e-110 | 9.53e-106 |
| Malignant epithelial | C1 Das ECs | EML4 | 0.806 | 0.413 | 0.266 | 3.18e-26 | 1.07e-21 |
| Malignant epithelial | C1 Das ECs | RBMX2 | 0.806 | 0.409 | 0.289 | 1.55e-20 | 5.23e-16 |
| Malignant epithelial | C1 Das ECs | EPHB2 | 0.806 | 0.213 | 0.138 | 5.26e-11 | 1.77e-06 |
| Malignant epithelial | C1 Das ECs | IGLC3 | 0.805 | 0.322 | 0.205 | 3.07e-18 | 1.04e-13 |
| Malignant epithelial | C1 Das ECs | SH3D19 | 0.805 | 0.202 | 0.129 | 6.81e-11 | 2.3e-06 |
| Malignant epithelial | C1 Das ECs | FAM104B | 0.804 | 0.299 | 0.188 | 3.58e-18 | 1.2e-13 |
| Malignant epithelial | C1 Das ECs | MCEE | 0.804 | 0.175 | 0.11 | 5.55e-10 | 1.87e-05 |
| Malignant epithelial | C1 Das ECs | GSTK1 | 0.801 | 0.845 | 0.682 | 1.63e-73 | 5.49e-69 |
| Malignant epithelial | C1 Das ECs | ALDH1A1 | 0.801 | 0.132 | 0.062 | 1.4e-14 | 4.73e-10 |
| Malignant epithelial | C1 Das ECs | WWC3 | 0.8 | 0.119 | 0.066 | 2.23e-09 | 7.51e-05 |
| Malignant epithelial | C1 Das ECs | PLAGL2 | 0.799 | 0.141 | 0.083 | 2.76e-09 | 9.3e-05 |
| Malignant epithelial | C1 Das ECs | IGLC2 | 0.798 | 0.686 | 0.511 | 2.11e-42 | 7.1e-38 |
| Malignant epithelial | C1 Das ECs | AKR1C3 | 0.798 | 0.424 | 0.274 | 6.55e-25 | 2.21e-20 |
| Malignant epithelial | C1 Das ECs | GABRE | 0.797 | 0.235 | 0.143 | 1.44e-14 | 4.84e-10 |
| Malignant epithelial | C1 Das ECs | NR1D2 | 0.797 | 0.171 | 0.119 | 4.5e-07 | 0.0152 |
| Malignant epithelial | C1 Das ECs | LIMK2 | 0.796 | 0.116 | 0.074 | 1.45e-06 | 0.0489 |
| Malignant epithelial | C1 Das ECs | ATP8B1 | 0.795 | 0.224 | 0.135 | 2.14e-14 | 7.22e-10 |
| Malignant epithelial | C1 Das ECs | EFNA3 | 0.793 | 0.281 | 0.166 | 1.52e-19 | 5.11e-15 |
| Malignant epithelial | C1 Das ECs | NEBL | 0.792 | 0.165 | 0.107 | 1.69e-08 | 0.000568 |
| Malignant epithelial | C1 Das ECs | PLEKHB1 | 0.792 | 0.14 | 0.086 | 3.15e-08 | 0.00106 |
| Malignant epithelial | C1 Das ECs | AP1S1 | 0.791 | 0.707 | 0.538 | 6.180000000000001e-47 | 2.08e-42 |
| Malignant epithelial | C1 Das ECs | CRLS1 | 0.791 | 0.523 | 0.351 | 7.44e-34 | 2.51e-29 |
| Malignant epithelial | C1 Das ECs | SMAD5 | 0.791 | 0.194 | 0.126 | 7.91e-10 | 2.67e-05 |
| Malignant epithelial | C1 Das ECs | CLCN3 | 0.789 | 0.286 | 0.184 | 6.18e-16 | 2.08e-11 |
| Malignant epithelial | C1 Das ECs | GPRC5C | 0.787 | 0.348 | 0.215 | 4.62e-22 | 1.56e-17 |
| Malignant epithelial | C1 Das ECs | VPS13A | 0.787 | 0.236 | 0.149 | 2e-13 | 6.74e-09 |
| Malignant epithelial | C1 Das ECs | PP7080 | 0.787 | 0.167 | 0.114 | 3.89e-07 | 0.0131 |
| Malignant epithelial | C1 Das ECs | IGHA2 | 0.785 | 0.324 | 0.193 | 2.59e-22 | 8.72e-18 |
| Malignant epithelial | C1 Das ECs | UTRN | 0.785 | 0.146 | 0.097 | 7.53e-07 | 0.0254 |
| Malignant epithelial | C1 Das ECs | RPL10 | 0.784 | 0.999 | 1.0 | 7.230000000000001e-213 | 2.44e-208 |
| Malignant epithelial | C1 Das ECs | CDC26 | 0.784 | 0.546 | 0.366 | 1.49e-37 | 5.02e-33 |
| Malignant epithelial | C1 Das ECs | TDP2 | 0.783 | 0.18 | 0.121 | 3.16e-08 | 0.00107 |
| Malignant epithelial | C1 Das ECs | YAE1D1 | 0.782 | 0.26 | 0.169 | 8.36e-14 | 2.82e-09 |
| Malignant epithelial | C1 Das ECs | ST3GAL1 | 0.782 | 0.198 | 0.125 | 1.06e-10 | 3.57e-06 |
| Malignant epithelial | C1 Das ECs | SPPL2B | 0.78 | 0.268 | 0.175 | 1.85e-14 | 6.24e-10 |
| Malignant epithelial | C1 Das ECs | ILVBL | 0.779 | 0.365 | 0.24 | 1.67e-20 | 5.63e-16 |
| Malignant epithelial | C1 Das ECs | DAB2 | 0.779 | 0.151 | 0.066 | 3.66e-19 | 1.23e-14 |
| Malignant epithelial | C1 Das ECs | ARSD | 0.779 | 0.33 | 0.212 | 1.15e-18 | 3.88e-14 |
| Malignant epithelial | C1 Das ECs | CLN3 | 0.779 | 0.325 | 0.211 | 1.22e-17 | 4.11e-13 |
| Malignant epithelial | C1 Das ECs | RP5-1159O4.1 | 0.779 | 0.117 | 0.071 | 1.92e-07 | 0.00647 |
| Malignant epithelial | C1 Das ECs | MXD4 | 0.777 | 0.173 | 0.108 | 6.32e-10 | 2.13e-05 |
| Malignant epithelial | C1 Das ECs | GALNT7 | 0.775 | 0.173 | 0.111 | 3.08e-09 | 0.000104 |
| Malignant epithelial | C1 Das ECs | SDHAF1 | 0.774 | 0.251 | 0.178 | 1.18e-09 | 3.98e-05 |
| Malignant epithelial | C1 Das ECs | HLA-DPA1 | 0.773 | 0.683 | 0.51 | 1.53e-38 | 5.15e-34 |
| Malignant epithelial | C1 Das ECs | SLC37A1 | 0.773 | 0.229 | 0.144 | 2.88e-13 | 9.7e-09 |
| Malignant epithelial | C1 Das ECs | EFNB1 | 0.772 | 0.333 | 0.194 | 6.93e-25 | 2.34e-20 |
| Malignant epithelial | C1 Das ECs | PREP | 0.772 | 0.23 | 0.149 | 4.83e-12 | 1.63e-07 |
| Malignant epithelial | C1 Das ECs | ATP5G1 | 0.771 | 0.869 | 0.715 | 1.42e-69 | 4.79e-65 |
| Malignant epithelial | C1 Das ECs | FAT1 | 0.771 | 0.27 | 0.175 | 2.55e-14 | 8.58e-10 |
| Malignant epithelial | C1 Das ECs | ARL14 | 0.77 | 0.199 | 0.138 | 2.26e-08 | 0.00076 |
| Malignant epithelial | C1 Das ECs | TRIM24 | 0.767 | 0.167 | 0.102 | 5.16e-10 | 1.74e-05 |
| Malignant epithelial | C1 Das ECs | PUDP | 0.767 | 0.147 | 0.094 | 7.46e-08 | 0.00252 |
| Malignant epithelial | C1 Das ECs | QPCT | 0.766 | 0.291 | 0.185 | 1.14e-16 | 3.85e-12 |
| Malignant epithelial | C1 Das ECs | ERP44 | 0.765 | 0.573 | 0.4 | 7.63e-36 | 2.57e-31 |
| Malignant epithelial | C1 Das ECs | DLD | 0.765 | 0.289 | 0.19 | 3.34e-15 | 1.12e-10 |
| Malignant epithelial | C1 Das ECs | LAIR2 | 0.764 | 0.125 | 0.065 | 1.63e-11 | 5.48e-07 |
| Malignant epithelial | C1 Das ECs | CBX4 | 0.761 | 0.252 | 0.168 | 5.33e-12 | 1.8e-07 |
| Malignant epithelial | C1 Das ECs | TAZ | 0.759 | 0.185 | 0.126 | 5.77e-08 | 0.00194 |
| Malignant epithelial | C1 Das ECs | SLC31A1 | 0.758 | 0.191 | 0.115 | 5.79e-12 | 1.95e-07 |
| Malignant epithelial | C1 Das ECs | CASP3 | 0.757 | 0.161 | 0.109 | 5.55e-07 | 0.0187 |
| Malignant epithelial | C1 Das ECs | FARSB | 0.753 | 0.304 | 0.206 | 2.12e-14 | 7.14e-10 |
| Malignant epithelial | C1 Das ECs | TFDP2 | 0.753 | 0.27 | 0.18 | 7.93e-13 | 2.67e-08 |

| compartment | subcluster | gene | avg_log2FC | pct.1 | pct.2 | p_val | p_val_adj |
|---|---|---|---|---|---|---|---|
| Malignant epithelial | C1 Das ECs | HLA-DPB1 | 0.752 | 0.598 | 0.434 | 8.450000000000001e-31 | 2.85e-26 |
| Malignant epithelial | C1 Das ECs | MGST2 | 0.751 | 0.832 | 0.691 | 2.79e-56 | 9.389999999999999e-52 |
| Malignant epithelial | C1 Das ECs | LGALS9 | 0.751 | 0.433 | 0.283 | 8.01e-25 | 2.7e-20 |
| Malignant epithelial | C1 Das ECs | HADH | 0.75 | 0.326 | 0.208 | 4.76e-19 | 1.6e-14 |
| Malignant epithelial | C1 Das ECs | ALG14 | 0.75 | 0.161 | 0.102 | 6.54e-09 | 0.00022 |
| Malignant epithelial | C1 Das ECs | ERBB2 | 0.749 | 0.376 | 0.254 | 1.58e-18 | 5.33e-14 |
| Malignant epithelial | C1 Das ECs | RP11-166P13.4 | 0.749 | 0.155 | 0.087 | 1.67e-11 | 5.63e-07 |
| Malignant epithelial | C1 Das ECs | MNX1-AS1 | 0.749 | 0.2 | 0.125 | 5e-11 | 1.69e-06 |
| Malignant epithelial | C1 Das ECs | IGHA1 | 0.748 | 0.7 | 0.522 | 2.78e-47 | 9.38e-43 |
| Malignant epithelial | C1 Das ECs | TBC1D1 | 0.748 | 0.123 | 0.076 | 2.15e-07 | 0.00726 |
| Malignant epithelial | C1 Das ECs | VGLL4 | 0.746 | 0.305 | 0.202 | 9.02e-16 | 3.04e-11 |
| Malignant epithelial | C1 Das ECs | ARAF | 0.746 | 0.286 | 0.187 | 5.91e-15 | 1.99e-10 |
| Malignant epithelial | C1 Das ECs | PLCB3 | 0.744 | 0.266 | 0.173 | 5.93e-14 | 2e-09 |
| Malignant epithelial | C1 Das ECs | CFAP97 | 0.744 | 0.27 | 0.182 | 5.98e-13 | 2.01e-08 |
| Malignant epithelial | C1 Das ECs | FNIP2 | 0.744 | 0.145 | 0.089 | 1.26e-08 | 0.000425 |
| Malignant epithelial | C1 Das ECs | RBM47 | 0.743 | 0.293 | 0.202 | 8.99e-13 | 3.03e-08 |
| Malignant epithelial | C1 Das ECs | TMA16 | 0.741 | 0.393 | 0.279 | 2.21e-17 | 7.45e-13 |
| Malignant epithelial | C1 Das ECs | PER2 | 0.741 | 0.272 | 0.178 | 8.66e-14 | 2.92e-09 |
| Malignant epithelial | C1 Das ECs | GALNT3 | 0.739 | 0.371 | 0.242 | 1.28e-21 | 4.33e-17 |
| Malignant epithelial | C1 Das ECs | ALDH9A1 | 0.739 | 0.235 | 0.161 | 2.16e-10 | 7.29e-06 |
| Malignant epithelial | C1 Das ECs | TMEM80 | 0.738 | 0.181 | 0.12 | 1.55e-08 | 0.000521 |
| Malignant epithelial | C1 Das ECs | RASEF | 0.737 | 0.188 | 0.119 | 1.9e-10 | 6.41e-06 |
| Malignant epithelial | C1 Das ECs | CANT1 | 0.736 | 0.298 | 0.204 | 9.84e-14 | 3.32e-09 |
| Malignant epithelial | C1 Das ECs | ELK1 | 0.736 | 0.152 | 0.102 | 8.28e-07 | 0.0279 |
| Malignant epithelial | C1 Das ECs | ZNF717 | 0.736 | 0.11 | 0.068 | 1.16e-06 | 0.0391 |
| Malignant epithelial | C1 Das ECs | NEK3 | 0.735 | 0.136 | 0.086 | 1.41e-07 | 0.00475 |
| Malignant epithelial | C1 Das ECs | IRF8 | 0.734 | 0.202 | 0.11 | 4.39e-16 | 1.48e-11 |
| Malignant epithelial | C1 Das ECs | PLGRKT | 0.732 | 0.49 | 0.341 | 1.34e-27 | 4.5e-23 |
| Malignant epithelial | C1 Das ECs | TMEM97 | 0.732 | 0.335 | 0.225 | 2.1e-16 | 7.08e-12 |
| Malignant epithelial | C1 Das ECs | RIN2 | 0.732 | 0.307 | 0.206 | 3.12e-14 | 1.05e-09 |
| Malignant epithelial | C1 Das ECs | BATF | 0.732 | 0.224 | 0.138 | 1.98e-13 | 6.65e-09 |
| Malignant epithelial | C1 Das ECs | HCCS | 0.731 | 0.296 | 0.2 | 2.16e-14 | 7.29e-10 |
| Malignant epithelial | C1 Das ECs | PCK2 | 0.731 | 0.257 | 0.17 | 2.46e-12 | 8.28e-08 |
| Malignant epithelial | C1 Das ECs | CMTM8 | 0.73 | 0.439 | 0.294 | 2.35e-24 | 7.92e-20 |
| Malignant epithelial | C1 Das ECs | NOL4L | 0.73 | 0.243 | 0.157 | 1.85e-12 | 6.23e-08 |
| Malignant epithelial | C1 Das ECs | TNFSF10 | 0.729 | 0.549 | 0.312 | 9.81e-50 | 3.3e-45 |
| Malignant epithelial | C1 Das ECs | PCYT2 | 0.728 | 0.221 | 0.142 | 1.34e-11 | 4.52e-07 |
| Malignant epithelial | C1 Das ECs | FRG1 | 0.727 | 0.4 | 0.276 | 1.65e-19 | 5.56e-15 |
| Malignant epithelial | C1 Das ECs | CCNG1 | 0.727 | 0.423 | 0.299 | 2.57e-19 | 8.65e-15 |
| Malignant epithelial | C1 Das ECs | SURF1 | 0.726 | 0.474 | 0.349 | 8.04e-22 | 2.71e-17 |
| Malignant epithelial | C1 Das ECs | NADK | 0.726 | 0.225 | 0.158 | 4.53e-09 | 0.000153 |
| Malignant epithelial | C1 Das ECs | TTC32 | 0.726 | 0.17 | 0.11 | 8.53e-09 | 0.000287 |
| Malignant epithelial | C1 Das ECs | NUDT2 | 0.725 | 0.51 | 0.382 | 1.49e-21 | 5.03e-17 |
| Malignant epithelial | C1 Das ECs | RBM18 | 0.725 | 0.219 | 0.15 | 1.65e-09 | 5.55e-05 |
| Malignant epithelial | C1 Das ECs | MDH2 | 0.724 | 0.829 | 0.695 | 6.04e-63 | 2.03e-58 |
| Malignant epithelial | C1 Das ECs | BCAP31 | 0.723 | 0.854 | 0.702 | 1.34e-66 | 4.499999999999999e-62 |
| Malignant epithelial | C1 Das ECs | TSR2 | 0.723 | 0.405 | 0.289 | 7.19e-18 | 2.42e-13 |
| Malignant epithelial | C1 Das ECs | IGHG4 | 0.722 | 0.743 | 0.6 | 6.02e-45 | 2.03e-40 |
| Malignant epithelial | C1 Das ECs | TOR2A | 0.722 | 0.249 | 0.167 | 6.45e-12 | 2.17e-07 |
| Malignant epithelial | C1 Das ECs | GALT | 0.722 | 0.183 | 0.127 | 2.63e-07 | 0.00885 |
| Malignant epithelial | C1 Das ECs | SLC35D2 | 0.721 | 0.308 | 0.223 | 3.73e-12 | 1.26e-07 |
| Malignant epithelial | C1 Das ECs | AHCYL2 | 0.721 | 0.159 | 0.093 | 1.09e-10 | 3.68e-06 |
| Malignant epithelial | C1 Das ECs | TMEM5 | 0.721 | 0.246 | 0.17 | 4.38e-10 | 1.47e-05 |
| Malignant epithelial | C1 Das ECs | TUSC1 | 0.719 | 0.205 | 0.13 | 1.05e-10 | 3.53e-06 |
| Malignant epithelial | C1 Das ECs | VRK3 | 0.718 | 0.267 | 0.188 | 5.84e-11 | 1.97e-06 |
| Malignant epithelial | C1 Das ECs | NADSYN1 | 0.718 | 0.216 | 0.144 | 2.6e-10 | 8.76e-06 |
| Malignant epithelial | C1 Das ECs | CTB-113I20.2 | 0.718 | 0.153 | 0.101 | 3.4e-07 | 0.0115 |
| Malignant epithelial | C1 Das ECs | THRA | 0.717 | 0.27 | 0.189 | 1e-10 | 3.38e-06 |
| Malignant epithelial | C1 Das ECs | LPIN1 | 0.716 | 0.162 | 0.111 | 1.28e-06 | 0.043 |
| Malignant epithelial | C1 Das ECs | MCTS1 | 0.715 | 0.683 | 0.534 | 7.9e-37 | 2.66e-32 |
| Malignant epithelial | C1 Das ECs | PRR5 | 0.715 | 0.207 | 0.149 | 1.84e-07 | 0.00619 |
| Malignant epithelial | C1 Das ECs | GLYCTK | 0.714 | 0.155 | 0.102 | 1.57e-07 | 0.00529 |
| Malignant epithelial | C1 Das ECs | PSMB8-AS1 | 0.712 | 0.236 | 0.165 | 2.2e-09 | 7.42e-05 |
| Malignant epithelial | C1 Das ECs | GSKIP | 0.711 | 0.363 | 0.244 | 6.61e-19 | 2.23e-14 |

| compartment | subcluster | gene | avg_log2FC | pct.1 | pct.2 | p_val | p_val_adj |
|---|---|---|---|---|---|---|---|
| Malignant epithelial | C1 Das ECs | FUT2 | 0.71 | 0.243 | 0.158 | 3.62e-12 | 1.22e-07 |
| Malignant epithelial | C1 Das ECs | RP11-357H14.17 | 0.709 | 0.635 | 0.492 | 1.04e-27 | 3.509999999999999e-23 |
| Malignant epithelial | C1 Das ECs | TSC22D4 | 0.709 | 0.481 | 0.339 | 9.88e-25 | 3.33e-20 |
| Malignant epithelial | C1 Das ECs | SLC9A3R1 | 0.709 | 0.424 | 0.308 | 8.16e-18 | 2.75e-13 |
| Malignant epithelial | C1 Das ECs | VPS13C | 0.708 | 0.166 | 0.109 | 3.66e-08 | 0.00123 |
| Malignant epithelial | C1 Das ECs | ZNF444 | 0.707 | 0.207 | 0.142 | 1.8e-08 | 0.000608 |
| Malignant epithelial | C1 Das ECs | SRI | 0.706 | 0.887 | 0.775 | 3.4e-64 | 1.15e-59 |
| Malignant epithelial | C1 Das ECs | LRRC47 | 0.706 | 0.235 | 0.154 | 1.04e-11 | 3.52e-07 |
| Malignant epithelial | C1 Das ECs | HBEGF | 0.704 | 0.514 | 0.351 | 2.43e-28 | 8.19e-24 |
| Malignant epithelial | C1 Das ECs | SLC35E2B | 0.704 | 0.151 | 0.094 | 1.33e-08 | 0.000447 |
| Malignant epithelial | C1 Das ECs | EPHB3 | 0.704 | 0.154 | 0.101 | 1.18e-07 | 0.00396 |
| Malignant epithelial | C1 Das ECs | GRIPAP1 | 0.703 | 0.216 | 0.148 | 5.71e-09 | 0.000192 |
| Malignant epithelial | C1 Das ECs | H6PD | 0.703 | 0.113 | 0.066 | 1.35e-07 | 0.00456 |
| Malignant epithelial | C1 Das ECs | KRTCAP3 | 0.702 | 0.659 | 0.482 | 3.17e-40 | 1.07e-35 |
| Malignant epithelial | C1 Das ECs | CMBL | 0.701 | 0.386 | 0.27 | 4.58e-17 | 1.54e-12 |
| Malignant epithelial | C1 Das ECs | CRIM1 | 0.701 | 0.196 | 0.132 | 1.34e-08 | 0.000453 |
| Malignant epithelial | C1 Das ECs | IMPA2 | 0.7 | 0.287 | 0.185 | 8.18e-16 | 2.76e-11 |
| Malignant epithelial | C1 Das ECs | NKAP | 0.7 | 0.363 | 0.264 | 4.12e-14 | 1.39e-09 |
| Malignant epithelial | C1 Das ECs | CREB3L2 | 0.7 | 0.22 | 0.134 | 2.32e-13 | 7.81e-09 |
| Malignant epithelial | C1 Das ECs | CAAP1 | 0.7 | 0.173 | 0.107 | 6.6e-10 | 2.23e-05 |
| Malignant epithelial | C1 Das ECs | SHPRH | 0.7 | 0.118 | 0.072 | 2.86e-07 | 0.00963 |
| Malignant epithelial | C1 Das ECs | ALDH18A1 | 0.699 | 0.19 | 0.126 | 7.23e-09 | 0.000244 |
| Malignant epithelial | C1 Das ECs | PHF6 | 0.698 | 0.17 | 0.119 | 1.04e-06 | 0.0351 |
| Malignant epithelial | C1 Das ECs | MARVELD3 | 0.697 | 0.397 | 0.269 | 8.6e-20 | 2.9e-15 |
| Malignant epithelial | C1 Das ECs | MLEC | 0.696 | 0.668 | 0.506 | 2.24e-34 | 7.549999999999999e-30 |
| Malignant epithelial | C1 Das ECs | FCGRT | 0.694 | 0.735 | 0.506 | 4.92e-48 | 1.66e-43 |
| Malignant epithelial | C1 Das ECs | C7orf55 | 0.693 | 0.6 | 0.447 | 4.43e-29 | 1.49e-24 |
| Malignant epithelial | C1 Das ECs | PDCD4 | 0.692 | 0.342 | 0.236 | 1.12e-15 | 3.77e-11 |
| Malignant epithelial | C1 Das ECs | CDK11B | 0.692 | 0.276 | 0.185 | 2.51e-13 | 8.46e-09 |
| Malignant epithelial | C1 Das ECs | SLC37A3 | 0.692 | 0.201 | 0.144 | 4.38e-07 | 0.0148 |
| Malignant epithelial | C1 Das ECs | TSPAN6 | 0.691 | 0.465 | 0.32 | 3.41e-24 | 1.15e-19 |
| Malignant epithelial | C1 Das ECs | DKC1 | 0.691 | 0.477 | 0.361 | 5.08e-18 | 1.71e-13 |
| Malignant epithelial | C1 Das ECs | SSBP3 | 0.691 | 0.181 | 0.115 | 6.78e-10 | 2.28e-05 |
| Malignant epithelial | C1 Das ECs | PAG1 | 0.689 | 0.124 | 0.071 | 5.06e-09 | 0.00017 |
| Malignant epithelial | C1 Das ECs | TMEM125 | 0.689 | 0.188 | 0.123 | 8e-09 | 0.000269 |
| Malignant epithelial | C1 Das ECs | CDK11A | 0.688 | 0.188 | 0.13 | 1.39e-07 | 0.0047 |
| Malignant epithelial | C1 Das ECs | EPSTI1 | 0.686 | 0.71 | 0.546 | 1.11e-39 | 3.75e-35 |
| Malignant epithelial | C1 Das ECs | FGD5-AS1 | 0.686 | 0.251 | 0.188 | 3.81e-08 | 0.00128 |
| Malignant epithelial | C1 Das ECs | FRK | 0.686 | 0.129 | 0.083 | 1.35e-06 | 0.0455 |
| Malignant epithelial | C1 Das ECs | ATP1B1 | 0.685 | 0.747 | 0.608 | 1.45e-35 | 4.89e-31 |
| Malignant epithelial | C1 Das ECs | NBL1 | 0.685 | 0.589 | 0.43 | 1.23e-29 | 4.13e-25 |
| Malignant epithelial | C1 Das ECs | DUSP10 | 0.685 | 0.196 | 0.128 | 1.06e-09 | 3.56e-05 |
| Malignant epithelial | C1 Das ECs | ZKSCAN1 | 0.682 | 0.427 | 0.283 | 2.2e-23 | 7.42e-19 |
| Malignant epithelial | C1 Das ECs | ATP6AP1 | 0.682 | 0.501 | 0.367 | 3.09e-23 | 1.04e-18 |
| Malignant epithelial | C1 Das ECs | TM9SF3 | 0.681 | 0.67 | 0.519 | 5.82e-34 | 1.96e-29 |
| Malignant epithelial | C1 Das ECs | EPB41L2 | 0.681 | 0.252 | 0.172 | 2.02e-10 | 6.79e-06 |
| Malignant epithelial | C1 Das ECs | RNF24 | 0.68 | 0.14 | 0.087 | 1.02e-07 | 0.00343 |
| Malignant epithelial | C1 Das ECs | TMPRSS3 | 0.678 | 0.253 | 0.191 | 1.48e-07 | 0.005 |
| Malignant epithelial | C1 Das ECs | PDHA1 | 0.677 | 0.431 | 0.302 | 6.180000000000001e-21 | 2.08e-16 |
| Malignant epithelial | C1 Das ECs | PBDC1 | 0.676 | 0.429 | 0.317 | 1.46e-16 | 4.93e-12 |
| Malignant epithelial | C1 Das ECs | DHRSX | 0.676 | 0.24 | 0.172 | 1.12e-08 | 0.000379 |
| Malignant epithelial | C1 Das ECs | SLC25A38 | 0.676 | 0.243 | 0.175 | 1.59e-08 | 0.000534 |
| Malignant epithelial | C1 Das ECs | ORC5 | 0.676 | 0.139 | 0.089 | 2.56e-07 | 0.00861 |
| Malignant epithelial | C1 Das ECs | CCDC64B | 0.675 | 0.418 | 0.27 | 1.01e-23 | 3.41e-19 |
| Malignant epithelial | C1 Das ECs | SDSL | 0.675 | 0.207 | 0.142 | 1.56e-08 | 0.000525 |
| Malignant epithelial | C1 Das ECs | NOC2L | 0.674 | 0.364 | 0.257 | 3.11e-15 | 1.05e-10 |
| Malignant epithelial | C1 Das ECs | TULP4 | 0.674 | 0.169 | 0.115 | 2.72e-07 | 0.00915 |
| Malignant epithelial | C1 Das ECs | RPIA | 0.673 | 0.404 | 0.277 | 3.11e-19 | 1.05e-14 |
| Malignant epithelial | C1 Das ECs | DHRS11 | 0.672 | 0.162 | 0.11 | 1.06e-06 | 0.0357 |
| Malignant epithelial | C1 Das ECs | POM121C | 0.671 | 0.174 | 0.121 | 6.25e-07 | 0.0211 |
| Malignant epithelial | C1 Das ECs | CC2D1A | 0.669 | 0.219 | 0.155 | 3.92e-08 | 0.00132 |
| Malignant epithelial | C1 Das ECs | GPKOW | 0.668 | 0.207 | 0.15 | 4.09e-07 | 0.0138 |
| Malignant epithelial | C1 Das ECs | SLC37A4 | 0.668 | 0.216 | 0.161 | 1.21e-06 | 0.0408 |
| Malignant epithelial | C1 Das ECs | HIP1R | 0.668 | 0.164 | 0.114 | 1.47e-06 | 0.0495 |

| compartment | subcluster | gene | avg_log2FC | pct.1 | pct.2 | p_val | p_val_adj |
|---|---|---|---|---|---|---|---|
| Malignant epithelial | C1 Das ECs | UPF3B | 0.667 | 0.319 | 0.222 | 6.02e-14 | 2.03e-09 |
| Malignant epithelial | C1 Das ECs | ARHGEF16 | 0.667 | 0.199 | 0.136 | 3.61e-08 | 0.00122 |
| Malignant epithelial | C1 Das ECs | TMEM261 | 0.666 | 0.76 | 0.611 | 3.99e-42 | 1.35e-37 |
| Malignant epithelial | C1 Das ECs | CHPT1 | 0.666 | 0.363 | 0.229 | 5.180000000000001e-21 | 1.74e-16 |
| Malignant epithelial | C1 Das ECs | TRIM54 | 0.666 | 0.4 | 0.273 | 3.52e-19 | 1.19e-14 |
| Malignant epithelial | C1 Das ECs | CBWD1 | 0.666 | 0.417 | 0.316 | 3.5e-14 | 1.18e-09 |
| Malignant epithelial | C1 Das ECs | CBWD5 | 0.665 | 0.422 | 0.321 | 8.01e-14 | 2.7e-09 |
| Malignant epithelial | C1 Das ECs | TOPORS-AS1 | 0.665 | 0.212 | 0.148 | 2.81e-08 | 0.000947 |
| Malignant epithelial | C1 Das ECs | ATP2C2 | 0.665 | 0.132 | 0.082 | 8.22e-08 | 0.00277 |
| Malignant epithelial | C1 Das ECs | LAMTOR4 | 0.664 | 0.908 | 0.796 | 8.24e-76 | 2.77e-71 |
| Malignant epithelial | C1 Das ECs | MUC12 | 0.663 | 0.322 | 0.188 | 4.07e-24 | 1.37e-19 |
| Malignant epithelial | C1 Das ECs | ZBTB7A | 0.663 | 0.485 | 0.351 | 9.87e-22 | 3.32e-17 |
| Malignant epithelial | C1 Das ECs | REXO4 | 0.661 | 0.273 | 0.195 | 4.98e-10 | 1.68e-05 |
| Malignant epithelial | C1 Das ECs | TPRG1L | 0.661 | 0.177 | 0.111 | 1.01e-09 | 3.41e-05 |
| Malignant epithelial | C1 Das ECs | DUSP18 | 0.661 | 0.171 | 0.116 | 2.27e-07 | 0.00764 |
| Malignant epithelial | C1 Das ECs | POLD2 | 0.66 | 0.522 | 0.39 | 4.3e-24 | 1.45e-19 |
| Malignant epithelial | C1 Das ECs | CXorf40B | 0.66 | 0.161 | 0.107 | 4.06e-07 | 0.0137 |
| Malignant epithelial | C1 Das ECs | IGKC | 0.659 | 0.934 | 0.87 | 1.31e-79 | 4.43e-75 |
| Malignant epithelial | C1 Das ECs | THEM6 | 0.658 | 0.302 | 0.213 | 1.35e-11 | 4.53e-07 |
| Malignant epithelial | C1 Das ECs | JAGN1 | 0.657 | 0.413 | 0.295 | 1.45e-17 | 4.87e-13 |
| Malignant epithelial | C1 Das ECs | RNASEH2B | 0.657 | 0.261 | 0.178 | 4.42e-11 | 1.49e-06 |
| Malignant epithelial | C1 Das ECs | GRTP1 | 0.657 | 0.242 | 0.17 | 3.86e-09 | 0.00013 |
| Malignant epithelial | C1 Das ECs | MT-CYB | 0.656 | 1.0 | 0.981 | 4.28e-118 | 1.44e-113 |
| Malignant epithelial | C1 Das ECs | HLA-DRB1 | 0.656 | 0.744 | 0.58 | 9.95e-38 | 3.35e-33 |
| Malignant epithelial | C1 Das ECs | CCDC28A | 0.656 | 0.177 | 0.12 | 2.01e-07 | 0.00676 |
| Malignant epithelial | C1 Das ECs | UXT | 0.655 | 0.878 | 0.778 | 4.04e-67 | 1.36e-62 |
| Malignant epithelial | C1 Das ECs | EEF1G | 0.655 | 0.49 | 0.38 | 3.37e-19 | 1.14e-14 |
| Malignant epithelial | C1 Das ECs | POLQ | 0.654 | 0.143 | 0.094 | 8.66e-07 | 0.0292 |
| Malignant epithelial | C1 Das ECs | WBP5 | 0.653 | 0.546 | 0.39 | 6.82e-28 | 2.3e-23 |
| Malignant epithelial | C1 Das ECs | C9orf114 | 0.653 | 0.209 | 0.155 | 1.06e-06 | 0.0357 |
| Malignant epithelial | C1 Das ECs | ANXA4 | 0.651 | 0.695 | 0.553 | 1.02e-29 | 3.45e-25 |
| Malignant epithelial | C1 Das ECs | PLEKHJ1 | 0.651 | 0.604 | 0.473 | 4.08e-27 | 1.37e-22 |
| Malignant epithelial | C1 Das ECs | CMIP | 0.649 | 0.276 | 0.201 | 2.17e-09 | 7.3e-05 |
| Malignant epithelial | C1 Das ECs | UBE2J2 | 0.648 | 0.303 | 0.222 | 8.26e-11 | 2.78e-06 |
| Malignant epithelial | C1 Das ECs | UBL4A | 0.646 | 0.238 | 0.173 | 4.19e-08 | 0.00141 |
| Malignant epithelial | C1 Das ECs | ICA1 | 0.645 | 0.553 | 0.399 | 1.32e-27 | 4.45e-23 |
| Malignant epithelial | C1 Das ECs | CYB5A | 0.644 | 0.581 | 0.424 | 2.46e-28 | 8.28e-24 |
| Malignant epithelial | C1 Das ECs | CDC42SE2 | 0.644 | 0.384 | 0.279 | 1.25e-14 | 4.21e-10 |
| Malignant epithelial | C1 Das ECs | SDF4 | 0.643 | 0.58 | 0.425 | 6.300000000000001e-28 | 2.12e-23 |
| Malignant epithelial | C1 Das ECs | PDXDC1 | 0.642 | 0.331 | 0.229 | 2.38e-14 | 8.02e-10 |
| Malignant epithelial | C1 Das ECs | FAM195A | 0.641 | 0.611 | 0.461 | 7.67e-29 | 2.59e-24 |
| Malignant epithelial | C1 Das ECs | PSMB10 | 0.64 | 0.57 | 0.414 | 6.31e-27 | 2.13e-22 |
| Malignant epithelial | C1 Das ECs | HPRT1 | 0.64 | 0.496 | 0.382 | 1.54e-18 | 5.18e-14 |
| Malignant epithelial | C1 Das ECs | ABO | 0.64 | 0.22 | 0.151 | 6.65e-09 | 0.000224 |
| Malignant epithelial | C1 Das ECs | INSR | 0.639 | 0.274 | 0.199 | 4.31e-09 | 0.000145 |
| Malignant epithelial | C1 Das ECs | FUNDC2 | 0.638 | 0.47 | 0.35 | 2.47e-19 | 8.31e-15 |
| Malignant epithelial | C1 Das ECs | FAM58A | 0.638 | 0.315 | 0.224 | 4.72e-12 | 1.59e-07 |
| Malignant epithelial | C1 Das ECs | PDZK1IP1 | 0.636 | 0.733 | 0.477 | 1.84e-53 | 6.21e-49 |
| Malignant epithelial | C1 Das ECs | PIN4 | 0.636 | 0.713 | 0.579 | 8.63e-34 | 2.91e-29 |
| Malignant epithelial | C1 Das ECs | PRDX4 | 0.635 | 0.74 | 0.582 | 1.68e-38 | 5.669999999999999e-34 |
| Malignant epithelial | C1 Das ECs | TMPRSS4 | 0.635 | 0.424 | 0.27 | 1.01e-24 | 3.41e-20 |
| Malignant epithelial | C1 Das ECs | PLPP5 | 0.635 | 0.459 | 0.333 | 8.15e-20 | 2.74e-15 |
| Malignant epithelial | C1 Das ECs | C9orf3 | 0.635 | 0.385 | 0.291 | 1.46e-12 | 4.93e-08 |
| Malignant epithelial | C1 Das ECs | EBAG9 | 0.634 | 0.49 | 0.356 | 1.38e-20 | 4.64e-16 |
| Malignant epithelial | C1 Das ECs | TFAP4 | 0.634 | 0.176 | 0.121 | 4.15e-07 | 0.014 |
| Malignant epithelial | C1 Das ECs | MMGT1 | 0.634 | 0.18 | 0.127 | 1.05e-06 | 0.0355 |
| Malignant epithelial | C1 Das ECs | DPM2 | 0.633 | 0.572 | 0.429 | 4.6e-26 | 1.55e-21 |
| Malignant epithelial | C1 Das ECs | CCNL2 | 0.633 | 0.333 | 0.227 | 2.07e-15 | 6.97e-11 |
| Malignant epithelial | C1 Das ECs | SCRN2 | 0.633 | 0.244 | 0.179 | 1.16e-07 | 0.00391 |
| Malignant epithelial | C1 Das ECs | RABEPK | 0.633 | 0.196 | 0.14 | 4.05e-07 | 0.0137 |
| Malignant epithelial | C1 Das ECs | NFX1 | 0.633 | 0.162 | 0.111 | 1.28e-06 | 0.0431 |
| Malignant epithelial | C1 Das ECs | PEX16 | 0.632 | 0.384 | 0.286 | 2.09e-13 | 7.05e-09 |
| Malignant epithelial | C1 Das ECs | NMRK1 | 0.632 | 0.319 | 0.227 | 5.34e-12 | 1.8e-07 |
| Malignant epithelial | C1 Das ECs | SURF6 | 0.632 | 0.31 | 0.226 | 6.75e-11 | 2.27e-06 |

| compartment | subcluster | gene | avg_log2FC | pct.1 | pct.2 | p_val | p_val_adj |
|---|---|---|---|---|---|---|---|
| Malignant epithelial | C1 Das ECs | SH3RF2 | 0.63 | 0.16 | 0.105 | 2.01e-07 | 0.00677 |
| Malignant epithelial | C1 Das ECs | VAMP3 | 0.629 | 0.44 | 0.33 | 9.87e-16 | 3.33e-11 |
| Malignant epithelial | C1 Das ECs | BORCS7 | 0.628 | 0.31 | 0.22 | 2.84e-12 | 9.58e-08 |
| Malignant epithelial | C1 Das ECs | ASB9 | 0.627 | 0.134 | 0.086 | 1.08e-06 | 0.0362 |
| Malignant epithelial | C1 Das ECs | CD44 | 0.626 | 0.693 | 0.537 | 1.7e-33 | 5.710000000000001e-29 |
| Malignant epithelial | C1 Das ECs | PDZD8 | 0.626 | 0.352 | 0.264 | 1.09e-10 | 3.66e-06 |
| Malignant epithelial | C1 Das ECs | CBR1 | 0.625 | 0.45 | 0.33 | 2.71e-17 | 9.12e-13 |
| Malignant epithelial | C1 Das ECs | TBL1X | 0.624 | 0.129 | 0.082 | 7.11e-07 | 0.024 |
| Malignant epithelial | C1 Das ECs | ACOT7 | 0.623 | 0.298 | 0.197 | 8.81e-15 | 2.97e-10 |
| Malignant epithelial | C1 Das ECs | RILPL2 | 0.622 | 0.227 | 0.165 | 1.35e-07 | 0.00457 |
| Malignant epithelial | C1 Das ECs | NT5DC1 | 0.62 | 0.321 | 0.228 | 1.21e-12 | 4.08e-08 |
| Malignant epithelial | C1 Das ECs | SYT13 | 0.62 | 0.245 | 0.172 | 2.78e-09 | 9.37e-05 |
| Malignant epithelial | C1 Das ECs | CDK10 | 0.619 | 0.354 | 0.271 | 5.53e-11 | 1.86e-06 |
| Malignant epithelial | C1 Das ECs | SMPDL3A | 0.618 | 0.181 | 0.118 | 9.96e-09 | 0.000336 |
| Malignant epithelial | C1 Das ECs | ADD3 | 0.616 | 0.436 | 0.305 | 2.94e-19 | 9.89e-15 |
| Malignant epithelial | C1 Das ECs | LENG9 | 0.616 | 0.182 | 0.114 | 4.36e-10 | 1.47e-05 |
| Malignant epithelial | C1 Das ECs | PSMD10 | 0.616 | 0.194 | 0.14 | 1.19e-06 | 0.0401 |
| Malignant epithelial | C1 Das ECs | LINC01184 | 0.615 | 0.215 | 0.157 | 4.79e-07 | 0.0161 |
| Malignant epithelial | C1 Das ECs | KIAA2026 | 0.614 | 0.248 | 0.165 | 1.64e-11 | 5.52e-07 |
| Malignant epithelial | C1 Das ECs | NR2F2 | 0.614 | 0.292 | 0.212 | 8.84e-10 | 2.98e-05 |
| Malignant epithelial | C1 Das ECs | NEDD4L | 0.613 | 0.198 | 0.144 | 1.35e-06 | 0.0455 |
| Malignant epithelial | C1 Das ECs | IL2RG | 0.612 | 0.313 | 0.198 | 1.01e-17 | 3.39e-13 |
| Malignant epithelial | C1 Das ECs | MAP1LC3A | 0.611 | 0.376 | 0.219 | 3.4e-27 | 1.15e-22 |
| Malignant epithelial | C1 Das ECs | CXCL16 | 0.611 | 0.396 | 0.273 | 2.12e-18 | 7.14e-14 |
| Malignant epithelial | C1 Das ECs | ARHGAP8 | 0.61 | 0.194 | 0.133 | 4.39e-08 | 0.00148 |
| Malignant epithelial | C1 Das ECs | STOML2 | 0.609 | 0.723 | 0.59 | 3.54e-34 | 1.19e-29 |
| Malignant epithelial | C1 Das ECs | WDR24 | 0.607 | 0.454 | 0.331 | 5.11e-17 | 1.72e-12 |
| Malignant epithelial | C1 Das ECs | SOD3 | 0.607 | 0.382 | 0.271 | 2.82e-14 | 9.51e-10 |
| Malignant epithelial | C1 Das ECs | STXBP6 | 0.607 | 0.214 | 0.152 | 6.21e-08 | 0.00209 |
| Malignant epithelial | C1 Das ECs | SNHG15 | 0.606 | 0.584 | 0.441 | 7.73e-25 | 2.6e-20 |
| Malignant epithelial | C1 Das ECs | PPP1R16A | 0.606 | 0.429 | 0.313 | 1.7e-16 | 5.71e-12 |
| Malignant epithelial | C1 Das ECs | MRPS33 | 0.605 | 0.779 | 0.649 | 1.98e-41 | 6.68e-37 |
| Malignant epithelial | C1 Das ECs | MAP7 | 0.605 | 0.308 | 0.205 | 7.99e-15 | 2.69e-10 |
| Malignant epithelial | C1 Das ECs | HUWE1 | 0.605 | 0.252 | 0.191 | 1.54e-07 | 0.0052 |
| Malignant epithelial | C1 Das ECs | RPS27L | 0.604 | 0.93 | 0.817 | 6.48e-73 | 2.18e-68 |
| Malignant epithelial | C1 Das ECs | C21orf2 | 0.604 | 0.212 | 0.15 | 1.66e-07 | 0.00558 |
| Malignant epithelial | C1 Das ECs | TMEM8B | 0.604 | 0.133 | 0.083 | 2.32e-07 | 0.00783 |
| Malignant epithelial | C1 Das ECs | TRUB2 | 0.602 | 0.258 | 0.199 | 6.33e-07 | 0.0213 |
| Malignant epithelial | C1 Das ECs | LINC01420 | 0.6 | 0.665 | 0.512 | 2.36e-32 | 7.96e-28 |
| Malignant epithelial | C1 Das ECs | SH3BGRL | 0.6 | 0.485 | 0.371 | 1.26e-16 | 4.24e-12 |
| Malignant epithelial | C1 Das ECs | COX7B | 0.599 | 0.979 | 0.936 | 3.48e-78 | 1.17e-73 |
| Malignant epithelial | C1 Das ECs | PQBP1 | 0.599 | 0.681 | 0.56 | 7.33e-29 | 2.47e-24 |
| Malignant epithelial | C1 Das ECs | PDPK1 | 0.599 | 0.248 | 0.173 | 9.64e-10 | 3.25e-05 |
| Malignant epithelial | C1 Das ECs | ITM2C | 0.598 | 0.384 | 0.284 | 4.41e-13 | 1.49e-08 |
| Malignant epithelial | C1 Das ECs | SLC6A8 | 0.596 | 0.246 | 0.146 | 1.42e-16 | 4.79e-12 |
| Malignant epithelial | C1 Das ECs | RP11-1094M14.11 | 0.595 | 0.298 | 0.223 | 1.89e-09 | 6.37e-05 |
| Malignant epithelial | C1 Das ECs | CISD3 | 0.594 | 0.735 | 0.586 | 2.76e-33 | 9.3e-29 |
| Malignant epithelial | C1 Das ECs | PGRMC1 | 0.594 | 0.641 | 0.505 | 1.39e-27 | 4.679999999999999e-23 |
| Malignant epithelial | C1 Das ECs | CHCHD3 | 0.594 | 0.464 | 0.341 | 5.37e-19 | 1.81e-14 |
| Malignant epithelial | C1 Das ECs | CYB561D2 | 0.594 | 0.213 | 0.145 | 6.49e-09 | 0.000219 |
| Malignant epithelial | C1 Das ECs | IFNAR2 | 0.594 | 0.194 | 0.129 | 6.53e-09 | 0.00022 |
| Malignant epithelial | C1 Das ECs | TTYH3 | 0.594 | 0.234 | 0.174 | 6.35e-07 | 0.0214 |
| Malignant epithelial | C1 Das ECs | TRAPPC6A | 0.593 | 0.604 | 0.451 | 4.15e-25 | 1.4e-20 |
| Malignant epithelial | C1 Das ECs | CAMTA1 | 0.593 | 0.662 | 0.553 | 3.01e-24 | 1.01e-19 |
| Malignant epithelial | C1 Das ECs | NCOA7 | 0.589 | 0.676 | 0.469 | 5.72e-39 | 1.93e-34 |
| Malignant epithelial | C1 Das ECs | AKAP9 | 0.589 | 0.609 | 0.444 | 9.79e-28 | 3.3e-23 |
| Malignant epithelial | C1 Das ECs | NAA10 | 0.589 | 0.72 | 0.605 | 3.89e-27 | 1.31e-22 |
| Malignant epithelial | C1 Das ECs | ENTPD6 | 0.588 | 0.417 | 0.283 | 2.66e-19 | 8.97e-15 |
| Malignant epithelial | C1 Das ECs | CLRN3 | 0.588 | 0.237 | 0.163 | 3.1e-09 | 0.000104 |
| Malignant epithelial | C1 Das ECs | ACSL5 | 0.587 | 0.517 | 0.348 | 7.900000000000001e-28 | 2.66e-23 |
| Malignant epithelial | C1 Das ECs | PSMB1 | 0.586 | 0.868 | 0.732 | 3.76e-53 | 1.27e-48 |
| Malignant epithelial | C1 Das ECs | NFE2L3 | 0.586 | 0.294 | 0.212 | 4.67e-10 | 1.57e-05 |
| Malignant epithelial | C1 Das ECs | EPRS | 0.586 | 0.299 | 0.23 | 4.34e-08 | 0.00146 |
| Malignant epithelial | C1 Das ECs | THOC2 | 0.584 | 0.503 | 0.408 | 6.86e-13 | 2.31e-08 |

| compartment | subcluster | gene | avg_log2FC | pct.1 | pct.2 | p_val | p_val_adj |
|---|---|---|---|---|---|---|---|
| Malignant epithelial | C1 Das ECs | PAK1 | 0.584 | 0.329 | 0.247 | 2.73e-10 | 9.21e-06 |
| Malignant epithelial | C1 Das ECs | OTUD5 | 0.584 | 0.226 | 0.163 | 8.21e-08 | 0.00277 |
| Malignant epithelial | C1 Das ECs | RPUSD3 | 0.584 | 0.214 | 0.155 | 4e-07 | 0.0135 |
| Malignant epithelial | C1 Das ECs | DIO3OS | 0.583 | 0.359 | 0.277 | 1.06e-09 | 3.58e-05 |
| Malignant epithelial | C1 Das ECs | EPHB4 | 0.582 | 0.189 | 0.128 | 4.92e-08 | 0.00166 |
| Malignant epithelial | C1 Das ECs | CTBP2 | 0.581 | 0.438 | 0.34 | 8e-14 | 2.7e-09 |
| Malignant epithelial | C1 Das ECs | IFT20 | 0.58 | 0.427 | 0.331 | 1.84e-12 | 6.21e-08 |
| Malignant epithelial | C1 Das ECs | CREB3 | 0.579 | 0.318 | 0.233 | 2.93e-10 | 9.87e-06 |
| Malignant epithelial | C1 Das ECs | SORL1 | 0.579 | 0.311 | 0.227 | 4.05e-10 | 1.37e-05 |
| Malignant epithelial | C1 Das ECs | SOD1 | 0.578 | 0.931 | 0.855 | 2.47e-64 | 8.319999999999999e-60 |
| Malignant epithelial | C1 Das ECs | PPFIA3 | 0.578 | 0.216 | 0.154 | 1.13e-07 | 0.00382 |
| Malignant epithelial | C1 Das ECs | LAGE3 | 0.577 | 0.571 | 0.444 | 3.28e-22 | 1.1e-17 |
| Malignant epithelial | C1 Das ECs | SPATA13 | 0.577 | 0.193 | 0.138 | 9.93e-07 | 0.0335 |
| Malignant epithelial | C1 Das ECs | C15orf40 | 0.574 | 0.308 | 0.242 | 7.68e-08 | 0.00259 |
| Malignant epithelial | C1 Das ECs | DLG3 | 0.573 | 0.221 | 0.162 | 4.93e-07 | 0.0166 |
| Malignant epithelial | C1 Das ECs | ASNSD1 | 0.573 | 0.258 | 0.198 | 9.39e-07 | 0.0316 |
| Malignant epithelial | C1 Das ECs | SNHG7 | 0.572 | 0.826 | 0.745 | 1.36e-30 | 4.59e-26 |
| Malignant epithelial | C1 Das ECs | PNKD | 0.572 | 0.559 | 0.449 | 3.95e-18 | 1.33e-13 |
| Malignant epithelial | C1 Das ECs | ZNF277 | 0.572 | 0.312 | 0.236 | 2.65e-09 | 8.94e-05 |
| Malignant epithelial | C1 Das ECs | ESD | 0.571 | 0.71 | 0.557 | 2.11e-31 | 7.1e-27 |
| Malignant epithelial | C1 Das ECs | EBP | 0.571 | 0.624 | 0.496 | 2.31e-22 | 7.77e-18 |
| Malignant epithelial | C1 Das ECs | PDCD7 | 0.571 | 0.269 | 0.193 | 2.29e-09 | 7.73e-05 |
| Malignant epithelial | C1 Das ECs | RAD50 | 0.57 | 0.324 | 0.248 | 8.44e-09 | 0.000284 |
| Malignant epithelial | C1 Das ECs | IRAK1 | 0.569 | 0.35 | 0.27 | 6.83e-10 | 2.3e-05 |
| Malignant epithelial | C1 Das ECs | MRRF | 0.569 | 0.341 | 0.264 | 1.08e-09 | 3.63e-05 |
| Malignant epithelial | C1 Das ECs | GLA | 0.569 | 0.21 | 0.146 | 4.85e-08 | 0.00163 |
| Malignant epithelial | C1 Das ECs | NDUFA1 | 0.568 | 0.931 | 0.865 | 6.39e-55 | 2.15e-50 |
| Malignant epithelial | C1 Das ECs | LINC01003 | 0.567 | 0.3 | 0.219 | 1.87e-10 | 6.31e-06 |
| Malignant epithelial | C1 Das ECs | RPL22 | 0.566 | 0.992 | 0.981 | 3.15e-118 | 1.06e-113 |
| Malignant epithelial | C1 Das ECs | TMEM54 | 0.566 | 0.758 | 0.624 | 3.12e-38 | 1.05e-33 |
| Malignant epithelial | C1 Das ECs | EPHA2 | 0.566 | 0.456 | 0.33 | 3.32e-17 | 1.12e-12 |
| Malignant epithelial | C1 Das ECs | SEC13 | 0.566 | 0.345 | 0.26 | 1.44e-10 | 4.85e-06 |
| Malignant epithelial | C1 Das ECs | MTIF2 | 0.564 | 0.229 | 0.17 | 5.2e-07 | 0.0175 |
| Malignant epithelial | C1 Das ECs | ZNF800 | 0.563 | 0.221 | 0.159 | 8.41e-08 | 0.00283 |
| Malignant epithelial | C1 Das ECs | EFNA4 | 0.563 | 0.253 | 0.188 | 1.82e-07 | 0.00612 |
| Malignant epithelial | C1 Das ECs | NSDHL | 0.563 | 0.193 | 0.136 | 8.99e-07 | 0.0303 |
| Malignant epithelial | C1 Das ECs | PRKAG1 | 0.562 | 0.308 | 0.226 | 2.3e-10 | 7.77e-06 |
| Malignant epithelial | C1 Das ECs | GATAD1 | 0.562 | 0.305 | 0.242 | 3.27e-07 | 0.011 |
| Malignant epithelial | C1 Das ECs | SDCBP2 | 0.561 | 0.397 | 0.239 | 1.22e-26 | 4.1e-22 |
| Malignant epithelial | C1 Das ECs | FH | 0.561 | 0.229 | 0.165 | 8.05e-08 | 0.00271 |
| Malignant epithelial | C1 Das ECs | KRT20 | 0.56 | 0.438 | 0.281 | 9.64e-26 | 3.25e-21 |
| Malignant epithelial | C1 Das ECs | PPA2 | 0.56 | 0.484 | 0.378 | 1.22e-15 | 4.13e-11 |
| Malignant epithelial | C1 Das ECs | NDUFA8 | 0.559 | 0.664 | 0.539 | 2.11e-26 | 7.11e-22 |
| Malignant epithelial | C1 Das ECs | LAMP2 | 0.559 | 0.41 | 0.302 | 2.6e-15 | 8.75e-11 |
| Malignant epithelial | C1 Das ECs | NSMF | 0.559 | 0.202 | 0.141 | 1.26e-07 | 0.00425 |
| Malignant epithelial | C1 Das ECs | SPTBN1 | 0.556 | 0.572 | 0.417 | 2.98e-25 | 1e-20 |
| Malignant epithelial | C1 Das ECs | PYCR1 | 0.556 | 0.378 | 0.285 | 1.22e-11 | 4.1e-07 |
| Malignant epithelial | C1 Das ECs | EMD | 0.556 | 0.425 | 0.345 | 3.48e-11 | 1.17e-06 |
| Malignant epithelial | C1 Das ECs | RCL1 | 0.556 | 0.242 | 0.175 | 2.43e-08 | 0.00082 |
| Malignant epithelial | C1 Das ECs | SLC25A6 | 0.555 | 0.958 | 0.919 | 8.01e-63 | 2.7e-58 |
| Malignant epithelial | C1 Das ECs | ARF5 | 0.555 | 0.731 | 0.614 | 2.03e-29 | 6.83e-25 |
| Malignant epithelial | C1 Das ECs | HSD17B11 | 0.553 | 0.603 | 0.472 | 3.9e-20 | 1.31e-15 |
| Malignant epithelial | C1 Das ECs | LYRM5 | 0.553 | 0.304 | 0.231 | 2.99e-08 | 0.00101 |
| Malignant epithelial | C1 Das ECs | CCDC174 | 0.553 | 0.213 | 0.151 | 1.54e-07 | 0.00519 |
| Malignant epithelial | C1 Das ECs | CCNB1IP1 | 0.552 | 0.379 | 0.291 | 8.21e-11 | 2.77e-06 |
| Malignant epithelial | C1 Das ECs | HOXA11-AS | 0.552 | 0.263 | 0.181 | 1.25e-10 | 4.21e-06 |
| Malignant epithelial | C1 Das ECs | FUCA1 | 0.551 | 0.181 | 0.119 | 2.43e-08 | 0.000819 |
| Malignant epithelial | C1 Das ECs | MKRN1 | 0.548 | 0.545 | 0.402 | 1.19e-23 | 4.02e-19 |
| Malignant epithelial | C1 Das ECs | SEC11C | 0.547 | 0.51 | 0.369 | 3.69e-21 | 1.24e-16 |
| Malignant epithelial | C1 Das ECs | HTATSF1 | 0.547 | 0.284 | 0.211 | 9.04e-09 | 0.000305 |
| Malignant epithelial | C1 Das ECs | NFIX | 0.547 | 0.225 | 0.158 | 4.71e-08 | 0.00159 |
| Malignant epithelial | C1 Das ECs | C6orf203 | 0.547 | 0.194 | 0.135 | 1.83e-07 | 0.00618 |
| Malignant epithelial | C1 Das ECs | BAIAP2L1 | 0.546 | 0.516 | 0.403 | 3.51e-16 | 1.18e-11 |
| Malignant epithelial | C1 Das ECs | ATRX | 0.546 | 0.479 | 0.377 | 1.36e-13 | 4.6e-09 |

| compartment | subcluster | gene | avg_log2FC | pct.1 | pct.2 | p_val | p_val_adj |
|---|---|---|---|---|---|---|---|
| Malignant epithelial | C1 Das ECs | EXOC4 | 0.545 | 0.212 | 0.147 | 7.24e-08 | 0.00244 |
| Malignant epithelial | C1 Das ECs | SMU1 | 0.545 | 0.23 | 0.173 | 1.35e-06 | 0.0456 |
| Malignant epithelial | C1 Das ECs | RNF167 | 0.544 | 0.302 | 0.233 | 6.62e-08 | 0.00223 |
| Malignant epithelial | C1 Das ECs | HLA-DMA | 0.543 | 0.582 | 0.446 | 9.64e-21 | 3.25e-16 |
| Malignant epithelial | C1 Das ECs | MMP7 | 0.543 | 0.513 | 0.393 | 7.16e-14 | 2.41e-09 |
| Malignant epithelial | C1 Das ECs | TMEM60 | 0.541 | 0.297 | 0.22 | 4.23e-09 | 0.000142 |
| Malignant epithelial | C1 Das ECs | MMAB | 0.541 | 0.31 | 0.24 | 1.34e-08 | 0.000452 |
| Malignant epithelial | C1 Das ECs | SLC35C1 | 0.54 | 0.2 | 0.139 | 2.04e-07 | 0.00689 |
| Malignant epithelial | C1 Das ECs | FAM120AOS | 0.539 | 0.448 | 0.34 | 7.35e-15 | 2.47e-10 |
| Malignant epithelial | C1 Das ECs | SAT2 | 0.539 | 0.39 | 0.285 | 1.01e-12 | 3.4e-08 |
| Malignant epithelial | C1 Das ECs | AZGP1 | 0.537 | 0.673 | 0.479 | 6.82e-34 | 2.3e-29 |
| Malignant epithelial | C1 Das ECs | EML3 | 0.537 | 0.174 | 0.122 | 1.4e-06 | 0.047 |
| Malignant epithelial | C1 Das ECs | GDF15 | 0.536 | 0.597 | 0.333 | 6.51e-62 | 2.19e-57 |
| Malignant epithelial | C1 Das ECs | DEFB1 | 0.534 | 0.302 | 0.164 | 4.19e-24 | 1.41e-19 |
| Malignant epithelial | C1 Das ECs | STAG2 | 0.534 | 0.319 | 0.242 | 3.43e-09 | 0.000116 |
| Malignant epithelial | C1 Das ECs | PSMD5-AS1 | 0.534 | 0.242 | 0.169 | 1.47e-08 | 0.000494 |
| Malignant epithelial | C1 Das ECs | LPP | 0.534 | 0.366 | 0.294 | 3.64e-08 | 0.00123 |
| Malignant epithelial | C1 Das ECs | MT-CO2 | 0.532 | 1.0 | 0.996 | 1.73e-86 | 5.84e-82 |
| Malignant epithelial | C1 Das ECs | LAP3 | 0.532 | 0.285 | 0.221 | 3.93e-07 | 0.0132 |
| Malignant epithelial | C1 Das ECs | SLC12A7 | 0.531 | 0.209 | 0.138 | 1.08e-09 | 3.63e-05 |
| Malignant epithelial | C1 Das ECs | POLR1D | 0.53 | 0.814 | 0.685 | 1.81e-40 | 6.1e-36 |
| Malignant epithelial | C1 Das ECs | HLA-F | 0.529 | 0.537 | 0.401 | 6.27e-19 | 2.11e-14 |
| Malignant epithelial | C1 Das ECs | IGF1R | 0.527 | 0.132 | 0.082 | 1.57e-07 | 0.00528 |
| Malignant epithelial | C1 Das ECs | COQ4 | 0.526 | 0.477 | 0.381 | 8.46e-14 | 2.85e-09 |
| Malignant epithelial | C1 Das ECs | IFT57 | 0.526 | 0.356 | 0.275 | 1.03e-09 | 3.47e-05 |
| Malignant epithelial | C1 Das ECs | PROSC | 0.526 | 0.352 | 0.273 | 1.12e-09 | 3.78e-05 |
| Malignant epithelial | C1 Das ECs | RPL32 | 0.525 | 1.0 | 1.0 | 4.51e-121 | 1.52e-116 |
| Malignant epithelial | C1 Das ECs | NDUFB11 | 0.525 | 0.954 | 0.891 | 1.17e-64 | 3.93e-60 |
| Malignant epithelial | C1 Das ECs | OFD1 | 0.525 | 0.456 | 0.361 | 6.02e-13 | 2.03e-08 |
| Malignant epithelial | C1 Das ECs | PPID | 0.524 | 0.27 | 0.202 | 1.31e-07 | 0.0044 |
| Malignant epithelial | C1 Das ECs | ALG13 | 0.523 | 0.365 | 0.276 | 7.92e-11 | 2.67e-06 |
| Malignant epithelial | C1 Das ECs | CD46 | 0.522 | 0.677 | 0.548 | 1.64e-23 | 5.52e-19 |
| Malignant epithelial | C1 Das ECs | AHCY | 0.522 | 0.614 | 0.495 | 9.36e-20 | 3.15e-15 |
| Malignant epithelial | C1 Das ECs | DUSP23 | 0.522 | 0.582 | 0.465 | 2.41e-18 | 8.13e-14 |
| Malignant epithelial | C1 Das ECs | PROCR | 0.521 | 0.332 | 0.23 | 3.44e-13 | 1.16e-08 |
| Malignant epithelial | C1 Das ECs | NFATC3 | 0.521 | 0.428 | 0.326 | 4.5e-12 | 1.52e-07 |
| Malignant epithelial | C1 Das ECs | OGT | 0.52 | 0.345 | 0.268 | 1.24e-08 | 0.000419 |
| Malignant epithelial | C1 Das ECs | BFAR | 0.519 | 0.487 | 0.386 | 3.82e-13 | 1.29e-08 |
| Malignant epithelial | C1 Das ECs | HLA-DRA | 0.518 | 0.829 | 0.737 | 1.03e-25 | 3.480000000000001e-21 |
| Malignant epithelial | C1 Das ECs | VCP | 0.518 | 0.482 | 0.381 | 1.31e-13 | 4.4e-09 |
| Malignant epithelial | C1 Das ECs | PPP6C | 0.518 | 0.363 | 0.288 | 8.03e-09 | 0.000271 |
| Malignant epithelial | C1 Das ECs | WDR45 | 0.514 | 0.368 | 0.274 | 6.7e-12 | 2.26e-07 |
| Malignant epithelial | C1 Das ECs | ARL2BP | 0.513 | 0.312 | 0.236 | 1.49e-08 | 0.000501 |
| Malignant epithelial | C1 Das ECs | IL13RA1 | 0.513 | 0.288 | 0.215 | 1.66e-08 | 0.00056 |
| Malignant epithelial | C1 Das ECs | ZNF394 | 0.513 | 0.271 | 0.209 | 1.15e-06 | 0.0388 |
| Malignant epithelial | C1 Das ECs | SMARCD2 | 0.512 | 0.28 | 0.21 | 3.94e-08 | 0.00133 |
| Malignant epithelial | C1 Das ECs | MYO1D | 0.512 | 0.228 | 0.165 | 2.08e-07 | 0.00701 |
| Malignant epithelial | C1 Das ECs | MRPS25 | 0.511 | 0.337 | 0.257 | 1.48e-09 | 4.98e-05 |
| Malignant epithelial | C1 Das ECs | RPL36A | 0.51 | 0.998 | 0.991 | 7.769999999999999e-90 | 2.62e-85 |
| Malignant epithelial | C1 Das ECs | DCXR | 0.51 | 0.582 | 0.457 | 1.12e-18 | 3.78e-14 |
| Malignant epithelial | C1 Das ECs | TMEM9B | 0.509 | 0.415 | 0.326 | 3.2e-11 | 1.08e-06 |
| Malignant epithelial | C1 Das ECs | KIF9 | 0.507 | 0.317 | 0.247 | 4.42e-08 | 0.00149 |
| Malignant epithelial | C1 Das ECs | FDX1 | 0.506 | 0.378 | 0.29 | 8.15e-11 | 2.75e-06 |
| Malignant epithelial | C1 Das ECs | MT-ND4 | 0.505 | 1.0 | 0.993 | 4.73e-80 | 1.59e-75 |
| Malignant epithelial | C1 Das ECs | KLK1 | 0.505 | 0.365 | 0.259 | 4.48e-13 | 1.51e-08 |
| Malignant epithelial | C1 Das ECs | TCEAL8 | 0.505 | 0.408 | 0.311 | 1.22e-11 | 4.11e-07 |
| Malignant epithelial | C1 Das ECs | RPS6 | 0.504 | 0.999 | 0.998 | 2.16e-100 | 7.28e-96 |
| Malignant epithelial | C1 Das ECs | DNPEP | 0.504 | 0.555 | 0.443 | 3.42e-15 | 1.15e-10 |
| Malignant epithelial | C1 Das ECs | COA3 | 0.503 | 0.684 | 0.577 | 9.5e-21 | 3.2e-16 |
| Malignant epithelial | C1 Das ECs | CPNE3 | 0.503 | 0.472 | 0.372 | 4.81e-13 | 1.62e-08 |
| Malignant epithelial | C1 Das ECs | HOXB9 | 0.503 | 0.385 | 0.309 | 8.37e-09 | 0.000282 |
| Malignant epithelial | C1 Das ECs | RPL7A | 0.502 | 0.998 | 0.992 | 1.35e-91 | 4.56e-87 |
| Malignant epithelial | C1 Das ECs | RBBP7 | 0.501 | 0.438 | 0.337 | 6.85e-13 | 2.31e-08 |
| Malignant epithelial | C1 Das ECs | POLE3 | 0.501 | 0.389 | 0.3 | 3.08e-10 | 1.04e-05 |

| compartment | subcluster | gene | avg_log2FC | pct.1 | pct.2 | p_val | p_val_adj |
|---|---|---|---|---|---|---|---|
| Malignant epithelial | C1 Das ECs | CYSTM1 | 0.5 | 0.845 | 0.73 | 2.51e-38 | 8.44e-34 |
| Malignant epithelial | C1 Das ECs | EMP2 | 0.5 | 0.638 | 0.462 | 1.55e-27 | 5.23e-23 |
| Malignant epithelial | C1 Das ECs | MLLT4 | 0.498 | 0.319 | 0.237 | 1.28e-09 | 4.33e-05 |
| Malignant epithelial | C1 Das ECs | MYO6 | 0.497 | 0.445 | 0.325 | 6.33e-16 | 2.13e-11 |
| Malignant epithelial | C1 Das ECs | MTMR11 | 0.497 | 0.344 | 0.238 | 4.78e-13 | 1.61e-08 |
| Malignant epithelial | C1 Das ECs | DYNLT1 | 0.495 | 0.789 | 0.667 | 2.29e-29 | 7.72e-25 |
| Malignant epithelial | C1 Das ECs | CD82 | 0.495 | 0.513 | 0.336 | 1.49e-28 | 5.009999999999999e-24 |
| Malignant epithelial | C1 Das ECs | ZFP36L2 | 0.493 | 0.865 | 0.722 | 2.73e-39 | 9.18e-35 |
| Malignant epithelial | C1 Das ECs | SNHG19 | 0.493 | 0.366 | 0.285 | 1.29e-09 | 4.33e-05 |
| Malignant epithelial | C1 Das ECs | ST14 | 0.492 | 0.655 | 0.52 | 7.01e-24 | 2.36e-19 |
| Malignant epithelial | C1 Das ECs | SYAP1 | 0.491 | 0.606 | 0.5 | 1.17e-18 | 3.93e-14 |
| Malignant epithelial | C1 Das ECs | HOXB6 | 0.491 | 0.359 | 0.277 | 4.75e-09 | 0.00016 |
| Malignant epithelial | C1 Das ECs | TFRC | 0.491 | 0.261 | 0.197 | 4.2e-07 | 0.0141 |
| Malignant epithelial | C1 Das ECs | PRKCZ | 0.488 | 0.195 | 0.132 | 2.71e-08 | 0.000913 |
| Malignant epithelial | C1 Das ECs | SETD9 | 0.488 | 0.234 | 0.172 | 5.69e-07 | 0.0192 |
| Malignant epithelial | C1 Das ECs | SLC25A25 | 0.487 | 0.424 | 0.321 | 1.17e-12 | 3.94e-08 |
| Malignant epithelial | C1 Das ECs | CPSF3L | 0.487 | 0.326 | 0.243 | 1.12e-09 | 3.79e-05 |
| Malignant epithelial | C1 Das ECs | WASL | 0.487 | 0.318 | 0.242 | 1.37e-08 | 0.000463 |
| Malignant epithelial | C1 Das ECs | NFU1 | 0.487 | 0.468 | 0.402 | 1.7e-08 | 0.000573 |
| Malignant epithelial | C1 Das ECs | LETMD1 | 0.486 | 0.404 | 0.307 | 3.49e-11 | 1.18e-06 |
| Malignant epithelial | C1 Das ECs | TXNDC12 | 0.485 | 0.452 | 0.349 | 5.49e-13 | 1.85e-08 |
| Malignant epithelial | C1 Das ECs | TMEM45B | 0.484 | 0.436 | 0.27 | 6.45e-25 | 2.17e-20 |
| Malignant epithelial | C1 Das ECs | CREG1 | 0.484 | 0.421 | 0.321 | 1.23e-12 | 4.14e-08 |
| Malignant epithelial | C1 Das ECs | FTL | 0.483 | 0.999 | 0.998 | 8.050000000000001e-85 | 2.71e-80 |
| Malignant epithelial | C1 Das ECs | MRPL20 | 0.483 | 0.768 | 0.652 | 4.72e-27 | 1.59e-22 |
| Malignant epithelial | C1 Das ECs | MAPKAPK5-AS1 | 0.483 | 0.343 | 0.262 | 4.37e-09 | 0.000147 |
| Malignant epithelial | C1 Das ECs | RPS3 | 0.482 | 1.0 | 0.997 | 1.42e-109 | 4.8e-105 |
| Malignant epithelial | C1 Das ECs | COX11 | 0.482 | 0.337 | 0.266 | 3.21e-08 | 0.00108 |
| Malignant epithelial | C1 Das ECs | PDRG1 | 0.482 | 0.314 | 0.244 | 6.64e-08 | 0.00224 |
| Malignant epithelial | C1 Das ECs | RSBN1L | 0.481 | 0.528 | 0.418 | 8.16e-17 | 2.75e-12 |
| Malignant epithelial | C1 Das ECs | C9orf78 | 0.481 | 0.488 | 0.411 | 1.36e-10 | 4.6e-06 |
| Malignant epithelial | C1 Das ECs | TRAPPC2 | 0.481 | 0.197 | 0.139 | 2.79e-07 | 0.0094 |
| Malignant epithelial | C1 Das ECs | RPL39 | 0.48 | 1.0 | 0.999 | 2.79e-99 | 9.419999999999999e-95 |
| Malignant epithelial | C1 Das ECs | EBPL | 0.48 | 0.69 | 0.574 | 2.45e-22 | 8.27e-18 |
| Malignant epithelial | C1 Das ECs | SHROOM1 | 0.48 | 0.215 | 0.153 | 3.5e-07 | 0.0118 |
| Malignant epithelial | C1 Das ECs | HOXA9 | 0.478 | 0.319 | 0.241 | 8.82e-09 | 0.000297 |
| Malignant epithelial | C1 Das ECs | RPS12 | 0.477 | 1.0 | 0.998 | 3.93e-116 | 1.32e-111 |
| Malignant epithelial | C1 Das ECs | CMPK1 | 0.477 | 0.556 | 0.43 | 1.68e-19 | 5.66e-15 |
| Malignant epithelial | C1 Das ECs | ISOC2 | 0.477 | 0.555 | 0.448 | 8.39e-15 | 2.83e-10 |
| Malignant epithelial | C1 Das ECs | ERBB3 | 0.476 | 0.592 | 0.431 | 6.96e-24 | 2.34e-19 |
| Malignant epithelial | C1 Das ECs | NDUFAF4 | 0.476 | 0.372 | 0.296 | 9.42e-09 | 0.000317 |
| Malignant epithelial | C1 Das ECs | MRPL12 | 0.475 | 0.665 | 0.552 | 9.960000000000001e-21 | 3.36e-16 |
| Malignant epithelial | C1 Das ECs | MPZL1 | 0.475 | 0.329 | 0.258 | 5.05e-08 | 0.0017 |
| Malignant epithelial | C1 Das ECs | TMEM173 | 0.475 | 0.22 | 0.153 | 5.62e-08 | 0.00189 |
| Malignant epithelial | C1 Das ECs | GLTSCR2 | 0.474 | 0.938 | 0.884 | 2.52e-39 | 8.51e-35 |
| Malignant epithelial | C1 Das ECs | FAM98C | 0.474 | 0.321 | 0.255 | 1.81e-07 | 0.00609 |
| Malignant epithelial | C1 Das ECs | NGFRAP1 | 0.473 | 0.637 | 0.537 | 5.32e-12 | 1.79e-07 |
| Malignant epithelial | C1 Das ECs | PGRMC2 | 0.473 | 0.411 | 0.33 | 1.87e-09 | 6.32e-05 |
| Malignant epithelial | C1 Das ECs | OCLN | 0.473 | 0.266 | 0.194 | 3.03e-08 | 0.00102 |
| Malignant epithelial | C1 Das ECs | TADA3 | 0.473 | 0.326 | 0.256 | 1.34e-07 | 0.00451 |
| Malignant epithelial | C1 Das ECs | SIGMAR1 | 0.471 | 0.497 | 0.396 | 8.46e-13 | 2.85e-08 |
| Malignant epithelial | C1 Das ECs | BECN1 | 0.471 | 0.273 | 0.207 | 1.26e-07 | 0.00423 |
| Malignant epithelial | C1 Das ECs | ATP6AP2 | 0.47 | 0.572 | 0.446 | 8.69e-19 | 2.93e-14 |
| Malignant epithelial | C1 Das ECs | RPS5 | 0.468 | 0.997 | 0.989 | 1.67e-88 | 5.629999999999999e-84 |
| Malignant epithelial | C1 Das ECs | CAPZA2 | 0.468 | 0.678 | 0.558 | 6.98e-22 | 2.35e-17 |
| Malignant epithelial | C1 Das ECs | SH3YL1 | 0.468 | 0.68 | 0.575 | 2.89e-20 | 9.73e-16 |
| Malignant epithelial | C1 Das ECs | ACHE | 0.468 | 0.141 | 0.068 | 1.09e-14 | 3.67e-10 |
| Malignant epithelial | C1 Das ECs | HSD17B12 | 0.468 | 0.502 | 0.403 | 1.68e-12 | 5.66e-08 |
| Malignant epithelial | C1 Das ECs | EXOSC5 | 0.468 | 0.387 | 0.297 | 9.37e-11 | 3.16e-06 |
| Malignant epithelial | C1 Das ECs | LLGL2 | 0.465 | 0.52 | 0.385 | 8.43e-19 | 2.84e-14 |
| Malignant epithelial | C1 Das ECs | PMPCB | 0.465 | 0.445 | 0.363 | 4.25e-10 | 1.43e-05 |
| Malignant epithelial | C1 Das ECs | GTF2H5 | 0.464 | 0.619 | 0.522 | 1.69e-14 | 5.68e-10 |
| Malignant epithelial | C1 Das ECs | HRSP12 | 0.463 | 0.315 | 0.247 | 4.64e-07 | 0.0156 |
| Malignant epithelial | C1 Das ECs | DHRS4L2 | 0.463 | 0.312 | 0.246 | 1.06e-06 | 0.0357 |

| compartment | subcluster | gene | avg_log2FC | pct.1 | pct.2 | p_val | p_val_adj |
|---|---|---|---|---|---|---|---|
| Malignant epithelial | C1 Das ECs | MT-ND1 | 0.462 | 1.0 | 0.989 | 2.44e-71 | 8.21e-67 |
| Malignant epithelial | C1 Das ECs | MRPS31 | 0.462 | 0.242 | 0.183 | 1.15e-06 | 0.0389 |
| Malignant epithelial | C1 Das ECs | FAM96A | 0.46 | 0.465 | 0.386 | 1.37e-10 | 4.61e-06 |
| Malignant epithelial | C1 Das ECs | ABHD17C | 0.46 | 0.272 | 0.193 | 9.46e-10 | 3.19e-05 |
| Malignant epithelial | C1 Das ECs | HLA-DQB1 | 0.459 | 0.296 | 0.221 | 7.38e-08 | 0.00249 |
| Malignant epithelial | C1 Das ECs | MACROD1 | 0.457 | 0.521 | 0.407 | 3.43e-14 | 1.15e-09 |
| Malignant epithelial | C1 Das ECs | FAM127B | 0.457 | 0.365 | 0.282 | 7.12e-09 | 0.00024 |
| Malignant epithelial | C1 Das ECs | RPL15 | 0.456 | 0.999 | 0.993 | 6.690000000000001e-73 | 2.25e-68 |
| Malignant epithelial | C1 Das ECs | NDUFB2 | 0.456 | 0.932 | 0.85 | 7.17e-44 | 2.42e-39 |
| Malignant epithelial | C1 Das ECs | DNPH1 | 0.456 | 0.77 | 0.652 | 2.59e-25 | 8.740000000000001e-21 |
| Malignant epithelial | C1 Das ECs | SEPP1 | 0.455 | 0.496 | 0.336 | 3.5e-24 | 1.18e-19 |
| Malignant epithelial | C1 Das ECs | UQCRC2 | 0.454 | 0.688 | 0.578 | 5.6e-19 | 1.89e-14 |
| Malignant epithelial | C1 Das ECs | PFKL | 0.454 | 0.415 | 0.319 | 5.69e-12 | 1.92e-07 |
| Malignant epithelial | C1 Das ECs | NUDT16L1 | 0.454 | 0.354 | 0.286 | 1.41e-07 | 0.00475 |
| Malignant epithelial | C1 Das ECs | ESRRA | 0.452 | 0.372 | 0.302 | 1.89e-08 | 0.000638 |
| Malignant epithelial | C1 Das ECs | PAXIP1-AS1 | 0.452 | 0.273 | 0.205 | 1.11e-07 | 0.00374 |
| Malignant epithelial | C1 Das ECs | LINC00467 | 0.452 | 0.328 | 0.259 | 1.62e-07 | 0.00544 |
| Malignant epithelial | C1 Das ECs | MRPL32 | 0.451 | 0.575 | 0.476 | 8.1e-15 | 2.73e-10 |
| Malignant epithelial | C1 Das ECs | TPD52 | 0.45 | 0.63 | 0.486 | 2.76e-23 | 9.31e-19 |
| Malignant epithelial | C1 Das ECs | JCHAIN | 0.45 | 0.295 | 0.185 | 2.68e-17 | 9.04e-13 |
| Malignant epithelial | C1 Das ECs | ACSS2 | 0.45 | 0.225 | 0.165 | 1.26e-06 | 0.0424 |
| Malignant epithelial | C1 Das ECs | IGHG1 | 0.449 | 0.405 | 0.294 | 2.96e-15 | 9.99e-11 |
| Malignant epithelial | C1 Das ECs | CCDC186 | 0.447 | 0.337 | 0.263 | 7.22e-08 | 0.00243 |
| Malignant epithelial | C1 Das ECs | NOB1 | 0.442 | 0.59 | 0.507 | 1.32e-11 | 4.44e-07 |
| Malignant epithelial | C1 Das ECs | EEF1B2 | 0.441 | 0.976 | 0.925 | 8.19e-50 | 2.76e-45 |
| Malignant epithelial | C1 Das ECs | TJP3 | 0.44 | 0.335 | 0.251 | 1.74e-09 | 5.85e-05 |
| Malignant epithelial | C1 Das ECs | SNHG25 | 0.44 | 0.445 | 0.37 | 2.37e-08 | 0.000797 |
| Malignant epithelial | C1 Das ECs | NDUFB5 | 0.438 | 0.725 | 0.652 | 1.37e-18 | 4.61e-14 |
| Malignant epithelial | C1 Das ECs | PPP1CC | 0.438 | 0.622 | 0.53 | 7.08e-14 | 2.38e-09 |
| Malignant epithelial | C1 Das ECs | ALG5 | 0.438 | 0.439 | 0.332 | 8.65e-13 | 2.91e-08 |
| Malignant epithelial | C1 Das ECs | KRCC1 | 0.438 | 0.368 | 0.303 | 6.28e-07 | 0.0212 |
| Malignant epithelial | C1 Das ECs | SDHB | 0.436 | 0.467 | 0.403 | 5.91e-08 | 0.00199 |
| Malignant epithelial | C1 Das ECs | TTC3 | 0.433 | 0.514 | 0.419 | 5.22e-12 | 1.76e-07 |
| Malignant epithelial | C1 Das ECs | TMEM205 | 0.432 | 0.584 | 0.503 | 7.67e-12 | 2.58e-07 |
| Malignant epithelial | C1 Das ECs | MPC1 | 0.432 | 0.377 | 0.286 | 3.05e-10 | 1.03e-05 |
| Malignant epithelial | C1 Das ECs | VDAC1 | 0.43 | 0.888 | 0.809 | 3.48e-34 | 1.17e-29 |
| Malignant epithelial | C1 Das ECs | MPHOSPH8 | 0.429 | 0.475 | 0.358 | 4e-15 | 1.35e-10 |
| Malignant epithelial | C1 Das ECs | DCTPP1 | 0.429 | 0.538 | 0.42 | 6.25e-14 | 2.11e-09 |
| Malignant epithelial | C1 Das ECs | YBEY | 0.429 | 0.407 | 0.331 | 2.48e-08 | 0.000834 |
| Malignant epithelial | C1 Das ECs | TSTA3 | 0.428 | 0.561 | 0.471 | 1.51e-11 | 5.08e-07 |
| Malignant epithelial | C1 Das ECs | C21orf59 | 0.428 | 0.396 | 0.301 | 1.25e-10 | 4.2e-06 |
| Malignant epithelial | C1 Das ECs | MTHFD2 | 0.428 | 0.403 | 0.321 | 1.92e-09 | 6.46e-05 |
| Malignant epithelial | C1 Das ECs | MRPL24 | 0.427 | 0.48 | 0.398 | 9.72e-10 | 3.27e-05 |
| Malignant epithelial | C1 Das ECs | TMEM183A | 0.427 | 0.41 | 0.332 | 4.81e-09 | 0.000162 |
| Malignant epithelial | C1 Das ECs | SRSF8 | 0.427 | 0.383 | 0.313 | 1.2e-07 | 0.00405 |
| Malignant epithelial | C1 Das ECs | NT5C3A | 0.426 | 0.706 | 0.597 | 8.2e-20 | 2.76e-15 |
| Malignant epithelial | C1 Das ECs | EPN1 | 0.426 | 0.511 | 0.428 | 2.08e-11 | 7.02e-07 |
| Malignant epithelial | C1 Das ECs | C19orf60 | 0.423 | 0.795 | 0.696 | 1.04e-21 | 3.51e-17 |
| Malignant epithelial | C1 Das ECs | MFF | 0.423 | 0.453 | 0.378 | 2.3e-09 | 7.76e-05 |
| Malignant epithelial | C1 Das ECs | CYC1 | 0.422 | 0.841 | 0.744 | 6.67e-29 | 2.25e-24 |
| Malignant epithelial | C1 Das ECs | JTB | 0.422 | 0.76 | 0.683 | 2.9e-22 | 9.77e-18 |
| Malignant epithelial | C1 Das ECs | SSU72 | 0.422 | 0.742 | 0.655 | 2.29e-19 | 7.73e-15 |
| Malignant epithelial | C1 Das ECs | OSTF1 | 0.422 | 0.491 | 0.394 | 3.3e-12 | 1.11e-07 |
| Malignant epithelial | C1 Das ECs | PLBD1 | 0.422 | 0.425 | 0.349 | 3.12e-08 | 0.00105 |
| Malignant epithelial | C1 Das ECs | TXN | 0.419 | 0.995 | 0.983 | 2.78e-46 | 9.38e-42 |
| Malignant epithelial | C1 Das ECs | TMC4 | 0.419 | 0.375 | 0.295 | 1.18e-08 | 0.000396 |
| Malignant epithelial | C1 Das ECs | SYPL1 | 0.418 | 0.742 | 0.62 | 1.78e-21 | 6e-17 |
| Malignant epithelial | C1 Das ECs | TBRG4 | 0.418 | 0.33 | 0.254 | 1.15e-08 | 0.000389 |
| Malignant epithelial | C1 Das ECs | SUMF2 | 0.417 | 0.539 | 0.44 | 7.44e-13 | 2.51e-08 |
| Malignant epithelial | C1 Das ECs | RYBP | 0.416 | 0.257 | 0.19 | 2.31e-07 | 0.00779 |
| Malignant epithelial | C1 Das ECs | SCCPDH | 0.415 | 0.379 | 0.294 | 5.57e-09 | 0.000188 |
| Malignant epithelial | C1 Das ECs | CALCOCO2 | 0.415 | 0.369 | 0.293 | 2.01e-08 | 0.000678 |
| Malignant epithelial | C1 Das ECs | PPP1R35 | 0.414 | 0.442 | 0.36 | 7.48e-10 | 2.52e-05 |
| Malignant epithelial | C1 Das ECs | USH1C | 0.413 | 0.498 | 0.383 | 5.45e-14 | 1.83e-09 |

| compartment | subcluster | gene | avg_log2FC | pct.1 | pct.2 | p_val | p_val_adj |
|---|---|---|---|---|---|---|---|
| Malignant epithelial | C1 Das ECs | ALKBH7 | 0.412 | 0.808 | 0.712 | 3.75e-25 | 1.26e-20 |
| Malignant epithelial | C1 Das ECs | ANAPC13 | 0.412 | 0.448 | 0.379 | 3.01e-07 | 0.0102 |
| Malignant epithelial | C1 Das ECs | MRPL4 | 0.411 | 0.522 | 0.44 | 8.74e-11 | 2.94e-06 |
| Malignant epithelial | C1 Das ECs | TRAF4 | 0.411 | 0.479 | 0.399 | 3.03e-09 | 0.000102 |
| Malignant epithelial | C1 Das ECs | MDK | 0.41 | 0.931 | 0.782 | 1.15e-49 | 3.88e-45 |
| Malignant epithelial | C1 Das ECs | DDRGK1 | 0.41 | 0.65 | 0.54 | 3.78e-16 | 1.27e-11 |
| Malignant epithelial | C1 Das ECs | MRPS35 | 0.41 | 0.463 | 0.379 | 3e-10 | 1.01e-05 |
| Malignant epithelial | C1 Das ECs | SNHG8 | 0.409 | 0.907 | 0.827 | 8.23e-35 | 2.77e-30 |
| Malignant epithelial | C1 Das ECs | WBSCR22 | 0.409 | 0.615 | 0.519 | 3.79e-14 | 1.28e-09 |
| Malignant epithelial | C1 Das ECs | GUSB | 0.409 | 0.276 | 0.2 | 1.07e-08 | 0.000362 |
| Malignant epithelial | C1 Das ECs | RFK | 0.408 | 0.282 | 0.206 | 1.29e-08 | 0.000435 |
| Malignant epithelial | C1 Das ECs | NUB1 | 0.408 | 0.374 | 0.304 | 1.19e-07 | 0.004 |
| Malignant epithelial | C1 Das ECs | ECSIT | 0.406 | 0.377 | 0.307 | 1.45e-07 | 0.00489 |
| Malignant epithelial | C1 Das ECs | AK3 | 0.405 | 0.407 | 0.333 | 8.72e-08 | 0.00294 |
| Malignant epithelial | C1 Das ECs | MT-ND3 | 0.403 | 0.998 | 0.97 | 1.82e-41 | 6.14e-37 |
| Malignant epithelial | C1 Das ECs | GSS | 0.403 | 0.394 | 0.314 | 3.14e-08 | 0.00106 |
| Malignant epithelial | C1 Das ECs | CST3 | 0.402 | 0.988 | 0.972 | 8.32e-44 | 2.8e-39 |
| Malignant epithelial | C1 Das ECs | SF3B5 | 0.4 | 0.851 | 0.742 | 6.41e-28 | 2.16e-23 |
| Malignant epithelial | C1 Das ECs | MT-ND2 | 0.399 | 1.0 | 0.993 | 6.46e-55 | 2.18e-50 |
| Malignant epithelial | C1 Das ECs | MYC | 0.398 | 0.589 | 0.478 | 6.54e-15 | 2.2e-10 |
| Malignant epithelial | C1 Das ECs | PPP1R14D | 0.398 | 0.467 | 0.343 | 5.28e-14 | 1.78e-09 |
| Malignant epithelial | C1 Das ECs | RPL21 | 0.397 | 0.997 | 0.989 | 2.94e-68 | 9.92e-64 |
| Malignant epithelial | C1 Das ECs | ZNF593 | 0.397 | 0.565 | 0.473 | 4.68e-12 | 1.58e-07 |
| Malignant epithelial | C1 Das ECs | TMEM70 | 0.397 | 0.354 | 0.279 | 6.96e-08 | 0.00234 |
| Malignant epithelial | C1 Das ECs | AURKAIP1 | 0.396 | 0.837 | 0.747 | 3.69e-25 | 1.24e-20 |
| Malignant epithelial | C1 Das ECs | POLR2J | 0.396 | 0.82 | 0.73 | 1.4e-22 | 4.72e-18 |
| Malignant epithelial | C1 Das ECs | FBXL15 | 0.396 | 0.398 | 0.331 | 1.47e-07 | 0.00496 |
| Malignant epithelial | C1 Das ECs | FIS1 | 0.395 | 0.852 | 0.788 | 3.63e-23 | 1.22e-18 |
| Malignant epithelial | C1 Das ECs | SUCLG2 | 0.395 | 0.484 | 0.411 | 4.16e-09 | 0.00014 |
| Malignant epithelial | C1 Das ECs | ZNHIT1 | 0.394 | 0.868 | 0.777 | 2.98e-27 | 1e-22 |
| Malignant epithelial | C1 Das ECs | DPM3 | 0.394 | 0.574 | 0.476 | 4.39e-13 | 1.48e-08 |
| Malignant epithelial | C1 Das ECs | KMT2E | 0.393 | 0.619 | 0.489 | 1.26e-17 | 4.26e-13 |
| Malignant epithelial | C1 Das ECs | EIF2S3 | 0.393 | 0.638 | 0.535 | 1.22e-14 | 4.1e-10 |
| Malignant epithelial | C1 Das ECs | MRPS26 | 0.392 | 0.594 | 0.496 | 9.17e-13 | 3.09e-08 |
| Malignant epithelial | C1 Das ECs | TKT | 0.391 | 0.76 | 0.679 | 4.92e-17 | 1.66e-12 |
| Malignant epithelial | C1 Das ECs | ARPC4 | 0.388 | 0.58 | 0.484 | 2.94e-14 | 9.91e-10 |
| Malignant epithelial | C1 Das ECs | ALG3 | 0.388 | 0.439 | 0.367 | 2.58e-07 | 0.00868 |
| Malignant epithelial | C1 Das ECs | IGFBP4 | 0.387 | 0.353 | 0.27 | 1.13e-08 | 0.000381 |
| Malignant epithelial | C1 Das ECs | YIF1A | 0.386 | 0.657 | 0.562 | 1.95e-15 | 6.55e-11 |
| Malignant epithelial | C1 Das ECs | GRHPR | 0.386 | 0.591 | 0.506 | 3.3e-11 | 1.11e-06 |
| Malignant epithelial | C1 Das ECs | XRN2 | 0.386 | 0.506 | 0.419 | 8.59e-10 | 2.9e-05 |
| Malignant epithelial | C1 Das ECs | COX14 | 0.385 | 0.697 | 0.605 | 5.21e-14 | 1.76e-09 |
| Malignant epithelial | C1 Das ECs | AMN | 0.385 | 0.454 | 0.334 | 1.05e-13 | 3.53e-09 |
| Malignant epithelial | C1 Das ECs | LSM8 | 0.383 | 0.762 | 0.666 | 4.64e-18 | 1.56e-13 |
| Malignant epithelial | C1 Das ECs | TIMM17B | 0.383 | 0.628 | 0.551 | 1.19e-11 | 4.02e-07 |
| Malignant epithelial | C1 Das ECs | SRPK1 | 0.383 | 0.484 | 0.409 | 1.61e-08 | 0.000542 |
| Malignant epithelial | C1 Das ECs | SIRT7 | 0.381 | 0.386 | 0.313 | 4.99e-07 | 0.0168 |
| Malignant epithelial | C1 Das ECs | DCTN3 | 0.38 | 0.587 | 0.497 | 1.64e-11 | 5.52e-07 |
| Malignant epithelial | C1 Das ECs | GCHFR | 0.38 | 0.377 | 0.297 | 4.21e-08 | 0.00142 |
| Malignant epithelial | C1 Das ECs | HNRNPH2 | 0.38 | 0.386 | 0.324 | 1.45e-06 | 0.049 |
| Malignant epithelial | C1 Das ECs | NBEAL1 | 0.378 | 0.942 | 0.873 | 1.84e-36 | 6.19e-32 |
| Malignant epithelial | C1 Das ECs | LRRFIP2 | 0.378 | 0.349 | 0.285 | 9.35e-07 | 0.0315 |
| Malignant epithelial | C1 Das ECs | NABP2 | 0.378 | 0.339 | 0.275 | 1.35e-06 | 0.0456 |
| Malignant epithelial | C1 Das ECs | ANP32B | 0.377 | 0.86 | 0.788 | 7.66e-24 | 2.58e-19 |
| Malignant epithelial | C1 Das ECs | EIF2A | 0.376 | 0.549 | 0.461 | 3.88e-10 | 1.31e-05 |
| Malignant epithelial | C1 Das ECs | ATPIF1 | 0.375 | 0.875 | 0.766 | 3.47e-25 | 1.17e-20 |
| Malignant epithelial | C1 Das ECs | EIF3K | 0.374 | 0.927 | 0.854 | 1.38e-33 | 4.66e-29 |
| Malignant epithelial | C1 Das ECs | COMMD7 | 0.374 | 0.475 | 0.4 | 4.24e-08 | 0.00143 |
| Malignant epithelial | C1 Das ECs | SMDT1 | 0.373 | 0.78 | 0.731 | 1.6e-16 | 5.38e-12 |
| Malignant epithelial | C1 Das ECs | RER1 | 0.373 | 0.589 | 0.514 | 3.78e-12 | 1.27e-07 |
| Malignant epithelial | C1 Das ECs | BAD | 0.372 | 0.438 | 0.366 | 4e-08 | 0.00135 |
| Malignant epithelial | C1 Das ECs | MRPL36 | 0.371 | 0.67 | 0.593 | 5.07e-11 | 1.71e-06 |
| Malignant epithelial | C1 Das ECs | RPL12 | 0.369 | 1.0 | 0.999 | 2.57e-69 | 8.66e-65 |
| Malignant epithelial | C1 Das ECs | ERGIC3 | 0.369 | 0.848 | 0.754 | 6.47e-24 | 2.18e-19 |

| compartment | subcluster | gene | avg_log2FC | pct.1 | pct.2 | p_val | p_val_adj |
|---|---|---|---|---|---|---|---|
| Malignant epithelial | C1 Das ECs | TSPAN15 | 0.369 | 0.443 | 0.354 | 2.36e-09 | 7.95e-05 |
| Malignant epithelial | C1 Das ECs | MTPN | 0.367 | 0.52 | 0.446 | 5.79e-08 | 0.00195 |
| Malignant epithelial | C1 Das ECs | NONO | 0.366 | 0.687 | 0.61 | 4.74e-13 | 1.6e-08 |
| Malignant epithelial | C1 Das ECs | ISCA1 | 0.366 | 0.407 | 0.331 | 4.21e-08 | 0.00142 |
| Malignant epithelial | C1 Das ECs | WTAP | 0.363 | 0.476 | 0.375 | 4.04e-12 | 1.36e-07 |
| Malignant epithelial | C1 Das ECs | VPS51 | 0.363 | 0.434 | 0.36 | 7.31e-08 | 0.00246 |
| Malignant epithelial | C1 Das ECs | RPL18 | 0.362 | 1.0 | 0.997 | 1.22e-95 | 4.09e-91 |
| Malignant epithelial | C1 Das ECs | MT-CO1 | 0.362 | 1.0 | 0.996 | 6.550000000000001e-47 | 2.21e-42 |
| Malignant epithelial | C1 Das ECs | GTF3A | 0.362 | 0.74 | 0.592 | 1.8e-27 | 6.079999999999999e-23 |
| Malignant epithelial | C1 Das ECs | DAP | 0.361 | 0.438 | 0.328 | 3.79e-13 | 1.28e-08 |
| Malignant epithelial | C1 Das ECs | EIF5 | 0.36 | 0.785 | 0.685 | 6.05e-19 | 2.04e-14 |
| Malignant epithelial | C1 Das ECs | PSMD3 | 0.36 | 0.418 | 0.348 | 2.01e-07 | 0.00677 |
| Malignant epithelial | C1 Das ECs | UQCRFS1 | 0.357 | 0.722 | 0.633 | 8.53e-16 | 2.87e-11 |
| Malignant epithelial | C1 Das ECs | ATP6V0D1 | 0.357 | 0.432 | 0.35 | 1.48e-08 | 0.000499 |
| Malignant epithelial | C1 Das ECs | WDR13 | 0.357 | 0.419 | 0.357 | 8.67e-07 | 0.0292 |
| Malignant epithelial | C1 Das ECs | RPL23A | 0.356 | 0.999 | 0.995 | 2.35e-75 | 7.92e-71 |
| Malignant epithelial | C1 Das ECs | PARK7 | 0.356 | 0.795 | 0.74 | 8.73e-15 | 2.94e-10 |
| Malignant epithelial | C1 Das ECs | MTIF3 | 0.356 | 0.524 | 0.405 | 1.13e-13 | 3.82e-09 |
| Malignant epithelial | C1 Das ECs | CAMLG | 0.356 | 0.606 | 0.519 | 4.96e-11 | 1.67e-06 |
| Malignant epithelial | C1 Das ECs | MYO10 | 0.356 | 0.354 | 0.272 | 9.04e-09 | 0.000304 |
| Malignant epithelial | C1 Das ECs | C19orf48 | 0.355 | 0.556 | 0.47 | 1.7e-09 | 5.73e-05 |
| Malignant epithelial | C1 Das ECs | HIST1H2AC | 0.355 | 0.422 | 0.346 | 2.03e-07 | 0.00684 |
| Malignant epithelial | C1 Das ECs | NDUFV1 | 0.354 | 0.531 | 0.458 | 9.59e-10 | 3.23e-05 |
| Malignant epithelial | C1 Das ECs | MEST | 0.353 | 0.339 | 0.271 | 9e-07 | 0.0303 |
| Malignant epithelial | C1 Das ECs | FAM60A | 0.352 | 0.491 | 0.411 | 2.5e-08 | 0.000843 |
| Malignant epithelial | C1 Das ECs | RPS2 | 0.351 | 1.0 | 0.998 | 6.29e-57 | 2.12e-52 |
| Malignant epithelial | C1 Das ECs | NAPA | 0.351 | 0.433 | 0.361 | 6.78e-08 | 0.00229 |
| Malignant epithelial | C1 Das ECs | NOP16 | 0.351 | 0.434 | 0.366 | 1.13e-06 | 0.0382 |
| Malignant epithelial | C1 Das ECs | BRI3 | 0.35 | 0.708 | 0.599 | 1.51e-19 | 5.08e-15 |
| Malignant epithelial | C1 Das ECs | ITM2B | 0.349 | 0.878 | 0.753 | 6.77e-27 | 2.28e-22 |
| Malignant epithelial | C1 Das ECs | CPNE1 | 0.349 | 0.601 | 0.516 | 3.7e-10 | 1.25e-05 |
| Malignant epithelial | C1 Das ECs | ETHE1 | 0.347 | 0.677 | 0.575 | 5.08e-15 | 1.71e-10 |
| Malignant epithelial | C1 Das ECs | TMEM179B | 0.346 | 0.571 | 0.489 | 3.96e-09 | 0.000134 |
| Malignant epithelial | C1 Das ECs | EIF2AK1 | 0.345 | 0.451 | 0.383 | 1.28e-06 | 0.043 |
| Malignant epithelial | C1 Das ECs | MAL2 | 0.343 | 0.694 | 0.582 | 3.78e-18 | 1.28e-13 |
| Malignant epithelial | C1 Das ECs | SYF2 | 0.343 | 0.675 | 0.583 | 6.74e-11 | 2.27e-06 |
| Malignant epithelial | C1 Das ECs | TMEM248 | 0.343 | 0.466 | 0.398 | 2.39e-07 | 0.00807 |
| Malignant epithelial | C1 Das ECs | EIF1AX | 0.342 | 0.773 | 0.72 | 9.01e-15 | 3.04e-10 |
| Malignant epithelial | C1 Das ECs | NDUFS7 | 0.341 | 0.728 | 0.648 | 7.94e-13 | 2.67e-08 |
| Malignant epithelial | C1 Das ECs | MRPS18B | 0.341 | 0.521 | 0.435 | 8.7e-09 | 0.000293 |
| Malignant epithelial | C1 Das ECs | TAOK3 | 0.341 | 0.425 | 0.357 | 1.46e-06 | 0.0491 |
| Malignant epithelial | C1 Das ECs | FABP5 | 0.34 | 0.645 | 0.526 | 7.52e-16 | 2.53e-11 |
| Malignant epithelial | C1 Das ECs | STRA13 | 0.34 | 0.633 | 0.541 | 2.07e-11 | 6.97e-07 |
| Malignant epithelial | C1 Das ECs | CLNS1A | 0.34 | 0.651 | 0.595 | 1.81e-10 | 6.1e-06 |
| Malignant epithelial | C1 Das ECs | GSDMD | 0.34 | 0.619 | 0.527 | 2.42e-10 | 8.16e-06 |
| Malignant epithelial | C1 Das ECs | RPL11 | 0.339 | 1.0 | 0.997 | 1.75e-51 | 5.89e-47 |
| Malignant epithelial | C1 Das ECs | UCHL3 | 0.338 | 0.486 | 0.381 | 2.11e-11 | 7.13e-07 |
| Malignant epithelial | C1 Das ECs | NDUFA10 | 0.338 | 0.574 | 0.488 | 4.46e-10 | 1.5e-05 |
| Malignant epithelial | C1 Das ECs | RPL8 | 0.335 | 1.0 | 1.0 | 8.11e-74 | 2.73e-69 |
| Malignant epithelial | C1 Das ECs | ACTN1 | 0.335 | 0.46 | 0.356 | 8.83e-11 | 2.98e-06 |
| Malignant epithelial | C1 Das ECs | SSNA1 | 0.335 | 0.702 | 0.633 | 1.22e-10 | 4.11e-06 |
| Malignant epithelial | C1 Das ECs | RPS10 | 0.334 | 0.996 | 0.982 | 5.27e-50 | 1.78e-45 |
| Malignant epithelial | C1 Das ECs | MINOS1 | 0.333 | 0.833 | 0.76 | 4.69e-15 | 1.58e-10 |
| Malignant epithelial | C1 Das ECs | ABHD12 | 0.333 | 0.469 | 0.388 | 1.53e-08 | 0.000517 |
| Malignant epithelial | C1 Das ECs | SSBP1 | 0.331 | 0.887 | 0.817 | 6.51e-20 | 2.19e-15 |
| Malignant epithelial | C1 Das ECs | COMMD6 | 0.33 | 0.931 | 0.877 | 1.11e-26 | 3.74e-22 |
| Malignant epithelial | C1 Das ECs | EFHD2 | 0.33 | 0.52 | 0.429 | 2.39e-10 | 8.05e-06 |
| Malignant epithelial | C1 Das ECs | LGALS3 | 0.329 | 0.979 | 0.967 | 1.24e-29 | 4.19e-25 |
| Malignant epithelial | C1 Das ECs | SND1 | 0.328 | 0.393 | 0.324 | 1.46e-06 | 0.0491 |
| Malignant epithelial | C1 Das ECs | CD74 | 0.327 | 0.933 | 0.887 | 1.4e-19 | 4.71e-15 |
| Malignant epithelial | C1 Das ECs | MARCKSL1 | 0.325 | 0.864 | 0.692 | 4.98e-31 | 1.68e-26 |
| Malignant epithelial | C1 Das ECs | WDR45B | 0.325 | 0.421 | 0.346 | 1.71e-07 | 0.00576 |
| Malignant epithelial | C1 Das ECs | RAB4A | 0.325 | 0.484 | 0.415 | 4.07e-07 | 0.0137 |
| Malignant epithelial | C1 Das ECs | CHMP4B | 0.324 | 0.886 | 0.768 | 5.27e-23 | 1.77e-18 |

| compartment | subcluster | gene | avg_log2FC | pct.1 | pct.2 | p_val | p_val_adj |
|---|---|---|---|---|---|---|---|
| Malignant epithelial | C1 Das ECs | MGLL | 0.323 | 0.332 | 0.244 | 4.57e-10 | 1.54e-05 |
| Malignant epithelial | C1 Das ECs | EAPP | 0.323 | 0.445 | 0.371 | 3.05e-07 | 0.0103 |
| Malignant epithelial | C1 Das ECs | CD164 | 0.322 | 0.817 | 0.717 | 1.84e-25 | 6.21e-21 |
| Malignant epithelial | C1 Das ECs | C6orf48 | 0.322 | 0.89 | 0.839 | 2.84e-16 | 9.55e-12 |
| Malignant epithelial | C1 Das ECs | HDDC2 | 0.322 | 0.477 | 0.403 | 2.63e-07 | 0.00886 |
| Malignant epithelial | C1 Das ECs | PPA1 | 0.321 | 0.94 | 0.886 | 2.25e-25 | 7.580000000000001e-21 |
| Malignant epithelial | C1 Das ECs | TMC5 | 0.321 | 0.59 | 0.497 | 6.57e-09 | 0.000221 |
| Malignant epithelial | C1 Das ECs | PSMD7 | 0.32 | 0.586 | 0.476 | 4.62e-12 | 1.56e-07 |
| Malignant epithelial | C1 Das ECs | IFNGR2 | 0.319 | 0.51 | 0.43 | 1.49e-08 | 0.000501 |
| Malignant epithelial | C1 Das ECs | CLTA | 0.317 | 0.915 | 0.859 | 5.33e-24 | 1.79e-19 |
| Malignant epithelial | C1 Das ECs | YDJC | 0.316 | 0.529 | 0.454 | 7.81e-08 | 0.00263 |
| Malignant epithelial | C1 Das ECs | CUTA | 0.315 | 0.899 | 0.829 | 5e-21 | 1.68e-16 |
| Malignant epithelial | C1 Das ECs | PHF14 | 0.315 | 0.641 | 0.547 | 1.48e-11 | 5e-07 |
| Malignant epithelial | C1 Das ECs | SMAGP | 0.315 | 0.619 | 0.542 | 4.46e-10 | 1.5e-05 |
| Malignant epithelial | C1 Das ECs | ATP5G3 | 0.314 | 0.919 | 0.865 | 1.31e-20 | 4.42e-16 |
| Malignant epithelial | C1 Das ECs | NDUFB9 | 0.313 | 0.919 | 0.867 | 4.429999999999999e-23 | 1.49e-18 |
| Malignant epithelial | C1 Das ECs | ATP5C1 | 0.312 | 0.777 | 0.671 | 8.78e-15 | 2.96e-10 |
| Malignant epithelial | C1 Das ECs | CD47 | 0.312 | 0.606 | 0.508 | 1.04e-11 | 3.51e-07 |
| Malignant epithelial | C1 Das ECs | PYCARD | 0.311 | 0.819 | 0.75 | 1.07e-13 | 3.59e-09 |
| Malignant epithelial | C1 Das ECs | NTMT1 | 0.311 | 0.483 | 0.412 | 1.13e-06 | 0.0381 |
| Malignant epithelial | C1 Das ECs | RPS9 | 0.309 | 0.999 | 0.993 | 4.96e-52 | 1.67e-47 |
| Malignant epithelial | C1 Das ECs | RWDD1 | 0.309 | 0.635 | 0.563 | 2.94e-09 | 9.9e-05 |
| Malignant epithelial | C1 Das ECs | BET1 | 0.302 | 0.382 | 0.311 | 1.4e-06 | 0.0472 |
| Malignant epithelial | C1 Das ECs | NHP2 | 0.301 | 0.778 | 0.697 | 7.39e-13 | 2.49e-08 |
| Malignant epithelial | C1 Das ECs | SF3B2 | 0.3 | 0.66 | 0.588 | 9.25e-09 | 0.000312 |
| Malignant epithelial | C1 Das ECs | PHPT1 | 0.298 | 0.817 | 0.76 | 7.47e-15 | 2.52e-10 |
| Malignant epithelial | C1 Das ECs | COPS6 | 0.298 | 0.706 | 0.637 | 1.73e-10 | 5.82e-06 |
| Malignant epithelial | C1 Das ECs | IMPDH2 | 0.297 | 0.63 | 0.552 | 9.08e-09 | 0.000306 |
| Malignant epithelial | C1 Das ECs | SBDS | 0.295 | 0.797 | 0.702 | 1.01e-17 | 3.4e-13 |
| Malignant epithelial | C1 Das ECs | PSMB8 | 0.295 | 0.796 | 0.72 | 1.42e-12 | 4.8e-08 |
| Malignant epithelial | C1 Das ECs | ITPA | 0.295 | 0.587 | 0.499 | 2.12e-09 | 7.16e-05 |
| Malignant epithelial | C1 Das ECs | SQSTM1 | 0.291 | 0.878 | 0.779 | 2.24e-21 | 7.54e-17 |
| Malignant epithelial | C1 Das ECs | EIF3E | 0.29 | 0.93 | 0.893 | 1.23e-16 | 4.15e-12 |
| Malignant epithelial | C1 Das ECs | ENSA | 0.29 | 0.675 | 0.605 | 4.38e-09 | 0.000148 |
| Malignant epithelial | C1 Das ECs | C11orf1 | 0.29 | 0.486 | 0.416 | 3.12e-07 | 0.0105 |
| Malignant epithelial | C1 Das ECs | ATP5B | 0.287 | 0.797 | 0.73 | 2.24e-15 | 7.53e-11 |
| Malignant epithelial | C1 Das ECs | KLF5 | 0.286 | 0.701 | 0.572 | 2.25e-15 | 7.58e-11 |
| Malignant epithelial | C1 Das ECs | CMC1 | 0.286 | 0.405 | 0.332 | 1.04e-06 | 0.035 |
| Malignant epithelial | C1 Das ECs | ENO1 | 0.284 | 0.847 | 0.784 | 2.78e-12 | 9.36e-08 |
| Malignant epithelial | C1 Das ECs | LINC00998 | 0.284 | 0.531 | 0.45 | 8.87e-08 | 0.00299 |
| Malignant epithelial | C1 Das ECs | RPL13A | 0.283 | 1.0 | 0.999 | 1.49e-47 | 5.02e-43 |
| Malignant epithelial | C1 Das ECs | RARRES1 | 0.282 | 0.336 | 0.225 | 3.15e-14 | 1.06e-09 |
| Malignant epithelial | C1 Das ECs | DUSP2 | 0.281 | 0.42 | 0.306 | 1.24e-12 | 4.19e-08 |
| Malignant epithelial | C1 Das ECs | IMP3 | 0.28 | 0.526 | 0.465 | 8.74e-07 | 0.0295 |
| Malignant epithelial | C1 Das ECs | RPS19 | 0.279 | 1.0 | 1.0 | 7.4e-47 | 2.49e-42 |
| Malignant epithelial | C1 Das ECs | PIGT | 0.278 | 0.51 | 0.425 | 1.01e-08 | 0.000339 |
| Malignant epithelial | C1 Das ECs | KRT10 | 0.277 | 0.778 | 0.687 | 3.89e-13 | 1.31e-08 |
| Malignant epithelial | C1 Das ECs | RGS1 | 0.276 | 0.507 | 0.38 | 8.51e-15 | 2.87e-10 |
| Malignant epithelial | C1 Das ECs | SDHD | 0.275 | 0.562 | 0.5 | 5.59e-07 | 0.0188 |
| Malignant epithelial | C1 Das ECs | DECR1 | 0.275 | 0.551 | 0.484 | 7.59e-07 | 0.0256 |
| Malignant epithelial | C1 Das ECs | BUD31 | 0.274 | 0.785 | 0.7 | 8.71e-15 | 2.93e-10 |
| Malignant epithelial | C1 Das ECs | ATP1A1 | 0.274 | 0.622 | 0.528 | 1.18e-11 | 3.97e-07 |
| Malignant epithelial | C1 Das ECs | VIL1 | 0.273 | 0.441 | 0.353 | 8.08e-08 | 0.00272 |
| Malignant epithelial | C1 Das ECs | CCT6A | 0.272 | 0.688 | 0.589 | 9.82e-12 | 3.31e-07 |
| Malignant epithelial | C1 Das ECs | COX5B | 0.271 | 0.989 | 0.972 | 2.49e-31 | 8.4e-27 |
| Malignant epithelial | C1 Das ECs | TFF1 | 0.271 | 0.253 | 0.146 | 3.26e-17 | 1.1e-12 |
| Malignant epithelial | C1 Das ECs | MISP | 0.271 | 0.688 | 0.574 | 1.84e-12 | 6.21e-08 |
| Malignant epithelial | C1 Das ECs | MRPS12 | 0.271 | 0.629 | 0.544 | 6.83e-10 | 2.3e-05 |
| Malignant epithelial | C1 Das ECs | GBP2 | 0.271 | 0.333 | 0.259 | 9.71e-07 | 0.0327 |
| Malignant epithelial | C1 Das ECs | SEPW1 | 0.269 | 0.885 | 0.817 | 1.73e-15 | 5.82e-11 |
| Malignant epithelial | C1 Das ECs | UBE2L6 | 0.269 | 0.58 | 0.488 | 5.63e-09 | 0.00019 |
| Malignant epithelial | C1 Das ECs | NAPRT | 0.269 | 0.685 | 0.617 | 6.98e-08 | 0.00235 |
| Malignant epithelial | C1 Das ECs | PNPLA2 | 0.269 | 0.404 | 0.325 | 1.85e-07 | 0.00624 |
| Malignant epithelial | C1 Das ECs | IP6K2 | 0.266 | 0.52 | 0.457 | 9.7e-07 | 0.0327 |

| compartment | subcluster | gene | avg_log2FC | pct.1 | pct.2 | p_val | p_val_adj |
|---|---|---|---|---|---|---|---|
| Malignant epithelial | C1 Das ECs | KXD1 | 0.266 | 0.571 | 0.52 | 1.25e-06 | 0.0422 |
| Malignant epithelial | C1 Das ECs | EEF2 | 0.265 | 0.955 | 0.94 | 1.25e-20 | 4.22e-16 |
| Malignant epithelial | C1 Das ECs | HEBP2 | 0.264 | 0.804 | 0.727 | 3.16e-14 | 1.06e-09 |
| Malignant epithelial | C1 Das ECs | PSME2 | 0.26 | 0.896 | 0.828 | 2.41e-13 | 8.11e-09 |
| Malignant epithelial | C1 Das ECs | MT-ATP6 | 0.259 | 0.999 | 0.979 | 1.79e-31 | 6.040000000000001e-27 |
| Malignant epithelial | C1 Das ECs | HLA-C | 0.259 | 0.967 | 0.942 | 1.43e-27 | 4.809999999999999e-23 |
| Malignant epithelial | C1 Das ECs | C14orf166 | 0.259 | 0.813 | 0.742 | 8.35e-12 | 2.81e-07 |
| Malignant epithelial | C1 Das ECs | PLA2G16 | 0.259 | 0.707 | 0.604 | 4.24e-11 | 1.43e-06 |
| Malignant epithelial | C1 Das ECs | EIF3F | 0.259 | 0.84 | 0.795 | 4.11e-10 | 1.39e-05 |
| Malignant epithelial | C1 Das ECs | PSMB7 | 0.257 | 0.777 | 0.714 | 1.39e-11 | 4.7e-07 |
| Malignant epithelial | C1 Das ECs | RAB25 | 0.257 | 0.748 | 0.679 | 8.02e-09 | 0.00027 |
| Malignant epithelial | C1 Das ECs | TST | 0.257 | 0.418 | 0.338 | 2.17e-07 | 0.00732 |
| Malignant epithelial | C1 Das ECs | CHCHD2 | 0.256 | 0.981 | 0.952 | 1.98e-39 | 6.68e-35 |
| Malignant epithelial | C1 Das ECs | TMEM219 | 0.255 | 0.74 | 0.683 | 1.39e-08 | 0.000467 |
| Malignant epithelial | C1 Das ECs | MAF1 | 0.254 | 0.625 | 0.54 | 1.24e-08 | 0.000418 |
| Malignant epithelial | C1 Das ECs | MRPL57 | 0.253 | 0.807 | 0.726 | 6.74e-10 | 2.27e-05 |
| Malignant epithelial | C1 Das ECs | LSM7 | 0.252 | 0.921 | 0.867 | 7.2e-17 | 2.43e-12 |
| Malignant epithelial | C1 Das ECs | EID1 | 0.252 | 0.658 | 0.587 | 5.23e-08 | 0.00176 |
| Malignant epithelial | C1 Das ECs | SHFM1 | 0.251 | 0.964 | 0.944 | 4.47e-21 | 1.51e-16 |
| Malignant epithelial | C1 Das ECs | SYNGR2 | 0.251 | 0.791 | 0.709 | 5.86e-11 | 1.97e-06 |
| Malignant epithelial | C2 WNT ECs | SEMA3F | 6.061 | 0.185 | 0.005 | 8.720000000000001e-156 | 2.94e-151 |
| Malignant epithelial | C2 WNT ECs | PRSS2 | 6.057 | 0.115 | 0.005 | 6.62e-84 | 2.23e-79 |
| Malignant epithelial | C2 WNT ECs | WIF1 | 6.038 | 0.143 | 0.003 | 7.630000000000001e-123 | 2.57e-118 |
| Malignant epithelial | C2 WNT ECs | DES | 5.933 | 0.11 | 0.002 | 1.33e-98 | 4.49e-94 |
| Malignant epithelial | C2 WNT ECs | APCDD1 | 5.84 | 0.187 | 0.003 | 1.55e-169 | 5.23e-165 |
| Malignant epithelial | C2 WNT ECs | INHBB | 5.745 | 0.211 | 0.004 | 2.2e-187 | 7.4e-183 |
| Malignant epithelial | C2 WNT ECs | NKD1 | 5.742 | 0.617 | 0.021 | 0.0 | 0.0 |
| Malignant epithelial | C2 WNT ECs | PARD6G-AS1 | 5.657 | 0.165 | 0.005 | 1.61e-135 | 5.44e-131 |
| Malignant epithelial | C2 WNT ECs | MME | 5.576 | 0.39 | 0.007 | 0.0 | 0.0 |
| Malignant epithelial | C2 WNT ECs | EPHB6 | 5.54 | 0.35 | 0.009 | 2.23e-300 | 7.52e-296 |
| Malignant epithelial | C2 WNT ECs | FGF19 | 5.461 | 0.304 | 0.018 | 1.11e-211 | 3.74e-207 |
| Malignant epithelial | C2 WNT ECs | NR0B2 | 5.451 | 0.157 | 0.004 | 4.93e-131 | 1.66e-126 |
| Malignant epithelial | C2 WNT ECs | GATA2 | 5.432 | 0.11 | 0.002 | 1.35e-95 | 4.54e-91 |
| Malignant epithelial | C2 WNT ECs | SP9 | 5.324 | 0.11 | 0.002 | 4.56e-100 | 1.54e-95 |
| Malignant epithelial | C2 WNT ECs | NOTUM | 5.285 | 0.497 | 0.024 | 0.0 | 0.0 |
| Malignant epithelial | C2 WNT ECs | CES1 | 5.251 | 0.304 | 0.015 | 4.37e-225 | 1.47e-220 |
| Malignant epithelial | C2 WNT ECs | SMOC2 | 5.239 | 0.169 | 0.015 | 3.52e-98 | 1.19e-93 |
| Malignant epithelial | C2 WNT ECs | CTD-2147F2.1 | 5.197 | 0.202 | 0.005 | 1.46e-170 | 4.91e-166 |
| Malignant epithelial | C2 WNT ECs | TSPAN32 | 5.093 | 0.202 | 0.008 | 3.68e-157 | 1.24e-152 |
| Malignant epithelial | C2 WNT ECs | DUSP27 | 5.078 | 0.214 | 0.004 | 2.49e-188 | 8.37e-184 |
| Malignant epithelial | C2 WNT ECs | KRTAP3-1 | 5.038 | 0.11 | 0.005 | 6.27e-81 | 2.11e-76 |
| Malignant epithelial | C2 WNT ECs | AC006262.5 | 5.013 | 0.422 | 0.026 | 4.03e-296 | 1.36e-291 |
| Malignant epithelial | C2 WNT ECs | RP11-11N5.3 | 4.98 | 0.223 | 0.005 | 4.6e-192 | 1.55e-187 |
| Malignant epithelial | C2 WNT ECs | DPYSL3 | 4.98 | 0.191 | 0.005 | 3.02e-162 | 1.02e-157 |
| Malignant epithelial | C2 WNT ECs | KLRC2 | 4.857 | 0.249 | 0.01 | 2.99e-194 | 1.01e-189 |
| Malignant epithelial | C2 WNT ECs | SOX15 | 4.838 | 0.231 | 0.004 | 6.89e-206 | 2.32e-201 |
| Malignant epithelial | C2 WNT ECs | PDLIM4 | 4.781 | 0.651 | 0.025 | 0.0 | 0.0 |
| Malignant epithelial | C2 WNT ECs | DLX3 | 4.765 | 0.103 | 0.002 | 1.02e-91 | 3.43e-87 |
| Malignant epithelial | C2 WNT ECs | ZNF83 | 4.685 | 0.326 | 0.013 | 8.07e-252 | 2.72e-247 |
| Malignant epithelial | C2 WNT ECs | A4GALT | 4.638 | 0.153 | 0.004 | 1.96e-126 | 6.61e-122 |
| Malignant epithelial | C2 WNT ECs | PCSK1N | 4.621 | 0.206 | 0.017 | 5.4e-122 | 1.82e-117 |
| Malignant epithelial | C2 WNT ECs | PCP4 | 4.572 | 0.116 | 0.004 | 4.86e-92 | 1.64e-87 |
| Malignant epithelial | C2 WNT ECs | ID4 | 4.569 | 0.383 | 0.023 | 3.34e-265 | 1.13e-260 |
| Malignant epithelial | C2 WNT ECs | KCNMB2-AS1 | 4.556 | 0.175 | 0.005 | 9.79e-143 | 3.3e-138 |
| Malignant epithelial | C2 WNT ECs | CEL | 4.536 | 0.284 | 0.015 | 1.86e-204 | 6.270000000000001e-200 |
| Malignant epithelial | C2 WNT ECs | CRYAB | 4.53 | 0.341 | 0.026 | 4.67e-213 | 1.57e-208 |
| Malignant epithelial | C2 WNT ECs | MLF1 | 4.508 | 0.185 | 0.01 | 2.27e-131 | 7.65e-127 |
| Malignant epithelial | C2 WNT ECs | WDR72 | 4.487 | 0.415 | 0.015 | 0.0 | 0.0 |
| Malignant epithelial | C2 WNT ECs | DACH1 | 4.45 | 0.115 | 0.006 | 7e-82 | 2.36e-77 |
| Malignant epithelial | C2 WNT ECs | RGL3 | 4.432 | 0.101 | 0.004 | 1.32e-78 | 4.46e-74 |
| Malignant epithelial | C2 WNT ECs | VAV3 | 4.419 | 0.214 | 0.015 | 3.78e-134 | 1.27e-129 |
| Malignant epithelial | C2 WNT ECs | HSPA1A | 4.387 | 0.817 | 0.135 | 0.0 | 0.0 |
| Malignant epithelial | C2 WNT ECs | EDAR | 4.384 | 0.138 | 0.006 | 7.160000000000001e-105 | 2.41e-100 |
| Malignant epithelial | C2 WNT ECs | HSPA12A | 4.358 | 0.146 | 0.006 | 2.74e-109 | 9.24e-105 |

| compartment | subcluster | gene | avg_log2FC | pct.1 | pct.2 | p_val | p_val_adj |
|---|---|---|---|---|---|---|---|
| Malignant epithelial | C2 WNT ECs | RP11-382A20.3 | 4.339 | 0.212 | 0.007 | 4.889999999999999e-172 | 1.65e-167 |
| Malignant epithelial | C2 WNT ECs | ECM1 | 4.303 | 0.342 | 0.049 | 2.76e-151 | 9.289999999999999e-147 |
| Malignant epithelial | C2 WNT ECs | ANKRD22 | 4.273 | 0.254 | 0.011 | 3.08e-193 | 1.04e-188 |
| Malignant epithelial | C2 WNT ECs | SLC38A11 | 4.261 | 0.111 | 0.006 | 5.66e-78 | 1.91e-73 |
| Malignant epithelial | C2 WNT ECs | TDGF1 | 4.207 | 0.187 | 0.012 | 2.26e-123 | 7.619999999999999e-119 |
| Malignant epithelial | C2 WNT ECs | LINC00176 | 4.176 | 0.294 | 0.051 | 5.3e-112 | 1.79e-107 |
| Malignant epithelial | C2 WNT ECs | EVA1A | 4.153 | 0.107 | 0.006 | 3.01e-74 | 1.02e-69 |
| Malignant epithelial | C2 WNT ECs | PTK7 | 4.115 | 0.426 | 0.046 | 2.2e-233 | 7.41e-229 |
| Malignant epithelial | C2 WNT ECs | IGFL4 | 4.109 | 0.107 | 0.005 | 2.64e-79 | 8.89e-75 |
| Malignant epithelial | C2 WNT ECs | FKBP1B | 4.081 | 0.257 | 0.01 | 1.35e-199 | 4.53e-195 |
| Malignant epithelial | C2 WNT ECs | GAPLINC | 4.016 | 0.303 | 0.019 | 2.15e-204 | 7.250000000000001e-200 |
| Malignant epithelial | C2 WNT ECs | RGS17 | 4.015 | 0.11 | 0.006 | 1.03e-76 | 3.46e-72 |
| Malignant epithelial | C2 WNT ECs | DPEP1 | 4.002 | 0.876 | 0.078 | 0.0 | 0.0 |
| Malignant epithelial | C2 WNT ECs | MINPP1 | 4.0 | 0.107 | 0.005 | 4.88e-76 | 1.64e-71 |
| Malignant epithelial | C2 WNT ECs | BAMBI | 3.979 | 0.788 | 0.118 | 0.0 | 0.0 |
| Malignant epithelial | C2 WNT ECs | CAB39L | 3.972 | 0.342 | 0.025 | 4.860000000000001e-217 | 1.64e-212 |
| Malignant epithelial | C2 WNT ECs | RBP1 | 3.95 | 0.133 | 0.007 | 2.24e-93 | 7.55e-89 |
| Malignant epithelial | C2 WNT ECs | SPTSSB | 3.919 | 0.145 | 0.011 | 9.73e-86 | 3.28e-81 |
| Malignant epithelial | C2 WNT ECs | KRT23 | 3.907 | 0.819 | 0.075 | 0.0 | 0.0 |
| Malignant epithelial | C2 WNT ECs | SORBS1 | 3.901 | 0.214 | 0.016 | 2.11e-132 | 7.11e-128 |
| Malignant epithelial | C2 WNT ECs | PLCB1 | 3.9 | 0.173 | 0.012 | 1.11e-108 | 3.73e-104 |
| Malignant epithelial | C2 WNT ECs | PSORS1C2 | 3.884 | 0.184 | 0.01 | 3.5e-126 | 1.18e-121 |
| Malignant epithelial | C2 WNT ECs | LINC00704 | 3.884 | 0.127 | 0.01 | 2.68e-77 | 9.030000000000001e-73 |
| Malignant epithelial | C2 WNT ECs | PTEN | 3.867 | 0.333 | 0.018 | 2.45e-234 | 8.24e-230 |
| Malignant epithelial | C2 WNT ECs | ID3 | 3.866 | 0.645 | 0.099 | 2.06e-310 | 6.94e-306 |
| Malignant epithelial | C2 WNT ECs | ANO6 | 3.866 | 0.137 | 0.006 | 4.99e-99 | 1.68e-94 |
| Malignant epithelial | C2 WNT ECs | IL11 | 3.858 | 0.108 | 0.01 | 2.3e-58 | 7.759999999999999e-54 |
| Malignant epithelial | C2 WNT ECs | TMSB4Y | 3.833 | 0.103 | 0.003 | 2.99e-82 | 1.01e-77 |
| Malignant epithelial | C2 WNT ECs | TAC1 | 3.827 | 0.135 | 0.01 | 1.97e-84 | 6.65e-80 |
| Malignant epithelial | C2 WNT ECs | ORM1 | 3.826 | 0.1 | 0.005 | 6.67e-72 | 2.25e-67 |
| Malignant epithelial | C2 WNT ECs | CASC15 | 3.803 | 0.108 | 0.006 | 3.49e-75 | 1.18e-70 |
| Malignant epithelial | C2 WNT ECs | COL6A1 | 3.802 | 0.287 | 0.029 | 3.8e-154 | 1.28e-149 |
| Malignant epithelial | C2 WNT ECs | SERPIND1 | 3.799 | 0.134 | 0.009 | 6.99e-86 | 2.35e-81 |
| Malignant epithelial | C2 WNT ECs | EPDR1 | 3.787 | 0.18 | 0.009 | 2.43e-129 | 8.200000000000001e-125 |
| Malignant epithelial | C2 WNT ECs | ATAD1 | 3.771 | 0.327 | 0.018 | 6.189999999999999e-229 | 2.08e-224 |
| Malignant epithelial | C2 WNT ECs | LBH | 3.748 | 0.341 | 0.03 | 2.8e-197 | 9.43e-193 |
| Malignant epithelial | C2 WNT ECs | LARP6 | 3.745 | 0.281 | 0.011 | 9.79e-214 | 3.3e-209 |
| Malignant epithelial | C2 WNT ECs | NUTM2A-AS1 | 3.735 | 0.271 | 0.02 | 3.450000000000001e-167 | 1.16e-162 |
| Malignant epithelial | C2 WNT ECs | PROX1 | 3.731 | 0.363 | 0.031 | 3.170000000000001e-213 | 1.07e-208 |
| Malignant epithelial | C2 WNT ECs | GSTM3 | 3.727 | 0.199 | 0.012 | 4.45e-132 | 1.5e-127 |
| Malignant epithelial | C2 WNT ECs | FAM25A | 3.725 | 0.13 | 0.011 | 1.39e-73 | 4.68e-69 |
| Malignant epithelial | C2 WNT ECs | PANK1 | 3.724 | 0.108 | 0.005 | 2.91e-77 | 9.810000000000001e-73 |
| Malignant epithelial | C2 WNT ECs | MAN1A1 | 3.646 | 0.271 | 0.022 | 2.41e-162 | 8.11e-158 |
| Malignant epithelial | C2 WNT ECs | IGFL1 | 3.633 | 0.103 | 0.006 | 3.5e-70 | 1.18e-65 |
| Malignant epithelial | C2 WNT ECs | XXbac-BPG32J3.19 | 3.631 | 0.371 | 0.023 | 1.3e-250 | 4.39e-246 |
| Malignant epithelial | C2 WNT ECs | SLC2A1-AS1 | 3.628 | 0.184 | 0.014 | 1.66e-112 | 5.61e-108 |
| Malignant epithelial | C2 WNT ECs | KREMEN1 | 3.626 | 0.139 | 0.011 | 1.57e-82 | 5.28e-78 |
| Malignant epithelial | C2 WNT ECs | DENND2C | 3.616 | 0.157 | 0.016 | 1.2e-80 | 4.05e-76 |
| Malignant epithelial | C2 WNT ECs | ZBTB7C | 3.613 | 0.21 | 0.018 | 1.17e-120 | 3.95e-116 |
| Malignant epithelial | C2 WNT ECs | C11orf21 | 3.609 | 0.147 | 0.011 | 4.7e-91 | 1.58e-86 |
| Malignant epithelial | C2 WNT ECs | RASD1 | 3.604 | 0.294 | 0.029 | 1.96e-158 | 6.62e-154 |
| Malignant epithelial | C2 WNT ECs | LY6G6C | 3.586 | 0.249 | 0.019 | 2.73e-150 | 9.2e-146 |
| Malignant epithelial | C2 WNT ECs | EPHA4 | 3.563 | 0.1 | 0.008 | 1.55e-58 | 5.22e-54 |
| Malignant epithelial | C2 WNT ECs | TUBB6 | 3.559 | 0.152 | 0.012 | 6.01e-90 | 2.03e-85 |
| Malignant epithelial | C2 WNT ECs | AXIN2 | 3.55 | 0.413 | 0.037 | 1.83e-240 | 6.17e-236 |
| Malignant epithelial | C2 WNT ECs | CGREF1 | 3.547 | 0.104 | 0.004 | 1e-78 | 3.37e-74 |
| Malignant epithelial | C2 WNT ECs | KRT7 | 3.536 | 0.589 | 0.153 | 6.72e-196 | 2.26e-191 |
| Malignant epithelial | C2 WNT ECs | WNT11 | 3.532 | 0.299 | 0.03 | 4.49e-161 | 1.51e-156 |
| Malignant epithelial | C2 WNT ECs | RARRES2 | 3.528 | 0.287 | 0.026 | 4.57e-161 | 1.54e-156 |
| Malignant epithelial | C2 WNT ECs | KLK12 | 3.506 | 0.177 | 0.018 | 1.53e-93 | 5.16e-89 |
| Malignant epithelial | C2 WNT ECs | KRT16 | 3.466 | 0.192 | 0.057 | 1.41e-37 | 4.75e-33 |
| Malignant epithelial | C2 WNT ECs | FN1 | 3.442 | 0.38 | 0.082 | 2.13e-123 | 7.17e-119 |
| Malignant epithelial | C2 WNT ECs | AMACR | 3.44 | 0.15 | 0.015 | 1.06e-79 | 3.57e-75 |
| Malignant epithelial | C2 WNT ECs | MXRA8 | 3.437 | 0.141 | 0.014 | 5.45e-75 | 1.83e-70 |

| compartment | subcluster | gene | avg_log2FC | pct.1 | pct.2 | p_val | p_val_adj |
|---|---|---|---|---|---|---|---|
| Malignant epithelial | C2 WNT ECs | RPL39L | 3.437 | 0.124 | 0.01 | 3.3e-73 | 1.11e-68 |
| Malignant epithelial | C2 WNT ECs | LGR5 | 3.428 | 0.207 | 0.023 | 1.16e-103 | 3.89e-99 |
| Malignant epithelial | C2 WNT ECs | MAP7D2 | 3.426 | 0.158 | 0.012 | 1.41e-96 | 4.77e-92 |
| Malignant epithelial | C2 WNT ECs | SMAD4 | 3.425 | 0.107 | 0.007 | 4.48e-70 | 1.51e-65 |
| Malignant epithelial | C2 WNT ECs | CKMT2 | 3.422 | 0.107 | 0.006 | 5.36e-74 | 1.8e-69 |
| Malignant epithelial | C2 WNT ECs | GOLGA8A | 3.371 | 0.161 | 0.011 | 2.55e-101 | 8.590000000000001e-97 |
| Malignant epithelial | C2 WNT ECs | WFS1 | 3.349 | 0.361 | 0.044 | 5.67e-177 | 1.91e-172 |
| Malignant epithelial | C2 WNT ECs | RAMP2 | 3.345 | 0.169 | 0.02 | 6.93e-81 | 2.34e-76 |
| Malignant epithelial | C2 WNT ECs | C6orf15 | 3.316 | 0.558 | 0.174 | 6.2e-134 | 2.09e-129 |
| Malignant epithelial | C2 WNT ECs | ABCB1 | 3.295 | 0.177 | 0.016 | 2.43e-98 | 8.189999999999999e-94 |
| Malignant epithelial | C2 WNT ECs | LAT | 3.282 | 0.391 | 0.052 | 1.52e-184 | 5.11e-180 |
| Malignant epithelial | C2 WNT ECs | STAMBPL1 | 3.279 | 0.142 | 0.013 | 2.21e-79 | 7.44e-75 |
| Malignant epithelial | C2 WNT ECs | GAL | 3.25 | 0.249 | 0.017 | 2.25e-159 | 7.57e-155 |
| Malignant epithelial | C2 WNT ECs | GPX3 | 3.245 | 0.509 | 0.062 | 1.06e-261 | 3.58e-257 |
| Malignant epithelial | C2 WNT ECs | ITGB7 | 3.24 | 0.204 | 0.024 | 1.91e-98 | 6.43e-94 |
| Malignant epithelial | C2 WNT ECs | FLNC | 3.231 | 0.11 | 0.007 | 7.77e-70 | 2.62e-65 |
| Malignant epithelial | C2 WNT ECs | MFAP2 | 3.227 | 0.327 | 0.038 | 6.26e-162 | 2.11e-157 |
| Malignant epithelial | C2 WNT ECs | MMP23B | 3.224 | 0.137 | 0.012 | 2.13e-75 | 7.18e-71 |
| Malignant epithelial | C2 WNT ECs | MTSS1 | 3.22 | 0.221 | 0.017 | 1.51e-132 | 5.1e-128 |
| Malignant epithelial | C2 WNT ECs | ATP6V1C2 | 3.198 | 0.118 | 0.011 | 7.29e-63 | 2.46e-58 |
| Malignant epithelial | C2 WNT ECs | PRAC1 | 3.182 | 0.43 | 0.027 | 1.06e-287 | 3.56e-283 |
| Malignant epithelial | C2 WNT ECs | GNG4 | 3.167 | 0.382 | 0.022 | 4.28e-262 | 1.44e-257 |
| Malignant epithelial | C2 WNT ECs | FIBCD1 | 3.154 | 0.16 | 0.024 | 4.34e-64 | 1.46e-59 |
| Malignant epithelial | C2 WNT ECs | ANXA6 | 3.149 | 0.472 | 0.065 | 2.95e-220 | 9.949999999999999e-216 |
| Malignant epithelial | C2 WNT ECs | SP5 | 3.107 | 0.288 | 0.033 | 7.499999999999999e-143 | 2.53e-138 |
| Malignant epithelial | C2 WNT ECs | AOAH | 3.097 | 0.24 | 0.021 | 2.94e-134 | 9.92e-130 |
| Malignant epithelial | C2 WNT ECs | SLC2A1 | 3.088 | 0.694 | 0.266 | 1.64e-168 | 5.53e-164 |
| Malignant epithelial | C2 WNT ECs | KISS1 | 3.083 | 0.149 | 0.027 | 9.7e-50 | 3.27e-45 |
| Malignant epithelial | C2 WNT ECs | FGFR3 | 3.075 | 0.108 | 0.013 | 4.22e-51 | 1.42e-46 |
| Malignant epithelial | C2 WNT ECs | DDX3Y | 3.06 | 0.199 | 0.026 | 6.549999999999999e-89 | 2.21e-84 |
| Malignant epithelial | C2 WNT ECs | MAOB | 3.055 | 0.218 | 0.016 | 1.41e-134 | 4.75e-130 |
| Malignant epithelial | C2 WNT ECs | GOLGA8B | 3.04 | 0.158 | 0.017 | 1.22e-79 | 4.1e-75 |
| Malignant epithelial | C2 WNT ECs | EIF1AY | 3.039 | 0.345 | 0.033 | 2.61e-191 | 8.79e-187 |
| Malignant epithelial | C2 WNT ECs | LPCAT2 | 3.037 | 0.497 | 0.073 | 6.13e-222 | 2.06e-217 |
| Malignant epithelial | C2 WNT ECs | ANKH | 3.036 | 0.512 | 0.077 | 1.53e-225 | 5.15e-221 |
| Malignant epithelial | C2 WNT ECs | CAPS | 3.032 | 0.566 | 0.077 | 2.19e-268 | 7.39e-264 |
| Malignant epithelial | C2 WNT ECs | PRAP1 | 3.027 | 0.909 | 0.27 | 0.0 | 0.0 |
| Malignant epithelial | C2 WNT ECs | QPRT | 3.025 | 0.494 | 0.09 | 2.18e-193 | 7.349999999999999e-189 |
| Malignant epithelial | C2 WNT ECs | ARL4D | 3.007 | 0.242 | 0.034 | 9.87e-102 | 3.33e-97 |
| Malignant epithelial | C2 WNT ECs | PAQR3 | 3.007 | 0.18 | 0.02 | 1.85e-87 | 6.24e-83 |
| Malignant epithelial | C2 WNT ECs | GYLTL1B | 2.982 | 0.161 | 0.014 | 3.05e-89 | 1.03e-84 |
| Malignant epithelial | C2 WNT ECs | EPHX4 | 2.98 | 0.218 | 0.02 | 4.38e-118 | 1.48e-113 |
| Malignant epithelial | C2 WNT ECs | DNAJC15 | 2.975 | 0.958 | 0.527 | 8.15e-290 | 2.74e-285 |
| Malignant epithelial | C2 WNT ECs | HSPH1 | 2.949 | 0.876 | 0.437 | 4.04e-197 | 1.36e-192 |
| Malignant epithelial | C2 WNT ECs | MSX1 | 2.949 | 0.184 | 0.033 | 2.6e-64 | 8.77e-60 |
| Malignant epithelial | C2 WNT ECs | COL9A3 | 2.942 | 0.476 | 0.097 | 8.54e-165 | 2.88e-160 |
| Malignant epithelial | C2 WNT ECs | ZNF341-AS1 | 2.941 | 0.103 | 0.011 | 1.1e-52 | 3.72e-48 |
| Malignant epithelial | C2 WNT ECs | LDHB | 2.94 | 0.627 | 0.098 | 2.55e-284 | 8.59e-280 |
| Malignant epithelial | C2 WNT ECs | ZNF611 | 2.923 | 0.114 | 0.02 | 1.89e-40 | 6.36e-36 |
| Malignant epithelial | C2 WNT ECs | GGH | 2.917 | 0.827 | 0.32 | 6.29e-235 | 2.12e-230 |
| Malignant epithelial | C2 WNT ECs | FUZ | 2.91 | 0.126 | 0.012 | 3.91e-68 | 1.32e-63 |
| Malignant epithelial | C2 WNT ECs | C15orf62 | 2.893 | 0.111 | 0.013 | 4.24e-53 | 1.43e-48 |
| Malignant epithelial | C2 WNT ECs | SPRY2 | 2.892 | 0.378 | 0.056 | 2.68e-159 | 9.040000000000001e-155 |
| Malignant epithelial | C2 WNT ECs | SERP2 | 2.887 | 0.135 | 0.016 | 1.04e-63 | 3.5e-59 |
| Malignant epithelial | C2 WNT ECs | RP4-781K5.4 | 2.885 | 0.187 | 0.034 | 2.09e-64 | 7.04e-60 |
| Malignant epithelial | C2 WNT ECs | LRCH1 | 2.883 | 0.406 | 0.07 | 7.8e-156 | 2.63e-151 |
| Malignant epithelial | C2 WNT ECs | CADM4 | 2.877 | 0.122 | 0.014 | 1.78e-58 | 5.989999999999999e-54 |
| Malignant epithelial | C2 WNT ECs | SYCE1L | 2.871 | 0.112 | 0.01 | 3.27e-61 | 1.1e-56 |
| Malignant epithelial | C2 WNT ECs | AIFM3 | 2.863 | 0.13 | 0.014 | 6.39e-65 | 2.15e-60 |
| Malignant epithelial | C2 WNT ECs | LRP8 | 2.858 | 0.106 | 0.015 | 2.42e-43 | 8.14e-39 |
| Malignant epithelial | C2 WNT ECs | GPSM2 | 2.845 | 0.447 | 0.072 | 1.78e-183 | 6.010000000000001e-179 |
| Malignant epithelial | C2 WNT ECs | G0S2 | 2.841 | 0.319 | 0.05 | 1.41e-126 | 4.74e-122 |
| Malignant epithelial | C2 WNT ECs | PPP1R14A | 2.833 | 0.152 | 0.015 | 6.94e-80 | 2.34e-75 |
| Malignant epithelial | C2 WNT ECs | RGMB | 2.817 | 0.219 | 0.044 | 1.58e-68 | 5.32e-64 |

| compartment | subcluster | gene | avg_log2FC | pct.1 | pct.2 | p_val | p_val_adj |
|---|---|---|---|---|---|---|---|
| Malignant epithelial | C2 WNT ECs | CLU | 2.809 | 0.309 | 0.057 | 6.35e-109 | 2.14e-104 |
| Malignant epithelial | C2 WNT ECs | MYCL | 2.808 | 0.154 | 0.019 | 7.3e-70 | 2.46e-65 |
| Malignant epithelial | C2 WNT ECs | CYP39A1 | 2.803 | 0.101 | 0.013 | 2.49e-45 | 8.38e-41 |
| Malignant epithelial | C2 WNT ECs | ZNF503 | 2.797 | 0.448 | 0.088 | 2.03e-155 | 6.82e-151 |
| Malignant epithelial | C2 WNT ECs | FAM3B | 2.795 | 0.156 | 0.023 | 1.52e-63 | 5.12e-59 |
| Malignant epithelial | C2 WNT ECs | ALDH1A1 | 2.794 | 0.26 | 0.048 | 2.58e-90 | 8.68e-86 |
| Malignant epithelial | C2 WNT ECs | SESN1 | 2.794 | 0.245 | 0.048 | 2.12e-80 | 7.14e-76 |
| Malignant epithelial | C2 WNT ECs | CST7 | 2.793 | 0.429 | 0.074 | 3.27e-166 | 1.1e-161 |
| Malignant epithelial | C2 WNT ECs | CNPY4 | 2.77 | 0.103 | 0.013 | 1.36e-45 | 4.58e-41 |
| Malignant epithelial | C2 WNT ECs | DNAJB4 | 2.765 | 0.222 | 0.044 | 1.12e-70 | 3.78e-66 |
| Malignant epithelial | C2 WNT ECs | FGF18 | 2.743 | 0.187 | 0.025 | 4.92e-82 | 1.66e-77 |
| Malignant epithelial | C2 WNT ECs | SLC1A5 | 2.74 | 0.836 | 0.314 | 3.36e-237 | 1.13e-232 |
| Malignant epithelial | C2 WNT ECs | GPR155 | 2.734 | 0.107 | 0.019 | 1.84e-37 | 6.21e-33 |
| Malignant epithelial | C2 WNT ECs | TMEM86A | 2.73 | 0.13 | 0.022 | 1.36e-46 | 4.58e-42 |
| Malignant epithelial | C2 WNT ECs | PPARD | 2.722 | 0.204 | 0.047 | 2.15e-55 | 7.23e-51 |
| Malignant epithelial | C2 WNT ECs | LINC00659 | 2.694 | 0.233 | 0.043 | 5.64e-79 | 1.9e-74 |
| Malignant epithelial | C2 WNT ECs | RP11-23P13.6 | 2.688 | 0.122 | 0.013 | 1.36e-59 | 4.57e-55 |
| Malignant epithelial | C2 WNT ECs | TNNC1 | 2.682 | 0.187 | 0.021 | 3.15e-89 | 1.06e-84 |
| Malignant epithelial | C2 WNT ECs | HSPA6 | 2.666 | 0.539 | 0.219 | 3.96e-93 | 1.33e-88 |
| Malignant epithelial | C2 WNT ECs | ATG9B | 2.663 | 0.158 | 0.042 | 7.21e-37 | 2.43e-32 |
| Malignant epithelial | C2 WNT ECs | CHFR | 2.649 | 0.115 | 0.014 | 4.01e-52 | 1.35e-47 |
| Malignant epithelial | C2 WNT ECs | RNF39 | 2.646 | 0.287 | 0.056 | 7.29e-95 | 2.45e-90 |
| Malignant epithelial | C2 WNT ECs | ZNF185 | 2.634 | 0.199 | 0.04 | 2.1e-61 | 7.07e-57 |
| Malignant epithelial | C2 WNT ECs | PCSK9 | 2.627 | 0.116 | 0.021 | 2.09e-39 | 7.04e-35 |
| Malignant epithelial | C2 WNT ECs | CH507-154B10.1 | 2.625 | 0.106 | 0.017 | 5.739999999999999e-40 | 1.94e-35 |
| Malignant epithelial | C2 WNT ECs | GULP1 | 2.614 | 0.353 | 0.076 | 7.97e-111 | 2.68e-106 |
| Malignant epithelial | C2 WNT ECs | ZNF165 | 2.614 | 0.23 | 0.065 | 2.69e-49 | 9.06e-45 |
| Malignant epithelial | C2 WNT ECs | KRT17 | 2.612 | 0.582 | 0.205 | 3.53e-121 | 1.19e-116 |
| Malignant epithelial | C2 WNT ECs | IL23A | 2.609 | 0.35 | 0.069 | 4.89e-119 | 1.65e-114 |
| Malignant epithelial | C2 WNT ECs | AFAP1L2 | 2.6 | 0.304 | 0.055 | 9.969999999999999e-108 | 3.36e-103 |
| Malignant epithelial | C2 WNT ECs | ATAD3C | 2.593 | 0.103 | 0.012 | 8.189999999999999e-48 | 2.76e-43 |
| Malignant epithelial | C2 WNT ECs | ZNF480 | 2.591 | 0.106 | 0.016 | 6.28e-42 | 2.11e-37 |
| Malignant epithelial | C2 WNT ECs | SLC7A8 | 2.57 | 0.218 | 0.037 | 2.29e-78 | 7.719999999999999e-74 |
| Malignant epithelial | C2 WNT ECs | FADS1 | 2.565 | 0.103 | 0.013 | 9.4e-47 | 3.17e-42 |
| Malignant epithelial | C2 WNT ECs | GLUL | 2.562 | 0.655 | 0.206 | 9.15e-170 | 3.08e-165 |
| Malignant epithelial | C2 WNT ECs | NEU4 | 2.561 | 0.123 | 0.018 | 8.730000000000001e-49 | 2.94e-44 |
| Malignant epithelial | C2 WNT ECs | CEBPA | 2.551 | 0.246 | 0.05 | 3.11e-77 | 1.05e-72 |
| Malignant epithelial | C2 WNT ECs | SPON2 | 2.55 | 0.491 | 0.119 | 8.73e-145 | 2.94e-140 |
| Malignant epithelial | C2 WNT ECs | STT3B | 2.549 | 0.782 | 0.322 | 5.51e-177 | 1.86e-172 |
| Malignant epithelial | C2 WNT ECs | HEY1 | 2.545 | 0.115 | 0.015 | 1.65e-50 | 5.559999999999999e-46 |
| Malignant epithelial | C2 WNT ECs | BEX2 | 2.541 | 0.127 | 0.015 | 8.289999999999999e-60 | 2.79e-55 |
| Malignant epithelial | C2 WNT ECs | SLC9A3R2 | 2.54 | 0.476 | 0.145 | 1.5e-104 | 5.06e-100 |
| Malignant epithelial | C2 WNT ECs | SCEL | 2.536 | 0.227 | 0.068 | 5.51e-46 | 1.86e-41 |
| Malignant epithelial | C2 WNT ECs | ABHD2 | 2.535 | 0.632 | 0.291 | 2.2e-110 | 7.419999999999999e-106 |
| Malignant epithelial | C2 WNT ECs | TEAD2 | 2.532 | 0.199 | 0.034 | 1.95e-71 | 6.57e-67 |
| Malignant epithelial | C2 WNT ECs | PHLDB1 | 2.501 | 0.214 | 0.034 | 4.83e-81 | 1.63e-76 |
| Malignant epithelial | C2 WNT ECs | RPS4Y1 | 2.5 | 0.486 | 0.152 | 1.99e-119 | 6.689999999999999e-115 |
| Malignant epithelial | C2 WNT ECs | SOX4 | 2.486 | 0.917 | 0.449 | 1.24e-226 | 4.18e-222 |
| Malignant epithelial | C2 WNT ECs | SULT1A1 | 2.469 | 0.207 | 0.038 | 2.28e-69 | 7.68e-65 |
| Malignant epithelial | C2 WNT ECs | ZFAND2A | 2.467 | 0.754 | 0.47 | 5.18e-77 | 1.75e-72 |
| Malignant epithelial | C2 WNT ECs | TRPV3 | 2.453 | 0.1 | 0.023 | 2.91e-27 | 9.789999999999999e-23 |
| Malignant epithelial | C2 WNT ECs | MSLN | 2.451 | 0.606 | 0.261 | 2.08e-118 | 7.01e-114 |
| Malignant epithelial | C2 WNT ECs | PFN2 | 2.446 | 0.222 | 0.041 | 2.9e-74 | 9.760000000000001e-70 |
| Malignant epithelial | C2 WNT ECs | BTN3A2 | 2.442 | 0.183 | 0.033 | 7.68e-62 | 2.59e-57 |
| Malignant epithelial | C2 WNT ECs | FAM46B | 2.438 | 0.165 | 0.035 | 4.15e-48 | 1.4e-43 |
| Malignant epithelial | C2 WNT ECs | AKAP12 | 2.436 | 0.478 | 0.208 | 1.16e-67 | 3.92e-63 |
| Malignant epithelial | C2 WNT ECs | RTKN2 | 2.434 | 0.114 | 0.023 | 6.990000000000001e-35 | 2.35e-30 |
| Malignant epithelial | C2 WNT ECs | RTN2 | 2.432 | 0.179 | 0.031 | 1.51e-62 | 5.08e-58 |
| Malignant epithelial | C2 WNT ECs | RGS16 | 2.425 | 0.2 | 0.052 | 5.66e-48 | 1.91e-43 |
| Malignant epithelial | C2 WNT ECs | UBL3 | 2.423 | 0.605 | 0.159 | 1.66e-172 | 5.58e-168 |
| Malignant epithelial | C2 WNT ECs | SERPINF1 | 2.418 | 0.127 | 0.017 | 1.32e-53 | 4.46e-49 |
| Malignant epithelial | C2 WNT ECs | ASCL2 | 2.416 | 0.769 | 0.317 | 2.07e-162 | 6.96e-158 |
| Malignant epithelial | C2 WNT ECs | TP53INP2 | 2.412 | 0.256 | 0.044 | 1.22e-92 | 4.11e-88 |
| Malignant epithelial | C2 WNT ECs | DAB2 | 2.405 | 0.238 | 0.06 | 3.38e-60 | 1.14e-55 |

| compartment | subcluster | gene | avg_log2FC | pct.1 | pct.2 | p_val | p_val_adj |
|---|---|---|---|---|---|---|---|
| Malignant epithelial | C2 WNT ECs | F10 | 2.402 | 0.196 | 0.048 | 2.76e-49 | 9.3e-45 |
| Malignant epithelial | C2 WNT ECs | PROS1 | 2.402 | 0.139 | 0.025 | 4.37e-48 | 1.47e-43 |
| Malignant epithelial | C2 WNT ECs | CHI3L1 | 2.402 | 0.141 | 0.036 | 1.16e-33 | 3.92e-29 |
| Malignant epithelial | C2 WNT ECs | AP003774.1 | 2.398 | 0.103 | 0.015 | 5.04e-42 | 1.7e-37 |
| Malignant epithelial | C2 WNT ECs | S100A8 | 2.392 | 0.118 | 0.041 | 6.29e-18 | 2.12e-13 |
| Malignant epithelial | C2 WNT ECs | RUNX3 | 2.386 | 0.166 | 0.032 | 7.519999999999999e-54 | 2.54e-49 |
| Malignant epithelial | C2 WNT ECs | FLJ22763 | 2.381 | 0.108 | 0.015 | 1.93e-45 | 6.499999999999999e-41 |
| Malignant epithelial | C2 WNT ECs | ABLIM2 | 2.375 | 0.138 | 0.023 | 2.2e-50 | 7.41e-46 |
| Malignant epithelial | C2 WNT ECs | LCK | 2.37 | 0.336 | 0.082 | 1.1e-91 | 3.72e-87 |
| Malignant epithelial | C2 WNT ECs | DKK1 | 2.362 | 0.24 | 0.082 | 3.66e-40 | 1.23e-35 |
| Malignant epithelial | C2 WNT ECs | FMO5 | 2.36 | 0.122 | 0.021 | 1.43e-42 | 4.81e-38 |
| Malignant epithelial | C2 WNT ECs | AREG | 2.358 | 0.829 | 0.471 | 9.5e-142 | 3.2e-137 |
| Malignant epithelial | C2 WNT ECs | FAM105A | 2.354 | 0.225 | 0.056 | 3.73e-57 | 1.26e-52 |
| Malignant epithelial | C2 WNT ECs | PLEKHB1 | 2.349 | 0.298 | 0.065 | 8.77e-88 | 2.95e-83 |
| Malignant epithelial | C2 WNT ECs | MMP11 | 2.344 | 0.129 | 0.023 | 1.22e-44 | 4.11e-40 |
| Malignant epithelial | C2 WNT ECs | TNFRSF11B | 2.342 | 0.133 | 0.033 | 3.47e-33 | 1.17e-28 |
| Malignant epithelial | C2 WNT ECs | FBXL6 | 2.333 | 0.537 | 0.136 | 1.92e-153 | 6.479999999999999e-149 |
| Malignant epithelial | C2 WNT ECs | FAM46A | 2.313 | 0.617 | 0.184 | 1.75e-153 | 5.88e-149 |
| Malignant epithelial | C2 WNT ECs | SPATA13 | 2.304 | 0.432 | 0.104 | 3.27e-120 | 1.1e-115 |
| Malignant epithelial | C2 WNT ECs | LGALSL | 2.303 | 0.242 | 0.068 | 9.9e-52 | 3.33e-47 |
| Malignant epithelial | C2 WNT ECs | INTS6 | 2.301 | 0.678 | 0.194 | 5.05e-179 | 1.7e-174 |
| Malignant epithelial | C2 WNT ECs | CDX2 | 2.291 | 0.591 | 0.12 | 1.03e-201 | 3.48e-197 |
| Malignant epithelial | C2 WNT ECs | BAG3 | 2.284 | 0.539 | 0.266 | 1.33e-64 | 4.47e-60 |
| Malignant epithelial | C2 WNT ECs | NES | 2.284 | 0.111 | 0.02 | 1.49e-37 | 5.03e-33 |
| Malignant epithelial | C2 WNT ECs | KCNK5 | 2.281 | 0.18 | 0.038 | 2.67e-53 | 9e-49 |
| Malignant epithelial | C2 WNT ECs | PITX2 | 2.277 | 0.175 | 0.033 | 6.54e-56 | 2.2e-51 |
| Malignant epithelial | C2 WNT ECs | CST6 | 2.276 | 0.543 | 0.275 | 1.73e-67 | 5.84e-63 |
| Malignant epithelial | C2 WNT ECs | ARHGAP23 | 2.274 | 0.112 | 0.027 | 4.71e-29 | 1.59e-24 |
| Malignant epithelial | C2 WNT ECs | DUOX1 | 2.271 | 0.116 | 0.019 | 1.87e-43 | 6.29e-39 |
| Malignant epithelial | C2 WNT ECs | ADH1C | 2.267 | 0.131 | 0.025 | 1.12e-42 | 3.78e-38 |
| Malignant epithelial | C2 WNT ECs | SEMA5A | 2.264 | 0.111 | 0.022 | 4.54e-34 | 1.53e-29 |
| Malignant epithelial | C2 WNT ECs | ZNRF3 | 2.262 | 0.198 | 0.043 | 8.720000000000001e-56 | 2.94e-51 |
| Malignant epithelial | C2 WNT ECs | MYLIP | 2.257 | 0.524 | 0.146 | 7.190000000000001e-133 | 2.42e-128 |
| Malignant epithelial | C2 WNT ECs | LMO7 | 2.248 | 0.829 | 0.442 | 2.95e-137 | 9.950000000000001e-133 |
| Malignant epithelial | C2 WNT ECs | SIT1 | 2.245 | 0.111 | 0.018 | 1.81e-41 | 6.080000000000001e-37 |
| Malignant epithelial | C2 WNT ECs | S100A14 | 2.242 | 0.824 | 0.293 | 3.61e-202 | 1.22e-197 |
| Malignant epithelial | C2 WNT ECs | RANBP9 | 2.242 | 0.29 | 0.055 | 1.69e-94 | 5.699999999999999e-90 |
| Malignant epithelial | C2 WNT ECs | HMGCS2 | 2.231 | 0.192 | 0.043 | 3.45e-53 | 1.16e-48 |
| Malignant epithelial | C2 WNT ECs | TMEM98 | 2.228 | 0.372 | 0.129 | 1.58e-69 | 5.32e-65 |
| Malignant epithelial | C2 WNT ECs | PROM2 | 2.225 | 0.141 | 0.026 | 2.99e-46 | 1.01e-41 |
| Malignant epithelial | C2 WNT ECs | TMEM255B | 2.224 | 0.116 | 0.021 | 1.22e-38 | 4.12e-34 |
| Malignant epithelial | C2 WNT ECs | RP5-881L22.5 | 2.223 | 0.158 | 0.028 | 5.259999999999999e-54 | 1.77e-49 |
| Malignant epithelial | C2 WNT ECs | PTGR1 | 2.221 | 0.25 | 0.049 | 1.81e-78 | 6.100000000000001e-74 |
| Malignant epithelial | C2 WNT ECs | INSIG1 | 2.214 | 0.525 | 0.229 | 7.810000000000001e-72 | 2.63e-67 |
| Malignant epithelial | C2 WNT ECs | SETDB2 | 2.211 | 0.18 | 0.035 | 4.56e-57 | 1.54e-52 |
| Malignant epithelial | C2 WNT ECs | SATB1 | 2.184 | 0.206 | 0.038 | 2.63e-67 | 8.86e-63 |
| Malignant epithelial | C2 WNT ECs | WLS | 2.183 | 0.399 | 0.094 | 5.84e-112 | 1.97e-107 |
| Malignant epithelial | C2 WNT ECs | RDH10 | 2.183 | 0.456 | 0.151 | 5.019999999999999e-91 | 1.69e-86 |
| Malignant epithelial | C2 WNT ECs | DEPDC7 | 2.183 | 0.101 | 0.022 | 2.39e-29 | 8.06e-25 |
| Malignant epithelial | C2 WNT ECs | SMAD7 | 2.18 | 0.179 | 0.037 | 1.16e-52 | 3.91e-48 |
| Malignant epithelial | C2 WNT ECs | TSPAN12 | 2.179 | 0.261 | 0.065 | 4.14e-65 | 1.39e-60 |
| Malignant epithelial | C2 WNT ECs | HMOX1 | 2.175 | 0.15 | 0.037 | 3.26e-37 | 1.1e-32 |
| Malignant epithelial | C2 WNT ECs | CALCOCO1 | 2.173 | 0.327 | 0.085 | 1.07e-78 | 3.6e-74 |
| Malignant epithelial | C2 WNT ECs | UNC119 | 2.168 | 0.405 | 0.147 | 1.21e-66 | 4.06e-62 |
| Malignant epithelial | C2 WNT ECs | GPR153 | 2.156 | 0.214 | 0.043 | 3.81e-65 | 1.28e-60 |
| Malignant epithelial | C2 WNT ECs | HSPA2 | 2.152 | 0.222 | 0.048 | 2.98e-63 | 1e-58 |
| Malignant epithelial | C2 WNT ECs | ARHGEF4 | 2.152 | 0.147 | 0.028 | 4.38e-48 | 1.48e-43 |
| Malignant epithelial | C2 WNT ECs | DPYSL2 | 2.147 | 0.383 | 0.097 | 1.73e-99 | 5.83e-95 |
| Malignant epithelial | C2 WNT ECs | KRT6A | 2.142 | 0.246 | 0.074 | 1.67e-49 | 5.619999999999999e-45 |
| Malignant epithelial | C2 WNT ECs | PARD3B | 2.126 | 0.1 | 0.021 | 1.12e-29 | 3.78e-25 |
| Malignant epithelial | C2 WNT ECs | ABAT | 2.12 | 0.114 | 0.022 | 7.65e-36 | 2.58e-31 |
| Malignant epithelial | C2 WNT ECs | PTP4A3 | 2.117 | 0.263 | 0.105 | 4.69e-37 | 1.58e-32 |
| Malignant epithelial | C2 WNT ECs | GDPD5 | 2.112 | 0.207 | 0.046 | 2.09e-57 | 7.05e-53 |
| Malignant epithelial | C2 WNT ECs | MAFB | 2.109 | 0.192 | 0.044 | 4.1e-51 | 1.38e-46 |

| compartment | subcluster | gene | avg_log2FC | pct.1 | pct.2 | p_val | p_val_adj |
|---|---|---|---|---|---|---|---|
| Malignant epithelial | C2 WNT ECs | HES6 | 2.106 | 0.452 | 0.118 | 5.02e-118 | 1.69e-113 |
| Malignant epithelial | C2 WNT ECs | GLB1L2 | 2.105 | 0.273 | 0.062 | 4e-75 | 1.35e-70 |
| Malignant epithelial | C2 WNT ECs | MFGE8 | 2.101 | 0.384 | 0.161 | 1.22e-49 | 4.12e-45 |
| Malignant epithelial | C2 WNT ECs | NMNAT3 | 2.098 | 0.263 | 0.054 | 2.09e-78 | 7.03e-74 |
| Malignant epithelial | C2 WNT ECs | CITED2 | 2.095 | 0.639 | 0.287 | 8.91e-99 | 3e-94 |
| Malignant epithelial | C2 WNT ECs | ABHD6 | 2.092 | 0.112 | 0.027 | 5.38e-29 | 1.81e-24 |
| Malignant epithelial | C2 WNT ECs | SLC7A6 | 2.09 | 0.235 | 0.056 | 3.23e-61 | 1.09e-56 |
| Malignant epithelial | C2 WNT ECs | KBTBD7 | 2.079 | 0.165 | 0.037 | 3.51e-45 | 1.18e-40 |
| Malignant epithelial | C2 WNT ECs | C8orf4 | 2.078 | 0.131 | 0.04 | 2.37e-25 | 7.980000000000001e-21 |
| Malignant epithelial | C2 WNT ECs | C2orf54 | 2.075 | 0.149 | 0.033 | 1.89e-41 | 6.360000000000001e-37 |
| Malignant epithelial | C2 WNT ECs | TRIM29 | 2.073 | 0.445 | 0.129 | 1.35e-102 | 4.55e-98 |
| Malignant epithelial | C2 WNT ECs | UPK2 | 2.067 | 0.191 | 0.094 | 5.32e-16 | 1.79e-11 |
| Malignant epithelial | C2 WNT ECs | HOXA3 | 2.061 | 0.233 | 0.055 | 1.13e-60 | 3.79e-56 |
| Malignant epithelial | C2 WNT ECs | NUAK2 | 2.06 | 0.187 | 0.047 | 8.58e-45 | 2.89e-40 |
| Malignant epithelial | C2 WNT ECs | FBXO2 | 2.054 | 0.299 | 0.073 | 2.73e-78 | 9.19e-74 |
| Malignant epithelial | C2 WNT ECs | ATP8A1 | 2.049 | 0.104 | 0.022 | 1.46e-30 | 4.91e-26 |
| Malignant epithelial | C2 WNT ECs | CDKN1C | 2.043 | 0.428 | 0.176 | 2.72e-56 | 9.159999999999999e-52 |
| Malignant epithelial | C2 WNT ECs | FAM84A | 2.042 | 0.53 | 0.165 | 1.75e-113 | 5.91e-109 |
| Malignant epithelial | C2 WNT ECs | ADAMTSL5 | 2.019 | 0.184 | 0.038 | 3.45e-54 | 1.16e-49 |
| Malignant epithelial | C2 WNT ECs | RUNX2 | 2.012 | 0.133 | 0.031 | 3.78e-35 | 1.27e-30 |
| Malignant epithelial | C2 WNT ECs | NTHL1 | 2.009 | 0.545 | 0.311 | 2.22e-38 | 7.49e-34 |
| Malignant epithelial | C2 WNT ECs | ARG2 | 2.007 | 0.15 | 0.036 | 1.95e-38 | 6.55e-34 |
| Malignant epithelial | C2 WNT ECs | PRR36 | 2.007 | 0.127 | 0.027 | 4.24e-36 | 1.43e-31 |
| Malignant epithelial | C2 WNT ECs | RGCC | 2.005 | 0.717 | 0.29 | 2.45e-129 | 8.24e-125 |
| Malignant epithelial | C2 WNT ECs | ATXN1 | 2.001 | 0.315 | 0.065 | 2.87e-95 | 9.68e-91 |
| Malignant epithelial | C2 WNT ECs | FYN | 2.0 | 0.326 | 0.1 | 3.01e-64 | 1.01e-59 |
| Malignant epithelial | C2 WNT ECs | LIPA | 2.0 | 0.171 | 0.034 | 1.09e-51 | 3.69e-47 |
| Malignant epithelial | C2 WNT ECs | TMEM176A | 1.999 | 0.893 | 0.522 | 4.09e-182 | 1.38e-177 |
| Malignant epithelial | C2 WNT ECs | SNAI1 | 1.997 | 0.421 | 0.118 | 9.8e-98 | 3.3e-93 |
| Malignant epithelial | C2 WNT ECs | IL20RA | 1.997 | 0.533 | 0.208 | 3.32e-88 | 1.12e-83 |
| Malignant epithelial | C2 WNT ECs | ABTB1 | 1.996 | 0.442 | 0.13 | 7.55e-96 | 2.55e-91 |
| Malignant epithelial | C2 WNT ECs | ACADM | 1.995 | 0.267 | 0.067 | 8.150000000000001e-66 | 2.75e-61 |
| Malignant epithelial | C2 WNT ECs | KIAA0226L | 1.993 | 0.162 | 0.044 | 3.14e-36 | 1.06e-31 |
| Malignant epithelial | C2 WNT ECs | TSC22D1 | 1.992 | 0.951 | 0.586 | 3.63e-200 | 1.22e-195 |
| Malignant epithelial | C2 WNT ECs | KDM7A | 1.98 | 0.333 | 0.075 | 4.33e-92 | 1.46e-87 |
| Malignant epithelial | C2 WNT ECs | DLX2 | 1.98 | 0.103 | 0.019 | 8.689999999999999e-34 | 2.93e-29 |
| Malignant epithelial | C2 WNT ECs | SHISA4 | 1.975 | 0.127 | 0.027 | 1.04e-36 | 3.5e-32 |
| Malignant epithelial | C2 WNT ECs | GDF15 | 1.97 | 0.614 | 0.354 | 3.54e-57 | 1.19e-52 |
| Malignant epithelial | C2 WNT ECs | TMPRSS13 | 1.97 | 0.204 | 0.051 | 2.02e-49 | 6.8e-45 |
| Malignant epithelial | C2 WNT ECs | FHL3 | 1.97 | 0.142 | 0.033 | 9.85e-37 | 3.32e-32 |
| Malignant epithelial | C2 WNT ECs | SDR16C5 | 1.967 | 0.413 | 0.147 | 9.1e-73 | 3.06e-68 |
| Malignant epithelial | C2 WNT ECs | FAM134B | 1.962 | 0.204 | 0.065 | 2.96e-37 | 9.98e-33 |
| Malignant epithelial | C2 WNT ECs | YPEL3 | 1.96 | 0.729 | 0.273 | 4.08e-156 | 1.37e-151 |
| Malignant epithelial | C2 WNT ECs | PAQR5 | 1.957 | 0.181 | 0.038 | 2.45e-52 | 8.269999999999999e-48 |
| Malignant epithelial | C2 WNT ECs | ARRB1 | 1.953 | 0.153 | 0.033 | 2e-42 | 6.73e-38 |
| Malignant epithelial | C2 WNT ECs | LINC00649 | 1.953 | 0.135 | 0.029 | 5.26e-39 | 1.77e-34 |
| Malignant epithelial | C2 WNT ECs | LACC1 | 1.953 | 0.1 | 0.02 | 2.36e-31 | 7.970000000000001e-27 |
| Malignant epithelial | C2 WNT ECs | RP11-511P7.5 | 1.952 | 0.241 | 0.056 | 5.41e-63 | 1.82e-58 |
| Malignant epithelial | C2 WNT ECs | ARC | 1.949 | 0.153 | 0.041 | 6.11e-34 | 2.06e-29 |
| Malignant epithelial | C2 WNT ECs | PROM1 | 1.949 | 0.145 | 0.04 | 1.27e-30 | 4.27e-26 |
| Malignant epithelial | C2 WNT ECs | RASSF4 | 1.941 | 0.104 | 0.02 | 1.76e-33 | 5.93e-29 |
| Malignant epithelial | C2 WNT ECs | SUPT20H | 1.939 | 0.414 | 0.109 | 5.14e-99 | 1.73e-94 |
| Malignant epithelial | C2 WNT ECs | BMP4 | 1.933 | 0.593 | 0.369 | 1.77e-65 | 5.96e-61 |
| Malignant epithelial | C2 WNT ECs | SULT2B1 | 1.933 | 0.587 | 0.317 | 2.26e-65 | 7.609999999999999e-61 |
| Malignant epithelial | C2 WNT ECs | MIPEP | 1.931 | 0.183 | 0.043 | 4.07e-47 | 1.37e-42 |
| Malignant epithelial | C2 WNT ECs | NUDT4 | 1.926 | 0.855 | 0.481 | 2.7e-145 | 9.11e-141 |
| Malignant epithelial | C2 WNT ECs | TPT1-AS1 | 1.925 | 0.352 | 0.09 | 6.19e-86 | 2.09e-81 |
| Malignant epithelial | C2 WNT ECs | RNF6 | 1.922 | 0.321 | 0.076 | 3.11e-83 | 1.05e-78 |
| Malignant epithelial | C2 WNT ECs | CHAC1 | 1.919 | 0.138 | 0.04 | 6.44e-28 | 2.17e-23 |
| Malignant epithelial | C2 WNT ECs | PRSS22 | 1.913 | 0.652 | 0.284 | 9.23e-96 | 3.11e-91 |
| Malignant epithelial | C2 WNT ECs | SPHK2 | 1.907 | 0.248 | 0.067 | 5.159999999999999e-55 | 1.74e-50 |
| Malignant epithelial | C2 WNT ECs | USPL1 | 1.905 | 0.181 | 0.044 | 1.26e-45 | 4.24e-41 |
| Malignant epithelial | C2 WNT ECs | FGFRL1 | 1.895 | 0.271 | 0.08 | 2.73e-55 | 9.19e-51 |
| Malignant epithelial | C2 WNT ECs | LRRC8E | 1.891 | 0.112 | 0.026 | 1.28e-29 | 4.32e-25 |

| compartment | subcluster | gene | avg_log2FC | pct.1 | pct.2 | p_val | p_val_adj |
|---|---|---|---|---|---|---|---|
| Malignant epithelial | C2 WNT ECs | NAA16 | 1.889 | 0.233 | 0.058 | 4.31e-56 | 1.45e-51 |
| Malignant epithelial | C2 WNT ECs | TCEA2 | 1.884 | 0.147 | 0.04 | 1.21e-32 | 4.06e-28 |
| Malignant epithelial | C2 WNT ECs | TUBA1A | 1.881 | 0.356 | 0.146 | 1.63e-45 | 5.5e-41 |
| Malignant epithelial | C2 WNT ECs | HPGD | 1.875 | 0.387 | 0.146 | 1.11e-60 | 3.74e-56 |
| Malignant epithelial | C2 WNT ECs | CYR61 | 1.875 | 0.398 | 0.158 | 4.78e-57 | 1.61e-52 |
| Malignant epithelial | C2 WNT ECs | GOLGA7B | 1.869 | 0.184 | 0.04 | 8.11e-52 | 2.73e-47 |
| Malignant epithelial | C2 WNT ECs | CLIC3 | 1.862 | 0.539 | 0.305 | 9.939999999999999e-52 | 3.35e-47 |
| Malignant epithelial | C2 WNT ECs | RHOBTB3 | 1.859 | 0.528 | 0.148 | 5.1e-123 | 1.72e-118 |
| Malignant epithelial | C2 WNT ECs | KIF5B | 1.858 | 0.785 | 0.325 | 2.44e-158 | 8.21e-154 |
| Malignant epithelial | C2 WNT ECs | SHF | 1.858 | 0.199 | 0.045 | 5.15e-53 | 1.73e-48 |
| Malignant epithelial | C2 WNT ECs | GJB2 | 1.855 | 0.361 | 0.16 | 2.61e-43 | 8.789999999999999e-39 |
| Malignant epithelial | C2 WNT ECs | MGAT5 | 1.85 | 0.256 | 0.066 | 2.87e-60 | 9.68e-56 |
| Malignant epithelial | C2 WNT ECs | IER5 | 1.846 | 0.608 | 0.359 | 2.47e-55 | 8.33e-51 |
| Malignant epithelial | C2 WNT ECs | ZSCAN16 | 1.844 | 0.112 | 0.028 | 2.76e-27 | 9.3e-23 |
| Malignant epithelial | C2 WNT ECs | FOXP4-AS1 | 1.843 | 0.157 | 0.041 | 4.36e-36 | 1.47e-31 |
| Malignant epithelial | C2 WNT ECs | CTU1 | 1.837 | 0.164 | 0.033 | 1.44e-48 | 4.85e-44 |
| Malignant epithelial | C2 WNT ECs | HOXB13 | 1.837 | 0.126 | 0.046 | 1.66e-18 | 5.58e-14 |
| Malignant epithelial | C2 WNT ECs | PLA2R1 | 1.836 | 0.11 | 0.026 | 6.32e-28 | 2.13e-23 |
| Malignant epithelial | C2 WNT ECs | CHRNB1 | 1.834 | 0.156 | 0.042 | 1.47e-34 | 4.96e-30 |
| Malignant epithelial | C2 WNT ECs | NEK3 | 1.832 | 0.269 | 0.069 | 4.46e-64 | 1.5e-59 |
| Malignant epithelial | C2 WNT ECs | RAB11FIP5 | 1.83 | 0.129 | 0.033 | 1.03e-29 | 3.46e-25 |
| Malignant epithelial | C2 WNT ECs | ZNF704 | 1.821 | 0.122 | 0.032 | 3.74e-28 | 1.26e-23 |
| Malignant epithelial | C2 WNT ECs | RNF43 | 1.816 | 0.69 | 0.235 | 4.6e-147 | 1.55e-142 |
| Malignant epithelial | C2 WNT ECs | RP11-395A13.2 | 1.814 | 0.192 | 0.048 | 5.71e-46 | 1.93e-41 |
| Malignant epithelial | C2 WNT ECs | SAMD13 | 1.813 | 0.114 | 0.031 | 4.65e-25 | 1.57e-20 |
| Malignant epithelial | C2 WNT ECs | USP6NL | 1.812 | 0.253 | 0.063 | 1.22e-61 | 4.12e-57 |
| Malignant epithelial | C2 WNT ECs | AC007405.6 | 1.811 | 0.135 | 0.039 | 1.1e-27 | 3.72e-23 |
| Malignant epithelial | C2 WNT ECs | MPZL3 | 1.805 | 0.268 | 0.07 | 1.08e-62 | 3.63e-58 |
| Malignant epithelial | C2 WNT ECs | SC5D | 1.801 | 0.292 | 0.082 | 2.41e-62 | 8.11e-58 |
| Malignant epithelial | C2 WNT ECs | FBXL3 | 1.8 | 0.409 | 0.118 | 3.29e-88 | 1.11e-83 |
| Malignant epithelial | C2 WNT ECs | ELF1 | 1.799 | 0.652 | 0.223 | 3.14e-135 | 1.06e-130 |
| Malignant epithelial | C2 WNT ECs | SLC39A5 | 1.799 | 0.3 | 0.076 | 4.09e-73 | 1.38e-68 |
| Malignant epithelial | C2 WNT ECs | IHH | 1.796 | 0.11 | 0.032 | 2.23e-22 | 7.5e-18 |
| Malignant epithelial | C2 WNT ECs | ZC3H13 | 1.795 | 0.593 | 0.193 | 1.49e-122 | 5.030000000000001e-118 |
| Malignant epithelial | C2 WNT ECs | WBP4 | 1.795 | 0.541 | 0.162 | 7.629999999999999e-120 | 2.57e-115 |
| Malignant epithelial | C2 WNT ECs | GABARAPL1 | 1.794 | 0.363 | 0.106 | 1.3e-75 | 4.38e-71 |
| Malignant epithelial | C2 WNT ECs | TPPP3 | 1.793 | 0.31 | 0.126 | 2.73e-39 | 9.2e-35 |
| Malignant epithelial | C2 WNT ECs | DAPK2 | 1.793 | 0.134 | 0.03 | 2.24e-36 | 7.550000000000001e-32 |
| Malignant epithelial | C2 WNT ECs | YRDC | 1.792 | 0.273 | 0.064 | 2.57e-71 | 8.669999999999999e-67 |
| Malignant epithelial | C2 WNT ECs | SUCLA2 | 1.788 | 0.464 | 0.153 | 4.98e-88 | 1.68e-83 |
| Malignant epithelial | C2 WNT ECs | OTUD1 | 1.787 | 0.153 | 0.043 | 4.53e-32 | 1.53e-27 |
| Malignant epithelial | C2 WNT ECs | ESPN | 1.786 | 0.313 | 0.102 | 3.81e-56 | 1.28e-51 |
| Malignant epithelial | C2 WNT ECs | COBL | 1.784 | 0.352 | 0.103 | 1.67e-73 | 5.64e-69 |
| Malignant epithelial | C2 WNT ECs | SEMA4C | 1.784 | 0.171 | 0.045 | 2.96e-38 | 9.96e-34 |
| Malignant epithelial | C2 WNT ECs | CARKD | 1.78 | 0.355 | 0.103 | 7.57e-75 | 2.55e-70 |
| Malignant epithelial | C2 WNT ECs | FCGRT | 1.779 | 0.899 | 0.501 | 5.75e-176 | 1.94e-171 |
| Malignant epithelial | C2 WNT ECs | CMB9-22P13.1 | 1.778 | 0.179 | 0.073 | 1.76e-20 | 5.92e-16 |
| Malignant epithelial | C2 WNT ECs | EFNA3 | 1.776 | 0.474 | 0.146 | 1.27e-96 | 4.28e-92 |
| Malignant epithelial | C2 WNT ECs | SORT1 | 1.775 | 0.161 | 0.038 | 9.77e-41 | 3.29e-36 |
| Malignant epithelial | C2 WNT ECs | KLK11 | 1.774 | 0.365 | 0.185 | 2.42e-31 | 8.140000000000001e-27 |
| Malignant epithelial | C2 WNT ECs | SP6 | 1.773 | 0.111 | 0.026 | 2.04e-28 | 6.88e-24 |
| Malignant epithelial | C2 WNT ECs | ITGA2 | 1.769 | 0.311 | 0.174 | 5.230000000000001e-21 | 1.76e-16 |
| Malignant epithelial | C2 WNT ECs | GRB10 | 1.766 | 0.145 | 0.031 | 1.68e-40 | 5.66e-36 |
| Malignant epithelial | C2 WNT ECs | ZMYM5 | 1.764 | 0.403 | 0.117 | 2.61e-85 | 8.79e-81 |
| Malignant epithelial | C2 WNT ECs | NCOA4 | 1.763 | 0.471 | 0.149 | 9.389999999999999e-94 | 3.17e-89 |
| Malignant epithelial | C2 WNT ECs | MOSPD1 | 1.762 | 0.383 | 0.126 | 1.35e-69 | 4.55e-65 |
| Malignant epithelial | C2 WNT ECs | CEMIP | 1.759 | 0.161 | 0.039 | 2.47e-40 | 8.33e-36 |
| Malignant epithelial | C2 WNT ECs | PCCA | 1.758 | 0.181 | 0.056 | 3e-33 | 1.01e-28 |
| Malignant epithelial | C2 WNT ECs | N6AMT2 | 1.757 | 0.223 | 0.061 | 2.3e-48 | 7.77e-44 |
| Malignant epithelial | C2 WNT ECs | EFHC1 | 1.755 | 0.378 | 0.111 | 1.98e-77 | 6.67e-73 |
| Malignant epithelial | C2 WNT ECs | MSX2 | 1.753 | 0.315 | 0.1 | 5.49e-59 | 1.85e-54 |
| Malignant epithelial | C2 WNT ECs | TPMT | 1.752 | 0.429 | 0.126 | 5.98e-91 | 2.02e-86 |
| Malignant epithelial | C2 WNT ECs | FAM8A1 | 1.75 | 0.122 | 0.032 | 9.68e-28 | 3.259999999999999e-23 |
| Malignant epithelial | C2 WNT ECs | TGFBR3 | 1.749 | 0.112 | 0.026 | 1.31e-29 | 4.42e-25 |

| compartment | subcluster | gene | avg_log2FC | pct.1 | pct.2 | p_val | p_val_adj |
|---|---|---|---|---|---|---|---|
| Malignant epithelial | C2 WNT ECs | PROSER1 | 1.745 | 0.242 | 0.058 | 9.56e-62 | 3.22e-57 |
| Malignant epithelial | C2 WNT ECs | WDFY2 | 1.741 | 0.388 | 0.118 | 9.210000000000001e-77 | 3.1e-72 |
| Malignant epithelial | C2 WNT ECs | GSTA4 | 1.737 | 0.107 | 0.031 | 3.81e-21 | 1.28e-16 |
| Malignant epithelial | C2 WNT ECs | RPPH1 | 1.733 | 0.165 | 0.044 | 4.08e-36 | 1.38e-31 |
| Malignant epithelial | C2 WNT ECs | CYTH3 | 1.733 | 0.122 | 0.034 | 1.18e-25 | 3.97e-21 |
| Malignant epithelial | C2 WNT ECs | RP11-553L6.5 | 1.733 | 0.101 | 0.026 | 2.12e-23 | 7.16e-19 |
| Malignant epithelial | C2 WNT ECs | SEMA3B | 1.732 | 0.407 | 0.167 | 1.44e-53 | 4.86e-49 |
| Malignant epithelial | C2 WNT ECs | KHDRBS3 | 1.732 | 0.161 | 0.042 | 5.11e-37 | 1.72e-32 |
| Malignant epithelial | C2 WNT ECs | SYT7 | 1.73 | 0.26 | 0.069 | 2.59e-58 | 8.739999999999999e-54 |
| Malignant epithelial | C2 WNT ECs | SLC6A20 | 1.725 | 0.135 | 0.039 | 2.13e-27 | 7.169999999999999e-23 |
| Malignant epithelial | C2 WNT ECs | SERPINB5 | 1.723 | 0.475 | 0.286 | 9.25e-36 | 3.12e-31 |
| Malignant epithelial | C2 WNT ECs | JHDM1D-AS1 | 1.717 | 0.237 | 0.067 | 1.37e-48 | 4.61e-44 |
| Malignant epithelial | C2 WNT ECs | BMPR1A | 1.716 | 0.112 | 0.027 | 2.75e-28 | 9.27e-24 |
| Malignant epithelial | C2 WNT ECs | XG | 1.713 | 0.12 | 0.026 | 2.55e-34 | 8.600000000000001e-30 |
| Malignant epithelial | C2 WNT ECs | PHLDB3 | 1.712 | 0.196 | 0.049 | 4.52e-47 | 1.52e-42 |
| Malignant epithelial | C2 WNT ECs | SYBU | 1.712 | 0.18 | 0.06 | 5.02e-30 | 1.69e-25 |
| Malignant epithelial | C2 WNT ECs | FGFBP1 | 1.71 | 0.116 | 0.032 | 8.98e-25 | 3.03e-20 |
| Malignant epithelial | C2 WNT ECs | SLC39A8 | 1.708 | 0.1 | 0.029 | 7.16e-20 | 2.41e-15 |
| Malignant epithelial | C2 WNT ECs | LNX2 | 1.705 | 0.21 | 0.055 | 1.63e-47 | 5.49e-43 |
| Malignant epithelial | C2 WNT ECs | TGFBI | 1.704 | 0.693 | 0.395 | 4.94e-76 | 1.66e-71 |
| Malignant epithelial | C2 WNT ECs | CHPT1 | 1.703 | 0.578 | 0.206 | 2.12e-106 | 7.14e-102 |
| Malignant epithelial | C2 WNT ECs | NEIL1 | 1.703 | 0.208 | 0.05 | 3.07e-52 | 1.03e-47 |
| Malignant epithelial | C2 WNT ECs | CAPN12 | 1.703 | 0.225 | 0.058 | 1.33e-51 | 4.470000000000001e-47 |
| Malignant epithelial | C2 WNT ECs | MGA | 1.703 | 0.145 | 0.038 | 2.9e-32 | 9.76e-28 |
| Malignant epithelial | C2 WNT ECs | LSMEM1 | 1.698 | 0.108 | 0.032 | 3.51e-21 | 1.18e-16 |
| Malignant epithelial | C2 WNT ECs | TSPAN5 | 1.694 | 0.242 | 0.095 | 2.59e-32 | 8.74e-28 |
| Malignant epithelial | C2 WNT ECs | SH3PXD2B | 1.694 | 0.122 | 0.039 | 1.91e-21 | 6.43e-17 |
| Malignant epithelial | C2 WNT ECs | CHN2 | 1.691 | 0.127 | 0.039 | 1.04e-23 | 3.5e-19 |
| Malignant epithelial | C2 WNT ECs | SESN3 | 1.689 | 0.11 | 0.031 | 2.63e-22 | 8.86e-18 |
| Malignant epithelial | C2 WNT ECs | TRIM13 | 1.687 | 0.302 | 0.085 | 8.12e-64 | 2.73e-59 |
| Malignant epithelial | C2 WNT ECs | RASSF6 | 1.682 | 0.269 | 0.083 | 3.25e-50 | 1.1e-45 |
| Malignant epithelial | C2 WNT ECs | TTL | 1.682 | 0.139 | 0.034 | 6.079999999999999e-34 | 2.05e-29 |
| Malignant epithelial | C2 WNT ECs | LBX2 | 1.682 | 0.118 | 0.026 | 8.39e-33 | 2.83e-28 |
| Malignant epithelial | C2 WNT ECs | CLDN9 | 1.681 | 0.156 | 0.051 | 1.16e-26 | 3.92e-22 |
| Malignant epithelial | C2 WNT ECs | RIOK1 | 1.68 | 0.226 | 0.061 | 2.26e-49 | 7.63e-45 |
| Malignant epithelial | C2 WNT ECs | PLPP3 | 1.679 | 0.114 | 0.034 | 2.01e-22 | 6.77e-18 |
| Malignant epithelial | C2 WNT ECs | TMEM214 | 1.678 | 0.39 | 0.123 | 1.59e-74 | 5.35e-70 |
| Malignant epithelial | C2 WNT ECs | COG6 | 1.677 | 0.176 | 0.041 | 6.38e-45 | 2.15e-40 |
| Malignant epithelial | C2 WNT ECs | HOXA5 | 1.673 | 0.191 | 0.053 | 5.82e-40 | 1.96e-35 |
| Malignant epithelial | C2 WNT ECs | KCNMB4 | 1.671 | 0.134 | 0.034 | 5.47e-31 | 1.84e-26 |
| Malignant epithelial | C2 WNT ECs | PLK2 | 1.669 | 0.616 | 0.296 | 1.1e-76 | 3.72e-72 |
| Malignant epithelial | C2 WNT ECs | BRD7 | 1.669 | 0.518 | 0.209 | 4.92e-75 | 1.66e-70 |
| Malignant epithelial | C2 WNT ECs | KDM1B | 1.669 | 0.137 | 0.03 | 1.79e-37 | 6.03e-33 |
| Malignant epithelial | C2 WNT ECs | ZNF430 | 1.669 | 0.219 | 0.073 | 8.390000000000001e-37 | 2.83e-32 |
| Malignant epithelial | C2 WNT ECs | MICU2 | 1.666 | 0.421 | 0.132 | 5.91e-81 | 1.99e-76 |
| Malignant epithelial | C2 WNT ECs | EPS8L3 | 1.664 | 0.472 | 0.147 | 1.3e-95 | 4.37e-91 |
| Malignant epithelial | C2 WNT ECs | STX3 | 1.662 | 0.562 | 0.184 | 3.9e-107 | 1.31e-102 |
| Malignant epithelial | C2 WNT ECs | TUBB2A | 1.661 | 0.525 | 0.229 | 2.16e-67 | 7.27e-63 |
| Malignant epithelial | C2 WNT ECs | LRRC42 | 1.66 | 0.33 | 0.111 | 1.69e-56 | 5.679999999999999e-52 |
| Malignant epithelial | C2 WNT ECs | PERP | 1.659 | 0.995 | 0.879 | 5.899999999999999e-206 | 1.99e-201 |
| Malignant epithelial | C2 WNT ECs | DUSP5 | 1.659 | 0.445 | 0.319 | 2.82e-18 | 9.49e-14 |
| Malignant epithelial | C2 WNT ECs | DNAJA4 | 1.658 | 0.441 | 0.212 | 6.79e-45 | 2.29e-40 |
| Malignant epithelial | C2 WNT ECs | NEBL | 1.656 | 0.322 | 0.086 | 8.820000000000001e-73 | 2.97e-68 |
| Malignant epithelial | C2 WNT ECs | HSD17B7 | 1.655 | 0.219 | 0.059 | 1.15e-47 | 3.86e-43 |
| Malignant epithelial | C2 WNT ECs | TSPAN6 | 1.654 | 0.648 | 0.304 | 1.27e-86 | 4.29e-82 |
| Malignant epithelial | C2 WNT ECs | PVRL4 | 1.653 | 0.28 | 0.124 | 1.11e-29 | 3.75e-25 |
| Malignant epithelial | C2 WNT ECs | MAP1LC3A | 1.65 | 0.566 | 0.202 | 2.27e-100 | 7.66e-96 |
| Malignant epithelial | C2 WNT ECs | FBLIM1 | 1.648 | 0.254 | 0.079 | 2.31e-46 | 7.79e-42 |
| Malignant epithelial | C2 WNT ECs | MCF2L | 1.648 | 0.181 | 0.05 | 1.45e-38 | 4.9e-34 |
| Malignant epithelial | C2 WNT ECs | MAD1L1 | 1.648 | 0.267 | 0.106 | 1.68e-33 | 5.66e-29 |
| Malignant epithelial | C2 WNT ECs | SRPX2 | 1.646 | 0.15 | 0.046 | 6.05e-28 | 2.04e-23 |
| Malignant epithelial | C2 WNT ECs | CLDN23 | 1.645 | 0.131 | 0.045 | 8.530000000000001e-21 | 2.87e-16 |
| Malignant epithelial | C2 WNT ECs | SERPINB8 | 1.64 | 0.131 | 0.041 | 1.09e-23 | 3.69e-19 |
| Malignant epithelial | C2 WNT ECs | TMEM45B | 1.635 | 0.686 | 0.244 | 3.09e-135 | 1.04e-130 |

| compartment | subcluster | gene | avg_log2FC | pct.1 | pct.2 | p_val | p_val_adj |
|---|---|---|---|---|---|---|---|
| Malignant epithelial | C2 WNT ECs | AMPD3 | 1.634 | 0.107 | 0.031 | 6e-22 | 2.02e-17 |
| Malignant epithelial | C2 WNT ECs | TNFSF15 | 1.631 | 0.141 | 0.048 | 3.45e-22 | 1.16e-17 |
| Malignant epithelial | C2 WNT ECs | NUFIP1 | 1.629 | 0.245 | 0.077 | 1.96e-44 | 6.61e-40 |
| Malignant epithelial | C2 WNT ECs | GRAMD1C | 1.629 | 0.124 | 0.031 | 1.45e-29 | 4.88e-25 |
| Malignant epithelial | C2 WNT ECs | FAM83B | 1.628 | 0.134 | 0.04 | 2.64e-25 | 8.880000000000001e-21 |
| Malignant epithelial | C2 WNT ECs | MAPRE3 | 1.626 | 0.208 | 0.059 | 2.13e-42 | 7.159999999999999e-38 |
| Malignant epithelial | C2 WNT ECs | DUSP22 | 1.625 | 0.319 | 0.092 | 4.53e-66 | 1.53e-61 |
| Malignant epithelial | C2 WNT ECs | FNBP1L | 1.619 | 0.321 | 0.098 | 2.32e-60 | 7.820000000000001e-56 |
| Malignant epithelial | C2 WNT ECs | CBLB | 1.617 | 0.221 | 0.063 | 1.84e-44 | 6.21e-40 |
| Malignant epithelial | C2 WNT ECs | PJA1 | 1.614 | 0.177 | 0.047 | 3.28e-39 | 1.1e-34 |
| Malignant epithelial | C2 WNT ECs | FAM188A | 1.614 | 0.166 | 0.048 | 2.1e-33 | 7.06e-29 |
| Malignant epithelial | C2 WNT ECs | RGS2 | 1.613 | 0.417 | 0.164 | 7.53e-58 | 2.54e-53 |
| Malignant epithelial | C2 WNT ECs | GPATCH3 | 1.611 | 0.146 | 0.044 | 2.59e-27 | 8.729999999999999e-23 |
| Malignant epithelial | C2 WNT ECs | MAOA | 1.608 | 0.265 | 0.086 | 3.13e-45 | 1.05e-40 |
| Malignant epithelial | C2 WNT ECs | HECA | 1.607 | 0.245 | 0.084 | 8.77e-39 | 2.95e-34 |
| Malignant epithelial | C2 WNT ECs | PRAC2 | 1.607 | 0.115 | 0.031 | 4.69e-25 | 1.58e-20 |
| Malignant epithelial | C2 WNT ECs | NDFIP2 | 1.603 | 0.682 | 0.26 | 1.85e-124 | 6.24e-120 |
| Malignant epithelial | C2 WNT ECs | KCTD11 | 1.603 | 0.237 | 0.082 | 3.22e-37 | 1.09e-32 |
| Malignant epithelial | C2 WNT ECs | ACTR3C | 1.603 | 0.162 | 0.047 | 6.06e-32 | 2.04e-27 |
| Malignant epithelial | C2 WNT ECs | PTPRR | 1.6 | 0.122 | 0.031 | 6.67e-29 | 2.25e-24 |
| Malignant epithelial | C2 WNT ECs | IL1RN | 1.6 | 0.143 | 0.055 | 1.05e-18 | 3.54e-14 |
| Malignant epithelial | C2 WNT ECs | KCNQ1 | 1.599 | 0.279 | 0.084 | 1.53e-53 | 5.15e-49 |
| Malignant epithelial | C2 WNT ECs | FUCA1 | 1.598 | 0.321 | 0.102 | 3.27e-58 | 1.1e-53 |
| Malignant epithelial | C2 WNT ECs | N4BP2L1 | 1.596 | 0.21 | 0.07 | 2.48e-34 | 8.339999999999999e-30 |
| Malignant epithelial | C2 WNT ECs | ZNF823 | 1.595 | 0.133 | 0.044 | 6.79e-22 | 2.29e-17 |
| Malignant epithelial | C2 WNT ECs | TRIM15 | 1.594 | 0.28 | 0.091 | 1.03e-48 | 3.48e-44 |
| Malignant epithelial | C2 WNT ECs | TRIM36 | 1.594 | 0.124 | 0.038 | 7.489999999999999e-23 | 2.52e-18 |
| Malignant epithelial | C2 WNT ECs | DUSP6 | 1.593 | 0.558 | 0.235 | 1.42e-78 | 4.789999999999999e-74 |
| Malignant epithelial | C2 WNT ECs | EVI5 | 1.591 | 0.187 | 0.051 | 3.39e-40 | 1.14e-35 |
| Malignant epithelial | C2 WNT ECs | FRAT2 | 1.587 | 0.226 | 0.067 | 1.45e-43 | 4.89e-39 |
| Malignant epithelial | C2 WNT ECs | AIM1L | 1.586 | 0.161 | 0.05 | 4.56e-29 | 1.54e-24 |
| Malignant epithelial | C2 WNT ECs | DSG3 | 1.585 | 0.306 | 0.123 | 1.94e-39 | 6.55e-35 |
| Malignant epithelial | C2 WNT ECs | CYB5B | 1.583 | 0.89 | 0.596 | 1.37e-120 | 4.61e-116 |
| Malignant epithelial | C2 WNT ECs | PDGFB | 1.58 | 0.101 | 0.031 | 2.33e-19 | 7.85e-15 |
| Malignant epithelial | C2 WNT ECs | PLEKHA4 | 1.578 | 0.192 | 0.057 | 1.51e-37 | 5.08e-33 |
| Malignant epithelial | C2 WNT ECs | IDI1 | 1.577 | 0.514 | 0.217 | 1.75e-69 | 5.91e-65 |
| Malignant epithelial | C2 WNT ECs | CD24 | 1.575 | 0.951 | 0.411 | 4.56e-187 | 1.54e-182 |
| Malignant epithelial | C2 WNT ECs | GATA6 | 1.575 | 0.221 | 0.063 | 4.01e-45 | 1.35e-40 |
| Malignant epithelial | C2 WNT ECs | ODC1 | 1.574 | 0.714 | 0.403 | 2.6e-76 | 8.750000000000001e-72 |
| Malignant epithelial | C2 WNT ECs | PCMTD2 | 1.572 | 0.414 | 0.148 | 1.45e-67 | 4.88e-63 |
| Malignant epithelial | C2 WNT ECs | RP1-313I6.12 | 1.571 | 0.474 | 0.219 | 1.35e-51 | 4.55e-47 |
| Malignant epithelial | C2 WNT ECs | BPHL | 1.569 | 0.241 | 0.075 | 2.08e-43 | 7.01e-39 |
| Malignant epithelial | C2 WNT ECs | ABCA5 | 1.569 | 0.168 | 0.044 | 5.01e-38 | 1.69e-33 |
| Malignant epithelial | C2 WNT ECs | CNN3 | 1.568 | 0.479 | 0.188 | 2.5e-70 | 8.43e-66 |
| Malignant epithelial | C2 WNT ECs | PAN3 | 1.568 | 0.367 | 0.12 | 1.23e-64 | 4.15e-60 |
| Malignant epithelial | C2 WNT ECs | SNX9 | 1.567 | 0.432 | 0.171 | 5.44e-60 | 1.83e-55 |
| Malignant epithelial | C2 WNT ECs | KIZ | 1.566 | 0.183 | 0.053 | 1.24e-36 | 4.19e-32 |
| Malignant epithelial | C2 WNT ECs | TUBB3 | 1.566 | 0.106 | 0.028 | 1.91e-23 | 6.43e-19 |
| Malignant epithelial | C2 WNT ECs | GOLGB1 | 1.565 | 0.491 | 0.163 | 1.64e-89 | 5.520000000000001e-85 |
| Malignant epithelial | C2 WNT ECs | C11orf54 | 1.565 | 0.303 | 0.092 | 1.88e-57 | 6.33e-53 |
| Malignant epithelial | C2 WNT ECs | GAST | 1.563 | 0.135 | 0.036 | 2.33e-30 | 7.870000000000001e-26 |
| Malignant epithelial | C2 WNT ECs | RP11-540A21.2 | 1.563 | 0.137 | 0.044 | 1.38e-23 | 4.66e-19 |
| Malignant epithelial | C2 WNT ECs | DNASE1 | 1.561 | 0.23 | 0.061 | 1.14e-51 | 3.83e-47 |
| Malignant epithelial | C2 WNT ECs | B3GNT8 | 1.559 | 0.181 | 0.046 | 5.49e-42 | 1.85e-37 |
| Malignant epithelial | C2 WNT ECs | EGR2 | 1.559 | 0.101 | 0.033 | 6.64e-18 | 2.24e-13 |
| Malignant epithelial | C2 WNT ECs | CRYZ | 1.556 | 0.276 | 0.078 | 6.42e-57 | 2.16e-52 |
| Malignant epithelial | C2 WNT ECs | TIMP3 | 1.556 | 0.226 | 0.079 | 3.69e-34 | 1.24e-29 |
| Malignant epithelial | C2 WNT ECs | NUP58 | 1.552 | 0.252 | 0.084 | 1e-40 | 3.38e-36 |
| Malignant epithelial | C2 WNT ECs | DLL4 | 1.551 | 0.111 | 0.027 | 4.78e-28 | 1.61e-23 |
| Malignant epithelial | C2 WNT ECs | EFNB1 | 1.549 | 0.501 | 0.179 | 1.7e-80 | 5.72e-76 |
| Malignant epithelial | C2 WNT ECs | ACOT11 | 1.543 | 0.199 | 0.057 | 9.52e-40 | 3.21e-35 |
| Malignant epithelial | C2 WNT ECs | CLOCK | 1.543 | 0.187 | 0.061 | 1.4e-30 | 4.7e-26 |
| Malignant epithelial | C2 WNT ECs | ACSL5 | 1.542 | 0.723 | 0.33 | 1.96e-108 | 6.59e-104 |
| Malignant epithelial | C2 WNT ECs | HIST1H2AE | 1.54 | 0.295 | 0.112 | 3.01e-40 | 1.01e-35 |

| compartment | subcluster | gene | avg_log2FC | pct.1 | pct.2 | p_val | p_val_adj |
|---|---|---|---|---|---|---|---|
| Malignant epithelial | C2 WNT ECs | IFT88 | 1.54 | 0.202 | 0.059 | 5.18e-39 | 1.75e-34 |
| Malignant epithelial | C2 WNT ECs | CCDC126 | 1.537 | 0.184 | 0.06 | 7.74e-31 | 2.61e-26 |
| Malignant epithelial | C2 WNT ECs | N4BP2L2 | 1.536 | 0.843 | 0.46 | 6.320000000000001e-125 | 2.13e-120 |
| Malignant epithelial | C2 WNT ECs | RANGRF | 1.536 | 0.235 | 0.076 | 2.7e-40 | 9.1e-36 |
| Malignant epithelial | C2 WNT ECs | SUGT1 | 1.532 | 0.775 | 0.364 | 3.43e-124 | 1.16e-119 |
| Malignant epithelial | C2 WNT ECs | DUSP14 | 1.532 | 0.28 | 0.146 | 2.86e-20 | 9.65e-16 |
| Malignant epithelial | C2 WNT ECs | DSP | 1.53 | 0.767 | 0.398 | 8.63e-102 | 2.91e-97 |
| Malignant epithelial | C2 WNT ECs | SBF1 | 1.53 | 0.295 | 0.095 | 7.58e-52 | 2.55e-47 |
| Malignant epithelial | C2 WNT ECs | RORA | 1.528 | 0.2 | 0.065 | 2.59e-34 | 8.72e-30 |
| Malignant epithelial | C2 WNT ECs | FOXO1 | 1.528 | 0.135 | 0.051 | 2.75e-18 | 9.27e-14 |
| Malignant epithelial | C2 WNT ECs | GRTP1 | 1.526 | 0.419 | 0.148 | 5.059999999999999e-69 | 1.7e-64 |
| Malignant epithelial | C2 WNT ECs | VIL1 | 1.525 | 0.708 | 0.318 | 2.3e-108 | 7.74e-104 |
| Malignant epithelial | C2 WNT ECs | COX6B2 | 1.525 | 0.184 | 0.055 | 6.05e-35 | 2.04e-30 |
| Malignant epithelial | C2 WNT ECs | IFRD1 | 1.524 | 0.889 | 0.618 | 7.14e-112 | 2.41e-107 |
| Malignant epithelial | C2 WNT ECs | HES1 | 1.524 | 0.876 | 0.689 | 3.62e-41 | 1.22e-36 |
| Malignant epithelial | C2 WNT ECs | NMU | 1.524 | 0.225 | 0.088 | 3.12e-29 | 1.05e-24 |
| Malignant epithelial | C2 WNT ECs | MPHOSPH8 | 1.522 | 0.729 | 0.327 | 1.49e-107 | 5.010000000000001e-103 |
| Malignant epithelial | C2 WNT ECs | TNKS2 | 1.522 | 0.212 | 0.063 | 1.31e-40 | 4.41e-36 |
| Malignant epithelial | C2 WNT ECs | HDAC4 | 1.519 | 0.135 | 0.044 | 7.339999999999999e-23 | 2.47e-18 |
| Malignant epithelial | C2 WNT ECs | GGT7 | 1.518 | 0.237 | 0.088 | 1.66e-33 | 5.61e-29 |
| Malignant epithelial | C2 WNT ECs | SATB2 | 1.511 | 0.161 | 0.053 | 3.27e-27 | 1.1e-22 |
| Malignant epithelial | C2 WNT ECs | GFOD1 | 1.511 | 0.124 | 0.044 | 1.36e-18 | 4.57e-14 |
| Malignant epithelial | C2 WNT ECs | RNF11 | 1.51 | 0.407 | 0.144 | 1.3e-64 | 4.38e-60 |
| Malignant epithelial | C2 WNT ECs | CCDC24 | 1.51 | 0.18 | 0.05 | 2.34e-37 | 7.87e-33 |
| Malignant epithelial | C2 WNT ECs | C11orf49 | 1.506 | 0.273 | 0.083 | 2.1e-51 | 7.070000000000001e-47 |
| Malignant epithelial | C2 WNT ECs | CTGF | 1.506 | 0.365 | 0.153 | 1.79e-44 | 6.05e-40 |
| Malignant epithelial | C2 WNT ECs | FAM83G | 1.506 | 0.111 | 0.037 | 5.02e-18 | 1.69e-13 |
| Malignant epithelial | C2 WNT ECs | GTF2IRD1 | 1.505 | 0.346 | 0.113 | 8.55e-60 | 2.88e-55 |
| Malignant epithelial | C2 WNT ECs | IL22RA1 | 1.505 | 0.162 | 0.048 | 3.7e-31 | 1.25e-26 |
| Malignant epithelial | C2 WNT ECs | MTIF3 | 1.502 | 0.775 | 0.375 | 8.03e-118 | 2.71e-113 |
| Malignant epithelial | C2 WNT ECs | ABHD13 | 1.502 | 0.3 | 0.092 | 1.92e-56 | 6.46e-52 |
| Malignant epithelial | C2 WNT ECs | PIM1 | 1.501 | 0.524 | 0.247 | 1.95e-57 | 6.59e-53 |
| Malignant epithelial | C2 WNT ECs | CD99 | 1.5 | 0.821 | 0.456 | 9.720000000000001e-122 | 3.28e-117 |
| Malignant epithelial | C2 WNT ECs | CCL3L3 | 1.498 | 0.373 | 0.148 | 3.76e-47 | 1.27e-42 |
| Malignant epithelial | C2 WNT ECs | B3GNT2 | 1.496 | 0.353 | 0.13 | 4.529999999999999e-52 | 1.53e-47 |
| Malignant epithelial | C2 WNT ECs | ZNF608 | 1.494 | 0.116 | 0.034 | 7.4e-23 | 2.49e-18 |
| Malignant epithelial | C2 WNT ECs | AGO2 | 1.493 | 0.502 | 0.195 | 6.61e-75 | 2.23e-70 |
| Malignant epithelial | C2 WNT ECs | RNF219 | 1.493 | 0.124 | 0.043 | 2.06e-19 | 6.94e-15 |
| Malignant epithelial | C2 WNT ECs | MAGED2 | 1.492 | 0.564 | 0.238 | 2.9e-78 | 9.78e-74 |
| Malignant epithelial | C2 WNT ECs | SCD | 1.49 | 0.572 | 0.34 | 9.229999999999999e-40 | 3.11e-35 |
| Malignant epithelial | C2 WNT ECs | CCSER2 | 1.489 | 0.307 | 0.1 | 4.39e-53 | 1.48e-48 |
| Malignant epithelial | C2 WNT ECs | GRHL1 | 1.486 | 0.166 | 0.055 | 3.27e-27 | 1.1e-22 |
| Malignant epithelial | C2 WNT ECs | SERINC2 | 1.485 | 0.771 | 0.419 | 8.77e-102 | 2.95e-97 |
| Malignant epithelial | C2 WNT ECs | SNHG18 | 1.482 | 0.373 | 0.115 | 1.02e-69 | 3.44e-65 |
| Malignant epithelial | C2 WNT ECs | NAPG | 1.477 | 0.31 | 0.117 | 1.37e-42 | 4.61e-38 |
| Malignant epithelial | C2 WNT ECs | RP11-347P5.1 | 1.475 | 0.223 | 0.062 | 9.84e-47 | 3.31e-42 |
| Malignant epithelial | C2 WNT ECs | TMEM234 | 1.472 | 0.198 | 0.057 | 3.9e-39 | 1.31e-34 |
| Malignant epithelial | C2 WNT ECs | ZMYM2 | 1.471 | 0.553 | 0.222 | 6.569999999999999e-81 | 2.21e-76 |
| Malignant epithelial | C2 WNT ECs | CPNE2 | 1.471 | 0.198 | 0.063 | 7.639999999999999e-34 | 2.57e-29 |
| Malignant epithelial | C2 WNT ECs | FNDC3A | 1.47 | 0.317 | 0.096 | 2.09e-60 | 7.030000000000001e-56 |
| Malignant epithelial | C2 WNT ECs | RASSF5 | 1.469 | 0.123 | 0.043 | 7.96e-19 | 2.68e-14 |
| Malignant epithelial | C2 WNT ECs | DEK | 1.467 | 0.903 | 0.546 | 1.09e-137 | 3.68e-133 |
| Malignant epithelial | C2 WNT ECs | HERPUD1 | 1.466 | 0.854 | 0.59 | 1.09e-84 | 3.68e-80 |
| Malignant epithelial | C2 WNT ECs | HOXA-AS2 | 1.465 | 0.146 | 0.05 | 5.7e-23 | 1.92e-18 |
| Malignant epithelial | C2 WNT ECs | CLN5 | 1.464 | 0.292 | 0.09 | 2.2e-54 | 7.43e-50 |
| Malignant epithelial | C2 WNT ECs | DMTN | 1.462 | 0.111 | 0.034 | 5.880000000000001e-21 | 1.98e-16 |
| Malignant epithelial | C2 WNT ECs | PGF | 1.461 | 0.134 | 0.046 | 5.110000000000001e-21 | 1.72e-16 |
| Malignant epithelial | C2 WNT ECs | SMTN | 1.46 | 0.382 | 0.148 | 2.28e-53 | 7.68e-49 |
| Malignant epithelial | C2 WNT ECs | SNTB1 | 1.459 | 0.222 | 0.07 | 4.5e-39 | 1.52e-34 |
| Malignant epithelial | C2 WNT ECs | ULK1 | 1.458 | 0.21 | 0.081 | 2.41e-27 | 8.119999999999999e-23 |
| Malignant epithelial | C2 WNT ECs | C1orf210 | 1.456 | 0.283 | 0.1 | 4.06e-43 | 1.37e-38 |
| Malignant epithelial | C2 WNT ECs | TLE4 | 1.455 | 0.337 | 0.111 | 4.84e-56 | 1.63e-51 |
| Malignant epithelial | C2 WNT ECs | GJB4 | 1.454 | 0.161 | 0.049 | 1.7e-29 | 5.71e-25 |
| Malignant epithelial | C2 WNT ECs | RASA3 | 1.454 | 0.166 | 0.065 | 5.22e-21 | 1.76e-16 |

| compartment | subcluster | gene | avg_log2FC | pct.1 | pct.2 | p_val | p_val_adj |
|---|---|---|---|---|---|---|---|
| Malignant epithelial | C2 WNT ECs | GALNT6 | 1.452 | 0.214 | 0.075 | 2e-32 | 6.74e-28 |
| Malignant epithelial | C2 WNT ECs | PACSIN3 | 1.451 | 0.459 | 0.161 | 1.4e-74 | 4.72e-70 |
| Malignant epithelial | C2 WNT ECs | MGLL | 1.449 | 0.467 | 0.23 | 1.52e-42 | 5.109999999999999e-38 |
| Malignant epithelial | C2 WNT ECs | ALG5 | 1.446 | 0.674 | 0.303 | 4.79e-95 | 1.61e-90 |
| Malignant epithelial | C2 WNT ECs | PLA2G15 | 1.446 | 0.139 | 0.046 | 1.59e-23 | 5.35e-19 |
| Malignant epithelial | C2 WNT ECs | TMEM59 | 1.445 | 0.907 | 0.65 | 2.63e-112 | 8.85e-108 |
| Malignant epithelial | C2 WNT ECs | BCOR | 1.445 | 0.104 | 0.038 | 8.55e-15 | 2.88e-10 |
| Malignant epithelial | C2 WNT ECs | KDM1A | 1.443 | 0.384 | 0.141 | 5.24e-57 | 1.77e-52 |
| Malignant epithelial | C2 WNT ECs | LMBR1L | 1.441 | 0.198 | 0.059 | 4.3e-37 | 1.45e-32 |
| Malignant epithelial | C2 WNT ECs | PCK1 | 1.441 | 0.218 | 0.083 | 1.95e-29 | 6.56e-25 |
| Malignant epithelial | C2 WNT ECs | MPP1 | 1.44 | 0.219 | 0.071 | 2.7e-37 | 9.100000000000001e-33 |
| Malignant epithelial | C2 WNT ECs | NAA30 | 1.437 | 0.253 | 0.109 | 3.94e-27 | 1.33e-22 |
| Malignant epithelial | C2 WNT ECs | BCL2A1 | 1.437 | 0.12 | 0.047 | 5.18e-15 | 1.75e-10 |
| Malignant epithelial | C2 WNT ECs | DNAJC3 | 1.436 | 0.583 | 0.235 | 8.95e-84 | 3.010000000000001e-79 |
| Malignant epithelial | C2 WNT ECs | SH3BGRL2 | 1.436 | 0.338 | 0.111 | 3.86e-58 | 1.3e-53 |
| Malignant epithelial | C2 WNT ECs | ABHD3 | 1.436 | 0.376 | 0.152 | 2.51e-47 | 8.47e-43 |
| Malignant epithelial | C2 WNT ECs | PPL | 1.435 | 0.371 | 0.181 | 1.8e-32 | 6.06e-28 |
| Malignant epithelial | C2 WNT ECs | DBN1 | 1.434 | 0.322 | 0.119 | 1.22e-46 | 4.1e-42 |
| Malignant epithelial | C2 WNT ECs | PPP1R9A | 1.434 | 0.133 | 0.04 | 1.34e-24 | 4.52e-20 |
| Malignant epithelial | C2 WNT ECs | ALDH2 | 1.433 | 0.595 | 0.241 | 1.43e-88 | 4.82e-84 |
| Malignant epithelial | C2 WNT ECs | TMCO3 | 1.432 | 0.436 | 0.154 | 1.86e-71 | 6.26e-67 |
| Malignant epithelial | C2 WNT ECs | KPNA3 | 1.432 | 0.357 | 0.13 | 3.78e-53 | 1.27e-48 |
| Malignant epithelial | C2 WNT ECs | FGGY | 1.432 | 0.207 | 0.073 | 4.18e-31 | 1.41e-26 |
| Malignant epithelial | C2 WNT ECs | PHF11 | 1.431 | 0.345 | 0.126 | 7.79e-51 | 2.62e-46 |
| Malignant epithelial | C2 WNT ECs | TPT1 | 1.429 | 1.0 | 0.995 | 3.03e-282 | 1.02e-277 |
| Malignant epithelial | C2 WNT ECs | ZNF33B | 1.429 | 0.272 | 0.086 | 3.72e-48 | 1.25e-43 |
| Malignant epithelial | C2 WNT ECs | NUP153 | 1.429 | 0.164 | 0.05 | 4.14e-30 | 1.4e-25 |
| Malignant epithelial | C2 WNT ECs | TLE2 | 1.428 | 0.18 | 0.066 | 2.14e-25 | 7.21e-21 |
| Malignant epithelial | C2 WNT ECs | SIAH2 | 1.426 | 0.467 | 0.181 | 1.63e-66 | 5.51e-62 |
| Malignant epithelial | C2 WNT ECs | TNIK | 1.426 | 0.222 | 0.067 | 4e-41 | 1.35e-36 |
| Malignant epithelial | C2 WNT ECs | VAT1 | 1.425 | 0.369 | 0.115 | 1.87e-68 | 6.31e-64 |
| Malignant epithelial | C2 WNT ECs | ZNF426 | 1.424 | 0.118 | 0.041 | 3.73e-18 | 1.26e-13 |
| Malignant epithelial | C2 WNT ECs | LAMP1 | 1.423 | 0.903 | 0.556 | 5.78e-155 | 1.95e-150 |
| Malignant epithelial | C2 WNT ECs | AKR1C3 | 1.423 | 0.605 | 0.258 | 5.1e-85 | 1.72e-80 |
| Malignant epithelial | C2 WNT ECs | ZNF562 | 1.423 | 0.235 | 0.076 | 1.29e-40 | 4.33e-36 |
| Malignant epithelial | C2 WNT ECs | PEPD | 1.42 | 0.572 | 0.234 | 6.99e-83 | 2.36e-78 |
| Malignant epithelial | C2 WNT ECs | LPGAT1 | 1.419 | 0.378 | 0.165 | 3.559999999999999e-42 | 1.2e-37 |
| Malignant epithelial | C2 WNT ECs | EPB41L5 | 1.419 | 0.217 | 0.068 | 7.38e-38 | 2.49e-33 |
| Malignant epithelial | C2 WNT ECs | C18orf8 | 1.418 | 0.143 | 0.039 | 2.63e-30 | 8.87e-26 |
| Malignant epithelial | C2 WNT ECs | WDR1 | 1.416 | 0.673 | 0.308 | 6.28e-93 | 2.12e-88 |
| Malignant epithelial | C2 WNT ECs | GLRX3 | 1.416 | 0.667 | 0.338 | 1.82e-80 | 6.14e-76 |
| Malignant epithelial | C2 WNT ECs | ARID5B | 1.416 | 0.382 | 0.153 | 2.09e-49 | 7.03e-45 |
| Malignant epithelial | C2 WNT ECs | NPC1 | 1.415 | 0.138 | 0.041 | 1.3e-26 | 4.36e-22 |
| Malignant epithelial | C2 WNT ECs | CCDC64B | 1.414 | 0.595 | 0.254 | 8.71e-81 | 2.93e-76 |
| Malignant epithelial | C2 WNT ECs | HIST1H2BG | 1.414 | 0.414 | 0.186 | 6.36e-43 | 2.14e-38 |
| Malignant epithelial | C2 WNT ECs | ARID3B | 1.414 | 0.162 | 0.049 | 2.49e-30 | 8.4e-26 |
| Malignant epithelial | C2 WNT ECs | CAP1 | 1.412 | 0.75 | 0.393 | 1.23e-94 | 4.15e-90 |
| Malignant epithelial | C2 WNT ECs | IL1B | 1.412 | 0.221 | 0.08 | 1.12e-30 | 3.77e-26 |
| Malignant epithelial | C2 WNT ECs | GLUD1 | 1.41 | 0.566 | 0.262 | 2.75e-67 | 9.250000000000001e-63 |
| Malignant epithelial | C2 WNT ECs | ZNF57 | 1.409 | 0.137 | 0.053 | 1.17e-17 | 3.96e-13 |
| Malignant epithelial | C2 WNT ECs | ZDHHC20 | 1.408 | 0.386 | 0.139 | 4.460000000000001e-58 | 1.5e-53 |
| Malignant epithelial | C2 WNT ECs | PRKCZ | 1.405 | 0.321 | 0.117 | 2.14e-46 | 7.21e-42 |
| Malignant epithelial | C2 WNT ECs | DHRS12 | 1.405 | 0.279 | 0.094 | 4.65e-45 | 1.57e-40 |
| Malignant epithelial | C2 WNT ECs | CDK8 | 1.403 | 0.187 | 0.061 | 1.5e-30 | 5.06e-26 |
| Malignant epithelial | C2 WNT ECs | SREK1IP1 | 1.402 | 0.635 | 0.343 | 1.33e-58 | 4.49e-54 |
| Malignant epithelial | C2 WNT ECs | ZBTB20 | 1.401 | 0.3 | 0.1 | 9.14e-49 | 3.08e-44 |
| Malignant epithelial | C2 WNT ECs | FBXO3 | 1.4 | 0.29 | 0.119 | 1.8e-34 | 6.08e-30 |
| Malignant epithelial | C2 WNT ECs | MFN2 | 1.4 | 0.154 | 0.055 | 2.36e-22 | 7.94e-18 |
| Malignant epithelial | C2 WNT ECs | DNAJB1 | 1.399 | 0.931 | 0.84 | 2.66e-64 | 8.959999999999999e-60 |
| Malignant epithelial | C2 WNT ECs | PMM1 | 1.396 | 0.394 | 0.138 | 4.56e-62 | 1.53e-57 |
| Malignant epithelial | C2 WNT ECs | TBC1D4 | 1.396 | 0.106 | 0.046 | 4.13e-11 | 1.39e-06 |
| Malignant epithelial | C2 WNT ECs | KLF5 | 1.395 | 0.881 | 0.554 | 6.5e-117 | 2.19e-112 |
| Malignant epithelial | C2 WNT ECs | KLC4 | 1.395 | 0.378 | 0.134 | 1.02e-57 | 3.44e-53 |
| Malignant epithelial | C2 WNT ECs | ZNF385A | 1.395 | 0.179 | 0.052 | 4.34e-35 | 1.46e-30 |

| compartment | subcluster | gene | avg_log2FC | pct.1 | pct.2 | p_val | p_val_adj |
|---|---|---|---|---|---|---|---|
| Malignant epithelial | C2 WNT ECs | ABI1 | 1.393 | 0.567 | 0.217 | 2.9e-86 | 9.78e-82 |
| Malignant epithelial | C2 WNT ECs | RAP2A | 1.393 | 0.166 | 0.053 | 9.74e-29 | 3.28e-24 |
| Malignant epithelial | C2 WNT ECs | ZBTB17 | 1.393 | 0.116 | 0.041 | 1.08e-17 | 3.64e-13 |
| Malignant epithelial | C2 WNT ECs | TMEM184B | 1.392 | 0.261 | 0.092 | 4.15e-39 | 1.4e-34 |
| Malignant epithelial | C2 WNT ECs | CCDC61 | 1.392 | 0.23 | 0.079 | 1.11e-35 | 3.74e-31 |
| Malignant epithelial | C2 WNT ECs | ARHGAP12 | 1.388 | 0.337 | 0.121 | 2.8e-50 | 9.43e-46 |
| Malignant epithelial | C2 WNT ECs | STAM | 1.384 | 0.204 | 0.071 | 1.44e-31 | 4.850000000000001e-27 |
| Malignant epithelial | C2 WNT ECs | CBR4 | 1.383 | 0.242 | 0.087 | 3.42e-35 | 1.15e-30 |
| Malignant epithelial | C2 WNT ECs | MTMR6 | 1.383 | 0.153 | 0.045 | 6.63e-29 | 2.23e-24 |
| Malignant epithelial | C2 WNT ECs | CLRN3 | 1.382 | 0.361 | 0.149 | 1.75e-43 | 5.91e-39 |
| Malignant epithelial | C2 WNT ECs | COG3 | 1.381 | 0.349 | 0.123 | 2.41e-53 | 8.12e-49 |
| Malignant epithelial | C2 WNT ECs | FCHSD1 | 1.38 | 0.104 | 0.034 | 1.28e-17 | 4.3e-13 |
| Malignant epithelial | C2 WNT ECs | ARMCX3 | 1.379 | 0.478 | 0.198 | 3.61e-61 | 1.22e-56 |
| Malignant epithelial | C2 WNT ECs | MT-ATP8 | 1.378 | 0.198 | 0.067 | 6.09e-31 | 2.05e-26 |
| Malignant epithelial | C2 WNT ECs | LYRM1 | 1.376 | 0.472 | 0.195 | 8.22e-60 | 2.77e-55 |
| Malignant epithelial | C2 WNT ECs | HECTD3 | 1.374 | 0.122 | 0.033 | 1.48e-26 | 4.99e-22 |
| Malignant epithelial | C2 WNT ECs | PBXIP1 | 1.373 | 0.263 | 0.086 | 2.81e-43 | 9.47e-39 |
| Malignant epithelial | C2 WNT ECs | PLCD1 | 1.372 | 0.107 | 0.034 | 6.24e-19 | 2.1e-14 |
| Malignant epithelial | C2 WNT ECs | FAM35A | 1.369 | 0.153 | 0.052 | 3.42e-24 | 1.15e-19 |
| Malignant epithelial | C2 WNT ECs | ZNF513 | 1.369 | 0.115 | 0.038 | 3.78e-19 | 1.27e-14 |
| Malignant epithelial | C2 WNT ECs | BSDC1 | 1.368 | 0.342 | 0.124 | 8.56e-50 | 2.88e-45 |
| Malignant epithelial | C2 WNT ECs | RRAGC | 1.368 | 0.256 | 0.098 | 7.68e-34 | 2.59e-29 |
| Malignant epithelial | C2 WNT ECs | MAP1LC3B | 1.367 | 0.903 | 0.594 | 3.07e-126 | 1.03e-121 |
| Malignant epithelial | C2 WNT ECs | GHITM | 1.367 | 0.767 | 0.391 | 1.21e-102 | 4.08e-98 |
| Malignant epithelial | C2 WNT ECs | INPP5D | 1.366 | 0.24 | 0.077 | 3.66e-41 | 1.23e-36 |
| Malignant epithelial | C2 WNT ECs | GTF3A | 1.362 | 0.874 | 0.584 | 9.450000000000001e-110 | 3.18e-105 |
| Malignant epithelial | C2 WNT ECs | MKNK1 | 1.361 | 0.208 | 0.079 | 1.57e-27 | 5.28e-23 |
| Malignant epithelial | C2 WNT ECs | TRIM26 | 1.36 | 0.231 | 0.082 | 6.029999999999999e-34 | 2.03e-29 |
| Malignant epithelial | C2 WNT ECs | CAT | 1.359 | 0.36 | 0.144 | 1.46e-45 | 4.929999999999999e-41 |
| Malignant epithelial | C2 WNT ECs | URGCP | 1.359 | 0.249 | 0.094 | 1.33e-33 | 4.49e-29 |
| Malignant epithelial | C2 WNT ECs | RNF125 | 1.359 | 0.111 | 0.04 | 6.31e-16 | 2.12e-11 |
| Malignant epithelial | C2 WNT ECs | SLC5A6 | 1.356 | 0.252 | 0.092 | 1.15e-35 | 3.87e-31 |
| Malignant epithelial | C2 WNT ECs | FAM109A | 1.352 | 0.147 | 0.052 | 4.16e-22 | 1.4e-17 |
| Malignant epithelial | C2 WNT ECs | GPR143 | 1.352 | 0.104 | 0.034 | 3.45e-17 | 1.16e-12 |
| Malignant epithelial | C2 WNT ECs | PDK2 | 1.351 | 0.345 | 0.132 | 2.26e-46 | 7.62e-42 |
| Malignant epithelial | C2 WNT ECs | FMNL1 | 1.35 | 0.108 | 0.031 | 9.22e-22 | 3.11e-17 |
| Malignant epithelial | C2 WNT ECs | HIVEP2 | 1.35 | 0.107 | 0.04 | 1.34e-14 | 4.5e-10 |
| Malignant epithelial | C2 WNT ECs | TPP2 | 1.348 | 0.348 | 0.124 | 1.96e-52 | 6.6e-48 |
| Malignant epithelial | C2 WNT ECs | PHTF1 | 1.346 | 0.119 | 0.041 | 2.01e-18 | 6.76e-14 |
| Malignant epithelial | C2 WNT ECs | NABP1 | 1.345 | 0.271 | 0.107 | 5.52e-33 | 1.86e-28 |
| Malignant epithelial | C2 WNT ECs | ZMIZ1 | 1.345 | 0.162 | 0.058 | 1.24e-23 | 4.17e-19 |
| Malignant epithelial | C2 WNT ECs | TLE3 | 1.344 | 0.349 | 0.131 | 1.69e-49 | 5.68e-45 |
| Malignant epithelial | C2 WNT ECs | MBIP | 1.344 | 0.246 | 0.106 | 2.77e-27 | 9.319999999999999e-23 |
| Malignant epithelial | C2 WNT ECs | SPRY4 | 1.344 | 0.157 | 0.063 | 6.55e-19 | 2.21e-14 |
| Malignant epithelial | C2 WNT ECs | JUN | 1.343 | 0.965 | 0.902 | 4.45e-76 | 1.5e-71 |
| Malignant epithelial | C2 WNT ECs | LSS | 1.343 | 0.15 | 0.051 | 1.23e-23 | 4.16e-19 |
| Malignant epithelial | C2 WNT ECs | FBXO31 | 1.342 | 0.173 | 0.06 | 2.37e-26 | 7.98e-22 |
| Malignant epithelial | C2 WNT ECs | PIBF1 | 1.341 | 0.18 | 0.059 | 6.9e-30 | 2.33e-25 |
| Malignant epithelial | C2 WNT ECs | RIPK1 | 1.341 | 0.111 | 0.038 | 1.06e-17 | 3.56e-13 |
| Malignant epithelial | C2 WNT ECs | OVOL1 | 1.339 | 0.543 | 0.288 | 3.04e-49 | 1.03e-44 |
| Malignant epithelial | C2 WNT ECs | ARL8B | 1.338 | 0.402 | 0.152 | 1.18e-56 | 3.959999999999999e-52 |
| Malignant epithelial | C2 WNT ECs | FLJ37453 | 1.338 | 0.157 | 0.052 | 2.51e-25 | 8.44e-21 |
| Malignant epithelial | C2 WNT ECs | DPH2 | 1.336 | 0.181 | 0.06 | 3.57e-29 | 1.2e-24 |
| Malignant epithelial | C2 WNT ECs | CASZ1 | 1.336 | 0.179 | 0.061 | 9.94e-28 | 3.35e-23 |
| Malignant epithelial | C2 WNT ECs | PNPLA2 | 1.333 | 0.667 | 0.289 | 5.47e-92 | 1.84e-87 |
| Malignant epithelial | C2 WNT ECs | ROCK1 | 1.333 | 0.422 | 0.16 | 3.64e-59 | 1.23e-54 |
| Malignant epithelial | C2 WNT ECs | RSBN1 | 1.333 | 0.334 | 0.126 | 5.49e-46 | 1.85e-41 |
| Malignant epithelial | C2 WNT ECs | CTBS | 1.333 | 0.222 | 0.074 | 1.34e-35 | 4.510000000000001e-31 |
| Malignant epithelial | C2 WNT ECs | TMEM43 | 1.332 | 0.329 | 0.12 | 1.67e-47 | 5.620000000000001e-43 |
| Malignant epithelial | C2 WNT ECs | MUM1 | 1.331 | 0.394 | 0.158 | 7.75e-51 | 2.61e-46 |
| Malignant epithelial | C2 WNT ECs | DCLRE1C | 1.331 | 0.189 | 0.061 | 2.79e-31 | 9.39e-27 |
| Malignant epithelial | C2 WNT ECs | ACTN4 | 1.33 | 0.934 | 0.72 | 3.73e-115 | 1.26e-110 |
| Malignant epithelial | C2 WNT ECs | ZFHX3 | 1.329 | 0.349 | 0.132 | 2.5e-48 | 8.41e-44 |
| Malignant epithelial | C2 WNT ECs | ADM | 1.329 | 0.702 | 0.48 | 3.09e-43 | 1.04e-38 |

| compartment | subcluster | gene | avg_log2FC | pct.1 | pct.2 | p_val | p_val_adj |
|---|---|---|---|---|---|---|---|
| Malignant epithelial | C2 WNT ECs | CATSPER1 | 1.328 | 0.184 | 0.065 | 5.790000000000001e-27 | 1.95e-22 |
| Malignant epithelial | C2 WNT ECs | FOLR1 | 1.327 | 0.158 | 0.044 | 1.43e-32 | 4.81e-28 |
| Malignant epithelial | C2 WNT ECs | TBX3 | 1.327 | 0.191 | 0.07 | 5.900000000000001e-27 | 1.99e-22 |
| Malignant epithelial | C2 WNT ECs | IGF2R | 1.325 | 0.261 | 0.094 | 3.43e-38 | 1.16e-33 |
| Malignant epithelial | C2 WNT ECs | RP11-783K16.5 | 1.325 | 0.264 | 0.101 | 8.09e-35 | 2.73e-30 |
| Malignant epithelial | C2 WNT ECs | RP3-512B11.3 | 1.325 | 0.175 | 0.056 | 1.08e-29 | 3.62e-25 |
| Malignant epithelial | C2 WNT ECs | FOXP4 | 1.324 | 0.302 | 0.103 | 1.56e-47 | 5.27e-43 |
| Malignant epithelial | C2 WNT ECs | TANC1 | 1.324 | 0.212 | 0.076 | 5.74e-31 | 1.93e-26 |
| Malignant epithelial | C2 WNT ECs | MFSD6 | 1.324 | 0.192 | 0.068 | 2.38e-28 | 8.029999999999999e-24 |
| Malignant epithelial | C2 WNT ECs | ZCCHC14 | 1.324 | 0.152 | 0.06 | 7.75e-19 | 2.61e-14 |
| Malignant epithelial | C2 WNT ECs | SERPINH1 | 1.323 | 0.651 | 0.416 | 9.44e-40 | 3.18e-35 |
| Malignant epithelial | C2 WNT ECs | GNA15 | 1.322 | 0.221 | 0.073 | 7.53e-36 | 2.54e-31 |
| Malignant epithelial | C2 WNT ECs | TMTC4 | 1.322 | 0.203 | 0.069 | 9.790000000000001e-32 | 3.3e-27 |
| Malignant epithelial | C2 WNT ECs | KDM4A | 1.322 | 0.126 | 0.04 | 8.71e-22 | 2.94e-17 |
| Malignant epithelial | C2 WNT ECs | CRNDE | 1.321 | 0.327 | 0.144 | 5.37e-35 | 1.81e-30 |
| Malignant epithelial | C2 WNT ECs | SLC43A2 | 1.321 | 0.12 | 0.038 | 6.620000000000001e-21 | 2.23e-16 |
| Malignant epithelial | C2 WNT ECs | MED4 | 1.319 | 0.662 | 0.299 | 3.97e-87 | 1.34e-82 |
| Malignant epithelial | C2 WNT ECs | AKAP11 | 1.316 | 0.2 | 0.061 | 9.03e-37 | 3.04e-32 |
| Malignant epithelial | C2 WNT ECs | GMFG | 1.315 | 0.271 | 0.126 | 1.26e-25 | 4.25e-21 |
| Malignant epithelial | C2 WNT ECs | ECHDC1 | 1.313 | 0.426 | 0.151 | 7.450000000000001e-66 | 2.51e-61 |
| Malignant epithelial | C2 WNT ECs | FAM89B | 1.313 | 0.456 | 0.188 | 7.35e-58 | 2.48e-53 |
| Malignant epithelial | C2 WNT ECs | MFI2 | 1.313 | 0.332 | 0.129 | 9.67e-44 | 3.26e-39 |
| Malignant epithelial | C2 WNT ECs | LEMD1 | 1.313 | 0.284 | 0.163 | 5.43e-17 | 1.83e-12 |
| Malignant epithelial | C2 WNT ECs | DVL1 | 1.311 | 0.294 | 0.111 | 1.51e-39 | 5.090000000000001e-35 |
| Malignant epithelial | C2 WNT ECs | PINK1 | 1.31 | 0.413 | 0.161 | 4.24e-54 | 1.43e-49 |
| Malignant epithelial | C2 WNT ECs | SMIM3 | 1.309 | 0.227 | 0.086 | 1.74e-29 | 5.85e-25 |
| Malignant epithelial | C2 WNT ECs | GEM | 1.308 | 0.294 | 0.134 | 1.04e-27 | 3.49e-23 |
| Malignant epithelial | C2 WNT ECs | KLHL23 | 1.308 | 0.194 | 0.074 | 4.97e-25 | 1.68e-20 |
| Malignant epithelial | C2 WNT ECs | PYROXD1 | 1.306 | 0.199 | 0.066 | 6.25e-32 | 2.11e-27 |
| Malignant epithelial | C2 WNT ECs | PDS5B | 1.305 | 0.267 | 0.095 | 7.78e-39 | 2.62e-34 |
| Malignant epithelial | C2 WNT ECs | SPIRE2 | 1.303 | 0.336 | 0.132 | 8.170000000000001e-43 | 2.75e-38 |
| Malignant epithelial | C2 WNT ECs | FAM118A | 1.301 | 0.479 | 0.247 | 8.12e-43 | 2.74e-38 |
| Malignant epithelial | C2 WNT ECs | CDHR2 | 1.299 | 0.184 | 0.059 | 5.71e-31 | 1.92e-26 |
| Malignant epithelial | C2 WNT ECs | HDAC11 | 1.299 | 0.158 | 0.05 | 2.12e-27 | 7.149999999999999e-23 |
| Malignant epithelial | C2 WNT ECs | RHBDF1 | 1.297 | 0.361 | 0.142 | 9.71e-47 | 3.27e-42 |
| Malignant epithelial | C2 WNT ECs | GOLIM4 | 1.297 | 0.415 | 0.183 | 1.19e-45 | 4e-41 |
| Malignant epithelial | C2 WNT ECs | PPM1D | 1.297 | 0.135 | 0.051 | 7.05e-18 | 2.38e-13 |
| Malignant epithelial | C2 WNT ECs | EIF1AD | 1.296 | 0.306 | 0.117 | 5.66e-41 | 1.91e-36 |
| Malignant epithelial | C2 WNT ECs | TCP1 | 1.295 | 0.773 | 0.428 | 1.47e-88 | 4.959999999999999e-84 |
| Malignant epithelial | C2 WNT ECs | AOC1 | 1.295 | 0.629 | 0.283 | 1.02e-75 | 3.43e-71 |
| Malignant epithelial | C2 WNT ECs | MAFF | 1.295 | 0.696 | 0.449 | 1.53e-52 | 5.149999999999999e-48 |
| Malignant epithelial | C2 WNT ECs | SLC52A2 | 1.294 | 0.793 | 0.439 | 6.04e-100 | 2.04e-95 |
| Malignant epithelial | C2 WNT ECs | BTN2A1 | 1.294 | 0.158 | 0.054 | 1.37e-24 | 4.62e-20 |
| Malignant epithelial | C2 WNT ECs | MARCKSL1 | 1.293 | 0.942 | 0.695 | 5.05e-136 | 1.7e-131 |
| Malignant epithelial | C2 WNT ECs | KCTD5 | 1.293 | 0.383 | 0.173 | 4.21e-39 | 1.42e-34 |
| Malignant epithelial | C2 WNT ECs | HENMT1 | 1.293 | 0.161 | 0.063 | 5.04e-20 | 1.7e-15 |
| Malignant epithelial | C2 WNT ECs | LRCH4 | 1.292 | 0.367 | 0.144 | 2.79e-47 | 9.399999999999999e-43 |
| Malignant epithelial | C2 WNT ECs | MID1 | 1.29 | 0.108 | 0.047 | 6.89e-12 | 2.32e-07 |
| Malignant epithelial | C2 WNT ECs | EHBP1L1 | 1.288 | 0.271 | 0.103 | 1.92e-35 | 6.46e-31 |
| Malignant epithelial | C2 WNT ECs | WDR11 | 1.288 | 0.299 | 0.125 | 1.88e-34 | 6.32e-30 |
| Malignant epithelial | C2 WNT ECs | HID1 | 1.288 | 0.165 | 0.054 | 2.39e-27 | 8.049999999999999e-23 |
| Malignant epithelial | C2 WNT ECs | ADAM8 | 1.287 | 0.171 | 0.059 | 7.35e-26 | 2.48e-21 |
| Malignant epithelial | C2 WNT ECs | MRPL18 | 1.286 | 0.774 | 0.516 | 3.66e-52 | 1.23e-47 |
| Malignant epithelial | C2 WNT ECs | TGDS | 1.286 | 0.198 | 0.069 | 1.99e-29 | 6.69e-25 |
| Malignant epithelial | C2 WNT ECs | KRT18 | 1.285 | 1.0 | 0.991 | 2.5e-69 | 8.43e-65 |
| Malignant epithelial | C2 WNT ECs | RIOK3 | 1.284 | 0.514 | 0.215 | 3.49e-64 | 1.18e-59 |
| Malignant epithelial | C2 WNT ECs | SLC30A1 | 1.284 | 0.288 | 0.112 | 9.64e-37 | 3.25e-32 |
| Malignant epithelial | C2 WNT ECs | MYZAP | 1.282 | 0.122 | 0.04 | 6.34e-20 | 2.14e-15 |
| Malignant epithelial | C2 WNT ECs | UGDH | 1.28 | 0.425 | 0.165 | 5.9e-57 | 1.99e-52 |
| Malignant epithelial | C2 WNT ECs | OPLAH | 1.28 | 0.137 | 0.05 | 3.34e-19 | 1.13e-14 |
| Malignant epithelial | C2 WNT ECs | GSE1 | 1.279 | 0.359 | 0.137 | 2.96e-48 | 9.96e-44 |
| Malignant epithelial | C2 WNT ECs | FAM160B1 | 1.279 | 0.107 | 0.043 | 8.55e-13 | 2.88e-08 |
| Malignant epithelial | C2 WNT ECs | SEL1L3 | 1.277 | 0.246 | 0.086 | 1e-36 | 3.37e-32 |
| Malignant epithelial | C2 WNT ECs | WDTC1 | 1.277 | 0.173 | 0.052 | 4.67e-32 | 1.57e-27 |

| compartment | subcluster | gene | avg_log2FC | pct.1 | pct.2 | p_val | p_val_adj |
|---|---|---|---|---|---|---|---|
| Malignant epithelial | C2 WNT ECs | ZNF326 | 1.273 | 0.375 | 0.149 | 2.33e-48 | 7.86e-44 |
| Malignant epithelial | C2 WNT ECs | DHX32 | 1.273 | 0.198 | 0.08 | 1.87e-23 | 6.31e-19 |
| Malignant epithelial | C2 WNT ECs | UBA6 | 1.272 | 0.35 | 0.146 | 1.44e-40 | 4.85e-36 |
| Malignant epithelial | C2 WNT ECs | SLC35B3 | 1.272 | 0.242 | 0.085 | 1.67e-35 | 5.64e-31 |
| Malignant epithelial | C2 WNT ECs | CSDE1 | 1.271 | 0.756 | 0.407 | 2.23e-90 | 7.51e-86 |
| Malignant epithelial | C2 WNT ECs | PHGR1 | 1.269 | 0.909 | 0.505 | 8.18e-114 | 2.75e-109 |
| Malignant epithelial | C2 WNT ECs | RTKN | 1.268 | 0.501 | 0.212 | 6.67e-61 | 2.25e-56 |
| Malignant epithelial | C2 WNT ECs | PTOV1 | 1.267 | 0.637 | 0.272 | 2.79e-88 | 9.409999999999999e-84 |
| Malignant epithelial | C2 WNT ECs | PHF23 | 1.267 | 0.227 | 0.092 | 4.490000000000001e-27 | 1.51e-22 |
| Malignant epithelial | C2 WNT ECs | TMEM189 | 1.266 | 0.499 | 0.217 | 3.36e-58 | 1.13e-53 |
| Malignant epithelial | C2 WNT ECs | ATP6V0B | 1.265 | 0.851 | 0.504 | 1.37e-113 | 4.63e-109 |
| Malignant epithelial | C2 WNT ECs | ABCG1 | 1.265 | 0.223 | 0.084 | 4.01e-30 | 1.35e-25 |
| Malignant epithelial | C2 WNT ECs | TM9SF2 | 1.264 | 0.774 | 0.421 | 1.1e-90 | 3.72e-86 |
| Malignant epithelial | C2 WNT ECs | LRIF1 | 1.264 | 0.269 | 0.109 | 2.85e-32 | 9.59e-28 |
| Malignant epithelial | C2 WNT ECs | HSDL1 | 1.264 | 0.139 | 0.058 | 1.58e-15 | 5.33e-11 |
| Malignant epithelial | C2 WNT ECs | RELT | 1.264 | 0.106 | 0.045 | 1.95e-11 | 6.56e-07 |
| Malignant epithelial | C2 WNT ECs | ARHGEF2 | 1.263 | 0.265 | 0.105 | 6.38e-33 | 2.15e-28 |
| Malignant epithelial | C2 WNT ECs | SGK1 | 1.262 | 0.452 | 0.196 | 7.070000000000001e-51 | 2.38e-46 |
| Malignant epithelial | C2 WNT ECs | KIAA0355 | 1.262 | 0.23 | 0.083 | 6.11e-33 | 2.06e-28 |
| Malignant epithelial | C2 WNT ECs | PWWP2B | 1.262 | 0.229 | 0.091 | 1.42e-27 | 4.8e-23 |
| Malignant epithelial | C2 WNT ECs | GRAMD1A | 1.261 | 0.536 | 0.222 | 4.559999999999999e-68 | 1.54e-63 |
| Malignant epithelial | C2 WNT ECs | PQLC2 | 1.26 | 0.107 | 0.038 | 6.34e-16 | 2.14e-11 |
| Malignant epithelial | C2 WNT ECs | SMIM14 | 1.259 | 0.516 | 0.209 | 2.99e-65 | 1.01e-60 |
| Malignant epithelial | C2 WNT ECs | CPEB4 | 1.259 | 0.211 | 0.08 | 2.84e-28 | 9.56e-24 |
| Malignant epithelial | C2 WNT ECs | AVEN | 1.259 | 0.139 | 0.052 | 1.46e-18 | 4.93e-14 |
| Malignant epithelial | C2 WNT ECs | SSR1 | 1.258 | 0.541 | 0.232 | 4.779999999999999e-67 | 1.61e-62 |
| Malignant epithelial | C2 WNT ECs | THUMPD3-AS1 | 1.258 | 0.559 | 0.261 | 2.25e-60 | 7.599999999999999e-56 |
| Malignant epithelial | C2 WNT ECs | GCC2 | 1.256 | 0.69 | 0.338 | 7.3e-79 | 2.46e-74 |
| Malignant epithelial | C2 WNT ECs | TOX3 | 1.255 | 0.333 | 0.125 | 1.73e-45 | 5.82e-41 |
| Malignant epithelial | C2 WNT ECs | SREBF1 | 1.255 | 0.299 | 0.111 | 7.32e-41 | 2.47e-36 |
| Malignant epithelial | C2 WNT ECs | ADD1 | 1.255 | 0.344 | 0.147 | 2.37e-38 | 7.99e-34 |
| Malignant epithelial | C2 WNT ECs | CSNK1E | 1.254 | 0.536 | 0.27 | 4.14e-47 | 1.39e-42 |
| Malignant epithelial | C2 WNT ECs | TRPM2 | 1.253 | 0.185 | 0.076 | 2.7e-21 | 9.1e-17 |
| Malignant epithelial | C2 WNT ECs | CEP41 | 1.253 | 0.103 | 0.041 | 8.09e-13 | 2.72e-08 |
| Malignant epithelial | C2 WNT ECs | DGKA | 1.252 | 0.101 | 0.045 | 2.14e-10 | 7.21e-06 |
| Malignant epithelial | C2 WNT ECs | UCHL3 | 1.251 | 0.689 | 0.358 | 1e-77 | 3.370000000000001e-73 |
| Malignant epithelial | C2 WNT ECs | HS2ST1 | 1.251 | 0.185 | 0.063 | 1.67e-28 | 5.61e-24 |
| Malignant epithelial | C2 WNT ECs | PRDM4 | 1.25 | 0.147 | 0.054 | 1.59e-20 | 5.36e-16 |
| Malignant epithelial | C2 WNT ECs | FAM214B | 1.249 | 0.146 | 0.054 | 4.24e-20 | 1.43e-15 |
| Malignant epithelial | C2 WNT ECs | RMND5A | 1.248 | 0.436 | 0.197 | 1.21e-44 | 4.07e-40 |
| Malignant epithelial | C2 WNT ECs | SFN | 1.245 | 0.882 | 0.678 | 1.62e-63 | 5.460000000000001e-59 |
| Malignant epithelial | C2 WNT ECs | PIGU | 1.245 | 0.314 | 0.131 | 2.52e-36 | 8.5e-32 |
| Malignant epithelial | C2 WNT ECs | TMEM79 | 1.245 | 0.162 | 0.05 | 6.72e-29 | 2.27e-24 |
| Malignant epithelial | C2 WNT ECs | CDKAL1 | 1.245 | 0.107 | 0.028 | 9.73e-24 | 3.28e-19 |
| Malignant epithelial | C2 WNT ECs | TRAK2 | 1.245 | 0.134 | 0.049 | 6.18e-19 | 2.08e-14 |
| Malignant epithelial | C2 WNT ECs | MRPS31 | 1.244 | 0.387 | 0.164 | 2.78e-43 | 9.38e-39 |
| Malignant epithelial | C2 WNT ECs | WIPI1 | 1.244 | 0.211 | 0.078 | 3.67e-28 | 1.24e-23 |
| Malignant epithelial | C2 WNT ECs | POMP | 1.243 | 0.968 | 0.831 | 9.15e-151 | 3.08e-146 |
| Malignant epithelial | C2 WNT ECs | MSMO1 | 1.242 | 0.348 | 0.157 | 2.12e-34 | 7.16e-30 |
| Malignant epithelial | C2 WNT ECs | NOP14-AS1 | 1.242 | 0.118 | 0.046 | 9.73e-15 | 3.28e-10 |
| Malignant epithelial | C2 WNT ECs | VOPP1 | 1.24 | 0.594 | 0.305 | 3.63e-54 | 1.22e-49 |
| Malignant epithelial | C2 WNT ECs | DUSP10 | 1.24 | 0.237 | 0.127 | 4.72e-15 | 1.59e-10 |
| Malignant epithelial | C2 WNT ECs | ARMT1 | 1.238 | 0.225 | 0.084 | 3.3e-30 | 1.11e-25 |
| Malignant epithelial | C2 WNT ECs | GUCY2C | 1.237 | 0.116 | 0.041 | 1.67e-17 | 5.64e-13 |
| Malignant epithelial | C2 WNT ECs | FAM110C | 1.236 | 0.189 | 0.073 | 7.039999999999999e-24 | 2.37e-19 |
| Malignant epithelial | C2 WNT ECs | EFNA4 | 1.235 | 0.39 | 0.171 | 1.37e-41 | 4.62e-37 |
| Malignant epithelial | C2 WNT ECs | TTC9 | 1.235 | 0.116 | 0.049 | 6.02e-13 | 2.03e-08 |
| Malignant epithelial | C2 WNT ECs | PICK1 | 1.234 | 0.229 | 0.086 | 7.799999999999999e-31 | 2.63e-26 |
| Malignant epithelial | C2 WNT ECs | STRIP1 | 1.234 | 0.134 | 0.042 | 5.6e-24 | 1.89e-19 |
| Malignant epithelial | C2 WNT ECs | AMOTL2 | 1.233 | 0.226 | 0.096 | 1.98e-24 | 6.67e-20 |
| Malignant epithelial | C2 WNT ECs | MCUR1 | 1.232 | 0.49 | 0.205 | 6.2e-62 | 2.09e-57 |
| Malignant epithelial | C2 WNT ECs | HSPA1B | 1.232 | 0.931 | 0.819 | 3.19e-57 | 1.08e-52 |
| Malignant epithelial | C2 WNT ECs | GADD45A | 1.231 | 0.714 | 0.445 | 7.529999999999999e-55 | 2.54e-50 |
| Malignant epithelial | C2 WNT ECs | DNAJC12 | 1.231 | 0.11 | 0.052 | 5.28e-10 | 1.78e-05 |

| compartment | subcluster | gene | avg_log2FC | pct.1 | pct.2 | p_val | p_val_adj |
|---|---|---|---|---|---|---|---|
| Malignant epithelial | C2 WNT ECs | ATP13A2 | 1.23 | 0.175 | 0.061 | 1.34e-25 | 4.5e-21 |
| Malignant epithelial | C2 WNT ECs | LRRC41 | 1.229 | 0.28 | 0.097 | 1.94e-42 | 6.54e-38 |
| Malignant epithelial | C2 WNT ECs | RP1-39G22.7 | 1.229 | 0.118 | 0.048 | 1.65e-13 | 5.56e-09 |
| Malignant epithelial | C2 WNT ECs | ECI2 | 1.228 | 0.295 | 0.102 | 1.1e-44 | 3.71e-40 |
| Malignant epithelial | C2 WNT ECs | CDC16 | 1.227 | 0.405 | 0.166 | 2.04e-49 | 6.859999999999999e-45 |
| Malignant epithelial | C2 WNT ECs | SQLE | 1.226 | 0.415 | 0.223 | 4.010000000000001e-31 | 1.35e-26 |
| Malignant epithelial | C2 WNT ECs | HSPA8 | 1.225 | 0.969 | 0.811 | 6.76e-104 | 2.28e-99 |
| Malignant epithelial | C2 WNT ECs | SLC35E4 | 1.225 | 0.279 | 0.108 | 9.64e-36 | 3.25e-31 |
| Malignant epithelial | C2 WNT ECs | GSTM4 | 1.225 | 0.108 | 0.035 | 2.34e-18 | 7.88e-14 |
| Malignant epithelial | C2 WNT ECs | SLC2A3 | 1.225 | 0.187 | 0.09 | 3.18e-15 | 1.07e-10 |
| Malignant epithelial | C2 WNT ECs | TTC39A | 1.224 | 0.165 | 0.064 | 7.150000000000001e-21 | 2.41e-16 |
| Malignant epithelial | C2 WNT ECs | DHCR7 | 1.224 | 0.229 | 0.106 | 8.050000000000001e-21 | 2.71e-16 |
| Malignant epithelial | C2 WNT ECs | SOWAHC | 1.223 | 0.472 | 0.199 | 9.520000000000001e-56 | 3.21e-51 |
| Malignant epithelial | C2 WNT ECs | MGAT4A | 1.223 | 0.188 | 0.073 | 1.43e-23 | 4.8e-19 |
| Malignant epithelial | C2 WNT ECs | TMEM57 | 1.222 | 0.234 | 0.092 | 6.97e-29 | 2.35e-24 |
| Malignant epithelial | C2 WNT ECs | PHACTR2 | 1.221 | 0.349 | 0.175 | 1.8e-27 | 6.05e-23 |
| Malignant epithelial | C2 WNT ECs | FLCN | 1.218 | 0.139 | 0.056 | 1.37e-16 | 4.61e-12 |
| Malignant epithelial | C2 WNT ECs | RP1-122P22.4 | 1.217 | 0.273 | 0.1 | 4.66e-39 | 1.57e-34 |
| Malignant epithelial | C2 WNT ECs | RBM26 | 1.216 | 0.39 | 0.155 | 1.55e-49 | 5.23e-45 |
| Malignant epithelial | C2 WNT ECs | NRAS | 1.216 | 0.296 | 0.108 | 4.82e-41 | 1.63e-36 |
| Malignant epithelial | C2 WNT ECs | C1orf52 | 1.216 | 0.349 | 0.143 | 1.67e-40 | 5.63e-36 |
| Malignant epithelial | C2 WNT ECs | ACAA2 | 1.215 | 0.559 | 0.276 | 2.44e-56 | 8.22e-52 |
| Malignant epithelial | C2 WNT ECs | CRK | 1.214 | 0.401 | 0.169 | 1.11e-46 | 3.72e-42 |
| Malignant epithelial | C2 WNT ECs | AC013461.1 | 1.214 | 0.493 | 0.248 | 5.74e-44 | 1.93e-39 |
| Malignant epithelial | C2 WNT ECs | RAB5B | 1.214 | 0.33 | 0.17 | 5.699999999999999e-24 | 1.92e-19 |
| Malignant epithelial | C2 WNT ECs | ZNF720 | 1.213 | 0.237 | 0.093 | 4.39e-29 | 1.48e-24 |
| Malignant epithelial | C2 WNT ECs | GCC1 | 1.213 | 0.137 | 0.052 | 9.56e-18 | 3.22e-13 |
| Malignant epithelial | C2 WNT ECs | DCUN1D3 | 1.212 | 0.134 | 0.046 | 2.12e-20 | 7.16e-16 |
| Malignant epithelial | C2 WNT ECs | POLM | 1.211 | 0.256 | 0.091 | 6.610000000000001e-37 | 2.23e-32 |
| Malignant epithelial | C2 WNT ECs | PPP4R1 | 1.211 | 0.212 | 0.075 | 1.37e-31 | 4.62e-27 |
| Malignant epithelial | C2 WNT ECs | ENGASE | 1.211 | 0.218 | 0.084 | 2.72e-27 | 9.15e-23 |
| Malignant epithelial | C2 WNT ECs | PI4K2A | 1.211 | 0.11 | 0.04 | 1.1e-15 | 3.69e-11 |
| Malignant epithelial | C2 WNT ECs | ZSWIM7 | 1.21 | 0.26 | 0.102 | 2.62e-32 | 8.82e-28 |
| Malignant epithelial | C2 WNT ECs | BMI1 | 1.209 | 0.208 | 0.077 | 2.92e-28 | 9.84e-24 |
| Malignant epithelial | C2 WNT ECs | TNK2 | 1.207 | 0.322 | 0.138 | 1.14e-35 | 3.84e-31 |
| Malignant epithelial | C2 WNT ECs | MBNL2 | 1.206 | 0.398 | 0.172 | 3.62e-43 | 1.22e-38 |
| Malignant epithelial | C2 WNT ECs | USP12 | 1.205 | 0.286 | 0.129 | 2.77e-27 | 9.339999999999999e-23 |
| Malignant epithelial | C2 WNT ECs | NHLRC3 | 1.204 | 0.382 | 0.153 | 1.76e-47 | 5.93e-43 |
| Malignant epithelial | C2 WNT ECs | TM7SF2 | 1.204 | 0.383 | 0.154 | 1.13e-46 | 3.82e-42 |
| Malignant epithelial | C2 WNT ECs | ZNF322 | 1.204 | 0.166 | 0.058 | 1.23e-24 | 4.15e-20 |
| Malignant epithelial | C2 WNT ECs | HBS1L | 1.203 | 0.258 | 0.098 | 1.24e-33 | 4.17e-29 |
| Malignant epithelial | C2 WNT ECs | STPG1 | 1.203 | 0.13 | 0.049 | 3.89e-17 | 1.31e-12 |
| Malignant epithelial | C2 WNT ECs | C1orf116 | 1.202 | 0.194 | 0.079 | 3.14e-22 | 1.06e-17 |
| Malignant epithelial | C2 WNT ECs | HMGCS1 | 1.201 | 0.495 | 0.331 | 1.56e-20 | 5.25e-16 |
| Malignant epithelial | C2 WNT ECs | ACSL1 | 1.201 | 0.103 | 0.035 | 3.51e-16 | 1.18e-11 |
| Malignant epithelial | C2 WNT ECs | MYSM1 | 1.2 | 0.145 | 0.053 | 4.69e-20 | 1.58e-15 |
| Malignant epithelial | C2 WNT ECs | NOMO1 | 1.199 | 0.143 | 0.057 | 1.45e-17 | 4.89e-13 |
| Malignant epithelial | C2 WNT ECs | NAB1 | 1.197 | 0.365 | 0.158 | 4.18e-39 | 1.41e-34 |
| Malignant epithelial | C2 WNT ECs | CYP4F11 | 1.196 | 0.104 | 0.037 | 2.69e-15 | 9.05e-11 |
| Malignant epithelial | C2 WNT ECs | TFEB | 1.193 | 0.181 | 0.067 | 5.33e-25 | 1.79e-20 |
| Malignant epithelial | C2 WNT ECs | NRP2 | 1.193 | 0.268 | 0.132 | 8.94e-22 | 3.01e-17 |
| Malignant epithelial | C2 WNT ECs | DUSP18 | 1.19 | 0.277 | 0.103 | 4.73e-37 | 1.59e-32 |
| Malignant epithelial | C2 WNT ECs | LYPLA2 | 1.188 | 0.559 | 0.25 | 2.56e-66 | 8.62e-62 |
| Malignant epithelial | C2 WNT ECs | TIMM23 | 1.188 | 0.314 | 0.117 | 7.459999999999999e-42 | 2.51e-37 |
| Malignant epithelial | C2 WNT ECs | TRAF5 | 1.187 | 0.333 | 0.128 | 3.97e-43 | 1.34e-38 |
| Malignant epithelial | C2 WNT ECs | TBC1D17 | 1.187 | 0.292 | 0.117 | 1.93e-35 | 6.49e-31 |
| Malignant epithelial | C2 WNT ECs | FAM210A | 1.187 | 0.183 | 0.078 | 3.6e-19 | 1.21e-14 |
| Malignant epithelial | C2 WNT ECs | GCLM | 1.187 | 0.134 | 0.05 | 6.44e-18 | 2.17e-13 |
| Malignant epithelial | C2 WNT ECs | SPTY2D1 | 1.186 | 0.391 | 0.188 | 8.440000000000001e-35 | 2.84e-30 |
| Malignant epithelial | C2 WNT ECs | SEPT10 | 1.185 | 0.418 | 0.18 | 3.88e-47 | 1.31e-42 |
| Malignant epithelial | C2 WNT ECs | SCARB2 | 1.184 | 0.449 | 0.184 | 9.78e-56 | 3.3e-51 |
| Malignant epithelial | C2 WNT ECs | RNASEH2B | 1.184 | 0.383 | 0.166 | 5.31e-42 | 1.79e-37 |
| Malignant epithelial | C2 WNT ECs | SPRYD7 | 1.184 | 0.329 | 0.136 | 3.98e-38 | 1.34e-33 |
| Malignant epithelial | C2 WNT ECs | PDK3 | 1.184 | 0.157 | 0.057 | 1.75e-21 | 5.91e-17 |

| compartment | subcluster | gene | avg_log2FC | pct.1 | pct.2 | p_val | p_val_adj |
|---|---|---|---|---|---|---|---|
| Malignant epithelial | C2 WNT ECs | CPT2 | 1.183 | 0.104 | 0.039 | 2.7e-14 | 9.11e-10 |
| Malignant epithelial | C2 WNT ECs | SEPP1 | 1.182 | 0.678 | 0.32 | 2.24e-80 | 7.56e-76 |
| Malignant epithelial | C2 WNT ECs | DRAM2 | 1.181 | 0.464 | 0.204 | 4.33e-52 | 1.46e-47 |
| Malignant epithelial | C2 WNT ECs | PLK3 | 1.181 | 0.587 | 0.33 | 1.59e-43 | 5.35e-39 |
| Malignant epithelial | C2 WNT ECs | HIBCH | 1.18 | 0.357 | 0.151 | 2.19e-40 | 7.39e-36 |
| Malignant epithelial | C2 WNT ECs | PDGFA | 1.179 | 0.356 | 0.163 | 4.66e-34 | 1.57e-29 |
| Malignant epithelial | C2 WNT ECs | PTER | 1.179 | 0.142 | 0.057 | 4.02e-17 | 1.36e-12 |
| Malignant epithelial | C2 WNT ECs | SPSB3 | 1.178 | 0.66 | 0.32 | 5.62e-71 | 1.89e-66 |
| Malignant epithelial | C2 WNT ECs | MAPK13 | 1.178 | 0.641 | 0.339 | 4.07e-65 | 1.37e-60 |
| Malignant epithelial | C2 WNT ECs | AAK1 | 1.175 | 0.25 | 0.1 | 7.23e-30 | 2.44e-25 |
| Malignant epithelial | C2 WNT ECs | HNF4A | 1.174 | 0.482 | 0.197 | 3.47e-59 | 1.17e-54 |
| Malignant epithelial | C2 WNT ECs | INTS4 | 1.174 | 0.147 | 0.052 | 1e-21 | 3.37e-17 |
| Malignant epithelial | C2 WNT ECs | SPINK1 | 1.174 | 0.378 | 0.256 | 3.65e-15 | 1.23e-10 |
| Malignant epithelial | C2 WNT ECs | IYD | 1.173 | 0.126 | 0.04 | 5.700000000000001e-22 | 1.92e-17 |
| Malignant epithelial | C2 WNT ECs | VMAC | 1.171 | 0.108 | 0.039 | 2.68e-15 | 9.04e-11 |
| Malignant epithelial | C2 WNT ECs | YTHDF2 | 1.169 | 0.543 | 0.254 | 2.7e-55 | 9.11e-51 |
| Malignant epithelial | C2 WNT ECs | EXOC1 | 1.169 | 0.302 | 0.107 | 1.41e-44 | 4.74e-40 |
| Malignant epithelial | C2 WNT ECs | TOP2B | 1.169 | 0.294 | 0.108 | 5.17e-40 | 1.74e-35 |
| Malignant epithelial | C2 WNT ECs | CTD-2377D24.6 | 1.169 | 0.204 | 0.066 | 5.74e-34 | 1.94e-29 |
| Malignant epithelial | C2 WNT ECs | MTF2 | 1.168 | 0.318 | 0.128 | 1.58e-37 | 5.31e-33 |
| Malignant epithelial | C2 WNT ECs | PPP1R11 | 1.167 | 0.599 | 0.276 | 6.04e-68 | 2.04e-63 |
| Malignant epithelial | C2 WNT ECs | NIPBL | 1.167 | 0.425 | 0.187 | 4.31e-46 | 1.45e-41 |
| Malignant epithelial | C2 WNT ECs | THOP1 | 1.167 | 0.29 | 0.123 | 1.22e-31 | 4.1e-27 |
| Malignant epithelial | C2 WNT ECs | PARP4 | 1.165 | 0.307 | 0.126 | 9.45e-36 | 3.18e-31 |
| Malignant epithelial | C2 WNT ECs | HOOK3 | 1.164 | 0.409 | 0.169 | 1.45e-48 | 4.88e-44 |
| Malignant epithelial | C2 WNT ECs | ZNF292 | 1.163 | 0.475 | 0.232 | 3.69e-43 | 1.24e-38 |
| Malignant epithelial | C2 WNT ECs | JARID2 | 1.163 | 0.184 | 0.069 | 4.339999999999999e-24 | 1.46e-19 |
| Malignant epithelial | C2 WNT ECs | MOGAT3 | 1.163 | 0.198 | 0.088 | 3.85e-19 | 1.3e-14 |
| Malignant epithelial | C2 WNT ECs | SAP18 | 1.162 | 0.988 | 0.861 | 3.37e-158 | 1.14e-153 |
| Malignant epithelial | C2 WNT ECs | ATP9A | 1.162 | 0.345 | 0.13 | 1.87e-46 | 6.3e-42 |
| Malignant epithelial | C2 WNT ECs | TEAD3 | 1.162 | 0.267 | 0.105 | 8.500000000000001e-33 | 2.86e-28 |
| Malignant epithelial | C2 WNT ECs | SLC22A5 | 1.162 | 0.131 | 0.041 | 1.15e-22 | 3.87e-18 |
| Malignant epithelial | C2 WNT ECs | B4GALNT3 | 1.159 | 0.208 | 0.105 | 6.37e-16 | 2.15e-11 |
| Malignant epithelial | C2 WNT ECs | BORCS6 | 1.159 | 0.111 | 0.047 | 4.34e-12 | 1.46e-07 |
| Malignant epithelial | C2 WNT ECs | IDH1 | 1.158 | 0.505 | 0.213 | 6.059999999999999e-61 | 2.04e-56 |
| Malignant epithelial | C2 WNT ECs | ZNF622 | 1.158 | 0.487 | 0.234 | 2.58e-46 | 8.709999999999999e-42 |
| Malignant epithelial | C2 WNT ECs | ACAT2 | 1.158 | 0.253 | 0.129 | 1.31e-18 | 4.4e-14 |
| Malignant epithelial | C2 WNT ECs | RP11-10C24.3 | 1.157 | 0.103 | 0.038 | 2.75e-14 | 9.28e-10 |
| Malignant epithelial | C2 WNT ECs | KIFAP3 | 1.156 | 0.222 | 0.088 | 6.330000000000001e-27 | 2.13e-22 |
| Malignant epithelial | C2 WNT ECs | TMEM191B | 1.155 | 0.47 | 0.244 | 3.83e-40 | 1.29e-35 |
| Malignant epithelial | C2 WNT ECs | TNFRSF12A | 1.154 | 0.84 | 0.599 | 3.15e-61 | 1.06e-56 |
| Malignant epithelial | C2 WNT ECs | TH | 1.154 | 0.177 | 0.053 | 1.79e-32 | 6.019999999999999e-28 |
| Malignant epithelial | C2 WNT ECs | NKD2 | 1.152 | 0.192 | 0.066 | 3.13e-29 | 1.05e-24 |
| Malignant epithelial | C2 WNT ECs | ITM2B | 1.149 | 0.955 | 0.752 | 1.9e-120 | 6.4e-116 |
| Malignant epithelial | C2 WNT ECs | ZBTB7B | 1.148 | 0.233 | 0.105 | 1.56e-22 | 5.25e-18 |
| Malignant epithelial | C2 WNT ECs | CLTB | 1.147 | 0.827 | 0.689 | 3.84e-38 | 1.29e-33 |
| Malignant epithelial | C2 WNT ECs | MFSD14A | 1.147 | 0.189 | 0.075 | 3.72e-23 | 1.25e-18 |
| Malignant epithelial | C2 WNT ECs | RBMS1 | 1.146 | 0.376 | 0.158 | 7.6e-43 | 2.56e-38 |
| Malignant epithelial | C2 WNT ECs | UBE2J1 | 1.145 | 0.555 | 0.246 | 1.93e-63 | 6.520000000000001e-59 |
| Malignant epithelial | C2 WNT ECs | MAK16 | 1.145 | 0.129 | 0.045 | 7.27e-19 | 2.45e-14 |
| Malignant epithelial | C2 WNT ECs | RP5-1085F17.3 | 1.145 | 0.138 | 0.062 | 2.23e-13 | 7.51e-09 |
| Malignant epithelial | C2 WNT ECs | SERINC3 | 1.144 | 0.701 | 0.39 | 1.13e-65 | 3.789999999999999e-61 |
| Malignant epithelial | C2 WNT ECs | HBP1 | 1.144 | 0.474 | 0.214 | 5.12e-49 | 1.72e-44 |
| Malignant epithelial | C2 WNT ECs | SKAP2 | 1.143 | 0.518 | 0.245 | 5.299999999999999e-53 | 1.79e-48 |
| Malignant epithelial | C2 WNT ECs | GPBP1L1 | 1.143 | 0.425 | 0.185 | 4.6e-46 | 1.55e-41 |
| Malignant epithelial | C2 WNT ECs | IGSF8 | 1.143 | 0.313 | 0.129 | 2.08e-35 | 6.99e-31 |
| Malignant epithelial | C2 WNT ECs | SMIM2-AS1 | 1.141 | 0.107 | 0.034 | 2.61e-18 | 8.81e-14 |
| Malignant epithelial | C2 WNT ECs | UPF3A | 1.138 | 0.71 | 0.369 | 9.510000000000001e-79 | 3.2e-74 |
| Malignant epithelial | C2 WNT ECs | TMEM39A | 1.138 | 0.2 | 0.084 | 9.9e-22 | 3.33e-17 |
| Malignant epithelial | C2 WNT ECs | RHOB | 1.137 | 0.958 | 0.832 | 1.4e-81 | 4.71e-77 |
| Malignant epithelial | C2 WNT ECs | KLHL42 | 1.137 | 0.264 | 0.099 | 3.48e-35 | 1.17e-30 |
| Malignant epithelial | C2 WNT ECs | BCL11A | 1.135 | 0.154 | 0.061 | 2.57e-18 | 8.64e-14 |
| Malignant epithelial | C2 WNT ECs | NFIA | 1.135 | 0.18 | 0.08 | 1.96e-17 | 6.62e-13 |
| Malignant epithelial | C2 WNT ECs | MAD2L1BP | 1.134 | 0.286 | 0.127 | 1.66e-28 | 5.59e-24 |

| compartment | subcluster | gene | avg_log2FC | pct.1 | pct.2 | p_val | p_val_adj |
|---|---|---|---|---|---|---|---|
| Malignant epithelial | C2 WNT ECs | UFL1 | 1.133 | 0.405 | 0.169 | 1.21e-45 | 4.08e-41 |
| Malignant epithelial | C2 WNT ECs | PITPNA | 1.131 | 0.18 | 0.067 | 6.26e-24 | 2.11e-19 |
| Malignant epithelial | C2 WNT ECs | AMMECR1L | 1.13 | 0.161 | 0.061 | 5.5e-21 | 1.85e-16 |
| Malignant epithelial | C2 WNT ECs | DVL3 | 1.129 | 0.271 | 0.104 | 9.44e-35 | 3.18e-30 |
| Malignant epithelial | C2 WNT ECs | FBXO44 | 1.129 | 0.135 | 0.045 | 9.85e-22 | 3.32e-17 |
| Malignant epithelial | C2 WNT ECs | PRDM2 | 1.129 | 0.206 | 0.088 | 1.39e-21 | 4.69e-17 |
| Malignant epithelial | C2 WNT ECs | FXR2 | 1.129 | 0.104 | 0.04 | 2.45e-13 | 8.24e-09 |
| Malignant epithelial | C2 WNT ECs | ZMPSTE24 | 1.126 | 0.273 | 0.11 | 2.77e-32 | 9.339999999999999e-28 |
| Malignant epithelial | C2 WNT ECs | MFSD10 | 1.125 | 0.683 | 0.369 | 1.06e-66 | 3.57e-62 |
| Malignant epithelial | C2 WNT ECs | OGDH | 1.125 | 0.319 | 0.135 | 1.76e-34 | 5.93e-30 |
| Malignant epithelial | C2 WNT ECs | KATNA1 | 1.125 | 0.206 | 0.08 | 9.830000000000001e-26 | 3.31e-21 |
| Malignant epithelial | C2 WNT ECs | PDSS1 | 1.125 | 0.119 | 0.048 | 2.21e-14 | 7.45e-10 |
| Malignant epithelial | C2 WNT ECs | RNF4 | 1.125 | 0.166 | 0.08 | 2.93e-14 | 9.88e-10 |
| Malignant epithelial | C2 WNT ECs | C1orf122 | 1.124 | 0.524 | 0.238 | 2.45e-55 | 8.26e-51 |
| Malignant epithelial | C2 WNT ECs | HSPA5 | 1.123 | 0.888 | 0.686 | 5.779999999999999e-67 | 1.95e-62 |
| Malignant epithelial | C2 WNT ECs | RAB3IP | 1.123 | 0.357 | 0.162 | 2.59e-35 | 8.71e-31 |
| Malignant epithelial | C2 WNT ECs | SLC19A2 | 1.122 | 0.111 | 0.039 | 7.02e-17 | 2.36e-12 |
| Malignant epithelial | C2 WNT ECs | ACBD5 | 1.121 | 0.215 | 0.081 | 6.18e-28 | 2.08e-23 |
| Malignant epithelial | C2 WNT ECs | SERTAD1 | 1.12 | 0.76 | 0.556 | 3.97e-46 | 1.34e-41 |
| Malignant epithelial | C2 WNT ECs | DPP3 | 1.12 | 0.302 | 0.121 | 4.1e-36 | 1.38e-31 |
| Malignant epithelial | C2 WNT ECs | EXOSC10 | 1.119 | 0.165 | 0.061 | 2.94e-22 | 9.92e-18 |
| Malignant epithelial | C2 WNT ECs | TAF13 | 1.118 | 0.164 | 0.06 | 1.46e-22 | 4.92e-18 |
| Malignant epithelial | C2 WNT ECs | CCDC93 | 1.118 | 0.185 | 0.073 | 2.19e-22 | 7.39e-18 |
| Malignant epithelial | C2 WNT ECs | C15orf39 | 1.118 | 0.204 | 0.086 | 7.66e-22 | 2.58e-17 |
| Malignant epithelial | C2 WNT ECs | DGKQ | 1.118 | 0.114 | 0.04 | 5.91e-17 | 1.99e-12 |
| Malignant epithelial | C2 WNT ECs | IGHMBP2 | 1.118 | 0.1 | 0.037 | 9.14e-14 | 3.08e-09 |
| Malignant epithelial | C2 WNT ECs | ADGRG1 | 1.117 | 0.594 | 0.283 | 3.85e-59 | 1.3e-54 |
| Malignant epithelial | C2 WNT ECs | PLXND1 | 1.117 | 0.127 | 0.051 | 3.04e-15 | 1.02e-10 |
| Malignant epithelial | C2 WNT ECs | BAZ2B | 1.116 | 0.399 | 0.177 | 5.94e-41 | 2e-36 |
| Malignant epithelial | C2 WNT ECs | MMP14 | 1.116 | 0.476 | 0.242 | 1.3e-38 | 4.38e-34 |
| Malignant epithelial | C2 WNT ECs | ARRB2 | 1.115 | 0.172 | 0.068 | 1.8e-20 | 6.05e-16 |
| Malignant epithelial | C2 WNT ECs | C10orf10 | 1.114 | 0.355 | 0.161 | 3.94e-34 | 1.33e-29 |
| Malignant epithelial | C2 WNT ECs | CNKSR1 | 1.114 | 0.276 | 0.108 | 6.12e-33 | 2.06e-28 |
| Malignant epithelial | C2 WNT ECs | ZNF431 | 1.114 | 0.176 | 0.086 | 4.95e-14 | 1.67e-09 |
| Malignant epithelial | C2 WNT ECs | ARL5A | 1.113 | 0.601 | 0.311 | 6.429999999999999e-54 | 2.17e-49 |
| Malignant epithelial | C2 WNT ECs | CRELD1 | 1.113 | 0.235 | 0.094 | 7.49e-28 | 2.52e-23 |
| Malignant epithelial | C2 WNT ECs | EIF5A2 | 1.113 | 0.156 | 0.063 | 7.21e-18 | 2.43e-13 |
| Malignant epithelial | C2 WNT ECs | PAQR8 | 1.112 | 0.133 | 0.057 | 7.73e-14 | 2.61e-09 |
| Malignant epithelial | C2 WNT ECs | UTP11L | 1.111 | 0.39 | 0.165 | 2.5e-44 | 8.43e-40 |
| Malignant epithelial | C2 WNT ECs | PLEKHM1 | 1.111 | 0.129 | 0.062 | 1.53e-10 | 5.14e-06 |
| Malignant epithelial | C2 WNT ECs | SH3BP2 | 1.108 | 0.137 | 0.049 | 8.67e-20 | 2.92e-15 |
| Malignant epithelial | C2 WNT ECs | ELL2 | 1.108 | 0.157 | 0.074 | 2.21e-13 | 7.45e-09 |
| Malignant epithelial | C2 WNT ECs | RCHY1 | 1.105 | 0.21 | 0.086 | 1.43e-23 | 4.81e-19 |
| Malignant epithelial | C2 WNT ECs | NDFIP1 | 1.102 | 0.805 | 0.547 | 8.680000000000001e-59 | 2.92e-54 |
| Malignant epithelial | C2 WNT ECs | MICAL1 | 1.102 | 0.212 | 0.082 | 3.2e-26 | 1.08e-21 |
| Malignant epithelial | C2 WNT ECs | CNOT6L | 1.102 | 0.153 | 0.065 | 4.75e-16 | 1.6e-11 |
| Malignant epithelial | C2 WNT ECs | C3orf58 | 1.102 | 0.116 | 0.048 | 3.99e-13 | 1.34e-08 |
| Malignant epithelial | C2 WNT ECs | FOXA2 | 1.101 | 0.265 | 0.133 | 7.460000000000001e-21 | 2.51e-16 |
| Malignant epithelial | C2 WNT ECs | LRRC8A | 1.1 | 0.46 | 0.264 | 9.240000000000001e-29 | 3.11e-24 |
| Malignant epithelial | C2 WNT ECs | ZNF143 | 1.099 | 0.138 | 0.057 | 1.56e-15 | 5.25e-11 |
| Malignant epithelial | C2 WNT ECs | TRA2B | 1.098 | 0.842 | 0.58 | 1.94e-61 | 6.53e-57 |
| Malignant epithelial | C2 WNT ECs | CHORDC1 | 1.098 | 0.513 | 0.266 | 5.799999999999999e-43 | 1.95e-38 |
| Malignant epithelial | C2 WNT ECs | NT5C3B | 1.098 | 0.338 | 0.151 | 2.66e-33 | 8.97e-29 |
| Malignant epithelial | C2 WNT ECs | SNN | 1.098 | 0.111 | 0.048 | 6.78e-12 | 2.29e-07 |
| Malignant epithelial | C2 WNT ECs | GTF2F2 | 1.097 | 0.647 | 0.311 | 2.51e-71 | 8.439999999999999e-67 |
| Malignant epithelial | C2 WNT ECs | ODF2L | 1.097 | 0.337 | 0.143 | 2.88e-36 | 9.710000000000001e-32 |
| Malignant epithelial | C2 WNT ECs | IGF1R | 1.097 | 0.184 | 0.078 | 1.5e-19 | 5.06e-15 |
| Malignant epithelial | C2 WNT ECs | TTC33 | 1.097 | 0.112 | 0.042 | 4.92e-15 | 1.66e-10 |
| Malignant epithelial | C2 WNT ECs | APP | 1.096 | 0.87 | 0.616 | 8.9e-70 | 3e-65 |
| Malignant epithelial | C2 WNT ECs | CTB-55O6.8 | 1.096 | 0.131 | 0.046 | 3.65e-19 | 1.23e-14 |
| Malignant epithelial | C2 WNT ECs | PELI1 | 1.095 | 0.391 | 0.179 | 1.33e-36 | 4.47e-32 |
| Malignant epithelial | C2 WNT ECs | TAMM41 | 1.095 | 0.114 | 0.041 | 2.47e-16 | 8.32e-12 |
| Malignant epithelial | C2 WNT ECs | USO1 | 1.094 | 0.31 | 0.126 | 6.36e-36 | 2.14e-31 |
| Malignant epithelial | C2 WNT ECs | ATAT1 | 1.094 | 0.172 | 0.066 | 1.54e-21 | 5.17e-17 |

| compartment | subcluster | gene | avg_log2FC | pct.1 | pct.2 | p_val | p_val_adj |
|---|---|---|---|---|---|---|---|
| Malignant epithelial | C2 WNT ECs | SPHK1 | 1.094 | 0.112 | 0.052 | 1.99e-10 | 6.71e-06 |
| Malignant epithelial | C2 WNT ECs | SEH1L | 1.093 | 0.238 | 0.096 | 1.6e-27 | 5.369999999999999e-23 |
| Malignant epithelial | C2 WNT ECs | OSBPL9 | 1.093 | 0.214 | 0.091 | 2.83e-22 | 9.55e-18 |
| Malignant epithelial | C2 WNT ECs | ST7 | 1.093 | 0.129 | 0.056 | 3.54e-13 | 1.19e-08 |
| Malignant epithelial | C2 WNT ECs | FAM65A | 1.092 | 0.191 | 0.075 | 5.03e-23 | 1.7e-18 |
| Malignant epithelial | C2 WNT ECs | RPL21 | 1.091 | 0.999 | 0.99 | 7.49e-157 | 2.52e-152 |
| Malignant epithelial | C2 WNT ECs | TUFT1 | 1.091 | 0.342 | 0.158 | 1.67e-31 | 5.62e-27 |
| Malignant epithelial | C2 WNT ECs | FAAH | 1.091 | 0.187 | 0.072 | 6.519999999999999e-23 | 2.2e-18 |
| Malignant epithelial | C2 WNT ECs | PSPC1 | 1.09 | 0.325 | 0.134 | 8.570000000000001e-37 | 2.89e-32 |
| Malignant epithelial | C2 WNT ECs | UTP14C | 1.09 | 0.123 | 0.044 | 5.85e-18 | 1.97e-13 |
| Malignant epithelial | C2 WNT ECs | RAB18 | 1.089 | 0.471 | 0.208 | 9.52e-51 | 3.21e-46 |
| Malignant epithelial | C2 WNT ECs | SESN2 | 1.089 | 0.149 | 0.065 | 3.24e-15 | 1.09e-10 |
| Malignant epithelial | C2 WNT ECs | RPF1 | 1.088 | 0.41 | 0.169 | 3.04e-47 | 1.02e-42 |
| Malignant epithelial | C2 WNT ECs | ARID3A | 1.088 | 0.388 | 0.175 | 5.07e-39 | 1.71e-34 |
| Malignant epithelial | C2 WNT ECs | AMIGO2 | 1.088 | 0.298 | 0.128 | 1.88e-31 | 6.350000000000001e-27 |
| Malignant epithelial | C2 WNT ECs | ERCC5 | 1.088 | 0.221 | 0.094 | 3.35e-23 | 1.13e-18 |
| Malignant epithelial | C2 WNT ECs | SPRYD3 | 1.088 | 0.18 | 0.071 | 5.71e-22 | 1.92e-17 |
| Malignant epithelial | C2 WNT ECs | ZBTB2 | 1.088 | 0.154 | 0.074 | 8.85e-13 | 2.98e-08 |
| Malignant epithelial | C2 WNT ECs | HNRNPF | 1.087 | 0.829 | 0.507 | 1.37e-82 | 4.63e-78 |
| Malignant epithelial | C2 WNT ECs | ACAP3 | 1.086 | 0.162 | 0.068 | 1.31e-17 | 4.41e-13 |
| Malignant epithelial | C2 WNT ECs | C3orf52 | 1.085 | 0.164 | 0.075 | 5.98e-15 | 2.02e-10 |
| Malignant epithelial | C2 WNT ECs | DECR2 | 1.083 | 0.329 | 0.138 | 8.120000000000001e-37 | 2.73e-32 |
| Malignant epithelial | C2 WNT ECs | CASP3 | 1.083 | 0.246 | 0.1 | 4.22e-28 | 1.42e-23 |
| Malignant epithelial | C2 WNT ECs | RAP1GAP2 | 1.082 | 0.127 | 0.051 | 3.66e-15 | 1.23e-10 |
| Malignant epithelial | C2 WNT ECs | TRABD2A | 1.082 | 0.336 | 0.221 | 1.54e-14 | 5.17e-10 |
| Malignant epithelial | C2 WNT ECs | ZNF117 | 1.082 | 0.126 | 0.057 | 4.94e-12 | 1.66e-07 |
| Malignant epithelial | C2 WNT ECs | SARAF | 1.081 | 0.727 | 0.439 | 7.26e-57 | 2.44e-52 |
| Malignant epithelial | C2 WNT ECs | ZCCHC17 | 1.081 | 0.357 | 0.167 | 2.64e-31 | 8.9e-27 |
| Malignant epithelial | C2 WNT ECs | MVP | 1.08 | 0.769 | 0.493 | 2.85e-59 | 9.589999999999999e-55 |
| Malignant epithelial | C2 WNT ECs | HMGCR | 1.08 | 0.225 | 0.103 | 2.59e-20 | 8.73e-16 |
| Malignant epithelial | C2 WNT ECs | BRI3 | 1.079 | 0.876 | 0.581 | 4.15e-93 | 1.4e-88 |
| Malignant epithelial | C2 WNT ECs | GMPPA | 1.079 | 0.31 | 0.132 | 9.98e-33 | 3.36e-28 |
| Malignant epithelial | C2 WNT ECs | RNF186 | 1.079 | 0.139 | 0.05 | 8.63e-20 | 2.91e-15 |
| Malignant epithelial | C2 WNT ECs | MTRR | 1.079 | 0.134 | 0.055 | 3.55e-15 | 1.2e-10 |
| Malignant epithelial | C2 WNT ECs | WWOX | 1.079 | 0.104 | 0.045 | 4.86e-11 | 1.64e-06 |
| Malignant epithelial | C2 WNT ECs | DBNDD1 | 1.078 | 0.143 | 0.053 | 3.73e-19 | 1.26e-14 |
| Malignant epithelial | C2 WNT ECs | TMPRSS4 | 1.077 | 0.51 | 0.27 | 9.42e-40 | 3.17e-35 |
| Malignant epithelial | C2 WNT ECs | FAM21C | 1.077 | 0.127 | 0.047 | 2.08e-17 | 7.01e-13 |
| Malignant epithelial | C2 WNT ECs | CTDP1 | 1.077 | 0.118 | 0.046 | 6.73e-15 | 2.27e-10 |
| Malignant epithelial | C2 WNT ECs | ATP1A1 | 1.076 | 0.781 | 0.511 | 4.38e-59 | 1.48e-54 |
| Malignant epithelial | C2 WNT ECs | ROCK2 | 1.076 | 0.441 | 0.212 | 9.53e-40 | 3.21e-35 |
| Malignant epithelial | C2 WNT ECs | CCDC120 | 1.076 | 0.153 | 0.056 | 2.89e-21 | 9.72e-17 |
| Malignant epithelial | C2 WNT ECs | STK40 | 1.076 | 0.183 | 0.074 | 3.88e-21 | 1.31e-16 |
| Malignant epithelial | C2 WNT ECs | PLAGL2 | 1.076 | 0.173 | 0.083 | 1.71e-14 | 5.75e-10 |
| Malignant epithelial | C2 WNT ECs | AZI2 | 1.075 | 0.295 | 0.125 | 5.6e-31 | 1.89e-26 |
| Malignant epithelial | C2 WNT ECs | ZNF746 | 1.075 | 0.127 | 0.05 | 9.77e-16 | 3.29e-11 |
| Malignant epithelial | C2 WNT ECs | ERBB3 | 1.074 | 0.743 | 0.421 | 5.64e-66 | 1.9e-61 |
| Malignant epithelial | C2 WNT ECs | POR | 1.074 | 0.548 | 0.272 | 1.23e-50 | 4.13e-46 |
| Malignant epithelial | C2 WNT ECs | SLU7 | 1.074 | 0.464 | 0.209 | 6.37e-48 | 2.15e-43 |
| Malignant epithelial | C2 WNT ECs | CDC34 | 1.074 | 0.52 | 0.289 | 3.96e-36 | 1.33e-31 |
| Malignant epithelial | C2 WNT ECs | SMPDL3A | 1.074 | 0.272 | 0.108 | 3.46e-32 | 1.17e-27 |
| Malignant epithelial | C2 WNT ECs | CCDC122 | 1.074 | 0.138 | 0.049 | 9.62e-20 | 3.24e-15 |
| Malignant epithelial | C2 WNT ECs | GBAS | 1.072 | 0.479 | 0.252 | 1.1e-36 | 3.7e-32 |
| Malignant epithelial | C2 WNT ECs | NINJ2 | 1.071 | 0.123 | 0.049 | 6e-15 | 2.02e-10 |
| Malignant epithelial | C2 WNT ECs | CCDC64 | 1.071 | 0.104 | 0.043 | 6.94e-12 | 2.34e-07 |
| Malignant epithelial | C2 WNT ECs | TRIM2 | 1.07 | 0.553 | 0.232 | 6.83e-65 | 2.3e-60 |
| Malignant epithelial | C2 WNT ECs | POLR2A | 1.07 | 0.379 | 0.165 | 2.05e-39 | 6.89e-35 |
| Malignant epithelial | C2 WNT ECs | FAM114A1 | 1.07 | 0.327 | 0.146 | 9.89e-32 | 3.33e-27 |
| Malignant epithelial | C2 WNT ECs | FNIP2 | 1.07 | 0.194 | 0.086 | 9.02e-19 | 3.04e-14 |
| Malignant epithelial | C2 WNT ECs | PRMT5 | 1.069 | 0.307 | 0.14 | 7.160000000000001e-29 | 2.41e-24 |
| Malignant epithelial | C2 WNT ECs | RAB22A | 1.068 | 0.655 | 0.338 | 1.39e-59 | 4.67e-55 |
| Malignant epithelial | C2 WNT ECs | DHCR24 | 1.068 | 0.204 | 0.1 | 9.53e-17 | 3.21e-12 |
| Malignant epithelial | C2 WNT ECs | RP11-660L16.2 | 1.067 | 0.152 | 0.052 | 4.159999999999999e-23 | 1.4e-18 |
| Malignant epithelial | C2 WNT ECs | KCNQ1OT1 | 1.066 | 0.618 | 0.362 | 1.91e-39 | 6.440000000000001e-35 |

| compartment | subcluster | gene | avg_log2FC | pct.1 | pct.2 | p_val | p_val_adj |
|---|---|---|---|---|---|---|---|
| Malignant epithelial | C2 WNT ECs | MACF1 | 1.066 | 0.372 | 0.162 | 4.35e-38 | 1.47e-33 |
| Malignant epithelial | C2 WNT ECs | SLC16A1 | 1.066 | 0.313 | 0.139 | 2.2e-30 | 7.420000000000001e-26 |
| Malignant epithelial | C2 WNT ECs | GPALPP1 | 1.066 | 0.248 | 0.099 | 6.48e-29 | 2.18e-24 |
| Malignant epithelial | C2 WNT ECs | TMEM106B | 1.065 | 0.543 | 0.291 | 1.34e-43 | 4.52e-39 |
| Malignant epithelial | C2 WNT ECs | RECQL | 1.065 | 0.164 | 0.07 | 5.92e-17 | 1.99e-12 |
| Malignant epithelial | C2 WNT ECs | SERPINB6 | 1.064 | 0.697 | 0.385 | 4.47e-64 | 1.5e-59 |
| Malignant epithelial | C2 WNT ECs | TINAGL1 | 1.063 | 0.762 | 0.418 | 1.38e-71 | 4.659999999999999e-67 |
| Malignant epithelial | C2 WNT ECs | BOD1L1 | 1.063 | 0.25 | 0.102 | 7.33e-28 | 2.47e-23 |
| Malignant epithelial | C2 WNT ECs | ECHS1 | 1.062 | 0.705 | 0.427 | 5.14e-61 | 1.73e-56 |
| Malignant epithelial | C2 WNT ECs | RP1-34B20.21 | 1.062 | 0.225 | 0.1 | 2.85e-21 | 9.62e-17 |
| Malignant epithelial | C2 WNT ECs | RABEP2 | 1.06 | 0.263 | 0.113 | 7.46e-27 | 2.51e-22 |
| Malignant epithelial | C2 WNT ECs | STAM2 | 1.06 | 0.141 | 0.053 | 1.66e-18 | 5.59e-14 |
| Malignant epithelial | C2 WNT ECs | MT-ATP6 | 1.059 | 0.997 | 0.981 | 7.49e-143 | 2.52e-138 |
| Malignant epithelial | C2 WNT ECs | PRPF38B | 1.058 | 0.643 | 0.349 | 4.33e-54 | 1.46e-49 |
| Malignant epithelial | C2 WNT ECs | MEAF6 | 1.058 | 0.555 | 0.279 | 5.88e-50 | 1.98e-45 |
| Malignant epithelial | C2 WNT ECs | VSNL1 | 1.058 | 0.25 | 0.113 | 9.84e-24 | 3.32e-19 |
| Malignant epithelial | C2 WNT ECs | FOXA3 | 1.058 | 0.202 | 0.11 | 2.13e-12 | 7.17e-08 |
| Malignant epithelial | C2 WNT ECs | SARS | 1.057 | 0.549 | 0.272 | 6.94e-50 | 2.34e-45 |
| Malignant epithelial | C2 WNT ECs | RAB32 | 1.057 | 0.409 | 0.195 | 3.55e-37 | 1.2e-32 |
| Malignant epithelial | C2 WNT ECs | PPIL4 | 1.057 | 0.296 | 0.124 | 9.56e-32 | 3.22e-27 |
| Malignant epithelial | C2 WNT ECs | B3GAT2 | 1.057 | 0.2 | 0.081 | 4.14e-23 | 1.39e-18 |
| Malignant epithelial | C2 WNT ECs | METRN | 1.056 | 0.276 | 0.114 | 2.74e-31 | 9.22e-27 |
| Malignant epithelial | C2 WNT ECs | PKIG | 1.056 | 0.264 | 0.108 | 4.4e-30 | 1.48e-25 |
| Malignant epithelial | C2 WNT ECs | TMEM181 | 1.056 | 0.227 | 0.092 | 5.63e-26 | 1.9e-21 |
| Malignant epithelial | C2 WNT ECs | MAD2L2 | 1.056 | 0.284 | 0.138 | 4.409999999999999e-23 | 1.49e-18 |
| Malignant epithelial | C2 WNT ECs | C4orf48 | 1.054 | 0.915 | 0.662 | 2.68e-91 | 9.05e-87 |
| Malignant epithelial | C2 WNT ECs | SLCO4A1-AS1 | 1.054 | 0.281 | 0.1 | 1.32e-41 | 4.44e-37 |
| Malignant epithelial | C2 WNT ECs | AKAP17A | 1.054 | 0.3 | 0.122 | 2.67e-34 | 8.989999999999999e-30 |
| Malignant epithelial | C2 WNT ECs | WAPL | 1.054 | 0.254 | 0.103 | 1.11e-29 | 3.75e-25 |
| Malignant epithelial | C2 WNT ECs | MIER1 | 1.053 | 0.287 | 0.123 | 1.33e-29 | 4.49e-25 |
| Malignant epithelial | C2 WNT ECs | PVR | 1.053 | 0.194 | 0.08 | 1.74e-21 | 5.85e-17 |
| Malignant epithelial | C2 WNT ECs | ACADVL | 1.052 | 0.744 | 0.408 | 1.09e-70 | 3.67e-66 |
| Malignant epithelial | C2 WNT ECs | LGMN | 1.052 | 0.521 | 0.247 | 9.65e-49 | 3.25e-44 |
| Malignant epithelial | C2 WNT ECs | FOXO3 | 1.052 | 0.524 | 0.295 | 3.3e-37 | 1.11e-32 |
| Malignant epithelial | C2 WNT ECs | SETD5 | 1.051 | 0.43 | 0.186 | 5.39e-47 | 1.82e-42 |
| Malignant epithelial | C2 WNT ECs | CACNA2D4 | 1.051 | 0.314 | 0.156 | 2.1e-25 | 7.06e-21 |
| Malignant epithelial | C2 WNT ECs | PID1 | 1.051 | 0.137 | 0.052 | 6.76e-18 | 2.28e-13 |
| Malignant epithelial | C2 WNT ECs | WAC | 1.05 | 0.547 | 0.257 | 4.46e-53 | 1.5e-48 |
| Malignant epithelial | C2 WNT ECs | KDF1 | 1.049 | 0.2 | 0.081 | 1.04e-22 | 3.51e-18 |
| Malignant epithelial | C2 WNT ECs | DYNLT3 | 1.047 | 0.614 | 0.337 | 5.39e-51 | 1.81e-46 |
| Malignant epithelial | C2 WNT ECs | ANKRD12 | 1.047 | 0.487 | 0.224 | 1.71e-48 | 5.77e-44 |
| Malignant epithelial | C2 WNT ECs | CARS | 1.047 | 0.311 | 0.135 | 1e-31 | 3.37e-27 |
| Malignant epithelial | C2 WNT ECs | SLC3A2 | 1.046 | 0.832 | 0.565 | 1.62e-65 | 5.44e-61 |
| Malignant epithelial | C2 WNT ECs | PKNOX1 | 1.046 | 0.172 | 0.079 | 1.93e-15 | 6.49e-11 |
| Malignant epithelial | C2 WNT ECs | SLC7A1 | 1.045 | 0.254 | 0.112 | 2.17e-25 | 7.320000000000001e-21 |
| Malignant epithelial | C2 WNT ECs | KB-1732A1.1 | 1.044 | 0.23 | 0.112 | 1.43e-18 | 4.81e-14 |
| Malignant epithelial | C2 WNT ECs | FAM131A | 1.044 | 0.141 | 0.062 | 5.22e-14 | 1.76e-09 |
| Malignant epithelial | C2 WNT ECs | TNFAIP8L1 | 1.042 | 0.146 | 0.065 | 4.23e-14 | 1.43e-09 |
| Malignant epithelial | C2 WNT ECs | C6orf62 | 1.04 | 0.396 | 0.168 | 1.51e-43 | 5.1e-39 |
| Malignant epithelial | C2 WNT ECs | NELFA | 1.04 | 0.302 | 0.12 | 2.99e-36 | 1.01e-31 |
| Malignant epithelial | C2 WNT ECs | CORO1A | 1.04 | 0.302 | 0.127 | 3.45e-32 | 1.16e-27 |
| Malignant epithelial | C2 WNT ECs | ARAP2 | 1.039 | 0.157 | 0.056 | 2.89e-22 | 9.73e-18 |
| Malignant epithelial | C2 WNT ECs | APOM | 1.039 | 0.124 | 0.049 | 2.22e-15 | 7.47e-11 |
| Malignant epithelial | C2 WNT ECs | PRR5 | 1.037 | 0.307 | 0.138 | 1.33e-29 | 4.47e-25 |
| Malignant epithelial | C2 WNT ECs | CUL4A | 1.035 | 0.555 | 0.262 | 2.1e-55 | 7.08e-51 |
| Malignant epithelial | C2 WNT ECs | DOK4 | 1.035 | 0.365 | 0.164 | 2.54e-35 | 8.56e-31 |
| Malignant epithelial | C2 WNT ECs | BTG2 | 1.035 | 0.763 | 0.546 | 1.94e-33 | 6.55e-29 |
| Malignant epithelial | C2 WNT ECs | POLR1D | 1.034 | 0.916 | 0.68 | 6.099999999999999e-96 | 2.06e-91 |
| Malignant epithelial | C2 WNT ECs | CHD6 | 1.034 | 0.403 | 0.199 | 6.52e-33 | 2.2e-28 |
| Malignant epithelial | C2 WNT ECs | CCDC28A | 1.033 | 0.25 | 0.114 | 3.56e-23 | 1.2e-18 |
| Malignant epithelial | C2 WNT ECs | OPN3 | 1.033 | 0.217 | 0.1 | 2.63e-19 | 8.86e-15 |
| Malignant epithelial | C2 WNT ECs | SEP15 | 1.032 | 0.792 | 0.48 | 2.37e-73 | 8e-69 |
| Malignant epithelial | C2 WNT ECs | GRIN2B | 1.032 | 0.134 | 0.055 | 6.71e-15 | 2.26e-10 |
| Malignant epithelial | C2 WNT ECs | CHPF | 1.031 | 0.648 | 0.34 | 3.97e-56 | 1.34e-51 |

| compartment | subcluster | gene | avg_log2FC | pct.1 | pct.2 | p_val | p_val_adj |
|---|---|---|---|---|---|---|---|
| Malignant epithelial | C2 WNT ECs | DCAF13 | 1.03 | 0.51 | 0.286 | 6.24e-35 | 2.1e-30 |
| Malignant epithelial | C2 WNT ECs | SNRNP48 | 1.03 | 0.176 | 0.073 | 2.72e-19 | 9.17e-15 |
| Malignant epithelial | C2 WNT ECs | RB1CC1 | 1.029 | 0.543 | 0.275 | 7.479999999999999e-48 | 2.52e-43 |
| Malignant epithelial | C2 WNT ECs | PREB | 1.029 | 0.398 | 0.172 | 1.11e-42 | 3.73e-38 |
| Malignant epithelial | C2 WNT ECs | KRT80 | 1.029 | 0.199 | 0.098 | 1.09e-15 | 3.67e-11 |
| Malignant epithelial | C2 WNT ECs | PSMD7 | 1.028 | 0.74 | 0.46 | 8.83e-59 | 2.97e-54 |
| Malignant epithelial | C2 WNT ECs | PRSS8 | 1.028 | 0.705 | 0.418 | 2.16e-54 | 7.270000000000001e-50 |
| Malignant epithelial | C2 WNT ECs | ABT1 | 1.028 | 0.384 | 0.174 | 1.54e-36 | 5.18e-32 |
| Malignant epithelial | C2 WNT ECs | AJUBA | 1.028 | 0.191 | 0.085 | 2.6e-18 | 8.76e-14 |
| Malignant epithelial | C2 WNT ECs | FIG4 | 1.028 | 0.107 | 0.04 | 2.38e-14 | 8.01e-10 |
| Malignant epithelial | C2 WNT ECs | UFM1 | 1.027 | 0.793 | 0.466 | 1.5e-68 | 5.06e-64 |
| Malignant epithelial | C2 WNT ECs | GNAI3 | 1.027 | 0.529 | 0.257 | 4.02e-49 | 1.36e-44 |
| Malignant epithelial | C2 WNT ECs | MYO6 | 1.027 | 0.593 | 0.312 | 8.94e-49 | 3.01e-44 |
| Malignant epithelial | C2 WNT ECs | BIVM | 1.025 | 0.119 | 0.049 | 1.19e-13 | 3.99e-09 |
| Malignant epithelial | C2 WNT ECs | ADIRF | 1.025 | 0.691 | 0.887 | 3.67e-07 | 0.0124 |
| Malignant epithelial | C2 WNT ECs | SERINC1 | 1.023 | 0.487 | 0.229 | 6.659999999999999e-45 | 2.24e-40 |
| Malignant epithelial | C2 WNT ECs | DGKZ | 1.023 | 0.395 | 0.171 | 3.59e-41 | 1.21e-36 |
| Malignant epithelial | C2 WNT ECs | AMPD2 | 1.022 | 0.123 | 0.048 | 3.77e-15 | 1.27e-10 |
| Malignant epithelial | C2 WNT ECs | LINC01006 | 1.022 | 0.168 | 0.079 | 1.01e-14 | 3.4e-10 |
| Malignant epithelial | C2 WNT ECs | DR1 | 1.021 | 0.31 | 0.142 | 3.68e-28 | 1.24e-23 |
| Malignant epithelial | C2 WNT ECs | RHOF | 1.021 | 0.472 | 0.306 | 4.78e-21 | 1.61e-16 |
| Malignant epithelial | C2 WNT ECs | DBNL | 1.02 | 0.629 | 0.356 | 5.69e-48 | 1.92e-43 |
| Malignant epithelial | C2 WNT ECs | USP33 | 1.019 | 0.189 | 0.076 | 6.61e-22 | 2.23e-17 |
| Malignant epithelial | C2 WNT ECs | ZC3H12A | 1.018 | 0.524 | 0.296 | 3.05e-34 | 1.03e-29 |
| Malignant epithelial | C2 WNT ECs | WASF2 | 1.016 | 0.618 | 0.337 | 6.49e-49 | 2.19e-44 |
| Malignant epithelial | C2 WNT ECs | HOOK1 | 1.016 | 0.183 | 0.07 | 3.919999999999999e-23 | 1.32e-18 |
| Malignant epithelial | C2 WNT ECs | RAPGEF2 | 1.016 | 0.134 | 0.063 | 9.87e-12 | 3.32e-07 |
| Malignant epithelial | C2 WNT ECs | TESC | 1.015 | 0.433 | 0.179 | 2.35e-50 | 7.92e-46 |
| Malignant epithelial | C2 WNT ECs | NUDT9 | 1.015 | 0.181 | 0.078 | 1.65e-18 | 5.55e-14 |
| Malignant epithelial | C2 WNT ECs | ING1 | 1.014 | 0.491 | 0.259 | 4.84e-36 | 1.63e-31 |
| Malignant epithelial | C2 WNT ECs | TUBGCP2 | 1.014 | 0.336 | 0.155 | 1.72e-30 | 5.8e-26 |
| Malignant epithelial | C2 WNT ECs | CDC42EP3 | 1.013 | 0.421 | 0.21 | 9.179999999999999e-34 | 3.09e-29 |
| Malignant epithelial | C2 WNT ECs | TMEM125 | 1.013 | 0.267 | 0.116 | 2.84e-26 | 9.58e-22 |
| Malignant epithelial | C2 WNT ECs | SERPINB1 | 1.012 | 0.801 | 0.608 | 7.01e-42 | 2.36e-37 |
| Malignant epithelial | C2 WNT ECs | SCNN1A | 1.012 | 0.471 | 0.233 | 2.15e-38 | 7.239999999999999e-34 |
| Malignant epithelial | C2 WNT ECs | MTMR14 | 1.012 | 0.24 | 0.105 | 1.91e-23 | 6.44e-19 |
| Malignant epithelial | C2 WNT ECs | WBSCR27 | 1.011 | 0.368 | 0.17 | 1.69e-34 | 5.68e-30 |
| Malignant epithelial | C2 WNT ECs | FUOM | 1.011 | 0.444 | 0.25 | 7.310000000000001e-31 | 2.46e-26 |
| Malignant epithelial | C2 WNT ECs | TRIM4 | 1.011 | 0.164 | 0.066 | 9.15e-19 | 3.08e-14 |
| Malignant epithelial | C2 WNT ECs | PURB | 1.011 | 0.244 | 0.121 | 1.44e-18 | 4.84e-14 |
| Malignant epithelial | C2 WNT ECs | FAM73A | 1.011 | 0.133 | 0.053 | 6.47e-16 | 2.18e-11 |
| Malignant epithelial | C2 WNT ECs | C16orf70 | 1.011 | 0.103 | 0.047 | 6.84e-10 | 2.3e-05 |
| Malignant epithelial | C2 WNT ECs | CCAT1 | 1.011 | 0.123 | 0.068 | 2.02e-07 | 0.0068 |
| Malignant epithelial | C2 WNT ECs | XBP1 | 1.009 | 0.763 | 0.495 | 1.05e-50 | 3.54e-46 |
| Malignant epithelial | C2 WNT ECs | RNF10 | 1.009 | 0.463 | 0.225 | 4.02e-41 | 1.36e-36 |
| Malignant epithelial | C2 WNT ECs | DOCK7 | 1.009 | 0.107 | 0.042 | 4.45e-13 | 1.5e-08 |
| Malignant epithelial | C2 WNT ECs | SNX7 | 1.008 | 0.372 | 0.18 | 9.84e-32 | 3.320000000000001e-27 |
| Malignant epithelial | C2 WNT ECs | EPHX2 | 1.007 | 0.134 | 0.051 | 1.86e-17 | 6.27e-13 |
| Malignant epithelial | C2 WNT ECs | PTPRE | 1.006 | 0.217 | 0.093 | 6.02e-22 | 2.03e-17 |
| Malignant epithelial | C2 WNT ECs | EAF1 | 1.006 | 0.185 | 0.075 | 1.92e-21 | 6.46e-17 |
| Malignant epithelial | C2 WNT ECs | ELOVL1 | 1.004 | 0.553 | 0.273 | 4e-50 | 1.35e-45 |
| Malignant epithelial | C2 WNT ECs | PSORS1C1 | 1.004 | 0.237 | 0.095 | 2.01e-27 | 6.779999999999999e-23 |
| Malignant epithelial | C2 WNT ECs | MKNK2 | 1.003 | 0.472 | 0.263 | 3.26e-32 | 1.1e-27 |
| Malignant epithelial | C2 WNT ECs | KLHL24 | 1.003 | 0.333 | 0.154 | 1.4e-30 | 4.71e-26 |
| Malignant epithelial | C2 WNT ECs | KLHDC3 | 1.002 | 0.39 | 0.209 | 6.130000000000001e-27 | 2.06e-22 |
| Malignant epithelial | C2 WNT ECs | LIMK2 | 1.002 | 0.185 | 0.067 | 1.39e-25 | 4.680000000000001e-21 |
| Malignant epithelial | C2 WNT ECs | DST | 1.001 | 0.617 | 0.341 | 3.73e-48 | 1.26e-43 |
| Malignant epithelial | C2 WNT ECs | SVBP | 1.001 | 0.402 | 0.18 | 3.37e-39 | 1.14e-34 |
| Malignant epithelial | C2 WNT ECs | TMEM33 | 1.001 | 0.399 | 0.181 | 2.64e-38 | 8.89e-34 |
| Malignant epithelial | C2 WNT ECs | GRB7 | 1.001 | 0.448 | 0.217 | 8.28e-38 | 2.79e-33 |
| Malignant epithelial | C2 WNT ECs | KLHL21 | 1.001 | 0.268 | 0.145 | 1.82e-17 | 6.13e-13 |
| Malignant epithelial | C2 WNT ECs | PDLIM7 | 1.0 | 0.43 | 0.237 | 4.37e-28 | 1.47e-23 |
| Malignant epithelial | C2 WNT ECs | MYO7B | 1.0 | 0.141 | 0.057 | 2.26e-16 | 7.6e-12 |
| Malignant epithelial | C2 WNT ECs | LRIG1 | 1.0 | 0.12 | 0.051 | 5.72e-13 | 1.93e-08 |

| compartment | subcluster | gene | avg_log2FC | pct.1 | pct.2 | p_val | p_val_adj |
|---|---|---|---|---|---|---|---|
| Malignant epithelial | C2 WNT ECs | STARD4 | 1.0 | 0.101 | 0.045 | 3.63e-10 | 1.22e-05 |
| Malignant epithelial | C2 WNT ECs | STIP1 | 0.999 | 0.625 | 0.331 | 4.56e-51 | 1.54e-46 |
| Malignant epithelial | C2 WNT ECs | CPTP | 0.999 | 0.248 | 0.104 | 2.3e-26 | 7.76e-22 |
| Malignant epithelial | C2 WNT ECs | IMMP2L | 0.999 | 0.26 | 0.113 | 1.13e-25 | 3.81e-21 |
| Malignant epithelial | C2 WNT ECs | BANP | 0.998 | 0.154 | 0.068 | 2.7e-15 | 9.09e-11 |
| Malignant epithelial | C2 WNT ECs | TMBIM1 | 0.997 | 0.709 | 0.456 | 7.33e-44 | 2.47e-39 |
| Malignant epithelial | C2 WNT ECs | CHD7 | 0.997 | 0.475 | 0.225 | 1.19e-43 | 4.02e-39 |
| Malignant epithelial | C2 WNT ECs | TWF2 | 0.997 | 0.502 | 0.253 | 1.63e-41 | 5.48e-37 |
| Malignant epithelial | C2 WNT ECs | NXF1 | 0.997 | 0.368 | 0.179 | 1.76e-30 | 5.94e-26 |
| Malignant epithelial | C2 WNT ECs | TRIM33 | 0.997 | 0.276 | 0.123 | 6.66e-27 | 2.24e-22 |
| Malignant epithelial | C2 WNT ECs | HSD17B4 | 0.997 | 0.249 | 0.117 | 1.02e-21 | 3.43e-17 |
| Malignant epithelial | C2 WNT ECs | PAK1IP1 | 0.996 | 0.275 | 0.114 | 3.03e-30 | 1.02e-25 |
| Malignant epithelial | C2 WNT ECs | CDYL | 0.996 | 0.184 | 0.077 | 7.92e-20 | 2.67e-15 |
| Malignant epithelial | C2 WNT ECs | ITGB5 | 0.995 | 0.333 | 0.146 | 4.44e-32 | 1.5e-27 |
| Malignant epithelial | C2 WNT ECs | TMC7 | 0.995 | 0.198 | 0.087 | 2.18e-19 | 7.35e-15 |
| Malignant epithelial | C2 WNT ECs | YIPF3 | 0.993 | 0.732 | 0.454 | 5.64e-57 | 1.9e-52 |
| Malignant epithelial | C2 WNT ECs | FARP1 | 0.993 | 0.369 | 0.161 | 3.39e-37 | 1.14e-32 |
| Malignant epithelial | C2 WNT ECs | TDRD3 | 0.993 | 0.146 | 0.056 | 1.65e-18 | 5.57e-14 |
| Malignant epithelial | C2 WNT ECs | NFYA | 0.993 | 0.171 | 0.077 | 9.33e-16 | 3.14e-11 |
| Malignant epithelial | C2 WNT ECs | SLC20A2 | 0.992 | 0.417 | 0.226 | 6.400000000000001e-27 | 2.16e-22 |
| Malignant epithelial | C2 WNT ECs | MRS2 | 0.992 | 0.15 | 0.063 | 3.4e-16 | 1.15e-11 |
| Malignant epithelial | C2 WNT ECs | TMPRSS2 | 0.991 | 0.48 | 0.199 | 7.109999999999999e-54 | 2.4e-49 |
| Malignant epithelial | C2 WNT ECs | RP11-14N7.2 | 0.991 | 0.33 | 0.159 | 7.66e-27 | 2.58e-22 |
| Malignant epithelial | C2 WNT ECs | INTS8 | 0.991 | 0.24 | 0.112 | 3.9e-21 | 1.31e-16 |
| Malignant epithelial | C2 WNT ECs | PCMT1 | 0.99 | 0.7 | 0.393 | 8.280000000000001e-56 | 2.79e-51 |
| Malignant epithelial | C2 WNT ECs | HBEGF | 0.99 | 0.602 | 0.351 | 2.25e-41 | 7.570000000000001e-37 |
| Malignant epithelial | C2 WNT ECs | LRRC1 | 0.99 | 0.356 | 0.155 | 7.32e-36 | 2.47e-31 |
| Malignant epithelial | C2 WNT ECs | ANKRD49 | 0.99 | 0.24 | 0.105 | 1.04e-23 | 3.52e-19 |
| Malignant epithelial | C2 WNT ECs | MAGIX | 0.99 | 0.124 | 0.045 | 1.04e-17 | 3.49e-13 |
| Malignant epithelial | C2 WNT ECs | RSRP1 | 0.989 | 0.616 | 0.328 | 5.17e-49 | 1.74e-44 |
| Malignant epithelial | C2 WNT ECs | PNPLA8 | 0.989 | 0.281 | 0.122 | 1.17e-28 | 3.949999999999999e-24 |
| Malignant epithelial | C2 WNT ECs | PITHD1 | 0.988 | 0.432 | 0.214 | 6.530000000000001e-35 | 2.2e-30 |
| Malignant epithelial | C2 WNT ECs | PRKCD | 0.988 | 0.299 | 0.131 | 3.1e-29 | 1.04e-24 |
| Malignant epithelial | C2 WNT ECs | PGM2L1 | 0.988 | 0.372 | 0.184 | 5.350000000000001e-29 | 1.8e-24 |
| Malignant epithelial | C2 WNT ECs | YTHDF3 | 0.988 | 0.35 | 0.17 | 1.58e-28 | 5.33e-24 |
| Malignant epithelial | C2 WNT ECs | OSBPL3 | 0.988 | 0.344 | 0.169 | 1.34e-27 | 4.52e-23 |
| Malignant epithelial | C2 WNT ECs | GNB1L | 0.988 | 0.192 | 0.077 | 4.33e-22 | 1.46e-17 |
| Malignant epithelial | C2 WNT ECs | HOMER1 | 0.988 | 0.123 | 0.054 | 3.27e-12 | 1.1e-07 |
| Malignant epithelial | C2 WNT ECs | EGFR | 0.987 | 0.264 | 0.133 | 9.73e-19 | 3.28e-14 |
| Malignant epithelial | C2 WNT ECs | BAG2 | 0.987 | 0.157 | 0.069 | 2.96e-15 | 9.97e-11 |
| Malignant epithelial | C2 WNT ECs | AKIRIN1 | 0.986 | 0.375 | 0.172 | 2.3e-35 | 7.74e-31 |
| Malignant epithelial | C2 WNT ECs | TFE3 | 0.986 | 0.18 | 0.079 | 8.03e-18 | 2.7e-13 |
| Malignant epithelial | C2 WNT ECs | WDR45 | 0.985 | 0.518 | 0.258 | 1.94e-44 | 6.52e-40 |
| Malignant epithelial | C2 WNT ECs | DEDD2 | 0.985 | 0.614 | 0.367 | 3.81e-39 | 1.28e-34 |
| Malignant epithelial | C2 WNT ECs | TRIM27 | 0.985 | 0.304 | 0.129 | 3.22e-32 | 1.09e-27 |
| Malignant epithelial | C2 WNT ECs | CDS1 | 0.985 | 0.217 | 0.094 | 8.86e-22 | 2.99e-17 |
| Malignant epithelial | C2 WNT ECs | PHF10 | 0.985 | 0.146 | 0.057 | 3.27e-18 | 1.1e-13 |
| Malignant epithelial | C2 WNT ECs | SRSF11 | 0.984 | 0.736 | 0.414 | 3.31e-64 | 1.12e-59 |
| Malignant epithelial | C2 WNT ECs | MYO1C | 0.984 | 0.311 | 0.141 | 2.7e-29 | 9.11e-25 |
| Malignant epithelial | C2 WNT ECs | UBTD2 | 0.984 | 0.124 | 0.062 | 2.07e-09 | 6.98e-05 |
| Malignant epithelial | C2 WNT ECs | TMEM150A | 0.983 | 0.227 | 0.092 | 2.01e-25 | 6.77e-21 |
| Malignant epithelial | C2 WNT ECs | KIAA0319L | 0.983 | 0.221 | 0.092 | 3.76e-23 | 1.27e-18 |
| Malignant epithelial | C2 WNT ECs | NEU1 | 0.982 | 0.72 | 0.415 | 1.98e-55 | 6.69e-51 |
| Malignant epithelial | C2 WNT ECs | PPP1R18 | 0.982 | 0.353 | 0.169 | 2.53e-30 | 8.52e-26 |
| Malignant epithelial | C2 WNT ECs | NEAT1 | 0.982 | 0.972 | 0.958 | 5.68e-20 | 1.91e-15 |
| Malignant epithelial | C2 WNT ECs | NUDT15 | 0.981 | 0.47 | 0.241 | 8.189999999999999e-38 | 2.76e-33 |
| Malignant epithelial | C2 WNT ECs | HMGA1 | 0.981 | 0.931 | 0.833 | 2.84e-32 | 9.55e-28 |
| Malignant epithelial | C2 WNT ECs | INPP5K | 0.981 | 0.143 | 0.051 | 1.38e-20 | 4.64e-16 |
| Malignant epithelial | C2 WNT ECs | MED26 | 0.981 | 0.141 | 0.068 | 2.32e-11 | 7.81e-07 |
| Malignant epithelial | C2 WNT ECs | ZNF274 | 0.979 | 0.198 | 0.098 | 3.25e-15 | 1.09e-10 |
| Malignant epithelial | C2 WNT ECs | PTCD3 | 0.978 | 0.368 | 0.162 | 3.39e-36 | 1.14e-31 |
| Malignant epithelial | C2 WNT ECs | CCPG1 | 0.978 | 0.258 | 0.119 | 1.87e-23 | 6.31e-19 |
| Malignant epithelial | C2 WNT ECs | GALNT1 | 0.978 | 0.211 | 0.1 | 7.87e-18 | 2.65e-13 |
| Malignant epithelial | C2 WNT ECs | TMEM9 | 0.976 | 0.436 | 0.207 | 9.34e-38 | 3.15e-33 |

| compartment | subcluster | gene | avg_log2FC | pct.1 | pct.2 | p_val | p_val_adj |
|---|---|---|---|---|---|---|---|
| Malignant epithelial | C2 WNT ECs | AC073283.4 | 0.976 | 0.104 | 0.048 | 1.28e-09 | 4.32e-05 |
| Malignant epithelial | C2 WNT ECs | EPHB3 | 0.975 | 0.208 | 0.096 | 2.72e-18 | 9.17e-14 |
| Malignant epithelial | C2 WNT ECs | ABHD17B | 0.975 | 0.172 | 0.08 | 1.06e-14 | 3.58e-10 |
| Malignant epithelial | C2 WNT ECs | RELL2 | 0.975 | 0.1 | 0.047 | 6e-09 | 0.000202 |
| Malignant epithelial | C2 WNT ECs | TOM1L1 | 0.974 | 0.359 | 0.155 | 1.7e-36 | 5.740000000000001e-32 |
| Malignant epithelial | C2 WNT ECs | TTC38 | 0.974 | 0.184 | 0.073 | 9.91e-22 | 3.34e-17 |
| Malignant epithelial | C2 WNT ECs | VPS36 | 0.973 | 0.457 | 0.224 | 6.79e-39 | 2.29e-34 |
| Malignant epithelial | C2 WNT ECs | AKTIP | 0.973 | 0.296 | 0.124 | 3.73e-32 | 1.26e-27 |
| Malignant epithelial | C2 WNT ECs | ZMIZ2 | 0.972 | 0.355 | 0.167 | 6.130000000000001e-31 | 2.07e-26 |
| Malignant epithelial | C2 WNT ECs | CYB5R1 | 0.972 | 0.23 | 0.105 | 3.38e-20 | 1.14e-15 |
| Malignant epithelial | C2 WNT ECs | LRP6 | 0.972 | 0.141 | 0.061 | 3.14e-14 | 1.06e-09 |
| Malignant epithelial | C2 WNT ECs | ARFGAP1 | 0.971 | 0.326 | 0.151 | 1.08e-28 | 3.63e-24 |
| Malignant epithelial | C2 WNT ECs | REST | 0.971 | 0.298 | 0.152 | 1.61e-21 | 5.44e-17 |
| Malignant epithelial | C2 WNT ECs | ATP6V1B2 | 0.971 | 0.191 | 0.078 | 1.67e-21 | 5.62e-17 |
| Malignant epithelial | C2 WNT ECs | CPOX | 0.971 | 0.152 | 0.069 | 7.34e-14 | 2.47e-09 |
| Malignant epithelial | C2 WNT ECs | TRIT1 | 0.97 | 0.1 | 0.046 | 2.34e-09 | 7.9e-05 |
| Malignant epithelial | C2 WNT ECs | CTR9 | 0.969 | 0.311 | 0.144 | 4.98e-27 | 1.68e-22 |
| Malignant epithelial | C2 WNT ECs | LINC00543 | 0.969 | 0.101 | 0.039 | 4.17e-13 | 1.41e-08 |
| Malignant epithelial | C2 WNT ECs | ECHDC2 | 0.968 | 0.28 | 0.12 | 2.85e-28 | 9.589999999999999e-24 |
| Malignant epithelial | C2 WNT ECs | GNB1 | 0.967 | 0.512 | 0.29 | 2.24e-33 | 7.560000000000001e-29 |
| Malignant epithelial | C2 WNT ECs | TPRA1 | 0.967 | 0.329 | 0.158 | 1.71e-27 | 5.759999999999999e-23 |
| Malignant epithelial | C2 WNT ECs | CORO7 | 0.967 | 0.171 | 0.07 | 1.16e-18 | 3.91e-14 |
| Malignant epithelial | C2 WNT ECs | APMAP | 0.967 | 0.507 | 0.384 | 1.03e-14 | 3.48e-10 |
| Malignant epithelial | C2 WNT ECs | GDA | 0.966 | 0.116 | 0.047 | 1.31e-13 | 4.42e-09 |
| Malignant epithelial | C2 WNT ECs | RTCA | 0.965 | 0.325 | 0.15 | 1.59e-28 | 5.35e-24 |
| Malignant epithelial | C2 WNT ECs | RBPMS | 0.965 | 0.185 | 0.072 | 1.92e-22 | 6.45e-18 |
| Malignant epithelial | C2 WNT ECs | FAM107B | 0.965 | 0.175 | 0.1 | 2.04e-09 | 6.86e-05 |
| Malignant epithelial | C2 WNT ECs | CCND1 | 0.964 | 0.819 | 0.598 | 4.31e-36 | 1.45e-31 |
| Malignant epithelial | C2 WNT ECs | PISD | 0.964 | 0.263 | 0.123 | 4.04e-22 | 1.36e-17 |
| Malignant epithelial | C2 WNT ECs | ANAPC4 | 0.964 | 0.12 | 0.044 | 5.4e-17 | 1.82e-12 |
| Malignant epithelial | C2 WNT ECs | JMY | 0.964 | 0.176 | 0.085 | 5.93e-14 | 2e-09 |
| Malignant epithelial | C2 WNT ECs | TMEM41B | 0.963 | 0.437 | 0.22 | 1.39e-33 | 4.68e-29 |
| Malignant epithelial | C2 WNT ECs | GPN2 | 0.963 | 0.16 | 0.067 | 8.2e-17 | 2.76e-12 |
| Malignant epithelial | C2 WNT ECs | TXNL4B | 0.963 | 0.153 | 0.073 | 1.42e-12 | 4.78e-08 |
| Malignant epithelial | C2 WNT ECs | DNAJB2 | 0.962 | 0.497 | 0.251 | 1.07e-39 | 3.6e-35 |
| Malignant epithelial | C2 WNT ECs | MVD | 0.962 | 0.401 | 0.218 | 1.44e-25 | 4.84e-21 |
| Malignant epithelial | C2 WNT ECs | TBC1D23 | 0.962 | 0.173 | 0.073 | 1.51e-18 | 5.08e-14 |
| Malignant epithelial | C2 WNT ECs | ARHGEF12 | 0.961 | 0.317 | 0.158 | 9.27e-24 | 3.12e-19 |
| Malignant epithelial | C2 WNT ECs | DOCK9 | 0.961 | 0.138 | 0.075 | 3e-08 | 0.00101 |
| Malignant epithelial | C2 WNT ECs | LEPROT | 0.96 | 0.528 | 0.276 | 1.06e-39 | 3.58e-35 |
| Malignant epithelial | C2 WNT ECs | RNF24 | 0.96 | 0.169 | 0.087 | 5.38e-12 | 1.81e-07 |
| Malignant epithelial | C2 WNT ECs | PRPF4B | 0.958 | 0.44 | 0.203 | 4.24e-41 | 1.43e-36 |
| Malignant epithelial | C2 WNT ECs | YTHDC1 | 0.958 | 0.486 | 0.252 | 2.4e-37 | 8.09e-33 |
| Malignant epithelial | C2 WNT ECs | FURIN | 0.958 | 0.217 | 0.082 | 1.73e-27 | 5.83e-23 |
| Malignant epithelial | C2 WNT ECs | ESRP2 | 0.958 | 0.249 | 0.107 | 3.6e-25 | 1.21e-20 |
| Malignant epithelial | C2 WNT ECs | PRDX1 | 0.957 | 0.947 | 0.802 | 2.18e-106 | 7.35e-102 |
| Malignant epithelial | C2 WNT ECs | ABHD5 | 0.957 | 0.367 | 0.185 | 1.88e-26 | 6.34e-22 |
| Malignant epithelial | C2 WNT ECs | USH1C | 0.956 | 0.674 | 0.364 | 3.06e-54 | 1.03e-49 |
| Malignant epithelial | C2 WNT ECs | RAF1 | 0.956 | 0.299 | 0.139 | 3.36e-26 | 1.13e-21 |
| Malignant epithelial | C2 WNT ECs | ALDH18A1 | 0.956 | 0.263 | 0.12 | 1.58e-23 | 5.31e-19 |
| Malignant epithelial | C2 WNT ECs | ALDOC | 0.955 | 0.249 | 0.14 | 1.31e-13 | 4.4e-09 |
| Malignant epithelial | C2 WNT ECs | DIS3 | 0.954 | 0.279 | 0.119 | 1.17e-28 | 3.93e-24 |
| Malignant epithelial | C2 WNT ECs | MKRN2 | 0.954 | 0.145 | 0.063 | 1.65e-14 | 5.56e-10 |
| Malignant epithelial | C2 WNT ECs | GMDS-AS1 | 0.954 | 0.106 | 0.044 | 6.69e-12 | 2.25e-07 |
| Malignant epithelial | C2 WNT ECs | TFDP2 | 0.953 | 0.368 | 0.172 | 7.83e-33 | 2.64e-28 |
| Malignant epithelial | C2 WNT ECs | CCDC174 | 0.953 | 0.31 | 0.141 | 7.87e-28 | 2.65e-23 |
| Malignant epithelial | C2 WNT ECs | LARP4B | 0.953 | 0.252 | 0.129 | 3.5e-17 | 1.18e-12 |
| Malignant epithelial | C2 WNT ECs | TAF1C | 0.953 | 0.153 | 0.075 | 7.4e-12 | 2.49e-07 |
| Malignant epithelial | C2 WNT ECs | ARID1A | 0.952 | 0.272 | 0.128 | 4.21e-23 | 1.42e-18 |
| Malignant epithelial | C2 WNT ECs | EPS8L1 | 0.952 | 0.452 | 0.286 | 1.11e-19 | 3.76e-15 |
| Malignant epithelial | C2 WNT ECs | TBC1D14 | 0.952 | 0.12 | 0.056 | 6.78e-11 | 2.28e-06 |
| Malignant epithelial | C2 WNT ECs | HDAC1 | 0.951 | 0.589 | 0.315 | 1.69e-45 | 5.699999999999999e-41 |
| Malignant epithelial | C2 WNT ECs | YIPF1 | 0.951 | 0.181 | 0.077 | 7.23e-19 | 2.43e-14 |
| Malignant epithelial | C2 WNT ECs | C7orf43 | 0.95 | 0.11 | 0.051 | 6.99e-10 | 2.35e-05 |

| compartment | subcluster | gene | avg_log2FC | pct.1 | pct.2 | p_val | p_val_adj |
|---|---|---|---|---|---|---|---|
| Malignant epithelial | C2 WNT ECs | DDAH2 | 0.949 | 0.865 | 0.602 | 9.15e-67 | 3.08e-62 |
| Malignant epithelial | C2 WNT ECs | NUDC | 0.949 | 0.724 | 0.434 | 1.99e-53 | 6.7e-49 |
| Malignant epithelial | C2 WNT ECs | SCML1 | 0.949 | 0.317 | 0.15 | 7.11e-27 | 2.4e-22 |
| Malignant epithelial | C2 WNT ECs | NCBP3 | 0.949 | 0.225 | 0.098 | 3.42e-22 | 1.15e-17 |
| Malignant epithelial | C2 WNT ECs | SYK | 0.949 | 0.164 | 0.076 | 1.13e-14 | 3.82e-10 |
| Malignant epithelial | C2 WNT ECs | RPS26 | 0.948 | 0.976 | 0.968 | 7.41e-50 | 2.5e-45 |
| Malignant epithelial | C2 WNT ECs | SBNO1 | 0.948 | 0.327 | 0.165 | 8.55e-25 | 2.88e-20 |
| Malignant epithelial | C2 WNT ECs | NR1D2 | 0.948 | 0.246 | 0.111 | 1.55e-22 | 5.21e-18 |
| Malignant epithelial | C2 WNT ECs | PRCP | 0.948 | 0.184 | 0.095 | 9.36e-13 | 3.15e-08 |
| Malignant epithelial | C2 WNT ECs | GNPTG | 0.947 | 0.33 | 0.15 | 1.64e-30 | 5.51e-26 |
| Malignant epithelial | C2 WNT ECs | IMPA1 | 0.947 | 0.253 | 0.107 | 2.8e-26 | 9.43e-22 |
| Malignant epithelial | C2 WNT ECs | LDLRAP1 | 0.947 | 0.138 | 0.057 | 1.6e-15 | 5.39e-11 |
| Malignant epithelial | C2 WNT ECs | PDE4D | 0.946 | 0.13 | 0.057 | 6.53e-13 | 2.2e-08 |
| Malignant epithelial | C2 WNT ECs | TARS | 0.945 | 0.483 | 0.239 | 5.52e-40 | 1.86e-35 |
| Malignant epithelial | C2 WNT ECs | REEP6 | 0.945 | 0.317 | 0.137 | 8.830000000000001e-32 | 2.98e-27 |
| Malignant epithelial | C2 WNT ECs | ADCK5 | 0.945 | 0.271 | 0.126 | 3.769999999999999e-23 | 1.27e-18 |
| Malignant epithelial | C2 WNT ECs | GNG12 | 0.944 | 0.378 | 0.179 | 2.42e-32 | 8.15e-28 |
| Malignant epithelial | C2 WNT ECs | SF3A3 | 0.943 | 0.296 | 0.136 | 4.19e-26 | 1.41e-21 |
| Malignant epithelial | C2 WNT ECs | CHMP4B | 0.942 | 0.957 | 0.767 | 3.22e-112 | 1.08e-107 |
| Malignant epithelial | C2 WNT ECs | CTBP1 | 0.942 | 0.39 | 0.199 | 1.92e-28 | 6.47e-24 |
| Malignant epithelial | C2 WNT ECs | FIP1L1 | 0.942 | 0.29 | 0.136 | 2.37e-24 | 7.97e-20 |
| Malignant epithelial | C2 WNT ECs | PHYH | 0.941 | 0.384 | 0.179 | 1.34e-33 | 4.51e-29 |
| Malignant epithelial | C2 WNT ECs | MAP7D1 | 0.941 | 0.398 | 0.202 | 9.57e-30 | 3.22e-25 |
| Malignant epithelial | C2 WNT ECs | ETNK1 | 0.941 | 0.417 | 0.224 | 7.79e-27 | 2.63e-22 |
| Malignant epithelial | C2 WNT ECs | TRAF6 | 0.941 | 0.103 | 0.044 | 7.9e-11 | 2.66e-06 |
| Malignant epithelial | C2 WNT ECs | DTNBP1 | 0.94 | 0.246 | 0.097 | 8.13e-29 | 2.74e-24 |
| Malignant epithelial | C2 WNT ECs | ZKSCAN1 | 0.939 | 0.53 | 0.28 | 6.559999999999999e-40 | 2.21e-35 |
| Malignant epithelial | C2 WNT ECs | SPPL2B | 0.939 | 0.357 | 0.169 | 6.23e-30 | 2.1e-25 |
| Malignant epithelial | C2 WNT ECs | SLC35E2B | 0.939 | 0.203 | 0.091 | 5.47e-19 | 1.84e-14 |
| Malignant epithelial | C2 WNT ECs | TSPYL2 | 0.939 | 0.268 | 0.14 | 2.03e-18 | 6.84e-14 |
| Malignant epithelial | C2 WNT ECs | RP2 | 0.939 | 0.101 | 0.052 | 1.67e-07 | 0.00564 |
| Malignant epithelial | C2 WNT ECs | MAPK3 | 0.938 | 0.575 | 0.307 | 4.69e-43 | 1.58e-38 |
| Malignant epithelial | C2 WNT ECs | TMCC1 | 0.938 | 0.203 | 0.084 | 7.19e-22 | 2.42e-17 |
| Malignant epithelial | C2 WNT ECs | PIP4K2C | 0.938 | 0.208 | 0.099 | 1.14e-17 | 3.86e-13 |
| Malignant epithelial | C2 WNT ECs | RAD54L2 | 0.938 | 0.168 | 0.072 | 4.11e-17 | 1.38e-12 |
| Malignant epithelial | C2 WNT ECs | TULP4 | 0.937 | 0.238 | 0.108 | 8.300000000000001e-22 | 2.8e-17 |
| Malignant epithelial | C2 WNT ECs | HILPDA | 0.937 | 0.273 | 0.166 | 1.12e-12 | 3.78e-08 |
| Malignant epithelial | C2 WNT ECs | HDAC2 | 0.936 | 0.743 | 0.45 | 9.87e-57 | 3.32e-52 |
| Malignant epithelial | C2 WNT ECs | APPL2 | 0.936 | 0.241 | 0.1 | 2.13e-26 | 7.19e-22 |
| Malignant epithelial | C2 WNT ECs | ALKBH5 | 0.936 | 0.254 | 0.128 | 1.16e-18 | 3.91e-14 |
| Malignant epithelial | C2 WNT ECs | PBX4 | 0.936 | 0.131 | 0.053 | 3.74e-15 | 1.26e-10 |
| Malignant epithelial | C2 WNT ECs | LATS1 | 0.936 | 0.122 | 0.05 | 1.05e-13 | 3.54e-09 |
| Malignant epithelial | C2 WNT ECs | KIF1C | 0.935 | 0.302 | 0.134 | 4.29e-29 | 1.45e-24 |
| Malignant epithelial | C2 WNT ECs | LINC00869 | 0.935 | 0.382 | 0.198 | 1.16e-27 | 3.909999999999999e-23 |
| Malignant epithelial | C2 WNT ECs | INO80C | 0.935 | 0.425 | 0.268 | 2.24e-19 | 7.56e-15 |
| Malignant epithelial | C2 WNT ECs | MED24 | 0.935 | 0.218 | 0.103 | 9.23e-18 | 3.11e-13 |
| Malignant epithelial | C2 WNT ECs | IP6K2 | 0.934 | 0.71 | 0.431 | 2.38e-49 | 8.01e-45 |
| Malignant epithelial | C2 WNT ECs | EZR | 0.934 | 0.893 | 0.707 | 9.16e-44 | 3.09e-39 |
| Malignant epithelial | C2 WNT ECs | HTT | 0.934 | 0.142 | 0.068 | 8.34e-12 | 2.81e-07 |
| Malignant epithelial | C2 WNT ECs | PIGT | 0.933 | 0.656 | 0.409 | 3.36e-40 | 1.13e-35 |
| Malignant epithelial | C2 WNT ECs | USP22 | 0.932 | 0.303 | 0.148 | 9.69e-24 | 3.26e-19 |
| Malignant epithelial | C2 WNT ECs | ATP11A | 0.932 | 0.249 | 0.113 | 1.37e-22 | 4.63e-18 |
| Malignant epithelial | C2 WNT ECs | C1GALT1C1 | 0.932 | 0.342 | 0.188 | 2.85e-21 | 9.61e-17 |
| Malignant epithelial | C2 WNT ECs | CLIC4 | 0.932 | 0.183 | 0.095 | 2.38e-12 | 8.03e-08 |
| Malignant epithelial | C2 WNT ECs | NCLN | 0.931 | 0.39 | 0.203 | 5.74e-29 | 1.93e-24 |
| Malignant epithelial | C2 WNT ECs | AFTPH | 0.931 | 0.309 | 0.143 | 1.01e-27 | 3.42e-23 |
| Malignant epithelial | C2 WNT ECs | AMFR | 0.931 | 0.269 | 0.119 | 7.050000000000001e-26 | 2.38e-21 |
| Malignant epithelial | C2 WNT ECs | DMAP1 | 0.931 | 0.21 | 0.092 | 3.55e-20 | 1.19e-15 |
| Malignant epithelial | C2 WNT ECs | HOXB8 | 0.93 | 0.499 | 0.284 | 1.64e-29 | 5.51e-25 |
| Malignant epithelial | C2 WNT ECs | MBOAT2 | 0.93 | 0.361 | 0.196 | 7.91e-24 | 2.67e-19 |
| Malignant epithelial | C2 WNT ECs | ATG9A | 0.93 | 0.223 | 0.095 | 5.38e-23 | 1.81e-18 |
| Malignant epithelial | C2 WNT ECs | TBC1D7 | 0.93 | 0.203 | 0.085 | 1.21e-21 | 4.06e-17 |
| Malignant epithelial | C2 WNT ECs | XRRA1 | 0.93 | 0.139 | 0.054 | 5.32e-17 | 1.79e-12 |
| Malignant epithelial | C2 WNT ECs | ABCD3 | 0.93 | 0.187 | 0.085 | 1.02e-16 | 3.45e-12 |

| compartment | subcluster | gene | avg_log2FC | pct.1 | pct.2 | p_val | p_val_adj |
|---|---|---|---|---|---|---|---|
| Malignant epithelial | C2 WNT ECs | GABPA | 0.93 | 0.124 | 0.052 | 1.75e-13 | 5.91e-09 |
| Malignant epithelial | C2 WNT ECs | EXOSC8 | 0.929 | 0.521 | 0.285 | 3.26e-36 | 1.1e-31 |
| Malignant epithelial | C2 WNT ECs | MAPK14 | 0.929 | 0.33 | 0.156 | 2.67e-27 | 8.979999999999999e-23 |
| Malignant epithelial | C2 WNT ECs | CERK | 0.929 | 0.11 | 0.05 | 2.57e-10 | 8.65e-06 |
| Malignant epithelial | C2 WNT ECs | VTA1 | 0.928 | 0.269 | 0.126 | 1.73e-23 | 5.83e-19 |
| Malignant epithelial | C2 WNT ECs | DGKD | 0.928 | 0.131 | 0.059 | 1.68e-12 | 5.65e-08 |
| Malignant epithelial | C2 WNT ECs | OARD1 | 0.927 | 0.553 | 0.309 | 1.73e-38 | 5.84e-34 |
| Malignant epithelial | C2 WNT ECs | TMED5 | 0.927 | 0.268 | 0.125 | 9.26e-23 | 3.12e-18 |
| Malignant epithelial | C2 WNT ECs | NOSIP | 0.926 | 0.755 | 0.501 | 8.47e-50 | 2.85e-45 |
| Malignant epithelial | C2 WNT ECs | TGFBR2 | 0.926 | 0.388 | 0.194 | 6.22e-30 | 2.1e-25 |
| Malignant epithelial | C2 WNT ECs | ARHGEF16 | 0.926 | 0.287 | 0.128 | 4.850000000000001e-27 | 1.64e-22 |
| Malignant epithelial | C2 WNT ECs | COMMD6 | 0.925 | 0.959 | 0.877 | 4.960000000000001e-118 | 1.67e-113 |
| Malignant epithelial | C2 WNT ECs | RAB5A | 0.925 | 0.497 | 0.244 | 6.81e-42 | 2.29e-37 |
| Malignant epithelial | C2 WNT ECs | SIAH1 | 0.925 | 0.346 | 0.181 | 1.68e-23 | 5.67e-19 |
| Malignant epithelial | C2 WNT ECs | ACY1 | 0.925 | 0.129 | 0.05 | 4.68e-16 | 1.58e-11 |
| Malignant epithelial | C2 WNT ECs | ACER3 | 0.925 | 0.147 | 0.066 | 5.76e-14 | 1.94e-09 |
| Malignant epithelial | C2 WNT ECs | MAFG | 0.924 | 0.342 | 0.168 | 2.37e-26 | 7.99e-22 |
| Malignant epithelial | C2 WNT ECs | HIST1H2BH | 0.924 | 0.168 | 0.064 | 2e-21 | 6.73e-17 |
| Malignant epithelial | C2 WNT ECs | SSBP3 | 0.923 | 0.237 | 0.112 | 9.43e-20 | 3.18e-15 |
| Malignant epithelial | C2 WNT ECs | GMEB1 | 0.923 | 0.104 | 0.044 | 3.74e-11 | 1.26e-06 |
| Malignant epithelial | C2 WNT ECs | PCGF3 | 0.921 | 0.181 | 0.082 | 1.48e-16 | 5e-12 |
| Malignant epithelial | C2 WNT ECs | FAM83F | 0.921 | 0.142 | 0.062 | 5.37e-14 | 1.81e-09 |
| Malignant epithelial | C2 WNT ECs | SLC12A6 | 0.92 | 0.133 | 0.058 | 1.54e-13 | 5.18e-09 |
| Malignant epithelial | C2 WNT ECs | SND1 | 0.919 | 0.548 | 0.305 | 1.56e-37 | 5.25e-33 |
| Malignant epithelial | C2 WNT ECs | AHCYL1 | 0.919 | 0.299 | 0.14 | 5.02e-26 | 1.69e-21 |
| Malignant epithelial | C2 WNT ECs | PGM3 | 0.919 | 0.207 | 0.084 | 3.789999999999999e-23 | 1.28e-18 |
| Malignant epithelial | C2 WNT ECs | NLN | 0.919 | 0.214 | 0.09 | 3.65e-22 | 1.23e-17 |
| Malignant epithelial | C2 WNT ECs | CTTNBP2NL | 0.919 | 0.191 | 0.086 | 1.72e-17 | 5.81e-13 |
| Malignant epithelial | C2 WNT ECs | H2AFY | 0.918 | 0.886 | 0.64 | 1.78e-68 | 5.99e-64 |
| Malignant epithelial | C2 WNT ECs | ACLY | 0.918 | 0.302 | 0.141 | 1.13e-26 | 3.8e-22 |
| Malignant epithelial | C2 WNT ECs | HSP90AB1 | 0.917 | 0.992 | 0.962 | 1.18e-107 | 3.97e-103 |
| Malignant epithelial | C2 WNT ECs | AKAP9 | 0.917 | 0.717 | 0.441 | 1.37e-47 | 4.61e-43 |
| Malignant epithelial | C2 WNT ECs | PRKAB1 | 0.917 | 0.261 | 0.105 | 2.47e-29 | 8.34e-25 |
| Malignant epithelial | C2 WNT ECs | SREBF2 | 0.917 | 0.291 | 0.135 | 6.94e-26 | 2.34e-21 |
| Malignant epithelial | C2 WNT ECs | SGPL1 | 0.917 | 0.302 | 0.143 | 7.53e-25 | 2.54e-20 |
| Malignant epithelial | C2 WNT ECs | MAEA | 0.916 | 0.361 | 0.168 | 1.82e-31 | 6.12e-27 |
| Malignant epithelial | C2 WNT ECs | HYOU1 | 0.916 | 0.256 | 0.119 | 5.23e-22 | 1.76e-17 |
| Malignant epithelial | C2 WNT ECs | KIAA0232 | 0.916 | 0.181 | 0.088 | 3.87e-14 | 1.31e-09 |
| Malignant epithelial | C2 WNT ECs | GMIP | 0.916 | 0.119 | 0.049 | 2.71e-13 | 9.12e-09 |
| Malignant epithelial | C2 WNT ECs | ZCCHC11 | 0.916 | 0.168 | 0.084 | 2.84e-12 | 9.56e-08 |
| Malignant epithelial | C2 WNT ECs | ARHGAP21 | 0.915 | 0.288 | 0.141 | 3.56e-22 | 1.2e-17 |
| Malignant epithelial | C2 WNT ECs | DUSP16 | 0.915 | 0.145 | 0.063 | 2.1e-14 | 7.07e-10 |
| Malignant epithelial | C2 WNT ECs | ELF4 | 0.915 | 0.104 | 0.048 | 1.14e-09 | 3.83e-05 |
| Malignant epithelial | C2 WNT ECs | MYL12B | 0.913 | 0.972 | 0.88 | 6.1e-126 | 2.05e-121 |
| Malignant epithelial | C2 WNT ECs | VPS4B | 0.913 | 0.419 | 0.213 | 2.14e-31 | 7.2e-27 |
| Malignant epithelial | C2 WNT ECs | TMEM14C | 0.912 | 0.848 | 0.58 | 2.55e-72 | 8.58e-68 |
| Malignant epithelial | C2 WNT ECs | DUSP2 | 0.912 | 0.455 | 0.311 | 4.66e-14 | 1.57e-09 |
| Malignant epithelial | C2 WNT ECs | ZNF212 | 0.912 | 0.129 | 0.062 | 1.77e-10 | 5.95e-06 |
| Malignant epithelial | C2 WNT ECs | SELM | 0.911 | 0.432 | 0.281 | 1.05e-19 | 3.54e-15 |
| Malignant epithelial | C2 WNT ECs | PDIA5 | 0.911 | 0.192 | 0.086 | 5.39e-18 | 1.82e-13 |
| Malignant epithelial | C2 WNT ECs | A1CF | 0.911 | 0.154 | 0.068 | 6.81e-15 | 2.3e-10 |
| Malignant epithelial | C2 WNT ECs | CTSB | 0.91 | 0.824 | 0.572 | 5.88e-57 | 1.98e-52 |
| Malignant epithelial | C2 WNT ECs | CASC3 | 0.91 | 0.284 | 0.138 | 1.69e-22 | 5.69e-18 |
| Malignant epithelial | C2 WNT ECs | GAN | 0.91 | 0.231 | 0.119 | 9.54e-16 | 3.21e-11 |
| Malignant epithelial | C2 WNT ECs | CCDC50 | 0.909 | 0.497 | 0.257 | 1.93e-37 | 6.52e-33 |
| Malignant epithelial | C2 WNT ECs | SH3BP1 | 0.909 | 0.175 | 0.08 | 8.88e-16 | 2.99e-11 |
| Malignant epithelial | C2 WNT ECs | FDFT1 | 0.908 | 0.513 | 0.277 | 3.04e-36 | 1.03e-31 |
| Malignant epithelial | C2 WNT ECs | CDKN1B | 0.908 | 0.315 | 0.174 | 8.97e-19 | 3.02e-14 |
| Malignant epithelial | C2 WNT ECs | MIB1 | 0.908 | 0.115 | 0.048 | 1.05e-12 | 3.53e-08 |
| Malignant epithelial | C2 WNT ECs | LUC7L | 0.907 | 0.365 | 0.167 | 1.13e-33 | 3.8e-29 |
| Malignant epithelial | C2 WNT ECs | KDM5B | 0.907 | 0.28 | 0.134 | 1.89e-22 | 6.36e-18 |
| Malignant epithelial | C2 WNT ECs | UGGT2 | 0.907 | 0.175 | 0.078 | 2.69e-16 | 9.07e-12 |
| Malignant epithelial | C2 WNT ECs | GTPBP2 | 0.907 | 0.127 | 0.051 | 6.9e-15 | 2.32e-10 |
| Malignant epithelial | C2 WNT ECs | CD151 | 0.906 | 0.934 | 0.691 | 6.11e-83 | 2.06e-78 |

| compartment | subcluster | gene | avg_log2FC | pct.1 | pct.2 | p_val | p_val_adj |
|---|---|---|---|---|---|---|---|
| Malignant epithelial | C2 WNT ECs | CDIPT | 0.906 | 0.549 | 0.291 | 9.769999999999999e-40 | 3.29e-35 |
| Malignant epithelial | C2 WNT ECs | MRPS25 | 0.906 | 0.471 | 0.243 | 2.94e-35 | 9.91e-31 |
| Malignant epithelial | C2 WNT ECs | C22orf29 | 0.906 | 0.154 | 0.068 | 7.95e-15 | 2.68e-10 |
| Malignant epithelial | C2 WNT ECs | RUFY3 | 0.906 | 0.104 | 0.049 | 3.14e-09 | 0.000106 |
| Malignant epithelial | C2 WNT ECs | CACYBP | 0.905 | 0.756 | 0.527 | 6.27e-37 | 2.11e-32 |
| Malignant epithelial | C2 WNT ECs | ZBED1 | 0.905 | 0.177 | 0.084 | 1.06e-14 | 3.58e-10 |
| Malignant epithelial | C2 WNT ECs | TERF2IP | 0.904 | 0.562 | 0.306 | 1.08e-40 | 3.64e-36 |
| Malignant epithelial | C2 WNT ECs | USB1 | 0.904 | 0.287 | 0.126 | 2.77e-28 | 9.329999999999999e-24 |
| Malignant epithelial | C2 WNT ECs | PSMD14 | 0.903 | 0.564 | 0.307 | 1.4e-39 | 4.71e-35 |
| Malignant epithelial | C2 WNT ECs | KRAS | 0.903 | 0.428 | 0.255 | 8.61e-20 | 2.9e-15 |
| Malignant epithelial | C2 WNT ECs | CSRNP2 | 0.903 | 0.112 | 0.054 | 2.04e-09 | 6.88e-05 |
| Malignant epithelial | C2 WNT ECs | TRIM16 | 0.902 | 0.299 | 0.143 | 3.9e-24 | 1.31e-19 |
| Malignant epithelial | C2 WNT ECs | EGR1 | 0.902 | 0.846 | 0.752 | 2.53e-21 | 8.53e-17 |
| Malignant epithelial | C2 WNT ECs | IQCB1 | 0.902 | 0.161 | 0.073 | 8.22e-15 | 2.77e-10 |
| Malignant epithelial | C2 WNT ECs | ACAD8 | 0.902 | 0.123 | 0.053 | 1.32e-12 | 4.45e-08 |
| Malignant epithelial | C2 WNT ECs | PAIP1 | 0.899 | 0.658 | 0.393 | 1.1e-42 | 3.72e-38 |
| Malignant epithelial | C2 WNT ECs | KIN | 0.899 | 0.268 | 0.116 | 7.410000000000001e-26 | 2.5e-21 |
| Malignant epithelial | C2 WNT ECs | SLC15A4 | 0.899 | 0.124 | 0.059 | 8.96e-11 | 3.02e-06 |
| Malignant epithelial | C2 WNT ECs | EGR3 | 0.899 | 0.1 | 0.052 | 2.42e-07 | 0.00816 |
| Malignant epithelial | C2 WNT ECs | CHUK | 0.898 | 0.11 | 0.052 | 1.09e-09 | 3.68e-05 |
| Malignant epithelial | C2 WNT ECs | CHIC2 | 0.897 | 0.276 | 0.133 | 1.02e-21 | 3.45e-17 |
| Malignant epithelial | C2 WNT ECs | COPB2 | 0.896 | 0.418 | 0.197 | 4.44e-37 | 1.5e-32 |
| Malignant epithelial | C2 WNT ECs | BCL10 | 0.896 | 0.346 | 0.172 | 1.54e-26 | 5.2e-22 |
| Malignant epithelial | C2 WNT ECs | PIK3IP1 | 0.896 | 0.35 | 0.192 | 4.82e-21 | 1.62e-16 |
| Malignant epithelial | C2 WNT ECs | YES1 | 0.896 | 0.196 | 0.091 | 8.21e-17 | 2.77e-12 |
| Malignant epithelial | C2 WNT ECs | SIRT6 | 0.895 | 0.333 | 0.168 | 1.4e-24 | 4.71e-20 |
| Malignant epithelial | C2 WNT ECs | VPS51 | 0.894 | 0.608 | 0.338 | 3.97e-43 | 1.34e-38 |
| Malignant epithelial | C2 WNT ECs | GCLC | 0.893 | 0.318 | 0.151 | 3.3e-27 | 1.11e-22 |
| Malignant epithelial | C2 WNT ECs | VPS11 | 0.893 | 0.126 | 0.054 | 3e-13 | 1.01e-08 |
| Malignant epithelial | C2 WNT ECs | HSP90AA1 | 0.892 | 0.997 | 0.968 | 4.16e-50 | 1.4e-45 |
| Malignant epithelial | C2 WNT ECs | SLC37A3 | 0.892 | 0.279 | 0.136 | 2.7e-21 | 9.09e-17 |
| Malignant epithelial | C2 WNT ECs | ZNF770 | 0.891 | 0.276 | 0.149 | 1.27e-17 | 4.28e-13 |
| Malignant epithelial | C2 WNT ECs | KIAA0430 | 0.891 | 0.175 | 0.081 | 5.54e-15 | 1.87e-10 |
| Malignant epithelial | C2 WNT ECs | CHD1L | 0.891 | 0.176 | 0.082 | 1.02e-14 | 3.44e-10 |
| Malignant epithelial | C2 WNT ECs | CHMP4C | 0.89 | 0.581 | 0.331 | 4.03e-38 | 1.36e-33 |
| Malignant epithelial | C2 WNT ECs | TMEM8A | 0.89 | 0.275 | 0.124 | 1.88e-24 | 6.32e-20 |
| Malignant epithelial | C2 WNT ECs | PHC2 | 0.89 | 0.221 | 0.104 | 3.01e-18 | 1.01e-13 |
| Malignant epithelial | C2 WNT ECs | MISP | 0.889 | 0.794 | 0.567 | 8.350000000000001e-43 | 2.81e-38 |
| Malignant epithelial | C2 WNT ECs | PAAF1 | 0.889 | 0.212 | 0.088 | 1.02e-22 | 3.43e-18 |
| Malignant epithelial | C2 WNT ECs | RNF185 | 0.889 | 0.127 | 0.054 | 7.78e-14 | 2.62e-09 |
| Malignant epithelial | C2 WNT ECs | HMGB1 | 0.888 | 0.992 | 0.926 | 2.26e-93 | 7.6e-89 |
| Malignant epithelial | C2 WNT ECs | SLC41A3 | 0.888 | 0.326 | 0.148 | 2.95e-29 | 9.94e-25 |
| Malignant epithelial | C2 WNT ECs | RASA2 | 0.888 | 0.229 | 0.111 | 5.1e-18 | 1.72e-13 |
| Malignant epithelial | C2 WNT ECs | PHACTR4 | 0.888 | 0.234 | 0.118 | 1.2e-16 | 4.06e-12 |
| Malignant epithelial | C2 WNT ECs | ABCA1 | 0.888 | 0.119 | 0.053 | 1.06e-11 | 3.56e-07 |
| Malignant epithelial | C2 WNT ECs | GLCE | 0.886 | 0.304 | 0.146 | 1.97e-24 | 6.64e-20 |
| Malignant epithelial | C2 WNT ECs | UBR2 | 0.886 | 0.225 | 0.105 | 4.87e-19 | 1.64e-14 |
| Malignant epithelial | C2 WNT ECs | PEA15 | 0.886 | 0.286 | 0.153 | 1.41e-17 | 4.75e-13 |
| Malignant epithelial | C2 WNT ECs | TSG101 | 0.885 | 0.62 | 0.349 | 3.06e-44 | 1.03e-39 |
| Malignant epithelial | C2 WNT ECs | TTC19 | 0.885 | 0.287 | 0.149 | 1.33e-19 | 4.48e-15 |
| Malignant epithelial | C2 WNT ECs | FBXO32 | 0.885 | 0.372 | 0.218 | 2.17e-19 | 7.31e-15 |
| Malignant epithelial | C2 WNT ECs | ZNF24 | 0.885 | 0.234 | 0.114 | 5.57e-18 | 1.88e-13 |
| Malignant epithelial | C2 WNT ECs | UBE2E1 | 0.884 | 0.51 | 0.263 | 4.82e-39 | 1.62e-34 |
| Malignant epithelial | C2 WNT ECs | KPNA1 | 0.884 | 0.267 | 0.111 | 4.19e-28 | 1.41e-23 |
| Malignant epithelial | C2 WNT ECs | AP5Z1 | 0.884 | 0.273 | 0.126 | 1.69e-23 | 5.71e-19 |
| Malignant epithelial | C2 WNT ECs | GTF2B | 0.884 | 0.3 | 0.149 | 5.88e-23 | 1.98e-18 |
| Malignant epithelial | C2 WNT ECs | CAPN8 | 0.883 | 0.231 | 0.113 | 7.9e-18 | 2.66e-13 |
| Malignant epithelial | C2 WNT ECs | BAIAP2L2 | 0.883 | 0.244 | 0.125 | 1.46e-16 | 4.92e-12 |
| Malignant epithelial | C2 WNT ECs | CAPN1 | 0.882 | 0.746 | 0.475 | 9.470000000000001e-51 | 3.19e-46 |
| Malignant epithelial | C2 WNT ECs | ANO9 | 0.882 | 0.368 | 0.182 | 4.66e-28 | 1.57e-23 |
| Malignant epithelial | C2 WNT ECs | CDC42BPA | 0.882 | 0.29 | 0.149 | 1.63e-20 | 5.48e-16 |
| Malignant epithelial | C2 WNT ECs | RPP30 | 0.881 | 0.303 | 0.145 | 2.59e-25 | 8.720000000000001e-21 |
| Malignant epithelial | C2 WNT ECs | EPPK1 | 0.881 | 0.183 | 0.076 | 1.14e-19 | 3.84e-15 |
| Malignant epithelial | C2 WNT ECs | LMBRD1 | 0.88 | 0.392 | 0.187 | 2.66e-33 | 8.97e-29 |

| compartment | subcluster | gene | avg_log2FC | pct.1 | pct.2 | p_val | p_val_adj |
|---|---|---|---|---|---|---|---|
| Malignant epithelial | C2 WNT ECs | COPG1 | 0.88 | 0.333 | 0.157 | 2.56e-28 | 8.639999999999999e-24 |
| Malignant epithelial | C2 WNT ECs | CD276 | 0.88 | 0.252 | 0.113 | 7.29e-23 | 2.46e-18 |
| Malignant epithelial | C2 WNT ECs | PLCD3 | 0.88 | 0.322 | 0.176 | 3.83e-20 | 1.29e-15 |
| Malignant epithelial | C2 WNT ECs | PODXL2 | 0.88 | 0.234 | 0.119 | 1.02e-16 | 3.45e-12 |
| Malignant epithelial | C2 WNT ECs | UBXN4 | 0.879 | 0.823 | 0.542 | 1.65e-59 | 5.57e-55 |
| Malignant epithelial | C2 WNT ECs | BCAR3 | 0.879 | 0.206 | 0.107 | 1.86e-13 | 6.26e-09 |
| Malignant epithelial | C2 WNT ECs | QARS | 0.877 | 0.628 | 0.377 | 2.11e-40 | 7.100000000000001e-36 |
| Malignant epithelial | C2 WNT ECs | RAP1A | 0.877 | 0.464 | 0.237 | 3.95e-34 | 1.33e-29 |
| Malignant epithelial | C2 WNT ECs | RRAD | 0.877 | 0.16 | 0.099 | 4.55e-07 | 0.0153 |
| Malignant epithelial | C2 WNT ECs | CHD4 | 0.876 | 0.579 | 0.347 | 4.71e-35 | 1.59e-30 |
| Malignant epithelial | C2 WNT ECs | REPIN1 | 0.876 | 0.559 | 0.339 | 7.13e-30 | 2.4e-25 |
| Malignant epithelial | C2 WNT ECs | ATG16L1 | 0.876 | 0.198 | 0.094 | 3.72e-16 | 1.25e-11 |
| Malignant epithelial | C2 WNT ECs | HSD11B2 | 0.875 | 0.28 | 0.14 | 1.71e-20 | 5.77e-16 |
| Malignant epithelial | C2 WNT ECs | AIFM2 | 0.875 | 0.227 | 0.108 | 3.98e-18 | 1.34e-13 |
| Malignant epithelial | C2 WNT ECs | RNF128 | 0.874 | 0.447 | 0.193 | 6.439999999999999e-45 | 2.17e-40 |
| Malignant epithelial | C2 WNT ECs | SEC63 | 0.874 | 0.532 | 0.283 | 2.85e-38 | 9.61e-34 |
| Malignant epithelial | C2 WNT ECs | YARS | 0.874 | 0.295 | 0.136 | 9.33e-26 | 3.14e-21 |
| Malignant epithelial | C2 WNT ECs | UPP1 | 0.874 | 0.463 | 0.282 | 4.12e-21 | 1.39e-16 |
| Malignant epithelial | C2 WNT ECs | NBEAL2 | 0.874 | 0.23 | 0.109 | 3.79e-19 | 1.28e-14 |
| Malignant epithelial | C2 WNT ECs | MRFAP1L1 | 0.874 | 0.208 | 0.1 | 1.8e-16 | 6.06e-12 |
| Malignant epithelial | C2 WNT ECs | VPS4A | 0.873 | 0.474 | 0.23 | 7.34e-38 | 2.47e-33 |
| Malignant epithelial | C2 WNT ECs | PHIP | 0.873 | 0.518 | 0.274 | 1.05e-36 | 3.52e-32 |
| Malignant epithelial | C2 WNT ECs | MT-ND3 | 0.872 | 0.992 | 0.974 | 2.21e-98 | 7.439999999999999e-94 |
| Malignant epithelial | C2 WNT ECs | SPNS1 | 0.872 | 0.436 | 0.228 | 1.1e-31 | 3.7e-27 |
| Malignant epithelial | C2 WNT ECs | MAGEF1 | 0.872 | 0.386 | 0.199 | 1.03e-26 | 3.46e-22 |
| Malignant epithelial | C2 WNT ECs | HOMER3 | 0.872 | 0.217 | 0.1 | 2.59e-18 | 8.74e-14 |
| Malignant epithelial | C2 WNT ECs | RAB15 | 0.872 | 0.122 | 0.046 | 3.8e-16 | 1.28e-11 |
| Malignant epithelial | C2 WNT ECs | HARS2 | 0.872 | 0.146 | 0.069 | 3.18e-12 | 1.07e-07 |
| Malignant epithelial | C2 WNT ECs | LGALS4 | 0.871 | 0.923 | 0.43 | 2.6e-112 | 8.75e-108 |
| Malignant epithelial | C2 WNT ECs | IDH2 | 0.871 | 0.666 | 0.413 | 3.87e-40 | 1.31e-35 |
| Malignant epithelial | C2 WNT ECs | MKRN1 | 0.871 | 0.629 | 0.401 | 5.179999999999999e-34 | 1.75e-29 |
| Malignant epithelial | C2 WNT ECs | DDOST | 0.871 | 0.629 | 0.408 | 9.150000000000001e-32 | 3.08e-27 |
| Malignant epithelial | C2 WNT ECs | THAP9-AS1 | 0.871 | 0.348 | 0.159 | 8.47e-31 | 2.86e-26 |
| Malignant epithelial | C2 WNT ECs | NARS2 | 0.871 | 0.12 | 0.053 | 7.48e-12 | 2.52e-07 |
| Malignant epithelial | C2 WNT ECs | COPG2 | 0.871 | 0.13 | 0.06 | 1.24e-11 | 4.19e-07 |
| Malignant epithelial | C2 WNT ECs | DNAJB6 | 0.87 | 0.877 | 0.648 | 5.54e-56 | 1.87e-51 |
| Malignant epithelial | C2 WNT ECs | INADL | 0.87 | 0.269 | 0.133 | 3.48e-20 | 1.17e-15 |
| Malignant epithelial | C2 WNT ECs | GCH1 | 0.87 | 0.171 | 0.08 | 3.36e-14 | 1.13e-09 |
| Malignant epithelial | C2 WNT ECs | PRKCA | 0.868 | 0.31 | 0.151 | 2.36e-24 | 7.95e-20 |
| Malignant epithelial | C2 WNT ECs | TRAF7 | 0.867 | 0.471 | 0.241 | 2.12e-35 | 7.130000000000001e-31 |
| Malignant epithelial | C2 WNT ECs | RP11-140K17.3 | 0.866 | 0.21 | 0.096 | 2.34e-18 | 7.87e-14 |
| Malignant epithelial | C2 WNT ECs | SIRT5 | 0.866 | 0.175 | 0.075 | 9.63e-18 | 3.25e-13 |
| Malignant epithelial | C2 WNT ECs | PLEKHG3 | 0.866 | 0.222 | 0.115 | 1.42e-14 | 4.8e-10 |
| Malignant epithelial | C2 WNT ECs | SOX13 | 0.866 | 0.119 | 0.055 | 7.27e-11 | 2.45e-06 |
| Malignant epithelial | C2 WNT ECs | SLC25A14 | 0.866 | 0.111 | 0.052 | 8.46e-10 | 2.85e-05 |
| Malignant epithelial | C2 WNT ECs | EIF2B5 | 0.865 | 0.313 | 0.164 | 4.49e-20 | 1.51e-15 |
| Malignant epithelial | C2 WNT ECs | TPTEP1 | 0.865 | 0.189 | 0.086 | 1.93e-17 | 6.5e-13 |
| Malignant epithelial | C2 WNT ECs | PABPC1L | 0.865 | 0.13 | 0.054 | 1.85e-14 | 6.23e-10 |
| Malignant epithelial | C2 WNT ECs | SNX11 | 0.864 | 0.123 | 0.062 | 3.26e-09 | 0.00011 |
| Malignant epithelial | C2 WNT ECs | ST7L | 0.864 | 0.106 | 0.05 | 6.05e-09 | 0.000204 |
| Malignant epithelial | C2 WNT ECs | CDC5L | 0.863 | 0.461 | 0.239 | 4.149999999999999e-34 | 1.4e-29 |
| Malignant epithelial | C2 WNT ECs | MON1B | 0.863 | 0.212 | 0.096 | 5.67e-19 | 1.91e-14 |
| Malignant epithelial | C2 WNT ECs | SCYL2 | 0.863 | 0.23 | 0.115 | 9.52e-17 | 3.21e-12 |
| Malignant epithelial | C2 WNT ECs | RAB1A | 0.862 | 0.813 | 0.569 | 6.89e-47 | 2.32e-42 |
| Malignant epithelial | C2 WNT ECs | BAG6 | 0.862 | 0.541 | 0.29 | 4.2e-38 | 1.41e-33 |
| Malignant epithelial | C2 WNT ECs | TRIM24 | 0.862 | 0.215 | 0.1 | 3.59e-18 | 1.21e-13 |
| Malignant epithelial | C2 WNT ECs | ASPSCR1 | 0.862 | 0.417 | 0.267 | 1.03e-13 | 3.46e-09 |
| Malignant epithelial | C2 WNT ECs | SPEN | 0.861 | 0.276 | 0.12 | 2.38e-27 | 8.02e-23 |
| Malignant epithelial | C2 WNT ECs | CCDC68 | 0.861 | 0.181 | 0.088 | 3.44e-14 | 1.16e-09 |
| Malignant epithelial | C2 WNT ECs | GNL1 | 0.86 | 0.39 | 0.211 | 5.659999999999999e-24 | 1.91e-19 |
| Malignant epithelial | C2 WNT ECs | PPARA | 0.86 | 0.162 | 0.075 | 4.77e-14 | 1.61e-09 |
| Malignant epithelial | C2 WNT ECs | UEVLD | 0.86 | 0.118 | 0.05 | 1.63e-12 | 5.48e-08 |
| Malignant epithelial | C2 WNT ECs | EPB41 | 0.86 | 0.108 | 0.054 | 2.4e-08 | 0.000808 |
| Malignant epithelial | C2 WNT ECs | PBRM1 | 0.859 | 0.288 | 0.138 | 1.47e-23 | 4.96e-19 |

| compartment | subcluster | gene | avg_log2FC | pct.1 | pct.2 | p_val | p_val_adj |
|---|---|---|---|---|---|---|---|
| Malignant epithelial | C2 WNT ECs | OTUD7B | 0.859 | 0.206 | 0.101 | 1.07e-15 | 3.62e-11 |
| Malignant epithelial | C2 WNT ECs | HMBOX1 | 0.859 | 0.122 | 0.055 | 1.77e-11 | 5.96e-07 |
| Malignant epithelial | C2 WNT ECs | ENOX2 | 0.859 | 0.12 | 0.06 | 2.87e-09 | 9.67e-05 |
| Malignant epithelial | C2 WNT ECs | RBBP8NL | 0.859 | 0.122 | 0.062 | 6.08e-09 | 0.000205 |
| Malignant epithelial | C2 WNT ECs | CRBN | 0.858 | 0.327 | 0.17 | 2.15e-22 | 7.23e-18 |
| Malignant epithelial | C2 WNT ECs | MARCH9 | 0.858 | 0.196 | 0.094 | 3.43e-16 | 1.16e-11 |
| Malignant epithelial | C2 WNT ECs | PPP1R16A | 0.857 | 0.562 | 0.302 | 4.05e-39 | 1.36e-34 |
| Malignant epithelial | C2 WNT ECs | RND3 | 0.857 | 0.77 | 0.611 | 2.98e-24 | 1e-19 |
| Malignant epithelial | C2 WNT ECs | RAB40C | 0.857 | 0.281 | 0.138 | 7.77e-22 | 2.62e-17 |
| Malignant epithelial | C2 WNT ECs | YKT6 | 0.856 | 0.409 | 0.213 | 2.07e-28 | 6.96e-24 |
| Malignant epithelial | C2 WNT ECs | OXR1 | 0.856 | 0.394 | 0.207 | 5.12e-27 | 1.73e-22 |
| Malignant epithelial | C2 WNT ECs | FAM122B | 0.856 | 0.294 | 0.164 | 2.45e-16 | 8.25e-12 |
| Malignant epithelial | C2 WNT ECs | GPD2 | 0.856 | 0.199 | 0.095 | 6.88e-16 | 2.32e-11 |
| Malignant epithelial | C2 WNT ECs | TMEM158 | 0.856 | 0.115 | 0.059 | 4.06e-08 | 0.00137 |
| Malignant epithelial | C2 WNT ECs | OSBPL2 | 0.854 | 0.445 | 0.213 | 3.71e-37 | 1.25e-32 |
| Malignant epithelial | C2 WNT ECs | PVRL1 | 0.854 | 0.225 | 0.101 | 1.6e-20 | 5.38e-16 |
| Malignant epithelial | C2 WNT ECs | SLC39A13 | 0.853 | 0.187 | 0.085 | 8.25e-17 | 2.78e-12 |
| Malignant epithelial | C2 WNT ECs | CTNNA1 | 0.852 | 0.763 | 0.473 | 1.69e-51 | 5.71e-47 |
| Malignant epithelial | C2 WNT ECs | KIF1B | 0.852 | 0.165 | 0.078 | 9.12e-14 | 3.07e-09 |
| Malignant epithelial | C2 WNT ECs | PRKACB | 0.851 | 0.106 | 0.056 | 3.37e-07 | 0.0113 |
| Malignant epithelial | C2 WNT ECs | RNPEP | 0.85 | 0.418 | 0.2 | 1.97e-33 | 6.63e-29 |
| Malignant epithelial | C2 WNT ECs | STXBP3 | 0.85 | 0.272 | 0.11 | 3.76e-30 | 1.27e-25 |
| Malignant epithelial | C2 WNT ECs | MLLT10 | 0.85 | 0.277 | 0.128 | 2.19e-24 | 7.39e-20 |
| Malignant epithelial | C2 WNT ECs | CTD-2186M15.3 | 0.85 | 0.169 | 0.082 | 3.76e-13 | 1.27e-08 |
| Malignant epithelial | C2 WNT ECs | RGL2 | 0.849 | 0.375 | 0.176 | 4.47e-31 | 1.51e-26 |
| Malignant epithelial | C2 WNT ECs | UROD | 0.849 | 0.241 | 0.117 | 1.96e-18 | 6.59e-14 |
| Malignant epithelial | C2 WNT ECs | MRE11A | 0.849 | 0.185 | 0.092 | 4.74e-14 | 1.6e-09 |
| Malignant epithelial | C2 WNT ECs | TIA1 | 0.848 | 0.394 | 0.196 | 1.87e-30 | 6.31e-26 |
| Malignant epithelial | C2 WNT ECs | ZNF644 | 0.847 | 0.267 | 0.14 | 1.32e-17 | 4.44e-13 |
| Malignant epithelial | C2 WNT ECs | PUM1 | 0.846 | 0.379 | 0.173 | 2.42e-34 | 8.16e-30 |
| Malignant epithelial | C2 WNT ECs | LMTK3 | 0.846 | 0.396 | 0.256 | 1.23e-15 | 4.15e-11 |
| Malignant epithelial | C2 WNT ECs | RPS6KA1 | 0.846 | 0.164 | 0.075 | 2.19e-14 | 7.38e-10 |
| Malignant epithelial | C2 WNT ECs | PEBP1 | 0.845 | 0.858 | 0.705 | 2.8e-51 | 9.45e-47 |
| Malignant epithelial | C2 WNT ECs | IMPA2 | 0.845 | 0.35 | 0.184 | 9.11e-24 | 3.07e-19 |
| Malignant epithelial | C2 WNT ECs | DVL2 | 0.845 | 0.16 | 0.077 | 7.95e-13 | 2.68e-08 |
| Malignant epithelial | C2 WNT ECs | RHOT2 | 0.844 | 0.463 | 0.235 | 5.629999999999999e-34 | 1.9e-29 |
| Malignant epithelial | C2 WNT ECs | PDPK1 | 0.844 | 0.332 | 0.167 | 1.46e-23 | 4.91e-19 |
| Malignant epithelial | C2 WNT ECs | CUL4B | 0.844 | 0.21 | 0.095 | 4.54e-19 | 1.53e-14 |
| Malignant epithelial | C2 WNT ECs | ZNF22 | 0.844 | 0.275 | 0.146 | 7.3e-18 | 2.46e-13 |
| Malignant epithelial | C2 WNT ECs | MAGED1 | 0.843 | 0.212 | 0.1 | 1.06e-17 | 3.56e-13 |
| Malignant epithelial | C2 WNT ECs | RASSF10 | 0.843 | 0.171 | 0.107 | 3.59e-07 | 0.0121 |
| Malignant epithelial | C2 WNT ECs | HNRNPH2 | 0.842 | 0.549 | 0.303 | 1.09e-36 | 3.67e-32 |
| Malignant epithelial | C2 WNT ECs | SCARB1 | 0.842 | 0.303 | 0.15 | 2.07e-23 | 6.98e-19 |
| Malignant epithelial | C2 WNT ECs | SPSB1 | 0.842 | 0.129 | 0.066 | 3.74e-09 | 0.000126 |
| Malignant epithelial | C2 WNT ECs | TNNC2 | 0.841 | 0.664 | 0.355 | 4.29e-54 | 1.45e-49 |
| Malignant epithelial | C2 WNT ECs | PCGF5 | 0.841 | 0.403 | 0.196 | 2.35e-31 | 7.930000000000001e-27 |
| Malignant epithelial | C2 WNT ECs | YAP1 | 0.841 | 0.448 | 0.233 | 1.58e-30 | 5.31e-26 |
| Malignant epithelial | C2 WNT ECs | LPCAT1 | 0.841 | 0.122 | 0.062 | 6.71e-09 | 0.000226 |
| Malignant epithelial | C2 WNT ECs | AKR7A2 | 0.84 | 0.437 | 0.227 | 2.83e-30 | 9.520000000000001e-26 |
| Malignant epithelial | C2 WNT ECs | ALDH7A1 | 0.84 | 0.296 | 0.151 | 1e-20 | 3.38e-16 |
| Malignant epithelial | C2 WNT ECs | CMTR2 | 0.84 | 0.299 | 0.155 | 8.99e-20 | 3.03e-15 |
| Malignant epithelial | C2 WNT ECs | SHPRH | 0.84 | 0.157 | 0.07 | 4.39e-15 | 1.48e-10 |
| Malignant epithelial | C2 WNT ECs | PPP2R5D | 0.84 | 0.138 | 0.071 | 9.03e-10 | 3.04e-05 |
| Malignant epithelial | C2 WNT ECs | TSPAN8 | 0.839 | 0.946 | 0.504 | 5.3e-99 | 1.79e-94 |
| Malignant epithelial | C2 WNT ECs | NCK2 | 0.839 | 0.25 | 0.12 | 9.1e-20 | 3.07e-15 |
| Malignant epithelial | C2 WNT ECs | DERL2 | 0.838 | 0.325 | 0.159 | 2.68e-24 | 9.04e-20 |
| Malignant epithelial | C2 WNT ECs | DDAH1 | 0.838 | 0.221 | 0.11 | 5.66e-16 | 1.91e-11 |
| Malignant epithelial | C2 WNT ECs | UMAD1 | 0.838 | 0.199 | 0.101 | 6.2e-14 | 2.09e-09 |
| Malignant epithelial | C2 WNT ECs | SP2 | 0.838 | 0.123 | 0.057 | 6.55e-11 | 2.21e-06 |
| Malignant epithelial | C2 WNT ECs | GOLGA7 | 0.837 | 0.659 | 0.411 | 5.87e-36 | 1.98e-31 |
| Malignant epithelial | C2 WNT ECs | AFG3L2 | 0.837 | 0.275 | 0.137 | 3.1e-20 | 1.04e-15 |
| Malignant epithelial | C2 WNT ECs | AGAP3 | 0.837 | 0.275 | 0.138 | 4.45e-20 | 1.5e-15 |
| Malignant epithelial | C2 WNT ECs | MRPS28 | 0.836 | 0.475 | 0.238 | 9.64e-37 | 3.25e-32 |
| Malignant epithelial | C2 WNT ECs | SEC14L2 | 0.836 | 0.139 | 0.082 | 5.38e-07 | 0.0181 |

| compartment | subcluster | gene | avg_log2FC | pct.1 | pct.2 | p_val | p_val_adj |
|---|---|---|---|---|---|---|---|
| Malignant epithelial | C2 WNT ECs | TUBA4A | 0.835 | 0.671 | 0.411 | 1.34e-39 | 4.52e-35 |
| Malignant epithelial | C2 WNT ECs | COG5 | 0.835 | 0.225 | 0.103 | 4.19e-19 | 1.41e-14 |
| Malignant epithelial | C2 WNT ECs | PRRG1 | 0.835 | 0.176 | 0.089 | 1.55e-12 | 5.23e-08 |
| Malignant epithelial | C2 WNT ECs | SCAF8 | 0.835 | 0.118 | 0.054 | 1.48e-10 | 4.98e-06 |
| Malignant epithelial | C2 WNT ECs | KRT8 | 0.834 | 0.999 | 0.997 | 5.41e-47 | 1.82e-42 |
| Malignant epithelial | C2 WNT ECs | OTULIN | 0.834 | 0.249 | 0.106 | 1.01e-24 | 3.4e-20 |
| Malignant epithelial | C2 WNT ECs | C2 | 0.834 | 0.299 | 0.174 | 8.2e-16 | 2.76e-11 |
| Malignant epithelial | C2 WNT ECs | CCDC91 | 0.834 | 0.273 | 0.165 | 2.43e-11 | 8.18e-07 |
| Malignant epithelial | C2 WNT ECs | GABARAP | 0.833 | 0.633 | 0.347 | 4.27e-45 | 1.44e-40 |
| Malignant epithelial | C2 WNT ECs | DAP | 0.833 | 0.555 | 0.319 | 9.6e-34 | 3.23e-29 |
| Malignant epithelial | C2 WNT ECs | GNPDA1 | 0.833 | 0.336 | 0.174 | 2.94e-23 | 9.89e-19 |
| Malignant epithelial | C2 WNT ECs | POLRMT | 0.833 | 0.242 | 0.114 | 1.06e-19 | 3.59e-15 |
| Malignant epithelial | C2 WNT ECs | HIC2 | 0.833 | 0.181 | 0.08 | 3.46e-17 | 1.17e-12 |
| Malignant epithelial | C2 WNT ECs | TMEM51 | 0.833 | 0.192 | 0.095 | 3.31e-14 | 1.12e-09 |
| Malignant epithelial | C2 WNT ECs | CBX8 | 0.833 | 0.119 | 0.06 | 8.44e-09 | 0.000284 |
| Malignant epithelial | C2 WNT ECs | LINC00657 | 0.832 | 0.785 | 0.516 | 2.1e-48 | 7.08e-44 |
| Malignant epithelial | C2 WNT ECs | CCDC71L | 0.831 | 0.237 | 0.122 | 7.12e-16 | 2.4e-11 |
| Malignant epithelial | C2 WNT ECs | SLC25A29 | 0.831 | 0.175 | 0.079 | 1.53e-15 | 5.14e-11 |
| Malignant epithelial | C2 WNT ECs | TYW1 | 0.831 | 0.173 | 0.088 | 4.17e-12 | 1.4e-07 |
| Malignant epithelial | C2 WNT ECs | CCZ1 | 0.83 | 0.576 | 0.307 | 5.74e-41 | 1.94e-36 |
| Malignant epithelial | C2 WNT ECs | POLR2B | 0.83 | 0.329 | 0.163 | 4.8799999999999999e-24 | 1.64e-19 |
| Malignant epithelial | C2 WNT ECs | VPS39 | 0.83 | 0.116 | 0.055 | 5.38e-10 | 1.81e-05 |
| Malignant epithelial | C2 WNT ECs | C6orf1 | 0.829 | 0.402 | 0.208 | 2.37e-27 | 7.9899999999999999e-23 |
| Malignant epithelial | C2 WNT ECs | ALDH3A2 | 0.829 | 0.225 | 0.124 | 5.34e-13 | 1.8e-08 |
| Malignant epithelial | C2 WNT ECs | TRPC4AP | 0.828 | 0.384 | 0.203 | 1.3e-25 | 4.39e-21 |
| Malignant epithelial | C2 WNT ECs | COG4 | 0.828 | 0.21 | 0.095 | 5.43e-19 | 1.83e-14 |
| Malignant epithelial | C2 WNT ECs | HMGCL | 0.827 | 0.307 | 0.153 | 1.94e-22 | 6.54e-18 |
| Malignant epithelial | C2 WNT ECs | COPS4 | 0.827 | 0.221 | 0.094 | 9.07e-22 | 3.06e-17 |
| Malignant epithelial | C2 WNT ECs | NEO1 | 0.827 | 0.124 | 0.06 | 4.84e-10 | 1.63e-05 |
| Malignant epithelial | C2 WNT ECs | WDR77 | 0.826 | 0.248 | 0.121 | 2.33e-18 | 7.85e-14 |
| Malignant epithelial | C2 WNT ECs | PPP1R14B | 0.825 | 0.782 | 0.572 | 2.49e-40 | 8.39e-36 |
| Malignant epithelial | C2 WNT ECs | MICAL2 | 0.825 | 0.16 | 0.087 | 1.55e-09 | 5.23e-05 |
| Malignant epithelial | C2 WNT ECs | SEC24C | 0.823 | 0.219 | 0.109 | 2.2e-16 | 7.42e-12 |
| Malignant epithelial | C2 WNT ECs | ENOPH1 | 0.822 | 0.342 | 0.175 | 3.04e-23 | 1.03e-18 |
| Malignant epithelial | C2 WNT ECs | GLB1 | 0.822 | 0.323 | 0.177 | 2.69e-19 | 9.07e-15 |
| Malignant epithelial | C2 WNT ECs | NCOA5 | 0.822 | 0.106 | 0.053 | 5.01e-08 | 0.00169 |
| Malignant epithelial | C2 WNT ECs | PLEKHA1 | 0.821 | 0.47 | 0.259 | 5.65e-29 | 1.9e-24 |
| Malignant epithelial | C2 WNT ECs | PRPF38A | 0.821 | 0.364 | 0.177 | 7.8499999999999999e-28 | 2.64e-23 |
| Malignant epithelial | C2 WNT ECs | OPTN | 0.821 | 0.57 | 0.392 | 5.1700000000000001e-21 | 1.74e-16 |
| Malignant epithelial | C2 WNT ECs | FDX1 | 0.82 | 0.51 | 0.276 | 1.82e-33 | 6.12e-29 |
| Malignant epithelial | C2 WNT ECs | GUSB | 0.82 | 0.371 | 0.192 | 1.15e-25 | 3.89e-21 |
| Malignant epithelial | C2 WNT ECs | SLC44A3 | 0.82 | 0.189 | 0.106 | 1.94e-10 | 6.53e-06 |
| Malignant epithelial | C2 WNT ECs | MED17 | 0.819 | 0.33 | 0.162 | 1.25e-24 | 4.22e-20 |
| Malignant epithelial | C2 WNT ECs | GCHFR | 0.819 | 0.478 | 0.287 | 4.78e-23 | 1.61e-18 |
| Malignant epithelial | C2 WNT ECs | DGKH | 0.819 | 0.233 | 0.11 | 5.68e-19 | 1.91e-14 |
| Malignant epithelial | C2 WNT ECs | TMEM260 | 0.819 | 0.123 | 0.065 | 3.2e-08 | 0.00108 |
| Malignant epithelial | C2 WNT ECs | HSPA14 | 0.818 | 0.229 | 0.111 | 1.36e-17 | 4.57e-13 |
| Malignant epithelial | C2 WNT ECs | GGA2 | 0.818 | 0.192 | 0.092 | 2.21e-15 | 7.45e-11 |
| Malignant epithelial | C2 WNT ECs | GAS8 | 0.818 | 0.122 | 0.052 | 5.89e-13 | 1.98e-08 |
| Malignant epithelial | C2 WNT ECs | PDIA6 | 0.817 | 0.825 | 0.657 | 9.83e-35 | 3.31e-30 |
| Malignant epithelial | C2 WNT ECs | RSPRY1 | 0.817 | 0.298 | 0.143 | 1.06e-23 | 3.57e-19 |
| Malignant epithelial | C2 WNT ECs | HUS1 | 0.817 | 0.257 | 0.123 | 4.78e-20 | 1.61e-15 |
| Malignant epithelial | C2 WNT ECs | H2AFX | 0.816 | 0.578 | 0.312 | 3.64e-39 | 1.23e-34 |
| Malignant epithelial | C2 WNT ECs | SACM1L | 0.816 | 0.142 | 0.067 | 6.11e-12 | 2.06e-07 |
| Malignant epithelial | C2 WNT ECs | CLPB | 0.816 | 0.11 | 0.049 | 1.24e-10 | 4.17e-06 |
| Malignant epithelial | C2 WNT ECs | STRAP | 0.815 | 0.683 | 0.445 | 1.13e-36 | 3.79e-32 |
| Malignant epithelial | C2 WNT ECs | PSEN2 | 0.815 | 0.129 | 0.058 | 3.84e-12 | 1.29e-07 |
| Malignant epithelial | C2 WNT ECs | NFAT5 | 0.814 | 0.585 | 0.366 | 4.7700000000000001e-27 | 1.61e-22 |
| Malignant epithelial | C2 WNT ECs | ARL8A | 0.814 | 0.33 | 0.165 | 9.69e-25 | 3.27e-20 |
| Malignant epithelial | C2 WNT ECs | RNFT1 | 0.814 | 0.139 | 0.063 | 1.31e-12 | 4.41e-08 |
| Malignant epithelial | C2 WNT ECs | BOK | 0.814 | 0.214 | 0.121 | 2.73e-11 | 9.2e-07 |
| Malignant epithelial | C2 WNT ECs | MBOAT7 | 0.813 | 0.547 | 0.318 | 1.99e-32 | 6.69e-28 |
| Malignant epithelial | C2 WNT ECs | FZR1 | 0.813 | 0.235 | 0.122 | 1.06e-15 | 3.58e-11 |
| Malignant epithelial | C2 WNT ECs | YWHAQ | 0.812 | 0.919 | 0.722 | 2.34e-60 | 7.8999999999999999e-56 |

| compartment | subcluster | gene | avg_log2FC | pct.1 | pct.2 | p_val | p_val_adj |
|---|---|---|---|---|---|---|---|
| Malignant epithelial | C2 WNT ECs | NADK2 | 0.812 | 0.13 | 0.057 | 1.06e-12 | 3.58e-08 |
| Malignant epithelial | C2 WNT ECs | VPS37A | 0.812 | 0.138 | 0.066 | 3.08e-11 | 1.04e-06 |
| Malignant epithelial | C2 WNT ECs | KPNA6 | 0.811 | 0.25 | 0.122 | 5.59e-19 | 1.88e-14 |
| Malignant epithelial | C2 WNT ECs | RDX | 0.811 | 0.359 | 0.213 | 5.73e-17 | 1.93e-12 |
| Malignant epithelial | C2 WNT ECs | ATAD3B | 0.811 | 0.15 | 0.069 | 3.71e-13 | 1.25e-08 |
| Malignant epithelial | C2 WNT ECs | CLDND2 | 0.811 | 0.169 | 0.083 | 8.4e-13 | 2.83e-08 |
| Malignant epithelial | C2 WNT ECs | FRMD8 | 0.811 | 0.218 | 0.121 | 1.78e-12 | 5.98e-08 |
| Malignant epithelial | C2 WNT ECs | MOB4 | 0.81 | 0.556 | 0.303 | 1.44e-36 | 4.86e-32 |
| Malignant epithelial | C2 WNT ECs | API5 | 0.81 | 0.267 | 0.134 | 5.75e-19 | 1.94e-14 |
| Malignant epithelial | C2 WNT ECs | CCBL2 | 0.809 | 0.147 | 0.062 | 3.38e-15 | 1.14e-10 |
| Malignant epithelial | C2 WNT ECs | UBXN7 | 0.809 | 0.199 | 0.099 | 1.52e-14 | 5.11e-10 |
| Malignant epithelial | C2 WNT ECs | KCTD10 | 0.809 | 0.149 | 0.076 | 1.11e-10 | 3.75e-06 |
| Malignant epithelial | C2 WNT ECs | TMEM168 | 0.809 | 0.141 | 0.071 | 2.53e-10 | 8.52e-06 |
| Malignant epithelial | C2 WNT ECs | EIF3I | 0.808 | 0.821 | 0.554 | 2.17e-57 | 7.32e-53 |
| Malignant epithelial | C2 WNT ECs | TXNDC12 | 0.808 | 0.563 | 0.341 | 4.34e-31 | 1.46e-26 |
| Malignant epithelial | C2 WNT ECs | SLC29A1 | 0.808 | 0.172 | 0.104 | 7.8e-08 | 0.00263 |
| Malignant epithelial | C2 WNT ECs | MT-ND4L | 0.807 | 0.832 | 0.619 | 1.01e-41 | 3.41e-37 |
| Malignant epithelial | C2 WNT ECs | TMEM165 | 0.807 | 0.572 | 0.321 | 1.25e-33 | 4.21e-29 |
| Malignant epithelial | C2 WNT ECs | MED8 | 0.807 | 0.298 | 0.14 | 1.06e-24 | 3.59e-20 |
| Malignant epithelial | C2 WNT ECs | PCYT2 | 0.806 | 0.268 | 0.142 | 4.42e-17 | 1.49e-12 |
| Malignant epithelial | C2 WNT ECs | RGS19 | 0.806 | 0.229 | 0.126 | 4.88e-13 | 1.64e-08 |
| Malignant epithelial | C2 WNT ECs | SLC9A1 | 0.805 | 0.129 | 0.066 | 5.66e-09 | 0.000191 |
| Malignant epithelial | C2 WNT ECs | COMMD3 | 0.804 | 0.493 | 0.266 | 2.6e-32 | 8.76e-28 |
| Malignant epithelial | C2 WNT ECs | PBX1 | 0.804 | 0.138 | 0.066 | 1.62e-11 | 5.47e-07 |
| Malignant epithelial | C2 WNT ECs | GTF2A1 | 0.804 | 0.131 | 0.065 | 3.54e-10 | 1.19e-05 |
| Malignant epithelial | C2 WNT ECs | MRFAP1 | 0.803 | 0.702 | 0.435 | 2.87e-42 | 9.65e-38 |
| Malignant epithelial | C2 WNT ECs | SRSF8 | 0.803 | 0.524 | 0.296 | 1.15e-31 | 3.87e-27 |
| Malignant epithelial | C2 WNT ECs | ZFP36L2 | 0.802 | 0.893 | 0.731 | 2.72e-38 | 9.16e-34 |
| Malignant epithelial | C2 WNT ECs | ESCO1 | 0.802 | 0.147 | 0.067 | 1.36e-13 | 4.58e-09 |
| Malignant epithelial | C2 WNT ECs | DYNC1I2 | 0.801 | 0.779 | 0.508 | 5.139999999999999e-48 | 1.73e-43 |
| Malignant epithelial | C2 WNT ECs | ERCC3 | 0.801 | 0.189 | 0.086 | 3.3e-17 | 1.11e-12 |
| Malignant epithelial | C2 WNT ECs | NGLY1 | 0.801 | 0.2 | 0.096 | 5.11e-16 | 1.72e-11 |
| Malignant epithelial | C2 WNT ECs | MTFR1L | 0.801 | 0.177 | 0.08 | 5.3e-16 | 1.78e-11 |
| Malignant epithelial | C2 WNT ECs | IL17RC | 0.801 | 0.162 | 0.082 | 1.49e-11 | 5.04e-07 |
| Malignant epithelial | C2 WNT ECs | C6orf120 | 0.801 | 0.11 | 0.051 | 1.14e-09 | 3.83e-05 |
| Malignant epithelial | C2 WNT ECs | GPRC5C | 0.8 | 0.438 | 0.213 | 1.87e-35 | 6.299999999999999e-31 |
| Malignant epithelial | C2 WNT ECs | ERICH1 | 0.8 | 0.313 | 0.165 | 7.98e-20 | 2.69e-15 |
| Malignant epithelial | C2 WNT ECs | NSUN6 | 0.8 | 0.18 | 0.084 | 2.37e-15 | 7.99e-11 |
| Malignant epithelial | C2 WNT ECs | PRMT3 | 0.8 | 0.157 | 0.076 | 1.84e-12 | 6.19e-08 |
| Malignant epithelial | C2 WNT ECs | CCNYL1 | 0.8 | 0.122 | 0.062 | 7.61e-09 | 0.000257 |
| Malignant epithelial | C2 WNT ECs | PRPF18 | 0.799 | 0.283 | 0.134 | 1.17e-22 | 3.96e-18 |
| Malignant epithelial | C2 WNT ECs | MSRB1 | 0.798 | 0.535 | 0.305 | 7.81e-32 | 2.63e-27 |
| Malignant epithelial | C2 WNT ECs | NRBP1 | 0.798 | 0.447 | 0.24 | 4.58e-28 | 1.54e-23 |
| Malignant epithelial | C2 WNT ECs | TTYH3 | 0.798 | 0.304 | 0.168 | 1.56e-17 | 5.25e-13 |
| Malignant epithelial | C2 WNT ECs | CARS2 | 0.797 | 0.403 | 0.206 | 4.32e-29 | 1.46e-24 |
| Malignant epithelial | C2 WNT ECs | YAF2 | 0.797 | 0.417 | 0.223 | 1.13e-25 | 3.790000000000001e-21 |
| Malignant epithelial | C2 WNT ECs | CBFA2T2 | 0.797 | 0.198 | 0.101 | 9.25e-14 | 3.12e-09 |
| Malignant epithelial | C2 WNT ECs | MAPRE1 | 0.796 | 0.667 | 0.445 | 3.2e-32 | 1.08e-27 |
| Malignant epithelial | C2 WNT ECs | SKI | 0.796 | 0.135 | 0.068 | 3.3e-10 | 1.11e-05 |
| Malignant epithelial | C2 WNT ECs | TDG | 0.795 | 0.451 | 0.244 | 5.2e-29 | 1.75e-24 |
| Malignant epithelial | C2 WNT ECs | WRNIP1 | 0.794 | 0.252 | 0.117 | 5.08e-21 | 1.71e-16 |
| Malignant epithelial | C2 WNT ECs | PPIF | 0.794 | 0.503 | 0.327 | 4.21e-17 | 1.42e-12 |
| Malignant epithelial | C2 WNT ECs | AGO3 | 0.794 | 0.166 | 0.075 | 3.81e-15 | 1.28e-10 |
| Malignant epithelial | C2 WNT ECs | EMC3 | 0.793 | 0.482 | 0.25 | 3.58e-33 | 1.21e-28 |
| Malignant epithelial | C2 WNT ECs | PGAP3 | 0.793 | 0.114 | 0.051 | 6.37e-11 | 2.15e-06 |
| Malignant epithelial | C2 WNT ECs | OAT | 0.792 | 0.486 | 0.265 | 1.84e-29 | 6.2e-25 |
| Malignant epithelial | C2 WNT ECs | LAMP2 | 0.792 | 0.513 | 0.295 | 8.34e-29 | 2.81e-24 |
| Malignant epithelial | C2 WNT ECs | TMEM120A | 0.792 | 0.391 | 0.204 | 1.36e-26 | 4.570000000000001e-22 |
| Malignant epithelial | C2 WNT ECs | SPATC1L | 0.792 | 0.235 | 0.104 | 1.43e-21 | 4.81e-17 |
| Malignant epithelial | C2 WNT ECs | SPATA2L | 0.792 | 0.219 | 0.109 | 6.51e-16 | 2.19e-11 |
| Malignant epithelial | C2 WNT ECs | SCPEP1 | 0.792 | 0.127 | 0.071 | 1.83e-07 | 0.00618 |
| Malignant epithelial | C2 WNT ECs | SNAP23 | 0.791 | 0.396 | 0.205 | 9.030000000000001e-27 | 3.04e-22 |
| Malignant epithelial | C2 WNT ECs | RNF146 | 0.791 | 0.214 | 0.104 | 2.15e-16 | 7.26e-12 |
| Malignant epithelial | C2 WNT ECs | IRF2BP1 | 0.791 | 0.16 | 0.076 | 5.21e-13 | 1.76e-08 |

| compartment | subcluster | gene | avg_log2FC | pct.1 | pct.2 | p_val | p_val_adj |
|---|---|---|---|---|---|---|---|
| Malignant epithelial | C2 WNT ECs | ZNHIT6 | 0.79 | 0.114 | 0.054 | 7.47e-10 | 2.52e-05 |
| Malignant epithelial | C2 WNT ECs | MYH14 | 0.789 | 0.629 | 0.377 | 9.06e-36 | 3.05e-31 |
| Malignant epithelial | C2 WNT ECs | EPHA2 | 0.789 | 0.518 | 0.331 | 1.14e-21 | 3.86e-17 |
| Malignant epithelial | C2 WNT ECs | MARS | 0.789 | 0.254 | 0.121 | 2.61e-20 | 8.79e-16 |
| Malignant epithelial | C2 WNT ECs | C12orf49 | 0.789 | 0.321 | 0.168 | 6.5e-20 | 2.19e-15 |
| Malignant epithelial | C2 WNT ECs | PTK6 | 0.789 | 0.356 | 0.205 | 4.55e-18 | 1.53e-13 |
| Malignant epithelial | C2 WNT ECs | ZUFSP | 0.789 | 0.111 | 0.056 | 2.09e-08 | 0.000705 |
| Malignant epithelial | C2 WNT ECs | CD9 | 0.788 | 0.977 | 0.917 | 3.46e-49 | 1.17e-44 |
| Malignant epithelial | C2 WNT ECs | ACTL6A | 0.788 | 0.475 | 0.272 | 3.38e-26 | 1.14e-21 |
| Malignant epithelial | C2 WNT ECs | PEX3 | 0.788 | 0.192 | 0.081 | 2.51e-19 | 8.45e-15 |
| Malignant epithelial | C2 WNT ECs | VPS8 | 0.788 | 0.157 | 0.073 | 1.53e-13 | 5.15e-09 |
| Malignant epithelial | C2 WNT ECs | CLN8 | 0.788 | 0.196 | 0.102 | 7.11e-13 | 2.4e-08 |
| Malignant epithelial | C2 WNT ECs | NARS | 0.787 | 0.43 | 0.225 | 1.08e-29 | 3.65e-25 |
| Malignant epithelial | C2 WNT ECs | IPO5 | 0.787 | 0.36 | 0.194 | 6.67e-22 | 2.25e-17 |
| Malignant epithelial | C2 WNT ECs | MYL12A | 0.786 | 0.984 | 0.877 | 1.65e-82 | 5.55e-78 |
| Malignant epithelial | C2 WNT ECs | HSPB11 | 0.786 | 0.54 | 0.31 | 3.96e-33 | 1.34e-28 |
| Malignant epithelial | C2 WNT ECs | CD81 | 0.786 | 0.578 | 0.359 | 4.79e-29 | 1.61e-24 |
| Malignant epithelial | C2 WNT ECs | FO538757.2 | 0.786 | 0.258 | 0.116 | 1.28e-23 | 4.32e-19 |
| Malignant epithelial | C2 WNT ECs | RB1 | 0.786 | 0.215 | 0.101 | 4.54e-18 | 1.53e-13 |
| Malignant epithelial | C2 WNT ECs | CEPT1 | 0.786 | 0.143 | 0.062 | 3.17e-14 | 1.07e-09 |
| Malignant epithelial | C2 WNT ECs | S100P | 0.785 | 0.838 | 0.508 | 1.01e-61 | 3.39e-57 |
| Malignant epithelial | C2 WNT ECs | SCP2 | 0.785 | 0.717 | 0.445 | 4.53e-44 | 1.53e-39 |
| Malignant epithelial | C2 WNT ECs | DDB1 | 0.785 | 0.457 | 0.241 | 1.69e-30 | 5.69e-26 |
| Malignant epithelial | C2 WNT ECs | EIF4G3 | 0.785 | 0.187 | 0.093 | 1.27e-13 | 4.28e-09 |
| Malignant epithelial | C2 WNT ECs | OPA3 | 0.784 | 0.218 | 0.122 | 8.66e-12 | 2.92e-07 |
| Malignant epithelial | C2 WNT ECs | ABHD16A | 0.784 | 0.108 | 0.054 | 1.72e-08 | 0.00058 |
| Malignant epithelial | C2 WNT ECs | EIF2B4 | 0.783 | 0.318 | 0.156 | 9.96e-24 | 3.35e-19 |
| Malignant epithelial | C2 WNT ECs | CMAS | 0.783 | 0.464 | 0.286 | 8.69e-20 | 2.93e-15 |
| Malignant epithelial | C2 WNT ECs | STX7 | 0.782 | 0.414 | 0.216 | 3e-28 | 1.01e-23 |
| Malignant epithelial | C2 WNT ECs | ASMTL | 0.782 | 0.23 | 0.096 | 4.22e-24 | 1.42e-19 |
| Malignant epithelial | C2 WNT ECs | KCNK6 | 0.782 | 0.372 | 0.199 | 2.44e-23 | 8.22e-19 |
| Malignant epithelial | C2 WNT ECs | XRCC5 | 0.781 | 0.744 | 0.475 | 1.09e-42 | 3.66e-38 |
| Malignant epithelial | C2 WNT ECs | SLC30A9 | 0.78 | 0.245 | 0.115 | 6.29e-20 | 2.12e-15 |
| Malignant epithelial | C2 WNT ECs | PSME4 | 0.78 | 0.258 | 0.136 | 2.19e-16 | 7.36e-12 |
| Malignant epithelial | C2 WNT ECs | ZNF263 | 0.78 | 0.227 | 0.125 | 1.4e-12 | 4.73e-08 |
| Malignant epithelial | C2 WNT ECs | SFPQ | 0.779 | 0.804 | 0.541 | 2.61e-46 | 8.79e-42 |
| Malignant epithelial | C2 WNT ECs | GLRX | 0.779 | 0.54 | 0.277 | 8.790000000000001e-43 | 2.96e-38 |
| Malignant epithelial | C2 WNT ECs | CLIC5 | 0.779 | 0.172 | 0.079 | 1.62e-15 | 5.47e-11 |
| Malignant epithelial | C2 WNT ECs | TFB2M | 0.779 | 0.196 | 0.095 | 2.45e-15 | 8.25e-11 |
| Malignant epithelial | C2 WNT ECs | PPP1R2 | 0.778 | 0.652 | 0.394 | 1.96e-34 | 6.61e-30 |
| Malignant epithelial | C2 WNT ECs | FBRSL1 | 0.778 | 0.206 | 0.101 | 1.19e-15 | 4.01e-11 |
| Malignant epithelial | C2 WNT ECs | FGFR4 | 0.778 | 0.211 | 0.112 | 3.84e-13 | 1.29e-08 |
| Malignant epithelial | C2 WNT ECs | CCT6A | 0.777 | 0.812 | 0.578 | 8.700000000000001e-48 | 2.93e-43 |
| Malignant epithelial | C2 WNT ECs | SMAD2 | 0.777 | 0.268 | 0.141 | 2.39e-17 | 8.06e-13 |
| Malignant epithelial | C2 WNT ECs | STAU2 | 0.777 | 0.225 | 0.111 | 1.16e-16 | 3.9e-12 |
| Malignant epithelial | C2 WNT ECs | HIST1H2BN | 0.777 | 0.137 | 0.056 | 3.89e-15 | 1.31e-10 |
| Malignant epithelial | C2 WNT ECs | WSB1 | 0.776 | 0.789 | 0.545 | 1.05e-34 | 3.55e-30 |
| Malignant epithelial | C2 WNT ECs | BRD2 | 0.776 | 0.816 | 0.634 | 4.7299999999999999e-34 | 1.59e-29 |
| Malignant epithelial | C2 WNT ECs | GJB1 | 0.776 | 0.3 | 0.152 | 2.23e-21 | 7.51e-17 |
| Malignant epithelial | C2 WNT ECs | TRIM38 | 0.776 | 0.195 | 0.086 | 1.37e-18 | 4.6e-14 |
| Malignant epithelial | C2 WNT ECs | CDADC1 | 0.776 | 0.179 | 0.086 | 5.7e-14 | 1.92e-09 |
| Malignant epithelial | C2 WNT ECs | DNMT3A | 0.776 | 0.137 | 0.069 | 3.36e-10 | 1.13e-05 |
| Malignant epithelial | C2 WNT ECs | CCNDBP1 | 0.775 | 0.432 | 0.23 | 3.22e-27 | 1.09e-22 |
| Malignant epithelial | C2 WNT ECs | PLEKHA3 | 0.775 | 0.33 | 0.168 | 7.9499999999999999e-23 | 2.68e-18 |
| Malignant epithelial | C2 WNT ECs | CDK11A | 0.775 | 0.263 | 0.123 | 8.84e-22 | 2.98e-17 |
| Malignant epithelial | C2 WNT ECs | AASDHPPT | 0.775 | 0.298 | 0.151 | 1.94e-20 | 6.55e-16 |
| Malignant epithelial | C2 WNT ECs | MFN1 | 0.775 | 0.154 | 0.068 | 7.16e-15 | 2.41e-10 |
| Malignant epithelial | C2 WNT ECs | TGFA | 0.775 | 0.156 | 0.088 | 2.53e-08 | 0.000853 |
| Malignant epithelial | C2 WNT ECs | RAB40B | 0.774 | 0.35 | 0.183 | 1.69e-23 | 5.7e-19 |
| Malignant epithelial | C2 WNT ECs | NFATC2IP | 0.774 | 0.391 | 0.216 | 5.4299999999999999e-23 | 1.83e-18 |
| Malignant epithelial | C2 WNT ECs | NCSTN | 0.774 | 0.265 | 0.138 | 3.76e-17 | 1.27e-12 |
| Malignant epithelial | C2 WNT ECs | RPN2 | 0.772 | 0.748 | 0.546 | 7.2999999999999999e-31 | 2.46e-26 |
| Malignant epithelial | C2 WNT ECs | USP14 | 0.772 | 0.288 | 0.144 | 1.04e-20 | 3.51e-16 |
| Malignant epithelial | C2 WNT ECs | DDX52 | 0.772 | 0.333 | 0.188 | 5.33e-18 | 1.8e-13 |

| compartment | subcluster | gene | avg_log2FC | pct.1 | pct.2 | p_val | p_val_adj |
|---|---|---|---|---|---|---|---|
| Malignant epithelial | C2 WNT ECs | CDKN1A | 0.772 | 0.589 | 0.424 | 1.86e-17 | 6.25e-13 |
| Malignant epithelial | C2 WNT ECs | MOSPD2 | 0.772 | 0.166 | 0.081 | 6.55e-13 | 2.21e-08 |
| Malignant epithelial | C2 WNT ECs | UBE2E3 | 0.771 | 0.639 | 0.402 | 2.68e-30 | 9.050000000000001e-26 |
| Malignant epithelial | C2 WNT ECs | LINC00685 | 0.771 | 0.143 | 0.078 | 7.59e-09 | 0.000256 |
| Malignant epithelial | C2 WNT ECs | VGLL4 | 0.77 | 0.364 | 0.202 | 5.13e-20 | 1.73e-15 |
| Malignant epithelial | C2 WNT ECs | ITPR3 | 0.77 | 0.349 | 0.223 | 2.05e-13 | 6.92e-09 |
| Malignant epithelial | C2 WNT ECs | MOAP1 | 0.769 | 0.223 | 0.097 | 1.19e-21 | 4.01e-17 |
| Malignant epithelial | C2 WNT ECs | GPN1 | 0.769 | 0.364 | 0.2 | 3.04e-21 | 1.03e-16 |
| Malignant epithelial | C2 WNT ECs | TAGLN | 0.769 | 0.111 | 0.035 | 1.06e-19 | 3.56e-15 |
| Malignant epithelial | C2 WNT ECs | TBC1D10A | 0.769 | 0.157 | 0.083 | 8.12e-10 | 2.74e-05 |
| Malignant epithelial | C2 WNT ECs | DNAJC21 | 0.768 | 0.36 | 0.185 | 3.79e-24 | 1.28e-19 |
| Malignant epithelial | C2 WNT ECs | HSF2 | 0.768 | 0.104 | 0.048 | 2.3e-09 | 7.74e-05 |
| Malignant epithelial | C2 WNT ECs | COPB1 | 0.767 | 0.371 | 0.195 | 1.71e-24 | 5.78e-20 |
| Malignant epithelial | C2 WNT ECs | UAP1 | 0.766 | 0.359 | 0.202 | 2.25e-19 | 7.59e-15 |
| Malignant epithelial | C2 WNT ECs | ETS2 | 0.766 | 0.526 | 0.358 | 9.5e-18 | 3.2e-13 |
| Malignant epithelial | C2 WNT ECs | STX18 | 0.766 | 0.222 | 0.107 | 1.86e-17 | 6.27e-13 |
| Malignant epithelial | C2 WNT ECs | CLK4 | 0.766 | 0.2 | 0.092 | 4.62e-17 | 1.56e-12 |
| Malignant epithelial | C2 WNT ECs | ALAS1 | 0.766 | 0.295 | 0.165 | 6.41e-16 | 2.16e-11 |
| Malignant epithelial | C2 WNT ECs | STX12 | 0.766 | 0.207 | 0.102 | 1.96e-15 | 6.59e-11 |
| Malignant epithelial | C2 WNT ECs | NR1D1 | 0.766 | 0.364 | 0.226 | 1.52e-14 | 5.11e-10 |
| Malignant epithelial | C2 WNT ECs | CEP63 | 0.766 | 0.179 | 0.09 | 8.12e-13 | 2.74e-08 |
| Malignant epithelial | C2 WNT ECs | GPR183 | 0.766 | 0.154 | 0.086 | 2.95e-08 | 0.000993 |
| Malignant epithelial | C2 WNT ECs | OS9 | 0.765 | 0.65 | 0.405 | 9.01e-35 | 3.04e-30 |
| Malignant epithelial | C2 WNT ECs | KMT5B | 0.765 | 0.342 | 0.171 | 2.91e-25 | 9.82e-21 |
| Malignant epithelial | C2 WNT ECs | AIF1 | 0.765 | 0.114 | 0.063 | 9.16e-07 | 0.0309 |
| Malignant epithelial | C2 WNT ECs | IST1 | 0.764 | 0.589 | 0.312 | 2.64e-40 | 8.88e-36 |
| Malignant epithelial | C2 WNT ECs | PSMD1 | 0.764 | 0.395 | 0.226 | 6.360000000000001e-21 | 2.14e-16 |
| Malignant epithelial | C2 WNT ECs | ZNF33A | 0.764 | 0.279 | 0.139 | 3.68e-20 | 1.24e-15 |
| Malignant epithelial | C2 WNT ECs | AUTS2 | 0.764 | 0.137 | 0.069 | 1.13e-09 | 3.8e-05 |
| Malignant epithelial | C2 WNT ECs | ETV4 | 0.763 | 0.379 | 0.182 | 2.55e-29 | 8.59e-25 |
| Malignant epithelial | C2 WNT ECs | TMEM80 | 0.763 | 0.208 | 0.121 | 3.63e-10 | 1.22e-05 |
| Malignant epithelial | C2 WNT ECs | ZMYND8 | 0.762 | 0.487 | 0.276 | 3.96e-29 | 1.34e-24 |
| Malignant epithelial | C2 WNT ECs | TRADD | 0.762 | 0.36 | 0.187 | 3.36e-23 | 1.13e-18 |
| Malignant epithelial | C2 WNT ECs | RMND1 | 0.762 | 0.196 | 0.09 | 9.89e-17 | 3.33e-12 |
| Malignant epithelial | C2 WNT ECs | PPP2R2D | 0.762 | 0.2 | 0.101 | 4.05e-14 | 1.36e-09 |
| Malignant epithelial | C2 WNT ECs | IDE | 0.762 | 0.111 | 0.048 | 1.54e-11 | 5.2e-07 |
| Malignant epithelial | C2 WNT ECs | TMEM176B | 0.761 | 0.919 | 0.774 | 2.53e-33 | 8.520000000000001e-29 |
| Malignant epithelial | C2 WNT ECs | FUCA2 | 0.761 | 0.493 | 0.269 | 5.09e-31 | 1.71e-26 |
| Malignant epithelial | C2 WNT ECs | MOB2 | 0.761 | 0.433 | 0.234 | 2.88e-26 | 9.7e-22 |
| Malignant epithelial | C2 WNT ECs | KIDINS220 | 0.761 | 0.261 | 0.13 | 9.02e-19 | 3.04e-14 |
| Malignant epithelial | C2 WNT ECs | ABR | 0.761 | 0.225 | 0.115 | 2.07e-15 | 6.96e-11 |
| Malignant epithelial | C2 WNT ECs | SYVN1 | 0.761 | 0.199 | 0.103 | 3.46e-13 | 1.17e-08 |
| Malignant epithelial | C2 WNT ECs | PRIM2 | 0.761 | 0.16 | 0.083 | 1.31e-10 | 4.4e-06 |
| Malignant epithelial | C2 WNT ECs | NETO2 | 0.761 | 0.143 | 0.072 | 1.33e-10 | 4.47e-06 |
| Malignant epithelial | C2 WNT ECs | AIG1 | 0.76 | 0.671 | 0.42 | 2.89e-34 | 9.729999999999999e-30 |
| Malignant epithelial | C2 WNT ECs | SH3GLB1 | 0.76 | 0.567 | 0.341 | 5.25e-31 | 1.77e-26 |
| Malignant epithelial | C2 WNT ECs | SFXN3 | 0.76 | 0.35 | 0.19 | 9.39e-22 | 3.16e-17 |
| Malignant epithelial | C2 WNT ECs | INTS5 | 0.76 | 0.108 | 0.057 | 2.88e-07 | 0.0097 |
| Malignant epithelial | C2 WNT ECs | DDX5 | 0.759 | 0.954 | 0.863 | 4.6e-49 | 1.55e-44 |
| Malignant epithelial | C2 WNT ECs | PNMA1 | 0.759 | 0.25 | 0.128 | 2.21e-17 | 7.46e-13 |
| Malignant epithelial | C2 WNT ECs | QPCT | 0.759 | 0.3 | 0.193 | 4.62e-11 | 1.56e-06 |
| Malignant epithelial | C2 WNT ECs | PAFAH1B2 | 0.758 | 0.591 | 0.338 | 1.82e-35 | 6.14e-31 |
| Malignant epithelial | C2 WNT ECs | NFE2L2 | 0.758 | 0.627 | 0.383 | 2.89e-31 | 9.73e-27 |
| Malignant epithelial | C2 WNT ECs | SLBP | 0.758 | 0.453 | 0.241 | 4.13e-29 | 1.39e-24 |
| Malignant epithelial | C2 WNT ECs | KCNE3 | 0.758 | 0.177 | 0.079 | 2.94e-16 | 9.9e-12 |
| Malignant epithelial | C2 WNT ECs | GPR160 | 0.757 | 0.356 | 0.209 | 8.95e-18 | 3.02e-13 |
| Malignant epithelial | C2 WNT ECs | SORBS2 | 0.757 | 0.107 | 0.055 | 1.16e-07 | 0.00391 |
| Malignant epithelial | C2 WNT ECs | AKR1A1 | 0.756 | 0.593 | 0.335 | 1.6e-37 | 5.4e-33 |
| Malignant epithelial | C2 WNT ECs | DNAJC3-AS1 | 0.756 | 0.172 | 0.073 | 4.91e-18 | 1.65e-13 |
| Malignant epithelial | C2 WNT ECs | NNT-AS1 | 0.756 | 0.227 | 0.119 | 3.3e-14 | 1.11e-09 |
| Malignant epithelial | C2 WNT ECs | TSEN2 | 0.756 | 0.116 | 0.048 | 9.3e-13 | 3.13e-08 |
| Malignant epithelial | C2 WNT ECs | FRS2 | 0.756 | 0.107 | 0.051 | 6.15e-09 | 0.000207 |
| Malignant epithelial | C2 WNT ECs | CNST | 0.756 | 0.104 | 0.054 | 2.47e-07 | 0.00833 |
| Malignant epithelial | C2 WNT ECs | BCL2L1 | 0.755 | 0.647 | 0.446 | 1.13e-22 | 3.82e-18 |

| compartment | subcluster | gene | avg_log2FC | pct.1 | pct.2 | p_val | p_val_adj |
|---|---|---|---|---|---|---|---|
| Malignant epithelial | C2 WNT ECs | UBE2E2 | 0.755 | 0.217 | 0.108 | 1.43e-15 | 4.82e-11 |
| Malignant epithelial | C2 WNT ECs | PNRC2 | 0.754 | 0.609 | 0.362 | 8.2e-31 | 2.76e-26 |
| Malignant epithelial | C2 WNT ECs | MRPL1 | 0.754 | 0.379 | 0.191 | 2.4e-26 | 8.08e-22 |
| Malignant epithelial | C2 WNT ECs | DEXI | 0.754 | 0.356 | 0.179 | 6.89e-25 | 2.32e-20 |
| Malignant epithelial | C2 WNT ECs | INTS10 | 0.754 | 0.234 | 0.117 | 4.83e-16 | 1.63e-11 |
| Malignant epithelial | C2 WNT ECs | TSKU | 0.753 | 0.157 | 0.078 | 2.36e-11 | 7.95e-07 |
| Malignant epithelial | C2 WNT ECs | SRSF10 | 0.752 | 0.697 | 0.421 | 3.44e-39 | 1.16e-34 |
| Malignant epithelial | C2 WNT ECs | ELP2 | 0.752 | 0.233 | 0.113 | 4.82e-18 | 1.62e-13 |
| Malignant epithelial | C2 WNT ECs | LUZP1 | 0.752 | 0.345 | 0.206 | 3.89e-16 | 1.31e-11 |
| Malignant epithelial | C2 WNT ECs | GSAP | 0.752 | 0.208 | 0.105 | 2.13e-14 | 7.19e-10 |
| Malignant epithelial | C2 WNT ECs | MAN2A1 | 0.752 | 0.194 | 0.098 | 1.95e-13 | 6.56e-09 |
| Malignant epithelial | C2 WNT ECs | TMEM127 | 0.752 | 0.25 | 0.148 | 6.99e-12 | 2.36e-07 |
| Malignant epithelial | C2 WNT ECs | GGT6 | 0.751 | 0.106 | 0.046 | 7.87e-11 | 2.65e-06 |
| Malignant epithelial | C2 WNT ECs | ATMIN | 0.751 | 0.142 | 0.076 | 7.65e-09 | 0.000258 |
| Malignant epithelial | C2 WNT ECs | FDPS | 0.75 | 0.671 | 0.455 | 8.810000000000001e-27 | 2.97e-22 |
| Malignant epithelial | C2 WNT ECs | BLOC1S4 | 0.75 | 0.361 | 0.185 | 2.46e-24 | 8.28e-20 |
| Malignant epithelial | C2 WNT ECs | SLC48A1 | 0.75 | 0.16 | 0.085 | 7.3e-10 | 2.46e-05 |
| Malignant epithelial | C2 WNT ECs | ICA1 | 0.749 | 0.667 | 0.394 | 1.03e-38 | 3.46e-34 |
| Malignant epithelial | C2 WNT ECs | PRPF6 | 0.749 | 0.654 | 0.448 | 3.73e-29 | 1.26e-24 |
| Malignant epithelial | C2 WNT ECs | MBTD1 | 0.749 | 0.106 | 0.052 | 3.04e-08 | 0.00102 |
| Malignant epithelial | C2 WNT ECs | HSP90B1 | 0.748 | 0.936 | 0.78 | 1.19e-54 | 4e-50 |
| Malignant epithelial | C2 WNT ECs | TMUB1 | 0.748 | 0.685 | 0.397 | 3.8299999999999999e-42 | 1.29e-37 |
| Malignant epithelial | C2 WNT ECs | GAS6 | 0.748 | 0.522 | 0.334 | 7.43e-22 | 2.5e-17 |
| Malignant epithelial | C2 WNT ECs | PCSK7 | 0.748 | 0.541 | 0.366 | 2.06e-18 | 6.95e-14 |
| Malignant epithelial | C2 WNT ECs | ANKRA2 | 0.748 | 0.211 | 0.112 | 4.05e-13 | 1.37e-08 |
| Malignant epithelial | C2 WNT ECs | ABCC1 | 0.748 | 0.118 | 0.062 | 6.48e-08 | 0.00218 |
| Malignant epithelial | C2 WNT ECs | RABGGTB | 0.747 | 0.506 | 0.27 | 1.19e-32 | 4.01e-28 |
| Malignant epithelial | C2 WNT ECs | DNAJB11 | 0.747 | 0.613 | 0.39 | 3.98e-30 | 1.34e-25 |
| Malignant epithelial | C2 WNT ECs | ZNF146 | 0.747 | 0.275 | 0.157 | 1.87e-14 | 6.29e-10 |
| Malignant epithelial | C2 WNT ECs | CD83 | 0.747 | 0.146 | 0.085 | 1.86e-07 | 0.00628 |
| Malignant epithelial | C2 WNT ECs | PRELID3B | 0.745 | 0.844 | 0.579 | 1.33e-46 | 4.49e-42 |
| Malignant epithelial | C2 WNT ECs | LAPTM4A | 0.745 | 0.892 | 0.749 | 4.49e-42 | 1.51e-37 |
| Malignant epithelial | C2 WNT ECs | FAM63B | 0.745 | 0.279 | 0.14 | 3.41e-19 | 1.15e-14 |
| Malignant epithelial | C2 WNT ECs | SECISBP2L | 0.745 | 0.168 | 0.082 | 1.09e-12 | 3.66e-08 |
| Malignant epithelial | C2 WNT ECs | CDH1 | 0.744 | 0.685 | 0.436 | 4.94e-32 | 1.66e-27 |
| Malignant epithelial | C2 WNT ECs | LIMS1 | 0.744 | 0.505 | 0.295 | 4.57e-26 | 1.54e-21 |
| Malignant epithelial | C2 WNT ECs | TARDBP | 0.744 | 0.41 | 0.229 | 1.53e-22 | 5.15e-18 |
| Malignant epithelial | C2 WNT ECs | TMEM63B | 0.744 | 0.272 | 0.13 | 6.94e-22 | 2.34e-17 |
| Malignant epithelial | C2 WNT ECs | ANKRD27 | 0.744 | 0.143 | 0.073 | 3.11e-10 | 1.05e-05 |
| Malignant epithelial | C2 WNT ECs | CDX1 | 0.743 | 0.453 | 0.205 | 1.51e-39 | 5.1e-35 |
| Malignant epithelial | C2 WNT ECs | ST14 | 0.743 | 0.725 | 0.521 | 1.15e-31 | 3.880000000000001e-27 |
| Malignant epithelial | C2 WNT ECs | EPC2 | 0.743 | 0.129 | 0.063 | 4.83e-10 | 1.63e-05 |
| Malignant epithelial | C2 WNT ECs | AHDC1 | 0.743 | 0.1 | 0.049 | 4.41e-08 | 0.00149 |
| Malignant epithelial | C2 WNT ECs | GDI2 | 0.742 | 0.82 | 0.603 | 1.62e-47 | 5.46e-43 |
| Malignant epithelial | C2 WNT ECs | CLPX | 0.742 | 0.261 | 0.116 | 1.72e-24 | 5.78e-20 |
| Malignant epithelial | C2 WNT ECs | PCBP4 | 0.742 | 0.337 | 0.192 | 9.19e-18 | 3.1e-13 |
| Malignant epithelial | C2 WNT ECs | ZNF668 | 0.742 | 0.12 | 0.057 | 4.87e-10 | 1.64e-05 |
| Malignant epithelial | C2 WNT ECs | RP11-1275H24.1 | 0.742 | 0.116 | 0.055 | 7.33e-10 | 2.47e-05 |
| Malignant epithelial | C2 WNT ECs | YME1L1 | 0.741 | 0.547 | 0.29 | 1.07e-35 | 3.59e-31 |
| Malignant epithelial | C2 WNT ECs | ARHGEF1 | 0.741 | 0.411 | 0.248 | 6.33e-18 | 2.13e-13 |
| Malignant epithelial | C2 WNT ECs | UBALD1 | 0.741 | 0.29 | 0.157 | 1.01e-16 | 3.4e-12 |
| Malignant epithelial | C2 WNT ECs | TFPT | 0.741 | 0.203 | 0.099 | 1.85e-15 | 6.23e-11 |
| Malignant epithelial | C2 WNT ECs | C10orf54 | 0.741 | 0.302 | 0.188 | 1.78e-12 | 5.98e-08 |
| Malignant epithelial | C2 WNT ECs | KIAA1217 | 0.741 | 0.141 | 0.066 | 4.16e-12 | 1.4e-07 |
| Malignant epithelial | C2 WNT ECs | NOP14 | 0.741 | 0.141 | 0.075 | 5.31e-09 | 0.000179 |
| Malignant epithelial | C2 WNT ECs | GK | 0.741 | 0.118 | 0.059 | 6.25e-09 | 0.000211 |
| Malignant epithelial | C2 WNT ECs | SPOP | 0.74 | 0.367 | 0.205 | 1.71e-19 | 5.78e-15 |
| Malignant epithelial | C2 WNT ECs | TBL3 | 0.74 | 0.288 | 0.153 | 3.22e-18 | 1.09e-13 |
| Malignant epithelial | C2 WNT ECs | ZDHHC7 | 0.74 | 0.214 | 0.108 | 1.71e-14 | 5.77e-10 |
| Malignant epithelial | C2 WNT ECs | HMHA1 | 0.739 | 0.33 | 0.161 | 8.37e-25 | 2.82e-20 |
| Malignant epithelial | C2 WNT ECs | SLC39A10 | 0.739 | 0.233 | 0.119 | 1.39e-15 | 4.69e-11 |
| Malignant epithelial | C2 WNT ECs | LFNG | 0.739 | 0.25 | 0.134 | 5.26e-15 | 1.77e-10 |
| Malignant epithelial | C2 WNT ECs | LAMA5 | 0.739 | 0.245 | 0.134 | 5.36e-14 | 1.81e-09 |
| Malignant epithelial | C2 WNT ECs | CRY1 | 0.739 | 0.139 | 0.067 | 4.46e-11 | 1.5e-06 |

| compartment | subcluster | gene | avg_log2FC | pct.1 | pct.2 | p_val | p_val_adj |
|---|---|---|---|---|---|---|---|
| Malignant epithelial | C2 WNT ECs | SIRT7 | 0.738 | 0.54 | 0.294 | 3.52e-33 | 1.19e-28 |
| Malignant epithelial | C2 WNT ECs | HP1BP3 | 0.738 | 0.589 | 0.35 | 5.750000000000001e-32 | 1.94e-27 |
| Malignant epithelial | C2 WNT ECs | SRM | 0.738 | 0.401 | 0.231 | 1.6e-20 | 5.4e-16 |
| Malignant epithelial | C2 WNT ECs | SMIM1 | 0.738 | 0.183 | 0.077 | 1.42e-18 | 4.78e-14 |
| Malignant epithelial | C2 WNT ECs | EHD1 | 0.738 | 0.406 | 0.246 | 6.52e-17 | 2.2e-12 |
| Malignant epithelial | C2 WNT ECs | MYO1F | 0.738 | 0.112 | 0.059 | 2.23e-07 | 0.00752 |
| Malignant epithelial | C2 WNT ECs | SEPHS1 | 0.737 | 0.235 | 0.126 | 3.65e-14 | 1.23e-09 |
| Malignant epithelial | C2 WNT ECs | IFT52 | 0.737 | 0.26 | 0.146 | 8.05e-14 | 2.71e-09 |
| Malignant epithelial | C2 WNT ECs | PIKFYVE | 0.737 | 0.11 | 0.049 | 1.13e-10 | 3.81e-06 |
| Malignant epithelial | C2 WNT ECs | FLII | 0.736 | 0.267 | 0.15 | 2.56e-14 | 8.61e-10 |
| Malignant epithelial | C2 WNT ECs | ARHGAP32 | 0.735 | 0.372 | 0.221 | 2.83e-17 | 9.53e-13 |
| Malignant epithelial | C2 WNT ECs | MYC | 0.735 | 0.636 | 0.481 | 8.22e-15 | 2.77e-10 |
| Malignant epithelial | C2 WNT ECs | PEX14 | 0.735 | 0.139 | 0.065 | 1.01e-11 | 3.4e-07 |
| Malignant epithelial | C2 WNT ECs | FUT3 | 0.735 | 0.2 | 0.111 | 1.99e-11 | 6.72e-07 |
| Malignant epithelial | C2 WNT ECs | LMO4 | 0.735 | 0.177 | 0.101 | 4.36e-09 | 0.000147 |
| Malignant epithelial | C2 WNT ECs | PCYOX1 | 0.734 | 0.229 | 0.119 | 4.62e-15 | 1.56e-10 |
| Malignant epithelial | C2 WNT ECs | RCN1 | 0.734 | 0.595 | 0.501 | 5.03e-10 | 1.69e-05 |
| Malignant epithelial | C2 WNT ECs | SMG6 | 0.734 | 0.103 | 0.051 | 3.24e-08 | 0.00109 |
| Malignant epithelial | C2 WNT ECs | SRSF4 | 0.733 | 0.457 | 0.245 | 5.77e-29 | 1.94e-24 |
| Malignant epithelial | C2 WNT ECs | RAE1 | 0.733 | 0.449 | 0.242 | 1.39e-27 | 4.669999999999999e-23 |
| Malignant epithelial | C2 WNT ECs | DLD | 0.733 | 0.355 | 0.188 | 6.77e-22 | 2.28e-17 |
| Malignant epithelial | C2 WNT ECs | FLYWCH2 | 0.733 | 0.338 | 0.178 | 1.39e-21 | 4.67e-17 |
| Malignant epithelial | C2 WNT ECs | FYTTD1 | 0.733 | 0.307 | 0.162 | 1.6e-19 | 5.38e-15 |
| Malignant epithelial | C2 WNT ECs | PRR14 | 0.733 | 0.287 | 0.149 | 6.07e-19 | 2.05e-14 |
| Malignant epithelial | C2 WNT ECs | ACYP2 | 0.733 | 0.248 | 0.12 | 7.17e-19 | 2.42e-14 |
| Malignant epithelial | C2 WNT ECs | MAGI1 | 0.733 | 0.175 | 0.089 | 3.02e-12 | 1.02e-07 |
| Malignant epithelial | C2 WNT ECs | GTPBP4 | 0.732 | 0.44 | 0.254 | 9.96e-24 | 3.36e-19 |
| Malignant epithelial | C2 WNT ECs | GLT8D1 | 0.732 | 0.294 | 0.153 | 1.09e-18 | 3.68e-14 |
| Malignant epithelial | C2 WNT ECs | HCG18 | 0.732 | 0.173 | 0.086 | 2.17e-12 | 7.31e-08 |
| Malignant epithelial | C2 WNT ECs | ZDHHC13 | 0.732 | 0.187 | 0.1 | 1.86e-11 | 6.25e-07 |
| Malignant epithelial | C2 WNT ECs | SPAST | 0.732 | 0.12 | 0.061 | 1.15e-08 | 0.000387 |
| Malignant epithelial | C2 WNT ECs | CANX | 0.731 | 0.783 | 0.569 | 2.1e-40 | 7.060000000000001e-36 |
| Malignant epithelial | C2 WNT ECs | FAM221A | 0.731 | 0.166 | 0.069 | 2.19e-17 | 7.37e-13 |
| Malignant epithelial | C2 WNT ECs | AK1 | 0.731 | 0.471 | 0.318 | 4.6e-17 | 1.55e-12 |
| Malignant epithelial | C2 WNT ECs | ZNF92 | 0.731 | 0.103 | 0.053 | 2.33e-07 | 0.00784 |
| Malignant epithelial | C2 WNT ECs | OSTM1 | 0.731 | 0.118 | 0.064 | 2.71e-07 | 0.00913 |
| Malignant epithelial | C2 WNT ECs | CFL1 | 0.73 | 0.995 | 0.972 | 2.44e-107 | 8.230000000000001e-103 |
| Malignant epithelial | C2 WNT ECs | RPUSD3 | 0.73 | 0.28 | 0.15 | 1.33e-16 | 4.49e-12 |
| Malignant epithelial | C2 WNT ECs | NT5DC2 | 0.73 | 0.141 | 0.06 | 1.4e-14 | 4.73e-10 |
| Malignant epithelial | C2 WNT ECs | BCR | 0.729 | 0.187 | 0.093 | 1.34e-13 | 4.53e-09 |
| Malignant epithelial | C2 WNT ECs | LRPAP1 | 0.728 | 0.582 | 0.337 | 1.36e-32 | 4.59e-28 |
| Malignant epithelial | C2 WNT ECs | EEF1E1 | 0.728 | 0.475 | 0.271 | 1.27e-27 | 4.279999999999999e-23 |
| Malignant epithelial | C2 WNT ECs | BCAS2 | 0.728 | 0.401 | 0.202 | 1.41e-27 | 4.74e-23 |
| Malignant epithelial | C2 WNT ECs | SLC35B2 | 0.728 | 0.46 | 0.291 | 1.83e-19 | 6.15e-15 |
| Malignant epithelial | C2 WNT ECs | LPCAT3 | 0.728 | 0.323 | 0.184 | 5.07e-17 | 1.71e-12 |
| Malignant epithelial | C2 WNT ECs | LRP1 | 0.728 | 0.18 | 0.082 | 9.52e-16 | 3.21e-11 |
| Malignant epithelial | C2 WNT ECs | USP1 | 0.728 | 0.276 | 0.15 | 1.21e-15 | 4.09e-11 |
| Malignant epithelial | C2 WNT ECs | SNX4 | 0.728 | 0.199 | 0.096 | 1.73e-15 | 5.83e-11 |
| Malignant epithelial | C2 WNT ECs | FN3KRP | 0.728 | 0.246 | 0.133 | 2.01e-14 | 6.78e-10 |
| Malignant epithelial | C2 WNT ECs | TUBE1 | 0.728 | 0.171 | 0.082 | 4.09e-13 | 1.38e-08 |
| Malignant epithelial | C2 WNT ECs | NUPR1 | 0.728 | 0.207 | 0.114 | 5.72e-12 | 1.93e-07 |
| Malignant epithelial | C2 WNT ECs | TOM1L2 | 0.728 | 0.147 | 0.079 | 4.21e-09 | 0.000142 |
| Malignant epithelial | C2 WNT ECs | TNFRSF1B | 0.728 | 0.118 | 0.061 | 3.53e-08 | 0.00119 |
| Malignant epithelial | C2 WNT ECs | GAK | 0.727 | 0.263 | 0.128 | 1.41e-19 | 4.76e-15 |
| Malignant epithelial | C2 WNT ECs | ALG14 | 0.727 | 0.202 | 0.101 | 1.91e-14 | 6.44e-10 |
| Malignant epithelial | C2 WNT ECs | SMIM13 | 0.727 | 0.138 | 0.06 | 1.69e-13 | 5.7e-09 |
| Malignant epithelial | C2 WNT ECs | PSMD3 | 0.726 | 0.548 | 0.333 | 1.71e-27 | 5.77e-23 |
| Malignant epithelial | C2 WNT ECs | NXN | 0.726 | 0.115 | 0.062 | 1.9e-07 | 0.00642 |
| Malignant epithelial | C2 WNT ECs | RSRC2 | 0.725 | 0.797 | 0.574 | 9.680000000000001e-37 | 3.26e-32 |
| Malignant epithelial | C2 WNT ECs | RTCB | 0.725 | 0.47 | 0.256 | 5.37e-28 | 1.81e-23 |
| Malignant epithelial | C2 WNT ECs | TXNDC11 | 0.725 | 0.28 | 0.144 | 1.13e-18 | 3.8e-14 |
| Malignant epithelial | C2 WNT ECs | CXXC5 | 0.724 | 0.65 | 0.409 | 1.06e-32 | 3.55e-28 |
| Malignant epithelial | C2 WNT ECs | PRDX5 | 0.724 | 0.915 | 0.869 | 3.4e-14 | 1.14e-09 |
| Malignant epithelial | C2 WNT ECs | FLYWCH1 | 0.724 | 0.126 | 0.06 | 3.14e-10 | 1.06e-05 |

| compartment | subcluster | gene | avg_log2FC | pct.1 | pct.2 | p_val | p_val_adj |
|---|---|---|---|---|---|---|---|
| Malignant epithelial | C2 WNT ECs | UCKL1 | 0.723 | 0.414 | 0.215 | 1.59e-27 | 5.36e-23 |
| Malignant epithelial | C2 WNT ECs | VWA1 | 0.723 | 0.432 | 0.255 | 2.71e-21 | 9.13e-17 |
| Malignant epithelial | C2 WNT ECs | ZNF3 | 0.723 | 0.3 | 0.158 | 5.93e-19 | 2e-14 |
| Malignant epithelial | C2 WNT ECs | THUMPD3 | 0.723 | 0.211 | 0.114 | 2.66e-12 | 8.95e-08 |
| Malignant epithelial | C2 WNT ECs | HACL1 | 0.722 | 0.137 | 0.071 | 2.85e-09 | 9.6e-05 |
| Malignant epithelial | C2 WNT ECs | RAB7A | 0.721 | 0.861 | 0.644 | 3.65e-40 | 1.23e-35 |
| Malignant epithelial | C2 WNT ECs | SCYL1 | 0.721 | 0.323 | 0.182 | 2.13e-16 | 7.19e-12 |
| Malignant epithelial | C2 WNT ECs | WDR75 | 0.721 | 0.198 | 0.094 | 6.99e-16 | 2.36e-11 |
| Malignant epithelial | C2 WNT ECs | MUT | 0.721 | 0.166 | 0.08 | 2.47e-13 | 8.31e-09 |
| Malignant epithelial | C2 WNT ECs | ZXDC | 0.721 | 0.108 | 0.043 | 5.54e-13 | 1.87e-08 |
| Malignant epithelial | C2 WNT ECs | HOTAIRM1 | 0.72 | 0.329 | 0.164 | 1.43e-23 | 4.82e-19 |
| Malignant epithelial | C2 WNT ECs | GNAI2 | 0.72 | 0.589 | 0.392 | 1.69e-21 | 5.68e-17 |
| Malignant epithelial | C2 WNT ECs | CNOT8 | 0.72 | 0.257 | 0.14 | 4.75e-15 | 1.6e-10 |
| Malignant epithelial | C2 WNT ECs | PLIN3 | 0.719 | 0.608 | 0.38 | 1.67e-28 | 5.62e-24 |
| Malignant epithelial | C2 WNT ECs | MARCH5 | 0.719 | 0.353 | 0.185 | 2.36e-22 | 7.97e-18 |
| Malignant epithelial | C2 WNT ECs | MXD4 | 0.719 | 0.206 | 0.109 | 7.1e-13 | 2.39e-08 |
| Malignant epithelial | C2 WNT ECs | ZSWIM6 | 0.719 | 0.13 | 0.061 | 6.71e-11 | 2.26e-06 |
| Malignant epithelial | C2 WNT ECs | PHF13 | 0.719 | 0.116 | 0.053 | 1.06e-10 | 3.59e-06 |
| Malignant epithelial | C2 WNT ECs | KCTD20 | 0.719 | 0.152 | 0.089 | 1.77e-07 | 0.00597 |
| Malignant epithelial | C2 WNT ECs | CAMK2N1 | 0.718 | 0.643 | 0.41 | 2.13e-28 | 7.18e-24 |
| Malignant epithelial | C2 WNT ECs | HGD | 0.718 | 0.133 | 0.063 | 3.81e-11 | 1.28e-06 |
| Malignant epithelial | C2 WNT ECs | TRAPPC9 | 0.718 | 0.122 | 0.057 | 8.43e-11 | 2.84e-06 |
| Malignant epithelial | C2 WNT ECs | ANKRD46 | 0.718 | 0.141 | 0.075 | 5.07e-09 | 0.000171 |
| Malignant epithelial | C2 WNT ECs | PDLIM1 | 0.717 | 0.839 | 0.58 | 1.86e-47 | 6.260000000000001e-43 |
| Malignant epithelial | C2 WNT ECs | CRACR2B | 0.717 | 0.429 | 0.231 | 1.29e-26 | 4.34e-22 |
| Malignant epithelial | C2 WNT ECs | TM4SF5 | 0.717 | 0.311 | 0.15 | 6.6e-24 | 2.22e-19 |
| Malignant epithelial | C2 WNT ECs | FAM168B | 0.717 | 0.233 | 0.115 | 5.99e-17 | 2.02e-12 |
| Malignant epithelial | C2 WNT ECs | GRAMD3 | 0.717 | 0.235 | 0.131 | 8.54e-13 | 2.88e-08 |
| Malignant epithelial | C2 WNT ECs | GTF3C1 | 0.717 | 0.16 | 0.084 | 2.76e-10 | 9.3e-06 |
| Malignant epithelial | C2 WNT ECs | DOPEY2 | 0.717 | 0.119 | 0.061 | 1.3e-08 | 0.000439 |
| Malignant epithelial | C2 WNT ECs | LRRC59 | 0.716 | 0.62 | 0.387 | 4.31e-30 | 1.45e-25 |
| Malignant epithelial | C2 WNT ECs | HOXB9 | 0.716 | 0.501 | 0.297 | 2.43e-24 | 8.2e-20 |
| Malignant epithelial | C2 WNT ECs | PPID | 0.716 | 0.344 | 0.196 | 6.49e-18 | 2.19e-13 |
| Malignant epithelial | C2 WNT ECs | OGFOD1 | 0.716 | 0.158 | 0.087 | 3.92e-09 | 0.000132 |
| Malignant epithelial | C2 WNT ECs | GRN | 0.715 | 0.748 | 0.595 | 3.29e-24 | 1.11e-19 |
| Malignant epithelial | C2 WNT ECs | LYRM4 | 0.714 | 0.479 | 0.267 | 3.92e-28 | 1.32e-23 |
| Malignant epithelial | C2 WNT ECs | GTF3C5 | 0.714 | 0.38 | 0.206 | 1.6e-21 | 5.4e-17 |
| Malignant epithelial | C2 WNT ECs | TMEM128 | 0.714 | 0.245 | 0.138 | 5.41e-13 | 1.82e-08 |
| Malignant epithelial | C2 WNT ECs | CMPK1 | 0.713 | 0.651 | 0.425 | 4.17e-29 | 1.41e-24 |
| Malignant epithelial | C2 WNT ECs | ZNF32 | 0.713 | 0.238 | 0.129 | 9.1e-14 | 3.07e-09 |
| Malignant epithelial | C2 WNT ECs | RPGR | 0.713 | 0.158 | 0.074 | 3.64e-13 | 1.23e-08 |
| Malignant epithelial | C2 WNT ECs | ATPAF1 | 0.713 | 0.222 | 0.124 | 4.52e-12 | 1.52e-07 |
| Malignant epithelial | C2 WNT ECs | PLEKHB2 | 0.712 | 0.499 | 0.289 | 6.2e-26 | 2.09e-21 |
| Malignant epithelial | C2 WNT ECs | GRSF1 | 0.712 | 0.457 | 0.257 | 2.86e-25 | 9.63e-21 |
| Malignant epithelial | C2 WNT ECs | CRYL1 | 0.712 | 0.323 | 0.154 | 9.18e-25 | 3.09e-20 |
| Malignant epithelial | C2 WNT ECs | LZTS2 | 0.712 | 0.183 | 0.102 | 5.76e-10 | 1.94e-05 |
| Malignant epithelial | C2 WNT ECs | SBDS | 0.711 | 0.896 | 0.694 | 7.909999999999999e-45 | 2.66e-40 |
| Malignant epithelial | C2 WNT ECs | CCDC53 | 0.711 | 0.543 | 0.327 | 1.18e-27 | 3.98e-23 |
| Malignant epithelial | C2 WNT ECs | TSNAX | 0.711 | 0.28 | 0.159 | 3.01e-14 | 1.01e-09 |
| Malignant epithelial | C2 WNT ECs | MGMT | 0.709 | 0.67 | 0.411 | 5.64e-37 | 1.9e-32 |
| Malignant epithelial | C2 WNT ECs | TMEM167B | 0.709 | 0.21 | 0.113 | 1.73e-12 | 5.82e-08 |
| Malignant epithelial | C2 WNT ECs | STK38L | 0.709 | 0.194 | 0.105 | 2.19e-11 | 7.38e-07 |
| Malignant epithelial | C2 WNT ECs | CACNB3 | 0.709 | 0.143 | 0.074 | 9.16e-10 | 3.09e-05 |
| Malignant epithelial | C2 WNT ECs | TMEM14B | 0.708 | 0.756 | 0.504 | 1.63e-39 | 5.51e-35 |
| Malignant epithelial | C2 WNT ECs | RBBP4 | 0.708 | 0.455 | 0.256 | 1.26e-25 | 4.240000000000001e-21 |
| Malignant epithelial | C2 WNT ECs | MAFK | 0.708 | 0.352 | 0.215 | 3.87e-14 | 1.3e-09 |
| Malignant epithelial | C2 WNT ECs | MCCC1 | 0.708 | 0.134 | 0.065 | 1.51e-10 | 5.09e-06 |
| Malignant epithelial | C2 WNT ECs | TRMT13 | 0.708 | 0.112 | 0.051 | 3.48e-10 | 1.17e-05 |
| Malignant epithelial | C2 WNT ECs | SPINT1 | 0.707 | 0.88 | 0.635 | 1.29e-45 | 4.34e-41 |
| Malignant epithelial | C2 WNT ECs | HNRNPA0 | 0.707 | 0.88 | 0.693 | 1.29e-44 | 4.35e-40 |
| Malignant epithelial | C2 WNT ECs | VASP | 0.707 | 0.763 | 0.538 | 1.08e-35 | 3.63e-31 |
| Malignant epithelial | C2 WNT ECs | CYTH2 | 0.707 | 0.639 | 0.399 | 7.65e-31 | 2.58e-26 |
| Malignant epithelial | C2 WNT ECs | SMAP1 | 0.707 | 0.478 | 0.276 | 9.010000000000001e-26 | 3.04e-21 |
| Malignant epithelial | C2 WNT ECs | PIH1D1 | 0.707 | 0.53 | 0.319 | 2.22e-25 | 7.47e-21 |

| compartment | subcluster | gene | avg_log2FC | pct.1 | pct.2 | p_val | p_val_adj |
|---|---|---|---|---|---|---|---|
| Malignant epithelial | C2 WNT ECs | ITGB1 | 0.705 | 0.835 | 0.613 | 1.49e-35 | 5.03e-31 |
| Malignant epithelial | C2 WNT ECs | CDC123 | 0.705 | 0.575 | 0.336 | 5.039999999999999e-30 | 1.7e-25 |
| Malignant epithelial | C2 WNT ECs | TES | 0.705 | 0.59 | 0.389 | 3.91e-24 | 1.32e-19 |
| Malignant epithelial | C2 WNT ECs | SLC6A6 | 0.705 | 0.161 | 0.086 | 4.62e-10 | 1.56e-05 |
| Malignant epithelial | C2 WNT ECs | RFWD3 | 0.705 | 0.122 | 0.066 | 1.54e-07 | 0.00517 |
| Malignant epithelial | C2 WNT ECs | AC093323.3 | 0.704 | 0.134 | 0.073 | 3.63e-08 | 0.00122 |
| Malignant epithelial | C2 WNT ECs | ACTN1 | 0.703 | 0.56 | 0.349 | 4.72e-26 | 1.59e-21 |
| Malignant epithelial | C2 WNT ECs | TTC32 | 0.703 | 0.204 | 0.109 | 5.54e-12 | 1.87e-07 |
| Malignant epithelial | C2 WNT ECs | STXBP1 | 0.703 | 0.131 | 0.073 | 1.78e-07 | 0.00599 |
| Malignant epithelial | C2 WNT ECs | NET1 | 0.702 | 0.767 | 0.54 | 4.33e-32 | 1.46e-27 |
| Malignant epithelial | C2 WNT ECs | RERE | 0.702 | 0.33 | 0.157 | 8.18e-26 | 2.76e-21 |
| Malignant epithelial | C2 WNT ECs | TYW3 | 0.702 | 0.231 | 0.12 | 4.74e-15 | 1.6e-10 |
| Malignant epithelial | C2 WNT ECs | SF3A1 | 0.701 | 0.311 | 0.161 | 4.65e-20 | 1.57e-15 |
| Malignant epithelial | C2 WNT ECs | SH3BP5L | 0.701 | 0.103 | 0.05 | 1.94e-08 | 0.000654 |
| Malignant epithelial | C2 WNT ECs | POLR2C | 0.7 | 0.616 | 0.405 | 2.6e-26 | 8.77e-22 |
| Malignant epithelial | C2 WNT ECs | PSMC2 | 0.699 | 0.513 | 0.299 | 4.4e-26 | 1.48e-21 |
| Malignant epithelial | C2 WNT ECs | RPIA | 0.699 | 0.448 | 0.282 | 8.75e-19 | 2.95e-14 |
| Malignant epithelial | C2 WNT ECs | COG7 | 0.699 | 0.153 | 0.069 | 9.89e-14 | 3.33e-09 |
| Malignant epithelial | C2 WNT ECs | BMS1 | 0.699 | 0.199 | 0.105 | 1.52e-12 | 5.14e-08 |
| Malignant epithelial | C2 WNT ECs | CREM | 0.698 | 0.352 | 0.197 | 2.19e-18 | 7.38e-14 |
| Malignant epithelial | C2 WNT ECs | ATF1 | 0.698 | 0.31 | 0.167 | 2.93e-18 | 9.87e-14 |
| Malignant epithelial | C2 WNT ECs | WTAP | 0.697 | 0.613 | 0.362 | 1.59e-32 | 5.37e-28 |
| Malignant epithelial | C2 WNT ECs | ZBTB7A | 0.697 | 0.582 | 0.348 | 1.54e-28 | 5.19e-24 |
| Malignant epithelial | C2 WNT ECs | SUPT7L | 0.697 | 0.204 | 0.102 | 1.91e-14 | 6.43e-10 |
| Malignant epithelial | C2 WNT ECs | SPAG1 | 0.697 | 0.2 | 0.113 | 1.48e-10 | 5e-06 |
| Malignant epithelial | C2 WNT ECs | MUC20 | 0.697 | 0.138 | 0.068 | 2.1e-10 | 7.07e-06 |
| Malignant epithelial | C2 WNT ECs | XPC | 0.697 | 0.146 | 0.081 | 2.43e-08 | 0.000819 |
| Malignant epithelial | C2 WNT ECs | FUBP1 | 0.696 | 0.355 | 0.193 | 2.88e-20 | 9.7e-16 |
| Malignant epithelial | C2 WNT ECs | TMEM115 | 0.696 | 0.333 | 0.178 | 2.92e-19 | 9.86e-15 |
| Malignant epithelial | C2 WNT ECs | CCDC84 | 0.696 | 0.189 | 0.107 | 4.39e-10 | 1.48e-05 |
| Malignant epithelial | C2 WNT ECs | DYRK1B | 0.696 | 0.15 | 0.081 | 3.68e-09 | 0.000124 |
| Malignant epithelial | C2 WNT ECs | DDX19B | 0.695 | 0.38 | 0.211 | 5.02e-21 | 1.69e-16 |
| Malignant epithelial | C2 WNT ECs | FEM1C | 0.695 | 0.277 | 0.154 | 5.22e-15 | 1.76e-10 |
| Malignant epithelial | C2 WNT ECs | SLC26A6 | 0.695 | 0.214 | 0.111 | 6.96e-14 | 2.34e-09 |
| Malignant epithelial | C2 WNT ECs | RIOK2 | 0.695 | 0.141 | 0.071 | 2.84e-10 | 9.56e-06 |
| Malignant epithelial | C2 WNT ECs | MACC1 | 0.694 | 0.361 | 0.178 | 2.39e-26 | 8.04e-22 |
| Malignant epithelial | C2 WNT ECs | ITGA6 | 0.694 | 0.667 | 0.45 | 1.36e-25 | 4.58e-21 |
| Malignant epithelial | C2 WNT ECs | CMTM8 | 0.694 | 0.509 | 0.296 | 1.45e-25 | 4.89e-21 |
| Malignant epithelial | C2 WNT ECs | SLC39A7 | 0.694 | 0.405 | 0.23 | 3.35e-21 | 1.13e-16 |
| Malignant epithelial | C2 WNT ECs | TMEM222 | 0.694 | 0.279 | 0.145 | 1.09e-17 | 3.67e-13 |
| Malignant epithelial | C2 WNT ECs | DNAJB9 | 0.693 | 0.34 | 0.197 | 6.6e-18 | 2.22e-13 |
| Malignant epithelial | C2 WNT ECs | SYF2 | 0.692 | 0.827 | 0.567 | 3.73e-48 | 1.26e-43 |
| Malignant epithelial | C2 WNT ECs | MTCH1 | 0.692 | 0.809 | 0.588 | 3.88e-35 | 1.31e-30 |
| Malignant epithelial | C2 WNT ECs | NRDC | 0.692 | 0.309 | 0.167 | 8.35e-18 | 2.81e-13 |
| Malignant epithelial | C2 WNT ECs | TNFRSF10B | 0.692 | 0.218 | 0.119 | 9.84e-13 | 3.32e-08 |
| Malignant epithelial | C2 WNT ECs | ADCK3 | 0.692 | 0.231 | 0.139 | 1.29e-10 | 4.33e-06 |
| Malignant epithelial | C2 WNT ECs | RAD50 | 0.691 | 0.415 | 0.24 | 1.06e-21 | 3.57e-17 |
| Malignant epithelial | C2 WNT ECs | SVIP | 0.69 | 0.659 | 0.412 | 5.84e-32 | 1.97e-27 |
| Malignant epithelial | C2 WNT ECs | VPS54 | 0.69 | 0.107 | 0.054 | 7.02e-08 | 0.00236 |
| Malignant epithelial | C2 WNT ECs | MAP7 | 0.689 | 0.373 | 0.204 | 8.28e-21 | 2.79e-16 |
| Malignant epithelial | C2 WNT ECs | AARS | 0.689 | 0.267 | 0.138 | 2.21e-17 | 7.45e-13 |
| Malignant epithelial | C2 WNT ECs | PTPN3 | 0.689 | 0.219 | 0.118 | 6.64e-13 | 2.24e-08 |
| Malignant epithelial | C2 WNT ECs | WIPI2 | 0.688 | 0.609 | 0.368 | 1.82e-31 | 6.14e-27 |
| Malignant epithelial | C2 WNT ECs | RBM7 | 0.688 | 0.456 | 0.266 | 2.26e-21 | 7.62e-17 |
| Malignant epithelial | C2 WNT ECs | UBE4B | 0.688 | 0.146 | 0.063 | 1.11e-14 | 3.74e-10 |
| Malignant epithelial | C2 WNT ECs | SH2B1 | 0.688 | 0.2 | 0.101 | 1.17e-13 | 3.94e-09 |
| Malignant epithelial | C2 WNT ECs | CLDN1 | 0.687 | 0.501 | 0.282 | 4.22e-28 | 1.42e-23 |
| Malignant epithelial | C2 WNT ECs | DNM2 | 0.687 | 0.509 | 0.287 | 6.17e-28 | 2.08e-23 |
| Malignant epithelial | C2 WNT ECs | SEC13 | 0.687 | 0.455 | 0.25 | 2.09e-26 | 7.05e-22 |
| Malignant epithelial | C2 WNT ECs | TDP2 | 0.687 | 0.246 | 0.115 | 3.47e-20 | 1.17e-15 |
| Malignant epithelial | C2 WNT ECs | DNTTIP2 | 0.687 | 0.31 | 0.161 | 1.04e-19 | 3.5e-15 |
| Malignant epithelial | C2 WNT ECs | N4BP1 | 0.687 | 0.185 | 0.101 | 1.22e-10 | 4.11e-06 |
| Malignant epithelial | C2 WNT ECs | DUSP8 | 0.687 | 0.168 | 0.089 | 1.66e-10 | 5.58e-06 |
| Malignant epithelial | C2 WNT ECs | RPL5 | 0.686 | 0.968 | 0.956 | 7.03e-61 | 2.37e-56 |

| compartment | subcluster | gene | avg_log2FC | pct.1 | pct.2 | p_val | p_val_adj |
|---|---|---|---|---|---|---|---|
| Malignant epithelial | C2 WNT ECs | ATE1 | 0.686 | 0.161 | 0.083 | 1.1e-10 | 3.7e-06 |
| Malignant epithelial | C2 WNT ECs | RP11-51J9.5 | 0.686 | 0.123 | 0.061 | 1.5e-09 | 5.07e-05 |
| Malignant epithelial | C2 WNT ECs | GABARAPL2 | 0.685 | 0.859 | 0.652 | 9.139999999999999e-42 | 3.08e-37 |
| Malignant epithelial | C2 WNT ECs | ATP6V0D1 | 0.685 | 0.567 | 0.335 | 1.19e-28 | 4.02e-24 |
| Malignant epithelial | C2 WNT ECs | RP4-568C11.4 | 0.685 | 0.225 | 0.13 | 4.99e-11 | 1.68e-06 |
| Malignant epithelial | C2 WNT ECs | VPS35 | 0.684 | 0.612 | 0.378 | 1.03e-29 | 3.46e-25 |
| Malignant epithelial | C2 WNT ECs | HGH1 | 0.683 | 0.169 | 0.08 | 1.88e-13 | 6.32e-09 |
| Malignant epithelial | C2 WNT ECs | EEA1 | 0.683 | 0.244 | 0.136 | 3.2e-13 | 1.08e-08 |
| Malignant epithelial | C2 WNT ECs | LTBP4 | 0.683 | 0.392 | 0.274 | 6.44e-11 | 2.17e-06 |
| Malignant epithelial | C2 WNT ECs | CD164 | 0.682 | 0.896 | 0.713 | 3.38e-40 | 1.14e-35 |
| Malignant epithelial | C2 WNT ECs | TMED4 | 0.682 | 0.709 | 0.443 | 6.94e-36 | 2.34e-31 |
| Malignant epithelial | C2 WNT ECs | MALAT1 | 0.682 | 0.997 | 0.998 | 8.08e-32 | 2.72e-27 |
| Malignant epithelial | C2 WNT ECs | MYO1D | 0.682 | 0.304 | 0.159 | 6.94e-19 | 2.34e-14 |
| Malignant epithelial | C2 WNT ECs | PRR15 | 0.681 | 0.456 | 0.249 | 4.87e-27 | 1.64e-22 |
| Malignant epithelial | C2 WNT ECs | PSMD2 | 0.681 | 0.522 | 0.314 | 1.39e-25 | 4.680000000000001e-21 |
| Malignant epithelial | C2 WNT ECs | C18orf32 | 0.681 | 0.253 | 0.121 | 3.45e-19 | 1.16e-14 |
| Malignant epithelial | C2 WNT ECs | GRK6 | 0.681 | 0.198 | 0.099 | 9.7e-14 | 3.27e-09 |
| Malignant epithelial | C2 WNT ECs | RPA1 | 0.681 | 0.185 | 0.097 | 8.78e-12 | 2.96e-07 |
| Malignant epithelial | C2 WNT ECs | ATXN10 | 0.68 | 0.424 | 0.241 | 2.13e-22 | 7.19e-18 |
| Malignant epithelial | C2 WNT ECs | DDA1 | 0.68 | 0.54 | 0.364 | 1.99e-19 | 6.69e-15 |
| Malignant epithelial | C2 WNT ECs | RBL2 | 0.68 | 0.227 | 0.106 | 6.29e-19 | 2.12e-14 |
| Malignant epithelial | C2 WNT ECs | HOXA10 | 0.68 | 0.494 | 0.324 | 2.18e-17 | 7.35e-13 |
| Malignant epithelial | C2 WNT ECs | FOXJ3 | 0.68 | 0.145 | 0.07 | 2.69e-11 | 9.08e-07 |
| Malignant epithelial | C2 WNT ECs | CENPC | 0.68 | 0.114 | 0.054 | 1.48e-09 | 5e-05 |
| Malignant epithelial | C2 WNT ECs | GCAT | 0.68 | 0.138 | 0.072 | 5.66e-09 | 0.000191 |
| Malignant epithelial | C2 WNT ECs | ACTR3 | 0.679 | 0.714 | 0.474 | 6.88e-33 | 2.32e-28 |
| Malignant epithelial | C2 WNT ECs | PRPF8 | 0.679 | 0.249 | 0.127 | 8.57e-17 | 2.89e-12 |
| Malignant epithelial | C2 WNT ECs | ARHGEF5 | 0.679 | 0.292 | 0.164 | 3.96e-15 | 1.33e-10 |
| Malignant epithelial | C2 WNT ECs | GALNS | 0.679 | 0.188 | 0.097 | 1.56e-12 | 5.25e-08 |
| Malignant epithelial | C2 WNT ECs | QDPR | 0.678 | 0.341 | 0.18 | 9.260000000000001e-21 | 3.12e-16 |
| Malignant epithelial | C2 WNT ECs | MAPKAPK3 | 0.678 | 0.344 | 0.205 | 5.03e-16 | 1.7e-11 |
| Malignant epithelial | C2 WNT ECs | C1orf27 | 0.678 | 0.196 | 0.099 | 1.45e-13 | 4.89e-09 |
| Malignant epithelial | C2 WNT ECs | ZBED5-AS1 | 0.678 | 0.198 | 0.106 | 3.91e-12 | 1.32e-07 |
| Malignant epithelial | C2 WNT ECs | TSN | 0.677 | 0.438 | 0.263 | 1.38e-20 | 4.67e-16 |
| Malignant epithelial | C2 WNT ECs | ATP6V0A2 | 0.677 | 0.101 | 0.047 | 3.44e-09 | 0.000116 |
| Malignant epithelial | C2 WNT ECs | CMTM6 | 0.676 | 0.654 | 0.434 | 1.18e-29 | 3.97e-25 |
| Malignant epithelial | C2 WNT ECs | CCDC47 | 0.675 | 0.438 | 0.262 | 4.77e-21 | 1.61e-16 |
| Malignant epithelial | C2 WNT ECs | BDH2 | 0.675 | 0.187 | 0.092 | 1.15e-13 | 3.88e-09 |
| Malignant epithelial | C2 WNT ECs | SPRED2 | 0.675 | 0.164 | 0.085 | 7.74e-11 | 2.61e-06 |
| Malignant epithelial | C2 WNT ECs | GPC1 | 0.675 | 0.212 | 0.129 | 5.32e-09 | 0.000179 |
| Malignant epithelial | C2 WNT ECs | ZYX | 0.674 | 0.457 | 0.259 | 2.47e-24 | 8.32e-20 |
| Malignant epithelial | C2 WNT ECs | NFRKB | 0.674 | 0.154 | 0.075 | 7.07e-12 | 2.38e-07 |
| Malignant epithelial | C2 WNT ECs | TEAD1 | 0.674 | 0.191 | 0.105 | 9.8e-11 | 3.3e-06 |
| Malignant epithelial | C2 WNT ECs | PLEKHG5 | 0.674 | 0.103 | 0.048 | 7.37e-09 | 0.000248 |
| Malignant epithelial | C2 WNT ECs | MALT1 | 0.674 | 0.129 | 0.067 | 1.16e-08 | 0.00039 |
| Malignant epithelial | C2 WNT ECs | MEF2D | 0.674 | 0.208 | 0.127 | 1.43e-08 | 0.000481 |
| Malignant epithelial | C2 WNT ECs | CALU | 0.673 | 0.529 | 0.32 | 8.119999999999999e-24 | 2.73e-19 |
| Malignant epithelial | C2 WNT ECs | ITFG1 | 0.673 | 0.399 | 0.24 | 9.37e-19 | 3.16e-14 |
| Malignant epithelial | C2 WNT ECs | SLC35A2 | 0.673 | 0.345 | 0.197 | 1.04e-17 | 3.51e-13 |
| Malignant epithelial | C2 WNT ECs | CBL | 0.673 | 0.123 | 0.056 | 3.16e-11 | 1.07e-06 |
| Malignant epithelial | C2 WNT ECs | SEC11C | 0.672 | 0.586 | 0.369 | 1.14e-26 | 3.83e-22 |
| Malignant epithelial | C2 WNT ECs | HSPA4 | 0.672 | 0.463 | 0.261 | 4.27e-24 | 1.44e-19 |
| Malignant epithelial | C2 WNT ECs | RAB10 | 0.672 | 0.708 | 0.507 | 2.26e-21 | 7.63e-17 |
| Malignant epithelial | C2 WNT ECs | RAB2A | 0.67 | 0.919 | 0.742 | 1.32e-45 | 4.43e-41 |
| Malignant epithelial | C2 WNT ECs | SF3B1 | 0.67 | 0.62 | 0.399 | 1.64e-25 | 5.51e-21 |
| Malignant epithelial | C2 WNT ECs | IRF6 | 0.67 | 0.396 | 0.211 | 7.22e-24 | 2.43e-19 |
| Malignant epithelial | C2 WNT ECs | EIF2B3 | 0.67 | 0.107 | 0.056 | 3.85e-07 | 0.013 |
| Malignant epithelial | C2 WNT ECs | C8orf82 | 0.669 | 0.512 | 0.318 | 4.45e-22 | 1.5e-17 |
| Malignant epithelial | C2 WNT ECs | CHTF8 | 0.669 | 0.36 | 0.207 | 1.1e-18 | 3.71e-14 |
| Malignant epithelial | C2 WNT ECs | LRRC47 | 0.669 | 0.284 | 0.153 | 3.07e-16 | 1.03e-11 |
| Malignant epithelial | C2 WNT ECs | FSCN1 | 0.669 | 0.488 | 0.36 | 9.25e-13 | 3.12e-08 |
| Malignant epithelial | C2 WNT ECs | MYO9B | 0.669 | 0.214 | 0.124 | 1.47e-10 | 4.96e-06 |
| Malignant epithelial | C2 WNT ECs | USP38 | 0.669 | 0.15 | 0.077 | 2.19e-10 | 7.39e-06 |
| Malignant epithelial | C2 WNT ECs | MROH1 | 0.668 | 0.309 | 0.17 | 3.03e-17 | 1.02e-12 |

| compartment | subcluster | gene | avg_log2FC | pct.1 | pct.2 | p_val | p_val_adj |
|---|---|---|---|---|---|---|---|
| Malignant epithelial | C2 WNT ECs | DERA | 0.668 | 0.39 | 0.231 | 3.77e-17 | 1.27e-12 |
| Malignant epithelial | C2 WNT ECs | SCAF4 | 0.668 | 0.321 | 0.182 | 1.07e-15 | 3.59e-11 |
| Malignant epithelial | C2 WNT ECs | DAPK3 | 0.668 | 0.294 | 0.165 | 2.53e-15 | 8.54e-11 |
| Malignant epithelial | C2 WNT ECs | ST6GAL1 | 0.668 | 0.138 | 0.067 | 1.18e-10 | 3.96e-06 |
| Malignant epithelial | C2 WNT ECs | HSPBAP1 | 0.668 | 0.116 | 0.054 | 5.77e-10 | 1.94e-05 |
| Malignant epithelial | C2 WNT ECs | CDR2 | 0.668 | 0.138 | 0.072 | 2.71e-09 | 9.15e-05 |
| Malignant epithelial | C2 WNT ECs | ZFAND3 | 0.667 | 0.424 | 0.234 | 1.08e-23 | 3.65e-19 |
| Malignant epithelial | C2 WNT ECs | MEST | 0.667 | 0.396 | 0.267 | 3.84e-13 | 1.29e-08 |
| Malignant epithelial | C2 WNT ECs | SNX14 | 0.666 | 0.342 | 0.176 | 3.47e-22 | 1.17e-17 |
| Malignant epithelial | C2 WNT ECs | FAM200B | 0.666 | 0.298 | 0.159 | 4.48e-18 | 1.51e-13 |
| Malignant epithelial | C2 WNT ECs | WAC-AS1 | 0.666 | 0.206 | 0.12 | 5.74e-10 | 1.93e-05 |
| Malignant epithelial | C2 WNT ECs | DHTKD1 | 0.666 | 0.126 | 0.067 | 3.84e-08 | 0.00129 |
| Malignant epithelial | C2 WNT ECs | TPP1 | 0.665 | 0.387 | 0.228 | 4.7e-18 | 1.58e-13 |
| Malignant epithelial | C2 WNT ECs | SUMF1 | 0.665 | 0.242 | 0.125 | 1.67e-15 | 5.63e-11 |
| Malignant epithelial | C2 WNT ECs | PXN | 0.665 | 0.407 | 0.266 | 1.53e-14 | 5.16e-10 |
| Malignant epithelial | C2 WNT ECs | FBXL5 | 0.665 | 0.162 | 0.087 | 6.7e-10 | 2.26e-05 |
| Malignant epithelial | C2 WNT ECs | RAB20 | 0.664 | 0.359 | 0.199 | 2.92e-19 | 9.84e-15 |
| Malignant epithelial | C2 WNT ECs | TRIP12 | 0.664 | 0.272 | 0.156 | 4.02e-13 | 1.36e-08 |
| Malignant epithelial | C2 WNT ECs | NAGK | 0.664 | 0.271 | 0.158 | 5.89e-13 | 1.99e-08 |
| Malignant epithelial | C2 WNT ECs | RBM3 | 0.663 | 0.922 | 0.792 | 4.44e-52 | 1.5e-47 |
| Malignant epithelial | C2 WNT ECs | NOL7 | 0.663 | 0.687 | 0.436 | 8.99e-35 | 3.03e-30 |
| Malignant epithelial | C2 WNT ECs | CD55 | 0.663 | 0.59 | 0.386 | 4.22e-24 | 1.42e-19 |
| Malignant epithelial | C2 WNT ECs | PDSS2 | 0.663 | 0.137 | 0.075 | 6.68e-08 | 0.00225 |
| Malignant epithelial | C2 WNT ECs | KCTD17 | 0.663 | 0.124 | 0.071 | 6.62e-07 | 0.0223 |
| Malignant epithelial | C2 WNT ECs | FABP1 | 0.662 | 0.188 | 0.068 | 3.84e-27 | 1.29e-22 |
| Malignant epithelial | C2 WNT ECs | SCAMP2 | 0.662 | 0.414 | 0.238 | 8.95e-20 | 3.02e-15 |
| Malignant epithelial | C2 WNT ECs | SMOX | 0.662 | 0.264 | 0.163 | 1.94e-10 | 6.52e-06 |
| Malignant epithelial | C2 WNT ECs | EPCAM | 0.661 | 0.986 | 0.956 | 2.81e-52 | 9.479999999999999e-48 |
| Malignant epithelial | C2 WNT ECs | VAPA | 0.661 | 0.788 | 0.56 | 5.52e-35 | 1.86e-30 |
| Malignant epithelial | C2 WNT ECs | ACTR1A | 0.661 | 0.407 | 0.229 | 1.35e-20 | 4.56e-16 |
| Malignant epithelial | C2 WNT ECs | DNAJC5 | 0.661 | 0.313 | 0.199 | 2.18e-11 | 7.33e-07 |
| Malignant epithelial | C2 WNT ECs | DDX27 | 0.66 | 0.613 | 0.4 | 4.97e-27 | 1.68e-22 |
| Malignant epithelial | C2 WNT ECs | PLXNB1 | 0.66 | 0.336 | 0.177 | 1.26e-20 | 4.26e-16 |
| Malignant epithelial | C2 WNT ECs | SLC25A33 | 0.66 | 0.296 | 0.167 | 5.5e-15 | 1.85e-10 |
| Malignant epithelial | C2 WNT ECs | MAP2K6 | 0.66 | 0.145 | 0.075 | 1.67e-09 | 5.63e-05 |
| Malignant epithelial | C2 WNT ECs | HNF1A | 0.66 | 0.103 | 0.052 | 6.65e-08 | 0.00224 |
| Malignant epithelial | C2 WNT ECs | ARGLU1 | 0.659 | 0.823 | 0.63 | 2.94e-31 | 9.920000000000001e-27 |
| Malignant epithelial | C2 WNT ECs | RCN2 | 0.659 | 0.604 | 0.391 | 1.28e-24 | 4.31e-20 |
| Malignant epithelial | C2 WNT ECs | ITGAE | 0.659 | 0.318 | 0.161 | 2.89e-21 | 9.74e-17 |
| Malignant epithelial | C2 WNT ECs | CDK10 | 0.659 | 0.442 | 0.264 | 1.97e-20 | 6.64e-16 |
| Malignant epithelial | C2 WNT ECs | NASP | 0.659 | 0.478 | 0.3 | 2.5e-19 | 8.43e-15 |
| Malignant epithelial | C2 WNT ECs | SOS1 | 0.659 | 0.123 | 0.067 | 1.28e-07 | 0.00431 |
| Malignant epithelial | C2 WNT ECs | CNOT2 | 0.658 | 0.464 | 0.287 | 1.34e-18 | 4.53e-14 |
| Malignant epithelial | C2 WNT ECs | JMJD6 | 0.658 | 0.36 | 0.208 | 1.77e-17 | 5.95e-13 |
| Malignant epithelial | C2 WNT ECs | BOLA2B | 0.658 | 0.194 | 0.099 | 5.71e-13 | 1.93e-08 |
| Malignant epithelial | C2 WNT ECs | ID2 | 0.657 | 0.888 | 0.751 | 5.44e-29 | 1.83e-24 |
| Malignant epithelial | C2 WNT ECs | TADA3 | 0.657 | 0.421 | 0.246 | 4.9e-21 | 1.65e-16 |
| Malignant epithelial | C2 WNT ECs | CCNG2 | 0.657 | 0.231 | 0.122 | 4.24e-14 | 1.43e-09 |
| Malignant epithelial | C2 WNT ECs | ACSS1 | 0.657 | 0.231 | 0.13 | 2.61e-12 | 8.79e-08 |
| Malignant epithelial | C2 WNT ECs | EDEM1 | 0.657 | 0.112 | 0.05 | 6.7e-11 | 2.26e-06 |
| Malignant epithelial | C2 WNT ECs | LAMTOR5 | 0.656 | 0.881 | 0.669 | 6.62e-51 | 2.23e-46 |
| Malignant epithelial | C2 WNT ECs | STX5 | 0.656 | 0.382 | 0.215 | 3.56e-20 | 1.2e-15 |
| Malignant epithelial | C2 WNT ECs | PAFAH1B1 | 0.656 | 0.338 | 0.193 | 2.04e-17 | 6.86e-13 |
| Malignant epithelial | C2 WNT ECs | POLR3E | 0.656 | 0.244 | 0.127 | 3.03e-15 | 1.02e-10 |
| Malignant epithelial | C2 WNT ECs | PRKD2 | 0.656 | 0.28 | 0.159 | 7.72e-15 | 2.6e-10 |
| Malignant epithelial | C2 WNT ECs | OGG1 | 0.656 | 0.139 | 0.064 | 5.87e-12 | 1.98e-07 |
| Malignant epithelial | C2 WNT ECs | AAMP | 0.655 | 0.717 | 0.485 | 2.08e-30 | 7.020000000000001e-26 |
| Malignant epithelial | C2 WNT ECs | ASF1A | 0.655 | 0.238 | 0.135 | 3.25e-12 | 1.1e-07 |
| Malignant epithelial | C2 WNT ECs | TXNDC15 | 0.655 | 0.217 | 0.124 | 8.59e-11 | 2.89e-06 |
| Malignant epithelial | C2 WNT ECs | TMX3 | 0.655 | 0.112 | 0.059 | 1.94e-07 | 0.00653 |
| Malignant epithelial | C2 WNT ECs | JAK1 | 0.654 | 0.367 | 0.201 | 1.71e-20 | 5.76e-16 |
| Malignant epithelial | C2 WNT ECs | CERS6 | 0.654 | 0.357 | 0.216 | 2.82e-15 | 9.5e-11 |
| Malignant epithelial | C2 WNT ECs | NDUFS1 | 0.654 | 0.269 | 0.153 | 6.96e-14 | 2.34e-09 |
| Malignant epithelial | C2 WNT ECs | OSER1 | 0.653 | 0.582 | 0.371 | 2.42e-23 | 8.15e-19 |

| compartment | subcluster | gene | avg_log2FC | pct.1 | pct.2 | p_val | p_val_adj |
|---|---|---|---|---|---|---|---|
| Malignant epithelial | C2 WNT ECs | SLC35A5 | 0.653 | 0.107 | 0.052 | 1.22e-08 | 0.000412 |
| Malignant epithelial | C2 WNT ECs | PPP2CA | 0.652 | 0.613 | 0.411 | 2.09e-24 | 7.04e-20 |
| Malignant epithelial | C2 WNT ECs | LLGL2 | 0.652 | 0.593 | 0.385 | 8.719999999999999e-23 | 2.94e-18 |
| Malignant epithelial | C2 WNT ECs | NECAP2 | 0.652 | 0.417 | 0.237 | 1.5e-20 | 5.05e-16 |
| Malignant epithelial | C2 WNT ECs | NDEL1 | 0.652 | 0.187 | 0.098 | 9.97e-12 | 3.36e-07 |
| Malignant epithelial | C2 WNT ECs | PPP1R8 | 0.652 | 0.195 | 0.105 | 1.35e-11 | 4.53e-07 |
| Malignant epithelial | C2 WNT ECs | PLCG1 | 0.652 | 0.11 | 0.057 | 1.01e-07 | 0.00341 |
| Malignant epithelial | C2 WNT ECs | HRH1 | 0.652 | 0.103 | 0.053 | 2.97e-07 | 0.01 |
| Malignant epithelial | C2 WNT ECs | MLYCD | 0.652 | 0.119 | 0.065 | 3.44e-07 | 0.0116 |
| Malignant epithelial | C2 WNT ECs | MYPOP | 0.652 | 0.11 | 0.059 | 9.79e-07 | 0.033 |
| Malignant epithelial | C2 WNT ECs | KMT2E | 0.651 | 0.724 | 0.483 | 2.23e-30 | 7.51e-26 |
| Malignant epithelial | C2 WNT ECs | STK24 | 0.651 | 0.552 | 0.394 | 7.13e-17 | 2.4e-12 |
| Malignant epithelial | C2 WNT ECs | PSMG1 | 0.651 | 0.426 | 0.284 | 8.39e-14 | 2.83e-09 |
| Malignant epithelial | C2 WNT ECs | ADNP | 0.651 | 0.35 | 0.222 | 1.65e-12 | 5.55e-08 |
| Malignant epithelial | C2 WNT ECs | DPP9 | 0.651 | 0.249 | 0.147 | 2.2e-11 | 7.42e-07 |
| Malignant epithelial | C2 WNT ECs | NR6A1 | 0.651 | 0.221 | 0.127 | 1.29e-10 | 4.35e-06 |
| Malignant epithelial | C2 WNT ECs | GFPT1 | 0.65 | 0.387 | 0.203 | 4.15e-24 | 1.4e-19 |
| Malignant epithelial | C2 WNT ECs | HEXIM1 | 0.65 | 0.571 | 0.379 | 4.41e-21 | 1.48e-16 |
| Malignant epithelial | C2 WNT ECs | CAB39 | 0.65 | 0.355 | 0.199 | 2.45e-18 | 8.26e-14 |
| Malignant epithelial | C2 WNT ECs | YIF1B | 0.65 | 0.252 | 0.13 | 7.66e-17 | 2.58e-12 |
| Malignant epithelial | C2 WNT ECs | BYSL | 0.65 | 0.242 | 0.127 | 8.66e-15 | 2.92e-10 |
| Malignant epithelial | C2 WNT ECs | ZNF766 | 0.65 | 0.177 | 0.088 | 3.68e-13 | 1.24e-08 |
| Malignant epithelial | C2 WNT ECs | HSDL2 | 0.65 | 0.225 | 0.132 | 2.84e-10 | 9.56e-06 |
| Malignant epithelial | C2 WNT ECs | NOL10 | 0.65 | 0.189 | 0.107 | 8.87e-10 | 2.99e-05 |
| Malignant epithelial | C2 WNT ECs | PSPH | 0.65 | 0.172 | 0.101 | 3.46e-08 | 0.00117 |
| Malignant epithelial | C2 WNT ECs | TPGS1 | 0.649 | 0.365 | 0.23 | 4.03e-14 | 1.36e-09 |
| Malignant epithelial | C2 WNT ECs | HOXA7 | 0.649 | 0.217 | 0.12 | 1.05e-11 | 3.55e-07 |
| Malignant epithelial | C2 WNT ECs | KLHDC10 | 0.649 | 0.164 | 0.084 | 3.68e-11 | 1.24e-06 |
| Malignant epithelial | C2 WNT ECs | ATP6AP2 | 0.648 | 0.663 | 0.442 | 1.6e-27 | 5.38e-23 |
| Malignant epithelial | C2 WNT ECs | PDCD6IP | 0.648 | 0.518 | 0.31 | 5.06e-25 | 1.71e-20 |
| Malignant epithelial | C2 WNT ECs | ZNF195 | 0.648 | 0.219 | 0.117 | 3.4e-13 | 1.15e-08 |
| Malignant epithelial | C2 WNT ECs | THAP1 | 0.648 | 0.203 | 0.11 | 8.38e-12 | 2.82e-07 |
| Malignant epithelial | C2 WNT ECs | GANC | 0.648 | 0.104 | 0.05 | 8.16e-09 | 0.000275 |
| Malignant epithelial | C2 WNT ECs | ARHGEF3 | 0.648 | 0.134 | 0.074 | 1.03e-07 | 0.00347 |
| Malignant epithelial | C2 WNT ECs | RBM47 | 0.647 | 0.375 | 0.197 | 8.62e-24 | 2.9e-19 |
| Malignant epithelial | C2 WNT ECs | RNF170 | 0.647 | 0.311 | 0.175 | 3.46e-16 | 1.17e-11 |
| Malignant epithelial | C2 WNT ECs | GMCL1 | 0.647 | 0.275 | 0.164 | 2.62e-12 | 8.84e-08 |
| Malignant epithelial | C2 WNT ECs | RNF168 | 0.647 | 0.175 | 0.09 | 1.12e-11 | 3.77e-07 |
| Malignant epithelial | C2 WNT ECs | OSER1-AS1 | 0.647 | 0.158 | 0.08 | 3.23e-11 | 1.09e-06 |
| Malignant epithelial | C2 WNT ECs | HERC4 | 0.647 | 0.352 | 0.239 | 1.87e-10 | 6.28e-06 |
| Malignant epithelial | C2 WNT ECs | HNRNPL | 0.646 | 0.491 | 0.264 | 5.4e-30 | 1.82e-25 |
| Malignant epithelial | C2 WNT ECs | GALE | 0.646 | 0.449 | 0.249 | 8.5e-25 | 2.87e-20 |
| Malignant epithelial | C2 WNT ECs | ZNF451 | 0.646 | 0.314 | 0.152 | 2.62e-23 | 8.82e-19 |
| Malignant epithelial | C2 WNT ECs | PTPRF | 0.646 | 0.475 | 0.284 | 2.76e-22 | 9.3e-18 |
| Malignant epithelial | C2 WNT ECs | TNIP2 | 0.646 | 0.321 | 0.163 | 5.15e-21 | 1.73e-16 |
| Malignant epithelial | C2 WNT ECs | IL17RE | 0.646 | 0.337 | 0.185 | 1.87e-18 | 6.29e-14 |
| Malignant epithelial | C2 WNT ECs | PDHX | 0.645 | 0.196 | 0.103 | 3.03e-12 | 1.02e-07 |
| Malignant epithelial | C2 WNT ECs | TSPAN14 | 0.644 | 0.401 | 0.238 | 9.7e-18 | 3.27e-13 |
| Malignant epithelial | C2 WNT ECs | C16orf58 | 0.643 | 0.245 | 0.128 | 2.83e-15 | 9.53e-11 |
| Malignant epithelial | C2 WNT ECs | ATG13 | 0.643 | 0.227 | 0.124 | 9.17e-13 | 3.09e-08 |
| Malignant epithelial | C2 WNT ECs | WRB | 0.643 | 0.226 | 0.128 | 2.31e-11 | 7.77e-07 |
| Malignant epithelial | C2 WNT ECs | POU2F1 | 0.643 | 0.165 | 0.09 | 4.63e-10 | 1.56e-05 |
| Malignant epithelial | C2 WNT ECs | SRF | 0.643 | 0.179 | 0.1 | 1.43e-09 | 4.82e-05 |
| Malignant epithelial | C2 WNT ECs | TUBGCP3 | 0.643 | 0.124 | 0.066 | 2.96e-08 | 0.000996 |
| Malignant epithelial | C2 WNT ECs | ATG5 | 0.642 | 0.422 | 0.234 | 2.61e-23 | 8.81e-19 |
| Malignant epithelial | C2 WNT ECs | GORASP2 | 0.642 | 0.497 | 0.303 | 7.44e-22 | 2.51e-17 |
| Malignant epithelial | C2 WNT ECs | KLHL36 | 0.642 | 0.25 | 0.136 | 2.14e-14 | 7.21e-10 |
| Malignant epithelial | C2 WNT ECs | CHPF2 | 0.642 | 0.165 | 0.083 | 7.46e-12 | 2.51e-07 |
| Malignant epithelial | C2 WNT ECs | ARID5A | 0.642 | 0.2 | 0.12 | 4.22e-09 | 0.000142 |
| Malignant epithelial | C2 WNT ECs | BIN3 | 0.642 | 0.18 | 0.107 | 3.07e-08 | 0.00104 |
| Malignant epithelial | C2 WNT ECs | GADD45B | 0.641 | 0.862 | 0.735 | 5.95e-21 | 2.01e-16 |
| Malignant epithelial | C2 WNT ECs | PDXDC1 | 0.641 | 0.396 | 0.228 | 1.9e-19 | 6.4e-15 |
| Malignant epithelial | C2 WNT ECs | ICK | 0.641 | 0.12 | 0.065 | 1.07e-07 | 0.00362 |
| Malignant epithelial | C2 WNT ECs | PGRMC1 | 0.64 | 0.75 | 0.499 | 8.26e-37 | 2.78e-32 |

| compartment | subcluster | gene | avg_log2FC | pct.1 | pct.2 | p_val | p_val_adj |
|---|---|---|---|---|---|---|---|
| Malignant epithelial | C2 WNT ECs | UBE2K | 0.64 | 0.521 | 0.29 | 1.14e-28 | 3.84e-24 |
| Malignant epithelial | C2 WNT ECs | TRNT1 | 0.64 | 0.177 | 0.078 | 2.34e-16 | 7.89e-12 |
| Malignant epithelial | C2 WNT ECs | MCOLN1 | 0.64 | 0.147 | 0.076 | 3.7e-10 | 1.25e-05 |
| Malignant epithelial | C2 WNT ECs | GMDS | 0.639 | 0.505 | 0.287 | 1.03e-26 | 3.48e-22 |
| Malignant epithelial | C2 WNT ECs | EIF3L | 0.639 | 0.794 | 0.618 | 3.33e-26 | 1.12e-21 |
| Malignant epithelial | C2 WNT ECs | MLLT4 | 0.639 | 0.417 | 0.229 | 9.069999999999999e-23 | 3.06e-18 |
| Malignant epithelial | C2 WNT ECs | SND1-IT1 | 0.639 | 0.162 | 0.083 | 3.51e-11 | 1.18e-06 |
| Malignant epithelial | C2 WNT ECs | ATP11B | 0.639 | 0.199 | 0.116 | 1.89e-09 | 6.38e-05 |
| Malignant epithelial | C2 WNT ECs | KHDRBS1 | 0.638 | 0.748 | 0.508 | 2.69e-33 | 9.05e-29 |
| Malignant epithelial | C2 WNT ECs | VPS26A | 0.638 | 0.533 | 0.325 | 3.5e-23 | 1.18e-18 |
| Malignant epithelial | C2 WNT ECs | DYNLL2 | 0.638 | 0.476 | 0.299 | 3.63e-19 | 1.22e-14 |
| Malignant epithelial | C2 WNT ECs | DCTN1 | 0.638 | 0.281 | 0.153 | 8.88e-16 | 2.99e-11 |
| Malignant epithelial | C2 WNT ECs | TAB2 | 0.638 | 0.161 | 0.083 | 6.69e-11 | 2.25e-06 |
| Malignant epithelial | C2 WNT ECs | KBTBD3 | 0.637 | 0.175 | 0.09 | 1.6e-11 | 5.38e-07 |
| Malignant epithelial | C2 WNT ECs | TMEM14A | 0.636 | 0.64 | 0.414 | 4e-27 | 1.35e-22 |
| Malignant epithelial | C2 WNT ECs | CD2BP2 | 0.636 | 0.533 | 0.325 | 1.93e-23 | 6.49e-19 |
| Malignant epithelial | C2 WNT ECs | CEP104 | 0.636 | 0.124 | 0.067 | 7.27e-08 | 0.00245 |
| Malignant epithelial | C2 WNT ECs | SIKE1 | 0.635 | 0.241 | 0.125 | 1.69e-15 | 5.68e-11 |
| Malignant epithelial | C2 WNT ECs | KIAA1522 | 0.635 | 0.287 | 0.163 | 2.79e-14 | 9.41e-10 |
| Malignant epithelial | C2 WNT ECs | MPP6 | 0.635 | 0.181 | 0.092 | 3.24e-12 | 1.09e-07 |
| Malignant epithelial | C2 WNT ECs | CHD3 | 0.635 | 0.165 | 0.09 | 2.43e-09 | 8.19e-05 |
| Malignant epithelial | C2 WNT ECs | MTMR12 | 0.635 | 0.104 | 0.052 | 6.67e-08 | 0.00225 |
| Malignant epithelial | C2 WNT ECs | CCDC117 | 0.634 | 0.119 | 0.055 | 2.12e-10 | 7.14e-06 |
| Malignant epithelial | C2 WNT ECs | NUP98 | 0.633 | 0.192 | 0.113 | 4.72e-09 | 0.000159 |
| Malignant epithelial | C2 WNT ECs | CXADR | 0.632 | 0.545 | 0.319 | 9.370000000000001e-27 | 3.16e-22 |
| Malignant epithelial | C2 WNT ECs | SEC61A1 | 0.632 | 0.486 | 0.32 | 4.08e-17 | 1.37e-12 |
| Malignant epithelial | C2 WNT ECs | RREB1 | 0.632 | 0.235 | 0.132 | 2.51e-12 | 8.47e-08 |
| Malignant epithelial | C2 WNT ECs | ILK | 0.631 | 0.587 | 0.364 | 3.08e-26 | 1.04e-21 |
| Malignant epithelial | C2 WNT ECs | HEXB | 0.631 | 0.512 | 0.356 | 1.12e-15 | 3.78e-11 |
| Malignant epithelial | C2 WNT ECs | ME2 | 0.631 | 0.126 | 0.066 | 2.24e-08 | 0.000756 |
| Malignant epithelial | C2 WNT ECs | ESYT2 | 0.63 | 0.364 | 0.222 | 4.98e-15 | 1.68e-10 |
| Malignant epithelial | C2 WNT ECs | ACSS2 | 0.63 | 0.268 | 0.163 | 2.97e-11 | 1e-06 |
| Malignant epithelial | C2 WNT ECs | TMEM192 | 0.63 | 0.191 | 0.104 | 5.87e-11 | 1.98e-06 |
| Malignant epithelial | C2 WNT ECs | STX1A | 0.63 | 0.177 | 0.096 | 1.27e-10 | 4.27e-06 |
| Malignant epithelial | C2 WNT ECs | NAE1 | 0.629 | 0.457 | 0.268 | 5.46e-22 | 1.84e-17 |
| Malignant epithelial | C2 WNT ECs | CYB561A3 | 0.629 | 0.233 | 0.132 | 4.65e-12 | 1.57e-07 |
| Malignant epithelial | C2 WNT ECs | CYP27A1 | 0.629 | 0.13 | 0.069 | 1.32e-08 | 0.000445 |
| Malignant epithelial | C2 WNT ECs | MAF1 | 0.628 | 0.723 | 0.532 | 1.54e-23 | 5.18e-19 |
| Malignant epithelial | C2 WNT ECs | HSPD1 | 0.628 | 0.919 | 0.826 | 1.15e-22 | 3.89e-18 |
| Malignant epithelial | C2 WNT ECs | VSIG10 | 0.628 | 0.367 | 0.216 | 9.52e-17 | 3.21e-12 |
| Malignant epithelial | C2 WNT ECs | USP7 | 0.627 | 0.36 | 0.208 | 3.9e-17 | 1.32e-12 |
| Malignant epithelial | C2 WNT ECs | LTV1 | 0.627 | 0.245 | 0.126 | 9.3e-16 | 3.13e-11 |
| Malignant epithelial | C2 WNT ECs | GFOD2 | 0.627 | 0.147 | 0.082 | 2.18e-08 | 0.000736 |
| Malignant epithelial | C2 WNT ECs | PSMD13 | 0.626 | 0.608 | 0.4 | 3.7e-23 | 1.25e-18 |
| Malignant epithelial | C2 WNT ECs | KIAA1191 | 0.626 | 0.375 | 0.233 | 7.22e-15 | 2.43e-10 |
| Malignant epithelial | C2 WNT ECs | VEZF1 | 0.626 | 0.268 | 0.148 | 1.14e-14 | 3.85e-10 |
| Malignant epithelial | C2 WNT ECs | NT5DC3 | 0.626 | 0.114 | 0.061 | 2.49e-07 | 0.00837 |
| Malignant epithelial | C2 WNT ECs | RSPH3 | 0.625 | 0.166 | 0.091 | 1.37e-09 | 4.6e-05 |
| Malignant epithelial | C2 WNT ECs | DDR1 | 0.624 | 0.621 | 0.39 | 4.39e-27 | 1.48e-22 |
| Malignant epithelial | C2 WNT ECs | EPHB2 | 0.624 | 0.249 | 0.138 | 1.81e-13 | 6.09e-09 |
| Malignant epithelial | C2 WNT ECs | LINC00623 | 0.624 | 0.153 | 0.079 | 4.52e-10 | 1.52e-05 |
| Malignant epithelial | C2 WNT ECs | TMEM134 | 0.623 | 0.673 | 0.45 | 4.51e-27 | 1.52e-22 |
| Malignant epithelial | C2 WNT ECs | TSTA3 | 0.623 | 0.689 | 0.458 | 6.96e-26 | 2.35e-21 |
| Malignant epithelial | C2 WNT ECs | BECN1 | 0.623 | 0.349 | 0.201 | 8.32e-17 | 2.8e-12 |
| Malignant epithelial | C2 WNT ECs | PEF1 | 0.623 | 0.326 | 0.183 | 1.8e-16 | 6.08e-12 |
| Malignant epithelial | C2 WNT ECs | DIP2B | 0.623 | 0.204 | 0.1 | 4.28e-15 | 1.44e-10 |
| Malignant epithelial | C2 WNT ECs | VILL | 0.623 | 0.192 | 0.102 | 1.05e-11 | 3.52e-07 |
| Malignant epithelial | C2 WNT ECs | ITCH | 0.623 | 0.211 | 0.123 | 5.25e-10 | 1.77e-05 |
| Malignant epithelial | C2 WNT ECs | ZNF738 | 0.623 | 0.1 | 0.054 | 1.32e-06 | 0.0445 |
| Malignant epithelial | C2 WNT ECs | TMEM63A | 0.622 | 0.44 | 0.242 | 3.22e-24 | 1.08e-19 |
| Malignant epithelial | C2 WNT ECs | RHPN2 | 0.622 | 0.281 | 0.156 | 5.94e-15 | 2e-10 |
| Malignant epithelial | C2 WNT ECs | CXXC1 | 0.622 | 0.185 | 0.091 | 1.41e-13 | 4.75e-09 |
| Malignant epithelial | C2 WNT ECs | THOC5 | 0.622 | 0.158 | 0.081 | 9.55e-11 | 3.22e-06 |
| Malignant epithelial | C2 WNT ECs | ZNRD1 | 0.621 | 0.535 | 0.319 | 2.67e-24 | 9.01e-20 |

| compartment | subcluster | gene | avg_log2FC | pct.1 | pct.2 | p_val | p_val_adj |
|---|---|---|---|---|---|---|---|
| Malignant epithelial | C2 WNT ECs | RNF8 | 0.621 | 0.265 | 0.141 | 1.16e-15 | 3.92e-11 |
| Malignant epithelial | C2 WNT ECs | METTL21A | 0.62 | 0.207 | 0.115 | 7.64e-11 | 2.58e-06 |
| Malignant epithelial | C2 WNT ECs | LAPTM4B | 0.62 | 0.461 | 0.358 | 9.8e-10 | 3.3e-05 |
| Malignant epithelial | C2 WNT ECs | NADK | 0.619 | 0.283 | 0.154 | 1.09e-15 | 3.67e-11 |
| Malignant epithelial | C2 WNT ECs | CREG1 | 0.619 | 0.474 | 0.322 | 1.78e-14 | 6.01e-10 |
| Malignant epithelial | C2 WNT ECs | ANKRD17 | 0.619 | 0.218 | 0.113 | 2.41e-14 | 8.12e-10 |
| Malignant epithelial | C2 WNT ECs | AXIN1 | 0.619 | 0.199 | 0.104 | 1.79e-12 | 6.03e-08 |
| Malignant epithelial | C2 WNT ECs | DDIT3 | 0.619 | 0.617 | 0.478 | 2.98e-12 | 1e-07 |
| Malignant epithelial | C2 WNT ECs | RP11-156K23.3 | 0.619 | 0.124 | 0.065 | 1.44e-08 | 0.000486 |
| Malignant epithelial | C2 WNT ECs | TSEN34 | 0.618 | 0.631 | 0.406 | 5.570000000000001e-27 | 1.88e-22 |
| Malignant epithelial | C2 WNT ECs | PAWR | 0.618 | 0.612 | 0.398 | 5.27e-23 | 1.77e-18 |
| Malignant epithelial | C2 WNT ECs | RBM5 | 0.618 | 0.387 | 0.213 | 4.52e-21 | 1.52e-16 |
| Malignant epithelial | C2 WNT ECs | GON4L | 0.618 | 0.291 | 0.151 | 4.04e-18 | 1.36e-13 |
| Malignant epithelial | C2 WNT ECs | ZNF664 | 0.618 | 0.379 | 0.224 | 2.81e-17 | 9.45e-13 |
| Malignant epithelial | C2 WNT ECs | POLR2D | 0.618 | 0.226 | 0.127 | 5.36e-12 | 1.81e-07 |
| Malignant epithelial | C2 WNT ECs | ARIH1 | 0.617 | 0.373 | 0.207 | 2.26e-20 | 7.63e-16 |
| Malignant epithelial | C2 WNT ECs | PYCR1 | 0.617 | 0.442 | 0.283 | 6.79e-16 | 2.29e-11 |
| Malignant epithelial | C2 WNT ECs | TBCC | 0.617 | 0.359 | 0.238 | 1.29e-11 | 4.35e-07 |
| Malignant epithelial | C2 WNT ECs | ARL5B | 0.617 | 0.379 | 0.251 | 3.28e-11 | 1.1e-06 |
| Malignant epithelial | C2 WNT ECs | HSPG2 | 0.617 | 0.237 | 0.137 | 3.54e-11 | 1.19e-06 |
| Malignant epithelial | C2 WNT ECs | EMC10 | 0.616 | 0.706 | 0.483 | 9.570000000000001e-27 | 3.23e-22 |
| Malignant epithelial | C2 WNT ECs | CYP2S1 | 0.616 | 0.311 | 0.157 | 2.07e-20 | 6.97e-16 |
| Malignant epithelial | C2 WNT ECs | CTA-292E10.6 | 0.616 | 0.135 | 0.065 | 5.4e-11 | 1.82e-06 |
| Malignant epithelial | C2 WNT ECs | AP3D1 | 0.615 | 0.478 | 0.307 | 7.1e-18 | 2.39e-13 |
| Malignant epithelial | C2 WNT ECs | RNF103 | 0.615 | 0.302 | 0.165 | 3.55e-16 | 1.2e-11 |
| Malignant epithelial | C2 WNT ECs | SLC12A7 | 0.615 | 0.257 | 0.137 | 1.26e-15 | 4.24e-11 |
| Malignant epithelial | C2 WNT ECs | C11orf80 | 0.615 | 0.241 | 0.134 | 1.45e-12 | 4.9e-08 |
| Malignant epithelial | C2 WNT ECs | MYO1B | 0.615 | 0.188 | 0.097 | 2.59e-12 | 8.74e-08 |
| Malignant epithelial | C2 WNT ECs | DDI2 | 0.615 | 0.158 | 0.084 | 7.34e-10 | 2.47e-05 |
| Malignant epithelial | C2 WNT ECs | CREB3L4 | 0.615 | 0.149 | 0.088 | 3.59e-07 | 0.0121 |
| Malignant epithelial | C2 WNT ECs | PDIA4 | 0.614 | 0.723 | 0.546 | 3.46e-21 | 1.17e-16 |
| Malignant epithelial | C2 WNT ECs | OAF | 0.614 | 0.276 | 0.143 | 1.72e-17 | 5.8e-13 |
| Malignant epithelial | C2 WNT ECs | EFCAB14 | 0.614 | 0.26 | 0.133 | 3.79e-17 | 1.28e-12 |
| Malignant epithelial | C2 WNT ECs | RBPJ | 0.614 | 0.365 | 0.215 | 7.99e-17 | 2.69e-12 |
| Malignant epithelial | C2 WNT ECs | EXT2 | 0.614 | 0.161 | 0.097 | 3.83e-07 | 0.0129 |
| Malignant epithelial | C2 WNT ECs | STARD10 | 0.613 | 0.788 | 0.5 | 2.63e-44 | 8.86e-40 |
| Malignant epithelial | C2 WNT ECs | MED28 | 0.613 | 0.506 | 0.284 | 3.49e-26 | 1.18e-21 |
| Malignant epithelial | C2 WNT ECs | CAPZA1 | 0.613 | 0.602 | 0.388 | 1.37e-23 | 4.61e-19 |
| Malignant epithelial | C2 WNT ECs | PPM1B | 0.613 | 0.304 | 0.179 | 1.79e-13 | 6.05e-09 |
| Malignant epithelial | C2 WNT ECs | POGLUT1 | 0.613 | 0.111 | 0.056 | 3.87e-08 | 0.0013 |
| Malignant epithelial | C2 WNT ECs | PLXNA1 | 0.613 | 0.134 | 0.078 | 6.94e-07 | 0.0234 |
| Malignant epithelial | C2 WNT ECs | TFG | 0.612 | 0.66 | 0.435 | 1.35e-26 | 4.53e-22 |
| Malignant epithelial | C2 WNT ECs | TAF3 | 0.612 | 0.206 | 0.118 | 3.13e-10 | 1.06e-05 |
| Malignant epithelial | C2 WNT ECs | MYL6B | 0.611 | 0.556 | 0.364 | 1.21e-21 | 4.07e-17 |
| Malignant epithelial | C2 WNT ECs | MBTPS1 | 0.611 | 0.299 | 0.158 | 3.73e-18 | 1.26e-13 |
| Malignant epithelial | C2 WNT ECs | SLC17A5 | 0.611 | 0.143 | 0.081 | 1.36e-07 | 0.00458 |
| Malignant epithelial | C2 WNT ECs | DHRS3 | 0.61 | 0.413 | 0.233 | 1.53e-20 | 5.15e-16 |
| Malignant epithelial | C2 WNT ECs | THRAP3 | 0.61 | 0.456 | 0.298 | 5.38e-16 | 1.81e-11 |
| Malignant epithelial | C2 WNT ECs | PPFIBP1 | 0.61 | 0.244 | 0.151 | 1.98e-09 | 6.68e-05 |
| Malignant epithelial | C2 WNT ECs | SFT2D1 | 0.609 | 0.779 | 0.531 | 1.46e-34 | 4.92e-30 |
| Malignant epithelial | C2 WNT ECs | SLC38A2 | 0.609 | 0.742 | 0.523 | 3.689999999999999e-24 | 1.24e-19 |
| Malignant epithelial | C2 WNT ECs | CHDH | 0.609 | 0.26 | 0.144 | 1.07e-13 | 3.6e-09 |
| Malignant epithelial | C2 WNT ECs | ARHGAP27 | 0.609 | 0.417 | 0.267 | 3.38e-13 | 1.14e-08 |
| Malignant epithelial | C2 WNT ECs | MPG | 0.608 | 0.739 | 0.536 | 9.64e-28 | 3.25e-23 |
| Malignant epithelial | C2 WNT ECs | ZNF131 | 0.608 | 0.401 | 0.235 | 4.14e-19 | 1.39e-14 |
| Malignant epithelial | C2 WNT ECs | GPAT4 | 0.608 | 0.313 | 0.167 | 9.26e-19 | 3.12e-14 |
| Malignant epithelial | C2 WNT ECs | POMGNT1 | 0.608 | 0.176 | 0.084 | 9.09e-14 | 3.06e-09 |
| Malignant epithelial | C2 WNT ECs | DHX36 | 0.607 | 0.54 | 0.337 | 4.76e-23 | 1.6e-18 |
| Malignant epithelial | C2 WNT ECs | FKBP14 | 0.607 | 0.107 | 0.057 | 5.8e-07 | 0.0195 |
| Malignant epithelial | C2 WNT ECs | MED10 | 0.606 | 0.747 | 0.504 | 2.04e-32 | 6.89e-28 |
| Malignant epithelial | C2 WNT ECs | ARPP19 | 0.606 | 0.591 | 0.396 | 1.02e-19 | 3.42e-15 |
| Malignant epithelial | C2 WNT ECs | RAPGEFL1 | 0.604 | 0.203 | 0.105 | 7.17e-13 | 2.42e-08 |
| Malignant epithelial | C2 WNT ECs | DDX19A | 0.604 | 0.165 | 0.092 | 3.31e-09 | 0.000112 |
| Malignant epithelial | C2 WNT ECs | TSPAN1 | 0.604 | 0.616 | 0.53 | 3.46e-08 | 0.00117 |

| compartment | subcluster | gene | avg_log2FC | pct.1 | pct.2 | p_val | p_val_adj |
|---|---|---|---|---|---|---|---|
| Malignant epithelial | C2 WNT ECs | CLK1 | 0.603 | 0.723 | 0.496 | 1.91e-27 | 6.44e-23 |
| Malignant epithelial | C2 WNT ECs | CUX1 | 0.603 | 0.487 | 0.287 | 8.18e-22 | 2.75e-17 |
| Malignant epithelial | C2 WNT ECs | CCNC | 0.603 | 0.499 | 0.31 | 5.01e-20 | 1.69e-15 |
| Malignant epithelial | C2 WNT ECs | SMARCC2 | 0.603 | 0.311 | 0.176 | 5.29e-16 | 1.78e-11 |
| Malignant epithelial | C2 WNT ECs | TMEM170A | 0.603 | 0.405 | 0.255 | 8.77e-15 | 2.96e-10 |
| Malignant epithelial | C2 WNT ECs | ANKZF1 | 0.603 | 0.24 | 0.134 | 9.78e-13 | 3.3e-08 |
| Malignant epithelial | C2 WNT ECs | CASP6 | 0.603 | 0.319 | 0.199 | 1.88e-12 | 6.35e-08 |
| Malignant epithelial | C2 WNT ECs | PPP2R1A | 0.602 | 0.775 | 0.547 | 2.52e-27 | 8.489999999999999e-23 |
| Malignant epithelial | C2 WNT ECs | APEH | 0.602 | 0.369 | 0.209 | 1.56e-18 | 5.25e-14 |
| Malignant epithelial | C2 WNT ECs | EEPD1 | 0.602 | 0.245 | 0.149 | 2.5e-10 | 8.43e-06 |
| Malignant epithelial | C2 WNT ECs | HNRNPM | 0.601 | 0.754 | 0.526 | 5.51e-27 | 1.86e-22 |
| Malignant epithelial | C2 WNT ECs | BRMS1 | 0.601 | 0.521 | 0.318 | 3.26e-22 | 1.1e-17 |
| Malignant epithelial | C2 WNT ECs | DHX15 | 0.601 | 0.39 | 0.216 | 7.25e-21 | 2.44e-16 |
| Malignant epithelial | C2 WNT ECs | TOP1MT | 0.601 | 0.346 | 0.219 | 1.94e-12 | 6.52e-08 |
| Malignant epithelial | C2 WNT ECs | NDUFAF7 | 0.601 | 0.114 | 0.062 | 9.99e-07 | 0.0337 |
| Malignant epithelial | C2 WNT ECs | CAPZB | 0.6 | 0.8 | 0.566 | 1.15e-33 | 3.86e-29 |
| Malignant epithelial | C2 WNT ECs | AP2M1 | 0.6 | 0.742 | 0.531 | 2.29e-28 | 7.729999999999999e-24 |
| Malignant epithelial | C2 WNT ECs | PRRG2 | 0.6 | 0.242 | 0.129 | 5.56e-14 | 1.87e-09 |
| Malignant epithelial | C2 WNT ECs | DCP1A | 0.6 | 0.28 | 0.16 | 1.56e-13 | 5.25e-09 |
| Malignant epithelial | C2 WNT ECs | CCDC66 | 0.6 | 0.189 | 0.094 | 1.8e-13 | 6.06e-09 |
| Malignant epithelial | C2 WNT ECs | COMMD8 | 0.6 | 0.223 | 0.125 | 3.65e-11 | 1.23e-06 |
| Malignant epithelial | C2 WNT ECs | ZBED6 | 0.6 | 0.195 | 0.11 | 4.17e-10 | 1.4e-05 |
| Malignant epithelial | C2 WNT ECs | TVP23B | 0.6 | 0.126 | 0.071 | 1.12e-06 | 0.0377 |
| Malignant epithelial | C2 WNT ECs | ARL4A | 0.599 | 0.913 | 0.758 | 5.95e-33 | 2e-28 |
| Malignant epithelial | C2 WNT ECs | ETFB | 0.599 | 0.827 | 0.683 | 2.02e-23 | 6.8e-19 |
| Malignant epithelial | C2 WNT ECs | PCAT6 | 0.599 | 0.141 | 0.056 | 2.63e-16 | 8.85e-12 |
| Malignant epithelial | C2 WNT ECs | WDR70 | 0.599 | 0.25 | 0.134 | 1.08e-14 | 3.63e-10 |
| Malignant epithelial | C2 WNT ECs | CD58 | 0.599 | 0.317 | 0.189 | 9.09e-14 | 3.06e-09 |
| Malignant epithelial | C2 WNT ECs | CAPN15 | 0.599 | 0.219 | 0.129 | 7.89e-10 | 2.66e-05 |
| Malignant epithelial | C2 WNT ECs | SEPT2 | 0.598 | 0.689 | 0.46 | 3.16e-27 | 1.07e-22 |
| Malignant epithelial | C2 WNT ECs | ARID4B | 0.598 | 0.555 | 0.355 | 9.62e-22 | 3.24e-17 |
| Malignant epithelial | C2 WNT ECs | GTPBP6 | 0.598 | 0.355 | 0.191 | 1.28e-19 | 4.3e-15 |
| Malignant epithelial | C2 WNT ECs | SUN2 | 0.598 | 0.337 | 0.193 | 1.26e-16 | 4.23e-12 |
| Malignant epithelial | C2 WNT ECs | CUTC | 0.598 | 0.176 | 0.1 | 7.38e-09 | 0.000249 |
| Malignant epithelial | C2 WNT ECs | RHOA | 0.597 | 0.913 | 0.766 | 5.270000000000001e-32 | 1.78e-27 |
| Malignant epithelial | C2 WNT ECs | G3BP2 | 0.597 | 0.391 | 0.218 | 1.94e-20 | 6.54e-16 |
| Malignant epithelial | C2 WNT ECs | LRP11 | 0.597 | 0.332 | 0.193 | 1.27e-15 | 4.26e-11 |
| Malignant epithelial | C2 WNT ECs | TMUB2 | 0.597 | 0.329 | 0.191 | 7.29e-15 | 2.45e-10 |
| Malignant epithelial | C2 WNT ECs | PRKAR2A | 0.597 | 0.323 | 0.198 | 3.14e-13 | 1.06e-08 |
| Malignant epithelial | C2 WNT ECs | SLC39A6 | 0.597 | 0.181 | 0.096 | 2.74e-11 | 9.23e-07 |
| Malignant epithelial | C2 WNT ECs | C1orf50 | 0.597 | 0.179 | 0.094 | 5.7e-11 | 1.92e-06 |
| Malignant epithelial | C2 WNT ECs | NDRG2 | 0.597 | 0.188 | 0.103 | 1.44e-10 | 4.85e-06 |
| Malignant epithelial | C2 WNT ECs | UBIAD1 | 0.597 | 0.147 | 0.079 | 4.06e-09 | 0.000137 |
| Malignant epithelial | C2 WNT ECs | NELFCD | 0.596 | 0.614 | 0.404 | 7.29e-25 | 2.46e-20 |
| Malignant epithelial | C2 WNT ECs | AKAP1 | 0.596 | 0.509 | 0.329 | 1.76e-18 | 5.93e-14 |
| Malignant epithelial | C2 WNT ECs | PILRB | 0.596 | 0.263 | 0.143 | 6.49e-15 | 2.19e-10 |
| Malignant epithelial | C2 WNT ECs | PLRG1 | 0.596 | 0.248 | 0.134 | 6.8e-14 | 2.29e-09 |
| Malignant epithelial | C2 WNT ECs | RNF139 | 0.596 | 0.41 | 0.277 | 2.52e-12 | 8.5e-08 |
| Malignant epithelial | C2 WNT ECs | CIB2 | 0.596 | 0.164 | 0.102 | 1.34e-06 | 0.0452 |
| Malignant epithelial | C2 WNT ECs | CAPN7 | 0.596 | 0.124 | 0.071 | 1.46e-06 | 0.0492 |
| Malignant epithelial | C2 WNT ECs | C4orf32 | 0.595 | 0.254 | 0.138 | 4.03e-14 | 1.36e-09 |
| Malignant epithelial | C2 WNT ECs | UNC45A | 0.595 | 0.272 | 0.153 | 1.8e-13 | 6.06e-09 |
| Malignant epithelial | C2 WNT ECs | DLGAP1-AS1 | 0.595 | 0.171 | 0.09 | 5.09e-11 | 1.72e-06 |
| Malignant epithelial | C2 WNT ECs | TMEM107 | 0.595 | 0.191 | 0.106 | 1.28e-10 | 4.3e-06 |
| Malignant epithelial | C2 WNT ECs | RP11-386I14.4 | 0.595 | 0.147 | 0.084 | 9.71e-08 | 0.00327 |
| Malignant epithelial | C2 WNT ECs | TRIP4 | 0.595 | 0.152 | 0.088 | 2.44e-07 | 0.00823 |
| Malignant epithelial | C2 WNT ECs | PABPC1 | 0.594 | 0.988 | 0.963 | 3.06e-50 | 1.03e-45 |
| Malignant epithelial | C2 WNT ECs | DNMBP | 0.594 | 0.165 | 0.096 | 4.33e-08 | 0.00146 |
| Malignant epithelial | C2 WNT ECs | USP32 | 0.594 | 0.112 | 0.059 | 1.44e-07 | 0.00486 |
| Malignant epithelial | C2 WNT ECs | ESF1 | 0.593 | 0.441 | 0.333 | 2.79e-09 | 9.41e-05 |
| Malignant epithelial | C2 WNT ECs | DCK | 0.593 | 0.149 | 0.083 | 4.4e-08 | 0.00148 |
| Malignant epithelial | C2 WNT ECs | UBE2D1 | 0.592 | 0.51 | 0.325 | 2.31e-18 | 7.78e-14 |
| Malignant epithelial | C2 WNT ECs | FBXO18 | 0.592 | 0.231 | 0.123 | 1.15e-13 | 3.87e-09 |
| Malignant epithelial | C2 WNT ECs | PKD1 | 0.592 | 0.15 | 0.071 | 2.62e-12 | 8.82e-08 |

| compartment | subcluster | gene | avg_log2FC | pct.1 | pct.2 | p_val | p_val_adj |
|---|---|---|---|---|---|---|---|
| Malignant epithelial | C2 WNT ECs | RP11-352M15.2 | 0.592 | 0.192 | 0.119 | 1.07e-07 | 0.00362 |
| Malignant epithelial | C2 WNT ECs | UBE2I | 0.591 | 0.889 | 0.701 | 3.11e-41 | 1.05e-36 |
| Malignant epithelial | C2 WNT ECs | HSD17B12 | 0.591 | 0.598 | 0.396 | 5.57e-22 | 1.88e-17 |
| Malignant epithelial | C2 WNT ECs | ZPR1 | 0.591 | 0.43 | 0.251 | 1.14e-19 | 3.84e-15 |
| Malignant epithelial | C2 WNT ECs | CD68 | 0.591 | 0.392 | 0.244 | 8.01e-16 | 2.7e-11 |
| Malignant epithelial | C2 WNT ECs | LTBP3 | 0.591 | 0.421 | 0.271 | 8.92e-15 | 3.01e-10 |
| Malignant epithelial | C2 WNT ECs | ARHGAP17 | 0.591 | 0.191 | 0.116 | 7.19e-08 | 0.00242 |
| Malignant epithelial | C2 WNT ECs | LTN1 | 0.591 | 0.141 | 0.083 | 7.08e-07 | 0.0238 |
| Malignant epithelial | C2 WNT ECs | MIEN1 | 0.59 | 0.802 | 0.579 | 5.18e-32 | 1.75e-27 |
| Malignant epithelial | C2 WNT ECs | GOLPH3 | 0.59 | 0.405 | 0.243 | 1.9e-17 | 6.4e-13 |
| Malignant epithelial | C2 WNT ECs | PTPN23 | 0.59 | 0.187 | 0.1 | 5.13e-11 | 1.73e-06 |
| Malignant epithelial | C2 WNT ECs | MADD | 0.59 | 0.137 | 0.071 | 3.3e-09 | 0.000111 |
| Malignant epithelial | C2 WNT ECs | YIPF2 | 0.589 | 0.483 | 0.283 | 9.47e-24 | 3.19e-19 |
| Malignant epithelial | C2 WNT ECs | NKTR | 0.589 | 0.606 | 0.397 | 1.97e-23 | 6.65e-19 |
| Malignant epithelial | C2 WNT ECs | TRAPPC3 | 0.589 | 0.52 | 0.312 | 2.18e-23 | 7.35e-19 |
| Malignant epithelial | C2 WNT ECs | TGS1 | 0.589 | 0.368 | 0.214 | 3.63e-17 | 1.22e-12 |
| Malignant epithelial | C2 WNT ECs | ADIPOR2 | 0.589 | 0.304 | 0.21 | 2.54e-08 | 0.000857 |
| Malignant epithelial | C2 WNT ECs | GGA1 | 0.587 | 0.44 | 0.237 | 3.43e-25 | 1.15e-20 |
| Malignant epithelial | C2 WNT ECs | DNAJC7 | 0.587 | 0.724 | 0.512 | 1.75e-24 | 5.89e-20 |
| Malignant epithelial | C2 WNT ECs | GARS | 0.587 | 0.493 | 0.312 | 6.8e-19 | 2.29e-14 |
| Malignant epithelial | C2 WNT ECs | P4HA2 | 0.587 | 0.283 | 0.153 | 1.01e-15 | 3.42e-11 |
| Malignant epithelial | C2 WNT ECs | AGPS | 0.587 | 0.196 | 0.094 | 3.31e-15 | 1.12e-10 |
| Malignant epithelial | C2 WNT ECs | QRSL1 | 0.587 | 0.207 | 0.113 | 2.9e-11 | 9.76e-07 |
| Malignant epithelial | C2 WNT ECs | TJAP1 | 0.587 | 0.166 | 0.088 | 1.9e-10 | 6.39e-06 |
| Malignant epithelial | C2 WNT ECs | CWC22 | 0.587 | 0.157 | 0.088 | 2.24e-08 | 0.000755 |
| Malignant epithelial | C2 WNT ECs | WHSC1 | 0.587 | 0.145 | 0.081 | 5.07e-08 | 0.00171 |
| Malignant epithelial | C2 WNT ECs | SPP1 | 0.586 | 0.33 | 0.19 | 1.66e-16 | 5.6e-12 |
| Malignant epithelial | C2 WNT ECs | TSC22D2 | 0.586 | 0.403 | 0.251 | 3.34e-15 | 1.13e-10 |
| Malignant epithelial | C2 WNT ECs | CIAPIN1 | 0.586 | 0.332 | 0.195 | 7.89e-15 | 2.66e-10 |
| Malignant epithelial | C2 WNT ECs | MTG2 | 0.586 | 0.321 | 0.19 | 1.03e-13 | 3.48e-09 |
| Malignant epithelial | C2 WNT ECs | THNSL2 | 0.586 | 0.141 | 0.069 | 1.6e-10 | 5.38e-06 |
| Malignant epithelial | C2 WNT ECs | MVK | 0.586 | 0.138 | 0.077 | 5.33e-08 | 0.00179 |
| Malignant epithelial | C2 WNT ECs | TMEM65 | 0.586 | 0.129 | 0.073 | 4.87e-07 | 0.0164 |
| Malignant epithelial | C2 WNT ECs | TP53I13 | 0.585 | 0.359 | 0.22 | 2.67e-14 | 8.98e-10 |
| Malignant epithelial | C2 WNT ECs | PNP | 0.585 | 0.337 | 0.209 | 5.76e-13 | 1.94e-08 |
| Malignant epithelial | C2 WNT ECs | GLA | 0.585 | 0.249 | 0.145 | 4.06e-11 | 1.37e-06 |
| Malignant epithelial | C2 WNT ECs | AP1S2 | 0.585 | 0.199 | 0.126 | 4.89e-07 | 0.0165 |
| Malignant epithelial | C2 WNT ECs | TBRG4 | 0.584 | 0.434 | 0.244 | 6.98e-22 | 2.35e-17 |
| Malignant epithelial | C2 WNT ECs | CCNG1 | 0.584 | 0.48 | 0.301 | 1.92e-19 | 6.48e-15 |
| Malignant epithelial | C2 WNT ECs | NSUN2 | 0.584 | 0.248 | 0.139 | 4.76e-13 | 1.61e-08 |
| Malignant epithelial | C2 WNT ECs | RAB6A | 0.583 | 0.622 | 0.386 | 1.6e-29 | 5.41e-25 |
| Malignant epithelial | C2 WNT ECs | GNAS | 0.582 | 0.932 | 0.797 | 2.08e-31 | 7.030000000000001e-27 |
| Malignant epithelial | C2 WNT ECs | ERGIC3 | 0.582 | 0.892 | 0.755 | 7.310000000000001e-27 | 2.46e-22 |
| Malignant epithelial | C2 WNT ECs | DGAT1 | 0.582 | 0.399 | 0.237 | 4.8e-18 | 1.62e-13 |
| Malignant epithelial | C2 WNT ECs | UBE2G1 | 0.582 | 0.3 | 0.173 | 3.69e-14 | 1.24e-09 |
| Malignant epithelial | C2 WNT ECs | ANKRD10 | 0.581 | 0.708 | 0.486 | 1.34e-24 | 4.52e-20 |
| Malignant epithelial | C2 WNT ECs | DNAJC8 | 0.581 | 0.536 | 0.333 | 2.63e-21 | 8.85e-17 |
| Malignant epithelial | C2 WNT ECs | MYH9 | 0.581 | 0.667 | 0.476 | 3.78e-20 | 1.27e-15 |
| Malignant epithelial | C2 WNT ECs | KIF21A | 0.581 | 0.253 | 0.163 | 1.55e-08 | 0.000522 |
| Malignant epithelial | C2 WNT ECs | ATG2A | 0.58 | 0.296 | 0.174 | 3.28e-12 | 1.1e-07 |
| Malignant epithelial | C2 WNT ECs | PIK3C2A | 0.58 | 0.214 | 0.119 | 3e-11 | 1.01e-06 |
| Malignant epithelial | C2 WNT ECs | ULK3 | 0.58 | 0.181 | 0.102 | 1.72e-09 | 5.78e-05 |
| Malignant epithelial | C2 WNT ECs | NDE1 | 0.58 | 0.204 | 0.12 | 3.79e-09 | 0.000128 |
| Malignant epithelial | C2 WNT ECs | UBE2B | 0.579 | 0.861 | 0.643 | 4.63e-36 | 1.56e-31 |
| Malignant epithelial | C2 WNT ECs | WDR46 | 0.579 | 0.336 | 0.201 | 1.16e-14 | 3.92e-10 |
| Malignant epithelial | C2 WNT ECs | LUC7L3 | 0.578 | 0.651 | 0.441 | 8.63e-24 | 2.91e-19 |
| Malignant epithelial | C2 WNT ECs | CEBPG | 0.578 | 0.535 | 0.331 | 7.27e-22 | 2.45e-17 |
| Malignant epithelial | C2 WNT ECs | BRIX1 | 0.578 | 0.512 | 0.33 | 2.48e-16 | 8.37e-12 |
| Malignant epithelial | C2 WNT ECs | ORMDL3 | 0.578 | 0.294 | 0.183 | 3.69e-11 | 1.24e-06 |
| Malignant epithelial | C2 WNT ECs | CARHSP1 | 0.577 | 0.862 | 0.651 | 8.800000000000001e-30 | 2.97e-25 |
| Malignant epithelial | C2 WNT ECs | PABPC4 | 0.577 | 0.502 | 0.307 | 1.65e-21 | 5.57e-17 |
| Malignant epithelial | C2 WNT ECs | DCTN5 | 0.577 | 0.333 | 0.188 | 2.62e-16 | 8.81e-12 |
| Malignant epithelial | C2 WNT ECs | PTDSS2 | 0.577 | 0.187 | 0.099 | 2.42e-11 | 8.15e-07 |
| Malignant epithelial | C2 WNT ECs | ZMAT5 | 0.577 | 0.292 | 0.18 | 6.7e-11 | 2.26e-06 |

| compartment | subcluster | gene | avg_log2FC | pct.1 | pct.2 | p_val | p_val_adj |
|---|---|---|---|---|---|---|---|
| Malignant epithelial | C2 WNT ECs | PSMB1 | 0.576 | 0.907 | 0.738 | 1.64e-41 | 5.540000000000001e-37 |
| Malignant epithelial | C2 WNT ECs | SUMF2 | 0.576 | 0.628 | 0.435 | 8.04e-20 | 2.71e-15 |
| Malignant epithelial | C2 WNT ECs | RFK | 0.576 | 0.356 | 0.201 | 6.54e-18 | 2.2e-13 |
| Malignant epithelial | C2 WNT ECs | CCNL2 | 0.576 | 0.384 | 0.229 | 1.25e-16 | 4.21e-12 |
| Malignant epithelial | C2 WNT ECs | MARCH7 | 0.576 | 0.28 | 0.174 | 1.18e-10 | 3.97e-06 |
| Malignant epithelial | C2 WNT ECs | TWF1 | 0.575 | 0.499 | 0.319 | 1.39e-18 | 4.67e-14 |
| Malignant epithelial | C2 WNT ECs | LRWD1 | 0.575 | 0.181 | 0.094 | 1.26e-11 | 4.23e-07 |
| Malignant epithelial | C2 WNT ECs | SOCS1 | 0.575 | 0.292 | 0.18 | 2.89e-11 | 9.74e-07 |
| Malignant epithelial | C2 WNT ECs | SKP1 | 0.574 | 0.98 | 0.884 | 5.13e-60 | 1.73e-55 |
| Malignant epithelial | C2 WNT ECs | ZNF800 | 0.574 | 0.276 | 0.156 | 1.14e-13 | 3.85e-09 |
| Malignant epithelial | C2 WNT ECs | HOXA11 | 0.574 | 0.138 | 0.076 | 5.68e-08 | 0.00191 |
| Malignant epithelial | C2 WNT ECs | GTF2I | 0.573 | 0.732 | 0.494 | 4.33e-27 | 1.46e-22 |
| Malignant epithelial | C2 WNT ECs | CPNE1 | 0.573 | 0.713 | 0.505 | 3.12e-24 | 1.05e-19 |
| Malignant epithelial | C2 WNT ECs | HAGH | 0.573 | 0.461 | 0.275 | 2.54e-20 | 8.55e-16 |
| Malignant epithelial | C2 WNT ECs | DDX1 | 0.573 | 0.528 | 0.337 | 4.17e-20 | 1.4e-15 |
| Malignant epithelial | C2 WNT ECs | TMEM160 | 0.573 | 0.723 | 0.601 | 9.47e-14 | 3.19e-09 |
| Malignant epithelial | C2 WNT ECs | ZNF160 | 0.573 | 0.154 | 0.084 | 5.38e-09 | 0.000181 |
| Malignant epithelial | C2 WNT ECs | CTSZ | 0.572 | 0.731 | 0.519 | 1.62e-23 | 5.46e-19 |
| Malignant epithelial | C2 WNT ECs | GLYR1 | 0.572 | 0.326 | 0.178 | 8.97e-18 | 3.02e-13 |
| Malignant epithelial | C2 WNT ECs | IMPDH1 | 0.572 | 0.361 | 0.228 | 4.62e-13 | 1.56e-08 |
| Malignant epithelial | C2 WNT ECs | KMT5C | 0.572 | 0.244 | 0.135 | 6.89e-13 | 2.32e-08 |
| Malignant epithelial | C2 WNT ECs | TMEM68 | 0.571 | 0.35 | 0.205 | 6.25e-16 | 2.11e-11 |
| Malignant epithelial | C2 WNT ECs | NGEF | 0.571 | 0.263 | 0.14 | 8.36e-16 | 2.82e-11 |
| Malignant epithelial | C2 WNT ECs | TIMM10B | 0.571 | 0.352 | 0.207 | 1.64e-15 | 5.53e-11 |
| Malignant epithelial | C2 WNT ECs | SPOPL | 0.571 | 0.152 | 0.078 | 3.78e-10 | 1.27e-05 |
| Malignant epithelial | C2 WNT ECs | ENDOD1 | 0.571 | 0.184 | 0.103 | 1.03e-09 | 3.47e-05 |
| Malignant epithelial | C2 WNT ECs | MANF | 0.57 | 0.748 | 0.529 | 4.21e-28 | 1.42e-23 |
| Malignant epithelial | C2 WNT ECs | PSMB2 | 0.57 | 0.744 | 0.518 | 5.21e-26 | 1.76e-21 |
| Malignant epithelial | C2 WNT ECs | CHMP2B | 0.57 | 0.712 | 0.495 | 6.92e-25 | 2.33e-20 |
| Malignant epithelial | C2 WNT ECs | EBNA1BP2 | 0.57 | 0.479 | 0.298 | 8.83e-18 | 2.97e-13 |
| Malignant epithelial | C2 WNT ECs | FAT1 | 0.57 | 0.3 | 0.178 | 3.66e-13 | 1.23e-08 |
| Malignant epithelial | C2 WNT ECs | PTDSS1 | 0.57 | 0.309 | 0.197 | 5.93e-11 | 2e-06 |
| Malignant epithelial | C2 WNT ECs | ATP13A3 | 0.57 | 0.203 | 0.113 | 1.08e-10 | 3.63e-06 |
| Malignant epithelial | C2 WNT ECs | RAB11B | 0.569 | 0.674 | 0.459 | 7.509999999999999e-24 | 2.53e-19 |
| Malignant epithelial | C2 WNT ECs | SERINC5 | 0.569 | 0.585 | 0.378 | 2.09e-22 | 7.04e-18 |
| Malignant epithelial | C2 WNT ECs | USP8 | 0.569 | 0.428 | 0.24 | 7.58e-22 | 2.55e-17 |
| Malignant epithelial | C2 WNT ECs | CNEP1R1 | 0.569 | 0.101 | 0.042 | 4.9e-11 | 1.65e-06 |
| Malignant epithelial | C2 WNT ECs | TBPL1 | 0.568 | 0.318 | 0.195 | 2.92e-12 | 9.84e-08 |
| Malignant epithelial | C2 WNT ECs | NISCH | 0.568 | 0.204 | 0.113 | 2.7e-11 | 9.1e-07 |
| Malignant epithelial | C2 WNT ECs | ATP2B4 | 0.568 | 0.126 | 0.067 | 4.73e-08 | 0.00159 |
| Malignant epithelial | C2 WNT ECs | FUT4 | 0.568 | 0.16 | 0.092 | 5.68e-08 | 0.00191 |
| Malignant epithelial | C2 WNT ECs | NSMCE1 | 0.567 | 0.641 | 0.389 | 1.26e-30 | 4.26e-26 |
| Malignant epithelial | C2 WNT ECs | HAT1 | 0.567 | 0.321 | 0.195 | 7.03e-13 | 2.37e-08 |
| Malignant epithelial | C2 WNT ECs | TPRG1L | 0.567 | 0.198 | 0.114 | 2.65e-09 | 8.94e-05 |
| Malignant epithelial | C2 WNT ECs | ARMCX5 | 0.567 | 0.108 | 0.056 | 1.9e-07 | 0.0064 |
| Malignant epithelial | C2 WNT ECs | ZNF44 | 0.567 | 0.13 | 0.074 | 4.67e-07 | 0.0157 |
| Malignant epithelial | C2 WNT ECs | PHF20L1 | 0.566 | 0.46 | 0.257 | 7.78e-24 | 2.62e-19 |
| Malignant epithelial | C2 WNT ECs | ZBTB1 | 0.566 | 0.187 | 0.089 | 2.19e-14 | 7.39e-10 |
| Malignant epithelial | C2 WNT ECs | PPME1 | 0.566 | 0.229 | 0.12 | 4.32e-14 | 1.46e-09 |
| Malignant epithelial | C2 WNT ECs | HS1BP3 | 0.566 | 0.183 | 0.093 | 1.58e-12 | 5.31e-08 |
| Malignant epithelial | C2 WNT ECs | GPR180 | 0.566 | 0.135 | 0.072 | 1.73e-08 | 0.000582 |
| Malignant epithelial | C2 WNT ECs | CDK11B | 0.565 | 0.348 | 0.181 | 7.52e-21 | 2.53e-16 |
| Malignant epithelial | C2 WNT ECs | KBTBD2 | 0.565 | 0.229 | 0.138 | 1.29e-09 | 4.36e-05 |
| Malignant epithelial | C2 WNT ECs | ITGB4 | 0.565 | 0.563 | 0.444 | 8.55e-09 | 0.000288 |
| Malignant epithelial | C2 WNT ECs | KHK | 0.565 | 0.16 | 0.088 | 9.34e-09 | 0.000315 |
| Malignant epithelial | C2 WNT ECs | ANXA3 | 0.564 | 0.647 | 0.452 | 4.47e-19 | 1.51e-14 |
| Malignant epithelial | C2 WNT ECs | ATP6AP1 | 0.563 | 0.563 | 0.369 | 3.79e-20 | 1.28e-15 |
| Malignant epithelial | C2 WNT ECs | NARF | 0.563 | 0.363 | 0.207 | 3.36e-17 | 1.13e-12 |
| Malignant epithelial | C2 WNT ECs | CREB1 | 0.563 | 0.212 | 0.128 | 6.33e-09 | 0.000213 |
| Malignant epithelial | C2 WNT ECs | ARPC4 | 0.562 | 0.698 | 0.473 | 1.01e-25 | 3.420000000000001e-21 |
| Malignant epithelial | C2 WNT ECs | CIAO1 | 0.562 | 0.363 | 0.209 | 1.87e-17 | 6.29e-13 |
| Malignant epithelial | C2 WNT ECs | CHKA | 0.562 | 0.242 | 0.147 | 5.24e-10 | 1.77e-05 |
| Malignant epithelial | C2 WNT ECs | TONSL | 0.562 | 0.184 | 0.106 | 4.63e-09 | 0.000156 |
| Malignant epithelial | C2 WNT ECs | ACTR2 | 0.561 | 0.687 | 0.477 | 4.81e-24 | 1.62e-19 |

| compartment | subcluster | gene | avg_log2FC | pct.1 | pct.2 | p_val | p_val_adj |
|---|---|---|---|---|---|---|---|
| Malignant epithelial | C2 WNT ECs | SMARCC1 | 0.561 | 0.503 | 0.314 | 1.83e-20 | 6.17e-16 |
| Malignant epithelial | C2 WNT ECs | CANT1 | 0.561 | 0.348 | 0.204 | 5.62e-15 | 1.89e-10 |
| Malignant epithelial | C2 WNT ECs | SLC25A24 | 0.561 | 0.225 | 0.122 | 9.01e-13 | 3.04e-08 |
| Malignant epithelial | C2 WNT ECs | CCDC82 | 0.561 | 0.24 | 0.135 | 3.26e-12 | 1.1e-07 |
| Malignant epithelial | C2 WNT ECs | MOSPD3 | 0.561 | 0.332 | 0.208 | 4.66e-12 | 1.57e-07 |
| Malignant epithelial | C2 WNT ECs | RNGTT | 0.561 | 0.106 | 0.056 | 4.48e-07 | 0.0151 |
| Malignant epithelial | C2 WNT ECs | PIGF | 0.56 | 0.296 | 0.155 | 4.34e-18 | 1.46e-13 |
| Malignant epithelial | C2 WNT ECs | WSB2 | 0.56 | 0.264 | 0.15 | 1.89e-13 | 6.38e-09 |
| Malignant epithelial | C2 WNT ECs | NCOR2 | 0.56 | 0.348 | 0.218 | 8.02e-13 | 2.7e-08 |
| Malignant epithelial | C2 WNT ECs | CCT5 | 0.559 | 0.743 | 0.517 | 1.97e-26 | 6.650000000000001e-22 |
| Malignant epithelial | C2 WNT ECs | TRAM1 | 0.559 | 0.687 | 0.5 | 8.670000000000001e-21 | 2.92e-16 |
| Malignant epithelial | C2 WNT ECs | COPA | 0.559 | 0.375 | 0.214 | 6.14e-18 | 2.07e-13 |
| Malignant epithelial | C2 WNT ECs | TROVE2 | 0.559 | 0.326 | 0.193 | 2.38e-14 | 8.01e-10 |
| Malignant epithelial | C2 WNT ECs | KLHL7 | 0.559 | 0.263 | 0.156 | 2.87e-11 | 9.66e-07 |
| Malignant epithelial | C2 WNT ECs | CHMP1B | 0.558 | 0.75 | 0.505 | 4.37e-30 | 1.47e-25 |
| Malignant epithelial | C2 WNT ECs | HSPA9 | 0.558 | 0.629 | 0.434 | 1.49e-21 | 5.01e-17 |
| Malignant epithelial | C2 WNT ECs | CUL3 | 0.558 | 0.315 | 0.183 | 4.59e-14 | 1.55e-09 |
| Malignant epithelial | C2 WNT ECs | TFIP11 | 0.558 | 0.129 | 0.066 | 5.5e-09 | 0.000185 |
| Malignant epithelial | C2 WNT ECs | MT-CYB | 0.557 | 0.997 | 0.983 | 8.31e-41 | 2.8e-36 |
| Malignant epithelial | C2 WNT ECs | ZDHHC9 | 0.557 | 0.376 | 0.214 | 8.56e-19 | 2.89e-14 |
| Malignant epithelial | C2 WNT ECs | RUFY1 | 0.557 | 0.207 | 0.113 | 2.7e-11 | 9.1e-07 |
| Malignant epithelial | C2 WNT ECs | PPTC7 | 0.557 | 0.257 | 0.161 | 2.67e-09 | 8.98e-05 |
| Malignant epithelial | C2 WNT ECs | VHL | 0.557 | 0.139 | 0.082 | 1.3e-06 | 0.0439 |
| Malignant epithelial | C2 WNT ECs | TRAPPC4 | 0.556 | 0.465 | 0.276 | 2.67e-20 | 9e-16 |
| Malignant epithelial | C2 WNT ECs | CLK3 | 0.556 | 0.363 | 0.206 | 7.7e-18 | 2.59e-13 |
| Malignant epithelial | C2 WNT ECs | RNPS1 | 0.555 | 0.838 | 0.642 | 2.69e-28 | 9.069999999999999e-24 |
| Malignant epithelial | C2 WNT ECs | QTRT1 | 0.555 | 0.551 | 0.344 | 7.119999999999999e-23 | 2.4e-18 |
| Malignant epithelial | C2 WNT ECs | CBLL1 | 0.555 | 0.175 | 0.1 | 1.01e-08 | 0.00034 |
| Malignant epithelial | C2 WNT ECs | DUSP4 | 0.554 | 0.378 | 0.257 | 3.24e-11 | 1.09e-06 |
| Malignant epithelial | C2 WNT ECs | MIR22HG | 0.554 | 0.195 | 0.111 | 9.83e-10 | 3.31e-05 |
| Malignant epithelial | C2 WNT ECs | RP11-327P2.5 | 0.554 | 0.135 | 0.071 | 7.28e-09 | 0.000245 |
| Malignant epithelial | C2 WNT ECs | PHOSPHO2 | 0.554 | 0.13 | 0.071 | 1.47e-07 | 0.00497 |
| Malignant epithelial | C2 WNT ECs | MDK | 0.553 | 0.924 | 0.797 | 6.16e-29 | 2.08e-24 |
| Malignant epithelial | C2 WNT ECs | PTP4A2 | 0.553 | 0.742 | 0.509 | 1.34e-27 | 4.51e-23 |
| Malignant epithelial | C2 WNT ECs | USP3 | 0.553 | 0.314 | 0.178 | 5.82e-15 | 1.96e-10 |
| Malignant epithelial | C2 WNT ECs | FBXO21 | 0.553 | 0.264 | 0.158 | 5.98e-11 | 2.01e-06 |
| Malignant epithelial | C2 WNT ECs | ST13 | 0.552 | 0.893 | 0.755 | 6.15e-30 | 2.07e-25 |
| Malignant epithelial | C2 WNT ECs | PARVA | 0.552 | 0.246 | 0.141 | 3.09e-12 | 1.04e-07 |
| Malignant epithelial | C2 WNT ECs | SLC39A14 | 0.551 | 0.185 | 0.111 | 3.23e-08 | 0.00109 |
| Malignant epithelial | C2 WNT ECs | ATP5A1 | 0.55 | 0.608 | 0.425 | 3.34e-19 | 1.13e-14 |
| Malignant epithelial | C2 WNT ECs | MARCH6 | 0.55 | 0.494 | 0.315 | 1.24e-17 | 4.19e-13 |
| Malignant epithelial | C2 WNT ECs | PUM2 | 0.55 | 0.302 | 0.182 | 3.27e-12 | 1.1e-07 |
| Malignant epithelial | C2 WNT ECs | MAPK8 | 0.55 | 0.21 | 0.117 | 1.1e-10 | 3.71e-06 |
| Malignant epithelial | C2 WNT ECs | FGFR1OP | 0.55 | 0.137 | 0.071 | 4.14e-09 | 0.000139 |
| Malignant epithelial | C2 WNT ECs | DSC2 | 0.55 | 0.494 | 0.376 | 4.73e-09 | 0.000159 |
| Malignant epithelial | C2 WNT ECs | DLAT | 0.55 | 0.138 | 0.08 | 9.08e-07 | 0.0306 |
| Malignant epithelial | C2 WNT ECs | PDCD10 | 0.549 | 0.608 | 0.393 | 1.02e-22 | 3.43e-18 |
| Malignant epithelial | C2 WNT ECs | LEMD2 | 0.549 | 0.248 | 0.139 | 1.72e-12 | 5.81e-08 |
| Malignant epithelial | C2 WNT ECs | ZSWIM8 | 0.549 | 0.169 | 0.084 | 2.68e-12 | 9.04e-08 |
| Malignant epithelial | C2 WNT ECs | SDAD1 | 0.549 | 0.235 | 0.131 | 2.81e-12 | 9.48e-08 |
| Malignant epithelial | C2 WNT ECs | TSC22D4 | 0.548 | 0.53 | 0.343 | 1.87e-17 | 6.29e-13 |
| Malignant epithelial | C2 WNT ECs | HACD3 | 0.548 | 0.455 | 0.298 | 2.13e-14 | 7.18e-10 |
| Malignant epithelial | C2 WNT ECs | STT3A | 0.548 | 0.277 | 0.156 | 3.96e-14 | 1.33e-09 |
| Malignant epithelial | C2 WNT ECs | TOMM34 | 0.548 | 0.349 | 0.218 | 5.73e-13 | 1.93e-08 |
| Malignant epithelial | C2 WNT ECs | SNRNP40 | 0.548 | 0.287 | 0.183 | 1.92e-09 | 6.47e-05 |
| Malignant epithelial | C2 WNT ECs | PGD | 0.547 | 0.277 | 0.146 | 2.86e-16 | 9.63e-12 |
| Malignant epithelial | C2 WNT ECs | USP11 | 0.547 | 0.227 | 0.12 | 1.75e-13 | 5.9e-09 |
| Malignant epithelial | C2 WNT ECs | MPZL2 | 0.547 | 0.318 | 0.205 | 3.56e-10 | 1.2e-05 |
| Malignant epithelial | C2 WNT ECs | SFI1 | 0.547 | 0.15 | 0.085 | 7.58e-08 | 0.00255 |
| Malignant epithelial | C2 WNT ECs | CTBP2 | 0.546 | 0.529 | 0.334 | 9.67e-20 | 3.26e-15 |
| Malignant epithelial | C2 WNT ECs | CORO1C | 0.546 | 0.482 | 0.307 | 1.36e-16 | 4.59e-12 |
| Malignant epithelial | C2 WNT ECs | EXOC5 | 0.546 | 0.172 | 0.102 | 1.13e-07 | 0.00379 |
| Malignant epithelial | C2 WNT ECs | PRR15L | 0.545 | 0.441 | 0.248 | 1.75e-22 | 5.89e-18 |
| Malignant epithelial | C2 WNT ECs | GOLGA5 | 0.545 | 0.198 | 0.104 | 6.4e-12 | 2.16e-07 |

| compartment | subcluster | gene | avg_log2FC | pct.1 | pct.2 | p_val | p_val_adj |
|---|---|---|---|---|---|---|---|
| Malignant epithelial | C2 WNT ECs | MLXIPL | 0.545 | 0.162 | 0.092 | 2.2e-08 | 0.00074 |
| Malignant epithelial | C2 WNT ECs | RHBDD3 | 0.545 | 0.158 | 0.095 | 7.6e-07 | 0.0256 |
| Malignant epithelial | C2 WNT ECs | BBIP1 | 0.544 | 0.223 | 0.104 | 3.79e-18 | 1.28e-13 |
| Malignant epithelial | C2 WNT ECs | RNMT | 0.544 | 0.364 | 0.21 | 4.06e-16 | 1.37e-11 |
| Malignant epithelial | C2 WNT ECs | OTUD6B-AS1 | 0.544 | 0.562 | 0.395 | 3.83e-15 | 1.29e-10 |
| Malignant epithelial | C2 WNT ECs | ADPGK | 0.544 | 0.298 | 0.174 | 1.84e-13 | 6.2e-09 |
| Malignant epithelial | C2 WNT ECs | C9orf152 | 0.544 | 0.223 | 0.122 | 4e-12 | 1.35e-07 |
| Malignant epithelial | C2 WNT ECs | BTBD7 | 0.544 | 0.242 | 0.145 | 2.39e-10 | 8.04e-06 |
| Malignant epithelial | C2 WNT ECs | SLC25A32 | 0.544 | 0.181 | 0.106 | 1.68e-08 | 0.000566 |
| Malignant epithelial | C2 WNT ECs | UBC | 0.543 | 0.993 | 0.981 | 3.33e-31 | 1.12e-26 |
| Malignant epithelial | C2 WNT ECs | ARL3 | 0.543 | 0.405 | 0.229 | 1.72e-19 | 5.8e-15 |
| Malignant epithelial | C2 WNT ECs | PPM1A | 0.543 | 0.307 | 0.187 | 3.47e-12 | 1.17e-07 |
| Malignant epithelial | C2 WNT ECs | RXRB | 0.543 | 0.226 | 0.127 | 1.75e-11 | 5.9e-07 |
| Malignant epithelial | C2 WNT ECs | GPS2 | 0.543 | 0.13 | 0.069 | 3.94e-08 | 0.00133 |
| Malignant epithelial | C2 WNT ECs | SRRM2 | 0.542 | 0.839 | 0.653 | 1.33e-25 | 4.490000000000001e-21 |
| Malignant epithelial | C2 WNT ECs | UBAC2 | 0.542 | 0.487 | 0.311 | 3.67e-18 | 1.24e-13 |
| Malignant epithelial | C2 WNT ECs | RP11-345J4.5 | 0.542 | 0.188 | 0.101 | 8.44e-11 | 2.84e-06 |
| Malignant epithelial | C2 WNT ECs | NUP54 | 0.542 | 0.219 | 0.127 | 2.52e-10 | 8.49e-06 |
| Malignant epithelial | C2 WNT ECs | GORASP1 | 0.542 | 0.147 | 0.075 | 5.01e-10 | 1.69e-05 |
| Malignant epithelial | C2 WNT ECs | CCNI | 0.541 | 0.806 | 0.617 | 3.36e-23 | 1.13e-18 |
| Malignant epithelial | C2 WNT ECs | CAPN2 | 0.541 | 0.618 | 0.433 | 6.4e-16 | 2.16e-11 |
| Malignant epithelial | C2 WNT ECs | GALNT10 | 0.541 | 0.11 | 0.059 | 5.93e-07 | 0.02 |
| Malignant epithelial | C2 WNT ECs | MRPS30 | 0.54 | 0.441 | 0.263 | 3.75e-19 | 1.26e-14 |
| Malignant epithelial | C2 WNT ECs | SLC39A4 | 0.54 | 0.893 | 0.776 | 6.59e-17 | 2.22e-12 |
| Malignant epithelial | C2 WNT ECs | BAIAP2 | 0.54 | 0.479 | 0.314 | 3.4e-16 | 1.15e-11 |
| Malignant epithelial | C2 WNT ECs | CCND3 | 0.54 | 0.365 | 0.218 | 6.81e-15 | 2.3e-10 |
| Malignant epithelial | C2 WNT ECs | AMD1 | 0.539 | 0.709 | 0.515 | 1.19e-21 | 4.01e-17 |
| Malignant epithelial | C2 WNT ECs | HIST1H2AC | 0.539 | 0.532 | 0.335 | 4.82e-19 | 1.62e-14 |
| Malignant epithelial | C2 WNT ECs | FAM21A | 0.539 | 0.175 | 0.088 | 5.13e-12 | 1.73e-07 |
| Malignant epithelial | C2 WNT ECs | PDLIM2 | 0.539 | 0.253 | 0.16 | 5.92e-09 | 0.000199 |
| Malignant epithelial | C2 WNT ECs | CARD10 | 0.539 | 0.171 | 0.105 | 6.31e-07 | 0.0213 |
| Malignant epithelial | C2 WNT ECs | ARF4 | 0.538 | 0.899 | 0.742 | 4.76e-25 | 1.6e-20 |
| Malignant epithelial | C2 WNT ECs | ATP6V1E1 | 0.538 | 0.617 | 0.432 | 1.61e-16 | 5.43e-12 |
| Malignant epithelial | C2 WNT ECs | BRD9 | 0.538 | 0.34 | 0.197 | 2.7e-15 | 9.1e-11 |
| Malignant epithelial | C2 WNT ECs | DUSP11 | 0.538 | 0.277 | 0.155 | 1.66e-14 | 5.59e-10 |
| Malignant epithelial | C2 WNT ECs | PARD6B | 0.538 | 0.442 | 0.288 | 4.21e-14 | 1.42e-09 |
| Malignant epithelial | C2 WNT ECs | USP9X | 0.538 | 0.3 | 0.178 | 3.85e-13 | 1.3e-08 |
| Malignant epithelial | C2 WNT ECs | MCFD2 | 0.537 | 0.451 | 0.269 | 4.21e-19 | 1.42e-14 |
| Malignant epithelial | C2 WNT ECs | STARD3 | 0.537 | 0.275 | 0.154 | 5.27e-14 | 1.78e-09 |
| Malignant epithelial | C2 WNT ECs | EPHX1 | 0.537 | 0.33 | 0.206 | 4.49e-12 | 1.51e-07 |
| Malignant epithelial | C2 WNT ECs | ZFAND6 | 0.536 | 0.598 | 0.394 | 4.43e-19 | 1.49e-14 |
| Malignant epithelial | C2 WNT ECs | TOM1 | 0.536 | 0.271 | 0.152 | 1.45e-13 | 4.9e-09 |
| Malignant epithelial | C2 WNT ECs | ANKRD9 | 0.536 | 0.233 | 0.128 | 2.31e-12 | 7.79e-08 |
| Malignant epithelial | C2 WNT ECs | FANCF | 0.536 | 0.217 | 0.128 | 1.7e-09 | 5.73e-05 |
| Malignant epithelial | C2 WNT ECs | BCAP31 | 0.535 | 0.903 | 0.708 | 5.19e-37 | 1.75e-32 |
| Malignant epithelial | C2 WNT ECs | CMIP | 0.535 | 0.315 | 0.202 | 4.13e-11 | 1.39e-06 |
| Malignant epithelial | C2 WNT ECs | ZNF672 | 0.535 | 0.126 | 0.07 | 3.35e-07 | 0.0113 |
| Malignant epithelial | C2 WNT ECs | NBR1 | 0.534 | 0.307 | 0.184 | 1.01e-12 | 3.41e-08 |
| Malignant epithelial | C2 WNT ECs | SLC10A3 | 0.534 | 0.269 | 0.163 | 9.93e-11 | 3.35e-06 |
| Malignant epithelial | C2 WNT ECs | CD63 | 0.533 | 0.985 | 0.96 | 2.97e-48 | 1e-43 |
| Malignant epithelial | C2 WNT ECs | GSTP1 | 0.533 | 0.981 | 0.958 | 4.06e-44 | 1.37e-39 |
| Malignant epithelial | C2 WNT ECs | PPP4C | 0.533 | 0.838 | 0.623 | 1.31e-29 | 4.41e-25 |
| Malignant epithelial | C2 WNT ECs | AKR7A3 | 0.533 | 0.24 | 0.136 | 1.69e-11 | 5.69e-07 |
| Malignant epithelial | C2 WNT ECs | SSB | 0.532 | 0.793 | 0.59 | 6.919999999999999e-28 | 2.33e-23 |
| Malignant epithelial | C2 WNT ECs | EIF4A1 | 0.532 | 0.823 | 0.643 | 2.63e-25 | 8.85e-21 |
| Malignant epithelial | C2 WNT ECs | ATRX | 0.532 | 0.56 | 0.374 | 5.46e-19 | 1.84e-14 |
| Malignant epithelial | C2 WNT ECs | ACAA1 | 0.532 | 0.589 | 0.395 | 2.36e-18 | 7.96e-14 |
| Malignant epithelial | C2 WNT ECs | S100A16 | 0.532 | 0.834 | 0.757 | 1.07e-14 | 3.6e-10 |
| Malignant epithelial | C2 WNT ECs | RNF121 | 0.532 | 0.16 | 0.09 | 2.54e-08 | 0.000856 |
| Malignant epithelial | C2 WNT ECs | NAA50 | 0.531 | 0.505 | 0.319 | 7.09e-18 | 2.39e-13 |
| Malignant epithelial | C2 WNT ECs | TXNL1 | 0.531 | 0.53 | 0.365 | 1.97e-14 | 6.63e-10 |
| Malignant epithelial | C2 WNT ECs | ARSA | 0.531 | 0.222 | 0.127 | 1.08e-10 | 3.66e-06 |
| Malignant epithelial | C2 WNT ECs | GBF1 | 0.531 | 0.16 | 0.089 | 9.01e-09 | 0.000304 |
| Malignant epithelial | C2 WNT ECs | G2E3 | 0.531 | 0.153 | 0.09 | 3.5e-07 | 0.0118 |

| compartment | subcluster | gene | avg_log2FC | pct.1 | pct.2 | p_val | p_val_adj |
|---|---|---|---|---|---|---|---|
| Malignant epithelial | C2 WNT ECs | EIF3D | 0.53 | 0.811 | 0.623 | 6.049999999999999e-24 | 2.04e-19 |
| Malignant epithelial | C2 WNT ECs | SMC3 | 0.53 | 0.461 | 0.287 | 6.02e-18 | 2.03e-13 |
| Malignant epithelial | C2 WNT ECs | DENND3 | 0.53 | 0.246 | 0.167 | 8.96e-07 | 0.0302 |
| Malignant epithelial | C2 WNT ECs | MAP2K2 | 0.529 | 0.434 | 0.275 | 4.83e-17 | 1.63e-12 |
| Malignant epithelial | C2 WNT ECs | ERBB2IP | 0.529 | 0.334 | 0.206 | 1.11e-12 | 3.73e-08 |
| Malignant epithelial | C2 WNT ECs | ATP6V0A1 | 0.529 | 0.221 | 0.12 | 2.56e-12 | 8.62e-08 |
| Malignant epithelial | C2 WNT ECs | ABLIM1 | 0.529 | 0.273 | 0.175 | 1.11e-09 | 3.72e-05 |
| Malignant epithelial | C2 WNT ECs | CCDC90B | 0.528 | 0.517 | 0.331 | 1.17e-18 | 3.94e-14 |
| Malignant epithelial | C2 WNT ECs | RPS6KB2 | 0.528 | 0.409 | 0.241 | 5.65e-17 | 1.9e-12 |
| Malignant epithelial | C2 WNT ECs | NHSL1 | 0.528 | 0.175 | 0.096 | 1.01e-09 | 3.39e-05 |
| Malignant epithelial | C2 WNT ECs | HINT3 | 0.528 | 0.198 | 0.122 | 1.1e-07 | 0.0037 |
| Malignant epithelial | C2 WNT ECs | CNNM4 | 0.528 | 0.134 | 0.077 | 6.7e-07 | 0.0226 |
| Malignant epithelial | C2 WNT ECs | UTP18 | 0.527 | 0.344 | 0.213 | 1.03e-12 | 3.48e-08 |
| Malignant epithelial | C2 WNT ECs | NECAB3 | 0.527 | 0.23 | 0.124 | 1.53e-12 | 5.14e-08 |
| Malignant epithelial | C2 WNT ECs | UBA3 | 0.527 | 0.294 | 0.177 | 9.72e-12 | 3.27e-07 |
| Malignant epithelial | C2 WNT ECs | IGFBP2 | 0.527 | 0.309 | 0.217 | 4.24e-08 | 0.00143 |
| Malignant epithelial | C2 WNT ECs | MUL1 | 0.527 | 0.133 | 0.075 | 3.23e-07 | 0.0109 |
| Malignant epithelial | C2 WNT ECs | GJB3 | 0.526 | 0.405 | 0.268 | 1.99e-12 | 6.69e-08 |
| Malignant epithelial | C2 WNT ECs | DOCK5 | 0.526 | 0.203 | 0.116 | 3.55e-10 | 1.19e-05 |
| Malignant epithelial | C2 WNT ECs | B3GALNT2 | 0.526 | 0.158 | 0.085 | 1.2e-09 | 4.06e-05 |
| Malignant epithelial | C2 WNT ECs | STARD3NL | 0.525 | 0.345 | 0.208 | 7.1e-14 | 2.39e-09 |
| Malignant epithelial | C2 WNT ECs | RNF44 | 0.525 | 0.296 | 0.171 | 2.13e-13 | 7.18e-09 |
| Malignant epithelial | C2 WNT ECs | RARA | 0.525 | 0.233 | 0.125 | 4.93e-13 | 1.66e-08 |
| Malignant epithelial | C2 WNT ECs | MSI2 | 0.525 | 0.348 | 0.219 | 2.31e-12 | 7.79e-08 |
| Malignant epithelial | C2 WNT ECs | NFYC | 0.525 | 0.234 | 0.135 | 7.76e-11 | 2.61e-06 |
| Malignant epithelial | C2 WNT ECs | SHKBP1 | 0.524 | 0.39 | 0.24 | 8.25e-15 | 2.78e-10 |
| Malignant epithelial | C2 WNT ECs | SRP72 | 0.524 | 0.452 | 0.297 | 8.87e-15 | 2.99e-10 |
| Malignant epithelial | C2 WNT ECs | BIN1 | 0.524 | 0.311 | 0.202 | 4.45e-10 | 1.5e-05 |
| Malignant epithelial | C2 WNT ECs | COG2 | 0.524 | 0.154 | 0.088 | 5.47e-08 | 0.00184 |
| Malignant epithelial | C2 WNT ECs | RALBP1 | 0.523 | 0.617 | 0.417 | 3.08e-21 | 1.04e-16 |
| Malignant epithelial | C2 WNT ECs | MYO5B | 0.523 | 0.341 | 0.211 | 4.48e-13 | 1.51e-08 |
| Malignant epithelial | C2 WNT ECs | RCC2 | 0.523 | 0.309 | 0.188 | 1.21e-11 | 4.07e-07 |
| Malignant epithelial | C2 WNT ECs | KAT5 | 0.523 | 0.311 | 0.193 | 2.32e-11 | 7.82e-07 |
| Malignant epithelial | C2 WNT ECs | UBA5 | 0.523 | 0.179 | 0.11 | 6.37e-07 | 0.0214 |
| Malignant epithelial | C2 WNT ECs | TAX1BP1 | 0.522 | 0.859 | 0.686 | 5.76e-26 | 1.94e-21 |
| Malignant epithelial | C2 WNT ECs | C16orf72 | 0.522 | 0.436 | 0.28 | 1.21e-14 | 4.06e-10 |
| Malignant epithelial | C2 WNT ECs | MPC2 | 0.522 | 0.668 | 0.5 | 1.07e-13 | 3.61e-09 |
| Malignant epithelial | C2 WNT ECs | SS18 | 0.522 | 0.212 | 0.125 | 1.22e-09 | 4.12e-05 |
| Malignant epithelial | C2 WNT ECs | PFKFB3 | 0.522 | 0.133 | 0.07 | 1.73e-08 | 0.000582 |
| Malignant epithelial | C2 WNT ECs | INTS1 | 0.521 | 0.264 | 0.153 | 3.19e-12 | 1.07e-07 |
| Malignant epithelial | C2 WNT ECs | MYO5C | 0.521 | 0.173 | 0.1 | 3.19e-08 | 0.00108 |
| Malignant epithelial | C2 WNT ECs | TYROBP | 0.521 | 0.271 | 0.18 | 1.54e-07 | 0.0052 |
| Malignant epithelial | C2 WNT ECs | MLH3 | 0.521 | 0.175 | 0.106 | 1.92e-07 | 0.00646 |
| Malignant epithelial | C2 WNT ECs | HEBP2 | 0.52 | 0.857 | 0.726 | 2.74e-22 | 9.24e-18 |
| Malignant epithelial | C2 WNT ECs | ACP5 | 0.52 | 0.166 | 0.098 | 9.03e-08 | 0.00304 |
| Malignant epithelial | C2 WNT ECs | MMADHC | 0.519 | 0.705 | 0.478 | 1.15e-25 | 3.88e-21 |
| Malignant epithelial | C2 WNT ECs | TSPYL1 | 0.519 | 0.357 | 0.213 | 4.35e-15 | 1.47e-10 |
| Malignant epithelial | C2 WNT ECs | PEX2 | 0.519 | 0.449 | 0.305 | 3.65e-12 | 1.23e-07 |
| Malignant epithelial | C2 WNT ECs | MED30 | 0.519 | 0.444 | 0.32 | 6.23e-11 | 2.1e-06 |
| Malignant epithelial | C2 WNT ECs | TNRC6A | 0.519 | 0.273 | 0.165 | 7.82e-11 | 2.64e-06 |
| Malignant epithelial | C2 WNT ECs | BCL9L | 0.519 | 0.241 | 0.155 | 4.95e-08 | 0.00167 |
| Malignant epithelial | C2 WNT ECs | KLF6 | 0.519 | 0.947 | 0.877 | 8.1e-08 | 0.00273 |
| Malignant epithelial | C2 WNT ECs | TMEM259 | 0.518 | 0.598 | 0.401 | 1.5e-18 | 5.05e-14 |
| Malignant epithelial | C2 WNT ECs | KIAA0907 | 0.518 | 0.375 | 0.234 | 7.4e-14 | 2.49e-09 |
| Malignant epithelial | C2 WNT ECs | NFS1 | 0.518 | 0.206 | 0.116 | 4.02e-10 | 1.35e-05 |
| Malignant epithelial | C2 WNT ECs | CPSF1 | 0.518 | 0.235 | 0.149 | 1.49e-08 | 0.000503 |
| Malignant epithelial | C2 WNT ECs | ZNF821 | 0.518 | 0.139 | 0.076 | 2.96e-08 | 0.000998 |
| Malignant epithelial | C2 WNT ECs | RAB35 | 0.518 | 0.207 | 0.128 | 5.87e-08 | 0.00198 |
| Malignant epithelial | C2 WNT ECs | PLOD3 | 0.517 | 0.566 | 0.365 | 7.89e-20 | 2.66e-15 |
| Malignant epithelial | C2 WNT ECs | DNAJA2 | 0.517 | 0.528 | 0.36 | 1.14e-14 | 3.86e-10 |
| Malignant epithelial | C2 WNT ECs | SUGP1 | 0.517 | 0.145 | 0.079 | 3.86e-08 | 0.0013 |
| Malignant epithelial | C2 WNT ECs | SON | 0.516 | 0.804 | 0.618 | 8.679999999999999e-24 | 2.92e-19 |
| Malignant epithelial | C2 WNT ECs | FNIP1 | 0.516 | 0.643 | 0.452 | 3.08e-19 | 1.04e-14 |
| Malignant epithelial | C2 WNT ECs | DDX6 | 0.516 | 0.457 | 0.306 | 1.02e-13 | 3.43e-09 |

| compartment | subcluster | gene | avg_log2FC | pct.1 | pct.2 | p_val | p_val_adj |
|---|---|---|---|---|---|---|---|
| Malignant epithelial | C2 WNT ECs | KAT8 | 0.516 | 0.257 | 0.147 | 1.97e-12 | 6.63e-08 |
| Malignant epithelial | C2 WNT ECs | PITRM1 | 0.516 | 0.138 | 0.079 | 2.82e-07 | 0.00949 |
| Malignant epithelial | C2 WNT ECs | ATF4 | 0.515 | 0.92 | 0.8 | 1.01e-25 | 3.420000000000001e-21 |
| Malignant epithelial | C2 WNT ECs | CCDC186 | 0.515 | 0.438 | 0.253 | 1.96e-20 | 6.6e-16 |
| Malignant epithelial | C2 WNT ECs | PCID2 | 0.515 | 0.361 | 0.212 | 1.16e-16 | 3.91e-12 |
| Malignant epithelial | C2 WNT ECs | DYNC1H1 | 0.515 | 0.43 | 0.286 | 8.48e-14 | 2.86e-09 |
| Malignant epithelial | C2 WNT ECs | STK19 | 0.515 | 0.21 | 0.121 | 6.64e-10 | 2.24e-05 |
| Malignant epithelial | C2 WNT ECs | ALS2CL | 0.515 | 0.194 | 0.114 | 7.23e-09 | 0.000244 |
| Malignant epithelial | C2 WNT ECs | PRKRA | 0.514 | 0.461 | 0.269 | 1.6e-20 | 5.4e-16 |
| Malignant epithelial | C2 WNT ECs | SLC39A1 | 0.514 | 0.483 | 0.307 | 2.21e-17 | 7.44e-13 |
| Malignant epithelial | C2 WNT ECs | PEMT | 0.514 | 0.227 | 0.132 | 2.66e-10 | 8.97e-06 |
| Malignant epithelial | C2 WNT ECs | FUNDC2 | 0.513 | 0.536 | 0.35 | 1.13e-17 | 3.8e-13 |
| Malignant epithelial | C2 WNT ECs | CYB5A | 0.513 | 0.613 | 0.433 | 6.94e-17 | 2.34e-12 |
| Malignant epithelial | C2 WNT ECs | YTHDC2 | 0.512 | 0.123 | 0.067 | 2.5e-07 | 0.00843 |
| Malignant epithelial | C2 WNT ECs | PURA | 0.511 | 0.353 | 0.221 | 4.6e-13 | 1.55e-08 |
| Malignant epithelial | C2 WNT ECs | AP4M1 | 0.511 | 0.191 | 0.103 | 4.79e-11 | 1.61e-06 |
| Malignant epithelial | C2 WNT ECs | ACP2 | 0.511 | 0.208 | 0.12 | 5.48e-10 | 1.85e-05 |
| Malignant epithelial | C2 WNT ECs | UBN2 | 0.511 | 0.143 | 0.083 | 3.01e-07 | 0.0101 |
| Malignant epithelial | C2 WNT ECs | VIMP | 0.51 | 0.809 | 0.6 | 5.62e-27 | 1.89e-22 |
| Malignant epithelial | C2 WNT ECs | DECR1 | 0.51 | 0.633 | 0.476 | 2.88e-13 | 9.71e-09 |
| Malignant epithelial | C2 WNT ECs | USP10 | 0.51 | 0.287 | 0.176 | 4.52e-11 | 1.52e-06 |
| Malignant epithelial | C2 WNT ECs | AK3 | 0.509 | 0.506 | 0.323 | 1.62e-16 | 5.45e-12 |
| Malignant epithelial | C2 WNT ECs | COMMD7 | 0.509 | 0.529 | 0.398 | 3.32e-11 | 1.12e-06 |
| Malignant epithelial | C2 WNT ECs | C2orf76 | 0.509 | 0.138 | 0.072 | 7.24e-09 | 0.000244 |
| Malignant epithelial | C2 WNT ECs | HECTD4 | 0.509 | 0.133 | 0.07 | 1.92e-08 | 0.000649 |
| Malignant epithelial | C2 WNT ECs | CDK5RAP3 | 0.508 | 0.478 | 0.302 | 4.25e-17 | 1.43e-12 |
| Malignant epithelial | C2 WNT ECs | MARK3 | 0.508 | 0.46 | 0.303 | 1.03e-13 | 3.48e-09 |
| Malignant epithelial | C2 WNT ECs | WNK1 | 0.508 | 0.402 | 0.265 | 7.58e-13 | 2.55e-08 |
| Malignant epithelial | C2 WNT ECs | NUP88 | 0.508 | 0.15 | 0.084 | 5.05e-08 | 0.0017 |
| Malignant epithelial | C2 WNT ECs | AGPAT3 | 0.508 | 0.192 | 0.116 | 7.39e-08 | 0.00249 |
| Malignant epithelial | C2 WNT ECs | VEGFA | 0.507 | 0.512 | 0.325 | 3.15e-19 | 1.06e-14 |
| Malignant epithelial | C2 WNT ECs | RANGAP1 | 0.507 | 0.298 | 0.174 | 2.19e-13 | 7.38e-09 |
| Malignant epithelial | C2 WNT ECs | CSAD | 0.507 | 0.189 | 0.105 | 3.27e-10 | 1.1e-05 |
| Malignant epithelial | C2 WNT ECs | RP11-708J19.1 | 0.507 | 0.124 | 0.069 | 3.06e-07 | 0.0103 |
| Malignant epithelial | C2 WNT ECs | SLC30A7 | 0.507 | 0.146 | 0.085 | 3.51e-07 | 0.0118 |
| Malignant epithelial | C2 WNT ECs | ATP5C1 | 0.506 | 0.862 | 0.667 | 2.65e-32 | 8.92e-28 |
| Malignant epithelial | C2 WNT ECs | FNTA | 0.506 | 0.69 | 0.493 | 3.37e-20 | 1.14e-15 |
| Malignant epithelial | C2 WNT ECs | AIFM1 | 0.506 | 0.221 | 0.138 | 2.5e-08 | 0.000842 |
| Malignant epithelial | C2 WNT ECs | LEMD3 | 0.506 | 0.118 | 0.064 | 6.52e-07 | 0.022 |
| Malignant epithelial | C2 WNT ECs | PNISR | 0.505 | 0.777 | 0.575 | 1.82e-23 | 6.14e-19 |
| Malignant epithelial | C2 WNT ECs | GOLGA4 | 0.505 | 0.586 | 0.381 | 2.99e-21 | 1.01e-16 |
| Malignant epithelial | C2 WNT ECs | SMARCE1 | 0.505 | 0.493 | 0.335 | 1.61e-13 | 5.42e-09 |
| Malignant epithelial | C2 WNT ECs | PDCD4 | 0.505 | 0.379 | 0.24 | 1.69e-13 | 5.71e-09 |
| Malignant epithelial | C2 WNT ECs | PDRG1 | 0.505 | 0.38 | 0.24 | 2.34e-13 | 7.9e-09 |
| Malignant epithelial | C2 WNT ECs | CPPED1 | 0.505 | 0.257 | 0.163 | 4.23e-09 | 0.000142 |
| Malignant epithelial | C2 WNT ECs | POLB | 0.505 | 0.153 | 0.09 | 3.26e-07 | 0.011 |
| Malignant epithelial | C2 WNT ECs | DDX56 | 0.504 | 0.432 | 0.261 | 4.03e-17 | 1.36e-12 |
| Malignant epithelial | C2 WNT ECs | PPP5C | 0.504 | 0.261 | 0.146 | 2.74e-13 | 9.24e-09 |
| Malignant epithelial | C2 WNT ECs | CPSF7 | 0.504 | 0.254 | 0.141 | 3.95e-13 | 1.33e-08 |
| Malignant epithelial | C2 WNT ECs | ACAT1 | 0.504 | 0.281 | 0.192 | 6.5e-08 | 0.00219 |
| Malignant epithelial | C2 WNT ECs | SLC2A8 | 0.504 | 0.156 | 0.091 | 2.17e-07 | 0.00732 |
| Malignant epithelial | C2 WNT ECs | PRPF40A | 0.503 | 0.647 | 0.444 | 5.41e-19 | 1.82e-14 |
| Malignant epithelial | C2 WNT ECs | ARFRP1 | 0.503 | 0.455 | 0.286 | 1.79e-15 | 6.02e-11 |
| Malignant epithelial | C2 WNT ECs | ARMCX6 | 0.503 | 0.394 | 0.255 | 5.55e-13 | 1.87e-08 |
| Malignant epithelial | C2 WNT ECs | SERBP1 | 0.502 | 0.801 | 0.624 | 2.61e-24 | 8.79e-20 |
| Malignant epithelial | C2 WNT ECs | CCT2 | 0.502 | 0.748 | 0.537 | 4.76e-21 | 1.6e-16 |
| Malignant epithelial | C2 WNT ECs | TCF7 | 0.502 | 0.451 | 0.262 | 6.360000000000001e-21 | 2.14e-16 |
| Malignant epithelial | C2 WNT ECs | GRPEL1 | 0.502 | 0.425 | 0.276 | 1.11e-13 | 3.75e-09 |
| Malignant epithelial | C2 WNT ECs | SLC35F5 | 0.502 | 0.298 | 0.171 | 1.16e-13 | 3.91e-09 |
| Malignant epithelial | C2 WNT ECs | CWC27 | 0.502 | 0.322 | 0.198 | 2.66e-12 | 8.98e-08 |
| Malignant epithelial | C2 WNT ECs | DMWD | 0.502 | 0.172 | 0.088 | 2.04e-11 | 6.86e-07 |
| Malignant epithelial | C2 WNT ECs | SGSM3 | 0.502 | 0.191 | 0.121 | 7.29e-07 | 0.0246 |
| Malignant epithelial | C2 WNT ECs | ATP5F1 | 0.501 | 0.71 | 0.509 | 7.5e-23 | 2.53e-18 |
| Malignant epithelial | C2 WNT ECs | VARS | 0.501 | 0.283 | 0.16 | 5.14e-14 | 1.73e-09 |

| compartment | subcluster | gene | avg_log2FC | pct.1 | pct.2 | p_val | p_val_adj |
|---|---|---|---|---|---|---|---|
| Malignant epithelial | C2 WNT ECs | UBQLN2 | 0.501 | 0.24 | 0.142 | 2.5e-10 | 8.42e-06 |
| Malignant epithelial | C2 WNT ECs | NEDD9 | 0.501 | 0.208 | 0.126 | 1.53e-08 | 0.000516 |
| Malignant epithelial | C2 WNT ECs | CHMP7 | 0.501 | 0.11 | 0.059 | 8.33e-07 | 0.0281 |
| Malignant epithelial | C2 WNT ECs | XRCC6 | 0.5 | 0.739 | 0.55 | 2.62e-20 | 8.82e-16 |
| Malignant epithelial | C2 WNT ECs | DYNC1LI1 | 0.5 | 0.306 | 0.188 | 8.98e-12 | 3.03e-07 |
| Malignant epithelial | C2 WNT ECs | RANBP2 | 0.5 | 0.238 | 0.136 | 1.45e-11 | 4.9e-07 |
| Malignant epithelial | C2 WNT ECs | KDM2B | 0.5 | 0.112 | 0.06 | 3.74e-07 | 0.0126 |
| Malignant epithelial | C2 WNT ECs | RAD1 | 0.499 | 0.264 | 0.161 | 4.36e-10 | 1.47e-05 |
| Malignant epithelial | C2 WNT ECs | HTATIP2 | 0.498 | 0.71 | 0.531 | 4.28e-20 | 1.44e-15 |
| Malignant epithelial | C2 WNT ECs | MTCH2 | 0.498 | 0.64 | 0.44 | 2.23e-19 | 7.5e-15 |
| Malignant epithelial | C2 WNT ECs | YWHAG | 0.498 | 0.635 | 0.434 | 3.79e-17 | 1.28e-12 |
| Malignant epithelial | C2 WNT ECs | ASH1L | 0.498 | 0.378 | 0.233 | 5.15e-15 | 1.73e-10 |
| Malignant epithelial | C2 WNT ECs | HDAC3 | 0.498 | 0.35 | 0.213 | 1.82e-13 | 6.12e-09 |
| Malignant epithelial | C2 WNT ECs | DRG2 | 0.498 | 0.192 | 0.113 | 1.65e-08 | 0.000556 |
| Malignant epithelial | C2 WNT ECs | CCAR1 | 0.497 | 0.413 | 0.252 | 3.61e-16 | 1.22e-11 |
| Malignant epithelial | C2 WNT ECs | CBFB | 0.497 | 0.363 | 0.233 | 8.8e-12 | 2.96e-07 |
| Malignant epithelial | C2 WNT ECs | MAN1A2 | 0.497 | 0.229 | 0.138 | 2.23e-09 | 7.51e-05 |
| Malignant epithelial | C2 WNT ECs | SPPL2A | 0.497 | 0.272 | 0.174 | 5.16e-09 | 0.000174 |
| Malignant epithelial | C2 WNT ECs | PAIP2 | 0.496 | 0.678 | 0.473 | 7.1e-20 | 2.39e-15 |
| Malignant epithelial | C2 WNT ECs | CHMP1A | 0.496 | 0.539 | 0.361 | 1.23e-15 | 4.16e-11 |
| Malignant epithelial | C2 WNT ECs | FAM234A | 0.496 | 0.363 | 0.225 | 4.46e-13 | 1.5e-08 |
| Malignant epithelial | C2 WNT ECs | GABRE | 0.496 | 0.258 | 0.148 | 7.08e-13 | 2.38e-08 |
| Malignant epithelial | C2 WNT ECs | NMT2 | 0.496 | 0.183 | 0.1 | 3e-10 | 1.01e-05 |
| Malignant epithelial | C2 WNT ECs | RP11-390E23.6 | 0.496 | 0.219 | 0.138 | 5.96e-08 | 0.00201 |
| Malignant epithelial | C2 WNT ECs | MSRB2 | 0.495 | 0.658 | 0.422 | 4.19e-23 | 1.41e-18 |
| Malignant epithelial | C2 WNT ECs | JUP | 0.495 | 0.867 | 0.723 | 2.3e-14 | 7.75e-10 |
| Malignant epithelial | C2 WNT ECs | FBRS | 0.495 | 0.273 | 0.159 | 5.57e-12 | 1.88e-07 |
| Malignant epithelial | C2 WNT ECs | SHOC2 | 0.495 | 0.33 | 0.211 | 1.12e-10 | 3.76e-06 |
| Malignant epithelial | C2 WNT ECs | AP5M1 | 0.495 | 0.198 | 0.115 | 4.31e-09 | 0.000145 |
| Malignant epithelial | C2 WNT ECs | BLZF1 | 0.495 | 0.21 | 0.134 | 2.54e-07 | 0.00855 |
| Malignant epithelial | C2 WNT ECs | MAP2K3 | 0.494 | 0.413 | 0.262 | 2.14e-14 | 7.22e-10 |
| Malignant epithelial | C2 WNT ECs | HGSNAT | 0.494 | 0.338 | 0.209 | 1.78e-12 | 5.99e-08 |
| Malignant epithelial | C2 WNT ECs | D2HGDH | 0.494 | 0.196 | 0.108 | 2.22e-10 | 7.47e-06 |
| Malignant epithelial | C2 WNT ECs | APPBP2 | 0.494 | 0.137 | 0.071 | 4.04e-09 | 0.000136 |
| Malignant epithelial | C2 WNT ECs | INPPL1 | 0.494 | 0.162 | 0.09 | 5.09e-09 | 0.000171 |
| Malignant epithelial | C2 WNT ECs | ARAP1 | 0.494 | 0.139 | 0.074 | 9.94e-09 | 0.000335 |
| Malignant epithelial | C2 WNT ECs | TMEM179B | 0.493 | 0.67 | 0.48 | 5.4e-21 | 1.82e-16 |
| Malignant epithelial | C2 WNT ECs | VAMP2 | 0.493 | 0.548 | 0.359 | 1.05e-17 | 3.55e-13 |
| Malignant epithelial | C2 WNT ECs | SLC33A1 | 0.493 | 0.176 | 0.079 | 1.28e-15 | 4.32e-11 |
| Malignant epithelial | C2 WNT ECs | SAT2 | 0.493 | 0.445 | 0.286 | 1.1e-14 | 3.7e-10 |
| Malignant epithelial | C2 WNT ECs | ARHGEF7 | 0.493 | 0.256 | 0.139 | 3.82e-14 | 1.29e-09 |
| Malignant epithelial | C2 WNT ECs | LRP5 | 0.493 | 0.304 | 0.177 | 1.28e-13 | 4.31e-09 |
| Malignant epithelial | C2 WNT ECs | INPP1 | 0.493 | 0.338 | 0.218 | 6.57e-11 | 2.21e-06 |
| Malignant epithelial | C2 WNT ECs | DYNC1LI2 | 0.492 | 0.514 | 0.334 | 7.9e-16 | 2.66e-11 |
| Malignant epithelial | C2 WNT ECs | HPCAL1 | 0.491 | 0.572 | 0.384 | 4.2e-18 | 1.41e-13 |
| Malignant epithelial | C2 WNT ECs | METTL9 | 0.491 | 0.579 | 0.435 | 1.85e-11 | 6.24e-07 |
| Malignant epithelial | C2 WNT ECs | G6PD | 0.491 | 0.252 | 0.153 | 3.85e-10 | 1.3e-05 |
| Malignant epithelial | C2 WNT ECs | DCAF5 | 0.491 | 0.168 | 0.098 | 7.11e-08 | 0.0024 |
| Malignant epithelial | C2 WNT ECs | ZZZ3 | 0.491 | 0.127 | 0.07 | 2.04e-07 | 0.00688 |
| Malignant epithelial | C2 WNT ECs | UBE2H | 0.49 | 0.518 | 0.341 | 3.92e-15 | 1.32e-10 |
| Malignant epithelial | C2 WNT ECs | TSC2 | 0.49 | 0.204 | 0.119 | 1.22e-09 | 4.1e-05 |
| Malignant epithelial | C2 WNT ECs | MAPK6 | 0.49 | 0.337 | 0.232 | 1.13e-08 | 0.000381 |
| Malignant epithelial | C2 WNT ECs | KDM5C | 0.49 | 0.202 | 0.12 | 1.22e-08 | 0.00041 |
| Malignant epithelial | C2 WNT ECs | ELAC2 | 0.49 | 0.138 | 0.081 | 1.35e-06 | 0.0453 |
| Malignant epithelial | C2 WNT ECs | SPINT2 | 0.489 | 0.984 | 0.932 | 8.21e-43 | 2.77e-38 |
| Malignant epithelial | C2 WNT ECs | KDELR1 | 0.488 | 0.882 | 0.743 | 2.65e-29 | 8.94e-25 |
| Malignant epithelial | C2 WNT ECs | COQ10B | 0.488 | 0.414 | 0.27 | 4.2e-12 | 1.42e-07 |
| Malignant epithelial | C2 WNT ECs | UBE4A | 0.488 | 0.198 | 0.107 | 6.48e-11 | 2.18e-06 |
| Malignant epithelial | C2 WNT ECs | PSMD10 | 0.488 | 0.234 | 0.139 | 6.38e-10 | 2.15e-05 |
| Malignant epithelial | C2 WNT ECs | DIDO1 | 0.488 | 0.286 | 0.181 | 8.38e-10 | 2.83e-05 |
| Malignant epithelial | C2 WNT ECs | MYO10 | 0.488 | 0.387 | 0.274 | 1.08e-08 | 0.000364 |
| Malignant epithelial | C2 WNT ECs | CEACAM6 | 0.487 | 0.629 | 0.391 | 2.02e-29 | 6.8e-25 |
| Malignant epithelial | C2 WNT ECs | SLC25A36 | 0.487 | 0.432 | 0.265 | 5.94e-16 | 2e-11 |
| Malignant epithelial | C2 WNT ECs | PRNP | 0.487 | 0.495 | 0.342 | 2.86e-13 | 9.63e-09 |

| compartment | subcluster | gene | avg_log2FC | pct.1 | pct.2 | p_val | p_val_adj |
|---|---|---|---|---|---|---|---|
| Malignant epithelial | C2 WNT ECs | WIPF2 | 0.487 | 0.318 | 0.199 | 4.6e-11 | 1.55e-06 |
| Malignant epithelial | C2 WNT ECs | ZNF76 | 0.487 | 0.139 | 0.077 | 6.33e-08 | 0.00213 |
| Malignant epithelial | C2 WNT ECs | PHF14 | 0.486 | 0.748 | 0.538 | 9.69e-23 | 3.27e-18 |
| Malignant epithelial | C2 WNT ECs | TGOLN2 | 0.486 | 0.558 | 0.371 | 2.39e-17 | 8.07e-13 |
| Malignant epithelial | C2 WNT ECs | PDCD2 | 0.486 | 0.524 | 0.337 | 3.41e-17 | 1.15e-12 |
| Malignant epithelial | C2 WNT ECs | DNAJC10 | 0.486 | 0.283 | 0.161 | 1.49e-13 | 5e-09 |
| Malignant epithelial | C2 WNT ECs | MAT2B | 0.486 | 0.346 | 0.211 | 5.1e-13 | 1.72e-08 |
| Malignant epithelial | C2 WNT ECs | JMJD8 | 0.486 | 0.315 | 0.19 | 9.05e-13 | 3.05e-08 |
| Malignant epithelial | C2 WNT ECs | ARFGEF1 | 0.486 | 0.271 | 0.174 | 8.29e-09 | 0.000279 |
| Malignant epithelial | C2 WNT ECs | UBQLN4 | 0.486 | 0.166 | 0.094 | 2.29e-08 | 0.00077 |
| Malignant epithelial | C2 WNT ECs | PRR4 | 0.486 | 0.118 | 0.064 | 3.85e-07 | 0.013 |
| Malignant epithelial | C2 WNT ECs | TTC3 | 0.485 | 0.635 | 0.408 | 8.35e-23 | 2.81e-18 |
| Malignant epithelial | C2 WNT ECs | XIAP | 0.485 | 0.36 | 0.24 | 2.01e-10 | 6.77e-06 |
| Malignant epithelial | C2 WNT ECs | GOLGA3 | 0.485 | 0.211 | 0.121 | 5.11e-10 | 1.72e-05 |
| Malignant epithelial | C2 WNT ECs | ZNF600 | 0.485 | 0.161 | 0.095 | 1.54e-07 | 0.00519 |
| Malignant epithelial | C2 WNT ECs | BRAP | 0.485 | 0.135 | 0.077 | 3.55e-07 | 0.012 |
| Malignant epithelial | C2 WNT ECs | AFF4 | 0.484 | 0.491 | 0.295 | 1.44e-20 | 4.86e-16 |
| Malignant epithelial | C2 WNT ECs | ZFP91 | 0.484 | 0.263 | 0.159 | 1.99e-10 | 6.71e-06 |
| Malignant epithelial | C2 WNT ECs | PLAA | 0.484 | 0.18 | 0.099 | 1.23e-09 | 4.15e-05 |
| Malignant epithelial | C2 WNT ECs | APPL1 | 0.484 | 0.264 | 0.169 | 6.61e-09 | 0.000223 |
| Malignant epithelial | C2 WNT ECs | FAM98A | 0.484 | 0.15 | 0.085 | 4.44e-08 | 0.0015 |
| Malignant epithelial | C2 WNT ECs | ACTR10 | 0.483 | 0.445 | 0.269 | 2.12e-18 | 7.14e-14 |
| Malignant epithelial | C2 WNT ECs | EHD4 | 0.483 | 0.474 | 0.343 | 1.1e-09 | 3.71e-05 |
| Malignant epithelial | C2 WNT ECs | NR1H3 | 0.483 | 0.156 | 0.085 | 7.6e-09 | 0.000256 |
| Malignant epithelial | C2 WNT ECs | SKIV2L | 0.483 | 0.145 | 0.084 | 4.21e-07 | 0.0142 |
| Malignant epithelial | C2 WNT ECs | MBD1 | 0.483 | 0.138 | 0.08 | 8.92e-07 | 0.0301 |
| Malignant epithelial | C2 WNT ECs | ASAH1 | 0.482 | 0.525 | 0.354 | 3.32e-15 | 1.12e-10 |
| Malignant epithelial | C2 WNT ECs | SLC35B1 | 0.482 | 0.478 | 0.316 | 4.93e-14 | 1.66e-09 |
| Malignant epithelial | C2 WNT ECs | FHIT | 0.482 | 0.147 | 0.075 | 7.12e-10 | 2.4e-05 |
| Malignant epithelial | C2 WNT ECs | CCDC130 | 0.482 | 0.23 | 0.141 | 4.11e-09 | 0.000138 |
| Malignant epithelial | C2 WNT ECs | KDSR | 0.481 | 0.363 | 0.223 | 1.58e-13 | 5.33e-09 |
| Malignant epithelial | C2 WNT ECs | GLS | 0.481 | 0.502 | 0.346 | 5.42e-13 | 1.83e-08 |
| Malignant epithelial | C2 WNT ECs | PLXNA3 | 0.481 | 0.234 | 0.128 | 5.71e-13 | 1.92e-08 |
| Malignant epithelial | C2 WNT ECs | INSIG2 | 0.481 | 0.302 | 0.181 | 1.96e-12 | 6.62e-08 |
| Malignant epithelial | C2 WNT ECs | C3orf17 | 0.481 | 0.191 | 0.116 | 6.86e-08 | 0.00231 |
| Malignant epithelial | C2 WNT ECs | BZW1 | 0.48 | 0.724 | 0.521 | 2.41e-21 | 8.11e-17 |
| Malignant epithelial | C2 WNT ECs | RBBP7 | 0.48 | 0.506 | 0.335 | 8.22e-15 | 2.77e-10 |
| Malignant epithelial | C2 WNT ECs | LSM14A | 0.48 | 0.59 | 0.411 | 1.17e-14 | 3.93e-10 |
| Malignant epithelial | C2 WNT ECs | TNPO1 | 0.48 | 0.337 | 0.201 | 2.43e-13 | 8.2e-09 |
| Malignant epithelial | C2 WNT ECs | MED13 | 0.48 | 0.254 | 0.143 | 1.38e-12 | 4.66e-08 |
| Malignant epithelial | C2 WNT ECs | PDHB | 0.48 | 0.39 | 0.253 | 2.97e-12 | 1e-07 |
| Malignant epithelial | C2 WNT ECs | SARS2 | 0.48 | 0.118 | 0.064 | 5.55e-07 | 0.0187 |
| Malignant epithelial | C2 WNT ECs | TPR | 0.479 | 0.524 | 0.331 | 1.69e-18 | 5.69e-14 |
| Malignant epithelial | C2 WNT ECs | AHI1 | 0.479 | 0.311 | 0.182 | 3.2e-13 | 1.08e-08 |
| Malignant epithelial | C2 WNT ECs | MGEA5 | 0.479 | 0.549 | 0.396 | 5.96e-13 | 2.01e-08 |
| Malignant epithelial | C2 WNT ECs | TXNRD1 | 0.479 | 0.402 | 0.296 | 1.3e-06 | 0.0436 |
| Malignant epithelial | C2 WNT ECs | RNF167 | 0.478 | 0.401 | 0.224 | 4.95e-19 | 1.67e-14 |
| Malignant epithelial | C2 WNT ECs | ZNF428 | 0.478 | 0.503 | 0.349 | 1.37e-14 | 4.6e-10 |
| Malignant epithelial | C2 WNT ECs | SPIN2B | 0.478 | 0.147 | 0.077 | 1.57e-09 | 5.31e-05 |
| Malignant epithelial | C2 WNT ECs | UPF2 | 0.477 | 0.558 | 0.344 | 6.03e-21 | 2.03e-16 |
| Malignant epithelial | C2 WNT ECs | TAX1BP3 | 0.477 | 0.65 | 0.451 | 6.43e-18 | 2.17e-13 |
| Malignant epithelial | C2 WNT ECs | ADIPOR1 | 0.477 | 0.544 | 0.358 | 1.4e-17 | 4.72e-13 |
| Malignant epithelial | C2 WNT ECs | BRAT1 | 0.477 | 0.29 | 0.172 | 3.07e-12 | 1.03e-07 |
| Malignant epithelial | C2 WNT ECs | MPV17L2 | 0.477 | 0.169 | 0.093 | 3.65e-09 | 0.000123 |
| Malignant epithelial | C2 WNT ECs | SORBS3 | 0.477 | 0.195 | 0.117 | 5.23e-08 | 0.00176 |
| Malignant epithelial | C2 WNT ECs | SLC45A4 | 0.477 | 0.135 | 0.077 | 2.8e-07 | 0.00943 |
| Malignant epithelial | C2 WNT ECs | QSOX1 | 0.476 | 0.474 | 0.267 | 2.5e-23 | 8.41e-19 |
| Malignant epithelial | C2 WNT ECs | DSG2 | 0.476 | 0.602 | 0.428 | 4.43e-16 | 1.49e-11 |
| Malignant epithelial | C2 WNT ECs | ZBTB38 | 0.476 | 0.452 | 0.292 | 9.56e-15 | 3.22e-10 |
| Malignant epithelial | C2 WNT ECs | MFSD11 | 0.476 | 0.235 | 0.141 | 1.5e-09 | 5.07e-05 |
| Malignant epithelial | C2 WNT ECs | UBA1 | 0.475 | 0.559 | 0.348 | 1.22e-20 | 4.1e-16 |
| Malignant epithelial | C2 WNT ECs | ADAP1 | 0.475 | 0.568 | 0.403 | 1.75e-14 | 5.9e-10 |
| Malignant epithelial | C2 WNT ECs | SNX17 | 0.475 | 0.41 | 0.28 | 9.84e-11 | 3.32e-06 |
| Malignant epithelial | C2 WNT ECs | CKMT1A | 0.475 | 0.119 | 0.061 | 2.56e-08 | 0.000862 |

| compartment | subcluster | gene | avg_log2FC | pct.1 | pct.2 | p_val | p_val_adj |
|---|---|---|---|---|---|---|---|
| Malignant epithelial | C2 WNT ECs | HMGB3 | 0.474 | 0.499 | 0.307 | 3.71e-19 | 1.25e-14 |
| Malignant epithelial | C2 WNT ECs | TRAF4 | 0.474 | 0.587 | 0.389 | 1.37e-17 | 4.6e-13 |
| Malignant epithelial | C2 WNT ECs | IFNGR1 | 0.474 | 0.342 | 0.206 | 2e-13 | 6.75e-09 |
| Malignant epithelial | C2 WNT ECs | SLC35F6 | 0.474 | 0.272 | 0.154 | 3.82e-13 | 1.29e-08 |
| Malignant epithelial | C2 WNT ECs | PRKAR1B | 0.474 | 0.267 | 0.162 | 1.93e-10 | 6.51e-06 |
| Malignant epithelial | C2 WNT ECs | CNOT3 | 0.474 | 0.241 | 0.143 | 7.51e-10 | 2.53e-05 |
| Malignant epithelial | C2 WNT ECs | UBP1 | 0.474 | 0.217 | 0.132 | 1.38e-08 | 0.000465 |
| Malignant epithelial | C2 WNT ECs | SLC38A7 | 0.474 | 0.114 | 0.06 | 1.69e-07 | 0.0057 |
| Malignant epithelial | C2 WNT ECs | OCIAD1 | 0.473 | 0.678 | 0.448 | 3.9099999999999999e-23 | 1.32e-18 |
| Malignant epithelial | C2 WNT ECs | MYCBP2 | 0.473 | 0.304 | 0.169 | 1.72e-15 | 5.79e-11 |
| Malignant epithelial | C2 WNT ECs | STX8 | 0.473 | 0.281 | 0.162 | 6.5e-13 | 2.19e-08 |
| Malignant epithelial | C2 WNT ECs | RBM19 | 0.473 | 0.133 | 0.072 | 7.57e-08 | 0.00255 |
| Malignant epithelial | C2 WNT ECs | MBD6 | 0.473 | 0.194 | 0.117 | 9.19e-08 | 0.0031 |
| Malignant epithelial | C2 WNT ECs | MST1R | 0.473 | 0.21 | 0.137 | 7.6e-07 | 0.0256 |
| Malignant epithelial | C2 WNT ECs | C20orf24 | 0.472 | 0.861 | 0.679 | 1.35e-23 | 4.54e-19 |
| Malignant epithelial | C2 WNT ECs | FGD5-AS1 | 0.472 | 0.303 | 0.186 | 2.95e-11 | 9.95e-07 |
| Malignant epithelial | C2 WNT ECs | C21orf2 | 0.472 | 0.249 | 0.149 | 2.51e-10 | 8.46e-06 |
| Malignant epithelial | C2 WNT ECs | TMEM199 | 0.472 | 0.276 | 0.181 | 1.3e-08 | 0.000439 |
| Malignant epithelial | C2 WNT ECs | TBC1D15 | 0.472 | 0.223 | 0.138 | 1.81e-08 | 0.000611 |
| Malignant epithelial | C2 WNT ECs | PGM2 | 0.472 | 0.114 | 0.061 | 3.8e-07 | 0.0128 |
| Malignant epithelial | C2 WNT ECs | PPP1R12A | 0.471 | 0.464 | 0.275 | 5.55e-20 | 1.87e-15 |
| Malignant epithelial | C2 WNT ECs | EIF2AK1 | 0.47 | 0.553 | 0.372 | 2.97e-17 | 1e-12 |
| Malignant epithelial | C2 WNT ECs | SELENBP1 | 0.47 | 0.199 | 0.112 | 3.51e-10 | 1.18e-05 |
| Malignant epithelial | C2 WNT ECs | EMP2 | 0.469 | 0.701 | 0.468 | 2.36e-23 | 7.96e-19 |
| Malignant epithelial | C2 WNT ECs | RARA-AS1 | 0.469 | 0.149 | 0.086 | 2.39e-07 | 0.00805 |
| Malignant epithelial | C2 WNT ECs | CDC42EP5 | 0.468 | 0.271 | 0.146 | 2.73e-15 | 9.18e-11 |
| Malignant epithelial | C2 WNT ECs | TCEAL1 | 0.468 | 0.257 | 0.143 | 1.77e-13 | 5.97e-09 |
| Malignant epithelial | C2 WNT ECs | SPG7 | 0.468 | 0.41 | 0.27 | 7.65e-12 | 2.58e-07 |
| Malignant epithelial | C2 WNT ECs | KIAA0196 | 0.468 | 0.184 | 0.106 | 4.9e-09 | 0.000165 |
| Malignant epithelial | C2 WNT ECs | KDM4B | 0.468 | 0.168 | 0.094 | 6.74e-09 | 0.000227 |
| Malignant epithelial | C2 WNT ECs | ZNF587 | 0.468 | 0.116 | 0.06 | 6.96e-08 | 0.00235 |
| Malignant epithelial | C2 WNT ECs | CTNND1 | 0.467 | 0.602 | 0.409 | 3.22e-17 | 1.08e-12 |
| Malignant epithelial | C2 WNT ECs | PPP1CB | 0.467 | 0.719 | 0.565 | 8.07e-13 | 2.72e-08 |
| Malignant epithelial | C2 WNT ECs | GNL2 | 0.467 | 0.234 | 0.134 | 5.28e-11 | 1.78e-06 |
| Malignant epithelial | C2 WNT ECs | FNBP1 | 0.467 | 0.158 | 0.086 | 3.44e-09 | 0.000116 |
| Malignant epithelial | C2 WNT ECs | IL4R | 0.467 | 0.229 | 0.14 | 5.73e-09 | 0.000193 |
| Malignant epithelial | C2 WNT ECs | ZFYVE27 | 0.467 | 0.191 | 0.116 | 8.45e-08 | 0.00285 |
| Malignant epithelial | C2 WNT ECs | CNPPD1 | 0.466 | 0.419 | 0.251 | 7.32e-17 | 2.47e-12 |
| Malignant epithelial | C2 WNT ECs | CYP20A1 | 0.466 | 0.288 | 0.169 | 7.06e-13 | 2.38e-08 |
| Malignant epithelial | C2 WNT ECs | RALA | 0.465 | 0.754 | 0.577 | 7.03e-18 | 2.37e-13 |
| Malignant epithelial | C2 WNT ECs | NAP1L4 | 0.465 | 0.543 | 0.36 | 2.99e-16 | 1.01e-11 |
| Malignant epithelial | C2 WNT ECs | KLHDC2 | 0.465 | 0.448 | 0.285 | 1.16e-15 | 3.9e-11 |
| Malignant epithelial | C2 WNT ECs | SCRN2 | 0.465 | 0.3 | 0.175 | 5.08e-13 | 1.71e-08 |
| Malignant epithelial | C2 WNT ECs | SPTAN1 | 0.465 | 0.512 | 0.376 | 1.43e-10 | 4.82e-06 |
| Malignant epithelial | C2 WNT ECs | OSR2 | 0.465 | 0.263 | 0.18 | 7.48e-07 | 0.0252 |
| Malignant epithelial | C2 WNT ECs | CCDC115 | 0.464 | 0.452 | 0.288 | 1.3e-14 | 4.37e-10 |
| Malignant epithelial | C2 WNT ECs | CNN2 | 0.464 | 0.675 | 0.575 | 6e-10 | 2.02e-05 |
| Malignant epithelial | C2 WNT ECs | CTDSPL | 0.464 | 0.218 | 0.133 | 1.02e-08 | 0.000344 |
| Malignant epithelial | C2 WNT ECs | HSPE1 | 0.463 | 0.972 | 0.931 | 1.26e-21 | 4.24e-17 |
| Malignant epithelial | C2 WNT ECs | MRPL3 | 0.463 | 0.585 | 0.407 | 1.13e-15 | 3.79e-11 |
| Malignant epithelial | C2 WNT ECs | USP47 | 0.463 | 0.277 | 0.153 | 4.73e-14 | 1.59e-09 |
| Malignant epithelial | C2 WNT ECs | LRRFIP1 | 0.463 | 0.759 | 0.596 | 1.55e-13 | 5.23e-09 |
| Malignant epithelial | C2 WNT ECs | SUPT4H1 | 0.463 | 0.67 | 0.504 | 1.94e-13 | 6.53e-09 |
| Malignant epithelial | C2 WNT ECs | PODXL | 0.463 | 0.168 | 0.088 | 1.45e-10 | 4.9e-06 |
| Malignant epithelial | C2 WNT ECs | NSDHL | 0.463 | 0.221 | 0.137 | 2.49e-08 | 0.000838 |
| Malignant epithelial | C2 WNT ECs | LETM1 | 0.463 | 0.204 | 0.126 | 6.4e-08 | 0.00216 |
| Malignant epithelial | C2 WNT ECs | ZBED5 | 0.463 | 0.18 | 0.109 | 1.66e-07 | 0.00558 |
| Malignant epithelial | C2 WNT ECs | VCP | 0.462 | 0.566 | 0.376 | 4.11e-17 | 1.39e-12 |
| Malignant epithelial | C2 WNT ECs | THEM4 | 0.462 | 0.16 | 0.094 | 1.81e-07 | 0.0061 |
| Malignant epithelial | C2 WNT ECs | PPARG | 0.462 | 0.218 | 0.14 | 3.23e-07 | 0.0109 |
| Malignant epithelial | C2 WNT ECs | FARP2 | 0.462 | 0.177 | 0.111 | 9e-07 | 0.0303 |
| Malignant epithelial | C2 WNT ECs | LUC7L2 | 0.461 | 0.449 | 0.281 | 5.25e-16 | 1.77e-11 |
| Malignant epithelial | C2 WNT ECs | TFDP1 | 0.461 | 0.365 | 0.226 | 3.28e-13 | 1.11e-08 |
| Malignant epithelial | C2 WNT ECs | ITSN1 | 0.461 | 0.177 | 0.096 | 3.99e-10 | 1.34e-05 |

| compartment | subcluster | gene | avg_log2FC | pct.1 | pct.2 | p_val | p_val_adj |
|---|---|---|---|---|---|---|---|
| Malignant epithelial | C2 WNT ECs | MARCH3 | 0.461 | 0.18 | 0.106 | 5.51e-08 | 0.00186 |
| Malignant epithelial | C2 WNT ECs | HNRNPR | 0.46 | 0.578 | 0.384 | 5.8e-18 | 1.95e-13 |
| Malignant epithelial | C2 WNT ECs | TMF1 | 0.46 | 0.33 | 0.192 | 6.27e-15 | 2.11e-10 |
| Malignant epithelial | C2 WNT ECs | GLG1 | 0.46 | 0.484 | 0.326 | 1.94e-13 | 6.54e-09 |
| Malignant epithelial | C2 WNT ECs | ARF1 | 0.459 | 0.924 | 0.764 | 9.87e-28 | 3.33e-23 |
| Malignant epithelial | C2 WNT ECs | MED15 | 0.459 | 0.414 | 0.258 | 1.02e-14 | 3.44e-10 |
| Malignant epithelial | C2 WNT ECs | ATG4B | 0.459 | 0.413 | 0.261 | 5.4e-14 | 1.82e-09 |
| Malignant epithelial | C2 WNT ECs | IMMT | 0.459 | 0.378 | 0.241 | 1.63e-12 | 5.5e-08 |
| Malignant epithelial | C2 WNT ECs | IMMP1L | 0.458 | 0.3 | 0.184 | 3.62e-11 | 1.22e-06 |
| Malignant epithelial | C2 WNT ECs | RSAD1 | 0.458 | 0.135 | 0.073 | 5.08e-08 | 0.00171 |
| Malignant epithelial | C2 WNT ECs | RHOT1 | 0.458 | 0.192 | 0.12 | 3.15e-07 | 0.0106 |
| Malignant epithelial | C2 WNT ECs | PSMD8 | 0.456 | 0.843 | 0.655 | 3.82e-22 | 1.29e-17 |
| Malignant epithelial | C2 WNT ECs | PAK1 | 0.456 | 0.39 | 0.244 | 5.13e-13 | 1.73e-08 |
| Malignant epithelial | C2 WNT ECs | GTF2H1 | 0.456 | 0.225 | 0.138 | 8.74e-09 | 0.000294 |
| Malignant epithelial | C2 WNT ECs | PEX16 | 0.454 | 0.457 | 0.283 | 6.31e-17 | 2.13e-12 |
| Malignant epithelial | C2 WNT ECs | SLC12A2 | 0.454 | 0.46 | 0.329 | 1.71e-10 | 5.76e-06 |
| Malignant epithelial | C2 WNT ECs | ADCK2 | 0.454 | 0.185 | 0.113 | 1.77e-07 | 0.00598 |
| Malignant epithelial | C2 WNT ECs | MOGS | 0.453 | 0.257 | 0.149 | 3e-11 | 1.01e-06 |
| Malignant epithelial | C2 WNT ECs | ATF6 | 0.453 | 0.207 | 0.129 | 1.4e-07 | 0.00472 |
| Malignant epithelial | C2 WNT ECs | PPP1R9B | 0.453 | 0.129 | 0.072 | 4.83e-07 | 0.0163 |
| Malignant epithelial | C2 WNT ECs | PSAP | 0.452 | 0.854 | 0.711 | 4.06e-20 | 1.37e-15 |
| Malignant epithelial | C2 WNT ECs | RP9 | 0.452 | 0.313 | 0.197 | 1.73e-10 | 5.82e-06 |
| Malignant epithelial | C2 WNT ECs | MBD4 | 0.452 | 0.298 | 0.19 | 1.22e-09 | 4.11e-05 |
| Malignant epithelial | C2 WNT ECs | DCAF16 | 0.452 | 0.158 | 0.084 | 1.43e-09 | 4.81e-05 |
| Malignant epithelial | C2 WNT ECs | RARG | 0.452 | 0.222 | 0.139 | 6.32e-08 | 0.00213 |
| Malignant epithelial | C2 WNT ECs | UFC1 | 0.451 | 0.819 | 0.648 | 1.49e-21 | 5.03e-17 |
| Malignant epithelial | C2 WNT ECs | PDDC1 | 0.451 | 0.195 | 0.109 | 2.92e-10 | 9.85e-06 |
| Malignant epithelial | C2 WNT ECs | MLXIP | 0.45 | 0.461 | 0.298 | 1.22e-14 | 4.11e-10 |
| Malignant epithelial | C2 WNT ECs | UGP2 | 0.45 | 0.518 | 0.355 | 8.49e-14 | 2.86e-09 |
| Malignant epithelial | C2 WNT ECs | CUL1 | 0.45 | 0.242 | 0.136 | 2.85e-12 | 9.6e-08 |
| Malignant epithelial | C2 WNT ECs | SRRT | 0.45 | 0.29 | 0.175 | 1.47e-11 | 4.95e-07 |
| Malignant epithelial | C2 WNT ECs | CDK5RAP1 | 0.45 | 0.226 | 0.135 | 9.76e-10 | 3.29e-05 |
| Malignant epithelial | C2 WNT ECs | ERV3-1 | 0.45 | 0.286 | 0.183 | 1.11e-09 | 3.72e-05 |
| Malignant epithelial | C2 WNT ECs | CSTF1 | 0.45 | 0.206 | 0.133 | 1.27e-06 | 0.0427 |
| Malignant epithelial | C2 WNT ECs | EIF4G2 | 0.449 | 0.871 | 0.71 | 1.45e-21 | 4.87e-17 |
| Malignant epithelial | C2 WNT ECs | PAICS | 0.449 | 0.432 | 0.3 | 8.78e-11 | 2.96e-06 |
| Malignant epithelial | C2 WNT ECs | MZT1 | 0.448 | 0.471 | 0.269 | 1.53e-21 | 5.15e-17 |
| Malignant epithelial | C2 WNT ECs | NCKAP1 | 0.448 | 0.432 | 0.276 | 7.09e-14 | 2.39e-09 |
| Malignant epithelial | C2 WNT ECs | IFT57 | 0.448 | 0.41 | 0.274 | 4.36e-11 | 1.47e-06 |
| Malignant epithelial | C2 WNT ECs | COPS3 | 0.448 | 0.238 | 0.148 | 1.26e-08 | 0.000424 |
| Malignant epithelial | C2 WNT ECs | GDPD3 | 0.448 | 0.147 | 0.082 | 6.56e-08 | 0.00221 |
| Malignant epithelial | C2 WNT ECs | ADRM1 | 0.447 | 0.821 | 0.638 | 7.43e-22 | 2.5e-17 |
| Malignant epithelial | C2 WNT ECs | FUS | 0.446 | 0.949 | 0.844 | 8.240000000000001e-21 | 2.78e-16 |
| Malignant epithelial | C2 WNT ECs | SPTBN1 | 0.446 | 0.589 | 0.429 | 1.3e-13 | 4.38e-09 |
| Malignant epithelial | C2 WNT ECs | HOXB3 | 0.446 | 0.222 | 0.124 | 1.38e-11 | 4.64e-07 |
| Malignant epithelial | C2 WNT ECs | ATL3 | 0.446 | 0.296 | 0.186 | 2.39e-10 | 8.04e-06 |
| Malignant epithelial | C2 WNT ECs | PSMA2 | 0.445 | 0.89 | 0.757 | 2.16e-25 | 7.270000000000001e-21 |
| Malignant epithelial | C2 WNT ECs | APLP2 | 0.445 | 0.85 | 0.677 | 4.740000000000001e-21 | 1.6e-16 |
| Malignant epithelial | C2 WNT ECs | PPP2R2A | 0.445 | 0.394 | 0.253 | 4.62e-13 | 1.56e-08 |
| Malignant epithelial | C2 WNT ECs | UBL7 | 0.445 | 0.345 | 0.212 | 4.43e-12 | 1.49e-07 |
| Malignant epithelial | C2 WNT ECs | CCT4 | 0.444 | 0.811 | 0.636 | 5.52e-18 | 1.86e-13 |
| Malignant epithelial | C2 WNT ECs | CAPN5 | 0.444 | 0.23 | 0.11 | 6.4e-17 | 2.16e-12 |
| Malignant epithelial | C2 WNT ECs | PYURF | 0.444 | 0.594 | 0.426 | 3.23e-14 | 1.09e-09 |
| Malignant epithelial | C2 WNT ECs | BRE | 0.444 | 0.268 | 0.164 | 7.91e-10 | 2.67e-05 |
| Malignant epithelial | C2 WNT ECs | ZFYVE16 | 0.444 | 0.191 | 0.112 | 8.07e-09 | 0.000272 |
| Malignant epithelial | C2 WNT ECs | JAGN1 | 0.443 | 0.475 | 0.296 | 1.95e-17 | 6.58e-13 |
| Malignant epithelial | C2 WNT ECs | CLCN7 | 0.443 | 0.23 | 0.123 | 7.38e-13 | 2.49e-08 |
| Malignant epithelial | C2 WNT ECs | PPIL2 | 0.443 | 0.221 | 0.133 | 5.46e-09 | 0.000184 |
| Malignant epithelial | C2 WNT ECs | ERAP1 | 0.443 | 0.141 | 0.078 | 7.96e-08 | 0.00268 |
| Malignant epithelial | C2 WNT ECs | CD82 | 0.442 | 0.521 | 0.351 | 1.3e-12 | 4.37e-08 |
| Malignant epithelial | C2 WNT ECs | ERI3 | 0.442 | 0.31 | 0.188 | 1.58e-11 | 5.34e-07 |
| Malignant epithelial | C2 WNT ECs | TNPO3 | 0.442 | 0.314 | 0.196 | 7.21e-11 | 2.43e-06 |
| Malignant epithelial | C2 WNT ECs | ACADS | 0.442 | 0.25 | 0.151 | 3.77e-10 | 1.27e-05 |
| Malignant epithelial | C2 WNT ECs | ABCF3 | 0.442 | 0.162 | 0.098 | 4.12e-07 | 0.0139 |

| compartment | subcluster | gene | avg_log2FC | pct.1 | pct.2 | p_val | p_val_adj |
|---|---|---|---|---|---|---|---|
| Malignant epithelial | C2 WNT ECs | RAB11A | 0.441 | 0.867 | 0.712 | 5.84e-19 | 1.97e-14 |
| Malignant epithelial | C2 WNT ECs | ILF2 | 0.441 | 0.673 | 0.47 | 2.8e-18 | 9.42e-14 |
| Malignant epithelial | C2 WNT ECs | PKP3 | 0.441 | 0.763 | 0.615 | 3.68e-16 | 1.24e-11 |
| Malignant epithelial | C2 WNT ECs | MAT2A | 0.441 | 0.441 | 0.297 | 2.97e-12 | 1e-07 |
| Malignant epithelial | C2 WNT ECs | RNF220 | 0.441 | 0.249 | 0.144 | 2.08e-11 | 7e-07 |
| Malignant epithelial | C2 WNT ECs | ATP6V1D | 0.441 | 0.516 | 0.381 | 1e-09 | 3.37e-05 |
| Malignant epithelial | C2 WNT ECs | ARMC1 | 0.441 | 0.28 | 0.187 | 4.69e-08 | 0.00158 |
| Malignant epithelial | C2 WNT ECs | SEC61G | 0.44 | 0.98 | 0.949 | 1.94e-27 | 6.539999999999999e-23 |
| Malignant epithelial | C2 WNT ECs | ANXA4 | 0.44 | 0.781 | 0.552 | 5.48e-26 | 1.85e-21 |
| Malignant epithelial | C2 WNT ECs | BCLAF1 | 0.44 | 0.636 | 0.423 | 1.77e-19 | 5.95e-15 |
| Malignant epithelial | C2 WNT ECs | SH3BGRL3 | 0.439 | 0.935 | 0.878 | 2.49e-21 | 8.38e-17 |
| Malignant epithelial | C2 WNT ECs | ZC3H15 | 0.439 | 0.781 | 0.594 | 7.08e-18 | 2.38e-13 |
| Malignant epithelial | C2 WNT ECs | DPM1 | 0.439 | 0.719 | 0.545 | 8.06e-15 | 2.72e-10 |
| Malignant epithelial | C2 WNT ECs | IRF3 | 0.439 | 0.53 | 0.354 | 1.18e-14 | 3.99e-10 |
| Malignant epithelial | C2 WNT ECs | ELMO3 | 0.439 | 0.341 | 0.205 | 2.3e-13 | 7.76e-09 |
| Malignant epithelial | C2 WNT ECs | ACOT9 | 0.439 | 0.307 | 0.198 | 3.52e-10 | 1.19e-05 |
| Malignant epithelial | C2 WNT ECs | DPP8 | 0.439 | 0.203 | 0.123 | 1.92e-08 | 0.000645 |
| Malignant epithelial | C2 WNT ECs | CSNK2A2 | 0.439 | 0.245 | 0.156 | 3.15e-08 | 0.00106 |
| Malignant epithelial | C2 WNT ECs | PIP5K1A | 0.439 | 0.152 | 0.09 | 7.37e-07 | 0.0248 |
| Malignant epithelial | C2 WNT ECs | CLASP1 | 0.438 | 0.119 | 0.063 | 1.57e-07 | 0.0053 |
| Malignant epithelial | C2 WNT ECs | RAB5C | 0.437 | 0.709 | 0.527 | 3.9e-17 | 1.32e-12 |
| Malignant epithelial | C2 WNT ECs | MED25 | 0.437 | 0.352 | 0.205 | 3.09e-15 | 1.04e-10 |
| Malignant epithelial | C2 WNT ECs | ARHGAP8 | 0.437 | 0.227 | 0.133 | 3.8e-10 | 1.28e-05 |
| Malignant epithelial | C2 WNT ECs | WASH1 | 0.436 | 0.291 | 0.186 | 1.9e-09 | 6.41e-05 |
| Malignant epithelial | C2 WNT ECs | RIT1 | 0.436 | 0.164 | 0.091 | 1.09e-08 | 0.000367 |
| Malignant epithelial | C2 WNT ECs | CUL2 | 0.436 | 0.168 | 0.097 | 5.49e-08 | 0.00185 |
| Malignant epithelial | C2 WNT ECs | FADS3 | 0.436 | 0.254 | 0.164 | 6.31e-08 | 0.00213 |
| Malignant epithelial | C2 WNT ECs | STYXL1 | 0.435 | 0.337 | 0.203 | 1.01e-12 | 3.42e-08 |
| Malignant epithelial | C2 WNT ECs | RASSF7 | 0.434 | 0.602 | 0.379 | 4.680000000000001e-21 | 1.58e-16 |
| Malignant epithelial | C2 WNT ECs | REXO2 | 0.434 | 0.664 | 0.482 | 3.38e-18 | 1.14e-13 |
| Malignant epithelial | C2 WNT ECs | MTRF1L | 0.434 | 0.221 | 0.126 | 4e-10 | 1.35e-05 |
| Malignant epithelial | C2 WNT ECs | PSD4 | 0.434 | 0.257 | 0.159 | 1.79e-09 | 6.05e-05 |
| Malignant epithelial | C2 WNT ECs | TRIP11 | 0.434 | 0.214 | 0.129 | 1.57e-08 | 0.000529 |
| Malignant epithelial | C2 WNT ECs | ATP1B3 | 0.433 | 0.704 | 0.526 | 1.73e-18 | 5.83e-14 |
| Malignant epithelial | C2 WNT ECs | EPS8L2 | 0.433 | 0.621 | 0.433 | 1.17e-15 | 3.95e-11 |
| Malignant epithelial | C2 WNT ECs | CNOT7 | 0.433 | 0.365 | 0.239 | 6.24e-11 | 2.1e-06 |
| Malignant epithelial | C2 WNT ECs | CENPT | 0.433 | 0.203 | 0.121 | 2.64e-08 | 0.000891 |
| Malignant epithelial | C2 WNT ECs | TRPM4 | 0.433 | 0.221 | 0.138 | 5.61e-08 | 0.00189 |
| Malignant epithelial | C2 WNT ECs | CPSF3L | 0.432 | 0.383 | 0.241 | 1.99e-13 | 6.71e-09 |
| Malignant epithelial | C2 WNT ECs | PDCD7 | 0.432 | 0.313 | 0.193 | 2.39e-11 | 8.04e-07 |
| Malignant epithelial | C2 WNT ECs | WDR82 | 0.432 | 0.3 | 0.183 | 2.83e-11 | 9.55e-07 |
| Malignant epithelial | C2 WNT ECs | ADSL | 0.432 | 0.346 | 0.223 | 1.38e-10 | 4.66e-06 |
| Malignant epithelial | C2 WNT ECs | ADRBK1 | 0.432 | 0.29 | 0.188 | 6e-09 | 0.000202 |
| Malignant epithelial | C2 WNT ECs | CTPS2 | 0.432 | 0.124 | 0.069 | 4.68e-07 | 0.0158 |
| Malignant epithelial | C2 WNT ECs | MAP3K7 | 0.43 | 0.199 | 0.108 | 2.98e-11 | 1e-06 |
| Malignant epithelial | C2 WNT ECs | RPL7L1 | 0.43 | 0.52 | 0.377 | 5.31e-11 | 1.79e-06 |
| Malignant epithelial | C2 WNT ECs | C14orf1 | 0.43 | 0.535 | 0.389 | 4.87e-10 | 1.64e-05 |
| Malignant epithelial | C2 WNT ECs | TFRC | 0.43 | 0.304 | 0.196 | 5.92e-10 | 2e-05 |
| Malignant epithelial | C2 WNT ECs | ARFGAP3 | 0.43 | 0.287 | 0.186 | 5.12e-09 | 0.000172 |
| Malignant epithelial | C2 WNT ECs | DAGLB | 0.43 | 0.149 | 0.081 | 1.18e-08 | 0.000397 |
| Malignant epithelial | C2 WNT ECs | ZNF655 | 0.43 | 0.258 | 0.171 | 3.14e-07 | 0.0106 |
| Malignant epithelial | C2 WNT ECs | TM2D1 | 0.429 | 0.421 | 0.243 | 1.47e-18 | 4.96e-14 |
| Malignant epithelial | C2 WNT ECs | IDS | 0.429 | 0.419 | 0.29 | 7.66e-11 | 2.58e-06 |
| Malignant epithelial | C2 WNT ECs | RAB11FIP4 | 0.429 | 0.141 | 0.075 | 1.46e-08 | 0.000492 |
| Malignant epithelial | C2 WNT ECs | WDSUB1 | 0.429 | 0.119 | 0.067 | 1.33e-06 | 0.0447 |
| Malignant epithelial | C2 WNT ECs | NDUFV2 | 0.428 | 0.754 | 0.563 | 1.71e-20 | 5.76e-16 |
| Malignant epithelial | C2 WNT ECs | PAM | 0.428 | 0.273 | 0.153 | 6.46e-14 | 2.18e-09 |
| Malignant epithelial | C2 WNT ECs | FAM3A | 0.428 | 0.422 | 0.269 | 6.87e-14 | 2.31e-09 |
| Malignant epithelial | C2 WNT ECs | NCOR1 | 0.428 | 0.479 | 0.32 | 4.72e-13 | 1.59e-08 |
| Malignant epithelial | C2 WNT ECs | BCKDK | 0.428 | 0.322 | 0.204 | 1.71e-10 | 5.75e-06 |
| Malignant epithelial | C2 WNT ECs | PPT1 | 0.428 | 0.311 | 0.207 | 1.45e-08 | 0.000487 |
| Malignant epithelial | C2 WNT ECs | ZNF593 | 0.427 | 0.655 | 0.467 | 5.76e-17 | 1.94e-12 |
| Malignant epithelial | C2 WNT ECs | RAB17 | 0.427 | 0.3 | 0.187 | 3.52e-10 | 1.18e-05 |
| Malignant epithelial | C2 WNT ECs | UBL4A | 0.427 | 0.268 | 0.174 | 1.82e-08 | 0.000614 |

| compartment | subcluster | gene | avg_log2FC | pct.1 | pct.2 | p_val | p_val_adj |
|---|---|---|---|---|---|---|---|
| Malignant epithelial | C2 WNT ECs | LENG8 | 0.427 | 0.384 | 0.281 | 3.42e-07 | 0.0115 |
| Malignant epithelial | C2 WNT ECs | SFT2D3 | 0.427 | 0.149 | 0.086 | 3.92e-07 | 0.0132 |
| Malignant epithelial | C2 WNT ECs | ASXL2 | 0.427 | 0.173 | 0.106 | 7.47e-07 | 0.0252 |
| Malignant epithelial | C2 WNT ECs | YBX1 | 0.426 | 0.963 | 0.904 | 1.23e-31 | 4.14e-27 |
| Malignant epithelial | C2 WNT ECs | AP2S1 | 0.426 | 0.858 | 0.718 | 8.270000000000001e-21 | 2.79e-16 |
| Malignant epithelial | C2 WNT ECs | C21orf59 | 0.426 | 0.472 | 0.297 | 4.15e-15 | 1.4e-10 |
| Malignant epithelial | C2 WNT ECs | CIR1 | 0.426 | 0.556 | 0.378 | 5.46e-15 | 1.84e-10 |
| Malignant epithelial | C2 WNT ECs | PPP1R21 | 0.426 | 0.154 | 0.081 | 1.53e-09 | 5.16e-05 |
| Malignant epithelial | C2 WNT ECs | PPWD1 | 0.426 | 0.147 | 0.08 | 9.82e-09 | 0.000331 |
| Malignant epithelial | C2 WNT ECs | C1orf198 | 0.426 | 0.18 | 0.112 | 5.84e-07 | 0.0197 |
| Malignant epithelial | C2 WNT ECs | WDR55 | 0.426 | 0.218 | 0.141 | 8.07e-07 | 0.0272 |
| Malignant epithelial | C2 WNT ECs | YWHAB | 0.425 | 0.935 | 0.809 | 2.69e-20 | 9.05e-16 |
| Malignant epithelial | C2 WNT ECs | DCTN2 | 0.425 | 0.594 | 0.397 | 4.66e-18 | 1.57e-13 |
| Malignant epithelial | C2 WNT ECs | IGF2BP2 | 0.425 | 0.594 | 0.412 | 1.62e-14 | 5.45e-10 |
| Malignant epithelial | C2 WNT ECs | HNMT | 0.425 | 0.536 | 0.383 | 2.15e-12 | 7.25e-08 |
| Malignant epithelial | C2 WNT ECs | HSD17B11 | 0.425 | 0.635 | 0.479 | 3.18e-12 | 1.07e-07 |
| Malignant epithelial | C2 WNT ECs | CBX1 | 0.425 | 0.376 | 0.245 | 2.71e-11 | 9.14e-07 |
| Malignant epithelial | C2 WNT ECs | ADPRHL2 | 0.425 | 0.221 | 0.127 | 3.52e-10 | 1.19e-05 |
| Malignant epithelial | C2 WNT ECs | SUCO | 0.425 | 0.235 | 0.14 | 6.97e-10 | 2.35e-05 |
| Malignant epithelial | C2 WNT ECs | AAR2 | 0.425 | 0.188 | 0.115 | 1.54e-07 | 0.0052 |
| Malignant epithelial | C2 WNT ECs | WBSCR22 | 0.424 | 0.708 | 0.513 | 2.36e-18 | 7.96e-14 |
| Malignant epithelial | C2 WNT ECs | ORC4 | 0.424 | 0.284 | 0.165 | 1.47e-12 | 4.95e-08 |
| Malignant epithelial | C2 WNT ECs | BCL2L12 | 0.424 | 0.401 | 0.261 | 2.77e-12 | 9.32e-08 |
| Malignant epithelial | C2 WNT ECs | PIAS1 | 0.424 | 0.226 | 0.145 | 2.16e-07 | 0.00727 |
| Malignant epithelial | C2 WNT ECs | ZNF692 | 0.424 | 0.16 | 0.095 | 3.71e-07 | 0.0125 |
| Malignant epithelial | C2 WNT ECs | EHF | 0.424 | 0.536 | 0.429 | 4.19e-07 | 0.0141 |
| Malignant epithelial | C2 WNT ECs | TUT1 | 0.424 | 0.108 | 0.058 | 9.51e-07 | 0.032 |
| Malignant epithelial | C2 WNT ECs | SH3BGRL | 0.423 | 0.516 | 0.377 | 3.72e-10 | 1.25e-05 |
| Malignant epithelial | C2 WNT ECs | FBXO7 | 0.422 | 0.513 | 0.332 | 3.91e-16 | 1.32e-11 |
| Malignant epithelial | C2 WNT ECs | CELF1 | 0.422 | 0.49 | 0.314 | 3.01e-15 | 1.02e-10 |
| Malignant epithelial | C2 WNT ECs | FKBP9 | 0.422 | 0.495 | 0.329 | 5.93e-14 | 2e-09 |
| Malignant epithelial | C2 WNT ECs | SRPRA | 0.422 | 0.329 | 0.203 | 1.5e-11 | 5.04e-07 |
| Malignant epithelial | C2 WNT ECs | GART | 0.422 | 0.275 | 0.17 | 5.99e-10 | 2.02e-05 |
| Malignant epithelial | C2 WNT ECs | AACS | 0.422 | 0.157 | 0.086 | 1.17e-08 | 0.000393 |
| Malignant epithelial | C2 WNT ECs | CTCF | 0.422 | 0.277 | 0.184 | 3.58e-08 | 0.00121 |
| Malignant epithelial | C2 WNT ECs | PHF1 | 0.422 | 0.271 | 0.177 | 4.25e-08 | 0.00143 |
| Malignant epithelial | C2 WNT ECs | TLDC1 | 0.422 | 0.196 | 0.124 | 4.21e-07 | 0.0142 |
| Malignant epithelial | C2 WNT ECs | SUPT5H | 0.421 | 0.399 | 0.272 | 2.02e-09 | 6.81e-05 |
| Malignant epithelial | C2 WNT ECs | DHRS1 | 0.421 | 0.292 | 0.195 | 4.93e-08 | 0.00166 |
| Malignant epithelial | C2 WNT ECs | MYCBP | 0.421 | 0.204 | 0.126 | 1.08e-07 | 0.00364 |
| Malignant epithelial | C2 WNT ECs | FOSL2 | 0.421 | 0.414 | 0.301 | 1.11e-07 | 0.00375 |
| Malignant epithelial | C2 WNT ECs | GCN1 | 0.421 | 0.138 | 0.081 | 1.44e-06 | 0.0484 |
| Malignant epithelial | C2 WNT ECs | TMEM87B | 0.42 | 0.253 | 0.137 | 4.87e-14 | 1.64e-09 |
| Malignant epithelial | C2 WNT ECs | CERS2 | 0.42 | 0.484 | 0.338 | 1.38e-11 | 4.65e-07 |
| Malignant epithelial | C2 WNT ECs | CDC40 | 0.42 | 0.295 | 0.182 | 1.69e-10 | 5.69e-06 |
| Malignant epithelial | C2 WNT ECs | OSBP | 0.42 | 0.203 | 0.12 | 7.45e-09 | 0.000251 |
| Malignant epithelial | C2 WNT ECs | PHKB | 0.42 | 0.233 | 0.149 | 9.65e-08 | 0.00325 |
| Malignant epithelial | C2 WNT ECs | CRY2 | 0.42 | 0.114 | 0.062 | 7.97e-07 | 0.0268 |
| Malignant epithelial | C2 WNT ECs | AP2A1 | 0.419 | 0.284 | 0.18 | 9.19e-10 | 3.1e-05 |
| Malignant epithelial | C2 WNT ECs | CRTC2 | 0.419 | 0.146 | 0.077 | 3.73e-09 | 0.000126 |
| Malignant epithelial | C2 WNT ECs | PUSL1 | 0.419 | 0.152 | 0.083 | 1.36e-08 | 0.000459 |
| Malignant epithelial | C2 WNT ECs | RBM25 | 0.418 | 0.729 | 0.534 | 6.94e-17 | 2.34e-12 |
| Malignant epithelial | C2 WNT ECs | ARL1 | 0.418 | 0.521 | 0.343 | 3.54e-14 | 1.19e-09 |
| Malignant epithelial | C2 WNT ECs | ARL2BP | 0.418 | 0.356 | 0.236 | 5.89e-10 | 1.99e-05 |
| Malignant epithelial | C2 WNT ECs | MEPCE | 0.418 | 0.217 | 0.138 | 2.58e-07 | 0.0087 |
| Malignant epithelial | C2 WNT ECs | PPP1CA | 0.417 | 0.851 | 0.687 | 1.48e-20 | 4.98e-16 |
| Malignant epithelial | C2 WNT ECs | ZDHHC6 | 0.417 | 0.383 | 0.23 | 1.49e-14 | 5.02e-10 |
| Malignant epithelial | C2 WNT ECs | FAM210B | 0.417 | 0.447 | 0.286 | 1.28e-13 | 4.3e-09 |
| Malignant epithelial | C2 WNT ECs | NOL12 | 0.417 | 0.211 | 0.118 | 1.73e-10 | 5.81e-06 |
| Malignant epithelial | C2 WNT ECs | DAAM1 | 0.417 | 0.321 | 0.214 | 1.05e-08 | 0.000352 |
| Malignant epithelial | C2 WNT ECs | GOLT1A | 0.417 | 0.198 | 0.124 | 3.93e-07 | 0.0132 |
| Malignant epithelial | C2 WNT ECs | ANKRD26 | 0.417 | 0.103 | 0.055 | 1.48e-06 | 0.0499 |
| Malignant epithelial | C2 WNT ECs | ARPC5 | 0.416 | 0.813 | 0.618 | 7.58e-18 | 2.55e-13 |
| Malignant epithelial | C2 WNT ECs | PPHLN1 | 0.416 | 0.506 | 0.343 | 2.57e-12 | 8.67e-08 |

| compartment | subcluster | gene | avg_log2FC | pct.1 | pct.2 | p_val | p_val_adj |
|---|---|---|---|---|---|---|---|
| Malignant epithelial | C2 WNT ECs | ZNF444 | 0.416 | 0.242 | 0.142 | 2.11e-10 | 7.12e-06 |
| Malignant epithelial | C2 WNT ECs | CNBD2 | 0.416 | 0.245 | 0.159 | 1.57e-07 | 0.00529 |
| Malignant epithelial | C2 WNT ECs | STK26 | 0.416 | 0.171 | 0.105 | 1.01e-06 | 0.0341 |
| Malignant epithelial | C2 WNT ECs | HNRNPLL | 0.415 | 0.245 | 0.148 | 1.27e-09 | 4.27e-05 |
| Malignant epithelial | C2 WNT ECs | MT-ND2 | 0.414 | 0.997 | 0.994 | 4.11e-19 | 1.38e-14 |
| Malignant epithelial | C2 WNT ECs | IDH3A | 0.414 | 0.277 | 0.173 | 5.47e-10 | 1.84e-05 |
| Malignant epithelial | C2 WNT ECs | CRKL | 0.414 | 0.195 | 0.11 | 1.17e-09 | 3.95e-05 |
| Malignant epithelial | C2 WNT ECs | PNO1 | 0.414 | 0.322 | 0.209 | 1.86e-09 | 6.27e-05 |
| Malignant epithelial | C2 WNT ECs | DNPEP | 0.413 | 0.637 | 0.439 | 2.25e-16 | 7.58e-12 |
| Malignant epithelial | C2 WNT ECs | EIF5 | 0.413 | 0.853 | 0.683 | 5.2e-15 | 1.75e-10 |
| Malignant epithelial | C2 WNT ECs | SQSTM1 | 0.413 | 0.901 | 0.784 | 6.02e-13 | 2.03e-08 |
| Malignant epithelial | C2 WNT ECs | VAMP7 | 0.413 | 0.173 | 0.103 | 2.12e-07 | 0.00714 |
| Malignant epithelial | C2 WNT ECs | IBTK | 0.413 | 0.229 | 0.151 | 7.04e-07 | 0.0237 |
| Malignant epithelial | C2 WNT ECs | DNAJA1 | 0.413 | 0.899 | 0.805 | 7.89e-07 | 0.0266 |
| Malignant epithelial | C2 WNT ECs | HDAC5 | 0.413 | 0.185 | 0.118 | 1.41e-06 | 0.0475 |
| Malignant epithelial | C2 WNT ECs | RNF5 | 0.412 | 0.589 | 0.394 | 2.98e-16 | 1e-11 |
| Malignant epithelial | C2 WNT ECs | ESD | 0.412 | 0.721 | 0.569 | 4.56e-16 | 1.54e-11 |
| Malignant epithelial | C2 WNT ECs | RHBDD2 | 0.412 | 0.548 | 0.387 | 8.02e-13 | 2.7e-08 |
| Malignant epithelial | C2 WNT ECs | MAPKAPK5 | 0.412 | 0.323 | 0.203 | 3.84e-11 | 1.29e-06 |
| Malignant epithelial | C2 WNT ECs | TMEM184A | 0.412 | 0.276 | 0.174 | 2.73e-09 | 9.19e-05 |
| Malignant epithelial | C2 WNT ECs | TMEM263 | 0.412 | 0.16 | 0.094 | 2.99e-07 | 0.0101 |
| Malignant epithelial | C2 WNT ECs | LONP2 | 0.411 | 0.352 | 0.219 | 4.46e-12 | 1.5e-07 |
| Malignant epithelial | C2 WNT ECs | AKIRIN2 | 0.411 | 0.43 | 0.285 | 6.91e-11 | 2.33e-06 |
| Malignant epithelial | C2 WNT ECs | ACOT13 | 0.411 | 0.315 | 0.195 | 1.84e-10 | 6.19e-06 |
| Malignant epithelial | C2 WNT ECs | NPRL3 | 0.411 | 0.244 | 0.149 | 5.89e-09 | 0.000198 |
| Malignant epithelial | C2 WNT ECs | MCMBP | 0.411 | 0.26 | 0.167 | 1.66e-08 | 0.00056 |
| Malignant epithelial | C2 WNT ECs | SAMD8 | 0.41 | 0.154 | 0.085 | 2.02e-08 | 0.00068 |
| Malignant epithelial | C2 WNT ECs | CHRAC1 | 0.409 | 0.572 | 0.389 | 1.37e-15 | 4.61e-11 |
| Malignant epithelial | C2 WNT ECs | SLC11A2 | 0.409 | 0.169 | 0.092 | 1.5e-09 | 5.05e-05 |
| Malignant epithelial | C2 WNT ECs | ALG1 | 0.409 | 0.176 | 0.102 | 3.51e-08 | 0.00118 |
| Malignant epithelial | C2 WNT ECs | CAPN10 | 0.409 | 0.183 | 0.111 | 1.95e-07 | 0.00657 |
| Malignant epithelial | C2 WNT ECs | CORO1B | 0.408 | 0.697 | 0.52 | 4.66e-18 | 1.57e-13 |
| Malignant epithelial | C2 WNT ECs | EPB41L1 | 0.408 | 0.319 | 0.199 | 1.1e-11 | 3.71e-07 |
| Malignant epithelial | C2 WNT ECs | SLAIN2 | 0.408 | 0.176 | 0.096 | 1.35e-09 | 4.56e-05 |
| Malignant epithelial | C2 WNT ECs | SEPT11 | 0.408 | 0.267 | 0.174 | 2.49e-08 | 0.000839 |
| Malignant epithelial | C2 WNT ECs | CDC37 | 0.407 | 0.763 | 0.611 | 1.56e-16 | 5.26e-12 |
| Malignant epithelial | C2 WNT ECs | BLOC1S6 | 0.407 | 0.254 | 0.149 | 3.74e-11 | 1.26e-06 |
| Malignant epithelial | C2 WNT ECs | ZBTB11 | 0.407 | 0.161 | 0.086 | 1.04e-09 | 3.49e-05 |
| Malignant epithelial | C2 WNT ECs | MARK2 | 0.407 | 0.292 | 0.188 | 3.56e-09 | 0.00012 |
| Malignant epithelial | C2 WNT ECs | ARIH2 | 0.407 | 0.314 | 0.215 | 7.94e-08 | 0.00267 |
| Malignant epithelial | C2 WNT ECs | SMARCD1 | 0.406 | 0.284 | 0.175 | 8.01e-11 | 2.7e-06 |
| Malignant epithelial | C2 WNT ECs | ZNF148 | 0.406 | 0.28 | 0.183 | 1.58e-08 | 0.000534 |
| Malignant epithelial | C2 WNT ECs | NAPA | 0.405 | 0.547 | 0.349 | 8.13e-17 | 2.74e-12 |
| Malignant epithelial | C2 WNT ECs | PSMA5 | 0.404 | 0.685 | 0.477 | 3.91e-17 | 1.32e-12 |
| Malignant epithelial | C2 WNT ECs | SAMM50 | 0.404 | 0.357 | 0.224 | 4.04e-12 | 1.36e-07 |
| Malignant epithelial | C2 WNT ECs | JOSD1 | 0.404 | 0.373 | 0.244 | 1.11e-11 | 3.73e-07 |
| Malignant epithelial | C2 WNT ECs | ACOX1 | 0.404 | 0.246 | 0.143 | 3.77e-11 | 1.27e-06 |
| Malignant epithelial | C2 WNT ECs | FAM134A | 0.404 | 0.421 | 0.295 | 5.34e-10 | 1.8e-05 |
| Malignant epithelial | C2 WNT ECs | DHX16 | 0.404 | 0.177 | 0.103 | 4.11e-08 | 0.00139 |
| Malignant epithelial | C2 WNT ECs | EEF1G | 0.403 | 0.547 | 0.381 | 7.38e-14 | 2.49e-09 |
| Malignant epithelial | C2 WNT ECs | TRIOBP | 0.403 | 0.384 | 0.248 | 1.54e-12 | 5.18e-08 |
| Malignant epithelial | C2 WNT ECs | OAZ2 | 0.403 | 0.413 | 0.27 | 5.1e-12 | 1.72e-07 |
| Malignant epithelial | C2 WNT ECs | SMPD2 | 0.403 | 0.217 | 0.12 | 7.94e-11 | 2.68e-06 |
| Malignant epithelial | C2 WNT ECs | OTUD5 | 0.403 | 0.268 | 0.162 | 1.4e-10 | 4.73e-06 |
| Malignant epithelial | C2 WNT ECs | PCYT1A | 0.403 | 0.191 | 0.118 | 4.26e-07 | 0.0144 |
| Malignant epithelial | C2 WNT ECs | NDUFC2 | 0.402 | 0.899 | 0.758 | 9.140000000000001e-22 | 3.08e-17 |
| Malignant epithelial | C2 WNT ECs | CDC42 | 0.402 | 0.877 | 0.686 | 1.95e-20 | 6.56e-16 |
| Malignant epithelial | C2 WNT ECs | MPST | 0.402 | 0.811 | 0.653 | 2.14e-19 | 7.22e-15 |
| Malignant epithelial | C2 WNT ECs | U2AF2 | 0.402 | 0.434 | 0.281 | 9.65e-13 | 3.25e-08 |
| Malignant epithelial | C2 WNT ECs | CD46 | 0.402 | 0.702 | 0.555 | 4.06e-10 | 1.37e-05 |
| Malignant epithelial | C2 WNT ECs | MANSC1 | 0.402 | 0.279 | 0.182 | 2.29e-08 | 0.00077 |
| Malignant epithelial | C2 WNT ECs | FBXO8 | 0.402 | 0.119 | 0.064 | 2.77e-07 | 0.00934 |
| Malignant epithelial | C2 WNT ECs | TM2D3 | 0.401 | 0.357 | 0.212 | 3.74e-14 | 1.26e-09 |
| Malignant epithelial | C2 WNT ECs | TMEM50A | 0.4 | 0.8 | 0.598 | 2.83e-20 | 9.54e-16 |

| compartment | subcluster | gene | avg_log2FC | pct.1 | pct.2 | p_val | p_val_adj |
|---|---|---|---|---|---|---|---|
| Malignant epithelial | C2 WNT ECs | ATP6V1C1 | 0.4 | 0.315 | 0.199 | 5.1e-11 | 1.72e-06 |
| Malignant epithelial | C2 WNT ECs | VAPB | 0.4 | 0.311 | 0.216 | 2.41e-07 | 0.00812 |
| Malignant epithelial | C2 WNT ECs | TOR1A | 0.4 | 0.245 | 0.165 | 8.95e-07 | 0.0301 |
| Malignant epithelial | C2 WNT ECs | FAM96B | 0.399 | 0.877 | 0.732 | 8.559999999999999e-23 | 2.89e-18 |
| Malignant epithelial | C2 WNT ECs | FAM192A | 0.399 | 0.482 | 0.318 | 1.28e-13 | 4.32e-09 |
| Malignant epithelial | C2 WNT ECs | DNASE2 | 0.399 | 0.349 | 0.217 | 5.28e-12 | 1.78e-07 |
| Malignant epithelial | C2 WNT ECs | ANKRD39 | 0.399 | 0.25 | 0.159 | 3.46e-08 | 0.00117 |
| Malignant epithelial | C2 WNT ECs | UBB | 0.398 | 0.985 | 0.957 | 8.39e-23 | 2.83e-18 |
| Malignant epithelial | C2 WNT ECs | PSMD11 | 0.398 | 0.549 | 0.367 | 1.7e-15 | 5.72e-11 |
| Malignant epithelial | C2 WNT ECs | OFD1 | 0.398 | 0.526 | 0.358 | 9.7e-14 | 3.27e-09 |
| Malignant epithelial | C2 WNT ECs | SEC62 | 0.398 | 0.797 | 0.656 | 1.29e-12 | 4.34e-08 |
| Malignant epithelial | C2 WNT ECs | ZFR | 0.398 | 0.306 | 0.198 | 2.04e-09 | 6.89e-05 |
| Malignant epithelial | C2 WNT ECs | UBFD1 | 0.398 | 0.198 | 0.119 | 4.04e-08 | 0.00136 |
| Malignant epithelial | C2 WNT ECs | CMTM4 | 0.398 | 0.127 | 0.069 | 2.16e-07 | 0.00728 |
| Malignant epithelial | C2 WNT ECs | PSMG2 | 0.397 | 0.639 | 0.464 | 1.09e-14 | 3.66e-10 |
| Malignant epithelial | C2 WNT ECs | C8orf76 | 0.397 | 0.225 | 0.139 | 2.19e-08 | 0.000738 |
| Malignant epithelial | C2 WNT ECs | PI4K2B | 0.397 | 0.16 | 0.091 | 4.92e-08 | 0.00166 |
| Malignant epithelial | C2 WNT ECs | POLG | 0.397 | 0.158 | 0.095 | 6.73e-07 | 0.0227 |
| Malignant epithelial | C2 WNT ECs | GDE1 | 0.396 | 0.585 | 0.396 | 5.52e-15 | 1.86e-10 |
| Malignant epithelial | C2 WNT ECs | CTSA | 0.396 | 0.769 | 0.593 | 7.3e-14 | 2.46e-09 |
| Malignant epithelial | C2 WNT ECs | USP34 | 0.396 | 0.313 | 0.194 | 7.13e-11 | 2.4e-06 |
| Malignant epithelial | C2 WNT ECs | HYAL2 | 0.396 | 0.421 | 0.297 | 8.57e-09 | 0.000289 |
| Malignant epithelial | C2 WNT ECs | TSR1 | 0.396 | 0.168 | 0.097 | 4.54e-08 | 0.00153 |
| Malignant epithelial | C2 WNT ECs | HNRNPDL | 0.395 | 0.931 | 0.804 | 2.56e-21 | 8.62e-17 |
| Malignant epithelial | C2 WNT ECs | EIF4B | 0.395 | 0.712 | 0.554 | 1.01e-14 | 3.42e-10 |
| Malignant epithelial | C2 WNT ECs | C2orf49 | 0.395 | 0.256 | 0.148 | 1.84e-11 | 6.22e-07 |
| Malignant epithelial | C2 WNT ECs | EML4 | 0.395 | 0.418 | 0.279 | 6.52e-11 | 2.2e-06 |
| Malignant epithelial | C2 WNT ECs | DRG1 | 0.395 | 0.434 | 0.295 | 3.17e-10 | 1.07e-05 |
| Malignant epithelial | C2 WNT ECs | MRPS35 | 0.395 | 0.518 | 0.378 | 1.87e-09 | 6.3e-05 |
| Malignant epithelial | C2 WNT ECs | KIAA1033 | 0.395 | 0.233 | 0.142 | 4.62e-09 | 0.000156 |
| Malignant epithelial | C2 WNT ECs | RDH14 | 0.395 | 0.248 | 0.157 | 6.08e-08 | 0.00205 |
| Malignant epithelial | C2 WNT ECs | ZNF638 | 0.395 | 0.307 | 0.206 | 7.45e-08 | 0.00251 |
| Malignant epithelial | C2 WNT ECs | YWHAE | 0.394 | 0.888 | 0.723 | 9.97e-22 | 3.36e-17 |
| Malignant epithelial | C2 WNT ECs | UTP4 | 0.394 | 0.286 | 0.176 | 3.16e-10 | 1.06e-05 |
| Malignant epithelial | C2 WNT ECs | STK17B | 0.394 | 0.329 | 0.22 | 8.9e-09 | 0.0003 |
| Malignant epithelial | C2 WNT ECs | IGBP1 | 0.393 | 0.746 | 0.544 | 4.65e-20 | 1.57e-15 |
| Malignant epithelial | C2 WNT ECs | ANAPC15 | 0.393 | 0.46 | 0.279 | 5.35e-16 | 1.8e-11 |
| Malignant epithelial | C2 WNT ECs | E2F4 | 0.393 | 0.368 | 0.244 | 1.24e-10 | 4.17e-06 |
| Malignant epithelial | C2 WNT ECs | NFKBIB | 0.393 | 0.341 | 0.218 | 2.95e-10 | 9.94e-06 |
| Malignant epithelial | C2 WNT ECs | SNX1 | 0.393 | 0.248 | 0.154 | 1.16e-08 | 0.00039 |
| Malignant epithelial | C2 WNT ECs | PHKG1 | 0.393 | 0.179 | 0.105 | 4.61e-08 | 0.00155 |
| Malignant epithelial | C2 WNT ECs | HSPB1 | 0.393 | 0.993 | 0.978 | 4.74e-07 | 0.016 |
| Malignant epithelial | C2 WNT ECs | SLC9A3R1 | 0.392 | 0.472 | 0.311 | 1.95e-13 | 6.57e-09 |
| Malignant epithelial | C2 WNT ECs | ACAD9 | 0.392 | 0.15 | 0.082 | 1.46e-08 | 0.000492 |
| Malignant epithelial | C2 WNT ECs | RPL9 | 0.391 | 0.992 | 0.986 | 5.73e-45 | 1.93e-40 |
| Malignant epithelial | C2 WNT ECs | PON2 | 0.391 | 0.727 | 0.526 | 6.26e-17 | 2.11e-12 |
| Malignant epithelial | C2 WNT ECs | ABRACL | 0.391 | 0.762 | 0.592 | 9.42e-17 | 3.18e-12 |
| Malignant epithelial | C2 WNT ECs | PRKCI | 0.391 | 0.291 | 0.173 | 1.11e-11 | 3.73e-07 |
| Malignant epithelial | C2 WNT ECs | AES | 0.391 | 0.681 | 0.543 | 1.07e-09 | 3.59e-05 |
| Malignant epithelial | C2 WNT ECs | ISG20L2 | 0.391 | 0.281 | 0.18 | 3.47e-09 | 0.000117 |
| Malignant epithelial | C2 WNT ECs | FAM60A | 0.391 | 0.54 | 0.41 | 2.9e-08 | 0.000978 |
| Malignant epithelial | C2 WNT ECs | SMARCB1 | 0.39 | 0.513 | 0.345 | 7.48e-13 | 2.52e-08 |
| Malignant epithelial | C2 WNT ECs | RIC8A | 0.39 | 0.345 | 0.216 | 2.44e-11 | 8.22e-07 |
| Malignant epithelial | C2 WNT ECs | TCEB2 | 0.389 | 0.992 | 0.979 | 9.46e-13 | 3.19e-08 |
| Malignant epithelial | C2 WNT ECs | TPD52L2 | 0.389 | 0.604 | 0.443 | 2.63e-11 | 8.88e-07 |
| Malignant epithelial | C2 WNT ECs | ACSL3 | 0.389 | 0.364 | 0.243 | 1.31e-09 | 4.42e-05 |
| Malignant epithelial | C2 WNT ECs | MAPKAPK2 | 0.389 | 0.302 | 0.195 | 2.7e-09 | 9.11e-05 |
| Malignant epithelial | C2 WNT ECs | PPP1R15A | 0.389 | 0.922 | 0.898 | 4.53e-09 | 0.000153 |
| Malignant epithelial | C2 WNT ECs | TRIM8 | 0.389 | 0.292 | 0.188 | 4.75e-09 | 0.00016 |
| Malignant epithelial | C2 WNT ECs | IKBKB | 0.389 | 0.202 | 0.121 | 2.08e-08 | 0.000702 |
| Malignant epithelial | C2 WNT ECs | DCAF8 | 0.389 | 0.233 | 0.145 | 4.39e-08 | 0.00148 |
| Malignant epithelial | C2 WNT ECs | RABAC1 | 0.388 | 0.811 | 0.64 | 7.53e-16 | 2.54e-11 |
| Malignant epithelial | C2 WNT ECs | PFKL | 0.388 | 0.476 | 0.318 | 5.65e-13 | 1.9e-08 |
| Malignant epithelial | C2 WNT ECs | SFT2D2 | 0.388 | 0.202 | 0.128 | 3.98e-07 | 0.0134 |

| compartment | subcluster | gene | avg_log2FC | pct.1 | pct.2 | p_val | p_val_adj |
|---|---|---|---|---|---|---|---|
| Malignant epithelial | C2 WNT ECs | UTP3 | 0.388 | 0.141 | 0.083 | 1.33e-06 | 0.0448 |
| Malignant epithelial | C2 WNT ECs | WDR6 | 0.387 | 0.338 | 0.217 | 1.18e-10 | 3.99e-06 |
| Malignant epithelial | C2 WNT ECs | SSH3 | 0.387 | 0.269 | 0.165 | 3.67e-10 | 1.24e-05 |
| Malignant epithelial | C2 WNT ECs | ACTG1 | 0.386 | 0.999 | 0.981 | 1.14e-22 | 3.83e-18 |
| Malignant epithelial | C2 WNT ECs | FAM133B | 0.386 | 0.779 | 0.563 | 2.54e-20 | 8.57e-16 |
| Malignant epithelial | C2 WNT ECs | GANAB | 0.386 | 0.465 | 0.305 | 3.62e-13 | 1.22e-08 |
| Malignant epithelial | C2 WNT ECs | UBN1 | 0.386 | 0.34 | 0.225 | 1.82e-09 | 6.14e-05 |
| Malignant epithelial | C2 WNT ECs | NSMCE3 | 0.386 | 0.176 | 0.106 | 4.25e-07 | 0.0143 |
| Malignant epithelial | C2 WNT ECs | PARN | 0.386 | 0.181 | 0.114 | 1.34e-06 | 0.0451 |
| Malignant epithelial | C2 WNT ECs | FXR1 | 0.385 | 0.506 | 0.325 | 1.62e-15 | 5.47e-11 |
| Malignant epithelial | C2 WNT ECs | PICALM | 0.385 | 0.359 | 0.242 | 1.57e-09 | 5.28e-05 |
| Malignant epithelial | C2 WNT ECs | SLC25A46 | 0.385 | 0.202 | 0.117 | 2.67e-09 | 9.01e-05 |
| Malignant epithelial | C2 WNT ECs | CERS5 | 0.385 | 0.175 | 0.106 | 4.32e-07 | 0.0145 |
| Malignant epithelial | C2 WNT ECs | SNAP29 | 0.384 | 0.363 | 0.23 | 4.6e-11 | 1.55e-06 |
| Malignant epithelial | C2 WNT ECs | MGAT1 | 0.384 | 0.405 | 0.27 | 1.19e-10 | 4e-06 |
| Malignant epithelial | C2 WNT ECs | SCFD1 | 0.384 | 0.346 | 0.221 | 2.09e-10 | 7.03e-06 |
| Malignant epithelial | C2 WNT ECs | TERF1 | 0.384 | 0.364 | 0.248 | 1.72e-09 | 5.81e-05 |
| Malignant epithelial | C2 WNT ECs | PIEZO1 | 0.384 | 0.257 | 0.174 | 3.01e-07 | 0.0101 |
| Malignant epithelial | C2 WNT ECs | RAP1B | 0.383 | 0.437 | 0.319 | 3.35e-08 | 0.00113 |
| Malignant epithelial | C2 WNT ECs | MORF4L2 | 0.382 | 0.794 | 0.598 | 4.94e-19 | 1.66e-14 |
| Malignant epithelial | C2 WNT ECs | UBE2J2 | 0.382 | 0.342 | 0.223 | 2.05e-09 | 6.91e-05 |
| Malignant epithelial | C2 WNT ECs | LZIC | 0.382 | 0.207 | 0.13 | 3.16e-07 | 0.0106 |
| Malignant epithelial | C2 WNT ECs | TIMM44 | 0.382 | 0.352 | 0.251 | 6.2e-07 | 0.0209 |
| Malignant epithelial | C2 WNT ECs | BIRC6 | 0.381 | 0.284 | 0.184 | 1.3e-08 | 0.000436 |
| Malignant epithelial | C2 WNT ECs | EIF1AX | 0.38 | 0.859 | 0.711 | 2.31e-16 | 7.78e-12 |
| Malignant epithelial | C2 WNT ECs | PSME3 | 0.38 | 0.355 | 0.226 | 2.97e-11 | 1e-06 |
| Malignant epithelial | C2 WNT ECs | RNF115 | 0.38 | 0.273 | 0.163 | 7.41e-11 | 2.5e-06 |
| Malignant epithelial | C2 WNT ECs | C1orf131 | 0.38 | 0.265 | 0.173 | 4.17e-08 | 0.00141 |
| Malignant epithelial | C2 WNT ECs | DPAGT1 | 0.38 | 0.218 | 0.135 | 6.34e-08 | 0.00214 |
| Malignant epithelial | C2 WNT ECs | CMC1 | 0.379 | 0.478 | 0.327 | 2.8e-12 | 9.45e-08 |
| Malignant epithelial | C2 WNT ECs | SEPW1 | 0.379 | 0.9 | 0.821 | 4.69e-12 | 1.58e-07 |
| Malignant epithelial | C2 WNT ECs | TMEM139 | 0.379 | 0.309 | 0.199 | 1.35e-09 | 4.54e-05 |
| Malignant epithelial | C2 WNT ECs | STK38 | 0.379 | 0.204 | 0.124 | 4.79e-08 | 0.00161 |
| Malignant epithelial | C2 WNT ECs | MTFR1 | 0.379 | 0.273 | 0.182 | 8.87e-08 | 0.00299 |
| Malignant epithelial | C2 WNT ECs | NKIRAS2 | 0.378 | 0.249 | 0.158 | 2.21e-08 | 0.000744 |
| Malignant epithelial | C2 WNT ECs | PRRC1 | 0.378 | 0.206 | 0.13 | 2.72e-07 | 0.00916 |
| Malignant epithelial | C2 WNT ECs | CFAP97 | 0.377 | 0.31 | 0.183 | 6.33e-13 | 2.13e-08 |
| Malignant epithelial | C2 WNT ECs | OCLN | 0.377 | 0.315 | 0.193 | 1.52e-11 | 5.11e-07 |
| Malignant epithelial | C2 WNT ECs | SMG1 | 0.377 | 0.419 | 0.277 | 3.99e-11 | 1.34e-06 |
| Malignant epithelial | C2 WNT ECs | RPP21 | 0.377 | 0.314 | 0.196 | 1.03e-10 | 3.47e-06 |
| Malignant epithelial | C2 WNT ECs | CUL5 | 0.377 | 0.238 | 0.154 | 1.77e-07 | 0.00595 |
| Malignant epithelial | C2 WNT ECs | TUFM | 0.376 | 0.827 | 0.67 | 1.34e-18 | 4.52e-14 |
| Malignant epithelial | C2 WNT ECs | STUB1 | 0.376 | 0.777 | 0.6 | 8.21e-18 | 2.76e-13 |
| Malignant epithelial | C2 WNT ECs | SRSF6 | 0.376 | 0.668 | 0.504 | 8.59e-12 | 2.9e-07 |
| Malignant epithelial | C2 WNT ECs | HCCS | 0.376 | 0.33 | 0.203 | 3.22e-11 | 1.09e-06 |
| Malignant epithelial | C2 WNT ECs | NOL4L | 0.376 | 0.257 | 0.163 | 1.53e-08 | 0.000516 |
| Malignant epithelial | C2 WNT ECs | TBK1 | 0.376 | 0.154 | 0.09 | 3.2e-07 | 0.0108 |
| Malignant epithelial | C2 WNT ECs | BAZ1B | 0.375 | 0.411 | 0.262 | 2.01e-13 | 6.78e-09 |
| Malignant epithelial | C2 WNT ECs | MIB2 | 0.375 | 0.292 | 0.17 | 1.23e-12 | 4.13e-08 |
| Malignant epithelial | C2 WNT ECs | ZBTB44 | 0.375 | 0.273 | 0.169 | 1.27e-09 | 4.29e-05 |
| Malignant epithelial | C2 WNT ECs | MIIP | 0.375 | 0.175 | 0.109 | 1.47e-06 | 0.0494 |
| Malignant epithelial | C2 WNT ECs | USP48 | 0.374 | 0.256 | 0.144 | 5.78e-12 | 1.95e-07 |
| Malignant epithelial | C2 WNT ECs | CAMK2G | 0.374 | 0.349 | 0.218 | 1.06e-11 | 3.58e-07 |
| Malignant epithelial | C2 WNT ECs | ETF1 | 0.374 | 0.464 | 0.322 | 3.45e-11 | 1.16e-06 |
| Malignant epithelial | C2 WNT ECs | ADAM15 | 0.374 | 0.329 | 0.207 | 4.11e-11 | 1.38e-06 |
| Malignant epithelial | C2 WNT ECs | HIST1H4C | 0.372 | 0.64 | 0.422 | 9.359999999999999e-23 | 3.15e-18 |
| Malignant epithelial | C2 WNT ECs | BUD31 | 0.372 | 0.865 | 0.694 | 9.93e-19 | 3.35e-14 |
| Malignant epithelial | C2 WNT ECs | ZNF277 | 0.372 | 0.372 | 0.233 | 5.03e-12 | 1.69e-07 |
| Malignant epithelial | C2 WNT ECs | MRPL33 | 0.371 | 0.819 | 0.639 | 3.69e-17 | 1.24e-12 |
| Malignant epithelial | C2 WNT ECs | SAFB | 0.371 | 0.352 | 0.238 | 1.28e-08 | 0.000432 |
| Malignant epithelial | C2 WNT ECs | HDDC2 | 0.37 | 0.574 | 0.394 | 1.28e-13 | 4.31e-09 |
| Malignant epithelial | C2 WNT ECs | TTLL12 | 0.37 | 0.267 | 0.167 | 5.55e-09 | 0.000187 |
| Malignant epithelial | C2 WNT ECs | TEX261 | 0.37 | 0.214 | 0.131 | 3.54e-08 | 0.00119 |
| Malignant epithelial | C2 WNT ECs | SPCS2 | 0.369 | 0.846 | 0.727 | 1.3e-14 | 4.38e-10 |

| compartment | subcluster | gene | avg_log2FC | pct.1 | pct.2 | p_val | p_val_adj |
|---|---|---|---|---|---|---|---|
| Malignant epithelial | C2 WNT ECs | SHROOM1 | 0.369 | 0.253 | 0.153 | 5.82e-10 | 1.96e-05 |
| Malignant epithelial | C2 WNT ECs | ATP2C1 | 0.369 | 0.313 | 0.208 | 3.44e-08 | 0.00116 |
| Malignant epithelial | C2 WNT ECs | MIDN | 0.369 | 0.667 | 0.548 | 3.08e-07 | 0.0104 |
| Malignant epithelial | C2 WNT ECs | LGALS8 | 0.369 | 0.177 | 0.111 | 1.41e-06 | 0.0476 |
| Malignant epithelial | C2 WNT ECs | RPL15 | 0.368 | 0.995 | 0.994 | 2.37e-43 | 7.98e-39 |
| Malignant epithelial | C2 WNT ECs | ATP6V1F | 0.368 | 0.87 | 0.772 | 4.62e-15 | 1.56e-10 |
| Malignant epithelial | C2 WNT ECs | RNF187 | 0.368 | 0.242 | 0.147 | 2.55e-09 | 8.6e-05 |
| Malignant epithelial | C2 WNT ECs | RARS2 | 0.368 | 0.29 | 0.186 | 4.06e-09 | 0.000137 |
| Malignant epithelial | C2 WNT ECs | DXO | 0.368 | 0.187 | 0.109 | 2.38e-08 | 0.000802 |
| Malignant epithelial | C2 WNT ECs | CTTN | 0.367 | 0.668 | 0.5 | 1.1e-13 | 3.72e-09 |
| Malignant epithelial | C2 WNT ECs | USP5 | 0.367 | 0.267 | 0.169 | 4.76e-09 | 0.000161 |
| Malignant epithelial | C2 WNT ECs | CCDC25 | 0.367 | 0.292 | 0.193 | 3.54e-08 | 0.00119 |
| Malignant epithelial | C2 WNT ECs | CHCHD2 | 0.366 | 0.988 | 0.953 | 7.15e-33 | 2.41e-28 |
| Malignant epithelial | C2 WNT ECs | EPS8 | 0.366 | 0.574 | 0.409 | 5.34e-13 | 1.8e-08 |
| Malignant epithelial | C2 WNT ECs | COPS5 | 0.366 | 0.563 | 0.416 | 1.31e-10 | 4.4e-06 |
| Malignant epithelial | C2 WNT ECs | IARS2 | 0.366 | 0.364 | 0.236 | 1.36e-10 | 4.58e-06 |
| Malignant epithelial | C2 WNT ECs | RP11-83A24.2 | 0.366 | 0.214 | 0.132 | 5.07e-08 | 0.00171 |
| Malignant epithelial | C2 WNT ECs | NDUFS5 | 0.365 | 0.931 | 0.836 | 5.68e-20 | 1.91e-15 |
| Malignant epithelial | C2 WNT ECs | SMG5 | 0.365 | 0.158 | 0.096 | 1.1e-06 | 0.037 |
| Malignant epithelial | C2 WNT ECs | RBM39 | 0.364 | 0.958 | 0.868 | 2.54e-17 | 8.57e-13 |
| Malignant epithelial | C2 WNT ECs | AKT1S1 | 0.364 | 0.402 | 0.264 | 4.97e-11 | 1.67e-06 |
| Malignant epithelial | C2 WNT ECs | POLR2K | 0.363 | 0.893 | 0.763 | 7.65e-14 | 2.58e-09 |
| Malignant epithelial | C2 WNT ECs | CWF19L2 | 0.363 | 0.283 | 0.169 | 1.51e-11 | 5.1e-07 |
| Malignant epithelial | C2 WNT ECs | AK6 | 0.363 | 0.428 | 0.286 | 1.66e-11 | 5.59e-07 |
| Malignant epithelial | C2 WNT ECs | GDI1 | 0.363 | 0.403 | 0.265 | 2.01e-11 | 6.77e-07 |
| Malignant epithelial | C2 WNT ECs | NUCB1 | 0.363 | 0.61 | 0.46 | 3.14e-10 | 1.06e-05 |
| Malignant epithelial | C2 WNT ECs | AP1G2 | 0.363 | 0.252 | 0.164 | 1.18e-07 | 0.00399 |
| Malignant epithelial | C2 WNT ECs | MAGOH | 0.362 | 0.643 | 0.443 | 9.82e-16 | 3.31e-11 |
| Malignant epithelial | C2 WNT ECs | GSK3B | 0.362 | 0.294 | 0.199 | 1.92e-07 | 0.00645 |
| Malignant epithelial | C2 WNT ECs | NONO | 0.361 | 0.782 | 0.601 | 6.73e-14 | 2.27e-09 |
| Malignant epithelial | C2 WNT ECs | MAPK1 | 0.361 | 0.325 | 0.22 | 2.58e-08 | 0.00087 |
| Malignant epithelial | C2 WNT ECs | TMEM216 | 0.361 | 0.258 | 0.17 | 3e-07 | 0.0101 |
| Malignant epithelial | C2 WNT ECs | MED31 | 0.361 | 0.211 | 0.136 | 8.83e-07 | 0.0298 |
| Malignant epithelial | C2 WNT ECs | SZRD1 | 0.36 | 0.356 | 0.222 | 2.18e-11 | 7.35e-07 |
| Malignant epithelial | C2 WNT ECs | EIF3A | 0.36 | 0.67 | 0.523 | 2.85e-10 | 9.62e-06 |
| Malignant epithelial | C2 WNT ECs | ERN2 | 0.36 | 0.118 | 0.065 | 1.18e-06 | 0.0399 |
| Malignant epithelial | C2 WNT ECs | PFDN4 | 0.359 | 0.871 | 0.729 | 3.12e-16 | 1.05e-11 |
| Malignant epithelial | C2 WNT ECs | BAZ2A | 0.359 | 0.438 | 0.29 | 2.42e-12 | 8.15e-08 |
| Malignant epithelial | C2 WNT ECs | SNAPC2 | 0.359 | 0.261 | 0.154 | 1.37e-10 | 4.62e-06 |
| Malignant epithelial | C2 WNT ECs | KIAA2013 | 0.359 | 0.265 | 0.169 | 2.71e-08 | 0.000913 |
| Malignant epithelial | C2 WNT ECs | SLC7A5 | 0.359 | 0.355 | 0.251 | 5.08e-08 | 0.00171 |
| Malignant epithelial | C2 WNT ECs | EEF1A1 | 0.358 | 0.999 | 1.0 | 6.97e-35 | 2.35e-30 |
| Malignant epithelial | C2 WNT ECs | ANAPC5 | 0.358 | 0.483 | 0.349 | 4.18e-09 | 0.000141 |
| Malignant epithelial | C2 WNT ECs | PDIA3 | 0.357 | 0.893 | 0.781 | 1.49e-13 | 5.03e-09 |
| Malignant epithelial | C2 WNT ECs | FAM160A2 | 0.357 | 0.357 | 0.22 | 1.25e-12 | 4.22e-08 |
| Malignant epithelial | C2 WNT ECs | PDHA1 | 0.357 | 0.436 | 0.313 | 1.73e-08 | 0.000584 |
| Malignant epithelial | C2 WNT ECs | GNG5 | 0.355 | 0.912 | 0.794 | 1.04e-21 | 3.51e-17 |
| Malignant epithelial | C2 WNT ECs | FAM53C | 0.355 | 0.399 | 0.261 | 6.02e-12 | 2.03e-07 |
| Malignant epithelial | C2 WNT ECs | NFATC3 | 0.355 | 0.467 | 0.329 | 5.69e-10 | 1.92e-05 |
| Malignant epithelial | C2 WNT ECs | MPDU1 | 0.355 | 0.279 | 0.181 | 2.77e-08 | 0.000933 |
| Malignant epithelial | C2 WNT ECs | ZFC3H1 | 0.355 | 0.26 | 0.169 | 6.62e-08 | 0.00223 |
| Malignant epithelial | C2 WNT ECs | ASNSD1 | 0.355 | 0.29 | 0.198 | 2.86e-07 | 0.00965 |
| Malignant epithelial | C2 WNT ECs | BTF3L4 | 0.354 | 0.486 | 0.333 | 1.15e-11 | 3.89e-07 |
| Malignant epithelial | C2 WNT ECs | RPP38 | 0.354 | 0.291 | 0.178 | 1.19e-10 | 4.01e-06 |
| Malignant epithelial | C2 WNT ECs | SNRNP200 | 0.354 | 0.419 | 0.278 | 3.08e-10 | 1.04e-05 |
| Malignant epithelial | C2 WNT ECs | EIF3E | 0.354 | 0.938 | 0.895 | 9.01e-09 | 0.000303 |
| Malignant epithelial | C2 WNT ECs | EDEM2 | 0.354 | 0.277 | 0.183 | 4.96e-08 | 0.00167 |
| Malignant epithelial | C2 WNT ECs | ETV2 | 0.354 | 0.112 | 0.057 | 5.49e-08 | 0.00185 |
| Malignant epithelial | C2 WNT ECs | CXorf56 | 0.354 | 0.15 | 0.089 | 1.11e-06 | 0.0375 |
| Malignant epithelial | C2 WNT ECs | EIF5B | 0.353 | 0.808 | 0.6 | 1.28e-18 | 4.3e-14 |
| Malignant epithelial | C2 WNT ECs | GSS | 0.353 | 0.455 | 0.311 | 6.36e-11 | 2.14e-06 |
| Malignant epithelial | C2 WNT ECs | SNHG9 | 0.353 | 0.463 | 0.322 | 2.16e-10 | 7.29e-06 |
| Malignant epithelial | C2 WNT ECs | MRGBP | 0.353 | 0.46 | 0.315 | 1.03e-09 | 3.46e-05 |
| Malignant epithelial | C2 WNT ECs | VPS33A | 0.353 | 0.133 | 0.07 | 2.63e-08 | 0.000887 |

| compartment | subcluster | gene | avg_log2FC | pct.1 | pct.2 | p_val | p_val_adj |
|---|---|---|---|---|---|---|---|
| Malignant epithelial | C2 WNT ECs | NPTN | 0.352 | 0.254 | 0.169 | 2.66e-07 | 0.00898 |
| Malignant epithelial | C2 WNT ECs | RALGAPB | 0.352 | 0.177 | 0.11 | 6.69e-07 | 0.0225 |
| Malignant epithelial | C2 WNT ECs | CDK2AP2 | 0.351 | 0.72 | 0.557 | 1.08e-12 | 3.65e-08 |
| Malignant epithelial | C2 WNT ECs | SEC31A | 0.351 | 0.39 | 0.25 | 1.13e-11 | 3.79e-07 |
| Malignant epithelial | C2 WNT ECs | DERL1 | 0.351 | 0.516 | 0.365 | 1.07e-10 | 3.61e-06 |
| Malignant epithelial | C2 WNT ECs | VRK3 | 0.351 | 0.306 | 0.189 | 5.11e-10 | 1.72e-05 |
| Malignant epithelial | C2 WNT ECs | HTATSF1 | 0.351 | 0.325 | 0.211 | 6.48e-10 | 2.18e-05 |
| Malignant epithelial | C2 WNT ECs | PCIF1 | 0.351 | 0.314 | 0.209 | 1.18e-08 | 0.000397 |
| Malignant epithelial | C2 WNT ECs | DBP | 0.351 | 0.214 | 0.13 | 3.42e-08 | 0.00115 |
| Malignant epithelial | C2 WNT ECs | MMGT1 | 0.351 | 0.203 | 0.128 | 6.4e-07 | 0.0216 |
| Malignant epithelial | C2 WNT ECs | AKAP8L | 0.35 | 0.371 | 0.235 | 3.38e-11 | 1.14e-06 |
| Malignant epithelial | C2 WNT ECs | CSNK1A1 | 0.349 | 0.747 | 0.569 | 9.67e-12 | 3.26e-07 |
| Malignant epithelial | C2 WNT ECs | NUDCD3 | 0.349 | 0.323 | 0.205 | 2.84e-10 | 9.57e-06 |
| Malignant epithelial | C2 WNT ECs | BDH1 | 0.349 | 0.212 | 0.128 | 1.67e-08 | 0.000564 |
| Malignant epithelial | C2 WNT ECs | PPP1R13L | 0.349 | 0.392 | 0.28 | 8.45e-08 | 0.00285 |
| Malignant epithelial | C2 WNT ECs | PHACTR3 | 0.349 | 0.156 | 0.092 | 4.16e-07 | 0.014 |
| Malignant epithelial | C2 WNT ECs | MSL3 | 0.349 | 0.176 | 0.11 | 1.09e-06 | 0.0368 |
| Malignant epithelial | C2 WNT ECs | MAP3K11 | 0.348 | 0.395 | 0.26 | 4.24e-11 | 1.43e-06 |
| Malignant epithelial | C2 WNT ECs | SELT | 0.348 | 0.666 | 0.503 | 9.32e-11 | 3.14e-06 |
| Malignant epithelial | C2 WNT ECs | RPAIN | 0.348 | 0.414 | 0.278 | 1.96e-10 | 6.62e-06 |
| Malignant epithelial | C2 WNT ECs | MTURN | 0.348 | 0.277 | 0.179 | 2.68e-08 | 0.000904 |
| Malignant epithelial | C2 WNT ECs | HIP1R | 0.348 | 0.189 | 0.115 | 1.31e-07 | 0.00441 |
| Malignant epithelial | C2 WNT ECs | CHMP3 | 0.347 | 0.743 | 0.558 | 5.59e-13 | 1.88e-08 |
| Malignant epithelial | C2 WNT ECs | FAM134C | 0.347 | 0.222 | 0.139 | 6.75e-08 | 0.00228 |
| Malignant epithelial | C2 WNT ECs | SLC38A1 | 0.346 | 0.471 | 0.323 | 3.38e-11 | 1.14e-06 |
| Malignant epithelial | C2 WNT ECs | TAF7 | 0.346 | 0.829 | 0.698 | 1.48e-10 | 4.98e-06 |
| Malignant epithelial | C2 WNT ECs | GTF3C3 | 0.346 | 0.183 | 0.105 | 1.29e-08 | 0.000434 |
| Malignant epithelial | C2 WNT ECs | AP2A2 | 0.344 | 0.256 | 0.158 | 1.41e-09 | 4.76e-05 |
| Malignant epithelial | C2 WNT ECs | RDH11 | 0.344 | 0.315 | 0.221 | 3.68e-07 | 0.0124 |
| Malignant epithelial | C2 WNT ECs | DYNC2LI1 | 0.344 | 0.192 | 0.12 | 5.14e-07 | 0.0173 |
| Malignant epithelial | C2 WNT ECs | TP53RK | 0.343 | 0.344 | 0.229 | 4.26e-09 | 0.000143 |
| Malignant epithelial | C2 WNT ECs | GIGYF2 | 0.342 | 0.242 | 0.159 | 2.56e-07 | 0.00861 |
| Malignant epithelial | C2 WNT ECs | UBA2 | 0.34 | 0.46 | 0.32 | 2.08e-10 | 7.02e-06 |
| Malignant epithelial | C2 WNT ECs | UBE2A | 0.34 | 0.62 | 0.46 | 2.19e-10 | 7.37e-06 |
| Malignant epithelial | C2 WNT ECs | PTRF | 0.34 | 0.26 | 0.175 | 4.51e-07 | 0.0152 |
| Malignant epithelial | C2 WNT ECs | MECP2 | 0.34 | 0.3 | 0.209 | 1.12e-06 | 0.0378 |
| Malignant epithelial | C2 WNT ECs | FTSJ1 | 0.339 | 0.264 | 0.165 | 9.63e-09 | 0.000324 |
| Malignant epithelial | C2 WNT ECs | GNS | 0.339 | 0.18 | 0.105 | 4.74e-08 | 0.0016 |
| Malignant epithelial | C2 WNT ECs | CITED4 | 0.339 | 0.161 | 0.094 | 1.85e-07 | 0.00625 |
| Malignant epithelial | C2 WNT ECs | HSF1 | 0.339 | 0.605 | 0.486 | 2.88e-07 | 0.0097 |
| Malignant epithelial | C2 WNT ECs | SRSF3 | 0.338 | 0.942 | 0.834 | 1.3e-13 | 4.37e-09 |
| Malignant epithelial | C2 WNT ECs | WDR33 | 0.338 | 0.501 | 0.357 | 5.07e-10 | 1.71e-05 |
| Malignant epithelial | C2 WNT ECs | UFD1L | 0.338 | 0.562 | 0.42 | 3.41e-09 | 0.000115 |
| Malignant epithelial | C2 WNT ECs | SORL1 | 0.338 | 0.342 | 0.23 | 8.47e-09 | 0.000285 |
| Malignant epithelial | C2 WNT ECs | PIGQ | 0.338 | 0.202 | 0.128 | 1.31e-06 | 0.0441 |
| Malignant epithelial | C2 WNT ECs | PSMB6 | 0.337 | 0.728 | 0.576 | 4.13e-11 | 1.39e-06 |
| Malignant epithelial | C2 WNT ECs | KAT6A | 0.337 | 0.283 | 0.172 | 1.83e-10 | 6.16e-06 |
| Malignant epithelial | C2 WNT ECs | ETHE1 | 0.337 | 0.701 | 0.58 | 5.69e-10 | 1.92e-05 |
| Malignant epithelial | C2 WNT ECs | PCM1 | 0.337 | 0.267 | 0.166 | 2.29e-09 | 7.72e-05 |
| Malignant epithelial | C2 WNT ECs | LCMT1 | 0.337 | 0.403 | 0.277 | 4.63e-09 | 0.000156 |
| Malignant epithelial | C2 WNT ECs | CSNK1D | 0.337 | 0.448 | 0.335 | 6.93e-07 | 0.0234 |
| Malignant epithelial | C2 WNT ECs | PCBP1 | 0.336 | 0.897 | 0.761 | 7.15e-16 | 2.41e-11 |
| Malignant epithelial | C2 WNT ECs | RTN4 | 0.336 | 0.892 | 0.744 | 5.21e-14 | 1.76e-09 |
| Malignant epithelial | C2 WNT ECs | CLPTM1 | 0.336 | 0.421 | 0.282 | 4.12e-10 | 1.39e-05 |
| Malignant epithelial | C2 WNT ECs | PRKAG1 | 0.336 | 0.34 | 0.228 | 1.16e-08 | 0.00039 |
| Malignant epithelial | C2 WNT ECs | SLC35C2 | 0.335 | 0.414 | 0.276 | 5.68e-11 | 1.91e-06 |
| Malignant epithelial | C2 WNT ECs | SAFB2 | 0.335 | 0.405 | 0.274 | 1.13e-09 | 3.8e-05 |
| Malignant epithelial | C2 WNT ECs | GUCD1 | 0.335 | 0.318 | 0.216 | 6.84e-08 | 0.0023 |
| Malignant epithelial | C2 WNT ECs | EWSR1 | 0.334 | 0.659 | 0.496 | 1.73e-10 | 5.82e-06 |
| Malignant epithelial | C2 WNT ECs | MCCC2 | 0.334 | 0.272 | 0.167 | 6.51e-10 | 2.19e-05 |
| Malignant epithelial | C2 WNT ECs | RICTOR | 0.334 | 0.361 | 0.245 | 2.48e-08 | 0.000835 |
| Malignant epithelial | C2 WNT ECs | RBM42 | 0.333 | 0.694 | 0.495 | 2.12e-15 | 7.13e-11 |
| Malignant epithelial | C2 WNT ECs | RALY | 0.333 | 0.832 | 0.705 | 5.4e-13 | 1.82e-08 |
| Malignant epithelial | C2 WNT ECs | PES1 | 0.333 | 0.332 | 0.211 | 1.97e-10 | 6.63e-06 |

| compartment | subcluster | gene | avg_log2FC | pct.1 | pct.2 | p_val | p_val_adj |
|---|---|---|---|---|---|---|---|
| Malignant epithelial | C2 WNT ECs | HGS | 0.333 | 0.405 | 0.272 | 1.82e-09 | 6.14e-05 |
| Malignant epithelial | C2 WNT ECs | TXNDC9 | 0.332 | 0.456 | 0.313 | 7.77e-11 | 2.62e-06 |
| Malignant epithelial | C2 WNT ECs | NPEPPS | 0.332 | 0.452 | 0.307 | 8.89e-11 | 3e-06 |
| Malignant epithelial | C2 WNT ECs | BCAP29 | 0.332 | 0.398 | 0.276 | 1.89e-08 | 0.000637 |
| Malignant epithelial | C2 WNT ECs | MRPL49 | 0.332 | 0.313 | 0.211 | 5.13e-08 | 0.00173 |
| Malignant epithelial | C2 WNT ECs | FBXO11 | 0.332 | 0.233 | 0.153 | 7.9e-07 | 0.0266 |
| Malignant epithelial | C2 WNT ECs | RBM17 | 0.331 | 0.654 | 0.453 | 1.64e-14 | 5.51e-10 |
| Malignant epithelial | C2 WNT ECs | NOC2L | 0.331 | 0.405 | 0.261 | 7.93e-12 | 2.67e-07 |
| Malignant epithelial | C2 WNT ECs | PUS7L | 0.331 | 0.214 | 0.134 | 1.44e-07 | 0.00486 |
| Malignant epithelial | C2 WNT ECs | POFUT2 | 0.331 | 0.133 | 0.075 | 8.25e-07 | 0.0278 |
| Malignant epithelial | C2 WNT ECs | SF3B2 | 0.33 | 0.755 | 0.579 | 2.68e-11 | 9.03e-07 |
| Malignant epithelial | C2 WNT ECs | IWS1 | 0.33 | 0.307 | 0.195 | 1.13e-09 | 3.79e-05 |
| Malignant epithelial | C2 WNT ECs | LINC00667 | 0.33 | 0.208 | 0.132 | 4.86e-07 | 0.0164 |
| Malignant epithelial | C2 WNT ECs | FBXL4 | 0.33 | 0.13 | 0.074 | 1.08e-06 | 0.0365 |
| Malignant epithelial | C2 WNT ECs | PTP4A1 | 0.328 | 0.679 | 0.483 | 7.38e-14 | 2.49e-09 |
| Malignant epithelial | C2 WNT ECs | SLC25A11 | 0.328 | 0.311 | 0.199 | 8.5e-10 | 2.86e-05 |
| Malignant epithelial | C2 WNT ECs | PPP1R10 | 0.328 | 0.364 | 0.26 | 4.27e-07 | 0.0144 |
| Malignant epithelial | C2 WNT ECs | LSR | 0.327 | 0.885 | 0.752 | 4.23e-15 | 1.43e-10 |
| Malignant epithelial | C2 WNT ECs | RTFDC1 | 0.327 | 0.674 | 0.525 | 3.03e-10 | 1.02e-05 |
| Malignant epithelial | C2 WNT ECs | RSBN1L | 0.327 | 0.575 | 0.42 | 3.39e-10 | 1.14e-05 |
| Malignant epithelial | C2 WNT ECs | PPP3CA | 0.327 | 0.461 | 0.333 | 8.09e-09 | 0.000273 |
| Malignant epithelial | C2 WNT ECs | TOB2 | 0.327 | 0.528 | 0.392 | 8.94e-09 | 0.000301 |
| Malignant epithelial | C2 WNT ECs | ZFAND2B | 0.327 | 0.379 | 0.258 | 8.95e-09 | 0.000302 |
| Malignant epithelial | C2 WNT ECs | TOR1AIP2 | 0.327 | 0.422 | 0.294 | 9.63e-09 | 0.000324 |
| Malignant epithelial | C2 WNT ECs | RFWD2 | 0.327 | 0.226 | 0.143 | 8.63e-08 | 0.00291 |
| Malignant epithelial | C2 WNT ECs | ZRSR2 | 0.327 | 0.18 | 0.11 | 4.13e-07 | 0.0139 |
| Malignant epithelial | C2 WNT ECs | CRNKL1 | 0.327 | 0.222 | 0.147 | 1.1e-06 | 0.0371 |
| Malignant epithelial | C2 WNT ECs | TMEM9B | 0.326 | 0.472 | 0.325 | 2.66e-11 | 8.98e-07 |
| Malignant epithelial | C2 WNT ECs | NFE2L1 | 0.326 | 0.313 | 0.193 | 1.58e-10 | 5.33e-06 |
| Malignant epithelial | C2 WNT ECs | CHP1 | 0.326 | 0.537 | 0.388 | 1.92e-10 | 6.46e-06 |
| Malignant epithelial | C2 WNT ECs | ZRANB2 | 0.326 | 0.363 | 0.236 | 3.11e-10 | 1.05e-05 |
| Malignant epithelial | C2 WNT ECs | PRPF31 | 0.326 | 0.478 | 0.341 | 9.98e-10 | 3.36e-05 |
| Malignant epithelial | C2 WNT ECs | HCFC1 | 0.326 | 0.246 | 0.155 | 2.96e-08 | 0.000996 |
| Malignant epithelial | C2 WNT ECs | PCDH1 | 0.326 | 0.349 | 0.244 | 1.21e-07 | 0.00408 |
| Malignant epithelial | C2 WNT ECs | IRS2 | 0.325 | 0.581 | 0.423 | 3.19e-11 | 1.08e-06 |
| Malignant epithelial | C2 WNT ECs | MTHFD1L | 0.325 | 0.215 | 0.137 | 3.49e-07 | 0.0118 |
| Malignant epithelial | C2 WNT ECs | PRPSAP1 | 0.325 | 0.281 | 0.195 | 1.48e-06 | 0.0499 |
| Malignant epithelial | C2 WNT ECs | SRSF2 | 0.324 | 0.901 | 0.792 | 7.64e-11 | 2.58e-06 |
| Malignant epithelial | C2 WNT ECs | EFNA1 | 0.323 | 0.705 | 0.526 | 1.43e-13 | 4.81e-09 |
| Malignant epithelial | C2 WNT ECs | TMEM129 | 0.323 | 0.185 | 0.112 | 2.07e-07 | 0.00697 |
| Malignant epithelial | C2 WNT ECs | PLCB3 | 0.322 | 0.281 | 0.179 | 6.86e-09 | 0.000231 |
| Malignant epithelial | C2 WNT ECs | PDS5A | 0.322 | 0.283 | 0.184 | 2.96e-08 | 0.000997 |
| Malignant epithelial | C2 WNT ECs | TMX2 | 0.321 | 0.474 | 0.318 | 7.68e-12 | 2.59e-07 |
| Malignant epithelial | C2 WNT ECs | PFDN1 | 0.321 | 0.624 | 0.48 | 5.9e-09 | 0.000199 |
| Malignant epithelial | C2 WNT ECs | DCAF7 | 0.321 | 0.334 | 0.222 | 9.16e-09 | 0.000309 |
| Malignant epithelial | C2 WNT ECs | MPZL1 | 0.32 | 0.396 | 0.254 | 1.49e-11 | 5.03e-07 |
| Malignant epithelial | C2 WNT ECs | GOT2 | 0.32 | 0.365 | 0.24 | 8.06e-10 | 2.71e-05 |
| Malignant epithelial | C2 WNT ECs | LARP4 | 0.32 | 0.219 | 0.129 | 1.83e-09 | 6.18e-05 |
| Malignant epithelial | C2 WNT ECs | RNASEK | 0.32 | 0.13 | 0.066 | 5.83e-09 | 0.000196 |
| Malignant epithelial | C2 WNT ECs | EBPL | 0.32 | 0.69 | 0.585 | 8.61e-08 | 0.0029 |
| Malignant epithelial | C2 WNT ECs | TXNDC17 | 0.319 | 0.792 | 0.631 | 3.81e-12 | 1.29e-07 |
| Malignant epithelial | C2 WNT ECs | SYNE4 | 0.319 | 0.185 | 0.111 | 1.84e-07 | 0.00621 |
| Malignant epithelial | C2 WNT ECs | MARVELD3 | 0.318 | 0.399 | 0.28 | 5.91e-08 | 0.00199 |
| Malignant epithelial | C2 WNT ECs | ARCN1 | 0.318 | 0.288 | 0.195 | 7.85e-08 | 0.00264 |
| Malignant epithelial | C2 WNT ECs | SDF4 | 0.317 | 0.62 | 0.432 | 1.23e-14 | 4.16e-10 |
| Malignant epithelial | C2 WNT ECs | GSKIP | 0.317 | 0.391 | 0.25 | 2.16e-11 | 7.29e-07 |
| Malignant epithelial | C2 WNT ECs | EIF4A3 | 0.317 | 0.57 | 0.421 | 1.18e-08 | 0.000397 |
| Malignant epithelial | C2 WNT ECs | SHC1 | 0.317 | 0.265 | 0.18 | 1.17e-06 | 0.0396 |
| Malignant epithelial | C2 WNT ECs | TMBIM6 | 0.316 | 0.968 | 0.884 | 1.88e-14 | 6.34e-10 |
| Malignant epithelial | C2 WNT ECs | SDCBP | 0.316 | 0.69 | 0.548 | 5.96e-11 | 2.01e-06 |
| Malignant epithelial | C2 WNT ECs | DEF8 | 0.316 | 0.334 | 0.22 | 2.47e-09 | 8.31e-05 |
| Malignant epithelial | C2 WNT ECs | TALDO1 | 0.315 | 0.85 | 0.695 | 3.74e-16 | 1.26e-11 |
| Malignant epithelial | C2 WNT ECs | BNIP2 | 0.315 | 0.379 | 0.241 | 6.96e-11 | 2.35e-06 |
| Malignant epithelial | C2 WNT ECs | IK | 0.315 | 0.594 | 0.423 | 8.41e-10 | 2.83e-05 |

| compartment | subcluster | gene | avg_log2FC | pct.1 | pct.2 | p_val | p_val_adj |
|---|---|---|---|---|---|---|---|
| Malignant epithelial | C2 WNT ECs | PPP4R2 | 0.315 | 0.437 | 0.31 | 5.5e-09 | 0.000185 |
| Malignant epithelial | C2 WNT ECs | VPS72 | 0.315 | 0.326 | 0.216 | 5.6e-09 | 0.000189 |
| Malignant epithelial | C2 WNT ECs | PRDX3 | 0.315 | 0.558 | 0.43 | 3.37e-08 | 0.00114 |
| Malignant epithelial | C2 WNT ECs | WBSCR16 | 0.315 | 0.249 | 0.165 | 4.84e-07 | 0.0163 |
| Malignant epithelial | C2 WNT ECs | MUS81 | 0.314 | 0.211 | 0.124 | 4.24e-09 | 0.000143 |
| Malignant epithelial | C2 WNT ECs | NMD3 | 0.314 | 0.295 | 0.2 | 1.47e-07 | 0.00496 |
| Malignant epithelial | C2 WNT ECs | USP16 | 0.313 | 0.52 | 0.377 | 9.74e-10 | 3.28e-05 |
| Malignant epithelial | C2 WNT ECs | GID8 | 0.313 | 0.696 | 0.554 | 3.93e-09 | 0.000132 |
| Malignant epithelial | C2 WNT ECs | MXI1 | 0.313 | 0.265 | 0.165 | 5.46e-09 | 0.000184 |
| Malignant epithelial | C2 WNT ECs | TMED2 | 0.313 | 0.84 | 0.743 | 9.26e-09 | 0.000312 |
| Malignant epithelial | C2 WNT ECs | RBM22 | 0.313 | 0.341 | 0.236 | 2.11e-07 | 0.0071 |
| Malignant epithelial | C2 WNT ECs | ATXN2L | 0.312 | 0.365 | 0.246 | 9.96e-09 | 0.000336 |
| Malignant epithelial | C2 WNT ECs | TRABD | 0.312 | 0.529 | 0.4 | 1.67e-08 | 0.000563 |
| Malignant epithelial | C2 WNT ECs | LAMTOR3 | 0.312 | 0.38 | 0.265 | 3.89e-08 | 0.00131 |
| Malignant epithelial | C2 WNT ECs | NUDT16L1 | 0.312 | 0.398 | 0.285 | 7.87e-08 | 0.00265 |
| Malignant epithelial | C2 WNT ECs | BNIP3L | 0.312 | 0.419 | 0.302 | 3.83e-07 | 0.0129 |
| Malignant epithelial | C2 WNT ECs | RHOC | 0.312 | 0.827 | 0.719 | 9.94e-07 | 0.0335 |
| Malignant epithelial | C2 WNT ECs | DAZAP2 | 0.311 | 0.861 | 0.686 | 4.5e-14 | 1.52e-09 |
| Malignant epithelial | C2 WNT ECs | DPH3 | 0.311 | 0.363 | 0.241 | 6.69e-09 | 0.000226 |
| Malignant epithelial | C2 WNT ECs | ARID2 | 0.31 | 0.168 | 0.097 | 9.08e-08 | 0.00306 |
| Malignant epithelial | C2 WNT ECs | HNRNPA1 | 0.309 | 0.982 | 0.957 | 1.1e-14 | 3.71e-10 |
| Malignant epithelial | C2 WNT ECs | MRPL37 | 0.309 | 0.411 | 0.289 | 2.08e-08 | 0.000701 |
| Malignant epithelial | C2 WNT ECs | SP3 | 0.309 | 0.349 | 0.243 | 2.81e-07 | 0.00947 |
| Malignant epithelial | C2 WNT ECs | CNBP | 0.308 | 0.855 | 0.731 | 1.05e-10 | 3.52e-06 |
| Malignant epithelial | C2 WNT ECs | CGGBP1 | 0.308 | 0.395 | 0.269 | 4.49e-09 | 0.000151 |
| Malignant epithelial | C2 WNT ECs | CTNNBL1 | 0.308 | 0.44 | 0.314 | 3.74e-08 | 0.00126 |
| Malignant epithelial | C2 WNT ECs | DDX18 | 0.307 | 0.655 | 0.496 | 1.41e-10 | 4.74e-06 |
| Malignant epithelial | C2 WNT ECs | UBE2F | 0.307 | 0.464 | 0.324 | 1.38e-09 | 4.65e-05 |
| Malignant epithelial | C2 WNT ECs | MAP2K1 | 0.307 | 0.291 | 0.188 | 1.66e-08 | 0.000558 |
| Malignant epithelial | C2 WNT ECs | ARFIP2 | 0.307 | 0.346 | 0.237 | 4.36e-08 | 0.00147 |
| Malignant epithelial | C2 WNT ECs | RRBP1 | 0.307 | 0.765 | 0.629 | 7.35e-08 | 0.00248 |
| Malignant epithelial | C2 WNT ECs | SHMT2 | 0.307 | 0.387 | 0.275 | 8.24e-08 | 0.00278 |
| Malignant epithelial | C2 WNT ECs | MINK1 | 0.307 | 0.143 | 0.083 | 6.03e-07 | 0.0203 |
| Malignant epithelial | C2 WNT ECs | NUDT14 | 0.306 | 0.563 | 0.406 | 9.57e-11 | 3.23e-06 |
| Malignant epithelial | C2 WNT ECs | MPHOSPH10 | 0.306 | 0.345 | 0.236 | 4.05e-08 | 0.00136 |
| Malignant epithelial | C2 WNT ECs | ICMT | 0.306 | 0.203 | 0.124 | 7.89e-08 | 0.00266 |
| Malignant epithelial | C2 WNT ECs | ASXL1 | 0.306 | 0.275 | 0.187 | 6.71e-07 | 0.0226 |
| Malignant epithelial | C2 WNT ECs | CYSTM1 | 0.305 | 0.873 | 0.736 | 1.8e-14 | 6.05e-10 |
| Malignant epithelial | C2 WNT ECs | CCT8 | 0.305 | 0.72 | 0.557 | 2.15e-10 | 7.26e-06 |
| Malignant epithelial | C2 WNT ECs | RBBP6 | 0.305 | 0.618 | 0.498 | 5.1e-08 | 0.00172 |
| Malignant epithelial | C2 WNT ECs | CAND1 | 0.305 | 0.407 | 0.287 | 5.19e-08 | 0.00175 |
| Malignant epithelial | C2 WNT ECs | RAB21 | 0.305 | 0.384 | 0.279 | 2.24e-07 | 0.00753 |
| Malignant epithelial | C2 WNT ECs | VAMP3 | 0.304 | 0.488 | 0.332 | 2.83e-11 | 9.53e-07 |
| Malignant epithelial | C2 WNT ECs | E4F1 | 0.304 | 0.207 | 0.114 | 1.89e-10 | 6.35e-06 |
| Malignant epithelial | C2 WNT ECs | ISY1 | 0.304 | 0.298 | 0.202 | 6.48e-07 | 0.0218 |
| Malignant epithelial | C2 WNT ECs | CDH17 | 0.303 | 0.493 | 0.277 | 9.759999999999999e-23 | 3.29e-18 |
| Malignant epithelial | C2 WNT ECs | GSTK1 | 0.303 | 0.839 | 0.698 | 2.03e-14 | 6.83e-10 |
| Malignant epithelial | C2 WNT ECs | CLDND1 | 0.303 | 0.317 | 0.203 | 1.09e-09 | 3.66e-05 |
| Malignant epithelial | C2 WNT ECs | TMX4 | 0.303 | 0.333 | 0.218 | 6.43e-09 | 0.000217 |
| Malignant epithelial | C2 WNT ECs | SLC30A5 | 0.303 | 0.245 | 0.156 | 3.09e-08 | 0.00104 |
| Malignant epithelial | C2 WNT ECs | TMEM69 | 0.303 | 0.275 | 0.181 | 1.87e-07 | 0.00629 |
| Malignant epithelial | C2 WNT ECs | HNRNPA2B1 | 0.302 | 0.966 | 0.889 | 9.94e-16 | 3.35e-11 |
| Malignant epithelial | C2 WNT ECs | TCF25 | 0.302 | 0.721 | 0.548 | 1.34e-12 | 4.5e-08 |
| Malignant epithelial | C2 WNT ECs | SLC25A3 | 0.302 | 0.9 | 0.785 | 6.67e-12 | 2.25e-07 |
| Malignant epithelial | C2 WNT ECs | PDCL3 | 0.302 | 0.391 | 0.255 | 9.23e-11 | 3.11e-06 |
| Malignant epithelial | C2 WNT ECs | SLC35A3 | 0.302 | 0.268 | 0.174 | 7.62e-08 | 0.00257 |
| Malignant epithelial | C2 WNT ECs | KDM2A | 0.302 | 0.391 | 0.282 | 6.93e-07 | 0.0234 |
| Malignant epithelial | C2 WNT ECs | TMEM55B | 0.301 | 0.337 | 0.22 | 3.36e-09 | 0.000113 |
| Malignant epithelial | C2 WNT ECs | ANKLE2 | 0.301 | 0.461 | 0.327 | 4.45e-09 | 0.00015 |
| Malignant epithelial | C2 WNT ECs | CNP | 0.301 | 0.357 | 0.241 | 1.38e-08 | 0.000465 |
| Malignant epithelial | C2 WNT ECs | SYMPK | 0.301 | 0.309 | 0.205 | 3.4e-08 | 0.00115 |
| Malignant epithelial | C2 WNT ECs | RELA | 0.3 | 0.355 | 0.229 | 6.21e-10 | 2.09e-05 |
| Malignant epithelial | C2 WNT ECs | KDM3A | 0.3 | 0.171 | 0.102 | 2.12e-07 | 0.00714 |
| Malignant epithelial | C2 WNT ECs | AZIN1 | 0.3 | 0.585 | 0.462 | 4.59e-07 | 0.0155 |

| compartment | subcluster | gene | avg_log2FC | pct.1 | pct.2 | p_val | p_val_adj |
|---|---|---|---|---|---|---|---|
| Malignant epithelial | C2 WNT ECs | DCXR | 0.299 | 0.613 | 0.463 | 2.19e-08 | 0.000738 |
| Malignant epithelial | C2 WNT ECs | CC2D1A | 0.299 | 0.241 | 0.157 | 2.75e-07 | 0.00927 |
| Malignant epithelial | C2 WNT ECs | RPL7 | 0.298 | 0.995 | 0.998 | 5.37e-18 | 1.81e-13 |
| Malignant epithelial | C2 WNT ECs | HEBP1 | 0.298 | 0.513 | 0.36 | 2.38e-10 | 8.02e-06 |
| Malignant epithelial | C2 WNT ECs | PPP1R37 | 0.298 | 0.258 | 0.162 | 1.05e-08 | 0.000353 |
| Malignant epithelial | C2 WNT ECs | FRG1 | 0.297 | 0.434 | 0.282 | 8.44e-12 | 2.84e-07 |
| Malignant epithelial | C2 WNT ECs | MRPS10 | 0.297 | 0.556 | 0.397 | 1.13e-10 | 3.79e-06 |
| Malignant epithelial | C2 WNT ECs | WDR74 | 0.297 | 0.529 | 0.378 | 2.49e-09 | 8.39e-05 |
| Malignant epithelial | C2 WNT ECs | IGFBP4 | 0.297 | 0.387 | 0.272 | 3.95e-08 | 0.00133 |
| Malignant epithelial | C2 WNT ECs | ROGDI | 0.297 | 0.234 | 0.151 | 3.18e-07 | 0.0107 |
| Malignant epithelial | C2 WNT ECs | ARFIP1 | 0.297 | 0.237 | 0.153 | 3.52e-07 | 0.0118 |
| Malignant epithelial | C2 WNT ECs | PHF3 | 0.296 | 0.322 | 0.201 | 2.74e-10 | 9.24e-06 |
| Malignant epithelial | C2 WNT ECs | C1GALT1 | 0.296 | 0.365 | 0.251 | 3.62e-08 | 0.00122 |
| Malignant epithelial | C2 WNT ECs | RTN3 | 0.296 | 0.732 | 0.594 | 5.04e-08 | 0.0017 |
| Malignant epithelial | C2 WNT ECs | KATNBL1 | 0.296 | 0.344 | 0.242 | 8.52e-07 | 0.0287 |
| Malignant epithelial | C2 WNT ECs | NABP2 | 0.295 | 0.398 | 0.271 | 6.62e-09 | 0.000223 |
| Malignant epithelial | C2 WNT ECs | ZNF335 | 0.295 | 0.183 | 0.108 | 7.91e-08 | 0.00267 |
| Malignant epithelial | C2 WNT ECs | HMCES | 0.295 | 0.325 | 0.225 | 4e-07 | 0.0135 |
| Malignant epithelial | C2 WNT ECs | RNF14 | 0.295 | 0.158 | 0.095 | 9.55e-07 | 0.0322 |
| Malignant epithelial | C2 WNT ECs | MRPL57 | 0.294 | 0.835 | 0.729 | 1.78e-10 | 6.01e-06 |
| Malignant epithelial | C2 WNT ECs | MLEC | 0.294 | 0.667 | 0.521 | 1.12e-09 | 3.76e-05 |
| Malignant epithelial | C2 WNT ECs | SLC35A1 | 0.294 | 0.268 | 0.174 | 5.73e-08 | 0.00193 |
| Malignant epithelial | C2 WNT ECs | TSPAN13 | 0.293 | 0.494 | 0.336 | 1.12e-11 | 3.76e-07 |
| Malignant epithelial | C2 WNT ECs | YIPF4 | 0.293 | 0.558 | 0.388 | 3.51e-11 | 1.18e-06 |
| Malignant epithelial | C2 WNT ECs | SF3B3 | 0.293 | 0.277 | 0.176 | 9.62e-09 | 0.000324 |
| Malignant epithelial | C2 WNT ECs | BACE2 | 0.293 | 0.507 | 0.382 | 5.2e-08 | 0.00175 |
| Malignant epithelial | C2 WNT ECs | SDF2L1 | 0.292 | 0.686 | 0.494 | 1.11e-13 | 3.75e-09 |
| Malignant epithelial | C2 WNT ECs | PSMA7 | 0.292 | 0.957 | 0.893 | 1.91e-11 | 6.45e-07 |
| Malignant epithelial | C2 WNT ECs | DCAF11 | 0.292 | 0.3 | 0.189 | 1.71e-09 | 5.77e-05 |
| Malignant epithelial | C2 WNT ECs | IP6K1 | 0.292 | 0.184 | 0.11 | 1.25e-07 | 0.00422 |
| Malignant epithelial | C2 WNT ECs | PPIG | 0.291 | 0.735 | 0.547 | 7.68e-13 | 2.59e-08 |
| Malignant epithelial | C2 WNT ECs | DYNLRB1 | 0.291 | 0.963 | 0.908 | 7.97e-11 | 2.68e-06 |
| Malignant epithelial | C2 WNT ECs | MTHFD2L | 0.291 | 0.271 | 0.162 | 4.79e-10 | 1.61e-05 |
| Malignant epithelial | C2 WNT ECs | DGCR2 | 0.291 | 0.279 | 0.176 | 7.96e-09 | 0.000268 |
| Malignant epithelial | C2 WNT ECs | RAD9A | 0.291 | 0.344 | 0.235 | 4.49e-08 | 0.00151 |
| Malignant epithelial | C2 WNT ECs | SLCO4A1 | 0.291 | 0.372 | 0.27 | 4.02e-07 | 0.0135 |
| Malignant epithelial | C2 WNT ECs | HADHB | 0.29 | 0.474 | 0.331 | 8.91e-10 | 3e-05 |
| Malignant epithelial | C2 WNT ECs | NAA60 | 0.289 | 0.378 | 0.239 | 3.62e-11 | 1.22e-06 |
| Malignant epithelial | C2 WNT ECs | CLTC | 0.289 | 0.391 | 0.261 | 1.51e-09 | 5.09e-05 |
| Malignant epithelial | C2 WNT ECs | ATXN3 | 0.289 | 0.256 | 0.164 | 5.38e-08 | 0.00181 |
| Malignant epithelial | C2 WNT ECs | PDLIM5 | 0.289 | 0.44 | 0.315 | 1e-07 | 0.00338 |
| Malignant epithelial | C2 WNT ECs | ATP6V1H | 0.289 | 0.472 | 0.349 | 7.38e-07 | 0.0249 |
| Malignant epithelial | C2 WNT ECs | FLOT1 | 0.288 | 0.663 | 0.491 | 1.77e-10 | 5.96e-06 |
| Malignant epithelial | C2 WNT ECs | SSRP1 | 0.288 | 0.493 | 0.367 | 4.74e-08 | 0.0016 |
| Malignant epithelial | C2 WNT ECs | KPNA4 | 0.288 | 0.382 | 0.275 | 1.74e-07 | 0.00585 |
| Malignant epithelial | C2 WNT ECs | ERGIC1 | 0.288 | 0.375 | 0.266 | 1.86e-07 | 0.00628 |
| Malignant epithelial | C2 WNT ECs | RHBDF2 | 0.287 | 0.261 | 0.155 | 2.85e-10 | 9.61e-06 |
| Malignant epithelial | C2 WNT ECs | STAMBP | 0.287 | 0.409 | 0.285 | 3.68e-08 | 0.00124 |
| Malignant epithelial | C2 WNT ECs | SPPL3 | 0.287 | 0.31 | 0.208 | 2.1e-07 | 0.00709 |
| Malignant epithelial | C2 WNT ECs | ARHGEF35 | 0.286 | 0.356 | 0.247 | 9.48e-08 | 0.0032 |
| Malignant epithelial | C2 WNT ECs | SAE1 | 0.285 | 0.401 | 0.277 | 2.24e-08 | 0.000754 |
| Malignant epithelial | C2 WNT ECs | RCE1 | 0.285 | 0.336 | 0.226 | 2.58e-08 | 0.000869 |
| Malignant epithelial | C2 WNT ECs | AP3B1 | 0.285 | 0.208 | 0.131 | 4.82e-07 | 0.0163 |
| Malignant epithelial | C2 WNT ECs | TBL1XR1 | 0.284 | 0.463 | 0.324 | 2e-09 | 6.73e-05 |
| Malignant epithelial | C2 WNT ECs | STAG2 | 0.284 | 0.361 | 0.242 | 3.46e-09 | 0.000117 |
| Malignant epithelial | C2 WNT ECs | TUBA1C | 0.284 | 0.832 | 0.688 | 3.41e-08 | 0.00115 |
| Malignant epithelial | C2 WNT ECs | BMPR2 | 0.284 | 0.204 | 0.128 | 3.98e-07 | 0.0134 |
| Malignant epithelial | C2 WNT ECs | KRT20 | 0.283 | 0.463 | 0.291 | 7.19e-16 | 2.42e-11 |
| Malignant epithelial | C2 WNT ECs | CCT7 | 0.283 | 0.7 | 0.561 | 1.42e-08 | 0.000477 |
| Malignant epithelial | C2 WNT ECs | SPAG7 | 0.283 | 0.461 | 0.33 | 1.47e-08 | 0.000495 |
| Malignant epithelial | C2 WNT ECs | HYI | 0.283 | 0.321 | 0.215 | 4.07e-08 | 0.00137 |
| Malignant epithelial | C2 WNT ECs | ZNF703 | 0.282 | 0.667 | 0.483 | 2.7e-11 | 9.09e-07 |
| Malignant epithelial | C2 WNT ECs | UXS1 | 0.282 | 0.345 | 0.215 | 6.15e-11 | 2.07e-06 |
| Malignant epithelial | C2 WNT ECs | DCTPP1 | 0.282 | 0.556 | 0.428 | 4.79e-08 | 0.00161 |

| compartment | subcluster | gene | avg_log2FC | pct.1 | pct.2 | p_val | p_val_adj |
|---|---|---|---|---|---|---|---|
| Malignant epithelial | C2 WNT ECs | MPRIP | 0.282 | 0.223 | 0.142 | 1.66e-07 | 0.00559 |
| Malignant epithelial | C2 WNT ECs | CTDSP2 | 0.282 | 0.363 | 0.259 | 1.32e-06 | 0.0444 |
| Malignant epithelial | C2 WNT ECs | ZDHHC4 | 0.28 | 0.49 | 0.352 | 5.21e-09 | 0.000176 |
| Malignant epithelial | C2 WNT ECs | HDAC10 | 0.28 | 0.219 | 0.138 | 2.45e-07 | 0.00827 |
| Malignant epithelial | C2 WNT ECs | AATF | 0.28 | 0.395 | 0.282 | 3.83e-07 | 0.0129 |
| Malignant epithelial | C2 WNT ECs | HNRNPUL2 | 0.279 | 0.355 | 0.246 | 1.63e-07 | 0.0055 |
| Malignant epithelial | C2 WNT ECs | DCTN4 | 0.279 | 0.2 | 0.125 | 3.1e-07 | 0.0104 |
| Malignant epithelial | C2 WNT ECs | EIF4H | 0.278 | 0.74 | 0.561 | 8.4e-12 | 2.83e-07 |
| Malignant epithelial | C2 WNT ECs | EIF2S2 | 0.278 | 0.911 | 0.803 | 2.03e-11 | 6.85e-07 |
| Malignant epithelial | C2 WNT ECs | TBRG1 | 0.278 | 0.286 | 0.179 | 4.47e-09 | 0.000151 |
| Malignant epithelial | C2 WNT ECs | EGLN2 | 0.278 | 0.365 | 0.256 | 1.67e-07 | 0.00562 |
| Malignant epithelial | C2 WNT ECs | ATP9B | 0.278 | 0.103 | 0.054 | 9.69e-07 | 0.0326 |
| Malignant epithelial | C2 WNT ECs | CLN3 | 0.278 | 0.318 | 0.222 | 9.93e-07 | 0.0335 |
| Malignant epithelial | C2 WNT ECs | SLC25A6 | 0.276 | 0.961 | 0.922 | 1.06e-18 | 3.58e-14 |
| Malignant epithelial | C2 WNT ECs | WWP2 | 0.276 | 0.153 | 0.088 | 2.17e-07 | 0.00732 |
| Malignant epithelial | C2 WNT ECs | SLC4A2 | 0.276 | 0.287 | 0.194 | 2.19e-07 | 0.00739 |
| Malignant epithelial | C2 WNT ECs | HM13 | 0.275 | 0.714 | 0.574 | 1.23e-08 | 0.000414 |
| Malignant epithelial | C2 WNT ECs | DPH5 | 0.274 | 0.191 | 0.11 | 1.57e-08 | 0.000529 |
| Malignant epithelial | C2 WNT ECs | CAPZA2 | 0.273 | 0.708 | 0.564 | 9.32e-08 | 0.00314 |
| Malignant epithelial | C2 WNT ECs | TMEM208 | 0.272 | 0.747 | 0.589 | 8.59e-12 | 2.89e-07 |
| Malignant epithelial | C2 WNT ECs | USP15 | 0.272 | 0.353 | 0.225 | 1.3e-10 | 4.38e-06 |
| Malignant epithelial | C2 WNT ECs | RPS19BP1 | 0.272 | 0.767 | 0.609 | 3.36e-10 | 1.13e-05 |
| Malignant epithelial | C2 WNT ECs | POLR2G | 0.272 | 0.597 | 0.433 | 4.2e-10 | 1.41e-05 |
| Malignant epithelial | C2 WNT ECs | NUP62 | 0.272 | 0.275 | 0.178 | 2.11e-08 | 0.000712 |
| Malignant epithelial | C2 WNT ECs | EPC1 | 0.272 | 0.582 | 0.444 | 1.9e-07 | 0.00641 |
| Malignant epithelial | C2 WNT ECs | PVRL2 | 0.272 | 0.574 | 0.432 | 2.74e-07 | 0.00924 |
| Malignant epithelial | C2 WNT ECs | TOP1 | 0.271 | 0.709 | 0.546 | 8.15e-10 | 2.75e-05 |
| Malignant epithelial | C2 WNT ECs | OTUB1 | 0.271 | 0.618 | 0.468 | 8.42e-09 | 0.000284 |
| Malignant epithelial | C2 WNT ECs | PPIL3 | 0.271 | 0.336 | 0.232 | 7.08e-07 | 0.0239 |
| Malignant epithelial | C2 WNT ECs | AHSA1 | 0.27 | 0.605 | 0.45 | 5.38e-09 | 0.000181 |
| Malignant epithelial | C2 WNT ECs | HMGN3 | 0.27 | 0.43 | 0.315 | 8.68e-08 | 0.00292 |
| Malignant epithelial | C2 WNT ECs | USP39 | 0.27 | 0.244 | 0.157 | 2.56e-07 | 0.00861 |
| Malignant epithelial | C2 WNT ECs | IMPDH2 | 0.27 | 0.677 | 0.552 | 5.38e-07 | 0.0181 |
| Malignant epithelial | C2 WNT ECs | F11R | 0.269 | 0.459 | 0.334 | 4.17e-08 | 0.00141 |
| Malignant epithelial | C2 WNT ECs | FNDC3B | 0.269 | 0.321 | 0.224 | 5.71e-07 | 0.0192 |
| Malignant epithelial | C2 WNT ECs | LENG1 | 0.268 | 0.311 | 0.208 | 1.27e-07 | 0.00427 |
| Malignant epithelial | C2 WNT ECs | ELMSAN1 | 0.268 | 0.28 | 0.191 | 6.51e-07 | 0.0219 |
| Malignant epithelial | C2 WNT ECs | CDHR5 | 0.267 | 0.279 | 0.156 | 1.57e-13 | 5.28e-09 |
| Malignant epithelial | C2 WNT ECs | ZNF281 | 0.266 | 0.231 | 0.15 | 5.2e-07 | 0.0175 |
| Malignant epithelial | C2 WNT ECs | EIF4EBP2 | 0.265 | 0.428 | 0.278 | 1.29e-11 | 4.33e-07 |
| Malignant epithelial | C2 WNT ECs | PTK2 | 0.265 | 0.396 | 0.254 | 1.79e-11 | 6.02e-07 |
| Malignant epithelial | C2 WNT ECs | UBE2N | 0.265 | 0.7 | 0.557 | 4.35e-09 | 0.000147 |
| Malignant epithelial | C2 WNT ECs | HNRNPUL1 | 0.264 | 0.51 | 0.348 | 4.05e-11 | 1.36e-06 |
| Malignant epithelial | C2 WNT ECs | WDR60 | 0.264 | 0.35 | 0.232 | 3.9e-09 | 0.000132 |
| Malignant epithelial | C2 WNT ECs | TUSC2 | 0.264 | 0.509 | 0.365 | 9.35e-09 | 0.000315 |
| Malignant epithelial | C2 WNT ECs | CALCOCO2 | 0.264 | 0.417 | 0.292 | 3.43e-08 | 0.00115 |
| Malignant epithelial | C2 WNT ECs | TCERG1 | 0.264 | 0.373 | 0.257 | 5.81e-08 | 0.00196 |
| Malignant epithelial | C2 WNT ECs | AFMID | 0.264 | 0.304 | 0.209 | 3.46e-07 | 0.0116 |
| Malignant epithelial | C2 WNT ECs | ITGB1BP1 | 0.263 | 0.517 | 0.37 | 1.04e-09 | 3.51e-05 |
| Malignant epithelial | C2 WNT ECs | C11orf73 | 0.263 | 0.563 | 0.406 | 3.3e-09 | 0.000111 |
| Malignant epithelial | C2 WNT ECs | SMURF2 | 0.263 | 0.254 | 0.165 | 1.35e-07 | 0.00455 |
| Malignant epithelial | C2 WNT ECs | URI1 | 0.263 | 0.283 | 0.188 | 3.2e-07 | 0.0108 |
| Malignant epithelial | C2 WNT ECs | C6orf89 | 0.261 | 0.325 | 0.215 | 8.12e-09 | 0.000274 |
| Malignant epithelial | C2 WNT ECs | PMPCB | 0.261 | 0.502 | 0.361 | 1.3e-08 | 0.000439 |
| Malignant epithelial | C2 WNT ECs | AHCY | 0.261 | 0.631 | 0.504 | 1.62e-07 | 0.00547 |
| Malignant epithelial | C2 WNT ECs | PUF60 | 0.261 | 0.678 | 0.551 | 4.18e-07 | 0.0141 |
| Malignant epithelial | C2 WNT ECs | ELAVL1 | 0.26 | 0.514 | 0.368 | 1.36e-09 | 4.6e-05 |
| Malignant epithelial | C2 WNT ECs | CBX3 | 0.26 | 0.862 | 0.755 | 3.37e-09 | 0.000114 |
| Malignant epithelial | C2 WNT ECs | CEACAM5 | 0.259 | 0.606 | 0.342 | 1.97e-29 | 6.63e-25 |
| Malignant epithelial | C2 WNT ECs | MRPL32 | 0.259 | 0.633 | 0.475 | 5.4e-09 | 0.000182 |
| Malignant epithelial | C2 WNT ECs | CKAP4 | 0.259 | 0.263 | 0.167 | 3.12e-08 | 0.00105 |
| Malignant epithelial | C2 WNT ECs | PCF11 | 0.259 | 0.39 | 0.272 | 3.87e-08 | 0.0013 |
| Malignant epithelial | C2 WNT ECs | TMEM30A | 0.258 | 0.318 | 0.218 | 2.35e-07 | 0.00793 |
| Malignant epithelial | C2 WNT ECs | KMT2C | 0.257 | 0.327 | 0.225 | 1.59e-07 | 0.00536 |

| compartment | subcluster | gene | avg_log2FC | pct.1 | pct.2 | p_val | p_val_adj |
|---|---|---|---|---|---|---|---|
| Malignant epithelial | C2 WNT ECs | PTBP1 | 0.256 | 0.74 | 0.567 | 4.75e-09 | 0.00016 |
| Malignant epithelial | C2 WNT ECs | THYN1 | 0.255 | 0.442 | 0.298 | 9.37e-10 | 3.16e-05 |
| Malignant epithelial | C2 WNT ECs | RAC1 | 0.254 | 0.995 | 0.973 | 1.48e-14 | 4.98e-10 |
| Malignant epithelial | C2 WNT ECs | UBXN1 | 0.253 | 0.858 | 0.706 | 4.06e-12 | 1.37e-07 |
| Malignant epithelial | C2 WNT ECs | RPL17 | 0.253 | 0.9 | 0.848 | 1e-10 | 3.38e-06 |
| Malignant epithelial | C2 WNT ECs | PCMTD1 | 0.253 | 0.512 | 0.38 | 5.38e-08 | 0.00181 |
| Malignant epithelial | C2 WNT ECs | LRRC28 | 0.252 | 0.162 | 0.094 | 2.97e-07 | 0.01 |
| Malignant epithelial | C2 WNT ECs | RSF1 | 0.251 | 0.567 | 0.398 | 2.71e-10 | 9.15e-06 |
| Malignant epithelial | C2 WNT ECs | CCT3 | 0.25 | 0.737 | 0.604 | 4.55e-08 | 0.00153 |
| Malignant epithelial | C3 Mp ECs | KIF14 | 5.071 | 0.189 | 0.006 | 1.89e-150 | 6.37e-146 |
| Malignant epithelial | C3 Mp ECs | DEPDC1 | 5.011 | 0.16 | 0.006 | 4.02e-123 | 1.35e-118 |
| Malignant epithelial | C3 Mp ECs | DLGAP5 | 4.893 | 0.319 | 0.015 | 2.72e-239 | 9.160000000000001e-235 |
| Malignant epithelial | C3 Mp ECs | KIF18B | 4.881 | 0.132 | 0.006 | 1.29e-99 | 4.33e-95 |
| Malignant epithelial | C3 Mp ECs | CENPA | 4.861 | 0.433 | 0.025 | 3.23e-310 | 1.09e-305 |
| Malignant epithelial | C3 Mp ECs | NUF2 | 4.841 | 0.375 | 0.017 | 5.75e-285 | 1.94e-280 |
| Malignant epithelial | C3 Mp ECs | GTSE1 | 4.727 | 0.388 | 0.024 | 3.22e-270 | 1.09e-265 |
| Malignant epithelial | C3 Mp ECs | HJURP | 4.7 | 0.327 | 0.018 | 2.19e-232 | 7.389999999999999e-228 |
| Malignant epithelial | C3 Mp ECs | ASPM | 4.698 | 0.371 | 0.022 | 7.52e-260 | 2.53e-255 |
| Malignant epithelial | C3 Mp ECs | CDC25C | 4.651 | 0.175 | 0.009 | 7.249999999999999e-126 | 2.44e-121 |
| Malignant epithelial | C3 Mp ECs | PIF1 | 4.645 | 0.132 | 0.006 | 3.71e-97 | 1.25e-92 |
| Malignant epithelial | C3 Mp ECs | CDCA3 | 4.582 | 0.6 | 0.041 | 0.0 | 0.0 |
| Malignant epithelial | C3 Mp ECs | PLK1 | 4.539 | 0.412 | 0.032 | 1.12e-263 | 3.77e-259 |
| Malignant epithelial | C3 Mp ECs | NEK2 | 4.524 | 0.303 | 0.017 | 2.52e-211 | 8.49e-207 |
| Malignant epithelial | C3 Mp ECs | FAM64A | 4.523 | 0.171 | 0.009 | 7.36e-120 | 2.48e-115 |
| Malignant epithelial | C3 Mp ECs | CENPE | 4.479 | 0.338 | 0.026 | 2.69e-213 | 9.07e-209 |
| Malignant epithelial | C3 Mp ECs | CCNA2 | 4.436 | 0.407 | 0.024 | 1.61e-283 | 5.42e-279 |
| Malignant epithelial | C3 Mp ECs | KIF20A | 4.427 | 0.193 | 0.011 | 3.2e-132 | 1.08e-127 |
| Malignant epithelial | C3 Mp ECs | CKAP2L | 4.423 | 0.206 | 0.01 | 9.18e-150 | 3.09e-145 |
| Malignant epithelial | C3 Mp ECs | KIF23 | 4.414 | 0.342 | 0.022 | 5.14e-232 | 1.73e-227 |
| Malignant epithelial | C3 Mp ECs | CEP55 | 4.4 | 0.483 | 0.034 | 0.0 | 0.0 |
| Malignant epithelial | C3 Mp ECs | SPAG5 | 4.293 | 0.204 | 0.013 | 7.52e-134 | 2.53e-129 |
| Malignant epithelial | C3 Mp ECs | TOP2A | 4.289 | 0.495 | 0.045 | 2.65e-299 | 8.93e-295 |
| Malignant epithelial | C3 Mp ECs | ARHGAP11A | 4.277 | 0.354 | 0.026 | 5.65e-225 | 1.9e-220 |
| Malignant epithelial | C3 Mp ECs | CDCA2 | 4.267 | 0.101 | 0.007 | 6.530000000000001e-64 | 2.2e-59 |
| Malignant epithelial | C3 Mp ECs | AURKB | 4.239 | 0.454 | 0.031 | 7.44999999999999e-304 | 2.51e-299 |
| Malignant epithelial | C3 Mp ECs | CCNB1 | 4.234 | 0.662 | 0.095 | 0.0 | 0.0 |
| Malignant epithelial | C3 Mp ECs | HMMR | 4.228 | 0.317 | 0.024 | 9.45e-198 | 3.18e-193 |
| Malignant epithelial | C3 Mp ECs | CENPF | 4.21 | 0.636 | 0.08 | 0.0 | 0.0 |
| Malignant epithelial | C3 Mp ECs | DEPDC1B | 4.175 | 0.263 | 0.016 | 2.49e-176 | 8.39e-172 |
| Malignant epithelial | C3 Mp ECs | SKA1 | 4.096 | 0.124 | 0.008 | 1.07e-80 | 3.61e-76 |
| Malignant epithelial | C3 Mp ECs | TPX2 | 4.069 | 0.67 | 0.072 | 0.0 | 0.0 |
| Malignant epithelial | C3 Mp ECs | AURKA | 4.062 | 0.477 | 0.063 | 2.68e-236 | 9.04e-232 |
| Malignant epithelial | C3 Mp ECs | NUSAP1 | 4.033 | 0.552 | 0.057 | 4.33e-319 | 1.46e-314 |
| Malignant epithelial | C3 Mp ECs | KIF15 | 4.032 | 0.134 | 0.007 | 2.3e-91 | 7.75e-87 |
| Malignant epithelial | C3 Mp ECs | NDC80 | 4.004 | 0.15 | 0.011 | 3.76e-93 | 1.27e-88 |
| Malignant epithelial | C3 Mp ECs | KIF11 | 3.99 | 0.25 | 0.016 | 4.380000000000001e-163 | 1.47e-158 |
| Malignant epithelial | C3 Mp ECs | CIT | 3.989 | 0.219 | 0.015 | 3.4e-138 | 1.15e-133 |
| Malignant epithelial | C3 Mp ECs | BUB1 | 3.953 | 0.24 | 0.015 | 4.190000000000001e-160 | 1.41e-155 |
| Malignant epithelial | C3 Mp ECs | KIF4A | 3.937 | 0.25 | 0.018 | 5.71e-158 | 1.92e-153 |
| Malignant epithelial | C3 Mp ECs | SPC25 | 3.772 | 0.375 | 0.026 | 4.4e-242 | 1.48e-237 |
| Malignant epithelial | C3 Mp ECs | CDK1 | 3.763 | 0.434 | 0.042 | 5.51e-247 | 1.86e-242 |
| Malignant epithelial | C3 Mp ECs | IQGAP3 | 3.753 | 0.228 | 0.017 | 4.43e-138 | 1.49e-133 |
| Malignant epithelial | C3 Mp ECs | PSRC1 | 3.729 | 0.159 | 0.014 | 2.03e-89 | 6.84e-85 |
| Malignant epithelial | C3 Mp ECs | CCNB2 | 3.723 | 0.596 | 0.094 | 1.86e-278 | 6.28e-274 |
| Malignant epithelial | C3 Mp ECs | CDCA8 | 3.698 | 0.261 | 0.025 | 1.51e-143 | 5.09e-139 |
| Malignant epithelial | C3 Mp ECs | NCAPG | 3.694 | 0.159 | 0.011 | 8.81e-99 | 2.97e-94 |
| Malignant epithelial | C3 Mp ECs | OIP5 | 3.686 | 0.305 | 0.031 | 1.09e-163 | 3.66e-159 |
| Malignant epithelial | C3 Mp ECs | KIF2C | 3.677 | 0.153 | 0.012 | 4.2e-92 | 1.41e-87 |
| Malignant epithelial | C3 Mp ECs | PBK | 3.666 | 0.263 | 0.019 | 1.67e-163 | 5.62e-159 |
| Malignant epithelial | C3 Mp ECs | MKI67 | 3.658 | 0.589 | 0.071 | 8.53e-311 | 2.87e-306 |
| Malignant epithelial | C3 Mp ECs | PRR11 | 3.654 | 0.301 | 0.03 | 1.13e-163 | 3.82e-159 |
| Malignant epithelial | C3 Mp ECs | PRC1 | 3.612 | 0.531 | 0.061 | 3e-286 | 1.01e-281 |
| Malignant epithelial | C3 Mp ECs | KIFC1 | 3.581 | 0.392 | 0.039 | 2.52e-216 | 8.48e-212 |
| Malignant epithelial | C3 Mp ECs | BIRC5 | 3.577 | 0.741 | 0.103 | 0.0 | 0.0 |

| compartment | subcluster | gene | avg_log2FC | pct.1 | pct.2 | p_val | p_val_adj |
|---|---|---|---|---|---|---|---|
| Malignant epithelial | C3 Mp ECs | CDKN3 | 3.566 | 0.703 | 0.115 | 0.0 | 0.0 |
| Malignant epithelial | C3 Mp ECs | FOXM1 | 3.556 | 0.487 | 0.051 | 2.39e-270 | 8.04e-266 |
| Malignant epithelial | C3 Mp ECs | KNSTRN | 3.538 | 0.463 | 0.064 | 2.86e-220 | 9.649999999999999e-216 |
| Malignant epithelial | C3 Mp ECs | BUB1B | 3.483 | 0.142 | 0.013 | 7.53e-78 | 2.54e-73 |
| Malignant epithelial | C3 Mp ECs | MXD3 | 3.473 | 0.33 | 0.037 | 8.07e-169 | 2.72e-164 |
| Malignant epithelial | C3 Mp ECs | SGOL2 | 3.473 | 0.257 | 0.029 | 1.64e-128 | 5.520000000000001e-124 |
| Malignant epithelial | C3 Mp ECs | PTTG1 | 3.467 | 0.833 | 0.237 | 3.14e-315 | 1.06e-310 |
| Malignant epithelial | C3 Mp ECs | TTK | 3.466 | 0.154 | 0.015 | 2.08e-81 | 7.01e-77 |
| Malignant epithelial | C3 Mp ECs | SHCBP1 | 3.462 | 0.123 | 0.013 | 2.79e-63 | 9.390000000000001e-59 |
| Malignant epithelial | C3 Mp ECs | CASC5 | 3.446 | 0.197 | 0.017 | 1.3e-110 | 4.37e-106 |
| Malignant epithelial | C3 Mp ECs | CDC20 | 3.388 | 0.566 | 0.085 | 3.79e-261 | 1.28e-256 |
| Malignant epithelial | C3 Mp ECs | UBE2C | 3.385 | 0.909 | 0.299 | 6.94e-313 | 2.34e-308 |
| Malignant epithelial | C3 Mp ECs | SPC24 | 3.346 | 0.183 | 0.023 | 1.22e-83 | 4.1e-79 |
| Malignant epithelial | C3 Mp ECs | NCAPH | 3.323 | 0.153 | 0.015 | 1.29e-79 | 4.36e-75 |
| Malignant epithelial | C3 Mp ECs | KIF18A | 3.3 | 0.137 | 0.011 | 6.64e-79 | 2.24e-74 |
| Malignant epithelial | C3 Mp ECs | INCENP | 3.296 | 0.116 | 0.011 | 2.19e-61 | 7.38e-57 |
| Malignant epithelial | C3 Mp ECs | KIAA1524 | 3.225 | 0.171 | 0.019 | 1.91e-85 | 6.45e-81 |
| Malignant epithelial | C3 Mp ECs | UBE2T | 3.204 | 0.648 | 0.109 | 2.21e-290 | 7.449999999999999e-286 |
| Malignant epithelial | C3 Mp ECs | RRM2 | 3.185 | 0.401 | 0.043 | 1.11e-210 | 3.75e-206 |
| Malignant epithelial | C3 Mp ECs | TACC3 | 3.111 | 0.259 | 0.032 | 5e-121 | 1.69e-116 |
| Malignant epithelial | C3 Mp ECs | MELK | 3.104 | 0.258 | 0.032 | 1.83e-118 | 6.18e-114 |
| Malignant epithelial | C3 Mp ECs | RACGAP1 | 3.085 | 0.342 | 0.044 | 4.900000000000001e-158 | 1.65e-153 |
| Malignant epithelial | C3 Mp ECs | CDCA5 | 3.08 | 0.319 | 0.037 | 1.24e-155 | 4.16e-151 |
| Malignant epithelial | C3 Mp ECs | RAD51AP1 | 3.066 | 0.389 | 0.051 | 1.07e-181 | 3.6e-177 |
| Malignant epithelial | C3 Mp ECs | TRAIP | 3.058 | 0.117 | 0.014 | 3.87e-54 | 1.3e-49 |
| Malignant epithelial | C3 Mp ECs | TROAP | 3.054 | 0.6 | 0.098 | 1.63e-263 | 5.49e-259 |
| Malignant epithelial | C3 Mp ECs | MAD2L1 | 3.045 | 0.687 | 0.125 | 4.96e-295 | 1.67e-290 |
| Malignant epithelial | C3 Mp ECs | SGOL1 | 3.042 | 0.225 | 0.025 | 1.61e-109 | 5.44e-105 |
| Malignant epithelial | C3 Mp ECs | ARHGEF39 | 3.034 | 0.204 | 0.02 | 3.68e-107 | 1.24e-102 |
| Malignant epithelial | C3 Mp ECs | MYBL2 | 3.024 | 0.277 | 0.033 | 1.48e-133 | 5e-129 |
| Malignant epithelial | C3 Mp ECs | ZWINT | 2.998 | 0.623 | 0.104 | 5.26e-272 | 1.77e-267 |
| Malignant epithelial | C3 Mp ECs | ANLN | 2.974 | 0.549 | 0.093 | 8.03e-228 | 2.71e-223 |
| Malignant epithelial | C3 Mp ECs | LMNB1 | 2.972 | 0.43 | 0.07 | 5.3e-173 | 1.78e-168 |
| Malignant epithelial | C3 Mp ECs | PKMYT1 | 2.915 | 0.236 | 0.032 | 1.08e-101 | 3.64e-97 |
| Malignant epithelial | C3 Mp ECs | UBE2S | 2.905 | 0.93 | 0.465 | 3.24e-273 | 1.09e-268 |
| Malignant epithelial | C3 Mp ECs | ARHGAP11B | 2.9 | 0.281 | 0.039 | 3.39e-121 | 1.14e-116 |
| Malignant epithelial | C3 Mp ECs | RAD51 | 2.88 | 0.189 | 0.026 | 1.22e-80 | 4.12e-76 |
| Malignant epithelial | C3 Mp ECs | NCAPD2 | 2.874 | 0.338 | 0.055 | 1.52e-131 | 5.13e-127 |
| Malignant epithelial | C3 Mp ECs | CENPM | 2.869 | 0.503 | 0.082 | 5.429999999999999e-207 | 1.83e-202 |
| Malignant epithelial | C3 Mp ECs | ASF1B | 2.861 | 0.374 | 0.047 | 1.88e-174 | 6.349999999999999e-170 |
| Malignant epithelial | C3 Mp ECs | CCNF | 2.86 | 0.121 | 0.014 | 1.37e-57 | 4.61e-53 |
| Malignant epithelial | C3 Mp ECs | CLSPN | 2.86 | 0.141 | 0.021 | 1.33e-56 | 4.48e-52 |
| Malignant epithelial | C3 Mp ECs | FAM83D | 2.841 | 0.279 | 0.048 | 5.12e-102 | 1.73e-97 |
| Malignant epithelial | C3 Mp ECs | SKA3 | 2.825 | 0.185 | 0.026 | 5.22e-78 | 1.76e-73 |
| Malignant epithelial | C3 Mp ECs | HMGB2 | 2.816 | 0.931 | 0.429 | 3.71e-281 | 1.25e-276 |
| Malignant epithelial | C3 Mp ECs | SPDL1 | 2.788 | 0.201 | 0.031 | 3.99e-79 | 1.34e-74 |
| Malignant epithelial | C3 Mp ECs | C21orf58 | 2.771 | 0.211 | 0.033 | 1.69e-82 | 5.7e-78 |
| Malignant epithelial | C3 Mp ECs | TYMS | 2.761 | 0.39 | 0.059 | 2.61e-162 | 8.779999999999999e-158 |
| Malignant epithelial | C3 Mp ECs | MND1 | 2.726 | 0.394 | 0.063 | 4.09e-158 | 1.38e-153 |
| Malignant epithelial | C3 Mp ECs | POC1A | 2.707 | 0.277 | 0.047 | 5.79e-102 | 1.95e-97 |
| Malignant epithelial | C3 Mp ECs | BORA | 2.65 | 0.131 | 0.021 | 2.62e-48 | 8.82e-44 |
| Malignant epithelial | C3 Mp ECs | RECQL4 | 2.638 | 0.19 | 0.032 | 1.69e-68 | 5.68e-64 |
| Malignant epithelial | C3 Mp ECs | TK1 | 2.637 | 0.768 | 0.185 | 1.87e-275 | 6.3e-271 |
| Malignant epithelial | C3 Mp ECs | HIST1H1D | 2.616 | 0.262 | 0.051 | 6.28e-85 | 2.11e-80 |
| Malignant epithelial | C3 Mp ECs | CENPK | 2.613 | 0.509 | 0.093 | 6.45e-192 | 2.17e-187 |
| Malignant epithelial | C3 Mp ECs | CKAP2 | 2.577 | 0.295 | 0.087 | 5.59e-64 | 1.88e-59 |
| Malignant epithelial | C3 Mp ECs | DTL | 2.575 | 0.103 | 0.015 | 6.79e-42 | 2.29e-37 |
| Malignant epithelial | C3 Mp ECs | ORC6 | 2.55 | 0.377 | 0.067 | 4.57e-136 | 1.54e-131 |
| Malignant epithelial | C3 Mp ECs | FANCI | 2.51 | 0.244 | 0.037 | 1.65e-97 | 5.57e-93 |
| Malignant epithelial | C3 Mp ECs | GINS1 | 2.496 | 0.12 | 0.019 | 5.64e-46 | 1.9e-41 |
| Malignant epithelial | C3 Mp ECs | CDC45 | 2.477 | 0.116 | 0.019 | 5.36e-43 | 1.81e-38 |
| Malignant epithelial | C3 Mp ECs | TRIP13 | 2.441 | 0.268 | 0.047 | 5.04e-95 | 1.7e-90 |
| Malignant epithelial | C3 Mp ECs | CTB-193M12.5 | 2.441 | 0.166 | 0.037 | 5.97e-46 | 2.01e-41 |
| Malignant epithelial | C3 Mp ECs | SMC4 | 2.426 | 0.734 | 0.23 | 2.37e-206 | 7.97e-202 |

| compartment | subcluster | gene | avg_log2FC | pct.1 | pct.2 | p_val | p_val_adj |
|---|---|---|---|---|---|---|---|
| Malignant epithelial | C3 Mp ECs | POLE2 | 2.377 | 0.123 | 0.023 | 3.71e-40 | 1.25e-35 |
| Malignant epithelial | C3 Mp ECs | HIST2H2AC | 2.369 | 0.417 | 0.111 | 1.21e-105 | 4.08e-101 |
| Malignant epithelial | C3 Mp ECs | KIAA0101 | 2.353 | 0.699 | 0.205 | 1.98e-194 | 6.67e-190 |
| Malignant epithelial | C3 Mp ECs | CENPO | 2.332 | 0.114 | 0.02 | 1.82e-40 | 6.14e-36 |
| Malignant epithelial | C3 Mp ECs | UHRF1 | 2.324 | 0.244 | 0.045 | 6.41e-82 | 2.16e-77 |
| Malignant epithelial | C3 Mp ECs | KPNA2 | 2.289 | 0.586 | 0.193 | 2.17e-135 | 7.3e-131 |
| Malignant epithelial | C3 Mp ECs | PLK4 | 2.289 | 0.119 | 0.017 | 9.76e-48 | 3.29e-43 |
| Malignant epithelial | C3 Mp ECs | PARPBP | 2.284 | 0.215 | 0.043 | 6.3e-66 | 2.12e-61 |
| Malignant epithelial | C3 Mp ECs | ECT2 | 2.281 | 0.357 | 0.084 | 2.38e-99 | 8.029999999999999e-95 |
| Malignant epithelial | C3 Mp ECs | TCF19 | 2.276 | 0.167 | 0.032 | 8.919999999999999e-54 | 3e-49 |
| Malignant epithelial | C3 Mp ECs | TUBA1B | 2.241 | 0.982 | 0.708 | 7.61e-273 | 2.56e-268 |
| Malignant epithelial | C3 Mp ECs | BRCA1 | 2.24 | 0.201 | 0.036 | 7.32e-69 | 2.47e-64 |
| Malignant epithelial | C3 Mp ECs | CENPN | 2.235 | 0.672 | 0.195 | 1.68e-183 | 5.650000000000001e-179 |
| Malignant epithelial | C3 Mp ECs | PHF19 | 2.193 | 0.579 | 0.18 | 1.01e-140 | 3.39e-136 |
| Malignant epithelial | C3 Mp ECs | DTYMK | 2.169 | 0.73 | 0.237 | 1.26e-191 | 4.23e-187 |
| Malignant epithelial | C3 Mp ECs | CSF2 | 2.165 | 0.156 | 0.043 | 3.41e-33 | 1.15e-28 |
| Malignant epithelial | C3 Mp ECs | CENPP | 2.142 | 0.106 | 0.026 | 3.72e-26 | 1.25e-21 |
| Malignant epithelial | C3 Mp ECs | NEMP1 | 2.128 | 0.117 | 0.025 | 4.62e-33 | 1.56e-28 |
| Malignant epithelial | C3 Mp ECs | SAPCD2 | 2.119 | 0.455 | 0.137 | 9.01e-101 | 3.03e-96 |
| Malignant epithelial | C3 Mp ECs | RP11-596C23.2 | 2.108 | 0.112 | 0.035 | 3.93e-20 | 1.32e-15 |
| Malignant epithelial | C3 Mp ECs | KNTC1 | 2.105 | 0.12 | 0.028 | 1.23e-31 | 4.160000000000001e-27 |
| Malignant epithelial | C3 Mp ECs | ATAD2 | 2.089 | 0.345 | 0.079 | 5.33e-96 | 1.8e-91 |
| Malignant epithelial | C3 Mp ECs | CDKN2D | 2.089 | 0.313 | 0.088 | 1.68e-69 | 5.66e-65 |
| Malignant epithelial | C3 Mp ECs | PSMC3IP | 2.069 | 0.119 | 0.024 | 2.84e-36 | 9.570000000000001e-32 |
| Malignant epithelial | C3 Mp ECs | VRK1 | 2.062 | 0.388 | 0.101 | 5.52e-96 | 1.86e-91 |
| Malignant epithelial | C3 Mp ECs | CNTRL | 2.06 | 0.189 | 0.042 | 1.39e-52 | 4.69e-48 |
| Malignant epithelial | C3 Mp ECs | CENPW | 2.056 | 0.88 | 0.371 | 2.56e-215 | 8.64e-211 |
| Malignant epithelial | C3 Mp ECs | IKBIP | 2.045 | 0.232 | 0.06 | 1.34e-54 | 4.51e-50 |
| Malignant epithelial | C3 Mp ECs | SMC2 | 2.033 | 0.396 | 0.109 | 3.81e-92 | 1.28e-87 |
| Malignant epithelial | C3 Mp ECs | GAS2L3 | 2.025 | 0.121 | 0.031 | 7.09e-29 | 2.39e-24 |
| Malignant epithelial | C3 Mp ECs | MNS1 | 2.02 | 0.152 | 0.032 | 3.36e-43 | 1.13e-38 |
| Malignant epithelial | C3 Mp ECs | FEN1 | 2.017 | 0.302 | 0.07 | 1.13e-80 | 3.8e-76 |
| Malignant epithelial | C3 Mp ECs | ANP32E | 1.993 | 0.564 | 0.18 | 7.71e-124 | 2.6e-119 |
| Malignant epithelial | C3 Mp ECs | CBR3 | 1.951 | 0.32 | 0.099 | 1.42e-63 | 4.79e-59 |
| Malignant epithelial | C3 Mp ECs | C16orf59 | 1.949 | 0.103 | 0.031 | 3.46e-20 | 1.16e-15 |
| Malignant epithelial | C3 Mp ECs | H2AFZ | 1.944 | 0.99 | 0.809 | 1.85e-300 | 6.23e-296 |
| Malignant epithelial | C3 Mp ECs | STMN1 | 1.942 | 0.952 | 0.498 | 1.17e-234 | 3.93e-230 |
| Malignant epithelial | C3 Mp ECs | CHTF18 | 1.942 | 0.196 | 0.052 | 2.99e-44 | 1.01e-39 |
| Malignant epithelial | C3 Mp ECs | GINS4 | 1.937 | 0.175 | 0.041 | 6.78e-45 | 2.28e-40 |
| Malignant epithelial | C3 Mp ECs | LMNB2 | 1.906 | 0.468 | 0.144 | 3.7e-99 | 1.25e-94 |
| Malignant epithelial | C3 Mp ECs | CDC6 | 1.904 | 0.141 | 0.034 | 4.65e-35 | 1.57e-30 |
| Malignant epithelial | C3 Mp ECs | CCDC18 | 1.897 | 0.105 | 0.023 | 4.05e-29 | 1.36e-24 |
| Malignant epithelial | C3 Mp ECs | PHLDB2 | 1.894 | 0.102 | 0.023 | 3.850000000000001e-27 | 1.3e-22 |
| Malignant epithelial | C3 Mp ECs | DSCC1 | 1.883 | 0.116 | 0.028 | 3.12e-29 | 1.05e-24 |
| Malignant epithelial | C3 Mp ECs | KIF22 | 1.871 | 0.473 | 0.152 | 1.98e-96 | 6.68e-92 |
| Malignant epithelial | C3 Mp ECs | CKAP5 | 1.869 | 0.279 | 0.085 | 6.01e-54 | 2.02e-49 |
| Malignant epithelial | C3 Mp ECs | SKA2 | 1.851 | 0.655 | 0.233 | 3.01e-134 | 1.02e-129 |
| Malignant epithelial | C3 Mp ECs | ZNF695 | 1.85 | 0.101 | 0.025 | 1.17e-24 | 3.95e-20 |
| Malignant epithelial | C3 Mp ECs | ATAD5 | 1.848 | 0.19 | 0.049 | 1.7e-43 | 5.72e-39 |
| Malignant epithelial | C3 Mp ECs | TIMELESS | 1.841 | 0.138 | 0.031 | 9.16e-37 | 3.09e-32 |
| Malignant epithelial | C3 Mp ECs | RP11-800A18.4 | 1.831 | 0.131 | 0.027 | 1.56e-38 | 5.26e-34 |
| Malignant epithelial | C3 Mp ECs | NCAPG2 | 1.831 | 0.126 | 0.029 | 6.25e-33 | 2.11e-28 |
| Malignant epithelial | C3 Mp ECs | MCM8 | 1.824 | 0.145 | 0.04 | 1.16e-30 | 3.92e-26 |
| Malignant epithelial | C3 Mp ECs | RHNO1 | 1.818 | 0.508 | 0.174 | 3.27e-97 | 1.1e-92 |
| Malignant epithelial | C3 Mp ECs | RP11-160O5.1 | 1.818 | 0.11 | 0.029 | 1.15e-24 | 3.86e-20 |
| Malignant epithelial | C3 Mp ECs | CEP152 | 1.808 | 0.153 | 0.039 | 6.1e-36 | 2.06e-31 |
| Malignant epithelial | C3 Mp ECs | DHFR | 1.806 | 0.452 | 0.148 | 8.640000000000001e-88 | 2.91e-83 |
| Malignant epithelial | C3 Mp ECs | CENPU | 1.787 | 0.182 | 0.046 | 1.07e-42 | 3.61e-38 |
| Malignant epithelial | C3 Mp ECs | TEX30 | 1.774 | 0.331 | 0.102 | 6.26e-64 | 2.11e-59 |
| Malignant epithelial | C3 Mp ECs | CH17-373J23.1 | 1.773 | 0.121 | 0.043 | 3.95e-18 | 1.33e-13 |
| Malignant epithelial | C3 Mp ECs | RNASEH2A | 1.77 | 0.49 | 0.164 | 8.89e-94 | 3e-89 |
| Malignant epithelial | C3 Mp ECs | TUBG1 | 1.767 | 0.371 | 0.116 | 1.95e-72 | 6.56e-68 |
| Malignant epithelial | C3 Mp ECs | LINC01468 | 1.767 | 0.317 | 0.084 | 6.760000000000001e-72 | 2.28e-67 |
| Malignant epithelial | C3 Mp ECs | HAS3 | 1.765 | 0.141 | 0.044 | 1.25e-25 | 4.2e-21 |

| compartment | subcluster | gene | avg_log2FC | pct.1 | pct.2 | p_val | p_val_adj |
|---|---|---|---|---|---|---|---|
| Malignant epithelial | C3 Mp ECs | CCDC74A | 1.754 | 0.19 | 0.051 | 1.71e-41 | 5.780000000000001e-37 |
| Malignant epithelial | C3 Mp ECs | MIS18BP1 | 1.753 | 0.345 | 0.115 | 6.44e-61 | 2.17e-56 |
| Malignant epithelial | C3 Mp ECs | HMGB3 | 1.745 | 0.673 | 0.279 | 2.8e-118 | 9.43e-114 |
| Malignant epithelial | C3 Mp ECs | RFC5 | 1.741 | 0.225 | 0.068 | 7.920000000000001e-43 | 2.67e-38 |
| Malignant epithelial | C3 Mp ECs | FANCB | 1.738 | 0.15 | 0.039 | 1.32e-34 | 4.45e-30 |
| Malignant epithelial | C3 Mp ECs | CHAF1A | 1.729 | 0.276 | 0.081 | 4.11e-55 | 1.38e-50 |
| Malignant epithelial | C3 Mp ECs | CKS2 | 1.724 | 0.939 | 0.683 | 5.55e-139 | 1.87e-134 |
| Malignant epithelial | C3 Mp ECs | E2F1 | 1.722 | 0.113 | 0.035 | 1.75e-20 | 5.88e-16 |
| Malignant epithelial | C3 Mp ECs | CEP70 | 1.717 | 0.192 | 0.062 | 1.45e-33 | 4.9e-29 |
| Malignant epithelial | C3 Mp ECs | TMPO-AS1 | 1.707 | 0.114 | 0.028 | 1.44e-28 | 4.84e-24 |
| Malignant epithelial | C3 Mp ECs | NCAPD3 | 1.702 | 0.185 | 0.055 | 6.31e-36 | 2.13e-31 |
| Malignant epithelial | C3 Mp ECs | PGP | 1.699 | 0.652 | 0.269 | 1.63e-114 | 5.49e-110 |
| Malignant epithelial | C3 Mp ECs | GINS2 | 1.691 | 0.317 | 0.099 | 1.29e-59 | 4.33e-55 |
| Malignant epithelial | C3 Mp ECs | TRIM59 | 1.687 | 0.134 | 0.038 | 1.07e-27 | 3.59e-23 |
| Malignant epithelial | C3 Mp ECs | CENPH | 1.681 | 0.385 | 0.126 | 5.35e-69 | 1.8e-64 |
| Malignant epithelial | C3 Mp ECs | TNFAIP8L1 | 1.681 | 0.183 | 0.06 | 1.37e-31 | 4.61e-27 |
| Malignant epithelial | C3 Mp ECs | FANCG | 1.659 | 0.143 | 0.043 | 2.27e-27 | 7.65e-23 |
| Malignant epithelial | C3 Mp ECs | DNMT1 | 1.653 | 0.508 | 0.187 | 2.18e-86 | 7.36e-82 |
| Malignant epithelial | C3 Mp ECs | MCM4 | 1.647 | 0.41 | 0.151 | 9.72e-65 | 3.28e-60 |
| Malignant epithelial | C3 Mp ECs | LRR1 | 1.646 | 0.212 | 0.069 | 7.100000000000001e-36 | 2.39e-31 |
| Malignant epithelial | C3 Mp ECs | DLEU2 | 1.644 | 0.132 | 0.037 | 1.91e-27 | 6.44e-23 |
| Malignant epithelial | C3 Mp ECs | PRKCDBP | 1.632 | 0.894 | 0.54 | 2.17e-134 | 7.31e-130 |
| Malignant epithelial | C3 Mp ECs | CDCA4 | 1.632 | 0.332 | 0.11 | 2.27e-58 | 7.63e-54 |
| Malignant epithelial | C3 Mp ECs | RFC3 | 1.631 | 0.143 | 0.046 | 1.59e-25 | 5.370000000000001e-21 |
| Malignant epithelial | C3 Mp ECs | REEP4 | 1.628 | 0.305 | 0.103 | 1.71e-51 | 5.78e-47 |
| Malignant epithelial | C3 Mp ECs | CDT1 | 1.627 | 0.352 | 0.116 | 9.529999999999999e-61 | 3.21e-56 |
| Malignant epithelial | C3 Mp ECs | NRM | 1.626 | 0.312 | 0.098 | 1.95e-57 | 6.57e-53 |
| Malignant epithelial | C3 Mp ECs | HIST1H4C | 1.625 | 0.716 | 0.41 | 2.85e-80 | 9.62e-76 |
| Malignant epithelial | C3 Mp ECs | MIS18A | 1.624 | 0.321 | 0.104 | 8.500000000000001e-57 | 2.87e-52 |
| Malignant epithelial | C3 Mp ECs | CKS1B | 1.619 | 0.901 | 0.586 | 1.03e-160 | 3.48e-156 |
| Malignant epithelial | C3 Mp ECs | CHEK1 | 1.617 | 0.281 | 0.09 | 1.47e-49 | 4.94e-45 |
| Malignant epithelial | C3 Mp ECs | ODF2 | 1.604 | 0.21 | 0.069 | 1.21e-35 | 4.070000000000001e-31 |
| Malignant epithelial | C3 Mp ECs | HELLS | 1.602 | 0.393 | 0.128 | 9.17e-71 | 3.09e-66 |
| Malignant epithelial | C3 Mp ECs | NUCKS1 | 1.601 | 0.952 | 0.662 | 2.18e-168 | 7.36e-164 |
| Malignant epithelial | C3 Mp ECs | C14orf80 | 1.586 | 0.345 | 0.109 | 3.49e-63 | 1.18e-58 |
| Malignant epithelial | C3 Mp ECs | CHAC2 | 1.586 | 0.146 | 0.044 | 2.03e-27 | 6.83e-23 |
| Malignant epithelial | C3 Mp ECs | HIST1H1E | 1.586 | 0.145 | 0.046 | 8.75e-26 | 2.95e-21 |
| Malignant epithelial | C3 Mp ECs | CENPL | 1.551 | 0.139 | 0.042 | 3.31e-26 | 1.11e-21 |
| Malignant epithelial | C3 Mp ECs | SCLT1 | 1.538 | 0.294 | 0.084 | 3.53e-61 | 1.19e-56 |
| Malignant epithelial | C3 Mp ECs | TMPO | 1.537 | 0.534 | 0.214 | 1.94e-79 | 6.53e-75 |
| Malignant epithelial | C3 Mp ECs | EMC9 | 1.534 | 0.452 | 0.175 | 9.63e-68 | 3.25e-63 |
| Malignant epithelial | C3 Mp ECs | CCDC88A | 1.531 | 0.112 | 0.03 | 5.83e-25 | 1.97e-20 |
| Malignant epithelial | C3 Mp ECs | PCNA | 1.527 | 0.63 | 0.311 | 1.77e-77 | 5.97e-73 |
| Malignant epithelial | C3 Mp ECs | MCM2 | 1.523 | 0.19 | 0.062 | 3.86e-32 | 1.3e-27 |
| Malignant epithelial | C3 Mp ECs | C16orf95 | 1.517 | 0.13 | 0.043 | 1.98e-21 | 6.66e-17 |
| Malignant epithelial | C3 Mp ECs | ARL6IP1 | 1.514 | 0.908 | 0.706 | 6.66e-101 | 2.25e-96 |
| Malignant epithelial | C3 Mp ECs | B3GNTL1 | 1.495 | 0.172 | 0.06 | 3.58e-26 | 1.21e-21 |
| Malignant epithelial | C3 Mp ECs | TUBB | 1.478 | 0.939 | 0.646 | 8.28e-151 | 2.79e-146 |
| Malignant epithelial | C3 Mp ECs | RNF26 | 1.465 | 0.36 | 0.14 | 3.94e-49 | 1.33e-44 |
| Malignant epithelial | C3 Mp ECs | ANPEP | 1.464 | 0.337 | 0.124 | 1.2e-47 | 4.04e-43 |
| Malignant epithelial | C3 Mp ECs | CEP57L1 | 1.462 | 0.113 | 0.038 | 1.99e-18 | 6.7e-14 |
| Malignant epithelial | C3 Mp ECs | CHKB-AS1 | 1.46 | 0.139 | 0.044 | 2.09e-24 | 7.03e-20 |
| Malignant epithelial | C3 Mp ECs | RFC4 | 1.434 | 0.214 | 0.077 | 1.21e-30 | 4.09e-26 |
| Malignant epithelial | C3 Mp ECs | ZWILCH | 1.431 | 0.188 | 0.067 | 3.850000000000001e-27 | 1.3e-22 |
| Malignant epithelial | C3 Mp ECs | DDX11 | 1.426 | 0.234 | 0.083 | 2.92e-34 | 9.829999999999999e-30 |
| Malignant epithelial | C3 Mp ECs | MPHOSPH9 | 1.425 | 0.119 | 0.038 | 1.09e-20 | 3.67e-16 |
| Malignant epithelial | C3 Mp ECs | CBX5 | 1.424 | 0.47 | 0.201 | 5.52e-60 | 1.86e-55 |
| Malignant epithelial | C3 Mp ECs | POLE | 1.416 | 0.113 | 0.034 | 4.71e-21 | 1.59e-16 |
| Malignant epithelial | C3 Mp ECs | MZT1 | 1.412 | 0.549 | 0.257 | 5.690000000000001e-66 | 1.92e-61 |
| Malignant epithelial | C3 Mp ECs | MESP1 | 1.409 | 0.459 | 0.188 | 8.39e-60 | 2.83e-55 |
| Malignant epithelial | C3 Mp ECs | H2AFX | 1.404 | 0.64 | 0.302 | 1.35e-80 | 4.54e-76 |
| Malignant epithelial | C3 Mp ECs | CALB2 | 1.403 | 0.575 | 0.239 | 4.33e-79 | 1.46e-74 |
| Malignant epithelial | C3 Mp ECs | C9orf40 | 1.402 | 0.199 | 0.074 | 6.440000000000001e-27 | 2.17e-22 |
| Malignant epithelial | C3 Mp ECs | NKILA | 1.397 | 0.199 | 0.075 | 5.36e-26 | 1.81e-21 |

| compartment | subcluster | gene | avg_log2FC | pct.1 | pct.2 | p_val | p_val_adj |
|---|---|---|---|---|---|---|---|
| Malignant epithelial | C3 Mp ECs | CEP78 | 1.388 | 0.13 | 0.041 | 9.469999999999999e-23 | 3.19e-18 |
| Malignant epithelial | C3 Mp ECs | RP11-588G21.2 | 1.383 | 0.142 | 0.052 | 7.48e-20 | 2.52e-15 |
| Malignant epithelial | C3 Mp ECs | MT2A | 1.382 | 0.917 | 0.684 | 3.75e-88 | 1.26e-83 |
| Malignant epithelial | C3 Mp ECs | NUDT1 | 1.38 | 0.719 | 0.339 | 2.38e-99 | 8.019999999999999e-95 |
| Malignant epithelial | C3 Mp ECs | RANGAP1 | 1.377 | 0.389 | 0.16 | 5.11e-50 | 1.72e-45 |
| Malignant epithelial | C3 Mp ECs | CHST14 | 1.376 | 0.103 | 0.039 | 7.27e-14 | 2.45e-09 |
| Malignant epithelial | C3 Mp ECs | DONSON | 1.375 | 0.126 | 0.045 | 1.39e-18 | 4.69e-14 |
| Malignant epithelial | C3 Mp ECs | CNTLN | 1.374 | 0.177 | 0.06 | 1.48e-27 | 4.99e-23 |
| Malignant epithelial | C3 Mp ECs | LINC00973 | 1.371 | 0.108 | 0.041 | 2.79e-14 | 9.4e-10 |
| Malignant epithelial | C3 Mp ECs | NDE1 | 1.369 | 0.277 | 0.108 | 3.7e-36 | 1.25e-31 |
| Malignant epithelial | C3 Mp ECs | GRK6 | 1.363 | 0.23 | 0.094 | 4.9e-27 | 1.65e-22 |
| Malignant epithelial | C3 Mp ECs | FAM216A | 1.36 | 0.139 | 0.047 | 9.98e-22 | 3.36e-17 |
| Malignant epithelial | C3 Mp ECs | TNS4 | 1.36 | 0.163 | 0.065 | 1.18e-19 | 3.96e-15 |
| Malignant epithelial | C3 Mp ECs | PRIM1 | 1.357 | 0.193 | 0.069 | 8.349999999999999e-28 | 2.81e-23 |
| Malignant epithelial | C3 Mp ECs | AXL | 1.341 | 0.434 | 0.192 | 8.110000000000001e-47 | 2.73e-42 |
| Malignant epithelial | C3 Mp ECs | DBF4 | 1.337 | 0.281 | 0.112 | 2.27e-35 | 7.64e-31 |
| Malignant epithelial | C3 Mp ECs | TRIM7 | 1.336 | 0.181 | 0.064 | 1.89e-26 | 6.38e-22 |
| Malignant epithelial | C3 Mp ECs | DDX39A | 1.332 | 0.697 | 0.369 | 3.74e-82 | 1.26e-77 |
| Malignant epithelial | C3 Mp ECs | FAM111A | 1.326 | 0.323 | 0.13 | 1.25e-39 | 4.22e-35 |
| Malignant epithelial | C3 Mp ECs | DNAJC9 | 1.325 | 0.626 | 0.305 | 2.4e-72 | 8.08e-68 |
| Malignant epithelial | C3 Mp ECs | PSIP1 | 1.315 | 0.589 | 0.267 | 6.14e-70 | 2.07e-65 |
| Malignant epithelial | C3 Mp ECs | LIG1 | 1.315 | 0.161 | 0.061 | 9.59e-22 | 3.23e-17 |
| Malignant epithelial | C3 Mp ECs | SNRNP25 | 1.31 | 0.53 | 0.234 | 2.44e-64 | 8.22e-60 |
| Malignant epithelial | C3 Mp ECs | NAV2 | 1.309 | 0.109 | 0.037 | 3.36e-17 | 1.13e-12 |
| Malignant epithelial | C3 Mp ECs | CD3EAP | 1.305 | 0.361 | 0.146 | 1.38e-44 | 4.65e-40 |
| Malignant epithelial | C3 Mp ECs | MCM5 | 1.303 | 0.279 | 0.115 | 7.9e-33 | 2.66e-28 |
| Malignant epithelial | C3 Mp ECs | WHSC1 | 1.301 | 0.21 | 0.071 | 1.13e-32 | 3.82e-28 |
| Malignant epithelial | C3 Mp ECs | GMNN | 1.298 | 0.244 | 0.09 | 6.67e-34 | 2.25e-29 |
| Malignant epithelial | C3 Mp ECs | TMEM237 | 1.298 | 0.154 | 0.056 | 1.84e-21 | 6.2e-17 |
| Malignant epithelial | C3 Mp ECs | TNC | 1.297 | 0.103 | 0.039 | 4.82e-14 | 1.62e-09 |
| Malignant epithelial | C3 Mp ECs | TTF2 | 1.293 | 0.108 | 0.036 | 4.07e-17 | 1.37e-12 |
| Malignant epithelial | C3 Mp ECs | EFCAB2 | 1.292 | 0.15 | 0.052 | 1.7e-22 | 5.74e-18 |
| Malignant epithelial | C3 Mp ECs | H2AFV | 1.286 | 0.952 | 0.702 | 9.930000000000001e-157 | 3.35e-152 |
| Malignant epithelial | C3 Mp ECs | CDK5RAP2 | 1.285 | 0.228 | 0.087 | 2.39e-29 | 8.07e-25 |
| Malignant epithelial | C3 Mp ECs | BUB3 | 1.283 | 0.748 | 0.439 | 1.31e-77 | 4.410000000000001e-73 |
| Malignant epithelial | C3 Mp ECs | LGALS1 | 1.274 | 0.939 | 0.743 | 2.4e-84 | 8.09e-80 |
| Malignant epithelial | C3 Mp ECs | IL1R2 | 1.27 | 0.221 | 0.079 | 2.17e-31 | 7.310000000000001e-27 |
| Malignant epithelial | C3 Mp ECs | TUBB4B | 1.269 | 0.964 | 0.802 | 9.22e-100 | 3.11e-95 |
| Malignant epithelial | C3 Mp ECs | UBALD2 | 1.263 | 0.687 | 0.383 | 4.25e-66 | 1.43e-61 |
| Malignant epithelial | C3 Mp ECs | WWTR1 | 1.263 | 0.248 | 0.1 | 7.62e-29 | 2.57e-24 |
| Malignant epithelial | C3 Mp ECs | RCCD1 | 1.254 | 0.244 | 0.099 | 1.35e-28 | 4.56e-24 |
| Malignant epithelial | C3 Mp ECs | HMGN2 | 1.251 | 0.949 | 0.741 | 3.81e-144 | 1.28e-139 |
| Malignant epithelial | C3 Mp ECs | RANBP1 | 1.247 | 0.941 | 0.716 | 2.72e-138 | 9.15e-134 |
| Malignant epithelial | C3 Mp ECs | RAC3 | 1.245 | 0.11 | 0.041 | 4.9e-15 | 1.65e-10 |
| Malignant epithelial | C3 Mp ECs | MAGOHB | 1.237 | 0.615 | 0.315 | 3.33e-63 | 1.12e-58 |
| Malignant epithelial | C3 Mp ECs | NMU | 1.233 | 0.229 | 0.088 | 1e-29 | 3.37e-25 |
| Malignant epithelial | C3 Mp ECs | EFCAB11 | 1.228 | 0.161 | 0.056 | 3.7e-24 | 1.25e-19 |
| Malignant epithelial | C3 Mp ECs | RMI2 | 1.227 | 0.196 | 0.073 | 6.56e-26 | 2.21e-21 |
| Malignant epithelial | C3 Mp ECs | AIM2 | 1.221 | 0.137 | 0.054 | 1.8e-16 | 6.06e-12 |
| Malignant epithelial | C3 Mp ECs | CCL5 | 1.217 | 0.352 | 0.224 | 3.42e-13 | 1.15e-08 |
| Malignant epithelial | C3 Mp ECs | PTMS | 1.216 | 0.981 | 0.819 | 2.98e-136 | 1e-131 |
| Malignant epithelial | C3 Mp ECs | SAC3D1 | 1.216 | 0.408 | 0.186 | 6.36e-42 | 2.14e-37 |
| Malignant epithelial | C3 Mp ECs | GGT5 | 1.216 | 0.255 | 0.094 | 6.5e-35 | 2.19e-30 |
| Malignant epithelial | C3 Mp ECs | KMT5A | 1.216 | 0.279 | 0.12 | 2.53e-29 | 8.53e-25 |
| Malignant epithelial | C3 Mp ECs | AC009005.2 | 1.216 | 0.128 | 0.043 | 1.13e-20 | 3.79e-16 |
| Malignant epithelial | C3 Mp ECs | HN1 | 1.215 | 0.96 | 0.791 | 2.27e-125 | 7.650000000000001e-121 |
| Malignant epithelial | C3 Mp ECs | PCBD2 | 1.215 | 0.415 | 0.183 | 9.27e-47 | 3.13e-42 |
| Malignant epithelial | C3 Mp ECs | CHRNA5 | 1.215 | 0.123 | 0.052 | 1.05e-13 | 3.52e-09 |
| Malignant epithelial | C3 Mp ECs | CAV1 | 1.205 | 0.843 | 0.485 | 5.94e-92 | 2e-87 |
| Malignant epithelial | C3 Mp ECs | POLR3G | 1.203 | 0.124 | 0.045 | 2.03e-17 | 6.82e-13 |
| Malignant epithelial | C3 Mp ECs | SPA17 | 1.201 | 0.204 | 0.084 | 5.7e-23 | 1.92e-18 |
| Malignant epithelial | C3 Mp ECs | RAD21 | 1.195 | 0.728 | 0.424 | 2.68e-69 | 9.04e-65 |
| Malignant epithelial | C3 Mp ECs | GZMA | 1.192 | 0.174 | 0.095 | 4.01e-10 | 1.35e-05 |
| Malignant epithelial | C3 Mp ECs | LBR | 1.185 | 0.647 | 0.351 | 5.52e-62 | 1.86e-57 |

| compartment | subcluster | gene | avg_log2FC | pct.1 | pct.2 | p_val | p_val_adj |
|---|---|---|---|---|---|---|---|
| Malignant epithelial | C3 Mp ECs | GOT1 | 1.183 | 0.354 | 0.163 | 2.34e-35 | 7.9e-31 |
| Malignant epithelial | C3 Mp ECs | TM4SF18 | 1.18 | 0.49 | 0.217 | 3.86e-53 | 1.3e-48 |
| Malignant epithelial | C3 Mp ECs | CFLAR-AS1 | 1.178 | 0.124 | 0.065 | 6.77e-09 | 0.000228 |
| Malignant epithelial | C3 Mp ECs | PLAU | 1.176 | 0.764 | 0.403 | 1.78e-82 | 5.99e-78 |
| Malignant epithelial | C3 Mp ECs | NUP37 | 1.174 | 0.539 | 0.283 | 2.77e-49 | 9.339999999999999e-45 |
| Malignant epithelial | C3 Mp ECs | NUDCD2 | 1.173 | 0.646 | 0.366 | 3.48e-59 | 1.17e-54 |
| Malignant epithelial | C3 Mp ECs | OVOS2 | 1.17 | 0.141 | 0.049 | 5.27e-21 | 1.77e-16 |
| Malignant epithelial | C3 Mp ECs | APITD1 | 1.17 | 0.166 | 0.069 | 4.89e-18 | 1.65e-13 |
| Malignant epithelial | C3 Mp ECs | ALDH1A3 | 1.166 | 0.332 | 0.127 | 4.16e-43 | 1.4e-38 |
| Malignant epithelial | C3 Mp ECs | CAD | 1.165 | 0.143 | 0.054 | 1.22e-18 | 4.11e-14 |
| Malignant epithelial | C3 Mp ECs | UCP2 | 1.164 | 0.374 | 0.165 | 1.61e-39 | 5.42e-35 |
| Malignant epithelial | C3 Mp ECs | ARL4C | 1.162 | 0.633 | 0.332 | 1.76e-59 | 5.939999999999999e-55 |
| Malignant epithelial | C3 Mp ECs | KRR1 | 1.161 | 0.669 | 0.346 | 2.6e-67 | 8.75e-63 |
| Malignant epithelial | C3 Mp ECs | MCM3 | 1.161 | 0.306 | 0.131 | 2.05e-33 | 6.91e-29 |
| Malignant epithelial | C3 Mp ECs | FOSL1 | 1.16 | 0.726 | 0.409 | 2.59e-69 | 8.71e-65 |
| Malignant epithelial | C3 Mp ECs | CXCL3 | 1.159 | 0.855 | 0.685 | 4.78e-57 | 1.61e-52 |
| Malignant epithelial | C3 Mp ECs | DUT | 1.157 | 0.767 | 0.561 | 2.83e-50 | 9.52e-46 |
| Malignant epithelial | C3 Mp ECs | SUN2 | 1.152 | 0.393 | 0.185 | 5e-38 | 1.69e-33 |
| Malignant epithelial | C3 Mp ECs | MCM6 | 1.15 | 0.134 | 0.052 | 1.13e-16 | 3.81e-12 |
| Malignant epithelial | C3 Mp ECs | NUP93 | 1.147 | 0.295 | 0.124 | 6.8e-33 | 2.29e-28 |
| Malignant epithelial | C3 Mp ECs | RRM1 | 1.143 | 0.241 | 0.095 | 2.98e-29 | 1e-24 |
| Malignant epithelial | C3 Mp ECs | SRGAP2C | 1.141 | 0.131 | 0.046 | 4.47e-19 | 1.51e-14 |
| Malignant epithelial | C3 Mp ECs | F2R | 1.141 | 0.106 | 0.038 | 3.59e-15 | 1.21e-10 |
| Malignant epithelial | C3 Mp ECs | RPA3 | 1.136 | 0.906 | 0.632 | 9.170000000000001e-117 | 3.09e-112 |
| Malignant epithelial | C3 Mp ECs | ALYREF | 1.136 | 0.237 | 0.095 | 5.62e-28 | 1.89e-23 |
| Malignant epithelial | C3 Mp ECs | CENPQ | 1.136 | 0.116 | 0.048 | 4.6e-13 | 1.55e-08 |
| Malignant epithelial | C3 Mp ECs | HIRIP3 | 1.127 | 0.179 | 0.077 | 1.5e-18 | 5.05e-14 |
| Malignant epithelial | C3 Mp ECs | MSN | 1.124 | 0.183 | 0.074 | 4.18e-21 | 1.41e-16 |
| Malignant epithelial | C3 Mp ECs | C11orf24 | 1.12 | 0.243 | 0.106 | 2.98e-24 | 1.01e-19 |
| Malignant epithelial | C3 Mp ECs | TCOF1 | 1.119 | 0.383 | 0.181 | 1.42e-35 | 4.78e-31 |
| Malignant epithelial | C3 Mp ECs | TDP1 | 1.118 | 0.13 | 0.053 | 5.81e-15 | 1.96e-10 |
| Malignant epithelial | C3 Mp ECs | TM4SF1-AS1 | 1.117 | 0.246 | 0.129 | 2.99e-16 | 1.01e-11 |
| Malignant epithelial | C3 Mp ECs | NDC1 | 1.113 | 0.138 | 0.055 | 1.81e-16 | 6.1e-12 |
| Malignant epithelial | C3 Mp ECs | RRP9 | 1.109 | 0.488 | 0.259 | 3.03e-38 | 1.02e-33 |
| Malignant epithelial | C3 Mp ECs | DNAAF2 | 1.108 | 0.543 | 0.282 | 8.71e-46 | 2.94e-41 |
| Malignant epithelial | C3 Mp ECs | XPO7 | 1.107 | 0.105 | 0.045 | 3.3e-11 | 1.11e-06 |
| Malignant epithelial | C3 Mp ECs | TFDP1 | 1.101 | 0.443 | 0.214 | 1.03e-40 | 3.46e-36 |
| Malignant epithelial | C3 Mp ECs | KDELC2 | 1.101 | 0.146 | 0.067 | 1.62e-13 | 5.45e-09 |
| Malignant epithelial | C3 Mp ECs | DSN1 | 1.096 | 0.189 | 0.076 | 5.78e-22 | 1.95e-17 |
| Malignant epithelial | C3 Mp ECs | MRPL51 | 1.093 | 0.97 | 0.829 | 3.44e-152 | 1.16e-147 |
| Malignant epithelial | C3 Mp ECs | CD3E | 1.093 | 0.112 | 0.052 | 2.92e-10 | 9.86e-06 |
| Malignant epithelial | C3 Mp ECs | CXCL1 | 1.086 | 0.804 | 0.543 | 2.26e-61 | 7.600000000000001e-57 |
| Malignant epithelial | C3 Mp ECs | MTAP | 1.082 | 0.63 | 0.359 | 3.76e-50 | 1.27e-45 |
| Malignant epithelial | C3 Mp ECs | PIK3AP1 | 1.08 | 0.127 | 0.058 | 6.95e-12 | 2.34e-07 |
| Malignant epithelial | C3 Mp ECs | S100A2 | 1.079 | 0.89 | 0.625 | 3.96e-74 | 1.33e-69 |
| Malignant epithelial | C3 Mp ECs | FANCA | 1.076 | 0.186 | 0.082 | 3.33e-18 | 1.12e-13 |
| Malignant epithelial | C3 Mp ECs | BAG2 | 1.076 | 0.152 | 0.071 | 3.07e-13 | 1.03e-08 |
| Malignant epithelial | C3 Mp ECs | G2E3 | 1.075 | 0.178 | 0.086 | 8.39e-15 | 2.83e-10 |
| Malignant epithelial | C3 Mp ECs | UNG | 1.074 | 0.273 | 0.155 | 7.66e-16 | 2.58e-11 |
| Malignant epithelial | C3 Mp ECs | CNIH4 | 1.073 | 0.84 | 0.583 | 3.6e-79 | 1.21e-74 |
| Malignant epithelial | C3 Mp ECs | RFC2 | 1.072 | 0.33 | 0.163 | 2.06e-27 | 6.929999999999999e-23 |
| Malignant epithelial | C3 Mp ECs | STEAP1 | 1.07 | 0.757 | 0.468 | 1.06e-59 | 3.59e-55 |
| Malignant epithelial | C3 Mp ECs | BCL2L12 | 1.07 | 0.498 | 0.246 | 2.61e-45 | 8.79e-41 |
| Malignant epithelial | C3 Mp ECs | CTPS1 | 1.065 | 0.101 | 0.047 | 4.51e-09 | 0.000152 |
| Malignant epithelial | C3 Mp ECs | CEP250 | 1.063 | 0.157 | 0.069 | 7.93e-16 | 2.67e-11 |
| Malignant epithelial | C3 Mp ECs | CCDC85C | 1.062 | 0.156 | 0.063 | 4.24e-18 | 1.43e-13 |
| Malignant epithelial | C3 Mp ECs | RAD51C | 1.058 | 0.246 | 0.112 | 2.66e-22 | 8.96e-18 |
| Malignant epithelial | C3 Mp ECs | H1FX | 1.054 | 0.701 | 0.472 | 9.61e-41 | 3.24e-36 |
| Malignant epithelial | C3 Mp ECs | CSE1L | 1.053 | 0.477 | 0.237 | 5.55e-41 | 1.87e-36 |
| Malignant epithelial | C3 Mp ECs | CDC27 | 1.05 | 0.23 | 0.109 | 3.68e-19 | 1.24e-14 |
| Malignant epithelial | C3 Mp ECs | PCNT | 1.048 | 0.124 | 0.052 | 1.22e-13 | 4.12e-09 |
| Malignant epithelial | C3 Mp ECs | CTSC | 1.046 | 0.905 | 0.62 | 1.7e-83 | 5.720000000000001e-79 |
| Malignant epithelial | C3 Mp ECs | CCSAP | 1.046 | 0.113 | 0.048 | 2.65e-12 | 8.94e-08 |
| Malignant epithelial | C3 Mp ECs | SRGAP2 | 1.042 | 0.171 | 0.074 | 4.48e-17 | 1.51e-12 |

| compartment | subcluster | gene | avg_log2FC | pct.1 | pct.2 | p_val | p_val_adj |
|---|---|---|---|---|---|---|---|
| Malignant epithelial | C3 Mp ECs | RP11-758H9.2 | 1.039 | 0.241 | 0.13 | 2.19e-15 | 7.39e-11 |
| Malignant epithelial | C3 Mp ECs | RFWD3 | 1.037 | 0.143 | 0.063 | 3.39e-14 | 1.14e-09 |
| Malignant epithelial | C3 Mp ECs | GTF2A2 | 1.036 | 0.799 | 0.53 | 1.57e-70 | 5.29e-66 |
| Malignant epithelial | C3 Mp ECs | DLGAP1-AS5 | 1.036 | 0.157 | 0.069 | 1.71e-15 | 5.77e-11 |
| Malignant epithelial | C3 Mp ECs | LSM5 | 1.031 | 0.986 | 0.84 | 6.340000000000001e-156 | 2.14e-151 |
| Malignant epithelial | C3 Mp ECs | VIM | 1.028 | 0.421 | 0.278 | 1.22e-15 | 4.12e-11 |
| Malignant epithelial | C3 Mp ECs | GAS2L1 | 1.023 | 0.192 | 0.088 | 1.55e-17 | 5.24e-13 |
| Malignant epithelial | C3 Mp ECs | USP18 | 1.019 | 0.215 | 0.088 | 4.25e-24 | 1.43e-19 |
| Malignant epithelial | C3 Mp ECs | SIVA1 | 1.018 | 0.923 | 0.719 | 2.66e-97 | 8.95e-93 |
| Malignant epithelial | C3 Mp ECs | RUVBL2 | 1.017 | 0.681 | 0.405 | 3.44e-56 | 1.16e-51 |
| Malignant epithelial | C3 Mp ECs | RNASEH2C | 1.016 | 0.801 | 0.515 | 2.36e-71 | 7.97e-67 |
| Malignant epithelial | C3 Mp ECs | MYEF2 | 1.016 | 0.211 | 0.093 | 1.45e-20 | 4.89e-16 |
| Malignant epithelial | C3 Mp ECs | METTL10 | 1.015 | 0.324 | 0.153 | 1.13e-28 | 3.8e-24 |
| Malignant epithelial | C3 Mp ECs | CXCL2 | 1.012 | 0.797 | 0.639 | 1.25e-41 | 4.220000000000001e-37 |
| Malignant epithelial | C3 Mp ECs | TSPAN4 | 1.011 | 0.279 | 0.126 | 2.86e-26 | 9.63e-22 |
| Malignant epithelial | C3 Mp ECs | HOMER3 | 1.01 | 0.214 | 0.101 | 1.38e-18 | 4.67e-14 |
| Malignant epithelial | C3 Mp ECs | ILF2 | 1.009 | 0.728 | 0.462 | 3.2e-55 | 1.08e-50 |
| Malignant epithelial | C3 Mp ECs | IMMP1L | 1.008 | 0.341 | 0.178 | 2.24e-24 | 7.55e-20 |
| Malignant epithelial | C3 Mp ECs | SEC22C | 1.007 | 0.201 | 0.087 | 4.75e-20 | 1.6e-15 |
| Malignant epithelial | C3 Mp ECs | SNAPC1 | 1.007 | 0.193 | 0.091 | 2.07e-16 | 6.96e-12 |
| Malignant epithelial | C3 Mp ECs | HACD1 | 1.003 | 0.379 | 0.169 | 1.68e-36 | 5.67e-32 |
| Malignant epithelial | C3 Mp ECs | CCDC34 | 1.002 | 0.574 | 0.296 | 9.67e-48 | 3.26e-43 |
| Malignant epithelial | C3 Mp ECs | PXMP2 | 1.002 | 0.422 | 0.206 | 1.89e-36 | 6.36e-32 |
| Malignant epithelial | C3 Mp ECs | NFYB | 1.002 | 0.39 | 0.183 | 2.72e-35 | 9.17e-31 |
| Malignant epithelial | C3 Mp ECs | OLFML2A | 0.998 | 0.127 | 0.053 | 5.14e-14 | 1.73e-09 |
| Malignant epithelial | C3 Mp ECs | CAV2 | 0.996 | 0.783 | 0.492 | 5.93e-65 | 2e-60 |
| Malignant epithelial | C3 Mp ECs | TRERF1 | 0.996 | 0.124 | 0.069 | 2.13e-07 | 0.00719 |
| Malignant epithelial | C3 Mp ECs | CBX1 | 0.995 | 0.443 | 0.235 | 1.06e-32 | 3.56e-28 |
| Malignant epithelial | C3 Mp ECs | EZH2 | 0.995 | 0.283 | 0.13 | 2.55e-25 | 8.59e-21 |
| Malignant epithelial | C3 Mp ECs | SLC2A4RG | 0.994 | 0.574 | 0.304 | 4.27e-45 | 1.44e-40 |
| Malignant epithelial | C3 Mp ECs | DIO3 | 0.993 | 0.469 | 0.222 | 1.07e-45 | 3.61e-41 |
| Malignant epithelial | C3 Mp ECs | SNRPB | 0.991 | 0.983 | 0.873 | 8.299999999999999e-141 | 2.8e-136 |
| Malignant epithelial | C3 Mp ECs | GLRX5 | 0.99 | 0.713 | 0.444 | 1.08e-55 | 3.63e-51 |
| Malignant epithelial | C3 Mp ECs | CMC2 | 0.989 | 0.603 | 0.335 | 1.09e-45 | 3.68e-41 |
| Malignant epithelial | C3 Mp ECs | SAE1 | 0.988 | 0.517 | 0.259 | 4.12e-44 | 1.39e-39 |
| Malignant epithelial | C3 Mp ECs | LOXL1 | 0.988 | 0.164 | 0.076 | 1.24e-14 | 4.19e-10 |
| Malignant epithelial | C3 Mp ECs | SMPD4 | 0.987 | 0.221 | 0.105 | 7.85e-19 | 2.65e-14 |
| Malignant epithelial | C3 Mp ECs | TOMM40 | 0.985 | 0.652 | 0.41 | 1.6e-43 | 5.39e-39 |
| Malignant epithelial | C3 Mp ECs | MCM7 | 0.984 | 0.469 | 0.25 | 7.4e-35 | 2.49e-30 |
| Malignant epithelial | C3 Mp ECs | ITGB3BP | 0.984 | 0.297 | 0.145 | 6.149999999999999e-23 | 2.07e-18 |
| Malignant epithelial | C3 Mp ECs | PYGL | 0.981 | 0.327 | 0.145 | 1.19e-31 | 4e-27 |
| Malignant epithelial | C3 Mp ECs | MTL5 | 0.98 | 0.312 | 0.137 | 1.47e-30 | 4.94e-26 |
| Malignant epithelial | C3 Mp ECs | BARD1 | 0.98 | 0.192 | 0.092 | 1.03e-15 | 3.46e-11 |
| Malignant epithelial | C3 Mp ECs | KIF1BP | 0.979 | 0.196 | 0.082 | 1.41e-20 | 4.77e-16 |
| Malignant epithelial | C3 Mp ECs | AIMP2 | 0.978 | 0.528 | 0.293 | 1.03e-39 | 3.47e-35 |
| Malignant epithelial | C3 Mp ECs | LYAR | 0.976 | 0.366 | 0.179 | 5.23e-30 | 1.76e-25 |
| Malignant epithelial | C3 Mp ECs | NCEH1 | 0.976 | 0.274 | 0.13 | 1.53e-23 | 5.16e-19 |
| Malignant epithelial | C3 Mp ECs | MMS22L | 0.975 | 0.186 | 0.083 | 5.35e-18 | 1.8e-13 |
| Malignant epithelial | C3 Mp ECs | NUP205 | 0.974 | 0.142 | 0.071 | 1.11e-10 | 3.75e-06 |
| Malignant epithelial | C3 Mp ECs | TFAM | 0.971 | 0.486 | 0.259 | 7.180000000000001e-37 | 2.42e-32 |
| Malignant epithelial | C3 Mp ECs | MAPK12 | 0.969 | 0.17 | 0.066 | 2.65e-20 | 8.93e-16 |
| Malignant epithelial | C3 Mp ECs | MPHOSPH6 | 0.967 | 0.36 | 0.188 | 9.97e-26 | 3.36e-21 |
| Malignant epithelial | C3 Mp ECs | C16orf74 | 0.966 | 0.408 | 0.201 | 4.28e-33 | 1.44e-28 |
| Malignant epithelial | C3 Mp ECs | SLC25A17 | 0.966 | 0.206 | 0.1 | 7.46e-16 | 2.51e-11 |
| Malignant epithelial | C3 Mp ECs | CDKN2AIPNL | 0.965 | 0.531 | 0.299 | 6.06e-37 | 2.04e-32 |
| Malignant epithelial | C3 Mp ECs | THOC3 | 0.965 | 0.4 | 0.201 | 1.21e-31 | 4.070000000000001e-27 |
| Malignant epithelial | C3 Mp ECs | CTD-2371O3.3 | 0.961 | 0.126 | 0.058 | 3.36e-11 | 1.13e-06 |
| Malignant epithelial | C3 Mp ECs | ARL17A | 0.955 | 0.13 | 0.06 | 1.1e-11 | 3.72e-07 |
| Malignant epithelial | C3 Mp ECs | WDR90 | 0.954 | 0.119 | 0.055 | 8.76e-11 | 2.95e-06 |
| Malignant epithelial | C3 Mp ECs | YWHAH | 0.949 | 0.846 | 0.621 | 2.23e-71 | 7.51e-67 |
| Malignant epithelial | C3 Mp ECs | SOCS2 | 0.949 | 0.113 | 0.055 | 2.2e-09 | 7.41e-05 |
| Malignant epithelial | C3 Mp ECs | RAN | 0.948 | 0.979 | 0.884 | 6.67e-137 | 2.25e-132 |
| Malignant epithelial | C3 Mp ECs | H2AFY2 | 0.948 | 0.383 | 0.203 | 3.1e-27 | 1.05e-22 |
| Malignant epithelial | C3 Mp ECs | PRIM2 | 0.948 | 0.177 | 0.08 | 3.11e-16 | 1.05e-11 |

| compartment | subcluster | gene | avg_log2FC | pct.1 | pct.2 | p_val | p_val_adj |
|---|---|---|---|---|---|---|---|
| Malignant epithelial | C3 Mp ECs | COPRS | 0.945 | 0.686 | 0.397 | 3.09e-53 | 1.04e-48 |
| Malignant epithelial | C3 Mp ECs | SHMT1 | 0.94 | 0.113 | 0.049 | 8.88e-12 | 2.99e-07 |
| Malignant epithelial | C3 Mp ECs | NACC1 | 0.936 | 0.254 | 0.115 | 3.38e-23 | 1.14e-18 |
| Malignant epithelial | C3 Mp ECs | ROBO3 | 0.934 | 0.126 | 0.054 | 7.77e-13 | 2.62e-08 |
| Malignant epithelial | C3 Mp ECs | KATNA1 | 0.934 | 0.161 | 0.087 | 3.85e-10 | 1.3e-05 |
| Malignant epithelial | C3 Mp ECs | HNRNPA2B1 | 0.932 | 0.992 | 0.886 | 1.01e-126 | 3.39e-122 |
| Malignant epithelial | C3 Mp ECs | TMEM19 | 0.931 | 0.168 | 0.088 | 2.28e-11 | 7.69e-07 |
| Malignant epithelial | C3 Mp ECs | BOLA3 | 0.93 | 0.789 | 0.49 | 1.19e-65 | 4.01e-61 |
| Malignant epithelial | C3 Mp ECs | NT5E | 0.93 | 0.72 | 0.42 | 1.25e-55 | 4.21e-51 |
| Malignant epithelial | C3 Mp ECs | SAAL1 | 0.926 | 0.261 | 0.124 | 2.17e-21 | 7.3e-17 |
| Malignant epithelial | C3 Mp ECs | BTG3 | 0.924 | 0.4 | 0.203 | 1.71e-29 | 5.77e-25 |
| Malignant epithelial | C3 Mp ECs | NT5DC2 | 0.922 | 0.13 | 0.062 | 1.08e-10 | 3.65e-06 |
| Malignant epithelial | C3 Mp ECs | MX1 | 0.919 | 0.567 | 0.324 | 1.96e-37 | 6.59e-33 |
| Malignant epithelial | C3 Mp ECs | WDR34 | 0.918 | 0.644 | 0.374 | 1.89e-46 | 6.36e-42 |
| Malignant epithelial | C3 Mp ECs | PAQR4 | 0.916 | 0.279 | 0.153 | 2.22e-17 | 7.49e-13 |
| Malignant epithelial | C3 Mp ECs | NEDD1 | 0.914 | 0.139 | 0.06 | 3.46e-14 | 1.16e-09 |
| Malignant epithelial | C3 Mp ECs | ZBP1 | 0.91 | 0.284 | 0.126 | 3.11e-27 | 1.05e-22 |
| Malignant epithelial | C3 Mp ECs | UBXN11 | 0.909 | 0.113 | 0.052 | 4.04e-10 | 1.36e-05 |
| Malignant epithelial | C3 Mp ECs | PHGDH | 0.908 | 0.345 | 0.162 | 9.92e-31 | 3.34e-26 |
| Malignant epithelial | C3 Mp ECs | HMG20B | 0.906 | 0.556 | 0.349 | 4.010000000000001e-31 | 1.35e-26 |
| Malignant epithelial | C3 Mp ECs | COMMD2 | 0.906 | 0.465 | 0.258 | 8.53e-30 | 2.87e-25 |
| Malignant epithelial | C3 Mp ECs | PAGR1 | 0.906 | 0.141 | 0.068 | 1.51e-11 | 5.09e-07 |
| Malignant epithelial | C3 Mp ECs | STMN3 | 0.904 | 0.668 | 0.377 | 3.45e-49 | 1.16e-44 |
| Malignant epithelial | C3 Mp ECs | LPXN | 0.904 | 0.156 | 0.067 | 1.61e-15 | 5.42e-11 |
| Malignant epithelial | C3 Mp ECs | RRP1B | 0.903 | 0.298 | 0.149 | 5.700000000000001e-22 | 1.92e-17 |
| Malignant epithelial | C3 Mp ECs | MMD | 0.901 | 0.228 | 0.121 | 9e-15 | 3.03e-10 |
| Malignant epithelial | C3 Mp ECs | ARHGAP29 | 0.9 | 0.159 | 0.072 | 1.39e-14 | 4.69e-10 |
| Malignant epithelial | C3 Mp ECs | ETS1 | 0.899 | 0.309 | 0.16 | 1.32e-20 | 4.43e-16 |
| Malignant epithelial | C3 Mp ECs | SOCS3 | 0.897 | 0.712 | 0.437 | 6e-48 | 2.02e-43 |
| Malignant epithelial | C3 Mp ECs | KLC3 | 0.895 | 0.193 | 0.093 | 3e-15 | 1.01e-10 |
| Malignant epithelial | C3 Mp ECs | CACNA1A | 0.894 | 0.305 | 0.143 | 6.52e-26 | 2.2e-21 |
| Malignant epithelial | C3 Mp ECs | GAR1 | 0.891 | 0.583 | 0.341 | 2.66e-37 | 8.96e-33 |
| Malignant epithelial | C3 Mp ECs | ANO1 | 0.891 | 0.287 | 0.144 | 2.13e-21 | 7.16e-17 |
| Malignant epithelial | C3 Mp ECs | LSM4 | 0.89 | 0.956 | 0.804 | 3.37e-107 | 1.14e-102 |
| Malignant epithelial | C3 Mp ECs | TUBA1C | 0.889 | 0.88 | 0.681 | 1.13e-60 | 3.82e-56 |
| Malignant epithelial | C3 Mp ECs | NUCB2 | 0.888 | 0.343 | 0.173 | 7.57e-26 | 2.55e-21 |
| Malignant epithelial | C3 Mp ECs | HMGB1 | 0.887 | 0.994 | 0.926 | 1.81e-123 | 6.109999999999999e-119 |
| Malignant epithelial | C3 Mp ECs | TOPBP1 | 0.885 | 0.188 | 0.093 | 7.39e-14 | 2.49e-09 |
| Malignant epithelial | C3 Mp ECs | MTHFD1 | 0.884 | 0.263 | 0.133 | 7.11e-19 | 2.39e-14 |
| Malignant epithelial | C3 Mp ECs | EMG1 | 0.883 | 0.632 | 0.375 | 1.34e-41 | 4.510000000000001e-37 |
| Malignant epithelial | C3 Mp ECs | UNC119B | 0.883 | 0.117 | 0.058 | 5.14e-09 | 0.000173 |
| Malignant epithelial | C3 Mp ECs | COQ2 | 0.881 | 0.168 | 0.084 | 1.87e-12 | 6.31e-08 |
| Malignant epithelial | C3 Mp ECs | BICD2 | 0.881 | 0.108 | 0.048 | 1.77e-10 | 5.95e-06 |
| Malignant epithelial | C3 Mp ECs | EXOSC9 | 0.878 | 0.197 | 0.09 | 1.08e-17 | 3.65e-13 |
| Malignant epithelial | C3 Mp ECs | CBX6 | 0.877 | 0.257 | 0.121 | 1.95e-21 | 6.58e-17 |
| Malignant epithelial | C3 Mp ECs | DDB2 | 0.875 | 0.29 | 0.146 | 3.35e-21 | 1.13e-16 |
| Malignant epithelial | C3 Mp ECs | F3 | 0.874 | 0.619 | 0.34 | 1.15e-49 | 3.89e-45 |
| Malignant epithelial | C3 Mp ECs | PLXNA1 | 0.874 | 0.142 | 0.077 | 7.8e-09 | 0.000263 |
| Malignant epithelial | C3 Mp ECs | CXCL8 | 0.873 | 0.808 | 0.667 | 1.64e-32 | 5.53e-28 |
| Malignant epithelial | C3 Mp ECs | CSTF2 | 0.873 | 0.156 | 0.077 | 1.15e-11 | 3.86e-07 |
| Malignant epithelial | C3 Mp ECs | GEMIN2 | 0.873 | 0.156 | 0.078 | 1.58e-11 | 5.34e-07 |
| Malignant epithelial | C3 Mp ECs | NCAPH2 | 0.872 | 0.287 | 0.143 | 2.73e-21 | 9.18e-17 |
| Malignant epithelial | C3 Mp ECs | COQ3 | 0.872 | 0.154 | 0.073 | 7.29e-13 | 2.46e-08 |
| Malignant epithelial | C3 Mp ECs | TRBC2 | 0.872 | 0.196 | 0.113 | 6.59e-10 | 2.22e-05 |
| Malignant epithelial | C3 Mp ECs | C20orf27 | 0.871 | 0.646 | 0.41 | 4.37e-40 | 1.47e-35 |
| Malignant epithelial | C3 Mp ECs | ALDH16A1 | 0.871 | 0.222 | 0.108 | 2.27e-17 | 7.66e-13 |
| Malignant epithelial | C3 Mp ECs | TIMM10 | 0.87 | 0.621 | 0.375 | 1.12e-39 | 3.78e-35 |
| Malignant epithelial | C3 Mp ECs | NUP210 | 0.869 | 0.13 | 0.062 | 5.63e-11 | 1.9e-06 |
| Malignant epithelial | C3 Mp ECs | EXOSC3 | 0.868 | 0.45 | 0.244 | 8.85e-30 | 2.98e-25 |
| Malignant epithelial | C3 Mp ECs | CMSS1 | 0.867 | 0.472 | 0.256 | 5e-31 | 1.68e-26 |
| Malignant epithelial | C3 Mp ECs | WDR5 | 0.865 | 0.254 | 0.142 | 4.91e-14 | 1.66e-09 |
| Malignant epithelial | C3 Mp ECs | RP11-796E2.4 | 0.863 | 0.167 | 0.082 | 5.47e-13 | 1.84e-08 |
| Malignant epithelial | C3 Mp ECs | FABP5 | 0.862 | 0.75 | 0.521 | 6.26e-39 | 2.11e-34 |
| Malignant epithelial | C3 Mp ECs | PARP2 | 0.861 | 0.124 | 0.056 | 1.03e-11 | 3.46e-07 |

| compartment | subcluster | gene | avg_log2FC | pct.1 | pct.2 | p_val | p_val_adj |
|---|---|---|---|---|---|---|---|
| Malignant epithelial | C3 Mp ECs | POLR3H | 0.86 | 0.327 | 0.177 | 7.82e-21 | 2.63e-16 |
| Malignant epithelial | C3 Mp ECs | CD109 | 0.86 | 0.142 | 0.066 | 2.14e-12 | 7.21e-08 |
| Malignant epithelial | C3 Mp ECs | RP11-152N13.5 | 0.857 | 0.105 | 0.053 | 7.32e-08 | 0.00247 |
| Malignant epithelial | C3 Mp ECs | F12 | 0.856 | 0.538 | 0.319 | 1.55e-31 | 5.240000000000001e-27 |
| Malignant epithelial | C3 Mp ECs | ARNTL2 | 0.856 | 0.268 | 0.128 | 1.1e-21 | 3.71e-17 |
| Malignant epithelial | C3 Mp ECs | HAUS6 | 0.855 | 0.157 | 0.072 | 1.87e-14 | 6.3e-10 |
| Malignant epithelial | C3 Mp ECs | AEN | 0.854 | 0.217 | 0.112 | 9.79e-15 | 3.3e-10 |
| Malignant epithelial | C3 Mp ECs | RNF157 | 0.851 | 0.105 | 0.051 | 1.3e-08 | 0.000439 |
| Malignant epithelial | C3 Mp ECs | SNRPG | 0.85 | 0.972 | 0.852 | 1.5e-101 | 5.04e-97 |
| Malignant epithelial | C3 Mp ECs | MAP4K4 | 0.85 | 0.513 | 0.29 | 3.07e-31 | 1.03e-26 |
| Malignant epithelial | C3 Mp ECs | RRP7A | 0.849 | 0.731 | 0.473 | 2.25e-49 | 7.57e-45 |
| Malignant epithelial | C3 Mp ECs | MAP7D3 | 0.848 | 0.179 | 0.081 | 1.13e-15 | 3.79e-11 |
| Malignant epithelial | C3 Mp ECs | BRD8 | 0.845 | 0.25 | 0.146 | 1.53e-12 | 5.17e-08 |
| Malignant epithelial | C3 Mp ECs | FJX1 | 0.844 | 0.137 | 0.065 | 3.25e-11 | 1.1e-06 |
| Malignant epithelial | C3 Mp ECs | CCDC85B | 0.843 | 0.895 | 0.702 | 3.95e-63 | 1.33e-58 |
| Malignant epithelial | C3 Mp ECs | SFR1 | 0.841 | 0.354 | 0.184 | 3.37e-24 | 1.13e-19 |
| Malignant epithelial | C3 Mp ECs | TOR3A | 0.84 | 0.299 | 0.154 | 4.13e-20 | 1.39e-15 |
| Malignant epithelial | C3 Mp ECs | BBS7 | 0.839 | 0.171 | 0.081 | 3.83e-14 | 1.29e-09 |
| Malignant epithelial | C3 Mp ECs | NME1 | 0.838 | 0.85 | 0.629 | 5.480000000000001e-56 | 1.85e-51 |
| Malignant epithelial | C3 Mp ECs | AP2B1 | 0.838 | 0.652 | 0.422 | 6.74e-35 | 2.27e-30 |
| Malignant epithelial | C3 Mp ECs | NOTCH2 | 0.837 | 0.25 | 0.125 | 7.04e-18 | 2.37e-13 |
| Malignant epithelial | C3 Mp ECs | PITRM1 | 0.836 | 0.152 | 0.077 | 7.64e-11 | 2.57e-06 |
| Malignant epithelial | C3 Mp ECs | RP11-61J19.5 | 0.835 | 0.12 | 0.065 | 1.62e-07 | 0.00547 |
| Malignant epithelial | C3 Mp ECs | SNHG10 | 0.834 | 0.175 | 0.092 | 2.36e-11 | 7.95e-07 |
| Malignant epithelial | C3 Mp ECs | ANTXR2 | 0.834 | 0.124 | 0.062 | 3.09e-09 | 0.000104 |
| Malignant epithelial | C3 Mp ECs | H2AFJ | 0.832 | 0.974 | 0.883 | 2.22e-74 | 7.480000000000001e-70 |
| Malignant epithelial | C3 Mp ECs | SUPT16H | 0.832 | 0.483 | 0.28 | 2.35e-27 | 7.919999999999999e-23 |
| Malignant epithelial | C3 Mp ECs | MGME1 | 0.831 | 0.327 | 0.169 | 1.51e-22 | 5.09e-18 |
| Malignant epithelial | C3 Mp ECs | CCDC109B | 0.831 | 0.326 | 0.169 | 5.5e-22 | 1.85e-17 |
| Malignant epithelial | C3 Mp ECs | TRIM65 | 0.831 | 0.138 | 0.071 | 1.55e-09 | 5.23e-05 |
| Malignant epithelial | C3 Mp ECs | PHLDA1 | 0.829 | 0.926 | 0.714 | 2.64e-67 | 8.9e-63 |
| Malignant epithelial | C3 Mp ECs | RP11-464F9.20 | 0.829 | 0.114 | 0.056 | 4.69e-09 | 0.000158 |
| Malignant epithelial | C3 Mp ECs | BIK | 0.828 | 0.81 | 0.583 | 8.48e-50 | 2.86e-45 |
| Malignant epithelial | C3 Mp ECs | NDUFAF3 | 0.824 | 0.891 | 0.685 | 1.54e-68 | 5.18e-64 |
| Malignant epithelial | C3 Mp ECs | PYCRL | 0.821 | 0.24 | 0.126 | 8.29e-16 | 2.79e-11 |
| Malignant epithelial | C3 Mp ECs | PSMC3 | 0.819 | 0.717 | 0.482 | 1.21e-43 | 4.09e-39 |
| Malignant epithelial | C3 Mp ECs | HSD3B7 | 0.819 | 0.377 | 0.204 | 1.08e-23 | 3.64e-19 |
| Malignant epithelial | C3 Mp ECs | PPP2R5D | 0.819 | 0.135 | 0.072 | 8.02e-09 | 0.00027 |
| Malignant epithelial | C3 Mp ECs | ACYP1 | 0.818 | 0.379 | 0.207 | 3.62e-24 | 1.22e-19 |
| Malignant epithelial | C3 Mp ECs | MLKL | 0.817 | 0.178 | 0.087 | 1.87e-13 | 6.31e-09 |
| Malignant epithelial | C3 Mp ECs | ALCAM | 0.815 | 0.466 | 0.264 | 7.720000000000001e-29 | 2.6e-24 |
| Malignant epithelial | C3 Mp ECs | MZT2B | 0.814 | 0.989 | 0.896 | 1.22e-114 | 4.09e-110 |
| Malignant epithelial | C3 Mp ECs | CSTA | 0.814 | 0.451 | 0.218 | 1.46e-36 | 4.93e-32 |
| Malignant epithelial | C3 Mp ECs | DCBLD2 | 0.812 | 0.709 | 0.399 | 2.06e-53 | 6.96e-49 |
| Malignant epithelial | C3 Mp ECs | KCNK1 | 0.809 | 0.446 | 0.249 | 3.69e-27 | 1.24e-22 |
| Malignant epithelial | C3 Mp ECs | ISYNA1 | 0.809 | 0.332 | 0.164 | 4.73e-25 | 1.59e-20 |
| Malignant epithelial | C3 Mp ECs | HRAS | 0.808 | 0.503 | 0.288 | 4.08e-28 | 1.37e-23 |
| Malignant epithelial | C3 Mp ECs | SMC3 | 0.808 | 0.484 | 0.284 | 1.3e-26 | 4.37e-22 |
| Malignant epithelial | C3 Mp ECs | PSMD9 | 0.806 | 0.541 | 0.347 | 1.32e-25 | 4.46e-21 |
| Malignant epithelial | C3 Mp ECs | AC005355.2 | 0.806 | 0.132 | 0.069 | 4.68e-09 | 0.000158 |
| Malignant epithelial | C3 Mp ECs | COQ7 | 0.805 | 0.437 | 0.257 | 5.969999999999999e-23 | 2.01e-18 |
| Malignant epithelial | C3 Mp ECs | PTRH2 | 0.805 | 0.364 | 0.2 | 2.19e-22 | 7.39e-18 |
| Malignant epithelial | C3 Mp ECs | CDK2 | 0.805 | 0.138 | 0.071 | 7.09e-10 | 2.39e-05 |
| Malignant epithelial | C3 Mp ECs | TOMM5 | 0.803 | 0.648 | 0.394 | 1.85e-40 | 6.24e-36 |
| Malignant epithelial | C3 Mp ECs | DCP2 | 0.803 | 0.294 | 0.166 | 3.51e-16 | 1.18e-11 |
| Malignant epithelial | C3 Mp ECs | SNRPD1 | 0.8 | 0.887 | 0.647 | 9.64e-67 | 3.25e-62 |
| Malignant epithelial | C3 Mp ECs | PITPNC1 | 0.799 | 0.417 | 0.222 | 3.9e-27 | 1.31e-22 |
| Malignant epithelial | C3 Mp ECs | CCL20 | 0.799 | 0.651 | 0.514 | 6.07e-14 | 2.05e-09 |
| Malignant epithelial | C3 Mp ECs | THOC6 | 0.797 | 0.451 | 0.281 | 9.2e-21 | 3.1e-16 |
| Malignant epithelial | C3 Mp ECs | CCDC137 | 0.797 | 0.236 | 0.123 | 2.13e-15 | 7.18e-11 |
| Malignant epithelial | C3 Mp ECs | IPO9 | 0.796 | 0.221 | 0.117 | 4.29e-14 | 1.45e-09 |
| Malignant epithelial | C3 Mp ECs | COMMD4 | 0.795 | 0.625 | 0.373 | 1.3e-38 | 4.38e-34 |
| Malignant epithelial | C3 Mp ECs | DAZAP1 | 0.794 | 0.677 | 0.434 | 6.74e-41 | 2.27e-36 |
| Malignant epithelial | C3 Mp ECs | UCK2 | 0.794 | 0.297 | 0.161 | 1.65e-18 | 5.55e-14 |

| compartment | subcluster | gene | avg_log2FC | pct.1 | pct.2 | p_val | p_val_adj |
|---|---|---|---|---|---|---|---|
| Malignant epithelial | C3 Mp ECs | TOR4A | 0.794 | 0.214 | 0.117 | 1.25e-12 | 4.2e-08 |
| Malignant epithelial | C3 Mp ECs | RFT1 | 0.794 | 0.167 | 0.095 | 8.08e-09 | 0.000272 |
| Malignant epithelial | C3 Mp ECs | CENPB | 0.792 | 0.39 | 0.208 | 6.75e-25 | 2.27e-20 |
| Malignant epithelial | C3 Mp ECs | CRAT | 0.792 | 0.188 | 0.101 | 2.38e-11 | 8.02e-07 |
| Malignant epithelial | C3 Mp ECs | TNNT1 | 0.791 | 0.754 | 0.465 | 3.11e-46 | 1.05e-41 |
| Malignant epithelial | C3 Mp ECs | EXOSC7 | 0.791 | 0.53 | 0.314 | 3.29e-30 | 1.11e-25 |
| Malignant epithelial | C3 Mp ECs | PDXK | 0.786 | 0.814 | 0.587 | 2.91e-47 | 9.81e-43 |
| Malignant epithelial | C3 Mp ECs | EMP2 | 0.786 | 0.67 | 0.473 | 4.25e-27 | 1.43e-22 |
| Malignant epithelial | C3 Mp ECs | CD69 | 0.786 | 0.23 | 0.145 | 9.32e-09 | 0.000314 |
| Malignant epithelial | C3 Mp ECs | SPATA33 | 0.784 | 0.379 | 0.199 | 3.73e-25 | 1.26e-20 |
| Malignant epithelial | C3 Mp ECs | RP11-489E7.4 | 0.784 | 0.274 | 0.146 | 2.19e-17 | 7.39e-13 |
| Malignant epithelial | C3 Mp ECs | ECE2 | 0.784 | 0.254 | 0.136 | 6.56e-16 | 2.21e-11 |
| Malignant epithelial | C3 Mp ECs | LYL1 | 0.784 | 0.132 | 0.065 | 5.09e-10 | 1.72e-05 |
| Malignant epithelial | C3 Mp ECs | TMEM67 | 0.784 | 0.121 | 0.069 | 1.25e-06 | 0.0422 |
| Malignant epithelial | C3 Mp ECs | AP3M2 | 0.783 | 0.186 | 0.095 | 1.12e-12 | 3.78e-08 |
| Malignant epithelial | C3 Mp ECs | ERH | 0.782 | 0.96 | 0.83 | 9.629999999999999e-82 | 3.24e-77 |
| Malignant epithelial | C3 Mp ECs | BCS1L | 0.782 | 0.188 | 0.114 | 2.26e-08 | 0.000762 |
| Malignant epithelial | C3 Mp ECs | CTNNAL1 | 0.781 | 0.259 | 0.124 | 1.46e-20 | 4.91e-16 |
| Malignant epithelial | C3 Mp ECs | RP11-132A1.4 | 0.78 | 0.11 | 0.052 | 1.42e-09 | 4.77e-05 |
| Malignant epithelial | C3 Mp ECs | ANAPC11 | 0.779 | 0.971 | 0.849 | 7.380000000000001e-100 | 2.49e-95 |
| Malignant epithelial | C3 Mp ECs | RBM8A | 0.779 | 0.919 | 0.741 | 1.86e-67 | 6.28e-63 |
| Malignant epithelial | C3 Mp ECs | EIF5AL1 | 0.779 | 0.472 | 0.259 | 1.55e-28 | 5.22e-24 |
| Malignant epithelial | C3 Mp ECs | LETM2 | 0.779 | 0.399 | 0.201 | 1.81e-28 | 6.11e-24 |
| Malignant epithelial | C3 Mp ECs | HAUS7 | 0.779 | 0.16 | 0.087 | 1.06e-09 | 3.57e-05 |
| Malignant epithelial | C3 Mp ECs | NIN | 0.778 | 0.196 | 0.099 | 8.19e-14 | 2.76e-09 |
| Malignant epithelial | C3 Mp ECs | EIF5A | 0.776 | 0.903 | 0.708 | 2e-60 | 6.75e-56 |
| Malignant epithelial | C3 Mp ECs | PA2G4 | 0.776 | 0.872 | 0.683 | 1.37e-54 | 4.61e-50 |
| Malignant epithelial | C3 Mp ECs | MB21D1 | 0.776 | 0.19 | 0.095 | 6.78e-14 | 2.29e-09 |
| Malignant epithelial | C3 Mp ECs | UHRF2 | 0.775 | 0.203 | 0.103 | 2.56e-14 | 8.64e-10 |
| Malignant epithelial | C3 Mp ECs | RCC1 | 0.775 | 0.223 | 0.121 | 9.73e-14 | 3.28e-09 |
| Malignant epithelial | C3 Mp ECs | NAT14 | 0.775 | 0.132 | 0.07 | 8.9e-09 | 0.0003 |
| Malignant epithelial | C3 Mp ECs | PXMP4 | 0.773 | 0.222 | 0.121 | 3.95e-13 | 1.33e-08 |
| Malignant epithelial | C3 Mp ECs | MZF1-AS1 | 0.773 | 0.124 | 0.063 | 4.49e-09 | 0.000151 |
| Malignant epithelial | C3 Mp ECs | AGMO | 0.772 | 0.229 | 0.108 | 1.7e-18 | 5.74e-14 |
| Malignant epithelial | C3 Mp ECs | EEF2KMT | 0.772 | 0.223 | 0.115 | 5.84e-15 | 1.97e-10 |
| Malignant epithelial | C3 Mp ECs | SLC39A3 | 0.771 | 0.47 | 0.274 | 1.36e-25 | 4.58e-21 |
| Malignant epithelial | C3 Mp ECs | ILVBL | 0.771 | 0.423 | 0.243 | 1.69e-23 | 5.7e-19 |
| Malignant epithelial | C3 Mp ECs | DOHH | 0.771 | 0.299 | 0.168 | 4.26e-17 | 1.44e-12 |
| Malignant epithelial | C3 Mp ECs | WDR4 | 0.771 | 0.186 | 0.092 | 7.72e-14 | 2.6e-09 |
| Malignant epithelial | C3 Mp ECs | IRS1 | 0.77 | 0.292 | 0.141 | 4.149999999999999e-23 | 1.4e-18 |
| Malignant epithelial | C3 Mp ECs | FRMD6-AS1 | 0.77 | 0.127 | 0.066 | 1.21e-08 | 0.000409 |
| Malignant epithelial | C3 Mp ECs | CEP295 | 0.769 | 0.105 | 0.049 | 4.06e-09 | 0.000137 |
| Malignant epithelial | C3 Mp ECs | CALB1 | 0.768 | 0.807 | 0.49 | 3.56e-55 | 1.2e-50 |
| Malignant epithelial | C3 Mp ECs | HNRNPUL1 | 0.768 | 0.557 | 0.341 | 1.18e-28 | 3.97e-24 |
| Malignant epithelial | C3 Mp ECs | UCHL5 | 0.768 | 0.252 | 0.145 | 6.32e-13 | 2.13e-08 |
| Malignant epithelial | C3 Mp ECs | CDC25B | 0.766 | 0.557 | 0.336 | 4.21e-31 | 1.42e-26 |
| Malignant epithelial | C3 Mp ECs | LIMD2 | 0.766 | 0.188 | 0.094 | 1.28e-13 | 4.32e-09 |
| Malignant epithelial | C3 Mp ECs | EXOSC2 | 0.766 | 0.237 | 0.131 | 2.06e-13 | 6.94e-09 |
| Malignant epithelial | C3 Mp ECs | RABEP1 | 0.766 | 0.181 | 0.095 | 2.69e-11 | 9.07e-07 |
| Malignant epithelial | C3 Mp ECs | HNRNPA3 | 0.765 | 0.931 | 0.793 | 1e-75 | 3.37e-71 |
| Malignant epithelial | C3 Mp ECs | SRSF7 | 0.765 | 0.914 | 0.737 | 3.46e-61 | 1.16e-56 |
| Malignant epithelial | C3 Mp ECs | PAEP | 0.765 | 0.206 | 0.098 | 1.06e-16 | 3.57e-12 |
| Malignant epithelial | C3 Mp ECs | TBCD | 0.765 | 0.23 | 0.126 | 1.36e-13 | 4.59e-09 |
| Malignant epithelial | C3 Mp ECs | SLC25A19 | 0.765 | 0.127 | 0.065 | 5.66e-09 | 0.000191 |
| Malignant epithelial | C3 Mp ECs | MARC1 | 0.763 | 0.121 | 0.061 | 7.63e-09 | 0.000257 |
| Malignant epithelial | C3 Mp ECs | FOPNL | 0.762 | 0.425 | 0.271 | 1.34e-17 | 4.53e-13 |
| Malignant epithelial | C3 Mp ECs | HNRNPAB | 0.761 | 0.761 | 0.541 | 2.64e-39 | 8.9e-35 |
| Malignant epithelial | C3 Mp ECs | PPIL1 | 0.76 | 0.27 | 0.149 | 2.22e-15 | 7.49e-11 |
| Malignant epithelial | C3 Mp ECs | CASP8AP2 | 0.759 | 0.159 | 0.084 | 3.65e-10 | 1.23e-05 |
| Malignant epithelial | C3 Mp ECs | BLMH | 0.759 | 0.11 | 0.056 | 4.67e-08 | 0.00157 |
| Malignant epithelial | C3 Mp ECs | CYB5R3 | 0.758 | 0.708 | 0.464 | 1.21e-38 | 4.08e-34 |
| Malignant epithelial | C3 Mp ECs | GPATCH4 | 0.758 | 0.396 | 0.239 | 9.01e-19 | 3.03e-14 |
| Malignant epithelial | C3 Mp ECs | LINC00936 | 0.756 | 0.29 | 0.179 | 3.18e-12 | 1.07e-07 |
| Malignant epithelial | C3 Mp ECs | KLK6 | 0.755 | 0.754 | 0.517 | 5.359999999999999e-42 | 1.8e-37 |

| compartment | subcluster | gene | avg_log2FC | pct.1 | pct.2 | p_val | p_val_adj |
|---|---|---|---|---|---|---|---|
| Malignant epithelial | C3 Mp ECs | VDAC3 | 0.755 | 0.754 | 0.52 | 2.75e-38 | 9.26e-34 |
| Malignant epithelial | C3 Mp ECs | CPSF3 | 0.755 | 0.234 | 0.124 | 8.98e-15 | 3.02e-10 |
| Malignant epithelial | C3 Mp ECs | IGFBP4 | 0.753 | 0.429 | 0.266 | 6.51e-19 | 2.19e-14 |
| Malignant epithelial | C3 Mp ECs | SDHA | 0.752 | 0.52 | 0.326 | 3.31e-24 | 1.12e-19 |
| Malignant epithelial | C3 Mp ECs | TRIB2 | 0.752 | 0.247 | 0.12 | 1.48e-18 | 4.97e-14 |
| Malignant epithelial | C3 Mp ECs | PRTFDC1 | 0.752 | 0.163 | 0.094 | 3.38e-08 | 0.00114 |
| Malignant epithelial | C3 Mp ECs | EMP3 | 0.751 | 0.698 | 0.452 | 2.8e-36 | 9.430000000000001e-32 |
| Malignant epithelial | C3 Mp ECs | PTMA | 0.75 | 1.0 | 0.999 | 7.979999999999999e-172 | 2.69e-167 |
| Malignant epithelial | C3 Mp ECs | PARP1 | 0.748 | 0.349 | 0.208 | 5.86e-17 | 1.97e-12 |
| Malignant epithelial | C3 Mp ECs | ARPC5L | 0.747 | 0.752 | 0.519 | 1.3e-43 | 4.39e-39 |
| Malignant epithelial | C3 Mp ECs | COX17 | 0.746 | 0.939 | 0.79 | 2.24e-61 | 7.54e-57 |
| Malignant epithelial | C3 Mp ECs | CH17-360D5.2 | 0.746 | 0.233 | 0.111 | 2.05e-18 | 6.91e-14 |
| Malignant epithelial | C3 Mp ECs | PRADC1 | 0.746 | 0.197 | 0.101 | 2.66e-13 | 8.96e-09 |
| Malignant epithelial | C3 Mp ECs | HLTF | 0.746 | 0.127 | 0.069 | 1.3e-07 | 0.00437 |
| Malignant epithelial | C3 Mp ECs | AIFM2 | 0.745 | 0.197 | 0.114 | 3.8e-10 | 1.28e-05 |
| Malignant epithelial | C3 Mp ECs | FUT8 | 0.744 | 0.394 | 0.206 | 2.7e-26 | 9.1e-22 |
| Malignant epithelial | C3 Mp ECs | FKBP5 | 0.744 | 0.166 | 0.082 | 4.76e-12 | 1.61e-07 |
| Malignant epithelial | C3 Mp ECs | SMC1A | 0.743 | 0.276 | 0.146 | 9.91e-18 | 3.34e-13 |
| Malignant epithelial | C3 Mp ECs | NUP35 | 0.743 | 0.217 | 0.107 | 6.33e-16 | 2.13e-11 |
| Malignant epithelial | C3 Mp ECs | CRAMP1 | 0.743 | 0.106 | 0.056 | 4.12e-07 | 0.0139 |
| Malignant epithelial | C3 Mp ECs | SNCG | 0.742 | 0.153 | 0.092 | 3.56e-07 | 0.012 |
| Malignant epithelial | C3 Mp ECs | CBX3 | 0.741 | 0.917 | 0.746 | 6.83e-66 | 2.3e-61 |
| Malignant epithelial | C3 Mp ECs | GAS6 | 0.741 | 0.501 | 0.338 | 2.63e-19 | 8.85e-15 |
| Malignant epithelial | C3 Mp ECs | ALDH7A1 | 0.741 | 0.274 | 0.155 | 2.36e-14 | 7.94e-10 |
| Malignant epithelial | C3 Mp ECs | PARVB | 0.741 | 0.132 | 0.069 | 5.64e-09 | 0.00019 |
| Malignant epithelial | C3 Mp ECs | PNO1 | 0.74 | 0.337 | 0.207 | 2.2e-14 | 7.42e-10 |
| Malignant epithelial | C3 Mp ECs | OTUD6B | 0.74 | 0.145 | 0.084 | 4.39e-07 | 0.0148 |
| Malignant epithelial | C3 Mp ECs | NUP85 | 0.737 | 0.178 | 0.093 | 1.58e-11 | 5.31e-07 |
| Malignant epithelial | C3 Mp ECs | CALM3 | 0.736 | 0.869 | 0.678 | 5.78e-48 | 1.95e-43 |
| Malignant epithelial | C3 Mp ECs | CCT5 | 0.735 | 0.748 | 0.517 | 1.23e-38 | 4.16e-34 |
| Malignant epithelial | C3 Mp ECs | LRRCC1 | 0.735 | 0.141 | 0.08 | 1.45e-07 | 0.0049 |
| Malignant epithelial | C3 Mp ECs | RP11-182L21.6 | 0.734 | 0.217 | 0.136 | 1.15e-08 | 0.000386 |
| Malignant epithelial | C3 Mp ECs | SLF1 | 0.734 | 0.113 | 0.062 | 4.2e-07 | 0.0142 |
| Malignant epithelial | C3 Mp ECs | DRAP1 | 0.733 | 0.934 | 0.803 | 1.41e-61 | 4.75e-57 |
| Malignant epithelial | C3 Mp ECs | DTD1 | 0.732 | 0.585 | 0.388 | 1.57e-25 | 5.3e-21 |
| Malignant epithelial | C3 Mp ECs | LIF | 0.732 | 0.457 | 0.281 | 7.66e-21 | 2.58e-16 |
| Malignant epithelial | C3 Mp ECs | MKKS | 0.731 | 0.801 | 0.547 | 6.91e-45 | 2.33e-40 |
| Malignant epithelial | C3 Mp ECs | AHCTF1 | 0.731 | 0.163 | 0.079 | 1.82e-12 | 6.14e-08 |
| Malignant epithelial | C3 Mp ECs | POLR1E | 0.731 | 0.152 | 0.084 | 1.05e-08 | 0.000355 |
| Malignant epithelial | C3 Mp ECs | MZT2A | 0.73 | 0.971 | 0.815 | 1.58e-76 | 5.32e-72 |
| Malignant epithelial | C3 Mp ECs | AGAP2-AS1 | 0.73 | 0.225 | 0.121 | 2.29e-13 | 7.7e-09 |
| Malignant epithelial | C3 Mp ECs | FOXL1 | 0.73 | 0.114 | 0.053 | 5.07e-10 | 1.71e-05 |
| Malignant epithelial | C3 Mp ECs | EID3 | 0.729 | 0.178 | 0.099 | 6.21e-10 | 2.09e-05 |
| Malignant epithelial | C3 Mp ECs | ASPH | 0.727 | 0.84 | 0.621 | 1.14e-46 | 3.85e-42 |
| Malignant epithelial | C3 Mp ECs | ANP32A | 0.727 | 0.766 | 0.542 | 3.88e-39 | 1.31e-34 |
| Malignant epithelial | C3 Mp ECs | MRPL35 | 0.727 | 0.381 | 0.213 | 9.050000000000001e-22 | 3.05e-17 |
| Malignant epithelial | C3 Mp ECs | SLC27A3 | 0.727 | 0.132 | 0.061 | 1.54e-11 | 5.17e-07 |
| Malignant epithelial | C3 Mp ECs | LRFN4 | 0.726 | 0.259 | 0.154 | 4.83e-12 | 1.63e-07 |
| Malignant epithelial | C3 Mp ECs | LINC01589 | 0.725 | 0.117 | 0.062 | 1.19e-07 | 0.004 |
| Malignant epithelial | C3 Mp ECs | JMJD4 | 0.721 | 0.164 | 0.093 | 8.56e-09 | 0.000288 |
| Malignant epithelial | C3 Mp ECs | RP11-386I14.4 | 0.72 | 0.166 | 0.081 | 1.46e-12 | 4.91e-08 |
| Malignant epithelial | C3 Mp ECs | BICD1 | 0.717 | 0.157 | 0.084 | 8.66e-10 | 2.92e-05 |
| Malignant epithelial | C3 Mp ECs | CACYBP | 0.716 | 0.796 | 0.522 | 2.12e-48 | 7.14e-44 |
| Malignant epithelial | C3 Mp ECs | RRP36 | 0.716 | 0.248 | 0.142 | 1.27e-12 | 4.29e-08 |
| Malignant epithelial | C3 Mp ECs | PUS1 | 0.716 | 0.265 | 0.159 | 2.26e-12 | 7.61e-08 |
| Malignant epithelial | C3 Mp ECs | DSE | 0.716 | 0.121 | 0.055 | 3.13e-11 | 1.05e-06 |
| Malignant epithelial | C3 Mp ECs | PRKDC | 0.714 | 0.433 | 0.281 | 1.45e-16 | 4.88e-12 |
| Malignant epithelial | C3 Mp ECs | NIP7 | 0.713 | 0.426 | 0.268 | 9.25e-18 | 3.12e-13 |
| Malignant epithelial | C3 Mp ECs | HMBS | 0.713 | 0.314 | 0.18 | 3.29e-16 | 1.11e-11 |
| Malignant epithelial | C3 Mp ECs | YIF1B | 0.713 | 0.228 | 0.134 | 5.38e-11 | 1.81e-06 |
| Malignant epithelial | C3 Mp ECs | USP1 | 0.712 | 0.262 | 0.153 | 6.35e-13 | 2.14e-08 |
| Malignant epithelial | C3 Mp ECs | TPST1 | 0.712 | 0.153 | 0.086 | 2.49e-08 | 0.000838 |
| Malignant epithelial | C3 Mp ECs | SFTA2 | 0.711 | 0.661 | 0.441 | 8.75e-31 | 2.95e-26 |
| Malignant epithelial | C3 Mp ECs | HAUS1 | 0.711 | 0.214 | 0.129 | 3.38e-09 | 0.000114 |

| compartment | subcluster | gene | avg_log2FC | pct.1 | pct.2 | p_val | p_val_adj |
|---|---|---|---|---|---|---|---|
| Malignant epithelial | C3 Mp ECs | RP11-54H7.4 | 0.709 | 0.581 | 0.33 | 1.97e-34 | 6.64e-30 |
| Malignant epithelial | C3 Mp ECs | EREG | 0.709 | 0.724 | 0.515 | 2.65e-28 | 8.939999999999999e-24 |
| Malignant epithelial | C3 Mp ECs | NBPF14 | 0.708 | 0.237 | 0.123 | 1.21e-15 | 4.09e-11 |
| Malignant epithelial | C3 Mp ECs | SLC7A11 | 0.707 | 0.313 | 0.181 | 2.7e-15 | 9.09e-11 |
| Malignant epithelial | C3 Mp ECs | METTL4 | 0.707 | 0.103 | 0.049 | 3.56e-09 | 0.00012 |
| Malignant epithelial | C3 Mp ECs | DCPS | 0.706 | 0.255 | 0.14 | 6.97e-14 | 2.35e-09 |
| Malignant epithelial | C3 Mp ECs | MTFMT | 0.706 | 0.215 | 0.119 | 3.46e-12 | 1.16e-07 |
| Malignant epithelial | C3 Mp ECs | SEPHS1 | 0.705 | 0.225 | 0.128 | 6.94e-12 | 2.34e-07 |
| Malignant epithelial | C3 Mp ECs | XRCC1 | 0.704 | 0.252 | 0.134 | 1.76e-15 | 5.95e-11 |
| Malignant epithelial | C3 Mp ECs | CXCL11 | 0.703 | 0.105 | 0.052 | 2.75e-08 | 0.000926 |
| Malignant epithelial | C3 Mp ECs | TRIM69 | 0.702 | 0.225 | 0.127 | 1.07e-11 | 3.61e-07 |
| Malignant epithelial | C3 Mp ECs | MRPS16 | 0.701 | 0.855 | 0.647 | 5.1e-47 | 1.72e-42 |
| Malignant epithelial | C3 Mp ECs | NFKBIA | 0.701 | 0.95 | 0.873 | 6.37e-35 | 2.15e-30 |
| Malignant epithelial | C3 Mp ECs | TIGAR | 0.7 | 0.163 | 0.092 | 1.96e-08 | 0.000659 |
| Malignant epithelial | C3 Mp ECs | TYMP | 0.699 | 0.901 | 0.661 | 1.16e-50 | 3.89e-46 |
| Malignant epithelial | C3 Mp ECs | TEAD4 | 0.699 | 0.284 | 0.167 | 1.57e-13 | 5.28e-09 |
| Malignant epithelial | C3 Mp ECs | POLD3 | 0.698 | 0.113 | 0.056 | 1.56e-08 | 0.000527 |
| Malignant epithelial | C3 Mp ECs | RP11-110G21.1 | 0.698 | 0.167 | 0.099 | 5.84e-08 | 0.00197 |
| Malignant epithelial | C3 Mp ECs | SEPT7 | 0.697 | 0.843 | 0.673 | 9.779999999999999e-39 | 3.29e-34 |
| Malignant epithelial | C3 Mp ECs | EIF2S1 | 0.697 | 0.494 | 0.304 | 1.66e-23 | 5.59e-19 |
| Malignant epithelial | C3 Mp ECs | GPN3 | 0.697 | 0.262 | 0.146 | 3.18e-14 | 1.07e-09 |
| Malignant epithelial | C3 Mp ECs | NAP1L4 | 0.696 | 0.568 | 0.356 | 2.42e-27 | 8.15e-23 |
| Malignant epithelial | C3 Mp ECs | CEP170 | 0.696 | 0.121 | 0.063 | 3.5e-08 | 0.00118 |
| Malignant epithelial | C3 Mp ECs | LEO1 | 0.695 | 0.185 | 0.101 | 5.74e-11 | 1.93e-06 |
| Malignant epithelial | C3 Mp ECs | R3HDM1 | 0.694 | 0.188 | 0.101 | 2.18e-11 | 7.35e-07 |
| Malignant epithelial | C3 Mp ECs | FANCL | 0.694 | 0.215 | 0.124 | 1.29e-10 | 4.35e-06 |
| Malignant epithelial | C3 Mp ECs | XRCC4 | 0.694 | 0.175 | 0.103 | 3.13e-08 | 0.00105 |
| Malignant epithelial | C3 Mp ECs | PSME2 | 0.693 | 0.961 | 0.824 | 7.72e-62 | 2.6e-57 |
| Malignant epithelial | C3 Mp ECs | GLIPR1 | 0.693 | 0.619 | 0.34 | 7.35e-44 | 2.48e-39 |
| Malignant epithelial | C3 Mp ECs | CALM2 | 0.691 | 0.982 | 0.929 | 6.03e-51 | 2.03e-46 |
| Malignant epithelial | C3 Mp ECs | GMFB | 0.691 | 0.326 | 0.199 | 2.17e-14 | 7.32e-10 |
| Malignant epithelial | C3 Mp ECs | CPSF1 | 0.691 | 0.243 | 0.148 | 1.79e-10 | 6.04e-06 |
| Malignant epithelial | C3 Mp ECs | C2orf82 | 0.69 | 0.581 | 0.365 | 9.7e-27 | 3.27e-22 |
| Malignant epithelial | C3 Mp ECs | RGS14 | 0.69 | 0.212 | 0.133 | 4.44e-08 | 0.00149 |
| Malignant epithelial | C3 Mp ECs | TPRKB | 0.689 | 0.688 | 0.47 | 4.37e-32 | 1.47e-27 |
| Malignant epithelial | C3 Mp ECs | CLN6 | 0.689 | 0.437 | 0.261 | 6.45e-22 | 2.17e-17 |
| Malignant epithelial | C3 Mp ECs | METTL3 | 0.689 | 0.146 | 0.082 | 3.78e-08 | 0.00127 |
| Malignant epithelial | C3 Mp ECs | TIFA | 0.688 | 0.179 | 0.093 | 1.21e-11 | 4.07e-07 |
| Malignant epithelial | C3 Mp ECs | PAFAH1B3 | 0.687 | 0.745 | 0.55 | 7.689999999999999e-34 | 2.59e-29 |
| Malignant epithelial | C3 Mp ECs | MFSD14B | 0.687 | 0.236 | 0.128 | 4.57e-14 | 1.54e-09 |
| Malignant epithelial | C3 Mp ECs | LENG1 | 0.687 | 0.332 | 0.205 | 7.04e-14 | 2.37e-09 |
| Malignant epithelial | C3 Mp ECs | PLEK2 | 0.686 | 0.577 | 0.371 | 3.62e-25 | 1.22e-20 |
| Malignant epithelial | C3 Mp ECs | SMYD2 | 0.685 | 0.28 | 0.159 | 8.7e-15 | 2.93e-10 |
| Malignant epithelial | C3 Mp ECs | KCTD9 | 0.685 | 0.124 | 0.071 | 7.57e-07 | 0.0255 |
| Malignant epithelial | C3 Mp ECs | UQCC3 | 0.684 | 0.815 | 0.595 | 1.21e-44 | 4.08e-40 |
| Malignant epithelial | C3 Mp ECs | JADE2 | 0.683 | 0.259 | 0.141 | 5.67e-15 | 1.91e-10 |
| Malignant epithelial | C3 Mp ECs | BIRC3 | 0.682 | 0.625 | 0.383 | 2.25e-32 | 7.57e-28 |
| Malignant epithelial | C3 Mp ECs | HAGH | 0.682 | 0.452 | 0.277 | 1.1e-20 | 3.71e-16 |
| Malignant epithelial | C3 Mp ECs | RPP25 | 0.682 | 0.512 | 0.33 | 1.14e-20 | 3.85e-16 |
| Malignant epithelial | C3 Mp ECs | NOM1 | 0.682 | 0.301 | 0.174 | 4.16e-15 | 1.4e-10 |
| Malignant epithelial | C3 Mp ECs | WDR54 | 0.682 | 0.302 | 0.18 | 1.18e-13 | 3.98e-09 |
| Malignant epithelial | C3 Mp ECs | MRPL19 | 0.681 | 0.448 | 0.278 | 1.95e-19 | 6.58e-15 |
| Malignant epithelial | C3 Mp ECs | NTAN1 | 0.681 | 0.35 | 0.207 | 1.7e-16 | 5.72e-12 |
| Malignant epithelial | C3 Mp ECs | BNIP1 | 0.681 | 0.163 | 0.094 | 2.21e-08 | 0.000744 |
| Malignant epithelial | C3 Mp ECs | NQO2 | 0.68 | 0.27 | 0.169 | 8.67e-11 | 2.92e-06 |
| Malignant epithelial | C3 Mp ECs | KHSRP | 0.678 | 0.487 | 0.275 | 8.97e-28 | 3.02e-23 |
| Malignant epithelial | C3 Mp ECs | PRRC2A | 0.678 | 0.454 | 0.267 | 6.559999999999999e-23 | 2.21e-18 |
| Malignant epithelial | C3 Mp ECs | ADSL | 0.677 | 0.359 | 0.222 | 2.93e-15 | 9.88e-11 |
| Malignant epithelial | C3 Mp ECs | LTV1 | 0.677 | 0.215 | 0.131 | 3.05e-09 | 0.000103 |
| Malignant epithelial | C3 Mp ECs | ASB9 | 0.676 | 0.16 | 0.087 | 1.28e-09 | 4.32e-05 |
| Malignant epithelial | C3 Mp ECs | MORC4 | 0.676 | 0.127 | 0.071 | 4.6e-07 | 0.0155 |
| Malignant epithelial | C3 Mp ECs | POLR2F | 0.675 | 0.897 | 0.719 | 9.940000000000001e-51 | 3.35e-46 |
| Malignant epithelial | C3 Mp ECs | TOMM34 | 0.675 | 0.366 | 0.216 | 8.98e-18 | 3.03e-13 |
| Malignant epithelial | C3 Mp ECs | ELP6 | 0.675 | 0.259 | 0.157 | 2.1e-11 | 7.09e-07 |

| compartment | subcluster | gene | avg_log2FC | pct.1 | pct.2 | p_val | p_val_adj |
|---|---|---|---|---|---|---|---|
| Malignant epithelial | C3 Mp ECs | MDH1 | 0.673 | 0.652 | 0.457 | 1.01e-27 | 3.42e-23 |
| Malignant epithelial | C3 Mp ECs | TMEM109 | 0.673 | 0.378 | 0.226 | 2.61e-16 | 8.78e-12 |
| Malignant epithelial | C3 Mp ECs | SUZ12 | 0.672 | 0.341 | 0.201 | 7.44e-17 | 2.51e-12 |
| Malignant epithelial | C3 Mp ECs | RPS6KA4 | 0.672 | 0.276 | 0.169 | 6.14e-12 | 2.07e-07 |
| Malignant epithelial | C3 Mp ECs | SOAT1 | 0.672 | 0.121 | 0.062 | 9.95e-09 | 0.000335 |
| Malignant epithelial | C3 Mp ECs | PSMB9 | 0.671 | 0.927 | 0.753 | 4.449999999999999e-48 | 1.5e-43 |
| Malignant epithelial | C3 Mp ECs | SFPQ | 0.67 | 0.767 | 0.548 | 1.58e-34 | 5.33e-30 |
| Malignant epithelial | C3 Mp ECs | PPM1G | 0.67 | 0.712 | 0.485 | 1.96e-32 | 6.59e-28 |
| Malignant epithelial | C3 Mp ECs | BRI3BP | 0.67 | 0.188 | 0.106 | 2.49e-10 | 8.4e-06 |
| Malignant epithelial | C3 Mp ECs | MRTO4 | 0.669 | 0.39 | 0.229 | 5.58e-19 | 1.88e-14 |
| Malignant epithelial | C3 Mp ECs | PVRL1 | 0.667 | 0.185 | 0.108 | 4.85e-09 | 0.000163 |
| Malignant epithelial | C3 Mp ECs | MYO19 | 0.666 | 0.308 | 0.17 | 5.45e-17 | 1.83e-12 |
| Malignant epithelial | C3 Mp ECs | HNRNPH3 | 0.663 | 0.741 | 0.527 | 4.43e-32 | 1.49e-27 |
| Malignant epithelial | C3 Mp ECs | AK6 | 0.663 | 0.447 | 0.284 | 9.77e-18 | 3.29e-13 |
| Malignant epithelial | C3 Mp ECs | STX2 | 0.663 | 0.146 | 0.077 | 1.89e-09 | 6.36e-05 |
| Malignant epithelial | C3 Mp ECs | TIPIN | 0.663 | 0.109 | 0.052 | 5.14e-09 | 0.000173 |
| Malignant epithelial | C3 Mp ECs | UBTF | 0.662 | 0.306 | 0.186 | 3.15e-13 | 1.06e-08 |
| Malignant epithelial | C3 Mp ECs | IDH3A | 0.662 | 0.281 | 0.173 | 5.36e-12 | 1.8e-07 |
| Malignant epithelial | C3 Mp ECs | METTL13 | 0.662 | 0.138 | 0.082 | 1.02e-06 | 0.0343 |
| Malignant epithelial | C3 Mp ECs | SIGMAR1 | 0.661 | 0.588 | 0.391 | 6.71e-25 | 2.26e-20 |
| Malignant epithelial | C3 Mp ECs | SH2D2A | 0.661 | 0.138 | 0.072 | 4.99e-09 | 0.000168 |
| Malignant epithelial | C3 Mp ECs | ZNF814 | 0.661 | 0.116 | 0.061 | 7.71e-08 | 0.0026 |
| Malignant epithelial | C3 Mp ECs | CCDC86 | 0.66 | 0.274 | 0.16 | 2e-13 | 6.73e-09 |
| Malignant epithelial | C3 Mp ECs | SCRIB | 0.66 | 0.283 | 0.173 | 3.65e-12 | 1.23e-07 |
| Malignant epithelial | C3 Mp ECs | SNRPF | 0.659 | 0.946 | 0.817 | 2.53e-62 | 8.52e-58 |
| Malignant epithelial | C3 Mp ECs | ARHGDIB | 0.659 | 0.67 | 0.447 | 3.1e-28 | 1.05e-23 |
| Malignant epithelial | C3 Mp ECs | ZADH2 | 0.659 | 0.131 | 0.075 | 5.44e-07 | 0.0183 |
| Malignant epithelial | C3 Mp ECs | UFD1L | 0.658 | 0.611 | 0.412 | 3.43e-25 | 1.15e-20 |
| Malignant epithelial | C3 Mp ECs | RUVBL1 | 0.657 | 0.408 | 0.235 | 4.84e-21 | 1.63e-16 |
| Malignant epithelial | C3 Mp ECs | C12orf65 | 0.657 | 0.252 | 0.139 | 4.28e-14 | 1.44e-09 |
| Malignant epithelial | C3 Mp ECs | SEMA3A | 0.657 | 0.163 | 0.083 | 5.84e-11 | 1.97e-06 |
| Malignant epithelial | C3 Mp ECs | C1orf174 | 0.657 | 0.228 | 0.143 | 1.03e-08 | 0.000346 |
| Malignant epithelial | C3 Mp ECs | CCP110 | 0.656 | 0.124 | 0.066 | 6.35e-08 | 0.00214 |
| Malignant epithelial | C3 Mp ECs | CHKB | 0.656 | 0.13 | 0.071 | 1.23e-07 | 0.00413 |
| Malignant epithelial | C3 Mp ECs | CDK4 | 0.655 | 0.63 | 0.477 | 1.46e-20 | 4.9e-16 |
| Malignant epithelial | C3 Mp ECs | LSM6 | 0.654 | 0.745 | 0.527 | 3.08e-32 | 1.04e-27 |
| Malignant epithelial | C3 Mp ECs | CRNDE | 0.654 | 0.284 | 0.151 | 4.2e-17 | 1.42e-12 |
| Malignant epithelial | C3 Mp ECs | SLC38A6 | 0.654 | 0.197 | 0.106 | 1.24e-11 | 4.18e-07 |
| Malignant epithelial | C3 Mp ECs | C2orf69 | 0.654 | 0.177 | 0.107 | 7.46e-08 | 0.00251 |
| Malignant epithelial | C3 Mp ECs | UQCC2 | 0.653 | 0.741 | 0.525 | 5.759999999999999e-34 | 1.94e-29 |
| Malignant epithelial | C3 Mp ECs | CTSV | 0.653 | 0.274 | 0.138 | 2.04e-19 | 6.88e-15 |
| Malignant epithelial | C3 Mp ECs | HAUS5 | 0.653 | 0.123 | 0.064 | 2.99e-08 | 0.00101 |
| Malignant epithelial | C3 Mp ECs | MRPL22 | 0.652 | 0.743 | 0.528 | 4.55e-31 | 1.53e-26 |
| Malignant epithelial | C3 Mp ECs | SYNCRIP | 0.652 | 0.674 | 0.483 | 1.74e-24 | 5.86e-20 |
| Malignant epithelial | C3 Mp ECs | NFIC | 0.652 | 0.401 | 0.218 | 1.5e-23 | 5.06e-19 |
| Malignant epithelial | C3 Mp ECs | CBWD2 | 0.652 | 0.212 | 0.127 | 1.96e-09 | 6.62e-05 |
| Malignant epithelial | C3 Mp ECs | SURF2 | 0.65 | 0.564 | 0.358 | 6.26e-24 | 2.11e-19 |
| Malignant epithelial | C3 Mp ECs | BLVRA | 0.65 | 0.277 | 0.165 | 3.49e-12 | 1.18e-07 |
| Malignant epithelial | C3 Mp ECs | HP1BP3 | 0.649 | 0.535 | 0.359 | 3.1e-19 | 1.05e-14 |
| Malignant epithelial | C3 Mp ECs | IRAK1 | 0.649 | 0.423 | 0.265 | 2.63e-17 | 8.87e-13 |
| Malignant epithelial | C3 Mp ECs | POLR3K | 0.648 | 0.375 | 0.233 | 1.67e-15 | 5.62e-11 |
| Malignant epithelial | C3 Mp ECs | FLNA | 0.647 | 0.607 | 0.365 | 1.11e-31 | 3.740000000000001e-27 |
| Malignant epithelial | C3 Mp ECs | CALD1 | 0.647 | 0.257 | 0.132 | 9.7e-17 | 3.27e-12 |
| Malignant epithelial | C3 Mp ECs | CRELD2 | 0.646 | 0.474 | 0.269 | 2.36e-26 | 7.94e-22 |
| Malignant epithelial | C3 Mp ECs | NFKBID | 0.646 | 0.183 | 0.104 | 1.06e-09 | 3.58e-05 |
| Malignant epithelial | C3 Mp ECs | TTLL5 | 0.646 | 0.121 | 0.069 | 1.44e-06 | 0.0487 |
| Malignant epithelial | C3 Mp ECs | NR2C2AP | 0.645 | 0.361 | 0.204 | 3.63e-19 | 1.22e-14 |
| Malignant epithelial | C3 Mp ECs | ANXA1 | 0.643 | 0.952 | 0.773 | 6.659999999999999e-52 | 2.24e-47 |
| Malignant epithelial | C3 Mp ECs | HNRNPD | 0.642 | 0.688 | 0.492 | 5.990000000000001e-27 | 2.02e-22 |
| Malignant epithelial | C3 Mp ECs | SNRK | 0.642 | 0.277 | 0.164 | 1.1e-12 | 3.71e-08 |
| Malignant epithelial | C3 Mp ECs | METTL2A | 0.642 | 0.21 | 0.116 | 1.14e-11 | 3.83e-07 |
| Malignant epithelial | C3 Mp ECs | TNPO2 | 0.642 | 0.186 | 0.104 | 4.89e-10 | 1.65e-05 |
| Malignant epithelial | C3 Mp ECs | YARS2 | 0.642 | 0.204 | 0.128 | 7.73e-08 | 0.0026 |
| Malignant epithelial | C3 Mp ECs | PFDN2 | 0.64 | 0.902 | 0.737 | 1.93e-44 | 6.51e-40 |

| compartment | subcluster | gene | avg_log2FC | pct.1 | pct.2 | p_val | p_val_adj |
|---|---|---|---|---|---|---|---|
| Malignant epithelial | C3 Mp ECs | ACTL6A | 0.64 | 0.452 | 0.276 | 2e-20 | 6.73e-16 |
| Malignant epithelial | C3 Mp ECs | HACD3 | 0.64 | 0.465 | 0.297 | 3.65e-19 | 1.23e-14 |
| Malignant epithelial | C3 Mp ECs | LINC00346 | 0.64 | 0.2 | 0.115 | 6.08e-10 | 2.05e-05 |
| Malignant epithelial | C3 Mp ECs | ZNF281 | 0.639 | 0.255 | 0.147 | 5.52e-13 | 1.86e-08 |
| Malignant epithelial | C3 Mp ECs | C16orf52 | 0.639 | 0.117 | 0.058 | 3.94e-09 | 0.000133 |
| Malignant epithelial | C3 Mp ECs | NASP | 0.637 | 0.474 | 0.301 | 1.29e-19 | 4.36e-15 |
| Malignant epithelial | C3 Mp ECs | TPM2 | 0.636 | 0.957 | 0.741 | 4.759999999999999e-54 | 1.6e-49 |
| Malignant epithelial | C3 Mp ECs | HDGF | 0.636 | 0.703 | 0.486 | 2.42e-29 | 8.16e-25 |
| Malignant epithelial | C3 Mp ECs | NCL | 0.635 | 0.909 | 0.777 | 4.07e-38 | 1.37e-33 |
| Malignant epithelial | C3 Mp ECs | HAS2 | 0.635 | 0.223 | 0.111 | 8.6e-16 | 2.9e-11 |
| Malignant epithelial | C3 Mp ECs | PRPF4 | 0.635 | 0.218 | 0.121 | 1.66e-11 | 5.59e-07 |
| Malignant epithelial | C3 Mp ECs | RRP1 | 0.633 | 0.452 | 0.288 | 7.65e-18 | 2.58e-13 |
| Malignant epithelial | C3 Mp ECs | FTSJ2 | 0.632 | 0.298 | 0.185 | 4.11e-12 | 1.39e-07 |
| Malignant epithelial | C3 Mp ECs | UBE2L3 | 0.631 | 0.876 | 0.695 | 1.26e-44 | 4.26e-40 |
| Malignant epithelial | C3 Mp ECs | MAZ | 0.631 | 0.501 | 0.293 | 2.87e-25 | 9.66e-21 |
| Malignant epithelial | C3 Mp ECs | GNPNAT1 | 0.631 | 0.313 | 0.18 | 3.04e-15 | 1.02e-10 |
| Malignant epithelial | C3 Mp ECs | NUDT15 | 0.63 | 0.418 | 0.25 | 7.11e-20 | 2.4e-15 |
| Malignant epithelial | C3 Mp ECs | ARMC6 | 0.63 | 0.219 | 0.127 | 2.36e-10 | 7.96e-06 |
| Malignant epithelial | C3 Mp ECs | MT1E | 0.629 | 0.714 | 0.446 | 1.17e-34 | 3.94e-30 |
| Malignant epithelial | C3 Mp ECs | ELOVL5 | 0.628 | 0.324 | 0.193 | 5.26e-14 | 1.77e-09 |
| Malignant epithelial | C3 Mp ECs | ARHGAP5-AS1 | 0.628 | 0.257 | 0.165 | 4.37e-09 | 0.000147 |
| Malignant epithelial | C3 Mp ECs | PACSIN2 | 0.627 | 0.356 | 0.219 | 3.35e-15 | 1.13e-10 |
| Malignant epithelial | C3 Mp ECs | CNOT10 | 0.627 | 0.161 | 0.098 | 4e-07 | 0.0135 |
| Malignant epithelial | C3 Mp ECs | SLIRP | 0.626 | 0.952 | 0.83 | 4.78e-54 | 1.61e-49 |
| Malignant epithelial | C3 Mp ECs | WBP11 | 0.626 | 0.484 | 0.312 | 2.23e-19 | 7.52e-15 |
| Malignant epithelial | C3 Mp ECs | PLGRKT | 0.626 | 0.527 | 0.349 | 1.26e-18 | 4.24e-14 |
| Malignant epithelial | C3 Mp ECs | PAICS | 0.626 | 0.462 | 0.296 | 1.82e-18 | 6.12e-14 |
| Malignant epithelial | C3 Mp ECs | ABCE1 | 0.626 | 0.417 | 0.266 | 6.5e-16 | 2.19e-11 |
| Malignant epithelial | C3 Mp ECs | RBM14 | 0.626 | 0.295 | 0.169 | 1.54e-14 | 5.17e-10 |
| Malignant epithelial | C3 Mp ECs | CSTB | 0.625 | 1.0 | 0.979 | 1.03e-49 | 3.48e-45 |
| Malignant epithelial | C3 Mp ECs | GSTO1 | 0.625 | 0.843 | 0.677 | 2.77e-37 | 9.35e-33 |
| Malignant epithelial | C3 Mp ECs | GAA | 0.625 | 0.27 | 0.149 | 3.29e-14 | 1.11e-09 |
| Malignant epithelial | C3 Mp ECs | H1F0 | 0.624 | 0.892 | 0.765 | 5.11e-26 | 1.72e-21 |
| Malignant epithelial | C3 Mp ECs | TSNARE1 | 0.623 | 0.217 | 0.121 | 6.35e-12 | 2.14e-07 |
| Malignant epithelial | C3 Mp ECs | KLHL18 | 0.622 | 0.252 | 0.145 | 1.71e-12 | 5.76e-08 |
| Malignant epithelial | C3 Mp ECs | ADCY7 | 0.622 | 0.181 | 0.093 | 4.35e-12 | 1.46e-07 |
| Malignant epithelial | C3 Mp ECs | BATF3 | 0.622 | 0.127 | 0.065 | 6.26e-09 | 0.000211 |
| Malignant epithelial | C3 Mp ECs | RCC2 | 0.621 | 0.305 | 0.189 | 5.18e-12 | 1.75e-07 |
| Malignant epithelial | C3 Mp ECs | TP53 | 0.621 | 0.292 | 0.179 | 5.23e-12 | 1.76e-07 |
| Malignant epithelial | C3 Mp ECs | ZC3H7B | 0.621 | 0.192 | 0.11 | 2.41e-09 | 8.13e-05 |
| Malignant epithelial | C3 Mp ECs | RBM12B | 0.621 | 0.131 | 0.07 | 3.2e-08 | 0.00108 |
| Malignant epithelial | C3 Mp ECs | HPS3 | 0.62 | 0.167 | 0.099 | 1.14e-07 | 0.00385 |
| Malignant epithelial | C3 Mp ECs | VKORC1L1 | 0.618 | 0.279 | 0.167 | 4.56e-12 | 1.54e-07 |
| Malignant epithelial | C3 Mp ECs | HMGA2 | 0.617 | 0.523 | 0.283 | 1.63e-32 | 5.48e-28 |
| Malignant epithelial | C3 Mp ECs | TAF9 | 0.617 | 0.71 | 0.502 | 2.11e-27 | 7.119999999999999e-23 |
| Malignant epithelial | C3 Mp ECs | DHX9 | 0.616 | 0.401 | 0.246 | 3.44e-17 | 1.16e-12 |
| Malignant epithelial | C3 Mp ECs | CXCR4 | 0.616 | 0.367 | 0.257 | 5.8e-09 | 0.000195 |
| Malignant epithelial | C3 Mp ECs | NLE1 | 0.616 | 0.223 | 0.138 | 9.47e-09 | 0.000319 |
| Malignant epithelial | C3 Mp ECs | SNX5 | 0.615 | 0.666 | 0.466 | 7.46e-25 | 2.51e-20 |
| Malignant epithelial | C3 Mp ECs | FKBP3 | 0.615 | 0.626 | 0.428 | 1.61e-24 | 5.41e-20 |
| Malignant epithelial | C3 Mp ECs | PPP1R1C | 0.615 | 0.255 | 0.137 | 1.51e-14 | 5.09e-10 |
| Malignant epithelial | C3 Mp ECs | STYX | 0.615 | 0.174 | 0.096 | 1.55e-09 | 5.22e-05 |
| Malignant epithelial | C3 Mp ECs | IP6K1 | 0.615 | 0.182 | 0.11 | 4.78e-08 | 0.00161 |
| Malignant epithelial | C3 Mp ECs | MRPL11 | 0.614 | 0.782 | 0.582 | 2.41e-31 | 8.140000000000001e-27 |
| Malignant epithelial | C3 Mp ECs | SNRPA1 | 0.614 | 0.618 | 0.424 | 2.27e-23 | 7.65e-19 |
| Malignant epithelial | C3 Mp ECs | AP1S3 | 0.614 | 0.16 | 0.088 | 4.18e-09 | 0.000141 |
| Malignant epithelial | C3 Mp ECs | SAR1A | 0.613 | 0.71 | 0.52 | 1.95e-26 | 6.58e-22 |
| Malignant epithelial | C3 Mp ECs | PPIH | 0.613 | 0.401 | 0.235 | 1.97e-19 | 6.63e-15 |
| Malignant epithelial | C3 Mp ECs | MTX1 | 0.613 | 0.494 | 0.326 | 2.15e-18 | 7.25e-14 |
| Malignant epithelial | C3 Mp ECs | ADAM19 | 0.613 | 0.292 | 0.157 | 6.38e-17 | 2.15e-12 |
| Malignant epithelial | C3 Mp ECs | UBR7 | 0.613 | 0.102 | 0.053 | 3.97e-07 | 0.0134 |
| Malignant epithelial | C3 Mp ECs | SRSF3 | 0.612 | 0.954 | 0.832 | 1.49e-60 | 5.02e-56 |
| Malignant epithelial | C3 Mp ECs | DMKN | 0.612 | 0.874 | 0.635 | 2.18e-46 | 7.339999999999999e-42 |
| Malignant epithelial | C3 Mp ECs | RNASEH1 | 0.612 | 0.352 | 0.224 | 3.52e-13 | 1.19e-08 |

| compartment | subcluster | gene | avg_log2FC | pct.1 | pct.2 | p_val | p_val_adj |
|---|---|---|---|---|---|---|---|
| Malignant epithelial | C3 Mp ECs | CCDC124 | 0.611 | 0.677 | 0.457 | 5.670000000000001e-29 | 1.91e-24 |
| Malignant epithelial | C3 Mp ECs | CCT2 | 0.611 | 0.743 | 0.539 | 1.68e-28 | 5.65e-24 |
| Malignant epithelial | C3 Mp ECs | POLE3 | 0.61 | 0.454 | 0.298 | 5.52e-17 | 1.86e-12 |
| Malignant epithelial | C3 Mp ECs | RDX | 0.61 | 0.36 | 0.213 | 9.13e-17 | 3.08e-12 |
| Malignant epithelial | C3 Mp ECs | IARS | 0.61 | 0.328 | 0.198 | 4.07e-14 | 1.37e-09 |
| Malignant epithelial | C3 Mp ECs | ITGB2 | 0.61 | 0.16 | 0.082 | 8.44e-11 | 2.84e-06 |
| Malignant epithelial | C3 Mp ECs | MAD2L2 | 0.609 | 0.247 | 0.144 | 3.81e-11 | 1.28e-06 |
| Malignant epithelial | C3 Mp ECs | UBE2N | 0.608 | 0.763 | 0.548 | 7.570000000000001e-32 | 2.55e-27 |
| Malignant epithelial | C3 Mp ECs | ECI1 | 0.608 | 0.611 | 0.412 | 2.06e-23 | 6.94e-19 |
| Malignant epithelial | C3 Mp ECs | SMS | 0.607 | 0.817 | 0.663 | 8.5e-26 | 2.86e-21 |
| Malignant epithelial | C3 Mp ECs | PTRF | 0.607 | 0.297 | 0.17 | 5.79e-15 | 1.95e-10 |
| Malignant epithelial | C3 Mp ECs | BCCIP | 0.607 | 0.363 | 0.237 | 1.9e-12 | 6.4e-08 |
| Malignant epithelial | C3 Mp ECs | CCZ1B | 0.607 | 0.277 | 0.166 | 4.06e-12 | 1.37e-07 |
| Malignant epithelial | C3 Mp ECs | PGAM5 | 0.607 | 0.182 | 0.107 | 1.06e-08 | 0.000359 |
| Malignant epithelial | C3 Mp ECs | COX8A | 0.604 | 0.983 | 0.931 | 1.19e-88 | 4.01e-84 |
| Malignant epithelial | C3 Mp ECs | DCAF7 | 0.604 | 0.343 | 0.221 | 2.32e-12 | 7.83e-08 |
| Malignant epithelial | C3 Mp ECs | TMEM97 | 0.603 | 0.364 | 0.23 | 8.55e-14 | 2.88e-09 |
| Malignant epithelial | C3 Mp ECs | AP000692.10 | 0.603 | 0.108 | 0.052 | 5.64e-09 | 0.00019 |
| Malignant epithelial | C3 Mp ECs | MRPL47 | 0.602 | 0.657 | 0.456 | 3.87e-23 | 1.31e-18 |
| Malignant epithelial | C3 Mp ECs | POP4 | 0.602 | 0.418 | 0.264 | 5.73e-16 | 1.93e-11 |
| Malignant epithelial | C3 Mp ECs | SNRNP70 | 0.601 | 0.69 | 0.509 | 2.78e-24 | 9.37e-20 |
| Malignant epithelial | C3 Mp ECs | MRPL50 | 0.601 | 0.51 | 0.332 | 2.72e-19 | 9.15e-15 |
| Malignant epithelial | C3 Mp ECs | SSRP1 | 0.601 | 0.539 | 0.36 | 6.42e-19 | 2.16e-14 |
| Malignant epithelial | C3 Mp ECs | ATAD3A | 0.601 | 0.197 | 0.118 | 1.1e-08 | 0.000369 |
| Malignant epithelial | C3 Mp ECs | PPP6R1 | 0.6 | 0.287 | 0.172 | 6.08e-12 | 2.05e-07 |
| Malignant epithelial | C3 Mp ECs | SLC35G2 | 0.6 | 0.142 | 0.072 | 7.29e-10 | 2.46e-05 |
| Malignant epithelial | C3 Mp ECs | TPI1 | 0.599 | 0.986 | 0.944 | 4.53e-60 | 1.53e-55 |
| Malignant epithelial | C3 Mp ECs | NOP56 | 0.599 | 0.739 | 0.533 | 3.570000000000001e-27 | 1.2e-22 |
| Malignant epithelial | C3 Mp ECs | SLBP | 0.599 | 0.379 | 0.253 | 3.63e-12 | 1.22e-07 |
| Malignant epithelial | C3 Mp ECs | EED | 0.599 | 0.16 | 0.093 | 5.34e-08 | 0.0018 |
| Malignant epithelial | C3 Mp ECs | NOP58 | 0.598 | 0.588 | 0.4 | 4.65e-22 | 1.57e-17 |
| Malignant epithelial | C3 Mp ECs | CMTM3 | 0.598 | 0.255 | 0.15 | 8.92e-12 | 3.01e-07 |
| Malignant epithelial | C3 Mp ECs | ITGB1BP1 | 0.597 | 0.549 | 0.365 | 1.06e-19 | 3.57e-15 |
| Malignant epithelial | C3 Mp ECs | PSMA4 | 0.596 | 0.89 | 0.751 | 1.81e-35 | 6.09e-31 |
| Malignant epithelial | C3 Mp ECs | TCEA1 | 0.596 | 0.854 | 0.656 | 2.2e-34 | 7.4e-30 |
| Malignant epithelial | C3 Mp ECs | MLF2 | 0.596 | 0.81 | 0.638 | 3.3e-32 | 1.11e-27 |
| Malignant epithelial | C3 Mp ECs | CISD2 | 0.595 | 0.748 | 0.513 | 4.37e-33 | 1.47e-28 |
| Malignant epithelial | C3 Mp ECs | BOP1 | 0.595 | 0.484 | 0.32 | 1.31e-17 | 4.41e-13 |
| Malignant epithelial | C3 Mp ECs | RASSF10 | 0.595 | 0.172 | 0.107 | 7.8e-07 | 0.0263 |
| Malignant epithelial | C3 Mp ECs | SPTSSA | 0.594 | 0.754 | 0.525 | 1.28e-32 | 4.31e-28 |
| Malignant epithelial | C3 Mp ECs | NPSR1 | 0.594 | 0.455 | 0.279 | 7.81e-20 | 2.63e-15 |
| Malignant epithelial | C3 Mp ECs | CIB2 | 0.594 | 0.174 | 0.1 | 1.32e-08 | 0.000444 |
| Malignant epithelial | C3 Mp ECs | HAUS4 | 0.594 | 0.172 | 0.109 | 1.38e-06 | 0.0466 |
| Malignant epithelial | C3 Mp ECs | DHRS9 | 0.592 | 0.273 | 0.142 | 2.02e-18 | 6.82e-14 |
| Malignant epithelial | C3 Mp ECs | SNX24 | 0.592 | 0.132 | 0.067 | 2.87e-09 | 9.66e-05 |
| Malignant epithelial | C3 Mp ECs | TUBGCP4 | 0.592 | 0.163 | 0.095 | 1.14e-07 | 0.00385 |
| Malignant epithelial | C3 Mp ECs | SNAI2 | 0.591 | 0.145 | 0.082 | 1.28e-07 | 0.00431 |
| Malignant epithelial | C3 Mp ECs | PDCD5 | 0.59 | 0.932 | 0.769 | 2.82e-47 | 9.510000000000001e-43 |
| Malignant epithelial | C3 Mp ECs | SLC38A1 | 0.589 | 0.468 | 0.324 | 5.42e-14 | 1.83e-09 |
| Malignant epithelial | C3 Mp ECs | TM2D2 | 0.588 | 0.494 | 0.325 | 2.28e-18 | 7.69e-14 |
| Malignant epithelial | C3 Mp ECs | ANAPC15 | 0.588 | 0.439 | 0.283 | 1.62e-16 | 5.45e-12 |
| Malignant epithelial | C3 Mp ECs | YEATS4 | 0.587 | 0.39 | 0.265 | 1.83e-12 | 6.17e-08 |
| Malignant epithelial | C3 Mp ECs | TMEM14A | 0.586 | 0.628 | 0.417 | 6.96e-25 | 2.34e-20 |
| Malignant epithelial | C3 Mp ECs | EIF4E | 0.586 | 0.648 | 0.467 | 8.100000000000001e-22 | 2.73e-17 |
| Malignant epithelial | C3 Mp ECs | RBM42 | 0.586 | 0.67 | 0.5 | 3.33e-20 | 1.12e-15 |
| Malignant epithelial | C3 Mp ECs | C12orf57 | 0.585 | 0.839 | 0.641 | 1.67e-34 | 5.61e-30 |
| Malignant epithelial | C3 Mp ECs | POLR2L | 0.584 | 0.968 | 0.87 | 3.49e-57 | 1.18e-52 |
| Malignant epithelial | C3 Mp ECs | CKLF | 0.584 | 0.785 | 0.572 | 4.43e-32 | 1.49e-27 |
| Malignant epithelial | C3 Mp ECs | TMEM18 | 0.584 | 0.548 | 0.354 | 3.790000000000001e-21 | 1.28e-16 |
| Malignant epithelial | C3 Mp ECs | CD53 | 0.584 | 0.135 | 0.073 | 2.89e-08 | 0.000973 |
| Malignant epithelial | C3 Mp ECs | SNRPD3 | 0.583 | 0.85 | 0.693 | 3.69e-34 | 1.24e-29 |
| Malignant epithelial | C3 Mp ECs | PKN1 | 0.583 | 0.268 | 0.149 | 2.14e-14 | 7.2e-10 |
| Malignant epithelial | C3 Mp ECs | PSMG3 | 0.582 | 0.684 | 0.493 | 1.64e-24 | 5.51e-20 |
| Malignant epithelial | C3 Mp ECs | FAM136A | 0.581 | 0.709 | 0.489 | 9.26e-29 | 3.12e-24 |

| compartment | subcluster | gene | avg_log2FC | pct.1 | pct.2 | p_val | p_val_adj |
|---|---|---|---|---|---|---|---|
| Malignant epithelial | C3 Mp ECs | PDLIM7 | 0.581 | 0.415 | 0.24 | 1.21e-20 | 4.07e-16 |
| Malignant epithelial | C3 Mp ECs | GBP1 | 0.581 | 0.288 | 0.163 | 1.03e-14 | 3.48e-10 |
| Malignant epithelial | C3 Mp ECs | GPSM3 | 0.581 | 0.228 | 0.137 | 1.29e-09 | 4.35e-05 |
| Malignant epithelial | C3 Mp ECs | SUGP2 | 0.58 | 0.229 | 0.134 | 2.08e-10 | 6.99e-06 |
| Malignant epithelial | C3 Mp ECs | GMPPB | 0.58 | 0.172 | 0.093 | 4.83e-10 | 1.63e-05 |
| Malignant epithelial | C3 Mp ECs | TCTEX1D2 | 0.579 | 0.39 | 0.224 | 2.14e-19 | 7.2e-15 |
| Malignant epithelial | C3 Mp ECs | MRPL9 | 0.579 | 0.564 | 0.398 | 4.38e-18 | 1.48e-13 |
| Malignant epithelial | C3 Mp ECs | SRRD | 0.579 | 0.142 | 0.084 | 9.15e-07 | 0.0308 |
| Malignant epithelial | C3 Mp ECs | NUP50 | 0.578 | 0.407 | 0.223 | 6.53e-23 | 2.2e-18 |
| Malignant epithelial | C3 Mp ECs | UNC13D | 0.578 | 0.41 | 0.235 | 2.82e-20 | 9.49e-16 |
| Malignant epithelial | C3 Mp ECs | GPSM2 | 0.578 | 0.197 | 0.113 | 9.24e-10 | 3.11e-05 |
| Malignant epithelial | C3 Mp ECs | RP11-44F21.5 | 0.578 | 0.164 | 0.091 | 7.96e-09 | 0.000268 |
| Malignant epithelial | C3 Mp ECs | ICAM3 | 0.577 | 0.317 | 0.211 | 7.01e-10 | 2.36e-05 |
| Malignant epithelial | C3 Mp ECs | DCAF15 | 0.577 | 0.177 | 0.099 | 1.59e-09 | 5.37e-05 |
| Malignant epithelial | C3 Mp ECs | WDR12 | 0.577 | 0.214 | 0.133 | 2.1e-08 | 0.000707 |
| Malignant epithelial | C3 Mp ECs | ABLIM3 | 0.576 | 0.137 | 0.068 | 3.13e-10 | 1.05e-05 |
| Malignant epithelial | C3 Mp ECs | ANP32B | 0.575 | 0.909 | 0.787 | 9.789999999999999e-40 | 3.3e-35 |
| Malignant epithelial | C3 Mp ECs | KRT6B | 0.575 | 0.404 | 0.278 | 2.76e-11 | 9.31e-07 |
| Malignant epithelial | C3 Mp ECs | RASSF1 | 0.575 | 0.246 | 0.145 | 3.96e-11 | 1.34e-06 |
| Malignant epithelial | C3 Mp ECs | SS18 | 0.575 | 0.211 | 0.125 | 2.5e-09 | 8.44e-05 |
| Malignant epithelial | C3 Mp ECs | SORD | 0.574 | 0.381 | 0.247 | 7.84e-14 | 2.64e-09 |
| Malignant epithelial | C3 Mp ECs | NOC3L | 0.574 | 0.137 | 0.074 | 4.12e-08 | 0.00139 |
| Malignant epithelial | C3 Mp ECs | SET | 0.573 | 0.938 | 0.825 | 3.38e-35 | 1.14e-30 |
| Malignant epithelial | C3 Mp ECs | POP7 | 0.573 | 0.629 | 0.443 | 1.51e-21 | 5.08e-17 |
| Malignant epithelial | C3 Mp ECs | HNRNPH1 | 0.573 | 0.785 | 0.616 | 3.51e-18 | 1.18e-13 |
| Malignant epithelial | C3 Mp ECs | PAF1 | 0.573 | 0.262 | 0.153 | 3.01e-12 | 1.02e-07 |
| Malignant epithelial | C3 Mp ECs | TCEB1 | 0.571 | 0.953 | 0.825 | 1.1e-46 | 3.689999999999999e-42 |
| Malignant epithelial | C3 Mp ECs | GADD45GIP1 | 0.571 | 0.93 | 0.798 | 2.71e-45 | 9.14e-41 |
| Malignant epithelial | C3 Mp ECs | SUMO3 | 0.571 | 0.709 | 0.546 | 4.500000000000001e-22 | 1.52e-17 |
| Malignant epithelial | C3 Mp ECs | NUDT8 | 0.571 | 0.389 | 0.256 | 2.52e-13 | 8.48e-09 |
| Malignant epithelial | C3 Mp ECs | RPL26L1 | 0.57 | 0.741 | 0.536 | 5.3e-29 | 1.79e-24 |
| Malignant epithelial | C3 Mp ECs | ELAVL1 | 0.57 | 0.546 | 0.363 | 4.04e-20 | 1.36e-15 |
| Malignant epithelial | C3 Mp ECs | IFRD2 | 0.57 | 0.549 | 0.381 | 5.24e-18 | 1.77e-13 |
| Malignant epithelial | C3 Mp ECs | NDUFB6 | 0.569 | 0.868 | 0.663 | 2.72e-37 | 9.17e-33 |
| Malignant epithelial | C3 Mp ECs | ENDOG | 0.569 | 0.233 | 0.135 | 4.98e-11 | 1.68e-06 |
| Malignant epithelial | C3 Mp ECs | FKBP4 | 0.568 | 0.698 | 0.502 | 1.91e-25 | 6.440000000000001e-21 |
| Malignant epithelial | C3 Mp ECs | CPSF4 | 0.568 | 0.222 | 0.138 | 2.46e-08 | 0.000829 |
| Malignant epithelial | C3 Mp ECs | LINC01588 | 0.568 | 0.179 | 0.112 | 4.02e-07 | 0.0136 |
| Malignant epithelial | C3 Mp ECs | RP11-158I9.8 | 0.568 | 0.117 | 0.066 | 1.46e-06 | 0.0492 |
| Malignant epithelial | C3 Mp ECs | FADD | 0.567 | 0.277 | 0.16 | 3.19e-13 | 1.07e-08 |
| Malignant epithelial | C3 Mp ECs | LMNA | 0.566 | 0.959 | 0.86 | 8.51e-44 | 2.87e-39 |
| Malignant epithelial | C3 Mp ECs | IFI16 | 0.566 | 0.524 | 0.32 | 1.14e-22 | 3.83e-18 |
| Malignant epithelial | C3 Mp ECs | MRPS11 | 0.566 | 0.531 | 0.343 | 4.3e-20 | 1.45e-15 |
| Malignant epithelial | C3 Mp ECs | SAYSD1 | 0.566 | 0.201 | 0.119 | 2.68e-09 | 9.03e-05 |
| Malignant epithelial | C3 Mp ECs | TRMU | 0.566 | 0.168 | 0.1 | 7.24e-08 | 0.00244 |
| Malignant epithelial | C3 Mp ECs | NDUFA2 | 0.565 | 0.91 | 0.778 | 7.949999999999999e-41 | 2.68e-36 |
| Malignant epithelial | C3 Mp ECs | HINT2 | 0.564 | 0.817 | 0.595 | 3.11e-33 | 1.05e-28 |
| Malignant epithelial | C3 Mp ECs | PHF5A | 0.564 | 0.625 | 0.468 | 4.56e-19 | 1.54e-14 |
| Malignant epithelial | C3 Mp ECs | KIAA1143 | 0.564 | 0.423 | 0.263 | 4.67e-17 | 1.57e-12 |
| Malignant epithelial | C3 Mp ECs | BCAS4 | 0.564 | 0.345 | 0.21 | 2.61e-14 | 8.81e-10 |
| Malignant epithelial | C3 Mp ECs | AGPAT5 | 0.564 | 0.154 | 0.092 | 4.1e-07 | 0.0138 |
| Malignant epithelial | C3 Mp ECs | MIR4435-2HG | 0.563 | 0.869 | 0.614 | 1.87e-45 | 6.309999999999999e-41 |
| Malignant epithelial | C3 Mp ECs | CD74 | 0.563 | 0.993 | 0.882 | 1.47e-40 | 4.96e-36 |
| Malignant epithelial | C3 Mp ECs | SRA1 | 0.563 | 0.552 | 0.369 | 2.69e-19 | 9.05e-15 |
| Malignant epithelial | C3 Mp ECs | UROS | 0.563 | 0.532 | 0.359 | 2.22e-18 | 7.49e-14 |
| Malignant epithelial | C3 Mp ECs | POP5 | 0.563 | 0.458 | 0.293 | 5.46e-18 | 1.84e-13 |
| Malignant epithelial | C3 Mp ECs | DCUN1D5 | 0.563 | 0.463 | 0.304 | 1.08e-15 | 3.65e-11 |
| Malignant epithelial | C3 Mp ECs | GANAB | 0.563 | 0.455 | 0.307 | 2.13e-14 | 7.18e-10 |
| Malignant epithelial | C3 Mp ECs | KRI1 | 0.563 | 0.172 | 0.087 | 6.65e-12 | 2.24e-07 |
| Malignant epithelial | C3 Mp ECs | PIN1 | 0.562 | 0.702 | 0.507 | 4.92e-26 | 1.66e-21 |
| Malignant epithelial | C3 Mp ECs | FAM174A | 0.562 | 0.374 | 0.245 | 1.44e-12 | 4.86e-08 |
| Malignant epithelial | C3 Mp ECs | HNRNPA1L2 | 0.562 | 0.294 | 0.185 | 8.37e-11 | 2.82e-06 |
| Malignant epithelial | C3 Mp ECs | MRPL46 | 0.562 | 0.303 | 0.199 | 5.57e-10 | 1.88e-05 |
| Malignant epithelial | C3 Mp ECs | LRRC23 | 0.562 | 0.181 | 0.111 | 3.51e-07 | 0.0118 |

| compartment | subcluster | gene | avg_log2FC | pct.1 | pct.2 | p_val | p_val_adj |
|---|---|---|---|---|---|---|---|
| Malignant epithelial | C3 Mp ECs | SDF2L1 | 0.561 | 0.699 | 0.493 | 1.19e-28 | 4.01e-24 |
| Malignant epithelial | C3 Mp ECs | FAM104A | 0.561 | 0.337 | 0.215 | 4.36e-12 | 1.47e-07 |
| Malignant epithelial | C3 Mp ECs | RGS1 | 0.561 | 0.492 | 0.395 | 1.26e-06 | 0.0426 |
| Malignant epithelial | C3 Mp ECs | RHEB | 0.56 | 0.903 | 0.718 | 2.9e-37 | 9.76e-33 |
| Malignant epithelial | C3 Mp ECs | FSCN1 | 0.56 | 0.56 | 0.349 | 6.02e-24 | 2.03e-19 |
| Malignant epithelial | C3 Mp ECs | MAGOH | 0.56 | 0.652 | 0.442 | 1.05e-23 | 3.53e-19 |
| Malignant epithelial | C3 Mp ECs | HADH | 0.559 | 0.352 | 0.215 | 2.55e-14 | 8.58e-10 |
| Malignant epithelial | C3 Mp ECs | RBM17 | 0.558 | 0.636 | 0.456 | 3.3e-20 | 1.11e-15 |
| Malignant epithelial | C3 Mp ECs | PSAT1 | 0.558 | 0.21 | 0.113 | 3.05e-12 | 1.03e-07 |
| Malignant epithelial | C3 Mp ECs | DESI2 | 0.558 | 0.259 | 0.169 | 8.77e-09 | 0.000295 |
| Malignant epithelial | C3 Mp ECs | SLC25A45 | 0.558 | 0.137 | 0.08 | 1.14e-06 | 0.0383 |
| Malignant epithelial | C3 Mp ECs | FARSA | 0.557 | 0.389 | 0.259 | 4.77e-13 | 1.61e-08 |
| Malignant epithelial | C3 Mp ECs | HLA-DQA1 | 0.556 | 0.291 | 0.187 | 1.05e-09 | 3.55e-05 |
| Malignant epithelial | C3 Mp ECs | SAMHD1 | 0.556 | 0.143 | 0.081 | 1.26e-07 | 0.00423 |
| Malignant epithelial | C3 Mp ECs | HNRNPM | 0.555 | 0.734 | 0.53 | 7.17e-26 | 2.42e-21 |
| Malignant epithelial | C3 Mp ECs | SMTN | 0.555 | 0.276 | 0.165 | 8.57e-12 | 2.89e-07 |
| Malignant epithelial | C3 Mp ECs | CBFB | 0.554 | 0.364 | 0.233 | 1.44e-13 | 4.84e-09 |
| Malignant epithelial | C3 Mp ECs | UBP1 | 0.554 | 0.212 | 0.133 | 4.31e-08 | 0.00145 |
| Malignant epithelial | C3 Mp ECs | OAS3 | 0.553 | 0.348 | 0.194 | 5.04e-18 | 1.7e-13 |
| Malignant epithelial | C3 Mp ECs | DEF8 | 0.553 | 0.366 | 0.216 | 1.09e-16 | 3.67e-12 |
| Malignant epithelial | C3 Mp ECs | SDHAF4 | 0.553 | 0.27 | 0.161 | 8.42e-12 | 2.84e-07 |
| Malignant epithelial | C3 Mp ECs | PRPF19 | 0.552 | 0.41 | 0.276 | 1.34e-12 | 4.52e-08 |
| Malignant epithelial | C3 Mp ECs | XPO1 | 0.551 | 0.48 | 0.308 | 5.85e-18 | 1.97e-13 |
| Malignant epithelial | C3 Mp ECs | MT1A | 0.551 | 0.194 | 0.098 | 2.63e-13 | 8.86e-09 |
| Malignant epithelial | C3 Mp ECs | USP5 | 0.551 | 0.279 | 0.167 | 9.78e-12 | 3.29e-07 |
| Malignant epithelial | C3 Mp ECs | DCBLD1 | 0.55 | 0.408 | 0.228 | 3.09e-22 | 1.04e-17 |
| Malignant epithelial | C3 Mp ECs | TAF15 | 0.55 | 0.41 | 0.268 | 1.25e-13 | 4.23e-09 |
| Malignant epithelial | C3 Mp ECs | RPGRIP1L | 0.55 | 0.156 | 0.087 | 2.86e-08 | 0.000962 |
| Malignant epithelial | C3 Mp ECs | EVA1C | 0.55 | 0.141 | 0.079 | 1.83e-07 | 0.00617 |
| Malignant epithelial | C3 Mp ECs | HAT1 | 0.549 | 0.323 | 0.196 | 1.31e-13 | 4.42e-09 |
| Malignant epithelial | C3 Mp ECs | DYNLL1 | 0.548 | 0.993 | 0.945 | 6.88e-66 | 2.32e-61 |
| Malignant epithelial | C3 Mp ECs | PSMB3 | 0.548 | 0.934 | 0.808 | 1.78e-43 | 5.99e-39 |
| Malignant epithelial | C3 Mp ECs | IFITM1 | 0.548 | 0.799 | 0.585 | 2.05e-29 | 6.91e-25 |
| Malignant epithelial | C3 Mp ECs | PPP2R5C | 0.547 | 0.639 | 0.438 | 1.82e-23 | 6.13e-19 |
| Malignant epithelial | C3 Mp ECs | CCDC88C | 0.547 | 0.193 | 0.121 | 3.66e-07 | 0.0123 |
| Malignant epithelial | C3 Mp ECs | COX20 | 0.546 | 0.783 | 0.582 | 1.55e-28 | 5.22e-24 |
| Malignant epithelial | C3 Mp ECs | WEE1 | 0.546 | 0.614 | 0.429 | 4.56e-18 | 1.54e-13 |
| Malignant epithelial | C3 Mp ECs | PTPN11 | 0.546 | 0.29 | 0.182 | 1.05e-10 | 3.53e-06 |
| Malignant epithelial | C3 Mp ECs | C10orf35 | 0.546 | 0.212 | 0.123 | 6.75e-10 | 2.27e-05 |
| Malignant epithelial | C3 Mp ECs | OSGIN2 | 0.546 | 0.193 | 0.112 | 3.38e-09 | 0.000114 |
| Malignant epithelial | C3 Mp ECs | USP14 | 0.546 | 0.24 | 0.152 | 1.15e-08 | 0.000388 |
| Malignant epithelial | C3 Mp ECs | NOL10 | 0.546 | 0.185 | 0.108 | 1.49e-08 | 0.000503 |
| Malignant epithelial | C3 Mp ECs | FBXW2 | 0.546 | 0.233 | 0.149 | 2.41e-08 | 0.000811 |
| Malignant epithelial | C3 Mp ECs | CYCS | 0.544 | 0.977 | 0.896 | 3.11e-56 | 1.05e-51 |
| Malignant epithelial | C3 Mp ECs | ADK | 0.544 | 0.455 | 0.292 | 1.04e-17 | 3.5e-13 |
| Malignant epithelial | C3 Mp ECs | AEBP2 | 0.544 | 0.179 | 0.111 | 3.7e-07 | 0.0125 |
| Malignant epithelial | C3 Mp ECs | NIPA1 | 0.544 | 0.116 | 0.065 | 1.26e-06 | 0.0424 |
| Malignant epithelial | C3 Mp ECs | NDUFB1 | 0.543 | 0.966 | 0.858 | 2.28e-49 | 7.689999999999999e-45 |
| Malignant epithelial | C3 Mp ECs | EHBP1 | 0.543 | 0.328 | 0.19 | 9.77e-16 | 3.29e-11 |
| Malignant epithelial | C3 Mp ECs | VBP1 | 0.542 | 0.541 | 0.371 | 2.68e-17 | 9.01e-13 |
| Malignant epithelial | C3 Mp ECs | NME4 | 0.542 | 0.634 | 0.464 | 3.01e-17 | 1.02e-12 |
| Malignant epithelial | C3 Mp ECs | SLC19A1 | 0.542 | 0.109 | 0.059 | 1.18e-06 | 0.0396 |
| Malignant epithelial | C3 Mp ECs | ISOC1 | 0.541 | 0.295 | 0.181 | 1.95e-11 | 6.55e-07 |
| Malignant epithelial | C3 Mp ECs | TGFB1 | 0.541 | 0.27 | 0.165 | 8.51e-11 | 2.87e-06 |
| Malignant epithelial | C3 Mp ECs | SMC6 | 0.541 | 0.217 | 0.132 | 8.07e-09 | 0.000272 |
| Malignant epithelial | C3 Mp ECs | MRPL55 | 0.54 | 0.743 | 0.563 | 1.02e-25 | 3.45e-21 |
| Malignant epithelial | C3 Mp ECs | GPAA1 | 0.54 | 0.539 | 0.372 | 3.07e-16 | 1.03e-11 |
| Malignant epithelial | C3 Mp ECs | BANF1 | 0.539 | 0.946 | 0.799 | 7.1e-49 | 2.39e-44 |
| Malignant epithelial | C3 Mp ECs | C22orf39 | 0.539 | 0.563 | 0.375 | 3.51e-19 | 1.18e-14 |
| Malignant epithelial | C3 Mp ECs | SCRN1 | 0.539 | 0.223 | 0.118 | 3.49e-13 | 1.18e-08 |
| Malignant epithelial | C3 Mp ECs | VPS9D1-AS1 | 0.538 | 0.159 | 0.091 | 5.5e-08 | 0.00185 |
| Malignant epithelial | C3 Mp ECs | LY6E | 0.537 | 0.974 | 0.791 | 1.47e-48 | 4.95e-44 |
| Malignant epithelial | C3 Mp ECs | SLC35F2 | 0.537 | 0.199 | 0.122 | 4.07e-08 | 0.00137 |
| Malignant epithelial | C3 Mp ECs | TPM3 | 0.536 | 0.876 | 0.725 | 3.39e-34 | 1.14e-29 |

| compartment | subcluster | gene | avg_log2FC | pct.1 | pct.2 | p_val | p_val_adj |
|---|---|---|---|---|---|---|---|
| Malignant epithelial | C3 Mp ECs | TTLL12 | 0.536 | 0.265 | 0.167 | 7.49e-10 | 2.52e-05 |
| Malignant epithelial | C3 Mp ECs | LINC01116 | 0.536 | 0.262 | 0.17 | 1.49e-08 | 0.000502 |
| Malignant epithelial | C3 Mp ECs | DHX38 | 0.536 | 0.208 | 0.135 | 3.73e-07 | 0.0126 |
| Malignant epithelial | C3 Mp ECs | GPX1 | 0.535 | 0.983 | 0.939 | 4.96e-59 | 1.67e-54 |
| Malignant epithelial | C3 Mp ECs | PDZD11 | 0.535 | 0.65 | 0.465 | 3.33e-20 | 1.12e-15 |
| Malignant epithelial | C3 Mp ECs | PRMT1 | 0.535 | 0.734 | 0.588 | 3.93e-20 | 1.33e-15 |
| Malignant epithelial | C3 Mp ECs | C12orf45 | 0.534 | 0.509 | 0.331 | 2.07e-18 | 6.99e-14 |
| Malignant epithelial | C3 Mp ECs | XAF1 | 0.534 | 0.377 | 0.221 | 1.12e-17 | 3.77e-13 |
| Malignant epithelial | C3 Mp ECs | UCK1 | 0.534 | 0.19 | 0.106 | 4.87e-10 | 1.64e-05 |
| Malignant epithelial | C3 Mp ECs | FAR1 | 0.534 | 0.217 | 0.129 | 3.1e-09 | 0.000105 |
| Malignant epithelial | C3 Mp ECs | PRPSAP1 | 0.533 | 0.287 | 0.194 | 4.1e-08 | 0.00138 |
| Malignant epithelial | C3 Mp ECs | CXCL10 | 0.533 | 0.143 | 0.087 | 1.13e-06 | 0.0381 |
| Malignant epithelial | C3 Mp ECs | MRPS23 | 0.532 | 0.566 | 0.373 | 1.45e-20 | 4.87e-16 |
| Malignant epithelial | C3 Mp ECs | CTDSPL2 | 0.532 | 0.179 | 0.105 | 3.06e-08 | 0.00103 |
| Malignant epithelial | C3 Mp ECs | PKP4 | 0.532 | 0.194 | 0.121 | 2.08e-07 | 0.00701 |
| Malignant epithelial | C3 Mp ECs | PGAM1 | 0.53 | 0.738 | 0.577 | 1.27e-22 | 4.28e-18 |
| Malignant epithelial | C3 Mp ECs | ODF3B | 0.53 | 0.473 | 0.286 | 7.460000000000001e-21 | 2.51e-16 |
| Malignant epithelial | C3 Mp ECs | HPRT1 | 0.53 | 0.56 | 0.383 | 2.4e-17 | 8.08e-13 |
| Malignant epithelial | C3 Mp ECs | MRPL13 | 0.529 | 0.752 | 0.565 | 1.07e-24 | 3.59e-20 |
| Malignant epithelial | C3 Mp ECs | DDX46 | 0.529 | 0.687 | 0.486 | 7.910000000000001e-21 | 2.67e-16 |
| Malignant epithelial | C3 Mp ECs | SERPINE2 | 0.529 | 0.48 | 0.307 | 2.44e-19 | 8.21e-15 |
| Malignant epithelial | C3 Mp ECs | ZDHHC12 | 0.529 | 0.665 | 0.5 | 3.96e-19 | 1.33e-14 |
| Malignant epithelial | C3 Mp ECs | MRPL41 | 0.528 | 0.881 | 0.682 | 8.63e-33 | 2.91e-28 |
| Malignant epithelial | C3 Mp ECs | CHCHD1 | 0.528 | 0.686 | 0.488 | 1.84e-21 | 6.18e-17 |
| Malignant epithelial | C3 Mp ECs | TIMM17A | 0.528 | 0.517 | 0.352 | 1.13e-17 | 3.8e-13 |
| Malignant epithelial | C3 Mp ECs | NOLC1 | 0.528 | 0.466 | 0.32 | 6.17e-14 | 2.08e-09 |
| Malignant epithelial | C3 Mp ECs | AKT1 | 0.527 | 0.301 | 0.181 | 1.04e-12 | 3.5e-08 |
| Malignant epithelial | C3 Mp ECs | DNAJC24 | 0.527 | 0.149 | 0.078 | 1.41e-09 | 4.74e-05 |
| Malignant epithelial | C3 Mp ECs | RCAN1 | 0.527 | 0.178 | 0.101 | 4e-09 | 0.000135 |
| Malignant epithelial | C3 Mp ECs | XPO6 | 0.527 | 0.206 | 0.131 | 1.94e-07 | 0.00654 |
| Malignant epithelial | C3 Mp ECs | NDUFA12 | 0.526 | 0.908 | 0.773 | 6.61e-38 | 2.23e-33 |
| Malignant epithelial | C3 Mp ECs | TAP2 | 0.526 | 0.49 | 0.292 | 3.41e-22 | 1.15e-17 |
| Malignant epithelial | C3 Mp ECs | BCL7C | 0.526 | 0.628 | 0.462 | 1.95e-20 | 6.57e-16 |
| Malignant epithelial | C3 Mp ECs | PTS | 0.526 | 0.557 | 0.413 | 8.73e-14 | 2.94e-09 |
| Malignant epithelial | C3 Mp ECs | BAP1 | 0.526 | 0.168 | 0.102 | 3.43e-07 | 0.0115 |
| Malignant epithelial | C3 Mp ECs | KBTBD2 | 0.526 | 0.212 | 0.141 | 1.35e-06 | 0.0456 |
| Malignant epithelial | C3 Mp ECs | DBI | 0.525 | 0.956 | 0.839 | 6.15e-44 | 2.07e-39 |
| Malignant epithelial | C3 Mp ECs | MRPL4 | 0.525 | 0.593 | 0.436 | 3.86e-16 | 1.3e-11 |
| Malignant epithelial | C3 Mp ECs | GLO1 | 0.524 | 0.629 | 0.458 | 3.54e-18 | 1.19e-13 |
| Malignant epithelial | C3 Mp ECs | BAZ1B | 0.524 | 0.399 | 0.264 | 2.34e-12 | 7.89e-08 |
| Malignant epithelial | C3 Mp ECs | LEPROTL1 | 0.524 | 0.41 | 0.282 | 7.49e-11 | 2.52e-06 |
| Malignant epithelial | C3 Mp ECs | SNUPN | 0.524 | 0.25 | 0.153 | 1.15e-09 | 3.88e-05 |
| Malignant epithelial | C3 Mp ECs | QSOX2 | 0.524 | 0.108 | 0.056 | 2.05e-07 | 0.00691 |
| Malignant epithelial | C3 Mp ECs | ACAT1 | 0.523 | 0.302 | 0.189 | 2.99e-11 | 1.01e-06 |
| Malignant epithelial | C3 Mp ECs | TSFM | 0.523 | 0.31 | 0.203 | 2.3e-10 | 7.76e-06 |
| Malignant epithelial | C3 Mp ECs | ANAPC7 | 0.523 | 0.243 | 0.147 | 8.82e-10 | 2.97e-05 |
| Malignant epithelial | C3 Mp ECs | MRPL16 | 0.522 | 0.447 | 0.312 | 5.09e-12 | 1.71e-07 |
| Malignant epithelial | C3 Mp ECs | FAIM | 0.522 | 0.247 | 0.16 | 2.13e-08 | 0.000716 |
| Malignant epithelial | C3 Mp ECs | TWISTNB | 0.521 | 0.513 | 0.34 | 3.17e-18 | 1.07e-13 |
| Malignant epithelial | C3 Mp ECs | WDFY1 | 0.521 | 0.243 | 0.159 | 1.05e-07 | 0.00354 |
| Malignant epithelial | C3 Mp ECs | MT-ND6 | 0.52 | 0.519 | 0.33 | 3.85e-20 | 1.3e-15 |
| Malignant epithelial | C3 Mp ECs | ICAM2 | 0.52 | 0.255 | 0.142 | 7.77e-13 | 2.62e-08 |
| Malignant epithelial | C3 Mp ECs | TPGS2 | 0.52 | 0.334 | 0.211 | 9.28e-12 | 3.13e-07 |
| Malignant epithelial | C3 Mp ECs | AAAS | 0.52 | 0.24 | 0.151 | 1.08e-08 | 0.000365 |
| Malignant epithelial | C3 Mp ECs | TRIM37 | 0.52 | 0.112 | 0.06 | 4.88e-07 | 0.0165 |
| Malignant epithelial | C3 Mp ECs | LINC00460 | 0.519 | 0.352 | 0.21 | 9.16e-15 | 3.09e-10 |
| Malignant epithelial | C3 Mp ECs | ATL2 | 0.519 | 0.201 | 0.121 | 9.75e-09 | 0.000328 |
| Malignant epithelial | C3 Mp ECs | C7orf50 | 0.518 | 0.754 | 0.578 | 9.159999999999999e-24 | 3.09e-19 |
| Malignant epithelial | C3 Mp ECs | NDUFA6 | 0.517 | 0.879 | 0.743 | 3.98e-35 | 1.34e-30 |
| Malignant epithelial | C3 Mp ECs | RALB | 0.517 | 0.549 | 0.367 | 2.99e-17 | 1.01e-12 |
| Malignant epithelial | C3 Mp ECs | DUSP12 | 0.517 | 0.244 | 0.155 | 6.53e-09 | 0.00022 |
| Malignant epithelial | C3 Mp ECs | PRPS2 | 0.516 | 0.382 | 0.247 | 6.96e-13 | 2.34e-08 |
| Malignant epithelial | C3 Mp ECs | SRRT | 0.516 | 0.28 | 0.177 | 3.27e-10 | 1.1e-05 |
| Malignant epithelial | C3 Mp ECs | SNRPC | 0.515 | 0.793 | 0.644 | 2.51e-26 | 8.45e-22 |

| compartment | subcluster | gene | avg_log2FC | pct.1 | pct.2 | p_val | p_val_adj |
|---|---|---|---|---|---|---|---|
| Malignant epithelial | C3 Mp ECs | GTF3C6 | 0.515 | 0.797 | 0.622 | 3.11e-24 | 1.05e-19 |
| Malignant epithelial | C3 Mp ECs | THOP1 | 0.515 | 0.232 | 0.132 | 2.07e-11 | 6.97e-07 |
| Malignant epithelial | C3 Mp ECs | CDCA7L | 0.515 | 0.274 | 0.174 | 2.48e-09 | 8.35e-05 |
| Malignant epithelial | C3 Mp ECs | AMN1 | 0.515 | 0.128 | 0.072 | 5.26e-07 | 0.0177 |
| Malignant epithelial | C3 Mp ECs | LSM3 | 0.514 | 0.778 | 0.589 | 1.41e-24 | 4.75e-20 |
| Malignant epithelial | C3 Mp ECs | ORMDL2 | 0.514 | 0.719 | 0.517 | 2.73e-24 | 9.19e-20 |
| Malignant epithelial | C3 Mp ECs | BRIX1 | 0.514 | 0.502 | 0.332 | 2.28e-17 | 7.69e-13 |
| Malignant epithelial | C3 Mp ECs | ARHGAP35 | 0.514 | 0.203 | 0.122 | 2.75e-08 | 0.000927 |
| Malignant epithelial | C3 Mp ECs | EMC4 | 0.513 | 0.756 | 0.578 | 9.51e-22 | 3.2e-17 |
| Malignant epithelial | C3 Mp ECs | ACBD6 | 0.513 | 0.415 | 0.266 | 3.38e-14 | 1.14e-09 |
| Malignant epithelial | C3 Mp ECs | TSEN15 | 0.513 | 0.276 | 0.16 | 1.07e-12 | 3.62e-08 |
| Malignant epithelial | C3 Mp ECs | GGCT | 0.512 | 0.721 | 0.563 | 1.69e-20 | 5.69e-16 |
| Malignant epithelial | C3 Mp ECs | TRIM47 | 0.511 | 0.45 | 0.288 | 2.29e-16 | 7.73e-12 |
| Malignant epithelial | C3 Mp ECs | PAK4 | 0.511 | 0.389 | 0.25 | 1.07e-13 | 3.6e-09 |
| Malignant epithelial | C3 Mp ECs | GMPS | 0.511 | 0.338 | 0.219 | 1.92e-11 | 6.46e-07 |
| Malignant epithelial | C3 Mp ECs | ANAPC1 | 0.511 | 0.19 | 0.116 | 7.31e-08 | 0.00246 |
| Malignant epithelial | C3 Mp ECs | STIP1 | 0.51 | 0.527 | 0.347 | 1.38e-17 | 4.65e-13 |
| Malignant epithelial | C3 Mp ECs | ARHGAP4 | 0.51 | 0.262 | 0.155 | 1.65e-11 | 5.56e-07 |
| Malignant epithelial | C3 Mp ECs | SNRNP40 | 0.51 | 0.288 | 0.183 | 6.87e-10 | 2.32e-05 |
| Malignant epithelial | C3 Mp ECs | QTRTD1 | 0.51 | 0.17 | 0.103 | 2.64e-07 | 0.00891 |
| Malignant epithelial | C3 Mp ECs | FAHD2A | 0.51 | 0.208 | 0.135 | 1.4e-06 | 0.0471 |
| Malignant epithelial | C3 Mp ECs | SLC7A5 | 0.509 | 0.385 | 0.246 | 1.92e-14 | 6.47e-10 |
| Malignant epithelial | C3 Mp ECs | HMCES | 0.509 | 0.354 | 0.22 | 1.9e-13 | 6.41e-09 |
| Malignant epithelial | C3 Mp ECs | PRR7 | 0.509 | 0.414 | 0.286 | 1.44e-11 | 4.84e-07 |
| Malignant epithelial | C3 Mp ECs | COTL1 | 0.508 | 0.655 | 0.477 | 7.03e-20 | 2.37e-15 |
| Malignant epithelial | C3 Mp ECs | PSMC5 | 0.507 | 0.726 | 0.56 | 7.37e-22 | 2.48e-17 |
| Malignant epithelial | C3 Mp ECs | APOL6 | 0.507 | 0.375 | 0.229 | 4.13e-15 | 1.39e-10 |
| Malignant epithelial | C3 Mp ECs | UBAC1 | 0.507 | 0.287 | 0.18 | 2.99e-10 | 1.01e-05 |
| Malignant epithelial | C3 Mp ECs | CD163L1 | 0.507 | 0.27 | 0.17 | 8.66e-10 | 2.92e-05 |
| Malignant epithelial | C3 Mp ECs | RP11-119B16.2 | 0.507 | 0.11 | 0.057 | 1.1e-07 | 0.00371 |
| Malignant epithelial | C3 Mp ECs | ZDHHC16 | 0.507 | 0.236 | 0.157 | 2.84e-07 | 0.00959 |
| Malignant epithelial | C3 Mp ECs | STAM | 0.507 | 0.141 | 0.081 | 4.72e-07 | 0.0159 |
| Malignant epithelial | C3 Mp ECs | CNN2 | 0.506 | 0.77 | 0.56 | 3.91e-26 | 1.32e-21 |
| Malignant epithelial | C3 Mp ECs | LINC00116 | 0.506 | 0.662 | 0.475 | 1.7e-19 | 5.72e-15 |
| Malignant epithelial | C3 Mp ECs | GRPEL1 | 0.506 | 0.418 | 0.278 | 2.51e-13 | 8.47e-09 |
| Malignant epithelial | C3 Mp ECs | ZDHHC6 | 0.506 | 0.363 | 0.234 | 2.21e-12 | 7.43e-08 |
| Malignant epithelial | C3 Mp ECs | DCTN3 | 0.505 | 0.654 | 0.494 | 3.62e-17 | 1.22e-12 |
| Malignant epithelial | C3 Mp ECs | C1orf35 | 0.505 | 0.527 | 0.358 | 9.11e-16 | 3.07e-11 |
| Malignant epithelial | C3 Mp ECs | BDP1 | 0.505 | 0.393 | 0.261 | 4.47e-12 | 1.51e-07 |
| Malignant epithelial | C3 Mp ECs | CREG2 | 0.505 | 0.11 | 0.055 | 4.64e-08 | 0.00156 |
| Malignant epithelial | C3 Mp ECs | TRIM28 | 0.504 | 0.534 | 0.364 | 2.5e-16 | 8.44e-12 |
| Malignant epithelial | C3 Mp ECs | COQ6 | 0.504 | 0.17 | 0.098 | 3.45e-08 | 0.00116 |
| Malignant epithelial | C3 Mp ECs | OSBPL5 | 0.504 | 0.143 | 0.079 | 5.84e-08 | 0.00197 |
| Malignant epithelial | C3 Mp ECs | RBM10 | 0.504 | 0.261 | 0.171 | 1.1e-07 | 0.00371 |
| Malignant epithelial | C3 Mp ECs | COA1 | 0.503 | 0.506 | 0.335 | 3.21e-16 | 1.08e-11 |
| Malignant epithelial | C3 Mp ECs | LMF2 | 0.503 | 0.314 | 0.213 | 2.88e-09 | 9.7e-05 |
| Malignant epithelial | C3 Mp ECs | TAF6 | 0.503 | 0.291 | 0.192 | 6.24e-09 | 0.00021 |
| Malignant epithelial | C3 Mp ECs | ILF3 | 0.502 | 0.661 | 0.461 | 2.04e-22 | 6.88e-18 |
| Malignant epithelial | C3 Mp ECs | HAUS2 | 0.502 | 0.234 | 0.14 | 3.36e-10 | 1.13e-05 |
| Malignant epithelial | C3 Mp ECs | ERAP2 | 0.502 | 0.252 | 0.156 | 1.45e-09 | 4.9e-05 |
| Malignant epithelial | C3 Mp ECs | EFTUD2 | 0.502 | 0.255 | 0.168 | 3.42e-08 | 0.00115 |
| Malignant epithelial | C3 Mp ECs | TAGLN2 | 0.501 | 0.948 | 0.804 | 1.21e-43 | 4.07e-39 |
| Malignant epithelial | C3 Mp ECs | TCF3 | 0.501 | 0.269 | 0.158 | 4.59e-12 | 1.55e-07 |
| Malignant epithelial | C3 Mp ECs | ARF6 | 0.5 | 0.868 | 0.737 | 6.01e-27 | 2.03e-22 |
| Malignant epithelial | C3 Mp ECs | RNF126 | 0.5 | 0.601 | 0.431 | 1.25e-16 | 4.21e-12 |
| Malignant epithelial | C3 Mp ECs | CAPRIN2 | 0.5 | 0.164 | 0.089 | 1.15e-09 | 3.86e-05 |
| Malignant epithelial | C3 Mp ECs | SF3B4 | 0.499 | 0.469 | 0.309 | 2.49e-15 | 8.4e-11 |
| Malignant epithelial | C3 Mp ECs | TRIOBP | 0.499 | 0.383 | 0.249 | 2.52e-12 | 8.48e-08 |
| Malignant epithelial | C3 Mp ECs | RP11-545E17.3 | 0.499 | 0.124 | 0.069 | 4.72e-07 | 0.0159 |
| Malignant epithelial | C3 Mp ECs | SRP9 | 0.498 | 0.847 | 0.688 | 3.55e-27 | 1.2e-22 |
| Malignant epithelial | C3 Mp ECs | POLE4 | 0.498 | 0.661 | 0.488 | 4.9e-19 | 1.65e-14 |
| Malignant epithelial | C3 Mp ECs | RAB8A | 0.498 | 0.526 | 0.335 | 6.58e-19 | 2.22e-14 |
| Malignant epithelial | C3 Mp ECs | ASH2L | 0.498 | 0.192 | 0.112 | 6.98e-09 | 0.000235 |
| Malignant epithelial | C3 Mp ECs | MIA | 0.497 | 0.817 | 0.54 | 2.74e-33 | 9.25e-29 |

| compartment | subcluster | gene | avg_log2FC | pct.1 | pct.2 | p_val | p_val_adj |
|---|---|---|---|---|---|---|---|
| Malignant epithelial | C3 Mp ECs | FBXO22 | 0.497 | 0.331 | 0.206 | 3.02e-12 | 1.02e-07 |
| Malignant epithelial | C3 Mp ECs | TOR1A | 0.497 | 0.258 | 0.163 | 7.13e-09 | 0.00024 |
| Malignant epithelial | C3 Mp ECs | HSPA14 | 0.496 | 0.188 | 0.118 | 5.24e-07 | 0.0177 |
| Malignant epithelial | C3 Mp ECs | PPIA | 0.495 | 1.0 | 0.982 | 7.7e-89 | 2.59e-84 |
| Malignant epithelial | C3 Mp ECs | EBP | 0.495 | 0.701 | 0.496 | 1.08e-23 | 3.64e-19 |
| Malignant epithelial | C3 Mp ECs | EIF2AK2 | 0.495 | 0.509 | 0.343 | 2.06e-15 | 6.94e-11 |
| Malignant epithelial | C3 Mp ECs | SNW1 | 0.495 | 0.561 | 0.393 | 3.55e-15 | 1.2e-10 |
| Malignant epithelial | C3 Mp ECs | TMEM171 | 0.495 | 0.186 | 0.106 | 1.68e-09 | 5.66e-05 |
| Malignant epithelial | C3 Mp ECs | SEPT8 | 0.495 | 0.201 | 0.13 | 4.95e-07 | 0.0167 |
| Malignant epithelial | C3 Mp ECs | HMGN1 | 0.494 | 0.963 | 0.89 | 5.200000000000001e-44 | 1.75e-39 |
| Malignant epithelial | C3 Mp ECs | DKC1 | 0.494 | 0.528 | 0.364 | 7.38e-15 | 2.49e-10 |
| Malignant epithelial | C3 Mp ECs | SLC16A3 | 0.494 | 0.516 | 0.362 | 1.09e-14 | 3.66e-10 |
| Malignant epithelial | C3 Mp ECs | PYCR2 | 0.494 | 0.287 | 0.175 | 2.04e-11 | 6.89e-07 |
| Malignant epithelial | C3 Mp ECs | UBE2Q2 | 0.494 | 0.288 | 0.181 | 3.08e-10 | 1.04e-05 |
| Malignant epithelial | C3 Mp ECs | AAED1 | 0.494 | 0.273 | 0.175 | 2.5e-09 | 8.43e-05 |
| Malignant epithelial | C3 Mp ECs | TMA7 | 0.492 | 0.99 | 0.959 | 6.55e-57 | 2.21e-52 |
| Malignant epithelial | C3 Mp ECs | EMC8 | 0.492 | 0.455 | 0.294 | 2.06e-15 | 6.93e-11 |
| Malignant epithelial | C3 Mp ECs | IDO1 | 0.491 | 0.254 | 0.152 | 6.61e-11 | 2.23e-06 |
| Malignant epithelial | C3 Mp ECs | RPP38 | 0.491 | 0.281 | 0.179 | 9.44e-10 | 3.18e-05 |
| Malignant epithelial | C3 Mp ECs | ISY1 | 0.491 | 0.303 | 0.202 | 3.22e-09 | 0.000109 |
| Malignant epithelial | C3 Mp ECs | ACACA | 0.491 | 0.152 | 0.087 | 8.94e-08 | 0.00301 |
| Malignant epithelial | C3 Mp ECs | C17orf89 | 0.49 | 0.743 | 0.599 | 5.29e-19 | 1.78e-14 |
| Malignant epithelial | C3 Mp ECs | NUDT5 | 0.49 | 0.585 | 0.422 | 1.11e-15 | 3.72e-11 |
| Malignant epithelial | C3 Mp ECs | IFI27L1 | 0.49 | 0.434 | 0.3 | 8.18e-12 | 2.76e-07 |
| Malignant epithelial | C3 Mp ECs | C19orf52 | 0.49 | 0.251 | 0.17 | 4.23e-07 | 0.0142 |
| Malignant epithelial | C3 Mp ECs | COA6 | 0.489 | 0.567 | 0.386 | 1.27e-16 | 4.28e-12 |
| Malignant epithelial | C3 Mp ECs | MRPS9 | 0.489 | 0.287 | 0.187 | 2.68e-09 | 9.03e-05 |
| Malignant epithelial | C3 Mp ECs | GSPT1 | 0.488 | 0.676 | 0.504 | 2.43e-18 | 8.17e-14 |
| Malignant epithelial | C3 Mp ECs | CD320 | 0.488 | 0.486 | 0.316 | 1.24e-17 | 4.17e-13 |
| Malignant epithelial | C3 Mp ECs | NOC4L | 0.488 | 0.31 | 0.2 | 3.85e-10 | 1.3e-05 |
| Malignant epithelial | C3 Mp ECs | DDX21 | 0.487 | 0.841 | 0.687 | 3.34e-25 | 1.12e-20 |
| Malignant epithelial | C3 Mp ECs | CLPP | 0.487 | 0.571 | 0.412 | 8.98e-15 | 3.03e-10 |
| Malignant epithelial | C3 Mp ECs | UBTD1 | 0.487 | 0.258 | 0.159 | 1.43e-09 | 4.81e-05 |
| Malignant epithelial | C3 Mp ECs | HLA-DQB1 | 0.487 | 0.33 | 0.222 | 2.01e-09 | 6.76e-05 |
| Malignant epithelial | C3 Mp ECs | ISCA2 | 0.486 | 0.448 | 0.299 | 6.79e-14 | 2.29e-09 |
| Malignant epithelial | C3 Mp ECs | SMARCB1 | 0.486 | 0.491 | 0.349 | 3.58e-13 | 1.21e-08 |
| Malignant epithelial | C3 Mp ECs | NDUFAF4 | 0.485 | 0.428 | 0.295 | 7.41e-11 | 2.5e-06 |
| Malignant epithelial | C3 Mp ECs | HYI | 0.485 | 0.328 | 0.214 | 7.45e-11 | 2.51e-06 |
| Malignant epithelial | C3 Mp ECs | TST | 0.485 | 0.457 | 0.339 | 3.12e-09 | 0.000105 |
| Malignant epithelial | C3 Mp ECs | ABHD14A | 0.485 | 0.302 | 0.199 | 4.13e-09 | 0.000139 |
| Malignant epithelial | C3 Mp ECs | CDKN2A | 0.484 | 0.916 | 0.739 | 1.45e-33 | 4.88e-29 |
| Malignant epithelial | C3 Mp ECs | MRPL17 | 0.484 | 0.657 | 0.474 | 6.58e-19 | 2.22e-14 |
| Malignant epithelial | C3 Mp ECs | PTPMT1 | 0.484 | 0.589 | 0.424 | 5.9e-17 | 1.99e-12 |
| Malignant epithelial | C3 Mp ECs | DNM1L | 0.484 | 0.246 | 0.157 | 1.3e-08 | 0.000439 |
| Malignant epithelial | C3 Mp ECs | SKAP1 | 0.484 | 0.19 | 0.116 | 9.49e-08 | 0.0032 |
| Malignant epithelial | C3 Mp ECs | SPON1 | 0.483 | 0.45 | 0.27 | 6.34e-19 | 2.14e-14 |
| Malignant epithelial | C3 Mp ECs | CASP8 | 0.483 | 0.419 | 0.271 | 2.51e-14 | 8.45e-10 |
| Malignant epithelial | C3 Mp ECs | MRPS10 | 0.482 | 0.542 | 0.4 | 2.93e-13 | 9.88e-09 |
| Malignant epithelial | C3 Mp ECs | WDR18 | 0.481 | 0.423 | 0.283 | 2.61e-13 | 8.81e-09 |
| Malignant epithelial | C3 Mp ECs | VKORC1 | 0.48 | 0.666 | 0.485 | 5.73e-18 | 1.93e-13 |
| Malignant epithelial | C3 Mp ECs | CPSF6 | 0.48 | 0.393 | 0.257 | 5e-12 | 1.68e-07 |
| Malignant epithelial | C3 Mp ECs | TARBP2 | 0.48 | 0.204 | 0.123 | 1.03e-08 | 0.000349 |
| Malignant epithelial | C3 Mp ECs | SH3PXD2A-AS1 | 0.48 | 0.152 | 0.086 | 1.16e-07 | 0.00392 |
| Malignant epithelial | C3 Mp ECs | TMSB4X | 0.479 | 1.0 | 1.0 | 5.229999999999999e-67 | 1.76e-62 |
| Malignant epithelial | C3 Mp ECs | NAA20 | 0.479 | 0.792 | 0.604 | 3.45e-24 | 1.16e-19 |
| Malignant epithelial | C3 Mp ECs | EXOSC8 | 0.479 | 0.434 | 0.299 | 9.86e-12 | 3.32e-07 |
| Malignant epithelial | C3 Mp ECs | SEMA4B | 0.479 | 0.294 | 0.194 | 3.49e-09 | 0.000117 |
| Malignant epithelial | C3 Mp ECs | SDC1 | 0.478 | 0.703 | 0.513 | 6.47e-19 | 2.18e-14 |
| Malignant epithelial | C3 Mp ECs | NSL1 | 0.478 | 0.234 | 0.147 | 1.8e-08 | 0.000605 |
| Malignant epithelial | C3 Mp ECs | LAMA3 | 0.477 | 0.586 | 0.388 | 2.2e-21 | 7.4e-17 |
| Malignant epithelial | C3 Mp ECs | HNRNPU | 0.476 | 0.84 | 0.667 | 7.729999999999999e-24 | 2.6e-19 |
| Malignant epithelial | C3 Mp ECs | SNAPC5 | 0.476 | 0.32 | 0.203 | 2.43e-11 | 8.18e-07 |
| Malignant epithelial | C3 Mp ECs | AIM1 | 0.476 | 0.185 | 0.11 | 2.16e-08 | 0.000727 |
| Malignant epithelial | C3 Mp ECs | NUP98 | 0.476 | 0.185 | 0.114 | 3.1e-07 | 0.0104 |

| compartment | subcluster | gene | avg_log2FC | pct.1 | pct.2 | p_val | p_val_adj |
|---|---|---|---|---|---|---|---|
| Malignant epithelial | C3 Mp ECs | FBL | 0.475 | 0.825 | 0.658 | 1.05e-23 | 3.55e-19 |
| Malignant epithelial | C3 Mp ECs | TRNAU1AP | 0.475 | 0.192 | 0.121 | 4.43e-07 | 0.0149 |
| Malignant epithelial | C3 Mp ECs | MDFI | 0.473 | 0.259 | 0.156 | 1.39e-10 | 4.68e-06 |
| Malignant epithelial | C3 Mp ECs | ANKRD13D | 0.473 | 0.251 | 0.161 | 8.69e-09 | 0.000293 |
| Malignant epithelial | C3 Mp ECs | NUDT3 | 0.472 | 0.461 | 0.316 | 1.58e-12 | 5.32e-08 |
| Malignant epithelial | C3 Mp ECs | KEAP1 | 0.472 | 0.308 | 0.202 | 2.56e-09 | 8.63e-05 |
| Malignant epithelial | C3 Mp ECs | STT3A | 0.472 | 0.243 | 0.162 | 5.07e-07 | 0.0171 |
| Malignant epithelial | C3 Mp ECs | EIF4EBP1 | 0.471 | 0.858 | 0.675 | 1.04e-28 | 3.51e-24 |
| Malignant epithelial | C3 Mp ECs | MRPL27 | 0.471 | 0.719 | 0.526 | 6.39e-21 | 2.15e-16 |
| Malignant epithelial | C3 Mp ECs | TAP1 | 0.471 | 0.695 | 0.507 | 5.27e-19 | 1.78e-14 |
| Malignant epithelial | C3 Mp ECs | RABEPK | 0.471 | 0.229 | 0.14 | 7.71e-09 | 0.00026 |
| Malignant epithelial | C3 Mp ECs | FAM43A | 0.471 | 0.177 | 0.101 | 8.25e-09 | 0.000278 |
| Malignant epithelial | C3 Mp ECs | MFAP1 | 0.471 | 0.239 | 0.153 | 2.24e-08 | 0.000754 |
| Malignant epithelial | C3 Mp ECs | PLOD1 | 0.471 | 0.243 | 0.157 | 5.2e-08 | 0.00175 |
| Malignant epithelial | C3 Mp ECs | EIF2AK4 | 0.471 | 0.281 | 0.194 | 2.52e-07 | 0.00849 |
| Malignant epithelial | C3 Mp ECs | LINC00152 | 0.47 | 0.829 | 0.59 | 2.7e-33 | 9.080000000000001e-29 |
| Malignant epithelial | C3 Mp ECs | RBX1 | 0.47 | 0.913 | 0.791 | 6.32e-32 | 2.13e-27 |
| Malignant epithelial | C3 Mp ECs | PPP1CA | 0.47 | 0.844 | 0.688 | 1.01e-28 | 3.41e-24 |
| Malignant epithelial | C3 Mp ECs | ZNF217 | 0.47 | 0.35 | 0.231 | 8.08e-11 | 2.72e-06 |
| Malignant epithelial | C3 Mp ECs | BCAP29 | 0.47 | 0.39 | 0.278 | 1.43e-09 | 4.83e-05 |
| Malignant epithelial | C3 Mp ECs | LSM2 | 0.469 | 0.822 | 0.647 | 7.329999999999999e-25 | 2.47e-20 |
| Malignant epithelial | C3 Mp ECs | FXN | 0.469 | 0.262 | 0.165 | 2.23e-09 | 7.5e-05 |
| Malignant epithelial | C3 Mp ECs | DDX24 | 0.468 | 0.646 | 0.466 | 1.06e-16 | 3.58e-12 |
| Malignant epithelial | C3 Mp ECs | NOP16 | 0.468 | 0.527 | 0.358 | 1.27e-14 | 4.29e-10 |
| Malignant epithelial | C3 Mp ECs | PDCD11 | 0.468 | 0.163 | 0.097 | 3.81e-07 | 0.0128 |
| Malignant epithelial | C3 Mp ECs | CCT7 | 0.467 | 0.732 | 0.556 | 9.25e-20 | 3.12e-15 |
| Malignant epithelial | C3 Mp ECs | ANXA5 | 0.467 | 0.674 | 0.508 | 1.24e-19 | 4.19e-15 |
| Malignant epithelial | C3 Mp ECs | COL17A1 | 0.467 | 0.688 | 0.487 | 1.55e-19 | 5.23e-15 |
| Malignant epithelial | C3 Mp ECs | NUDT21 | 0.467 | 0.648 | 0.468 | 3.25e-18 | 1.1e-13 |
| Malignant epithelial | C3 Mp ECs | UBE2V2 | 0.466 | 0.681 | 0.512 | 1.61e-18 | 5.43e-14 |
| Malignant epithelial | C3 Mp ECs | ASCC3 | 0.466 | 0.232 | 0.142 | 2.92e-09 | 9.85e-05 |
| Malignant epithelial | C3 Mp ECs | LAS1L | 0.466 | 0.236 | 0.15 | 2.45e-08 | 0.000825 |
| Malignant epithelial | C3 Mp ECs | MRPL15 | 0.465 | 0.712 | 0.548 | 2.09e-19 | 7.05e-15 |
| Malignant epithelial | C3 Mp ECs | XRCC6 | 0.465 | 0.724 | 0.553 | 9.39e-19 | 3.16e-14 |
| Malignant epithelial | C3 Mp ECs | NDUFA9 | 0.464 | 0.611 | 0.462 | 1.3e-15 | 4.37e-11 |
| Malignant epithelial | C3 Mp ECs | AAMDC | 0.464 | 0.514 | 0.375 | 7.01e-12 | 2.36e-07 |
| Malignant epithelial | C3 Mp ECs | PMPCA | 0.464 | 0.326 | 0.217 | 8.91e-10 | 3e-05 |
| Malignant epithelial | C3 Mp ECs | FAM208B | 0.464 | 0.214 | 0.131 | 2.08e-08 | 0.000699 |
| Malignant epithelial | C3 Mp ECs | RABL2B | 0.464 | 0.156 | 0.094 | 1.13e-06 | 0.038 |
| Malignant epithelial | C3 Mp ECs | PFN1 | 0.463 | 0.99 | 0.961 | 3.71e-62 | 1.25e-57 |
| Malignant epithelial | C3 Mp ECs | MRPL52 | 0.463 | 0.932 | 0.801 | 4.12e-34 | 1.39e-29 |
| Malignant epithelial | C3 Mp ECs | PRKAG2-AS1 | 0.463 | 0.149 | 0.086 | 3.3e-07 | 0.0111 |
| Malignant epithelial | C3 Mp ECs | NKG7 | 0.463 | 0.229 | 0.149 | 5.58e-07 | 0.0188 |
| Malignant epithelial | C3 Mp ECs | SRSF2 | 0.462 | 0.932 | 0.788 | 6.100000000000001e-38 | 2.06e-33 |
| Malignant epithelial | C3 Mp ECs | ARMC1 | 0.462 | 0.284 | 0.187 | 1.12e-08 | 0.000379 |
| Malignant epithelial | C3 Mp ECs | MTFP1 | 0.462 | 0.232 | 0.155 | 6.54e-07 | 0.022 |
| Malignant epithelial | C3 Mp ECs | RPP30 | 0.462 | 0.234 | 0.156 | 1.2e-06 | 0.0404 |
| Malignant epithelial | C3 Mp ECs | H3F3B | 0.461 | 0.997 | 0.98 | 7.06e-56 | 2.38e-51 |
| Malignant epithelial | C3 Mp ECs | IFITM3 | 0.461 | 0.99 | 0.942 | 2.95e-38 | 9.96e-34 |
| Malignant epithelial | C3 Mp ECs | RCE1 | 0.461 | 0.346 | 0.225 | 2.2e-11 | 7.41e-07 |
| Malignant epithelial | C3 Mp ECs | DPH6 | 0.461 | 0.262 | 0.167 | 9.26e-09 | 0.000312 |
| Malignant epithelial | C3 Mp ECs | ARL2 | 0.46 | 0.745 | 0.558 | 1.83e-20 | 6.16e-16 |
| Malignant epithelial | C3 Mp ECs | HN1L | 0.46 | 0.444 | 0.303 | 8.34e-13 | 2.81e-08 |
| Malignant epithelial | C3 Mp ECs | PSMD14 | 0.46 | 0.469 | 0.323 | 3.88e-12 | 1.31e-07 |
| Malignant epithelial | C3 Mp ECs | KLF2 | 0.459 | 0.592 | 0.388 | 1.31e-20 | 4.42e-16 |
| Malignant epithelial | C3 Mp ECs | MRPS18C | 0.459 | 0.539 | 0.38 | 2.14e-13 | 7.2e-09 |
| Malignant epithelial | C3 Mp ECs | CARM1 | 0.459 | 0.277 | 0.177 | 1.35e-09 | 4.54e-05 |
| Malignant epithelial | C3 Mp ECs | RBMX | 0.458 | 0.763 | 0.612 | 2.51e-18 | 8.46e-14 |
| Malignant epithelial | C3 Mp ECs | PHB2 | 0.458 | 0.714 | 0.58 | 2.4e-17 | 8.08e-13 |
| Malignant epithelial | C3 Mp ECs | SNRPA | 0.458 | 0.597 | 0.414 | 7.46e-17 | 2.51e-12 |
| Malignant epithelial | C3 Mp ECs | NR2C1 | 0.458 | 0.255 | 0.155 | 1.73e-10 | 5.82e-06 |
| Malignant epithelial | C3 Mp ECs | SNTB2 | 0.458 | 0.192 | 0.12 | 3.54e-07 | 0.0119 |
| Malignant epithelial | C3 Mp ECs | GAPDH | 0.457 | 1.0 | 0.998 | 7.06e-65 | 2.38e-60 |
| Malignant epithelial | C3 Mp ECs | VMP1 | 0.457 | 0.935 | 0.842 | 5.39e-24 | 1.82e-19 |

| compartment | subcluster | gene | avg_log2FC | pct.1 | pct.2 | p_val | p_val_adj |
|---|---|---|---|---|---|---|---|
| Malignant epithelial | C3 Mp ECs | ISOC2 | 0.457 | 0.61 | 0.449 | 8.24e-16 | 2.78e-11 |
| Malignant epithelial | C3 Mp ECs | COPS7A | 0.457 | 0.356 | 0.226 | 5.31e-12 | 1.79e-07 |
| Malignant epithelial | C3 Mp ECs | ATG3 | 0.457 | 0.49 | 0.347 | 1.02e-11 | 3.45e-07 |
| Malignant epithelial | C3 Mp ECs | WNT5B | 0.457 | 0.128 | 0.067 | 3.71e-08 | 0.00125 |
| Malignant epithelial | C3 Mp ECs | RQCD1 | 0.457 | 0.233 | 0.147 | 3.92e-08 | 0.00132 |
| Malignant epithelial | C3 Mp ECs | MRPS2 | 0.456 | 0.521 | 0.382 | 8.69e-12 | 2.93e-07 |
| Malignant epithelial | C3 Mp ECs | LYSMD2 | 0.456 | 0.343 | 0.222 | 1.67e-10 | 5.62e-06 |
| Malignant epithelial | C3 Mp ECs | FN3KRP | 0.456 | 0.212 | 0.139 | 1.11e-06 | 0.0372 |
| Malignant epithelial | C3 Mp ECs | VPS29 | 0.455 | 0.786 | 0.621 | 3.36e-22 | 1.13e-17 |
| Malignant epithelial | C3 Mp ECs | DRG1 | 0.455 | 0.425 | 0.297 | 1.06e-10 | 3.56e-06 |
| Malignant epithelial | C3 Mp ECs | CLPTM1 | 0.455 | 0.408 | 0.284 | 1.98e-10 | 6.68e-06 |
| Malignant epithelial | C3 Mp ECs | NOL8 | 0.455 | 0.25 | 0.158 | 6.87e-09 | 0.000231 |
| Malignant epithelial | C3 Mp ECs | AP1M1 | 0.455 | 0.183 | 0.108 | 1.88e-08 | 0.000633 |
| Malignant epithelial | C3 Mp ECs | TECR | 0.454 | 0.858 | 0.73 | 4.52e-24 | 1.52e-19 |
| Malignant epithelial | C3 Mp ECs | CYC1 | 0.454 | 0.863 | 0.75 | 1.01e-21 | 3.4e-17 |
| Malignant epithelial | C3 Mp ECs | PSMB6 | 0.454 | 0.753 | 0.572 | 1.32e-19 | 4.44e-15 |
| Malignant epithelial | C3 Mp ECs | LRRC59 | 0.454 | 0.575 | 0.395 | 6.65e-16 | 2.24e-11 |
| Malignant epithelial | C3 Mp ECs | MRPS12 | 0.454 | 0.706 | 0.54 | 1.79e-15 | 6.03e-11 |
| Malignant epithelial | C3 Mp ECs | EBNA1BP2 | 0.454 | 0.432 | 0.306 | 1.17e-11 | 3.96e-07 |
| Malignant epithelial | C3 Mp ECs | ARHGDIA | 0.453 | 0.793 | 0.624 | 1.64e-20 | 5.53e-16 |
| Malignant epithelial | C3 Mp ECs | ABCF1 | 0.453 | 0.517 | 0.357 | 6.2e-14 | 2.09e-09 |
| Malignant epithelial | C3 Mp ECs | PSMG1 | 0.453 | 0.432 | 0.283 | 7.24e-14 | 2.44e-09 |
| Malignant epithelial | C3 Mp ECs | PNKP | 0.453 | 0.411 | 0.283 | 3.45e-11 | 1.16e-06 |
| Malignant epithelial | C3 Mp ECs | STIM1 | 0.453 | 0.229 | 0.134 | 2.99e-10 | 1.01e-05 |
| Malignant epithelial | C3 Mp ECs | ARV1 | 0.453 | 0.228 | 0.138 | 4.54e-09 | 0.000153 |
| Malignant epithelial | C3 Mp ECs | CASP6 | 0.453 | 0.299 | 0.203 | 7.82e-08 | 0.00264 |
| Malignant epithelial | C3 Mp ECs | NAA15 | 0.452 | 0.281 | 0.188 | 3.54e-08 | 0.00119 |
| Malignant epithelial | C3 Mp ECs | ING2 | 0.452 | 0.32 | 0.223 | 6.41e-08 | 0.00216 |
| Malignant epithelial | C3 Mp ECs | ENO1 | 0.451 | 0.898 | 0.782 | 2.17e-25 | 7.320000000000001e-21 |
| Malignant epithelial | C3 Mp ECs | MARVELD1 | 0.451 | 0.248 | 0.157 | 8e-09 | 0.00027 |
| Malignant epithelial | C3 Mp ECs | TIMM8B | 0.45 | 0.799 | 0.656 | 1.94e-21 | 6.54e-17 |
| Malignant epithelial | C3 Mp ECs | HLA-DRB1 | 0.45 | 0.759 | 0.593 | 1.25e-18 | 4.2e-14 |
| Malignant epithelial | C3 Mp ECs | PRR34-AS1 | 0.45 | 0.474 | 0.32 | 5.84e-14 | 1.97e-09 |
| Malignant epithelial | C3 Mp ECs | TCERG1 | 0.45 | 0.379 | 0.256 | 1.05e-10 | 3.52e-06 |
| Malignant epithelial | C3 Mp ECs | PSME3 | 0.45 | 0.341 | 0.228 | 6.78e-10 | 2.28e-05 |
| Malignant epithelial | C3 Mp ECs | HSPE1 | 0.449 | 0.981 | 0.93 | 3.02e-40 | 1.02e-35 |
| Malignant epithelial | C3 Mp ECs | TM4SF1 | 0.448 | 0.963 | 0.885 | 8.38e-30 | 2.82e-25 |
| Malignant epithelial | C3 Mp ECs | NMI | 0.448 | 0.43 | 0.275 | 1.32e-14 | 4.45e-10 |
| Malignant epithelial | C3 Mp ECs | ALG8 | 0.448 | 0.317 | 0.205 | 3.22e-10 | 1.08e-05 |
| Malignant epithelial | C3 Mp ECs | TIMM8A | 0.448 | 0.292 | 0.192 | 7.38e-09 | 0.000249 |
| Malignant epithelial | C3 Mp ECs | BORCS8 | 0.448 | 0.214 | 0.136 | 1.81e-07 | 0.00611 |
| Malignant epithelial | C3 Mp ECs | BST2 | 0.447 | 0.913 | 0.726 | 2.29e-31 | 7.7e-27 |
| Malignant epithelial | C3 Mp ECs | NTMT1 | 0.447 | 0.571 | 0.405 | 2.32e-15 | 7.81e-11 |
| Malignant epithelial | C3 Mp ECs | DENR | 0.447 | 0.567 | 0.425 | 1.92e-12 | 6.49e-08 |
| Malignant epithelial | C3 Mp ECs | GART | 0.447 | 0.258 | 0.174 | 1.89e-07 | 0.00635 |
| Malignant epithelial | C3 Mp ECs | YY1AP1 | 0.446 | 0.244 | 0.157 | 4.44e-08 | 0.0015 |
| Malignant epithelial | C3 Mp ECs | MIF | 0.445 | 0.994 | 0.942 | 6.2e-50 | 2.09e-45 |
| Malignant epithelial | C3 Mp ECs | CCNH | 0.445 | 0.368 | 0.256 | 1.4e-09 | 4.72e-05 |
| Malignant epithelial | C3 Mp ECs | C12orf29 | 0.445 | 0.183 | 0.117 | 1.35e-06 | 0.0455 |
| Malignant epithelial | C3 Mp ECs | PHB | 0.444 | 0.83 | 0.716 | 2.66e-20 | 8.98e-16 |
| Malignant epithelial | C3 Mp ECs | RIF1 | 0.444 | 0.248 | 0.161 | 1.17e-07 | 0.00393 |
| Malignant epithelial | C3 Mp ECs | RBFOX2 | 0.444 | 0.27 | 0.182 | 1.36e-07 | 0.00457 |
| Malignant epithelial | C3 Mp ECs | MED27 | 0.444 | 0.171 | 0.103 | 2.38e-07 | 0.00803 |
| Malignant epithelial | C3 Mp ECs | CD52 | 0.444 | 0.326 | 0.234 | 4.83e-07 | 0.0163 |
| Malignant epithelial | C3 Mp ECs | MALSU1 | 0.443 | 0.546 | 0.388 | 1.07e-14 | 3.6e-10 |
| Malignant epithelial | C3 Mp ECs | SQLE | 0.443 | 0.37 | 0.231 | 1.11e-13 | 3.75e-09 |
| Malignant epithelial | C3 Mp ECs | DEF6 | 0.443 | 0.17 | 0.105 | 1.35e-06 | 0.0456 |
| Malignant epithelial | C3 Mp ECs | SNRPE | 0.442 | 0.937 | 0.841 | 1.62e-32 | 5.47e-28 |
| Malignant epithelial | C3 Mp ECs | IMP4 | 0.442 | 0.563 | 0.391 | 2.47e-14 | 8.31e-10 |
| Malignant epithelial | C3 Mp ECs | PRKRIR | 0.442 | 0.223 | 0.143 | 1.17e-07 | 0.00395 |
| Malignant epithelial | C3 Mp ECs | TXNL4A | 0.441 | 0.588 | 0.429 | 9.32e-15 | 3.14e-10 |
| Malignant epithelial | C3 Mp ECs | CFAP20 | 0.441 | 0.41 | 0.277 | 8.06e-12 | 2.72e-07 |
| Malignant epithelial | C3 Mp ECs | COLGALT1 | 0.441 | 0.297 | 0.185 | 6.2e-11 | 2.09e-06 |
| Malignant epithelial | C3 Mp ECs | ZNF830 | 0.441 | 0.284 | 0.189 | 2.83e-08 | 0.000953 |

| compartment | subcluster | gene | avg_log2FC | pct.1 | pct.2 | p_val | p_val_adj |
|---|---|---|---|---|---|---|---|
| Malignant epithelial | C3 Mp ECs | CEP89 | 0.441 | 0.182 | 0.112 | 4.72e-07 | 0.0159 |
| Malignant epithelial | C3 Mp ECs | PSMB7 | 0.44 | 0.865 | 0.706 | 9.97e-22 | 3.36e-17 |
| Malignant epithelial | C3 Mp ECs | FUBP1 | 0.439 | 0.301 | 0.203 | 3.1e-08 | 0.00104 |
| Malignant epithelial | C3 Mp ECs | CCDC94 | 0.439 | 0.214 | 0.131 | 3.69e-08 | 0.00124 |
| Malignant epithelial | C3 Mp ECs | REEP3 | 0.438 | 0.687 | 0.495 | 1.34e-19 | 4.52e-15 |
| Malignant epithelial | C3 Mp ECs | PNN | 0.437 | 0.639 | 0.485 | 1.14e-14 | 3.84e-10 |
| Malignant epithelial | C3 Mp ECs | NELFB | 0.437 | 0.328 | 0.222 | 4.12e-09 | 0.000139 |
| Malignant epithelial | C3 Mp ECs | UQCR10 | 0.436 | 0.963 | 0.896 | 6.87e-38 | 2.32e-33 |
| Malignant epithelial | C3 Mp ECs | DDX18 | 0.436 | 0.655 | 0.496 | 3.6e-13 | 1.21e-08 |
| Malignant epithelial | C3 Mp ECs | PRMT2 | 0.436 | 0.498 | 0.345 | 3.9e-13 | 1.31e-08 |
| Malignant epithelial | C3 Mp ECs | GPT2 | 0.436 | 0.168 | 0.103 | 9.8e-07 | 0.033 |
| Malignant epithelial | C3 Mp ECs | TRMT112 | 0.435 | 0.957 | 0.877 | 4.639999999999999e-34 | 1.56e-29 |
| Malignant epithelial | C3 Mp ECs | AHSA1 | 0.435 | 0.618 | 0.449 | 1.18e-15 | 3.98e-11 |
| Malignant epithelial | C3 Mp ECs | RPA2 | 0.434 | 0.208 | 0.136 | 1.21e-06 | 0.0408 |
| Malignant epithelial | C3 Mp ECs | BAX | 0.433 | 0.792 | 0.624 | 2.08e-21 | 7e-17 |
| Malignant epithelial | C3 Mp ECs | PMVK | 0.433 | 0.498 | 0.366 | 1.74e-11 | 5.86e-07 |
| Malignant epithelial | C3 Mp ECs | SRSF1 | 0.432 | 0.597 | 0.399 | 1.99e-19 | 6.71e-15 |
| Malignant epithelial | C3 Mp ECs | NPM3 | 0.432 | 0.703 | 0.519 | 2.36e-17 | 7.94e-13 |
| Malignant epithelial | C3 Mp ECs | MTHFS | 0.432 | 0.461 | 0.299 | 2.23e-14 | 7.52e-10 |
| Malignant epithelial | C3 Mp ECs | NFATC2IP | 0.432 | 0.335 | 0.226 | 2.18e-09 | 7.34e-05 |
| Malignant epithelial | C3 Mp ECs | MCMBP | 0.432 | 0.251 | 0.169 | 3.48e-07 | 0.0117 |
| Malignant epithelial | C3 Mp ECs | RPA1 | 0.432 | 0.166 | 0.101 | 6.85e-07 | 0.0231 |
| Malignant epithelial | C3 Mp ECs | SRGN | 0.431 | 0.55 | 0.45 | 6.35e-08 | 0.00214 |
| Malignant epithelial | C3 Mp ECs | PPP1R10 | 0.431 | 0.363 | 0.26 | 1.19e-07 | 0.004 |
| Malignant epithelial | C3 Mp ECs | TMSB10 | 0.43 | 1.0 | 1.0 | 2.94e-54 | 9.89e-50 |
| Malignant epithelial | C3 Mp ECs | IFT27 | 0.43 | 0.352 | 0.239 | 1.73e-09 | 5.84e-05 |
| Malignant epithelial | C3 Mp ECs | TIMM22 | 0.43 | 0.194 | 0.117 | 5.63e-08 | 0.0019 |
| Malignant epithelial | C3 Mp ECs | CMTM7 | 0.429 | 0.654 | 0.448 | 4.9e-21 | 1.65e-16 |
| Malignant epithelial | C3 Mp ECs | PSMA3 | 0.429 | 0.77 | 0.614 | 1.75e-17 | 5.91e-13 |
| Malignant epithelial | C3 Mp ECs | NELFE | 0.429 | 0.6 | 0.474 | 3.08e-13 | 1.04e-08 |
| Malignant epithelial | C3 Mp ECs | HMGXB4 | 0.429 | 0.371 | 0.246 | 3.65e-11 | 1.23e-06 |
| Malignant epithelial | C3 Mp ECs | RUSC1 | 0.429 | 0.4 | 0.273 | 5.16e-11 | 1.74e-06 |
| Malignant epithelial | C3 Mp ECs | PSMA6 | 0.429 | 0.284 | 0.182 | 9.64e-10 | 3.25e-05 |
| Malignant epithelial | C3 Mp ECs | OSBPL8 | 0.429 | 0.417 | 0.296 | 1.34e-09 | 4.53e-05 |
| Malignant epithelial | C3 Mp ECs | HLA-DRA | 0.428 | 0.881 | 0.737 | 8.650000000000001e-21 | 2.91e-16 |
| Malignant epithelial | C3 Mp ECs | UQCRC1 | 0.428 | 0.731 | 0.58 | 8.69e-15 | 2.93e-10 |
| Malignant epithelial | C3 Mp ECs | C3 | 0.428 | 0.346 | 0.207 | 2.26e-14 | 7.62e-10 |
| Malignant epithelial | C3 Mp ECs | PSMA5 | 0.427 | 0.623 | 0.487 | 8.03e-12 | 2.71e-07 |
| Malignant epithelial | C3 Mp ECs | RP11-1094M14.11 | 0.426 | 0.334 | 0.224 | 3.6e-09 | 0.000121 |
| Malignant epithelial | C3 Mp ECs | PSMD12 | 0.426 | 0.375 | 0.271 | 1.3e-08 | 0.000439 |
| Malignant epithelial | C3 Mp ECs | STK25 | 0.426 | 0.378 | 0.266 | 2.46e-08 | 0.000828 |
| Malignant epithelial | C3 Mp ECs | TMEM223 | 0.426 | 0.436 | 0.323 | 2.56e-08 | 0.000863 |
| Malignant epithelial | C3 Mp ECs | TMEM138 | 0.426 | 0.236 | 0.151 | 7.13e-08 | 0.0024 |
| Malignant epithelial | C3 Mp ECs | SIPA1L3 | 0.426 | 0.217 | 0.14 | 5.74e-07 | 0.0193 |
| Malignant epithelial | C3 Mp ECs | PBX3 | 0.426 | 0.137 | 0.078 | 6.82e-07 | 0.023 |
| Malignant epithelial | C3 Mp ECs | COPE | 0.425 | 0.928 | 0.815 | 1.32e-31 | 4.430000000000001e-27 |
| Malignant epithelial | C3 Mp ECs | TBCC | 0.425 | 0.36 | 0.239 | 4.5e-10 | 1.52e-05 |
| Malignant epithelial | C3 Mp ECs | HRCT1 | 0.425 | 0.171 | 0.094 | 2.43e-09 | 8.19e-05 |
| Malignant epithelial | C3 Mp ECs | TXNDC9 | 0.425 | 0.432 | 0.317 | 5.27e-09 | 0.000178 |
| Malignant epithelial | C3 Mp ECs | OAS2 | 0.425 | 0.23 | 0.142 | 9.23e-09 | 0.000311 |
| Malignant epithelial | C3 Mp ECs | PSMA7 | 0.424 | 0.975 | 0.89 | 3.66e-33 | 1.23e-28 |
| Malignant epithelial | C3 Mp ECs | ING1 | 0.424 | 0.393 | 0.275 | 6.75e-10 | 2.27e-05 |
| Malignant epithelial | C3 Mp ECs | IKBKE | 0.424 | 0.25 | 0.159 | 1.9e-08 | 0.000639 |
| Malignant epithelial | C3 Mp ECs | TMEM167A | 0.423 | 0.782 | 0.59 | 1.35e-21 | 4.56e-17 |
| Malignant epithelial | C3 Mp ECs | FH | 0.423 | 0.25 | 0.168 | 4.24e-07 | 0.0143 |
| Malignant epithelial | C3 Mp ECs | ISG15 | 0.421 | 0.954 | 0.834 | 1.97e-31 | 6.650000000000001e-27 |
| Malignant epithelial | C3 Mp ECs | UBE2G2 | 0.421 | 0.35 | 0.221 | 1.53e-12 | 5.16e-08 |
| Malignant epithelial | C3 Mp ECs | NBN | 0.421 | 0.292 | 0.187 | 8.57e-10 | 2.89e-05 |
| Malignant epithelial | C3 Mp ECs | DNTTIP1 | 0.42 | 0.77 | 0.62 | 2.45e-15 | 8.24e-11 |
| Malignant epithelial | C3 Mp ECs | PIEZO1 | 0.42 | 0.286 | 0.169 | 1.79e-12 | 6.04e-08 |
| Malignant epithelial | C3 Mp ECs | RRP15 | 0.42 | 0.316 | 0.209 | 3.39e-09 | 0.000114 |
| Malignant epithelial | C3 Mp ECs | ATP2B1 | 0.42 | 0.433 | 0.322 | 2.19e-08 | 0.000737 |
| Malignant epithelial | C3 Mp ECs | NHLRC2 | 0.42 | 0.141 | 0.078 | 9.62e-08 | 0.00324 |
| Malignant epithelial | C3 Mp ECs | ENY2 | 0.419 | 0.946 | 0.834 | 9.639999999999999e-34 | 3.25e-29 |

| compartment | subcluster | gene | avg_log2FC | pct.1 | pct.2 | p_val | p_val_adj |
|---|---|---|---|---|---|---|---|
| Malignant epithelial | C3 Mp ECs | FAM207A | 0.419 | 0.561 | 0.403 | 1.33e-13 | 4.47e-09 |
| Malignant epithelial | C3 Mp ECs | EML2 | 0.419 | 0.417 | 0.272 | 7.53e-13 | 2.54e-08 |
| Malignant epithelial | C3 Mp ECs | RSRC1 | 0.419 | 0.386 | 0.276 | 5.55e-09 | 0.000187 |
| Malignant epithelial | C3 Mp ECs | CNOT6 | 0.419 | 0.185 | 0.11 | 3.78e-08 | 0.00127 |
| Malignant epithelial | C3 Mp ECs | FTSJ1 | 0.419 | 0.254 | 0.167 | 1.73e-07 | 0.00585 |
| Malignant epithelial | C3 Mp ECs | SHC1 | 0.418 | 0.279 | 0.178 | 1.17e-09 | 3.93e-05 |
| Malignant epithelial | C3 Mp ECs | SEC11A | 0.417 | 0.902 | 0.771 | 2.28e-25 | 7.670000000000001e-21 |
| Malignant epithelial | C3 Mp ECs | NDUFAB1 | 0.417 | 0.87 | 0.734 | 9.81e-22 | 3.31e-17 |
| Malignant epithelial | C3 Mp ECs | MESDC2 | 0.417 | 0.607 | 0.437 | 2.11e-15 | 7.12e-11 |
| Malignant epithelial | C3 Mp ECs | NR2F6 | 0.417 | 0.527 | 0.362 | 3.55e-15 | 1.2e-10 |
| Malignant epithelial | C3 Mp ECs | NSMCE2 | 0.417 | 0.4 | 0.286 | 2.91e-09 | 9.81e-05 |
| Malignant epithelial | C3 Mp ECs | MRPL44 | 0.417 | 0.33 | 0.227 | 1.61e-08 | 0.000542 |
| Malignant epithelial | C3 Mp ECs | RP1-313I6.12 | 0.417 | 0.341 | 0.241 | 2.68e-08 | 0.000903 |
| Malignant epithelial | C3 Mp ECs | SAV1 | 0.417 | 0.33 | 0.232 | 2.71e-08 | 0.000914 |
| Malignant epithelial | C3 Mp ECs | FASN | 0.417 | 0.244 | 0.157 | 3.16e-08 | 0.00107 |
| Malignant epithelial | C3 Mp ECs | UTP4 | 0.417 | 0.263 | 0.18 | 5.13e-07 | 0.0173 |
| Malignant epithelial | C3 Mp ECs | EFNB2 | 0.416 | 0.647 | 0.482 | 8.65e-14 | 2.91e-09 |
| Malignant epithelial | C3 Mp ECs | SRSF9 | 0.415 | 0.888 | 0.782 | 6.199999999999999e-24 | 2.09e-19 |
| Malignant epithelial | C3 Mp ECs | SIGIRR | 0.415 | 0.673 | 0.495 | 5.4e-16 | 1.82e-11 |
| Malignant epithelial | C3 Mp ECs | PPP4R2 | 0.415 | 0.439 | 0.31 | 1.92e-10 | 6.47e-06 |
| Malignant epithelial | C3 Mp ECs | RMND5B | 0.415 | 0.228 | 0.148 | 3.02e-07 | 0.0102 |
| Malignant epithelial | C3 Mp ECs | PDE9A | 0.415 | 0.12 | 0.066 | 5.44e-07 | 0.0183 |
| Malignant epithelial | C3 Mp ECs | PKM | 0.414 | 0.982 | 0.898 | 3.26e-38 | 1.1e-33 |
| Malignant epithelial | C3 Mp ECs | INO80E | 0.414 | 0.319 | 0.215 | 1.5e-08 | 0.000504 |
| Malignant epithelial | C3 Mp ECs | RBM38 | 0.414 | 0.246 | 0.158 | 3.1e-08 | 0.00104 |
| Malignant epithelial | C3 Mp ECs | NDUFS8 | 0.413 | 0.821 | 0.669 | 1.04e-19 | 3.52e-15 |
| Malignant epithelial | C3 Mp ECs | CDK2AP2 | 0.413 | 0.723 | 0.557 | 1.76e-16 | 5.93e-12 |
| Malignant epithelial | C3 Mp ECs | CTSL | 0.413 | 0.305 | 0.199 | 1.8e-09 | 6.06e-05 |
| Malignant epithelial | C3 Mp ECs | PPP2R3C | 0.413 | 0.258 | 0.169 | 3.98e-08 | 0.00134 |
| Malignant epithelial | C3 Mp ECs | TRMT61A | 0.413 | 0.228 | 0.146 | 1.37e-07 | 0.0046 |
| Malignant epithelial | C3 Mp ECs | CSTF3 | 0.412 | 0.353 | 0.204 | 2.31e-16 | 7.77e-12 |
| Malignant epithelial | C3 Mp ECs | IQGAP1 | 0.412 | 0.706 | 0.531 | 2.35e-15 | 7.92e-11 |
| Malignant epithelial | C3 Mp ECs | BID | 0.412 | 0.652 | 0.497 | 3.78e-14 | 1.27e-09 |
| Malignant epithelial | C3 Mp ECs | DUSP1 | 0.412 | 0.91 | 0.891 | 1.29e-13 | 4.34e-09 |
| Malignant epithelial | C3 Mp ECs | VCL | 0.412 | 0.363 | 0.25 | 1.52e-08 | 0.000513 |
| Malignant epithelial | C3 Mp ECs | UBB | 0.411 | 0.99 | 0.957 | 2.29e-31 | 7.7e-27 |
| Malignant epithelial | C3 Mp ECs | TOP1 | 0.411 | 0.712 | 0.546 | 3.07e-15 | 1.03e-10 |
| Malignant epithelial | C3 Mp ECs | CFD | 0.411 | 0.51 | 0.342 | 4.42e-15 | 1.49e-10 |
| Malignant epithelial | C3 Mp ECs | RASA1 | 0.411 | 0.375 | 0.228 | 9.33e-15 | 3.14e-10 |
| Malignant epithelial | C3 Mp ECs | C19orf48 | 0.411 | 0.619 | 0.468 | 1.13e-13 | 3.8e-09 |
| Malignant epithelial | C3 Mp ECs | FIBP | 0.411 | 0.52 | 0.378 | 1.25e-11 | 4.21e-07 |
| Malignant epithelial | C3 Mp ECs | PSMD2 | 0.411 | 0.455 | 0.325 | 1.12e-10 | 3.79e-06 |
| Malignant epithelial | C3 Mp ECs | RP11-670E13.6 | 0.41 | 0.514 | 0.366 | 1.82e-12 | 6.13e-08 |
| Malignant epithelial | C3 Mp ECs | ADGRE5 | 0.41 | 0.412 | 0.296 | 1.77e-09 | 5.96e-05 |
| Malignant epithelial | C3 Mp ECs | SSBP4 | 0.409 | 0.53 | 0.364 | 1.01e-13 | 3.4e-09 |
| Malignant epithelial | C3 Mp ECs | ITPRIPL2 | 0.409 | 0.343 | 0.217 | 1.74e-11 | 5.87e-07 |
| Malignant epithelial | C3 Mp ECs | EEF1E1 | 0.409 | 0.421 | 0.28 | 4.84e-11 | 1.63e-06 |
| Malignant epithelial | C3 Mp ECs | ACOT9 | 0.409 | 0.301 | 0.199 | 1.04e-08 | 0.000351 |
| Malignant epithelial | C3 Mp ECs | MRPS15 | 0.408 | 0.611 | 0.458 | 1.4e-13 | 4.71e-09 |
| Malignant epithelial | C3 Mp ECs | ICT1 | 0.408 | 0.407 | 0.282 | 4.65e-10 | 1.57e-05 |
| Malignant epithelial | C3 Mp ECs | SERPINB9 | 0.408 | 0.221 | 0.135 | 1.48e-08 | 0.000497 |
| Malignant epithelial | C3 Mp ECs | AKIRIN2 | 0.408 | 0.4 | 0.291 | 2.68e-08 | 0.000904 |
| Malignant epithelial | C3 Mp ECs | FDPS | 0.406 | 0.65 | 0.459 | 7.03e-20 | 2.37e-15 |
| Malignant epithelial | C3 Mp ECs | POLR2E | 0.406 | 0.72 | 0.565 | 1.7e-15 | 5.73e-11 |
| Malignant epithelial | C3 Mp ECs | SLC25A39 | 0.406 | 0.738 | 0.603 | 4.36e-14 | 1.47e-09 |
| Malignant epithelial | C3 Mp ECs | SSSCA1 | 0.406 | 0.63 | 0.476 | 2.13e-13 | 7.17e-09 |
| Malignant epithelial | C3 Mp ECs | SH3GL1 | 0.406 | 0.361 | 0.239 | 6.41e-11 | 2.16e-06 |
| Malignant epithelial | C3 Mp ECs | RALBP1 | 0.406 | 0.568 | 0.426 | 1.69e-10 | 5.68e-06 |
| Malignant epithelial | C3 Mp ECs | MMAB | 0.406 | 0.357 | 0.239 | 1.94e-09 | 6.53e-05 |
| Malignant epithelial | C3 Mp ECs | TTF1 | 0.405 | 0.269 | 0.179 | 9.33e-08 | 0.00314 |
| Malignant epithelial | C3 Mp ECs | TRMT10C | 0.405 | 0.374 | 0.271 | 2.56e-07 | 0.00863 |
| Malignant epithelial | C3 Mp ECs | MANF | 0.404 | 0.716 | 0.534 | 2.86e-19 | 9.65e-15 |
| Malignant epithelial | C3 Mp ECs | SNF8 | 0.404 | 0.669 | 0.515 | 9.41e-15 | 3.17e-10 |
| Malignant epithelial | C3 Mp ECs | C6orf141 | 0.404 | 0.291 | 0.184 | 7.4e-10 | 2.49e-05 |

| compartment | subcluster | gene | avg_log2FC | pct.1 | pct.2 | p_val | p_val_adj |
|---|---|---|---|---|---|---|---|
| Malignant epithelial | C3 Mp ECs | MNAT1 | 0.404 | 0.283 | 0.18 | 2.24e-09 | 7.53e-05 |
| Malignant epithelial | C3 Mp ECs | SPATS2L | 0.403 | 0.801 | 0.624 | 3.36e-21 | 1.13e-16 |
| Malignant epithelial | C3 Mp ECs | MEA1 | 0.403 | 0.634 | 0.461 | 8.66e-15 | 2.92e-10 |
| Malignant epithelial | C3 Mp ECs | DPAGT1 | 0.403 | 0.223 | 0.135 | 3.59e-09 | 0.000121 |
| Malignant epithelial | C3 Mp ECs | STRN3 | 0.403 | 0.272 | 0.175 | 1.98e-08 | 0.000667 |
| Malignant epithelial | C3 Mp ECs | CTCF | 0.403 | 0.268 | 0.186 | 1.02e-06 | 0.0344 |
| Malignant epithelial | C3 Mp ECs | THOC7 | 0.402 | 0.719 | 0.557 | 1.04e-16 | 3.5e-12 |
| Malignant epithelial | C3 Mp ECs | TMEM106C | 0.402 | 0.465 | 0.31 | 8.36e-13 | 2.82e-08 |
| Malignant epithelial | C3 Mp ECs | IER3 | 0.402 | 0.948 | 0.893 | 2.59e-11 | 8.72e-07 |
| Malignant epithelial | C3 Mp ECs | C15orf61 | 0.401 | 0.479 | 0.326 | 7.74e-12 | 2.61e-07 |
| Malignant epithelial | C3 Mp ECs | CCM2 | 0.401 | 0.328 | 0.214 | 2.82e-10 | 9.51e-06 |
| Malignant epithelial | C3 Mp ECs | DNPH1 | 0.4 | 0.815 | 0.656 | 2.15e-17 | 7.24e-13 |
| Malignant epithelial | C3 Mp ECs | PSMD6 | 0.4 | 0.486 | 0.343 | 1.05e-11 | 3.52e-07 |
| Malignant epithelial | C3 Mp ECs | VPS26A | 0.4 | 0.472 | 0.336 | 2.64e-10 | 8.9e-06 |
| Malignant epithelial | C3 Mp ECs | PSMB8 | 0.399 | 0.834 | 0.721 | 3.34e-18 | 1.13e-13 |
| Malignant epithelial | C3 Mp ECs | ANKRD11 | 0.399 | 0.541 | 0.403 | 1.48e-10 | 4.98e-06 |
| Malignant epithelial | C3 Mp ECs | TMEM107 | 0.399 | 0.182 | 0.108 | 3.86e-08 | 0.0013 |
| Malignant epithelial | C3 Mp ECs | HNRNPUL2 | 0.399 | 0.345 | 0.248 | 1.34e-07 | 0.00452 |
| Malignant epithelial | C3 Mp ECs | TSPO | 0.398 | 0.981 | 0.932 | 1.92e-39 | 6.490000000000001e-35 |
| Malignant epithelial | C3 Mp ECs | MRPL23 | 0.398 | 0.861 | 0.696 | 1.32e-22 | 4.44e-18 |
| Malignant epithelial | C3 Mp ECs | LAMC2 | 0.398 | 0.52 | 0.35 | 8.39e-16 | 2.83e-11 |
| Malignant epithelial | C3 Mp ECs | CNPY3 | 0.398 | 0.581 | 0.422 | 7.54e-14 | 2.54e-09 |
| Malignant epithelial | C3 Mp ECs | ILK | 0.398 | 0.516 | 0.376 | 4.13e-11 | 1.39e-06 |
| Malignant epithelial | C3 Mp ECs | PWP1 | 0.398 | 0.414 | 0.301 | 1.07e-08 | 0.000359 |
| Malignant epithelial | C3 Mp ECs | TSSC1 | 0.398 | 0.319 | 0.216 | 4.59e-08 | 0.00155 |
| Malignant epithelial | C3 Mp ECs | HCFC1 | 0.398 | 0.239 | 0.157 | 2.62e-07 | 0.00883 |
| Malignant epithelial | C3 Mp ECs | NEDD8 | 0.397 | 0.943 | 0.856 | 4.5e-32 | 1.52e-27 |
| Malignant epithelial | C3 Mp ECs | NOP10 | 0.397 | 0.946 | 0.858 | 1.36e-29 | 4.58e-25 |
| Malignant epithelial | C3 Mp ECs | EI24 | 0.397 | 0.708 | 0.56 | 2.26e-14 | 7.61e-10 |
| Malignant epithelial | C3 Mp ECs | RPE | 0.397 | 0.281 | 0.181 | 7.93e-09 | 0.000267 |
| Malignant epithelial | C3 Mp ECs | IMPDH1 | 0.397 | 0.337 | 0.233 | 1.9e-08 | 0.000641 |
| Malignant epithelial | C3 Mp ECs | DRAM1 | 0.397 | 0.171 | 0.104 | 6.14e-07 | 0.0207 |
| Malignant epithelial | C3 Mp ECs | MGAT2 | 0.397 | 0.237 | 0.157 | 7.11e-07 | 0.024 |
| Malignant epithelial | C3 Mp ECs | DAXX | 0.397 | 0.292 | 0.207 | 1.33e-06 | 0.0449 |
| Malignant epithelial | C3 Mp ECs | MRPS26 | 0.396 | 0.67 | 0.493 | 3.27e-16 | 1.1e-11 |
| Malignant epithelial | C3 Mp ECs | NAE1 | 0.396 | 0.386 | 0.28 | 1.11e-08 | 0.000372 |
| Malignant epithelial | C3 Mp ECs | PDS5A | 0.396 | 0.28 | 0.185 | 2.12e-08 | 0.000715 |
| Malignant epithelial | C3 Mp ECs | EIF4A3 | 0.395 | 0.608 | 0.415 | 1.5e-18 | 5.05e-14 |
| Malignant epithelial | C3 Mp ECs | LARP1 | 0.395 | 0.541 | 0.406 | 4.27e-10 | 1.44e-05 |
| Malignant epithelial | C3 Mp ECs | NFKB2 | 0.395 | 0.328 | 0.226 | 2.77e-08 | 0.000934 |
| Malignant epithelial | C3 Mp ECs | RBM28 | 0.395 | 0.214 | 0.136 | 2.32e-07 | 0.00783 |
| Malignant epithelial | C3 Mp ECs | BRD3 | 0.394 | 0.291 | 0.2 | 3.65e-07 | 0.0123 |
| Malignant epithelial | C3 Mp ECs | CYBA | 0.393 | 0.99 | 0.97 | 2.3e-37 | 7.74e-33 |
| Malignant epithelial | C3 Mp ECs | HLA-DPA1 | 0.393 | 0.697 | 0.524 | 9.11e-14 | 3.07e-09 |
| Malignant epithelial | C3 Mp ECs | SGMS1 | 0.392 | 0.183 | 0.111 | 1.09e-07 | 0.00367 |
| Malignant epithelial | C3 Mp ECs | EGFL7 | 0.392 | 0.177 | 0.108 | 3.72e-07 | 0.0125 |
| Malignant epithelial | C3 Mp ECs | CTNNBIP1 | 0.391 | 0.563 | 0.405 | 2.12e-13 | 7.15e-09 |
| Malignant epithelial | C3 Mp ECs | CHD4 | 0.391 | 0.512 | 0.358 | 1.08e-11 | 3.64e-07 |
| Malignant epithelial | C3 Mp ECs | RUNX1 | 0.391 | 0.568 | 0.434 | 6.31e-10 | 2.12e-05 |
| Malignant epithelial | C3 Mp ECs | ILKAP | 0.39 | 0.314 | 0.207 | 7.37e-09 | 0.000248 |
| Malignant epithelial | C3 Mp ECs | PLP2 | 0.389 | 0.971 | 0.908 | 5.67e-27 | 1.91e-22 |
| Malignant epithelial | C3 Mp ECs | MTCH2 | 0.389 | 0.594 | 0.448 | 3.22e-11 | 1.09e-06 |
| Malignant epithelial | C3 Mp ECs | BZW2 | 0.389 | 0.592 | 0.462 | 4.1e-09 | 0.000138 |
| Malignant epithelial | C3 Mp ECs | TPR | 0.389 | 0.463 | 0.341 | 1.49e-08 | 0.000503 |
| Malignant epithelial | C3 Mp ECs | HSP90AA1 | 0.387 | 0.994 | 0.969 | 1.64e-36 | 5.53e-32 |
| Malignant epithelial | C3 Mp ECs | STAT1 | 0.387 | 0.615 | 0.438 | 1.65e-15 | 5.56e-11 |
| Malignant epithelial | C3 Mp ECs | MRPL12 | 0.387 | 0.698 | 0.558 | 2.65e-13 | 8.91e-09 |
| Malignant epithelial | C3 Mp ECs | TNFAIP3 | 0.387 | 0.589 | 0.448 | 1.19e-10 | 4.01e-06 |
| Malignant epithelial | C3 Mp ECs | HARS | 0.387 | 0.313 | 0.215 | 4.09e-08 | 0.00138 |
| Malignant epithelial | C3 Mp ECs | ZCCHC8 | 0.387 | 0.266 | 0.181 | 5.26e-07 | 0.0177 |
| Malignant epithelial | C3 Mp ECs | PLAUR | 0.386 | 0.846 | 0.698 | 5.39e-19 | 1.82e-14 |
| Malignant epithelial | C3 Mp ECs | C9orf142 | 0.386 | 0.701 | 0.534 | 3.17e-16 | 1.07e-11 |
| Malignant epithelial | C3 Mp ECs | HSPBP1 | 0.386 | 0.44 | 0.308 | 3.08e-10 | 1.04e-05 |
| Malignant epithelial | C3 Mp ECs | CERS2 | 0.386 | 0.469 | 0.341 | 2.03e-09 | 6.84e-05 |

| compartment | subcluster | gene | avg_log2FC | pct.1 | pct.2 | p_val | p_val_adj |
|---|---|---|---|---|---|---|---|
| Malignant epithelial | C3 Mp ECs | ANXA2 | 0.385 | 0.994 | 0.975 | 1.75e-40 | 5.91e-36 |
| Malignant epithelial | C3 Mp ECs | SFN | 0.385 | 0.84 | 0.685 | 3.89e-19 | 1.31e-14 |
| Malignant epithelial | C3 Mp ECs | SUDS3 | 0.385 | 0.488 | 0.35 | 3.06e-11 | 1.03e-06 |
| Malignant epithelial | C3 Mp ECs | INTS1 | 0.384 | 0.239 | 0.158 | 4.94e-07 | 0.0166 |
| Malignant epithelial | C3 Mp ECs | IFI6 | 0.383 | 0.825 | 0.604 | 1.97e-31 | 6.650000000000001e-27 |
| Malignant epithelial | C3 Mp ECs | PUF60 | 0.383 | 0.713 | 0.545 | 1.51e-15 | 5.07e-11 |
| Malignant epithelial | C3 Mp ECs | FAM92A1 | 0.383 | 0.277 | 0.185 | 1.03e-07 | 0.00347 |
| Malignant epithelial | C3 Mp ECs | RTN4 | 0.382 | 0.866 | 0.748 | 7.35e-19 | 2.48e-14 |
| Malignant epithelial | C3 Mp ECs | UACA | 0.382 | 0.356 | 0.239 | 1.59e-09 | 5.37e-05 |
| Malignant epithelial | C3 Mp ECs | PPAN | 0.382 | 0.312 | 0.206 | 1.1e-08 | 0.00037 |
| Malignant epithelial | C3 Mp ECs | CLIC1 | 0.381 | 0.986 | 0.964 | 7.11e-36 | 2.4e-31 |
| Malignant epithelial | C3 Mp ECs | MRPL28 | 0.381 | 0.599 | 0.444 | 4.61e-13 | 1.55e-08 |
| Malignant epithelial | C3 Mp ECs | EMC6 | 0.381 | 0.577 | 0.417 | 8.29e-13 | 2.79e-08 |
| Malignant epithelial | C3 Mp ECs | FDFT1 | 0.381 | 0.393 | 0.297 | 1.36e-06 | 0.0457 |
| Malignant epithelial | C3 Mp ECs | AKAP12 | 0.38 | 0.394 | 0.222 | 7.670000000000001e-21 | 2.58e-16 |
| Malignant epithelial | C3 Mp ECs | OAZ2 | 0.38 | 0.378 | 0.276 | 4.96e-08 | 0.00167 |
| Malignant epithelial | C3 Mp ECs | KNOP1 | 0.379 | 0.503 | 0.36 | 5.35e-11 | 1.8e-06 |
| Malignant epithelial | C3 Mp ECs | SOCS6 | 0.379 | 0.429 | 0.297 | 1.14e-10 | 3.86e-06 |
| Malignant epithelial | C3 Mp ECs | P3H4 | 0.379 | 0.265 | 0.17 | 2.21e-08 | 0.000743 |
| Malignant epithelial | C3 Mp ECs | LRRC14 | 0.379 | 0.255 | 0.167 | 9.65e-08 | 0.00325 |
| Malignant epithelial | C3 Mp ECs | TNPO1 | 0.379 | 0.297 | 0.208 | 1.29e-06 | 0.0436 |
| Malignant epithelial | C3 Mp ECs | MYADM | 0.378 | 0.72 | 0.564 | 1.6e-13 | 5.39e-09 |
| Malignant epithelial | C3 Mp ECs | PQLC1 | 0.378 | 0.223 | 0.146 | 4.77e-07 | 0.0161 |
| Malignant epithelial | C3 Mp ECs | ADH5 | 0.377 | 0.567 | 0.434 | 3.95e-10 | 1.33e-05 |
| Malignant epithelial | C3 Mp ECs | SNX8 | 0.377 | 0.314 | 0.205 | 1.67e-09 | 5.63e-05 |
| Malignant epithelial | C3 Mp ECs | KLF6 | 0.376 | 0.931 | 0.88 | 8.72e-17 | 2.94e-12 |
| Malignant epithelial | C3 Mp ECs | MRPS17 | 0.376 | 0.452 | 0.328 | 3.09e-09 | 0.000104 |
| Malignant epithelial | C3 Mp ECs | GOT2 | 0.376 | 0.354 | 0.242 | 9.47e-09 | 0.000319 |
| Malignant epithelial | C3 Mp ECs | ENDOV | 0.376 | 0.321 | 0.223 | 2.61e-07 | 0.0088 |
| Malignant epithelial | C3 Mp ECs | VTI1A | 0.376 | 0.197 | 0.125 | 6.74e-07 | 0.0227 |
| Malignant epithelial | C3 Mp ECs | GATAD2A | 0.376 | 0.288 | 0.204 | 9.14e-07 | 0.0308 |
| Malignant epithelial | C3 Mp ECs | PRSS21 | 0.375 | 0.538 | 0.347 | 8.2e-17 | 2.76e-12 |
| Malignant epithelial | C3 Mp ECs | SF3A2 | 0.375 | 0.481 | 0.323 | 5.5e-13 | 1.85e-08 |
| Malignant epithelial | C3 Mp ECs | CHCHD3 | 0.375 | 0.468 | 0.352 | 1.05e-08 | 0.000353 |
| Malignant epithelial | C3 Mp ECs | ICAM1 | 0.375 | 0.214 | 0.132 | 1.6e-08 | 0.00054 |
| Malignant epithelial | C3 Mp ECs | KMT2E-AS1 | 0.375 | 0.273 | 0.181 | 3.57e-08 | 0.0012 |
| Malignant epithelial | C3 Mp ECs | CALR | 0.374 | 0.961 | 0.836 | 6.54e-24 | 2.2e-19 |
| Malignant epithelial | C3 Mp ECs | MYH9 | 0.374 | 0.673 | 0.476 | 2.77e-17 | 9.34e-13 |
| Malignant epithelial | C3 Mp ECs | APOA1BP | 0.374 | 0.617 | 0.466 | 3.91e-12 | 1.32e-07 |
| Malignant epithelial | C3 Mp ECs | TYSND1 | 0.374 | 0.308 | 0.207 | 4.73e-08 | 0.00159 |
| Malignant epithelial | C3 Mp ECs | ATP5H | 0.373 | 0.935 | 0.851 | 2.28e-27 | 7.689999999999999e-23 |
| Malignant epithelial | C3 Mp ECs | NDUFC2 | 0.373 | 0.898 | 0.759 | 4.56e-20 | 1.54e-15 |
| Malignant epithelial | C3 Mp ECs | SDCCAG3 | 0.373 | 0.207 | 0.129 | 1.2e-07 | 0.00406 |
| Malignant epithelial | C3 Mp ECs | IAH1 | 0.372 | 0.581 | 0.436 | 6.54e-11 | 2.2e-06 |
| Malignant epithelial | C3 Mp ECs | NUP107 | 0.372 | 0.308 | 0.192 | 2.23e-10 | 7.52e-06 |
| Malignant epithelial | C3 Mp ECs | HIF1A | 0.372 | 0.281 | 0.183 | 5.34e-09 | 0.00018 |
| Malignant epithelial | C3 Mp ECs | NDUFC1 | 0.371 | 0.912 | 0.789 | 4.63e-24 | 1.56e-19 |
| Malignant epithelial | C3 Mp ECs | GSTO2 | 0.371 | 0.55 | 0.418 | 7.38e-11 | 2.49e-06 |
| Malignant epithelial | C3 Mp ECs | HYAL2 | 0.371 | 0.437 | 0.295 | 8.45e-11 | 2.85e-06 |
| Malignant epithelial | C3 Mp ECs | HKDC1 | 0.371 | 0.332 | 0.215 | 1.6e-10 | 5.4e-06 |
| Malignant epithelial | C3 Mp ECs | TIPRL | 0.371 | 0.337 | 0.243 | 1.25e-06 | 0.0421 |
| Malignant epithelial | C3 Mp ECs | LTBR | 0.37 | 0.662 | 0.494 | 6.84e-15 | 2.3e-10 |
| Malignant epithelial | C3 Mp ECs | EWSR1 | 0.37 | 0.644 | 0.499 | 4.77e-12 | 1.61e-07 |
| Malignant epithelial | C3 Mp ECs | NME2 | 0.37 | 0.323 | 0.201 | 6.13e-11 | 2.07e-06 |
| Malignant epithelial | C3 Mp ECs | GPS1 | 0.37 | 0.401 | 0.277 | 2.52e-10 | 8.5e-06 |
| Malignant epithelial | C3 Mp ECs | CTB-50L17.10 | 0.37 | 0.283 | 0.184 | 3.96e-09 | 0.000134 |
| Malignant epithelial | C3 Mp ECs | SERTAD2 | 0.37 | 0.327 | 0.222 | 1.62e-08 | 0.000546 |
| Malignant epithelial | C3 Mp ECs | DNAJA1 | 0.369 | 0.919 | 0.802 | 6.51e-20 | 2.19e-15 |
| Malignant epithelial | C3 Mp ECs | PFKP | 0.369 | 0.495 | 0.321 | 9.07e-15 | 3.05e-10 |
| Malignant epithelial | C3 Mp ECs | UBE2E3 | 0.369 | 0.552 | 0.417 | 2.72e-10 | 9.17e-06 |
| Malignant epithelial | C3 Mp ECs | PTGES3 | 0.368 | 0.908 | 0.829 | 1.89e-19 | 6.35e-15 |
| Malignant epithelial | C3 Mp ECs | RAD23A | 0.368 | 0.786 | 0.64 | 4.12e-16 | 1.39e-11 |
| Malignant epithelial | C3 Mp ECs | CLCF1 | 0.367 | 0.509 | 0.328 | 1.42e-15 | 4.79e-11 |
| Malignant epithelial | C3 Mp ECs | SERBP1 | 0.367 | 0.785 | 0.627 | 1.59e-15 | 5.34e-11 |

| compartment | subcluster | gene | avg_log2FC | pct.1 | pct.2 | p_val | p_val_adj |
|---|---|---|---|---|---|---|---|
| Malignant epithelial | C3 Mp ECs | ALKBH2 | 0.367 | 0.316 | 0.216 | 1.14e-07 | 0.00386 |
| Malignant epithelial | C3 Mp ECs | NUTF2 | 0.366 | 0.785 | 0.647 | 3.67e-16 | 1.24e-11 |
| Malignant epithelial | C3 Mp ECs | ZDHHC3 | 0.366 | 0.647 | 0.486 | 2.59e-13 | 8.71e-09 |
| Malignant epithelial | C3 Mp ECs | KPNB1 | 0.366 | 0.634 | 0.466 | 6.06e-13 | 2.04e-08 |
| Malignant epithelial | C3 Mp ECs | SNAPC3 | 0.366 | 0.225 | 0.143 | 1.14e-07 | 0.00386 |
| Malignant epithelial | C3 Mp ECs | FAM98B | 0.366 | 0.192 | 0.123 | 1.27e-06 | 0.0429 |
| Malignant epithelial | C3 Mp ECs | NXT1 | 0.365 | 0.556 | 0.426 | 1.56e-09 | 5.26e-05 |
| Malignant epithelial | C3 Mp ECs | GEMIN6 | 0.365 | 0.294 | 0.194 | 1.64e-08 | 0.000553 |
| Malignant epithelial | C3 Mp ECs | LDHA | 0.364 | 0.977 | 0.914 | 2.99e-23 | 1.01e-18 |
| Malignant epithelial | C3 Mp ECs | UBE2L6 | 0.364 | 0.641 | 0.487 | 6.2e-11 | 2.09e-06 |
| Malignant epithelial | C3 Mp ECs | GTF2E2 | 0.364 | 0.466 | 0.339 | 1.99e-09 | 6.72e-05 |
| Malignant epithelial | C3 Mp ECs | ITPK1 | 0.364 | 0.349 | 0.243 | 2.16e-08 | 0.000727 |
| Malignant epithelial | C3 Mp ECs | SSBP1 | 0.363 | 0.916 | 0.819 | 3.86e-20 | 1.3e-15 |
| Malignant epithelial | C3 Mp ECs | NHP2 | 0.363 | 0.847 | 0.694 | 3.05e-19 | 1.03e-14 |
| Malignant epithelial | C3 Mp ECs | DDHD2 | 0.363 | 0.145 | 0.085 | 9.55e-07 | 0.0322 |
| Malignant epithelial | C3 Mp ECs | ATP5G3 | 0.362 | 0.943 | 0.866 | 1.78e-22 | 5.98e-18 |
| Malignant epithelial | C3 Mp ECs | S100A16 | 0.362 | 0.887 | 0.748 | 2.49e-18 | 8.39e-14 |
| Malignant epithelial | C3 Mp ECs | SSNA1 | 0.362 | 0.764 | 0.63 | 1.69e-13 | 5.69e-09 |
| Malignant epithelial | C3 Mp ECs | AIDA | 0.362 | 0.35 | 0.242 | 1.94e-08 | 0.000655 |
| Malignant epithelial | C3 Mp ECs | U2SURP | 0.361 | 0.618 | 0.488 | 2.44e-10 | 8.23e-06 |
| Malignant epithelial | C3 Mp ECs | FAM83H | 0.361 | 0.408 | 0.282 | 3.21e-10 | 1.08e-05 |
| Malignant epithelial | C3 Mp ECs | ACLY | 0.361 | 0.23 | 0.153 | 8.42e-07 | 0.0284 |
| Malignant epithelial | C3 Mp ECs | HERPUD2 | 0.361 | 0.372 | 0.277 | 9.5e-07 | 0.032 |
| Malignant epithelial | C3 Mp ECs | S100A11 | 0.36 | 1.0 | 0.995 | 7.289999999999999e-45 | 2.46e-40 |
| Malignant epithelial | C3 Mp ECs | NDUFA7 | 0.36 | 0.731 | 0.581 | 1.35e-13 | 4.54e-09 |
| Malignant epithelial | C3 Mp ECs | EIF3J | 0.36 | 0.757 | 0.619 | 4.53e-13 | 1.53e-08 |
| Malignant epithelial | C3 Mp ECs | HAX1 | 0.36 | 0.583 | 0.438 | 4.42e-11 | 1.49e-06 |
| Malignant epithelial | C3 Mp ECs | PARP9 | 0.36 | 0.326 | 0.225 | 1.14e-07 | 0.00384 |
| Malignant epithelial | C3 Mp ECs | NDUFS3 | 0.359 | 0.677 | 0.53 | 2.71e-12 | 9.14e-08 |
| Malignant epithelial | C3 Mp ECs | ASXL1 | 0.359 | 0.286 | 0.186 | 6.85e-09 | 0.000231 |
| Malignant epithelial | C3 Mp ECs | COPS8 | 0.358 | 0.401 | 0.284 | 3.5e-09 | 0.000118 |
| Malignant epithelial | C3 Mp ECs | COMMD5 | 0.358 | 0.399 | 0.29 | 2.68e-08 | 0.000903 |
| Malignant epithelial | C3 Mp ECs | TIMM50 | 0.357 | 0.462 | 0.329 | 2.9e-10 | 9.76e-06 |
| Malignant epithelial | C3 Mp ECs | SNAPIN | 0.357 | 0.361 | 0.25 | 1.34e-08 | 0.000452 |
| Malignant epithelial | C3 Mp ECs | IFI27 | 0.356 | 0.997 | 0.935 | 2.4e-39 | 8.099999999999999e-35 |
| Malignant epithelial | C3 Mp ECs | SMIM15 | 0.356 | 0.531 | 0.405 | 2.29e-08 | 0.000771 |
| Malignant epithelial | C3 Mp ECs | MCU | 0.356 | 0.208 | 0.133 | 5.6e-07 | 0.0189 |
| Malignant epithelial | C3 Mp ECs | ACAT2 | 0.356 | 0.208 | 0.136 | 1.16e-06 | 0.0389 |
| Malignant epithelial | C3 Mp ECs | FKBP10 | 0.354 | 0.48 | 0.305 | 3.63e-15 | 1.22e-10 |
| Malignant epithelial | C3 Mp ECs | MRPL34 | 0.354 | 0.567 | 0.431 | 2.46e-10 | 8.29e-06 |
| Malignant epithelial | C3 Mp ECs | BZW1 | 0.354 | 0.659 | 0.532 | 3.07e-09 | 0.000104 |
| Malignant epithelial | C3 Mp ECs | ANKRD28 | 0.354 | 0.305 | 0.204 | 2.6e-08 | 0.000877 |
| Malignant epithelial | C3 Mp ECs | CDC123 | 0.353 | 0.483 | 0.352 | 3.76e-10 | 1.27e-05 |
| Malignant epithelial | C3 Mp ECs | TMX1 | 0.353 | 0.377 | 0.276 | 4.56e-07 | 0.0154 |
| Malignant epithelial | C3 Mp ECs | PDCD6 | 0.352 | 0.785 | 0.666 | 8.84e-13 | 2.98e-08 |
| Malignant epithelial | C3 Mp ECs | DNAJB11 | 0.352 | 0.546 | 0.401 | 1.1e-11 | 3.69e-07 |
| Malignant epithelial | C3 Mp ECs | G3BP1 | 0.352 | 0.497 | 0.359 | 8.89e-11 | 3e-06 |
| Malignant epithelial | C3 Mp ECs | PUM3 | 0.352 | 0.357 | 0.247 | 3.64e-08 | 0.00123 |
| Malignant epithelial | C3 Mp ECs | NDUFS6 | 0.351 | 0.932 | 0.839 | 1.42e-22 | 4.8e-18 |
| Malignant epithelial | C3 Mp ECs | LAMB3 | 0.351 | 0.759 | 0.61 | 4.87e-12 | 1.64e-07 |
| Malignant epithelial | C3 Mp ECs | DNAJC1 | 0.351 | 0.339 | 0.23 | 7.21e-09 | 0.000243 |
| Malignant epithelial | C3 Mp ECs | COX16 | 0.351 | 0.383 | 0.271 | 1.48e-08 | 0.000499 |
| Malignant epithelial | C3 Mp ECs | ELK3 | 0.351 | 0.201 | 0.129 | 1.18e-06 | 0.0397 |
| Malignant epithelial | C3 Mp ECs | MLEC | 0.35 | 0.648 | 0.525 | 1.62e-08 | 0.000546 |
| Malignant epithelial | C3 Mp ECs | HNRNPC | 0.349 | 0.931 | 0.832 | 5.83e-22 | 1.96e-17 |
| Malignant epithelial | C3 Mp ECs | C16orf91 | 0.349 | 0.375 | 0.264 | 1.47e-08 | 0.000494 |
| Malignant epithelial | C3 Mp ECs | E2F4 | 0.348 | 0.353 | 0.247 | 1.39e-07 | 0.00468 |
| Malignant epithelial | C3 Mp ECs | MYEOV2 | 0.347 | 0.874 | 0.748 | 3.67e-18 | 1.24e-13 |
| Malignant epithelial | C3 Mp ECs | RRS1 | 0.347 | 0.377 | 0.264 | 2.41e-08 | 0.000812 |
| Malignant epithelial | C3 Mp ECs | NDUFS5 | 0.346 | 0.932 | 0.836 | 8.419999999999999e-23 | 2.84e-18 |
| Malignant epithelial | C3 Mp ECs | AP2S1 | 0.346 | 0.834 | 0.722 | 8.04e-15 | 2.71e-10 |
| Malignant epithelial | C3 Mp ECs | C1QBP | 0.346 | 0.665 | 0.51 | 1.18e-12 | 3.96e-08 |
| Malignant epithelial | C3 Mp ECs | PSMB2 | 0.346 | 0.663 | 0.532 | 3.32e-11 | 1.12e-06 |
| Malignant epithelial | C3 Mp ECs | NAA50 | 0.346 | 0.447 | 0.329 | 1.23e-08 | 0.000415 |

| compartment | subcluster | gene | avg_log2FC | pct.1 | pct.2 | p_val | p_val_adj |
|---|---|---|---|---|---|---|---|
| Malignant epithelial | C3 Mp ECs | MRPL37 | 0.346 | 0.404 | 0.29 | 5.33e-08 | 0.0018 |
| Malignant epithelial | C3 Mp ECs | IRF7 | 0.346 | 0.377 | 0.27 | 9.57e-08 | 0.00322 |
| Malignant epithelial | C3 Mp ECs | MST1R | 0.346 | 0.214 | 0.137 | 5.9e-07 | 0.0199 |
| Malignant epithelial | C3 Mp ECs | NUDC | 0.345 | 0.629 | 0.45 | 2.33e-13 | 7.86e-09 |
| Malignant epithelial | C3 Mp ECs | YY1 | 0.345 | 0.679 | 0.544 | 1.37e-11 | 4.62e-07 |
| Malignant epithelial | C3 Mp ECs | PSMD4 | 0.345 | 0.691 | 0.563 | 5.92e-10 | 1.99e-05 |
| Malignant epithelial | C3 Mp ECs | ACSL3 | 0.345 | 0.353 | 0.245 | 3.08e-08 | 0.00104 |
| Malignant epithelial | C3 Mp ECs | TRABD | 0.344 | 0.556 | 0.397 | 2.76e-12 | 9.3e-08 |
| Malignant epithelial | C3 Mp ECs | UQCRFS1 | 0.344 | 0.77 | 0.634 | 9.92e-12 | 3.34e-07 |
| Malignant epithelial | C3 Mp ECs | COPS7B | 0.344 | 0.252 | 0.159 | 8.56e-09 | 0.000288 |
| Malignant epithelial | C3 Mp ECs | GEMIN7 | 0.344 | 0.341 | 0.242 | 2.54e-07 | 0.00856 |
| Malignant epithelial | C3 Mp ECs | POLD2 | 0.343 | 0.548 | 0.398 | 6.69e-11 | 2.25e-06 |
| Malignant epithelial | C3 Mp ECs | DDX42 | 0.343 | 0.331 | 0.239 | 1.11e-06 | 0.0374 |
| Malignant epithelial | C3 Mp ECs | PLIN2 | 0.342 | 0.681 | 0.513 | 2.62e-15 | 8.83e-11 |
| Malignant epithelial | C3 Mp ECs | RABL6 | 0.342 | 0.465 | 0.326 | 1.03e-10 | 3.49e-06 |
| Malignant epithelial | C3 Mp ECs | PCBP4 | 0.342 | 0.297 | 0.199 | 7.36e-08 | 0.00248 |
| Malignant epithelial | C3 Mp ECs | ITGA3 | 0.341 | 0.535 | 0.377 | 2.27e-10 | 7.65e-06 |
| Malignant epithelial | C3 Mp ECs | TUSC2 | 0.341 | 0.495 | 0.368 | 1.57e-08 | 0.000529 |
| Malignant epithelial | C3 Mp ECs | CFL2 | 0.341 | 0.217 | 0.131 | 1.61e-08 | 0.000544 |
| Malignant epithelial | C3 Mp ECs | TBL3 | 0.341 | 0.244 | 0.16 | 2.88e-07 | 0.00972 |
| Malignant epithelial | C3 Mp ECs | ANXA3 | 0.34 | 0.634 | 0.454 | 7.04e-16 | 2.37e-11 |
| Malignant epithelial | C3 Mp ECs | TXNDC17 | 0.34 | 0.783 | 0.633 | 6.53e-15 | 2.2e-10 |
| Malignant epithelial | C3 Mp ECs | ASNA1 | 0.34 | 0.615 | 0.437 | 1.23e-13 | 4.15e-09 |
| Malignant epithelial | C3 Mp ECs | HLA-DPB1 | 0.34 | 0.585 | 0.451 | 1.07e-09 | 3.59e-05 |
| Malignant epithelial | C3 Mp ECs | CSRP1 | 0.34 | 0.414 | 0.298 | 1.88e-08 | 0.000635 |
| Malignant epithelial | C3 Mp ECs | SMIM4 | 0.339 | 0.603 | 0.448 | 6.38e-12 | 2.15e-07 |
| Malignant epithelial | C3 Mp ECs | SS18L2 | 0.339 | 0.579 | 0.436 | 1.78e-10 | 6e-06 |
| Malignant epithelial | C3 Mp ECs | SRPK1 | 0.339 | 0.549 | 0.405 | 3.27e-10 | 1.1e-05 |
| Malignant epithelial | C3 Mp ECs | DCTPP1 | 0.339 | 0.53 | 0.433 | 8.14e-07 | 0.0274 |
| Malignant epithelial | C3 Mp ECs | COPS6 | 0.338 | 0.775 | 0.632 | 7.18e-14 | 2.42e-09 |
| Malignant epithelial | C3 Mp ECs | KXD1 | 0.338 | 0.672 | 0.509 | 2.93e-13 | 9.88e-09 |
| Malignant epithelial | C3 Mp ECs | PML | 0.338 | 0.399 | 0.274 | 9.77e-10 | 3.29e-05 |
| Malignant epithelial | C3 Mp ECs | RSU1 | 0.338 | 0.363 | 0.241 | 1.16e-09 | 3.91e-05 |
| Malignant epithelial | C3 Mp ECs | DNAJC8 | 0.338 | 0.458 | 0.346 | 1.59e-08 | 0.000536 |
| Malignant epithelial | C3 Mp ECs | GJB3 | 0.338 | 0.388 | 0.272 | 2.63e-08 | 0.000888 |
| Malignant epithelial | C3 Mp ECs | POLDIP2 | 0.338 | 0.444 | 0.327 | 1.24e-07 | 0.00419 |
| Malignant epithelial | C3 Mp ECs | NUBP1 | 0.338 | 0.301 | 0.205 | 3.8e-07 | 0.0128 |
| Malignant epithelial | C3 Mp ECs | DYRK4 | 0.337 | 0.313 | 0.191 | 1.2e-11 | 4.03e-07 |
| Malignant epithelial | C3 Mp ECs | MRPS14 | 0.337 | 0.55 | 0.415 | 1.65e-09 | 5.56e-05 |
| Malignant epithelial | C3 Mp ECs | CCAR1 | 0.337 | 0.371 | 0.259 | 2.45e-08 | 0.000825 |
| Malignant epithelial | C3 Mp ECs | TARDBP | 0.337 | 0.343 | 0.241 | 2.64e-07 | 0.0089 |
| Malignant epithelial | C3 Mp ECs | NDUFB8 | 0.336 | 0.902 | 0.799 | 4.07e-17 | 1.37e-12 |
| Malignant epithelial | C3 Mp ECs | ATN1 | 0.336 | 0.327 | 0.216 | 2.27e-09 | 7.65e-05 |
| Malignant epithelial | C3 Mp ECs | WDR43 | 0.336 | 0.388 | 0.283 | 1.5e-07 | 0.00507 |
| Malignant epithelial | C3 Mp ECs | HSPA8 | 0.335 | 0.913 | 0.82 | 1.31e-16 | 4.41e-12 |
| Malignant epithelial | C3 Mp ECs | CCDC12 | 0.335 | 0.706 | 0.557 | 1.11e-10 | 3.76e-06 |
| Malignant epithelial | C3 Mp ECs | GADD45A | 0.335 | 0.601 | 0.464 | 2.48e-10 | 8.34e-06 |
| Malignant epithelial | C3 Mp ECs | HNRNPR | 0.335 | 0.528 | 0.393 | 1.53e-09 | 5.15e-05 |
| Malignant epithelial | C3 Mp ECs | C8orf33 | 0.335 | 0.494 | 0.38 | 3.19e-08 | 0.00107 |
| Malignant epithelial | C3 Mp ECs | GNL3 | 0.334 | 0.526 | 0.383 | 8.17e-11 | 2.75e-06 |
| Malignant epithelial | C3 Mp ECs | PABPN1 | 0.334 | 0.618 | 0.48 | 8.07e-10 | 2.72e-05 |
| Malignant epithelial | C3 Mp ECs | ABHD17A | 0.334 | 0.324 | 0.228 | 2.82e-07 | 0.00951 |
| Malignant epithelial | C3 Mp ECs | NDUFA11 | 0.333 | 0.945 | 0.875 | 3.98e-24 | 1.34e-19 |
| Malignant epithelial | C3 Mp ECs | CD59 | 0.332 | 0.863 | 0.734 | 1.72e-17 | 5.8e-13 |
| Malignant epithelial | C3 Mp ECs | PLPP2 | 0.332 | 0.826 | 0.686 | 3.17e-15 | 1.07e-10 |
| Malignant epithelial | C3 Mp ECs | TOMM22 | 0.332 | 0.793 | 0.668 | 7.9e-13 | 2.66e-08 |
| Malignant epithelial | C3 Mp ECs | TPD52L2 | 0.332 | 0.601 | 0.444 | 6.66e-12 | 2.25e-07 |
| Malignant epithelial | C3 Mp ECs | SAP30BP | 0.332 | 0.425 | 0.301 | 5.74e-10 | 1.94e-05 |
| Malignant epithelial | C3 Mp ECs | COA5 | 0.332 | 0.479 | 0.341 | 6.74e-10 | 2.27e-05 |
| Malignant epithelial | C3 Mp ECs | EIF5B | 0.331 | 0.738 | 0.612 | 1.09e-08 | 0.000367 |
| Malignant epithelial | C3 Mp ECs | DNAJB12 | 0.33 | 0.436 | 0.309 | 2.75e-09 | 9.25e-05 |
| Malignant epithelial | C3 Mp ECs | DDIT4 | 0.329 | 0.917 | 0.796 | 5.61e-19 | 1.89e-14 |
| Malignant epithelial | C3 Mp ECs | SMARCA4 | 0.329 | 0.502 | 0.352 | 1.71e-11 | 5.75e-07 |
| Malignant epithelial | C3 Mp ECs | APOC1 | 0.329 | 0.513 | 0.366 | 4.3e-11 | 1.45e-06 |

| compartment | subcluster | gene | avg_log2FC | pct.1 | pct.2 | p_val | p_val_adj |
|---|---|---|---|---|---|---|---|
| Malignant epithelial | C3 Mp ECs | PRRG4 | 0.329 | 0.459 | 0.333 | 2.69e-09 | 9.07e-05 |
| Malignant epithelial | C3 Mp ECs | NCBP2 | 0.329 | 0.55 | 0.437 | 7.54e-08 | 0.00254 |
| Malignant epithelial | C3 Mp ECs | MET | 0.329 | 0.379 | 0.274 | 1.16e-07 | 0.00392 |
| Malignant epithelial | C3 Mp ECs | MT-CO3 | 0.328 | 1.0 | 0.994 | 1.81e-22 | 6.12e-18 |
| Malignant epithelial | C3 Mp ECs | CHCHD6 | 0.328 | 0.248 | 0.166 | 1.06e-06 | 0.0356 |
| Malignant epithelial | C3 Mp ECs | NDUFB7 | 0.327 | 0.957 | 0.877 | 1.28e-24 | 4.3e-20 |
| Malignant epithelial | C3 Mp ECs | S100A13 | 0.327 | 0.897 | 0.756 | 3.06e-19 | 1.03e-14 |
| Malignant epithelial | C3 Mp ECs | DEK | 0.327 | 0.774 | 0.567 | 5.74e-19 | 1.94e-14 |
| Malignant epithelial | C3 Mp ECs | MINOS1 | 0.327 | 0.909 | 0.754 | 9.46e-18 | 3.19e-13 |
| Malignant epithelial | C3 Mp ECs | RPL22L1 | 0.327 | 0.942 | 0.856 | 3e-15 | 1.01e-10 |
| Malignant epithelial | C3 Mp ECs | HSP90B1 | 0.327 | 0.899 | 0.786 | 3.61e-14 | 1.22e-09 |
| Malignant epithelial | C3 Mp ECs | PSMB5 | 0.327 | 0.774 | 0.675 | 5.27e-10 | 1.78e-05 |
| Malignant epithelial | C3 Mp ECs | MRPL3 | 0.327 | 0.527 | 0.416 | 6.65e-08 | 0.00224 |
| Malignant epithelial | C3 Mp ECs | IRF9 | 0.327 | 0.305 | 0.208 | 1.42e-07 | 0.0048 |
| Malignant epithelial | C3 Mp ECs | UBE2D2 | 0.326 | 0.804 | 0.666 | 6.47e-13 | 2.18e-08 |
| Malignant epithelial | C3 Mp ECs | ABL2 | 0.326 | 0.382 | 0.257 | 6.12e-10 | 2.06e-05 |
| Malignant epithelial | C3 Mp ECs | PPP1R7 | 0.325 | 0.575 | 0.433 | 7.82e-10 | 2.64e-05 |
| Malignant epithelial | C3 Mp ECs | SLTM | 0.325 | 0.571 | 0.423 | 1.05e-09 | 3.54e-05 |
| Malignant epithelial | C3 Mp ECs | RAC2 | 0.325 | 0.41 | 0.302 | 3.15e-07 | 0.0106 |
| Malignant epithelial | C3 Mp ECs | GPX4 | 0.324 | 0.982 | 0.936 | 1.93e-25 | 6.510000000000001e-21 |
| Malignant epithelial | C3 Mp ECs | ZNF706 | 0.324 | 0.953 | 0.873 | 1.88e-22 | 6.34e-18 |
| Malignant epithelial | C3 Mp ECs | IER3IP1 | 0.324 | 0.502 | 0.369 | 1.6e-09 | 5.41e-05 |
| Malignant epithelial | C3 Mp ECs | ITGB4 | 0.324 | 0.574 | 0.442 | 2.83e-09 | 9.53e-05 |
| Malignant epithelial | C3 Mp ECs | RPL7L1 | 0.324 | 0.513 | 0.379 | 3.99e-09 | 0.000134 |
| Malignant epithelial | C3 Mp ECs | KCTD5 | 0.324 | 0.279 | 0.191 | 3.17e-07 | 0.0107 |
| Malignant epithelial | C3 Mp ECs | HNRNPK | 0.323 | 0.909 | 0.809 | 4.52e-16 | 1.52e-11 |
| Malignant epithelial | C3 Mp ECs | EIF3B | 0.323 | 0.614 | 0.466 | 1.11e-11 | 3.74e-07 |
| Malignant epithelial | C3 Mp ECs | SNRNP27 | 0.323 | 0.388 | 0.279 | 2.79e-07 | 0.0094 |
| Malignant epithelial | C3 Mp ECs | MRPL40 | 0.322 | 0.668 | 0.498 | 2.97e-13 | 1e-08 |
| Malignant epithelial | C3 Mp ECs | CDCA7 | 0.322 | 0.317 | 0.219 | 1.03e-07 | 0.00345 |
| Malignant epithelial | C3 Mp ECs | TUBA4A | 0.32 | 0.589 | 0.425 | 8.96e-14 | 3.02e-09 |
| Malignant epithelial | C3 Mp ECs | PSMC6 | 0.32 | 0.614 | 0.444 | 7.72e-13 | 2.6e-08 |
| Malignant epithelial | C3 Mp ECs | RABIF | 0.32 | 0.359 | 0.261 | 8.11e-07 | 0.0273 |
| Malignant epithelial | C3 Mp ECs | PSMC1 | 0.319 | 0.582 | 0.448 | 8.12e-10 | 2.74e-05 |
| Malignant epithelial | C3 Mp ECs | COQ4 | 0.319 | 0.506 | 0.385 | 1.5e-08 | 0.000506 |
| Malignant epithelial | C3 Mp ECs | LSM7 | 0.318 | 0.946 | 0.868 | 3.71e-23 | 1.25e-18 |
| Malignant epithelial | C3 Mp ECs | MT-ND5 | 0.318 | 0.989 | 0.959 | 2.71e-19 | 9.12e-15 |
| Malignant epithelial | C3 Mp ECs | SMIM19 | 0.318 | 0.792 | 0.644 | 2.82e-12 | 9.49e-08 |
| Malignant epithelial | C3 Mp ECs | RBBP7 | 0.318 | 0.483 | 0.34 | 4.89e-10 | 1.65e-05 |
| Malignant epithelial | C3 Mp ECs | CDC42EP1 | 0.316 | 0.448 | 0.32 | 1.79e-09 | 6.02e-05 |
| Malignant epithelial | C3 Mp ECs | LYPLA1 | 0.316 | 0.651 | 0.548 | 2.41e-07 | 0.00811 |
| Malignant epithelial | C3 Mp ECs | PPIF | 0.315 | 0.466 | 0.334 | 7.23e-11 | 2.44e-06 |
| Malignant epithelial | C3 Mp ECs | KIAA1551 | 0.315 | 0.266 | 0.175 | 1.2e-07 | 0.00403 |
| Malignant epithelial | C3 Mp ECs | PMF1 | 0.315 | 0.4 | 0.299 | 9.11e-07 | 0.0307 |
| Malignant epithelial | C3 Mp ECs | ACTB | 0.314 | 1.0 | 0.998 | 1.03e-45 | 3.46e-41 |
| Malignant epithelial | C3 Mp ECs | TUFM | 0.314 | 0.801 | 0.675 | 2.13e-11 | 7.18e-07 |
| Malignant epithelial | C3 Mp ECs | TMEM203 | 0.314 | 0.396 | 0.284 | 3.32e-08 | 0.00112 |
| Malignant epithelial | C3 Mp ECs | GNAI2 | 0.313 | 0.534 | 0.402 | 2.17e-09 | 7.31e-05 |
| Malignant epithelial | C3 Mp ECs | S100A10 | 0.312 | 1.0 | 0.996 | 3.09e-43 | 1.04e-38 |
| Malignant epithelial | C3 Mp ECs | MORF4L2 | 0.312 | 0.749 | 0.606 | 2.69e-11 | 9.08e-07 |
| Malignant epithelial | C3 Mp ECs | SFXN1 | 0.312 | 0.31 | 0.195 | 7.36e-11 | 2.48e-06 |
| Malignant epithelial | C3 Mp ECs | PTRH1 | 0.312 | 0.444 | 0.33 | 2.07e-07 | 0.00699 |
| Malignant epithelial | C3 Mp ECs | SGTA | 0.312 | 0.343 | 0.244 | 3.35e-07 | 0.0113 |
| Malignant epithelial | C3 Mp ECs | HNRNPA1 | 0.311 | 0.988 | 0.956 | 1.53e-27 | 5.15e-23 |
| Malignant epithelial | C3 Mp ECs | MRPS5 | 0.311 | 0.583 | 0.432 | 3.13e-11 | 1.05e-06 |
| Malignant epithelial | C3 Mp ECs | PPP2CA | 0.311 | 0.579 | 0.417 | 4.08e-11 | 1.37e-06 |
| Malignant epithelial | C3 Mp ECs | PER1 | 0.311 | 0.286 | 0.186 | 1.28e-08 | 0.000431 |
| Malignant epithelial | C3 Mp ECs | F2RL1 | 0.31 | 0.486 | 0.337 | 2.49e-12 | 8.38e-08 |
| Malignant epithelial | C3 Mp ECs | DPM1 | 0.308 | 0.705 | 0.547 | 4.41e-13 | 1.49e-08 |
| Malignant epithelial | C3 Mp ECs | ADI1 | 0.308 | 0.666 | 0.546 | 1.61e-08 | 0.000541 |
| Malignant epithelial | C3 Mp ECs | FUS | 0.307 | 0.942 | 0.845 | 9.59e-19 | 3.23e-14 |
| Malignant epithelial | C3 Mp ECs | AURKAIP1 | 0.307 | 0.855 | 0.753 | 3.45e-12 | 1.16e-07 |
| Malignant epithelial | C3 Mp ECs | STOML2 | 0.307 | 0.712 | 0.604 | 2.32e-09 | 7.83e-05 |
| Malignant epithelial | C3 Mp ECs | ADAR | 0.307 | 0.516 | 0.38 | 3.17e-09 | 0.000107 |

| compartment | subcluster | gene | avg_log2FC | pct.1 | pct.2 | p_val | p_val_adj |
|---|---|---|---|---|---|---|---|
| Malignant epithelial | C3 Mp ECs | ZFYVE21 | 0.306 | 0.411 | 0.289 | 8.81e-09 | 0.000297 |
| Malignant epithelial | C3 Mp ECs | MAP4 | 0.305 | 0.479 | 0.351 | 3.09e-08 | 0.00104 |
| Malignant epithelial | C3 Mp ECs | ERCC1 | 0.305 | 0.513 | 0.396 | 1.54e-07 | 0.00519 |
| Malignant epithelial | C3 Mp ECs | PSMB8-AS1 | 0.305 | 0.254 | 0.169 | 7.37e-07 | 0.0248 |
| Malignant epithelial | C3 Mp ECs | PPIB | 0.304 | 0.957 | 0.876 | 1.73e-18 | 5.82e-14 |
| Malignant epithelial | C3 Mp ECs | PMAIP1 | 0.304 | 0.812 | 0.659 | 3.18e-14 | 1.07e-09 |
| Malignant epithelial | C3 Mp ECs | PRDM1 | 0.304 | 0.421 | 0.266 | 5.68e-14 | 1.91e-09 |
| Malignant epithelial | C3 Mp ECs | STRA13 | 0.304 | 0.674 | 0.543 | 1.31e-09 | 4.43e-05 |
| Malignant epithelial | C3 Mp ECs | CCT3 | 0.304 | 0.727 | 0.606 | 4.31e-09 | 0.000145 |
| Malignant epithelial | C3 Mp ECs | YDJC | 0.303 | 0.601 | 0.449 | 4.18e-11 | 1.41e-06 |
| Malignant epithelial | C3 Mp ECs | TAF10 | 0.303 | 0.586 | 0.46 | 1.64e-08 | 0.000551 |
| Malignant epithelial | C3 Mp ECs | CLTA | 0.302 | 0.935 | 0.861 | 8.4e-15 | 2.83e-10 |
| Malignant epithelial | C3 Mp ECs | HCFC1R1 | 0.302 | 0.589 | 0.452 | 1.74e-09 | 5.86e-05 |
| Malignant epithelial | C3 Mp ECs | NDUFV1 | 0.302 | 0.59 | 0.455 | 4.44e-09 | 0.00015 |
| Malignant epithelial | C3 Mp ECs | UBE2D3 | 0.301 | 0.914 | 0.797 | 2.21e-15 | 7.44e-11 |
| Malignant epithelial | C3 Mp ECs | ABHD12 | 0.301 | 0.545 | 0.384 | 5.33e-12 | 1.8e-07 |
| Malignant epithelial | C3 Mp ECs | CECR5 | 0.301 | 0.316 | 0.211 | 3.73e-08 | 0.00126 |
| Malignant epithelial | C3 Mp ECs | SNU13 | 0.3 | 0.919 | 0.857 | 3.84e-16 | 1.29e-11 |
| Malignant epithelial | C3 Mp ECs | FAM58A | 0.3 | 0.326 | 0.231 | 1.27e-06 | 0.0428 |
| Malignant epithelial | C3 Mp ECs | SYTL1 | 0.299 | 0.323 | 0.215 | 9.46e-09 | 0.000319 |
| Malignant epithelial | C3 Mp ECs | ALG3 | 0.299 | 0.492 | 0.365 | 1.14e-08 | 0.000383 |
| Malignant epithelial | C3 Mp ECs | MID1IP1 | 0.299 | 0.406 | 0.287 | 3.69e-08 | 0.00124 |
| Malignant epithelial | C3 Mp ECs | AK2 | 0.298 | 0.583 | 0.452 | 1.02e-09 | 3.44e-05 |
| Malignant epithelial | C3 Mp ECs | APEH | 0.297 | 0.31 | 0.219 | 1.17e-06 | 0.0395 |
| Malignant epithelial | C3 Mp ECs | TSN | 0.296 | 0.383 | 0.272 | 5.77e-08 | 0.00194 |
| Malignant epithelial | C3 Mp ECs | C7orf55 | 0.295 | 0.606 | 0.46 | 6.44e-10 | 2.17e-05 |
| Malignant epithelial | C3 Mp ECs | SPRY1 | 0.295 | 0.372 | 0.262 | 1.88e-08 | 0.000635 |
| Malignant epithelial | C3 Mp ECs | FAM204A | 0.295 | 0.506 | 0.38 | 7.96e-08 | 0.00268 |
| Malignant epithelial | C3 Mp ECs | MORN2 | 0.295 | 0.392 | 0.293 | 1.03e-06 | 0.0346 |
| Malignant epithelial | C3 Mp ECs | VAMP5 | 0.294 | 0.417 | 0.31 | 1.05e-06 | 0.0354 |
| Malignant epithelial | C3 Mp ECs | RTF1 | 0.293 | 0.701 | 0.547 | 2.78e-10 | 9.37e-06 |
| Malignant epithelial | C3 Mp ECs | C14orf119 | 0.293 | 0.506 | 0.378 | 1.71e-09 | 5.75e-05 |
| Malignant epithelial | C3 Mp ECs | MBD3 | 0.293 | 0.19 | 0.111 | 1.78e-08 | 0.0006 |
| Malignant epithelial | C3 Mp ECs | MBTPS1 | 0.293 | 0.248 | 0.166 | 5.07e-07 | 0.0171 |
| Malignant epithelial | C3 Mp ECs | TIMM44 | 0.293 | 0.349 | 0.252 | 8.12e-07 | 0.0274 |
| Malignant epithelial | C3 Mp ECs | MRPL24 | 0.292 | 0.543 | 0.396 | 2.21e-10 | 7.46e-06 |
| Malignant epithelial | C3 Mp ECs | IRF1 | 0.292 | 0.803 | 0.695 | 3.18e-10 | 1.07e-05 |
| Malignant epithelial | C3 Mp ECs | CYB561 | 0.292 | 0.374 | 0.273 | 5.54e-07 | 0.0187 |
| Malignant epithelial | C3 Mp ECs | TGM2 | 0.291 | 0.633 | 0.457 | 2.7e-12 | 9.1e-08 |
| Malignant epithelial | C3 Mp ECs | PSMC4 | 0.291 | 0.606 | 0.451 | 8.03e-11 | 2.71e-06 |
| Malignant epithelial | C3 Mp ECs | MDM2 | 0.29 | 0.153 | 0.092 | 1.31e-06 | 0.0442 |
| Malignant epithelial | C3 Mp ECs | SSFA2 | 0.289 | 0.669 | 0.479 | 1.35e-15 | 4.56e-11 |
| Malignant epithelial | C3 Mp ECs | PRELID1 | 0.289 | 0.883 | 0.785 | 4.02e-13 | 1.35e-08 |
| Malignant epithelial | C3 Mp ECs | C19orf53 | 0.287 | 0.971 | 0.9 | 2.49e-21 | 8.38e-17 |
| Malignant epithelial | C3 Mp ECs | CIB1 | 0.287 | 0.873 | 0.764 | 5.02e-13 | 1.69e-08 |
| Malignant epithelial | C3 Mp ECs | ABCA7 | 0.287 | 0.327 | 0.208 | 5.83e-11 | 1.97e-06 |
| Malignant epithelial | C3 Mp ECs | PDAP1 | 0.287 | 0.753 | 0.639 | 7.02e-10 | 2.37e-05 |
| Malignant epithelial | C3 Mp ECs | ANXA7 | 0.287 | 0.589 | 0.461 | 2.85e-08 | 0.000961 |
| Malignant epithelial | C3 Mp ECs | TSR3 | 0.287 | 0.575 | 0.448 | 8.38e-08 | 0.00283 |
| Malignant epithelial | C3 Mp ECs | KDELR3 | 0.287 | 0.338 | 0.237 | 5.65e-07 | 0.019 |
| Malignant epithelial | C3 Mp ECs | TXN | 0.286 | 0.996 | 0.984 | 5.040000000000001e-31 | 1.7e-26 |
| Malignant epithelial | C3 Mp ECs | NFKBIZ | 0.286 | 0.582 | 0.45 | 2.39e-08 | 0.000804 |
| Malignant epithelial | C3 Mp ECs | ATP5J2 | 0.285 | 0.981 | 0.927 | 1.87e-23 | 6.3e-19 |
| Malignant epithelial | C3 Mp ECs | YWHAE | 0.285 | 0.846 | 0.73 | 7.19e-12 | 2.42e-07 |
| Malignant epithelial | C3 Mp ECs | NDUFS7 | 0.285 | 0.778 | 0.647 | 2.18e-11 | 7.35e-07 |
| Malignant epithelial | C3 Mp ECs | FAM173A | 0.285 | 0.654 | 0.497 | 8.04e-11 | 2.71e-06 |
| Malignant epithelial | C3 Mp ECs | GTPBP4 | 0.285 | 0.368 | 0.266 | 4.01e-07 | 0.0135 |
| Malignant epithelial | C3 Mp ECs | CALM1 | 0.284 | 0.971 | 0.905 | 2.07e-19 | 6.99e-15 |
| Malignant epithelial | C3 Mp ECs | SUMO1 | 0.284 | 0.898 | 0.777 | 1.51e-16 | 5.09e-12 |
| Malignant epithelial | C3 Mp ECs | ARPC5 | 0.284 | 0.746 | 0.629 | 4.47e-10 | 1.51e-05 |
| Malignant epithelial | C3 Mp ECs | NDUFS4 | 0.284 | 0.746 | 0.61 | 5.72e-10 | 1.93e-05 |
| Malignant epithelial | C3 Mp ECs | POLR2G | 0.284 | 0.559 | 0.44 | 9.64e-08 | 0.00325 |
| Malignant epithelial | C3 Mp ECs | TPM4 | 0.283 | 0.841 | 0.735 | 2.21e-10 | 7.45e-06 |
| Malignant epithelial | C3 Mp ECs | IDH3B | 0.283 | 0.579 | 0.452 | 2.05e-08 | 0.00069 |

| compartment | subcluster | gene | avg_log2FC | pct.1 | pct.2 | p_val | p_val_adj |
|---|---|---|---|---|---|---|---|
| Malignant epithelial | C3 Mp ECs | ATIC | 0.283 | 0.526 | 0.412 | 6.07e-07 | 0.0204 |
| Malignant epithelial | C3 Mp ECs | TNFRSF12A | 0.282 | 0.768 | 0.611 | 7.53e-16 | 2.54e-11 |
| Malignant epithelial | C3 Mp ECs | PLIN3 | 0.282 | 0.509 | 0.396 | 2.21e-07 | 0.00746 |
| Malignant epithelial | C3 Mp ECs | ARPP19 | 0.281 | 0.549 | 0.403 | 5.96e-10 | 2.01e-05 |
| Malignant epithelial | C3 Mp ECs | MESDC1 | 0.281 | 0.337 | 0.229 | 1.98e-08 | 0.000666 |
| Malignant epithelial | C3 Mp ECs | FAM96A | 0.281 | 0.513 | 0.386 | 2.82e-08 | 0.000951 |
| Malignant epithelial | C3 Mp ECs | IGF2BP3 | 0.281 | 0.295 | 0.205 | 4.52e-07 | 0.0152 |
| Malignant epithelial | C3 Mp ECs | YBX1 | 0.28 | 0.968 | 0.904 | 2.41e-18 | 8.11e-14 |
| Malignant epithelial | C3 Mp ECs | TRAPPC1 | 0.28 | 0.662 | 0.535 | 3.72e-08 | 0.00125 |
| Malignant epithelial | C3 Mp ECs | PRDX3 | 0.28 | 0.549 | 0.431 | 8.89e-08 | 0.003 |
| Malignant epithelial | C3 Mp ECs | ANAPC16 | 0.279 | 0.935 | 0.819 | 4.86e-16 | 1.64e-11 |
| Malignant epithelial | C3 Mp ECs | DNAJC19 | 0.279 | 0.714 | 0.559 | 6.65e-11 | 2.24e-06 |
| Malignant epithelial | C3 Mp ECs | POLR2H | 0.279 | 0.658 | 0.51 | 1.71e-09 | 5.76e-05 |
| Malignant epithelial | C3 Mp ECs | NCLN | 0.279 | 0.324 | 0.214 | 6.78e-09 | 0.000228 |
| Malignant epithelial | C3 Mp ECs | CCT4 | 0.279 | 0.766 | 0.644 | 2.1e-08 | 0.000709 |
| Malignant epithelial | C3 Mp ECs | MRGBP | 0.278 | 0.437 | 0.319 | 5.56e-08 | 0.00187 |
| Malignant epithelial | C3 Mp ECs | FAM110A | 0.278 | 0.309 | 0.207 | 6.91e-08 | 0.00233 |
| Malignant epithelial | C3 Mp ECs | FLOT1 | 0.278 | 0.615 | 0.499 | 3.72e-07 | 0.0125 |
| Malignant epithelial | C3 Mp ECs | PSMC2 | 0.277 | 0.426 | 0.313 | 2.66e-07 | 0.00896 |
| Malignant epithelial | C3 Mp ECs | KRTCAP2 | 0.276 | 0.906 | 0.815 | 2.05e-13 | 6.9e-09 |
| Malignant epithelial | C3 Mp ECs | ECH1 | 0.276 | 0.763 | 0.627 | 3.9e-11 | 1.32e-06 |
| Malignant epithelial | C3 Mp ECs | CWC15 | 0.276 | 0.654 | 0.534 | 4.3e-08 | 0.00145 |
| Malignant epithelial | C3 Mp ECs | USF2 | 0.276 | 0.444 | 0.324 | 7.06e-08 | 0.00238 |
| Malignant epithelial | C3 Mp ECs | SELT | 0.276 | 0.626 | 0.51 | 1.27e-07 | 0.00427 |
| Malignant epithelial | C3 Mp ECs | PTBP1 | 0.275 | 0.738 | 0.568 | 2.34e-13 | 7.88e-09 |
| Malignant epithelial | C3 Mp ECs | ZCRB1 | 0.275 | 0.69 | 0.553 | 1.99e-09 | 6.71e-05 |
| Malignant epithelial | C3 Mp ECs | NIPA2 | 0.275 | 0.367 | 0.267 | 5.55e-07 | 0.0187 |
| Malignant epithelial | C3 Mp ECs | LSM8 | 0.274 | 0.793 | 0.67 | 8.24e-09 | 0.000278 |
| Malignant epithelial | C3 Mp ECs | SP100 | 0.273 | 0.506 | 0.374 | 5.14e-09 | 0.000173 |
| Malignant epithelial | C3 Mp ECs | PPP2CB | 0.273 | 0.45 | 0.325 | 8.08e-09 | 0.000272 |
| Malignant epithelial | C3 Mp ECs | EIF4G1 | 0.273 | 0.479 | 0.353 | 6.39e-08 | 0.00215 |
| Malignant epithelial | C3 Mp ECs | UBE2Z | 0.273 | 0.389 | 0.279 | 1.2e-07 | 0.00405 |
| Malignant epithelial | C3 Mp ECs | VAPB | 0.273 | 0.309 | 0.217 | 1.25e-06 | 0.0422 |
| Malignant epithelial | C3 Mp ECs | NAA38 | 0.272 | 0.626 | 0.487 | 3.8e-10 | 1.28e-05 |
| Malignant epithelial | C3 Mp ECs | PPP1CC | 0.272 | 0.658 | 0.533 | 7.7e-08 | 0.00259 |
| Malignant epithelial | C3 Mp ECs | ACP1 | 0.271 | 0.833 | 0.701 | 3.7e-12 | 1.25e-07 |
| Malignant epithelial | C3 Mp ECs | ENSA | 0.271 | 0.724 | 0.604 | 2.18e-09 | 7.36e-05 |
| Malignant epithelial | C3 Mp ECs | RNPS1 | 0.27 | 0.797 | 0.649 | 1.91e-10 | 6.44e-06 |
| Malignant epithelial | C3 Mp ECs | REEP5 | 0.27 | 0.796 | 0.668 | 3.22e-10 | 1.08e-05 |
| Malignant epithelial | C3 Mp ECs | PROSC | 0.27 | 0.386 | 0.275 | 1.22e-07 | 0.0041 |
| Malignant epithelial | C3 Mp ECs | SLC38A10 | 0.27 | 0.28 | 0.188 | 3.93e-07 | 0.0132 |
| Malignant epithelial | C3 Mp ECs | NSUN5 | 0.27 | 0.335 | 0.244 | 1.33e-06 | 0.0448 |
| Malignant epithelial | C3 Mp ECs | SNRPD2 | 0.269 | 0.983 | 0.955 | 2.53e-24 | 8.52e-20 |
| Malignant epithelial | C3 Mp ECs | GSDMD | 0.269 | 0.679 | 0.526 | 3.14e-10 | 1.06e-05 |
| Malignant epithelial | C3 Mp ECs | FAM103A1 | 0.269 | 0.462 | 0.341 | 7.34e-08 | 0.00247 |
| Malignant epithelial | C3 Mp ECs | CBY1 | 0.269 | 0.288 | 0.193 | 1.43e-07 | 0.0048 |
| Malignant epithelial | C3 Mp ECs | PPP1R14B | 0.268 | 0.745 | 0.578 | 2.11e-12 | 7.1e-08 |
| Malignant epithelial | C3 Mp ECs | SNRNP200 | 0.268 | 0.392 | 0.283 | 1.42e-07 | 0.00478 |
| Malignant epithelial | C3 Mp ECs | ADAM9 | 0.267 | 0.498 | 0.352 | 6.71e-10 | 2.26e-05 |
| Malignant epithelial | C3 Mp ECs | BABAM1 | 0.267 | 0.48 | 0.365 | 3.27e-07 | 0.011 |
| Malignant epithelial | C3 Mp ECs | UQCRQ | 0.266 | 0.99 | 0.963 | 2.13e-23 | 7.16e-19 |
| Malignant epithelial | C3 Mp ECs | DPY30 | 0.266 | 0.836 | 0.705 | 6.52e-13 | 2.2e-08 |
| Malignant epithelial | C3 Mp ECs | FAM96B | 0.266 | 0.861 | 0.735 | 8.97e-12 | 3.02e-07 |
| Malignant epithelial | C3 Mp ECs | LDLR | 0.266 | 0.623 | 0.476 | 8.23e-09 | 0.000277 |
| Malignant epithelial | C3 Mp ECs | MT-CO2 | 0.265 | 1.0 | 0.996 | 4.43e-19 | 1.49e-14 |
| Malignant epithelial | C3 Mp ECs | ADRM1 | 0.265 | 0.79 | 0.644 | 7.17e-11 | 2.42e-06 |
| Malignant epithelial | C3 Mp ECs | LYPD3 | 0.265 | 0.29 | 0.19 | 6.41e-09 | 0.000216 |
| Malignant epithelial | C3 Mp ECs | VTI1B | 0.265 | 0.578 | 0.45 | 5.24e-08 | 0.00177 |
| Malignant epithelial | C3 Mp ECs | TMEM160 | 0.264 | 0.748 | 0.597 | 5.34e-12 | 1.8e-07 |
| Malignant epithelial | C3 Mp ECs | PPHLN1 | 0.264 | 0.459 | 0.351 | 1.32e-06 | 0.0443 |
| Malignant epithelial | C3 Mp ECs | HSD17B10 | 0.263 | 0.701 | 0.56 | 8.4e-11 | 2.83e-06 |
| Malignant epithelial | C3 Mp ECs | LSM14A | 0.263 | 0.553 | 0.418 | 9.27e-09 | 0.000312 |
| Malignant epithelial | C3 Mp ECs | AC013461.1 | 0.263 | 0.378 | 0.267 | 6.07e-08 | 0.00205 |
| Malignant epithelial | C3 Mp ECs | PPP1R18 | 0.263 | 0.276 | 0.182 | 1.38e-07 | 0.00465 |

| compartment | subcluster | gene | avg_log2FC | pct.1 | pct.2 | p_val | p_val_adj |
|---|---|---|---|---|---|---|---|
| Malignant epithelial | C3 Mp ECs | AHCY | 0.262 | 0.625 | 0.505 | 1.29e-07 | 0.00436 |
| Malignant epithelial | C3 Mp ECs | CFLAR | 0.261 | 0.677 | 0.534 | 9.09e-10 | 3.06e-05 |
| Malignant epithelial | C3 Mp ECs | SCAMP3 | 0.261 | 0.343 | 0.235 | 1.65e-07 | 0.00555 |
| Malignant epithelial | C3 Mp ECs | SEPT9 | 0.261 | 0.567 | 0.454 | 5.5e-07 | 0.0185 |
| Malignant epithelial | C3 Mp ECs | FKBP2 | 0.26 | 0.898 | 0.798 | 1.48e-11 | 4.98e-07 |
| Malignant epithelial | C3 Mp ECs | ARPC2 | 0.26 | 0.903 | 0.835 | 5.51e-11 | 1.86e-06 |
| Malignant epithelial | C3 Mp ECs | SHISA5 | 0.26 | 0.59 | 0.458 | 4e-08 | 0.00135 |
| Malignant epithelial | C3 Mp ECs | DEGS1 | 0.26 | 0.276 | 0.179 | 4.59e-08 | 0.00155 |
| Malignant epithelial | C3 Mp ECs | S100A6 | 0.259 | 1.0 | 1.0 | 2.05e-24 | 6.91e-20 |
| Malignant epithelial | C3 Mp ECs | RNH1 | 0.259 | 0.759 | 0.642 | 6.74e-09 | 0.000227 |
| Malignant epithelial | C3 Mp ECs | GLOD4 | 0.259 | 0.331 | 0.23 | 3.24e-07 | 0.0109 |
| Malignant epithelial | C3 Mp ECs | PARL | 0.259 | 0.429 | 0.324 | 5.12e-07 | 0.0173 |
| Malignant epithelial | C3 Mp ECs | MOB1A | 0.259 | 0.505 | 0.39 | 6.57e-07 | 0.0221 |
| Malignant epithelial | C3 Mp ECs | YWHAB | 0.258 | 0.919 | 0.812 | 2.92e-14 | 9.82e-10 |
| Malignant epithelial | C3 Mp ECs | MRPL42 | 0.258 | 0.599 | 0.465 | 1.69e-09 | 5.69e-05 |
| Malignant epithelial | C3 Mp ECs | PIH1D1 | 0.258 | 0.461 | 0.33 | 1.4e-08 | 0.000471 |
| Malignant epithelial | C3 Mp ECs | CDK16 | 0.258 | 0.441 | 0.323 | 1.04e-07 | 0.00349 |
| Malignant epithelial | C3 Mp ECs | CISD1 | 0.258 | 0.637 | 0.512 | 2.49e-07 | 0.00839 |
| Malignant epithelial | C3 Mp ECs | ELFN1-AS1 | 0.257 | 0.462 | 0.309 | 2.33e-11 | 7.85e-07 |
| Malignant epithelial | C3 Mp ECs | CHORDC1 | 0.257 | 0.399 | 0.285 | 6.01e-08 | 0.00202 |
| Malignant epithelial | C3 Mp ECs | HSPB11 | 0.257 | 0.436 | 0.327 | 1.37e-06 | 0.0461 |
| Malignant epithelial | C3 Mp ECs | ZNHIT2 | 0.257 | 0.268 | 0.182 | 1.46e-06 | 0.0493 |
| Malignant epithelial | C3 Mp ECs | TMBIM6 | 0.256 | 0.959 | 0.885 | 2.85e-16 | 9.62e-12 |
| Malignant epithelial | C3 Mp ECs | PSMD8 | 0.256 | 0.781 | 0.665 | 2.23e-09 | 7.52e-05 |
| Malignant epithelial | C3 Mp ECs | C9orf3 | 0.256 | 0.4 | 0.298 | 7.43e-07 | 0.025 |
| Malignant epithelial | C3 Mp ECs | CETN3 | 0.256 | 0.294 | 0.203 | 1.17e-06 | 0.0395 |
| Malignant epithelial | C3 Mp ECs | AP2M1 | 0.255 | 0.684 | 0.541 | 9.61e-10 | 3.24e-05 |
| Malignant epithelial | C3 Mp ECs | HSBP1 | 0.254 | 0.948 | 0.866 | 2.69e-18 | 9.06e-14 |
| Malignant epithelial | C3 Mp ECs | HLA-DMA | 0.254 | 0.604 | 0.455 | 1.04e-08 | 0.000351 |
| Malignant epithelial | C3 Mp ECs | DARS | 0.254 | 0.557 | 0.424 | 6.23e-08 | 0.0021 |
| Malignant epithelial | C3 Mp ECs | CNOT1 | 0.254 | 0.37 | 0.262 | 2.37e-07 | 0.00799 |
| Malignant epithelial | C3 Mp ECs | DHX30 | 0.254 | 0.354 | 0.255 | 8.09e-07 | 0.0273 |
| Malignant epithelial | C3 Mp ECs | CUEDC2 | 0.254 | 0.378 | 0.276 | 1.29e-06 | 0.0433 |
| Malignant epithelial | C3 Mp ECs | UBE2M | 0.254 | 0.25 | 0.167 | 1.3e-06 | 0.0437 |
| Malignant epithelial | C3 Mp ECs | C11orf31 | 0.253 | 0.96 | 0.886 | 9.66e-16 | 3.26e-11 |
| Malignant epithelial | C3 Mp ECs | PCBP1 | 0.252 | 0.901 | 0.761 | 5.01e-14 | 1.69e-09 |
| Malignant epithelial | C3 Mp ECs | OGFR | 0.252 | 0.61 | 0.485 | 8.94e-08 | 0.00301 |
| Malignant epithelial | C3 Mp ECs | COX5A | 0.251 | 0.948 | 0.895 | 4.18e-16 | 1.41e-11 |
| Malignant epithelial | C3 Mp ECs | SULF2 | 0.251 | 0.603 | 0.425 | 5.62e-13 | 1.89e-08 |
| Malignant epithelial | C3 Mp ECs | LLPH | 0.251 | 0.513 | 0.377 | 3.41e-09 | 0.000115 |
| Malignant epithelial | C3 Mp ECs | NSMCE4A | 0.251 | 0.36 | 0.263 | 1.47e-06 | 0.0496 |
| Malignant epithelial | C3 Mp ECs | CCT8 | 0.251 | 0.669 | 0.565 | 1.48e-06 | 0.0499 |
| Malignant epithelial | C4 T/r-h ECs | IGFBP6 | 1.325 | 0.635 | 0.541 | 1.1e-14 | 3.69e-10 |
| Malignant epithelial | C4 T/r-h ECs | S100A4 | 0.879 | 0.894 | 0.881 | 2.02e-20 | 6.8e-16 |
| Malignant epithelial | C4 T/r-h ECs | ADIRF | 0.866 | 0.858 | 0.86 | 1.08e-09 | 3.62e-05 |
| Malignant epithelial | C4 T/r-h ECs | ZFAS1 | 0.834 | 0.954 | 0.987 | 6.559999999999999e-23 | 2.21e-18 |
| Malignant epithelial | C4 T/r-h ECs | S100A10 | 0.687 | 0.988 | 0.997 | 4.83e-11 | 1.63e-06 |
| Malignant epithelial | C4 T/r-h ECs | RPL21 | 0.682 | 0.983 | 0.992 | 1.73e-11 | 5.83e-07 |
| Malignant epithelial | C4 T/r-h ECs | RPL26 | 0.649 | 0.998 | 0.997 | 3.14e-61 | 1.06e-56 |
| Malignant epithelial | C4 T/r-h ECs | RPL31 | 0.609 | 0.996 | 0.998 | 5.64e-70 | 1.9e-65 |
| Malignant epithelial | C4 T/r-h ECs | RPL17 | 0.604 | 0.796 | 0.861 | 2.11e-17 | 7.12e-13 |
| Malignant epithelial | C4 T/r-h ECs | USMG5 | 0.571 | 0.883 | 0.959 | 9.02e-14 | 3.04e-09 |
| Malignant epithelial | C4 T/r-h ECs | LGALS1 | 0.566 | 0.802 | 0.767 | 3.33e-11 | 1.12e-06 |
| Malignant epithelial | C4 T/r-h ECs | RPL35A | 0.561 | 0.994 | 0.999 | 1.11e-62 | 3.74e-58 |
| Malignant epithelial | C4 T/r-h ECs | RPS17 | 0.555 | 0.998 | 0.998 | 7.46e-63 | 2.52e-58 |
| Malignant epithelial | C4 T/r-h ECs | RPL36 | 0.546 | 1.0 | 1.0 | 6.02e-66 | 2.03e-61 |
| Malignant epithelial | C4 T/r-h ECs | UQCRB | 0.543 | 0.946 | 0.981 | 1.24e-30 | 4.18e-26 |
| Malignant epithelial | C4 T/r-h ECs | C19orf33 | 0.537 | 0.952 | 0.955 | 8.540000000000001e-27 | 2.88e-22 |
| Malignant epithelial | C4 T/r-h ECs | DMKN | 0.534 | 0.675 | 0.667 | 3.96e-08 | 0.00133 |
| Malignant epithelial | C4 T/r-h ECs | PFDN5 | 0.527 | 0.935 | 0.979 | 2.18e-32 | 7.36e-28 |
| Malignant epithelial | C4 T/r-h ECs | OST4 | 0.522 | 0.912 | 0.972 | 2.71e-16 | 9.15e-12 |
| Malignant epithelial | C4 T/r-h ECs | TBCA | 0.518 | 0.85 | 0.913 | 1.36e-10 | 4.58e-06 |
| Malignant epithelial | C4 T/r-h ECs | RPL37A | 0.513 | 1.0 | 0.998 | 1.52e-66 | 5.13e-62 |
| Malignant epithelial | C4 T/r-h ECs | C4orf3 | 0.51 | 0.775 | 0.878 | 5.71e-08 | 0.00192 |

| compartment | subcluster | gene | avg_log2FC | pct.1 | pct.2 | p_val | p_val_adj |
|---|---|---|---|---|---|---|---|
| Malignant epithelial | C4 T/r-h ECs | RPL36A | 0.504 | 0.977 | 0.994 | 3.54e-41 | 1.19e-36 |
| Malignant epithelial | C4 T/r-h ECs | S100A2 | 0.504 | 0.706 | 0.657 | 1.23e-07 | 0.00414 |
| Malignant epithelial | C4 T/r-h ECs | RPL41 | 0.498 | 0.998 | 1.0 | 2.8e-61 | 9.42e-57 |
| Malignant epithelial | C4 T/r-h ECs | RPS24 | 0.497 | 1.0 | 0.997 | 1.09e-36 | 3.68e-32 |
| Malignant epithelial | C4 T/r-h ECs | RPS25 | 0.475 | 0.996 | 0.997 | 1.38e-60 | 4.65e-56 |
| Malignant epithelial | C4 T/r-h ECs | RPS28 | 0.473 | 1.0 | 0.998 | 4.749999999999999e-52 | 1.6e-47 |
| Malignant epithelial | C4 T/r-h ECs | TMSB10 | 0.471 | 1.0 | 1.0 | 3.48e-33 | 1.17e-28 |
| Malignant epithelial | C4 T/r-h ECs | RPL6 | 0.463 | 0.992 | 0.994 | 1.99e-38 | 6.71e-34 |
| Malignant epithelial | C4 T/r-h ECs | RPS15 | 0.459 | 1.0 | 0.999 | 1.42e-53 | 4.79e-49 |
| Malignant epithelial | C4 T/r-h ECs | RPL34 | 0.454 | 0.994 | 0.999 | 4.52e-43 | 1.52e-38 |
| Malignant epithelial | C4 T/r-h ECs | RPS29 | 0.452 | 1.0 | 0.999 | 1.12e-42 | 3.76e-38 |
| Malignant epithelial | C4 T/r-h ECs | RPS15A | 0.446 | 0.998 | 1.0 | 4.04e-48 | 1.36e-43 |
| Malignant epithelial | C4 T/r-h ECs | RPS13 | 0.443 | 0.99 | 0.995 | 5.62e-40 | 1.89e-35 |
| Malignant epithelial | C4 T/r-h ECs | RPL12 | 0.439 | 0.996 | 0.999 | 2.14e-36 | 7.2e-32 |
| Malignant epithelial | C4 T/r-h ECs | RPL29 | 0.437 | 0.998 | 0.999 | 6.56e-41 | 2.21e-36 |
| Malignant epithelial | C4 T/r-h ECs | MIF | 0.436 | 0.931 | 0.951 | 2.75e-18 | 9.28e-14 |
| Malignant epithelial | C4 T/r-h ECs | TOMM7 | 0.428 | 0.906 | 0.958 | 3.89e-15 | 1.31e-10 |
| Malignant epithelial | C4 T/r-h ECs | NBEAL1 | 0.423 | 0.794 | 0.898 | 2.26e-07 | 0.00763 |
| Malignant epithelial | C4 T/r-h ECs | RPL18A | 0.416 | 0.998 | 0.997 | 2.48e-32 | 8.36e-28 |
| Malignant epithelial | C4 T/r-h ECs | RPL23 | 0.413 | 0.971 | 0.993 | 5.95e-25 | 2e-20 |
| Malignant epithelial | C4 T/r-h ECs | RPL38 | 0.41 | 0.994 | 0.998 | 1.36e-35 | 4.59e-31 |
| Malignant epithelial | C4 T/r-h ECs | RPL7 | 0.407 | 0.998 | 0.998 | 7.52e-26 | 2.54e-21 |
| Malignant epithelial | C4 T/r-h ECs | RPS20 | 0.406 | 1.0 | 0.998 | 6.08e-41 | 2.05e-36 |
| Malignant epithelial | C4 T/r-h ECs | MZT2A | 0.404 | 0.758 | 0.845 | 1.29e-06 | 0.0434 |
| Malignant epithelial | C4 T/r-h ECs | SNRPD2 | 0.403 | 0.904 | 0.965 | 1.27e-13 | 4.28e-09 |
| Malignant epithelial | C4 T/r-h ECs | ATP5E | 0.401 | 0.992 | 0.996 | 1.67e-21 | 5.62e-17 |
| Malignant epithelial | C4 T/r-h ECs | RPL35 | 0.4 | 1.0 | 0.999 | 1.52e-40 | 5.13e-36 |
| Malignant epithelial | C4 T/r-h ECs | TCEB2 | 0.399 | 0.954 | 0.983 | 1.37e-08 | 0.00046 |
| Malignant epithelial | C4 T/r-h ECs | RPL14 | 0.397 | 0.996 | 0.998 | 7.54e-35 | 2.54e-30 |
| Malignant epithelial | C4 T/r-h ECs | RPL28 | 0.39 | 1.0 | 1.0 | 1.82e-38 | 6.13e-34 |
| Malignant epithelial | C4 T/r-h ECs | RPL24 | 0.387 | 0.994 | 0.996 | 2.83e-35 | 9.55e-31 |
| Malignant epithelial | C4 T/r-h ECs | RPLP0 | 0.386 | 0.981 | 0.995 | 1.7e-14 | 5.72e-10 |
| Malignant epithelial | C4 T/r-h ECs | RPL37 | 0.385 | 0.998 | 0.999 | 8.490000000000001e-37 | 2.86e-32 |
| Malignant epithelial | C4 T/r-h ECs | RPS19 | 0.384 | 1.0 | 1.0 | 7.339999999999999e-41 | 2.47e-36 |
| Malignant epithelial | C4 T/r-h ECs | RPLP2 | 0.382 | 1.0 | 1.0 | 1.03e-49 | 3.46e-45 |
| Malignant epithelial | C4 T/r-h ECs | RPS21 | 0.381 | 0.992 | 0.997 | 4.600000000000001e-22 | 1.55e-17 |
| Malignant epithelial | C4 T/r-h ECs | RPL39 | 0.376 | 1.0 | 0.999 | 6.33e-28 | 2.13e-23 |
| Malignant epithelial | C4 T/r-h ECs | RPL30 | 0.373 | 1.0 | 0.999 | 1.17e-27 | 3.95e-23 |
| Malignant epithelial | C4 T/r-h ECs | RPS18 | 0.372 | 1.0 | 1.0 | 8.26e-41 | 2.78e-36 |
| Malignant epithelial | C4 T/r-h ECs | UQCRH | 0.368 | 0.883 | 0.935 | 1.72e-10 | 5.8e-06 |
| Malignant epithelial | C4 T/r-h ECs | RPS8 | 0.366 | 0.99 | 0.997 | 3.6e-19 | 1.21e-14 |
| Malignant epithelial | C4 T/r-h ECs | RPL11 | 0.365 | 0.996 | 0.998 | 6.249999999999999e-34 | 2.11e-29 |
| Malignant epithelial | C4 T/r-h ECs | RPS27 | 0.365 | 0.998 | 1.0 | 5.079999999999999e-23 | 1.71e-18 |
| Malignant epithelial | C4 T/r-h ECs | GAPDH | 0.363 | 1.0 | 0.998 | 9.35e-16 | 3.15e-11 |
| Malignant epithelial | C4 T/r-h ECs | ATP5G2 | 0.363 | 0.923 | 0.978 | 8.16e-13 | 2.75e-08 |
| Malignant epithelial | C4 T/r-h ECs | RPL5 | 0.36 | 0.91 | 0.962 | 6.09e-09 | 0.000205 |
| Malignant epithelial | C4 T/r-h ECs | UBA52 | 0.355 | 0.998 | 0.996 | 1.89e-31 | 6.350000000000001e-27 |
| Malignant epithelial | C4 T/r-h ECs | RPS23 | 0.351 | 0.996 | 0.998 | 5.64e-31 | 1.9e-26 |
| Malignant epithelial | C4 T/r-h ECs | RPL27A | 0.349 | 0.996 | 0.998 | 3.46e-38 | 1.16e-33 |
| Malignant epithelial | C4 T/r-h ECs | RPS7 | 0.349 | 0.992 | 0.998 | 2.77e-27 | 9.339999999999999e-23 |
| Malignant epithelial | C4 T/r-h ECs | S100A6 | 0.34 | 1.0 | 1.0 | 1.35e-10 | 4.55e-06 |
| Malignant epithelial | C4 T/r-h ECs | TMEM258 | 0.332 | 0.892 | 0.945 | 6.01e-08 | 0.00202 |
| Malignant epithelial | C4 T/r-h ECs | RPL32 | 0.324 | 1.0 | 1.0 | 1.19e-24 | 4e-20 |
| Malignant epithelial | C4 T/r-h ECs | RPS9 | 0.324 | 0.992 | 0.995 | 4.48e-18 | 1.51e-13 |
| Malignant epithelial | C4 T/r-h ECs | RPL13A | 0.319 | 1.0 | 0.999 | 3.48e-19 | 1.17e-14 |
| Malignant epithelial | C4 T/r-h ECs | RPLP1 | 0.318 | 1.0 | 1.0 | 7.86e-32 | 2.65e-27 |
| Malignant epithelial | C4 T/r-h ECs | RPL13 | 0.318 | 1.0 | 1.0 | 1.95e-28 | 6.56e-24 |
| Malignant epithelial | C4 T/r-h ECs | ATP5L | 0.317 | 0.95 | 0.98 | 6.61e-12 | 2.23e-07 |
| Malignant epithelial | C4 T/r-h ECs | RPS3A | 0.316 | 0.988 | 0.996 | 3.6e-17 | 1.21e-12 |
| Malignant epithelial | C4 T/r-h ECs | RPL10A | 0.31 | 0.99 | 0.996 | 3.01e-13 | 1.01e-08 |
| Malignant epithelial | C4 T/r-h ECs | IFITM3 | 0.308 | 0.925 | 0.951 | 1.2e-06 | 0.0406 |
| Malignant epithelial | C4 T/r-h ECs | KIF1C | 0.306 | 0.065 | 0.167 | 5.9e-08 | 0.00199 |
| Malignant epithelial | C4 T/r-h ECs | RPS10 | 0.302 | 0.96 | 0.987 | 1.34e-09 | 4.53e-05 |
| Malignant epithelial | C4 T/r-h ECs | RPS27A | 0.301 | 0.988 | 0.998 | 4.159999999999999e-23 | 1.4e-18 |

| compartment | subcluster | gene | avg_log2FC | pct.1 | pct.2 | p_val | p_val_adj |
|---|---|---|---|---|---|---|---|
| Malignant epithelial | C4 T/r-h ECs | COX6B1 | 0.292 | 0.96 | 0.991 | 1.41e-09 | 4.75e-05 |
| Malignant epithelial | C4 T/r-h ECs | RPS12 | 0.291 | 1.0 | 0.999 | 2.36e-08 | 0.000795 |
| Malignant epithelial | C4 T/r-h ECs | RPL22 | 0.287 | 0.96 | 0.986 | 9.15e-12 | 3.08e-07 |
| Malignant epithelial | C4 T/r-h ECs | RPS6 | 0.284 | 0.996 | 0.999 | 1.21e-18 | 4.06e-14 |
| Malignant epithelial | C4 T/r-h ECs | RPL27 | 0.283 | 0.969 | 0.994 | 3.62e-15 | 1.22e-10 |
| Malignant epithelial | C4 T/r-h ECs | RPS5 | 0.281 | 0.983 | 0.991 | 7.74e-08 | 0.00261 |
| Malignant epithelial | C4 T/r-h ECs | RPL9 | 0.28 | 0.958 | 0.99 | 2.09e-10 | 7.06e-06 |
| Malignant epithelial | C4 T/r-h ECs | RPL23A | 0.276 | 0.996 | 0.996 | 1.5e-13 | 5.05e-09 |
| Malignant epithelial | C4 T/r-h ECs | HINT1 | 0.276 | 0.94 | 0.972 | 1.01e-07 | 0.00342 |
| Malignant epithelial | C4 T/r-h ECs | DNAJB2 | 0.274 | 0.152 | 0.299 | 5.76e-08 | 0.00194 |
| Malignant epithelial | C4 T/r-h ECs | COX7C | 0.273 | 0.96 | 0.992 | 7.66e-12 | 2.58e-07 |
| Malignant epithelial | C4 T/r-h ECs | SNRNP27 | 0.267 | 0.167 | 0.307 | 5.02e-07 | 0.0169 |
| Malignant epithelial | C4 T/r-h ECs | NUTM2B-AS1 | 0.266 | 0.221 | 0.394 | 1.36e-07 | 0.00457 |
| Malignant epithelial | C4 T/r-h ECs | NEMF | 0.265 | 0.167 | 0.327 | 4.03e-08 | 0.00136 |
| Malignant epithelial | C4 T/r-h ECs | NARS | 0.263 | 0.14 | 0.266 | 4.54e-07 | 0.0153 |
| Malignant epithelial | C4 T/r-h ECs | PLXNA3 | 0.259 | 0.06 | 0.151 | 3.78e-07 | 0.0128 |
| Malignant epithelial | C4 T/r-h ECs | PI3 | 0.258 | 0.069 | 0.166 | 5.69e-08 | 0.00192 |
| Malignant epithelial | C4 T/r-h ECs | RPS11 | 0.257 | 0.996 | 0.997 | 1.37e-17 | 4.62e-13 |
| Malignant epithelial | C4 T/r-h ECs | NACA | 0.257 | 0.977 | 0.995 | 5.75e-16 | 1.94e-11 |
| Malignant epithelial | C4 T/r-h ECs | FAU | 0.256 | 0.985 | 0.998 | 1.37e-24 | 4.6e-20 |
| Malignant epithelial | C5 Am/CEA-like ECs | CEACAM7 | 5.607 | 0.169 | 0.011 | 3e-74 | 1.01e-69 |
| Malignant epithelial | C5 Am/CEA-like ECs | SLC26A3 | 5.482 | 0.188 | 0.013 | 1.26e-81 | 4.25e-77 |
| Malignant epithelial | C5 Am/CEA-like ECs | FABP3 | 4.802 | 0.123 | 0.015 | 1.24e-34 | 4.18e-30 |
| Malignant epithelial | C5 Am/CEA-like ECs | ACHE | 4.297 | 0.517 | 0.061 | 6.03e-158 | 2.03e-153 |
| Malignant epithelial | C5 Am/CEA-like ECs | CPE | 4.21 | 0.169 | 0.017 | 1.66e-54 | 5.59e-50 |
| Malignant epithelial | C5 Am/CEA-like ECs | TM4SF20 | 4.176 | 0.157 | 0.022 | 3.61e-37 | 1.22e-32 |
| Malignant epithelial | C5 Am/CEA-like ECs | RBP4 | 3.734 | 0.13 | 0.019 | 5.31e-30 | 1.79e-25 |
| Malignant epithelial | C5 Am/CEA-like ECs | MUC17 | 3.687 | 0.517 | 0.099 | 6.580000000000001e-105 | 2.22e-100 |
| Malignant epithelial | C5 Am/CEA-like ECs | RSAD2 | 3.682 | 0.176 | 0.032 | 6.3e-33 | 2.12e-28 |
| Malignant epithelial | C5 Am/CEA-like ECs | PARM1 | 3.675 | 0.157 | 0.018 | 2.69e-46 | 9.079999999999999e-42 |
| Malignant epithelial | C5 Am/CEA-like ECs | PSTPIP1 | 3.622 | 0.142 | 0.026 | 1.12e-25 | 3.77e-21 |
| Malignant epithelial | C5 Am/CEA-like ECs | BTNL3 | 3.565 | 0.1 | 0.012 | 1.12e-27 | 3.76e-23 |
| Malignant epithelial | C5 Am/CEA-like ECs | MUC13 | 3.558 | 0.801 | 0.285 | 1.73e-110 | 5.84e-106 |
| Malignant epithelial | C5 Am/CEA-like ECs | TFF1 | 3.542 | 0.467 | 0.154 | 5.54e-47 | 1.87e-42 |
| Malignant epithelial | C5 Am/CEA-like ECs | PKIB | 3.341 | 0.199 | 0.039 | 5.01e-34 | 1.69e-29 |
| Malignant epithelial | C5 Am/CEA-like ECs | GUCA2A | 3.336 | 0.157 | 0.022 | 1.36e-38 | 4.58e-34 |
| Malignant epithelial | C5 Am/CEA-like ECs | BMP2 | 3.311 | 0.287 | 0.044 | 1.22e-64 | 4.12e-60 |
| Malignant epithelial | C5 Am/CEA-like ECs | BCAS1 | 3.278 | 0.36 | 0.065 | 2.51e-70 | 8.45e-66 |
| Malignant epithelial | C5 Am/CEA-like ECs | FABP2 | 3.235 | 0.115 | 0.019 | 3.29e-23 | 1.11e-18 |
| Malignant epithelial | C5 Am/CEA-like ECs | SLC40A1 | 3.203 | 0.59 | 0.172 | 4.2e-77 | 1.41e-72 |
| Malignant epithelial | C5 Am/CEA-like ECs | SDCBP2 | 3.199 | 0.805 | 0.246 | 5.719999999999999e-126 | 1.93e-121 |
| Malignant epithelial | C5 Am/CEA-like ECs | GPA33 | 3.195 | 0.287 | 0.067 | 2.26e-41 | 7.61e-37 |
| Malignant epithelial | C5 Am/CEA-like ECs | ABHD11-AS1 | 3.159 | 0.525 | 0.139 | 3.54e-73 | 1.19e-68 |
| Malignant epithelial | C5 Am/CEA-like ECs | ULBP2 | 3.131 | 0.199 | 0.035 | 2.16e-38 | 7.279999999999999e-34 |
| Malignant epithelial | C5 Am/CEA-like ECs | ENTPD8 | 3.062 | 0.176 | 0.035 | 3.5e-29 | 1.18e-24 |
| Malignant epithelial | C5 Am/CEA-like ECs | C4orf19 | 3.031 | 0.161 | 0.033 | 2.6e-26 | 8.76e-22 |
| Malignant epithelial | C5 Am/CEA-like ECs | CLIC5 | 3.029 | 0.333 | 0.079 | 3.11e-47 | 1.05e-42 |
| Malignant epithelial | C5 Am/CEA-like ECs | THRB | 3.024 | 0.103 | 0.02 | 3.22e-18 | 1.09e-13 |
| Malignant epithelial | C5 Am/CEA-like ECs | SLC6A8 | 3.021 | 0.441 | 0.153 | 7.919999999999999e-42 | 2.67e-37 |
| Malignant epithelial | C5 Am/CEA-like ECs | BTNL8 | 3.02 | 0.195 | 0.032 | 1.31e-39 | 4.41e-35 |
| Malignant epithelial | C5 Am/CEA-like ECs | CEACAM1 | 3.016 | 0.617 | 0.239 | 4.88e-57 | 1.64e-52 |
| Malignant epithelial | C5 Am/CEA-like ECs | CEACAM5 | 3.008 | 0.851 | 0.355 | 3.26e-103 | 1.1e-98 |
| Malignant epithelial | C5 Am/CEA-like ECs | SLC6A20 | 3.007 | 0.253 | 0.042 | 6.98e-51 | 2.35e-46 |
| Malignant epithelial | C5 Am/CEA-like ECs | LINC01207 | 2.988 | 0.441 | 0.091 | 2.08e-75 | 6.99e-71 |
| Malignant epithelial | C5 Am/CEA-like ECs | DHRS9 | 2.982 | 0.483 | 0.143 | 1.9e-57 | 6.399999999999999e-53 |
| Malignant epithelial | C5 Am/CEA-like ECs | CEACAM6 | 2.971 | 0.858 | 0.401 | 6.98e-94 | 2.35e-89 |
| Malignant epithelial | C5 Am/CEA-like ECs | LRRC19 | 2.961 | 0.123 | 0.022 | 2.53e-22 | 8.52e-18 |
| Malignant epithelial | C5 Am/CEA-like ECs | RP11-395B7.2 | 2.953 | 0.123 | 0.024 | 4.2e-21 | 1.42e-16 |
| Malignant epithelial | C5 Am/CEA-like ECs | TMEM56 | 2.914 | 0.123 | 0.025 | 1.85e-19 | 6.22e-15 |
| Malignant epithelial | C5 Am/CEA-like ECs | PAG1 | 2.904 | 0.28 | 0.072 | 9.099999999999999e-35 | 3.07e-30 |
| Malignant epithelial | C5 Am/CEA-like ECs | PTPRH | 2.894 | 0.287 | 0.072 | 1.8e-37 | 6.07e-33 |
| Malignant epithelial | C5 Am/CEA-like ECs | PPP2R5B | 2.89 | 0.268 | 0.056 | 6.01e-42 | 2.03e-37 |
| Malignant epithelial | C5 Am/CEA-like ECs | B3GNT7 | 2.884 | 0.157 | 0.04 | 3.32e-19 | 1.12e-14 |
| Malignant epithelial | C5 Am/CEA-like ECs | RNF186 | 2.875 | 0.241 | 0.053 | 1.88e-35 | 6.34e-31 |

| compartment | subcluster | gene | avg_log2FC | pct.1 | pct.2 | p_val | p_val_adj |
|---|---|---|---|---|---|---|---|
| Malignant epithelial | C5 Am/CEA-like ECs | PPP1R14C | 2.875 | 0.149 | 0.03 | 3.11e-24 | 1.05e-19 |
| Malignant epithelial | C5 Am/CEA-like ECs | HHLA2 | 2.859 | 0.287 | 0.057 | 2.52e-48 | 8.5e-44 |
| Malignant epithelial | C5 Am/CEA-like ECs | LINC01133 | 2.815 | 0.418 | 0.1 | 2.92e-59 | 9.86e-55 |
| Malignant epithelial | C5 Am/CEA-like ECs | HEPH | 2.815 | 0.379 | 0.093 | 1.75e-52 | 5.889999999999999e-48 |
| Malignant epithelial | C5 Am/CEA-like ECs | SCIN | 2.767 | 0.188 | 0.043 | 5.870000000000001e-26 | 1.98e-21 |
| Malignant epithelial | C5 Am/CEA-like ECs | MFSD2A | 2.761 | 0.111 | 0.03 | 1.68e-12 | 5.67e-08 |
| Malignant epithelial | C5 Am/CEA-like ECs | KRT20 | 2.746 | 0.678 | 0.296 | 5.96e-61 | 2.01e-56 |
| Malignant epithelial | C5 Am/CEA-like ECs | CAPN5 | 2.746 | 0.433 | 0.111 | 8.85e-58 | 2.98e-53 |
| Malignant epithelial | C5 Am/CEA-like ECs | HK2 | 2.741 | 0.349 | 0.095 | 3.03e-41 | 1.02e-36 |
| Malignant epithelial | C5 Am/CEA-like ECs | COL16A1 | 2.74 | 0.107 | 0.026 | 2.02e-14 | 6.8e-10 |
| Malignant epithelial | C5 Am/CEA-like ECs | LINC01559 | 2.733 | 0.157 | 0.043 | 1.54e-17 | 5.2e-13 |
| Malignant epithelial | C5 Am/CEA-like ECs | FABP1 | 2.727 | 0.287 | 0.074 | 8.78e-35 | 2.96e-30 |
| Malignant epithelial | C5 Am/CEA-like ECs | ASS1 | 2.721 | 0.743 | 0.437 | 5.71e-49 | 1.92e-44 |
| Malignant epithelial | C5 Am/CEA-like ECs | CREB3L1 | 2.66 | 0.349 | 0.08 | 6.26e-51 | 2.11e-46 |
| Malignant epithelial | C5 Am/CEA-like ECs | C4BPB | 2.644 | 0.199 | 0.048 | 6.34e-26 | 2.14e-21 |
| Malignant epithelial | C5 Am/CEA-like ECs | SMPD1 | 2.64 | 0.287 | 0.077 | 2.46e-34 | 8.3e-30 |
| Malignant epithelial | C5 Am/CEA-like ECs | TMPRSS2 | 2.595 | 0.605 | 0.219 | 6.13e-58 | 2.07e-53 |
| Malignant epithelial | C5 Am/CEA-like ECs | GK | 2.594 | 0.241 | 0.058 | 2.29e-32 | 7.72e-28 |
| Malignant epithelial | C5 Am/CEA-like ECs | CDHR5 | 2.589 | 0.533 | 0.154 | 3.82e-65 | 1.29e-60 |
| Malignant epithelial | C5 Am/CEA-like ECs | LINC00675 | 2.581 | 0.303 | 0.076 | 8.000000000000001e-39 | 2.69e-34 |
| Malignant epithelial | C5 Am/CEA-like ECs | C6orf222 | 2.525 | 0.314 | 0.077 | 1.36e-40 | 4.59e-36 |
| Malignant epithelial | C5 Am/CEA-like ECs | ZG16B | 2.519 | 0.494 | 0.152 | 1.03e-54 | 3.48e-50 |
| Malignant epithelial | C5 Am/CEA-like ECs | TP53INP2 | 2.516 | 0.276 | 0.063 | 2.23e-39 | 7.530000000000001e-35 |
| Malignant epithelial | C5 Am/CEA-like ECs | APOBEC1 | 2.504 | 0.207 | 0.068 | 7.86e-18 | 2.65e-13 |
| Malignant epithelial | C5 Am/CEA-like ECs | FUT3 | 2.498 | 0.41 | 0.108 | 2.93e-51 | 9.87e-47 |
| Malignant epithelial | C5 Am/CEA-like ECs | SEMA6A | 2.498 | 0.107 | 0.024 | 1.7e-15 | 5.73e-11 |
| Malignant epithelial | C5 Am/CEA-like ECs | AGR2 | 2.495 | 0.858 | 0.553 | 8.370000000000001e-63 | 2.82e-58 |
| Malignant epithelial | C5 Am/CEA-like ECs | GLRX | 2.492 | 0.605 | 0.299 | 6.85e-40 | 2.31e-35 |
| Malignant epithelial | C5 Am/CEA-like ECs | CD55 | 2.464 | 0.77 | 0.396 | 1.9e-64 | 6.42e-60 |
| Malignant epithelial | C5 Am/CEA-like ECs | DNAJB9 | 2.449 | 0.483 | 0.203 | 9.63e-33 | 3.24e-28 |
| Malignant epithelial | C5 Am/CEA-like ECs | RP1-122P22.4 | 2.447 | 0.364 | 0.112 | 3.55e-36 | 1.2e-31 |
| Malignant epithelial | C5 Am/CEA-like ECs | MOB3B | 2.442 | 0.172 | 0.044 | 1.23e-20 | 4.15e-16 |
| Malignant epithelial | C5 Am/CEA-like ECs | SEPP1 | 2.425 | 0.751 | 0.351 | 1.1e-60 | 3.71e-56 |
| Malignant epithelial | C5 Am/CEA-like ECs | SPINK1 | 2.423 | 0.682 | 0.252 | 7.78e-62 | 2.62e-57 |
| Malignant epithelial | C5 Am/CEA-like ECs | DLL4 | 2.422 | 0.107 | 0.035 | 1.99e-09 | 6.7e-05 |
| Malignant epithelial | C5 Am/CEA-like ECs | IGHM | 2.42 | 0.161 | 0.055 | 9.46e-13 | 3.19e-08 |
| Malignant epithelial | C5 Am/CEA-like ECs | SLC44A4 | 2.416 | 0.709 | 0.259 | 2.97e-75 | 9.99e-71 |
| Malignant epithelial | C5 Am/CEA-like ECs | CDHR2 | 2.409 | 0.238 | 0.068 | 1.89e-24 | 6.37e-20 |
| Malignant epithelial | C5 Am/CEA-like ECs | MIER3 | 2.4 | 0.142 | 0.04 | 5.2e-15 | 1.75e-10 |
| Malignant epithelial | C5 Am/CEA-like ECs | CYP4F3 | 2.385 | 0.375 | 0.1 | 1.16e-44 | 3.92e-40 |
| Malignant epithelial | C5 Am/CEA-like ECs | CHAC1 | 2.371 | 0.161 | 0.048 | 1.28e-15 | 4.31e-11 |
| Malignant epithelial | C5 Am/CEA-like ECs | RASGRP3 | 2.369 | 0.146 | 0.029 | 5.789999999999999e-24 | 1.95e-19 |
| Malignant epithelial | C5 Am/CEA-like ECs | MUC3A | 2.354 | 0.659 | 0.273 | 4.65e-59 | 1.57e-54 |
| Malignant epithelial | C5 Am/CEA-like ECs | CYP3A5 | 2.348 | 0.751 | 0.342 | 1.34e-65 | 4.5e-61 |
| Malignant epithelial | C5 Am/CEA-like ECs | ITM2C | 2.345 | 0.536 | 0.293 | 8.02e-28 | 2.7e-23 |
| Malignant epithelial | C5 Am/CEA-like ECs | CDH17 | 2.341 | 0.728 | 0.285 | 1.85e-70 | 6.240000000000001e-66 |
| Malignant epithelial | C5 Am/CEA-like ECs | GAST | 2.313 | 0.153 | 0.044 | 1.08e-15 | 3.63e-11 |
| Malignant epithelial | C5 Am/CEA-like ECs | PIP5K1B | 2.31 | 0.249 | 0.073 | 1.48e-25 | 4.97e-21 |
| Malignant epithelial | C5 Am/CEA-like ECs | OCLN | 2.306 | 0.506 | 0.194 | 7.68e-43 | 2.59e-38 |
| Malignant epithelial | C5 Am/CEA-like ECs | CXADR | 2.282 | 0.72 | 0.332 | 1.11e-57 | 3.73e-53 |
| Malignant epithelial | C5 Am/CEA-like ECs | FXYD3 | 2.282 | 0.912 | 0.867 | 1.47e-46 | 4.95e-42 |
| Malignant epithelial | C5 Am/CEA-like ECs | PLS1 | 2.276 | 0.529 | 0.188 | 5.03e-49 | 1.69e-44 |
| Malignant epithelial | C5 Am/CEA-like ECs | ATP10B | 2.275 | 0.452 | 0.162 | 1.58e-38 | 5.309999999999999e-34 |
| Malignant epithelial | C5 Am/CEA-like ECs | PTPRR | 2.27 | 0.119 | 0.04 | 7.4e-10 | 2.49e-05 |
| Malignant epithelial | C5 Am/CEA-like ECs | SMIM5 | 2.258 | 0.241 | 0.06 | 1.64e-30 | 5.53e-26 |
| Malignant epithelial | C5 Am/CEA-like ECs | C6orf223 | 2.257 | 0.176 | 0.056 | 7.42e-16 | 2.5e-11 |
| Malignant epithelial | C5 Am/CEA-like ECs | EIF2AK3 | 2.231 | 0.1 | 0.035 | 4.72e-08 | 0.00159 |
| Malignant epithelial | C5 Am/CEA-like ECs | B3GNT8 | 2.229 | 0.184 | 0.059 | 2.22e-16 | 7.47e-12 |
| Malignant epithelial | C5 Am/CEA-like ECs | KRT80 | 2.218 | 0.28 | 0.103 | 1.86e-20 | 6.27e-16 |
| Malignant epithelial | C5 Am/CEA-like ECs | FAM132A | 2.208 | 0.153 | 0.047 | 3.63e-14 | 1.22e-09 |
| Malignant epithelial | C5 Am/CEA-like ECs | TSPAN1 | 2.205 | 0.82 | 0.528 | 3.04e-55 | 1.02e-50 |
| Malignant epithelial | C5 Am/CEA-like ECs | ALOX5 | 2.196 | 0.119 | 0.031 | 4.03e-14 | 1.36e-09 |
| Malignant epithelial | C5 Am/CEA-like ECs | QSOX1 | 2.176 | 0.582 | 0.281 | 6.97e-37 | 2.35e-32 |
| Malignant epithelial | C5 Am/CEA-like ECs | KALRN | 2.169 | 0.153 | 0.056 | 4.86e-11 | 1.64e-06 |

| compartment | subcluster | gene | avg_log2FC | pct.1 | pct.2 | p_val | p_val_adj |
|---|---|---|---|---|---|---|---|
| Malignant epithelial | C5 Am/CEA-like ECs | SPRR1B | 2.166 | 0.103 | 0.035 | 9e-09 | 0.000303 |
| Malignant epithelial | C5 Am/CEA-like ECs | IL2RG | 2.164 | 0.494 | 0.209 | 8.509999999999999e-34 | 2.87e-29 |
| Malignant epithelial | C5 Am/CEA-like ECs | FA2H | 2.158 | 0.314 | 0.118 | 7.000000000000001e-23 | 2.36e-18 |
| Malignant epithelial | C5 Am/CEA-like ECs | CA12 | 2.152 | 0.134 | 0.038 | 4.9e-14 | 1.65e-09 |
| Malignant epithelial | C5 Am/CEA-like ECs | MFSD4 | 2.15 | 0.103 | 0.033 | 2.66e-09 | 8.95e-05 |
| Malignant epithelial | C5 Am/CEA-like ECs | JSRP1 | 2.138 | 0.134 | 0.039 | 9.94e-14 | 3.35e-09 |
| Malignant epithelial | C5 Am/CEA-like ECs | FAM13A | 2.132 | 0.123 | 0.043 | 2.93e-09 | 9.86e-05 |
| Malignant epithelial | C5 Am/CEA-like ECs | MSMO1 | 2.13 | 0.349 | 0.175 | 9.41e-16 | 3.17e-11 |
| Malignant epithelial | C5 Am/CEA-like ECs | PRSS8 | 2.126 | 0.789 | 0.441 | 3.48e-57 | 1.17e-52 |
| Malignant epithelial | C5 Am/CEA-like ECs | RNASE1 | 2.126 | 0.494 | 0.23 | 8.94e-27 | 3.01e-22 |
| Malignant epithelial | C5 Am/CEA-like ECs | FAM134B | 2.12 | 0.234 | 0.077 | 1.81e-19 | 6.1e-15 |
| Malignant epithelial | C5 Am/CEA-like ECs | ABCG1 | 2.115 | 0.287 | 0.094 | 2.96e-25 | 9.97e-21 |
| Malignant epithelial | C5 Am/CEA-like ECs | UTP14C | 2.112 | 0.126 | 0.051 | 7.34e-08 | 0.00247 |
| Malignant epithelial | C5 Am/CEA-like ECs | GALNT5 | 2.107 | 0.379 | 0.132 | 5.740000000000001e-32 | 1.93e-27 |
| Malignant epithelial | C5 Am/CEA-like ECs | PLOD2 | 2.096 | 0.23 | 0.11 | 9.05e-11 | 3.05e-06 |
| Malignant epithelial | C5 Am/CEA-like ECs | MUC12 | 2.079 | 0.448 | 0.205 | 5.5e-25 | 1.85e-20 |
| Malignant epithelial | C5 Am/CEA-like ECs | TFF3 | 2.075 | 0.835 | 0.615 | 3.68e-37 | 1.24e-32 |
| Malignant epithelial | C5 Am/CEA-like ECs | C9orf72 | 2.066 | 0.188 | 0.061 | 7.12e-16 | 2.4e-11 |
| Malignant epithelial | C5 Am/CEA-like ECs | PHGR1 | 2.063 | 0.847 | 0.547 | 2.98e-40 | 1e-35 |
| Malignant epithelial | C5 Am/CEA-like ECs | SLC6A14 | 2.061 | 0.215 | 0.064 | 1.27e-20 | 4.29e-16 |
| Malignant epithelial | C5 Am/CEA-like ECs | NTN4 | 2.054 | 0.103 | 0.041 | 1.07e-06 | 0.0361 |
| Malignant epithelial | C5 Am/CEA-like ECs | MAPRE2 | 2.05 | 0.103 | 0.042 | 1.42e-06 | 0.0478 |
| Malignant epithelial | C5 Am/CEA-like ECs | CD68 | 2.049 | 0.571 | 0.249 | 9.08e-41 | 3.06e-36 |
| Malignant epithelial | C5 Am/CEA-like ECs | CMPK2 | 2.048 | 0.13 | 0.048 | 2.74e-09 | 9.22e-05 |
| Malignant epithelial | C5 Am/CEA-like ECs | WIPI1 | 2.045 | 0.241 | 0.089 | 3.12e-17 | 1.05e-12 |
| Malignant epithelial | C5 Am/CEA-like ECs | CCRL2 | 2.045 | 0.107 | 0.039 | 5.15e-08 | 0.00174 |
| Malignant epithelial | C5 Am/CEA-like ECs | ATG4A | 2.043 | 0.103 | 0.039 | 2.67e-07 | 0.009 |
| Malignant epithelial | C5 Am/CEA-like ECs | VIPR1 | 2.037 | 0.111 | 0.033 | 3.7e-11 | 1.25e-06 |
| Malignant epithelial | C5 Am/CEA-like ECs | VILL | 2.033 | 0.303 | 0.104 | 3.01e-24 | 1.01e-19 |
| Malignant epithelial | C5 Am/CEA-like ECs | RNF39 | 2.031 | 0.276 | 0.078 | 4.47e-29 | 1.5e-24 |
| Malignant epithelial | C5 Am/CEA-like ECs | CPEB4 | 2.029 | 0.245 | 0.09 | 2.21e-17 | 7.43e-13 |
| Malignant epithelial | C5 Am/CEA-like ECs | LGR4 | 2.025 | 0.249 | 0.072 | 3.83e-25 | 1.29e-20 |
| Malignant epithelial | C5 Am/CEA-like ECs | B3GALT5 | 2.018 | 0.111 | 0.035 | 4.29e-10 | 1.44e-05 |
| Malignant epithelial | C5 Am/CEA-like ECs | GDA | 2.008 | 0.161 | 0.051 | 3.33e-14 | 1.12e-09 |
| Malignant epithelial | C5 Am/CEA-like ECs | SMPDL3A | 2.0 | 0.28 | 0.124 | 6.35e-15 | 2.14e-10 |
| Malignant epithelial | C5 Am/CEA-like ECs | SRD5A3 | 1.995 | 0.234 | 0.108 | 3.12e-11 | 1.05e-06 |
| Malignant epithelial | C5 Am/CEA-like ECs | CNKSR3 | 1.989 | 0.184 | 0.066 | 2.21e-13 | 7.45e-09 |
| Malignant epithelial | C5 Am/CEA-like ECs | MARCH3 | 1.986 | 0.295 | 0.107 | 2.49e-22 | 8.38e-18 |
| Malignant epithelial | C5 Am/CEA-like ECs | TRIB3 | 1.985 | 0.506 | 0.249 | 3.78e-27 | 1.27e-22 |
| Malignant epithelial | C5 Am/CEA-like ECs | TMBIM1 | 1.984 | 0.805 | 0.475 | 7.29e-57 | 2.46e-52 |
| Malignant epithelial | C5 Am/CEA-like ECs | FBXO32 | 1.984 | 0.525 | 0.225 | 2.02e-35 | 6.81e-31 |
| Malignant epithelial | C5 Am/CEA-like ECs | MYO7B | 1.983 | 0.169 | 0.063 | 1.25e-11 | 4.22e-07 |
| Malignant epithelial | C5 Am/CEA-like ECs | VSIG2 | 1.981 | 0.184 | 0.067 | 2.39e-13 | 8.05e-09 |
| Malignant epithelial | C5 Am/CEA-like ECs | GRB10 | 1.981 | 0.119 | 0.043 | 9.99e-09 | 0.000337 |
| Malignant epithelial | C5 Am/CEA-like ECs | ME1 | 1.979 | 0.153 | 0.041 | 5.3e-17 | 1.79e-12 |
| Malignant epithelial | C5 Am/CEA-like ECs | FAM3D | 1.971 | 0.594 | 0.222 | 7.74e-49 | 2.61e-44 |
| Malignant epithelial | C5 Am/CEA-like ECs | CCL15 | 1.969 | 0.318 | 0.129 | 8.21e-20 | 2.77e-15 |
| Malignant epithelial | C5 Am/CEA-like ECs | TM4SF5 | 1.966 | 0.352 | 0.164 | 1.92e-17 | 6.47e-13 |
| Malignant epithelial | C5 Am/CEA-like ECs | C10orf99 | 1.956 | 0.115 | 0.042 | 2.42e-08 | 0.000815 |
| Malignant epithelial | C5 Am/CEA-like ECs | DENND2A | 1.953 | 0.115 | 0.04 | 5.73e-09 | 0.000193 |
| Malignant epithelial | C5 Am/CEA-like ECs | SLC17A5 | 1.95 | 0.215 | 0.084 | 9.94e-14 | 3.35e-09 |
| Malignant epithelial | C5 Am/CEA-like ECs | S100P | 1.946 | 0.923 | 0.535 | 3.96e-73 | 1.33e-68 |
| Malignant epithelial | C5 Am/CEA-like ECs | ARFGEF3 | 1.942 | 0.1 | 0.037 | 4.01e-07 | 0.0135 |
| Malignant epithelial | C5 Am/CEA-like ECs | MMEL1 | 1.938 | 0.211 | 0.079 | 7.8e-14 | 2.63e-09 |
| Malignant epithelial | C5 Am/CEA-like ECs | FSTL3 | 1.937 | 0.356 | 0.157 | 1.12e-19 | 3.76e-15 |
| Malignant epithelial | C5 Am/CEA-like ECs | PRR5L | 1.932 | 0.188 | 0.062 | 1.2e-15 | 4.04e-11 |
| Malignant epithelial | C5 Am/CEA-like ECs | SLC9A1 | 1.93 | 0.195 | 0.068 | 5.92e-15 | 1.99e-10 |
| Malignant epithelial | C5 Am/CEA-like ECs | ARG2 | 1.908 | 0.13 | 0.048 | 4.2e-09 | 0.000141 |
| Malignant epithelial | C5 Am/CEA-like ECs | CXCL16 | 1.899 | 0.582 | 0.285 | 1.92e-36 | 6.46e-32 |
| Malignant epithelial | C5 Am/CEA-like ECs | CLDN9 | 1.899 | 0.146 | 0.061 | 4.14e-08 | 0.00139 |
| Malignant epithelial | C5 Am/CEA-like ECs | TMEM45B | 1.896 | 0.674 | 0.287 | 4.08e-52 | 1.37e-47 |
| Malignant epithelial | C5 Am/CEA-like ECs | TMEM144 | 1.896 | 0.165 | 0.062 | 4.32e-11 | 1.45e-06 |
| Malignant epithelial | C5 Am/CEA-like ECs | TSPAN3 | 1.895 | 0.724 | 0.443 | 5.08e-44 | 1.71e-39 |
| Malignant epithelial | C5 Am/CEA-like ECs | SNX9 | 1.895 | 0.46 | 0.194 | 7.77e-30 | 2.62e-25 |

| compartment | subcluster | gene | avg_log2FC | pct.1 | pct.2 | p_val | p_val_adj |
|---|---|---|---|---|---|---|---|
| Malignant epithelial | C5 Am/CEA-like ECs | AGR3 | 1.895 | 0.28 | 0.097 | 2.76e-21 | 9.3e-17 |
| Malignant epithelial | C5 Am/CEA-like ECs | SMIM2-AS1 | 1.895 | 0.107 | 0.041 | 2.47e-07 | 0.00831 |
| Malignant epithelial | C5 Am/CEA-like ECs | TTC9 | 1.89 | 0.13 | 0.055 | 1.65e-07 | 0.00555 |
| Malignant epithelial | C5 Am/CEA-like ECs | SYTL2 | 1.888 | 0.199 | 0.088 | 5.06e-10 | 1.71e-05 |
| Malignant epithelial | C5 Am/CEA-like ECs | TMEM106B | 1.885 | 0.579 | 0.314 | 2.43e-28 | 8.199999999999999e-24 |
| Malignant epithelial | C5 Am/CEA-like ECs | ISG20 | 1.883 | 0.659 | 0.402 | 2.01e-30 | 6.79e-26 |
| Malignant epithelial | C5 Am/CEA-like ECs | AHCYL2 | 1.882 | 0.249 | 0.1 | 2.2e-15 | 7.41e-11 |
| Malignant epithelial | C5 Am/CEA-like ECs | CBLB | 1.881 | 0.195 | 0.08 | 2.09e-11 | 7.05e-07 |
| Malignant epithelial | C5 Am/CEA-like ECs | MACC1 | 1.87 | 0.414 | 0.193 | 7.73e-22 | 2.6e-17 |
| Malignant epithelial | C5 Am/CEA-like ECs | ENTPD5 | 1.867 | 0.146 | 0.061 | 4.54e-08 | 0.00153 |
| Malignant epithelial | C5 Am/CEA-like ECs | APPL2 | 1.859 | 0.253 | 0.113 | 1.23e-12 | 4.13e-08 |
| Malignant epithelial | C5 Am/CEA-like ECs | ANKS4B | 1.858 | 0.157 | 0.06 | 3.24e-10 | 1.09e-05 |
| Malignant epithelial | C5 Am/CEA-like ECs | BHLHE41 | 1.852 | 0.146 | 0.052 | 8.87e-11 | 2.99e-06 |
| Malignant epithelial | C5 Am/CEA-like ECs | RMND5A | 1.851 | 0.46 | 0.218 | 8.66e-24 | 2.92e-19 |
| Malignant epithelial | C5 Am/CEA-like ECs | PI3 | 1.849 | 0.291 | 0.15 | 2.34e-11 | 7.88e-07 |
| Malignant epithelial | C5 Am/CEA-like ECs | GAN | 1.845 | 0.31 | 0.126 | 2.79e-19 | 9.4e-15 |
| Malignant epithelial | C5 Am/CEA-like ECs | LMO7 | 1.844 | 0.828 | 0.479 | 8.76e-54 | 2.95e-49 |
| Malignant epithelial | C5 Am/CEA-like ECs | CYP4F11 | 1.844 | 0.13 | 0.042 | 3.47e-11 | 1.17e-06 |
| Malignant epithelial | C5 Am/CEA-like ECs | RFK | 1.838 | 0.464 | 0.21 | 5.14e-26 | 1.73e-21 |
| Malignant epithelial | C5 Am/CEA-like ECs | NDRG1 | 1.833 | 0.617 | 0.317 | 1.57e-31 | 5.3e-27 |
| Malignant epithelial | C5 Am/CEA-like ECs | RIOK3 | 1.822 | 0.521 | 0.243 | 1.42e-30 | 4.8e-26 |
| Malignant epithelial | C5 Am/CEA-like ECs | STK40 | 1.821 | 0.218 | 0.082 | 7.17e-15 | 2.42e-10 |
| Malignant epithelial | C5 Am/CEA-like ECs | CYP4F12 | 1.821 | 0.157 | 0.062 | 1.49e-09 | 5.01e-05 |
| Malignant epithelial | C5 Am/CEA-like ECs | LNX2 | 1.821 | 0.169 | 0.072 | 3e-09 | 0.000101 |
| Malignant epithelial | C5 Am/CEA-like ECs | UPP1 | 1.818 | 0.559 | 0.295 | 1.64e-28 | 5.529999999999999e-24 |
| Malignant epithelial | C5 Am/CEA-like ECs | GFPT1 | 1.808 | 0.483 | 0.215 | 1.25e-29 | 4.22e-25 |
| Malignant epithelial | C5 Am/CEA-like ECs | PRRG1 | 1.804 | 0.195 | 0.096 | 2.81e-08 | 0.000948 |
| Malignant epithelial | C5 Am/CEA-like ECs | CTD-2396E7.11 | 1.803 | 0.199 | 0.089 | 5.6e-10 | 1.89e-05 |
| Malignant epithelial | C5 Am/CEA-like ECs | ABCA5 | 1.803 | 0.138 | 0.057 | 7.58e-08 | 0.00256 |
| Malignant epithelial | C5 Am/CEA-like ECs | GPR153 | 1.802 | 0.149 | 0.063 | 3.09e-08 | 0.00104 |
| Malignant epithelial | C5 Am/CEA-like ECs | OSGIN1 | 1.797 | 0.115 | 0.041 | 1.41e-08 | 0.000475 |
| Malignant epithelial | C5 Am/CEA-like ECs | NEDD9 | 1.793 | 0.303 | 0.129 | 2.85e-17 | 9.61e-13 |
| Malignant epithelial | C5 Am/CEA-like ECs | CDR2 | 1.792 | 0.161 | 0.077 | 3.91e-07 | 0.0132 |
| Malignant epithelial | C5 Am/CEA-like ECs | CES2 | 1.791 | 0.429 | 0.213 | 5.63e-21 | 1.9e-16 |
| Malignant epithelial | C5 Am/CEA-like ECs | RASEF | 1.785 | 0.284 | 0.127 | 2.87e-14 | 9.68e-10 |
| Malignant epithelial | C5 Am/CEA-like ECs | C17orf67 | 1.783 | 0.203 | 0.08 | 1.96e-12 | 6.6e-08 |
| Malignant epithelial | C5 Am/CEA-like ECs | C12orf75 | 1.78 | 0.812 | 0.563 | 5.35e-42 | 1.8e-37 |
| Malignant epithelial | C5 Am/CEA-like ECs | RNF103 | 1.78 | 0.398 | 0.173 | 1.52e-23 | 5.12e-19 |
| Malignant epithelial | C5 Am/CEA-like ECs | MAP1LC3A | 1.778 | 0.498 | 0.241 | 3.44e-25 | 1.16e-20 |
| Malignant epithelial | C5 Am/CEA-like ECs | HDAC11 | 1.776 | 0.149 | 0.061 | 5.52e-09 | 0.000186 |
| Malignant epithelial | C5 Am/CEA-like ECs | MIR22HG | 1.77 | 0.291 | 0.114 | 1.91e-18 | 6.42e-14 |
| Malignant epithelial | C5 Am/CEA-like ECs | RP11-462G2.1 | 1.769 | 0.632 | 0.316 | 4.17e-36 | 1.41e-31 |
| Malignant epithelial | C5 Am/CEA-like ECs | EAF2 | 1.767 | 0.138 | 0.046 | 3.06e-11 | 1.03e-06 |
| Malignant epithelial | C5 Am/CEA-like ECs | EPS8L1 | 1.763 | 0.571 | 0.296 | 2.25e-29 | 7.58e-25 |
| Malignant epithelial | C5 Am/CEA-like ECs | OSTM1 | 1.763 | 0.157 | 0.067 | 1.37e-08 | 0.00046 |
| Malignant epithelial | C5 Am/CEA-like ECs | RAB11FIP4 | 1.763 | 0.169 | 0.08 | 2.65e-07 | 0.00893 |
| Malignant epithelial | C5 Am/CEA-like ECs | MUC20 | 1.759 | 0.188 | 0.072 | 7.44e-12 | 2.51e-07 |
| Malignant epithelial | C5 Am/CEA-like ECs | USP53 | 1.755 | 0.529 | 0.254 | 4e-29 | 1.35e-24 |
| Malignant epithelial | C5 Am/CEA-like ECs | MGLL | 1.754 | 0.571 | 0.247 | 7.07e-39 | 2.38e-34 |
| Malignant epithelial | C5 Am/CEA-like ECs | PLXNA2 | 1.748 | 0.115 | 0.033 | 1.26e-11 | 4.25e-07 |
| Malignant epithelial | C5 Am/CEA-like ECs | TSPAN13 | 1.747 | 0.655 | 0.342 | 2.55e-37 | 8.59e-33 |
| Malignant epithelial | C5 Am/CEA-like ECs | MMP28 | 1.747 | 0.1 | 0.032 | 1.02e-08 | 0.000344 |
| Malignant epithelial | C5 Am/CEA-like ECs | TLE4 | 1.744 | 0.314 | 0.134 | 1.92e-17 | 6.47e-13 |
| Malignant epithelial | C5 Am/CEA-like ECs | DAPP1 | 1.743 | 0.149 | 0.057 | 1.1e-09 | 3.71e-05 |
| Malignant epithelial | C5 Am/CEA-like ECs | SH3RF1 | 1.735 | 0.261 | 0.115 | 1.62e-13 | 5.44e-09 |
| Malignant epithelial | C5 Am/CEA-like ECs | CTSS | 1.733 | 0.686 | 0.425 | 4.26e-34 | 1.44e-29 |
| Malignant epithelial | C5 Am/CEA-like ECs | MIR210HG | 1.73 | 0.161 | 0.073 | 7.21e-08 | 0.00243 |
| Malignant epithelial | C5 Am/CEA-like ECs | FAM46A | 1.726 | 0.498 | 0.231 | 1.8e-28 | 6.08e-24 |
| Malignant epithelial | C5 Am/CEA-like ECs | CORO2A | 1.72 | 0.276 | 0.122 | 3.3e-14 | 1.11e-09 |
| Malignant epithelial | C5 Am/CEA-like ECs | SDR16C5 | 1.718 | 0.421 | 0.172 | 2.41e-26 | 8.11e-22 |
| Malignant epithelial | C5 Am/CEA-like ECs | CLDN23 | 1.714 | 0.157 | 0.052 | 3.37e-13 | 1.13e-08 |
| Malignant epithelial | C5 Am/CEA-like ECs | PRSS3 | 1.711 | 0.916 | 0.687 | 5.96e-65 | 2.01e-60 |
| Malignant epithelial | C5 Am/CEA-like ECs | STARD4 | 1.705 | 0.13 | 0.049 | 9.55e-09 | 0.000322 |
| Malignant epithelial | C5 Am/CEA-like ECs | NDFIP2 | 1.703 | 0.598 | 0.304 | 1.17e-31 | 3.94e-27 |

| compartment | subcluster | gene | avg_log2FC | pct.1 | pct.2 | p_val | p_val_adj |
|---|---|---|---|---|---|---|---|
| Malignant epithelial | C5 Am/CEA-like ECs | STX3 | 1.701 | 0.444 | 0.227 | 2.12e-20 | 7.15e-16 |
| Malignant epithelial | C5 Am/CEA-like ECs | PDE4C | 1.7 | 0.487 | 0.284 | 1.51e-19 | 5.1e-15 |
| Malignant epithelial | C5 Am/CEA-like ECs | MOSPD1 | 1.698 | 0.291 | 0.155 | 1.28e-10 | 4.31e-06 |
| Malignant epithelial | C5 Am/CEA-like ECs | ENTPD2 | 1.697 | 0.268 | 0.112 | 3.49e-15 | 1.18e-10 |
| Malignant epithelial | C5 Am/CEA-like ECs | ATP13A2 | 1.689 | 0.176 | 0.072 | 3.68e-10 | 1.24e-05 |
| Malignant epithelial | C5 Am/CEA-like ECs | TMEM65 | 1.685 | 0.161 | 0.076 | 3.43e-07 | 0.0115 |
| Malignant epithelial | C5 Am/CEA-like ECs | TOR4A | 1.684 | 0.264 | 0.123 | 2.14e-12 | 7.22e-08 |
| Malignant epithelial | C5 Am/CEA-like ECs | EPS8L3 | 1.683 | 0.387 | 0.183 | 1.71e-18 | 5.77e-14 |
| Malignant epithelial | C5 Am/CEA-like ECs | GDPD3 | 1.683 | 0.218 | 0.085 | 1.05e-13 | 3.54e-09 |
| Malignant epithelial | C5 Am/CEA-like ECs | C1orf106 | 1.682 | 0.529 | 0.273 | 1.25e-25 | 4.21e-21 |
| Malignant epithelial | C5 Am/CEA-like ECs | CYP51A1 | 1.681 | 0.103 | 0.035 | 1.25e-08 | 0.000421 |
| Malignant epithelial | C5 Am/CEA-like ECs | BIN2 | 1.681 | 0.111 | 0.043 | 2.8e-07 | 0.00945 |
| Malignant epithelial | C5 Am/CEA-like ECs | FNIP2 | 1.673 | 0.234 | 0.094 | 3.58e-14 | 1.21e-09 |
| Malignant epithelial | C5 Am/CEA-like ECs | PLA2G10 | 1.672 | 0.261 | 0.101 | 1.01e-16 | 3.41e-12 |
| Malignant epithelial | C5 Am/CEA-like ECs | SLC31A1 | 1.672 | 0.241 | 0.126 | 7.28e-09 | 0.000245 |
| Malignant epithelial | C5 Am/CEA-like ECs | CD24 | 1.669 | 0.885 | 0.466 | 6.94e-58 | 2.34e-53 |
| Malignant epithelial | C5 Am/CEA-like ECs | GABARAPL1 | 1.669 | 0.295 | 0.134 | 4.78e-14 | 1.61e-09 |
| Malignant epithelial | C5 Am/CEA-like ECs | TMF1 | 1.665 | 0.372 | 0.203 | 1.47e-13 | 4.95e-09 |
| Malignant epithelial | C5 Am/CEA-like ECs | CDC42EP5 | 1.661 | 0.341 | 0.154 | 1.53e-17 | 5.17e-13 |
| Malignant epithelial | C5 Am/CEA-like ECs | EFNB1 | 1.658 | 0.441 | 0.213 | 4.73e-22 | 1.59e-17 |
| Malignant epithelial | C5 Am/CEA-like ECs | RBPMS | 1.657 | 0.211 | 0.081 | 4.84e-13 | 1.63e-08 |
| Malignant epithelial | C5 Am/CEA-like ECs | EPB41L4B | 1.655 | 0.192 | 0.09 | 1.36e-08 | 0.000458 |
| Malignant epithelial | C5 Am/CEA-like ECs | ARRDC4 | 1.654 | 0.226 | 0.097 | 4.19e-12 | 1.41e-07 |
| Malignant epithelial | C5 Am/CEA-like ECs | CCPG1 | 1.653 | 0.28 | 0.131 | 6.53e-13 | 2.2e-08 |
| Malignant epithelial | C5 Am/CEA-like ECs | MIR194-2HG | 1.651 | 0.142 | 0.054 | 2.04e-09 | 6.87e-05 |
| Malignant epithelial | C5 Am/CEA-like ECs | ZNF562 | 1.648 | 0.188 | 0.093 | 1.52e-07 | 0.00513 |
| Malignant epithelial | C5 Am/CEA-like ECs | MPZL3 | 1.644 | 0.184 | 0.093 | 3.64e-07 | 0.0123 |
| Malignant epithelial | C5 Am/CEA-like ECs | IHH | 1.643 | 0.111 | 0.039 | 1.97e-08 | 0.000665 |
| Malignant epithelial | C5 Am/CEA-like ECs | USP12 | 1.642 | 0.326 | 0.142 | 1.19e-17 | 4.02e-13 |
| Malignant epithelial | C5 Am/CEA-like ECs | DUSP8 | 1.642 | 0.211 | 0.094 | 2.63e-10 | 8.88e-06 |
| Malignant epithelial | C5 Am/CEA-like ECs | TRAC | 1.64 | 0.195 | 0.104 | 7.64e-07 | 0.0257 |
| Malignant epithelial | C5 Am/CEA-like ECs | MAL2 | 1.634 | 0.759 | 0.599 | 9.55e-30 | 3.22e-25 |
| Malignant epithelial | C5 Am/CEA-like ECs | SHROOM3 | 1.634 | 0.218 | 0.102 | 7.17e-10 | 2.42e-05 |
| Malignant epithelial | C5 Am/CEA-like ECs | MYO15B | 1.628 | 0.307 | 0.14 | 2.32e-14 | 7.82e-10 |
| Malignant epithelial | C5 Am/CEA-like ECs | C10orf54 | 1.626 | 0.425 | 0.193 | 6.22e-23 | 2.1e-18 |
| Malignant epithelial | C5 Am/CEA-like ECs | DAB2IP | 1.625 | 0.188 | 0.093 | 1.03e-07 | 0.00345 |
| Malignant epithelial | C5 Am/CEA-like ECs | FOXA3 | 1.623 | 0.264 | 0.116 | 1.01e-13 | 3.42e-09 |
| Malignant epithelial | C5 Am/CEA-like ECs | CLDN12 | 1.621 | 0.406 | 0.191 | 1.88e-20 | 6.34e-16 |
| Malignant epithelial | C5 Am/CEA-like ECs | PNPLA8 | 1.614 | 0.318 | 0.135 | 8.5e-18 | 2.86e-13 |
| Malignant epithelial | C5 Am/CEA-like ECs | GCNT1 | 1.614 | 0.138 | 0.057 | 7.93e-08 | 0.00267 |
| Malignant epithelial | C5 Am/CEA-like ECs | DST | 1.612 | 0.632 | 0.366 | 1.49e-28 | 5.03e-24 |
| Malignant epithelial | C5 Am/CEA-like ECs | AC009133.12 | 1.607 | 0.253 | 0.113 | 2.27e-12 | 7.64e-08 |
| Malignant epithelial | C5 Am/CEA-like ECs | VDR | 1.605 | 0.188 | 0.082 | 9.98e-10 | 3.36e-05 |
| Malignant epithelial | C5 Am/CEA-like ECs | ST3GAL1 | 1.604 | 0.276 | 0.134 | 6.95e-12 | 2.34e-07 |
| Malignant epithelial | C5 Am/CEA-like ECs | AOC1 | 1.603 | 0.521 | 0.322 | 2.4e-17 | 8.08e-13 |
| Malignant epithelial | C5 Am/CEA-like ECs | GALC | 1.603 | 0.165 | 0.065 | 3.84e-10 | 1.29e-05 |
| Malignant epithelial | C5 Am/CEA-like ECs | CD164 | 1.6 | 0.92 | 0.73 | 9.62e-49 | 3.24e-44 |
| Malignant epithelial | C5 Am/CEA-like ECs | SETD9 | 1.592 | 0.352 | 0.177 | 3.37e-15 | 1.13e-10 |
| Malignant epithelial | C5 Am/CEA-like ECs | MUC4 | 1.591 | 0.276 | 0.116 | 3.38e-15 | 1.14e-10 |
| Malignant epithelial | C5 Am/CEA-like ECs | MAP1LC3B | 1.589 | 0.854 | 0.626 | 1.72e-44 | 5.81e-40 |
| Malignant epithelial | C5 Am/CEA-like ECs | SLC12A7 | 1.588 | 0.257 | 0.148 | 2.57e-07 | 0.00865 |
| Malignant epithelial | C5 Am/CEA-like ECs | MLLT3 | 1.582 | 0.253 | 0.115 | 7.32e-12 | 2.47e-07 |
| Malignant epithelial | C5 Am/CEA-like ECs | TIMP2 | 1.573 | 0.437 | 0.252 | 1.12e-14 | 3.77e-10 |
| Malignant epithelial | C5 Am/CEA-like ECs | IFNAR2 | 1.57 | 0.303 | 0.135 | 2.91e-15 | 9.81e-11 |
| Malignant epithelial | C5 Am/CEA-like ECs | PRKCZ | 1.57 | 0.241 | 0.141 | 1.02e-06 | 0.0345 |
| Malignant epithelial | C5 Am/CEA-like ECs | IL13RA1 | 1.568 | 0.414 | 0.221 | 1.54e-16 | 5.2e-12 |
| Malignant epithelial | C5 Am/CEA-like ECs | IL1RN | 1.567 | 0.161 | 0.063 | 4.49e-10 | 1.51e-05 |
| Malignant epithelial | C5 Am/CEA-like ECs | TMEM54 | 1.56 | 0.805 | 0.645 | 4.61e-37 | 1.55e-32 |
| Malignant epithelial | C5 Am/CEA-like ECs | ARFGAP3 | 1.553 | 0.406 | 0.189 | 2.54e-21 | 8.55e-17 |
| Malignant epithelial | C5 Am/CEA-like ECs | SEC23B | 1.55 | 0.264 | 0.133 | 1.91e-10 | 6.44e-06 |
| Malignant epithelial | C5 Am/CEA-like ECs | LGALS4 | 1.549 | 0.874 | 0.48 | 1.46e-53 | 4.92e-49 |
| Malignant epithelial | C5 Am/CEA-like ECs | ERBB2IP | 1.545 | 0.341 | 0.218 | 1.05e-08 | 0.000353 |
| Malignant epithelial | C5 Am/CEA-like ECs | POF1B | 1.54 | 0.318 | 0.143 | 4.62e-16 | 1.56e-11 |
| Malignant epithelial | C5 Am/CEA-like ECs | ACSS2 | 1.536 | 0.36 | 0.168 | 1.2e-16 | 4.05e-12 |

| compartment | subcluster | gene | avg_log2FC | pct.1 | pct.2 | p_val | p_val_adj |
|---|---|---|---|---|---|---|---|
| Malignant epithelial | C5 Am/CEA-like ECs | ENPP4 | 1.536 | 0.126 | 0.053 | 3.88e-07 | 0.0131 |
| Malignant epithelial | C5 Am/CEA-like ECs | GPRC5C | 1.535 | 0.452 | 0.234 | 1.84e-19 | 6.19e-15 |
| Malignant epithelial | C5 Am/CEA-like ECs | TINAGL1 | 1.524 | 0.64 | 0.457 | 1.33e-18 | 4.47e-14 |
| Malignant epithelial | C5 Am/CEA-like ECs | PPARA | 1.524 | 0.176 | 0.083 | 5.82e-08 | 0.00196 |
| Malignant epithelial | C5 Am/CEA-like ECs | PNPLA2 | 1.522 | 0.579 | 0.33 | 5.57e-25 | 1.88e-20 |
| Malignant epithelial | C5 Am/CEA-like ECs | SERINC2 | 1.52 | 0.678 | 0.457 | 4.04e-24 | 1.36e-19 |
| Malignant epithelial | C5 Am/CEA-like ECs | C12orf49 | 1.519 | 0.352 | 0.181 | 1.14e-14 | 3.83e-10 |
| Malignant epithelial | C5 Am/CEA-like ECs | ANG | 1.517 | 0.46 | 0.231 | 1.17e-20 | 3.93e-16 |
| Malignant epithelial | C5 Am/CEA-like ECs | RORA | 1.517 | 0.172 | 0.079 | 5.23e-08 | 0.00176 |
| Malignant epithelial | C5 Am/CEA-like ECs | RHPN2 | 1.515 | 0.318 | 0.166 | 6.38e-12 | 2.15e-07 |
| Malignant epithelial | C5 Am/CEA-like ECs | WBP2 | 1.509 | 0.567 | 0.33 | 1.11e-23 | 3.75e-19 |
| Malignant epithelial | C5 Am/CEA-like ECs | TRIM25 | 1.5 | 0.261 | 0.131 | 4.29e-10 | 1.44e-05 |
| Malignant epithelial | C5 Am/CEA-like ECs | MGAT4A | 1.5 | 0.188 | 0.084 | 3.57e-09 | 0.00012 |
| Malignant epithelial | C5 Am/CEA-like ECs | ST6GALNAC1 | 1.499 | 0.203 | 0.07 | 2.1e-15 | 7.06e-11 |
| Malignant epithelial | C5 Am/CEA-like ECs | AC005550.3 | 1.497 | 0.153 | 0.058 | 3.05e-10 | 1.03e-05 |
| Malignant epithelial | C5 Am/CEA-like ECs | DNAJC3 | 1.496 | 0.441 | 0.275 | 1.31e-12 | 4.42e-08 |
| Malignant epithelial | C5 Am/CEA-like ECs | PEA15 | 1.495 | 0.326 | 0.163 | 2.49e-13 | 8.4e-09 |
| Malignant epithelial | C5 Am/CEA-like ECs | ERO1A | 1.494 | 0.697 | 0.544 | 7.83e-15 | 2.64e-10 |
| Malignant epithelial | C5 Am/CEA-like ECs | PIGH | 1.494 | 0.28 | 0.14 | 5.3e-11 | 1.78e-06 |
| Malignant epithelial | C5 Am/CEA-like ECs | ACE2 | 1.487 | 0.126 | 0.049 | 3.61e-08 | 0.00122 |
| Malignant epithelial | C5 Am/CEA-like ECs | LHFPL2 | 1.486 | 0.165 | 0.078 | 4.03e-07 | 0.0136 |
| Malignant epithelial | C5 Am/CEA-like ECs | EZR | 1.485 | 0.935 | 0.722 | 2.72e-60 | 9.17e-56 |
| Malignant epithelial | C5 Am/CEA-like ECs | TRIM31 | 1.485 | 0.648 | 0.341 | 1.75e-32 | 5.91e-28 |
| Malignant epithelial | C5 Am/CEA-like ECs | FAM122A | 1.483 | 0.234 | 0.113 | 8.44e-10 | 2.84e-05 |
| Malignant epithelial | C5 Am/CEA-like ECs | SH3BGRL3 | 1.482 | 0.962 | 0.882 | 2.32e-69 | 7.81e-65 |
| Malignant epithelial | C5 Am/CEA-like ECs | INSIG1 | 1.482 | 0.494 | 0.259 | 4.550000000000001e-21 | 1.53e-16 |
| Malignant epithelial | C5 Am/CEA-like ECs | TSPAN8 | 1.478 | 0.912 | 0.548 | 9.91e-49 | 3.34e-44 |
| Malignant epithelial | C5 Am/CEA-like ECs | MYO6 | 1.478 | 0.598 | 0.339 | 7.42e-25 | 2.5e-20 |
| Malignant epithelial | C5 Am/CEA-like ECs | SERPINB8 | 1.478 | 0.119 | 0.05 | 1.37e-06 | 0.0461 |
| Malignant epithelial | C5 Am/CEA-like ECs | PGM3 | 1.473 | 0.192 | 0.096 | 1.34e-07 | 0.00452 |
| Malignant epithelial | C5 Am/CEA-like ECs | RDH10 | 1.472 | 0.356 | 0.185 | 5.42e-13 | 1.83e-08 |
| Malignant epithelial | C5 Am/CEA-like ECs | UBR4 | 1.472 | 0.333 | 0.199 | 3.41e-09 | 0.000115 |
| Malignant epithelial | C5 Am/CEA-like ECs | RNASE4 | 1.471 | 0.161 | 0.078 | 1.21e-06 | 0.0409 |
| Malignant epithelial | C5 Am/CEA-like ECs | TRIM15 | 1.468 | 0.234 | 0.111 | 3.63e-10 | 1.22e-05 |
| Malignant epithelial | C5 Am/CEA-like ECs | FRK | 1.468 | 0.184 | 0.089 | 1.06e-07 | 0.00356 |
| Malignant epithelial | C5 Am/CEA-like ECs | DPH3 | 1.467 | 0.479 | 0.247 | 8.960000000000001e-22 | 3.02e-17 |
| Malignant epithelial | C5 Am/CEA-like ECs | LIMA1 | 1.466 | 0.506 | 0.281 | 4.06e-21 | 1.37e-16 |
| Malignant epithelial | C5 Am/CEA-like ECs | NEU1 | 1.463 | 0.69 | 0.446 | 2.86e-24 | 9.65e-20 |
| Malignant epithelial | C5 Am/CEA-like ECs | SEMA3B | 1.463 | 0.398 | 0.19 | 3.35e-18 | 1.13e-13 |
| Malignant epithelial | C5 Am/CEA-like ECs | PDZK1IP1 | 1.462 | 0.709 | 0.524 | 1.93e-22 | 6.51e-18 |
| Malignant epithelial | C5 Am/CEA-like ECs | SH3BGRL2 | 1.461 | 0.284 | 0.135 | 1.39e-12 | 4.69e-08 |
| Malignant epithelial | C5 Am/CEA-like ECs | TFE3 | 1.461 | 0.195 | 0.088 | 2.63e-09 | 8.85e-05 |
| Malignant epithelial | C5 Am/CEA-like ECs | MICAL1 | 1.46 | 0.203 | 0.095 | 8.6e-09 | 0.00029 |
| Malignant epithelial | C5 Am/CEA-like ECs | RAPGEF5 | 1.46 | 0.134 | 0.055 | 1.1e-07 | 0.0037 |
| Malignant epithelial | C5 Am/CEA-like ECs | RP5-1159O4.1 | 1.457 | 0.176 | 0.076 | 5.46e-09 | 0.000184 |
| Malignant epithelial | C5 Am/CEA-like ECs | EFCAB14 | 1.457 | 0.264 | 0.145 | 1.5e-08 | 0.000506 |
| Malignant epithelial | C5 Am/CEA-like ECs | PID1 | 1.456 | 0.138 | 0.06 | 3.03e-07 | 0.0102 |
| Malignant epithelial | C5 Am/CEA-like ECs | GDF15 | 1.455 | 0.648 | 0.378 | 1.22e-22 | 4.12e-18 |
| Malignant epithelial | C5 Am/CEA-like ECs | F11R | 1.452 | 0.563 | 0.34 | 1.3e-20 | 4.39e-16 |
| Malignant epithelial | C5 Am/CEA-like ECs | AAK1 | 1.449 | 0.241 | 0.115 | 1.59e-10 | 5.34e-06 |
| Malignant epithelial | C5 Am/CEA-like ECs | GABRE | 1.448 | 0.307 | 0.156 | 6.61e-12 | 2.23e-07 |
| Malignant epithelial | C5 Am/CEA-like ECs | CRYAB | 1.448 | 0.153 | 0.066 | 3.04e-08 | 0.00103 |
| Malignant epithelial | C5 Am/CEA-like ECs | FAM214B | 1.447 | 0.149 | 0.063 | 3.83e-08 | 0.00129 |
| Malignant epithelial | C5 Am/CEA-like ECs | TMEM125 | 1.444 | 0.287 | 0.129 | 5.97e-14 | 2.01e-09 |
| Malignant epithelial | C5 Am/CEA-like ECs | ABHD17C | 1.444 | 0.337 | 0.204 | 1.55e-09 | 5.23e-05 |
| Malignant epithelial | C5 Am/CEA-like ECs | CYSTM1 | 1.443 | 0.912 | 0.747 | 1.06e-56 | 3.57e-52 |
| Malignant epithelial | C5 Am/CEA-like ECs | TAF13 | 1.443 | 0.149 | 0.071 | 1.05e-06 | 0.0355 |
| Malignant epithelial | C5 Am/CEA-like ECs | P4HA2 | 1.442 | 0.307 | 0.164 | 1.11e-10 | 3.75e-06 |
| Malignant epithelial | C5 Am/CEA-like ECs | CEBPG | 1.44 | 0.575 | 0.348 | 2.6e-21 | 8.77e-17 |
| Malignant epithelial | C5 Am/CEA-like ECs | RRBP1 | 1.437 | 0.816 | 0.64 | 6.8e-38 | 2.29e-33 |
| Malignant epithelial | C5 Am/CEA-like ECs | TMPRSS4 | 1.436 | 0.563 | 0.29 | 1.68e-26 | 5.65e-22 |
| Malignant epithelial | C5 Am/CEA-like ECs | GSKIP | 1.434 | 0.448 | 0.26 | 2.48e-15 | 8.34e-11 |
| Malignant epithelial | C5 Am/CEA-like ECs | TMEM2 | 1.434 | 0.314 | 0.153 | 3.87e-13 | 1.3e-08 |
| Malignant epithelial | C5 Am/CEA-like ECs | SMIM24 | 1.433 | 0.314 | 0.163 | 2.31e-11 | 7.77e-07 |

| compartment | subcluster | gene | avg_log2FC | pct.1 | pct.2 | p_val | p_val_adj |
|---|---|---|---|---|---|---|---|
| Malignant epithelial | C5 Am/CEA-like ECs | TTC39B | 1.432 | 0.192 | 0.087 | 6.16e-09 | 0.000208 |
| Malignant epithelial | C5 Am/CEA-like ECs | PUDP | 1.426 | 0.195 | 0.101 | 3.67e-07 | 0.0124 |
| Malignant epithelial | C5 Am/CEA-like ECs | P2RX4 | 1.425 | 0.299 | 0.166 | 2.51e-09 | 8.46e-05 |
| Malignant epithelial | C5 Am/CEA-like ECs | MYO1B | 1.424 | 0.215 | 0.104 | 7e-09 | 0.000236 |
| Malignant epithelial | C5 Am/CEA-like ECs | DPP4 | 1.422 | 0.261 | 0.152 | 3.72e-07 | 0.0125 |
| Malignant epithelial | C5 Am/CEA-like ECs | TRIM69 | 1.42 | 0.245 | 0.135 | 8.37e-08 | 0.00282 |
| Malignant epithelial | C5 Am/CEA-like ECs | GPR160 | 1.418 | 0.437 | 0.219 | 4.73e-18 | 1.59e-13 |
| Malignant epithelial | C5 Am/CEA-like ECs | B4GALT4 | 1.418 | 0.264 | 0.133 | 5.79e-10 | 1.95e-05 |
| Malignant epithelial | C5 Am/CEA-like ECs | COG3 | 1.413 | 0.284 | 0.148 | 4.6e-10 | 1.55e-05 |
| Malignant epithelial | C5 Am/CEA-like ECs | KRT15 | 1.41 | 0.192 | 0.093 | 4.97e-08 | 0.00168 |
| Malignant epithelial | C5 Am/CEA-like ECs | SESTD1 | 1.407 | 0.203 | 0.102 | 9.64e-08 | 0.00325 |
| Malignant epithelial | C5 Am/CEA-like ECs | PDGFA | 1.406 | 0.36 | 0.182 | 1.44e-14 | 4.86e-10 |
| Malignant epithelial | C5 Am/CEA-like ECs | RHOC | 1.405 | 0.897 | 0.726 | 9.25e-51 | 3.12e-46 |
| Malignant epithelial | C5 Am/CEA-like ECs | TRAM1 | 1.405 | 0.713 | 0.516 | 1.58e-24 | 5.33e-20 |
| Malignant epithelial | C5 Am/CEA-like ECs | CDKN2B | 1.405 | 0.341 | 0.185 | 4.47e-12 | 1.51e-07 |
| Malignant epithelial | C5 Am/CEA-like ECs | TBC1D23 | 1.402 | 0.172 | 0.082 | 2.2e-07 | 0.00742 |
| Malignant epithelial | C5 Am/CEA-like ECs | TBC1D1 | 1.401 | 0.169 | 0.082 | 5.05e-07 | 0.017 |
| Malignant epithelial | C5 Am/CEA-like ECs | ITGA2 | 1.398 | 0.352 | 0.185 | 5.95e-13 | 2e-08 |
| Malignant epithelial | C5 Am/CEA-like ECs | GBP3 | 1.398 | 0.276 | 0.127 | 6.81e-13 | 2.29e-08 |
| Malignant epithelial | C5 Am/CEA-like ECs | ABHD17B | 1.398 | 0.199 | 0.088 | 7.12e-10 | 2.4e-05 |
| Malignant epithelial | C5 Am/CEA-like ECs | HLA-C | 1.395 | 0.973 | 0.946 | 8.319999999999999e-42 | 2.8e-37 |
| Malignant epithelial | C5 Am/CEA-like ECs | NBL1 | 1.394 | 0.701 | 0.453 | 1.05e-30 | 3.53e-26 |
| Malignant epithelial | C5 Am/CEA-like ECs | IFT172 | 1.394 | 0.552 | 0.316 | 1.41e-21 | 4.75e-17 |
| Malignant epithelial | C5 Am/CEA-like ECs | MYH14 | 1.39 | 0.648 | 0.4 | 3.34e-24 | 1.12e-19 |
| Malignant epithelial | C5 Am/CEA-like ECs | SURF4 | 1.39 | 0.628 | 0.421 | 8.210000000000001e-22 | 2.77e-17 |
| Malignant epithelial | C5 Am/CEA-like ECs | MOGAT3 | 1.387 | 0.23 | 0.096 | 2.33e-12 | 7.86e-08 |
| Malignant epithelial | C5 Am/CEA-like ECs | SAMSN1 | 1.387 | 0.153 | 0.071 | 4.21e-07 | 0.0142 |
| Malignant epithelial | C5 Am/CEA-like ECs | NCSTN | 1.385 | 0.28 | 0.15 | 1.57e-09 | 5.3e-05 |
| Malignant epithelial | C5 Am/CEA-like ECs | SMPD3 | 1.382 | 0.13 | 0.057 | 9.13e-07 | 0.0307 |
| Malignant epithelial | C5 Am/CEA-like ECs | SPTBN1 | 1.38 | 0.709 | 0.438 | 2.77e-29 | 9.35e-25 |
| Malignant epithelial | C5 Am/CEA-like ECs | FAM102A | 1.379 | 0.483 | 0.299 | 2.41e-15 | 8.11e-11 |
| Malignant epithelial | C5 Am/CEA-like ECs | RBM47 | 1.378 | 0.379 | 0.214 | 4.05e-12 | 1.37e-07 |
| Malignant epithelial | C5 Am/CEA-like ECs | PRR15L | 1.376 | 0.49 | 0.264 | 2.91e-19 | 9.79e-15 |
| Malignant epithelial | C5 Am/CEA-like ECs | CACFD1 | 1.375 | 0.188 | 0.08 | 8.29e-10 | 2.79e-05 |
| Malignant epithelial | C5 Am/CEA-like ECs | MAPK3 | 1.374 | 0.54 | 0.335 | 1.48e-18 | 4.99e-14 |
| Malignant epithelial | C5 Am/CEA-like ECs | RHOF | 1.371 | 0.556 | 0.318 | 9.22e-22 | 3.11e-17 |
| Malignant epithelial | C5 Am/CEA-like ECs | GEM | 1.371 | 0.261 | 0.151 | 2.13e-07 | 0.00716 |
| Malignant epithelial | C5 Am/CEA-like ECs | RGCC | 1.37 | 0.494 | 0.343 | 9.27e-11 | 3.12e-06 |
| Malignant epithelial | C5 Am/CEA-like ECs | DNM2 | 1.369 | 0.521 | 0.308 | 1.02e-17 | 3.42e-13 |
| Malignant epithelial | C5 Am/CEA-like ECs | CYTH1 | 1.368 | 0.245 | 0.129 | 2.14e-08 | 0.00072 |
| Malignant epithelial | C5 Am/CEA-like ECs | CDX1 | 1.366 | 0.506 | 0.226 | 2.4e-26 | 8.07e-22 |
| Malignant epithelial | C5 Am/CEA-like ECs | CLSTN1 | 1.365 | 0.471 | 0.3 | 3.46e-13 | 1.17e-08 |
| Malignant epithelial | C5 Am/CEA-like ECs | TPP1 | 1.362 | 0.414 | 0.242 | 1.16e-12 | 3.91e-08 |
| Malignant epithelial | C5 Am/CEA-like ECs | HDHD3 | 1.36 | 0.625 | 0.4 | 3.37e-23 | 1.14e-18 |
| Malignant epithelial | C5 Am/CEA-like ECs | CAB39 | 1.36 | 0.375 | 0.213 | 8.36e-13 | 2.82e-08 |
| Malignant epithelial | C5 Am/CEA-like ECs | LINC00926 | 1.359 | 0.203 | 0.094 | 6.53e-09 | 0.00022 |
| Malignant epithelial | C5 Am/CEA-like ECs | RAPGEFL1 | 1.358 | 0.222 | 0.114 | 2.87e-08 | 0.000967 |
| Malignant epithelial | C5 Am/CEA-like ECs | TOX3 | 1.353 | 0.31 | 0.146 | 2.76e-13 | 9.31e-09 |
| Malignant epithelial | C5 Am/CEA-like ECs | DEFB1 | 1.351 | 0.375 | 0.185 | 2.03e-16 | 6.83e-12 |
| Malignant epithelial | C5 Am/CEA-like ECs | GNA11 | 1.351 | 0.268 | 0.163 | 5.61e-07 | 0.0189 |
| Malignant epithelial | C5 Am/CEA-like ECs | APOL1 | 1.349 | 0.433 | 0.32 | 1.27e-06 | 0.0429 |
| Malignant epithelial | C5 Am/CEA-like ECs | C1orf116 | 1.348 | 0.211 | 0.089 | 6.06e-11 | 2.04e-06 |
| Malignant epithelial | C5 Am/CEA-like ECs | SLPI | 1.346 | 0.686 | 0.599 | 1.3e-09 | 4.39e-05 |
| Malignant epithelial | C5 Am/CEA-like ECs | SUSD6 | 1.344 | 0.218 | 0.104 | 2.35e-09 | 7.92e-05 |
| Malignant epithelial | C5 Am/CEA-like ECs | UBL3 | 1.343 | 0.368 | 0.214 | 5.42e-11 | 1.83e-06 |
| Malignant epithelial | C5 Am/CEA-like ECs | ATP6V0D1 | 1.334 | 0.602 | 0.356 | 6.24e-24 | 2.1e-19 |
| Malignant epithelial | C5 Am/CEA-like ECs | B3GALT4 | 1.333 | 0.234 | 0.112 | 4.41e-10 | 1.49e-05 |
| Malignant epithelial | C5 Am/CEA-like ECs | PRR15 | 1.331 | 0.475 | 0.268 | 6.5e-16 | 2.19e-11 |
| Malignant epithelial | C5 Am/CEA-like ECs | ATP1B1 | 1.331 | 0.762 | 0.632 | 9.48e-16 | 3.19e-11 |
| Malignant epithelial | C5 Am/CEA-like ECs | XIAP | 1.331 | 0.437 | 0.247 | 2.99e-14 | 1.01e-09 |
| Malignant epithelial | C5 Am/CEA-like ECs | SLC10A3 | 1.331 | 0.303 | 0.171 | 1.71e-09 | 5.78e-05 |
| Malignant epithelial | C5 Am/CEA-like ECs | LRP10 | 1.326 | 0.674 | 0.475 | 3.38e-24 | 1.14e-19 |
| Malignant epithelial | C5 Am/CEA-like ECs | TUFT1 | 1.323 | 0.314 | 0.177 | 8.73e-10 | 2.94e-05 |
| Malignant epithelial | C5 Am/CEA-like ECs | FBLIM1 | 1.315 | 0.192 | 0.099 | 1.08e-06 | 0.0365 |

| compartment | subcluster | gene | avg_log2FC | pct.1 | pct.2 | p_val | p_val_adj |
|---|---|---|---|---|---|---|---|
| Malignant epithelial | C5 Am/CEA-like ECs | CLDN3 | 1.311 | 0.962 | 0.908 | 5.22e-43 | 1.76e-38 |
| Malignant epithelial | C5 Am/CEA-like ECs | TAX1BP3 | 1.31 | 0.697 | 0.468 | 1.75e-25 | 5.89e-21 |
| Malignant epithelial | C5 Am/CEA-like ECs | ZYX | 1.309 | 0.464 | 0.278 | 1.48e-14 | 5e-10 |
| Malignant epithelial | C5 Am/CEA-like ECs | TNFRSF12A | 1.308 | 0.77 | 0.625 | 7.18e-20 | 2.42e-15 |
| Malignant epithelial | C5 Am/CEA-like ECs | STX12 | 1.305 | 0.23 | 0.111 | 1.26e-09 | 4.25e-05 |
| Malignant epithelial | C5 Am/CEA-like ECs | CLTB | 1.302 | 0.931 | 0.697 | 6.339999999999999e-55 | 2.14e-50 |
| Malignant epithelial | C5 Am/CEA-like ECs | PLPP5 | 1.3 | 0.548 | 0.351 | 9.55e-18 | 3.22e-13 |
| Malignant epithelial | C5 Am/CEA-like ECs | ARL8B | 1.299 | 0.337 | 0.18 | 1.11e-11 | 3.75e-07 |
| Malignant epithelial | C5 Am/CEA-like ECs | TDP2 | 1.299 | 0.226 | 0.129 | 1.3e-06 | 0.0437 |
| Malignant epithelial | C5 Am/CEA-like ECs | TPMT | 1.298 | 0.33 | 0.16 | 3.75e-14 | 1.26e-09 |
| Malignant epithelial | C5 Am/CEA-like ECs | LCOR | 1.298 | 0.295 | 0.163 | 2.44e-09 | 8.23e-05 |
| Malignant epithelial | C5 Am/CEA-like ECs | RRAGC | 1.297 | 0.222 | 0.115 | 5.89e-08 | 0.00198 |
| Malignant epithelial | C5 Am/CEA-like ECs | DGKH | 1.292 | 0.253 | 0.12 | 1.22e-10 | 4.11e-06 |
| Malignant epithelial | C5 Am/CEA-like ECs | GDI1 | 1.29 | 0.46 | 0.275 | 4.02e-13 | 1.36e-08 |
| Malignant epithelial | C5 Am/CEA-like ECs | CLDN7 | 1.287 | 0.916 | 0.782 | 1.28e-43 | 4.32e-39 |
| Malignant epithelial | C5 Am/CEA-like ECs | GSDMB | 1.287 | 0.241 | 0.123 | 9.63e-09 | 0.000324 |
| Malignant epithelial | C5 Am/CEA-like ECs | TJP3 | 1.285 | 0.437 | 0.26 | 4.81e-13 | 1.62e-08 |
| Malignant epithelial | C5 Am/CEA-like ECs | LAMC2 | 1.285 | 0.54 | 0.364 | 1.86e-11 | 6.27e-07 |
| Malignant epithelial | C5 Am/CEA-like ECs | KDM7A | 1.285 | 0.211 | 0.106 | 7.9e-08 | 0.00266 |
| Malignant epithelial | C5 Am/CEA-like ECs | CCNDBP1 | 1.284 | 0.406 | 0.251 | 1.86e-10 | 6.27e-06 |
| Malignant epithelial | C5 Am/CEA-like ECs | GNAQ | 1.284 | 0.383 | 0.248 | 3.14e-09 | 0.000106 |
| Malignant epithelial | C5 Am/CEA-like ECs | SMIM14 | 1.283 | 0.402 | 0.244 | 5.6e-11 | 1.89e-06 |
| Malignant epithelial | C5 Am/CEA-like ECs | DNAJC5 | 1.281 | 0.375 | 0.207 | 1.64e-12 | 5.54e-08 |
| Malignant epithelial | C5 Am/CEA-like ECs | CREB3L2 | 1.276 | 0.28 | 0.146 | 1.01e-09 | 3.42e-05 |
| Malignant epithelial | C5 Am/CEA-like ECs | S100A14 | 1.275 | 0.632 | 0.354 | 8.99e-25 | 3.03e-20 |
| Malignant epithelial | C5 Am/CEA-like ECs | APAF1 | 1.273 | 0.142 | 0.058 | 4.52e-08 | 0.00152 |
| Malignant epithelial | C5 Am/CEA-like ECs | TACC1 | 1.272 | 0.337 | 0.194 | 4.92e-10 | 1.66e-05 |
| Malignant epithelial | C5 Am/CEA-like ECs | PLEKHA6 | 1.272 | 0.192 | 0.095 | 1.86e-07 | 0.00625 |
| Malignant epithelial | C5 Am/CEA-like ECs | TMEM120A | 1.271 | 0.398 | 0.221 | 9.7e-14 | 3.27e-09 |
| Malignant epithelial | C5 Am/CEA-like ECs | WBSCR27 | 1.268 | 0.314 | 0.192 | 1.1e-07 | 0.00372 |
| Malignant epithelial | C5 Am/CEA-like ECs | BAIAP2L2 | 1.268 | 0.238 | 0.137 | 1.32e-06 | 0.0445 |
| Malignant epithelial | C5 Am/CEA-like ECs | SCARB2 | 1.267 | 0.375 | 0.213 | 5.44e-12 | 1.83e-07 |
| Malignant epithelial | C5 Am/CEA-like ECs | PTPN3 | 1.267 | 0.234 | 0.127 | 1.97e-07 | 0.00663 |
| Malignant epithelial | C5 Am/CEA-like ECs | UFM1 | 1.265 | 0.701 | 0.502 | 4.96e-22 | 1.67e-17 |
| Malignant epithelial | C5 Am/CEA-like ECs | WASL | 1.264 | 0.406 | 0.251 | 1.02e-11 | 3.44e-07 |
| Malignant epithelial | C5 Am/CEA-like ECs | TMEM127 | 1.264 | 0.291 | 0.155 | 4.86e-10 | 1.64e-05 |
| Malignant epithelial | C5 Am/CEA-like ECs | CREB3 | 1.264 | 0.395 | 0.244 | 5.14e-10 | 1.73e-05 |
| Malignant epithelial | C5 Am/CEA-like ECs | LGALS3 | 1.256 | 0.969 | 0.97 | 1.29e-30 | 4.34e-26 |
| Malignant epithelial | C5 Am/CEA-like ECs | PLAGL2 | 1.252 | 0.18 | 0.091 | 8.4e-07 | 0.0283 |
| Malignant epithelial | C5 Am/CEA-like ECs | HPGD | 1.25 | 0.395 | 0.168 | 2.46e-21 | 8.29e-17 |
| Malignant epithelial | C5 Am/CEA-like ECs | GALNT3 | 1.248 | 0.456 | 0.26 | 1.64e-15 | 5.52e-11 |
| Malignant epithelial | C5 Am/CEA-like ECs | BAIAP2L1 | 1.245 | 0.613 | 0.418 | 9.39e-18 | 3.16e-13 |
| Malignant epithelial | C5 Am/CEA-like ECs | INSR | 1.242 | 0.345 | 0.209 | 1.2e-08 | 0.000404 |
| Malignant epithelial | C5 Am/CEA-like ECs | PIP4K2C | 1.242 | 0.203 | 0.109 | 1.48e-06 | 0.0499 |
| Malignant epithelial | C5 Am/CEA-like ECs | GOLM1 | 1.24 | 0.479 | 0.363 | 1.07e-08 | 0.000359 |
| Malignant epithelial | C5 Am/CEA-like ECs | RASSF7 | 1.238 | 0.621 | 0.399 | 5.110000000000001e-21 | 1.72e-16 |
| Malignant epithelial | C5 Am/CEA-like ECs | HLA-B | 1.235 | 0.935 | 0.915 | 4.93e-32 | 1.66e-27 |
| Malignant epithelial | C5 Am/CEA-like ECs | KRAS | 1.235 | 0.433 | 0.271 | 6.29e-12 | 2.12e-07 |
| Malignant epithelial | C5 Am/CEA-like ECs | RASSF6 | 1.234 | 0.199 | 0.104 | 1.04e-06 | 0.0351 |
| Malignant epithelial | C5 Am/CEA-like ECs | CDX2 | 1.233 | 0.349 | 0.178 | 9.12e-13 | 3.07e-08 |
| Malignant epithelial | C5 Am/CEA-like ECs | BRI3 | 1.232 | 0.816 | 0.613 | 5.05e-33 | 1.7e-28 |
| Malignant epithelial | C5 Am/CEA-like ECs | LAMP2 | 1.228 | 0.513 | 0.316 | 2.8e-16 | 9.44e-12 |
| Malignant epithelial | C5 Am/CEA-like ECs | SLC22A18 | 1.228 | 0.609 | 0.451 | 4.17e-16 | 1.4e-11 |
| Malignant epithelial | C5 Am/CEA-like ECs | OTUD7B | 1.227 | 0.207 | 0.111 | 9.72e-07 | 0.0327 |
| Malignant epithelial | C5 Am/CEA-like ECs | VEGFA | 1.225 | 0.548 | 0.341 | 9.11e-16 | 3.07e-11 |
| Malignant epithelial | C5 Am/CEA-like ECs | ELF1 | 1.223 | 0.452 | 0.274 | 3.98e-12 | 1.34e-07 |
| Malignant epithelial | C5 Am/CEA-like ECs | LRCH4 | 1.222 | 0.303 | 0.168 | 2.27e-09 | 7.65e-05 |
| Malignant epithelial | C5 Am/CEA-like ECs | FZR1 | 1.222 | 0.241 | 0.133 | 1.28e-07 | 0.00433 |
| Malignant epithelial | C5 Am/CEA-like ECs | PHF1 | 1.221 | 0.322 | 0.183 | 1.07e-09 | 3.6e-05 |
| Malignant epithelial | C5 Am/CEA-like ECs | SIRT6 | 1.221 | 0.303 | 0.185 | 2.27e-07 | 0.00765 |
| Malignant epithelial | C5 Am/CEA-like ECs | MPC1 | 1.218 | 0.452 | 0.298 | 1.02e-10 | 3.43e-06 |
| Malignant epithelial | C5 Am/CEA-like ECs | PPP1R11 | 1.215 | 0.479 | 0.313 | 2.54e-12 | 8.55e-08 |
| Malignant epithelial | C5 Am/CEA-like ECs | TPRG1L | 1.215 | 0.218 | 0.121 | 1.23e-06 | 0.0415 |
| Malignant epithelial | C5 Am/CEA-like ECs | SPCS3 | 1.213 | 0.59 | 0.38 | 3.68e-18 | 1.24e-13 |

| compartment | subcluster | gene | avg_log2FC | pct.1 | pct.2 | p_val | p_val_adj |
|---|---|---|---|---|---|---|---|
| Malignant epithelial | C5 Am/CEA-like ECs | PIGR | 1.211 | 0.521 | 0.205 | 1.12e-33 | 3.78e-29 |
| Malignant epithelial | C5 Am/CEA-like ECs | RAB32 | 1.211 | 0.345 | 0.218 | 4.99e-08 | 0.00168 |
| Malignant epithelial | C5 Am/CEA-like ECs | BCL10 | 1.209 | 0.326 | 0.19 | 3.17e-09 | 0.000107 |
| Malignant epithelial | C5 Am/CEA-like ECs | CDH1 | 1.206 | 0.728 | 0.457 | 5.28e-30 | 1.78e-25 |
| Malignant epithelial | C5 Am/CEA-like ECs | SMCO4 | 1.205 | 0.295 | 0.165 | 4.21e-09 | 0.000142 |
| Malignant epithelial | C5 Am/CEA-like ECs | BCL2L1 | 1.202 | 0.644 | 0.465 | 2.78e-18 | 9.38e-14 |
| Malignant epithelial | C5 Am/CEA-like ECs | PITPNM1 | 1.202 | 0.249 | 0.13 | 1.1e-08 | 0.00037 |
| Malignant epithelial | C5 Am/CEA-like ECs | DGAT1 | 1.201 | 0.387 | 0.253 | 9.36e-09 | 0.000315 |
| Malignant epithelial | C5 Am/CEA-like ECs | ERP44 | 1.2 | 0.605 | 0.429 | 9.92e-16 | 3.34e-11 |
| Malignant epithelial | C5 Am/CEA-like ECs | MGAT1 | 1.199 | 0.46 | 0.28 | 7.58e-13 | 2.55e-08 |
| Malignant epithelial | C5 Am/CEA-like ECs | FAM3A | 1.195 | 0.441 | 0.283 | 2.02e-10 | 6.79e-06 |
| Malignant epithelial | C5 Am/CEA-like ECs | YKT6 | 1.193 | 0.41 | 0.232 | 3.84e-13 | 1.29e-08 |
| Malignant epithelial | C5 Am/CEA-like ECs | TMEM63B | 1.193 | 0.264 | 0.144 | 4.44e-08 | 0.0015 |
| Malignant epithelial | C5 Am/CEA-like ECs | TES | 1.192 | 0.598 | 0.408 | 1.72e-17 | 5.79e-13 |
| Malignant epithelial | C5 Am/CEA-like ECs | CGN | 1.192 | 0.28 | 0.147 | 2.61e-09 | 8.79e-05 |
| Malignant epithelial | C5 Am/CEA-like ECs | SMIM6 | 1.19 | 0.284 | 0.145 | 5.37e-10 | 1.81e-05 |
| Malignant epithelial | C5 Am/CEA-like ECs | RAP1B | 1.189 | 0.559 | 0.324 | 8.77e-20 | 2.95e-15 |
| Malignant epithelial | C5 Am/CEA-like ECs | DDIT3 | 1.188 | 0.651 | 0.489 | 9.92e-13 | 3.34e-08 |
| Malignant epithelial | C5 Am/CEA-like ECs | PPARD | 1.186 | 0.161 | 0.065 | 2.27e-09 | 7.64e-05 |
| Malignant epithelial | C5 Am/CEA-like ECs | OSTF1 | 1.184 | 0.609 | 0.406 | 1.29e-18 | 4.36e-14 |
| Malignant epithelial | C5 Am/CEA-like ECs | CTTNBP2NL | 1.184 | 0.195 | 0.096 | 1.47e-07 | 0.00496 |
| Malignant epithelial | C5 Am/CEA-like ECs | OTUD5 | 1.183 | 0.307 | 0.17 | 1.65e-09 | 5.58e-05 |
| Malignant epithelial | C5 Am/CEA-like ECs | UBE2J1 | 1.182 | 0.421 | 0.282 | 2.24e-09 | 7.54e-05 |
| Malignant epithelial | C5 Am/CEA-like ECs | PRSS23 | 1.182 | 0.379 | 0.252 | 8.21e-08 | 0.00277 |
| Malignant epithelial | C5 Am/CEA-like ECs | GSN | 1.181 | 0.805 | 0.65 | 4.939999999999999e-23 | 1.66e-18 |
| Malignant epithelial | C5 Am/CEA-like ECs | TJP2 | 1.179 | 0.513 | 0.339 | 4.88e-12 | 1.64e-07 |
| Malignant epithelial | C5 Am/CEA-like ECs | ABHD3 | 1.178 | 0.291 | 0.178 | 8.04e-07 | 0.0271 |
| Malignant epithelial | C5 Am/CEA-like ECs | KCNK6 | 1.177 | 0.406 | 0.214 | 1.25e-14 | 4.21e-10 |
| Malignant epithelial | C5 Am/CEA-like ECs | MYD88 | 1.177 | 0.333 | 0.204 | 1.76e-08 | 0.000594 |
| Malignant epithelial | C5 Am/CEA-like ECs | AMN | 1.176 | 0.571 | 0.35 | 8.33e-18 | 2.81e-13 |
| Malignant epithelial | C5 Am/CEA-like ECs | CTGF | 1.171 | 0.31 | 0.176 | 8.74e-09 | 0.000294 |
| Malignant epithelial | C5 Am/CEA-like ECs | PPIC | 1.17 | 0.525 | 0.386 | 3.75e-10 | 1.26e-05 |
| Malignant epithelial | C5 Am/CEA-like ECs | PCDH1 | 1.17 | 0.406 | 0.251 | 5.58e-10 | 1.88e-05 |
| Malignant epithelial | C5 Am/CEA-like ECs | IDS | 1.168 | 0.475 | 0.3 | 1.85e-11 | 6.24e-07 |
| Malignant epithelial | C5 Am/CEA-like ECs | RAB11FIP1 | 1.166 | 0.759 | 0.564 | 3.16e-22 | 1.06e-17 |
| Malignant epithelial | C5 Am/CEA-like ECs | GPCPD1 | 1.163 | 0.307 | 0.186 | 1.92e-07 | 0.00648 |
| Malignant epithelial | C5 Am/CEA-like ECs | FNDC3B | 1.158 | 0.368 | 0.23 | 6.4e-09 | 0.000216 |
| Malignant epithelial | C5 Am/CEA-like ECs | BATF2 | 1.158 | 0.249 | 0.136 | 1.27e-07 | 0.00428 |
| Malignant epithelial | C5 Am/CEA-like ECs | MEF2D | 1.157 | 0.257 | 0.132 | 5.39e-09 | 0.000182 |
| Malignant epithelial | C5 Am/CEA-like ECs | TPD52 | 1.155 | 0.686 | 0.509 | 6.29e-19 | 2.12e-14 |
| Malignant epithelial | C5 Am/CEA-like ECs | HEXB | 1.155 | 0.525 | 0.37 | 6.34e-11 | 2.14e-06 |
| Malignant epithelial | C5 Am/CEA-like ECs | YIPF6 | 1.152 | 0.525 | 0.359 | 1.4e-13 | 4.73e-09 |
| Malignant epithelial | C5 Am/CEA-like ECs | SAMD9 | 1.152 | 0.295 | 0.166 | 2.03e-08 | 0.000685 |
| Malignant epithelial | C5 Am/CEA-like ECs | B2M | 1.149 | 1.0 | 1.0 | 4.77e-39 | 1.61e-34 |
| Malignant epithelial | C5 Am/CEA-like ECs | GUCD1 | 1.149 | 0.356 | 0.223 | 1.93e-08 | 0.00065 |
| Malignant epithelial | C5 Am/CEA-like ECs | GTF2B | 1.148 | 0.287 | 0.164 | 6.97e-08 | 0.00235 |
| Malignant epithelial | C5 Am/CEA-like ECs | CDKN1A | 1.147 | 0.686 | 0.435 | 3.05e-23 | 1.03e-18 |
| Malignant epithelial | C5 Am/CEA-like ECs | ST14 | 1.146 | 0.716 | 0.541 | 1.16e-18 | 3.92e-14 |
| Malignant epithelial | C5 Am/CEA-like ECs | ATP2A2 | 1.145 | 0.51 | 0.318 | 2.73e-14 | 9.21e-10 |
| Malignant epithelial | C5 Am/CEA-like ECs | CMPK1 | 1.141 | 0.67 | 0.446 | 6.5e-22 | 2.19e-17 |
| Malignant epithelial | C5 Am/CEA-like ECs | DAAM1 | 1.14 | 0.356 | 0.222 | 3.15e-08 | 0.00106 |
| Malignant epithelial | C5 Am/CEA-like ECs | PDLIM5 | 1.137 | 0.483 | 0.324 | 9.66e-11 | 3.25e-06 |
| Malignant epithelial | C5 Am/CEA-like ECs | PPP1R14D | 1.136 | 0.567 | 0.36 | 6.2e-17 | 2.09e-12 |
| Malignant epithelial | C5 Am/CEA-like ECs | LACTB2 | 1.136 | 0.502 | 0.331 | 2.99e-13 | 1.01e-08 |
| Malignant epithelial | C5 Am/CEA-like ECs | COBL | 1.136 | 0.241 | 0.132 | 2.64e-07 | 0.0089 |
| Malignant epithelial | C5 Am/CEA-like ECs | TXNIP | 1.135 | 0.778 | 0.624 | 5.23e-19 | 1.76e-14 |
| Malignant epithelial | C5 Am/CEA-like ECs | PDIA4 | 1.135 | 0.648 | 0.567 | 1.86e-08 | 0.000626 |
| Malignant epithelial | C5 Am/CEA-like ECs | MARVELD3 | 1.13 | 0.41 | 0.291 | 3.87e-08 | 0.0013 |
| Malignant epithelial | C5 Am/CEA-like ECs | TMEM59 | 1.129 | 0.87 | 0.677 | 9.999999999999999e-33 | 3.38e-28 |
| Malignant epithelial | C5 Am/CEA-like ECs | TNFRSF14 | 1.128 | 0.429 | 0.29 | 5.52e-09 | 0.000186 |
| Malignant epithelial | C5 Am/CEA-like ECs | VIMP | 1.127 | 0.774 | 0.621 | 1.29e-21 | 4.36e-17 |
| Malignant epithelial | C5 Am/CEA-like ECs | DUSP10 | 1.126 | 0.253 | 0.137 | 3.07e-08 | 0.00104 |
| Malignant epithelial | C5 Am/CEA-like ECs | GRN | 1.124 | 0.755 | 0.609 | 3.39e-18 | 1.14e-13 |
| Malignant epithelial | C5 Am/CEA-like ECs | PSMD10 | 1.124 | 0.253 | 0.147 | 7.68e-07 | 0.0259 |

| compartment | subcluster | gene | avg_log2FC | pct.1 | pct.2 | p_val | p_val_adj |
|---|---|---|---|---|---|---|---|
| Malignant epithelial | C5 Am/CEA-like ECs | OSR2 | 1.123 | 0.307 | 0.185 | 9.23e-08 | 0.00311 |
| Malignant epithelial | C5 Am/CEA-like ECs | PINK1 | 1.122 | 0.31 | 0.19 | 4.19e-08 | 0.00141 |
| Malignant epithelial | C5 Am/CEA-like ECs | NEAT1 | 1.121 | 0.973 | 0.959 | 7.46e-39 | 2.51e-34 |
| Malignant epithelial | C5 Am/CEA-like ECs | MISP | 1.121 | 0.728 | 0.592 | 3.63e-19 | 1.22e-14 |
| Malignant epithelial | C5 Am/CEA-like ECs | NEBL | 1.12 | 0.222 | 0.114 | 1.77e-07 | 0.00596 |
| Malignant epithelial | C5 Am/CEA-like ECs | LINC00969 | 1.118 | 0.475 | 0.318 | 3.85e-10 | 1.3e-05 |
| Malignant epithelial | C5 Am/CEA-like ECs | PPM1A | 1.118 | 0.307 | 0.198 | 9.94e-07 | 0.0335 |
| Malignant epithelial | C5 Am/CEA-like ECs | SRI | 1.116 | 0.858 | 0.796 | 1.79e-22 | 6.04e-18 |
| Malignant epithelial | C5 Am/CEA-like ECs | PICALM | 1.116 | 0.379 | 0.252 | 1.23e-07 | 0.00414 |
| Malignant epithelial | C5 Am/CEA-like ECs | CYP2S1 | 1.115 | 0.284 | 0.173 | 1.02e-06 | 0.0344 |
| Malignant epithelial | C5 Am/CEA-like ECs | SEC11C | 1.114 | 0.533 | 0.393 | 2.07e-09 | 6.98e-05 |
| Malignant epithelial | C5 Am/CEA-like ECs | TM2D1 | 1.114 | 0.398 | 0.261 | 2.72e-08 | 0.000915 |
| Malignant epithelial | C5 Am/CEA-like ECs | SEC61A1 | 1.113 | 0.471 | 0.337 | 2.61e-09 | 8.81e-05 |
| Malignant epithelial | C5 Am/CEA-like ECs | JOSD1 | 1.111 | 0.425 | 0.253 | 3.3e-11 | 1.11e-06 |
| Malignant epithelial | C5 Am/CEA-like ECs | MUC5B | 1.111 | 0.195 | 0.103 | 1.31e-06 | 0.0443 |
| Malignant epithelial | C5 Am/CEA-like ECs | IL32 | 1.11 | 0.927 | 0.889 | 9.370000000000001e-27 | 3.16e-22 |
| Malignant epithelial | C5 Am/CEA-like ECs | TMEM106C | 1.109 | 0.533 | 0.321 | 3.9e-17 | 1.32e-12 |
| Malignant epithelial | C5 Am/CEA-like ECs | RAB5A | 1.108 | 0.452 | 0.27 | 5.14e-12 | 1.73e-07 |
| Malignant epithelial | C5 Am/CEA-like ECs | PLEKHA3 | 1.108 | 0.307 | 0.185 | 1.07e-07 | 0.0036 |
| Malignant epithelial | C5 Am/CEA-like ECs | COQ10B | 1.105 | 0.487 | 0.28 | 2.28e-15 | 7.7e-11 |
| Malignant epithelial | C5 Am/CEA-like ECs | MKRN1 | 1.105 | 0.598 | 0.425 | 1.47e-14 | 4.95e-10 |
| Malignant epithelial | C5 Am/CEA-like ECs | NUDT4 | 1.1 | 0.728 | 0.524 | 1.48e-22 | 4.98e-18 |
| Malignant epithelial | C5 Am/CEA-like ECs | PYGB | 1.1 | 0.529 | 0.396 | 4.01e-09 | 0.000135 |
| Malignant epithelial | C5 Am/CEA-like ECs | MTMR11 | 1.099 | 0.41 | 0.254 | 8.61e-11 | 2.9e-06 |
| Malignant epithelial | C5 Am/CEA-like ECs | DAB2 | 1.099 | 0.172 | 0.08 | 2.2e-07 | 0.00742 |
| Malignant epithelial | C5 Am/CEA-like ECs | SPPL2A | 1.098 | 0.318 | 0.181 | 2.82e-09 | 9.49e-05 |
| Malignant epithelial | C5 Am/CEA-like ECs | ASL | 1.093 | 0.51 | 0.371 | 1.16e-10 | 3.91e-06 |
| Malignant epithelial | C5 Am/CEA-like ECs | JAK1 | 1.093 | 0.352 | 0.218 | 6.5e-09 | 0.000219 |
| Malignant epithelial | C5 Am/CEA-like ECs | FHL2 | 1.092 | 0.69 | 0.506 | 3.45e-16 | 1.16e-11 |
| Malignant epithelial | C5 Am/CEA-like ECs | LINC00467 | 1.091 | 0.391 | 0.268 | 3.63e-07 | 0.0122 |
| Malignant epithelial | C5 Am/CEA-like ECs | RAB22A | 1.09 | 0.533 | 0.375 | 2.95e-12 | 9.93e-08 |
| Malignant epithelial | C5 Am/CEA-like ECs | GORASP2 | 1.085 | 0.467 | 0.323 | 2.85e-10 | 9.6e-06 |
| Malignant epithelial | C5 Am/CEA-like ECs | TPM1 | 1.083 | 0.739 | 0.592 | 3.3e-17 | 1.11e-12 |
| Malignant epithelial | C5 Am/CEA-like ECs | BACE2 | 1.083 | 0.579 | 0.391 | 9.26e-14 | 3.12e-09 |
| Malignant epithelial | C5 Am/CEA-like ECs | RPS6KA3 | 1.083 | 0.295 | 0.16 | 6.04e-09 | 0.000203 |
| Malignant epithelial | C5 Am/CEA-like ECs | MAX | 1.083 | 0.506 | 0.385 | 3.62e-08 | 0.00122 |
| Malignant epithelial | C5 Am/CEA-like ECs | LMBRD1 | 1.082 | 0.352 | 0.209 | 1.52e-09 | 5.11e-05 |
| Malignant epithelial | C5 Am/CEA-like ECs | CCDC186 | 1.081 | 0.398 | 0.273 | 7.18e-08 | 0.00242 |
| Malignant epithelial | C5 Am/CEA-like ECs | EPCAM | 1.08 | 0.973 | 0.959 | 1.55e-29 | 5.24e-25 |
| Malignant epithelial | C5 Am/CEA-like ECs | UMAD1 | 1.08 | 0.234 | 0.109 | 6.84e-10 | 2.3e-05 |
| Malignant epithelial | C5 Am/CEA-like ECs | UBE2B | 1.079 | 0.808 | 0.666 | 6.59e-21 | 2.22e-16 |
| Malignant epithelial | C5 Am/CEA-like ECs | RNASET2 | 1.079 | 0.789 | 0.66 | 6.7e-19 | 2.26e-14 |
| Malignant epithelial | C5 Am/CEA-like ECs | PGK1 | 1.079 | 0.862 | 0.771 | 1.7e-15 | 5.74e-11 |
| Malignant epithelial | C5 Am/CEA-like ECs | SH3GLB1 | 1.079 | 0.525 | 0.364 | 2.14e-12 | 7.2e-08 |
| Malignant epithelial | C5 Am/CEA-like ECs | GCNT3 | 1.078 | 0.571 | 0.448 | 3.23e-09 | 0.000109 |
| Malignant epithelial | C5 Am/CEA-like ECs | SDCBP | 1.074 | 0.716 | 0.56 | 4.81e-14 | 1.62e-09 |
| Malignant epithelial | C5 Am/CEA-like ECs | FOSL2 | 1.073 | 0.475 | 0.309 | 1.96e-10 | 6.61e-06 |
| Malignant epithelial | C5 Am/CEA-like ECs | NKIRAS2 | 1.072 | 0.28 | 0.165 | 2.85e-07 | 0.00961 |
| Malignant epithelial | C5 Am/CEA-like ECs | ACBD3 | 1.069 | 0.391 | 0.257 | 7.32e-08 | 0.00247 |
| Malignant epithelial | C5 Am/CEA-like ECs | LIPH | 1.067 | 0.563 | 0.424 | 5.16e-09 | 0.000174 |
| Malignant epithelial | C5 Am/CEA-like ECs | SPINT1 | 1.065 | 0.858 | 0.66 | 3.86e-33 | 1.3e-28 |
| Malignant epithelial | C5 Am/CEA-like ECs | MFGE8 | 1.065 | 0.314 | 0.186 | 2.55e-08 | 0.000859 |
| Malignant epithelial | C5 Am/CEA-like ECs | JMJD6 | 1.065 | 0.356 | 0.223 | 3.26e-08 | 0.0011 |
| Malignant epithelial | C5 Am/CEA-like ECs | LAMP1 | 1.063 | 0.693 | 0.6 | 1.09e-08 | 0.000367 |
| Malignant epithelial | C5 Am/CEA-like ECs | ATP1B3 | 1.061 | 0.659 | 0.545 | 8.86e-10 | 2.99e-05 |
| Malignant epithelial | C5 Am/CEA-like ECs | ADIPOR1 | 1.058 | 0.533 | 0.376 | 2.66e-11 | 8.98e-07 |
| Malignant epithelial | C5 Am/CEA-like ECs | RHEB | 1.056 | 0.904 | 0.735 | 1.36e-32 | 4.59e-28 |
| Malignant epithelial | C5 Am/CEA-like ECs | EPS8 | 1.054 | 0.579 | 0.425 | 2.18e-11 | 7.33e-07 |
| Malignant epithelial | C5 Am/CEA-like ECs | ATP6V0B | 1.053 | 0.705 | 0.545 | 8.5e-16 | 2.86e-11 |
| Malignant epithelial | C5 Am/CEA-like ECs | TM9SF3 | 1.053 | 0.682 | 0.546 | 2.55e-14 | 8.59e-10 |
| Malignant epithelial | C5 Am/CEA-like ECs | CCM2 | 1.053 | 0.345 | 0.224 | 2.58e-07 | 0.00869 |
| Malignant epithelial | C5 Am/CEA-like ECs | FRMD4B | 1.051 | 0.303 | 0.173 | 2.44e-08 | 0.000823 |
| Malignant epithelial | C5 Am/CEA-like ECs | FDX1 | 1.05 | 0.471 | 0.301 | 3.47e-11 | 1.17e-06 |
| Malignant epithelial | C5 Am/CEA-like ECs | RAB18 | 1.05 | 0.364 | 0.238 | 2.92e-07 | 0.00983 |

| compartment | subcluster | gene | avg_log2FC | pct.1 | pct.2 | p_val | p_val_adj |
|---|---|---|---|---|---|---|---|
| Malignant epithelial | C5 Am/CEA-like ECs | CHPF | 1.048 | 0.533 | 0.376 | 2.36e-12 | 7.97e-08 |
| Malignant epithelial | C5 Am/CEA-like ECs | OPTN | 1.048 | 0.563 | 0.409 | 1.48e-11 | 4.98e-07 |
| Malignant epithelial | C5 Am/CEA-like ECs | DEGS2 | 1.048 | 0.261 | 0.126 | 4e-10 | 1.35e-05 |
| Malignant epithelial | C5 Am/CEA-like ECs | ABHD12 | 1.042 | 0.521 | 0.4 | 3.57e-09 | 0.00012 |
| Malignant epithelial | C5 Am/CEA-like ECs | DERL2 | 1.042 | 0.295 | 0.177 | 2.7e-07 | 0.0091 |
| Malignant epithelial | C5 Am/CEA-like ECs | MAGT1 | 1.042 | 0.368 | 0.249 | 4.06e-07 | 0.0137 |
| Malignant epithelial | C5 Am/CEA-like ECs | LSR | 1.041 | 0.851 | 0.766 | 1.92e-23 | 6.49e-19 |
| Malignant epithelial | C5 Am/CEA-like ECs | CAMK2N1 | 1.039 | 0.621 | 0.433 | 6.73e-14 | 2.27e-09 |
| Malignant epithelial | C5 Am/CEA-like ECs | SCAMP2 | 1.037 | 0.391 | 0.256 | 4.11e-08 | 0.00139 |
| Malignant epithelial | C5 Am/CEA-like ECs | PRMT7 | 1.036 | 0.195 | 0.099 | 4.07e-07 | 0.0137 |
| Malignant epithelial | C5 Am/CEA-like ECs | PRELID3B | 1.034 | 0.77 | 0.608 | 3.8e-21 | 1.28e-16 |
| Malignant epithelial | C5 Am/CEA-like ECs | TMEM50B | 1.033 | 0.437 | 0.306 | 2.2e-08 | 0.000742 |
| Malignant epithelial | C5 Am/CEA-like ECs | STK24 | 1.032 | 0.586 | 0.407 | 3.92e-13 | 1.32e-08 |
| Malignant epithelial | C5 Am/CEA-like ECs | SELK | 1.031 | 0.885 | 0.775 | 4.17e-27 | 1.4e-22 |
| Malignant epithelial | C5 Am/CEA-like ECs | SSR4 | 1.029 | 0.946 | 0.921 | 1.16e-36 | 3.91e-32 |
| Malignant epithelial | C5 Am/CEA-like ECs | PPP2CB | 1.029 | 0.475 | 0.335 | 1.04e-09 | 3.52e-05 |
| Malignant epithelial | C5 Am/CEA-like ECs | RYBP | 1.029 | 0.337 | 0.198 | 2.46e-08 | 0.000829 |
| Malignant epithelial | C5 Am/CEA-like ECs | SLC9A3R1 | 1.026 | 0.479 | 0.326 | 1.1e-10 | 3.71e-06 |
| Malignant epithelial | C5 Am/CEA-like ECs | CALU | 1.026 | 0.494 | 0.342 | 3.95e-10 | 1.33e-05 |
| Malignant epithelial | C5 Am/CEA-like ECs | EMC3 | 1.024 | 0.402 | 0.276 | 6.31e-08 | 0.00213 |
| Malignant epithelial | C5 Am/CEA-like ECs | LRP11 | 1.024 | 0.326 | 0.206 | 5.77e-07 | 0.0195 |
| Malignant epithelial | C5 Am/CEA-like ECs | P4HB | 1.023 | 0.862 | 0.772 | 2.86e-23 | 9.64e-19 |
| Malignant epithelial | C5 Am/CEA-like ECs | CLCN3 | 1.022 | 0.352 | 0.198 | 2.8e-10 | 9.44e-06 |
| Malignant epithelial | C5 Am/CEA-like ECs | ARPC1A | 1.021 | 0.743 | 0.611 | 1.37e-18 | 4.63e-14 |
| Malignant epithelial | C5 Am/CEA-like ECs | SERINC1 | 1.021 | 0.406 | 0.258 | 4.3e-09 | 0.000145 |
| Malignant epithelial | C5 Am/CEA-like ECs | OSBPL2 | 1.018 | 0.375 | 0.238 | 1.59e-08 | 0.000536 |
| Malignant epithelial | C5 Am/CEA-like ECs | SQSTM1 | 1.016 | 0.927 | 0.794 | 2.23e-33 | 7.51e-29 |
| Malignant epithelial | C5 Am/CEA-like ECs | ANKRD9 | 1.016 | 0.245 | 0.138 | 4.84e-07 | 0.0163 |
| Malignant epithelial | C5 Am/CEA-like ECs | UGCG | 1.014 | 0.326 | 0.206 | 2.73e-07 | 0.0092 |
| Malignant epithelial | C5 Am/CEA-like ECs | DYNLT1 | 1.011 | 0.839 | 0.686 | 2.52e-27 | 8.5e-23 |
| Malignant epithelial | C5 Am/CEA-like ECs | PARD6B | 1.011 | 0.429 | 0.303 | 1.54e-07 | 0.00518 |
| Malignant epithelial | C5 Am/CEA-like ECs | TNFAIP1 | 1.011 | 0.222 | 0.121 | 6.7e-07 | 0.0226 |
| Malignant epithelial | C5 Am/CEA-like ECs | CHMP2B | 1.009 | 0.644 | 0.519 | 5.39e-12 | 1.82e-07 |
| Malignant epithelial | C5 Am/CEA-like ECs | WDR45B | 1.005 | 0.513 | 0.355 | 1.55e-10 | 5.21e-06 |
| Malignant epithelial | C5 Am/CEA-like ECs | LCN2 | 1.005 | 0.785 | 0.756 | 3.12e-07 | 0.0105 |
| Malignant epithelial | C5 Am/CEA-like ECs | NRBP1 | 1.003 | 0.391 | 0.263 | 4.69e-08 | 0.00158 |
| Malignant epithelial | C5 Am/CEA-like ECs | TIMP1 | 0.999 | 0.935 | 0.929 | 1.16e-22 | 3.91e-18 |
| Malignant epithelial | C5 Am/CEA-like ECs | SHOC2 | 0.999 | 0.349 | 0.221 | 1.05e-07 | 0.00353 |
| Malignant epithelial | C5 Am/CEA-like ECs | EIF4EBP2 | 0.998 | 0.41 | 0.293 | 5.17e-07 | 0.0174 |
| Malignant epithelial | C5 Am/CEA-like ECs | ALDOC | 0.993 | 0.261 | 0.15 | 3.94e-07 | 0.0133 |
| Malignant epithelial | C5 Am/CEA-like ECs | ZBTB43 | 0.991 | 0.498 | 0.345 | 2.02e-10 | 6.8e-06 |
| Malignant epithelial | C5 Am/CEA-like ECs | PLEC | 0.989 | 0.72 | 0.589 | 4.13e-14 | 1.39e-09 |
| Malignant epithelial | C5 Am/CEA-like ECs | WDR1 | 0.989 | 0.506 | 0.352 | 1.3e-09 | 4.38e-05 |
| Malignant epithelial | C5 Am/CEA-like ECs | PAK1 | 0.989 | 0.372 | 0.259 | 9.69e-07 | 0.0327 |
| Malignant epithelial | C5 Am/CEA-like ECs | KDELR3 | 0.988 | 0.368 | 0.245 | 1.78e-07 | 0.00601 |
| Malignant epithelial | C5 Am/CEA-like ECs | ZSWIM8 | 0.985 | 0.184 | 0.091 | 8.7e-07 | 0.0293 |
| Malignant epithelial | C5 Am/CEA-like ECs | SH3KBP1 | 0.983 | 0.544 | 0.45 | 7.22e-08 | 0.00243 |
| Malignant epithelial | C5 Am/CEA-like ECs | SMAP1 | 0.979 | 0.452 | 0.296 | 1.4e-10 | 4.71e-06 |
| Malignant epithelial | C5 Am/CEA-like ECs | DYNLT3 | 0.977 | 0.536 | 0.368 | 1.12e-11 | 3.78e-07 |
| Malignant epithelial | C5 Am/CEA-like ECs | HBEGF | 0.974 | 0.54 | 0.379 | 2.16e-09 | 7.28e-05 |
| Malignant epithelial | C5 Am/CEA-like ECs | RAB5B | 0.974 | 0.318 | 0.186 | 4.7e-08 | 0.00158 |
| Malignant epithelial | C5 Am/CEA-like ECs | MPC2 | 0.969 | 0.667 | 0.516 | 1.61e-13 | 5.44e-09 |
| Malignant epithelial | C5 Am/CEA-like ECs | FOXO3 | 0.969 | 0.479 | 0.319 | 5.12e-10 | 1.73e-05 |
| Malignant epithelial | C5 Am/CEA-like ECs | NAPA | 0.968 | 0.563 | 0.367 | 7.7e-14 | 2.6e-09 |
| Malignant epithelial | C5 Am/CEA-like ECs | SBDS | 0.962 | 0.843 | 0.716 | 7.58e-25 | 2.55e-20 |
| Malignant epithelial | C5 Am/CEA-like ECs | MORF4L2 | 0.962 | 0.736 | 0.62 | 9.4e-16 | 3.17e-11 |
| Malignant epithelial | C5 Am/CEA-like ECs | C1orf122 | 0.961 | 0.406 | 0.271 | 4.21e-08 | 0.00142 |
| Malignant epithelial | C5 Am/CEA-like ECs | GUK1 | 0.959 | 0.962 | 0.894 | 2.36e-40 | 7.94e-36 |
| Malignant epithelial | C5 Am/CEA-like ECs | SLC39A7 | 0.959 | 0.383 | 0.248 | 7.96e-08 | 0.00268 |
| Malignant epithelial | C5 Am/CEA-like ECs | PGRMC1 | 0.957 | 0.678 | 0.527 | 7.37e-14 | 2.48e-09 |
| Malignant epithelial | C5 Am/CEA-like ECs | ATP6V1G1 | 0.955 | 0.908 | 0.835 | 3.56e-34 | 1.2e-29 |
| Malignant epithelial | C5 Am/CEA-like ECs | GHITM | 0.955 | 0.552 | 0.438 | 8.64e-08 | 0.00291 |
| Malignant epithelial | C5 Am/CEA-like ECs | MT1X | 0.954 | 0.172 | 0.408 | 2.99e-13 | 1.01e-08 |
| Malignant epithelial | C5 Am/CEA-like ECs | IGHG3 | 0.954 | 0.536 | 0.494 | 4.67e-07 | 0.0157 |

| compartment | subcluster | gene | avg_log2FC | pct.1 | pct.2 | p_val | p_val_adj |
|---|---|---|---|---|---|---|---|
| Malignant epithelial | C5 Am/CEA-like ECs | CHP1 | 0.95 | 0.548 | 0.402 | 9.77e-10 | 3.29e-05 |
| Malignant epithelial | C5 Am/CEA-like ECs | IST1 | 0.948 | 0.494 | 0.343 | 4.19e-09 | 0.000141 |
| Malignant epithelial | C5 Am/CEA-like ECs | MSLN | 0.946 | 0.559 | 0.296 | 1.55e-21 | 5.21e-17 |
| Malignant epithelial | C5 Am/CEA-like ECs | HLA-A | 0.945 | 0.981 | 0.975 | 2.91e-23 | 9.79e-19 |
| Malignant epithelial | C5 Am/CEA-like ECs | CARD19 | 0.942 | 0.648 | 0.553 | 7.55e-09 | 0.000254 |
| Malignant epithelial | C5 Am/CEA-like ECs | ZDHHC12 | 0.941 | 0.655 | 0.516 | 3.45e-12 | 1.16e-07 |
| Malignant epithelial | C5 Am/CEA-like ECs | WTAP | 0.941 | 0.556 | 0.389 | 9.7e-12 | 3.27e-07 |
| Malignant epithelial | C5 Am/CEA-like ECs | PSAP | 0.938 | 0.862 | 0.725 | 1.69e-21 | 5.71e-17 |
| Malignant epithelial | C5 Am/CEA-like ECs | LLGL2 | 0.938 | 0.559 | 0.407 | 2.4e-10 | 8.08e-06 |
| Malignant epithelial | C5 Am/CEA-like ECs | LRRFIP2 | 0.936 | 0.406 | 0.294 | 8.65e-07 | 0.0291 |
| Malignant epithelial | C5 Am/CEA-like ECs | CAPZA2 | 0.934 | 0.759 | 0.575 | 7.71e-19 | 2.6e-14 |
| Malignant epithelial | C5 Am/CEA-like ECs | TNFRSF21 | 0.928 | 0.479 | 0.336 | 2.23e-08 | 0.000752 |
| Malignant epithelial | C5 Am/CEA-like ECs | PDLIM1 | 0.925 | 0.797 | 0.607 | 7.35e-23 | 2.48e-18 |
| Malignant epithelial | C5 Am/CEA-like ECs | BLCAP | 0.923 | 0.544 | 0.406 | 9.43e-09 | 0.000318 |
| Malignant epithelial | C5 Am/CEA-like ECs | HSPB1 | 0.921 | 0.996 | 0.979 | 3.05e-27 | 1.03e-22 |
| Malignant epithelial | C5 Am/CEA-like ECs | ABHD11 | 0.92 | 0.766 | 0.625 | 2.69e-16 | 9.06e-12 |
| Malignant epithelial | C5 Am/CEA-like ECs | RPN2 | 0.92 | 0.682 | 0.569 | 4.97e-08 | 0.00168 |
| Malignant epithelial | C5 Am/CEA-like ECs | ARHGAP27 | 0.919 | 0.429 | 0.281 | 3.23e-09 | 0.000109 |
| Malignant epithelial | C5 Am/CEA-like ECs | ORMDL3 | 0.919 | 0.314 | 0.192 | 2.13e-07 | 0.00716 |
| Malignant epithelial | C5 Am/CEA-like ECs | HSPA5 | 0.917 | 0.762 | 0.712 | 2.86e-09 | 9.63e-05 |
| Malignant epithelial | C5 Am/CEA-like ECs | PPP1CB | 0.916 | 0.732 | 0.579 | 8.28e-15 | 2.79e-10 |
| Malignant epithelial | C5 Am/CEA-like ECs | TAX1BP1 | 0.914 | 0.843 | 0.703 | 2.37e-23 | 8e-19 |
| Malignant epithelial | C5 Am/CEA-like ECs | CCDC64B | 0.914 | 0.475 | 0.293 | 4e-10 | 1.35e-05 |
| Malignant epithelial | C5 Am/CEA-like ECs | MXD1 | 0.911 | 0.648 | 0.469 | 5.83e-14 | 1.96e-09 |
| Malignant epithelial | C5 Am/CEA-like ECs | ATP6AP2 | 0.911 | 0.586 | 0.467 | 2.67e-09 | 8.99e-05 |
| Malignant epithelial | C5 Am/CEA-like ECs | TPT1 | 0.91 | 1.0 | 0.996 | 1.95e-55 | 6.58e-51 |
| Malignant epithelial | C5 Am/CEA-like ECs | SYAP1 | 0.908 | 0.69 | 0.514 | 5.89e-16 | 1.98e-11 |
| Malignant epithelial | C5 Am/CEA-like ECs | GABARAP | 0.907 | 0.548 | 0.379 | 6.7e-11 | 2.26e-06 |
| Malignant epithelial | C5 Am/CEA-like ECs | FAM50A | 0.906 | 0.54 | 0.391 | 1.27e-09 | 4.29e-05 |
| Malignant epithelial | C5 Am/CEA-like ECs | TMEM55B | 0.906 | 0.372 | 0.229 | 9.13e-09 | 0.000307 |
| Malignant epithelial | C5 Am/CEA-like ECs | PRR13 | 0.902 | 0.935 | 0.885 | 1.2e-29 | 4.04e-25 |
| Malignant epithelial | C5 Am/CEA-like ECs | ZFAND2B | 0.902 | 0.395 | 0.269 | 2.14e-07 | 0.0072 |
| Malignant epithelial | C5 Am/CEA-like ECs | KMT2E | 0.901 | 0.632 | 0.511 | 5.17e-11 | 1.74e-06 |
| Malignant epithelial | C5 Am/CEA-like ECs | UGP2 | 0.901 | 0.506 | 0.371 | 7.78e-08 | 0.00262 |
| Malignant epithelial | C5 Am/CEA-like ECs | MAFG | 0.901 | 0.299 | 0.187 | 1.46e-06 | 0.0491 |
| Malignant epithelial | C5 Am/CEA-like ECs | MAFK | 0.9 | 0.372 | 0.227 | 6.83e-08 | 0.0023 |
| Malignant epithelial | C5 Am/CEA-like ECs | PLAC8 | 0.9 | 0.533 | 0.419 | 3.35e-07 | 0.0113 |
| Malignant epithelial | C5 Am/CEA-like ECs | EMC7 | 0.899 | 0.548 | 0.429 | 6.46e-08 | 0.00218 |
| Malignant epithelial | C5 Am/CEA-like ECs | MLX | 0.897 | 0.437 | 0.296 | 6.18e-08 | 0.00208 |
| Malignant epithelial | C5 Am/CEA-like ECs | TNIP1 | 0.893 | 0.54 | 0.41 | 5.51e-08 | 0.00186 |
| Malignant epithelial | C5 Am/CEA-like ECs | RAB4B | 0.884 | 0.276 | 0.162 | 8.74e-07 | 0.0294 |
| Malignant epithelial | C5 Am/CEA-like ECs | GPX2 | 0.883 | 0.755 | 0.484 | 7.59e-21 | 2.56e-16 |
| Malignant epithelial | C5 Am/CEA-like ECs | HERPUD1 | 0.882 | 0.736 | 0.621 | 2.01e-10 | 6.77e-06 |
| Malignant epithelial | C5 Am/CEA-like ECs | SNHG18 | 0.881 | 0.268 | 0.145 | 4.21e-08 | 0.00142 |
| Malignant epithelial | C5 Am/CEA-like ECs | IGKC | 0.878 | 0.885 | 0.884 | 2.86e-11 | 9.65e-07 |
| Malignant epithelial | C5 Am/CEA-like ECs | MDK | 0.871 | 0.904 | 0.81 | 6.960000000000001e-21 | 2.34e-16 |
| Malignant epithelial | C5 Am/CEA-like ECs | MTIF3 | 0.87 | 0.567 | 0.424 | 3.46e-09 | 0.000117 |
| Malignant epithelial | C5 Am/CEA-like ECs | EPHA2 | 0.869 | 0.483 | 0.351 | 2.4e-08 | 0.000808 |
| Malignant epithelial | C5 Am/CEA-like ECs | FTL | 0.868 | 1.0 | 0.998 | 3.17e-31 | 1.07e-26 |
| Malignant epithelial | C5 Am/CEA-like ECs | DSG2 | 0.868 | 0.598 | 0.445 | 2.45e-10 | 8.26e-06 |
| Malignant epithelial | C5 Am/CEA-like ECs | SIRT7 | 0.868 | 0.437 | 0.323 | 1.34e-06 | 0.045 |
| Malignant epithelial | C5 Am/CEA-like ECs | RBCK1 | 0.865 | 0.801 | 0.779 | 1.15e-09 | 3.87e-05 |
| Malignant epithelial | C5 Am/CEA-like ECs | CD82 | 0.864 | 0.513 | 0.367 | 9.85e-10 | 3.32e-05 |
| Malignant epithelial | C5 Am/CEA-like ECs | TMEM248 | 0.861 | 0.525 | 0.407 | 5.79e-08 | 0.00195 |
| Malignant epithelial | C5 Am/CEA-like ECs | PTTG1IP | 0.86 | 0.805 | 0.663 | 3.66e-17 | 1.23e-12 |
| Malignant epithelial | C5 Am/CEA-like ECs | ARF4 | 0.86 | 0.866 | 0.759 | 8e-15 | 2.69e-10 |
| Malignant epithelial | C5 Am/CEA-like ECs | FAM84A | 0.858 | 0.379 | 0.207 | 2.46e-11 | 8.3e-07 |
| Malignant epithelial | C5 Am/CEA-like ECs | RHBDF1 | 0.857 | 0.28 | 0.167 | 1.03e-06 | 0.0345 |
| Malignant epithelial | C5 Am/CEA-like ECs | PPP1R16A | 0.856 | 0.475 | 0.331 | 4.84e-09 | 0.000163 |
| Malignant epithelial | C5 Am/CEA-like ECs | ATP6AP1 | 0.856 | 0.533 | 0.389 | 6.92e-09 | 0.000233 |
| Malignant epithelial | C5 Am/CEA-like ECs | KLF5 | 0.854 | 0.736 | 0.593 | 5.8e-13 | 1.95e-08 |
| Malignant epithelial | C5 Am/CEA-like ECs | TPM4 | 0.853 | 0.847 | 0.744 | 6.44e-20 | 2.17e-15 |
| Malignant epithelial | C5 Am/CEA-like ECs | CHMP5 | 0.852 | 0.728 | 0.605 | 1.01e-13 | 3.42e-09 |
| Malignant epithelial | C5 Am/CEA-like ECs | MPHOSPH8 | 0.85 | 0.533 | 0.376 | 8.28e-09 | 0.000279 |

| compartment | subcluster | gene | avg_log2FC | pct.1 | pct.2 | p_val | p_val_adj |
|---|---|---|---|---|---|---|---|
| Malignant epithelial | C5 Am/CEA-like ECs | TMED4 | 0.848 | 0.598 | 0.474 | 2.18e-09 | 7.35e-05 |
| Malignant epithelial | C5 Am/CEA-like ECs | CAP1 | 0.848 | 0.559 | 0.437 | 5.25e-08 | 0.00177 |
| Malignant epithelial | C5 Am/CEA-like ECs | HLA-E | 0.847 | 0.912 | 0.888 | 8.41e-17 | 2.83e-12 |
| Malignant epithelial | C5 Am/CEA-like ECs | ERBB3 | 0.847 | 0.632 | 0.458 | 3.35e-11 | 1.13e-06 |
| Malignant epithelial | C5 Am/CEA-like ECs | HIGD1A | 0.847 | 0.594 | 0.429 | 2.48e-10 | 8.34e-06 |
| Malignant epithelial | C5 Am/CEA-like ECs | TMC5 | 0.843 | 0.648 | 0.51 | 1.84e-09 | 6.21e-05 |
| Malignant epithelial | C5 Am/CEA-like ECs | PDCD6IP | 0.843 | 0.464 | 0.333 | 1.95e-07 | 0.00656 |
| Malignant epithelial | C5 Am/CEA-like ECs | CD2AP | 0.843 | 0.525 | 0.407 | 4.97e-07 | 0.0168 |
| Malignant epithelial | C5 Am/CEA-like ECs | DYNLRB1 | 0.842 | 0.95 | 0.914 | 1.12e-29 | 3.78e-25 |
| Malignant epithelial | C5 Am/CEA-like ECs | CTNNA1 | 0.842 | 0.67 | 0.506 | 4.8e-13 | 1.62e-08 |
| Malignant epithelial | C5 Am/CEA-like ECs | CAPN2 | 0.842 | 0.598 | 0.452 | 7.06e-09 | 0.000238 |
| Malignant epithelial | C5 Am/CEA-like ECs | RABAC1 | 0.84 | 0.766 | 0.659 | 9.13e-15 | 3.08e-10 |
| Malignant epithelial | C5 Am/CEA-like ECs | RER1 | 0.84 | 0.663 | 0.523 | 1.55e-10 | 5.23e-06 |
| Malignant epithelial | C5 Am/CEA-like ECs | TSC22D3 | 0.84 | 0.617 | 0.507 | 1.08e-08 | 0.000364 |
| Malignant epithelial | C5 Am/CEA-like ECs | LMAN2 | 0.833 | 0.793 | 0.7 | 2.09e-17 | 7.04e-13 |
| Malignant epithelial | C5 Am/CEA-like ECs | POLD4 | 0.833 | 0.582 | 0.449 | 3.85e-10 | 1.3e-05 |
| Malignant epithelial | C5 Am/CEA-like ECs | DAZAP2 | 0.83 | 0.847 | 0.703 | 1.51e-21 | 5.1e-17 |
| Malignant epithelial | C5 Am/CEA-like ECs | VAMP2 | 0.828 | 0.502 | 0.379 | 5.67e-07 | 0.0191 |
| Malignant epithelial | C5 Am/CEA-like ECs | ESRRA | 0.826 | 0.437 | 0.311 | 3.72e-07 | 0.0125 |
| Malignant epithelial | C5 Am/CEA-like ECs | ATP6V0E1 | 0.825 | 0.927 | 0.871 | 1.08e-21 | 3.63e-17 |
| Malignant epithelial | C5 Am/CEA-like ECs | CDC42 | 0.825 | 0.816 | 0.707 | 4.52e-15 | 1.52e-10 |
| Malignant epithelial | C5 Am/CEA-like ECs | CD59 | 0.823 | 0.839 | 0.748 | 8.65e-15 | 2.91e-10 |
| Malignant epithelial | C5 Am/CEA-like ECs | DAP | 0.815 | 0.471 | 0.346 | 1.44e-07 | 0.00484 |
| Malignant epithelial | C5 Am/CEA-like ECs | UBE2A | 0.81 | 0.571 | 0.478 | 1.78e-07 | 0.006 |
| Malignant epithelial | C5 Am/CEA-like ECs | POR | 0.808 | 0.441 | 0.304 | 9.5e-08 | 0.0032 |
| Malignant epithelial | C5 Am/CEA-like ECs | VAMP3 | 0.807 | 0.464 | 0.349 | 1.44e-06 | 0.0484 |
| Malignant epithelial | C5 Am/CEA-like ECs | RAB14 | 0.805 | 0.49 | 0.337 | 7.29e-09 | 0.000246 |
| Malignant epithelial | C5 Am/CEA-like ECs | IFNGR2 | 0.804 | 0.533 | 0.443 | 4.19e-07 | 0.0141 |
| Malignant epithelial | C5 Am/CEA-like ECs | TMEM179B | 0.799 | 0.613 | 0.501 | 6.23e-08 | 0.0021 |
| Malignant epithelial | C5 Am/CEA-like ECs | KRT7 | 0.796 | 0.387 | 0.205 | 2.27e-12 | 7.66e-08 |
| Malignant epithelial | C5 Am/CEA-like ECs | NAMPT | 0.795 | 0.617 | 0.504 | 8.13e-09 | 0.000274 |
| Malignant epithelial | C5 Am/CEA-like ECs | CST3 | 0.793 | 0.958 | 0.976 | 1.96e-17 | 6.59e-13 |
| Malignant epithelial | C5 Am/CEA-like ECs | ARPC5 | 0.791 | 0.762 | 0.639 | 4.11e-14 | 1.38e-09 |
| Malignant epithelial | C5 Am/CEA-like ECs | HEBP2 | 0.79 | 0.824 | 0.74 | 1.77e-21 | 5.97e-17 |
| Malignant epithelial | C5 Am/CEA-like ECs | CD151 | 0.785 | 0.805 | 0.721 | 6.46e-15 | 2.18e-10 |
| Malignant epithelial | C5 Am/CEA-like ECs | ZBTB7A | 0.78 | 0.483 | 0.375 | 1.35e-06 | 0.0456 |
| Malignant epithelial | C5 Am/CEA-like ECs | C20orf24 | 0.777 | 0.782 | 0.701 | 4.57e-14 | 1.54e-09 |
| Malignant epithelial | C5 Am/CEA-like ECs | VOPP1 | 0.777 | 0.464 | 0.339 | 6.52e-07 | 0.022 |
| Malignant epithelial | C5 Am/CEA-like ECs | TMED2 | 0.774 | 0.843 | 0.752 | 1.26e-15 | 4.25e-11 |
| Malignant epithelial | C5 Am/CEA-like ECs | SVIP | 0.773 | 0.59 | 0.439 | 2.47e-09 | 8.33e-05 |
| Malignant epithelial | C5 Am/CEA-like ECs | DEDD2 | 0.767 | 0.533 | 0.395 | 1.47e-08 | 0.000495 |
| Malignant epithelial | C5 Am/CEA-like ECs | CTSB | 0.767 | 0.678 | 0.603 | 1.39e-07 | 0.00467 |
| Malignant epithelial | C5 Am/CEA-like ECs | NT5C3A | 0.766 | 0.693 | 0.617 | 4.53e-08 | 0.00153 |
| Malignant epithelial | C5 Am/CEA-like ECs | POMP | 0.765 | 0.943 | 0.846 | 3.05e-26 | 1.03e-21 |
| Malignant epithelial | C5 Am/CEA-like ECs | ITGB1 | 0.765 | 0.728 | 0.64 | 4e-11 | 1.35e-06 |
| Malignant epithelial | C5 Am/CEA-like ECs | SERP1 | 0.762 | 0.889 | 0.866 | 3.9e-20 | 1.31e-15 |
| Malignant epithelial | C5 Am/CEA-like ECs | KDELR2 | 0.762 | 0.885 | 0.821 | 4.24e-17 | 1.43e-12 |
| Malignant epithelial | C5 Am/CEA-like ECs | SPINT2 | 0.761 | 0.966 | 0.938 | 2.96e-31 | 9.970000000000001e-27 |
| Malignant epithelial | C5 Am/CEA-like ECs | BUD31 | 0.76 | 0.831 | 0.712 | 1.98e-14 | 6.67e-10 |
| Malignant epithelial | C5 Am/CEA-like ECs | RAB5C | 0.759 | 0.678 | 0.546 | 2.58e-10 | 8.69e-06 |
| Malignant epithelial | C5 Am/CEA-like ECs | ATP6V1D | 0.759 | 0.51 | 0.394 | 1.29e-06 | 0.0434 |
| Malignant epithelial | C5 Am/CEA-like ECs | SDF4 | 0.758 | 0.59 | 0.452 | 1.14e-09 | 3.84e-05 |
| Malignant epithelial | C5 Am/CEA-like ECs | PRAP1 | 0.756 | 0.605 | 0.347 | 1.04e-19 | 3.52e-15 |
| Malignant epithelial | C5 Am/CEA-like ECs | AGPAT2 | 0.756 | 0.87 | 0.785 | 1.28e-19 | 4.32e-15 |
| Malignant epithelial | C5 Am/CEA-like ECs | WSB1 | 0.749 | 0.713 | 0.572 | 3.73e-10 | 1.26e-05 |
| Malignant epithelial | C5 Am/CEA-like ECs | STAU1 | 0.745 | 0.69 | 0.574 | 1.12e-09 | 3.76e-05 |
| Malignant epithelial | C5 Am/CEA-like ECs | ARF1 | 0.744 | 0.9 | 0.781 | 1.07e-22 | 3.59e-18 |
| Malignant epithelial | C5 Am/CEA-like ECs | GNB2 | 0.74 | 0.816 | 0.714 | 1.55e-16 | 5.22e-12 |
| Malignant epithelial | C5 Am/CEA-like ECs | CHMP4B | 0.739 | 0.877 | 0.789 | 4.01e-20 | 1.35e-15 |
| Malignant epithelial | C5 Am/CEA-like ECs | RAB1A | 0.739 | 0.686 | 0.599 | 6.44e-09 | 0.000217 |
| Malignant epithelial | C5 Am/CEA-like ECs | BZW1 | 0.737 | 0.682 | 0.542 | 5.16e-10 | 1.74e-05 |
| Malignant epithelial | C5 Am/CEA-like ECs | GID8 | 0.735 | 0.693 | 0.568 | 9.26e-10 | 3.12e-05 |
| Malignant epithelial | C5 Am/CEA-like ECs | TMBIM6 | 0.732 | 0.95 | 0.893 | 5.52e-21 | 1.86e-16 |
| Malignant epithelial | C5 Am/CEA-like ECs | CHMP2A | 0.732 | 0.82 | 0.753 | 5.12e-10 | 1.73e-05 |

| compartment | subcluster | gene | avg_log2FC | pct.1 | pct.2 | p_val | p_val_adj |
|---|---|---|---|---|---|---|---|
| Malignant epithelial | C5 Am/CEA-like ECs | RHBDD2 | 0.726 | 0.536 | 0.403 | 1.69e-08 | 0.00057 |
| Malignant epithelial | C5 Am/CEA-like ECs | YPEL3 | 0.726 | 0.471 | 0.33 | 6.08e-07 | 0.0205 |
| Malignant epithelial | C5 Am/CEA-like ECs | TMED9 | 0.724 | 0.808 | 0.729 | 9.03e-13 | 3.04e-08 |
| Malignant epithelial | C5 Am/CEA-like ECs | SEC61B | 0.723 | 0.908 | 0.898 | 2.3e-21 | 7.73e-17 |
| Malignant epithelial | C5 Am/CEA-like ECs | KLK10 | 0.722 | 0.54 | 0.45 | 9.88e-07 | 0.0333 |
| Malignant epithelial | C5 Am/CEA-like ECs | RRAS | 0.712 | 0.648 | 0.502 | 3.86e-10 | 1.3e-05 |
| Malignant epithelial | C5 Am/CEA-like ECs | TSC22D4 | 0.707 | 0.487 | 0.364 | 4.38e-07 | 0.0148 |
| Malignant epithelial | C5 Am/CEA-like ECs | ADAP1 | 0.707 | 0.533 | 0.42 | 7.09e-07 | 0.0239 |
| Malignant epithelial | C5 Am/CEA-like ECs | OS9 | 0.704 | 0.54 | 0.434 | 1.3e-06 | 0.0439 |
| Malignant epithelial | C5 Am/CEA-like ECs | KLF3 | 0.702 | 0.659 | 0.516 | 2.31e-09 | 7.78e-05 |
| Malignant epithelial | C5 Am/CEA-like ECs | SFT2D1 | 0.698 | 0.636 | 0.562 | 6.04e-08 | 0.00204 |
| Malignant epithelial | C5 Am/CEA-like ECs | NANS | 0.694 | 0.625 | 0.506 | 3.94e-08 | 0.00133 |
| Malignant epithelial | C5 Am/CEA-like ECs | SARAF | 0.692 | 0.575 | 0.475 | 3.67e-07 | 0.0124 |
| Malignant epithelial | C5 Am/CEA-like ECs | ANXA11 | 0.688 | 0.877 | 0.776 | 4.26e-16 | 1.43e-11 |
| Malignant epithelial | C5 Am/CEA-like ECs | GABARAPL2 | 0.684 | 0.724 | 0.679 | 5.39e-07 | 0.0182 |
| Malignant epithelial | C5 Am/CEA-like ECs | MYDGF | 0.682 | 0.808 | 0.767 | 1.26e-08 | 0.000425 |
| Malignant epithelial | C5 Am/CEA-like ECs | CD63 | 0.681 | 0.958 | 0.964 | 8.66e-20 | 2.92e-15 |
| Malignant epithelial | C5 Am/CEA-like ECs | VAPA | 0.676 | 0.674 | 0.588 | 5.07e-09 | 0.000171 |
| Malignant epithelial | C5 Am/CEA-like ECs | FTH1 | 0.664 | 1.0 | 1.0 | 7.66e-16 | 2.58e-11 |
| Malignant epithelial | C5 Am/CEA-like ECs | TSC22D1 | 0.66 | 0.747 | 0.631 | 1.25e-09 | 4.22e-05 |
| Malignant epithelial | C5 Am/CEA-like ECs | PPDPF | 0.657 | 0.966 | 0.989 | 4.49e-26 | 1.51e-21 |
| Malignant epithelial | C5 Am/CEA-like ECs | ADRM1 | 0.657 | 0.728 | 0.66 | 2.54e-09 | 8.57e-05 |
| Malignant epithelial | C5 Am/CEA-like ECs | MVP | 0.657 | 0.648 | 0.525 | 1.11e-08 | 0.000372 |
| Malignant epithelial | C5 Am/CEA-like ECs | C4orf48 | 0.653 | 0.808 | 0.692 | 1.9e-12 | 6.4e-08 |
| Malignant epithelial | C5 Am/CEA-like ECs | BAG1 | 0.648 | 0.655 | 0.585 | 7.84e-07 | 0.0264 |
| Malignant epithelial | C5 Am/CEA-like ECs | TMEM208 | 0.644 | 0.701 | 0.606 | 2.06e-07 | 0.00693 |
| Malignant epithelial | C5 Am/CEA-like ECs | CAST | 0.641 | 0.831 | 0.732 | 1.56e-12 | 5.27e-08 |
| Malignant epithelial | C5 Am/CEA-like ECs | SEP15 | 0.638 | 0.628 | 0.518 | 5.99e-08 | 0.00202 |
| Malignant epithelial | C5 Am/CEA-like ECs | CALR | 0.635 | 0.885 | 0.851 | 2.81e-08 | 0.000948 |
| Malignant epithelial | C5 Am/CEA-like ECs | ANXA4 | 0.635 | 0.674 | 0.579 | 3.14e-08 | 0.00106 |
| Malignant epithelial | C5 Am/CEA-like ECs | CIB1 | 0.634 | 0.854 | 0.775 | 4.36e-11 | 1.47e-06 |
| Malignant epithelial | C5 Am/CEA-like ECs | KRT10 | 0.632 | 0.762 | 0.704 | 1.52e-08 | 0.000511 |
| Malignant epithelial | C5 Am/CEA-like ECs | ARHGDIA | 0.631 | 0.72 | 0.643 | 1.06e-09 | 3.59e-05 |
| Malignant epithelial | C5 Am/CEA-like ECs | MMP24-AS1 | 0.624 | 0.736 | 0.675 | 2.15e-07 | 0.00724 |
| Malignant epithelial | C5 Am/CEA-like ECs | NDFIP1 | 0.62 | 0.67 | 0.578 | 1.16e-06 | 0.039 |
| Malignant epithelial | C5 Am/CEA-like ECs | MYL6 | 0.619 | 1.0 | 0.985 | 6.54e-30 | 2.2e-25 |
| Malignant epithelial | C5 Am/CEA-like ECs | ARPC4 | 0.618 | 0.605 | 0.5 | 1.05e-06 | 0.0354 |
| Malignant epithelial | C5 Am/CEA-like ECs | SEC61G | 0.617 | 0.969 | 0.953 | 4.07e-22 | 1.37e-17 |
| Malignant epithelial | C5 Am/CEA-like ECs | TPM3 | 0.613 | 0.839 | 0.741 | 6.56e-13 | 2.21e-08 |
| Malignant epithelial | C5 Am/CEA-like ECs | ATP6V1F | 0.601 | 0.812 | 0.785 | 3.58e-11 | 1.21e-06 |
| Malignant epithelial | C5 Am/CEA-like ECs | TMCO1 | 0.601 | 0.693 | 0.617 | 6.03e-07 | 0.0203 |
| Malignant epithelial | C5 Am/CEA-like ECs | GLUL | 0.599 | 0.406 | 0.262 | 5.35e-08 | 0.0018 |
| Malignant epithelial | C5 Am/CEA-like ECs | MALAT1 | 0.598 | 1.0 | 0.997 | 8.41e-17 | 2.84e-12 |
| Malignant epithelial | C5 Am/CEA-like ECs | IFRD1 | 0.598 | 0.751 | 0.651 | 2.69e-07 | 0.00908 |
| Malignant epithelial | C5 Am/CEA-like ECs | CCDC107 | 0.597 | 0.575 | 0.463 | 1.03e-06 | 0.0348 |
| Malignant epithelial | C5 Am/CEA-like ECs | RAB2A | 0.596 | 0.812 | 0.765 | 4.48e-08 | 0.00151 |
| Malignant epithelial | C5 Am/CEA-like ECs | FNIP1 | 0.596 | 0.598 | 0.473 | 1.05e-06 | 0.0354 |
| Malignant epithelial | C5 Am/CEA-like ECs | VAMP8 | 0.593 | 0.912 | 0.883 | 8.52e-18 | 2.87e-13 |
| Malignant epithelial | C5 Am/CEA-like ECs | RAB7A | 0.592 | 0.762 | 0.669 | 5.43e-09 | 0.000183 |
| Malignant epithelial | C5 Am/CEA-like ECs | CCNI | 0.588 | 0.762 | 0.637 | 1.81e-07 | 0.0061 |
| Malignant epithelial | C5 Am/CEA-like ECs | JUP | 0.586 | 0.797 | 0.741 | 7.17e-08 | 0.00242 |
| Malignant epithelial | C5 Am/CEA-like ECs | GNG5 | 0.585 | 0.854 | 0.808 | 6.91e-13 | 2.33e-08 |
| Malignant epithelial | C5 Am/CEA-like ECs | DNAJB6 | 0.584 | 0.739 | 0.677 | 2.68e-07 | 0.00902 |
| Malignant epithelial | C5 Am/CEA-like ECs | RPS27L | 0.578 | 0.858 | 0.841 | 2.31e-10 | 7.79e-06 |
| Malignant epithelial | C5 Am/CEA-like ECs | PSMB1 | 0.576 | 0.854 | 0.757 | 2.21e-12 | 7.44e-08 |
| Malignant epithelial | C5 Am/CEA-like ECs | SAP18 | 0.57 | 0.923 | 0.876 | 1e-12 | 3.38e-08 |
| Malignant epithelial | C5 Am/CEA-like ECs | GPRC5A | 0.568 | 0.9 | 0.817 | 2.37e-10 | 7.97e-06 |
| Malignant epithelial | C5 Am/CEA-like ECs | LAPTM4A | 0.565 | 0.854 | 0.765 | 2.22e-11 | 7.49e-07 |
| Malignant epithelial | C5 Am/CEA-like ECs | MYL12B | 0.564 | 0.908 | 0.892 | 1.12e-12 | 3.77e-08 |
| Malignant epithelial | C5 Am/CEA-like ECs | AUP1 | 0.563 | 0.713 | 0.633 | 6.38e-08 | 0.00215 |
| Malignant epithelial | C5 Am/CEA-like ECs | LSM3 | 0.563 | 0.732 | 0.609 | 2.51e-07 | 0.00846 |
| Malignant epithelial | C5 Am/CEA-like ECs | MYL12A | 0.562 | 0.916 | 0.891 | 1.72e-08 | 0.000578 |
| Malignant epithelial | C5 Am/CEA-like ECs | LASP1 | 0.56 | 0.559 | 0.442 | 1.14e-06 | 0.0385 |
| Malignant epithelial | C5 Am/CEA-like ECs | LAMTOR4 | 0.551 | 0.843 | 0.82 | 1.25e-06 | 0.0421 |

| compartment | subcluster | gene | avg_log2FC | pct.1 | pct.2 | p_val | p_val_adj |
|---|---|---|---|---|---|---|---|
| Malignant epithelial | C5 Am/CEA-like ECs | SSU72 | 0.543 | 0.736 | 0.671 | 5.29e-07 | 0.0178 |
| Malignant epithelial | C5 Am/CEA-like ECs | ALDOA | 0.542 | 0.977 | 0.965 | 3.99e-09 | 0.000134 |
| Malignant epithelial | C5 Am/CEA-like ECs | ZNHIT1 | 0.537 | 0.824 | 0.796 | 2.04e-10 | 6.87e-06 |
| Malignant epithelial | C5 Am/CEA-like ECs | ITM2B | 0.536 | 0.847 | 0.777 | 3.98e-10 | 1.34e-05 |
| Malignant epithelial | C5 Am/CEA-like ECs | RHOA | 0.534 | 0.843 | 0.784 | 6.64e-09 | 0.000224 |
| Malignant epithelial | C5 Am/CEA-like ECs | CTSD | 0.533 | 0.851 | 0.774 | 1.56e-12 | 5.26e-08 |
| Malignant epithelial | C5 Am/CEA-like ECs | PERP | 0.532 | 0.943 | 0.892 | 1.24e-14 | 4.18e-10 |
| Malignant epithelial | C5 Am/CEA-like ECs | CLDN4 | 0.523 | 0.985 | 0.967 | 4.07e-10 | 1.37e-05 |
| Malignant epithelial | C5 Am/CEA-like ECs | EDF1 | 0.509 | 0.95 | 0.957 | 6.34e-18 | 2.14e-13 |
| Malignant epithelial | C5 Am/CEA-like ECs | OAZ1 | 0.508 | 0.996 | 0.973 | 5.9e-22 | 1.99e-17 |
| Malignant epithelial | C5 Am/CEA-like ECs | ARPC1B | 0.498 | 0.789 | 0.765 | 5.18e-08 | 0.00175 |
| Malignant epithelial | C5 Am/CEA-like ECs | ACTN4 | 0.487 | 0.805 | 0.747 | 7.92e-08 | 0.00267 |
| Malignant epithelial | C5 Am/CEA-like ECs | YPEL5 | 0.486 | 0.793 | 0.625 | 3.13e-11 | 1.06e-06 |
| Malignant epithelial | C5 Am/CEA-like ECs | RALY | 0.481 | 0.801 | 0.719 | 3.37e-07 | 0.0114 |
| Malignant epithelial | C5 Am/CEA-like ECs | CFL1 | 0.474 | 0.985 | 0.974 | 1.67e-22 | 5.63e-18 |
| Malignant epithelial | C5 Am/CEA-like ECs | ACTB | 0.474 | 1.0 | 0.998 | 5.94e-16 | 2e-11 |
| Malignant epithelial | C5 Am/CEA-like ECs | SAT1 | 0.473 | 0.981 | 0.969 | 5.36e-11 | 1.81e-06 |
| Malignant epithelial | C5 Am/CEA-like ECs | RAC1 | 0.466 | 0.981 | 0.976 | 5.91e-28 | 1.99e-23 |
| Malignant epithelial | C5 Am/CEA-like ECs | SEPW1 | 0.458 | 0.858 | 0.831 | 3.48e-07 | 0.0117 |
| Malignant epithelial | C5 Am/CEA-like ECs | CHCHD2 | 0.457 | 0.985 | 0.957 | 3.29e-22 | 1.11e-17 |
| Malignant epithelial | C5 Am/CEA-like ECs | POLR2J | 0.445 | 0.805 | 0.747 | 9.59e-07 | 0.0323 |
| Malignant epithelial | C5 Am/CEA-like ECs | COX17 | 0.445 | 0.812 | 0.811 | 1.08e-06 | 0.0363 |
| Malignant epithelial | C5 Am/CEA-like ECs | VDAC2 | 0.439 | 0.904 | 0.844 | 7e-07 | 0.0236 |
| Malignant epithelial | C5 Am/CEA-like ECs | SKP1 | 0.43 | 0.939 | 0.895 | 1.32e-08 | 0.000445 |
| Malignant epithelial | C5 Am/CEA-like ECs | ARPC2 | 0.425 | 0.874 | 0.843 | 2.18e-08 | 0.000734 |
| Malignant epithelial | C5 Am/CEA-like ECs | COX7A2 | 0.419 | 0.989 | 0.972 | 4.38e-14 | 1.47e-09 |
| Malignant epithelial | C5 Am/CEA-like ECs | TMSB4X | 0.419 | 1.0 | 1.0 | 8.72e-09 | 0.000294 |
| Malignant epithelial | C5 Am/CEA-like ECs | ZFAND2A | 0.408 | 0.64 | 0.503 | 1.85e-08 | 0.000623 |
| Malignant epithelial | C5 Am/CEA-like ECs | CALM2 | 0.403 | 0.958 | 0.936 | 1.79e-09 | 6.02e-05 |
| Malignant epithelial | C5 Am/CEA-like ECs | YWHAZ | 0.401 | 0.962 | 0.944 | 3.78e-11 | 1.27e-06 |
| Malignant epithelial | C5 Am/CEA-like ECs | SERF2 | 0.391 | 0.992 | 0.996 | 1.54e-16 | 5.18e-12 |
| Malignant epithelial | C5 Am/CEA-like ECs | PSMA7 | 0.367 | 0.916 | 0.901 | 8.15e-07 | 0.0274 |
| Malignant epithelial | C5 Am/CEA-like ECs | NPC2 | 0.365 | 0.897 | 0.876 | 1.11e-08 | 0.000375 |
| Malignant epithelial | C5 Am/CEA-like ECs | RHOB | 0.365 | 0.881 | 0.848 | 5.34e-07 | 0.018 |
| Malignant epithelial | C5 Am/CEA-like ECs | C19orf33 | 0.316 | 0.973 | 0.954 | 6.1e-07 | 0.0206 |
| Malignant epithelial | C5 Am/CEA-like ECs | RPL36AL | 0.308 | 0.966 | 0.965 | 1.24e-07 | 0.00417 |
| Malignant epithelial | C5 Am/CEA-like ECs | H3F3B | 0.306 | 0.996 | 0.982 | 2.48e-08 | 0.000836 |
| Malignant epithelial | C6 DNA-r p ECs | LY6D | 5.827 | 0.565 | 0.057 | 3.16e-117 | 1.06e-112 |
| Malignant epithelial | C6 DNA-r p ECs | A2ML1 | 5.232 | 0.129 | 0.003 | 5.99e-71 | 2.02e-66 |
| Malignant epithelial | C6 DNA-r p ECs | DHRS2 | 4.801 | 0.177 | 0.024 | 1.18e-25 | 3.990000000000001e-21 |
| Malignant epithelial | C6 DNA-r p ECs | CCDC144NL-AS1 | 3.973 | 0.25 | 0.016 | 3.54e-72 | 1.19e-67 |
| Malignant epithelial | C6 DNA-r p ECs | RIMKLB | 3.653 | 0.105 | 0.007 | 1.75e-28 | 5.89e-24 |
| Malignant epithelial | C6 DNA-r p ECs | ZNF367 | 3.629 | 0.202 | 0.015 | 2.18e-49 | 7.33e-45 |
| Malignant epithelial | C6 DNA-r p ECs | TMEM40 | 3.617 | 0.129 | 0.007 | 5.08e-40 | 1.71e-35 |
| Malignant epithelial | C6 DNA-r p ECs | GPR87 | 3.597 | 0.169 | 0.012 | 6.039999999999999e-45 | 2.04e-40 |
| Malignant epithelial | C6 DNA-r p ECs | NRGN | 3.53 | 0.113 | 0.01 | 7.429999999999999e-24 | 2.51e-19 |
| Malignant epithelial | C6 DNA-r p ECs | RIBC2 | 3.498 | 0.113 | 0.011 | 2.1e-23 | 7.06e-19 |
| Malignant epithelial | C6 DNA-r p ECs | FAM83A | 3.297 | 0.395 | 0.063 | 1.16e-48 | 3.89e-44 |
| Malignant epithelial | C6 DNA-r p ECs | CRIP2 | 3.289 | 0.573 | 0.119 | 9.25e-57 | 3.12e-52 |
| Malignant epithelial | C6 DNA-r p ECs | RRM2 | 3.288 | 0.524 | 0.082 | 6.53e-67 | 2.2e-62 |
| Malignant epithelial | C6 DNA-r p ECs | FAM111B | 3.277 | 0.29 | 0.029 | 1.4e-54 | 4.71e-50 |
| Malignant epithelial | C6 DNA-r p ECs | CCNE2 | 3.265 | 0.169 | 0.015 | 2.06e-35 | 6.940000000000001e-31 |
| Malignant epithelial | C6 DNA-r p ECs | MIOX | 3.247 | 0.113 | 0.009 | 6.14e-26 | 2.07e-21 |
| Malignant epithelial | C6 DNA-r p ECs | CLSPN | 3.227 | 0.242 | 0.032 | 3.09e-34 | 1.04e-29 |
| Malignant epithelial | C6 DNA-r p ECs | DIAPH3 | 3.137 | 0.194 | 0.021 | 2.06e-34 | 6.94e-30 |
| Malignant epithelial | C6 DNA-r p ECs | PHACTR3 | 3.103 | 0.444 | 0.092 | 6.93e-41 | 2.33e-36 |
| Malignant epithelial | C6 DNA-r p ECs | MTFR2 | 3.077 | 0.194 | 0.017 | 1.2e-41 | 4.03e-37 |
| Malignant epithelial | C6 DNA-r p ECs | SERPINE1 | 3.055 | 0.194 | 0.045 | 1.89e-14 | 6.38e-10 |
| Malignant epithelial | C6 DNA-r p ECs | CDC45 | 3.051 | 0.226 | 0.027 | 4.42e-35 | 1.49e-30 |
| Malignant epithelial | C6 DNA-r p ECs | ESCO2 | 3.049 | 0.145 | 0.014 | 5.88e-28 | 1.98e-23 |
| Malignant epithelial | C6 DNA-r p ECs | KRT81 | 3.045 | 0.105 | 0.01 | 1.31e-21 | 4.4e-17 |
| Malignant epithelial | C6 DNA-r p ECs | ASF1B | 3.022 | 0.524 | 0.082 | 7.87e-66 | 2.65e-61 |
| Malignant epithelial | C6 DNA-r p ECs | IGFBP7 | 2.926 | 0.25 | 0.049 | 1.64e-23 | 5.53e-19 |
| Malignant epithelial | C6 DNA-r p ECs | UNC5B | 2.92 | 0.161 | 0.025 | 6.14e-20 | 2.07e-15 |

| compartment | subcluster | gene | avg_log2FC | pct.1 | pct.2 | p_val | p_val_adj |
|---|---|---|---|---|---|---|---|
| Malignant epithelial | C6 DNA-r p ECs | CDC6 | 2.918 | 0.298 | 0.043 | 1.81e-39 | 6.11e-35 |
| Malignant epithelial | C6 DNA-r p ECs | TCF19 | 2.917 | 0.298 | 0.044 | 9.34e-38 | 3.15e-33 |
| Malignant epithelial | C6 DNA-r p ECs | DSCC1 | 2.897 | 0.298 | 0.034 | 1.19e-49 | 4.01e-45 |
| Malignant epithelial | C6 DNA-r p ECs | NCAPH | 2.889 | 0.258 | 0.029 | 5.58e-43 | 1.88e-38 |
| Malignant epithelial | C6 DNA-r p ECs | C9orf24 | 2.881 | 0.145 | 0.019 | 4.620000000000001e-21 | 1.56e-16 |
| Malignant epithelial | C6 DNA-r p ECs | AP000251.3 | 2.872 | 0.105 | 0.011 | 3.01e-20 | 1.01e-15 |
| Malignant epithelial | C6 DNA-r p ECs | WDR76 | 2.852 | 0.185 | 0.026 | 8.13e-25 | 2.74e-20 |
| Malignant epithelial | C6 DNA-r p ECs | CENPU | 2.846 | 0.395 | 0.057 | 5.11e-52 | 1.72e-47 |
| Malignant epithelial | C6 DNA-r p ECs | DTL | 2.843 | 0.145 | 0.024 | 2.1e-16 | 7.08e-12 |
| Malignant epithelial | C6 DNA-r p ECs | PLAT | 2.829 | 0.185 | 0.034 | 2.23e-18 | 7.51e-14 |
| Malignant epithelial | C6 DNA-r p ECs | LINC01602 | 2.827 | 0.121 | 0.019 | 2.2e-14 | 7.42e-10 |
| Malignant epithelial | C6 DNA-r p ECs | E2F1 | 2.818 | 0.266 | 0.041 | 8e-33 | 2.7e-28 |
| Malignant epithelial | C6 DNA-r p ECs | CNTNAP2 | 2.817 | 0.185 | 0.024 | 2.36e-27 | 7.94e-23 |
| Malignant epithelial | C6 DNA-r p ECs | CDCA5 | 2.811 | 0.411 | 0.068 | 3.79e-47 | 1.28e-42 |
| Malignant epithelial | C6 DNA-r p ECs | VASN | 2.807 | 0.121 | 0.021 | 4.88e-13 | 1.64e-08 |
| Malignant epithelial | C6 DNA-r p ECs | BRCA2 | 2.804 | 0.185 | 0.024 | 3.31e-27 | 1.11e-22 |
| Malignant epithelial | C6 DNA-r p ECs | SPC25 | 2.799 | 0.411 | 0.066 | 2.12e-48 | 7.13e-44 |
| Malignant epithelial | C6 DNA-r p ECs | MELK | 2.727 | 0.387 | 0.056 | 1.31e-50 | 4.409999999999999e-46 |
| Malignant epithelial | C6 DNA-r p ECs | TH | 2.719 | 0.274 | 0.066 | 4.23e-20 | 1.42e-15 |
| Malignant epithelial | C6 DNA-r p ECs | ANXA8L1 | 2.715 | 0.105 | 0.013 | 1.69e-16 | 5.7e-12 |
| Malignant epithelial | C6 DNA-r p ECs | ANXA10 | 2.647 | 0.113 | 0.013 | 1.69e-19 | 5.71e-15 |
| Malignant epithelial | C6 DNA-r p ECs | CDT1 | 2.638 | 0.565 | 0.139 | 5.93e-44 | 2e-39 |
| Malignant epithelial | C6 DNA-r p ECs | RFC3 | 2.626 | 0.315 | 0.053 | 2.52e-34 | 8.48e-30 |
| Malignant epithelial | C6 DNA-r p ECs | PKMYT1 | 2.613 | 0.355 | 0.053 | 7.5e-44 | 2.53e-39 |
| Malignant epithelial | C6 DNA-r p ECs | CDK1 | 2.606 | 0.484 | 0.087 | 1.86e-50 | 6.26e-46 |
| Malignant epithelial | C6 DNA-r p ECs | TM4SF4 | 2.564 | 0.105 | 0.021 | 4.55e-10 | 1.53e-05 |
| Malignant epithelial | C6 DNA-r p ECs | SKA3 | 2.54 | 0.282 | 0.042 | 9.990000000000001e-35 | 3.37e-30 |
| Malignant epithelial | C6 DNA-r p ECs | RAD51 | 2.54 | 0.282 | 0.043 | 3.55e-34 | 1.19e-29 |
| Malignant epithelial | C6 DNA-r p ECs | POLA2 | 2.498 | 0.202 | 0.034 | 1.39e-21 | 4.68e-17 |
| Malignant epithelial | C6 DNA-r p ECs | DERL3 | 2.489 | 0.202 | 0.034 | 1.32e-21 | 4.44e-17 |
| Malignant epithelial | C6 DNA-r p ECs | KIAA0101 | 2.484 | 0.774 | 0.261 | 1.22e-48 | 4.12e-44 |
| Malignant epithelial | C6 DNA-r p ECs | FANCG | 2.482 | 0.331 | 0.05 | 5.33e-40 | 1.8e-35 |
| Malignant epithelial | C6 DNA-r p ECs | AC007325.4 | 2.48 | 0.242 | 0.064 | 7.05e-15 | 2.38e-10 |
| Malignant epithelial | C6 DNA-r p ECs | MNS1 | 2.452 | 0.266 | 0.043 | 1.12e-29 | 3.76e-25 |
| Malignant epithelial | C6 DNA-r p ECs | CENPP | 2.441 | 0.226 | 0.033 | 1.67e-28 | 5.63e-24 |
| Malignant epithelial | C6 DNA-r p ECs | NDC80 | 2.437 | 0.177 | 0.026 | 5e-22 | 1.68e-17 |
| Malignant epithelial | C6 DNA-r p ECs | TK1 | 2.427 | 0.79 | 0.252 | 5.01e-50 | 1.69e-45 |
| Malignant epithelial | C6 DNA-r p ECs | RAET1G | 2.425 | 0.121 | 0.026 | 2.06e-10 | 6.94e-06 |
| Malignant epithelial | C6 DNA-r p ECs | SHMT1 | 2.42 | 0.282 | 0.052 | 1.56e-27 | 5.26e-23 |
| Malignant epithelial | C6 DNA-r p ECs | WDR62 | 2.416 | 0.129 | 0.018 | 7.39e-18 | 2.49e-13 |
| Malignant epithelial | C6 DNA-r p ECs | NCAPG2 | 2.415 | 0.218 | 0.038 | 1.29e-22 | 4.35e-18 |
| Malignant epithelial | C6 DNA-r p ECs | TRIP13 | 2.414 | 0.323 | 0.071 | 5.85e-25 | 1.97e-20 |
| Malignant epithelial | C6 DNA-r p ECs | TYMS | 2.397 | 0.411 | 0.097 | 7.97e-31 | 2.69e-26 |
| Malignant epithelial | C6 DNA-r p ECs | GINS2 | 2.392 | 0.565 | 0.119 | 4.62e-50 | 1.56e-45 |
| Malignant epithelial | C6 DNA-r p ECs | C16orf59 | 2.389 | 0.226 | 0.036 | 1.48e-25 | 4.990000000000001e-21 |
| Malignant epithelial | C6 DNA-r p ECs | FEN1 | 2.373 | 0.411 | 0.094 | 7.06e-32 | 2.38e-27 |
| Malignant epithelial | C6 DNA-r p ECs | RP11-350J20.12 | 2.37 | 0.435 | 0.117 | 6.45e-28 | 2.17e-23 |
| Malignant epithelial | C6 DNA-r p ECs | RP11-554I8.2 | 2.368 | 0.177 | 0.034 | 1.14e-16 | 3.85e-12 |
| Malignant epithelial | C6 DNA-r p ECs | UPK3BL | 2.368 | 0.185 | 0.038 | 3.04e-16 | 1.02e-11 |
| Malignant epithelial | C6 DNA-r p ECs | GINS1 | 2.366 | 0.169 | 0.029 | 5.89e-18 | 1.99e-13 |
| Malignant epithelial | C6 DNA-r p ECs | ADRB2 | 2.338 | 0.153 | 0.028 | 2.87e-15 | 9.66e-11 |
| Malignant epithelial | C6 DNA-r p ECs | MCM10 | 2.317 | 0.129 | 0.017 | 8.93e-19 | 3.01e-14 |
| Malignant epithelial | C6 DNA-r p ECs | C5orf34 | 2.302 | 0.129 | 0.018 | 5.47e-17 | 1.84e-12 |
| Malignant epithelial | C6 DNA-r p ECs | PRIM1 | 2.297 | 0.379 | 0.079 | 7.19e-32 | 2.42e-27 |
| Malignant epithelial | C6 DNA-r p ECs | CKAP2L | 2.286 | 0.177 | 0.033 | 5.15e-17 | 1.73e-12 |
| Malignant epithelial | C6 DNA-r p ECs | FANCB | 2.276 | 0.266 | 0.049 | 8.7e-26 | 2.93e-21 |
| Malignant epithelial | C6 DNA-r p ECs | ITPKA | 2.272 | 0.137 | 0.031 | 8.18e-11 | 2.76e-06 |
| Malignant epithelial | C6 DNA-r p ECs | PCNA | 2.258 | 0.758 | 0.345 | 1.38e-32 | 4.65e-28 |
| Malignant epithelial | C6 DNA-r p ECs | MYBL2 | 2.248 | 0.323 | 0.06 | 5.5e-31 | 1.85e-26 |
| Malignant epithelial | C6 DNA-r p ECs | ATAD2 | 2.24 | 0.452 | 0.108 | 2.92e-33 | 9.85e-29 |
| Malignant epithelial | C6 DNA-r p ECs | NPB | 2.227 | 0.121 | 0.031 | 2.44e-08 | 0.000821 |
| Malignant epithelial | C6 DNA-r p ECs | SGOL1 | 2.223 | 0.25 | 0.048 | 5.61e-23 | 1.89e-18 |
| Malignant epithelial | C6 DNA-r p ECs | XRCC2 | 2.194 | 0.113 | 0.02 | 3.13e-12 | 1.05e-07 |
| Malignant epithelial | C6 DNA-r p ECs | SPC24 | 2.191 | 0.202 | 0.041 | 3.13e-17 | 1.05e-12 |

| compartment | subcluster | gene | avg_log2FC | pct.1 | pct.2 | p_val | p_val_adj |
|---|---|---|---|---|---|---|---|
| Malignant epithelial | C6 DNA-r p ECs | TCN1 | 2.179 | 0.21 | 0.027 | 2.14e-29 | 7.22e-25 |
| Malignant epithelial | C6 DNA-r p ECs | ATAD5 | 2.174 | 0.258 | 0.064 | 3.81e-17 | 1.28e-12 |
| Malignant epithelial | C6 DNA-r p ECs | ARHGEF39 | 2.17 | 0.177 | 0.042 | 9.73e-13 | 3.28e-08 |
| Malignant epithelial | C6 DNA-r p ECs | SERPINA4 | 2.15 | 0.113 | 0.025 | 3.63e-09 | 0.000122 |
| Malignant epithelial | C6 DNA-r p ECs | RP11-160O5.1 | 2.144 | 0.137 | 0.038 | 2.65e-08 | 0.000893 |
| Malignant epithelial | C6 DNA-r p ECs | XRCC3 | 2.143 | 0.153 | 0.028 | 2.43e-15 | 8.19e-11 |
| Malignant epithelial | C6 DNA-r p ECs | ZWINT | 2.14 | 0.605 | 0.165 | 7.42e-39 | 2.5e-34 |
| Malignant epithelial | C6 DNA-r p ECs | GDPD3 | 2.138 | 0.347 | 0.085 | 1.06e-23 | 3.56e-19 |
| Malignant epithelial | C6 DNA-r p ECs | RP11-680F8.1 | 2.138 | 0.29 | 0.066 | 1.95e-21 | 6.57e-17 |
| Malignant epithelial | C6 DNA-r p ECs | CENPK | 2.134 | 0.589 | 0.139 | 1.54e-43 | 5.2e-39 |
| Malignant epithelial | C6 DNA-r p ECs | IFITM10 | 2.124 | 0.137 | 0.038 | 2.52e-08 | 0.000848 |
| Malignant epithelial | C6 DNA-r p ECs | LIPT2 | 2.117 | 0.129 | 0.027 | 5.29e-11 | 1.78e-06 |
| Malignant epithelial | C6 DNA-r p ECs | MND1 | 2.104 | 0.476 | 0.1 | 1.23e-39 | 4.15e-35 |
| Malignant epithelial | C6 DNA-r p ECs | DSN1 | 2.103 | 0.331 | 0.085 | 2.22e-21 | 7.49e-17 |
| Malignant epithelial | C6 DNA-r p ECs | LRRCC1 | 2.101 | 0.355 | 0.082 | 3.78e-26 | 1.27e-21 |
| Malignant epithelial | C6 DNA-r p ECs | RRAD | 2.1 | 0.323 | 0.102 | 6.44e-16 | 2.17e-11 |
| Malignant epithelial | C6 DNA-r p ECs | BRCA1 | 2.098 | 0.234 | 0.054 | 5.03e-17 | 1.7e-12 |
| Malignant epithelial | C6 DNA-r p ECs | ACOT7 | 2.097 | 0.669 | 0.208 | 1.5e-38 | 5.06e-34 |
| Malignant epithelial | C6 DNA-r p ECs | KIF2C | 2.097 | 0.145 | 0.028 | 1.75e-13 | 5.89e-09 |
| Malignant epithelial | C6 DNA-r p ECs | CENPH | 2.094 | 0.516 | 0.153 | 1.22e-29 | 4.1e-25 |
| Malignant epithelial | C6 DNA-r p ECs | TMEM140 | 2.089 | 0.113 | 0.025 | 3.97e-09 | 0.000134 |
| Malignant epithelial | C6 DNA-r p ECs | RTKN2 | 2.086 | 0.194 | 0.032 | 2.78e-21 | 9.35e-17 |
| Malignant epithelial | C6 DNA-r p ECs | RECQL4 | 2.071 | 0.25 | 0.049 | 4.1e-22 | 1.38e-17 |
| Malignant epithelial | C6 DNA-r p ECs | SHCBP1 | 2.067 | 0.113 | 0.026 | 5.96e-09 | 0.000201 |
| Malignant epithelial | C6 DNA-r p ECs | MCM6 | 2.053 | 0.234 | 0.059 | 2.43e-15 | 8.18e-11 |
| Malignant epithelial | C6 DNA-r p ECs | SKA1 | 2.052 | 0.105 | 0.022 | 2.98e-09 | 0.0001 |
| Malignant epithelial | C6 DNA-r p ECs | KRT6B | 2.05 | 0.653 | 0.287 | 2.81e-24 | 9.45e-20 |
| Malignant epithelial | C6 DNA-r p ECs | ABLIM3 | 2.046 | 0.274 | 0.073 | 5.31e-17 | 1.79e-12 |
| Malignant epithelial | C6 DNA-r p ECs | ARHGEF19 | 2.023 | 0.169 | 0.045 | 8.01e-11 | 2.7e-06 |
| Malignant epithelial | C6 DNA-r p ECs | POLE2 | 2.02 | 0.153 | 0.034 | 3.46e-12 | 1.17e-07 |
| Malignant epithelial | C6 DNA-r p ECs | DHFR | 2.017 | 0.573 | 0.181 | 1.37e-30 | 4.62e-26 |
| Malignant epithelial | C6 DNA-r p ECs | HIRIP3 | 2.014 | 0.403 | 0.083 | 1.98e-33 | 6.67e-29 |
| Malignant epithelial | C6 DNA-r p ECs | LAMP3 | 2.012 | 0.105 | 0.022 | 5.45e-09 | 0.000184 |
| Malignant epithelial | C6 DNA-r p ECs | FBXO5 | 2.011 | 0.129 | 0.032 | 4.45e-09 | 0.00015 |
| Malignant epithelial | C6 DNA-r p ECs | ZNF385A | 1.985 | 0.21 | 0.066 | 3.88e-10 | 1.31e-05 |
| Malignant epithelial | C6 DNA-r p ECs | CENPM | 1.973 | 0.468 | 0.133 | 3.78e-26 | 1.28e-21 |
| Malignant epithelial | C6 DNA-r p ECs | SPHK1 | 1.971 | 0.21 | 0.057 | 1.71e-12 | 5.76e-08 |
| Malignant epithelial | C6 DNA-r p ECs | RP11-38M8.1 | 1.964 | 0.177 | 0.039 | 9.8e-14 | 3.3e-09 |
| Malignant epithelial | C6 DNA-r p ECs | UBE2T | 1.96 | 0.573 | 0.174 | 2.53e-31 | 8.540000000000001e-27 |
| Malignant epithelial | C6 DNA-r p ECs | PROM2 | 1.957 | 0.161 | 0.04 | 2.98e-11 | 1e-06 |
| Malignant epithelial | C6 DNA-r p ECs | WDHD1 | 1.951 | 0.113 | 0.027 | 2e-08 | 0.000673 |
| Malignant epithelial | C6 DNA-r p ECs | PC | 1.944 | 0.105 | 0.022 | 2.61e-09 | 8.8e-05 |
| Malignant epithelial | C6 DNA-r p ECs | NEURL1B | 1.94 | 0.153 | 0.03 | 6.67e-14 | 2.25e-09 |
| Malignant epithelial | C6 DNA-r p ECs | POLD1 | 1.934 | 0.25 | 0.055 | 1.52e-19 | 5.12e-15 |
| Malignant epithelial | C6 DNA-r p ECs | ZBED2 | 1.93 | 0.258 | 0.082 | 3.11e-12 | 1.05e-07 |
| Malignant epithelial | C6 DNA-r p ECs | NUSAP1 | 1.925 | 0.508 | 0.116 | 1.3e-38 | 4.369999999999999e-34 |
| Malignant epithelial | C6 DNA-r p ECs | XG | 1.919 | 0.169 | 0.036 | 5.39e-14 | 1.82e-09 |
| Malignant epithelial | C6 DNA-r p ECs | SERPINA5 | 1.916 | 0.218 | 0.061 | 1.3e-12 | 4.37e-08 |
| Malignant epithelial | C6 DNA-r p ECs | NRP1 | 1.912 | 0.202 | 0.047 | 1.88e-14 | 6.35e-10 |
| Malignant epithelial | C6 DNA-r p ECs | PSMC3IP | 1.91 | 0.129 | 0.034 | 2.8e-08 | 0.000945 |
| Malignant epithelial | C6 DNA-r p ECs | HELLS | 1.896 | 0.484 | 0.157 | 1.35e-23 | 4.54e-19 |
| Malignant epithelial | C6 DNA-r p ECs | MMP28 | 1.893 | 0.121 | 0.034 | 2.47e-07 | 0.00833 |
| Malignant epithelial | C6 DNA-r p ECs | RP5-884M6.1 | 1.889 | 0.153 | 0.036 | 2.22e-11 | 7.47e-07 |
| Malignant epithelial | C6 DNA-r p ECs | CDK2 | 1.881 | 0.323 | 0.074 | 3.33e-23 | 1.12e-18 |
| Malignant epithelial | C6 DNA-r p ECs | RAD51AP1 | 1.879 | 0.331 | 0.092 | 2.25e-18 | 7.57e-14 |
| Malignant epithelial | C6 DNA-r p ECs | MORC4 | 1.876 | 0.29 | 0.074 | 6.99e-19 | 2.36e-14 |
| Malignant epithelial | C6 DNA-r p ECs | RFC2 | 1.872 | 0.581 | 0.177 | 3.17e-31 | 1.07e-26 |
| Malignant epithelial | C6 DNA-r p ECs | PRR5L | 1.871 | 0.25 | 0.063 | 5.41e-16 | 1.82e-11 |
| Malignant epithelial | C6 DNA-r p ECs | SLCO4A1-AS1 | 1.863 | 0.395 | 0.119 | 7.22e-20 | 2.43e-15 |
| Malignant epithelial | C6 DNA-r p ECs | RFC4 | 1.863 | 0.315 | 0.091 | 2.81e-17 | 9.46e-13 |
| Malignant epithelial | C6 DNA-r p ECs | SMC2 | 1.861 | 0.444 | 0.142 | 1.18e-21 | 3.97e-17 |
| Malignant epithelial | C6 DNA-r p ECs | ST3GAL4 | 1.86 | 0.331 | 0.099 | 1.96e-17 | 6.61e-13 |
| Malignant epithelial | C6 DNA-r p ECs | AURKB | 1.854 | 0.339 | 0.084 | 1.8e-22 | 6.05e-18 |
| Malignant epithelial | C6 DNA-r p ECs | LAMB1 | 1.847 | 0.137 | 0.034 | 1.89e-09 | 6.36e-05 |

| compartment | subcluster | gene | avg_log2FC | pct.1 | pct.2 | p_val | p_val_adj |
|---|---|---|---|---|---|---|---|
| Malignant epithelial | C6 DNA-r p ECs | UHRF1 | 1.84 | 0.274 | 0.067 | 1.61e-18 | 5.42e-14 |
| Malignant epithelial | C6 DNA-r p ECs | MCM7 | 1.837 | 0.645 | 0.272 | 1.22e-25 | 4.110000000000001e-21 |
| Malignant epithelial | C6 DNA-r p ECs | MXD3 | 1.837 | 0.282 | 0.072 | 9.26e-18 | 3.12e-13 |
| Malignant epithelial | C6 DNA-r p ECs | DHDH | 1.829 | 0.121 | 0.023 | 1.19e-11 | 4.01e-07 |
| Malignant epithelial | C6 DNA-r p ECs | RNASEH2A | 1.827 | 0.589 | 0.2 | 2.08e-27 | 7.000000000000001e-23 |
| Malignant epithelial | C6 DNA-r p ECs | CHAF1B | 1.827 | 0.105 | 0.027 | 4.36e-07 | 0.0147 |
| Malignant epithelial | C6 DNA-r p ECs | LINC01137 | 1.826 | 0.129 | 0.035 | 6.3e-08 | 0.00212 |
| Malignant epithelial | C6 DNA-r p ECs | CD82 | 1.823 | 0.79 | 0.365 | 1.35e-26 | 4.540000000000001e-22 |
| Malignant epithelial | C6 DNA-r p ECs | HMGB2 | 1.817 | 0.863 | 0.489 | 1.65e-32 | 5.55e-28 |
| Malignant epithelial | C6 DNA-r p ECs | LMNB1 | 1.81 | 0.427 | 0.113 | 2.51e-26 | 8.47e-22 |
| Malignant epithelial | C6 DNA-r p ECs | PSCA | 1.81 | 0.121 | 0.022 | 1.4e-12 | 4.7e-08 |
| Malignant epithelial | C6 DNA-r p ECs | TACC3 | 1.809 | 0.258 | 0.058 | 3.59e-19 | 1.21e-14 |
| Malignant epithelial | C6 DNA-r p ECs | MDC1 | 1.807 | 0.169 | 0.038 | 7.88e-13 | 2.65e-08 |
| Malignant epithelial | C6 DNA-r p ECs | FANCD2 | 1.805 | 0.113 | 0.031 | 3.83e-07 | 0.0129 |
| Malignant epithelial | C6 DNA-r p ECs | HRASLS2 | 1.8 | 0.129 | 0.035 | 4.8e-08 | 0.00162 |
| Malignant epithelial | C6 DNA-r p ECs | CDCA4 | 1.799 | 0.435 | 0.134 | 1.21e-21 | 4.09e-17 |
| Malignant epithelial | C6 DNA-r p ECs | ALG1L | 1.797 | 0.274 | 0.08 | 2.42e-14 | 8.15e-10 |
| Malignant epithelial | C6 DNA-r p ECs | ULBP2 | 1.785 | 0.177 | 0.04 | 1.92e-13 | 6.46e-09 |
| Malignant epithelial | C6 DNA-r p ECs | L1CAM | 1.783 | 0.153 | 0.042 | 4.92e-09 | 0.000166 |
| Malignant epithelial | C6 DNA-r p ECs | PARP16 | 1.78 | 0.161 | 0.051 | 5.67e-08 | 0.00191 |
| Malignant epithelial | C6 DNA-r p ECs | ICMT | 1.779 | 0.46 | 0.127 | 7.38e-26 | 2.49e-21 |
| Malignant epithelial | C6 DNA-r p ECs | PDLIM7 | 1.777 | 0.532 | 0.258 | 1.12e-13 | 3.78e-09 |
| Malignant epithelial | C6 DNA-r p ECs | UNC5B-AS1 | 1.77 | 0.242 | 0.064 | 1.22e-14 | 4.11e-10 |
| Malignant epithelial | C6 DNA-r p ECs | ARHGAP10 | 1.77 | 0.185 | 0.049 | 1.38e-11 | 4.64e-07 |
| Malignant epithelial | C6 DNA-r p ECs | CDCA8 | 1.767 | 0.25 | 0.053 | 3.77e-20 | 1.27e-15 |
| Malignant epithelial | C6 DNA-r p ECs | RRM1 | 1.767 | 0.315 | 0.111 | 1.34e-12 | 4.5e-08 |
| Malignant epithelial | C6 DNA-r p ECs | CD109 | 1.764 | 0.266 | 0.072 | 1.52e-15 | 5.13e-11 |
| Malignant epithelial | C6 DNA-r p ECs | DLEU2 | 1.764 | 0.169 | 0.047 | 1.24e-09 | 4.19e-05 |
| Malignant epithelial | C6 DNA-r p ECs | MSH2 | 1.763 | 0.194 | 0.069 | 1.64e-07 | 0.00554 |
| Malignant epithelial | C6 DNA-r p ECs | ORC6 | 1.761 | 0.419 | 0.102 | 2.11e-27 | 7.119999999999999e-23 |
| Malignant epithelial | C6 DNA-r p ECs | RMI2 | 1.747 | 0.339 | 0.084 | 9.93e-22 | 3.35e-17 |
| Malignant epithelial | C6 DNA-r p ECs | HIST1H1D | 1.734 | 0.282 | 0.075 | 1.69e-16 | 5.7e-12 |
| Malignant epithelial | C6 DNA-r p ECs | EPHX3 | 1.734 | 0.226 | 0.072 | 2.76e-10 | 9.3e-06 |
| Malignant epithelial | C6 DNA-r p ECs | TSKU | 1.729 | 0.306 | 0.084 | 1.35e-17 | 4.56e-13 |
| Malignant epithelial | C6 DNA-r p ECs | OIP5 | 1.725 | 0.274 | 0.064 | 2.81e-19 | 9.47e-15 |
| Malignant epithelial | C6 DNA-r p ECs | TNFRSF11A | 1.719 | 0.153 | 0.035 | 1.69e-11 | 5.68e-07 |
| Malignant epithelial | C6 DNA-r p ECs | APITD1 | 1.717 | 0.266 | 0.078 | 3.83e-14 | 1.29e-09 |
| Malignant epithelial | C6 DNA-r p ECs | C4orf46 | 1.712 | 0.185 | 0.043 | 2.52e-13 | 8.49e-09 |
| Malignant epithelial | C6 DNA-r p ECs | TMEM106C | 1.711 | 0.742 | 0.322 | 1.32e-25 | 4.44e-21 |
| Malignant epithelial | C6 DNA-r p ECs | POLE | 1.71 | 0.161 | 0.042 | 4.75e-10 | 1.6e-05 |
| Malignant epithelial | C6 DNA-r p ECs | LRRC26 | 1.706 | 0.105 | 0.023 | 6.21e-09 | 0.000209 |
| Malignant epithelial | C6 DNA-r p ECs | ORC3 | 1.704 | 0.371 | 0.119 | 9.73e-17 | 3.28e-12 |
| Malignant epithelial | C6 DNA-r p ECs | RP11-362F19.1 | 1.703 | 0.234 | 0.065 | 6.91e-13 | 2.33e-08 |
| Malignant epithelial | C6 DNA-r p ECs | STARD5 | 1.703 | 0.121 | 0.029 | 7.59e-09 | 0.000256 |
| Malignant epithelial | C6 DNA-r p ECs | C1orf112 | 1.7 | 0.113 | 0.029 | 1.51e-07 | 0.00509 |
| Malignant epithelial | C6 DNA-r p ECs | PPP1R14C | 1.699 | 0.145 | 0.033 | 7.07e-11 | 2.38e-06 |
| Malignant epithelial | C6 DNA-r p ECs | EZH2 | 1.691 | 0.403 | 0.145 | 4.2e-16 | 1.42e-11 |
| Malignant epithelial | C6 DNA-r p ECs | MCM2 | 1.686 | 0.266 | 0.075 | 1.53e-14 | 5.16e-10 |
| Malignant epithelial | C6 DNA-r p ECs | PRADC1 | 1.678 | 0.323 | 0.109 | 1.41e-13 | 4.75e-09 |
| Malignant epithelial | C6 DNA-r p ECs | POC1A | 1.678 | 0.25 | 0.075 | 2.02e-12 | 6.79e-08 |
| Malignant epithelial | C6 DNA-r p ECs | FDXR | 1.675 | 0.323 | 0.085 | 5.01e-19 | 1.69e-14 |
| Malignant epithelial | C6 DNA-r p ECs | FAM46B | 1.674 | 0.177 | 0.051 | 6.97e-10 | 2.35e-05 |
| Malignant epithelial | C6 DNA-r p ECs | DONSON | 1.669 | 0.234 | 0.051 | 1.85e-17 | 6.25e-13 |
| Malignant epithelial | C6 DNA-r p ECs | CD163L1 | 1.667 | 0.516 | 0.176 | 4.939999999999999e-23 | 1.66e-18 |
| Malignant epithelial | C6 DNA-r p ECs | AGTRAP | 1.666 | 0.621 | 0.258 | 3.43e-21 | 1.15e-16 |
| Malignant epithelial | C6 DNA-r p ECs | SUV39H1 | 1.662 | 0.145 | 0.036 | 7.08e-10 | 2.39e-05 |
| Malignant epithelial | C6 DNA-r p ECs | FAM64A | 1.661 | 0.121 | 0.029 | 1.1e-08 | 0.00037 |
| Malignant epithelial | C6 DNA-r p ECs | FANCI | 1.658 | 0.242 | 0.061 | 2.63e-15 | 8.88e-11 |
| Malignant epithelial | C6 DNA-r p ECs | LMCD1 | 1.657 | 0.177 | 0.047 | 5.4e-11 | 1.82e-06 |
| Malignant epithelial | C6 DNA-r p ECs | TESC | 1.655 | 0.524 | 0.208 | 1.48e-18 | 5e-14 |
| Malignant epithelial | C6 DNA-r p ECs | RAD18 | 1.655 | 0.177 | 0.051 | 1.67e-09 | 5.62e-05 |
| Malignant epithelial | C6 DNA-r p ECs | NRM | 1.653 | 0.339 | 0.122 | 3e-13 | 1.01e-08 |
| Malignant epithelial | C6 DNA-r p ECs | KCNK9 | 1.651 | 0.25 | 0.065 | 2.99e-15 | 1.01e-10 |
| Malignant epithelial | C6 DNA-r p ECs | NCAPD3 | 1.646 | 0.242 | 0.069 | 3.35e-13 | 1.13e-08 |

| compartment | subcluster | gene | avg_log2FC | pct.1 | pct.2 | p_val | p_val_adj |
|---|---|---|---|---|---|---|---|
| Malignant epithelial | C6 DNA-r p ECs | RP11-152N13.5 | 1.641 | 0.226 | 0.056 | 1.78e-14 | 6e-10 |
| Malignant epithelial | C6 DNA-r p ECs | KIF11 | 1.638 | 0.185 | 0.045 | 1.69e-12 | 5.68e-08 |
| Malignant epithelial | C6 DNA-r p ECs | RBBP8 | 1.636 | 0.194 | 0.062 | 5.83e-09 | 0.000196 |
| Malignant epithelial | C6 DNA-r p ECs | MCM3 | 1.635 | 0.411 | 0.149 | 4.68e-16 | 1.58e-11 |
| Malignant epithelial | C6 DNA-r p ECs | RCCD1 | 1.632 | 0.363 | 0.113 | 8.11e-18 | 2.73e-13 |
| Malignant epithelial | C6 DNA-r p ECs | FAM174B | 1.631 | 0.194 | 0.044 | 4.12e-14 | 1.39e-09 |
| Malignant epithelial | C6 DNA-r p ECs | RP11-443B7.1 | 1.629 | 0.145 | 0.037 | 1.49e-09 | 5.01e-05 |
| Malignant epithelial | C6 DNA-r p ECs | AFAP1-AS1 | 1.624 | 0.202 | 0.054 | 5.69e-12 | 1.92e-07 |
| Malignant epithelial | C6 DNA-r p ECs | TRIM29 | 1.621 | 0.54 | 0.164 | 4.85e-28 | 1.63e-23 |
| Malignant epithelial | C6 DNA-r p ECs | LRRC45 | 1.619 | 0.25 | 0.072 | 3.06e-13 | 1.03e-08 |
| Malignant epithelial | C6 DNA-r p ECs | SYNE1 | 1.616 | 0.121 | 0.03 | 2.21e-08 | 0.000745 |
| Malignant epithelial | C6 DNA-r p ECs | C1QTNF9B-AS1 | 1.613 | 0.105 | 0.027 | 3.81e-07 | 0.0128 |
| Malignant epithelial | C6 DNA-r p ECs | RP6-65G23.3 | 1.607 | 0.202 | 0.065 | 2.35e-09 | 7.9e-05 |
| Malignant epithelial | C6 DNA-r p ECs | USP37 | 1.6 | 0.153 | 0.04 | 8.43e-10 | 2.84e-05 |
| Malignant epithelial | C6 DNA-r p ECs | RP11-150O12.3 | 1.598 | 0.161 | 0.045 | 3.21e-09 | 0.000108 |
| Malignant epithelial | C6 DNA-r p ECs | PRIMPOL | 1.598 | 0.145 | 0.041 | 3.45e-08 | 0.00116 |
| Malignant epithelial | C6 DNA-r p ECs | PARD6A | 1.595 | 0.282 | 0.075 | 3.06e-16 | 1.03e-11 |
| Malignant epithelial | C6 DNA-r p ECs | IL17RB | 1.594 | 0.226 | 0.067 | 1.99e-11 | 6.69e-07 |
| Malignant epithelial | C6 DNA-r p ECs | DHRS13 | 1.594 | 0.145 | 0.038 | 2.7e-09 | 9.11e-05 |
| Malignant epithelial | C6 DNA-r p ECs | CTSV | 1.586 | 0.484 | 0.149 | 1.73e-24 | 5.84e-20 |
| Malignant epithelial | C6 DNA-r p ECs | CBR3 | 1.585 | 0.395 | 0.123 | 3.45e-18 | 1.16e-13 |
| Malignant epithelial | C6 DNA-r p ECs | RMI1 | 1.585 | 0.145 | 0.04 | 1.06e-08 | 0.000357 |
| Malignant epithelial | C6 DNA-r p ECs | ST3GAL1 | 1.584 | 0.395 | 0.135 | 2.15e-16 | 7.24e-12 |
| Malignant epithelial | C6 DNA-r p ECs | YBX2 | 1.584 | 0.137 | 0.039 | 7.03e-08 | 0.00237 |
| Malignant epithelial | C6 DNA-r p ECs | EFNA5 | 1.583 | 0.137 | 0.034 | 2.09e-09 | 7.03e-05 |
| Malignant epithelial | C6 DNA-r p ECs | CTSD | 1.582 | 0.944 | 0.774 | 1.02e-16 | 3.45e-12 |
| Malignant epithelial | C6 DNA-r p ECs | TTF2 | 1.581 | 0.153 | 0.044 | 1.81e-08 | 0.000611 |
| Malignant epithelial | C6 DNA-r p ECs | ATG16L2 | 1.581 | 0.129 | 0.038 | 4.6e-07 | 0.0155 |
| Malignant epithelial | C6 DNA-r p ECs | MAD2L1 | 1.58 | 0.581 | 0.193 | 6.690000000000001e-27 | 2.25e-22 |
| Malignant epithelial | C6 DNA-r p ECs | CCNA2 | 1.574 | 0.315 | 0.071 | 1.14e-22 | 3.84e-18 |
| Malignant epithelial | C6 DNA-r p ECs | SCLY | 1.573 | 0.153 | 0.046 | 4.28e-08 | 0.00144 |
| Malignant epithelial | C6 DNA-r p ECs | AQP3 | 1.57 | 0.137 | 0.041 | 2.34e-07 | 0.00789 |
| Malignant epithelial | C6 DNA-r p ECs | POLD3 | 1.557 | 0.234 | 0.06 | 2.39e-14 | 8.06e-10 |
| Malignant epithelial | C6 DNA-r p ECs | AGFG2 | 1.555 | 0.274 | 0.073 | 1e-15 | 3.38e-11 |
| Malignant epithelial | C6 DNA-r p ECs | TRNP1 | 1.547 | 0.21 | 0.052 | 1.25e-13 | 4.2e-09 |
| Malignant epithelial | C6 DNA-r p ECs | FAM83D | 1.544 | 0.218 | 0.077 | 8.28e-09 | 0.000279 |
| Malignant epithelial | C6 DNA-r p ECs | ME3 | 1.543 | 0.234 | 0.058 | 8.02e-15 | 2.7e-10 |
| Malignant epithelial | C6 DNA-r p ECs | HAUS8 | 1.543 | 0.194 | 0.049 | 3.96e-12 | 1.34e-07 |
| Malignant epithelial | C6 DNA-r p ECs | PODXL | 1.543 | 0.282 | 0.095 | 8.4e-12 | 2.83e-07 |
| Malignant epithelial | C6 DNA-r p ECs | SMIM5 | 1.541 | 0.185 | 0.066 | 2.32e-07 | 0.00782 |
| Malignant epithelial | C6 DNA-r p ECs | PBK | 1.539 | 0.177 | 0.05 | 7.78e-10 | 2.62e-05 |
| Malignant epithelial | C6 DNA-r p ECs | CORO2A | 1.538 | 0.419 | 0.123 | 8.430000000000001e-21 | 2.84e-16 |
| Malignant epithelial | C6 DNA-r p ECs | MCM4 | 1.537 | 0.532 | 0.179 | 3.9199999999999999e-23 | 1.32e-18 |
| Malignant epithelial | C6 DNA-r p ECs | LIG1 | 1.537 | 0.274 | 0.07 | 8.91e-17 | 3e-12 |
| Malignant epithelial | C6 DNA-r p ECs | TUBA1B | 1.535 | 0.984 | 0.74 | 3.62e-31 | 1.22e-26 |
| Malignant epithelial | C6 DNA-r p ECs | KLK6 | 1.528 | 0.831 | 0.543 | 6.98e-17 | 2.35e-12 |
| Malignant epithelial | C6 DNA-r p ECs | TMEM154 | 1.528 | 0.194 | 0.065 | 2.23e-08 | 0.000751 |
| Malignant epithelial | C6 DNA-r p ECs | GCHFR | 1.526 | 0.71 | 0.305 | 4.97e-25 | 1.67e-20 |
| Malignant epithelial | C6 DNA-r p ECs | UNG | 1.526 | 0.379 | 0.166 | 3.02e-11 | 1.02e-06 |
| Malignant epithelial | C6 DNA-r p ECs | WDR34 | 1.525 | 0.774 | 0.403 | 1.5e-26 | 5.05e-22 |
| Malignant epithelial | C6 DNA-r p ECs | CENPQ | 1.525 | 0.185 | 0.054 | 1e-09 | 3.37e-05 |
| Malignant epithelial | C6 DNA-r p ECs | PLEKHS1 | 1.522 | 0.21 | 0.062 | 1.42e-10 | 4.78e-06 |
| Malignant epithelial | C6 DNA-r p ECs | DUT | 1.521 | 0.847 | 0.583 | 1.28e-23 | 4.3e-19 |
| Malignant epithelial | C6 DNA-r p ECs | DMRT2 | 1.519 | 0.121 | 0.031 | 6.04e-08 | 0.00204 |
| Malignant epithelial | C6 DNA-r p ECs | IL20RB | 1.516 | 0.194 | 0.055 | 1.37e-10 | 4.6e-06 |
| Malignant epithelial | C6 DNA-r p ECs | KIAA1841 | 1.509 | 0.161 | 0.047 | 1.81e-08 | 0.000611 |
| Malignant epithelial | C6 DNA-r p ECs | CENPN | 1.506 | 0.629 | 0.252 | 1.57e-22 | 5.28e-18 |
| Malignant epithelial | C6 DNA-r p ECs | ZWILCH | 1.506 | 0.234 | 0.08 | 1.1e-09 | 3.72e-05 |
| Malignant epithelial | C6 DNA-r p ECs | ATAD3A | 1.503 | 0.395 | 0.122 | 2.03e-18 | 6.83e-14 |
| Malignant epithelial | C6 DNA-r p ECs | MCM8 | 1.502 | 0.161 | 0.052 | 1.67e-07 | 0.00562 |
| Malignant epithelial | C6 DNA-r p ECs | PSAT1 | 1.501 | 0.339 | 0.121 | 1.65e-12 | 5.55e-08 |
| Malignant epithelial | C6 DNA-r p ECs | C4BPB | 1.498 | 0.202 | 0.052 | 2.13e-12 | 7.18e-08 |
| Malignant epithelial | C6 DNA-r p ECs | KCNS3 | 1.496 | 0.145 | 0.045 | 3.35e-07 | 0.0113 |
| Malignant epithelial | C6 DNA-r p ECs | C4orf27 | 1.492 | 0.298 | 0.098 | 7.01e-13 | 2.36e-08 |

| compartment | subcluster | gene | avg_log2FC | pct.1 | pct.2 | p_val | p_val_adj |
|---|---|---|---|---|---|---|---|
| Malignant epithelial | C6 DNA-r p ECs | RNASE6 | 1.492 | 0.113 | 0.027 | 2.88e-08 | 0.000972 |
| Malignant epithelial | C6 DNA-r p ECs | SCEL | 1.491 | 0.339 | 0.084 | 5.04e-22 | 1.7e-17 |
| Malignant epithelial | C6 DNA-r p ECs | CENPO | 1.49 | 0.113 | 0.031 | 5.75e-07 | 0.0194 |
| Malignant epithelial | C6 DNA-r p ECs | RHOBTB3 | 1.488 | 0.581 | 0.192 | 2.74e-25 | 9.240000000000001e-21 |
| Malignant epithelial | C6 DNA-r p ECs | TBC1D2 | 1.487 | 0.194 | 0.055 | 1.6e-10 | 5.38e-06 |
| Malignant epithelial | C6 DNA-r p ECs | DTYMK | 1.486 | 0.702 | 0.296 | 1.02e-24 | 3.45e-20 |
| Malignant epithelial | C6 DNA-r p ECs | CD14 | 1.483 | 0.242 | 0.074 | 1.13e-11 | 3.82e-07 |
| Malignant epithelial | C6 DNA-r p ECs | EHD2 | 1.483 | 0.21 | 0.066 | 1.14e-09 | 3.83e-05 |
| Malignant epithelial | C6 DNA-r p ECs | CDC23 | 1.482 | 0.258 | 0.077 | 8.47e-13 | 2.86e-08 |
| Malignant epithelial | C6 DNA-r p ECs | SNCG | 1.478 | 0.226 | 0.097 | 1.27e-06 | 0.0427 |
| Malignant epithelial | C6 DNA-r p ECs | RAD51C | 1.473 | 0.387 | 0.125 | 2.08e-17 | 7.01e-13 |
| Malignant epithelial | C6 DNA-r p ECs | DEFB1 | 1.471 | 0.492 | 0.187 | 1.48e-16 | 4.98e-12 |
| Malignant epithelial | C6 DNA-r p ECs | CH17-360D5.2 | 1.471 | 0.323 | 0.123 | 1.87e-10 | 6.3e-06 |
| Malignant epithelial | C6 DNA-r p ECs | WDR54 | 1.47 | 0.54 | 0.188 | 7.99e-22 | 2.69e-17 |
| Malignant epithelial | C6 DNA-r p ECs | HJURP | 1.469 | 0.218 | 0.057 | 2.41e-13 | 8.12e-09 |
| Malignant epithelial | C6 DNA-r p ECs | FAM120C | 1.468 | 0.137 | 0.038 | 5.28e-08 | 0.00178 |
| Malignant epithelial | C6 DNA-r p ECs | SLC41A2 | 1.467 | 0.137 | 0.042 | 3.86e-07 | 0.013 |
| Malignant epithelial | C6 DNA-r p ECs | MIB2 | 1.466 | 0.524 | 0.179 | 1.83e-20 | 6.16e-16 |
| Malignant epithelial | C6 DNA-r p ECs | PHF19 | 1.461 | 0.556 | 0.227 | 3.11e-18 | 1.05e-13 |
| Malignant epithelial | C6 DNA-r p ECs | FAM110C | 1.46 | 0.258 | 0.086 | 1.18e-10 | 3.99e-06 |
| Malignant epithelial | C6 DNA-r p ECs | SLC25A4 | 1.459 | 0.444 | 0.158 | 4.06e-17 | 1.37e-12 |
| Malignant epithelial | C6 DNA-r p ECs | MCM5 | 1.459 | 0.379 | 0.131 | 6.29e-15 | 2.12e-10 |
| Malignant epithelial | C6 DNA-r p ECs | SLC22A3 | 1.45 | 0.194 | 0.064 | 2.01e-08 | 0.000676 |
| Malignant epithelial | C6 DNA-r p ECs | FANCL | 1.439 | 0.387 | 0.131 | 7.94e-16 | 2.67e-11 |
| Malignant epithelial | C6 DNA-r p ECs | HIST1H4C | 1.436 | 0.677 | 0.447 | 1.74e-09 | 5.86e-05 |
| Malignant epithelial | C6 DNA-r p ECs | SFXN5 | 1.434 | 0.137 | 0.044 | 1.16e-06 | 0.0391 |
| Malignant epithelial | C6 DNA-r p ECs | PROCR | 1.43 | 0.556 | 0.245 | 1.42e-14 | 4.78e-10 |
| Malignant epithelial | C6 DNA-r p ECs | FAAP24 | 1.43 | 0.129 | 0.036 | 1.51e-07 | 0.00508 |
| Malignant epithelial | C6 DNA-r p ECs | PRC1 | 1.427 | 0.435 | 0.118 | 1.01e-24 | 3.41e-20 |
| Malignant epithelial | C6 DNA-r p ECs | SLF1 | 1.427 | 0.194 | 0.066 | 3.64e-08 | 0.00123 |
| Malignant epithelial | C6 DNA-r p ECs | ZNF213 | 1.426 | 0.129 | 0.037 | 2.37e-07 | 0.00797 |
| Malignant epithelial | C6 DNA-r p ECs | RELL2 | 1.423 | 0.153 | 0.052 | 1.33e-06 | 0.0449 |
| Malignant epithelial | C6 DNA-r p ECs | BUB1B | 1.422 | 0.113 | 0.029 | 1.42e-07 | 0.00478 |
| Malignant epithelial | C6 DNA-r p ECs | FIGNL1 | 1.421 | 0.121 | 0.032 | 9.23e-08 | 0.00311 |
| Malignant epithelial | C6 DNA-r p ECs | SMIM6 | 1.419 | 0.379 | 0.146 | 7.72e-13 | 2.6e-08 |
| Malignant epithelial | C6 DNA-r p ECs | RLN2 | 1.406 | 0.137 | 0.034 | 3.34e-09 | 0.000113 |
| Malignant epithelial | C6 DNA-r p ECs | HIST1H2BJ | 1.406 | 0.202 | 0.073 | 1.04e-07 | 0.00349 |
| Malignant epithelial | C6 DNA-r p ECs | PAQR4 | 1.399 | 0.427 | 0.164 | 2e-14 | 6.73e-10 |
| Malignant epithelial | C6 DNA-r p ECs | ACAT2 | 1.399 | 0.387 | 0.14 | 2.2e-14 | 7.4e-10 |
| Malignant epithelial | C6 DNA-r p ECs | TRIM22 | 1.398 | 0.306 | 0.088 | 1.37e-15 | 4.63e-11 |
| Malignant epithelial | C6 DNA-r p ECs | DNMT1 | 1.397 | 0.532 | 0.224 | 1.51e-16 | 5.07e-12 |
| Malignant epithelial | C6 DNA-r p ECs | BARX2 | 1.396 | 0.194 | 0.064 | 2.34e-08 | 0.000789 |
| Malignant epithelial | C6 DNA-r p ECs | TOP2A | 1.389 | 0.371 | 0.1 | 4.16e-21 | 1.4e-16 |
| Malignant epithelial | C6 DNA-r p ECs | POLM | 1.387 | 0.347 | 0.108 | 6.7e-16 | 2.26e-11 |
| Malignant epithelial | C6 DNA-r p ECs | DOK1 | 1.386 | 0.169 | 0.061 | 1.22e-06 | 0.0412 |
| Malignant epithelial | C6 DNA-r p ECs | RPA3 | 1.385 | 0.944 | 0.664 | 4.069999999999999e-30 | 1.37e-25 |
| Malignant epithelial | C6 DNA-r p ECs | CTNNAL1 | 1.385 | 0.395 | 0.136 | 1.78e-15 | 6e-11 |
| Malignant epithelial | C6 DNA-r p ECs | SYTL1 | 1.384 | 0.54 | 0.222 | 4.01e-16 | 1.35e-11 |
| Malignant epithelial | C6 DNA-r p ECs | BIRC5 | 1.382 | 0.556 | 0.182 | 1.79e-24 | 6.04e-20 |
| Malignant epithelial | C6 DNA-r p ECs | NEK2 | 1.382 | 0.202 | 0.053 | 4.51e-12 | 1.52e-07 |
| Malignant epithelial | C6 DNA-r p ECs | CDH3 | 1.38 | 0.339 | 0.157 | 9.14e-08 | 0.00308 |
| Malignant epithelial | C6 DNA-r p ECs | B9D1 | 1.379 | 0.331 | 0.103 | 1.26e-14 | 4.26e-10 |
| Malignant epithelial | C6 DNA-r p ECs | ASNS | 1.378 | 0.331 | 0.135 | 8.17e-10 | 2.75e-05 |
| Malignant epithelial | C6 DNA-r p ECs | C9orf40 | 1.375 | 0.258 | 0.087 | 1.18e-10 | 3.96e-06 |
| Malignant epithelial | C6 DNA-r p ECs | MMEL1 | 1.369 | 0.266 | 0.081 | 9.15e-13 | 3.08e-08 |
| Malignant epithelial | C6 DNA-r p ECs | VRK1 | 1.367 | 0.444 | 0.133 | 8.68e-21 | 2.92e-16 |
| Malignant epithelial | C6 DNA-r p ECs | ALDOB | 1.365 | 0.105 | 0.023 | 9.18e-09 | 0.000309 |
| Malignant epithelial | C6 DNA-r p ECs | BLVRA | 1.356 | 0.452 | 0.174 | 2.93e-15 | 9.89e-11 |
| Malignant epithelial | C6 DNA-r p ECs | FAM111A | 1.356 | 0.379 | 0.151 | 4.19e-12 | 1.41e-07 |
| Malignant epithelial | C6 DNA-r p ECs | CLDN2 | 1.354 | 0.169 | 0.039 | 2.81e-12 | 9.47e-08 |
| Malignant epithelial | C6 DNA-r p ECs | COL4A2 | 1.354 | 0.129 | 0.039 | 6.74e-07 | 0.0227 |
| Malignant epithelial | C6 DNA-r p ECs | IFI27L1 | 1.351 | 0.75 | 0.308 | 5.46e-25 | 1.84e-20 |
| Malignant epithelial | C6 DNA-r p ECs | CCNF | 1.351 | 0.105 | 0.027 | 4.89e-07 | 0.0165 |
| Malignant epithelial | C6 DNA-r p ECs | THEM6 | 1.349 | 0.492 | 0.226 | 3.31e-13 | 1.11e-08 |

| compartment | subcluster | gene | avg_log2FC | pct.1 | pct.2 | p_val | p_val_adj |
|---|---|---|---|---|---|---|---|
| Malignant epithelial | C6 DNA-r p ECs | GLIPR2 | 1.346 | 0.145 | 0.039 | 1.06e-08 | 0.000359 |
| Malignant epithelial | C6 DNA-r p ECs | COPS3 | 1.344 | 0.411 | 0.154 | 4.02e-14 | 1.36e-09 |
| Malignant epithelial | C6 DNA-r p ECs | WRAP73 | 1.342 | 0.298 | 0.102 | 8.59e-12 | 2.9e-07 |
| Malignant epithelial | C6 DNA-r p ECs | RFWD3 | 1.342 | 0.226 | 0.07 | 1.34e-10 | 4.53e-06 |
| Malignant epithelial | C6 DNA-r p ECs | PANK2 | 1.337 | 0.419 | 0.164 | 6.64e-14 | 2.24e-09 |
| Malignant epithelial | C6 DNA-r p ECs | RARRES3 | 1.335 | 0.935 | 0.628 | 1.89e-18 | 6.38e-14 |
| Malignant epithelial | C6 DNA-r p ECs | KIF23 | 1.335 | 0.218 | 0.062 | 1.46e-11 | 4.91e-07 |
| Malignant epithelial | C6 DNA-r p ECs | SEMA7A | 1.33 | 0.202 | 0.069 | 3.63e-08 | 0.00122 |
| Malignant epithelial | C6 DNA-r p ECs | CPTP | 1.329 | 0.282 | 0.12 | 3.85e-08 | 0.0013 |
| Malignant epithelial | C6 DNA-r p ECs | PRSS23 | 1.322 | 0.484 | 0.253 | 5.92e-09 | 0.0002 |
| Malignant epithelial | C6 DNA-r p ECs | LCLAT1 | 1.321 | 0.185 | 0.069 | 1.17e-06 | 0.0394 |
| Malignant epithelial | C6 DNA-r p ECs | ETV7 | 1.318 | 0.444 | 0.137 | 1.23e-19 | 4.15e-15 |
| Malignant epithelial | C6 DNA-r p ECs | EVA1C | 1.318 | 0.242 | 0.084 | 2.4e-09 | 8.08e-05 |
| Malignant epithelial | C6 DNA-r p ECs | RAD51D | 1.318 | 0.161 | 0.055 | 6.72e-07 | 0.0226 |
| Malignant epithelial | C6 DNA-r p ECs | IGFBP2 | 1.317 | 0.669 | 0.219 | 5.070000000000001e-29 | 1.71e-24 |
| Malignant epithelial | C6 DNA-r p ECs | MFI2 | 1.316 | 0.435 | 0.15 | 4.61e-17 | 1.55e-12 |
| Malignant epithelial | C6 DNA-r p ECs | HAUS1 | 1.316 | 0.363 | 0.135 | 1.69e-12 | 5.7e-08 |
| Malignant epithelial | C6 DNA-r p ECs | PARPBP | 1.316 | 0.177 | 0.064 | 1.21e-06 | 0.0406 |
| Malignant epithelial | C6 DNA-r p ECs | CKAP2 | 1.315 | 0.363 | 0.109 | 1.16e-17 | 3.9e-13 |
| Malignant epithelial | C6 DNA-r p ECs | ACYP1 | 1.311 | 0.548 | 0.223 | 1.17e-17 | 3.96e-13 |
| Malignant epithelial | C6 DNA-r p ECs | MFSD13A | 1.311 | 0.202 | 0.072 | 8.52e-08 | 0.00287 |
| Malignant epithelial | C6 DNA-r p ECs | DDC | 1.31 | 0.282 | 0.088 | 1.42e-12 | 4.79e-08 |
| Malignant epithelial | C6 DNA-r p ECs | PCOLCE | 1.309 | 0.113 | 0.032 | 1.43e-06 | 0.0481 |
| Malignant epithelial | C6 DNA-r p ECs | PSIP1 | 1.303 | 0.653 | 0.303 | 6.82e-18 | 2.3e-13 |
| Malignant epithelial | C6 DNA-r p ECs | UBE2C | 1.302 | 0.798 | 0.373 | 1.54e-23 | 5.19e-19 |
| Malignant epithelial | C6 DNA-r p ECs | MILR1 | 1.302 | 0.169 | 0.049 | 5.91e-09 | 0.000199 |
| Malignant epithelial | C6 DNA-r p ECs | MTHFD1 | 1.295 | 0.371 | 0.146 | 9.75e-12 | 3.28e-07 |
| Malignant epithelial | C6 DNA-r p ECs | NUP85 | 1.293 | 0.282 | 0.1 | 1.89e-10 | 6.38e-06 |
| Malignant epithelial | C6 DNA-r p ECs | EIF4EBP1 | 1.292 | 0.968 | 0.694 | 3.32e-28 | 1.12e-23 |
| Malignant epithelial | C6 DNA-r p ECs | B3GNTL1 | 1.292 | 0.226 | 0.072 | 4.52e-10 | 1.52e-05 |
| Malignant epithelial | C6 DNA-r p ECs | TMEM51 | 1.291 | 0.29 | 0.104 | 1.75e-10 | 5.89e-06 |
| Malignant epithelial | C6 DNA-r p ECs | TMEM80 | 1.289 | 0.323 | 0.128 | 5.63e-10 | 1.9e-05 |
| Malignant epithelial | C6 DNA-r p ECs | SAC3D1 | 1.288 | 0.508 | 0.209 | 1.42e-15 | 4.78e-11 |
| Malignant epithelial | C6 DNA-r p ECs | KAZALD1 | 1.286 | 0.234 | 0.082 | 5.17e-09 | 0.000174 |
| Malignant epithelial | C6 DNA-r p ECs | SLC29A2 | 1.286 | 0.194 | 0.063 | 1.33e-08 | 0.000447 |
| Malignant epithelial | C6 DNA-r p ECs | POLD2 | 1.283 | 0.75 | 0.411 | 4.02e-18 | 1.35e-13 |
| Malignant epithelial | C6 DNA-r p ECs | C2orf70 | 1.282 | 0.226 | 0.077 | 3.7e-09 | 0.000125 |
| Malignant epithelial | C6 DNA-r p ECs | ARHGAP6 | 1.281 | 0.234 | 0.092 | 2.64e-07 | 0.00891 |
| Malignant epithelial | C6 DNA-r p ECs | SDCBP2 | 1.279 | 0.629 | 0.265 | 3.38e-20 | 1.14e-15 |
| Malignant epithelial | C6 DNA-r p ECs | ORAI2 | 1.277 | 0.282 | 0.111 | 5.89e-09 | 0.000198 |
| Malignant epithelial | C6 DNA-r p ECs | SPTSSB | 1.276 | 0.105 | 0.028 | 1.42e-06 | 0.0477 |
| Malignant epithelial | C6 DNA-r p ECs | SUV39H2 | 1.276 | 0.121 | 0.036 | 1.43e-06 | 0.0482 |
| Malignant epithelial | C6 DNA-r p ECs | NEDD9 | 1.275 | 0.371 | 0.132 | 1.05e-13 | 3.55e-09 |
| Malignant epithelial | C6 DNA-r p ECs | FOXM1 | 1.273 | 0.315 | 0.106 | 1.16e-12 | 3.9e-08 |
| Malignant epithelial | C6 DNA-r p ECs | SMC4 | 1.272 | 0.71 | 0.289 | 6.66e-25 | 2.24e-20 |
| Malignant epithelial | C6 DNA-r p ECs | KALRN | 1.272 | 0.185 | 0.058 | 1.07e-08 | 0.000359 |
| Malignant epithelial | C6 DNA-r p ECs | CREB3L2 | 1.268 | 0.363 | 0.148 | 6.36e-11 | 2.14e-06 |
| Malignant epithelial | C6 DNA-r p ECs | ARL4D | 1.266 | 0.169 | 0.061 | 1.17e-06 | 0.0395 |
| Malignant epithelial | C6 DNA-r p ECs | SMC1A | 1.265 | 0.379 | 0.158 | 6.16e-11 | 2.08e-06 |
| Malignant epithelial | C6 DNA-r p ECs | RP11-54H7.4 | 1.265 | 0.556 | 0.36 | 2.33e-07 | 0.00784 |
| Malignant epithelial | C6 DNA-r p ECs | MDK | 1.264 | 0.984 | 0.81 | 2.06e-30 | 6.95e-26 |
| Malignant epithelial | C6 DNA-r p ECs | HPRT1 | 1.262 | 0.718 | 0.4 | 8.52e-16 | 2.87e-11 |
| Malignant epithelial | C6 DNA-r p ECs | RBPMS | 1.261 | 0.242 | 0.084 | 1.83e-09 | 6.17e-05 |
| Malignant epithelial | C6 DNA-r p ECs | CASC5 | 1.261 | 0.137 | 0.04 | 1.76e-07 | 0.00592 |
| Malignant epithelial | C6 DNA-r p ECs | ENO1 | 1.259 | 0.968 | 0.794 | 1.13e-28 | 3.8199999999999999e-24 |
| Malignant epithelial | C6 DNA-r p ECs | HERC6 | 1.254 | 0.169 | 0.052 | 4.13e-08 | 0.00139 |
| Malignant epithelial | C6 DNA-r p ECs | KLK11 | 1.252 | 0.492 | 0.204 | 2.15e-15 | 7.26e-11 |
| Malignant epithelial | C6 DNA-r p ECs | MAD2L2 | 1.252 | 0.371 | 0.153 | 1.12e-10 | 3.76e-06 |
| Malignant epithelial | C6 DNA-r p ECs | PIDD1 | 1.252 | 0.137 | 0.043 | 9.25e-07 | 0.0312 |
| Malignant epithelial | C6 DNA-r p ECs | TNFAIP8 | 1.251 | 0.331 | 0.121 | 2.07e-11 | 6.98e-07 |
| Malignant epithelial | C6 DNA-r p ECs | CPSF4 | 1.25 | 0.363 | 0.145 | 4.19e-11 | 1.41e-06 |
| Malignant epithelial | C6 DNA-r p ECs | GALNT12 | 1.25 | 0.282 | 0.099 | 1.04e-10 | 3.5e-06 |
| Malignant epithelial | C6 DNA-r p ECs | CCDC109B | 1.249 | 0.508 | 0.183 | 4.78e-18 | 1.61e-13 |
| Malignant epithelial | C6 DNA-r p ECs | RNASEH2B | 1.248 | 0.46 | 0.19 | 1.29e-13 | 4.33e-09 |

| compartment | subcluster | gene | avg_log2FC | pct.1 | pct.2 | p_val | p_val_adj |
|---|---|---|---|---|---|---|---|
| Malignant epithelial | C6 DNA-r p ECs | DNAJC9 | 1.245 | 0.653 | 0.342 | 1.03e-14 | 3.46e-10 |
| Malignant epithelial | C6 DNA-r p ECs | GMNN | 1.243 | 0.306 | 0.106 | 5.38e-12 | 1.81e-07 |
| Malignant epithelial | C6 DNA-r p ECs | PIK3AP1 | 1.242 | 0.177 | 0.064 | 1.22e-06 | 0.041 |
| Malignant epithelial | C6 DNA-r p ECs | MSH6 | 1.239 | 0.234 | 0.082 | 6.31e-09 | 0.000213 |
| Malignant epithelial | C6 DNA-r p ECs | NR2F2 | 1.238 | 0.46 | 0.224 | 9.28e-11 | 3.13e-06 |
| Malignant epithelial | C6 DNA-r p ECs | WDR90 | 1.238 | 0.177 | 0.061 | 2.34e-07 | 0.00787 |
| Malignant epithelial | C6 DNA-r p ECs | PRPF4 | 1.237 | 0.347 | 0.129 | 1.61e-11 | 5.41e-07 |
| Malignant epithelial | C6 DNA-r p ECs | RP5-1159O4.1 | 1.234 | 0.25 | 0.077 | 1.83e-11 | 6.18e-07 |
| Malignant epithelial | C6 DNA-r p ECs | ACP6 | 1.232 | 0.25 | 0.093 | 1.51e-08 | 0.00051 |
| Malignant epithelial | C6 DNA-r p ECs | UBR4 | 1.229 | 0.5 | 0.198 | 2.18e-15 | 7.33e-11 |
| Malignant epithelial | C6 DNA-r p ECs | BATF | 1.229 | 0.379 | 0.151 | 2.55e-11 | 8.59e-07 |
| Malignant epithelial | C6 DNA-r p ECs | FAM118B | 1.229 | 0.226 | 0.083 | 3.19e-08 | 0.00107 |
| Malignant epithelial | C6 DNA-r p ECs | USP1 | 1.227 | 0.411 | 0.162 | 1.37e-12 | 4.62e-08 |
| Malignant epithelial | C6 DNA-r p ECs | RPA2 | 1.227 | 0.339 | 0.142 | 2.38e-09 | 8.01e-05 |
| Malignant epithelial | C6 DNA-r p ECs | C14orf80 | 1.226 | 0.411 | 0.135 | 1.07e-16 | 3.59e-12 |
| Malignant epithelial | C6 DNA-r p ECs | CEP57L1 | 1.226 | 0.145 | 0.046 | 5.68e-07 | 0.0191 |
| Malignant epithelial | C6 DNA-r p ECs | H2AFZ | 1.224 | 0.96 | 0.831 | 1.74e-23 | 5.88e-19 |
| Malignant epithelial | C6 DNA-r p ECs | CCM2 | 1.224 | 0.573 | 0.222 | 1.97e-19 | 6.64e-15 |
| Malignant epithelial | C6 DNA-r p ECs | CCDC74A | 1.224 | 0.202 | 0.067 | 2.05e-08 | 0.000691 |
| Malignant epithelial | C6 DNA-r p ECs | ZNF185 | 1.223 | 0.169 | 0.06 | 1.01e-06 | 0.0341 |
| Malignant epithelial | C6 DNA-r p ECs | FABP5 | 1.221 | 0.831 | 0.546 | 4.43e-17 | 1.49e-12 |
| Malignant epithelial | C6 DNA-r p ECs | EXOSC8 | 1.22 | 0.653 | 0.31 | 6.49e-17 | 2.19e-12 |
| Malignant epithelial | C6 DNA-r p ECs | BHLHE41 | 1.22 | 0.169 | 0.054 | 9.19e-08 | 0.0031 |
| Malignant epithelial | C6 DNA-r p ECs | IFI44L | 1.219 | 0.468 | 0.176 | 3.82e-15 | 1.29e-10 |
| Malignant epithelial | C6 DNA-r p ECs | PLOD1 | 1.219 | 0.379 | 0.164 | 5.94e-10 | 2e-05 |
| Malignant epithelial | C6 DNA-r p ECs | ZDHHC12 | 1.217 | 0.839 | 0.515 | 2.56e-19 | 8.62e-15 |
| Malignant epithelial | C6 DNA-r p ECs | HAUS7 | 1.215 | 0.258 | 0.093 | 2.81e-09 | 9.46e-05 |
| Malignant epithelial | C6 DNA-r p ECs | ARHGEF16 | 1.214 | 0.347 | 0.145 | 5.04e-10 | 1.7e-05 |
| Malignant epithelial | C6 DNA-r p ECs | CCDC34 | 1.202 | 0.629 | 0.327 | 3.46e-14 | 1.16e-09 |
| Malignant epithelial | C6 DNA-r p ECs | IL1R2 | 1.202 | 0.258 | 0.095 | 5.19e-09 | 0.000175 |
| Malignant epithelial | C6 DNA-r p ECs | H2AFX | 1.198 | 0.677 | 0.341 | 8.15e-16 | 2.75e-11 |
| Malignant epithelial | C6 DNA-r p ECs | SEPT8 | 1.198 | 0.379 | 0.134 | 5.42e-14 | 1.82e-09 |
| Malignant epithelial | C6 DNA-r p ECs | EIF6 | 1.196 | 0.984 | 0.824 | 5.26e-21 | 1.77e-16 |
| Malignant epithelial | C6 DNA-r p ECs | UBL7-AS1 | 1.195 | 0.169 | 0.059 | 8.62e-07 | 0.029 |
| Malignant epithelial | C6 DNA-r p ECs | TPD52L1 | 1.194 | 0.548 | 0.27 | 2.01e-12 | 6.78e-08 |
| Malignant epithelial | C6 DNA-r p ECs | TGFBI | 1.194 | 0.645 | 0.432 | 1.15e-07 | 0.00388 |
| Malignant epithelial | C6 DNA-r p ECs | LRR1 | 1.193 | 0.25 | 0.085 | 5.31e-10 | 1.79e-05 |
| Malignant epithelial | C6 DNA-r p ECs | SP140 | 1.191 | 0.169 | 0.055 | 1.74e-07 | 0.00588 |
| Malignant epithelial | C6 DNA-r p ECs | AKR7A3 | 1.19 | 0.371 | 0.145 | 8.75e-12 | 2.95e-07 |
| Malignant epithelial | C6 DNA-r p ECs | OVOL2 | 1.189 | 0.234 | 0.09 | 6.87e-08 | 0.00232 |
| Malignant epithelial | C6 DNA-r p ECs | KHK | 1.184 | 0.25 | 0.094 | 1.97e-08 | 0.000662 |
| Malignant epithelial | C6 DNA-r p ECs | APEX2 | 1.181 | 0.315 | 0.103 | 5.95e-13 | 2.01e-08 |
| Malignant epithelial | C6 DNA-r p ECs | DDX39A | 1.179 | 0.718 | 0.406 | 4.95e-16 | 1.67e-11 |
| Malignant epithelial | C6 DNA-r p ECs | PXMP2 | 1.178 | 0.492 | 0.23 | 1.49e-11 | 5.02e-07 |
| Malignant epithelial | C6 DNA-r p ECs | CHEK1 | 1.176 | 0.331 | 0.111 | 1.05e-12 | 3.52e-08 |
| Malignant epithelial | C6 DNA-r p ECs | EED | 1.173 | 0.29 | 0.098 | 2.02e-11 | 6.81e-07 |
| Malignant epithelial | C6 DNA-r p ECs | ACE2 | 1.173 | 0.153 | 0.05 | 8.06e-07 | 0.0272 |
| Malignant epithelial | C6 DNA-r p ECs | TMEM97 | 1.172 | 0.508 | 0.243 | 9.5e-12 | 3.2e-07 |
| Malignant epithelial | C6 DNA-r p ECs | CHMP6 | 1.172 | 0.315 | 0.114 | 6.45e-11 | 2.17e-06 |
| Malignant epithelial | C6 DNA-r p ECs | GPN3 | 1.171 | 0.419 | 0.156 | 8.33e-14 | 2.81e-09 |
| Malignant epithelial | C6 DNA-r p ECs | TEX30 | 1.169 | 0.323 | 0.129 | 1.46e-09 | 4.91e-05 |
| Malignant epithelial | C6 DNA-r p ECs | TBXAS1 | 1.169 | 0.218 | 0.079 | 5.88e-08 | 0.00198 |
| Malignant epithelial | C6 DNA-r p ECs | SLC35A1 | 1.168 | 0.371 | 0.183 | 1.08e-07 | 0.00363 |
| Malignant epithelial | C6 DNA-r p ECs | ARC | 1.168 | 0.161 | 0.054 | 6.09e-07 | 0.0205 |
| Malignant epithelial | C6 DNA-r p ECs | EFHD2 | 1.167 | 0.742 | 0.442 | 6.9e-15 | 2.33e-10 |
| Malignant epithelial | C6 DNA-r p ECs | NUP62CL | 1.164 | 0.234 | 0.07 | 2.47e-11 | 8.33e-07 |
| Malignant epithelial | C6 DNA-r p ECs | SH3BGRL2 | 1.163 | 0.363 | 0.137 | 1.4e-11 | 4.73e-07 |
| Malignant epithelial | C6 DNA-r p ECs | GPX2 | 1.159 | 0.911 | 0.487 | 6.859999999999999e-23 | 2.31e-18 |
| Malignant epithelial | C6 DNA-r p ECs | KRT80 | 1.158 | 0.298 | 0.107 | 1.3e-10 | 4.38e-06 |
| Malignant epithelial | C6 DNA-r p ECs | BPGM | 1.158 | 0.218 | 0.084 | 2.63e-07 | 0.00887 |
| Malignant epithelial | C6 DNA-r p ECs | TMPRSS13 | 1.155 | 0.194 | 0.07 | 3.68e-07 | 0.0124 |
| Malignant epithelial | C6 DNA-r p ECs | TPM2 | 1.154 | 0.968 | 0.766 | 2.92e-24 | 9.84e-20 |
| Malignant epithelial | C6 DNA-r p ECs | RFC5 | 1.153 | 0.226 | 0.086 | 2.32e-07 | 0.00782 |
| Malignant epithelial | C6 DNA-r p ECs | LINC01133 | 1.149 | 0.339 | 0.111 | 8.37e-14 | 2.82e-09 |

| compartment | subcluster | gene | avg_log2FC | pct.1 | pct.2 | p_val | p_val_adj |
|---|---|---|---|---|---|---|---|
| Malignant epithelial | C6 DNA-r p ECs | HMMR | 1.148 | 0.226 | 0.061 | 8e-13 | 2.69e-08 |
| Malignant epithelial | C6 DNA-r p ECs | CDCA7L | 1.146 | 0.411 | 0.183 | 3.66e-10 | 1.23e-05 |
| Malignant epithelial | C6 DNA-r p ECs | RAB27A | 1.145 | 0.266 | 0.091 | 2.31e-10 | 7.79e-06 |
| Malignant epithelial | C6 DNA-r p ECs | SLC9A3R2 | 1.145 | 0.403 | 0.186 | 3.36e-10 | 1.13e-05 |
| Malignant epithelial | C6 DNA-r p ECs | NDRG2 | 1.142 | 0.274 | 0.111 | 4.65e-08 | 0.00157 |
| Malignant epithelial | C6 DNA-r p ECs | NOL9 | 1.141 | 0.202 | 0.071 | 1.25e-07 | 0.00421 |
| Malignant epithelial | C6 DNA-r p ECs | PRTFDC1 | 1.139 | 0.242 | 0.1 | 6.93e-07 | 0.0234 |
| Malignant epithelial | C6 DNA-r p ECs | SLC29A1 | 1.138 | 0.282 | 0.11 | 6.08e-09 | 0.000205 |
| Malignant epithelial | C6 DNA-r p ECs | RND1 | 1.138 | 0.266 | 0.115 | 4.62e-07 | 0.0156 |
| Malignant epithelial | C6 DNA-r p ECs | RAMP1 | 1.137 | 0.629 | 0.351 | 1.33e-11 | 4.48e-07 |
| Malignant epithelial | C6 DNA-r p ECs | GUSB | 1.136 | 0.484 | 0.21 | 7.67e-13 | 2.59e-08 |
| Malignant epithelial | C6 DNA-r p ECs | SLC9A3R1 | 1.132 | 0.661 | 0.325 | 2.88e-16 | 9.71e-12 |
| Malignant epithelial | C6 DNA-r p ECs | PITX1 | 1.13 | 0.839 | 0.448 | 8.3e-20 | 2.8e-15 |
| Malignant epithelial | C6 DNA-r p ECs | GALM | 1.129 | 0.331 | 0.13 | 2.64e-10 | 8.88e-06 |
| Malignant epithelial | C6 DNA-r p ECs | CTPS1 | 1.128 | 0.161 | 0.052 | 3.19e-07 | 0.0108 |
| Malignant epithelial | C6 DNA-r p ECs | PFKP | 1.126 | 0.629 | 0.339 | 2.68e-11 | 9.02e-07 |
| Malignant epithelial | C6 DNA-r p ECs | ISG20 | 1.126 | 0.702 | 0.408 | 6.84e-11 | 2.31e-06 |
| Malignant epithelial | C6 DNA-r p ECs | CDCA3 | 1.125 | 0.347 | 0.113 | 2.08e-14 | 7.02e-10 |
| Malignant epithelial | C6 DNA-r p ECs | SH3GLB2 | 1.125 | 0.492 | 0.208 | 1.69e-13 | 5.71e-09 |
| Malignant epithelial | C6 DNA-r p ECs | SYT1 | 1.125 | 0.218 | 0.08 | 7.96e-08 | 0.00268 |
| Malignant epithelial | C6 DNA-r p ECs | CENPW | 1.124 | 0.831 | 0.432 | 6.39e-21 | 2.15e-16 |
| Malignant epithelial | C6 DNA-r p ECs | AKR7A2 | 1.124 | 0.508 | 0.251 | 1.84e-11 | 6.21e-07 |
| Malignant epithelial | C6 DNA-r p ECs | RECQL | 1.124 | 0.25 | 0.079 | 8.36e-11 | 2.82e-06 |
| Malignant epithelial | C6 DNA-r p ECs | ZDHHC14 | 1.122 | 0.169 | 0.059 | 9.26e-07 | 0.0312 |
| Malignant epithelial | C6 DNA-r p ECs | POLR3K | 1.12 | 0.556 | 0.245 | 3.55e-14 | 1.2e-09 |
| Malignant epithelial | C6 DNA-r p ECs | AGA | 1.119 | 0.234 | 0.096 | 9.14e-07 | 0.0308 |
| Malignant epithelial | C6 DNA-r p ECs | TMEM243 | 1.118 | 0.629 | 0.258 | 1.27e-17 | 4.29e-13 |
| Malignant epithelial | C6 DNA-r p ECs | CYP4F3 | 1.118 | 0.29 | 0.109 | 3.03e-09 | 0.000102 |
| Malignant epithelial | C6 DNA-r p ECs | PARP2 | 1.117 | 0.194 | 0.062 | 1.63e-08 | 0.000548 |
| Malignant epithelial | C6 DNA-r p ECs | CKMT1B | 1.114 | 0.306 | 0.113 | 3.3e-10 | 1.11e-05 |
| Malignant epithelial | C6 DNA-r p ECs | FAHD2A | 1.112 | 0.347 | 0.14 | 5.87e-10 | 1.98e-05 |
| Malignant epithelial | C6 DNA-r p ECs | AQP1 | 1.111 | 0.371 | 0.14 | 1e-11 | 3.37e-07 |
| Malignant epithelial | C6 DNA-r p ECs | AIM1 | 1.111 | 0.315 | 0.115 | 9.38e-11 | 3.16e-06 |
| Malignant epithelial | C6 DNA-r p ECs | MGME1 | 1.107 | 0.435 | 0.185 | 8.63e-12 | 2.91e-07 |
| Malignant epithelial | C6 DNA-r p ECs | EPB41L4B | 1.105 | 0.226 | 0.092 | 1.08e-06 | 0.0365 |
| Malignant epithelial | C6 DNA-r p ECs | PLPP1 | 1.104 | 0.274 | 0.103 | 4.07e-09 | 0.000137 |
| Malignant epithelial | C6 DNA-r p ECs | OGFOD3 | 1.103 | 0.403 | 0.201 | 2.53e-08 | 0.000851 |
| Malignant epithelial | C6 DNA-r p ECs | UGT8 | 1.103 | 0.194 | 0.069 | 3.18e-07 | 0.0107 |
| Malignant epithelial | C6 DNA-r p ECs | DPM2 | 1.099 | 0.823 | 0.452 | 3.85e-18 | 1.3e-13 |
| Malignant epithelial | C6 DNA-r p ECs | CNTLN | 1.095 | 0.226 | 0.072 | 1.1e-09 | 3.71e-05 |
| Malignant epithelial | C6 DNA-r p ECs | PUSL1 | 1.095 | 0.242 | 0.089 | 1.88e-08 | 0.000634 |
| Malignant epithelial | C6 DNA-r p ECs | SIVA1 | 1.094 | 0.952 | 0.742 | 3.88e-19 | 1.31e-14 |
| Malignant epithelial | C6 DNA-r p ECs | G6PC3 | 1.093 | 0.476 | 0.209 | 1.73e-12 | 5.82e-08 |
| Malignant epithelial | C6 DNA-r p ECs | SMC3 | 1.092 | 0.54 | 0.306 | 4.22e-08 | 0.00142 |
| Malignant epithelial | C6 DNA-r p ECs | GIPC2 | 1.092 | 0.169 | 0.057 | 5.18e-07 | 0.0174 |
| Malignant epithelial | C6 DNA-r p ECs | CDKN3 | 1.091 | 0.5 | 0.189 | 2.88e-16 | 9.69e-12 |
| Malignant epithelial | C6 DNA-r p ECs | SPDEF | 1.089 | 0.21 | 0.078 | 2.62e-07 | 0.00884 |
| Malignant epithelial | C6 DNA-r p ECs | DLEU1 | 1.089 | 0.339 | 0.16 | 3.28e-07 | 0.011 |
| Malignant epithelial | C6 DNA-r p ECs | TUBB | 1.084 | 0.919 | 0.681 | 1.81e-18 | 6.11e-14 |
| Malignant epithelial | C6 DNA-r p ECs | MRPS11 | 1.084 | 0.669 | 0.362 | 7.53e-13 | 2.54e-08 |
| Malignant epithelial | C6 DNA-r p ECs | C12orf29 | 1.084 | 0.323 | 0.121 | 2.8e-10 | 9.45e-06 |
| Malignant epithelial | C6 DNA-r p ECs | IGFBP4 | 1.083 | 0.637 | 0.28 | 9.01e-16 | 3.04e-11 |
| Malignant epithelial | C6 DNA-r p ECs | CTGF | 1.079 | 0.411 | 0.178 | 4.85e-10 | 1.64e-05 |
| Malignant epithelial | C6 DNA-r p ECs | RRP36 | 1.079 | 0.323 | 0.153 | 1.23e-06 | 0.0415 |
| Malignant epithelial | C6 DNA-r p ECs | DKK1 | 1.077 | 0.306 | 0.099 | 3.64e-13 | 1.23e-08 |
| Malignant epithelial | C6 DNA-r p ECs | ATP13A2 | 1.077 | 0.202 | 0.074 | 3.22e-07 | 0.0108 |
| Malignant epithelial | C6 DNA-r p ECs | GPR153 | 1.075 | 0.177 | 0.065 | 1.15e-06 | 0.0388 |
| Malignant epithelial | C6 DNA-r p ECs | GALE | 1.074 | 0.524 | 0.272 | 7.19e-10 | 2.42e-05 |
| Malignant epithelial | C6 DNA-r p ECs | PEX10 | 1.073 | 0.411 | 0.166 | 6.22e-12 | 2.1e-07 |
| Malignant epithelial | C6 DNA-r p ECs | GOSR2 | 1.072 | 0.347 | 0.147 | 2.56e-09 | 8.62e-05 |
| Malignant epithelial | C6 DNA-r p ECs | MOCS2 | 1.072 | 0.371 | 0.171 | 8.78e-09 | 0.000296 |
| Malignant epithelial | C6 DNA-r p ECs | FKBP5 | 1.069 | 0.226 | 0.091 | 8.37e-07 | 0.0282 |
| Malignant epithelial | C6 DNA-r p ECs | EPN3 | 1.069 | 0.218 | 0.087 | 1.08e-06 | 0.0363 |
| Malignant epithelial | C6 DNA-r p ECs | BLCAP | 1.068 | 0.71 | 0.406 | 5.03e-13 | 1.69e-08 |

| compartment | subcluster | gene | avg_log2FC | pct.1 | pct.2 | p_val | p_val_adj |
|---|---|---|---|---|---|---|---|
| Malignant epithelial | C6 DNA-r p ECs | KIF22 | 1.067 | 0.435 | 0.19 | 4.19e-11 | 1.41e-06 |
| Malignant epithelial | C6 DNA-r p ECs | WDR5 | 1.066 | 0.387 | 0.151 | 1.71e-11 | 5.75e-07 |
| Malignant epithelial | C6 DNA-r p ECs | CSF2 | 1.065 | 0.185 | 0.056 | 2.29e-09 | 7.71e-05 |
| Malignant epithelial | C6 DNA-r p ECs | HAT1 | 1.065 | 0.395 | 0.209 | 4.18e-07 | 0.0141 |
| Malignant epithelial | C6 DNA-r p ECs | PTRF | 1.061 | 0.427 | 0.181 | 1.49e-11 | 5.01e-07 |
| Malignant epithelial | C6 DNA-r p ECs | DBF4 | 1.061 | 0.298 | 0.132 | 2.34e-07 | 0.00788 |
| Malignant epithelial | C6 DNA-r p ECs | SLC2A8 | 1.059 | 0.258 | 0.097 | 1.21e-08 | 0.000407 |
| Malignant epithelial | C6 DNA-r p ECs | OAS2 | 1.059 | 0.323 | 0.15 | 3.86e-07 | 0.013 |
| Malignant epithelial | C6 DNA-r p ECs | FADD | 1.057 | 0.395 | 0.171 | 4.17e-10 | 1.4e-05 |
| Malignant epithelial | C6 DNA-r p ECs | PLEK2 | 1.056 | 0.758 | 0.391 | 2.13e-16 | 7.18e-12 |
| Malignant epithelial | C6 DNA-r p ECs | PRPS2 | 1.056 | 0.516 | 0.26 | 1.42e-10 | 4.78e-06 |
| Malignant epithelial | C6 DNA-r p ECs | RNF121 | 1.056 | 0.258 | 0.096 | 1.1e-08 | 0.000369 |
| Malignant epithelial | C6 DNA-r p ECs | PBX3 | 1.055 | 0.242 | 0.083 | 2e-09 | 6.75e-05 |
| Malignant epithelial | C6 DNA-r p ECs | BAIAP2 | 1.054 | 0.589 | 0.331 | 1.19e-10 | 4e-06 |
| Malignant epithelial | C6 DNA-r p ECs | IFI44 | 1.052 | 0.331 | 0.152 | 7.56e-08 | 0.00255 |
| Malignant epithelial | C6 DNA-r p ECs | TFDP1 | 1.051 | 0.54 | 0.238 | 5.92e-14 | 1.99e-09 |
| Malignant epithelial | C6 DNA-r p ECs | C8orf76 | 1.05 | 0.331 | 0.147 | 9.14e-08 | 0.00308 |
| Malignant epithelial | C6 DNA-r p ECs | VGF | 1.047 | 0.258 | 0.108 | 2.94e-07 | 0.00992 |
| Malignant epithelial | C6 DNA-r p ECs | RNASET2 | 1.046 | 0.935 | 0.66 | 2.93e-17 | 9.86e-13 |
| Malignant epithelial | C6 DNA-r p ECs | CCL15 | 1.046 | 0.403 | 0.132 | 3.88e-16 | 1.31e-11 |
| Malignant epithelial | C6 DNA-r p ECs | TUBA4A | 1.046 | 0.758 | 0.44 | 1.29e-12 | 4.35e-08 |
| Malignant epithelial | C6 DNA-r p ECs | COX6B2 | 1.045 | 0.194 | 0.07 | 4.25e-07 | 0.0143 |
| Malignant epithelial | C6 DNA-r p ECs | DDB2 | 1.044 | 0.395 | 0.16 | 3.11e-11 | 1.05e-06 |
| Malignant epithelial | C6 DNA-r p ECs | DNAJC11 | 1.044 | 0.242 | 0.096 | 1.76e-07 | 0.00592 |
| Malignant epithelial | C6 DNA-r p ECs | HMGN3 | 1.042 | 0.669 | 0.323 | 4.44e-15 | 1.49e-10 |
| Malignant epithelial | C6 DNA-r p ECs | TCTEX1D2 | 1.042 | 0.54 | 0.24 | 5.96e-13 | 2.01e-08 |
| Malignant epithelial | C6 DNA-r p ECs | LAT2 | 1.042 | 0.194 | 0.074 | 1.28e-06 | 0.043 |
| Malignant epithelial | C6 DNA-r p ECs | LRCH4 | 1.041 | 0.363 | 0.171 | 8.46e-08 | 0.00285 |
| Malignant epithelial | C6 DNA-r p ECs | AP1M1 | 1.04 | 0.274 | 0.114 | 2.53e-07 | 0.00851 |
| Malignant epithelial | C6 DNA-r p ECs | USP18 | 1.039 | 0.25 | 0.102 | 2.7e-07 | 0.0091 |
| Malignant epithelial | C6 DNA-r p ECs | RBBP7 | 1.038 | 0.581 | 0.354 | 4.72e-10 | 1.59e-05 |
| Malignant epithelial | C6 DNA-r p ECs | CC2D1A | 1.038 | 0.379 | 0.164 | 5.25e-09 | 0.000177 |
| Malignant epithelial | C6 DNA-r p ECs | SLC7A1 | 1.037 | 0.331 | 0.127 | 2.63e-10 | 8.85e-06 |
| Malignant epithelial | C6 DNA-r p ECs | ANXA9 | 1.037 | 0.21 | 0.074 | 7.98e-08 | 0.00269 |
| Malignant epithelial | C6 DNA-r p ECs | UFSP2 | 1.036 | 0.274 | 0.108 | 2.35e-08 | 0.000791 |
| Malignant epithelial | C6 DNA-r p ECs | SDHB | 1.035 | 0.702 | 0.41 | 2.91e-13 | 9.79e-09 |
| Malignant epithelial | C6 DNA-r p ECs | DPP3 | 1.035 | 0.339 | 0.142 | 3.5e-09 | 0.000118 |
| Malignant epithelial | C6 DNA-r p ECs | ATP2C2 | 1.033 | 0.242 | 0.089 | 2.1e-08 | 0.000706 |
| Malignant epithelial | C6 DNA-r p ECs | PIGX | 1.032 | 0.403 | 0.18 | 1.03e-09 | 3.45e-05 |
| Malignant epithelial | C6 DNA-r p ECs | RGS3 | 1.032 | 0.266 | 0.093 | 1.03e-09 | 3.47e-05 |
| Malignant epithelial | C6 DNA-r p ECs | RNF223 | 1.032 | 0.323 | 0.154 | 4.07e-07 | 0.0137 |
| Malignant epithelial | C6 DNA-r p ECs | CDKN2A | 1.03 | 0.976 | 0.758 | 1.44e-21 | 4.85e-17 |
| Malignant epithelial | C6 DNA-r p ECs | NIPAL3 | 1.029 | 0.226 | 0.08 | 2.59e-08 | 0.000871 |
| Malignant epithelial | C6 DNA-r p ECs | C3orf14 | 1.028 | 0.298 | 0.112 | 1.76e-09 | 5.94e-05 |
| Malignant epithelial | C6 DNA-r p ECs | ZFYVE19 | 1.028 | 0.315 | 0.147 | 3.55e-07 | 0.012 |
| Malignant epithelial | C6 DNA-r p ECs | SLC27A5 | 1.025 | 0.21 | 0.082 | 9.2e-07 | 0.031 |
| Malignant epithelial | C6 DNA-r p ECs | SLBP | 1.024 | 0.508 | 0.265 | 1.18e-09 | 3.97e-05 |
| Malignant epithelial | C6 DNA-r p ECs | PPIF | 1.023 | 0.637 | 0.345 | 1.31e-11 | 4.41e-07 |
| Malignant epithelial | C6 DNA-r p ECs | MAP3K4 | 1.023 | 0.169 | 0.059 | 8.14e-07 | 0.0274 |
| Malignant epithelial | C6 DNA-r p ECs | TRIM69 | 1.022 | 0.323 | 0.136 | 1.44e-08 | 0.000485 |
| Malignant epithelial | C6 DNA-r p ECs | GTSE1 | 1.021 | 0.226 | 0.07 | 2.75e-10 | 9.26e-06 |
| Malignant epithelial | C6 DNA-r p ECs | TEX261 | 1.021 | 0.331 | 0.138 | 1.49e-08 | 0.000503 |
| Malignant epithelial | C6 DNA-r p ECs | MIS18A | 1.02 | 0.306 | 0.13 | 4.06e-08 | 0.00137 |
| Malignant epithelial | C6 DNA-r p ECs | UNK | 1.019 | 0.355 | 0.138 | 1.31e-10 | 4.4e-06 |
| Malignant epithelial | C6 DNA-r p ECs | MRTO4 | 1.019 | 0.5 | 0.245 | 3.62e-10 | 1.22e-05 |
| Malignant epithelial | C6 DNA-r p ECs | MAPK9 | 1.016 | 0.25 | 0.104 | 5.57e-07 | 0.0188 |
| Malignant epithelial | C6 DNA-r p ECs | KIFC1 | 1.015 | 0.25 | 0.084 | 6.13e-10 | 2.07e-05 |
| Malignant epithelial | C6 DNA-r p ECs | HMGN5 | 1.015 | 0.234 | 0.098 | 1.24e-06 | 0.0418 |
| Malignant epithelial | C6 DNA-r p ECs | CTSL | 1.014 | 0.411 | 0.209 | 1.56e-07 | 0.00524 |
| Malignant epithelial | C6 DNA-r p ECs | ACER3 | 1.014 | 0.202 | 0.074 | 4.41e-07 | 0.0149 |
| Malignant epithelial | C6 DNA-r p ECs | BUB3 | 1.013 | 0.782 | 0.474 | 2.66e-15 | 8.96e-11 |
| Malignant epithelial | C6 DNA-r p ECs | COA1 | 1.012 | 0.726 | 0.35 | 1.17e-16 | 3.93e-12 |
| Malignant epithelial | C6 DNA-r p ECs | NBL1 | 1.011 | 0.806 | 0.457 | 2.21e-17 | 7.43e-13 |
| Malignant epithelial | C6 DNA-r p ECs | ANLN | 1.011 | 0.427 | 0.149 | 2.72e-15 | 9.16e-11 |

| compartment | subcluster | gene | avg_log2FC | pct.1 | pct.2 | p_val | p_val_adj |
|---|---|---|---|---|---|---|---|
| Malignant epithelial | C6 DNA-r p ECs | UPK2 | 1.011 | 0.274 | 0.103 | 8.91e-10 | 3e-05 |
| Malignant epithelial | C6 DNA-r p ECs | FBP1 | 1.01 | 0.371 | 0.173 | 1.21e-07 | 0.00406 |
| Malignant epithelial | C6 DNA-r p ECs | CLSTN1 | 1.007 | 0.653 | 0.3 | 3.92e-15 | 1.32e-10 |
| Malignant epithelial | C6 DNA-r p ECs | C9orf152 | 1.005 | 0.347 | 0.131 | 5.21e-11 | 1.75e-06 |
| Malignant epithelial | C6 DNA-r p ECs | HAUS5 | 1.004 | 0.202 | 0.069 | 7.05e-08 | 0.00238 |
| Malignant epithelial | C6 DNA-r p ECs | CKAP5 | 1.003 | 0.274 | 0.108 | 3.88e-08 | 0.00131 |
| Malignant epithelial | C6 DNA-r p ECs | TMEM62 | 1.003 | 0.274 | 0.112 | 6.11e-08 | 0.00206 |
| Malignant epithelial | C6 DNA-r p ECs | ELOVL6 | 1.002 | 0.194 | 0.072 | 7.44e-07 | 0.0251 |
| Malignant epithelial | C6 DNA-r p ECs | ANO1 | 1.0 | 0.339 | 0.159 | 1.49e-07 | 0.00502 |
| Malignant epithelial | C6 DNA-r p ECs | GOT1 | 0.997 | 0.427 | 0.184 | 4.71e-11 | 1.59e-06 |
| Malignant epithelial | C6 DNA-r p ECs | TMX1 | 0.996 | 0.597 | 0.283 | 1.39e-13 | 4.69e-09 |
| Malignant epithelial | C6 DNA-r p ECs | GIPC1 | 0.996 | 0.863 | 0.612 | 3.47e-13 | 1.17e-08 |
| Malignant epithelial | C6 DNA-r p ECs | OAS3 | 0.996 | 0.411 | 0.211 | 2.7e-07 | 0.00908 |
| Malignant epithelial | C6 DNA-r p ECs | RACGAP1 | 0.993 | 0.226 | 0.082 | 5.36e-08 | 0.00181 |
| Malignant epithelial | C6 DNA-r p ECs | SPCS3 | 0.992 | 0.71 | 0.382 | 2.1e-14 | 7.07e-10 |
| Malignant epithelial | C6 DNA-r p ECs | TUBG1 | 0.992 | 0.339 | 0.147 | 3.07e-08 | 0.00104 |
| Malignant epithelial | C6 DNA-r p ECs | ARHGDIB | 0.988 | 0.758 | 0.471 | 2.46e-13 | 8.3e-09 |
| Malignant epithelial | C6 DNA-r p ECs | MAPK3 | 0.988 | 0.653 | 0.337 | 7.07e-12 | 2.38e-07 |
| Malignant epithelial | C6 DNA-r p ECs | RWDD4 | 0.988 | 0.427 | 0.204 | 8.29e-09 | 0.000279 |
| Malignant epithelial | C6 DNA-r p ECs | TPRG1L | 0.988 | 0.298 | 0.122 | 2.45e-08 | 0.000826 |
| Malignant epithelial | C6 DNA-r p ECs | PGAM1 | 0.987 | 0.871 | 0.593 | 4.6e-15 | 1.55e-10 |
| Malignant epithelial | C6 DNA-r p ECs | FANCA | 0.986 | 0.242 | 0.093 | 1.22e-07 | 0.00411 |
| Malignant epithelial | C6 DNA-r p ECs | SLC48A1 | 0.986 | 0.226 | 0.092 | 1.44e-06 | 0.0484 |
| Malignant epithelial | C6 DNA-r p ECs | HBEGF | 0.984 | 0.637 | 0.381 | 2.54e-09 | 8.57e-05 |
| Malignant epithelial | C6 DNA-r p ECs | DCTN3 | 0.983 | 0.863 | 0.508 | 2.76e-15 | 9.29e-11 |
| Malignant epithelial | C6 DNA-r p ECs | HPGD | 0.983 | 0.452 | 0.173 | 7.09e-15 | 2.39e-10 |
| Malignant epithelial | C6 DNA-r p ECs | TSPAN5 | 0.983 | 0.298 | 0.112 | 9.16e-10 | 3.09e-05 |
| Malignant epithelial | C6 DNA-r p ECs | PCYT2 | 0.983 | 0.347 | 0.155 | 5.34e-08 | 0.0018 |
| Malignant epithelial | C6 DNA-r p ECs | PSPH | 0.983 | 0.258 | 0.107 | 4.33e-07 | 0.0146 |
| Malignant epithelial | C6 DNA-r p ECs | ELP5 | 0.98 | 0.315 | 0.116 | 3.1e-10 | 1.04e-05 |
| Malignant epithelial | C6 DNA-r p ECs | RER1 | 0.978 | 0.815 | 0.523 | 1.65e-17 | 5.56e-13 |
| Malignant epithelial | C6 DNA-r p ECs | TRIM14 | 0.977 | 0.274 | 0.115 | 1.78e-07 | 0.006 |
| Malignant epithelial | C6 DNA-r p ECs | FAM24B | 0.976 | 0.25 | 0.098 | 1.16e-07 | 0.0039 |
| Malignant epithelial | C6 DNA-r p ECs | SMCO4 | 0.976 | 0.347 | 0.167 | 4.52e-07 | 0.0152 |
| Malignant epithelial | C6 DNA-r p ECs | HMGN2 | 0.975 | 0.968 | 0.765 | 5.46e-18 | 1.84e-13 |
| Malignant epithelial | C6 DNA-r p ECs | CHAF1A | 0.975 | 0.25 | 0.104 | 6.87e-07 | 0.0232 |
| Malignant epithelial | C6 DNA-r p ECs | FXYD5 | 0.974 | 0.992 | 0.891 | 5.73e-16 | 1.93e-11 |
| Malignant epithelial | C6 DNA-r p ECs | POP7 | 0.974 | 0.782 | 0.461 | 1.73e-15 | 5.83e-11 |
| Malignant epithelial | C6 DNA-r p ECs | EBP | 0.974 | 0.79 | 0.517 | 6.67e-15 | 2.25e-10 |
| Malignant epithelial | C6 DNA-r p ECs | CTNNBIP1 | 0.974 | 0.71 | 0.42 | 5.1e-11 | 1.72e-06 |
| Malignant epithelial | C6 DNA-r p ECs | CTSZ | 0.973 | 0.839 | 0.542 | 2.22e-14 | 7.46e-10 |
| Malignant epithelial | C6 DNA-r p ECs | SRA1 | 0.973 | 0.694 | 0.387 | 4.99e-12 | 1.68e-07 |
| Malignant epithelial | C6 DNA-r p ECs | FTSJ2 | 0.972 | 0.435 | 0.194 | 1.11e-09 | 3.73e-05 |
| Malignant epithelial | C6 DNA-r p ECs | ECE1 | 0.97 | 0.339 | 0.164 | 6.38e-07 | 0.0215 |
| Malignant epithelial | C6 DNA-r p ECs | SLC25A37 | 0.969 | 0.419 | 0.207 | 4.99e-08 | 0.00168 |
| Malignant epithelial | C6 DNA-r p ECs | UBAC1 | 0.968 | 0.411 | 0.19 | 2.8e-09 | 9.43e-05 |
| Malignant epithelial | C6 DNA-r p ECs | PPIC | 0.966 | 0.694 | 0.385 | 4.9e-13 | 1.65e-08 |
| Malignant epithelial | C6 DNA-r p ECs | EPHB4 | 0.965 | 0.323 | 0.137 | 1.27e-08 | 0.000428 |
| Malignant epithelial | C6 DNA-r p ECs | COMMD4 | 0.962 | 0.758 | 0.399 | 2.3e-15 | 7.74e-11 |
| Malignant epithelial | C6 DNA-r p ECs | NUDT1 | 0.962 | 0.71 | 0.384 | 1.49e-13 | 5.01e-09 |
| Malignant epithelial | C6 DNA-r p ECs | POLE4 | 0.959 | 0.847 | 0.504 | 4.09e-16 | 1.38e-11 |
| Malignant epithelial | C6 DNA-r p ECs | EPB41L2 | 0.959 | 0.387 | 0.185 | 4.76e-08 | 0.0016 |
| Malignant epithelial | C6 DNA-r p ECs | DYNLT1 | 0.958 | 0.968 | 0.687 | 1.02e-24 | 3.43e-20 |
| Malignant epithelial | C6 DNA-r p ECs | PLK1 | 0.957 | 0.234 | 0.081 | 4.44e-09 | 0.00015 |
| Malignant epithelial | C6 DNA-r p ECs | GINM1 | 0.957 | 0.427 | 0.208 | 4.91e-09 | 0.000165 |
| Malignant epithelial | C6 DNA-r p ECs | RSPO3 | 0.956 | 0.444 | 0.196 | 2.78e-10 | 9.36e-06 |
| Malignant epithelial | C6 DNA-r p ECs | HLA-DPA1 | 0.954 | 0.855 | 0.541 | 1.05e-15 | 3.55e-11 |
| Malignant epithelial | C6 DNA-r p ECs | CANT1 | 0.953 | 0.46 | 0.219 | 5.93e-10 | 2e-05 |
| Malignant epithelial | C6 DNA-r p ECs | SPAG1 | 0.952 | 0.29 | 0.121 | 9.8e-08 | 0.0033 |
| Malignant epithelial | C6 DNA-r p ECs | RB1 | 0.951 | 0.306 | 0.112 | 2.62e-10 | 8.83e-06 |
| Malignant epithelial | C6 DNA-r p ECs | DCPS | 0.951 | 0.355 | 0.151 | 4.84e-09 | 0.000163 |
| Malignant epithelial | C6 DNA-r p ECs | MKI67 | 0.95 | 0.387 | 0.137 | 7.5e-14 | 2.53e-09 |
| Malignant epithelial | C6 DNA-r p ECs | GDF15 | 0.947 | 0.685 | 0.384 | 6.64e-10 | 2.24e-05 |
| Malignant epithelial | C6 DNA-r p ECs | ACO1 | 0.947 | 0.258 | 0.109 | 8.78e-07 | 0.0296 |

| compartment | subcluster | gene | avg_log2FC | pct.1 | pct.2 | p_val | p_val_adj |
|---|---|---|---|---|---|---|---|
| Malignant epithelial | C6 DNA-r p ECs | MAL2 | 0.946 | 0.911 | 0.599 | 2e-21 | 6.74e-17 |
| Malignant epithelial | C6 DNA-r p ECs | HADH | 0.946 | 0.444 | 0.229 | 4.72e-08 | 0.00159 |
| Malignant epithelial | C6 DNA-r p ECs | NANS | 0.945 | 0.823 | 0.504 | 1.61e-16 | 5.43e-12 |
| Malignant epithelial | C6 DNA-r p ECs | POLE3 | 0.945 | 0.581 | 0.313 | 3.2e-10 | 1.08e-05 |
| Malignant epithelial | C6 DNA-r p ECs | IRF7 | 0.945 | 0.508 | 0.28 | 1.47e-08 | 0.000496 |
| Malignant epithelial | C6 DNA-r p ECs | ISG15 | 0.944 | 0.984 | 0.847 | 2.67e-13 | 9.01e-09 |
| Malignant epithelial | C6 DNA-r p ECs | SKA2 | 0.944 | 0.597 | 0.284 | 1.19e-12 | 4.01e-08 |
| Malignant epithelial | C6 DNA-r p ECs | CIAO1 | 0.943 | 0.427 | 0.226 | 3.8e-07 | 0.0128 |
| Malignant epithelial | C6 DNA-r p ECs | EXOSC3 | 0.942 | 0.54 | 0.266 | 4.55e-11 | 1.53e-06 |
| Malignant epithelial | C6 DNA-r p ECs | SNRNP40 | 0.942 | 0.403 | 0.193 | 3.2e-08 | 0.00108 |
| Malignant epithelial | C6 DNA-r p ECs | TECR | 0.941 | 0.927 | 0.743 | 2.48e-17 | 8.34e-13 |
| Malignant epithelial | C6 DNA-r p ECs | ATP5G1 | 0.94 | 0.935 | 0.744 | 1.62e-14 | 5.45e-10 |
| Malignant epithelial | C6 DNA-r p ECs | NEK6 | 0.939 | 0.411 | 0.18 | 1.19e-09 | 4.01e-05 |
| Malignant epithelial | C6 DNA-r p ECs | PGAP2 | 0.937 | 0.379 | 0.181 | 9.18e-08 | 0.00309 |
| Malignant epithelial | C6 DNA-r p ECs | B3GNT3 | 0.935 | 0.613 | 0.367 | 9.33e-09 | 0.000314 |
| Malignant epithelial | C6 DNA-r p ECs | PPP5C | 0.935 | 0.355 | 0.158 | 2.9e-08 | 0.000977 |
| Malignant epithelial | C6 DNA-r p ECs | SLC7A5 | 0.934 | 0.516 | 0.259 | 9.95e-09 | 0.000335 |
| Malignant epithelial | C6 DNA-r p ECs | ARHGEF10L | 0.934 | 0.21 | 0.081 | 8.5e-07 | 0.0286 |
| Malignant epithelial | C6 DNA-r p ECs | PPM1G | 0.933 | 0.79 | 0.509 | 2e-13 | 6.72e-09 |
| Malignant epithelial | C6 DNA-r p ECs | TCOF1 | 0.933 | 0.435 | 0.203 | 3.8e-09 | 0.000128 |
| Malignant epithelial | C6 DNA-r p ECs | STK17A | 0.933 | 0.452 | 0.231 | 7.86e-09 | 0.000265 |
| Malignant epithelial | C6 DNA-r p ECs | ING2 | 0.932 | 0.476 | 0.231 | 8.56e-10 | 2.88e-05 |
| Malignant epithelial | C6 DNA-r p ECs | CCND3 | 0.932 | 0.468 | 0.233 | 1.91e-09 | 6.43e-05 |
| Malignant epithelial | C6 DNA-r p ECs | LAP3 | 0.931 | 0.468 | 0.229 | 8.5e-10 | 2.86e-05 |
| Malignant epithelial | C6 DNA-r p ECs | ANAPC2 | 0.931 | 0.226 | 0.081 | 5.22e-08 | 0.00176 |
| Malignant epithelial | C6 DNA-r p ECs | GSDMB | 0.929 | 0.282 | 0.125 | 7.74e-07 | 0.0261 |
| Malignant epithelial | C6 DNA-r p ECs | MINOS1 | 0.927 | 0.968 | 0.771 | 7.550000000000001e-21 | 2.55e-16 |
| Malignant epithelial | C6 DNA-r p ECs | CEP55 | 0.925 | 0.298 | 0.091 | 1.94e-13 | 6.55e-09 |
| Malignant epithelial | C6 DNA-r p ECs | C9orf3 | 0.925 | 0.581 | 0.305 | 5.41e-10 | 1.82e-05 |
| Malignant epithelial | C6 DNA-r p ECs | RNF167 | 0.924 | 0.476 | 0.243 | 6.89e-09 | 0.000232 |
| Malignant epithelial | C6 DNA-r p ECs | VWA1 | 0.924 | 0.476 | 0.275 | 3.78e-07 | 0.0127 |
| Malignant epithelial | C6 DNA-r p ECs | ORC5 | 0.923 | 0.234 | 0.096 | 1.29e-06 | 0.0434 |
| Malignant epithelial | C6 DNA-r p ECs | PSMC2 | 0.922 | 0.621 | 0.322 | 2.56e-12 | 8.64e-08 |
| Malignant epithelial | C6 DNA-r p ECs | PSMB8-AS1 | 0.921 | 0.371 | 0.176 | 1.19e-07 | 0.00402 |
| Malignant epithelial | C6 DNA-r p ECs | SDF4 | 0.919 | 0.774 | 0.451 | 1.68e-14 | 5.65e-10 |
| Malignant epithelial | C6 DNA-r p ECs | NUCB2 | 0.918 | 0.427 | 0.191 | 2.86e-09 | 9.63e-05 |
| Malignant epithelial | C6 DNA-r p ECs | MTHFS | 0.917 | 0.589 | 0.315 | 1.3e-10 | 4.37e-06 |
| Malignant epithelial | C6 DNA-r p ECs | GGCT | 0.914 | 0.823 | 0.579 | 5.67e-14 | 1.91e-09 |
| Malignant epithelial | C6 DNA-r p ECs | SIGMAR1 | 0.914 | 0.718 | 0.411 | 2.92e-11 | 9.84e-07 |
| Malignant epithelial | C6 DNA-r p ECs | MTHFD1L | 0.913 | 0.331 | 0.144 | 5.01e-08 | 0.00169 |
| Malignant epithelial | C6 DNA-r p ECs | TSC22D3 | 0.913 | 0.718 | 0.508 | 8.85e-08 | 0.00298 |
| Malignant epithelial | C6 DNA-r p ECs | C12orf65 | 0.912 | 0.331 | 0.15 | 1.71e-07 | 0.00575 |
| Malignant epithelial | C6 DNA-r p ECs | STMN1 | 0.911 | 0.831 | 0.554 | 9.69e-13 | 3.26e-08 |
| Malignant epithelial | C6 DNA-r p ECs | RPS6KA4 | 0.91 | 0.387 | 0.179 | 2.3e-08 | 0.000775 |
| Malignant epithelial | C6 DNA-r p ECs | ACD | 0.91 | 0.355 | 0.17 | 2.9e-07 | 0.00977 |
| Malignant epithelial | C6 DNA-r p ECs | OSTF1 | 0.908 | 0.766 | 0.407 | 1.44e-14 | 4.86e-10 |
| Malignant epithelial | C6 DNA-r p ECs | MRPL20 | 0.907 | 0.944 | 0.671 | 1.2e-18 | 4.05e-14 |
| Malignant epithelial | C6 DNA-r p ECs | RASSF7 | 0.907 | 0.782 | 0.401 | 2.5e-17 | 8.42e-13 |
| Malignant epithelial | C6 DNA-r p ECs | AGRN | 0.906 | 0.508 | 0.249 | 5.19e-10 | 1.75e-05 |
| Malignant epithelial | C6 DNA-r p ECs | MRFAP1L1 | 0.906 | 0.258 | 0.112 | 1.18e-06 | 0.0397 |
| Malignant epithelial | C6 DNA-r p ECs | DBNL | 0.905 | 0.766 | 0.385 | 1.23e-15 | 4.14e-11 |
| Malignant epithelial | C6 DNA-r p ECs | MRPL17 | 0.905 | 0.839 | 0.491 | 1.46e-15 | 4.91e-11 |
| Malignant epithelial | C6 DNA-r p ECs | DLG5 | 0.905 | 0.331 | 0.158 | 5.97e-07 | 0.0201 |
| Malignant epithelial | C6 DNA-r p ECs | CDA | 0.905 | 0.734 | 0.635 | 1.2e-06 | 0.0403 |
| Malignant epithelial | C6 DNA-r p ECs | STARD3NL | 0.904 | 0.435 | 0.223 | 1.02e-07 | 0.00345 |
| Malignant epithelial | C6 DNA-r p ECs | LSM4 | 0.903 | 0.968 | 0.821 | 8.1e-26 | 2.73e-21 |
| Malignant epithelial | C6 DNA-r p ECs | HINT2 | 0.903 | 0.887 | 0.619 | 3.37e-15 | 1.13e-10 |
| Malignant epithelial | C6 DNA-r p ECs | TMPO | 0.903 | 0.492 | 0.252 | 1.42e-09 | 4.78e-05 |
| Malignant epithelial | C6 DNA-r p ECs | FAM76B | 0.903 | 0.202 | 0.076 | 6.88e-07 | 0.0232 |
| Malignant epithelial | C6 DNA-r p ECs | NAPRT | 0.902 | 0.944 | 0.625 | 1.31e-17 | 4.42e-13 |
| Malignant epithelial | C6 DNA-r p ECs | ADK | 0.902 | 0.556 | 0.309 | 5.99e-09 | 0.000202 |
| Malignant epithelial | C6 DNA-r p ECs | CKAP4 | 0.902 | 0.363 | 0.176 | 2.65e-07 | 0.00894 |
| Malignant epithelial | C6 DNA-r p ECs | DIMT1 | 0.902 | 0.355 | 0.174 | 1.03e-06 | 0.0346 |
| Malignant epithelial | C6 DNA-r p ECs | SNRNP25 | 0.901 | 0.621 | 0.266 | 1.73e-15 | 5.83e-11 |

| compartment | subcluster | gene | avg_log2FC | pct.1 | pct.2 | p_val | p_val_adj |
|---|---|---|---|---|---|---|---|
| Malignant epithelial | C6 DNA-r p ECs | NFIB | 0.9 | 0.476 | 0.218 | 2.59e-10 | 8.71e-06 |
| Malignant epithelial | C6 DNA-r p ECs | PXMP4 | 0.9 | 0.331 | 0.13 | 1.43e-09 | 4.82e-05 |
| Malignant epithelial | C6 DNA-r p ECs | SFR1 | 0.9 | 0.427 | 0.202 | 7.88e-09 | 0.000266 |
| Malignant epithelial | C6 DNA-r p ECs | RAD1 | 0.899 | 0.387 | 0.171 | 3.68e-09 | 0.000124 |
| Malignant epithelial | C6 DNA-r p ECs | SAE1 | 0.898 | 0.532 | 0.288 | 2.48e-09 | 8.35e-05 |
| Malignant epithelial | C6 DNA-r p ECs | VPS25 | 0.897 | 0.758 | 0.42 | 3.41e-13 | 1.15e-08 |
| Malignant epithelial | C6 DNA-r p ECs | EMP2 | 0.896 | 0.831 | 0.493 | 5.25e-16 | 1.77e-11 |
| Malignant epithelial | C6 DNA-r p ECs | MANSC1 | 0.896 | 0.395 | 0.191 | 4.28e-08 | 0.00144 |
| Malignant epithelial | C6 DNA-r p ECs | CPSF3 | 0.895 | 0.323 | 0.135 | 2.45e-08 | 0.000826 |
| Malignant epithelial | C6 DNA-r p ECs | NT5E | 0.894 | 0.782 | 0.453 | 3.94e-13 | 1.33e-08 |
| Malignant epithelial | C6 DNA-r p ECs | PNP | 0.894 | 0.427 | 0.222 | 2.98e-07 | 0.0101 |
| Malignant epithelial | C6 DNA-r p ECs | UBE2D4 | 0.894 | 0.298 | 0.133 | 5.35e-07 | 0.018 |
| Malignant epithelial | C6 DNA-r p ECs | BUD31 | 0.892 | 0.944 | 0.713 | 1.03e-19 | 3.49e-15 |
| Malignant epithelial | C6 DNA-r p ECs | LRP11 | 0.891 | 0.411 | 0.207 | 3.2e-08 | 0.00108 |
| Malignant epithelial | C6 DNA-r p ECs | H2AFV | 0.889 | 0.927 | 0.732 | 1.76e-16 | 5.94e-12 |
| Malignant epithelial | C6 DNA-r p ECs | LACTB2 | 0.889 | 0.621 | 0.333 | 4.51e-11 | 1.52e-06 |
| Malignant epithelial | C6 DNA-r p ECs | MTHFD2 | 0.889 | 0.573 | 0.333 | 1.99e-08 | 0.00067 |
| Malignant epithelial | C6 DNA-r p ECs | TP53I11 | 0.887 | 0.419 | 0.181 | 2.2e-10 | 7.41e-06 |
| Malignant epithelial | C6 DNA-r p ECs | CHCHD3 | 0.885 | 0.637 | 0.362 | 1.55e-10 | 5.22e-06 |
| Malignant epithelial | C6 DNA-r p ECs | PRNP | 0.885 | 0.605 | 0.358 | 4.63e-08 | 0.00156 |
| Malignant epithelial | C6 DNA-r p ECs | CREB3 | 0.882 | 0.476 | 0.246 | 3.19e-08 | 0.00108 |
| Malignant epithelial | C6 DNA-r p ECs | SMARCA2 | 0.882 | 0.258 | 0.108 | 7.92e-07 | 0.0267 |
| Malignant epithelial | C6 DNA-r p ECs | LY6E | 0.877 | 1.0 | 0.812 | 2.39e-14 | 8.06e-10 |
| Malignant epithelial | C6 DNA-r p ECs | NQO1 | 0.877 | 0.903 | 0.735 | 4.84e-09 | 0.000163 |
| Malignant epithelial | C6 DNA-r p ECs | KPNA2 | 0.876 | 0.435 | 0.243 | 1.08e-06 | 0.0365 |
| Malignant epithelial | C6 DNA-r p ECs | C12orf75 | 0.875 | 0.879 | 0.568 | 1.96e-17 | 6.61e-13 |
| Malignant epithelial | C6 DNA-r p ECs | CPVL | 0.875 | 0.298 | 0.121 | 2.53e-08 | 0.000853 |
| Malignant epithelial | C6 DNA-r p ECs | TSPAN15 | 0.874 | 0.677 | 0.366 | 1.51e-11 | 5.07e-07 |
| Malignant epithelial | C6 DNA-r p ECs | SQRDL | 0.873 | 0.685 | 0.424 | 1.09e-09 | 3.68e-05 |
| Malignant epithelial | C6 DNA-r p ECs | SZRD1 | 0.873 | 0.468 | 0.235 | 4.42e-09 | 0.000149 |
| Malignant epithelial | C6 DNA-r p ECs | TNFRSF12A | 0.872 | 0.871 | 0.627 | 3.63e-10 | 1.22e-05 |
| Malignant epithelial | C6 DNA-r p ECs | AURKAIP1 | 0.871 | 0.968 | 0.762 | 3.08e-20 | 1.04e-15 |
| Malignant epithelial | C6 DNA-r p ECs | NMRAL1 | 0.871 | 0.685 | 0.404 | 2.58e-10 | 8.7e-06 |
| Malignant epithelial | C6 DNA-r p ECs | KRT19 | 0.87 | 1.0 | 0.977 | 2.56e-12 | 8.61e-08 |
| Malignant epithelial | C6 DNA-r p ECs | SHROOM3 | 0.867 | 0.266 | 0.104 | 6.76e-08 | 0.00228 |
| Malignant epithelial | C6 DNA-r p ECs | TMEM242 | 0.866 | 0.395 | 0.209 | 5.57e-07 | 0.0188 |
| Malignant epithelial | C6 DNA-r p ECs | PCMT1 | 0.864 | 0.742 | 0.428 | 1.16e-12 | 3.92e-08 |
| Malignant epithelial | C6 DNA-r p ECs | PCDH1 | 0.864 | 0.508 | 0.253 | 2.17e-09 | 7.32e-05 |
| Malignant epithelial | C6 DNA-r p ECs | GNB2 | 0.863 | 0.976 | 0.713 | 8.9e-19 | 3e-14 |
| Malignant epithelial | C6 DNA-r p ECs | PARK7 | 0.863 | 0.935 | 0.747 | 9.85e-19 | 3.32e-14 |
| Malignant epithelial | C6 DNA-r p ECs | HNRNPAB | 0.863 | 0.766 | 0.566 | 1.31e-10 | 4.4e-06 |
| Malignant epithelial | C6 DNA-r p ECs | EFTUD2 | 0.861 | 0.371 | 0.175 | 1.51e-07 | 0.00508 |
| Malignant epithelial | C6 DNA-r p ECs | APIP | 0.859 | 0.468 | 0.272 | 1.48e-06 | 0.05 |
| Malignant epithelial | C6 DNA-r p ECs | TMEM92 | 0.858 | 0.347 | 0.132 | 2.15e-10 | 7.26e-06 |
| Malignant epithelial | C6 DNA-r p ECs | NOC2L | 0.858 | 0.532 | 0.275 | 7.82e-09 | 0.000264 |
| Malignant epithelial | C6 DNA-r p ECs | PSME2 | 0.857 | 0.992 | 0.839 | 2.47e-21 | 8.34e-17 |
| Malignant epithelial | C6 DNA-r p ECs | CPSF3L | 0.857 | 0.532 | 0.255 | 1.52e-10 | 5.12e-06 |
| Malignant epithelial | C6 DNA-r p ECs | C19orf48 | 0.856 | 0.734 | 0.483 | 3.29e-09 | 0.000111 |
| Malignant epithelial | C6 DNA-r p ECs | UQCC1 | 0.856 | 0.306 | 0.135 | 2.03e-07 | 0.00684 |
| Malignant epithelial | C6 DNA-r p ECs | FAM98B | 0.856 | 0.29 | 0.128 | 4.44e-07 | 0.015 |
| Malignant epithelial | C6 DNA-r p ECs | PDLIM1 | 0.855 | 0.919 | 0.609 | 2.18e-17 | 7.33e-13 |
| Malignant epithelial | C6 DNA-r p ECs | ANKHD1 | 0.855 | 0.298 | 0.137 | 1.32e-06 | 0.0445 |
| Malignant epithelial | C6 DNA-r p ECs | CALU | 0.854 | 0.605 | 0.343 | 7.75e-10 | 2.61e-05 |
| Malignant epithelial | C6 DNA-r p ECs | OGDH | 0.854 | 0.331 | 0.157 | 6.66e-07 | 0.0224 |
| Malignant epithelial | C6 DNA-r p ECs | RHEB | 0.853 | 0.895 | 0.74 | 1.94e-15 | 6.55e-11 |
| Malignant epithelial | C6 DNA-r p ECs | SRM | 0.853 | 0.452 | 0.25 | 5.39e-07 | 0.0182 |
| Malignant epithelial | C6 DNA-r p ECs | IFT52 | 0.853 | 0.331 | 0.157 | 9.06e-07 | 0.0305 |
| Malignant epithelial | C6 DNA-r p ECs | SYT13 | 0.852 | 0.419 | 0.183 | 6.03e-10 | 2.03e-05 |
| Malignant epithelial | C6 DNA-r p ECs | DERA | 0.851 | 0.5 | 0.247 | 1.59e-09 | 5.37e-05 |
| Malignant epithelial | C6 DNA-r p ECs | HDHD3 | 0.85 | 0.766 | 0.402 | 3.72e-15 | 1.25e-10 |
| Malignant epithelial | C6 DNA-r p ECs | NUDT8 | 0.849 | 0.589 | 0.267 | 1.07e-12 | 3.6e-08 |
| Malignant epithelial | C6 DNA-r p ECs | FKBP3 | 0.849 | 0.742 | 0.449 | 9.04e-11 | 3.05e-06 |
| Malignant epithelial | C6 DNA-r p ECs | CST6 | 0.849 | 0.516 | 0.307 | 8.29e-09 | 0.000279 |
| Malignant epithelial | C6 DNA-r p ECs | SRD5A1 | 0.846 | 0.379 | 0.162 | 2.51e-09 | 8.47e-05 |

| compartment | subcluster | gene | avg_log2FC | pct.1 | pct.2 | p_val | p_val_adj |
|---|---|---|---|---|---|---|---|
| Malignant epithelial | C6 DNA-r p ECs | C1orf174 | 0.846 | 0.331 | 0.15 | 1.74e-07 | 0.00587 |
| Malignant epithelial | C6 DNA-r p ECs | EPB41L1 | 0.845 | 0.435 | 0.21 | 1.74e-08 | 0.000587 |
| Malignant epithelial | C6 DNA-r p ECs | NASP | 0.844 | 0.613 | 0.318 | 1.77e-10 | 5.96e-06 |
| Malignant epithelial | C6 DNA-r p ECs | ERI3 | 0.843 | 0.387 | 0.2 | 8.94e-07 | 0.0301 |
| Malignant epithelial | C6 DNA-r p ECs | PPIH | 0.842 | 0.516 | 0.251 | 5.12e-10 | 1.72e-05 |
| Malignant epithelial | C6 DNA-r p ECs | HMGA1 | 0.84 | 0.968 | 0.843 | 4.95e-17 | 1.67e-12 |
| Malignant epithelial | C6 DNA-r p ECs | MCFD2 | 0.839 | 0.5 | 0.289 | 9.82e-07 | 0.0331 |
| Malignant epithelial | C6 DNA-r p ECs | CLTB | 0.836 | 0.935 | 0.703 | 8.61e-16 | 2.9e-11 |
| Malignant epithelial | C6 DNA-r p ECs | PRIM2 | 0.836 | 0.226 | 0.091 | 1.25e-06 | 0.042 |
| Malignant epithelial | C6 DNA-r p ECs | CARHSP1 | 0.835 | 0.927 | 0.675 | 1.58e-15 | 5.31e-11 |
| Malignant epithelial | C6 DNA-r p ECs | TBRG4 | 0.833 | 0.548 | 0.264 | 3.41e-10 | 1.15e-05 |
| Malignant epithelial | C6 DNA-r p ECs | MMAB | 0.833 | 0.5 | 0.249 | 1.32e-09 | 4.44e-05 |
| Malignant epithelial | C6 DNA-r p ECs | RNASE1 | 0.832 | 0.548 | 0.235 | 7.81e-14 | 2.63e-09 |
| Malignant epithelial | C6 DNA-r p ECs | SUPT16H | 0.832 | 0.556 | 0.302 | 8.59e-09 | 0.000289 |
| Malignant epithelial | C6 DNA-r p ECs | SMUG1 | 0.832 | 0.581 | 0.321 | 8.96e-09 | 0.000302 |
| Malignant epithelial | C6 DNA-r p ECs | RALY | 0.83 | 0.952 | 0.717 | 2.38e-18 | 8.02e-14 |
| Malignant epithelial | C6 DNA-r p ECs | YEATS4 | 0.83 | 0.524 | 0.276 | 1.39e-08 | 0.000469 |
| Malignant epithelial | C6 DNA-r p ECs | LYPLA2 | 0.829 | 0.54 | 0.288 | 1.31e-08 | 0.000442 |
| Malignant epithelial | C6 DNA-r p ECs | TMEM203 | 0.829 | 0.54 | 0.294 | 6.7e-08 | 0.00226 |
| Malignant epithelial | C6 DNA-r p ECs | ACAD9 | 0.827 | 0.226 | 0.088 | 5.7e-07 | 0.0192 |
| Malignant epithelial | C6 DNA-r p ECs | ZHX1 | 0.826 | 0.331 | 0.139 | 3.31e-08 | 0.00112 |
| Malignant epithelial | C6 DNA-r p ECs | VAPB | 0.826 | 0.427 | 0.225 | 6.16e-07 | 0.0208 |
| Malignant epithelial | C6 DNA-r p ECs | TALDO1 | 0.825 | 0.944 | 0.712 | 1.5e-18 | 5.05e-14 |
| Malignant epithelial | C6 DNA-r p ECs | IFI35 | 0.824 | 0.661 | 0.391 | 1.12e-09 | 3.79e-05 |
| Malignant epithelial | C6 DNA-r p ECs | CRIP1 | 0.822 | 0.968 | 0.789 | 6.6e-10 | 2.22e-05 |
| Malignant epithelial | C6 DNA-r p ECs | SH2D4A | 0.822 | 0.339 | 0.16 | 6.3e-07 | 0.0212 |
| Malignant epithelial | C6 DNA-r p ECs | COPS6 | 0.821 | 0.903 | 0.646 | 3.66e-15 | 1.23e-10 |
| Malignant epithelial | C6 DNA-r p ECs | FIBP | 0.821 | 0.677 | 0.391 | 8.15e-10 | 2.75e-05 |
| Malignant epithelial | C6 DNA-r p ECs | DDX56 | 0.819 | 0.508 | 0.279 | 3.42e-07 | 0.0115 |
| Malignant epithelial | C6 DNA-r p ECs | PGD | 0.819 | 0.331 | 0.16 | 1.46e-06 | 0.0491 |
| Malignant epithelial | C6 DNA-r p ECs | CDK11B | 0.817 | 0.427 | 0.199 | 4.83e-09 | 0.000163 |
| Malignant epithelial | C6 DNA-r p ECs | RAB8A | 0.815 | 0.653 | 0.354 | 3.39e-10 | 1.14e-05 |
| Malignant epithelial | C6 DNA-r p ECs | LRRC47 | 0.815 | 0.355 | 0.167 | 3.7e-07 | 0.0125 |
| Malignant epithelial | C6 DNA-r p ECs | TMA16 | 0.814 | 0.565 | 0.298 | 2.87e-10 | 9.68e-06 |
| Malignant epithelial | C6 DNA-r p ECs | NUDT5 | 0.814 | 0.677 | 0.439 | 9.28e-09 | 0.000313 |
| Malignant epithelial | C6 DNA-r p ECs | SSU72 | 0.813 | 0.919 | 0.668 | 1.53e-16 | 5.15e-12 |
| Malignant epithelial | C6 DNA-r p ECs | AP1S1 | 0.812 | 0.879 | 0.567 | 8.94e-14 | 3.01e-09 |
| Malignant epithelial | C6 DNA-r p ECs | PSMD14 | 0.812 | 0.589 | 0.337 | 5.47e-08 | 0.00184 |
| Malignant epithelial | C6 DNA-r p ECs | PTPRN2 | 0.812 | 0.274 | 0.114 | 2.75e-07 | 0.00928 |
| Malignant epithelial | C6 DNA-r p ECs | COMMD8 | 0.811 | 0.298 | 0.135 | 9.8e-07 | 0.033 |
| Malignant epithelial | C6 DNA-r p ECs | BARD1 | 0.81 | 0.266 | 0.102 | 4.01e-08 | 0.00135 |
| Malignant epithelial | C6 DNA-r p ECs | LLGL2 | 0.808 | 0.718 | 0.407 | 7.09e-11 | 2.39e-06 |
| Malignant epithelial | C6 DNA-r p ECs | GSTZ1 | 0.804 | 0.444 | 0.192 | 5.01e-10 | 1.69e-05 |
| Malignant epithelial | C6 DNA-r p ECs | YIPF2 | 0.802 | 0.54 | 0.305 | 5.36e-08 | 0.00181 |
| Malignant epithelial | C6 DNA-r p ECs | CDC42 | 0.801 | 0.903 | 0.708 | 3.55e-12 | 1.2e-07 |
| Malignant epithelial | C6 DNA-r p ECs | CAPG | 0.801 | 0.952 | 0.772 | 4.45e-10 | 1.5e-05 |
| Malignant epithelial | C6 DNA-r p ECs | GMPS | 0.799 | 0.435 | 0.23 | 3.35e-07 | 0.0113 |
| Malignant epithelial | C6 DNA-r p ECs | AGR2 | 0.798 | 0.96 | 0.558 | 1.59e-22 | 5.37e-18 |
| Malignant epithelial | C6 DNA-r p ECs | ECI1 | 0.797 | 0.742 | 0.432 | 1.64e-11 | 5.51e-07 |
| Malignant epithelial | C6 DNA-r p ECs | AHCY | 0.797 | 0.79 | 0.515 | 1.25e-10 | 4.22e-06 |
| Malignant epithelial | C6 DNA-r p ECs | PPP1R7 | 0.797 | 0.71 | 0.447 | 4.35e-09 | 0.000146 |
| Malignant epithelial | C6 DNA-r p ECs | NMI | 0.797 | 0.532 | 0.291 | 1.33e-08 | 0.000448 |
| Malignant epithelial | C6 DNA-r p ECs | NAGK | 0.796 | 0.347 | 0.17 | 1.38e-06 | 0.0466 |
| Malignant epithelial | C6 DNA-r p ECs | S100P | 0.795 | 0.927 | 0.546 | 2.81e-16 | 9.46e-12 |
| Malignant epithelial | C6 DNA-r p ECs | KPNB1 | 0.793 | 0.726 | 0.483 | 7.64e-09 | 0.000257 |
| Malignant epithelial | C6 DNA-r p ECs | UBE2H | 0.793 | 0.597 | 0.361 | 9.65e-08 | 0.00325 |
| Malignant epithelial | C6 DNA-r p ECs | PROSC | 0.793 | 0.5 | 0.285 | 6.35e-07 | 0.0214 |
| Malignant epithelial | C6 DNA-r p ECs | ITGB4 | 0.792 | 0.774 | 0.453 | 8.2e-12 | 2.76e-07 |
| Malignant epithelial | C6 DNA-r p ECs | TPX2 | 0.789 | 0.419 | 0.148 | 2.47e-14 | 8.32e-10 |
| Malignant epithelial | C6 DNA-r p ECs | CDK4 | 0.789 | 0.75 | 0.492 | 1.49e-09 | 5.04e-05 |
| Malignant epithelial | C6 DNA-r p ECs | KNSTRN | 0.789 | 0.298 | 0.114 | 4.8e-09 | 0.000162 |
| Malignant epithelial | C6 DNA-r p ECs | SWI5 | 0.789 | 0.476 | 0.248 | 7.33e-08 | 0.00247 |
| Malignant epithelial | C6 DNA-r p ECs | CSE1L | 0.789 | 0.492 | 0.265 | 8.79e-08 | 0.00296 |
| Malignant epithelial | C6 DNA-r p ECs | USP48 | 0.789 | 0.331 | 0.156 | 7.29e-07 | 0.0246 |

| compartment | subcluster | gene | avg_log2FC | pct.1 | pct.2 | p_val | p_val_adj |
|---|---|---|---|---|---|---|---|
| Malignant epithelial | C6 DNA-r p ECs | RMND5B | 0.788 | 0.339 | 0.155 | 2.85e-07 | 0.00962 |
| Malignant epithelial | C6 DNA-r p ECs | XRCC1 | 0.787 | 0.339 | 0.146 | 4.21e-08 | 0.00142 |
| Malignant epithelial | C6 DNA-r p ECs | TMEM55B | 0.787 | 0.46 | 0.231 | 5.76e-08 | 0.00194 |
| Malignant epithelial | C6 DNA-r p ECs | PSMB9 | 0.786 | 0.976 | 0.772 | 1.42e-15 | 4.8e-11 |
| Malignant epithelial | C6 DNA-r p ECs | TTC7A | 0.786 | 0.242 | 0.098 | 5.82e-07 | 0.0196 |
| Malignant epithelial | C6 DNA-r p ECs | PSMC5 | 0.784 | 0.823 | 0.577 | 1.64e-10 | 5.53e-06 |
| Malignant epithelial | C6 DNA-r p ECs | ITGB1BP1 | 0.784 | 0.653 | 0.384 | 5.19e-09 | 0.000175 |
| Malignant epithelial | C6 DNA-r p ECs | MAEA | 0.784 | 0.387 | 0.191 | 3.99e-07 | 0.0134 |
| Malignant epithelial | C6 DNA-r p ECs | HLA-DRB1 | 0.783 | 0.903 | 0.609 | 5.12e-13 | 1.73e-08 |
| Malignant epithelial | C6 DNA-r p ECs | FRMD4B | 0.783 | 0.363 | 0.175 | 2.49e-07 | 0.00838 |
| Malignant epithelial | C6 DNA-r p ECs | ANXA13 | 0.782 | 0.25 | 0.099 | 2.36e-07 | 0.00796 |
| Malignant epithelial | C6 DNA-r p ECs | ZDHHC16 | 0.782 | 0.339 | 0.164 | 9.29e-07 | 0.0313 |
| Malignant epithelial | C6 DNA-r p ECs | CBR1 | 0.781 | 0.669 | 0.349 | 6.25e-11 | 2.11e-06 |
| Malignant epithelial | C6 DNA-r p ECs | CDH17 | 0.779 | 0.581 | 0.301 | 4.19e-10 | 1.41e-05 |
| Malignant epithelial | C6 DNA-r p ECs | TBCD | 0.775 | 0.323 | 0.136 | 4.39e-08 | 0.00148 |
| Malignant epithelial | C6 DNA-r p ECs | FHL2 | 0.774 | 0.847 | 0.507 | 2.38e-13 | 8.01e-09 |
| Malignant epithelial | C6 DNA-r p ECs | GMDS | 0.774 | 0.605 | 0.31 | 2.27e-10 | 7.66e-06 |
| Malignant epithelial | C6 DNA-r p ECs | CMIP | 0.774 | 0.444 | 0.212 | 2.13e-08 | 0.000719 |
| Malignant epithelial | C6 DNA-r p ECs | GSDMD | 0.773 | 0.863 | 0.54 | 2.6e-13 | 8.75e-09 |
| Malignant epithelial | C6 DNA-r p ECs | NUF2 | 0.773 | 0.202 | 0.063 | 7.27e-09 | 0.000245 |
| Malignant epithelial | C6 DNA-r p ECs | CENPA | 0.773 | 0.21 | 0.078 | 5.76e-07 | 0.0194 |
| Malignant epithelial | C6 DNA-r p ECs | PTGES2 | 0.771 | 0.815 | 0.449 | 2.89e-13 | 9.75e-09 |
| Malignant epithelial | C6 DNA-r p ECs | KXD1 | 0.771 | 0.831 | 0.524 | 4.61e-11 | 1.55e-06 |
| Malignant epithelial | C6 DNA-r p ECs | PDZD11 | 0.771 | 0.742 | 0.484 | 1.02e-09 | 3.43e-05 |
| Malignant epithelial | C6 DNA-r p ECs | TMEM70 | 0.77 | 0.532 | 0.29 | 4.89e-08 | 0.00165 |
| Malignant epithelial | C6 DNA-r p ECs | SH3KBP1 | 0.769 | 0.758 | 0.447 | 1.1e-10 | 3.7e-06 |
| Malignant epithelial | C6 DNA-r p ECs | IFI27 | 0.768 | 1.0 | 0.942 | 1.2e-15 | 4.04e-11 |
| Malignant epithelial | C6 DNA-r p ECs | NUB1 | 0.768 | 0.573 | 0.313 | 2.52e-08 | 0.000848 |
| Malignant epithelial | C6 DNA-r p ECs | PPT1 | 0.768 | 0.419 | 0.216 | 3.76e-07 | 0.0127 |
| Malignant epithelial | C6 DNA-r p ECs | SNRPB | 0.767 | 0.992 | 0.886 | 1.5e-20 | 5.04e-16 |
| Malignant epithelial | C6 DNA-r p ECs | UACA | 0.767 | 0.524 | 0.249 | 1.32e-10 | 4.45e-06 |
| Malignant epithelial | C6 DNA-r p ECs | SNRPA1 | 0.767 | 0.685 | 0.445 | 1.32e-08 | 0.000446 |
| Malignant epithelial | C6 DNA-r p ECs | GOPC | 0.767 | 0.484 | 0.246 | 2.42e-08 | 0.000817 |
| Malignant epithelial | C6 DNA-r p ECs | PPID | 0.767 | 0.419 | 0.212 | 2.12e-07 | 0.00714 |
| Malignant epithelial | C6 DNA-r p ECs | MIS18BP1 | 0.766 | 0.315 | 0.142 | 5.69e-07 | 0.0192 |
| Malignant epithelial | C6 DNA-r p ECs | C1orf21 | 0.765 | 0.516 | 0.257 | 2.67e-09 | 9.01e-05 |
| Malignant epithelial | C6 DNA-r p ECs | LRRC1 | 0.764 | 0.379 | 0.179 | 8.56e-08 | 0.00289 |
| Malignant epithelial | C6 DNA-r p ECs | UNC13D | 0.764 | 0.476 | 0.254 | 9.77e-08 | 0.00329 |
| Malignant epithelial | C6 DNA-r p ECs | VKORC1L1 | 0.764 | 0.379 | 0.178 | 1.63e-07 | 0.0055 |
| Malignant epithelial | C6 DNA-r p ECs | EPSTI1 | 0.763 | 0.887 | 0.574 | 1.05e-12 | 3.55e-08 |
| Malignant epithelial | C6 DNA-r p ECs | LGALS3BP | 0.763 | 0.919 | 0.741 | 9.2e-12 | 3.1e-07 |
| Malignant epithelial | C6 DNA-r p ECs | PLGRKT | 0.763 | 0.621 | 0.368 | 4.74e-08 | 0.0016 |
| Malignant epithelial | C6 DNA-r p ECs | CCT5 | 0.762 | 0.855 | 0.542 | 4.85e-12 | 1.63e-07 |
| Malignant epithelial | C6 DNA-r p ECs | GJB2 | 0.762 | 0.387 | 0.184 | 9.14e-08 | 0.00308 |
| Malignant epithelial | C6 DNA-r p ECs | PARP1 | 0.762 | 0.435 | 0.222 | 3.82e-07 | 0.0129 |
| Malignant epithelial | C6 DNA-r p ECs | CALM1 | 0.761 | 0.992 | 0.912 | 3.1e-23 | 1.04e-18 |
| Malignant epithelial | C6 DNA-r p ECs | FKBP1A | 0.761 | 0.96 | 0.824 | 5.74e-15 | 1.94e-10 |
| Malignant epithelial | C6 DNA-r p ECs | NELFB | 0.761 | 0.516 | 0.23 | 3.19e-11 | 1.07e-06 |
| Malignant epithelial | C6 DNA-r p ECs | MALL | 0.761 | 0.718 | 0.463 | 4.52e-09 | 0.000152 |
| Malignant epithelial | C6 DNA-r p ECs | DCTPP1 | 0.761 | 0.694 | 0.44 | 3.04e-08 | 0.00102 |
| Malignant epithelial | C6 DNA-r p ECs | RAB20 | 0.757 | 0.484 | 0.215 | 1.93e-10 | 6.5e-06 |
| Malignant epithelial | C6 DNA-r p ECs | ASPH | 0.756 | 0.903 | 0.645 | 1.79e-09 | 6.02e-05 |
| Malignant epithelial | C6 DNA-r p ECs | ADGRG1 | 0.756 | 0.589 | 0.32 | 5.72e-09 | 0.000193 |
| Malignant epithelial | C6 DNA-r p ECs | PDE4C | 0.755 | 0.54 | 0.288 | 5.32e-09 | 0.000179 |
| Malignant epithelial | C6 DNA-r p ECs | GSN | 0.755 | 0.895 | 0.652 | 7.75e-09 | 0.000261 |
| Malignant epithelial | C6 DNA-r p ECs | TPM1 | 0.754 | 0.879 | 0.592 | 4.04e-13 | 1.36e-08 |
| Malignant epithelial | C6 DNA-r p ECs | HRAS | 0.753 | 0.589 | 0.311 | 5.06e-09 | 0.00017 |
| Malignant epithelial | C6 DNA-r p ECs | MED7 | 0.753 | 0.274 | 0.116 | 6.78e-07 | 0.0229 |
| Malignant epithelial | C6 DNA-r p ECs | STRA13 | 0.752 | 0.815 | 0.555 | 6.04e-11 | 2.04e-06 |
| Malignant epithelial | C6 DNA-r p ECs | MZT1 | 0.752 | 0.556 | 0.291 | 7.96e-10 | 2.68e-05 |
| Malignant epithelial | C6 DNA-r p ECs | CAPZB | 0.752 | 0.847 | 0.593 | 2.81e-09 | 9.47e-05 |
| Malignant epithelial | C6 DNA-r p ECs | PRPF19 | 0.75 | 0.548 | 0.288 | 4.41e-09 | 0.000148 |
| Malignant epithelial | C6 DNA-r p ECs | SMIM15 | 0.749 | 0.71 | 0.415 | 4.18e-10 | 1.41e-05 |
| Malignant epithelial | C6 DNA-r p ECs | BCL2L12 | 0.747 | 0.516 | 0.275 | 1.07e-07 | 0.00362 |

| compartment | subcluster | gene | avg_log2FC | pct.1 | pct.2 | p_val | p_val_adj |
|---|---|---|---|---|---|---|---|
| Malignant epithelial | C6 DNA-r p ECs | PSMC3 | 0.746 | 0.815 | 0.507 | 5.15e-12 | 1.74e-07 |
| Malignant epithelial | C6 DNA-r p ECs | PLA2G16 | 0.746 | 0.927 | 0.619 | 7.55e-12 | 2.54e-07 |
| Malignant epithelial | C6 DNA-r p ECs | PSMA3 | 0.746 | 0.895 | 0.629 | 1.71e-10 | 5.75e-06 |
| Malignant epithelial | C6 DNA-r p ECs | HSPB11 | 0.746 | 0.589 | 0.336 | 1.32e-08 | 0.000444 |
| Malignant epithelial | C6 DNA-r p ECs | ABHD12 | 0.746 | 0.629 | 0.401 | 1.51e-08 | 0.000508 |
| Malignant epithelial | C6 DNA-r p ECs | LGALS4 | 0.745 | 0.935 | 0.489 | 6.28e-19 | 2.12e-14 |
| Malignant epithelial | C6 DNA-r p ECs | DUSP6 | 0.745 | 0.5 | 0.275 | 3.05e-07 | 0.0103 |
| Malignant epithelial | C6 DNA-r p ECs | GSTO1 | 0.744 | 0.944 | 0.694 | 5.88e-13 | 1.98e-08 |
| Malignant epithelial | C6 DNA-r p ECs | PCK2 | 0.744 | 0.379 | 0.185 | 4.61e-07 | 0.0155 |
| Malignant epithelial | C6 DNA-r p ECs | RNASEH2C | 0.741 | 0.823 | 0.548 | 1.27e-11 | 4.29e-07 |
| Malignant epithelial | C6 DNA-r p ECs | DDX41 | 0.739 | 0.435 | 0.222 | 5.38e-07 | 0.0181 |
| Malignant epithelial | C6 DNA-r p ECs | VOPP1 | 0.736 | 0.605 | 0.339 | 1.24e-08 | 0.000418 |
| Malignant epithelial | C6 DNA-r p ECs | SSRP1 | 0.735 | 0.621 | 0.379 | 4.91e-08 | 0.00165 |
| Malignant epithelial | C6 DNA-r p ECs | COBL | 0.735 | 0.298 | 0.134 | 9.6e-07 | 0.0323 |
| Malignant epithelial | C6 DNA-r p ECs | TSR2 | 0.734 | 0.581 | 0.308 | 2.12e-09 | 7.15e-05 |
| Malignant epithelial | C6 DNA-r p ECs | ISCA2 | 0.734 | 0.565 | 0.314 | 1.74e-07 | 0.00585 |
| Malignant epithelial | C6 DNA-r p ECs | KRT6A | 0.733 | 0.355 | 0.092 | 1.32e-22 | 4.46e-18 |
| Malignant epithelial | C6 DNA-r p ECs | PSMB7 | 0.733 | 0.96 | 0.722 | 2.82e-15 | 9.5e-11 |
| Malignant epithelial | C6 DNA-r p ECs | STOML2 | 0.733 | 0.879 | 0.613 | 5.95e-12 | 2e-07 |
| Malignant epithelial | C6 DNA-r p ECs | TRIM16 | 0.733 | 0.355 | 0.161 | 8.32e-08 | 0.0028 |
| Malignant epithelial | C6 DNA-r p ECs | SPTSSA | 0.732 | 0.815 | 0.55 | 1.08e-08 | 0.000364 |
| Malignant epithelial | C6 DNA-r p ECs | CCDC47 | 0.731 | 0.516 | 0.282 | 2.26e-07 | 0.00763 |
| Malignant epithelial | C6 DNA-r p ECs | CRACR2B | 0.73 | 0.492 | 0.253 | 4.21e-08 | 0.00142 |
| Malignant epithelial | C6 DNA-r p ECs | UBE2L6 | 0.729 | 0.806 | 0.501 | 1.79e-10 | 6.03e-06 |
| Malignant epithelial | C6 DNA-r p ECs | CNP | 0.729 | 0.5 | 0.252 | 6.87e-09 | 0.000231 |
| Malignant epithelial | C6 DNA-r p ECs | ANP32B | 0.728 | 0.935 | 0.801 | 3.83e-15 | 1.29e-10 |
| Malignant epithelial | C6 DNA-r p ECs | PSMB10 | 0.728 | 0.766 | 0.44 | 2.31e-11 | 7.79e-07 |
| Malignant epithelial | C6 DNA-r p ECs | MAGOHB | 0.727 | 0.669 | 0.349 | 9.93e-11 | 3.35e-06 |
| Malignant epithelial | C6 DNA-r p ECs | THOC6 | 0.726 | 0.573 | 0.298 | 9.79e-10 | 3.3e-05 |
| Malignant epithelial | C6 DNA-r p ECs | COLGALT1 | 0.726 | 0.419 | 0.195 | 3.5e-08 | 0.00118 |
| Malignant epithelial | C6 DNA-r p ECs | CLN6 | 0.726 | 0.484 | 0.28 | 7.75e-07 | 0.0261 |
| Malignant epithelial | C6 DNA-r p ECs | TMEM14A | 0.724 | 0.71 | 0.44 | 5.51e-09 | 0.000186 |
| Malignant epithelial | C6 DNA-r p ECs | SERPINB5 | 0.723 | 0.629 | 0.305 | 4.11e-13 | 1.39e-08 |
| Malignant epithelial | C6 DNA-r p ECs | PPP6R1 | 0.723 | 0.371 | 0.183 | 7.72e-07 | 0.026 |
| Malignant epithelial | C6 DNA-r p ECs | HLA-DRA | 0.721 | 0.935 | 0.753 | 5.01e-11 | 1.69e-06 |
| Malignant epithelial | C6 DNA-r p ECs | MAPK13 | 0.721 | 0.685 | 0.374 | 5.07e-10 | 1.71e-05 |
| Malignant epithelial | C6 DNA-r p ECs | NDUFS8 | 0.72 | 0.911 | 0.685 | 1.72e-11 | 5.78e-07 |
| Malignant epithelial | C6 DNA-r p ECs | MYO19 | 0.72 | 0.379 | 0.185 | 3.31e-07 | 0.0112 |
| Malignant epithelial | C6 DNA-r p ECs | PTTG1 | 0.719 | 0.573 | 0.313 | 1.08e-09 | 3.64e-05 |
| Malignant epithelial | C6 DNA-r p ECs | ZCCHC9 | 0.719 | 0.444 | 0.22 | 2.93e-07 | 0.00989 |
| Malignant epithelial | C6 DNA-r p ECs | HMGB1 | 0.718 | 1.0 | 0.934 | 4.75e-19 | 1.6e-14 |
| Malignant epithelial | C6 DNA-r p ECs | GLYCTK | 0.718 | 0.266 | 0.11 | 6.68e-07 | 0.0225 |
| Malignant epithelial | C6 DNA-r p ECs | RHBDD2 | 0.717 | 0.669 | 0.403 | 1.31e-08 | 0.000442 |
| Malignant epithelial | C6 DNA-r p ECs | MPC1 | 0.717 | 0.54 | 0.3 | 4.33e-08 | 0.00146 |
| Malignant epithelial | C6 DNA-r p ECs | ASCC3 | 0.717 | 0.323 | 0.15 | 1.14e-06 | 0.0384 |
| Malignant epithelial | C6 DNA-r p ECs | RAD23A | 0.716 | 0.887 | 0.654 | 3.52e-13 | 1.18e-08 |
| Malignant epithelial | C6 DNA-r p ECs | HNRNPD | 0.716 | 0.79 | 0.512 | 5.31e-10 | 1.79e-05 |
| Malignant epithelial | C6 DNA-r p ECs | TPRKB | 0.716 | 0.758 | 0.493 | 1.15e-09 | 3.87e-05 |
| Malignant epithelial | C6 DNA-r p ECs | CD74 | 0.715 | 0.992 | 0.895 | 7.21e-15 | 2.43e-10 |
| Malignant epithelial | C6 DNA-r p ECs | POLR2G | 0.715 | 0.694 | 0.45 | 1.08e-07 | 0.00364 |
| Malignant epithelial | C6 DNA-r p ECs | CAMTA1 | 0.714 | 0.815 | 0.572 | 3.21e-11 | 1.08e-06 |
| Malignant epithelial | C6 DNA-r p ECs | ACTN1 | 0.713 | 0.637 | 0.373 | 1.72e-08 | 0.000579 |
| Malignant epithelial | C6 DNA-r p ECs | ARPC1B | 0.712 | 0.927 | 0.762 | 2.73e-12 | 9.2e-08 |
| Malignant epithelial | C6 DNA-r p ECs | H2AFY | 0.711 | 0.903 | 0.669 | 7.49e-12 | 2.52e-07 |
| Malignant epithelial | C6 DNA-r p ECs | NPSR1 | 0.709 | 0.548 | 0.297 | 2.57e-08 | 0.000866 |
| Malignant epithelial | C6 DNA-r p ECs | AP3B1 | 0.709 | 0.306 | 0.138 | 8.5e-07 | 0.0287 |
| Malignant epithelial | C6 DNA-r p ECs | UBXN2B | 0.708 | 0.266 | 0.111 | 6.55e-07 | 0.0221 |
| Malignant epithelial | C6 DNA-r p ECs | UBE2J2 | 0.706 | 0.46 | 0.234 | 1.19e-07 | 0.00401 |
| Malignant epithelial | C6 DNA-r p ECs | CDCA7 | 0.705 | 0.452 | 0.227 | 9.82e-08 | 0.00331 |
| Malignant epithelial | C6 DNA-r p ECs | NFYB | 0.705 | 0.419 | 0.207 | 1.88e-07 | 0.00634 |
| Malignant epithelial | C6 DNA-r p ECs | BST2 | 0.704 | 0.968 | 0.747 | 6.38e-12 | 2.15e-07 |
| Malignant epithelial | C6 DNA-r p ECs | LRRC59 | 0.704 | 0.702 | 0.413 | 1.27e-09 | 4.3e-05 |
| Malignant epithelial | C6 DNA-r p ECs | TOMM40 | 0.704 | 0.702 | 0.438 | 9.51e-09 | 0.00032 |
| Malignant epithelial | C6 DNA-r p ECs | FUCA2 | 0.704 | 0.54 | 0.295 | 1.18e-07 | 0.00397 |

| compartment | subcluster | gene | avg_log2FC | pct.1 | pct.2 | p_val | p_val_adj |
|---|---|---|---|---|---|---|---|
| Malignant epithelial | C6 DNA-r p ECs | MRPL22 | 0.701 | 0.79 | 0.552 | 7.1e-09 | 0.000239 |
| Malignant epithelial | C6 DNA-r p ECs | CMSS1 | 0.701 | 0.516 | 0.28 | 9.43e-08 | 0.00318 |
| Malignant epithelial | C6 DNA-r p ECs | ADRM1 | 0.699 | 0.903 | 0.658 | 5.56e-13 | 1.87e-08 |
| Malignant epithelial | C6 DNA-r p ECs | TUBA1C | 0.699 | 0.903 | 0.704 | 8.18e-11 | 2.76e-06 |
| Malignant epithelial | C6 DNA-r p ECs | UTP23 | 0.699 | 0.452 | 0.229 | 2.68e-07 | 0.00902 |
| Malignant epithelial | C6 DNA-r p ECs | ANXA4 | 0.695 | 0.806 | 0.579 | 1.81e-09 | 6.11e-05 |
| Malignant epithelial | C6 DNA-r p ECs | NQO2 | 0.695 | 0.395 | 0.178 | 2.06e-08 | 0.000693 |
| Malignant epithelial | C6 DNA-r p ECs | CYC1 | 0.694 | 0.984 | 0.76 | 3.44e-13 | 1.16e-08 |
| Malignant epithelial | C6 DNA-r p ECs | TES | 0.694 | 0.661 | 0.412 | 6.5e-07 | 0.0219 |
| Malignant epithelial | C6 DNA-r p ECs | ITM2B | 0.692 | 0.976 | 0.776 | 1.27e-13 | 4.27e-09 |
| Malignant epithelial | C6 DNA-r p ECs | OBFC1 | 0.69 | 0.452 | 0.21 | 1.05e-08 | 0.000352 |
| Malignant epithelial | C6 DNA-r p ECs | CTDSP1 | 0.69 | 0.508 | 0.26 | 7.69e-08 | 0.00259 |
| Malignant epithelial | C6 DNA-r p ECs | APLP2 | 0.689 | 0.903 | 0.697 | 1.11e-06 | 0.0374 |
| Malignant epithelial | C6 DNA-r p ECs | EFNB1 | 0.688 | 0.484 | 0.218 | 1.92e-10 | 6.48e-06 |
| Malignant epithelial | C6 DNA-r p ECs | CHRAC1 | 0.688 | 0.669 | 0.409 | 2.19e-08 | 0.000737 |
| Malignant epithelial | C6 DNA-r p ECs | C15orf40 | 0.688 | 0.492 | 0.251 | 7.53e-08 | 0.00254 |
| Malignant epithelial | C6 DNA-r p ECs | SH3GL1 | 0.687 | 0.508 | 0.25 | 6.74e-09 | 0.000227 |
| Malignant epithelial | C6 DNA-r p ECs | RPS27L | 0.686 | 0.968 | 0.839 | 2.82e-15 | 9.52e-11 |
| Malignant epithelial | C6 DNA-r p ECs | ATP6V1D | 0.686 | 0.653 | 0.394 | 3.02e-08 | 0.00102 |
| Malignant epithelial | C6 DNA-r p ECs | FUT3 | 0.686 | 0.282 | 0.12 | 3.94e-07 | 0.0133 |
| Malignant epithelial | C6 DNA-r p ECs | PA2G4 | 0.685 | 0.927 | 0.703 | 6.97e-13 | 2.35e-08 |
| Malignant epithelial | C6 DNA-r p ECs | UBE2S | 0.685 | 0.774 | 0.523 | 1.61e-10 | 5.43e-06 |
| Malignant epithelial | C6 DNA-r p ECs | TAGLN2 | 0.683 | 0.944 | 0.821 | 2.08e-13 | 7.02e-09 |
| Malignant epithelial | C6 DNA-r p ECs | BCAP29 | 0.683 | 0.565 | 0.287 | 1.12e-09 | 3.78e-05 |
| Malignant epithelial | C6 DNA-r p ECs | CST3 | 0.682 | 1.0 | 0.975 | 3.65e-18 | 1.23e-13 |
| Malignant epithelial | C6 DNA-r p ECs | SLC25A5 | 0.682 | 0.968 | 0.885 | 5.46e-09 | 0.000184 |
| Malignant epithelial | C6 DNA-r p ECs | MLXIP | 0.682 | 0.597 | 0.314 | 8.16e-09 | 0.000275 |
| Malignant epithelial | C6 DNA-r p ECs | PPP2CA | 0.682 | 0.677 | 0.433 | 3.78e-08 | 0.00127 |
| Malignant epithelial | C6 DNA-r p ECs | EIF4H | 0.679 | 0.847 | 0.58 | 1.08e-09 | 3.63e-05 |
| Malignant epithelial | C6 DNA-r p ECs | TXN | 0.678 | 1.0 | 0.986 | 8.01e-19 | 2.7e-14 |
| Malignant epithelial | C6 DNA-r p ECs | DDOST | 0.678 | 0.694 | 0.433 | 2.33e-08 | 0.000784 |
| Malignant epithelial | C6 DNA-r p ECs | PKM | 0.677 | 0.992 | 0.908 | 2.51e-13 | 8.45e-09 |
| Malignant epithelial | C6 DNA-r p ECs | SAR1B | 0.677 | 0.597 | 0.292 | 4.56e-10 | 1.54e-05 |
| Malignant epithelial | C6 DNA-r p ECs | THYN1 | 0.677 | 0.54 | 0.313 | 3.57e-07 | 0.012 |
| Malignant epithelial | C6 DNA-r p ECs | VPS29 | 0.674 | 0.887 | 0.638 | 1.31e-10 | 4.43e-06 |
| Malignant epithelial | C6 DNA-r p ECs | GANAB | 0.673 | 0.556 | 0.322 | 2.2e-07 | 0.00742 |
| Malignant epithelial | C6 DNA-r p ECs | TMEM223 | 0.673 | 0.556 | 0.333 | 1.25e-06 | 0.042 |
| Malignant epithelial | C6 DNA-r p ECs | VMA21 | 0.672 | 0.605 | 0.348 | 1.25e-07 | 0.00423 |
| Malignant epithelial | C6 DNA-r p ECs | LSM3 | 0.671 | 0.831 | 0.609 | 5.28e-09 | 0.000178 |
| Malignant epithelial | C6 DNA-r p ECs | FGFR4 | 0.67 | 0.282 | 0.122 | 1.19e-06 | 0.0401 |
| Malignant epithelial | C6 DNA-r p ECs | LRRC8A | 0.667 | 0.508 | 0.286 | 1.36e-06 | 0.0459 |
| Malignant epithelial | C6 DNA-r p ECs | COPE | 0.666 | 0.984 | 0.827 | 2.99e-16 | 1.01e-11 |
| Malignant epithelial | C6 DNA-r p ECs | ERP44 | 0.665 | 0.718 | 0.431 | 6.53e-10 | 2.2e-05 |
| Malignant epithelial | C6 DNA-r p ECs | TMEM50A | 0.665 | 0.831 | 0.622 | 3.73e-08 | 0.00126 |
| Malignant epithelial | C6 DNA-r p ECs | CD151 | 0.664 | 0.968 | 0.719 | 4.12e-13 | 1.39e-08 |
| Malignant epithelial | C6 DNA-r p ECs | STK25 | 0.663 | 0.508 | 0.276 | 4.05e-07 | 0.0136 |
| Malignant epithelial | C6 DNA-r p ECs | NECAP2 | 0.66 | 0.492 | 0.257 | 9.26e-08 | 0.00312 |
| Malignant epithelial | C6 DNA-r p ECs | TRIM47 | 0.66 | 0.54 | 0.305 | 4.03e-07 | 0.0136 |
| Malignant epithelial | C6 DNA-r p ECs | PPIA | 0.659 | 1.0 | 0.984 | 1.99e-24 | 6.7e-20 |
| Malignant epithelial | C6 DNA-r p ECs | CCNB2 | 0.659 | 0.403 | 0.157 | 6.34e-12 | 2.14e-07 |
| Malignant epithelial | C6 DNA-r p ECs | MRPL12 | 0.659 | 0.855 | 0.57 | 1.88e-09 | 6.33e-05 |
| Malignant epithelial | C6 DNA-r p ECs | EXOSC1 | 0.659 | 0.605 | 0.348 | 2.85e-08 | 0.000961 |
| Malignant epithelial | C6 DNA-r p ECs | CKLF | 0.657 | 0.871 | 0.595 | 3.23e-12 | 1.09e-07 |
| Malignant epithelial | C6 DNA-r p ECs | ELMSAN1 | 0.657 | 0.419 | 0.198 | 8.29e-08 | 0.00279 |
| Malignant epithelial | C6 DNA-r p ECs | FAM129B | 0.654 | 0.685 | 0.425 | 1.17e-07 | 0.00393 |
| Malignant epithelial | C6 DNA-r p ECs | ICT1 | 0.654 | 0.54 | 0.293 | 1.68e-07 | 0.00567 |
| Malignant epithelial | C6 DNA-r p ECs | JAGN1 | 0.653 | 0.573 | 0.315 | 7.65e-08 | 0.00258 |
| Malignant epithelial | C6 DNA-r p ECs | CKS2 | 0.652 | 0.919 | 0.713 | 2.48e-11 | 8.36e-07 |
| Malignant epithelial | C6 DNA-r p ECs | SLC4A2 | 0.652 | 0.411 | 0.202 | 2.76e-07 | 0.0093 |
| Malignant epithelial | C6 DNA-r p ECs | INO80C | 0.651 | 0.532 | 0.284 | 8.15e-08 | 0.00275 |
| Malignant epithelial | C6 DNA-r p ECs | MT2A | 0.651 | 0.935 | 0.711 | 8.78e-08 | 0.00296 |
| Malignant epithelial | C6 DNA-r p ECs | PYCARD | 0.65 | 0.935 | 0.761 | 2.22e-09 | 7.49e-05 |
| Malignant epithelial | C6 DNA-r p ECs | NAMPT | 0.649 | 0.742 | 0.504 | 4.63e-07 | 0.0156 |
| Malignant epithelial | C6 DNA-r p ECs | HSD17B10 | 0.648 | 0.806 | 0.574 | 9e-09 | 0.000303 |

| compartment | subcluster | gene | avg_log2FC | pct.1 | pct.2 | p_val | p_val_adj |
|---|---|---|---|---|---|---|---|
| Malignant epithelial | C6 DNA-r p ECs | NAP1L4 | 0.648 | 0.645 | 0.379 | 4.31e-08 | 0.00145 |
| Malignant epithelial | C6 DNA-r p ECs | NDUFV1 | 0.648 | 0.677 | 0.469 | 1.2e-06 | 0.0405 |
| Malignant epithelial | C6 DNA-r p ECs | TMEM248 | 0.647 | 0.677 | 0.407 | 7.81e-08 | 0.00263 |
| Malignant epithelial | C6 DNA-r p ECs | ATP6V0D1 | 0.647 | 0.597 | 0.362 | 4.39e-07 | 0.0148 |
| Malignant epithelial | C6 DNA-r p ECs | TRAPPC4 | 0.647 | 0.532 | 0.297 | 9.81e-07 | 0.033 |
| Malignant epithelial | C6 DNA-r p ECs | PHPT1 | 0.646 | 0.935 | 0.769 | 8.08e-12 | 2.72e-07 |
| Malignant epithelial | C6 DNA-r p ECs | TIMM10 | 0.645 | 0.645 | 0.403 | 1.44e-07 | 0.00486 |
| Malignant epithelial | C6 DNA-r p ECs | NFE2L3 | 0.645 | 0.427 | 0.225 | 1.33e-06 | 0.0447 |
| Malignant epithelial | C6 DNA-r p ECs | SURF2 | 0.644 | 0.661 | 0.38 | 1.76e-08 | 0.000592 |
| Malignant epithelial | C6 DNA-r p ECs | NSMCE4A | 0.644 | 0.492 | 0.271 | 4.29e-07 | 0.0144 |
| Malignant epithelial | C6 DNA-r p ECs | LARP7 | 0.644 | 0.532 | 0.305 | 8.3e-07 | 0.028 |
| Malignant epithelial | C6 DNA-r p ECs | TMUB1 | 0.643 | 0.702 | 0.431 | 4.64e-09 | 0.000156 |
| Malignant epithelial | C6 DNA-r p ECs | VAMP3 | 0.643 | 0.597 | 0.348 | 1.76e-07 | 0.00594 |
| Malignant epithelial | C6 DNA-r p ECs | ABCF2 | 0.643 | 0.339 | 0.158 | 9.48e-07 | 0.032 |
| Malignant epithelial | C6 DNA-r p ECs | RAB5C | 0.642 | 0.831 | 0.546 | 7.6e-10 | 2.56e-05 |
| Malignant epithelial | C6 DNA-r p ECs | PLPP5 | 0.642 | 0.613 | 0.355 | 3.97e-07 | 0.0134 |
| Malignant epithelial | C6 DNA-r p ECs | NCAPH2 | 0.64 | 0.355 | 0.158 | 1.54e-07 | 0.00518 |
| Malignant epithelial | C6 DNA-r p ECs | PRRG4 | 0.64 | 0.605 | 0.344 | 4.28e-07 | 0.0144 |
| Malignant epithelial | C6 DNA-r p ECs | ARPC5L | 0.639 | 0.815 | 0.544 | 7.9e-10 | 2.66e-05 |
| Malignant epithelial | C6 DNA-r p ECs | FAM213B | 0.639 | 0.476 | 0.222 | 4.79e-09 | 0.000161 |
| Malignant epithelial | C6 DNA-r p ECs | ASS1 | 0.639 | 0.71 | 0.446 | 1.44e-08 | 0.000485 |
| Malignant epithelial | C6 DNA-r p ECs | MLST8 | 0.639 | 0.508 | 0.274 | 1.37e-07 | 0.00463 |
| Malignant epithelial | C6 DNA-r p ECs | ZNF428 | 0.638 | 0.621 | 0.364 | 1.35e-07 | 0.00456 |
| Malignant epithelial | C6 DNA-r p ECs | NABP2 | 0.638 | 0.508 | 0.284 | 9.94e-07 | 0.0335 |
| Malignant epithelial | C6 DNA-r p ECs | KLK1 | 0.637 | 0.573 | 0.275 | 4.89e-11 | 1.65e-06 |
| Malignant epithelial | C6 DNA-r p ECs | MROH6 | 0.637 | 0.605 | 0.316 | 3.28e-09 | 0.00011 |
| Malignant epithelial | C6 DNA-r p ECs | REEP3 | 0.637 | 0.766 | 0.515 | 5.53e-07 | 0.0186 |
| Malignant epithelial | C6 DNA-r p ECs | SHKBP1 | 0.637 | 0.476 | 0.256 | 7.63e-07 | 0.0257 |
| Malignant epithelial | C6 DNA-r p ECs | TMED9 | 0.636 | 0.863 | 0.73 | 8.51e-09 | 0.000287 |
| Malignant epithelial | C6 DNA-r p ECs | NUDC | 0.636 | 0.718 | 0.469 | 8.02e-08 | 0.0027 |
| Malignant epithelial | C6 DNA-r p ECs | PDCD2 | 0.635 | 0.637 | 0.357 | 1.31e-07 | 0.0044 |
| Malignant epithelial | C6 DNA-r p ECs | S100A14 | 0.634 | 0.694 | 0.36 | 9.12e-12 | 3.07e-07 |
| Malignant epithelial | C6 DNA-r p ECs | STARD10 | 0.634 | 0.806 | 0.534 | 3.94e-09 | 0.000133 |
| Malignant epithelial | C6 DNA-r p ECs | MRPL27 | 0.634 | 0.806 | 0.547 | 1.2e-08 | 0.000406 |
| Malignant epithelial | C6 DNA-r p ECs | BCL2L15 | 0.633 | 0.379 | 0.17 | 4.95e-08 | 0.00167 |
| Malignant epithelial | C6 DNA-r p ECs | CHCHD2 | 0.632 | 0.984 | 0.958 | 2.7e-23 | 9.09e-19 |
| Malignant epithelial | C6 DNA-r p ECs | AURKA | 0.632 | 0.266 | 0.117 | 1.28e-06 | 0.0431 |
| Malignant epithelial | C6 DNA-r p ECs | TSPAN13 | 0.631 | 0.637 | 0.351 | 1.27e-09 | 4.28e-05 |
| Malignant epithelial | C6 DNA-r p ECs | DAP | 0.631 | 0.605 | 0.346 | 1.44e-07 | 0.00486 |
| Malignant epithelial | C6 DNA-r p ECs | TSTA3 | 0.629 | 0.79 | 0.483 | 2.63e-10 | 8.87e-06 |
| Malignant epithelial | C6 DNA-r p ECs | COMTD1 | 0.629 | 0.855 | 0.587 | 8.44e-09 | 0.000284 |
| Malignant epithelial | C6 DNA-r p ECs | RABL6 | 0.629 | 0.589 | 0.339 | 1.62e-07 | 0.00544 |
| Malignant epithelial | C6 DNA-r p ECs | RAB10 | 0.629 | 0.79 | 0.529 | 2.35e-07 | 0.00793 |
| Malignant epithelial | C6 DNA-r p ECs | MGAT4B | 0.628 | 0.589 | 0.308 | 3.49e-09 | 0.000118 |
| Malignant epithelial | C6 DNA-r p ECs | HLA-DPB1 | 0.627 | 0.718 | 0.464 | 7.16e-08 | 0.00241 |
| Malignant epithelial | C6 DNA-r p ECs | LGMN | 0.626 | 0.573 | 0.279 | 6.92e-10 | 2.33e-05 |
| Malignant epithelial | C6 DNA-r p ECs | PPP1CA | 0.624 | 0.919 | 0.705 | 6.95e-10 | 2.34e-05 |
| Malignant epithelial | C6 DNA-r p ECs | SAMD9L | 0.623 | 0.315 | 0.131 | 9.19e-08 | 0.00309 |
| Malignant epithelial | C6 DNA-r p ECs | ERH | 0.621 | 1.0 | 0.844 | 2.45e-15 | 8.24e-11 |
| Malignant epithelial | C6 DNA-r p ECs | P4HB | 0.618 | 0.992 | 0.771 | 3.62e-15 | 1.22e-10 |
| Malignant epithelial | C6 DNA-r p ECs | GORASP2 | 0.618 | 0.597 | 0.324 | 3.66e-08 | 0.00123 |
| Malignant epithelial | C6 DNA-r p ECs | SLC22A18 | 0.617 | 0.718 | 0.453 | 1.31e-08 | 0.000443 |
| Malignant epithelial | C6 DNA-r p ECs | TMEM11 | 0.617 | 0.452 | 0.23 | 3.17e-07 | 0.0107 |
| Malignant epithelial | C6 DNA-r p ECs | NOP56 | 0.616 | 0.815 | 0.555 | 1.67e-08 | 0.000562 |
| Malignant epithelial | C6 DNA-r p ECs | TUBB4B | 0.615 | 0.919 | 0.822 | 1.86e-09 | 6.27e-05 |
| Malignant epithelial | C6 DNA-r p ECs | HTATIP2 | 0.615 | 0.823 | 0.55 | 2.47e-09 | 8.32e-05 |
| Malignant epithelial | C6 DNA-r p ECs | SDC1 | 0.615 | 0.782 | 0.534 | 6.01e-07 | 0.0203 |
| Malignant epithelial | C6 DNA-r p ECs | EMD | 0.614 | 0.629 | 0.356 | 5.97e-08 | 0.00201 |
| Malignant epithelial | C6 DNA-r p ECs | PSMD7 | 0.614 | 0.742 | 0.494 | 7.92e-08 | 0.00267 |
| Malignant epithelial | C6 DNA-r p ECs | ASNA1 | 0.613 | 0.677 | 0.456 | 6.02e-07 | 0.0203 |
| Malignant epithelial | C6 DNA-r p ECs | PSMD1 | 0.613 | 0.46 | 0.245 | 6.31e-07 | 0.0212 |
| Malignant epithelial | C6 DNA-r p ECs | MESDC2 | 0.608 | 0.726 | 0.454 | 1.77e-07 | 0.00596 |
| Malignant epithelial | C6 DNA-r p ECs | ANKIB1 | 0.605 | 0.339 | 0.159 | 1.43e-06 | 0.0483 |
| Malignant epithelial | C6 DNA-r p ECs | TRAPPC1 | 0.6 | 0.758 | 0.547 | 1.16e-06 | 0.039 |

| compartment | subcluster | gene | avg_log2FC | pct.1 | pct.2 | p_val | p_val_adj |
|---|---|---|---|---|---|---|---|
| Malignant epithelial | C6 DNA-r p ECs | NDUFS3 | 0.598 | 0.823 | 0.543 | 6.23e-10 | 2.1e-05 |
| Malignant epithelial | C6 DNA-r p ECs | CAST | 0.597 | 0.944 | 0.732 | 7.18e-08 | 0.00242 |
| Malignant epithelial | C6 DNA-r p ECs | MTCH1 | 0.597 | 0.823 | 0.614 | 1.76e-07 | 0.00593 |
| Malignant epithelial | C6 DNA-r p ECs | POLDIP2 | 0.597 | 0.589 | 0.338 | 3.23e-07 | 0.0109 |
| Malignant epithelial | C6 DNA-r p ECs | KRT17 | 0.593 | 0.516 | 0.252 | 8.74e-10 | 2.95e-05 |
| Malignant epithelial | C6 DNA-r p ECs | FAAP20 | 0.593 | 0.742 | 0.482 | 5.04e-09 | 0.00017 |
| Malignant epithelial | C6 DNA-r p ECs | MRPL18 | 0.592 | 0.847 | 0.545 | 6.08e-13 | 2.05e-08 |
| Malignant epithelial | C6 DNA-r p ECs | TAP2 | 0.59 | 0.581 | 0.313 | 1.35e-08 | 0.000454 |
| Malignant epithelial | C6 DNA-r p ECs | ESD | 0.59 | 0.831 | 0.585 | 1.77e-08 | 0.000595 |
| Malignant epithelial | C6 DNA-r p ECs | TMOD3 | 0.59 | 0.548 | 0.319 | 7.08e-07 | 0.0239 |
| Malignant epithelial | C6 DNA-r p ECs | HN1 | 0.589 | 0.952 | 0.811 | 1.13e-09 | 3.82e-05 |
| Malignant epithelial | C6 DNA-r p ECs | NDUFB2 | 0.588 | 0.976 | 0.865 | 5.96e-15 | 2.01e-10 |
| Malignant epithelial | C6 DNA-r p ECs | C7orf50 | 0.587 | 0.847 | 0.597 | 1.97e-08 | 0.000665 |
| Malignant epithelial | C6 DNA-r p ECs | NDUFB8 | 0.584 | 0.976 | 0.81 | 8.09e-14 | 2.73e-09 |
| Malignant epithelial | C6 DNA-r p ECs | LMAN2 | 0.584 | 0.911 | 0.7 | 5.05e-09 | 0.00017 |
| Malignant epithelial | C6 DNA-r p ECs | ATG3 | 0.584 | 0.589 | 0.361 | 1.28e-06 | 0.0431 |
| Malignant epithelial | C6 DNA-r p ECs | PYGB | 0.582 | 0.669 | 0.396 | 8.53e-09 | 0.000287 |
| Malignant epithelial | C6 DNA-r p ECs | MRPL13 | 0.582 | 0.823 | 0.585 | 2.88e-08 | 0.000971 |
| Malignant epithelial | C6 DNA-r p ECs | CD47 | 0.579 | 0.839 | 0.522 | 1.93e-10 | 6.51e-06 |
| Malignant epithelial | C6 DNA-r p ECs | RPL26L1 | 0.578 | 0.855 | 0.557 | 1.4e-08 | 0.000471 |
| Malignant epithelial | C6 DNA-r p ECs | RANBP1 | 0.577 | 0.895 | 0.743 | 4.2e-08 | 0.00141 |
| Malignant epithelial | C6 DNA-r p ECs | PSMB1 | 0.576 | 0.968 | 0.757 | 2.95e-10 | 9.93e-06 |
| Malignant epithelial | C6 DNA-r p ECs | PDAP1 | 0.576 | 0.879 | 0.649 | 1.63e-08 | 0.00055 |
| Malignant epithelial | C6 DNA-r p ECs | SYPL1 | 0.575 | 0.863 | 0.641 | 8.92e-08 | 0.00301 |
| Malignant epithelial | C6 DNA-r p ECs | VDAC3 | 0.572 | 0.782 | 0.547 | 6.01e-07 | 0.0202 |
| Malignant epithelial | C6 DNA-r p ECs | EZR | 0.571 | 0.96 | 0.727 | 2.04e-14 | 6.88e-10 |
| Malignant epithelial | C6 DNA-r p ECs | PSMD13 | 0.571 | 0.71 | 0.422 | 2.29e-08 | 0.000772 |
| Malignant epithelial | C6 DNA-r p ECs | UCHL3 | 0.569 | 0.669 | 0.398 | 4.52e-08 | 0.00152 |
| Malignant epithelial | C6 DNA-r p ECs | CAPNS1 | 0.569 | 0.831 | 0.619 | 3.84e-07 | 0.0129 |
| Malignant epithelial | C6 DNA-r p ECs | CALM2 | 0.568 | 0.984 | 0.936 | 3.53e-13 | 1.19e-08 |
| Malignant epithelial | C6 DNA-r p ECs | ARF5 | 0.568 | 0.863 | 0.634 | 1.53e-07 | 0.00516 |
| Malignant epithelial | C6 DNA-r p ECs | GOT2 | 0.568 | 0.484 | 0.252 | 4.83e-07 | 0.0163 |
| Malignant epithelial | C6 DNA-r p ECs | PIGT | 0.567 | 0.726 | 0.437 | 3.8e-09 | 0.000128 |
| Malignant epithelial | C6 DNA-r p ECs | ALG5 | 0.567 | 0.597 | 0.349 | 1.93e-07 | 0.00651 |
| Malignant epithelial | C6 DNA-r p ECs | PSMD11 | 0.567 | 0.629 | 0.387 | 9.97e-07 | 0.0336 |
| Malignant epithelial | C6 DNA-r p ECs | BRMS1 | 0.567 | 0.556 | 0.341 | 1.48e-06 | 0.0499 |
| Malignant epithelial | C6 DNA-r p ECs | ASL | 0.566 | 0.661 | 0.371 | 1.37e-08 | 0.000463 |
| Malignant epithelial | C6 DNA-r p ECs | CALM3 | 0.565 | 0.887 | 0.7 | 1.64e-08 | 0.000551 |
| Malignant epithelial | C6 DNA-r p ECs | SSBP1 | 0.564 | 0.952 | 0.829 | 2.14e-11 | 7.22e-07 |
| Malignant epithelial | C6 DNA-r p ECs | CCT6A | 0.564 | 0.831 | 0.605 | 3.55e-08 | 0.00119 |
| Malignant epithelial | C6 DNA-r p ECs | MX1 | 0.564 | 0.637 | 0.351 | 8.01e-08 | 0.0027 |
| Malignant epithelial | C6 DNA-r p ECs | NXT1 | 0.564 | 0.677 | 0.439 | 6.08e-07 | 0.0205 |
| Malignant epithelial | C6 DNA-r p ECs | PSMA2 | 0.563 | 0.96 | 0.771 | 1.82e-11 | 6.13e-07 |
| Malignant epithelial | C6 DNA-r p ECs | ANXA2 | 0.562 | 1.0 | 0.977 | 6.76e-11 | 2.28e-06 |
| Malignant epithelial | C6 DNA-r p ECs | HLA-F | 0.562 | 0.685 | 0.424 | 9.07e-07 | 0.0306 |
| Malignant epithelial | C6 DNA-r p ECs | SH3BGRL3 | 0.561 | 0.976 | 0.884 | 8.51e-12 | 2.87e-07 |
| Malignant epithelial | C6 DNA-r p ECs | SNF8 | 0.561 | 0.782 | 0.531 | 2.98e-07 | 0.01 |
| Malignant epithelial | C6 DNA-r p ECs | CKS1B | 0.56 | 0.831 | 0.625 | 1.75e-08 | 0.000589 |
| Malignant epithelial | C6 DNA-r p ECs | TROAP | 0.559 | 0.363 | 0.162 | 1.17e-07 | 0.00395 |
| Malignant epithelial | C6 DNA-r p ECs | MMP7 | 0.559 | 0.677 | 0.413 | 4.57e-07 | 0.0154 |
| Malignant epithelial | C6 DNA-r p ECs | FAM96B | 0.558 | 0.927 | 0.748 | 8.73e-10 | 2.94e-05 |
| Malignant epithelial | C6 DNA-r p ECs | GTF3A | 0.557 | 0.887 | 0.618 | 4.08e-11 | 1.37e-06 |
| Malignant epithelial | C6 DNA-r p ECs | TSPAN3 | 0.557 | 0.71 | 0.451 | 3.56e-08 | 0.0012 |
| Malignant epithelial | C6 DNA-r p ECs | TXNDC17 | 0.556 | 0.895 | 0.648 | 1.44e-09 | 4.84e-05 |
| Malignant epithelial | C6 DNA-r p ECs | ACP1 | 0.555 | 0.919 | 0.715 | 5.91e-08 | 0.00199 |
| Malignant epithelial | C6 DNA-r p ECs | WBSCR22 | 0.555 | 0.766 | 0.535 | 1.42e-06 | 0.0478 |
| Malignant epithelial | C6 DNA-r p ECs | PSMB8 | 0.554 | 0.944 | 0.731 | 2.73e-11 | 9.18e-07 |
| Malignant epithelial | C6 DNA-r p ECs | PSMG3 | 0.552 | 0.766 | 0.514 | 1.36e-07 | 0.00457 |
| Malignant epithelial | C6 DNA-r p ECs | PSMF1 | 0.552 | 0.782 | 0.531 | 2.62e-07 | 0.00884 |
| Malignant epithelial | C6 DNA-r p ECs | LGALS3 | 0.55 | 1.0 | 0.969 | 6.36e-17 | 2.14e-12 |
| Malignant epithelial | C6 DNA-r p ECs | VDAC1 | 0.547 | 0.935 | 0.823 | 1.14e-09 | 3.84e-05 |
| Malignant epithelial | C6 DNA-r p ECs | INPP4B | 0.546 | 0.395 | 0.198 | 1.11e-06 | 0.0373 |
| Malignant epithelial | C6 DNA-r p ECs | ITPA | 0.546 | 0.75 | 0.513 | 1.24e-06 | 0.0417 |
| Malignant epithelial | C6 DNA-r p ECs | ADAP1 | 0.545 | 0.734 | 0.419 | 2.57e-08 | 0.000866 |

| compartment | subcluster | gene | avg_log2FC | pct.1 | pct.2 | p_val | p_val_adj |
|---|---|---|---|---|---|---|---|
| Malignant epithelial | C6 DNA-r p ECs | CHMP4A | 0.545 | 0.742 | 0.484 | 2.92e-07 | 0.00984 |
| Malignant epithelial | C6 DNA-r p ECs | NDUFB9 | 0.542 | 0.984 | 0.876 | 1.55e-10 | 5.22e-06 |
| Malignant epithelial | C6 DNA-r p ECs | SSNA1 | 0.542 | 0.847 | 0.644 | 3.76e-07 | 0.0127 |
| Malignant epithelial | C6 DNA-r p ECs | FASTK | 0.541 | 0.605 | 0.344 | 4.09e-07 | 0.0138 |
| Malignant epithelial | C6 DNA-r p ECs | PRDX4 | 0.54 | 0.831 | 0.612 | 5.41e-07 | 0.0182 |
| Malignant epithelial | C6 DNA-r p ECs | UCP2 | 0.54 | 0.387 | 0.189 | 8.65e-07 | 0.0291 |
| Malignant epithelial | C6 DNA-r p ECs | NDUFB11 | 0.535 | 0.984 | 0.903 | 1.13e-10 | 3.8e-06 |
| Malignant epithelial | C6 DNA-r p ECs | IFI27L2 | 0.535 | 0.935 | 0.669 | 8.02e-10 | 2.7e-05 |
| Malignant epithelial | C6 DNA-r p ECs | RAB11FIP1 | 0.535 | 0.815 | 0.568 | 2.99e-08 | 0.00101 |
| Malignant epithelial | C6 DNA-r p ECs | PRELID1 | 0.534 | 0.968 | 0.795 | 7.66e-10 | 2.58e-05 |
| Malignant epithelial | C6 DNA-r p ECs | LSM8 | 0.532 | 0.863 | 0.683 | 4.96e-08 | 0.00167 |
| Malignant epithelial | C6 DNA-r p ECs | HDDC3 | 0.532 | 0.532 | 0.292 | 4.36e-07 | 0.0147 |
| Malignant epithelial | C6 DNA-r p ECs | PSME1 | 0.53 | 1.0 | 0.891 | 1.24e-14 | 4.19e-10 |
| Malignant epithelial | C6 DNA-r p ECs | RRP7A | 0.528 | 0.758 | 0.503 | 6.94e-07 | 0.0234 |
| Malignant epithelial | C6 DNA-r p ECs | CCDC124 | 0.527 | 0.742 | 0.482 | 3.24e-07 | 0.0109 |
| Malignant epithelial | C6 DNA-r p ECs | C4orf48 | 0.526 | 0.944 | 0.691 | 3.85e-09 | 0.00013 |
| Malignant epithelial | C6 DNA-r p ECs | TMEM141 | 0.525 | 0.968 | 0.797 | 1.52e-08 | 0.000513 |
| Malignant epithelial | C6 DNA-r p ECs | ARHGDIA | 0.524 | 0.903 | 0.641 | 7.9e-09 | 0.000266 |
| Malignant epithelial | C6 DNA-r p ECs | CAPZA2 | 0.524 | 0.831 | 0.578 | 1.86e-08 | 0.000626 |
| Malignant epithelial | C6 DNA-r p ECs | TBCB | 0.522 | 0.798 | 0.571 | 7.2e-07 | 0.0243 |
| Malignant epithelial | C6 DNA-r p ECs | PSMA5 | 0.521 | 0.79 | 0.499 | 1.12e-07 | 0.00378 |
| Malignant epithelial | C6 DNA-r p ECs | BAIAP2L1 | 0.521 | 0.685 | 0.421 | 9.08e-07 | 0.0306 |
| Malignant epithelial | C6 DNA-r p ECs | AP2S1 | 0.52 | 0.935 | 0.733 | 3.35e-09 | 0.000113 |
| Malignant epithelial | C6 DNA-r p ECs | TMEM54 | 0.52 | 0.839 | 0.649 | 1.24e-07 | 0.00416 |
| Malignant epithelial | C6 DNA-r p ECs | PUF60 | 0.518 | 0.815 | 0.563 | 7.67e-07 | 0.0258 |
| Malignant epithelial | C6 DNA-r p ECs | RAB11A | 0.516 | 0.919 | 0.729 | 1.56e-08 | 0.000524 |
| Malignant epithelial | C6 DNA-r p ECs | DCTN2 | 0.516 | 0.694 | 0.418 | 2.54e-07 | 0.00857 |
| Malignant epithelial | C6 DNA-r p ECs | ATP5J2 | 0.514 | 0.992 | 0.933 | 1.71e-13 | 5.76e-09 |
| Malignant epithelial | C6 DNA-r p ECs | PSMB3 | 0.514 | 0.984 | 0.822 | 2.76e-10 | 9.3e-06 |
| Malignant epithelial | C6 DNA-r p ECs | EIF3M | 0.512 | 0.895 | 0.682 | 1.34e-07 | 0.00453 |
| Malignant epithelial | C6 DNA-r p ECs | CDC20 | 0.511 | 0.347 | 0.146 | 1.29e-08 | 0.000435 |
| Malignant epithelial | C6 DNA-r p ECs | DEK | 0.511 | 0.782 | 0.591 | 6.06e-07 | 0.0204 |
| Malignant epithelial | C6 DNA-r p ECs | CYSTM1 | 0.51 | 0.911 | 0.752 | 9.94e-09 | 0.000335 |
| Malignant epithelial | C6 DNA-r p ECs | TMED10 | 0.51 | 0.887 | 0.704 | 3.49e-07 | 0.0117 |
| Malignant epithelial | C6 DNA-r p ECs | NDUFB7 | 0.509 | 0.968 | 0.886 | 7.81e-08 | 0.00263 |
| Malignant epithelial | C6 DNA-r p ECs | EHF | 0.509 | 0.718 | 0.437 | 1.78e-07 | 0.00601 |
| Malignant epithelial | C6 DNA-r p ECs | RNH1 | 0.507 | 0.887 | 0.653 | 1.18e-06 | 0.0399 |
| Malignant epithelial | C6 DNA-r p ECs | NDUFA8 | 0.505 | 0.831 | 0.56 | 7.79e-07 | 0.0262 |
| Malignant epithelial | C6 DNA-r p ECs | NDUFS6 | 0.503 | 0.992 | 0.849 | 9.25e-10 | 3.12e-05 |
| Malignant epithelial | C6 DNA-r p ECs | PAIP2 | 0.503 | 0.758 | 0.495 | 2.52e-07 | 0.00848 |
| Malignant epithelial | C6 DNA-r p ECs | MRPL41 | 0.501 | 0.887 | 0.705 | 1.26e-07 | 0.00425 |
| Malignant epithelial | C6 DNA-r p ECs | TSTD1 | 0.497 | 0.895 | 0.658 | 8.73e-08 | 0.00294 |
| Malignant epithelial | C6 DNA-r p ECs | C2 | 0.496 | 0.387 | 0.186 | 1.12e-06 | 0.0379 |
| Malignant epithelial | C6 DNA-r p ECs | RAN | 0.495 | 0.992 | 0.894 | 4.52e-11 | 1.52e-06 |
| Malignant epithelial | C6 DNA-r p ECs | SUMF2 | 0.494 | 0.694 | 0.456 | 6.2e-07 | 0.0209 |
| Malignant epithelial | C6 DNA-r p ECs | CHCHD7 | 0.493 | 0.863 | 0.602 | 4.17e-08 | 0.0014 |
| Malignant epithelial | C6 DNA-r p ECs | RP11-462G2.1 | 0.492 | 0.637 | 0.324 | 4.33e-11 | 1.46e-06 |
| Malignant epithelial | C6 DNA-r p ECs | FAM49B | 0.492 | 0.677 | 0.435 | 5.96e-07 | 0.0201 |
| Malignant epithelial | C6 DNA-r p ECs | TPM4 | 0.49 | 0.935 | 0.745 | 7.7e-09 | 0.000259 |
| Malignant epithelial | C6 DNA-r p ECs | CDK5RAP3 | 0.488 | 0.581 | 0.32 | 2.97e-07 | 0.00999 |
| Malignant epithelial | C6 DNA-r p ECs | MDH2 | 0.487 | 0.903 | 0.72 | 2.76e-08 | 0.00093 |
| Malignant epithelial | C6 DNA-r p ECs | SUCLG1 | 0.487 | 0.766 | 0.461 | 6.62e-08 | 0.00223 |
| Malignant epithelial | C6 DNA-r p ECs | POMP | 0.486 | 0.984 | 0.847 | 3.42e-12 | 1.15e-07 |
| Malignant epithelial | C6 DNA-r p ECs | FIS1 | 0.486 | 0.968 | 0.798 | 5.87e-08 | 0.00198 |
| Malignant epithelial | C6 DNA-r p ECs | EIF5A | 0.485 | 0.927 | 0.731 | 2.99e-09 | 0.000101 |
| Malignant epithelial | C6 DNA-r p ECs | ITGA2 | 0.485 | 0.395 | 0.189 | 8.68e-08 | 0.00293 |
| Malignant epithelial | C6 DNA-r p ECs | CD44 | 0.485 | 0.823 | 0.565 | 2.42e-07 | 0.00815 |
| Malignant epithelial | C6 DNA-r p ECs | CENPB | 0.485 | 0.452 | 0.228 | 8.26e-07 | 0.0278 |
| Malignant epithelial | C6 DNA-r p ECs | RTN3 | 0.481 | 0.847 | 0.608 | 6.05e-07 | 0.0204 |
| Malignant epithelial | C6 DNA-r p ECs | TRAPPC2L | 0.48 | 0.806 | 0.563 | 5.16e-07 | 0.0174 |
| Malignant epithelial | C6 DNA-r p ECs | YWHAB | 0.477 | 0.968 | 0.823 | 5.74e-09 | 0.000193 |
| Malignant epithelial | C6 DNA-r p ECs | TFF3 | 0.474 | 0.871 | 0.62 | 1.87e-08 | 0.000631 |
| Malignant epithelial | C6 DNA-r p ECs | SUGT1 | 0.473 | 0.71 | 0.415 | 2.35e-07 | 0.00793 |
| Malignant epithelial | C6 DNA-r p ECs | ATP5C1 | 0.473 | 0.895 | 0.69 | 1.09e-06 | 0.0366 |

| compartment | subcluster | gene | avg_log2FC | pct.1 | pct.2 | p_val | p_val_adj |
|---|---|---|---|---|---|---|---|
| Malignant epithelial | C6 DNA-r p ECs | CSNK2B | 0.471 | 0.952 | 0.778 | 7.21e-09 | 0.000243 |
| Malignant epithelial | C6 DNA-r p ECs | MGST1 | 0.471 | 0.895 | 0.714 | 2.34e-07 | 0.00788 |
| Malignant epithelial | C6 DNA-r p ECs | NELFE | 0.47 | 0.766 | 0.485 | 5.38e-07 | 0.0181 |
| Malignant epithelial | C6 DNA-r p ECs | ANAPC11 | 0.468 | 0.96 | 0.863 | 3.73e-09 | 0.000126 |
| Malignant epithelial | C6 DNA-r p ECs | PSMD8 | 0.466 | 0.871 | 0.676 | 1.02e-06 | 0.0343 |
| Malignant epithelial | C6 DNA-r p ECs | MCTS1 | 0.465 | 0.806 | 0.561 | 8.36e-07 | 0.0282 |
| Malignant epithelial | C6 DNA-r p ECs | PSMB6 | 0.464 | 0.847 | 0.591 | 1.04e-06 | 0.0351 |
| Malignant epithelial | C6 DNA-r p ECs | RAB14 | 0.461 | 0.605 | 0.338 | 2.95e-07 | 0.00994 |
| Malignant epithelial | C6 DNA-r p ECs | PRSS3 | 0.46 | 0.952 | 0.692 | 1.38e-11 | 4.66e-07 |
| Malignant epithelial | C6 DNA-r p ECs | ZNHIT1 | 0.46 | 0.976 | 0.793 | 1.59e-08 | 0.000534 |
| Malignant epithelial | C6 DNA-r p ECs | DRAP1 | 0.459 | 0.984 | 0.817 | 1.46e-08 | 0.000492 |
| Malignant epithelial | C6 DNA-r p ECs | HNRNPA3 | 0.457 | 0.952 | 0.809 | 1.12e-08 | 0.000379 |
| Malignant epithelial | C6 DNA-r p ECs | CFL1 | 0.453 | 1.0 | 0.974 | 1.36e-18 | 4.59e-14 |
| Malignant epithelial | C6 DNA-r p ECs | SNRPG | 0.452 | 0.952 | 0.867 | 1.29e-07 | 0.00433 |
| Malignant epithelial | C6 DNA-r p ECs | ATP6V0E1 | 0.451 | 0.96 | 0.872 | 1.27e-08 | 0.000427 |
| Malignant epithelial | C6 DNA-r p ECs | PLP2 | 0.448 | 1.0 | 0.915 | 3.9e-09 | 0.000131 |
| Malignant epithelial | C6 DNA-r p ECs | PSMA7 | 0.447 | 1.0 | 0.9 | 7.04e-10 | 2.37e-05 |
| Malignant epithelial | C6 DNA-r p ECs | TAP1 | 0.447 | 0.766 | 0.528 | 8.95e-07 | 0.0301 |
| Malignant epithelial | C6 DNA-r p ECs | NDUFB6 | 0.446 | 0.863 | 0.687 | 3.94e-07 | 0.0133 |
| Malignant epithelial | C6 DNA-r p ECs | RWDD1 | 0.444 | 0.815 | 0.573 | 1.47e-06 | 0.0496 |
| Malignant epithelial | C6 DNA-r p ECs | UQCC3 | 0.442 | 0.879 | 0.619 | 6.5e-07 | 0.0219 |
| Malignant epithelial | C6 DNA-r p ECs | UQCRQ | 0.439 | 1.0 | 0.966 | 6.83e-13 | 2.3e-08 |
| Malignant epithelial | C6 DNA-r p ECs | SPTBN1 | 0.437 | 0.742 | 0.444 | 9.89e-08 | 0.00333 |
| Malignant epithelial | C6 DNA-r p ECs | MRPS34 | 0.437 | 0.903 | 0.687 | 5.3e-07 | 0.0179 |
| Malignant epithelial | C6 DNA-r p ECs | KRT20 | 0.433 | 0.556 | 0.31 | 1.49e-07 | 0.00501 |
| Malignant epithelial | C6 DNA-r p ECs | CLIC1 | 0.431 | 1.0 | 0.966 | 1.28e-11 | 4.33e-07 |
| Malignant epithelial | C6 DNA-r p ECs | H3F3B | 0.431 | 1.0 | 0.982 | 4.53e-07 | 0.0153 |
| Malignant epithelial | C6 DNA-r p ECs | SRI | 0.431 | 0.944 | 0.796 | 7.14e-07 | 0.0241 |
| Malignant epithelial | C6 DNA-r p ECs | SLC40A1 | 0.421 | 0.387 | 0.188 | 3.19e-07 | 0.0107 |
| Malignant epithelial | C6 DNA-r p ECs | C19orf43 | 0.418 | 0.968 | 0.844 | 6.38e-08 | 0.00215 |
| Malignant epithelial | C6 DNA-r p ECs | EDF1 | 0.408 | 1.0 | 0.955 | 6.44e-11 | 2.17e-06 |
| Malignant epithelial | C6 DNA-r p ECs | PSMA4 | 0.407 | 0.96 | 0.765 | 3.28e-07 | 0.0111 |
| Malignant epithelial | C6 DNA-r p ECs | PCBD1 | 0.406 | 0.976 | 0.847 | 1.06e-07 | 0.00358 |
| Malignant epithelial | C6 DNA-r p ECs | KLK10 | 0.405 | 0.702 | 0.449 | 1.23e-07 | 0.00414 |
| Malignant epithelial | C6 DNA-r p ECs | UFM1 | 0.405 | 0.75 | 0.506 | 1.42e-06 | 0.048 |
| Malignant epithelial | C6 DNA-r p ECs | LSR | 0.401 | 0.96 | 0.766 | 2.97e-07 | 0.00999 |
| Malignant epithelial | C6 DNA-r p ECs | PTGES3 | 0.4 | 0.952 | 0.837 | 3.04e-08 | 0.00103 |
| Malignant epithelial | C6 DNA-r p ECs | MT-ND4 | 0.398 | 1.0 | 0.994 | 7.15e-09 | 0.000241 |
| Malignant epithelial | C6 DNA-r p ECs | FXYD3 | 0.395 | 0.992 | 0.867 | 2.17e-12 | 7.32e-08 |
| Malignant epithelial | C6 DNA-r p ECs | PRSS8 | 0.395 | 0.782 | 0.45 | 6.77e-09 | 0.000228 |
| Malignant epithelial | C6 DNA-r p ECs | SRRM1 | 0.394 | 0.863 | 0.577 | 1.45e-06 | 0.0489 |
| Malignant epithelial | C6 DNA-r p ECs | ENY2 | 0.384 | 0.968 | 0.847 | 8.33e-07 | 0.0281 |
| Malignant epithelial | C6 DNA-r p ECs | EEF1D | 0.383 | 1.0 | 0.973 | 1.43e-07 | 0.00481 |
| Malignant epithelial | C6 DNA-r p ECs | MT-CO2 | 0.381 | 1.0 | 0.997 | 3.65e-08 | 0.00123 |
| Malignant epithelial | C6 DNA-r p ECs | IGHA2 | 0.377 | 0.476 | 0.215 | 1.3e-09 | 4.37e-05 |
| Malignant epithelial | C6 DNA-r p ECs | NPM1 | 0.375 | 0.984 | 0.961 | 1.64e-07 | 0.00554 |
| Malignant epithelial | C6 DNA-r p ECs | SRSF9 | 0.375 | 0.944 | 0.793 | 1.1e-06 | 0.037 |
| Malignant epithelial | C6 DNA-r p ECs | KRT8 | 0.372 | 1.0 | 0.997 | 3.76e-08 | 0.00127 |
| Malignant epithelial | C6 DNA-r p ECs | SHFM1 | 0.369 | 0.992 | 0.947 | 2.19e-09 | 7.39e-05 |
| Malignant epithelial | C6 DNA-r p ECs | CD24 | 0.368 | 0.887 | 0.477 | 4.87e-13 | 1.64e-08 |
| Malignant epithelial | C6 DNA-r p ECs | HLA-C | 0.362 | 0.984 | 0.946 | 4.16e-07 | 0.014 |
| Malignant epithelial | C6 DNA-r p ECs | COX8A | 0.357 | 0.992 | 0.937 | 8.03e-09 | 0.000271 |
| Malignant epithelial | C6 DNA-r p ECs | ALDOA | 0.346 | 1.0 | 0.965 | 2.21e-07 | 0.00744 |
| Malignant epithelial | C6 DNA-r p ECs | ATP5H | 0.346 | 0.96 | 0.86 | 1.28e-06 | 0.043 |
| Malignant epithelial | C6 DNA-r p ECs | MIA | 0.338 | 0.863 | 0.571 | 1.57e-07 | 0.0053 |
| Malignant epithelial | C6 DNA-r p ECs | GSTP1 | 0.333 | 0.992 | 0.96 | 1.15e-10 | 3.86e-06 |
| Malignant epithelial | C6 DNA-r p ECs | ARPC3 | 0.331 | 0.992 | 0.929 | 8.1e-07 | 0.0273 |
| Malignant epithelial | C6 DNA-r p ECs | SUMO2 | 0.326 | 0.984 | 0.937 | 1.43e-06 | 0.0482 |
| Malignant epithelial | C6 DNA-r p ECs | PGP | 0.318 | 0.581 | 0.315 | 1.24e-06 | 0.0418 |
| Malignant epithelial | C6 DNA-r p ECs | KRT18 | 0.304 | 0.992 | 0.992 | 1.84e-07 | 0.00619 |
| Malignant epithelial | C6 DNA-r p ECs | SLC44A4 | 0.302 | 0.524 | 0.275 | 3.94e-07 | 0.0133 |
| Malignant epithelial | C6 DNA-r p ECs | RPL22 | 0.293 | 1.0 | 0.983 | 8.88e-09 | 0.000299 |
| Malignant epithelial | C6 DNA-r p ECs | B2M | 0.293 | 1.0 | 1.0 | 1.53e-08 | 0.000517 |
| Malignant epithelial | C6 DNA-r p ECs | PFN1 | 0.284 | 1.0 | 0.964 | 4.32e-07 | 0.0145 |

| compartment | subcluster | gene | avg_log2FC | pct.1 | pct.2 | p_val | p_val_adj |
|---|---|---|---|---|---|---|---|
| Malignant epithelial | C6 DNA-r p ECs | HINT1 | 0.277 | 1.0 | 0.969 | 1.59e-07 | 0.00537 |
| Malignant epithelial | C6 DNA-r p ECs | SAP18 | 0.274 | 0.984 | 0.876 | 9.75e-07 | 0.0328 |
| Fibroblast | F0 SFRP2+ Fib | OMG | 4.855 | 0.133 | 0.006 | 1.69e-18 | 5.69e-14 |
| Fibroblast | F0 SFRP2+ Fib | SNCA | 4.772 | 0.13 | 0.008 | 2.12e-17 | 7.13e-13 |
| Fibroblast | F0 SFRP2+ Fib | KCNN3 | 4.46 | 0.13 | 0.011 | 2.79e-16 | 9.4e-12 |
| Fibroblast | F0 SFRP2+ Fib | THBS4 | 4.341 | 0.21 | 0.025 | 2.08e-23 | 7.01e-19 |
| Fibroblast | F0 SFRP2+ Fib | CD34 | 4.196 | 0.252 | 0.009 | 4.43e-36 | 1.49e-31 |
| Fibroblast | F0 SFRP2+ Fib | ABCA9 | 3.543 | 0.151 | 0.012 | 7.23e-19 | 2.44e-14 |
| Fibroblast | F0 SFRP2+ Fib | SFRP1 | 3.497 | 0.562 | 0.077 | 4.42e-67 | 1.49e-62 |
| Fibroblast | F0 SFRP2+ Fib | PLA2G2A | 3.467 | 0.56 | 0.055 | 1.94e-75 | 6.55e-71 |
| Fibroblast | F0 SFRP2+ Fib | OGN | 3.391 | 0.645 | 0.072 | 1.17e-84 | 3.94e-80 |
| Fibroblast | F0 SFRP2+ Fib | MGST1 | 3.385 | 0.615 | 0.067 | 5.77e-82 | 1.94e-77 |
| Fibroblast | F0 SFRP2+ Fib | PI16 | 3.32 | 0.194 | 0.021 | 1.26e-21 | 4.24e-17 |
| Fibroblast | F0 SFRP2+ Fib | CILP | 3.289 | 0.377 | 0.044 | 2.16e-43 | 7.28e-39 |
| Fibroblast | F0 SFRP2+ Fib | OSR1 | 3.268 | 0.101 | 0.009 | 2.64e-12 | 8.9e-08 |
| Fibroblast | F0 SFRP2+ Fib | DPT | 3.261 | 0.554 | 0.093 | 1.49e-61 | 5.02e-57 |
| Fibroblast | F0 SFRP2+ Fib | RSPO3 | 3.227 | 0.233 | 0.046 | 1.27e-20 | 4.29e-16 |
| Fibroblast | F0 SFRP2+ Fib | SFRP2 | 3.209 | 0.862 | 0.222 | 9.73e-101 | 3.28e-96 |
| Fibroblast | F0 SFRP2+ Fib | XPNPEP2 | 3.121 | 0.133 | 0.009 | 4.94e-17 | 1.66e-12 |
| Fibroblast | F0 SFRP2+ Fib | LEPR | 3.065 | 0.21 | 0.029 | 1.43e-21 | 4.81e-17 |
| Fibroblast | F0 SFRP2+ Fib | C7 | 3.056 | 0.785 | 0.133 | 2.87e-96 | 9.679999999999999e-92 |
| Fibroblast | F0 SFRP2+ Fib | ABLIM1 | 3.036 | 0.172 | 0.018 | 1.21e-19 | 4.07e-15 |
| Fibroblast | F0 SFRP2+ Fib | WISP2 | 3.018 | 0.263 | 0.043 | 6.17e-25 | 2.08e-20 |
| Fibroblast | F0 SFRP2+ Fib | HTR2B | 2.993 | 0.117 | 0.017 | 5.91e-12 | 1.99e-07 |
| Fibroblast | F0 SFRP2+ Fib | LMO3 | 2.974 | 0.146 | 0.009 | 5.29e-19 | 1.78e-14 |
| Fibroblast | F0 SFRP2+ Fib | AOX1 | 2.932 | 0.199 | 0.015 | 1.7e-24 | 5.74e-20 |
| Fibroblast | F0 SFRP2+ Fib | ABCA10 | 2.927 | 0.143 | 0.018 | 3.49e-15 | 1.18e-10 |
| Fibroblast | F0 SFRP2+ Fib | TNXB | 2.91 | 0.424 | 0.052 | 1.68e-48 | 5.65e-44 |
| Fibroblast | F0 SFRP2+ Fib | C16orf89 | 2.89 | 0.215 | 0.021 | 8.66e-25 | 2.92e-20 |
| Fibroblast | F0 SFRP2+ Fib | CHRDL2 | 2.888 | 0.175 | 0.02 | 3.03e-19 | 1.02e-14 |
| Fibroblast | F0 SFRP2+ Fib | ITM2A | 2.813 | 0.509 | 0.066 | 1.67e-58 | 5.63e-54 |
| Fibroblast | F0 SFRP2+ Fib | SHISA3 | 2.813 | 0.133 | 0.017 | 4.26e-14 | 1.44e-09 |
| Fibroblast | F0 SFRP2+ Fib | ABI3BP | 2.794 | 0.414 | 0.06 | 3.81e-44 | 1.28e-39 |
| Fibroblast | F0 SFRP2+ Fib | CRISPLD1 | 2.759 | 0.175 | 0.038 | 2.97e-14 | 1e-09 |
| Fibroblast | F0 SFRP2+ Fib | ADH1C | 2.759 | 0.125 | 0.017 | 5.75e-13 | 1.94e-08 |
| Fibroblast | F0 SFRP2+ Fib | CTSH | 2.756 | 0.56 | 0.123 | 2.3e-54 | 7.74e-50 |
| Fibroblast | F0 SFRP2+ Fib | SRPX | 2.706 | 0.682 | 0.156 | 8.72e-70 | 2.94e-65 |
| Fibroblast | F0 SFRP2+ Fib | CHRDL1 | 2.703 | 0.279 | 0.026 | 3.72e-33 | 1.26e-28 |
| Fibroblast | F0 SFRP2+ Fib | INMT | 2.703 | 0.294 | 0.06 | 6.37e-25 | 2.15e-20 |
| Fibroblast | F0 SFRP2+ Fib | DCLK1 | 2.685 | 0.215 | 0.04 | 5.65e-19 | 1.9e-14 |
| Fibroblast | F0 SFRP2+ Fib | KAZALD1 | 2.661 | 0.114 | 0.017 | 1.73e-11 | 5.82e-07 |
| Fibroblast | F0 SFRP2+ Fib | CDO1 | 2.653 | 0.215 | 0.032 | 5.47e-21 | 1.84e-16 |
| Fibroblast | F0 SFRP2+ Fib | FAM198A | 2.644 | 0.159 | 0.018 | 1.98e-17 | 6.67e-13 |
| Fibroblast | F0 SFRP2+ Fib | SCRG1 | 2.643 | 0.263 | 0.043 | 7.92e-25 | 2.67e-20 |
| Fibroblast | F0 SFRP2+ Fib | CPVL | 2.642 | 0.215 | 0.028 | 7.789999999999999e-23 | 2.63e-18 |
| Fibroblast | F0 SFRP2+ Fib | OMD | 2.638 | 0.363 | 0.057 | 3.04e-36 | 1.02e-31 |
| Fibroblast | F0 SFRP2+ Fib | BOC | 2.62 | 0.459 | 0.077 | 7.870000000000001e-47 | 2.65e-42 |
| Fibroblast | F0 SFRP2+ Fib | C3 | 2.615 | 0.894 | 0.231 | 2.26e-101 | 7.62e-97 |
| Fibroblast | F0 SFRP2+ Fib | ST3GAL5 | 2.557 | 0.117 | 0.017 | 8.43e-12 | 2.84e-07 |
| Fibroblast | F0 SFRP2+ Fib | IGF1 | 2.554 | 0.708 | 0.136 | 6.04e-78 | 2.03e-73 |
| Fibroblast | F0 SFRP2+ Fib | CAPN6 | 2.504 | 0.103 | 0.017 | 5.96e-10 | 2.01e-05 |
| Fibroblast | F0 SFRP2+ Fib | PID1 | 2.489 | 0.361 | 0.058 | 1.94e-35 | 6.55e-31 |
| Fibroblast | F0 SFRP2+ Fib | EFEMP1 | 2.479 | 0.764 | 0.21 | 4.22e-74 | 1.42e-69 |
| Fibroblast | F0 SFRP2+ Fib | SCN7A | 2.473 | 0.106 | 0.015 | 7.39e-11 | 2.49e-06 |
| Fibroblast | F0 SFRP2+ Fib | RARRES1 | 2.471 | 0.684 | 0.184 | 5.34e-61 | 1.8e-56 |
| Fibroblast | F0 SFRP2+ Fib | GFRA1 | 2.469 | 0.17 | 0.026 | 2.33e-16 | 7.86e-12 |
| Fibroblast | F0 SFRP2+ Fib | PDGFRL | 2.468 | 0.459 | 0.112 | 4.17e-37 | 1.41e-32 |
| Fibroblast | F0 SFRP2+ Fib | FLRT2 | 2.465 | 0.207 | 0.041 | 1.7e-17 | 5.74e-13 |
| Fibroblast | F0 SFRP2+ Fib | FMO2 | 2.463 | 0.133 | 0.025 | 8.89e-12 | 2.99e-07 |
| Fibroblast | F0 SFRP2+ Fib | FAM134B | 2.46 | 0.125 | 0.023 | 3.48e-11 | 1.17e-06 |
| Fibroblast | F0 SFRP2+ Fib | CLDN11 | 2.446 | 0.308 | 0.07 | 6.01e-24 | 2.02e-19 |
| Fibroblast | F0 SFRP2+ Fib | GPC3 | 2.445 | 0.334 | 0.067 | 6.05e-29 | 2.04e-24 |
| Fibroblast | F0 SFRP2+ Fib | CPXM2 | 2.379 | 0.164 | 0.035 | 1.91e-13 | 6.44e-09 |
| Fibroblast | F0 SFRP2+ Fib | MMP23B | 2.373 | 0.472 | 0.133 | 5.61e-35 | 1.89e-30 |

| compartment | subcluster | gene | avg_log2FC | pct.1 | pct.2 | p_val | p_val_adj |
|---|---|---|---|---|---|---|---|
| Fibroblast | F0 SFRP2+ Fib | JAM2 | 2.368 | 0.231 | 0.035 | 2.28e-22 | 7.69e-18 |
| Fibroblast | F0 SFRP2+ Fib | CPXM1 | 2.344 | 0.496 | 0.129 | 1.61e-40 | 5.44e-36 |
| Fibroblast | F0 SFRP2+ Fib | MGP | 2.317 | 0.984 | 0.796 | 6.139999999999999e-92 | 2.07e-87 |
| Fibroblast | F0 SFRP2+ Fib | NRXN2 | 2.307 | 0.18 | 0.028 | 2.85e-17 | 9.6e-13 |
| Fibroblast | F0 SFRP2+ Fib | SERPINE2 | 2.299 | 0.615 | 0.201 | 7.8e-43 | 2.63e-38 |
| Fibroblast | F0 SFRP2+ Fib | ABCA8 | 2.287 | 0.512 | 0.083 | 9.25e-52 | 3.12e-47 |
| Fibroblast | F0 SFRP2+ Fib | RAMP2 | 2.286 | 0.18 | 0.035 | 2.64e-15 | 8.91e-11 |
| Fibroblast | F0 SFRP2+ Fib | APOD | 2.284 | 0.775 | 0.233 | 2.39e-62 | 8.05e-58 |
| Fibroblast | F0 SFRP2+ Fib | PRICKLE1 | 2.26 | 0.154 | 0.034 | 3.07e-12 | 1.03e-07 |
| Fibroblast | F0 SFRP2+ Fib | COLEC12 | 2.234 | 0.416 | 0.092 | 4.49e-35 | 1.51e-30 |
| Fibroblast | F0 SFRP2+ Fib | CCDC80 | 2.223 | 0.95 | 0.484 | 1.19e-86 | 3.99e-82 |
| Fibroblast | F0 SFRP2+ Fib | FGL2 | 2.22 | 0.223 | 0.025 | 9.01e-25 | 3.04e-20 |
| Fibroblast | F0 SFRP2+ Fib | FBLN1 | 2.208 | 0.971 | 0.417 | 1.46e-96 | 4.91e-92 |
| Fibroblast | F0 SFRP2+ Fib | STEAP2 | 2.198 | 0.255 | 0.06 | 2.26e-19 | 7.62e-15 |
| Fibroblast | F0 SFRP2+ Fib | IL11RA | 2.196 | 0.265 | 0.054 | 4.08e-22 | 1.37e-17 |
| Fibroblast | F0 SFRP2+ Fib | HAS1 | 2.177 | 0.178 | 0.035 | 1.31e-14 | 4.41e-10 |
| Fibroblast | F0 SFRP2+ Fib | HSD11B1 | 2.165 | 0.273 | 0.066 | 4.32e-20 | 1.46e-15 |
| Fibroblast | F0 SFRP2+ Fib | GSN | 2.154 | 0.976 | 0.734 | 4.01e-62 | 1.35e-57 |
| Fibroblast | F0 SFRP2+ Fib | ADH1B | 2.146 | 0.544 | 0.1 | 3.02e-52 | 1.02e-47 |
| Fibroblast | F0 SFRP2+ Fib | SVEP1 | 2.136 | 0.231 | 0.049 | 1.27e-18 | 4.29e-14 |
| Fibroblast | F0 SFRP2+ Fib | RNF212 | 2.135 | 0.133 | 0.028 | 7.73e-11 | 2.61e-06 |
| Fibroblast | F0 SFRP2+ Fib | UBXN8 | 2.133 | 0.109 | 0.015 | 3.06e-11 | 1.03e-06 |
| Fibroblast | F0 SFRP2+ Fib | PODN | 2.123 | 0.584 | 0.133 | 1.22e-51 | 4.12e-47 |
| Fibroblast | F0 SFRP2+ Fib | C1QTNF3 | 2.12 | 0.273 | 0.055 | 1.68e-22 | 5.66e-18 |
| Fibroblast | F0 SFRP2+ Fib | LPAR1 | 2.119 | 0.231 | 0.052 | 9.91e-18 | 3.34e-13 |
| Fibroblast | F0 SFRP2+ Fib | ARHGEF26 | 2.118 | 0.109 | 0.017 | 1.03e-10 | 3.48e-06 |
| Fibroblast | F0 SFRP2+ Fib | FOXP2 | 2.114 | 0.109 | 0.02 | 1e-09 | 3.37e-05 |
| Fibroblast | F0 SFRP2+ Fib | PTGIS | 2.111 | 0.334 | 0.078 | 3.45e-26 | 1.16e-21 |
| Fibroblast | F0 SFRP2+ Fib | BDKRB1 | 2.11 | 0.111 | 0.032 | 2.47e-07 | 0.00833 |
| Fibroblast | F0 SFRP2+ Fib | MAPKAP1 | 2.076 | 0.297 | 0.113 | 1.82e-14 | 6.14e-10 |
| Fibroblast | F0 SFRP2+ Fib | CXCL12 | 2.054 | 0.775 | 0.292 | 8.61e-55 | 2.9e-50 |
| Fibroblast | F0 SFRP2+ Fib | ZFHX4 | 2.036 | 0.194 | 0.046 | 2.63e-14 | 8.87e-10 |
| Fibroblast | F0 SFRP2+ Fib | CES1 | 2.029 | 0.239 | 0.055 | 3.74e-18 | 1.26e-13 |
| Fibroblast | F0 SFRP2+ Fib | ALDH1A1 | 2.027 | 0.525 | 0.112 | 6.830000000000001e-47 | 2.3e-42 |
| Fibroblast | F0 SFRP2+ Fib | DIO3OS | 2.021 | 0.361 | 0.081 | 6.81e-28 | 2.3e-23 |
| Fibroblast | F0 SFRP2+ Fib | DEPTOR | 2.021 | 0.114 | 0.031 | 4.72e-08 | 0.00159 |
| Fibroblast | F0 SFRP2+ Fib | CSTA | 2.017 | 0.138 | 0.038 | 4.07e-09 | 0.000137 |
| Fibroblast | F0 SFRP2+ Fib | STEAP1 | 2.012 | 0.525 | 0.145 | 1.52e-40 | 5.11e-36 |
| Fibroblast | F0 SFRP2+ Fib | PDGFRA | 2.011 | 0.56 | 0.161 | 5.49e-43 | 1.85e-38 |
| Fibroblast | F0 SFRP2+ Fib | SPON1 | 1.999 | 0.435 | 0.103 | 1.84e-34 | 6.200000000000001e-30 |
| Fibroblast | F0 SFRP2+ Fib | GREM1 | 1.998 | 0.247 | 0.087 | 7.06e-13 | 2.38e-08 |
| Fibroblast | F0 SFRP2+ Fib | FBLN5 | 1.995 | 0.655 | 0.172 | 2.16e-55 | 7.29e-51 |
| Fibroblast | F0 SFRP2+ Fib | RNF144A | 1.987 | 0.119 | 0.037 | 2.47e-07 | 0.00834 |
| Fibroblast | F0 SFRP2+ Fib | GSTM5 | 1.98 | 0.284 | 0.051 | 2.8e-25 | 9.450000000000001e-21 |
| Fibroblast | F0 SFRP2+ Fib | NT5E | 1.973 | 0.241 | 0.069 | 9.99e-16 | 3.37e-11 |
| Fibroblast | F0 SFRP2+ Fib | GPNMB | 1.964 | 0.772 | 0.27 | 8.08e-59 | 2.72e-54 |
| Fibroblast | F0 SFRP2+ Fib | SAMD11 | 1.953 | 0.284 | 0.081 | 2.99e-18 | 1.01e-13 |
| Fibroblast | F0 SFRP2+ Fib | SERPINF1 | 1.952 | 0.984 | 0.513 | 9.17e-96 | 3.09e-91 |
| Fibroblast | F0 SFRP2+ Fib | TSHZ2 | 1.944 | 0.684 | 0.207 | 8.42e-54 | 2.84e-49 |
| Fibroblast | F0 SFRP2+ Fib | SH3D19 | 1.925 | 0.268 | 0.063 | 3.51e-20 | 1.18e-15 |
| Fibroblast | F0 SFRP2+ Fib | ANK2 | 1.919 | 0.215 | 0.046 | 8.78e-17 | 2.96e-12 |
| Fibroblast | F0 SFRP2+ Fib | CHEK2 | 1.918 | 0.143 | 0.028 | 5.09e-12 | 1.71e-07 |
| Fibroblast | F0 SFRP2+ Fib | SLIT2 | 1.91 | 0.324 | 0.075 | 1.38e-24 | 4.65e-20 |
| Fibroblast | F0 SFRP2+ Fib | SCN1B | 1.908 | 0.111 | 0.029 | 6.26e-08 | 0.00211 |
| Fibroblast | F0 SFRP2+ Fib | IGFBP4 | 1.901 | 0.963 | 0.706 | 3.08e-70 | 1.04e-65 |
| Fibroblast | F0 SFRP2+ Fib | OSR2 | 1.897 | 0.355 | 0.118 | 4.46e-20 | 1.5e-15 |
| Fibroblast | F0 SFRP2+ Fib | CPNE8 | 1.888 | 0.122 | 0.038 | 2.4e-07 | 0.00808 |
| Fibroblast | F0 SFRP2+ Fib | MAN1C1 | 1.886 | 0.156 | 0.051 | 4.49e-09 | 0.000151 |
| Fibroblast | F0 SFRP2+ Fib | DPEP1 | 1.886 | 0.111 | 0.035 | 1.07e-06 | 0.0359 |
| Fibroblast | F0 SFRP2+ Fib | CFD | 1.884 | 0.732 | 0.187 | 3.83e-65 | 1.29e-60 |
| Fibroblast | F0 SFRP2+ Fib | NGFR | 1.877 | 0.26 | 0.051 | 5.47e-22 | 1.84e-17 |
| Fibroblast | F0 SFRP2+ Fib | AKR1C1 | 1.859 | 0.265 | 0.063 | 2.03e-19 | 6.84e-15 |
| Fibroblast | F0 SFRP2+ Fib | IGF2 | 1.854 | 0.18 | 0.037 | 4.7e-15 | 1.58e-10 |
| Fibroblast | F0 SFRP2+ Fib | VIPR2 | 1.853 | 0.207 | 0.052 | 2.51e-14 | 8.45e-10 |

| compartment | subcluster | gene | avg_log2FC | pct.1 | pct.2 | p_val | p_val_adj |
|---|---|---|---|---|---|---|---|
| Fibroblast | F0 SFRP2+ Fib | PRELP | 1.849 | 0.658 | 0.234 | 1.42e-40 | 4.79e-36 |
| Fibroblast | F0 SFRP2+ Fib | LRRN4CL | 1.847 | 0.183 | 0.041 | 8.61e-14 | 2.9e-09 |
| Fibroblast | F0 SFRP2+ Fib | TWIST2 | 1.844 | 0.355 | 0.095 | 2.19e-24 | 7.39e-20 |
| Fibroblast | F0 SFRP2+ Fib | LAMA2 | 1.839 | 0.233 | 0.052 | 8.09e-18 | 2.72e-13 |
| Fibroblast | F0 SFRP2+ Fib | DCN | 1.832 | 0.989 | 0.658 | 1.53e-91 | 5.16e-87 |
| Fibroblast | F0 SFRP2+ Fib | RGL1 | 1.832 | 0.117 | 0.025 | 1.71e-09 | 5.76e-05 |
| Fibroblast | F0 SFRP2+ Fib | CHL1 | 1.83 | 0.117 | 0.035 | 2.84e-07 | 0.00958 |
| Fibroblast | F0 SFRP2+ Fib | PTPRN2 | 1.815 | 0.13 | 0.026 | 8.28e-11 | 2.79e-06 |
| Fibroblast | F0 SFRP2+ Fib | CFB | 1.803 | 0.268 | 0.075 | 2.12e-17 | 7.13e-13 |
| Fibroblast | F0 SFRP2+ Fib | IGFBP6 | 1.797 | 0.737 | 0.378 | 3.19e-33 | 1.08e-28 |
| Fibroblast | F0 SFRP2+ Fib | AKR1C3 | 1.796 | 0.26 | 0.06 | 3.55e-19 | 1.2e-14 |
| Fibroblast | F0 SFRP2+ Fib | ABCA6 | 1.792 | 0.271 | 0.077 | 1.27e-17 | 4.29e-13 |
| Fibroblast | F0 SFRP2+ Fib | PLBD1 | 1.785 | 0.403 | 0.121 | 1.69e-25 | 5.690000000000001e-21 |
| Fibroblast | F0 SFRP2+ Fib | TGFBR3 | 1.78 | 0.194 | 0.052 | 1.93e-12 | 6.51e-08 |
| Fibroblast | F0 SFRP2+ Fib | FGF7 | 1.779 | 0.761 | 0.251 | 1.33e-55 | 4.48e-51 |
| Fibroblast | F0 SFRP2+ Fib | APPL2 | 1.773 | 0.162 | 0.043 | 9.44e-11 | 3.18e-06 |
| Fibroblast | F0 SFRP2+ Fib | DHRS3 | 1.762 | 0.43 | 0.147 | 1.07e-24 | 3.6e-20 |
| Fibroblast | F0 SFRP2+ Fib | BST1 | 1.756 | 0.167 | 0.058 | 8.22e-09 | 0.000277 |
| Fibroblast | F0 SFRP2+ Fib | SDC3 | 1.737 | 0.101 | 0.023 | 5.43e-08 | 0.00183 |
| Fibroblast | F0 SFRP2+ Fib | CYP1B1 | 1.734 | 0.366 | 0.127 | 1.84e-19 | 6.22e-15 |
| Fibroblast | F0 SFRP2+ Fib | OLFML3 | 1.732 | 0.756 | 0.271 | 1.08e-53 | 3.63e-49 |
| Fibroblast | F0 SFRP2+ Fib | SEPP1 | 1.731 | 0.881 | 0.392 | 4.29e-61 | 1.45e-56 |
| Fibroblast | F0 SFRP2+ Fib | SAMHD1 | 1.712 | 0.3 | 0.093 | 2.08e-17 | 7.01e-13 |
| Fibroblast | F0 SFRP2+ Fib | SEMA3C | 1.712 | 0.194 | 0.058 | 1.2e-11 | 4.05e-07 |
| Fibroblast | F0 SFRP2+ Fib | FBLN2 | 1.708 | 0.607 | 0.21 | 2.3e-39 | 7.760000000000001e-35 |
| Fibroblast | F0 SFRP2+ Fib | MMP2 | 1.696 | 0.958 | 0.417 | 3.49e-72 | 1.18e-67 |
| Fibroblast | F0 SFRP2+ Fib | OAF | 1.691 | 0.645 | 0.23 | 1.03e-39 | 3.46e-35 |
| Fibroblast | F0 SFRP2+ Fib | SSPN | 1.687 | 0.653 | 0.262 | 8.62e-37 | 2.9e-32 |
| Fibroblast | F0 SFRP2+ Fib | BIN1 | 1.685 | 0.305 | 0.104 | 2.86e-16 | 9.62e-12 |
| Fibroblast | F0 SFRP2+ Fib | BNC2 | 1.682 | 0.294 | 0.086 | 3.22e-18 | 1.09e-13 |
| Fibroblast | F0 SFRP2+ Fib | LSAMP | 1.679 | 0.363 | 0.1 | 2.89e-24 | 9.74e-20 |
| Fibroblast | F0 SFRP2+ Fib | MARCH3 | 1.665 | 0.141 | 0.031 | 5.5e-11 | 1.85e-06 |
| Fibroblast | F0 SFRP2+ Fib | ITPKC | 1.661 | 0.117 | 0.032 | 8.59e-08 | 0.0029 |
| Fibroblast | F0 SFRP2+ Fib | MEDAG | 1.658 | 0.228 | 0.058 | 1.06e-15 | 3.58e-11 |
| Fibroblast | F0 SFRP2+ Fib | MOXD1 | 1.653 | 0.276 | 0.098 | 3.76e-14 | 1.27e-09 |
| Fibroblast | F0 SFRP2+ Fib | PCDH9 | 1.648 | 0.138 | 0.028 | 2.12e-11 | 7.14e-07 |
| Fibroblast | F0 SFRP2+ Fib | SFRP4 | 1.633 | 0.671 | 0.221 | 1.23e-45 | 4.14e-41 |
| Fibroblast | F0 SFRP2+ Fib | MBP | 1.629 | 0.239 | 0.07 | 3.86e-14 | 1.3e-09 |
| Fibroblast | F0 SFRP2+ Fib | MID1 | 1.622 | 0.13 | 0.035 | 1.16e-08 | 0.000389 |
| Fibroblast | F0 SFRP2+ Fib | SH3BP5 | 1.621 | 0.408 | 0.133 | 1.22e-23 | 4.11e-19 |
| Fibroblast | F0 SFRP2+ Fib | TCN2 | 1.611 | 0.233 | 0.054 | 3.83e-17 | 1.29e-12 |
| Fibroblast | F0 SFRP2+ Fib | RAI2 | 1.611 | 0.199 | 0.043 | 1.26e-15 | 4.25e-11 |
| Fibroblast | F0 SFRP2+ Fib | SULT1A1 | 1.603 | 0.233 | 0.06 | 5.93e-16 | 2e-11 |
| Fibroblast | F0 SFRP2+ Fib | DNM1 | 1.593 | 0.117 | 0.037 | 7e-07 | 0.0236 |
| Fibroblast | F0 SFRP2+ Fib | AR | 1.59 | 0.109 | 0.032 | 7.92e-07 | 0.0267 |
| Fibroblast | F0 SFRP2+ Fib | CYBRD1 | 1.582 | 0.727 | 0.288 | 4.09e-44 | 1.38e-39 |
| Fibroblast | F0 SFRP2+ Fib | SH3PXD2B | 1.574 | 0.225 | 0.08 | 2.56e-11 | 8.64e-07 |
| Fibroblast | F0 SFRP2+ Fib | SNED1 | 1.568 | 0.167 | 0.046 | 6.35e-11 | 2.14e-06 |
| Fibroblast | F0 SFRP2+ Fib | CLIC2 | 1.56 | 0.223 | 0.047 | 1.99e-17 | 6.72e-13 |
| Fibroblast | F0 SFRP2+ Fib | COL14A1 | 1.559 | 0.645 | 0.292 | 1.3e-31 | 4.37e-27 |
| Fibroblast | F0 SFRP2+ Fib | PLTP | 1.557 | 0.695 | 0.268 | 5.61e-41 | 1.89e-36 |
| Fibroblast | F0 SFRP2+ Fib | CELF2 | 1.556 | 0.265 | 0.08 | 6.47e-16 | 2.18e-11 |
| Fibroblast | F0 SFRP2+ Fib | MATN2 | 1.555 | 0.223 | 0.084 | 3.12e-10 | 1.05e-05 |
| Fibroblast | F0 SFRP2+ Fib | CFI | 1.537 | 0.358 | 0.123 | 7.4e-19 | 2.49e-14 |
| Fibroblast | F0 SFRP2+ Fib | CENPW | 1.536 | 0.135 | 0.037 | 6.5e-09 | 0.000219 |
| Fibroblast | F0 SFRP2+ Fib | PLSCR4 | 1.527 | 0.374 | 0.138 | 8.95e-19 | 3.02e-14 |
| Fibroblast | F0 SFRP2+ Fib | NID2 | 1.527 | 0.188 | 0.064 | 6.67e-10 | 2.25e-05 |
| Fibroblast | F0 SFRP2+ Fib | C1S | 1.516 | 0.987 | 0.609 | 5.03e-77 | 1.69e-72 |
| Fibroblast | F0 SFRP2+ Fib | CFH | 1.513 | 0.721 | 0.345 | 1.37e-32 | 4.62e-28 |
| Fibroblast | F0 SFRP2+ Fib | DOK5 | 1.513 | 0.164 | 0.066 | 3e-07 | 0.0101 |
| Fibroblast | F0 SFRP2+ Fib | PTGDS | 1.51 | 0.647 | 0.208 | 1.07e-42 | 3.6e-38 |
| Fibroblast | F0 SFRP2+ Fib | CBR3 | 1.504 | 0.199 | 0.07 | 6.48e-10 | 2.18e-05 |
| Fibroblast | F0 SFRP2+ Fib | ABHD14A | 1.503 | 0.3 | 0.109 | 3.97e-15 | 1.34e-10 |
| Fibroblast | F0 SFRP2+ Fib | RNASE4 | 1.501 | 0.31 | 0.118 | 2.77e-14 | 9.33e-10 |

| compartment | subcluster | gene | avg_log2FC | pct.1 | pct.2 | p_val | p_val_adj |
|---|---|---|---|---|---|---|---|
| Fibroblast | F0 SFRP2+ Fib | PRKAG2-AS1 | 1.501 | 0.183 | 0.04 | 5.43e-14 | 1.83e-09 |
| Fibroblast | F0 SFRP2+ Fib | BICC1 | 1.499 | 0.422 | 0.156 | 1.96e-21 | 6.59e-17 |
| Fibroblast | F0 SFRP2+ Fib | ADAM33 | 1.497 | 0.183 | 0.044 | 5.2e-13 | 1.75e-08 |
| Fibroblast | F0 SFRP2+ Fib | GFPT2 | 1.49 | 0.297 | 0.095 | 1.98e-16 | 6.68e-12 |
| Fibroblast | F0 SFRP2+ Fib | VWA5A | 1.488 | 0.103 | 0.029 | 7.62e-07 | 0.0257 |
| Fibroblast | F0 SFRP2+ Fib | FUCA1 | 1.471 | 0.188 | 0.06 | 1.43e-10 | 4.82e-06 |
| Fibroblast | F0 SFRP2+ Fib | ICAM1 | 1.469 | 0.488 | 0.219 | 1.17e-19 | 3.95e-15 |
| Fibroblast | F0 SFRP2+ Fib | TGS1 | 1.466 | 0.143 | 0.049 | 1.58e-07 | 0.00533 |
| Fibroblast | F0 SFRP2+ Fib | DAB2 | 1.464 | 0.594 | 0.225 | 9.200000000000001e-33 | 3.1e-28 |
| Fibroblast | F0 SFRP2+ Fib | PDGFD | 1.462 | 0.133 | 0.044 | 2.51e-07 | 0.00844 |
| Fibroblast | F0 SFRP2+ Fib | C1R | 1.46 | 0.968 | 0.642 | 1.24e-65 | 4.18e-61 |
| Fibroblast | F0 SFRP2+ Fib | LIPT1 | 1.452 | 0.103 | 0.028 | 3.3e-07 | 0.0111 |
| Fibroblast | F0 SFRP2+ Fib | APOL3 | 1.449 | 0.286 | 0.09 | 4.97e-16 | 1.67e-11 |
| Fibroblast | F0 SFRP2+ Fib | S1PR2 | 1.444 | 0.17 | 0.063 | 3.9e-08 | 0.00132 |
| Fibroblast | F0 SFRP2+ Fib | LXN | 1.439 | 0.472 | 0.156 | 1.74e-25 | 5.860000000000001e-21 |
| Fibroblast | F0 SFRP2+ Fib | MEOX2 | 1.43 | 0.164 | 0.051 | 1.61e-09 | 5.43e-05 |
| Fibroblast | F0 SFRP2+ Fib | ANGPTL2 | 1.429 | 0.416 | 0.156 | 5.750000000000001e-21 | 1.94e-16 |
| Fibroblast | F0 SFRP2+ Fib | PARP16 | 1.429 | 0.13 | 0.038 | 4.37e-08 | 0.00147 |
| Fibroblast | F0 SFRP2+ Fib | PLXDC2 | 1.422 | 0.438 | 0.165 | 3.980000000000001e-21 | 1.34e-16 |
| Fibroblast | F0 SFRP2+ Fib | MMP19 | 1.422 | 0.289 | 0.129 | 6.56e-11 | 2.21e-06 |
| Fibroblast | F0 SFRP2+ Fib | FMO1 | 1.42 | 0.103 | 0.026 | 1.6e-07 | 0.00538 |
| Fibroblast | F0 SFRP2+ Fib | EPB41L3 | 1.417 | 0.101 | 0.023 | 5.73e-08 | 0.00193 |
| Fibroblast | F0 SFRP2+ Fib | EXT1 | 1.413 | 0.156 | 0.054 | 3.37e-08 | 0.00114 |
| Fibroblast | F0 SFRP2+ Fib | COL6A3 | 1.402 | 0.881 | 0.407 | 3.6e-52 | 1.21e-47 |
| Fibroblast | F0 SFRP2+ Fib | SETD7 | 1.396 | 0.162 | 0.06 | 8.37e-08 | 0.00282 |
| Fibroblast | F0 SFRP2+ Fib | DHRS12 | 1.393 | 0.17 | 0.055 | 2.84e-09 | 9.58e-05 |
| Fibroblast | F0 SFRP2+ Fib | TGFBR2 | 1.389 | 0.302 | 0.124 | 4.28e-13 | 1.44e-08 |
| Fibroblast | F0 SFRP2+ Fib | CCDC109B | 1.378 | 0.393 | 0.144 | 2.84e-19 | 9.57e-15 |
| Fibroblast | F0 SFRP2+ Fib | C1orf21 | 1.376 | 0.552 | 0.239 | 4.67e-24 | 1.57e-19 |
| Fibroblast | F0 SFRP2+ Fib | TNFSF10 | 1.375 | 0.43 | 0.15 | 1.61e-22 | 5.43e-18 |
| Fibroblast | F0 SFRP2+ Fib | GAS7 | 1.374 | 0.191 | 0.064 | 3.29e-10 | 1.11e-05 |
| Fibroblast | F0 SFRP2+ Fib | ABCC9 | 1.366 | 0.151 | 0.041 | 1.09e-09 | 3.67e-05 |
| Fibroblast | F0 SFRP2+ Fib | GGT5 | 1.364 | 0.721 | 0.343 | 1.42e-34 | 4.78e-30 |
| Fibroblast | F0 SFRP2+ Fib | LUM | 1.358 | 0.987 | 0.489 | 2.36e-59 | 7.939999999999999e-55 |
| Fibroblast | F0 SFRP2+ Fib | MITF | 1.352 | 0.337 | 0.106 | 1.25e-18 | 4.22e-14 |
| Fibroblast | F0 SFRP2+ Fib | S100A13 | 1.338 | 0.857 | 0.484 | 6.48e-44 | 2.18e-39 |
| Fibroblast | F0 SFRP2+ Fib | CLU | 1.337 | 0.676 | 0.254 | 1.12e-38 | 3.78e-34 |
| Fibroblast | F0 SFRP2+ Fib | SCPEP1 | 1.336 | 0.663 | 0.322 | 5.57e-29 | 1.88e-24 |
| Fibroblast | F0 SFRP2+ Fib | DYNC2LI1 | 1.334 | 0.207 | 0.084 | 1.25e-08 | 0.000422 |
| Fibroblast | F0 SFRP2+ Fib | IDUA | 1.333 | 0.17 | 0.052 | 9.05e-10 | 3.05e-05 |
| Fibroblast | F0 SFRP2+ Fib | AHNAK2 | 1.333 | 0.167 | 0.058 | 1.82e-08 | 0.000612 |
| Fibroblast | F0 SFRP2+ Fib | DDX3Y | 1.318 | 0.239 | 0.096 | 8.6e-10 | 2.9e-05 |
| Fibroblast | F0 SFRP2+ Fib | METTL7A | 1.316 | 0.467 | 0.172 | 1.58e-23 | 5.34e-19 |
| Fibroblast | F0 SFRP2+ Fib | CD68 | 1.315 | 0.366 | 0.13 | 5.85e-18 | 1.97e-13 |
| Fibroblast | F0 SFRP2+ Fib | CST3 | 1.314 | 0.995 | 0.876 | 5.46e-73 | 1.84e-68 |
| Fibroblast | F0 SFRP2+ Fib | ADD3 | 1.311 | 0.475 | 0.168 | 3.15e-24 | 1.06e-19 |
| Fibroblast | F0 SFRP2+ Fib | TNFAIP2 | 1.309 | 0.284 | 0.103 | 9.39e-14 | 3.16e-09 |
| Fibroblast | F0 SFRP2+ Fib | NINJ2 | 1.309 | 0.167 | 0.063 | 5.62e-08 | 0.00189 |
| Fibroblast | F0 SFRP2+ Fib | OLFML1 | 1.308 | 0.326 | 0.135 | 1.77e-13 | 5.95e-09 |
| Fibroblast | F0 SFRP2+ Fib | TNFSF13B | 1.306 | 0.316 | 0.106 | 2.27e-16 | 7.64e-12 |
| Fibroblast | F0 SFRP2+ Fib | PLPP3 | 1.305 | 0.504 | 0.194 | 2.7e-24 | 9.11e-20 |
| Fibroblast | F0 SFRP2+ Fib | AGTR1 | 1.305 | 0.117 | 0.025 | 1.64e-09 | 5.53e-05 |
| Fibroblast | F0 SFRP2+ Fib | NR2F1 | 1.302 | 0.305 | 0.104 | 7.16e-16 | 2.41e-11 |
| Fibroblast | F0 SFRP2+ Fib | PXDC1 | 1.298 | 0.302 | 0.113 | 6.2e-14 | 2.09e-09 |
| Fibroblast | F0 SFRP2+ Fib | ARSA | 1.295 | 0.247 | 0.103 | 4.22e-10 | 1.42e-05 |
| Fibroblast | F0 SFRP2+ Fib | LRP1 | 1.293 | 0.751 | 0.335 | 2.16e-38 | 7.279999999999999e-34 |
| Fibroblast | F0 SFRP2+ Fib | NFIB | 1.287 | 0.594 | 0.237 | 3.1e-29 | 1.05e-24 |
| Fibroblast | F0 SFRP2+ Fib | CTSF | 1.287 | 0.605 | 0.303 | 3.34e-22 | 1.12e-17 |
| Fibroblast | F0 SFRP2+ Fib | KCTD12 | 1.285 | 0.263 | 0.109 | 8.41e-11 | 2.83e-06 |
| Fibroblast | F0 SFRP2+ Fib | BASP1 | 1.284 | 0.528 | 0.23 | 4.51e-21 | 1.52e-16 |
| Fibroblast | F0 SFRP2+ Fib | ACVRL1 | 1.284 | 0.26 | 0.096 | 5.19e-12 | 1.75e-07 |
| Fibroblast | F0 SFRP2+ Fib | ELN | 1.28 | 0.565 | 0.237 | 1.15e-24 | 3.88e-20 |
| Fibroblast | F0 SFRP2+ Fib | PHYHIP | 1.28 | 0.141 | 0.035 | 8.63e-10 | 2.91e-05 |
| Fibroblast | F0 SFRP2+ Fib | NOVA1 | 1.275 | 0.247 | 0.067 | 2.81e-15 | 9.45e-11 |

| compartment | subcluster | gene | avg_log2FC | pct.1 | pct.2 | p_val | p_val_adj |
|---|---|---|---|---|---|---|---|
| Fibroblast | F0 SFRP2+ Fib | MFAP5 | 1.265 | 0.332 | 0.121 | 4.19e-16 | 1.41e-11 |
| Fibroblast | F0 SFRP2+ Fib | COPS8 | 1.261 | 0.562 | 0.254 | 6.49e-22 | 2.19e-17 |
| Fibroblast | F0 SFRP2+ Fib | CLMP | 1.261 | 0.347 | 0.138 | 3.45e-15 | 1.16e-10 |
| Fibroblast | F0 SFRP2+ Fib | GPRC5A | 1.26 | 0.355 | 0.141 | 4.67e-15 | 1.57e-10 |
| Fibroblast | F0 SFRP2+ Fib | ZNF423 | 1.256 | 0.133 | 0.029 | 2.96e-10 | 9.98e-06 |
| Fibroblast | F0 SFRP2+ Fib | LPAR6 | 1.251 | 0.26 | 0.104 | 6.18e-11 | 2.08e-06 |
| Fibroblast | F0 SFRP2+ Fib | SSC5D | 1.251 | 0.218 | 0.077 | 1.02e-10 | 3.44e-06 |
| Fibroblast | F0 SFRP2+ Fib | BTN3A3 | 1.248 | 0.167 | 0.069 | 7.85e-07 | 0.0265 |
| Fibroblast | F0 SFRP2+ Fib | IL33 | 1.247 | 0.18 | 0.054 | 1.18e-10 | 3.96e-06 |
| Fibroblast | F0 SFRP2+ Fib | FBXO17 | 1.245 | 0.308 | 0.127 | 3.98e-12 | 1.34e-07 |
| Fibroblast | F0 SFRP2+ Fib | PRCP | 1.224 | 0.403 | 0.173 | 4.61e-16 | 1.55e-11 |
| Fibroblast | F0 SFRP2+ Fib | THNSL2 | 1.223 | 0.178 | 0.064 | 1.89e-08 | 0.000636 |
| Fibroblast | F0 SFRP2+ Fib | CTGF | 1.218 | 0.894 | 0.666 | 3.6e-20 | 1.21e-15 |
| Fibroblast | F0 SFRP2+ Fib | MAN1A1 | 1.218 | 0.271 | 0.104 | 9.34e-12 | 3.15e-07 |
| Fibroblast | F0 SFRP2+ Fib | VEGFB | 1.216 | 0.249 | 0.113 | 5.77e-09 | 0.000195 |
| Fibroblast | F0 SFRP2+ Fib | ADAMTSL4 | 1.216 | 0.122 | 0.034 | 5.9e-08 | 0.00199 |
| Fibroblast | F0 SFRP2+ Fib | PCOLCE | 1.213 | 0.857 | 0.505 | 6.21e-39 | 2.09e-34 |
| Fibroblast | F0 SFRP2+ Fib | TPBG | 1.207 | 0.385 | 0.172 | 7.59e-15 | 2.56e-10 |
| Fibroblast | F0 SFRP2+ Fib | NHLRC3 | 1.203 | 0.268 | 0.089 | 7.39e-14 | 2.49e-09 |
| Fibroblast | F0 SFRP2+ Fib | FCGRT | 1.195 | 0.838 | 0.438 | 6.36e-42 | 2.14e-37 |
| Fibroblast | F0 SFRP2+ Fib | PBX3 | 1.185 | 0.22 | 0.086 | 1.43e-09 | 4.83e-05 |
| Fibroblast | F0 SFRP2+ Fib | NFIA | 1.181 | 0.682 | 0.306 | 5.589999999999999e-30 | 1.88e-25 |
| Fibroblast | F0 SFRP2+ Fib | PLBD2 | 1.178 | 0.212 | 0.086 | 1.57e-08 | 0.00053 |
| Fibroblast | F0 SFRP2+ Fib | EMP1 | 1.176 | 0.785 | 0.4 | 1.41e-31 | 4.770000000000001e-27 |
| Fibroblast | F0 SFRP2+ Fib | VAT1 | 1.174 | 0.334 | 0.142 | 5.06e-13 | 1.7e-08 |
| Fibroblast | F0 SFRP2+ Fib | CTSK | 1.169 | 0.881 | 0.41 | 1.45e-45 | 4.88e-41 |
| Fibroblast | F0 SFRP2+ Fib | PTGS2 | 1.169 | 0.324 | 0.104 | 2.65e-17 | 8.93e-13 |
| Fibroblast | F0 SFRP2+ Fib | PLIN2 | 1.167 | 0.467 | 0.26 | 5.84e-12 | 1.97e-07 |
| Fibroblast | F0 SFRP2+ Fib | MPZL1 | 1.165 | 0.454 | 0.214 | 1.01e-15 | 3.39e-11 |
| Fibroblast | F0 SFRP2+ Fib | MEIS1 | 1.162 | 0.239 | 0.103 | 4.4e-09 | 0.000148 |
| Fibroblast | F0 SFRP2+ Fib | UAP1 | 1.161 | 0.414 | 0.172 | 8.92e-17 | 3e-12 |
| Fibroblast | F0 SFRP2+ Fib | FLCN | 1.16 | 0.186 | 0.066 | 5.48e-09 | 0.000185 |
| Fibroblast | F0 SFRP2+ Fib | RP11-1391J7.1 | 1.155 | 0.133 | 0.034 | 4.67e-09 | 0.000158 |
| Fibroblast | F0 SFRP2+ Fib | CYP27A1 | 1.153 | 0.297 | 0.109 | 1.55e-13 | 5.21e-09 |
| Fibroblast | F0 SFRP2+ Fib | ADCK3 | 1.149 | 0.154 | 0.047 | 1.23e-08 | 0.000413 |
| Fibroblast | F0 SFRP2+ Fib | CREB3L1 | 1.146 | 0.252 | 0.115 | 1.45e-08 | 0.000488 |
| Fibroblast | F0 SFRP2+ Fib | GAS6 | 1.145 | 0.645 | 0.351 | 3.78e-19 | 1.27e-14 |
| Fibroblast | F0 SFRP2+ Fib | NTPCR | 1.145 | 0.268 | 0.109 | 1.41e-10 | 4.75e-06 |
| Fibroblast | F0 SFRP2+ Fib | MFAP4 | 1.141 | 0.905 | 0.482 | 5.45e-42 | 1.84e-37 |
| Fibroblast | F0 SFRP2+ Fib | TXNIP | 1.136 | 0.838 | 0.524 | 3.38e-28 | 1.14e-23 |
| Fibroblast | F0 SFRP2+ Fib | SIL1 | 1.135 | 0.289 | 0.124 | 1.01e-10 | 3.4e-06 |
| Fibroblast | F0 SFRP2+ Fib | MIR99AHG | 1.133 | 0.257 | 0.083 | 2.4e-13 | 8.09e-09 |
| Fibroblast | F0 SFRP2+ Fib | FBN1 | 1.126 | 0.496 | 0.219 | 1.04e-19 | 3.52e-15 |
| Fibroblast | F0 SFRP2+ Fib | FIBIN | 1.125 | 0.329 | 0.155 | 2.54e-10 | 8.55e-06 |
| Fibroblast | F0 SFRP2+ Fib | MILR1 | 1.123 | 0.114 | 0.035 | 8.99e-07 | 0.0303 |
| Fibroblast | F0 SFRP2+ Fib | KAZN | 1.116 | 0.141 | 0.04 | 1.01e-08 | 0.000339 |
| Fibroblast | F0 SFRP2+ Fib | CRYL1 | 1.114 | 0.207 | 0.084 | 2.4e-08 | 0.000809 |
| Fibroblast | F0 SFRP2+ Fib | CCL19 | 1.11 | 0.149 | 0.046 | 1.21e-08 | 0.000408 |
| Fibroblast | F0 SFRP2+ Fib | CD302 | 1.106 | 0.44 | 0.168 | 7.39e-20 | 2.49e-15 |
| Fibroblast | F0 SFRP2+ Fib | ABL2 | 1.106 | 0.326 | 0.158 | 1.78e-10 | 5.99e-06 |
| Fibroblast | F0 SFRP2+ Fib | NFIX | 1.103 | 0.371 | 0.139 | 1.31e-16 | 4.4e-12 |
| Fibroblast | F0 SFRP2+ Fib | FAM110B | 1.102 | 0.18 | 0.077 | 6.95e-07 | 0.0234 |
| Fibroblast | F0 SFRP2+ Fib | MDK | 1.101 | 0.812 | 0.401 | 2.38e-34 | 8.03e-30 |
| Fibroblast | F0 SFRP2+ Fib | TAPBPL | 1.1 | 0.257 | 0.101 | 7.1e-11 | 2.39e-06 |
| Fibroblast | F0 SFRP2+ Fib | SLIT3 | 1.098 | 0.395 | 0.168 | 3.11e-15 | 1.05e-10 |
| Fibroblast | F0 SFRP2+ Fib | GNPDA2 | 1.096 | 0.167 | 0.069 | 8.84e-07 | 0.0298 |
| Fibroblast | F0 SFRP2+ Fib | ARAP1 | 1.091 | 0.212 | 0.075 | 3.31e-10 | 1.11e-05 |
| Fibroblast | F0 SFRP2+ Fib | XPNPEP1 | 1.089 | 0.204 | 0.09 | 1.9e-07 | 0.0064 |
| Fibroblast | F0 SFRP2+ Fib | ANXA1 | 1.088 | 0.873 | 0.59 | 5.35e-24 | 1.8e-19 |
| Fibroblast | F0 SFRP2+ Fib | LARP6 | 1.088 | 0.302 | 0.132 | 9.33e-11 | 3.14e-06 |
| Fibroblast | F0 SFRP2+ Fib | PLD3 | 1.083 | 0.692 | 0.337 | 1.44e-27 | 4.84e-23 |
| Fibroblast | F0 SFRP2+ Fib | TMEM45A | 1.083 | 0.477 | 0.242 | 9.79e-15 | 3.3e-10 |
| Fibroblast | F0 SFRP2+ Fib | SGCE | 1.082 | 0.552 | 0.256 | 1.51e-20 | 5.1e-16 |
| Fibroblast | F0 SFRP2+ Fib | EPHX1 | 1.082 | 0.565 | 0.259 | 2.32e-20 | 7.82e-16 |

| compartment | subcluster | gene | avg_log2FC | pct.1 | pct.2 | p_val | p_val_adj |
|---|---|---|---|---|---|---|---|
| Fibroblast | F0 SFRP2+ Fib | C2 | 1.082 | 0.403 | 0.176 | 3.31e-15 | 1.11e-10 |
| Fibroblast | F0 SFRP2+ Fib | LGALS3 | 1.08 | 0.905 | 0.536 | 3.66e-35 | 1.23e-30 |
| Fibroblast | F0 SFRP2+ Fib | CD320 | 1.08 | 0.252 | 0.098 | 1.37e-10 | 4.61e-06 |
| Fibroblast | F0 SFRP2+ Fib | TSPAN4 | 1.079 | 0.708 | 0.369 | 3.43e-26 | 1.16e-21 |
| Fibroblast | F0 SFRP2+ Fib | HSD17B11 | 1.074 | 0.549 | 0.253 | 5.27e-20 | 1.78e-15 |
| Fibroblast | F0 SFRP2+ Fib | GAS1 | 1.073 | 0.321 | 0.116 | 6.36e-15 | 2.14e-10 |
| Fibroblast | F0 SFRP2+ Fib | MEG3 | 1.069 | 0.756 | 0.364 | 4.51e-27 | 1.52e-22 |
| Fibroblast | F0 SFRP2+ Fib | PPP3CA | 1.068 | 0.387 | 0.176 | 2.59e-14 | 8.72e-10 |
| Fibroblast | F0 SFRP2+ Fib | PRRX2 | 1.068 | 0.133 | 0.047 | 1.44e-06 | 0.0487 |
| Fibroblast | F0 SFRP2+ Fib | ZCCHC24 | 1.067 | 0.21 | 0.081 | 4.89e-09 | 0.000165 |
| Fibroblast | F0 SFRP2+ Fib | MRC2 | 1.066 | 0.464 | 0.207 | 1.18e-17 | 3.97e-13 |
| Fibroblast | F0 SFRP2+ Fib | S100A10 | 1.062 | 0.878 | 0.501 | 1.38e-32 | 4.65e-28 |
| Fibroblast | F0 SFRP2+ Fib | HSPB6 | 1.062 | 0.435 | 0.165 | 1.36e-18 | 4.59e-14 |
| Fibroblast | F0 SFRP2+ Fib | SCARA3 | 1.058 | 0.188 | 0.063 | 1.88e-09 | 6.34e-05 |
| Fibroblast | F0 SFRP2+ Fib | PMP22 | 1.054 | 0.825 | 0.458 | 1.76e-31 | 5.930000000000001e-27 |
| Fibroblast | F0 SFRP2+ Fib | TMEM100 | 1.053 | 0.188 | 0.069 | 1.53e-08 | 0.000515 |
| Fibroblast | F0 SFRP2+ Fib | QSOX1 | 1.052 | 0.448 | 0.228 | 3.13e-13 | 1.06e-08 |
| Fibroblast | F0 SFRP2+ Fib | RUNX1T1 | 1.052 | 0.316 | 0.138 | 6.56e-11 | 2.21e-06 |
| Fibroblast | F0 SFRP2+ Fib | ZNF106 | 1.051 | 0.462 | 0.194 | 1.25e-18 | 4.22e-14 |
| Fibroblast | F0 SFRP2+ Fib | MDFIC | 1.048 | 0.21 | 0.093 | 2.48e-07 | 0.00837 |
| Fibroblast | F0 SFRP2+ Fib | UGDH | 1.046 | 0.408 | 0.179 | 1.88e-14 | 6.33e-10 |
| Fibroblast | F0 SFRP2+ Fib | RORA | 1.046 | 0.289 | 0.132 | 1.18e-09 | 3.96e-05 |
| Fibroblast | F0 SFRP2+ Fib | NEGR1 | 1.045 | 0.233 | 0.077 | 6.86e-12 | 2.31e-07 |
| Fibroblast | F0 SFRP2+ Fib | PTPRS | 1.044 | 0.252 | 0.121 | 4.7e-08 | 0.00158 |
| Fibroblast | F0 SFRP2+ Fib | PIK3R1 | 1.035 | 0.255 | 0.126 | 1.94e-07 | 0.00652 |
| Fibroblast | F0 SFRP2+ Fib | LGALS3BP | 1.032 | 0.899 | 0.545 | 7.22e-36 | 2.43e-31 |
| Fibroblast | F0 SFRP2+ Fib | TMEM119 | 1.031 | 0.294 | 0.112 | 1.64e-12 | 5.53e-08 |
| Fibroblast | F0 SFRP2+ Fib | EGR3 | 1.03 | 0.286 | 0.118 | 4.88e-11 | 1.64e-06 |
| Fibroblast | F0 SFRP2+ Fib | HOXA10 | 1.027 | 0.236 | 0.093 | 1.31e-09 | 4.42e-05 |
| Fibroblast | F0 SFRP2+ Fib | LHFPL2 | 1.024 | 0.186 | 0.07 | 3.97e-08 | 0.00134 |
| Fibroblast | F0 SFRP2+ Fib | IFNGR1 | 1.022 | 0.387 | 0.182 | 5.6e-13 | 1.89e-08 |
| Fibroblast | F0 SFRP2+ Fib | TGFB3 | 1.02 | 0.358 | 0.147 | 3.22e-14 | 1.09e-09 |
| Fibroblast | F0 SFRP2+ Fib | SLC16A4 | 1.016 | 0.172 | 0.072 | 9.96e-07 | 0.0336 |
| Fibroblast | F0 SFRP2+ Fib | NPDC1 | 1.014 | 0.459 | 0.211 | 7e-16 | 2.36e-11 |
| Fibroblast | F0 SFRP2+ Fib | PXK | 1.01 | 0.183 | 0.078 | 5.03e-07 | 0.017 |
| Fibroblast | F0 SFRP2+ Fib | KDSR | 1.009 | 0.427 | 0.205 | 5.9e-14 | 1.99e-09 |
| Fibroblast | F0 SFRP2+ Fib | CCDC71L | 1.009 | 0.289 | 0.115 | 5.66e-12 | 1.91e-07 |
| Fibroblast | F0 SFRP2+ Fib | IL15RA | 1.009 | 0.268 | 0.119 | 2.47e-09 | 8.32e-05 |
| Fibroblast | F0 SFRP2+ Fib | SLC35G2 | 1.008 | 0.122 | 0.035 | 1.41e-07 | 0.00474 |
| Fibroblast | F0 SFRP2+ Fib | COL16A1 | 1.006 | 0.313 | 0.152 | 2.2e-09 | 7.4e-05 |
| Fibroblast | F0 SFRP2+ Fib | PREB | 1.004 | 0.332 | 0.145 | 1.11e-11 | 3.73e-07 |
| Fibroblast | F0 SFRP2+ Fib | TMEM98 | 1.002 | 0.668 | 0.342 | 3.64e-22 | 1.23e-17 |
| Fibroblast | F0 SFRP2+ Fib | ANG | 1.002 | 0.292 | 0.107 | 7.42e-13 | 2.5e-08 |
| Fibroblast | F0 SFRP2+ Fib | IL1R1 | 0.999 | 0.366 | 0.165 | 1.05e-12 | 3.53e-08 |
| Fibroblast | F0 SFRP2+ Fib | EGR2 | 0.998 | 0.435 | 0.193 | 2.03e-15 | 6.84e-11 |
| Fibroblast | F0 SFRP2+ Fib | TMEM9B | 0.998 | 0.374 | 0.167 | 9.67e-13 | 3.26e-08 |
| Fibroblast | F0 SFRP2+ Fib | HACD3 | 0.998 | 0.305 | 0.144 | 5.98e-10 | 2.02e-05 |
| Fibroblast | F0 SFRP2+ Fib | GUSB | 0.992 | 0.358 | 0.164 | 3.74e-12 | 1.26e-07 |
| Fibroblast | F0 SFRP2+ Fib | EGFR | 0.992 | 0.175 | 0.061 | 2.2e-08 | 0.00074 |
| Fibroblast | F0 SFRP2+ Fib | GLT8D2 | 0.991 | 0.528 | 0.253 | 6.86e-18 | 2.31e-13 |
| Fibroblast | F0 SFRP2+ Fib | NFYB | 0.991 | 0.284 | 0.141 | 3.57e-08 | 0.0012 |
| Fibroblast | F0 SFRP2+ Fib | CLEC11A | 0.989 | 0.817 | 0.433 | 1.6e-26 | 5.41e-22 |
| Fibroblast | F0 SFRP2+ Fib | FAP | 0.989 | 0.483 | 0.21 | 5.79e-18 | 1.95e-13 |
| Fibroblast | F0 SFRP2+ Fib | DHRS7B | 0.989 | 0.183 | 0.075 | 3.37e-07 | 0.0113 |
| Fibroblast | F0 SFRP2+ Fib | SMARCA2 | 0.984 | 0.284 | 0.124 | 5e-10 | 1.68e-05 |
| Fibroblast | F0 SFRP2+ Fib | SLPI | 0.983 | 0.167 | 0.041 | 5.21e-12 | 1.76e-07 |
| Fibroblast | F0 SFRP2+ Fib | CHID1 | 0.981 | 0.568 | 0.266 | 1.85e-19 | 6.24e-15 |
| Fibroblast | F0 SFRP2+ Fib | LSP1 | 0.978 | 0.477 | 0.172 | 4.2e-23 | 1.42e-18 |
| Fibroblast | F0 SFRP2+ Fib | NNMT | 0.972 | 0.905 | 0.642 | 1.38e-24 | 4.65e-20 |
| Fibroblast | F0 SFRP2+ Fib | SPRY1 | 0.971 | 0.544 | 0.277 | 9.41e-17 | 3.17e-12 |
| Fibroblast | F0 SFRP2+ Fib | TPST1 | 0.971 | 0.403 | 0.181 | 1.73e-13 | 5.84e-09 |
| Fibroblast | F0 SFRP2+ Fib | ELMSAN1 | 0.97 | 0.252 | 0.124 | 4.13e-07 | 0.0139 |
| Fibroblast | F0 SFRP2+ Fib | SOD2 | 0.969 | 0.844 | 0.516 | 2.97e-29 | 1e-24 |
| Fibroblast | F0 SFRP2+ Fib | P4HA2 | 0.968 | 0.316 | 0.141 | 9.6e-11 | 3.23e-06 |

| compartment | subcluster | gene | avg_log2FC | pct.1 | pct.2 | p_val | p_val_adj |
|---|---|---|---|---|---|---|---|
| Fibroblast | F0 SFRP2+ Fib | SPG20 | 0.967 | 0.408 | 0.187 | 1.95e-14 | 6.58e-10 |
| Fibroblast | F0 SFRP2+ Fib | TMX4 | 0.966 | 0.318 | 0.144 | 4.1e-11 | 1.38e-06 |
| Fibroblast | F0 SFRP2+ Fib | CREG1 | 0.96 | 0.541 | 0.268 | 5.37e-18 | 1.81e-13 |
| Fibroblast | F0 SFRP2+ Fib | PTGFRN | 0.956 | 0.218 | 0.101 | 4.91e-07 | 0.0165 |
| Fibroblast | F0 SFRP2+ Fib | RP11-553L6.5 | 0.952 | 0.188 | 0.067 | 1.06e-08 | 0.000357 |
| Fibroblast | F0 SFRP2+ Fib | CPQ | 0.951 | 0.549 | 0.265 | 1.74e-18 | 5.86e-14 |
| Fibroblast | F0 SFRP2+ Fib | CDC27 | 0.951 | 0.188 | 0.084 | 1.33e-06 | 0.0448 |
| Fibroblast | F0 SFRP2+ Fib | CYB5A | 0.95 | 0.531 | 0.28 | 8.44e-15 | 2.84e-10 |
| Fibroblast | F0 SFRP2+ Fib | ALDH2 | 0.948 | 0.493 | 0.26 | 5.95e-14 | 2e-09 |
| Fibroblast | F0 SFRP2+ Fib | ANKRD35 | 0.948 | 0.244 | 0.115 | 7.75e-08 | 0.00261 |
| Fibroblast | F0 SFRP2+ Fib | P3H3 | 0.941 | 0.297 | 0.141 | 3.05e-09 | 0.000103 |
| Fibroblast | F0 SFRP2+ Fib | TCEAL7 | 0.941 | 0.321 | 0.161 | 9.56e-09 | 0.000322 |
| Fibroblast | F0 SFRP2+ Fib | ZNF226 | 0.941 | 0.167 | 0.067 | 9.4e-07 | 0.0317 |
| Fibroblast | F0 SFRP2+ Fib | SLC38A10 | 0.939 | 0.294 | 0.13 | 5.72e-10 | 1.93e-05 |
| Fibroblast | F0 SFRP2+ Fib | PBX1 | 0.938 | 0.432 | 0.219 | 2.18e-12 | 7.34e-08 |
| Fibroblast | F0 SFRP2+ Fib | ITSN1 | 0.937 | 0.215 | 0.08 | 1.27e-09 | 4.27e-05 |
| Fibroblast | F0 SFRP2+ Fib | EPS15 | 0.937 | 0.236 | 0.11 | 1.74e-07 | 0.00587 |
| Fibroblast | F0 SFRP2+ Fib | SRGAP1 | 0.935 | 0.233 | 0.104 | 2.54e-08 | 0.000857 |
| Fibroblast | F0 SFRP2+ Fib | CLN5 | 0.935 | 0.276 | 0.147 | 7.17e-07 | 0.0242 |
| Fibroblast | F0 SFRP2+ Fib | QPRT | 0.933 | 0.199 | 0.092 | 1.01e-06 | 0.034 |
| Fibroblast | F0 SFRP2+ Fib | RHOBTB3 | 0.93 | 0.517 | 0.279 | 5.08e-14 | 1.71e-09 |
| Fibroblast | F0 SFRP2+ Fib | NDRG1 | 0.93 | 0.385 | 0.172 | 1.32e-13 | 4.46e-09 |
| Fibroblast | F0 SFRP2+ Fib | PAM | 0.93 | 0.44 | 0.225 | 9.82e-13 | 3.31e-08 |
| Fibroblast | F0 SFRP2+ Fib | CHPT1 | 0.93 | 0.355 | 0.173 | 1.38e-10 | 4.66e-06 |
| Fibroblast | F0 SFRP2+ Fib | CYGB | 0.929 | 0.541 | 0.248 | 3.96e-18 | 1.34e-13 |
| Fibroblast | F0 SFRP2+ Fib | NPC2 | 0.927 | 0.923 | 0.658 | 5.81e-36 | 1.96e-31 |
| Fibroblast | F0 SFRP2+ Fib | CHIC2 | 0.926 | 0.361 | 0.181 | 3.81e-10 | 1.28e-05 |
| Fibroblast | F0 SFRP2+ Fib | BAIAP2 | 0.926 | 0.215 | 0.086 | 1.24e-08 | 0.000417 |
| Fibroblast | F0 SFRP2+ Fib | H6PD | 0.923 | 0.196 | 0.086 | 3.02e-07 | 0.0102 |
| Fibroblast | F0 SFRP2+ Fib | ADAM17 | 0.921 | 0.289 | 0.145 | 7.92e-08 | 0.00267 |
| Fibroblast | F0 SFRP2+ Fib | ZNF330 | 0.921 | 0.225 | 0.106 | 5.41e-07 | 0.0182 |
| Fibroblast | F0 SFRP2+ Fib | PROCR | 0.919 | 0.395 | 0.198 | 2.5e-11 | 8.43e-07 |
| Fibroblast | F0 SFRP2+ Fib | ITIH5 | 0.913 | 0.215 | 0.089 | 4.35e-08 | 0.00147 |
| Fibroblast | F0 SFRP2+ Fib | PSAP | 0.91 | 0.936 | 0.683 | 4.180000000000001e-37 | 1.41e-32 |
| Fibroblast | F0 SFRP2+ Fib | THY1 | 0.909 | 0.822 | 0.446 | 9.999999999999999e-27 | 3.38e-22 |
| Fibroblast | F0 SFRP2+ Fib | SNUPN | 0.897 | 0.215 | 0.09 | 4.01e-08 | 0.00135 |
| Fibroblast | F0 SFRP2+ Fib | PROS1 | 0.896 | 0.379 | 0.159 | 1.23e-13 | 4.15e-09 |
| Fibroblast | F0 SFRP2+ Fib | RP11-47A8.5 | 0.894 | 0.154 | 0.055 | 2.68e-07 | 0.00902 |
| Fibroblast | F0 SFRP2+ Fib | RNF130 | 0.892 | 0.363 | 0.184 | 5.58e-10 | 1.88e-05 |
| Fibroblast | F0 SFRP2+ Fib | FKBP11 | 0.887 | 0.48 | 0.234 | 1.2e-14 | 4.05e-10 |
| Fibroblast | F0 SFRP2+ Fib | CERCAM | 0.886 | 0.562 | 0.279 | 2.34e-17 | 7.88e-13 |
| Fibroblast | F0 SFRP2+ Fib | PRRX1 | 0.885 | 0.512 | 0.23 | 7.46e-17 | 2.51e-12 |
| Fibroblast | F0 SFRP2+ Fib | CUTC | 0.883 | 0.244 | 0.109 | 3.08e-08 | 0.00104 |
| Fibroblast | F0 SFRP2+ Fib | SCARB2 | 0.881 | 0.507 | 0.282 | 1.4e-12 | 4.72e-08 |
| Fibroblast | F0 SFRP2+ Fib | RERG | 0.881 | 0.395 | 0.185 | 8.92e-12 | 3.01e-07 |
| Fibroblast | F0 SFRP2+ Fib | THBS3 | 0.878 | 0.218 | 0.096 | 1.27e-07 | 0.00428 |
| Fibroblast | F0 SFRP2+ Fib | LTBP3 | 0.877 | 0.501 | 0.236 | 5.89e-16 | 1.99e-11 |
| Fibroblast | F0 SFRP2+ Fib | DPYSL2 | 0.876 | 0.422 | 0.194 | 1.42e-14 | 4.77e-10 |
| Fibroblast | F0 SFRP2+ Fib | UCHL1 | 0.876 | 0.154 | 0.054 | 1.01e-07 | 0.00342 |
| Fibroblast | F0 SFRP2+ Fib | RNF146 | 0.873 | 0.337 | 0.172 | 6.65e-09 | 0.000224 |
| Fibroblast | F0 SFRP2+ Fib | GLIPR1 | 0.87 | 0.443 | 0.227 | 7.12e-13 | 2.4e-08 |
| Fibroblast | F0 SFRP2+ Fib | SYNGR1 | 0.869 | 0.22 | 0.103 | 4.19e-07 | 0.0141 |
| Fibroblast | F0 SFRP2+ Fib | GOSR1 | 0.866 | 0.202 | 0.092 | 8.41e-07 | 0.0283 |
| Fibroblast | F0 SFRP2+ Fib | PRNP | 0.864 | 0.674 | 0.34 | 8.82e-21 | 2.97e-16 |
| Fibroblast | F0 SFRP2+ Fib | GALNT11 | 0.863 | 0.26 | 0.113 | 3.18e-09 | 0.000107 |
| Fibroblast | F0 SFRP2+ Fib | GLT8D1 | 0.862 | 0.347 | 0.176 | 4.23e-09 | 0.000143 |
| Fibroblast | F0 SFRP2+ Fib | DIMT1 | 0.858 | 0.252 | 0.126 | 4.87e-07 | 0.0164 |
| Fibroblast | F0 SFRP2+ Fib | SCARA5 | 0.857 | 0.196 | 0.064 | 5.58e-10 | 1.88e-05 |
| Fibroblast | F0 SFRP2+ Fib | FSTL1 | 0.855 | 0.804 | 0.469 | 8.969999999999999e-23 | 3.02e-18 |
| Fibroblast | F0 SFRP2+ Fib | PLPP1 | 0.855 | 0.562 | 0.279 | 1.13e-18 | 3.81e-14 |
| Fibroblast | F0 SFRP2+ Fib | BDH2 | 0.855 | 0.446 | 0.219 | 6.67e-13 | 2.25e-08 |
| Fibroblast | F0 SFRP2+ Fib | TFPI | 0.854 | 0.634 | 0.322 | 1.35e-19 | 4.54e-15 |
| Fibroblast | F0 SFRP2+ Fib | SMIM15 | 0.851 | 0.279 | 0.141 | 7.6e-08 | 0.00256 |
| Fibroblast | F0 SFRP2+ Fib | NUCB2 | 0.848 | 0.491 | 0.271 | 8e-12 | 2.69e-07 |

| compartment | subcluster | gene | avg_log2FC | pct.1 | pct.2 | p_val | p_val_adj |
|---|---|---|---|---|---|---|---|
| Fibroblast | F0 SFRP2+ Fib | RBMS3 | 0.848 | 0.347 | 0.158 | 2.53e-11 | 8.52e-07 |
| Fibroblast | F0 SFRP2+ Fib | LAMB1 | 0.848 | 0.342 | 0.17 | 1.13e-09 | 3.82e-05 |
| Fibroblast | F0 SFRP2+ Fib | HEXB | 0.842 | 0.618 | 0.326 | 3.15e-17 | 1.06e-12 |
| Fibroblast | F0 SFRP2+ Fib | HIBADH | 0.84 | 0.257 | 0.126 | 1.43e-07 | 0.00482 |
| Fibroblast | F0 SFRP2+ Fib | ZNF385A | 0.839 | 0.215 | 0.084 | 1.66e-08 | 0.00056 |
| Fibroblast | F0 SFRP2+ Fib | FMOD | 0.838 | 0.401 | 0.204 | 3.21e-11 | 1.08e-06 |
| Fibroblast | F0 SFRP2+ Fib | HTRA1 | 0.837 | 0.682 | 0.312 | 1.02e-25 | 3.44e-21 |
| Fibroblast | F0 SFRP2+ Fib | ZC2HC1A | 0.835 | 0.263 | 0.126 | 9.79e-08 | 0.0033 |
| Fibroblast | F0 SFRP2+ Fib | MXRA8 | 0.834 | 0.846 | 0.492 | 3.03e-25 | 1.02e-20 |
| Fibroblast | F0 SFRP2+ Fib | THYN1 | 0.834 | 0.361 | 0.204 | 3.38e-08 | 0.00114 |
| Fibroblast | F0 SFRP2+ Fib | SERPING1 | 0.829 | 0.958 | 0.735 | 7.67e-34 | 2.58e-29 |
| Fibroblast | F0 SFRP2+ Fib | OLFML2B | 0.829 | 0.369 | 0.194 | 5.08e-09 | 0.000171 |
| Fibroblast | F0 SFRP2+ Fib | LAPTM4A | 0.828 | 0.95 | 0.769 | 6.21e-33 | 2.09e-28 |
| Fibroblast | F0 SFRP2+ Fib | PHYHD1 | 0.828 | 0.233 | 0.09 | 1.21e-09 | 4.08e-05 |
| Fibroblast | F0 SFRP2+ Fib | FAM198B | 0.823 | 0.271 | 0.126 | 1.82e-08 | 0.000613 |
| Fibroblast | F0 SFRP2+ Fib | SPON2 | 0.822 | 0.796 | 0.383 | 2.88e-27 | 9.7e-23 |
| Fibroblast | F0 SFRP2+ Fib | PLSCR1 | 0.814 | 0.454 | 0.205 | 6.57e-15 | 2.21e-10 |
| Fibroblast | F0 SFRP2+ Fib | ANTXR2 | 0.813 | 0.313 | 0.149 | 1.37e-09 | 4.63e-05 |
| Fibroblast | F0 SFRP2+ Fib | TMBIM4 | 0.809 | 0.748 | 0.438 | 2.46e-20 | 8.27e-16 |
| Fibroblast | F0 SFRP2+ Fib | LIMA1 | 0.809 | 0.541 | 0.254 | 2.89e-17 | 9.73e-13 |
| Fibroblast | F0 SFRP2+ Fib | CDC26 | 0.809 | 0.438 | 0.224 | 8.89e-12 | 3e-07 |
| Fibroblast | F0 SFRP2+ Fib | TBL1XR1 | 0.809 | 0.305 | 0.15 | 1.17e-08 | 0.000394 |
| Fibroblast | F0 SFRP2+ Fib | VCAN | 0.807 | 0.735 | 0.343 | 9.44e-27 | 3.18e-22 |
| Fibroblast | F0 SFRP2+ Fib | CBLB | 0.802 | 0.212 | 0.092 | 1.83e-07 | 0.00615 |
| Fibroblast | F0 SFRP2+ Fib | TIMP2 | 0.8 | 0.782 | 0.435 | 1.26e-22 | 4.26e-18 |
| Fibroblast | F0 SFRP2+ Fib | MARC2 | 0.8 | 0.178 | 0.075 | 1.19e-06 | 0.0402 |
| Fibroblast | F0 SFRP2+ Fib | CKS1B | 0.799 | 0.347 | 0.144 | 6.2e-13 | 2.09e-08 |
| Fibroblast | F0 SFRP2+ Fib | P4HTM | 0.798 | 0.268 | 0.138 | 4.65e-07 | 0.0157 |
| Fibroblast | F0 SFRP2+ Fib | GPX8 | 0.797 | 0.443 | 0.21 | 7.43e-13 | 2.5e-08 |
| Fibroblast | F0 SFRP2+ Fib | NRP1 | 0.794 | 0.35 | 0.152 | 5.99e-12 | 2.02e-07 |
| Fibroblast | F0 SFRP2+ Fib | ACAA2 | 0.794 | 0.289 | 0.144 | 8.05e-08 | 0.00271 |
| Fibroblast | F0 SFRP2+ Fib | ECHDC2 | 0.792 | 0.294 | 0.144 | 4.44e-08 | 0.0015 |
| Fibroblast | F0 SFRP2+ Fib | REV3L | 0.789 | 0.233 | 0.107 | 1.85e-07 | 0.00623 |
| Fibroblast | F0 SFRP2+ Fib | IL6ST | 0.787 | 0.554 | 0.286 | 2.26e-17 | 7.61e-13 |
| Fibroblast | F0 SFRP2+ Fib | LTBP2 | 0.786 | 0.239 | 0.116 | 4.63e-07 | 0.0156 |
| Fibroblast | F0 SFRP2+ Fib | HSD17B4 | 0.784 | 0.188 | 0.08 | 5.01e-07 | 0.0169 |
| Fibroblast | F0 SFRP2+ Fib | WRB | 0.783 | 0.292 | 0.152 | 1.73e-07 | 0.00583 |
| Fibroblast | F0 SFRP2+ Fib | RHBDD2 | 0.779 | 0.358 | 0.211 | 4.55e-07 | 0.0153 |
| Fibroblast | F0 SFRP2+ Fib | COL1A2 | 0.778 | 0.968 | 0.658 | 4.14e-33 | 1.39e-28 |
| Fibroblast | F0 SFRP2+ Fib | COMP | 0.777 | 0.225 | 0.103 | 9.29e-08 | 0.00313 |
| Fibroblast | F0 SFRP2+ Fib | NUCB1 | 0.775 | 0.674 | 0.424 | 3.91e-15 | 1.32e-10 |
| Fibroblast | F0 SFRP2+ Fib | RARRES3 | 0.774 | 0.615 | 0.329 | 1.7e-15 | 5.74e-11 |
| Fibroblast | F0 SFRP2+ Fib | APOL2 | 0.772 | 0.207 | 0.09 | 3.67e-07 | 0.0124 |
| Fibroblast | F0 SFRP2+ Fib | AMZ2 | 0.767 | 0.196 | 0.089 | 1.26e-06 | 0.0425 |
| Fibroblast | F0 SFRP2+ Fib | TMEM43 | 0.762 | 0.371 | 0.19 | 9.85e-10 | 3.32e-05 |
| Fibroblast | F0 SFRP2+ Fib | F3 | 0.762 | 0.297 | 0.141 | 3.8e-09 | 0.000128 |
| Fibroblast | F0 SFRP2+ Fib | C14orf1 | 0.762 | 0.265 | 0.135 | 5.29e-07 | 0.0178 |
| Fibroblast | F0 SFRP2+ Fib | RPA2 | 0.76 | 0.284 | 0.144 | 1.67e-07 | 0.00564 |
| Fibroblast | F0 SFRP2+ Fib | TPP1 | 0.755 | 0.329 | 0.167 | 1.18e-08 | 0.000399 |
| Fibroblast | F0 SFRP2+ Fib | SLC43A3 | 0.755 | 0.212 | 0.092 | 1.64e-07 | 0.00553 |
| Fibroblast | F0 SFRP2+ Fib | DAAM1 | 0.754 | 0.353 | 0.164 | 1.37e-10 | 4.6e-06 |
| Fibroblast | F0 SFRP2+ Fib | COL8A1 | 0.753 | 0.44 | 0.19 | 6.18e-15 | 2.08e-10 |
| Fibroblast | F0 SFRP2+ Fib | FAM3C | 0.752 | 0.599 | 0.332 | 2.94e-14 | 9.91e-10 |
| Fibroblast | F0 SFRP2+ Fib | IL10RB | 0.752 | 0.316 | 0.165 | 5.9e-08 | 0.00199 |
| Fibroblast | F0 SFRP2+ Fib | SMIM19 | 0.75 | 0.451 | 0.225 | 3.88e-12 | 1.31e-07 |
| Fibroblast | F0 SFRP2+ Fib | AZI2 | 0.745 | 0.406 | 0.188 | 1.06e-12 | 3.57e-08 |
| Fibroblast | F0 SFRP2+ Fib | DERL1 | 0.745 | 0.329 | 0.165 | 4.47e-09 | 0.000151 |
| Fibroblast | F0 SFRP2+ Fib | CRABP2 | 0.744 | 0.305 | 0.115 | 3.24e-13 | 1.09e-08 |
| Fibroblast | F0 SFRP2+ Fib | TMEM204 | 0.736 | 0.446 | 0.27 | 3.97e-08 | 0.00134 |
| Fibroblast | F0 SFRP2+ Fib | LAPTM4B | 0.727 | 0.297 | 0.158 | 2.76e-07 | 0.00929 |
| Fibroblast | F0 SFRP2+ Fib | DPP7 | 0.725 | 0.581 | 0.308 | 1.35e-14 | 4.53e-10 |
| Fibroblast | F0 SFRP2+ Fib | LHFP | 0.724 | 0.785 | 0.516 | 2.37e-16 | 8e-12 |
| Fibroblast | F0 SFRP2+ Fib | SLC44A1 | 0.72 | 0.353 | 0.184 | 5.32e-09 | 0.000179 |
| Fibroblast | F0 SFRP2+ Fib | PIGT | 0.718 | 0.493 | 0.274 | 1.4e-11 | 4.7e-07 |

| compartment | subcluster | gene | avg_log2FC | pct.1 | pct.2 | p_val | p_val_adj |
|---|---|---|---|---|---|---|---|
| Fibroblast | F0 SFRP2+ Fib | EBF2 | 0.718 | 0.111 | 0.026 | 2.87e-08 | 0.000967 |
| Fibroblast | F0 SFRP2+ Fib | COL6A2 | 0.717 | 0.976 | 0.809 | 3.42e-31 | 1.15e-26 |
| Fibroblast | F0 SFRP2+ Fib | AHNAK | 0.707 | 0.615 | 0.372 | 4.9e-12 | 1.65e-07 |
| Fibroblast | F0 SFRP2+ Fib | INSIG1 | 0.705 | 0.332 | 0.139 | 1.97e-11 | 6.65e-07 |
| Fibroblast | F0 SFRP2+ Fib | PGRMC2 | 0.704 | 0.435 | 0.234 | 3.56e-10 | 1.2e-05 |
| Fibroblast | F0 SFRP2+ Fib | NUPR1 | 0.701 | 0.947 | 0.753 | 5.690000000000001e-21 | 1.92e-16 |
| Fibroblast | F0 SFRP2+ Fib | TCEA3 | 0.701 | 0.263 | 0.13 | 1.87e-07 | 0.00629 |
| Fibroblast | F0 SFRP2+ Fib | SRPRB | 0.701 | 0.257 | 0.13 | 7.66e-07 | 0.0258 |
| Fibroblast | F0 SFRP2+ Fib | TMEM9 | 0.7 | 0.374 | 0.207 | 3.13e-08 | 0.00105 |
| Fibroblast | F0 SFRP2+ Fib | TTC3 | 0.698 | 0.541 | 0.323 | 1.74e-10 | 5.87e-06 |
| Fibroblast | F0 SFRP2+ Fib | SRP72 | 0.689 | 0.427 | 0.251 | 1.1e-07 | 0.0037 |
| Fibroblast | F0 SFRP2+ Fib | HTRA3 | 0.688 | 0.475 | 0.217 | 2.15e-15 | 7.23e-11 |
| Fibroblast | F0 SFRP2+ Fib | GOLGA7 | 0.687 | 0.342 | 0.201 | 1.24e-06 | 0.0418 |
| Fibroblast | F0 SFRP2+ Fib | CLEC2B | 0.685 | 0.647 | 0.337 | 9.9e-17 | 3.34e-12 |
| Fibroblast | F0 SFRP2+ Fib | ATF1 | 0.684 | 0.284 | 0.135 | 2.75e-08 | 0.000928 |
| Fibroblast | F0 SFRP2+ Fib | DNPH1 | 0.683 | 0.432 | 0.233 | 2.07e-10 | 6.98e-06 |
| Fibroblast | F0 SFRP2+ Fib | KDM6B | 0.683 | 0.353 | 0.191 | 5.42e-08 | 0.00183 |
| Fibroblast | F0 SFRP2+ Fib | TCF7L2 | 0.683 | 0.257 | 0.132 | 9.14e-07 | 0.0308 |
| Fibroblast | F0 SFRP2+ Fib | TRAM1 | 0.676 | 0.565 | 0.329 | 2.21e-11 | 7.45e-07 |
| Fibroblast | F0 SFRP2+ Fib | BORCS7 | 0.676 | 0.427 | 0.253 | 2.82e-08 | 0.000952 |
| Fibroblast | F0 SFRP2+ Fib | HEXA | 0.673 | 0.448 | 0.217 | 4.97e-13 | 1.67e-08 |
| Fibroblast | F0 SFRP2+ Fib | WEE1 | 0.672 | 0.268 | 0.118 | 8.24e-09 | 0.000278 |
| Fibroblast | F0 SFRP2+ Fib | TMED3 | 0.671 | 0.491 | 0.279 | 3.8e-10 | 1.28e-05 |
| Fibroblast | F0 SFRP2+ Fib | CXCL2 | 0.669 | 0.552 | 0.323 | 1.88e-12 | 6.33e-08 |
| Fibroblast | F0 SFRP2+ Fib | CTSO | 0.669 | 0.366 | 0.196 | 8.72e-09 | 0.000294 |
| Fibroblast | F0 SFRP2+ Fib | APOL1 | 0.668 | 0.353 | 0.211 | 1.25e-06 | 0.0422 |
| Fibroblast | F0 SFRP2+ Fib | EPB41L2 | 0.667 | 0.44 | 0.237 | 6.11e-10 | 2.06e-05 |
| Fibroblast | F0 SFRP2+ Fib | ITM2B | 0.662 | 0.971 | 0.867 | 3.78e-26 | 1.27e-21 |
| Fibroblast | F0 SFRP2+ Fib | CASP4 | 0.661 | 0.353 | 0.19 | 4.16e-08 | 0.0014 |
| Fibroblast | F0 SFRP2+ Fib | SPTBN1 | 0.658 | 0.48 | 0.279 | 1.49e-09 | 5.02e-05 |
| Fibroblast | F0 SFRP2+ Fib | SULF2 | 0.653 | 0.371 | 0.204 | 1.27e-08 | 0.000429 |
| Fibroblast | F0 SFRP2+ Fib | FRMD6 | 0.651 | 0.21 | 0.098 | 8.46e-07 | 0.0285 |
| Fibroblast | F0 SFRP2+ Fib | ANAPC13 | 0.647 | 0.387 | 0.211 | 1.1e-08 | 0.000371 |
| Fibroblast | F0 SFRP2+ Fib | TMEM222 | 0.646 | 0.265 | 0.127 | 1.64e-07 | 0.00554 |
| Fibroblast | F0 SFRP2+ Fib | FGFR1 | 0.645 | 0.501 | 0.282 | 1.27e-11 | 4.28e-07 |
| Fibroblast | F0 SFRP2+ Fib | TNPO1 | 0.643 | 0.255 | 0.127 | 8.96e-07 | 0.0302 |
| Fibroblast | F0 SFRP2+ Fib | GPR108 | 0.642 | 0.257 | 0.126 | 5.19e-07 | 0.0175 |
| Fibroblast | F0 SFRP2+ Fib | ARL6IP5 | 0.639 | 0.793 | 0.518 | 1.38e-16 | 4.66e-12 |
| Fibroblast | F0 SFRP2+ Fib | PPIC | 0.636 | 0.751 | 0.447 | 1.08e-15 | 3.64e-11 |
| Fibroblast | F0 SFRP2+ Fib | UNC50 | 0.636 | 0.353 | 0.185 | 3.17e-08 | 0.00107 |
| Fibroblast | F0 SFRP2+ Fib | TUBB2A | 0.636 | 0.411 | 0.256 | 1.08e-06 | 0.0364 |
| Fibroblast | F0 SFRP2+ Fib | SEC63 | 0.635 | 0.488 | 0.273 | 2.16e-10 | 7.28e-06 |
| Fibroblast | F0 SFRP2+ Fib | HBP1 | 0.635 | 0.414 | 0.233 | 2.33e-08 | 0.000786 |
| Fibroblast | F0 SFRP2+ Fib | LAMP2 | 0.633 | 0.493 | 0.26 | 3.66e-12 | 1.23e-07 |
| Fibroblast | F0 SFRP2+ Fib | SDCBP | 0.632 | 0.708 | 0.424 | 5.36e-15 | 1.81e-10 |
| Fibroblast | F0 SFRP2+ Fib | NDN | 0.63 | 0.581 | 0.357 | 2.38e-10 | 8.02e-06 |
| Fibroblast | F0 SFRP2+ Fib | CYR61 | 0.628 | 0.952 | 0.767 | 4.32e-12 | 1.46e-07 |
| Fibroblast | F0 SFRP2+ Fib | TMEM176A | 0.625 | 0.684 | 0.34 | 3.32e-20 | 1.12e-15 |
| Fibroblast | F0 SFRP2+ Fib | TMEM176B | 0.623 | 0.764 | 0.395 | 1.2e-22 | 4.03e-18 |
| Fibroblast | F0 SFRP2+ Fib | ATP6V0E1 | 0.623 | 0.905 | 0.616 | 3.82e-21 | 1.29e-16 |
| Fibroblast | F0 SFRP2+ Fib | ACP5 | 0.618 | 0.196 | 0.086 | 6.65e-07 | 0.0224 |
| Fibroblast | F0 SFRP2+ Fib | DDAH2 | 0.617 | 0.658 | 0.429 | 1.2e-10 | 4.04e-06 |
| Fibroblast | F0 SFRP2+ Fib | CD63 | 0.615 | 0.989 | 0.91 | 2.04e-31 | 6.88e-27 |
| Fibroblast | F0 SFRP2+ Fib | CDK5RAP3 | 0.614 | 0.305 | 0.156 | 7.53e-08 | 0.00254 |
| Fibroblast | F0 SFRP2+ Fib | BAG2 | 0.609 | 0.302 | 0.165 | 1.35e-06 | 0.0454 |
| Fibroblast | F0 SFRP2+ Fib | LATS2 | 0.608 | 0.191 | 0.083 | 9.21e-07 | 0.031 |
| Fibroblast | F0 SFRP2+ Fib | LY6E | 0.603 | 0.873 | 0.564 | 2.26e-20 | 7.63e-16 |
| Fibroblast | F0 SFRP2+ Fib | ENG | 0.6 | 0.414 | 0.221 | 1.39e-09 | 4.69e-05 |
| Fibroblast | F0 SFRP2+ Fib | ATRAID | 0.598 | 0.793 | 0.502 | 6.8e-15 | 2.29e-10 |
| Fibroblast | F0 SFRP2+ Fib | SMIM14 | 0.593 | 0.454 | 0.237 | 4.16e-11 | 1.4e-06 |
| Fibroblast | F0 SFRP2+ Fib | RNF13 | 0.593 | 0.353 | 0.176 | 4.3e-09 | 0.000145 |
| Fibroblast | F0 SFRP2+ Fib | RPS4Y1 | 0.592 | 0.655 | 0.413 | 2.8e-11 | 9.43e-07 |
| Fibroblast | F0 SFRP2+ Fib | PPCS | 0.59 | 0.35 | 0.198 | 7.29e-07 | 0.0246 |
| Fibroblast | F0 SFRP2+ Fib | PLAGL1 | 0.587 | 0.249 | 0.124 | 1.02e-06 | 0.0343 |

| compartment | subcluster | gene | avg_log2FC | pct.1 | pct.2 | p_val | p_val_adj |
|---|---|---|---|---|---|---|---|
| Fibroblast | F0 SFRP2+ Fib | IFI35 | 0.585 | 0.347 | 0.198 | 2.45e-07 | 0.00826 |
| Fibroblast | F0 SFRP2+ Fib | LRP10 | 0.584 | 0.491 | 0.286 | 2.34e-09 | 7.87e-05 |
| Fibroblast | F0 SFRP2+ Fib | MESDC2 | 0.58 | 0.401 | 0.221 | 9.5e-09 | 0.00032 |
| Fibroblast | F0 SFRP2+ Fib | ANKRD28 | 0.579 | 0.382 | 0.231 | 8.93e-07 | 0.0301 |
| Fibroblast | F0 SFRP2+ Fib | GNG11 | 0.573 | 0.711 | 0.423 | 3.18e-16 | 1.07e-11 |
| Fibroblast | F0 SFRP2+ Fib | REXO2 | 0.572 | 0.65 | 0.404 | 9.82e-12 | 3.31e-07 |
| Fibroblast | F0 SFRP2+ Fib | COL6A1 | 0.571 | 0.883 | 0.62 | 4.52e-16 | 1.52e-11 |
| Fibroblast | F0 SFRP2+ Fib | COL3A1 | 0.568 | 0.883 | 0.606 | 3.29e-16 | 1.11e-11 |
| Fibroblast | F0 SFRP2+ Fib | SSR2 | 0.568 | 0.889 | 0.625 | 6.54e-16 | 2.2e-11 |
| Fibroblast | F0 SFRP2+ Fib | IFI16 | 0.568 | 0.74 | 0.462 | 2.19e-14 | 7.38e-10 |
| Fibroblast | F0 SFRP2+ Fib | SNAI2 | 0.567 | 0.371 | 0.199 | 2.58e-08 | 0.000869 |
| Fibroblast | F0 SFRP2+ Fib | C19orf60 | 0.565 | 0.642 | 0.392 | 1.03e-11 | 3.48e-07 |
| Fibroblast | F0 SFRP2+ Fib | FSTL3 | 0.564 | 0.321 | 0.17 | 2.01e-07 | 0.00678 |
| Fibroblast | F0 SFRP2+ Fib | CEBPZOS | 0.564 | 0.241 | 0.115 | 4.57e-07 | 0.0154 |
| Fibroblast | F0 SFRP2+ Fib | TNFRSF1A | 0.562 | 0.597 | 0.361 | 2.06e-10 | 6.93e-06 |
| Fibroblast | F0 SFRP2+ Fib | C6orf62 | 0.562 | 0.406 | 0.239 | 1.49e-07 | 0.00503 |
| Fibroblast | F0 SFRP2+ Fib | SEP15 | 0.56 | 0.753 | 0.482 | 2.2e-13 | 7.41e-09 |
| Fibroblast | F0 SFRP2+ Fib | SDF4 | 0.558 | 0.69 | 0.429 | 2.52e-11 | 8.49e-07 |
| Fibroblast | F0 SFRP2+ Fib | NXT1 | 0.557 | 0.382 | 0.21 | 2.97e-08 | 0.001 |
| Fibroblast | F0 SFRP2+ Fib | STX4 | 0.555 | 0.3 | 0.152 | 1.13e-07 | 0.0038 |
| Fibroblast | F0 SFRP2+ Fib | UROD | 0.554 | 0.347 | 0.187 | 1.57e-07 | 0.00531 |
| Fibroblast | F0 SFRP2+ Fib | ANXA2 | 0.553 | 0.902 | 0.666 | 1.11e-16 | 3.75e-12 |
| Fibroblast | F0 SFRP2+ Fib | APH1A | 0.553 | 0.462 | 0.262 | 1.8e-09 | 6.07e-05 |
| Fibroblast | F0 SFRP2+ Fib | BLVRB | 0.552 | 0.462 | 0.245 | 1.16e-09 | 3.9e-05 |
| Fibroblast | F0 SFRP2+ Fib | GINM1 | 0.549 | 0.477 | 0.294 | 1.19e-07 | 0.00401 |
| Fibroblast | F0 SFRP2+ Fib | CSTB | 0.546 | 0.804 | 0.521 | 8.89e-16 | 3e-11 |
| Fibroblast | F0 SFRP2+ Fib | RTN4 | 0.546 | 0.857 | 0.657 | 6.03e-13 | 2.03e-08 |
| Fibroblast | F0 SFRP2+ Fib | ARRDC3 | 0.546 | 0.385 | 0.196 | 3.3e-09 | 0.000111 |
| Fibroblast | F0 SFRP2+ Fib | ERLEC1 | 0.545 | 0.448 | 0.257 | 1.42e-08 | 0.000477 |
| Fibroblast | F0 SFRP2+ Fib | NEXN | 0.544 | 0.488 | 0.296 | 5.58e-08 | 0.00188 |
| Fibroblast | F0 SFRP2+ Fib | TMEM126B | 0.544 | 0.385 | 0.228 | 5.52e-07 | 0.0186 |
| Fibroblast | F0 SFRP2+ Fib | APOE | 0.542 | 0.682 | 0.469 | 1.03e-10 | 3.47e-06 |
| Fibroblast | F0 SFRP2+ Fib | KCNQ1OT1 | 0.541 | 0.597 | 0.386 | 9.84e-09 | 0.000331 |
| Fibroblast | F0 SFRP2+ Fib | ALDH1A3 | 0.538 | 0.26 | 0.123 | 1.96e-07 | 0.00662 |
| Fibroblast | F0 SFRP2+ Fib | FUNDC2 | 0.537 | 0.544 | 0.329 | 9.4e-10 | 3.17e-05 |
| Fibroblast | F0 SFRP2+ Fib | DDOST | 0.53 | 0.427 | 0.25 | 5.9e-08 | 0.00199 |
| Fibroblast | F0 SFRP2+ Fib | TCF21 | 0.529 | 0.43 | 0.219 | 4.26e-11 | 1.44e-06 |
| Fibroblast | F0 SFRP2+ Fib | RTN3 | 0.527 | 0.44 | 0.26 | 4.92e-08 | 0.00166 |
| Fibroblast | F0 SFRP2+ Fib | SPG21 | 0.525 | 0.454 | 0.273 | 5.02e-08 | 0.00169 |
| Fibroblast | F0 SFRP2+ Fib | FOXO3 | 0.524 | 0.533 | 0.299 | 2.2e-11 | 7.42e-07 |
| Fibroblast | F0 SFRP2+ Fib | GYPC | 0.523 | 0.573 | 0.34 | 2.27e-10 | 7.64e-06 |
| Fibroblast | F0 SFRP2+ Fib | HEBP1 | 0.523 | 0.462 | 0.273 | 2.24e-07 | 0.00753 |
| Fibroblast | F0 SFRP2+ Fib | PRDX2 | 0.521 | 0.706 | 0.44 | 2.24e-11 | 7.56e-07 |
| Fibroblast | F0 SFRP2+ Fib | C1orf54 | 0.521 | 0.326 | 0.165 | 3.34e-08 | 0.00113 |
| Fibroblast | F0 SFRP2+ Fib | SFT2D1 | 0.518 | 0.43 | 0.25 | 3.25e-08 | 0.00109 |
| Fibroblast | F0 SFRP2+ Fib | LMAN2 | 0.515 | 0.647 | 0.409 | 2.98e-10 | 1.01e-05 |
| Fibroblast | F0 SFRP2+ Fib | LAMP1 | 0.515 | 0.676 | 0.447 | 5.88e-10 | 1.98e-05 |
| Fibroblast | F0 SFRP2+ Fib | PCBD1 | 0.515 | 0.419 | 0.247 | 1.61e-07 | 0.00542 |
| Fibroblast | F0 SFRP2+ Fib | EFEMP2 | 0.514 | 0.711 | 0.429 | 6.52e-12 | 2.2e-07 |
| Fibroblast | F0 SFRP2+ Fib | TMEM179B | 0.51 | 0.48 | 0.283 | 7.92e-09 | 0.000267 |
| Fibroblast | F0 SFRP2+ Fib | CCDC53 | 0.51 | 0.305 | 0.165 | 8.21e-07 | 0.0277 |
| Fibroblast | F0 SFRP2+ Fib | GSTM3 | 0.509 | 0.528 | 0.305 | 1.66e-09 | 5.58e-05 |
| Fibroblast | F0 SFRP2+ Fib | RPLP0 | 0.507 | 0.995 | 0.916 | 1.63e-22 | 5.48e-18 |
| Fibroblast | F0 SFRP2+ Fib | PDPN | 0.505 | 0.406 | 0.222 | 5.68e-08 | 0.00192 |
| Fibroblast | F0 SFRP2+ Fib | ERGIC1 | 0.505 | 0.292 | 0.156 | 9.25e-07 | 0.0312 |
| Fibroblast | F0 SFRP2+ Fib | ECHS1 | 0.503 | 0.456 | 0.268 | 7.91e-08 | 0.00267 |
| Fibroblast | F0 SFRP2+ Fib | CHPF | 0.5 | 0.403 | 0.239 | 9.33e-07 | 0.0314 |
| Fibroblast | F0 SFRP2+ Fib | ABHD14B | 0.497 | 0.385 | 0.227 | 3.71e-07 | 0.0125 |
| Fibroblast | F0 SFRP2+ Fib | CCS | 0.496 | 0.371 | 0.217 | 9.64e-07 | 0.0325 |
| Fibroblast | F0 SFRP2+ Fib | BEX4 | 0.494 | 0.347 | 0.185 | 1.57e-07 | 0.00528 |
| Fibroblast | F0 SFRP2+ Fib | RBMS1 | 0.493 | 0.541 | 0.328 | 1.84e-09 | 6.21e-05 |
| Fibroblast | F0 SFRP2+ Fib | PLAC9 | 0.488 | 0.737 | 0.571 | 2.52e-08 | 0.000848 |
| Fibroblast | F0 SFRP2+ Fib | NFKBIA | 0.486 | 0.973 | 0.818 | 3.74e-16 | 1.26e-11 |
| Fibroblast | F0 SFRP2+ Fib | CNN3 | 0.486 | 0.846 | 0.636 | 2.68e-09 | 9.04e-05 |

| compartment | subcluster | gene | avg_log2FC | pct.1 | pct.2 | p_val | p_val_adj |
|---|---|---|---|---|---|---|---|
| Fibroblast | F0 SFRP2+ Fib | TIMM9 | 0.486 | 0.363 | 0.196 | 3.72e-08 | 0.00125 |
| Fibroblast | F0 SFRP2+ Fib | PRDX4 | 0.483 | 0.599 | 0.348 | 1.15e-10 | 3.86e-06 |
| Fibroblast | F0 SFRP2+ Fib | NGFRAP1 | 0.483 | 0.836 | 0.6 | 2.03e-10 | 6.84e-06 |
| Fibroblast | F0 SFRP2+ Fib | PDLIM4 | 0.483 | 0.401 | 0.21 | 1.53e-09 | 5.15e-05 |
| Fibroblast | F0 SFRP2+ Fib | LMO4 | 0.483 | 0.592 | 0.387 | 1.38e-07 | 0.00464 |
| Fibroblast | F0 SFRP2+ Fib | FTL | 0.481 | 1.0 | 0.995 | 8.94e-33 | 3.01e-28 |
| Fibroblast | F0 SFRP2+ Fib | SQSTM1 | 0.481 | 0.796 | 0.556 | 1.06e-11 | 3.58e-07 |
| Fibroblast | F0 SFRP2+ Fib | OCIAD1 | 0.479 | 0.57 | 0.337 | 2.23e-09 | 7.51e-05 |
| Fibroblast | F0 SFRP2+ Fib | ARID5B | 0.478 | 0.851 | 0.616 | 1.5e-10 | 5.06e-06 |
| Fibroblast | F0 SFRP2+ Fib | RARRES2 | 0.473 | 0.971 | 0.668 | 5.18e-20 | 1.75e-15 |
| Fibroblast | F0 SFRP2+ Fib | ECH1 | 0.473 | 0.61 | 0.4 | 5.64e-08 | 0.0019 |
| Fibroblast | F0 SFRP2+ Fib | PHLDA3 | 0.471 | 0.61 | 0.415 | 1.5e-07 | 0.00507 |
| Fibroblast | F0 SFRP2+ Fib | TGIF1 | 0.471 | 0.432 | 0.26 | 3.19e-07 | 0.0107 |
| Fibroblast | F0 SFRP2+ Fib | THBS1 | 0.47 | 0.525 | 0.331 | 5.3e-08 | 0.00179 |
| Fibroblast | F0 SFRP2+ Fib | CDIPT | 0.468 | 0.477 | 0.294 | 6e-08 | 0.00202 |
| Fibroblast | F0 SFRP2+ Fib | RRBP1 | 0.468 | 0.602 | 0.397 | 7.46e-08 | 0.00251 |
| Fibroblast | F0 SFRP2+ Fib | NTAN1 | 0.468 | 0.337 | 0.19 | 1.18e-06 | 0.0397 |
| Fibroblast | F0 SFRP2+ Fib | NENF | 0.467 | 0.851 | 0.622 | 1.88e-11 | 6.32e-07 |
| Fibroblast | F0 SFRP2+ Fib | CXCL14 | 0.466 | 0.568 | 0.219 | 1.17e-25 | 3.94e-21 |
| Fibroblast | F0 SFRP2+ Fib | UGP2 | 0.463 | 0.541 | 0.346 | 2.99e-07 | 0.0101 |
| Fibroblast | F0 SFRP2+ Fib | GNG12 | 0.462 | 0.353 | 0.196 | 6.24e-07 | 0.021 |
| Fibroblast | F0 SFRP2+ Fib | PQLC3 | 0.458 | 0.419 | 0.228 | 3.15e-08 | 0.00106 |
| Fibroblast | F0 SFRP2+ Fib | BLOC1S2 | 0.457 | 0.289 | 0.15 | 3.96e-07 | 0.0133 |
| Fibroblast | F0 SFRP2+ Fib | COPZ2 | 0.456 | 0.31 | 0.165 | 3.92e-07 | 0.0132 |
| Fibroblast | F0 SFRP2+ Fib | SPAG7 | 0.455 | 0.416 | 0.248 | 6.27e-07 | 0.0211 |
| Fibroblast | F0 SFRP2+ Fib | SRI | 0.453 | 0.605 | 0.401 | 4.92e-08 | 0.00166 |
| Fibroblast | F0 SFRP2+ Fib | SEPT11 | 0.452 | 0.536 | 0.32 | 1.7e-08 | 0.000574 |
| Fibroblast | F0 SFRP2+ Fib | MTCH1 | 0.451 | 0.653 | 0.438 | 4.26e-08 | 0.00144 |
| Fibroblast | F0 SFRP2+ Fib | SLC2A3 | 0.449 | 0.597 | 0.36 | 5.38e-11 | 1.81e-06 |
| Fibroblast | F0 SFRP2+ Fib | DNASE2 | 0.449 | 0.366 | 0.205 | 2.92e-07 | 0.00983 |
| Fibroblast | F0 SFRP2+ Fib | EMILIN1 | 0.448 | 0.716 | 0.489 | 6.66e-09 | 0.000225 |
| Fibroblast | F0 SFRP2+ Fib | CTSC | 0.447 | 0.605 | 0.355 | 3.94e-13 | 1.33e-08 |
| Fibroblast | F0 SFRP2+ Fib | FAM162A | 0.447 | 0.515 | 0.303 | 5.68e-09 | 0.000192 |
| Fibroblast | F0 SFRP2+ Fib | NBL1 | 0.446 | 0.849 | 0.536 | 1.39e-15 | 4.68e-11 |
| Fibroblast | F0 SFRP2+ Fib | TMED10 | 0.445 | 0.66 | 0.412 | 1.51e-09 | 5.09e-05 |
| Fibroblast | F0 SFRP2+ Fib | G3BP1 | 0.442 | 0.342 | 0.188 | 2.44e-07 | 0.00823 |
| Fibroblast | F0 SFRP2+ Fib | ADH5 | 0.442 | 0.485 | 0.309 | 9.62e-07 | 0.0324 |
| Fibroblast | F0 SFRP2+ Fib | CD99 | 0.435 | 0.944 | 0.721 | 1e-15 | 3.37e-11 |
| Fibroblast | F0 SFRP2+ Fib | NAMPT | 0.435 | 0.491 | 0.306 | 1.77e-07 | 0.00596 |
| Fibroblast | F0 SFRP2+ Fib | TOR1AIP1 | 0.433 | 0.265 | 0.121 | 6.79e-08 | 0.00229 |
| Fibroblast | F0 SFRP2+ Fib | CTSL | 0.433 | 0.568 | 0.383 | 1.13e-06 | 0.0381 |
| Fibroblast | F0 SFRP2+ Fib | PPIB | 0.432 | 0.907 | 0.691 | 1.31e-12 | 4.42e-08 |
| Fibroblast | F0 SFRP2+ Fib | APP | 0.43 | 0.753 | 0.536 | 1.06e-07 | 0.00358 |
| Fibroblast | F0 SFRP2+ Fib | CD248 | 0.43 | 0.302 | 0.165 | 7.98e-07 | 0.0269 |
| Fibroblast | F0 SFRP2+ Fib | RAB34 | 0.425 | 0.66 | 0.447 | 1.13e-07 | 0.0038 |
| Fibroblast | F0 SFRP2+ Fib | ASPN | 0.423 | 0.592 | 0.332 | 1.84e-11 | 6.19e-07 |
| Fibroblast | F0 SFRP2+ Fib | TAX1BP3 | 0.423 | 0.645 | 0.449 | 2.13e-07 | 0.00717 |
| Fibroblast | F0 SFRP2+ Fib | GRN | 0.423 | 0.584 | 0.395 | 3.7e-07 | 0.0125 |
| Fibroblast | F0 SFRP2+ Fib | BZW1 | 0.418 | 0.552 | 0.337 | 1.84e-08 | 0.00062 |
| Fibroblast | F0 SFRP2+ Fib | NEU1 | 0.415 | 0.668 | 0.421 | 4.38e-11 | 1.48e-06 |
| Fibroblast | F0 SFRP2+ Fib | PRDX5 | 0.415 | 0.838 | 0.591 | 1.1e-09 | 3.69e-05 |
| Fibroblast | F0 SFRP2+ Fib | IER3 | 0.413 | 0.814 | 0.665 | 1.19e-07 | 0.004 |
| Fibroblast | F0 SFRP2+ Fib | KLF4 | 0.408 | 0.849 | 0.611 | 3.17e-15 | 1.07e-10 |
| Fibroblast | F0 SFRP2+ Fib | RAB7A | 0.406 | 0.698 | 0.467 | 7.46e-09 | 0.000251 |
| Fibroblast | F0 SFRP2+ Fib | SEC62 | 0.405 | 0.817 | 0.585 | 3.82e-09 | 0.000129 |
| Fibroblast | F0 SFRP2+ Fib | NFE2L2 | 0.405 | 0.578 | 0.372 | 1.46e-07 | 0.00492 |
| Fibroblast | F0 SFRP2+ Fib | AKAP13 | 0.404 | 0.568 | 0.357 | 4.19e-08 | 0.00141 |
| Fibroblast | F0 SFRP2+ Fib | CRTAP | 0.404 | 0.631 | 0.427 | 4.23e-08 | 0.00142 |
| Fibroblast | F0 SFRP2+ Fib | FXYD1 | 0.401 | 0.676 | 0.418 | 5.81e-09 | 0.000196 |
| Fibroblast | F0 SFRP2+ Fib | SNHG8 | 0.4 | 0.706 | 0.456 | 1.21e-09 | 4.08e-05 |
| Fibroblast | F0 SFRP2+ Fib | SSBP2 | 0.398 | 0.318 | 0.165 | 1.22e-06 | 0.0412 |
| Fibroblast | F0 SFRP2+ Fib | SEC11A | 0.394 | 0.775 | 0.508 | 6.36e-11 | 2.14e-06 |
| Fibroblast | F0 SFRP2+ Fib | DDIT3 | 0.393 | 0.509 | 0.322 | 1.21e-06 | 0.0408 |
| Fibroblast | F0 SFRP2+ Fib | AKAP12 | 0.383 | 0.507 | 0.311 | 6.08e-08 | 0.00205 |

| compartment | subcluster | gene | avg_log2FC | pct.1 | pct.2 | p_val | p_val_adj |
|---|---|---|---|---|---|---|---|
| Fibroblast | F0 SFRP2+ Fib | ZFP36L2 | 0.38 | 0.878 | 0.704 | 1.06e-09 | 3.57e-05 |
| Fibroblast | F0 SFRP2+ Fib | WDR83OS | 0.379 | 0.817 | 0.568 | 2.02e-08 | 0.000682 |
| Fibroblast | F0 SFRP2+ Fib | RPL36 | 0.378 | 0.997 | 0.968 | 3.07e-20 | 1.03e-15 |
| Fibroblast | F0 SFRP2+ Fib | FHL2 | 0.376 | 0.507 | 0.302 | 1.75e-08 | 0.000589 |
| Fibroblast | F0 SFRP2+ Fib | MYDGF | 0.375 | 0.698 | 0.418 | 2.25e-10 | 7.6e-06 |
| Fibroblast | F0 SFRP2+ Fib | C6orf48 | 0.375 | 0.814 | 0.614 | 1.62e-08 | 0.000544 |
| Fibroblast | F0 SFRP2+ Fib | CD81 | 0.375 | 0.584 | 0.374 | 2.92e-08 | 0.000984 |
| Fibroblast | F0 SFRP2+ Fib | COL1A1 | 0.369 | 0.931 | 0.735 | 2.27e-15 | 7.64e-11 |
| Fibroblast | F0 SFRP2+ Fib | GABARAPL2 | 0.369 | 0.714 | 0.524 | 5.09e-07 | 0.0172 |
| Fibroblast | F0 SFRP2+ Fib | METTL9 | 0.366 | 0.541 | 0.34 | 2.93e-07 | 0.00986 |
| Fibroblast | F0 SFRP2+ Fib | WSB1 | 0.364 | 0.793 | 0.568 | 1.58e-08 | 0.000532 |
| Fibroblast | F0 SFRP2+ Fib | SSR4 | 0.362 | 0.886 | 0.662 | 8.34e-10 | 2.81e-05 |
| Fibroblast | F0 SFRP2+ Fib | IFI27L2 | 0.361 | 0.721 | 0.493 | 1.02e-07 | 0.00343 |
| Fibroblast | F0 SFRP2+ Fib | TMBIM6 | 0.36 | 0.806 | 0.609 | 1.78e-07 | 0.006 |
| Fibroblast | F0 SFRP2+ Fib | CUTA | 0.359 | 0.769 | 0.539 | 1.3e-07 | 0.00438 |
| Fibroblast | F0 SFRP2+ Fib | MTDH | 0.359 | 0.724 | 0.522 | 1.31e-06 | 0.044 |
| Fibroblast | F0 SFRP2+ Fib | TCF4 | 0.356 | 0.61 | 0.421 | 5.16e-07 | 0.0174 |
| Fibroblast | F0 SFRP2+ Fib | RPSA | 0.348 | 0.979 | 0.855 | 2.61e-10 | 8.81e-06 |
| Fibroblast | F0 SFRP2+ Fib | GLTSCR2 | 0.348 | 0.915 | 0.729 | 1.04e-09 | 3.5e-05 |
| Fibroblast | F0 SFRP2+ Fib | KLF10 | 0.346 | 0.637 | 0.438 | 1.12e-07 | 0.00378 |
| Fibroblast | F0 SFRP2+ Fib | CD164 | 0.342 | 0.679 | 0.453 | 1.73e-07 | 0.00584 |
| Fibroblast | F0 SFRP2+ Fib | SCP2 | 0.34 | 0.658 | 0.421 | 1.83e-08 | 0.000616 |
| Fibroblast | F0 SFRP2+ Fib | CCNG2 | 0.34 | 0.324 | 0.168 | 1.43e-07 | 0.00481 |
| Fibroblast | F0 SFRP2+ Fib | KDELR2 | 0.332 | 0.732 | 0.493 | 2.48e-08 | 0.000835 |
| Fibroblast | F0 SFRP2+ Fib | TMEM59 | 0.322 | 0.886 | 0.678 | 2.71e-07 | 0.00911 |
| Fibroblast | F0 SFRP2+ Fib | DNAJB1 | 0.318 | 0.963 | 0.824 | 1.05e-09 | 3.53e-05 |
| Fibroblast | F0 SFRP2+ Fib | ARF4 | 0.318 | 0.777 | 0.541 | 4.02e-08 | 0.00135 |
| Fibroblast | F0 SFRP2+ Fib | RCN3 | 0.317 | 0.568 | 0.332 | 3.26e-09 | 0.00011 |
| Fibroblast | F0 SFRP2+ Fib | GPX4 | 0.317 | 0.875 | 0.7 | 1e-06 | 0.0338 |
| Fibroblast | F0 SFRP2+ Fib | VKORC1 | 0.306 | 0.767 | 0.544 | 8.15e-07 | 0.0275 |
| Fibroblast | F0 SFRP2+ Fib | NFKBIZ | 0.306 | 0.639 | 0.438 | 1.48e-06 | 0.0497 |
| Fibroblast | F0 SFRP2+ Fib | TMEM258 | 0.297 | 0.886 | 0.657 | 5.92e-08 | 0.002 |
| Fibroblast | F0 SFRP2+ Fib | SPCS2 | 0.287 | 0.756 | 0.541 | 1.13e-06 | 0.0382 |
| Fibroblast | F0 SFRP2+ Fib | SNX3 | 0.286 | 0.891 | 0.686 | 6.21e-07 | 0.0209 |
| Fibroblast | F0 SFRP2+ Fib | EEF1B2 | 0.277 | 0.907 | 0.726 | 1.1e-06 | 0.0372 |
| Fibroblast | F0 SFRP2+ Fib | RPS12 | 0.266 | 0.997 | 0.971 | 2.32e-11 | 7.81e-07 |
| Fibroblast | F0 SFRP2+ Fib | RPL28 | 0.263 | 1.0 | 0.971 | 2.32e-12 | 7.82e-08 |
| Fibroblast | F0 SFRP2+ Fib | ANGPTL4 | 0.259 | 0.347 | 0.19 | 4.56e-07 | 0.0154 |
| Fibroblast | F0 SFRP2+ Fib | IER2 | 0.256 | 0.995 | 0.885 | 6.72e-09 | 0.000226 |
| Fibroblast | F1 MYH11+ Fib | SBSPON | 3.017 | 0.125 | 0.023 | 3.45e-11 | 1.16e-06 |
| Fibroblast | F1 MYH11+ Fib | HRH2 | 2.824 | 0.121 | 0.026 | 1.71e-09 | 5.78e-05 |
| Fibroblast | F1 MYH11+ Fib | SPEG | 2.798 | 0.151 | 0.04 | 5.62e-10 | 1.89e-05 |
| Fibroblast | F1 MYH11+ Fib | HEXIM1 | 1.753 | 0.56 | 0.5 | 7.31e-09 | 0.000246 |
| Fibroblast | F1 MYH11+ Fib | JUN | 1.434 | 0.897 | 0.982 | 1.46e-09 | 4.9e-05 |
| Fibroblast | F1 MYH11+ Fib | MYH11 | 1.223 | 0.431 | 0.19 | 2.67e-14 | 8.98e-10 |
| Fibroblast | F1 MYH11+ Fib | GADD45B | 1.174 | 0.845 | 0.94 | 1.9e-08 | 0.000639 |
| Fibroblast | F1 MYH11+ Fib | ADIRF | 1.157 | 0.612 | 0.53 | 1.14e-08 | 0.000385 |
| Fibroblast | F1 MYH11+ Fib | HSPB1 | 1.095 | 0.94 | 0.971 | 6.19e-15 | 2.09e-10 |
| Fibroblast | F1 MYH11+ Fib | FOS | 1.027 | 0.909 | 0.98 | 1.54e-09 | 5.2e-05 |
| Fibroblast | F1 MYH11+ Fib | MALAT1 | 0.914 | 0.991 | 1.0 | 1.06e-17 | 3.57e-13 |
| Fibroblast | F1 MYH11+ Fib | H3F3B | 0.749 | 0.944 | 0.995 | 1.02e-09 | 3.43e-05 |
| Fibroblast | F1 MYH11+ Fib | MT-ATP6 | 0.615 | 0.991 | 0.997 | 1.81e-14 | 6.11e-10 |
| Fibroblast | F1 MYH11+ Fib | MT-ND2 | 0.573 | 0.991 | 0.996 | 1.43e-16 | 4.83e-12 |
| Fibroblast | F1 MYH11+ Fib | MT-ND1 | 0.501 | 0.966 | 0.992 | 3.05e-09 | 0.000103 |
| Fibroblast | F1 MYH11+ Fib | PTMA | 0.498 | 1.0 | 1.0 | 2.95e-15 | 9.93e-11 |
| Fibroblast | F1 MYH11+ Fib | MT-CYB | 0.497 | 0.987 | 0.997 | 3.5e-10 | 1.18e-05 |
| Fibroblast | F1 MYH11+ Fib | MT-CO1 | 0.48 | 1.0 | 0.999 | 2.84e-10 | 9.55e-06 |
| Fibroblast | F1 MYH11+ Fib | MT-CO3 | 0.47 | 0.996 | 0.997 | 1.33e-09 | 4.49e-05 |
| Fibroblast | F1 MYH11+ Fib | EIF1 | 0.438 | 0.978 | 1.0 | 7.16e-08 | 0.00241 |
| Fibroblast | F1 MYH11+ Fib | MT-ND4 | 0.38 | 0.996 | 0.999 | 1.44e-09 | 4.84e-05 |
| Fibroblast | F1 MYH11+ Fib | RAB13 | 0.375 | 0.31 | 0.61 | 9.15e-07 | 0.0308 |
| Fibroblast | F2 RGS5+ Fib | COX4I2 | 5.289 | 0.39 | 0.007 | 5.4e-70 | 1.82e-65 |
| Fibroblast | F2 RGS5+ Fib | GPR4 | 5.077 | 0.277 | 0.008 | 2.86e-46 | 9.619999999999999e-42 |
| Fibroblast | F2 RGS5+ Fib | RGS5 | 5.072 | 0.795 | 0.08 | 3.08e-114 | 1.04e-109 |

| compartment | subcluster | gene | avg_log2FC | pct.1 | pct.2 | p_val | p_val_adj |
|---|---|---|---|---|---|---|---|
| Fibroblast | F2 RGS5+ Fib | NDUFA4L2 | 4.618 | 0.841 | 0.129 | 7.5e-109 | 2.53e-104 |
| Fibroblast | F2 RGS5+ Fib | CDH6 | 4.31 | 0.318 | 0.016 | 2.17e-48 | 7.300000000000001e-44 |
| Fibroblast | F2 RGS5+ Fib | HRAT92 | 4.153 | 0.103 | 0.002 | 3.71e-18 | 1.25e-13 |
| Fibroblast | F2 RGS5+ Fib | CACNA1H | 4.117 | 0.221 | 0.017 | 2.29e-29 | 7.72e-25 |
| Fibroblast | F2 RGS5+ Fib | GUCY1A2 | 4.093 | 0.123 | 0.006 | 5.62e-19 | 1.89e-14 |
| Fibroblast | F2 RGS5+ Fib | ADGRF5 | 4.064 | 0.149 | 0.01 | 4.21e-21 | 1.42e-16 |
| Fibroblast | F2 RGS5+ Fib | GRIP2 | 4.018 | 0.113 | 0.006 | 4.3e-17 | 1.45e-12 |
| Fibroblast | F2 RGS5+ Fib | HIGD1B | 3.961 | 0.431 | 0.022 | 1.04e-65 | 3.51e-61 |
| Fibroblast | F2 RGS5+ Fib | SERPINI1 | 3.931 | 0.369 | 0.041 | 2.37e-43 | 7.99e-39 |
| Fibroblast | F2 RGS5+ Fib | KLHL23 | 3.914 | 0.313 | 0.019 | 1.19e-44 | 4e-40 |
| Fibroblast | F2 RGS5+ Fib | CABP1 | 3.883 | 0.164 | 0.01 | 9.84e-24 | 3.32e-19 |
| Fibroblast | F2 RGS5+ Fib | SEMA5B | 3.878 | 0.103 | 0.002 | 3.74e-18 | 1.26e-13 |
| Fibroblast | F2 RGS5+ Fib | ENPEP | 3.854 | 0.19 | 0.016 | 1.55e-24 | 5.23e-20 |
| Fibroblast | F2 RGS5+ Fib | EFHD1 | 3.818 | 0.513 | 0.048 | 2.36e-66 | 7.959999999999999e-62 |
| Fibroblast | F2 RGS5+ Fib | ADAP2 | 3.817 | 0.297 | 0.014 | 7.72e-45 | 2.6e-40 |
| Fibroblast | F2 RGS5+ Fib | MAP3K7CL | 3.771 | 0.554 | 0.063 | 5.79e-68 | 1.95e-63 |
| Fibroblast | F2 RGS5+ Fib | ACAN | 3.744 | 0.103 | 0.01 | 5.26e-13 | 1.77e-08 |
| Fibroblast | F2 RGS5+ Fib | NRGN | 3.677 | 0.313 | 0.025 | 7.52e-41 | 2.53e-36 |
| Fibroblast | F2 RGS5+ Fib | RERGL | 3.611 | 0.344 | 0.053 | 3.94e-33 | 1.33e-28 |
| Fibroblast | F2 RGS5+ Fib | ESAM | 3.559 | 0.482 | 0.037 | 1.03e-64 | 3.46e-60 |
| Fibroblast | F2 RGS5+ Fib | NRIP2 | 3.496 | 0.231 | 0.025 | 2.14e-26 | 7.21e-22 |
| Fibroblast | F2 RGS5+ Fib | ITGA3 | 3.474 | 0.185 | 0.022 | 2.12e-20 | 7.15e-16 |
| Fibroblast | F2 RGS5+ Fib | GJC1 | 3.386 | 0.287 | 0.026 | 2.2e-35 | 7.42e-31 |
| Fibroblast | F2 RGS5+ Fib | OR51E1 | 3.373 | 0.179 | 0.011 | 2.42e-25 | 8.150000000000001e-21 |
| Fibroblast | F2 RGS5+ Fib | GJA4 | 3.353 | 0.626 | 0.05 | 1.2e-85 | 4.04e-81 |
| Fibroblast | F2 RGS5+ Fib | OLFML2A | 3.342 | 0.205 | 0.025 | 4.93e-22 | 1.66e-17 |
| Fibroblast | F2 RGS5+ Fib | TMEM74B | 3.319 | 0.103 | 0.007 | 2.42e-14 | 8.15e-10 |
| Fibroblast | F2 RGS5+ Fib | ITGA7 | 3.292 | 0.359 | 0.04 | 4.61e-41 | 1.55e-36 |
| Fibroblast | F2 RGS5+ Fib | FAM162B | 3.245 | 0.236 | 0.031 | 2.61e-24 | 8.79e-20 |
| Fibroblast | F2 RGS5+ Fib | NFASC | 3.223 | 0.303 | 0.041 | 3.86e-31 | 1.3e-26 |
| Fibroblast | F2 RGS5+ Fib | GPAT2 | 3.21 | 0.144 | 0.025 | 1.49e-12 | 5.01e-08 |
| Fibroblast | F2 RGS5+ Fib | GPRC5C | 3.208 | 0.436 | 0.055 | 1.11e-47 | 3.73e-43 |
| Fibroblast | F2 RGS5+ Fib | WFDC1 | 3.207 | 0.349 | 0.049 | 3.28e-35 | 1.11e-30 |
| Fibroblast | F2 RGS5+ Fib | TESC | 3.198 | 0.164 | 0.013 | 1.95e-21 | 6.58e-17 |
| Fibroblast | F2 RGS5+ Fib | C17orf82 | 3.159 | 0.118 | 0.016 | 1.91e-12 | 6.43e-08 |
| Fibroblast | F2 RGS5+ Fib | TINAGL1 | 3.151 | 0.81 | 0.12 | 1.99e-92 | 6.7e-88 |
| Fibroblast | F2 RGS5+ Fib | EPS8L1 | 3.133 | 0.118 | 0.017 | 5.53e-12 | 1.86e-07 |
| Fibroblast | F2 RGS5+ Fib | BCAM | 3.123 | 0.662 | 0.108 | 7.23e-68 | 2.44e-63 |
| Fibroblast | F2 RGS5+ Fib | TBX2-AS1 | 3.102 | 0.477 | 0.084 | 1.01e-43 | 3.42e-39 |
| Fibroblast | F2 RGS5+ Fib | ADRA2B | 3.072 | 0.113 | 0.011 | 6.87e-14 | 2.31e-09 |
| Fibroblast | F2 RGS5+ Fib | MCAM | 3.064 | 0.646 | 0.077 | 9.940000000000001e-77 | 3.35e-72 |
| Fibroblast | F2 RGS5+ Fib | PTH1R | 3.013 | 0.128 | 0.023 | 3.36e-11 | 1.13e-06 |
| Fibroblast | F2 RGS5+ Fib | CRIP1 | 3.008 | 0.877 | 0.281 | 1.47e-76 | 4.97e-72 |
| Fibroblast | F2 RGS5+ Fib | KCNA5 | 2.993 | 0.221 | 0.018 | 3.58e-28 | 1.21e-23 |
| Fibroblast | F2 RGS5+ Fib | EGFLAM | 2.96 | 0.149 | 0.028 | 1.9e-12 | 6.41e-08 |
| Fibroblast | F2 RGS5+ Fib | HEY2 | 2.957 | 0.323 | 0.043 | 6.92e-33 | 2.33e-28 |
| Fibroblast | F2 RGS5+ Fib | MEF2C | 2.952 | 0.687 | 0.149 | 1.59e-62 | 5.34e-58 |
| Fibroblast | F2 RGS5+ Fib | LGI4 | 2.941 | 0.359 | 0.034 | 4.91e-43 | 1.65e-38 |
| Fibroblast | F2 RGS5+ Fib | PTP4A3 | 2.938 | 0.672 | 0.109 | 2e-68 | 6.740000000000001e-64 |
| Fibroblast | F2 RGS5+ Fib | ADCY5 | 2.895 | 0.123 | 0.016 | 4.03e-13 | 1.36e-08 |
| Fibroblast | F2 RGS5+ Fib | PGF | 2.89 | 0.687 | 0.184 | 2.19e-55 | 7.37e-51 |
| Fibroblast | F2 RGS5+ Fib | CAP2 | 2.839 | 0.21 | 0.047 | 2.37e-15 | 7.99e-11 |
| Fibroblast | F2 RGS5+ Fib | RASGRP2 | 2.837 | 0.374 | 0.074 | 7.630000000000001e-31 | 2.57e-26 |
| Fibroblast | F2 RGS5+ Fib | GPR20 | 2.834 | 0.303 | 0.03 | 2.78e-35 | 9.36e-31 |
| Fibroblast | F2 RGS5+ Fib | TBX2 | 2.827 | 0.456 | 0.077 | 2.14e-42 | 7.22e-38 |
| Fibroblast | F2 RGS5+ Fib | ISYNA1 | 2.824 | 0.641 | 0.153 | 3.81e-53 | 1.29e-48 |
| Fibroblast | F2 RGS5+ Fib | AFAP1L2 | 2.819 | 0.154 | 0.036 | 1.04e-10 | 3.52e-06 |
| Fibroblast | F2 RGS5+ Fib | INPP4B | 2.815 | 0.144 | 0.024 | 6.22e-13 | 2.1e-08 |
| Fibroblast | F2 RGS5+ Fib | NEURL1B | 2.806 | 0.19 | 0.029 | 6.38e-18 | 2.15e-13 |
| Fibroblast | F2 RGS5+ Fib | DMPK | 2.793 | 0.308 | 0.065 | 1.19e-23 | 4.02e-19 |
| Fibroblast | F2 RGS5+ Fib | HEYL | 2.779 | 0.385 | 0.078 | 1.96e-30 | 6.610000000000001e-26 |
| Fibroblast | F2 RGS5+ Fib | NOTCH3 | 2.777 | 0.826 | 0.22 | 8.49e-73 | 2.86e-68 |
| Fibroblast | F2 RGS5+ Fib | SEPT4 | 2.763 | 0.585 | 0.158 | 9.73e-44 | 3.28e-39 |
| Fibroblast | F2 RGS5+ Fib | PTK2 | 2.747 | 0.59 | 0.15 | 9.99e-47 | 3.359999999999999e-42 |

| compartment | subcluster | gene | avg_log2FC | pct.1 | pct.2 | p_val | p_val_adj |
|---|---|---|---|---|---|---|---|
| Fibroblast | F2 RGS5+ Fib | LDB3 | 2.73 | 0.174 | 0.02 | 7.76e-19 | 2.61e-14 |
| Fibroblast | F2 RGS5+ Fib | LAMA5 | 2.705 | 0.256 | 0.037 | 1.02e-24 | 3.44e-20 |
| Fibroblast | F2 RGS5+ Fib | PLN | 2.704 | 0.544 | 0.115 | 7.01e-44 | 2.36e-39 |
| Fibroblast | F2 RGS5+ Fib | CSRP2 | 2.694 | 0.759 | 0.205 | 2.54e-60 | 8.57e-56 |
| Fibroblast | F2 RGS5+ Fib | RCAN2 | 2.687 | 0.544 | 0.12 | 1.8e-44 | 6.07e-40 |
| Fibroblast | F2 RGS5+ Fib | NTF3 | 2.686 | 0.164 | 0.031 | 3.01e-13 | 1.02e-08 |
| Fibroblast | F2 RGS5+ Fib | CNNM2 | 2.685 | 0.108 | 0.018 | 6.6e-10 | 2.22e-05 |
| Fibroblast | F2 RGS5+ Fib | FOXS1 | 2.684 | 0.456 | 0.089 | 2.13e-38 | 7.18e-34 |
| Fibroblast | F2 RGS5+ Fib | SORT1 | 2.681 | 0.226 | 0.048 | 6.29e-17 | 2.12e-12 |
| Fibroblast | F2 RGS5+ Fib | ANO1 | 2.66 | 0.205 | 0.04 | 2.33e-16 | 7.86e-12 |
| Fibroblast | F2 RGS5+ Fib | ATP1B2 | 2.653 | 0.2 | 0.026 | 2.48e-20 | 8.36e-16 |
| Fibroblast | F2 RGS5+ Fib | TPPP3 | 2.635 | 0.692 | 0.152 | 1.15e-61 | 3.86e-57 |
| Fibroblast | F2 RGS5+ Fib | SSTR2 | 2.629 | 0.164 | 0.013 | 4.8e-21 | 1.62e-16 |
| Fibroblast | F2 RGS5+ Fib | PLCE1 | 2.628 | 0.277 | 0.046 | 4.97e-25 | 1.67e-20 |
| Fibroblast | F2 RGS5+ Fib | HRC | 2.613 | 0.179 | 0.016 | 4.24e-22 | 1.43e-17 |
| Fibroblast | F2 RGS5+ Fib | RP3-395M20.12 | 2.606 | 0.138 | 0.026 | 2.54e-11 | 8.55e-07 |
| Fibroblast | F2 RGS5+ Fib | MGLL | 2.602 | 0.405 | 0.087 | 4.66e-31 | 1.57e-26 |
| Fibroblast | F2 RGS5+ Fib | BCAS3 | 2.602 | 0.164 | 0.041 | 7.87e-11 | 2.65e-06 |
| Fibroblast | F2 RGS5+ Fib | CSPG4 | 2.601 | 0.215 | 0.025 | 2.49e-23 | 8.37e-19 |
| Fibroblast | F2 RGS5+ Fib | MYH11 | 2.597 | 0.723 | 0.133 | 3.11e-66 | 1.05e-61 |
| Fibroblast | F2 RGS5+ Fib | LGALSL | 2.552 | 0.159 | 0.031 | 1.36e-12 | 4.6e-08 |
| Fibroblast | F2 RGS5+ Fib | CPM | 2.551 | 0.231 | 0.077 | 4.18e-11 | 1.41e-06 |
| Fibroblast | F2 RGS5+ Fib | CCDC3 | 2.521 | 0.446 | 0.109 | 1.16e-31 | 3.9e-27 |
| Fibroblast | F2 RGS5+ Fib | CLMN | 2.512 | 0.318 | 0.073 | 1.7e-22 | 5.73e-18 |
| Fibroblast | F2 RGS5+ Fib | KIAA0040 | 2.494 | 0.292 | 0.067 | 1.2e-20 | 4.05e-16 |
| Fibroblast | F2 RGS5+ Fib | MRVI1 | 2.477 | 0.256 | 0.074 | 9.22e-15 | 3.11e-10 |
| Fibroblast | F2 RGS5+ Fib | CDK18 | 2.475 | 0.103 | 0.022 | 5.36e-08 | 0.0018 |
| Fibroblast | F2 RGS5+ Fib | MYO1B | 2.457 | 0.221 | 0.083 | 2.38e-09 | 8.01e-05 |
| Fibroblast | F2 RGS5+ Fib | CASQ2 | 2.454 | 0.128 | 0.014 | 1.87e-14 | 6.3e-10 |
| Fibroblast | F2 RGS5+ Fib | HOXB-AS1 | 2.445 | 0.385 | 0.095 | 4.02e-26 | 1.35e-21 |
| Fibroblast | F2 RGS5+ Fib | RBPMS2 | 2.445 | 0.185 | 0.036 | 1.4e-14 | 4.73e-10 |
| Fibroblast | F2 RGS5+ Fib | SGCA | 2.439 | 0.297 | 0.059 | 5.889999999999999e-23 | 1.99e-18 |
| Fibroblast | F2 RGS5+ Fib | PDGFA | 2.432 | 0.369 | 0.087 | 9.67e-26 | 3.26e-21 |
| Fibroblast | F2 RGS5+ Fib | CCDC102B | 2.393 | 0.318 | 0.087 | 6.01e-19 | 2.02e-14 |
| Fibroblast | F2 RGS5+ Fib | PACSIN2 | 2.377 | 0.338 | 0.095 | 1.55e-20 | 5.21e-16 |
| Fibroblast | F2 RGS5+ Fib | SLC7A2 | 2.365 | 0.169 | 0.038 | 8.7e-12 | 2.93e-07 |
| Fibroblast | F2 RGS5+ Fib | CRIP2 | 2.354 | 0.856 | 0.375 | 8.15e-60 | 2.74e-55 |
| Fibroblast | F2 RGS5+ Fib | C1QTNF1 | 2.35 | 0.477 | 0.123 | 3.34e-33 | 1.12e-28 |
| Fibroblast | F2 RGS5+ Fib | MTHFD2 | 2.346 | 0.503 | 0.169 | 4.46e-28 | 1.5e-23 |
| Fibroblast | F2 RGS5+ Fib | SUSD5 | 2.339 | 0.133 | 0.022 | 4.95e-12 | 1.67e-07 |
| Fibroblast | F2 RGS5+ Fib | USP2 | 2.338 | 0.133 | 0.024 | 2.69e-11 | 9.07e-07 |
| Fibroblast | F2 RGS5+ Fib | NTN4 | 2.326 | 0.262 | 0.061 | 7.65e-18 | 2.58e-13 |
| Fibroblast | F2 RGS5+ Fib | CHCHD10 | 2.324 | 0.677 | 0.256 | 3.23e-41 | 1.09e-36 |
| Fibroblast | F2 RGS5+ Fib | RCSD1 | 2.316 | 0.236 | 0.054 | 5.84e-16 | 1.97e-11 |
| Fibroblast | F2 RGS5+ Fib | ARHGEF17 | 2.31 | 0.292 | 0.089 | 5.6e-16 | 1.89e-11 |
| Fibroblast | F2 RGS5+ Fib | MYOCD | 2.31 | 0.113 | 0.014 | 6.17e-12 | 2.08e-07 |
| Fibroblast | F2 RGS5+ Fib | OAZ2 | 2.3 | 0.728 | 0.398 | 1.34e-38 | 4.51e-34 |
| Fibroblast | F2 RGS5+ Fib | NET1 | 2.3 | 0.462 | 0.176 | 4.91e-22 | 1.66e-17 |
| Fibroblast | F2 RGS5+ Fib | DAAM2 | 2.298 | 0.226 | 0.043 | 5.94e-18 | 2e-13 |
| Fibroblast | F2 RGS5+ Fib | PPP1R14A | 2.293 | 0.887 | 0.247 | 5.29e-70 | 1.78e-65 |
| Fibroblast | F2 RGS5+ Fib | KLHDC8B | 2.262 | 0.251 | 0.079 | 8.53e-13 | 2.88e-08 |
| Fibroblast | F2 RGS5+ Fib | RNF180 | 2.252 | 0.21 | 0.063 | 2.43e-11 | 8.18e-07 |
| Fibroblast | F2 RGS5+ Fib | ANGPT2 | 2.249 | 0.379 | 0.06 | 5.15e-35 | 1.73e-30 |
| Fibroblast | F2 RGS5+ Fib | ADIRF | 2.243 | 0.892 | 0.468 | 1.72e-51 | 5.790000000000001e-47 |
| Fibroblast | F2 RGS5+ Fib | FILIP1 | 2.232 | 0.369 | 0.08 | 1.64e-27 | 5.53e-23 |
| Fibroblast | F2 RGS5+ Fib | B3GNT2 | 2.225 | 0.421 | 0.119 | 7.33e-26 | 2.47e-21 |
| Fibroblast | F2 RGS5+ Fib | PLEKHH3 | 2.209 | 0.282 | 0.068 | 1.87e-18 | 6.29e-14 |
| Fibroblast | F2 RGS5+ Fib | TOX2 | 2.208 | 0.118 | 0.032 | 3.74e-07 | 0.0126 |
| Fibroblast | F2 RGS5+ Fib | DACT3 | 2.204 | 0.144 | 0.038 | 1.07e-08 | 0.000362 |
| Fibroblast | F2 RGS5+ Fib | PARM1 | 2.203 | 0.159 | 0.04 | 3.84e-10 | 1.29e-05 |
| Fibroblast | F2 RGS5+ Fib | FRMD4A | 2.191 | 0.2 | 0.065 | 4.67e-10 | 1.57e-05 |
| Fibroblast | F2 RGS5+ Fib | EPAS1 | 2.19 | 0.574 | 0.224 | 6.65e-29 | 2.24e-24 |
| Fibroblast | F2 RGS5+ Fib | PCDH1 | 2.188 | 0.123 | 0.018 | 5.66e-12 | 1.91e-07 |
| Fibroblast | F2 RGS5+ Fib | SYNM | 2.179 | 0.149 | 0.036 | 9.18e-10 | 3.09e-05 |

| compartment | subcluster | gene | avg_log2FC | pct.1 | pct.2 | p_val | p_val_adj |
|---|---|---|---|---|---|---|---|
| Fibroblast | F2 RGS5+ Fib | CAV2 | 2.173 | 0.59 | 0.219 | 1.16e-32 | 3.93e-28 |
| Fibroblast | F2 RGS5+ Fib | FHL5 | 2.171 | 0.133 | 0.025 | 1.01e-10 | 3.42e-06 |
| Fibroblast | F2 RGS5+ Fib | SPECC1 | 2.153 | 0.179 | 0.065 | 8.13e-08 | 0.00274 |
| Fibroblast | F2 RGS5+ Fib | NGF | 2.146 | 0.215 | 0.043 | 2.27e-16 | 7.65e-12 |
| Fibroblast | F2 RGS5+ Fib | AC133644.2 | 2.139 | 0.236 | 0.078 | 2.62e-11 | 8.82e-07 |
| Fibroblast | F2 RGS5+ Fib | SNCG | 2.131 | 0.179 | 0.053 | 1.13e-09 | 3.81e-05 |
| Fibroblast | F2 RGS5+ Fib | MYLK | 2.128 | 0.641 | 0.279 | 2.42e-30 | 8.14e-26 |
| Fibroblast | F2 RGS5+ Fib | TBC1D1 | 2.118 | 0.308 | 0.089 | 1.77e-17 | 5.98e-13 |
| Fibroblast | F2 RGS5+ Fib | ACTA2 | 2.117 | 0.933 | 0.622 | 9.93e-46 | 3.35e-41 |
| Fibroblast | F2 RGS5+ Fib | SORBS2 | 2.104 | 0.262 | 0.063 | 9.26e-17 | 3.12e-12 |
| Fibroblast | F2 RGS5+ Fib | MAPRE2 | 2.098 | 0.308 | 0.122 | 2.48e-12 | 8.36e-08 |
| Fibroblast | F2 RGS5+ Fib | EHD2 | 2.094 | 0.554 | 0.267 | 5.14e-23 | 1.73e-18 |
| Fibroblast | F2 RGS5+ Fib | COL18A1 | 2.083 | 0.887 | 0.51 | 2.1e-51 | 7.08e-47 |
| Fibroblast | F2 RGS5+ Fib | DGKH | 2.081 | 0.149 | 0.043 | 2.63e-08 | 0.000888 |
| Fibroblast | F2 RGS5+ Fib | NPNT | 2.063 | 0.164 | 0.034 | 2.75e-12 | 9.27e-08 |
| Fibroblast | F2 RGS5+ Fib | KIAA0895L | 2.06 | 0.108 | 0.029 | 1.05e-06 | 0.0352 |
| Fibroblast | F2 RGS5+ Fib | LINGO1 | 2.045 | 0.108 | 0.01 | 1.23e-13 | 4.13e-09 |
| Fibroblast | F2 RGS5+ Fib | NES | 2.023 | 0.236 | 0.074 | 7.27e-12 | 2.45e-07 |
| Fibroblast | F2 RGS5+ Fib | SNTA1 | 2.018 | 0.2 | 0.047 | 3.36e-13 | 1.13e-08 |
| Fibroblast | F2 RGS5+ Fib | TMEM38B | 2.01 | 0.236 | 0.086 | 5.03e-10 | 1.7e-05 |
| Fibroblast | F2 RGS5+ Fib | SYNGR2 | 1.995 | 0.267 | 0.097 | 1.17e-11 | 3.94e-07 |
| Fibroblast | F2 RGS5+ Fib | CRIM1 | 1.971 | 0.256 | 0.098 | 2.26e-10 | 7.62e-06 |
| Fibroblast | F2 RGS5+ Fib | JAG1 | 1.961 | 0.431 | 0.174 | 7e-18 | 2.36e-13 |
| Fibroblast | F2 RGS5+ Fib | FRZB | 1.957 | 0.785 | 0.387 | 1.77e-36 | 5.97e-32 |
| Fibroblast | F2 RGS5+ Fib | LZTS1 | 1.943 | 0.205 | 0.051 | 8.67e-13 | 2.92e-08 |
| Fibroblast | F2 RGS5+ Fib | CARMN | 1.942 | 0.4 | 0.116 | 2.28e-22 | 7.69e-18 |
| Fibroblast | F2 RGS5+ Fib | MYL9 | 1.938 | 0.969 | 0.794 | 2.32e-51 | 7.81e-47 |
| Fibroblast | F2 RGS5+ Fib | ARHGDIB | 1.932 | 0.554 | 0.29 | 2.28e-18 | 7.69e-14 |
| Fibroblast | F2 RGS5+ Fib | MPRIP | 1.929 | 0.303 | 0.143 | 2.37e-09 | 7.99e-05 |
| Fibroblast | F2 RGS5+ Fib | RASL12 | 1.923 | 0.374 | 0.125 | 1.13e-18 | 3.8e-14 |
| Fibroblast | F2 RGS5+ Fib | UBA2 | 1.918 | 0.497 | 0.223 | 1.28e-19 | 4.3e-15 |
| Fibroblast | F2 RGS5+ Fib | CCND1 | 1.917 | 0.282 | 0.128 | 4.6e-09 | 0.000155 |
| Fibroblast | F2 RGS5+ Fib | TMEM51 | 1.907 | 0.174 | 0.068 | 8.62e-07 | 0.029 |
| Fibroblast | F2 RGS5+ Fib | IMPA2 | 1.904 | 0.169 | 0.049 | 2.97e-09 | 0.0001 |
| Fibroblast | F2 RGS5+ Fib | NT5DC2 | 1.902 | 0.287 | 0.105 | 2.37e-12 | 7.98e-08 |
| Fibroblast | F2 RGS5+ Fib | ARHGAP29 | 1.881 | 0.369 | 0.114 | 3.09e-19 | 1.04e-14 |
| Fibroblast | F2 RGS5+ Fib | ARHGEF7 | 1.869 | 0.277 | 0.098 | 4.61e-12 | 1.55e-07 |
| Fibroblast | F2 RGS5+ Fib | FKBP4 | 1.86 | 0.333 | 0.134 | 4.52e-13 | 1.52e-08 |
| Fibroblast | F2 RGS5+ Fib | ENTPD1 | 1.85 | 0.313 | 0.114 | 1.18e-13 | 3.97e-09 |
| Fibroblast | F2 RGS5+ Fib | ITGA1 | 1.847 | 0.328 | 0.159 | 2.06e-09 | 6.93e-05 |
| Fibroblast | F2 RGS5+ Fib | AOC3 | 1.84 | 0.277 | 0.105 | 1.87e-11 | 6.31e-07 |
| Fibroblast | F2 RGS5+ Fib | CNN1 | 1.82 | 0.441 | 0.146 | 4.13e-22 | 1.39e-17 |
| Fibroblast | F2 RGS5+ Fib | AC013461.1 | 1.815 | 0.508 | 0.291 | 1.01e-15 | 3.4e-11 |
| Fibroblast | F2 RGS5+ Fib | CACNA1C | 1.805 | 0.185 | 0.075 | 1.17e-06 | 0.0395 |
| Fibroblast | F2 RGS5+ Fib | LRRC32 | 1.801 | 0.4 | 0.193 | 2.3e-12 | 7.74e-08 |
| Fibroblast | F2 RGS5+ Fib | PDE3A | 1.796 | 0.169 | 0.053 | 1.78e-08 | 0.000599 |
| Fibroblast | F2 RGS5+ Fib | STK11 | 1.794 | 0.154 | 0.054 | 7.49e-07 | 0.0252 |
| Fibroblast | F2 RGS5+ Fib | FAM129A | 1.791 | 0.251 | 0.083 | 1.17e-11 | 3.94e-07 |
| Fibroblast | F2 RGS5+ Fib | ANGPT1 | 1.791 | 0.215 | 0.067 | 8.3e-11 | 2.8e-06 |
| Fibroblast | F2 RGS5+ Fib | COBLL1 | 1.791 | 0.185 | 0.066 | 6.65e-08 | 0.00224 |
| Fibroblast | F2 RGS5+ Fib | ADAMTS4 | 1.759 | 0.359 | 0.164 | 9.87e-12 | 3.33e-07 |
| Fibroblast | F2 RGS5+ Fib | CAV1 | 1.751 | 0.754 | 0.406 | 5.04e-32 | 1.7e-27 |
| Fibroblast | F2 RGS5+ Fib | DSTN | 1.748 | 0.944 | 0.853 | 8.17e-36 | 2.75e-31 |
| Fibroblast | F2 RGS5+ Fib | RP11-394O4.5 | 1.748 | 0.113 | 0.026 | 8.7e-08 | 0.00293 |
| Fibroblast | F2 RGS5+ Fib | FOXC2 | 1.742 | 0.246 | 0.071 | 4.01e-13 | 1.35e-08 |
| Fibroblast | F2 RGS5+ Fib | EDNRA | 1.732 | 0.292 | 0.122 | 2.94e-10 | 9.89e-06 |
| Fibroblast | F2 RGS5+ Fib | LINC01197 | 1.729 | 0.118 | 0.034 | 9.67e-07 | 0.0326 |
| Fibroblast | F2 RGS5+ Fib | SOX13 | 1.725 | 0.128 | 0.04 | 9.51e-07 | 0.032 |
| Fibroblast | F2 RGS5+ Fib | ADAMTS9 | 1.723 | 0.185 | 0.044 | 5.83e-12 | 1.96e-07 |
| Fibroblast | F2 RGS5+ Fib | HDAC5 | 1.71 | 0.277 | 0.11 | 2.36e-10 | 7.94e-06 |
| Fibroblast | F2 RGS5+ Fib | GUCY1B3 | 1.708 | 0.4 | 0.16 | 4.68e-16 | 1.58e-11 |
| Fibroblast | F2 RGS5+ Fib | HIP1 | 1.707 | 0.236 | 0.104 | 1.22e-07 | 0.00412 |
| Fibroblast | F2 RGS5+ Fib | SLCO3A1 | 1.706 | 0.241 | 0.114 | 3.5e-07 | 0.0118 |
| Fibroblast | F2 RGS5+ Fib | C4orf32 | 1.697 | 0.226 | 0.08 | 1.54e-09 | 5.19e-05 |

| compartment | subcluster | gene | avg_log2FC | pct.1 | pct.2 | p_val | p_val_adj |
|---|---|---|---|---|---|---|---|
| Fibroblast | F2 RGS5+ Fib | ITGA8 | 1.695 | 0.103 | 0.025 | 7.21e-07 | 0.0243 |
| Fibroblast | F2 RGS5+ Fib | HACD1 | 1.692 | 0.169 | 0.056 | 6.06e-08 | 0.00204 |
| Fibroblast | F2 RGS5+ Fib | LMOD1 | 1.684 | 0.451 | 0.176 | 8.29e-19 | 2.79e-14 |
| Fibroblast | F2 RGS5+ Fib | CABIN1 | 1.68 | 0.262 | 0.12 | 3.72e-08 | 0.00125 |
| Fibroblast | F2 RGS5+ Fib | C2orf40 | 1.679 | 0.241 | 0.081 | 8.49e-11 | 2.86e-06 |
| Fibroblast | F2 RGS5+ Fib | PLXDC1 | 1.677 | 0.4 | 0.194 | 6.46e-13 | 2.18e-08 |
| Fibroblast | F2 RGS5+ Fib | COL4A1 | 1.673 | 0.477 | 0.358 | 7.05e-07 | 0.0237 |
| Fibroblast | F2 RGS5+ Fib | TLE1 | 1.66 | 0.349 | 0.134 | 6.31e-14 | 2.13e-09 |
| Fibroblast | F2 RGS5+ Fib | PDGFRB | 1.659 | 0.692 | 0.398 | 1.79e-24 | 6.02e-20 |
| Fibroblast | F2 RGS5+ Fib | KNOP1 | 1.652 | 0.303 | 0.15 | 2.01e-08 | 0.000677 |
| Fibroblast | F2 RGS5+ Fib | CTD-3193K9.4 | 1.645 | 0.174 | 0.046 | 2.25e-10 | 7.57e-06 |
| Fibroblast | F2 RGS5+ Fib | TPM2 | 1.634 | 0.974 | 0.811 | 2.9e-45 | 9.78e-41 |
| Fibroblast | F2 RGS5+ Fib | VASP | 1.624 | 0.477 | 0.23 | 3.09e-16 | 1.04e-11 |
| Fibroblast | F2 RGS5+ Fib | FABP4 | 1.623 | 0.179 | 0.061 | 3.38e-08 | 0.00114 |
| Fibroblast | F2 RGS5+ Fib | CAMK2G | 1.623 | 0.221 | 0.095 | 1.57e-07 | 0.00528 |
| Fibroblast | F2 RGS5+ Fib | PRKG1 | 1.621 | 0.19 | 0.075 | 5.7e-07 | 0.0192 |
| Fibroblast | F2 RGS5+ Fib | KRT18 | 1.609 | 0.349 | 0.19 | 4.3e-08 | 0.00145 |
| Fibroblast | F2 RGS5+ Fib | PPP1R12B | 1.608 | 0.354 | 0.135 | 4.74e-14 | 1.6e-09 |
| Fibroblast | F2 RGS5+ Fib | TNS1 | 1.604 | 0.385 | 0.187 | 1.36e-11 | 4.59e-07 |
| Fibroblast | F2 RGS5+ Fib | COL4A2 | 1.599 | 0.61 | 0.413 | 3.55e-13 | 1.2e-08 |
| Fibroblast | F2 RGS5+ Fib | UACA | 1.589 | 0.415 | 0.229 | 6.39e-11 | 2.15e-06 |
| Fibroblast | F2 RGS5+ Fib | ARPC1A | 1.565 | 0.559 | 0.338 | 3.94e-16 | 1.33e-11 |
| Fibroblast | F2 RGS5+ Fib | SYNE2 | 1.565 | 0.277 | 0.115 | 1.1e-09 | 3.72e-05 |
| Fibroblast | F2 RGS5+ Fib | NTRK2 | 1.564 | 0.282 | 0.149 | 7.42e-07 | 0.025 |
| Fibroblast | F2 RGS5+ Fib | NDRG2 | 1.559 | 0.579 | 0.325 | 3.55e-17 | 1.2e-12 |
| Fibroblast | F2 RGS5+ Fib | LBH | 1.558 | 0.59 | 0.325 | 4.71e-18 | 1.59e-13 |
| Fibroblast | F2 RGS5+ Fib | ACTG2 | 1.552 | 0.344 | 0.109 | 1.92e-16 | 6.48e-12 |
| Fibroblast | F2 RGS5+ Fib | CDC42EP4 | 1.538 | 0.318 | 0.137 | 1.2e-10 | 4.06e-06 |
| Fibroblast | F2 RGS5+ Fib | HES4 | 1.537 | 0.769 | 0.431 | 4.13e-30 | 1.39e-25 |
| Fibroblast | F2 RGS5+ Fib | ACTN4 | 1.534 | 0.579 | 0.321 | 4.5e-17 | 1.52e-12 |
| Fibroblast | F2 RGS5+ Fib | APOLD1 | 1.521 | 0.103 | 0.024 | 3.25e-07 | 0.0109 |
| Fibroblast | F2 RGS5+ Fib | MAP1B | 1.52 | 0.641 | 0.371 | 4.98e-19 | 1.68e-14 |
| Fibroblast | F2 RGS5+ Fib | RBPMS | 1.495 | 0.569 | 0.323 | 1.08e-16 | 3.63e-12 |
| Fibroblast | F2 RGS5+ Fib | TAGLN | 1.489 | 0.969 | 0.725 | 7.77e-37 | 2.62e-32 |
| Fibroblast | F2 RGS5+ Fib | SOD3 | 1.477 | 0.959 | 0.692 | 6.46e-41 | 2.18e-36 |
| Fibroblast | F2 RGS5+ Fib | PHLDA2 | 1.476 | 0.641 | 0.407 | 1.02e-14 | 3.42e-10 |
| Fibroblast | F2 RGS5+ Fib | SMTN | 1.476 | 0.4 | 0.218 | 2.57e-10 | 8.65e-06 |
| Fibroblast | F2 RGS5+ Fib | ARFGAP3 | 1.469 | 0.308 | 0.158 | 3.92e-08 | 0.00132 |
| Fibroblast | F2 RGS5+ Fib | SORBS3 | 1.468 | 0.385 | 0.251 | 4.41e-07 | 0.0149 |
| Fibroblast | F2 RGS5+ Fib | SMOC2 | 1.465 | 0.462 | 0.219 | 8.46e-15 | 2.85e-10 |
| Fibroblast | F2 RGS5+ Fib | MIR4435-2HG | 1.463 | 0.456 | 0.283 | 2.86e-09 | 9.62e-05 |
| Fibroblast | F2 RGS5+ Fib | CPE | 1.452 | 0.631 | 0.331 | 4.83e-17 | 1.63e-12 |
| Fibroblast | F2 RGS5+ Fib | KCNMB1 | 1.451 | 0.292 | 0.116 | 1.07e-10 | 3.6e-06 |
| Fibroblast | F2 RGS5+ Fib | EPS8 | 1.447 | 0.472 | 0.234 | 9.65e-15 | 3.25e-10 |
| Fibroblast | F2 RGS5+ Fib | CSRP1 | 1.442 | 0.636 | 0.425 | 2.08e-15 | 7.02e-11 |
| Fibroblast | F2 RGS5+ Fib | ABTB1 | 1.44 | 0.333 | 0.175 | 2.88e-08 | 0.000969 |
| Fibroblast | F2 RGS5+ Fib | SLC25A4 | 1.44 | 0.262 | 0.127 | 5.15e-07 | 0.0174 |
| Fibroblast | F2 RGS5+ Fib | GMFG | 1.439 | 0.277 | 0.126 | 2.64e-08 | 0.000891 |
| Fibroblast | F2 RGS5+ Fib | HAGH | 1.437 | 0.267 | 0.135 | 9.02e-07 | 0.0304 |
| Fibroblast | F2 RGS5+ Fib | RNF152 | 1.421 | 0.231 | 0.061 | 3.77e-13 | 1.27e-08 |
| Fibroblast | F2 RGS5+ Fib | PAWR | 1.42 | 0.344 | 0.181 | 9.07e-09 | 0.000306 |
| Fibroblast | F2 RGS5+ Fib | GAMT | 1.42 | 0.292 | 0.165 | 1.45e-06 | 0.0487 |
| Fibroblast | F2 RGS5+ Fib | NOL3 | 1.418 | 0.226 | 0.09 | 5.06e-08 | 0.0017 |
| Fibroblast | F2 RGS5+ Fib | PTGIR | 1.417 | 0.354 | 0.138 | 3.76e-13 | 1.27e-08 |
| Fibroblast | F2 RGS5+ Fib | MFGE8 | 1.411 | 0.877 | 0.696 | 7.860000000000001e-26 | 2.65e-21 |
| Fibroblast | F2 RGS5+ Fib | ARID5A | 1.405 | 0.405 | 0.19 | 1.72e-12 | 5.78e-08 |
| Fibroblast | F2 RGS5+ Fib | KCNS3 | 1.404 | 0.267 | 0.115 | 1.25e-08 | 0.00042 |
| Fibroblast | F2 RGS5+ Fib | SPARCL1 | 1.401 | 0.877 | 0.71 | 2.81e-21 | 9.46e-17 |
| Fibroblast | F2 RGS5+ Fib | TGFB1 | 1.387 | 0.385 | 0.235 | 4.95e-08 | 0.00167 |
| Fibroblast | F2 RGS5+ Fib | MIR22HG | 1.375 | 0.349 | 0.19 | 6.04e-08 | 0.00204 |
| Fibroblast | F2 RGS5+ Fib | HOXB2 | 1.355 | 0.374 | 0.208 | 1.77e-08 | 0.000597 |
| Fibroblast | F2 RGS5+ Fib | ARHGAP15 | 1.339 | 0.344 | 0.175 | 1.44e-08 | 0.000485 |
| Fibroblast | F2 RGS5+ Fib | PPP1R12A | 1.336 | 0.467 | 0.296 | 8.39e-10 | 2.83e-05 |
| Fibroblast | F2 RGS5+ Fib | LINC00152 | 1.331 | 0.533 | 0.382 | 2.26e-09 | 7.61e-05 |

| compartment | subcluster | gene | avg_log2FC | pct.1 | pct.2 | p_val | p_val_adj |
|---|---|---|---|---|---|---|---|
| Fibroblast | F2 RGS5+ Fib | UTRN | 1.324 | 0.297 | 0.156 | 4.66e-07 | 0.0157 |
| Fibroblast | F2 RGS5+ Fib | ROCK1 | 1.321 | 0.487 | 0.311 | 1.22e-10 | 4.11e-06 |
| Fibroblast | F2 RGS5+ Fib | RSU1 | 1.306 | 0.446 | 0.297 | 1.75e-07 | 0.00588 |
| Fibroblast | F2 RGS5+ Fib | S1PR3 | 1.302 | 0.21 | 0.073 | 7.57e-09 | 0.000255 |
| Fibroblast | F2 RGS5+ Fib | MSRB3 | 1.272 | 0.405 | 0.251 | 5.83e-08 | 0.00197 |
| Fibroblast | F2 RGS5+ Fib | AGT | 1.27 | 0.272 | 0.126 | 1.51e-07 | 0.00509 |
| Fibroblast | F2 RGS5+ Fib | ABI2 | 1.268 | 0.313 | 0.181 | 1.21e-06 | 0.0406 |
| Fibroblast | F2 RGS5+ Fib | SYNPO2 | 1.237 | 0.369 | 0.195 | 5.78e-09 | 0.000195 |
| Fibroblast | F2 RGS5+ Fib | SH3BGRL | 1.232 | 0.851 | 0.649 | 8.89e-27 | 3e-22 |
| Fibroblast | F2 RGS5+ Fib | CD46 | 1.225 | 0.344 | 0.194 | 2.86e-07 | 0.00965 |
| Fibroblast | F2 RGS5+ Fib | COX7A1 | 1.199 | 0.626 | 0.485 | 2.2e-11 | 7.43e-07 |
| Fibroblast | F2 RGS5+ Fib | TPM1 | 1.194 | 0.862 | 0.677 | 1.14e-20 | 3.86e-16 |
| Fibroblast | F2 RGS5+ Fib | TGFB1I1 | 1.185 | 0.615 | 0.432 | 2.12e-13 | 7.15e-09 |
| Fibroblast | F2 RGS5+ Fib | ID4 | 1.183 | 0.651 | 0.412 | 1.32e-13 | 4.44e-09 |
| Fibroblast | F2 RGS5+ Fib | ST6GALNAC6 | 1.183 | 0.379 | 0.243 | 6.56e-07 | 0.0221 |
| Fibroblast | F2 RGS5+ Fib | FADS3 | 1.174 | 0.421 | 0.224 | 3.83e-10 | 1.29e-05 |
| Fibroblast | F2 RGS5+ Fib | KANK2 | 1.17 | 0.415 | 0.263 | 5.61e-07 | 0.0189 |
| Fibroblast | F2 RGS5+ Fib | WDR1 | 1.168 | 0.492 | 0.368 | 1.01e-07 | 0.00342 |
| Fibroblast | F2 RGS5+ Fib | AXL | 1.165 | 0.441 | 0.304 | 8.53e-07 | 0.0287 |
| Fibroblast | F2 RGS5+ Fib | LRRFIP1 | 1.133 | 0.415 | 0.262 | 1.42e-07 | 0.0048 |
| Fibroblast | F2 RGS5+ Fib | PKIG | 1.128 | 0.656 | 0.402 | 1.84e-17 | 6.19e-13 |
| Fibroblast | F2 RGS5+ Fib | C1orf132 | 1.123 | 0.174 | 0.065 | 7.75e-07 | 0.0261 |
| Fibroblast | F2 RGS5+ Fib | ATP1B3 | 1.12 | 0.503 | 0.344 | 4.32e-09 | 0.000146 |
| Fibroblast | F2 RGS5+ Fib | SEPT7 | 1.116 | 0.8 | 0.672 | 5.27e-15 | 1.78e-10 |
| Fibroblast | F2 RGS5+ Fib | STOM | 1.099 | 0.615 | 0.436 | 1.53e-10 | 5.15e-06 |
| Fibroblast | F2 RGS5+ Fib | CALD1 | 1.079 | 1.0 | 0.947 | 2.49e-36 | 8.390000000000001e-32 |
| Fibroblast | F2 RGS5+ Fib | FLNA | 1.078 | 0.677 | 0.533 | 6.12e-10 | 2.06e-05 |
| Fibroblast | F2 RGS5+ Fib | ARPC5 | 1.065 | 0.631 | 0.501 | 4.64e-11 | 1.56e-06 |
| Fibroblast | F2 RGS5+ Fib | FBLIM1 | 1.062 | 0.369 | 0.226 | 7.74e-07 | 0.0261 |
| Fibroblast | F2 RGS5+ Fib | RNF5 | 1.053 | 0.4 | 0.255 | 3.17e-07 | 0.0107 |
| Fibroblast | F2 RGS5+ Fib | CYSTM1 | 1.037 | 0.595 | 0.43 | 2.3e-10 | 7.74e-06 |
| Fibroblast | F2 RGS5+ Fib | PRKCDBP | 1.018 | 0.769 | 0.628 | 2.79e-13 | 9.4e-09 |
| Fibroblast | F2 RGS5+ Fib | CALM2 | 1.011 | 0.954 | 0.885 | 5.89e-33 | 1.98e-28 |
| Fibroblast | F2 RGS5+ Fib | ZFHX3 | 1.008 | 0.369 | 0.204 | 6.44e-08 | 0.00217 |
| Fibroblast | F2 RGS5+ Fib | MYL6 | 0.982 | 0.985 | 0.958 | 1.54e-34 | 5.189999999999999e-30 |
| Fibroblast | F2 RGS5+ Fib | NR2F2 | 0.981 | 0.605 | 0.425 | 6.25e-11 | 2.11e-06 |
| Fibroblast | F2 RGS5+ Fib | PTMA | 0.979 | 1.0 | 1.0 | 3.38e-48 | 1.14e-43 |
| Fibroblast | F2 RGS5+ Fib | PMEPA1 | 0.972 | 0.338 | 0.194 | 1.27e-06 | 0.043 |
| Fibroblast | F2 RGS5+ Fib | CD151 | 0.951 | 0.718 | 0.6 | 1.69e-10 | 5.7e-06 |
| Fibroblast | F2 RGS5+ Fib | CRISPLD2 | 0.947 | 0.641 | 0.523 | 3.75e-08 | 0.00126 |
| Fibroblast | F2 RGS5+ Fib | ADAMTS1 | 0.933 | 0.615 | 0.351 | 6.51e-15 | 2.19e-10 |
| Fibroblast | F2 RGS5+ Fib | IGFBP7 | 0.925 | 1.0 | 0.97 | 6.27e-36 | 2.11e-31 |
| Fibroblast | F2 RGS5+ Fib | MRPL33 | 0.925 | 0.605 | 0.431 | 1.27e-10 | 4.28e-06 |
| Fibroblast | F2 RGS5+ Fib | PLS3 | 0.921 | 0.467 | 0.331 | 1.4e-06 | 0.0472 |
| Fibroblast | F2 RGS5+ Fib | NUDT4 | 0.914 | 0.646 | 0.408 | 1.31e-14 | 4.41e-10 |
| Fibroblast | F2 RGS5+ Fib | USMG5 | 0.912 | 0.774 | 0.642 | 1.3e-14 | 4.37e-10 |
| Fibroblast | F2 RGS5+ Fib | HCFC1R1 | 0.908 | 0.692 | 0.503 | 7.01e-12 | 2.36e-07 |
| Fibroblast | F2 RGS5+ Fib | HMGN3 | 0.891 | 0.579 | 0.444 | 1.48e-07 | 0.00498 |
| Fibroblast | F2 RGS5+ Fib | PTRF | 0.888 | 0.805 | 0.663 | 2.72e-14 | 9.15e-10 |
| Fibroblast | F2 RGS5+ Fib | NDUFA6 | 0.885 | 0.513 | 0.384 | 4.71e-07 | 0.0159 |
| Fibroblast | F2 RGS5+ Fib | COX7A2 | 0.879 | 0.887 | 0.757 | 3.92e-20 | 1.32e-15 |
| Fibroblast | F2 RGS5+ Fib | CRYAB | 0.857 | 0.687 | 0.438 | 3.49e-15 | 1.18e-10 |
| Fibroblast | F2 RGS5+ Fib | PHLDA1 | 0.85 | 0.774 | 0.604 | 3.62e-13 | 1.22e-08 |
| Fibroblast | F2 RGS5+ Fib | MGST3 | 0.846 | 0.785 | 0.717 | 3.02e-08 | 0.00102 |
| Fibroblast | F2 RGS5+ Fib | CYCS | 0.844 | 0.738 | 0.571 | 1.07e-12 | 3.59e-08 |
| Fibroblast | F2 RGS5+ Fib | CAMTA1 | 0.826 | 0.585 | 0.411 | 8.34e-10 | 2.81e-05 |
| Fibroblast | F2 RGS5+ Fib | DEK | 0.813 | 0.595 | 0.454 | 8.28e-08 | 0.00279 |
| Fibroblast | F2 RGS5+ Fib | PDLIM1 | 0.807 | 0.579 | 0.466 | 1.25e-06 | 0.042 |
| Fibroblast | F2 RGS5+ Fib | ITGB1 | 0.8 | 0.795 | 0.662 | 9.28e-10 | 3.13e-05 |
| Fibroblast | F2 RGS5+ Fib | GADD45GIP1 | 0.788 | 0.636 | 0.545 | 3.26e-07 | 0.011 |
| Fibroblast | F2 RGS5+ Fib | ACTN1 | 0.774 | 0.631 | 0.477 | 1.71e-07 | 0.00576 |
| Fibroblast | F2 RGS5+ Fib | COX7B | 0.771 | 0.759 | 0.632 | 8.01e-12 | 2.7e-07 |
| Fibroblast | F2 RGS5+ Fib | ID3 | 0.749 | 0.933 | 0.787 | 7.77e-19 | 2.62e-14 |
| Fibroblast | F2 RGS5+ Fib | RGS16 | 0.741 | 0.569 | 0.369 | 5.07e-10 | 1.71e-05 |

| compartment | subcluster | gene | avg_log2FC | pct.1 | pct.2 | p_val | p_val_adj |
|---|---|---|---|---|---|---|---|
| Fibroblast | F2 RGS5+ Fib | CAP1 | 0.737 | 0.595 | 0.472 | 2.88e-07 | 0.00972 |
| Fibroblast | F2 RGS5+ Fib | PFN1 | 0.727 | 0.974 | 0.889 | 1.58e-19 | 5.33e-15 |
| Fibroblast | F2 RGS5+ Fib | ANAPC16 | 0.723 | 0.677 | 0.587 | 4.43e-07 | 0.0149 |
| Fibroblast | F2 RGS5+ Fib | SBDS | 0.71 | 0.728 | 0.642 | 4.8e-09 | 0.000162 |
| Fibroblast | F2 RGS5+ Fib | C11orf96 | 0.7 | 0.836 | 0.713 | 5.12e-12 | 1.72e-07 |
| Fibroblast | F2 RGS5+ Fib | PDLIM3 | 0.674 | 0.569 | 0.371 | 1.01e-07 | 0.00339 |
| Fibroblast | F2 RGS5+ Fib | COX5B | 0.672 | 0.81 | 0.762 | 1.41e-10 | 4.75e-06 |
| Fibroblast | F2 RGS5+ Fib | YWHAZ | 0.668 | 0.677 | 0.581 | 1.25e-07 | 0.0042 |
| Fibroblast | F2 RGS5+ Fib | GPX3 | 0.662 | 0.59 | 0.378 | 1.27e-08 | 0.000429 |
| Fibroblast | F2 RGS5+ Fib | COX8A | 0.655 | 0.779 | 0.667 | 7.08e-10 | 2.39e-05 |
| Fibroblast | F2 RGS5+ Fib | IFITM2 | 0.647 | 0.944 | 0.831 | 2.11e-14 | 7.12e-10 |
| Fibroblast | F2 RGS5+ Fib | ATP6V1G1 | 0.637 | 0.744 | 0.668 | 7.51e-07 | 0.0253 |
| Fibroblast | F2 RGS5+ Fib | NDUFA1 | 0.636 | 0.759 | 0.634 | 2.33e-09 | 7.84e-05 |
| Fibroblast | F2 RGS5+ Fib | VAPA | 0.626 | 0.713 | 0.632 | 4.74e-07 | 0.016 |
| Fibroblast | F2 RGS5+ Fib | NDUFB7 | 0.625 | 0.744 | 0.613 | 2.24e-08 | 0.000754 |
| Fibroblast | F2 RGS5+ Fib | NDUFB1 | 0.62 | 0.708 | 0.613 | 1.33e-06 | 0.0448 |
| Fibroblast | F2 RGS5+ Fib | TSC22D1 | 0.617 | 0.897 | 0.79 | 8.61e-12 | 2.9e-07 |
| Fibroblast | F2 RGS5+ Fib | YBX1 | 0.61 | 0.923 | 0.823 | 3.79e-13 | 1.28e-08 |
| Fibroblast | F2 RGS5+ Fib | VAMP2 | 0.598 | 0.744 | 0.62 | 2.55e-08 | 0.000858 |
| Fibroblast | F2 RGS5+ Fib | NDUFB8 | 0.594 | 0.713 | 0.607 | 2.64e-07 | 0.00889 |
| Fibroblast | F2 RGS5+ Fib | PLAC9 | 0.575 | 0.759 | 0.602 | 4.23e-07 | 0.0142 |
| Fibroblast | F2 RGS5+ Fib | H3F3A | 0.565 | 0.964 | 0.92 | 6.27e-17 | 2.11e-12 |
| Fibroblast | F2 RGS5+ Fib | GNAS | 0.564 | 0.703 | 0.598 | 3.43e-07 | 0.0116 |
| Fibroblast | F2 RGS5+ Fib | ATP5G2 | 0.558 | 0.877 | 0.796 | 3.94e-11 | 1.33e-06 |
| Fibroblast | F2 RGS5+ Fib | ARPC2 | 0.558 | 0.795 | 0.695 | 3.11e-07 | 0.0105 |
| Fibroblast | F2 RGS5+ Fib | COX6A1 | 0.557 | 0.851 | 0.763 | 4.51e-09 | 0.000152 |
| Fibroblast | F2 RGS5+ Fib | MALAT1 | 0.551 | 1.0 | 0.998 | 5.570000000000001e-27 | 1.88e-22 |
| Fibroblast | F2 RGS5+ Fib | HSPB1 | 0.539 | 0.99 | 0.958 | 2.09e-19 | 7.04e-15 |
| Fibroblast | F2 RGS5+ Fib | ATP5I | 0.527 | 0.8 | 0.707 | 2.87e-07 | 0.00966 |
| Fibroblast | F2 RGS5+ Fib | GAPDH | 0.485 | 0.974 | 0.937 | 3.65e-12 | 1.23e-07 |
| Fibroblast | F2 RGS5+ Fib | ACTB | 0.472 | 0.995 | 0.986 | 1.78e-11 | 5.99e-07 |
| Fibroblast | F2 RGS5+ Fib | ALDOA | 0.469 | 0.846 | 0.783 | 8.27e-07 | 0.0279 |
| Fibroblast | F2 RGS5+ Fib | PPIA | 0.44 | 0.897 | 0.83 | 9.02e-08 | 0.00304 |
| Fibroblast | F2 RGS5+ Fib | HMGB1 | 0.439 | 0.938 | 0.884 | 5.08e-09 | 0.000171 |
| Fibroblast | F2 RGS5+ Fib | PTMS | 0.437 | 0.821 | 0.759 | 2.17e-07 | 0.00731 |
| Fibroblast | F2 RGS5+ Fib | CHCHD2 | 0.435 | 0.882 | 0.81 | 3.27e-07 | 0.011 |
| Fibroblast | F2 RGS5+ Fib | SKP1 | 0.427 | 0.913 | 0.85 | 1.19e-07 | 0.004 |
| Fibroblast | F2 RGS5+ Fib | COX6C | 0.426 | 0.856 | 0.778 | 4.45e-07 | 0.015 |
| Fibroblast | F2 RGS5+ Fib | LMNA | 0.422 | 0.903 | 0.874 | 7.95e-08 | 0.00268 |
| Fibroblast | F2 RGS5+ Fib | UQCRB | 0.419 | 0.908 | 0.865 | 1.08e-07 | 0.00363 |
| Fibroblast | F2 RGS5+ Fib | A2M | 0.405 | 0.687 | 0.469 | 2.29e-07 | 0.0077 |
| Fibroblast | F2 RGS5+ Fib | CFL1 | 0.38 | 0.938 | 0.886 | 1.35e-06 | 0.0456 |
| Fibroblast | F2 RGS5+ Fib | RPS7 | 0.373 | 0.99 | 0.957 | 2.99e-10 | 1.01e-05 |
| Fibroblast | F2 RGS5+ Fib | BTG1 | 0.347 | 0.877 | 0.843 | 3.77e-08 | 0.00127 |
| Fibroblast | F2 RGS5+ Fib | MT-CO1 | 0.309 | 1.0 | 0.999 | 2.16e-07 | 0.00729 |
| Fibroblast | F2 RGS5+ Fib | MT-CYB | 0.29 | 1.0 | 0.994 | 2.17e-07 | 0.00731 |
| Fibroblast | F2 RGS5+ Fib | MT-CO3 | 0.279 | 1.0 | 0.996 | 2.92e-07 | 0.00985 |
| Fibroblast | F3 COL11A1+ mCAF | IGFL2 | 6.73 | 0.303 | 0.007 | 4.02e-53 | 1.35e-48 |
| Fibroblast | F3 COL11A1+ mCAF | CALB2 | 6.071 | 0.108 | 0.001 | 1.24e-20 | 4.19e-16 |
| Fibroblast | F3 COL11A1+ mCAF | C5orf46 | 5.284 | 0.151 | 0.007 | 2.15e-23 | 7.23e-19 |
| Fibroblast | F3 COL11A1+ mCAF | STRA6 | 5.164 | 0.205 | 0.005 | 1.09e-35 | 3.66e-31 |
| Fibroblast | F3 COL11A1+ mCAF | MMP7 | 4.74 | 0.168 | 0.018 | 2.17e-19 | 7.32e-15 |
| Fibroblast | F3 COL11A1+ mCAF | GJB2 | 4.372 | 0.422 | 0.031 | 8.62e-58 | 2.9e-53 |
| Fibroblast | F3 COL11A1+ mCAF | CST1 | 4.364 | 0.189 | 0.015 | 2.29e-24 | 7.71e-20 |
| Fibroblast | F3 COL11A1+ mCAF | COL11A1 | 4.32 | 0.416 | 0.025 | 6.71e-61 | 2.26e-56 |
| Fibroblast | F3 COL11A1+ mCAF | SDC1 | 4.202 | 0.292 | 0.02 | 8.02e-40 | 2.7e-35 |
| Fibroblast | F3 COL11A1+ mCAF | EPYC | 4.195 | 0.135 | 0.015 | 3.84e-15 | 1.29e-10 |
| Fibroblast | F3 COL11A1+ mCAF | MMP11 | 4.186 | 0.584 | 0.131 | 1.24e-48 | 4.19e-44 |
| Fibroblast | F3 COL11A1+ mCAF | LRRC15 | 4.155 | 0.205 | 0.012 | 1.3e-29 | 4.37e-25 |
| Fibroblast | F3 COL11A1+ mCAF | PLPP4 | 4.009 | 0.33 | 0.024 | 1.1e-44 | 3.71e-40 |
| Fibroblast | F3 COL11A1+ mCAF | SLC12A8 | 3.852 | 0.378 | 0.031 | 2.56e-49 | 8.64e-45 |
| Fibroblast | F3 COL11A1+ mCAF | RP11-400N13.3 | 3.681 | 0.108 | 0.008 | 1.44e-14 | 4.84e-10 |
| Fibroblast | F3 COL11A1+ mCAF | CXCL13 | 3.504 | 0.151 | 0.033 | 1.09e-10 | 3.66e-06 |
| Fibroblast | F3 COL11A1+ mCAF | POSTN | 3.496 | 0.697 | 0.133 | 5.489999999999999e-67 | 1.85e-62 |

| compartment | subcluster | gene | avg_log2FC | pct.1 | pct.2 | p_val | p_val_adj |
|---|---|---|---|---|---|---|---|
| Fibroblast | F3 COL11A1+ mCAF | DSG2 | 3.486 | 0.13 | 0.008 | 2.47e-18 | 8.33e-14 |
| Fibroblast | F3 COL11A1+ mCAF | S100A2 | 3.48 | 0.119 | 0.024 | 2.18e-09 | 7.33e-05 |
| Fibroblast | F3 COL11A1+ mCAF | COL10A1 | 3.384 | 0.535 | 0.088 | 5.63e-52 | 1.9e-47 |
| Fibroblast | F3 COL11A1+ mCAF | SERINC2 | 3.294 | 0.151 | 0.014 | 2.58e-18 | 8.68e-14 |
| Fibroblast | F3 COL11A1+ mCAF | DMKN | 3.29 | 0.216 | 0.028 | 1.06e-21 | 3.56e-17 |
| Fibroblast | F3 COL11A1+ mCAF | CA12 | 3.261 | 0.168 | 0.037 | 5.04e-12 | 1.7e-07 |
| Fibroblast | F3 COL11A1+ mCAF | TMEM158 | 3.139 | 0.151 | 0.019 | 6.63e-16 | 2.23e-11 |
| Fibroblast | F3 COL11A1+ mCAF | NFE2L3 | 3.127 | 0.157 | 0.013 | 6.99e-20 | 2.36e-15 |
| Fibroblast | F3 COL11A1+ mCAF | PIEZO2 | 3.119 | 0.124 | 0.013 | 1.89e-14 | 6.38e-10 |
| Fibroblast | F3 COL11A1+ mCAF | NTM | 3.105 | 0.308 | 0.049 | 2.56e-28 | 8.62e-24 |
| Fibroblast | F3 COL11A1+ mCAF | BCAT1 | 3.103 | 0.222 | 0.031 | 7.76e-22 | 2.62e-17 |
| Fibroblast | F3 COL11A1+ mCAF | WNT2 | 3.088 | 0.308 | 0.033 | 4.08e-35 | 1.37e-30 |
| Fibroblast | F3 COL11A1+ mCAF | RBP4 | 3.036 | 0.168 | 0.024 | 1.93e-16 | 6.5e-12 |
| Fibroblast | F3 COL11A1+ mCAF | MIAT | 3.024 | 0.103 | 0.013 | 4.88e-11 | 1.64e-06 |
| Fibroblast | F3 COL11A1+ mCAF | LOXL2 | 2.955 | 0.405 | 0.05 | 6.49e-43 | 2.19e-38 |
| Fibroblast | F3 COL11A1+ mCAF | LEF1 | 2.95 | 0.205 | 0.025 | 1.7e-21 | 5.71e-17 |
| Fibroblast | F3 COL11A1+ mCAF | FAM19A5 | 2.939 | 0.108 | 0.018 | 6.03e-10 | 2.03e-05 |
| Fibroblast | F3 COL11A1+ mCAF | GLRB | 2.898 | 0.114 | 0.009 | 1.4e-14 | 4.73e-10 |
| Fibroblast | F3 COL11A1+ mCAF | SEZ6L2 | 2.891 | 0.151 | 0.021 | 1.03e-14 | 3.48e-10 |
| Fibroblast | F3 COL11A1+ mCAF | IGFBP3 | 2.885 | 0.524 | 0.115 | 5.56e-39 | 1.87e-34 |
| Fibroblast | F3 COL11A1+ mCAF | WNT5A | 2.884 | 0.168 | 0.028 | 2.25e-14 | 7.59e-10 |
| Fibroblast | F3 COL11A1+ mCAF | PLAU | 2.846 | 0.584 | 0.133 | 4.6e-44 | 1.55e-39 |
| Fibroblast | F3 COL11A1+ mCAF | MEST | 2.79 | 0.211 | 0.046 | 9.88e-15 | 3.33e-10 |
| Fibroblast | F3 COL11A1+ mCAF | PODNL1 | 2.765 | 0.368 | 0.064 | 1.89e-31 | 6.36e-27 |
| Fibroblast | F3 COL11A1+ mCAF | FN1 | 2.753 | 0.849 | 0.44 | 1.05e-41 | 3.54e-37 |
| Fibroblast | F3 COL11A1+ mCAF | SLC16A3 | 2.731 | 0.303 | 0.052 | 1.43e-25 | 4.82e-21 |
| Fibroblast | F3 COL11A1+ mCAF | WISP1 | 2.71 | 0.173 | 0.02 | 1.63e-18 | 5.51e-14 |
| Fibroblast | F3 COL11A1+ mCAF | PDGFC | 2.686 | 0.405 | 0.053 | 2.65e-40 | 8.92e-36 |
| Fibroblast | F3 COL11A1+ mCAF | CTHRC1 | 2.673 | 0.903 | 0.334 | 1.23e-63 | 4.16e-59 |
| Fibroblast | F3 COL11A1+ mCAF | GJA1 | 2.665 | 0.389 | 0.075 | 3.9e-31 | 1.31e-26 |
| Fibroblast | F3 COL11A1+ mCAF | SULF1 | 2.65 | 0.724 | 0.213 | 2.24e-50 | 7.549999999999999e-46 |
| Fibroblast | F3 COL11A1+ mCAF | SLC38A5 | 2.596 | 0.119 | 0.018 | 2.44e-11 | 8.22e-07 |
| Fibroblast | F3 COL11A1+ mCAF | LIMK2 | 2.58 | 0.13 | 0.022 | 5.11e-11 | 1.72e-06 |
| Fibroblast | F3 COL11A1+ mCAF | ADAM12 | 2.573 | 0.368 | 0.07 | 4.23e-29 | 1.43e-24 |
| Fibroblast | F3 COL11A1+ mCAF | TNFSF4 | 2.559 | 0.114 | 0.014 | 5.35e-12 | 1.8e-07 |
| Fibroblast | F3 COL11A1+ mCAF | ADAMTS12 | 2.53 | 0.303 | 0.053 | 2.86e-25 | 9.650000000000001e-21 |
| Fibroblast | F3 COL11A1+ mCAF | INHBA | 2.524 | 0.686 | 0.155 | 4.17e-56 | 1.4e-51 |
| Fibroblast | F3 COL11A1+ mCAF | ARL4C | 2.511 | 0.395 | 0.07 | 4.75e-33 | 1.6e-28 |
| Fibroblast | F3 COL11A1+ mCAF | TNFSF11 | 2.506 | 0.108 | 0.015 | 8.59e-11 | 2.89e-06 |
| Fibroblast | F3 COL11A1+ mCAF | MXRA5 | 2.447 | 0.622 | 0.156 | 4.3e-43 | 1.45e-38 |
| Fibroblast | F3 COL11A1+ mCAF | CDH13 | 2.428 | 0.108 | 0.024 | 7.07e-08 | 0.00238 |
| Fibroblast | F3 COL11A1+ mCAF | CLIC3 | 2.411 | 0.151 | 0.034 | 2.69e-10 | 9.07e-06 |
| Fibroblast | F3 COL11A1+ mCAF | SHISA2 | 2.362 | 0.124 | 0.021 | 1.16e-10 | 3.9e-06 |
| Fibroblast | F3 COL11A1+ mCAF | FLNB | 2.354 | 0.184 | 0.037 | 6.55e-14 | 2.21e-09 |
| Fibroblast | F3 COL11A1+ mCAF | CYB561 | 2.346 | 0.178 | 0.044 | 2.8e-11 | 9.45e-07 |
| Fibroblast | F3 COL11A1+ mCAF | ANKH | 2.342 | 0.411 | 0.099 | 9.030000000000001e-27 | 3.04e-22 |
| Fibroblast | F3 COL11A1+ mCAF | SPOCK1 | 2.32 | 0.303 | 0.073 | 3.02e-19 | 1.02e-14 |
| Fibroblast | F3 COL11A1+ mCAF | IFIH1 | 2.286 | 0.108 | 0.021 | 1.4e-08 | 0.000471 |
| Fibroblast | F3 COL11A1+ mCAF | KIAA1217 | 2.266 | 0.308 | 0.054 | 5.84e-25 | 1.97e-20 |
| Fibroblast | F3 COL11A1+ mCAF | THBS2 | 2.259 | 0.751 | 0.275 | 7.08e-44 | 2.39e-39 |
| Fibroblast | F3 COL11A1+ mCAF | COL7A1 | 2.259 | 0.146 | 0.015 | 1.45e-16 | 4.89e-12 |
| Fibroblast | F3 COL11A1+ mCAF | TMEM132A | 2.259 | 0.124 | 0.028 | 1.4e-08 | 0.000472 |
| Fibroblast | F3 COL11A1+ mCAF | IL27RA | 2.243 | 0.135 | 0.021 | 3.17e-12 | 1.07e-07 |
| Fibroblast | F3 COL11A1+ mCAF | RUNX2 | 2.235 | 0.276 | 0.062 | 1e-18 | 3.38e-14 |
| Fibroblast | F3 COL11A1+ mCAF | CHST15 | 2.229 | 0.13 | 0.03 | 6.38e-09 | 0.000215 |
| Fibroblast | F3 COL11A1+ mCAF | COL12A1 | 2.221 | 0.514 | 0.185 | 8.509999999999999e-23 | 2.87e-18 |
| Fibroblast | F3 COL11A1+ mCAF | RGS17 | 2.212 | 0.13 | 0.022 | 5.59e-11 | 1.88e-06 |
| Fibroblast | F3 COL11A1+ mCAF | HOPX | 2.188 | 0.459 | 0.142 | 3.949999999999999e-24 | 1.33e-19 |
| Fibroblast | F3 COL11A1+ mCAF | C1QTNF6 | 2.166 | 0.276 | 0.058 | 1.04e-19 | 3.51e-15 |
| Fibroblast | F3 COL11A1+ mCAF | NKD2 | 2.162 | 0.146 | 0.033 | 7.52e-10 | 2.53e-05 |
| Fibroblast | F3 COL11A1+ mCAF | C16orf95 | 2.157 | 0.13 | 0.024 | 1.27e-10 | 4.29e-06 |
| Fibroblast | F3 COL11A1+ mCAF | COL8A2 | 2.129 | 0.108 | 0.022 | 4.37e-08 | 0.00147 |
| Fibroblast | F3 COL11A1+ mCAF | IGF2BP2 | 2.117 | 0.168 | 0.031 | 2.29e-13 | 7.72e-09 |
| Fibroblast | F3 COL11A1+ mCAF | KLRD1 | 2.088 | 0.124 | 0.022 | 3.03e-10 | 1.02e-05 |

| compartment | subcluster | gene | avg_log2FC | pct.1 | pct.2 | p_val | p_val_adj |
|---|---|---|---|---|---|---|---|
| Fibroblast | F3 COL11A1+ mCAF | CDH2 | 2.077 | 0.114 | 0.014 | 6.14e-12 | 2.07e-07 |
| Fibroblast | F3 COL11A1+ mCAF | ADAM19 | 2.047 | 0.232 | 0.041 | 1.59e-18 | 5.36e-14 |
| Fibroblast | F3 COL11A1+ mCAF | TWIST1 | 2.043 | 0.243 | 0.064 | 6.82e-14 | 2.3e-09 |
| Fibroblast | F3 COL11A1+ mCAF | COL5A2 | 2.034 | 0.8 | 0.327 | 4.829999999999999e-42 | 1.63e-37 |
| Fibroblast | F3 COL11A1+ mCAF | PYCR1 | 2.031 | 0.308 | 0.077 | 1.77e-18 | 5.95e-14 |
| Fibroblast | F3 COL11A1+ mCAF | MFAP2 | 2.015 | 0.692 | 0.212 | 6.27e-40 | 2.11e-35 |
| Fibroblast | F3 COL11A1+ mCAF | MATN3 | 2.015 | 0.243 | 0.037 | 5.53e-22 | 1.86e-17 |
| Fibroblast | F3 COL11A1+ mCAF | APBA2 | 2.014 | 0.108 | 0.02 | 6.63e-09 | 0.000223 |
| Fibroblast | F3 COL11A1+ mCAF | TPST2 | 1.998 | 0.297 | 0.062 | 8.930000000000001e-21 | 3.01e-16 |
| Fibroblast | F3 COL11A1+ mCAF | PDPN | 1.994 | 0.649 | 0.211 | 2.08e-35 | 7.02e-31 |
| Fibroblast | F3 COL11A1+ mCAF | C15orf48 | 1.991 | 0.238 | 0.069 | 2.98e-12 | 1.01e-07 |
| Fibroblast | F3 COL11A1+ mCAF | HAGHL | 1.991 | 0.135 | 0.031 | 3.65e-09 | 0.000123 |
| Fibroblast | F3 COL11A1+ mCAF | BHLHE41 | 1.982 | 0.324 | 0.079 | 1.11e-19 | 3.73e-15 |
| Fibroblast | F3 COL11A1+ mCAF | LOXL1 | 1.973 | 0.595 | 0.211 | 1.51e-28 | 5.1e-24 |
| Fibroblast | F3 COL11A1+ mCAF | DERL3 | 1.966 | 0.13 | 0.037 | 4.02e-07 | 0.0136 |
| Fibroblast | F3 COL11A1+ mCAF | COL5A1 | 1.956 | 0.8 | 0.276 | 9.9e-45 | 3.34e-40 |
| Fibroblast | F3 COL11A1+ mCAF | LMO7 | 1.944 | 0.259 | 0.058 | 7.04e-17 | 2.37e-12 |
| Fibroblast | F3 COL11A1+ mCAF | LOX | 1.917 | 0.292 | 0.104 | 1.77e-11 | 5.96e-07 |
| Fibroblast | F3 COL11A1+ mCAF | ETV1 | 1.914 | 0.216 | 0.047 | 2.24e-14 | 7.55e-10 |
| Fibroblast | F3 COL11A1+ mCAF | ANTXR1 | 1.89 | 0.708 | 0.263 | 4.72e-34 | 1.59e-29 |
| Fibroblast | F3 COL11A1+ mCAF | SMCO4 | 1.887 | 0.281 | 0.059 | 1.1e-19 | 3.69e-15 |
| Fibroblast | F3 COL11A1+ mCAF | HTRA3 | 1.886 | 0.649 | 0.238 | 1.65e-28 | 5.57e-24 |
| Fibroblast | F3 COL11A1+ mCAF | COL1A1 | 1.877 | 0.995 | 0.766 | 3.1e-44 | 1.04e-39 |
| Fibroblast | F3 COL11A1+ mCAF | TSPAN5 | 1.87 | 0.162 | 0.03 | 6.86e-13 | 2.31e-08 |
| Fibroblast | F3 COL11A1+ mCAF | CNIH3 | 1.861 | 0.124 | 0.026 | 3.4e-09 | 0.000115 |
| Fibroblast | F3 COL11A1+ mCAF | SLC6A6 | 1.858 | 0.238 | 0.06 | 7.91e-14 | 2.67e-09 |
| Fibroblast | F3 COL11A1+ mCAF | RAI14 | 1.855 | 0.308 | 0.07 | 3.69e-20 | 1.24e-15 |
| Fibroblast | F3 COL11A1+ mCAF | SUGCT | 1.853 | 0.135 | 0.033 | 1.82e-08 | 0.000615 |
| Fibroblast | F3 COL11A1+ mCAF | FGD6 | 1.848 | 0.108 | 0.021 | 1.99e-08 | 0.000671 |
| Fibroblast | F3 COL11A1+ mCAF | GPC1 | 1.839 | 0.416 | 0.103 | 3.06e-25 | 1.03e-20 |
| Fibroblast | F3 COL11A1+ mCAF | CHST11 | 1.8 | 0.232 | 0.058 | 5.81e-14 | 1.96e-09 |
| Fibroblast | F3 COL11A1+ mCAF | P3H4 | 1.799 | 0.378 | 0.107 | 4.81e-20 | 1.62e-15 |
| Fibroblast | F3 COL11A1+ mCAF | MMP14 | 1.797 | 0.773 | 0.369 | 9.9e-34 | 3.33e-29 |
| Fibroblast | F3 COL11A1+ mCAF | NXN | 1.794 | 0.238 | 0.058 | 2.76e-14 | 9.31e-10 |
| Fibroblast | F3 COL11A1+ mCAF | PRDM1 | 1.786 | 0.146 | 0.045 | 3.09e-07 | 0.0104 |
| Fibroblast | F3 COL11A1+ mCAF | P4HA3 | 1.785 | 0.216 | 0.064 | 7.68e-11 | 2.59e-06 |
| Fibroblast | F3 COL11A1+ mCAF | CDH11 | 1.779 | 0.681 | 0.236 | 1.34e-33 | 4.51e-29 |
| Fibroblast | F3 COL11A1+ mCAF | HMGB3 | 1.773 | 0.124 | 0.032 | 1.63e-07 | 0.00548 |
| Fibroblast | F3 COL11A1+ mCAF | CDC42EP3 | 1.769 | 0.335 | 0.084 | 1.38e-19 | 4.66e-15 |
| Fibroblast | F3 COL11A1+ mCAF | NUP50 | 1.766 | 0.162 | 0.043 | 2.76e-09 | 9.3e-05 |
| Fibroblast | F3 COL11A1+ mCAF | IKBKE | 1.762 | 0.103 | 0.025 | 8.21e-07 | 0.0277 |
| Fibroblast | F3 COL11A1+ mCAF | DIO2 | 1.76 | 0.238 | 0.05 | 8.82e-17 | 2.97e-12 |
| Fibroblast | F3 COL11A1+ mCAF | GPR68 | 1.751 | 0.195 | 0.047 | 5.63e-12 | 1.9e-07 |
| Fibroblast | F3 COL11A1+ mCAF | CMTM3 | 1.75 | 0.735 | 0.253 | 7.93e-39 | 2.67e-34 |
| Fibroblast | F3 COL11A1+ mCAF | UBTD1 | 1.713 | 0.281 | 0.071 | 3.93e-16 | 1.33e-11 |
| Fibroblast | F3 COL11A1+ mCAF | FNDC1 | 1.711 | 0.368 | 0.072 | 6.22e-27 | 2.1e-22 |
| Fibroblast | F3 COL11A1+ mCAF | CTSB | 1.71 | 0.897 | 0.484 | 1.41e-36 | 4.73e-32 |
| Fibroblast | F3 COL11A1+ mCAF | MFSD5 | 1.701 | 0.124 | 0.028 | 2.42e-08 | 0.000815 |
| Fibroblast | F3 COL11A1+ mCAF | SYNDIG1 | 1.701 | 0.114 | 0.031 | 1.21e-06 | 0.0408 |
| Fibroblast | F3 COL11A1+ mCAF | TSKU | 1.7 | 0.2 | 0.06 | 1.3e-09 | 4.38e-05 |
| Fibroblast | F3 COL11A1+ mCAF | EVA1A | 1.698 | 0.227 | 0.063 | 5.84e-12 | 1.97e-07 |
| Fibroblast | F3 COL11A1+ mCAF | TSHZ3 | 1.697 | 0.119 | 0.027 | 5.07e-08 | 0.00171 |
| Fibroblast | F3 COL11A1+ mCAF | ZDHHC20 | 1.692 | 0.189 | 0.047 | 2.88e-11 | 9.7e-07 |
| Fibroblast | F3 COL11A1+ mCAF | CCL26 | 1.672 | 0.173 | 0.058 | 1.35e-07 | 0.00456 |
| Fibroblast | F3 COL11A1+ mCAF | ISLR | 1.671 | 0.67 | 0.305 | 2.64e-23 | 8.9e-19 |
| Fibroblast | F3 COL11A1+ mCAF | COL8A1 | 1.671 | 0.546 | 0.224 | 7.840000000000001e-21 | 2.64e-16 |
| Fibroblast | F3 COL11A1+ mCAF | COL3A1 | 1.662 | 0.978 | 0.649 | 6.48e-44 | 2.18e-39 |
| Fibroblast | F3 COL11A1+ mCAF | ZBED1 | 1.657 | 0.341 | 0.097 | 3.44e-17 | 1.16e-12 |
| Fibroblast | F3 COL11A1+ mCAF | RARB | 1.654 | 0.119 | 0.027 | 4.79e-08 | 0.00162 |
| Fibroblast | F3 COL11A1+ mCAF | ITGA11 | 1.653 | 0.243 | 0.056 | 1e-15 | 3.37e-11 |
| Fibroblast | F3 COL11A1+ mCAF | TTYH3 | 1.649 | 0.114 | 0.031 | 1.13e-06 | 0.0381 |
| Fibroblast | F3 COL11A1+ mCAF | GPX7 | 1.648 | 0.308 | 0.089 | 1.2e-15 | 4.04e-11 |
| Fibroblast | F3 COL11A1+ mCAF | CHPF | 1.632 | 0.595 | 0.234 | 3.93e-24 | 1.33e-19 |
| Fibroblast | F3 COL11A1+ mCAF | KDM1A | 1.628 | 0.222 | 0.067 | 1.46e-10 | 4.93e-06 |

| compartment | subcluster | gene | avg_log2FC | pct.1 | pct.2 | p_val | p_val_adj |
|---|---|---|---|---|---|---|---|
| Fibroblast | F3 COL11A1+ mCAF | TMEM200A | 1.628 | 0.168 | 0.04 | 2.39e-10 | 8.06e-06 |
| Fibroblast | F3 COL11A1+ mCAF | RAB31 | 1.626 | 0.686 | 0.289 | 6.84e-27 | 2.31e-22 |
| Fibroblast | F3 COL11A1+ mCAF | RIN2 | 1.624 | 0.346 | 0.142 | 4.48e-11 | 1.51e-06 |
| Fibroblast | F3 COL11A1+ mCAF | CDKN2B | 1.622 | 0.178 | 0.057 | 3.95e-08 | 0.00133 |
| Fibroblast | F3 COL11A1+ mCAF | FAM101B | 1.621 | 0.146 | 0.04 | 5.63e-08 | 0.0019 |
| Fibroblast | F3 COL11A1+ mCAF | CD55 | 1.613 | 0.595 | 0.236 | 1.89e-22 | 6.37e-18 |
| Fibroblast | F3 COL11A1+ mCAF | GOLM1 | 1.601 | 0.503 | 0.166 | 3.45e-23 | 1.16e-18 |
| Fibroblast | F3 COL11A1+ mCAF | LGALS9 | 1.595 | 0.314 | 0.103 | 6.98e-14 | 2.35e-09 |
| Fibroblast | F3 COL11A1+ mCAF | ALDH1A3 | 1.592 | 0.303 | 0.144 | 4.16e-08 | 0.0014 |
| Fibroblast | F3 COL11A1+ mCAF | KDELR3 | 1.581 | 0.416 | 0.159 | 7.55e-16 | 2.54e-11 |
| Fibroblast | F3 COL11A1+ mCAF | SLC7A7 | 1.58 | 0.146 | 0.04 | 4.64e-08 | 0.00156 |
| Fibroblast | F3 COL11A1+ mCAF | COMP | 1.579 | 0.281 | 0.118 | 3.72e-08 | 0.00125 |
| Fibroblast | F3 COL11A1+ mCAF | ADAMTS2 | 1.575 | 0.303 | 0.09 | 9.43e-15 | 3.18e-10 |
| Fibroblast | F3 COL11A1+ mCAF | F2R | 1.572 | 0.524 | 0.169 | 1.29e-24 | 4.35e-20 |
| Fibroblast | F3 COL11A1+ mCAF | NUAK1 | 1.564 | 0.216 | 0.056 | 3.91e-12 | 1.32e-07 |
| Fibroblast | F3 COL11A1+ mCAF | VKORC1L1 | 1.562 | 0.151 | 0.046 | 2.01e-07 | 0.00677 |
| Fibroblast | F3 COL11A1+ mCAF | NREP | 1.548 | 0.449 | 0.136 | 9.46e-22 | 3.19e-17 |
| Fibroblast | F3 COL11A1+ mCAF | DPYSL3 | 1.546 | 0.53 | 0.188 | 3.22e-23 | 1.09e-18 |
| Fibroblast | F3 COL11A1+ mCAF | UNC5B | 1.535 | 0.216 | 0.057 | 7.3e-12 | 2.46e-07 |
| Fibroblast | F3 COL11A1+ mCAF | HSBP1L1 | 1.534 | 0.265 | 0.082 | 2.79e-12 | 9.41e-08 |
| Fibroblast | F3 COL11A1+ mCAF | ITGA2 | 1.533 | 0.103 | 0.017 | 2.18e-09 | 7.35e-05 |
| Fibroblast | F3 COL11A1+ mCAF | AEBP1 | 1.527 | 0.941 | 0.617 | 1.16e-28 | 3.92e-24 |
| Fibroblast | F3 COL11A1+ mCAF | S100A16 | 1.522 | 0.584 | 0.231 | 7.4e-24 | 2.49e-19 |
| Fibroblast | F3 COL11A1+ mCAF | OCIAD2 | 1.52 | 0.238 | 0.067 | 3.39e-12 | 1.14e-07 |
| Fibroblast | F3 COL11A1+ mCAF | TSPAN9 | 1.517 | 0.265 | 0.089 | 3.91e-11 | 1.32e-06 |
| Fibroblast | F3 COL11A1+ mCAF | ATG7 | 1.511 | 0.135 | 0.038 | 2.27e-07 | 0.00764 |
| Fibroblast | F3 COL11A1+ mCAF | LMCD1 | 1.495 | 0.492 | 0.169 | 2.18e-20 | 7.33e-16 |
| Fibroblast | F3 COL11A1+ mCAF | IQSEC2 | 1.491 | 0.13 | 0.032 | 5e-08 | 0.00168 |
| Fibroblast | F3 COL11A1+ mCAF | IL32 | 1.485 | 0.892 | 0.464 | 8.82e-36 | 2.97e-31 |
| Fibroblast | F3 COL11A1+ mCAF | C6orf136 | 1.485 | 0.13 | 0.034 | 1.69e-07 | 0.00568 |
| Fibroblast | F3 COL11A1+ mCAF | COL1A2 | 1.481 | 0.995 | 0.723 | 3.22e-39 | 1.09e-34 |
| Fibroblast | F3 COL11A1+ mCAF | PXDN | 1.466 | 0.319 | 0.109 | 5.83e-13 | 1.97e-08 |
| Fibroblast | F3 COL11A1+ mCAF | MEIS3 | 1.461 | 0.146 | 0.044 | 2.2e-07 | 0.0074 |
| Fibroblast | F3 COL11A1+ mCAF | WARS | 1.448 | 0.357 | 0.116 | 9.42e-16 | 3.17e-11 |
| Fibroblast | F3 COL11A1+ mCAF | TP53I3 | 1.448 | 0.405 | 0.155 | 8.33e-15 | 2.81e-10 |
| Fibroblast | F3 COL11A1+ mCAF | MTHFD1L | 1.444 | 0.146 | 0.044 | 2.6e-07 | 0.00875 |
| Fibroblast | F3 COL11A1+ mCAF | NBL1 | 1.44 | 0.946 | 0.586 | 2.54e-34 | 8.55e-30 |
| Fibroblast | F3 COL11A1+ mCAF | VCAN | 1.44 | 0.816 | 0.414 | 4.320000000000001e-27 | 1.46e-22 |
| Fibroblast | F3 COL11A1+ mCAF | LGALS1 | 1.437 | 1.0 | 0.944 | 1.3e-59 | 4.38e-55 |
| Fibroblast | F3 COL11A1+ mCAF | ANO10 | 1.437 | 0.141 | 0.036 | 2.03e-08 | 0.000685 |
| Fibroblast | F3 COL11A1+ mCAF | GMPPB | 1.433 | 0.227 | 0.071 | 2.53e-10 | 8.51e-06 |
| Fibroblast | F3 COL11A1+ mCAF | SRPX2 | 1.433 | 0.254 | 0.099 | 9.55e-09 | 0.000322 |
| Fibroblast | F3 COL11A1+ mCAF | UBAC2 | 1.431 | 0.47 | 0.169 | 3.82e-18 | 1.29e-13 |
| Fibroblast | F3 COL11A1+ mCAF | CDK2AP1 | 1.424 | 0.238 | 0.08 | 9.21e-10 | 3.1e-05 |
| Fibroblast | F3 COL11A1+ mCAF | ERAP2 | 1.417 | 0.243 | 0.072 | 1.05e-11 | 3.55e-07 |
| Fibroblast | F3 COL11A1+ mCAF | KCTD17 | 1.411 | 0.205 | 0.065 | 2.4e-09 | 8.08e-05 |
| Fibroblast | F3 COL11A1+ mCAF | XXYLT1 | 1.41 | 0.146 | 0.044 | 2.84e-07 | 0.00957 |
| Fibroblast | F3 COL11A1+ mCAF | RP5-1172A22.1 | 1.407 | 0.173 | 0.06 | 4.32e-07 | 0.0146 |
| Fibroblast | F3 COL11A1+ mCAF | DUS1L | 1.404 | 0.135 | 0.033 | 2.24e-08 | 0.000755 |
| Fibroblast | F3 COL11A1+ mCAF | HAPLN3 | 1.402 | 0.432 | 0.149 | 7.1e-18 | 2.39e-13 |
| Fibroblast | F3 COL11A1+ mCAF | PCDH7 | 1.401 | 0.27 | 0.09 | 2.53e-11 | 8.53e-07 |
| Fibroblast | F3 COL11A1+ mCAF | INSIG2 | 1.399 | 0.265 | 0.099 | 2.43e-09 | 8.18e-05 |
| Fibroblast | F3 COL11A1+ mCAF | CALU | 1.396 | 0.746 | 0.412 | 1.08e-22 | 3.63e-18 |
| Fibroblast | F3 COL11A1+ mCAF | EGLN3 | 1.396 | 0.205 | 0.058 | 1.72e-10 | 5.78e-06 |
| Fibroblast | F3 COL11A1+ mCAF | GPR176 | 1.396 | 0.157 | 0.039 | 2.14e-09 | 7.22e-05 |
| Fibroblast | F3 COL11A1+ mCAF | CTNNBIP1 | 1.383 | 0.249 | 0.071 | 2.3e-12 | 7.76e-08 |
| Fibroblast | F3 COL11A1+ mCAF | GALNT1 | 1.374 | 0.265 | 0.097 | 1.03e-09 | 3.47e-05 |
| Fibroblast | F3 COL11A1+ mCAF | PTK7 | 1.368 | 0.427 | 0.149 | 5.75e-17 | 1.94e-12 |
| Fibroblast | F3 COL11A1+ mCAF | ANKS3 | 1.367 | 0.211 | 0.066 | 1.68e-09 | 5.65e-05 |
| Fibroblast | F3 COL11A1+ mCAF | ALDH1B1 | 1.366 | 0.157 | 0.052 | 6.07e-07 | 0.0204 |
| Fibroblast | F3 COL11A1+ mCAF | SGPL1 | 1.365 | 0.168 | 0.049 | 1.84e-08 | 0.000619 |
| Fibroblast | F3 COL11A1+ mCAF | CD2AP | 1.364 | 0.146 | 0.045 | 4.91e-07 | 0.0165 |
| Fibroblast | F3 COL11A1+ mCAF | PIGS | 1.361 | 0.157 | 0.051 | 5.53e-07 | 0.0186 |
| Fibroblast | F3 COL11A1+ mCAF | COLGALT1 | 1.359 | 0.351 | 0.118 | 1.44e-14 | 4.87e-10 |

| compartment | subcluster | gene | avg_log2FC | pct.1 | pct.2 | p_val | p_val_adj |
|---|---|---|---|---|---|---|---|
| Fibroblast | F3 COL11A1+ mCAF | ACSL4 | 1.358 | 0.151 | 0.049 | 4.7e-07 | 0.0158 |
| Fibroblast | F3 COL11A1+ mCAF | YKT6 | 1.357 | 0.238 | 0.072 | 5.49e-11 | 1.85e-06 |
| Fibroblast | F3 COL11A1+ mCAF | RCN3 | 1.356 | 0.714 | 0.354 | 1.95e-22 | 6.57e-18 |
| Fibroblast | F3 COL11A1+ mCAF | SPHK1 | 1.352 | 0.443 | 0.188 | 1.67e-14 | 5.62e-10 |
| Fibroblast | F3 COL11A1+ mCAF | MFSD7 | 1.351 | 0.157 | 0.049 | 2.22e-07 | 0.00748 |
| Fibroblast | F3 COL11A1+ mCAF | TNFAIP6 | 1.35 | 0.335 | 0.135 | 1.68e-10 | 5.68e-06 |
| Fibroblast | F3 COL11A1+ mCAF | SERPINH1 | 1.343 | 0.941 | 0.581 | 1.11e-34 | 3.74e-30 |
| Fibroblast | F3 COL11A1+ mCAF | ROBO1 | 1.338 | 0.184 | 0.062 | 6.66e-08 | 0.00224 |
| Fibroblast | F3 COL11A1+ mCAF | AKT1 | 1.337 | 0.151 | 0.049 | 5.37e-07 | 0.0181 |
| Fibroblast | F3 COL11A1+ mCAF | PERP | 1.329 | 0.443 | 0.161 | 1.42e-16 | 4.77e-12 |
| Fibroblast | F3 COL11A1+ mCAF | MORN4 | 1.325 | 0.119 | 0.032 | 7.88e-07 | 0.0266 |
| Fibroblast | F3 COL11A1+ mCAF | FAP | 1.323 | 0.616 | 0.243 | 2e-22 | 6.74e-18 |
| Fibroblast | F3 COL11A1+ mCAF | VAMP7 | 1.321 | 0.227 | 0.079 | 8.82e-09 | 0.000297 |
| Fibroblast | F3 COL11A1+ mCAF | PLOD2 | 1.318 | 0.454 | 0.175 | 1.66e-16 | 5.58e-12 |
| Fibroblast | F3 COL11A1+ mCAF | UCK2 | 1.313 | 0.114 | 0.027 | 2.58e-07 | 0.0087 |
| Fibroblast | F3 COL11A1+ mCAF | LLNLR-245B6.1 | 1.309 | 0.308 | 0.101 | 5.81e-13 | 1.96e-08 |
| Fibroblast | F3 COL11A1+ mCAF | TMEM167A | 1.307 | 0.578 | 0.246 | 3.03e-19 | 1.02e-14 |
| Fibroblast | F3 COL11A1+ mCAF | SULF2 | 1.305 | 0.53 | 0.207 | 1.49e-17 | 5e-13 |
| Fibroblast | F3 COL11A1+ mCAF | XAF1 | 1.305 | 0.27 | 0.108 | 1.03e-08 | 0.000348 |
| Fibroblast | F3 COL11A1+ mCAF | SLFN5 | 1.305 | 0.151 | 0.05 | 9.9e-07 | 0.0333 |
| Fibroblast | F3 COL11A1+ mCAF | XYLT2 | 1.302 | 0.151 | 0.049 | 6.68e-07 | 0.0225 |
| Fibroblast | F3 COL11A1+ mCAF | COL15A1 | 1.295 | 0.357 | 0.143 | 2.84e-11 | 9.57e-07 |
| Fibroblast | F3 COL11A1+ mCAF | GAS1 | 1.295 | 0.33 | 0.161 | 8.76e-08 | 0.00295 |
| Fibroblast | F3 COL11A1+ mCAF | SPARC | 1.292 | 0.989 | 0.809 | 1.28e-35 | 4.33e-31 |
| Fibroblast | F3 COL11A1+ mCAF | SURF4 | 1.284 | 0.568 | 0.253 | 8.51e-17 | 2.87e-12 |
| Fibroblast | F3 COL11A1+ mCAF | DAP | 1.284 | 0.459 | 0.195 | 2.37e-14 | 8e-10 |
| Fibroblast | F3 COL11A1+ mCAF | MYO10 | 1.282 | 0.124 | 0.031 | 8.64e-08 | 0.00291 |
| Fibroblast | F3 COL11A1+ mCAF | TGFBI | 1.281 | 0.595 | 0.245 | 1.22e-17 | 4.1e-13 |
| Fibroblast | F3 COL11A1+ mCAF | IFI27 | 1.281 | 0.741 | 0.492 | 3.13e-14 | 1.05e-09 |
| Fibroblast | F3 COL11A1+ mCAF | PLAT | 1.279 | 0.557 | 0.273 | 1.06e-14 | 3.56e-10 |
| Fibroblast | F3 COL11A1+ mCAF | GSKIP | 1.275 | 0.184 | 0.065 | 2.88e-07 | 0.00969 |
| Fibroblast | F3 COL11A1+ mCAF | SPATS2L | 1.274 | 0.681 | 0.317 | 3.35e-20 | 1.13e-15 |
| Fibroblast | F3 COL11A1+ mCAF | SCARF2 | 1.272 | 0.276 | 0.08 | 2.53e-13 | 8.53e-09 |
| Fibroblast | F3 COL11A1+ mCAF | FZD1 | 1.272 | 0.378 | 0.141 | 3.27e-13 | 1.1e-08 |
| Fibroblast | F3 COL11A1+ mCAF | RELB | 1.267 | 0.254 | 0.084 | 2.03e-10 | 6.83e-06 |
| Fibroblast | F3 COL11A1+ mCAF | PARP14 | 1.267 | 0.249 | 0.098 | 2.83e-08 | 0.000955 |
| Fibroblast | F3 COL11A1+ mCAF | GPX8 | 1.263 | 0.519 | 0.246 | 2.08e-14 | 7.01e-10 |
| Fibroblast | F3 COL11A1+ mCAF | NAAA | 1.263 | 0.168 | 0.057 | 4.11e-07 | 0.0139 |
| Fibroblast | F3 COL11A1+ mCAF | MARCKS | 1.262 | 0.838 | 0.422 | 4.13e-28 | 1.39e-23 |
| Fibroblast | F3 COL11A1+ mCAF | HTRA1 | 1.261 | 0.67 | 0.399 | 2.86e-13 | 9.62e-09 |
| Fibroblast | F3 COL11A1+ mCAF | EPDR1 | 1.261 | 0.281 | 0.116 | 1.8e-08 | 0.000606 |
| Fibroblast | F3 COL11A1+ mCAF | CNN2 | 1.26 | 0.659 | 0.29 | 9.52e-21 | 3.21e-16 |
| Fibroblast | F3 COL11A1+ mCAF | MBOAT2 | 1.26 | 0.13 | 0.034 | 1.82e-07 | 0.00612 |
| Fibroblast | F3 COL11A1+ mCAF | GPR107 | 1.256 | 0.157 | 0.041 | 5.94e-09 | 0.0002 |
| Fibroblast | F3 COL11A1+ mCAF | ERO1A | 1.254 | 0.195 | 0.06 | 6.21e-09 | 0.000209 |
| Fibroblast | F3 COL11A1+ mCAF | RARRES2 | 1.249 | 0.989 | 0.733 | 1.37e-34 | 4.61e-30 |
| Fibroblast | F3 COL11A1+ mCAF | DHRSX | 1.242 | 0.259 | 0.099 | 6.19e-09 | 0.000209 |
| Fibroblast | F3 COL11A1+ mCAF | TMEM119 | 1.237 | 0.314 | 0.149 | 6.5e-08 | 0.00219 |
| Fibroblast | F3 COL11A1+ mCAF | CTSZ | 1.234 | 0.573 | 0.285 | 1.87e-15 | 6.31e-11 |
| Fibroblast | F3 COL11A1+ mCAF | SERPINE1 | 1.229 | 0.308 | 0.134 | 1.94e-08 | 0.000653 |
| Fibroblast | F3 COL11A1+ mCAF | TAP1 | 1.223 | 0.395 | 0.151 | 1.15e-12 | 3.87e-08 |
| Fibroblast | F3 COL11A1+ mCAF | TMEM44 | 1.223 | 0.103 | 0.022 | 2.38e-07 | 0.00801 |
| Fibroblast | F3 COL11A1+ mCAF | H2AFY2 | 1.221 | 0.211 | 0.073 | 3.43e-08 | 0.00116 |
| Fibroblast | F3 COL11A1+ mCAF | LAMA4 | 1.215 | 0.584 | 0.282 | 1.33e-15 | 4.47e-11 |
| Fibroblast | F3 COL11A1+ mCAF | BMP1 | 1.214 | 0.411 | 0.166 | 1.11e-12 | 3.75e-08 |
| Fibroblast | F3 COL11A1+ mCAF | MRPL23 | 1.212 | 0.449 | 0.172 | 3.04e-15 | 1.02e-10 |
| Fibroblast | F3 COL11A1+ mCAF | ROR2 | 1.212 | 0.216 | 0.076 | 1.54e-08 | 0.000518 |
| Fibroblast | F3 COL11A1+ mCAF | CD82 | 1.21 | 0.378 | 0.144 | 4.97e-13 | 1.68e-08 |
| Fibroblast | F3 COL11A1+ mCAF | CKAP4 | 1.208 | 0.449 | 0.183 | 2.54e-14 | 8.57e-10 |
| Fibroblast | F3 COL11A1+ mCAF | FAM63B | 1.205 | 0.357 | 0.143 | 2.83e-11 | 9.54e-07 |
| Fibroblast | F3 COL11A1+ mCAF | FOXO1 | 1.193 | 0.211 | 0.078 | 1.78e-07 | 0.00601 |
| Fibroblast | F3 COL11A1+ mCAF | HPCAL1 | 1.193 | 0.205 | 0.078 | 3e-07 | 0.0101 |
| Fibroblast | F3 COL11A1+ mCAF | SPON2 | 1.191 | 0.854 | 0.464 | 3.35e-25 | 1.13e-20 |
| Fibroblast | F3 COL11A1+ mCAF | CCDC74A | 1.191 | 0.184 | 0.058 | 2.4e-08 | 0.000809 |

| compartment | subcluster | gene | avg_log2FC | pct.1 | pct.2 | p_val | p_val_adj |
|---|---|---|---|---|---|---|---|
| Fibroblast | F3 COL11A1+ mCAF | ALG3 | 1.188 | 0.281 | 0.107 | 1.31e-09 | 4.41e-05 |
| Fibroblast | F3 COL11A1+ mCAF | CREB3L1 | 1.186 | 0.33 | 0.129 | 9.87e-11 | 3.33e-06 |
| Fibroblast | F3 COL11A1+ mCAF | WDR34 | 1.185 | 0.259 | 0.097 | 3.68e-09 | 0.000124 |
| Fibroblast | F3 COL11A1+ mCAF | P4HB | 1.182 | 0.908 | 0.57 | 1.44e-25 | 4.860000000000001e-21 |
| Fibroblast | F3 COL11A1+ mCAF | NOX4 | 1.181 | 0.173 | 0.046 | 2.33e-09 | 7.86e-05 |
| Fibroblast | F3 COL11A1+ mCAF | LIMD2 | 1.18 | 0.254 | 0.072 | 2.07e-12 | 6.96e-08 |
| Fibroblast | F3 COL11A1+ mCAF | ITPR2 | 1.18 | 0.178 | 0.051 | 4.35e-09 | 0.000146 |
| Fibroblast | F3 COL11A1+ mCAF | AP2S1 | 1.174 | 0.805 | 0.449 | 2.07e-21 | 6.97e-17 |
| Fibroblast | F3 COL11A1+ mCAF | MAF | 1.173 | 0.314 | 0.125 | 7.58e-10 | 2.56e-05 |
| Fibroblast | F3 COL11A1+ mCAF | IRAK1 | 1.173 | 0.168 | 0.054 | 1.89e-07 | 0.00635 |
| Fibroblast | F3 COL11A1+ mCAF | RNF34 | 1.168 | 0.195 | 0.064 | 2.75e-08 | 0.000926 |
| Fibroblast | F3 COL11A1+ mCAF | PFN2 | 1.167 | 0.476 | 0.234 | 5.33e-11 | 1.79e-06 |
| Fibroblast | F3 COL11A1+ mCAF | MDFI | 1.167 | 0.281 | 0.112 | 8.99e-09 | 0.000303 |
| Fibroblast | F3 COL11A1+ mCAF | MYDGF | 1.162 | 0.811 | 0.457 | 6.61e-22 | 2.23e-17 |
| Fibroblast | F3 COL11A1+ mCAF | MARVELD1 | 1.153 | 0.265 | 0.093 | 5.89e-10 | 1.99e-05 |
| Fibroblast | F3 COL11A1+ mCAF | FTH1 | 1.152 | 1.0 | 0.999 | 6.39e-09 | 0.000215 |
| Fibroblast | F3 COL11A1+ mCAF | KLHL5 | 1.151 | 0.168 | 0.054 | 2.23e-07 | 0.00751 |
| Fibroblast | F3 COL11A1+ mCAF | TAPBP | 1.15 | 0.654 | 0.297 | 4.77e-19 | 1.61e-14 |
| Fibroblast | F3 COL11A1+ mCAF | MAGED1 | 1.143 | 0.405 | 0.175 | 2.34e-11 | 7.9e-07 |
| Fibroblast | F3 COL11A1+ mCAF | MT1F | 1.14 | 0.232 | 0.079 | 3.65e-09 | 0.000123 |
| Fibroblast | F3 COL11A1+ mCAF | PPFIBP1 | 1.139 | 0.373 | 0.153 | 3.8e-11 | 1.28e-06 |
| Fibroblast | F3 COL11A1+ mCAF | SLC52A2 | 1.134 | 0.276 | 0.108 | 3.21e-09 | 0.000108 |
| Fibroblast | F3 COL11A1+ mCAF | HS2ST1 | 1.133 | 0.173 | 0.056 | 1.67e-07 | 0.00564 |
| Fibroblast | F3 COL11A1+ mCAF | PTGFRN | 1.128 | 0.281 | 0.114 | 8.69e-09 | 0.000293 |
| Fibroblast | F3 COL11A1+ mCAF | PTTG1IP | 1.126 | 0.627 | 0.32 | 3.8e-16 | 1.28e-11 |
| Fibroblast | F3 COL11A1+ mCAF | C4orf48 | 1.124 | 0.681 | 0.308 | 9.240000000000001e-21 | 3.11e-16 |
| Fibroblast | F3 COL11A1+ mCAF | CARD16 | 1.123 | 0.33 | 0.146 | 1.62e-08 | 0.000546 |
| Fibroblast | F3 COL11A1+ mCAF | RGS10 | 1.122 | 0.395 | 0.162 | 9.93e-12 | 3.35e-07 |
| Fibroblast | F3 COL11A1+ mCAF | AGAP2-AS1 | 1.116 | 0.249 | 0.092 | 9.27e-09 | 0.000312 |
| Fibroblast | F3 COL11A1+ mCAF | DCBLD1 | 1.114 | 0.238 | 0.069 | 2.32e-11 | 7.83e-07 |
| Fibroblast | F3 COL11A1+ mCAF | SMG9 | 1.106 | 0.189 | 0.072 | 1.32e-06 | 0.0445 |
| Fibroblast | F3 COL11A1+ mCAF | YIF1B | 1.101 | 0.254 | 0.09 | 1.14e-09 | 3.83e-05 |
| Fibroblast | F3 COL11A1+ mCAF | TPST1 | 1.096 | 0.465 | 0.218 | 3.07e-12 | 1.04e-07 |
| Fibroblast | F3 COL11A1+ mCAF | MYH9 | 1.095 | 0.795 | 0.408 | 4.67e-22 | 1.57e-17 |
| Fibroblast | F3 COL11A1+ mCAF | C17orf49 | 1.095 | 0.184 | 0.06 | 7.48e-08 | 0.00252 |
| Fibroblast | F3 COL11A1+ mCAF | RARS2 | 1.095 | 0.254 | 0.116 | 1.19e-06 | 0.04 |
| Fibroblast | F3 COL11A1+ mCAF | CHN1 | 1.091 | 0.27 | 0.082 | 1.84e-12 | 6.19e-08 |
| Fibroblast | F3 COL11A1+ mCAF | RHOD | 1.091 | 0.238 | 0.085 | 1.15e-08 | 0.000388 |
| Fibroblast | F3 COL11A1+ mCAF | FHL2 | 1.088 | 0.6 | 0.328 | 5.94e-11 | 2e-06 |
| Fibroblast | F3 COL11A1+ mCAF | MOXD1 | 1.087 | 0.308 | 0.131 | 1.49e-08 | 0.000501 |
| Fibroblast | F3 COL11A1+ mCAF | BID | 1.08 | 0.178 | 0.06 | 1.94e-07 | 0.00654 |
| Fibroblast | F3 COL11A1+ mCAF | CTSK | 1.078 | 0.903 | 0.512 | 2.12e-21 | 7.14e-17 |
| Fibroblast | F3 COL11A1+ mCAF | FAM114A1 | 1.078 | 0.503 | 0.207 | 3.61e-15 | 1.22e-10 |
| Fibroblast | F3 COL11A1+ mCAF | TRAC | 1.078 | 0.357 | 0.122 | 3.75e-13 | 1.26e-08 |
| Fibroblast | F3 COL11A1+ mCAF | BGN | 1.072 | 0.973 | 0.768 | 2.79e-27 | 9.39e-23 |
| Fibroblast | F3 COL11A1+ mCAF | ITGBL1 | 1.072 | 0.341 | 0.128 | 5.79e-12 | 1.95e-07 |
| Fibroblast | F3 COL11A1+ mCAF | PVRL2 | 1.071 | 0.589 | 0.267 | 4.51e-16 | 1.52e-11 |
| Fibroblast | F3 COL11A1+ mCAF | LMAN1 | 1.071 | 0.514 | 0.258 | 3e-11 | 1.01e-06 |
| Fibroblast | F3 COL11A1+ mCAF | NPM3 | 1.068 | 0.346 | 0.156 | 7.79e-09 | 0.000262 |
| Fibroblast | F3 COL11A1+ mCAF | PLXNB2 | 1.067 | 0.27 | 0.11 | 2.66e-08 | 0.000896 |
| Fibroblast | F3 COL11A1+ mCAF | PLEC | 1.064 | 0.438 | 0.205 | 5.42e-11 | 1.83e-06 |
| Fibroblast | F3 COL11A1+ mCAF | FSCN1 | 1.056 | 0.362 | 0.149 | 2.76e-11 | 9.31e-07 |
| Fibroblast | F3 COL11A1+ mCAF | CMTM8 | 1.056 | 0.173 | 0.062 | 1.31e-06 | 0.0442 |
| Fibroblast | F3 COL11A1+ mCAF | ANKLE2 | 1.054 | 0.249 | 0.107 | 5.05e-07 | 0.017 |
| Fibroblast | F3 COL11A1+ mCAF | GOLT1B | 1.053 | 0.362 | 0.134 | 2.23e-12 | 7.52e-08 |
| Fibroblast | F3 COL11A1+ mCAF | SLC25A37 | 1.053 | 0.422 | 0.192 | 1.63e-10 | 5.5e-06 |
| Fibroblast | F3 COL11A1+ mCAF | UBE2L6 | 1.047 | 0.616 | 0.289 | 9.65e-15 | 3.25e-10 |
| Fibroblast | F3 COL11A1+ mCAF | ITGB5 | 1.044 | 0.373 | 0.151 | 6.8e-11 | 2.29e-06 |
| Fibroblast | F3 COL11A1+ mCAF | SDF2L1 | 1.04 | 0.476 | 0.218 | 2.38e-12 | 8.02e-08 |
| Fibroblast | F3 COL11A1+ mCAF | SMIM3 | 1.038 | 0.53 | 0.253 | 1.83e-13 | 6.17e-09 |
| Fibroblast | F3 COL11A1+ mCAF | PPP1R13L | 1.034 | 0.157 | 0.045 | 6.02e-08 | 0.00203 |
| Fibroblast | F3 COL11A1+ mCAF | CARHSP1 | 1.032 | 0.692 | 0.337 | 4.55e-18 | 1.53e-13 |
| Fibroblast | F3 COL11A1+ mCAF | PLAUR | 1.032 | 0.6 | 0.292 | 1.66e-14 | 5.6e-10 |
| Fibroblast | F3 COL11A1+ mCAF | SLC39A13 | 1.032 | 0.378 | 0.156 | 1.8e-11 | 6.06e-07 |

| compartment | subcluster | gene | avg_log2FC | pct.1 | pct.2 | p_val | p_val_adj |
|---|---|---|---|---|---|---|---|
| Fibroblast | F3 COL11A1+ mCAF | TAOK3 | 1.03 | 0.324 | 0.128 | 4.94e-10 | 1.66e-05 |
| Fibroblast | F3 COL11A1+ mCAF | FAM20C | 1.029 | 0.276 | 0.125 | 1.03e-06 | 0.0349 |
| Fibroblast | F3 COL11A1+ mCAF | PKM | 1.028 | 0.881 | 0.567 | 1.38e-18 | 4.63e-14 |
| Fibroblast | F3 COL11A1+ mCAF | FKBP14 | 1.025 | 0.216 | 0.086 | 5.21e-07 | 0.0176 |
| Fibroblast | F3 COL11A1+ mCAF | KIAA0930 | 1.021 | 0.292 | 0.111 | 1.27e-09 | 4.26e-05 |
| Fibroblast | F3 COL11A1+ mCAF | CMBL | 1.021 | 0.222 | 0.089 | 5.49e-07 | 0.0185 |
| Fibroblast | F3 COL11A1+ mCAF | SLC44A1 | 1.018 | 0.416 | 0.208 | 2.12e-08 | 0.000716 |
| Fibroblast | F3 COL11A1+ mCAF | RCN1 | 1.015 | 0.708 | 0.36 | 4.3e-17 | 1.45e-12 |
| Fibroblast | F3 COL11A1+ mCAF | ACBD3 | 1.015 | 0.249 | 0.101 | 1.49e-07 | 0.00503 |
| Fibroblast | F3 COL11A1+ mCAF | AKIP1 | 1.014 | 0.314 | 0.129 | 4.66e-09 | 0.000157 |
| Fibroblast | F3 COL11A1+ mCAF | SLC31A2 | 1.013 | 0.27 | 0.097 | 6.67e-10 | 2.25e-05 |
| Fibroblast | F3 COL11A1+ mCAF | ITPRIP | 1.013 | 0.238 | 0.103 | 1.12e-06 | 0.0378 |
| Fibroblast | F3 COL11A1+ mCAF | ARCN1 | 1.011 | 0.227 | 0.091 | 3.44e-07 | 0.0116 |
| Fibroblast | F3 COL11A1+ mCAF | OSTC | 1.008 | 0.827 | 0.537 | 6.94e-19 | 2.34e-14 |
| Fibroblast | F3 COL11A1+ mCAF | BIRC3 | 1.002 | 0.222 | 0.091 | 1.21e-06 | 0.0406 |
| Fibroblast | F3 COL11A1+ mCAF | PMAIP1 | 0.998 | 0.422 | 0.198 | 3.39e-10 | 1.14e-05 |
| Fibroblast | F3 COL11A1+ mCAF | B4GALT1 | 0.998 | 0.416 | 0.217 | 1.36e-07 | 0.00457 |
| Fibroblast | F3 COL11A1+ mCAF | ISOC2 | 0.997 | 0.53 | 0.234 | 4.77e-14 | 1.61e-09 |
| Fibroblast | F3 COL11A1+ mCAF | TDRD3 | 0.99 | 0.2 | 0.067 | 5.03e-08 | 0.00169 |
| Fibroblast | F3 COL11A1+ mCAF | DRAM1 | 0.988 | 0.238 | 0.088 | 2.27e-08 | 0.000764 |
| Fibroblast | F3 COL11A1+ mCAF | COL6A1 | 0.986 | 0.951 | 0.665 | 1.5e-19 | 5.06e-15 |
| Fibroblast | F3 COL11A1+ mCAF | BST2 | 0.984 | 0.827 | 0.566 | 2.26e-14 | 7.6e-10 |
| Fibroblast | F3 COL11A1+ mCAF | FIBP | 0.983 | 0.503 | 0.232 | 6.12e-13 | 2.06e-08 |
| Fibroblast | F3 COL11A1+ mCAF | LXN | 0.983 | 0.432 | 0.237 | 2.56e-08 | 0.000861 |
| Fibroblast | F3 COL11A1+ mCAF | PARP8 | 0.981 | 0.162 | 0.056 | 1.38e-06 | 0.0466 |
| Fibroblast | F3 COL11A1+ mCAF | PPIC | 0.979 | 0.843 | 0.496 | 2.31e-20 | 7.77e-16 |
| Fibroblast | F3 COL11A1+ mCAF | SSR3 | 0.973 | 0.751 | 0.418 | 2.62e-17 | 8.81e-13 |
| Fibroblast | F3 COL11A1+ mCAF | THY1 | 0.971 | 0.908 | 0.512 | 1.33e-22 | 4.47e-18 |
| Fibroblast | F3 COL11A1+ mCAF | TMEM101 | 0.968 | 0.254 | 0.112 | 7.53e-07 | 0.0254 |
| Fibroblast | F3 COL11A1+ mCAF | SMYD3 | 0.964 | 0.222 | 0.088 | 3.14e-07 | 0.0106 |
| Fibroblast | F3 COL11A1+ mCAF | MDFIC | 0.963 | 0.254 | 0.11 | 5.35e-07 | 0.018 |
| Fibroblast | F3 COL11A1+ mCAF | CAPZB | 0.962 | 0.897 | 0.621 | 1.62e-22 | 5.46e-18 |
| Fibroblast | F3 COL11A1+ mCAF | FAM234A | 0.96 | 0.303 | 0.122 | 7.37e-09 | 0.000248 |
| Fibroblast | F3 COL11A1+ mCAF | FAM89B | 0.959 | 0.389 | 0.18 | 1.41e-09 | 4.76e-05 |
| Fibroblast | F3 COL11A1+ mCAF | PPP2R4 | 0.958 | 0.362 | 0.137 | 1.2e-11 | 4.05e-07 |
| Fibroblast | F3 COL11A1+ mCAF | PDIA4 | 0.957 | 0.605 | 0.293 | 6.77e-15 | 2.28e-10 |
| Fibroblast | F3 COL11A1+ mCAF | LACTB | 0.954 | 0.27 | 0.112 | 1.04e-07 | 0.00351 |
| Fibroblast | F3 COL11A1+ mCAF | ADAM10 | 0.949 | 0.222 | 0.084 | 1.18e-07 | 0.00396 |
| Fibroblast | F3 COL11A1+ mCAF | IFI6 | 0.948 | 0.692 | 0.425 | 4.7e-11 | 1.58e-06 |
| Fibroblast | F3 COL11A1+ mCAF | P3H1 | 0.948 | 0.357 | 0.151 | 4.75e-10 | 1.6e-05 |
| Fibroblast | F3 COL11A1+ mCAF | CTNND1 | 0.947 | 0.249 | 0.093 | 1.35e-08 | 0.000456 |
| Fibroblast | F3 COL11A1+ mCAF | IMPDH1 | 0.946 | 0.211 | 0.083 | 9.13e-07 | 0.0308 |
| Fibroblast | F3 COL11A1+ mCAF | SHISA5 | 0.94 | 0.557 | 0.245 | 1.53e-14 | 5.14e-10 |
| Fibroblast | F3 COL11A1+ mCAF | MSC | 0.94 | 0.384 | 0.159 | 1.31e-10 | 4.4e-06 |
| Fibroblast | F3 COL11A1+ mCAF | LY6E | 0.931 | 0.919 | 0.624 | 4.54e-18 | 1.53e-13 |
| Fibroblast | F3 COL11A1+ mCAF | TM4SF1 | 0.93 | 0.362 | 0.167 | 3.55e-08 | 0.0012 |
| Fibroblast | F3 COL11A1+ mCAF | ENO1 | 0.928 | 0.8 | 0.512 | 2.46e-14 | 8.27e-10 |
| Fibroblast | F3 COL11A1+ mCAF | SLC39A14 | 0.928 | 0.222 | 0.08 | 4.57e-08 | 0.00154 |
| Fibroblast | F3 COL11A1+ mCAF | RALA | 0.926 | 0.378 | 0.155 | 3.25e-11 | 1.09e-06 |
| Fibroblast | F3 COL11A1+ mCAF | SEC61A1 | 0.924 | 0.416 | 0.185 | 6.8e-11 | 2.29e-06 |
| Fibroblast | F3 COL11A1+ mCAF | MICAL2 | 0.921 | 0.2 | 0.047 | 7.51e-12 | 2.53e-07 |
| Fibroblast | F3 COL11A1+ mCAF | IKBIP | 0.919 | 0.362 | 0.166 | 6.69e-09 | 0.000226 |
| Fibroblast | F3 COL11A1+ mCAF | NANS | 0.919 | 0.351 | 0.172 | 5.43e-08 | 0.00183 |
| Fibroblast | F3 COL11A1+ mCAF | ARF4 | 0.917 | 0.87 | 0.574 | 6.14e-17 | 2.07e-12 |
| Fibroblast | F3 COL11A1+ mCAF | RGS3 | 0.915 | 0.4 | 0.175 | 2.48e-10 | 8.36e-06 |
| Fibroblast | F3 COL11A1+ mCAF | VGLL4 | 0.914 | 0.573 | 0.256 | 2.47e-14 | 8.31e-10 |
| Fibroblast | F3 COL11A1+ mCAF | PDLIM2 | 0.912 | 0.708 | 0.404 | 4.41e-14 | 1.49e-09 |
| Fibroblast | F3 COL11A1+ mCAF | MRPL17 | 0.911 | 0.286 | 0.127 | 1.48e-07 | 0.005 |
| Fibroblast | F3 COL11A1+ mCAF | WIPI1 | 0.91 | 0.27 | 0.12 | 4.91e-07 | 0.0166 |
| Fibroblast | F3 COL11A1+ mCAF | LTBP2 | 0.906 | 0.292 | 0.133 | 4.01e-07 | 0.0135 |
| Fibroblast | F3 COL11A1+ mCAF | SPSB1 | 0.905 | 0.4 | 0.179 | 6.51e-10 | 2.19e-05 |
| Fibroblast | F3 COL11A1+ mCAF | GSTO1 | 0.904 | 0.724 | 0.417 | 2.23e-13 | 7.52e-09 |
| Fibroblast | F3 COL11A1+ mCAF | FKBP10 | 0.904 | 0.697 | 0.399 | 6.47e-13 | 2.18e-08 |
| Fibroblast | F3 COL11A1+ mCAF | NRIP1 | 0.903 | 0.232 | 0.088 | 5.15e-08 | 0.00174 |

| compartment | subcluster | gene | avg_log2FC | pct.1 | pct.2 | p_val | p_val_adj |
|---|---|---|---|---|---|---|---|
| Fibroblast | F3 COL11A1+ mCAF | SLC35B2 | 0.902 | 0.373 | 0.144 | 4.05e-11 | 1.36e-06 |
| Fibroblast | F3 COL11A1+ mCAF | OSBPL8 | 0.901 | 0.497 | 0.225 | 1.11e-12 | 3.74e-08 |
| Fibroblast | F3 COL11A1+ mCAF | TMED3 | 0.901 | 0.589 | 0.305 | 1.07e-11 | 3.6e-07 |
| Fibroblast | F3 COL11A1+ mCAF | ITGA5 | 0.9 | 0.314 | 0.149 | 3.41e-07 | 0.0115 |
| Fibroblast | F3 COL11A1+ mCAF | NRP2 | 0.899 | 0.33 | 0.11 | 7.76e-13 | 2.61e-08 |
| Fibroblast | F3 COL11A1+ mCAF | EIF4EBP1 | 0.897 | 0.357 | 0.172 | 4.03e-08 | 0.00136 |
| Fibroblast | F3 COL11A1+ mCAF | DBN1 | 0.897 | 0.286 | 0.121 | 4.49e-08 | 0.00151 |
| Fibroblast | F3 COL11A1+ mCAF | PDLIM4 | 0.895 | 0.508 | 0.23 | 2.95e-13 | 9.94e-09 |
| Fibroblast | F3 COL11A1+ mCAF | TGM2 | 0.893 | 0.324 | 0.12 | 8.45e-11 | 2.85e-06 |
| Fibroblast | F3 COL11A1+ mCAF | TMEM123 | 0.892 | 0.627 | 0.318 | 4.47e-14 | 1.51e-09 |
| Fibroblast | F3 COL11A1+ mCAF | TMED9 | 0.892 | 0.773 | 0.497 | 1.43e-13 | 4.82e-09 |
| Fibroblast | F3 COL11A1+ mCAF | NEK6 | 0.89 | 0.254 | 0.093 | 4.65e-09 | 0.000157 |
| Fibroblast | F3 COL11A1+ mCAF | HSPG2 | 0.884 | 0.53 | 0.22 | 6.02e-15 | 2.03e-10 |
| Fibroblast | F3 COL11A1+ mCAF | COL16A1 | 0.883 | 0.384 | 0.173 | 2.35e-09 | 7.93e-05 |
| Fibroblast | F3 COL11A1+ mCAF | TYMP | 0.882 | 0.8 | 0.419 | 1.17e-17 | 3.94e-13 |
| Fibroblast | F3 COL11A1+ mCAF | BPGM | 0.881 | 0.281 | 0.115 | 2.33e-08 | 0.000784 |
| Fibroblast | F3 COL11A1+ mCAF | GRN | 0.88 | 0.708 | 0.411 | 5.41e-12 | 1.82e-07 |
| Fibroblast | F3 COL11A1+ mCAF | EFEMP2 | 0.879 | 0.811 | 0.471 | 3.32e-17 | 1.12e-12 |
| Fibroblast | F3 COL11A1+ mCAF | B4GALT7 | 0.879 | 0.254 | 0.104 | 2.14e-07 | 0.00721 |
| Fibroblast | F3 COL11A1+ mCAF | TMSB10 | 0.877 | 1.0 | 0.97 | 7.39e-38 | 2.49e-33 |
| Fibroblast | F3 COL11A1+ mCAF | ITGAV | 0.876 | 0.324 | 0.162 | 1.04e-06 | 0.0351 |
| Fibroblast | F3 COL11A1+ mCAF | VPS25 | 0.875 | 0.389 | 0.172 | 3.92e-10 | 1.32e-05 |
| Fibroblast | F3 COL11A1+ mCAF | LAP3 | 0.869 | 0.362 | 0.192 | 9.04e-07 | 0.0305 |
| Fibroblast | F3 COL11A1+ mCAF | MDK | 0.866 | 0.8 | 0.497 | 2.82e-14 | 9.52e-10 |
| Fibroblast | F3 COL11A1+ mCAF | PRAF2 | 0.864 | 0.449 | 0.236 | 1.99e-08 | 0.00067 |
| Fibroblast | F3 COL11A1+ mCAF | SUMO3 | 0.863 | 0.632 | 0.344 | 1.36e-12 | 4.57e-08 |
| Fibroblast | F3 COL11A1+ mCAF | LUZP1 | 0.859 | 0.368 | 0.176 | 2.65e-08 | 0.000894 |
| Fibroblast | F3 COL11A1+ mCAF | SEC22B | 0.857 | 0.373 | 0.187 | 1.22e-07 | 0.00411 |
| Fibroblast | F3 COL11A1+ mCAF | RNPEP | 0.857 | 0.286 | 0.13 | 7.55e-07 | 0.0254 |
| Fibroblast | F3 COL11A1+ mCAF | PSMB10 | 0.855 | 0.319 | 0.134 | 1.06e-08 | 0.000358 |
| Fibroblast | F3 COL11A1+ mCAF | MYOF | 0.851 | 0.227 | 0.091 | 4.27e-07 | 0.0144 |
| Fibroblast | F3 COL11A1+ mCAF | YIPF2 | 0.848 | 0.384 | 0.204 | 5.02e-07 | 0.0169 |
| Fibroblast | F3 COL11A1+ mCAF | TRAF4 | 0.847 | 0.222 | 0.086 | 4.96e-07 | 0.0167 |
| Fibroblast | F3 COL11A1+ mCAF | TIMP2 | 0.846 | 0.832 | 0.503 | 1.09e-14 | 3.67e-10 |
| Fibroblast | F3 COL11A1+ mCAF | PHLDA3 | 0.845 | 0.73 | 0.433 | 2.49e-12 | 8.39e-08 |
| Fibroblast | F3 COL11A1+ mCAF | RAMP1 | 0.844 | 0.351 | 0.131 | 1.14e-11 | 3.85e-07 |
| Fibroblast | F3 COL11A1+ mCAF | BASP1 | 0.843 | 0.616 | 0.278 | 6.69e-16 | 2.25e-11 |
| Fibroblast | F3 COL11A1+ mCAF | COL6A3 | 0.842 | 0.914 | 0.508 | 4.55e-18 | 1.53e-13 |
| Fibroblast | F3 COL11A1+ mCAF | HLA-B | 0.841 | 0.995 | 0.872 | 3.96e-23 | 1.33e-18 |
| Fibroblast | F3 COL11A1+ mCAF | PLOD3 | 0.84 | 0.259 | 0.111 | 4.77e-07 | 0.0161 |
| Fibroblast | F3 COL11A1+ mCAF | YIPF5 | 0.839 | 0.368 | 0.199 | 8.21e-07 | 0.0276 |
| Fibroblast | F3 COL11A1+ mCAF | SNRPA1 | 0.838 | 0.308 | 0.131 | 2.99e-08 | 0.00101 |
| Fibroblast | F3 COL11A1+ mCAF | NISCH | 0.838 | 0.265 | 0.117 | 9.99e-07 | 0.0336 |
| Fibroblast | F3 COL11A1+ mCAF | GALNS | 0.835 | 0.249 | 0.099 | 1.1e-07 | 0.00369 |
| Fibroblast | F3 COL11A1+ mCAF | TUBB | 0.832 | 0.941 | 0.671 | 1.09e-16 | 3.66e-12 |
| Fibroblast | F3 COL11A1+ mCAF | APOL1 | 0.83 | 0.486 | 0.214 | 2.19e-12 | 7.38e-08 |
| Fibroblast | F3 COL11A1+ mCAF | FUOM | 0.825 | 0.195 | 0.065 | 6.7e-08 | 0.00226 |
| Fibroblast | F3 COL11A1+ mCAF | EIF2AK1 | 0.825 | 0.249 | 0.099 | 1.44e-07 | 0.00484 |
| Fibroblast | F3 COL11A1+ mCAF | YIF1A | 0.824 | 0.589 | 0.327 | 2.39e-09 | 8.05e-05 |
| Fibroblast | F3 COL11A1+ mCAF | DCUN1D5 | 0.824 | 0.292 | 0.122 | 3.39e-08 | 0.00114 |
| Fibroblast | F3 COL11A1+ mCAF | H2AFY | 0.823 | 0.605 | 0.301 | 1.39e-12 | 4.68e-08 |
| Fibroblast | F3 COL11A1+ mCAF | PLXDC2 | 0.819 | 0.454 | 0.224 | 3.71e-09 | 0.000125 |
| Fibroblast | F3 COL11A1+ mCAF | PLPP5 | 0.818 | 0.281 | 0.127 | 5.89e-07 | 0.0198 |
| Fibroblast | F3 COL11A1+ mCAF | PRELID1 | 0.816 | 0.751 | 0.427 | 4.64e-13 | 1.56e-08 |
| Fibroblast | F3 COL11A1+ mCAF | BHLHE40 | 0.816 | 0.422 | 0.233 | 7.1e-07 | 0.0239 |
| Fibroblast | F3 COL11A1+ mCAF | TIMP1 | 0.815 | 1.0 | 0.928 | 1.1e-12 | 3.7e-08 |
| Fibroblast | F3 COL11A1+ mCAF | LUM | 0.814 | 0.978 | 0.604 | 1.53e-20 | 5.16e-16 |
| Fibroblast | F3 COL11A1+ mCAF | GOLGA4 | 0.814 | 0.368 | 0.189 | 2.52e-07 | 0.00851 |
| Fibroblast | F3 COL11A1+ mCAF | MXRA8 | 0.813 | 0.859 | 0.569 | 1.26e-13 | 4.26e-09 |
| Fibroblast | F3 COL11A1+ mCAF | PLOD1 | 0.813 | 0.384 | 0.179 | 2.88e-08 | 0.00097 |
| Fibroblast | F3 COL11A1+ mCAF | STAT1 | 0.812 | 0.395 | 0.193 | 1.76e-08 | 0.000594 |
| Fibroblast | F3 COL11A1+ mCAF | STMN3 | 0.812 | 0.373 | 0.18 | 4.25e-08 | 0.00143 |
| Fibroblast | F3 COL11A1+ mCAF | CKLF | 0.808 | 0.47 | 0.236 | 8.52e-10 | 2.87e-05 |
| Fibroblast | F3 COL11A1+ mCAF | BICC1 | 0.808 | 0.443 | 0.212 | 1.51e-09 | 5.08e-05 |

| compartment | subcluster | gene | avg_log2FC | pct.1 | pct.2 | p_val | p_val_adj |
|---|---|---|---|---|---|---|---|
| Fibroblast | F3 COL11A1+ mCAF | MRPL12 | 0.806 | 0.389 | 0.18 | 9.03e-09 | 0.000304 |
| Fibroblast | F3 COL11A1+ mCAF | CNPY4 | 0.806 | 0.308 | 0.129 | 1.71e-08 | 0.000575 |
| Fibroblast | F3 COL11A1+ mCAF | CLMP | 0.805 | 0.378 | 0.179 | 3.17e-08 | 0.00107 |
| Fibroblast | F3 COL11A1+ mCAF | PPIB | 0.804 | 0.951 | 0.73 | 1.74e-17 | 5.85e-13 |
| Fibroblast | F3 COL11A1+ mCAF | CISD1 | 0.804 | 0.611 | 0.29 | 9.4e-14 | 3.17e-09 |
| Fibroblast | F3 COL11A1+ mCAF | S100A11 | 0.803 | 0.962 | 0.808 | 1.4e-22 | 4.72e-18 |
| Fibroblast | F3 COL11A1+ mCAF | CD276 | 0.803 | 0.362 | 0.17 | 2.33e-08 | 0.000786 |
| Fibroblast | F3 COL11A1+ mCAF | PBDC1 | 0.802 | 0.335 | 0.149 | 3.29e-08 | 0.00111 |
| Fibroblast | F3 COL11A1+ mCAF | SNAP23 | 0.802 | 0.346 | 0.16 | 6.53e-08 | 0.0022 |
| Fibroblast | F3 COL11A1+ mCAF | IL10RB | 0.802 | 0.373 | 0.187 | 3.39e-07 | 0.0114 |
| Fibroblast | F3 COL11A1+ mCAF | UROS | 0.799 | 0.389 | 0.176 | 4.94e-09 | 0.000167 |
| Fibroblast | F3 COL11A1+ mCAF | FAM198B | 0.799 | 0.33 | 0.146 | 1.53e-08 | 0.000516 |
| Fibroblast | F3 COL11A1+ mCAF | NCLN | 0.799 | 0.195 | 0.072 | 6.36e-07 | 0.0214 |
| Fibroblast | F3 COL11A1+ mCAF | HDLBP | 0.798 | 0.578 | 0.314 | 1.53e-10 | 5.17e-06 |
| Fibroblast | F3 COL11A1+ mCAF | ARF5 | 0.797 | 0.611 | 0.325 | 8.5e-11 | 2.86e-06 |
| Fibroblast | F3 COL11A1+ mCAF | BRI3 | 0.796 | 0.686 | 0.387 | 2.12e-12 | 7.14e-08 |
| Fibroblast | F3 COL11A1+ mCAF | WBSCR16 | 0.793 | 0.281 | 0.128 | 1.13e-06 | 0.0381 |
| Fibroblast | F3 COL11A1+ mCAF | CERCAM | 0.792 | 0.616 | 0.331 | 3.94e-11 | 1.33e-06 |
| Fibroblast | F3 COL11A1+ mCAF | AKR1A1 | 0.792 | 0.584 | 0.314 | 8.39e-11 | 2.83e-06 |
| Fibroblast | F3 COL11A1+ mCAF | PLEKHJ1 | 0.792 | 0.324 | 0.147 | 1.6e-07 | 0.00539 |
| Fibroblast | F3 COL11A1+ mCAF | RPL26L1 | 0.79 | 0.427 | 0.174 | 5.91e-12 | 1.99e-07 |
| Fibroblast | F3 COL11A1+ mCAF | LAMB1 | 0.79 | 0.4 | 0.196 | 3.54e-08 | 0.00119 |
| Fibroblast | F3 COL11A1+ mCAF | PSMB9 | 0.786 | 0.589 | 0.338 | 9.1e-09 | 0.000306 |
| Fibroblast | F3 COL11A1+ mCAF | GTF3A | 0.786 | 0.519 | 0.286 | 1.09e-08 | 0.000368 |
| Fibroblast | F3 COL11A1+ mCAF | NCOR2 | 0.785 | 0.286 | 0.121 | 1.14e-07 | 0.00383 |
| Fibroblast | F3 COL11A1+ mCAF | ABRACL | 0.785 | 0.314 | 0.148 | 2.16e-07 | 0.00729 |
| Fibroblast | F3 COL11A1+ mCAF | VIMP | 0.784 | 0.714 | 0.407 | 8.28e-12 | 2.79e-07 |
| Fibroblast | F3 COL11A1+ mCAF | CAPG | 0.784 | 0.281 | 0.125 | 1.69e-07 | 0.00569 |
| Fibroblast | F3 COL11A1+ mCAF | IFNGR2 | 0.781 | 0.438 | 0.237 | 1.2e-07 | 0.00406 |
| Fibroblast | F3 COL11A1+ mCAF | BACE2 | 0.781 | 0.362 | 0.188 | 8.64e-07 | 0.0291 |
| Fibroblast | F3 COL11A1+ mCAF | FMOD | 0.777 | 0.459 | 0.236 | 1.87e-08 | 0.000631 |
| Fibroblast | F3 COL11A1+ mCAF | HN1 | 0.777 | 0.378 | 0.185 | 5.28e-08 | 0.00178 |
| Fibroblast | F3 COL11A1+ mCAF | FSTL1 | 0.776 | 0.876 | 0.529 | 1.46e-16 | 4.9e-12 |
| Fibroblast | F3 COL11A1+ mCAF | PDIA6 | 0.775 | 0.697 | 0.433 | 1.31e-11 | 4.4e-07 |
| Fibroblast | F3 COL11A1+ mCAF | FKBP9 | 0.774 | 0.481 | 0.25 | 7.96e-09 | 0.000268 |
| Fibroblast | F3 COL11A1+ mCAF | NOP10 | 0.773 | 0.805 | 0.492 | 1.41e-12 | 4.77e-08 |
| Fibroblast | F3 COL11A1+ mCAF | ITPRIPL2 | 0.767 | 0.243 | 0.107 | 1.26e-06 | 0.0426 |
| Fibroblast | F3 COL11A1+ mCAF | PRRX1 | 0.766 | 0.524 | 0.291 | 3.09e-09 | 0.000104 |
| Fibroblast | F3 COL11A1+ mCAF | SLC20A1 | 0.765 | 0.346 | 0.163 | 8.14e-08 | 0.00274 |
| Fibroblast | F3 COL11A1+ mCAF | CDK6 | 0.764 | 0.141 | 0.041 | 6.41e-07 | 0.0216 |
| Fibroblast | F3 COL11A1+ mCAF | TMEM54 | 0.762 | 0.297 | 0.125 | 3.77e-08 | 0.00127 |
| Fibroblast | F3 COL11A1+ mCAF | RPS27L | 0.761 | 0.941 | 0.741 | 3.74e-15 | 1.26e-10 |
| Fibroblast | F3 COL11A1+ mCAF | PSME2 | 0.761 | 0.719 | 0.402 | 1.46e-12 | 4.91e-08 |
| Fibroblast | F3 COL11A1+ mCAF | CCND1 | 0.761 | 0.33 | 0.12 | 6.83e-11 | 2.3e-06 |
| Fibroblast | F3 COL11A1+ mCAF | EVA1B | 0.756 | 0.465 | 0.226 | 1.51e-09 | 5.08e-05 |
| Fibroblast | F3 COL11A1+ mCAF | CAPZA1 | 0.756 | 0.459 | 0.253 | 1.71e-07 | 0.00577 |
| Fibroblast | F3 COL11A1+ mCAF | EIF4E2 | 0.755 | 0.427 | 0.205 | 1.37e-08 | 0.000462 |
| Fibroblast | F3 COL11A1+ mCAF | SLC39A7 | 0.753 | 0.454 | 0.226 | 1.08e-08 | 0.000364 |
| Fibroblast | F3 COL11A1+ mCAF | SAR1A | 0.752 | 0.524 | 0.303 | 1.45e-08 | 0.000488 |
| Fibroblast | F3 COL11A1+ mCAF | STIP1 | 0.752 | 0.292 | 0.131 | 3.93e-07 | 0.0133 |
| Fibroblast | F3 COL11A1+ mCAF | GAPDH | 0.75 | 0.989 | 0.934 | 9.41e-22 | 3.17e-17 |
| Fibroblast | F3 COL11A1+ mCAF | NEDD9 | 0.75 | 0.189 | 0.067 | 4.31e-07 | 0.0145 |
| Fibroblast | F3 COL11A1+ mCAF | EMILIN1 | 0.749 | 0.822 | 0.517 | 4.75e-13 | 1.6e-08 |
| Fibroblast | F3 COL11A1+ mCAF | SVIL | 0.749 | 0.503 | 0.259 | 4.94e-09 | 0.000166 |
| Fibroblast | F3 COL11A1+ mCAF | TMEM45A | 0.748 | 0.573 | 0.275 | 1.14e-11 | 3.83e-07 |
| Fibroblast | F3 COL11A1+ mCAF | PEMT | 0.744 | 0.227 | 0.093 | 1.24e-06 | 0.0418 |
| Fibroblast | F3 COL11A1+ mCAF | MIF | 0.742 | 0.876 | 0.62 | 1.24e-12 | 4.19e-08 |
| Fibroblast | F3 COL11A1+ mCAF | DUSP10 | 0.742 | 0.222 | 0.08 | 6.46e-08 | 0.00218 |
| Fibroblast | F3 COL11A1+ mCAF | PSEN1 | 0.741 | 0.211 | 0.076 | 1.5e-07 | 0.00504 |
| Fibroblast | F3 COL11A1+ mCAF | CD99 | 0.738 | 0.924 | 0.776 | 7.45e-12 | 2.51e-07 |
| Fibroblast | F3 COL11A1+ mCAF | AGTRAP | 0.737 | 0.411 | 0.199 | 2.26e-08 | 0.000762 |
| Fibroblast | F3 COL11A1+ mCAF | CTSA | 0.737 | 0.541 | 0.321 | 3.03e-08 | 0.00102 |
| Fibroblast | F3 COL11A1+ mCAF | MYL12A | 0.735 | 0.924 | 0.746 | 3.83e-14 | 1.29e-09 |
| Fibroblast | F3 COL11A1+ mCAF | LYPLA1 | 0.735 | 0.281 | 0.125 | 8.97e-07 | 0.0302 |

| compartment | subcluster | gene | avg_log2FC | pct.1 | pct.2 | p_val | p_val_adj |
|---|---|---|---|---|---|---|---|
| Fibroblast | F3 COL11A1+ mCAF | MRPL36 | 0.734 | 0.438 | 0.22 | 2.37e-08 | 0.000797 |
| Fibroblast | F3 COL11A1+ mCAF | SH3BGRL3 | 0.733 | 0.87 | 0.63 | 2.58e-11 | 8.69e-07 |
| Fibroblast | F3 COL11A1+ mCAF | FAM118A | 0.733 | 0.308 | 0.141 | 4.52e-07 | 0.0152 |
| Fibroblast | F3 COL11A1+ mCAF | TPM1 | 0.732 | 0.849 | 0.682 | 1.93e-08 | 0.00065 |
| Fibroblast | F3 COL11A1+ mCAF | KDELR2 | 0.731 | 0.778 | 0.537 | 4.41e-10 | 1.49e-05 |
| Fibroblast | F3 COL11A1+ mCAF | CASC4 | 0.729 | 0.443 | 0.212 | 5.07e-09 | 0.000171 |
| Fibroblast | F3 COL11A1+ mCAF | TNFRSF12A | 0.729 | 0.557 | 0.324 | 1.57e-07 | 0.00528 |
| Fibroblast | F3 COL11A1+ mCAF | LY96 | 0.729 | 0.432 | 0.232 | 3.88e-07 | 0.0131 |
| Fibroblast | F3 COL11A1+ mCAF | IL1R1 | 0.725 | 0.395 | 0.205 | 3.5e-07 | 0.0118 |
| Fibroblast | F3 COL11A1+ mCAF | GLT8D2 | 0.724 | 0.589 | 0.302 | 8e-11 | 2.7e-06 |
| Fibroblast | F3 COL11A1+ mCAF | UBL7 | 0.724 | 0.297 | 0.124 | 5.18e-08 | 0.00174 |
| Fibroblast | F3 COL11A1+ mCAF | PDIA3 | 0.722 | 0.881 | 0.612 | 2.22e-14 | 7.49e-10 |
| Fibroblast | F3 COL11A1+ mCAF | COPZ1 | 0.721 | 0.508 | 0.279 | 1.42e-08 | 0.000478 |
| Fibroblast | F3 COL11A1+ mCAF | ATOX1 | 0.716 | 0.757 | 0.466 | 1.78e-12 | 6.01e-08 |
| Fibroblast | F3 COL11A1+ mCAF | LEPROTL1 | 0.716 | 0.422 | 0.198 | 3.49e-09 | 0.000118 |
| Fibroblast | F3 COL11A1+ mCAF | CYB5A | 0.715 | 0.638 | 0.314 | 2.25e-12 | 7.6e-08 |
| Fibroblast | F3 COL11A1+ mCAF | RCN2 | 0.712 | 0.595 | 0.32 | 5.15e-10 | 1.74e-05 |
| Fibroblast | F3 COL11A1+ mCAF | IFT22 | 0.708 | 0.33 | 0.151 | 1.63e-07 | 0.00549 |
| Fibroblast | F3 COL11A1+ mCAF | PLIN3 | 0.707 | 0.476 | 0.25 | 1.02e-08 | 0.000342 |
| Fibroblast | F3 COL11A1+ mCAF | ETHE1 | 0.705 | 0.405 | 0.207 | 1.75e-07 | 0.00591 |
| Fibroblast | F3 COL11A1+ mCAF | TPM4 | 0.703 | 0.903 | 0.66 | 1.46e-13 | 4.92e-09 |
| Fibroblast | F3 COL11A1+ mCAF | LSM2 | 0.703 | 0.595 | 0.31 | 8.6e-11 | 2.9e-06 |
| Fibroblast | F3 COL11A1+ mCAF | AGPAT2 | 0.702 | 0.373 | 0.189 | 1.11e-06 | 0.0376 |
| Fibroblast | F3 COL11A1+ mCAF | CALR | 0.701 | 0.908 | 0.703 | 1e-12 | 3.38e-08 |
| Fibroblast | F3 COL11A1+ mCAF | PDLIM7 | 0.699 | 0.681 | 0.418 | 6.11e-09 | 0.000206 |
| Fibroblast | F3 COL11A1+ mCAF | FAM32A | 0.699 | 0.497 | 0.264 | 9.67e-09 | 0.000326 |
| Fibroblast | F3 COL11A1+ mCAF | PYCARD | 0.697 | 0.368 | 0.187 | 4.12e-07 | 0.0139 |
| Fibroblast | F3 COL11A1+ mCAF | PGAM1 | 0.693 | 0.741 | 0.439 | 1.65e-11 | 5.56e-07 |
| Fibroblast | F3 COL11A1+ mCAF | POLR2G | 0.692 | 0.568 | 0.305 | 2.46e-09 | 8.28e-05 |
| Fibroblast | F3 COL11A1+ mCAF | TAX1BP3 | 0.688 | 0.768 | 0.466 | 4.04e-11 | 1.36e-06 |
| Fibroblast | F3 COL11A1+ mCAF | DEF8 | 0.687 | 0.314 | 0.129 | 9.43e-09 | 0.000318 |
| Fibroblast | F3 COL11A1+ mCAF | PRSS23 | 0.687 | 0.568 | 0.343 | 1.51e-07 | 0.00509 |
| Fibroblast | F3 COL11A1+ mCAF | TMED2 | 0.686 | 0.849 | 0.566 | 6.86e-12 | 2.31e-07 |
| Fibroblast | F3 COL11A1+ mCAF | RHBDD2 | 0.686 | 0.47 | 0.22 | 9.23e-10 | 3.11e-05 |
| Fibroblast | F3 COL11A1+ mCAF | GRINA | 0.684 | 0.481 | 0.254 | 4.73e-08 | 0.00159 |
| Fibroblast | F3 COL11A1+ mCAF | TWF2 | 0.684 | 0.373 | 0.18 | 1.16e-07 | 0.0039 |
| Fibroblast | F3 COL11A1+ mCAF | COA3 | 0.681 | 0.47 | 0.266 | 6.38e-07 | 0.0215 |
| Fibroblast | F3 COL11A1+ mCAF | CLEC11A | 0.678 | 0.827 | 0.518 | 5.87e-13 | 1.98e-08 |
| Fibroblast | F3 COL11A1+ mCAF | RAB32 | 0.674 | 0.378 | 0.193 | 3.94e-07 | 0.0133 |
| Fibroblast | F3 COL11A1+ mCAF | PCOLCE | 0.671 | 0.892 | 0.578 | 1.14e-12 | 3.83e-08 |
| Fibroblast | F3 COL11A1+ mCAF | TOM1 | 0.67 | 0.286 | 0.122 | 1.76e-07 | 0.00592 |
| Fibroblast | F3 COL11A1+ mCAF | TPBG | 0.669 | 0.427 | 0.211 | 4.54e-08 | 0.00153 |
| Fibroblast | F3 COL11A1+ mCAF | NDUFC2 | 0.667 | 0.832 | 0.579 | 8.52e-09 | 0.000287 |
| Fibroblast | F3 COL11A1+ mCAF | SEPT11 | 0.667 | 0.595 | 0.356 | 2.91e-08 | 0.000979 |
| Fibroblast | F3 COL11A1+ mCAF | AZIN1 | 0.667 | 0.368 | 0.185 | 3.88e-07 | 0.0131 |
| Fibroblast | F3 COL11A1+ mCAF | C9orf16 | 0.666 | 0.741 | 0.476 | 3.1e-10 | 1.04e-05 |
| Fibroblast | F3 COL11A1+ mCAF | HLA-C | 0.664 | 0.978 | 0.884 | 5.44e-17 | 1.83e-12 |
| Fibroblast | F3 COL11A1+ mCAF | ITGB1 | 0.664 | 0.87 | 0.647 | 1.94e-10 | 6.52e-06 |
| Fibroblast | F3 COL11A1+ mCAF | DYNC1LI2 | 0.664 | 0.384 | 0.193 | 1.09e-07 | 0.00368 |
| Fibroblast | F3 COL11A1+ mCAF | DYNLT1 | 0.663 | 0.73 | 0.446 | 3.79e-10 | 1.28e-05 |
| Fibroblast | F3 COL11A1+ mCAF | MPZL1 | 0.663 | 0.465 | 0.266 | 1.43e-06 | 0.0482 |
| Fibroblast | F3 COL11A1+ mCAF | EPSTI1 | 0.661 | 0.503 | 0.29 | 2.81e-07 | 0.00946 |
| Fibroblast | F3 COL11A1+ mCAF | SH3GLB1 | 0.66 | 0.578 | 0.307 | 5.55e-10 | 1.87e-05 |
| Fibroblast | F3 COL11A1+ mCAF | TMEM141 | 0.658 | 0.427 | 0.217 | 9.31e-08 | 0.00314 |
| Fibroblast | F3 COL11A1+ mCAF | HDGF | 0.656 | 0.341 | 0.169 | 1.1e-06 | 0.0369 |
| Fibroblast | F3 COL11A1+ mCAF | TMEM176A | 0.655 | 0.681 | 0.419 | 4.17e-08 | 0.00141 |
| Fibroblast | F3 COL11A1+ mCAF | EMC10 | 0.653 | 0.589 | 0.329 | 4.57e-09 | 0.000154 |
| Fibroblast | F3 COL11A1+ mCAF | TNS3 | 0.653 | 0.243 | 0.09 | 2.55e-08 | 0.000861 |
| Fibroblast | F3 COL11A1+ mCAF | PEA15 | 0.649 | 0.422 | 0.221 | 1.9e-07 | 0.0064 |
| Fibroblast | F3 COL11A1+ mCAF | GREM1 | 0.648 | 0.281 | 0.116 | 6.04e-08 | 0.00203 |
| Fibroblast | F3 COL11A1+ mCAF | HSP90B1 | 0.646 | 0.876 | 0.604 | 1.23e-09 | 4.14e-05 |
| Fibroblast | F3 COL11A1+ mCAF | PDLIM3 | 0.646 | 0.578 | 0.372 | 6.06e-07 | 0.0204 |
| Fibroblast | F3 COL11A1+ mCAF | NAA50 | 0.645 | 0.378 | 0.189 | 4.22e-07 | 0.0142 |
| Fibroblast | F3 COL11A1+ mCAF | MAP4K4 | 0.645 | 0.303 | 0.142 | 4.31e-07 | 0.0145 |

| compartment | subcluster | gene | avg_log2FC | pct.1 | pct.2 | p_val | p_val_adj |
|---|---|---|---|---|---|---|---|
| Fibroblast | F3 COL11A1+ mCAF | GPC6 | 0.644 | 0.33 | 0.159 | 9.25e-07 | 0.0312 |
| Fibroblast | F3 COL11A1+ mCAF | CASP4 | 0.642 | 0.4 | 0.217 | 1.14e-06 | 0.0385 |
| Fibroblast | F3 COL11A1+ mCAF | REEP3 | 0.641 | 0.384 | 0.187 | 1.28e-07 | 0.00432 |
| Fibroblast | F3 COL11A1+ mCAF | ERCC1 | 0.641 | 0.465 | 0.264 | 3.12e-07 | 0.0105 |
| Fibroblast | F3 COL11A1+ mCAF | ATP6V0B | 0.64 | 0.692 | 0.413 | 4.3e-09 | 0.000145 |
| Fibroblast | F3 COL11A1+ mCAF | FAT1 | 0.64 | 0.205 | 0.07 | 5.9e-08 | 0.00199 |
| Fibroblast | F3 COL11A1+ mCAF | NRBP1 | 0.64 | 0.465 | 0.262 | 8.95e-07 | 0.0302 |
| Fibroblast | F3 COL11A1+ mCAF | RPN2 | 0.64 | 0.519 | 0.312 | 1.28e-06 | 0.0433 |
| Fibroblast | F3 COL11A1+ mCAF | LRP10 | 0.639 | 0.557 | 0.318 | 1.74e-07 | 0.00587 |
| Fibroblast | F3 COL11A1+ mCAF | GARS | 0.638 | 0.303 | 0.138 | 6.84e-07 | 0.0231 |
| Fibroblast | F3 COL11A1+ mCAF | NME1 | 0.637 | 0.486 | 0.254 | 9.29e-09 | 0.000313 |
| Fibroblast | F3 COL11A1+ mCAF | SEC61G | 0.637 | 0.892 | 0.682 | 2.12e-08 | 0.000713 |
| Fibroblast | F3 COL11A1+ mCAF | SEC23A | 0.637 | 0.362 | 0.18 | 4.83e-07 | 0.0163 |
| Fibroblast | F3 COL11A1+ mCAF | SNX17 | 0.636 | 0.524 | 0.279 | 2.11e-08 | 0.000711 |
| Fibroblast | F3 COL11A1+ mCAF | PALLD | 0.632 | 0.681 | 0.417 | 1.18e-09 | 3.98e-05 |
| Fibroblast | F3 COL11A1+ mCAF | ECM1 | 0.628 | 0.503 | 0.292 | 2.46e-07 | 0.00828 |
| Fibroblast | F3 COL11A1+ mCAF | YIPF3 | 0.626 | 0.562 | 0.32 | 3.03e-08 | 0.00102 |
| Fibroblast | F3 COL11A1+ mCAF | VKORC1 | 0.625 | 0.854 | 0.575 | 1.77e-11 | 5.97e-07 |
| Fibroblast | F3 COL11A1+ mCAF | TXNDC17 | 0.625 | 0.476 | 0.245 | 2.53e-08 | 0.000854 |
| Fibroblast | F3 COL11A1+ mCAF | DNAJC3 | 0.622 | 0.438 | 0.219 | 3.85e-08 | 0.0013 |
| Fibroblast | F3 COL11A1+ mCAF | PSMD7 | 0.619 | 0.535 | 0.276 | 5.03e-09 | 0.00017 |
| Fibroblast | F3 COL11A1+ mCAF | BAX | 0.618 | 0.638 | 0.386 | 4.96e-08 | 0.00167 |
| Fibroblast | F3 COL11A1+ mCAF | TMEM176B | 0.616 | 0.789 | 0.473 | 1.82e-10 | 6.15e-06 |
| Fibroblast | F3 COL11A1+ mCAF | MMP2 | 0.613 | 0.957 | 0.54 | 4.12e-15 | 1.39e-10 |
| Fibroblast | F3 COL11A1+ mCAF | SPCS3 | 0.613 | 0.368 | 0.157 | 4.01e-09 | 0.000135 |
| Fibroblast | F3 COL11A1+ mCAF | CCPG1 | 0.613 | 0.443 | 0.253 | 1.42e-06 | 0.0478 |
| Fibroblast | F3 COL11A1+ mCAF | CNIH1 | 0.612 | 0.6 | 0.325 | 1.21e-09 | 4.06e-05 |
| Fibroblast | F3 COL11A1+ mCAF | RNF181 | 0.612 | 0.665 | 0.378 | 2.25e-09 | 7.6e-05 |
| Fibroblast | F3 COL11A1+ mCAF | S100A10 | 0.611 | 0.897 | 0.582 | 1.55e-14 | 5.24e-10 |
| Fibroblast | F3 COL11A1+ mCAF | TXNL1 | 0.609 | 0.578 | 0.328 | 3.51e-08 | 0.00118 |
| Fibroblast | F3 COL11A1+ mCAF | NNMT | 0.607 | 0.941 | 0.693 | 2.1e-15 | 7.09e-11 |
| Fibroblast | F3 COL11A1+ mCAF | SND1 | 0.607 | 0.4 | 0.189 | 1.14e-08 | 0.000386 |
| Fibroblast | F3 COL11A1+ mCAF | TMEM126A | 0.607 | 0.378 | 0.183 | 2.56e-07 | 0.00864 |
| Fibroblast | F3 COL11A1+ mCAF | SHC1 | 0.607 | 0.324 | 0.15 | 2.69e-07 | 0.00908 |
| Fibroblast | F3 COL11A1+ mCAF | SMARCB1 | 0.606 | 0.422 | 0.226 | 4.37e-07 | 0.0147 |
| Fibroblast | F3 COL11A1+ mCAF | CUEDC2 | 0.605 | 0.486 | 0.276 | 8.79e-07 | 0.0296 |
| Fibroblast | F3 COL11A1+ mCAF | CREB3 | 0.604 | 0.303 | 0.14 | 6.51e-07 | 0.0219 |
| Fibroblast | F3 COL11A1+ mCAF | AUP1 | 0.603 | 0.573 | 0.304 | 2.72e-09 | 9.16e-05 |
| Fibroblast | F3 COL11A1+ mCAF | CAPNS1 | 0.602 | 0.481 | 0.245 | 1.8e-08 | 0.000605 |
| Fibroblast | F3 COL11A1+ mCAF | CLNS1A | 0.602 | 0.443 | 0.241 | 2.62e-07 | 0.00882 |
| Fibroblast | F3 COL11A1+ mCAF | CETN2 | 0.601 | 0.476 | 0.265 | 3.06e-07 | 0.0103 |
| Fibroblast | F3 COL11A1+ mCAF | SSU72 | 0.6 | 0.665 | 0.389 | 1.49e-09 | 5.02e-05 |
| Fibroblast | F3 COL11A1+ mCAF | C7orf73 | 0.599 | 0.622 | 0.342 | 1.39e-09 | 4.67e-05 |
| Fibroblast | F3 COL11A1+ mCAF | PHPT1 | 0.598 | 0.822 | 0.605 | 6.8e-09 | 0.000229 |
| Fibroblast | F3 COL11A1+ mCAF | PPP1CC | 0.598 | 0.659 | 0.413 | 5.22e-07 | 0.0176 |
| Fibroblast | F3 COL11A1+ mCAF | HSPA5 | 0.598 | 0.816 | 0.613 | 8.74e-07 | 0.0295 |
| Fibroblast | F3 COL11A1+ mCAF | UQCC2 | 0.596 | 0.438 | 0.244 | 9.55e-07 | 0.0322 |
| Fibroblast | F3 COL11A1+ mCAF | ARPC2 | 0.594 | 0.903 | 0.672 | 5.84e-12 | 1.97e-07 |
| Fibroblast | F3 COL11A1+ mCAF | FEZ2 | 0.593 | 0.373 | 0.181 | 3.28e-07 | 0.0111 |
| Fibroblast | F3 COL11A1+ mCAF | JTB | 0.591 | 0.832 | 0.55 | 1.64e-09 | 5.52e-05 |
| Fibroblast | F3 COL11A1+ mCAF | ALKBH7 | 0.591 | 0.6 | 0.393 | 1.13e-06 | 0.0381 |
| Fibroblast | F3 COL11A1+ mCAF | GNAI2 | 0.591 | 0.514 | 0.299 | 1.26e-06 | 0.0425 |
| Fibroblast | F3 COL11A1+ mCAF | ATP5E | 0.59 | 0.989 | 0.867 | 3.48e-20 | 1.17e-15 |
| Fibroblast | F3 COL11A1+ mCAF | EMP3 | 0.59 | 0.816 | 0.569 | 7.56e-11 | 2.55e-06 |
| Fibroblast | F3 COL11A1+ mCAF | HEBP1 | 0.59 | 0.524 | 0.302 | 9.58e-08 | 0.00323 |
| Fibroblast | F3 COL11A1+ mCAF | TMEM160 | 0.586 | 0.481 | 0.259 | 9.74e-08 | 0.00328 |
| Fibroblast | F3 COL11A1+ mCAF | PRDX4 | 0.585 | 0.643 | 0.395 | 1.96e-08 | 0.00066 |
| Fibroblast | F3 COL11A1+ mCAF | LINC01420 | 0.584 | 0.589 | 0.337 | 5.24e-08 | 0.00177 |
| Fibroblast | F3 COL11A1+ mCAF | XRN2 | 0.584 | 0.432 | 0.227 | 4.02e-07 | 0.0135 |
| Fibroblast | F3 COL11A1+ mCAF | COL6A2 | 0.583 | 0.989 | 0.844 | 7.17e-10 | 2.42e-05 |
| Fibroblast | F3 COL11A1+ mCAF | DNAJC1 | 0.582 | 0.411 | 0.22 | 1.22e-06 | 0.0412 |
| Fibroblast | F3 COL11A1+ mCAF | SLC39A1 | 0.579 | 0.584 | 0.354 | 3.1e-07 | 0.0104 |
| Fibroblast | F3 COL11A1+ mCAF | HEXA | 0.579 | 0.47 | 0.265 | 9.59e-07 | 0.0323 |
| Fibroblast | F3 COL11A1+ mCAF | ACTN1 | 0.571 | 0.714 | 0.46 | 3.33e-08 | 0.00112 |

| compartment | subcluster | gene | avg_log2FC | pct.1 | pct.2 | p_val | p_val_adj |
|---|---|---|---|---|---|---|---|
| Fibroblast | F3 COL11A1+ mCAF | ALG5 | 0.571 | 0.4 | 0.205 | 4.8e-07 | 0.0162 |
| Fibroblast | F3 COL11A1+ mCAF | MRPL14 | 0.567 | 0.47 | 0.26 | 1.82e-07 | 0.00612 |
| Fibroblast | F3 COL11A1+ mCAF | PSMD8 | 0.558 | 0.67 | 0.4 | 9.57e-09 | 0.000323 |
| Fibroblast | F3 COL11A1+ mCAF | HMGN1 | 0.558 | 0.795 | 0.518 | 1.33e-08 | 0.000448 |
| Fibroblast | F3 COL11A1+ mCAF | SFXN3 | 0.557 | 0.259 | 0.107 | 3.87e-07 | 0.013 |
| Fibroblast | F3 COL11A1+ mCAF | FAM195B | 0.555 | 0.627 | 0.383 | 5.15e-08 | 0.00173 |
| Fibroblast | F3 COL11A1+ mCAF | TMEM98 | 0.554 | 0.703 | 0.408 | 1.83e-09 | 6.18e-05 |
| Fibroblast | F3 COL11A1+ mCAF | ARPC5L | 0.554 | 0.368 | 0.173 | 1.04e-07 | 0.0035 |
| Fibroblast | F3 COL11A1+ mCAF | MARCKSL1 | 0.552 | 0.632 | 0.35 | 9.5e-11 | 3.2e-06 |
| Fibroblast | F3 COL11A1+ mCAF | TSR3 | 0.552 | 0.373 | 0.181 | 3.7e-07 | 0.0125 |
| Fibroblast | F3 COL11A1+ mCAF | FXYD5 | 0.55 | 0.573 | 0.315 | 2.8e-08 | 0.000944 |
| Fibroblast | F3 COL11A1+ mCAF | COMT | 0.549 | 0.616 | 0.376 | 3.62e-08 | 0.00122 |
| Fibroblast | F3 COL11A1+ mCAF | MPLKIP | 0.546 | 0.449 | 0.241 | 7.37e-07 | 0.0248 |
| Fibroblast | F3 COL11A1+ mCAF | NUTF2 | 0.546 | 0.486 | 0.283 | 9.94e-07 | 0.0335 |
| Fibroblast | F3 COL11A1+ mCAF | ARHGDIA | 0.545 | 0.741 | 0.431 | 9.85e-10 | 3.32e-05 |
| Fibroblast | F3 COL11A1+ mCAF | CACYBP | 0.543 | 0.551 | 0.293 | 1.99e-09 | 6.69e-05 |
| Fibroblast | F3 COL11A1+ mCAF | VCP | 0.543 | 0.514 | 0.282 | 1.99e-07 | 0.00671 |
| Fibroblast | F3 COL11A1+ mCAF | NDUFA4 | 0.541 | 0.957 | 0.794 | 2.33e-11 | 7.86e-07 |
| Fibroblast | F3 COL11A1+ mCAF | DAD1 | 0.541 | 0.859 | 0.622 | 3.14e-09 | 0.000106 |
| Fibroblast | F3 COL11A1+ mCAF | FIBIN | 0.541 | 0.368 | 0.186 | 6.65e-07 | 0.0224 |
| Fibroblast | F3 COL11A1+ mCAF | FBN1 | 0.54 | 0.497 | 0.282 | 5.27e-07 | 0.0178 |
| Fibroblast | F3 COL11A1+ mCAF | LYPLA2 | 0.538 | 0.211 | 0.082 | 1.03e-06 | 0.0347 |
| Fibroblast | F3 COL11A1+ mCAF | ANXA5 | 0.534 | 0.892 | 0.65 | 2.59e-10 | 8.72e-06 |
| Fibroblast | F3 COL11A1+ mCAF | ENC1 | 0.534 | 0.286 | 0.121 | 2.19e-07 | 0.00739 |
| Fibroblast | F3 COL11A1+ mCAF | WDR1 | 0.531 | 0.6 | 0.346 | 2.81e-08 | 0.000948 |
| Fibroblast | F3 COL11A1+ mCAF | MRPL41 | 0.529 | 0.676 | 0.404 | 1.88e-08 | 0.000633 |
| Fibroblast | F3 COL11A1+ mCAF | CDC42 | 0.529 | 0.827 | 0.567 | 6.41e-08 | 0.00216 |
| Fibroblast | F3 COL11A1+ mCAF | APRT | 0.527 | 0.778 | 0.467 | 1.05e-08 | 0.000352 |
| Fibroblast | F3 COL11A1+ mCAF | AP3S1 | 0.527 | 0.703 | 0.469 | 6.64e-07 | 0.0224 |
| Fibroblast | F3 COL11A1+ mCAF | TM9SF3 | 0.526 | 0.443 | 0.239 | 8.81e-07 | 0.0297 |
| Fibroblast | F3 COL11A1+ mCAF | CDK4 | 0.526 | 0.557 | 0.33 | 1.24e-06 | 0.0419 |
| Fibroblast | F3 COL11A1+ mCAF | ZNF706 | 0.525 | 0.703 | 0.47 | 2e-07 | 0.00674 |
| Fibroblast | F3 COL11A1+ mCAF | PGLS | 0.525 | 0.573 | 0.35 | 1.19e-06 | 0.0401 |
| Fibroblast | F3 COL11A1+ mCAF | UBE2L3 | 0.523 | 0.659 | 0.394 | 2.05e-07 | 0.0069 |
| Fibroblast | F3 COL11A1+ mCAF | CNIH4 | 0.522 | 0.497 | 0.262 | 1.12e-07 | 0.00376 |
| Fibroblast | F3 COL11A1+ mCAF | DBI | 0.52 | 0.789 | 0.497 | 4.45e-10 | 1.5e-05 |
| Fibroblast | F3 COL11A1+ mCAF | SSR2 | 0.519 | 0.914 | 0.679 | 1.17e-10 | 3.94e-06 |
| Fibroblast | F3 COL11A1+ mCAF | CBX1 | 0.518 | 0.416 | 0.224 | 1.34e-06 | 0.0453 |
| Fibroblast | F3 COL11A1+ mCAF | PSMA1 | 0.517 | 0.757 | 0.459 | 1.54e-09 | 5.19e-05 |
| Fibroblast | F3 COL11A1+ mCAF | ANXA2 | 0.516 | 0.924 | 0.715 | 5.71e-10 | 1.93e-05 |
| Fibroblast | F3 COL11A1+ mCAF | CFL1 | 0.515 | 0.978 | 0.878 | 6.22e-12 | 2.1e-07 |
| Fibroblast | F3 COL11A1+ mCAF | HLA-A | 0.515 | 0.995 | 0.893 | 3.21e-10 | 1.08e-05 |
| Fibroblast | F3 COL11A1+ mCAF | OST4 | 0.514 | 0.946 | 0.825 | 1.84e-11 | 6.19e-07 |
| Fibroblast | F3 COL11A1+ mCAF | GNB1 | 0.513 | 0.362 | 0.17 | 1.93e-07 | 0.00651 |
| Fibroblast | F3 COL11A1+ mCAF | GTF3C6 | 0.51 | 0.535 | 0.277 | 4.47e-08 | 0.00151 |
| Fibroblast | F3 COL11A1+ mCAF | RRBP1 | 0.509 | 0.681 | 0.426 | 7.07e-08 | 0.00238 |
| Fibroblast | F3 COL11A1+ mCAF | TMED10 | 0.508 | 0.686 | 0.463 | 4.48e-07 | 0.0151 |
| Fibroblast | F3 COL11A1+ mCAF | EDIL3 | 0.504 | 0.308 | 0.135 | 3.02e-07 | 0.0102 |
| Fibroblast | F3 COL11A1+ mCAF | B2M | 0.502 | 1.0 | 0.999 | 2.37e-18 | 8e-14 |
| Fibroblast | F3 COL11A1+ mCAF | LAMTOR1 | 0.501 | 0.708 | 0.465 | 6.34e-07 | 0.0214 |
| Fibroblast | F3 COL11A1+ mCAF | PFN1 | 0.499 | 0.984 | 0.888 | 4.72e-13 | 1.59e-08 |
| Fibroblast | F3 COL11A1+ mCAF | SH3PXD2A | 0.498 | 0.351 | 0.167 | 2.68e-07 | 0.00903 |
| Fibroblast | F3 COL11A1+ mCAF | CBX3 | 0.497 | 0.551 | 0.336 | 6.06e-07 | 0.0204 |
| Fibroblast | F3 COL11A1+ mCAF | TMEM147 | 0.496 | 0.686 | 0.407 | 3.33e-08 | 0.00112 |
| Fibroblast | F3 COL11A1+ mCAF | HSPA6 | 0.496 | 0.454 | 0.241 | 6.32e-08 | 0.00213 |
| Fibroblast | F3 COL11A1+ mCAF | SF3B5 | 0.495 | 0.805 | 0.534 | 1.77e-09 | 5.95e-05 |
| Fibroblast | F3 COL11A1+ mCAF | GOLGB1 | 0.494 | 0.368 | 0.176 | 1.21e-07 | 0.00408 |
| Fibroblast | F3 COL11A1+ mCAF | CLIC4 | 0.494 | 0.503 | 0.278 | 3.99e-07 | 0.0134 |
| Fibroblast | F3 COL11A1+ mCAF | RHOC | 0.493 | 0.865 | 0.637 | 1.26e-08 | 0.000423 |
| Fibroblast | F3 COL11A1+ mCAF | YWHAB | 0.492 | 0.789 | 0.531 | 1.44e-07 | 0.00486 |
| Fibroblast | F3 COL11A1+ mCAF | TMEM248 | 0.492 | 0.503 | 0.279 | 1.35e-06 | 0.0456 |
| Fibroblast | F3 COL11A1+ mCAF | MINOS1 | 0.49 | 0.865 | 0.606 | 2.12e-08 | 0.000714 |
| Fibroblast | F3 COL11A1+ mCAF | RAB2A | 0.488 | 0.778 | 0.516 | 5.07e-07 | 0.0171 |
| Fibroblast | F3 COL11A1+ mCAF | MSRB2 | 0.487 | 0.557 | 0.307 | 5.05e-07 | 0.017 |

| compartment | subcluster | gene | avg_log2FC | pct.1 | pct.2 | p_val | p_val_adj |
|---|---|---|---|---|---|---|---|
| Fibroblast | F3 COL11A1+ mCAF | APP | 0.485 | 0.811 | 0.573 | 6.64e-08 | 0.00224 |
| Fibroblast | F3 COL11A1+ mCAF | C19orf24 | 0.481 | 0.459 | 0.254 | 1.33e-06 | 0.0449 |
| Fibroblast | F3 COL11A1+ mCAF | CSTB | 0.48 | 0.843 | 0.576 | 2.88e-08 | 0.00097 |
| Fibroblast | F3 COL11A1+ mCAF | MYL12B | 0.479 | 0.957 | 0.756 | 2.33e-08 | 0.000785 |
| Fibroblast | F3 COL11A1+ mCAF | DKK3 | 0.477 | 0.622 | 0.367 | 2.26e-07 | 0.00761 |
| Fibroblast | F3 COL11A1+ mCAF | ATP6V1F | 0.477 | 0.735 | 0.466 | 3.58e-07 | 0.0121 |
| Fibroblast | F3 COL11A1+ mCAF | PSMA7 | 0.468 | 0.876 | 0.646 | 9.87e-09 | 0.000333 |
| Fibroblast | F3 COL11A1+ mCAF | PHF14 | 0.467 | 0.535 | 0.298 | 5.16e-07 | 0.0174 |
| Fibroblast | F3 COL11A1+ mCAF | TCEB2 | 0.465 | 0.946 | 0.788 | 7.36e-10 | 2.48e-05 |
| Fibroblast | F3 COL11A1+ mCAF | TFG | 0.463 | 0.524 | 0.293 | 8.09e-07 | 0.0273 |
| Fibroblast | F3 COL11A1+ mCAF | MRPL28 | 0.463 | 0.416 | 0.217 | 8.09e-07 | 0.0273 |
| Fibroblast | F3 COL11A1+ mCAF | SFRP4 | 0.46 | 0.578 | 0.343 | 2.01e-07 | 0.00677 |
| Fibroblast | F3 COL11A1+ mCAF | EDF1 | 0.459 | 0.951 | 0.802 | 1.45e-10 | 4.89e-06 |
| Fibroblast | F3 COL11A1+ mCAF | METTL9 | 0.459 | 0.605 | 0.372 | 8.54e-07 | 0.0288 |
| Fibroblast | F3 COL11A1+ mCAF | CYP1B1 | 0.457 | 0.368 | 0.181 | 3.46e-07 | 0.0117 |
| Fibroblast | F3 COL11A1+ mCAF | RPL28 | 0.456 | 1.0 | 0.978 | 7.57e-19 | 2.55e-14 |
| Fibroblast | F3 COL11A1+ mCAF | LAMTOR2 | 0.456 | 0.578 | 0.335 | 4.45e-07 | 0.015 |
| Fibroblast | F3 COL11A1+ mCAF | ARL2BP | 0.447 | 0.481 | 0.253 | 2.25e-07 | 0.00758 |
| Fibroblast | F3 COL11A1+ mCAF | C20orf24 | 0.442 | 0.449 | 0.239 | 9.58e-07 | 0.0323 |
| Fibroblast | F3 COL11A1+ mCAF | C11orf31 | 0.44 | 0.784 | 0.537 | 2.72e-07 | 0.00916 |
| Fibroblast | F3 COL11A1+ mCAF | TXN | 0.438 | 0.935 | 0.67 | 8.88e-08 | 0.00299 |
| Fibroblast | F3 COL11A1+ mCAF | GDI2 | 0.436 | 0.784 | 0.486 | 1.47e-07 | 0.00497 |
| Fibroblast | F3 COL11A1+ mCAF | HSPE1 | 0.436 | 0.854 | 0.595 | 2.18e-07 | 0.00736 |
| Fibroblast | F3 COL11A1+ mCAF | ATP5J2 | 0.436 | 0.789 | 0.54 | 4.77e-07 | 0.0161 |
| Fibroblast | F3 COL11A1+ mCAF | POLR2L | 0.43 | 0.908 | 0.647 | 4.22e-10 | 1.42e-05 |
| Fibroblast | F3 COL11A1+ mCAF | RABAC1 | 0.43 | 0.892 | 0.702 | 2.77e-07 | 0.00934 |
| Fibroblast | F3 COL11A1+ mCAF | TMEM258 | 0.426 | 0.935 | 0.698 | 9.74e-09 | 0.000328 |
| Fibroblast | F3 COL11A1+ mCAF | IFI27L2 | 0.426 | 0.789 | 0.53 | 7.02e-08 | 0.00236 |
| Fibroblast | F3 COL11A1+ mCAF | FKBP1A | 0.426 | 0.859 | 0.622 | 7.24e-07 | 0.0244 |
| Fibroblast | F3 COL11A1+ mCAF | PPP1CA | 0.425 | 0.524 | 0.31 | 1.46e-06 | 0.049 |
| Fibroblast | F3 COL11A1+ mCAF | SUMO1 | 0.424 | 0.768 | 0.498 | 2.46e-07 | 0.00827 |
| Fibroblast | F3 COL11A1+ mCAF | UBXN4 | 0.419 | 0.649 | 0.38 | 4.89e-07 | 0.0165 |
| Fibroblast | F3 COL11A1+ mCAF | GNG5 | 0.419 | 0.8 | 0.533 | 1.29e-06 | 0.0436 |
| Fibroblast | F3 COL11A1+ mCAF | DNAJC15 | 0.418 | 0.6 | 0.362 | 4.43e-07 | 0.0149 |
| Fibroblast | F3 COL11A1+ mCAF | MRPL51 | 0.417 | 0.768 | 0.505 | 1.82e-07 | 0.00612 |
| Fibroblast | F3 COL11A1+ mCAF | SERF2 | 0.416 | 1.0 | 0.948 | 1.37e-12 | 4.61e-08 |
| Fibroblast | F3 COL11A1+ mCAF | KRTCAP2 | 0.416 | 0.784 | 0.529 | 6.95e-08 | 0.00234 |
| Fibroblast | F3 COL11A1+ mCAF | ARPC1B | 0.416 | 0.768 | 0.512 | 1e-06 | 0.0338 |
| Fibroblast | F3 COL11A1+ mCAF | SF3B1 | 0.41 | 0.535 | 0.298 | 2.95e-07 | 0.00995 |
| Fibroblast | F3 COL11A1+ mCAF | SEC61B | 0.402 | 0.903 | 0.676 | 2.3e-07 | 0.00775 |
| Fibroblast | F3 COL11A1+ mCAF | ISG15 | 0.394 | 0.703 | 0.441 | 6.14e-08 | 0.00207 |
| Fibroblast | F3 COL11A1+ mCAF | FBXW5 | 0.391 | 0.454 | 0.251 | 1.22e-06 | 0.041 |
| Fibroblast | F3 COL11A1+ mCAF | PSMA2 | 0.39 | 0.751 | 0.525 | 5.48e-07 | 0.0185 |
| Fibroblast | F3 COL11A1+ mCAF | AURKAIP1 | 0.387 | 0.703 | 0.447 | 5.48e-07 | 0.0185 |
| Fibroblast | F3 COL11A1+ mCAF | LRP1 | 0.368 | 0.719 | 0.437 | 3.53e-07 | 0.0119 |
| Fibroblast | F3 COL11A1+ mCAF | GUK1 | 0.367 | 0.897 | 0.735 | 3.15e-08 | 0.00106 |
| Fibroblast | F3 COL11A1+ mCAF | RAC1 | 0.362 | 0.978 | 0.819 | 3.44e-08 | 0.00116 |
| Fibroblast | F3 COL11A1+ mCAF | SSR4 | 0.362 | 0.935 | 0.702 | 5.62e-07 | 0.0189 |
| Fibroblast | F3 COL11A1+ mCAF | CLIC1 | 0.361 | 0.924 | 0.714 | 2.78e-07 | 0.00937 |
| Fibroblast | F3 COL11A1+ mCAF | CD63 | 0.344 | 1.0 | 0.925 | 4.01e-08 | 0.00135 |
| Fibroblast | F3 COL11A1+ mCAF | RBX1 | 0.342 | 0.773 | 0.528 | 1.4e-06 | 0.0471 |
| Fibroblast | F3 COL11A1+ mCAF | LINC00152 | 0.324 | 0.605 | 0.368 | 2.38e-07 | 0.00803 |
| Fibroblast | F3 COL11A1+ mCAF | MEG3 | 0.299 | 0.762 | 0.452 | 4.01e-09 | 0.000135 |
| Fibroblast | F3 COL11A1+ mCAF | RPS18 | 0.294 | 1.0 | 0.982 | 1.79e-10 | 6.02e-06 |
| Fibroblast | F3 COL11A1+ mCAF | RPS19 | 0.288 | 1.0 | 0.986 | 3.41e-08 | 0.00115 |
| Fibroblast | F3 COL11A1+ mCAF | COL4A2 | 0.282 | 0.638 | 0.409 | 7.03e-07 | 0.0237 |
| Fibroblast | F3 COL11A1+ mCAF | RPL35 | 0.265 | 1.0 | 0.961 | 2.72e-08 | 0.000917 |
| Fibroblast | F3 COL11A1+ mCAF | RPS17 | 0.264 | 1.0 | 0.957 | 3.6e-08 | 0.00121 |
| Fibroblast | F4 ADAMDEC1+ Fib | ADAMDEC1 | 6.603 | 0.976 | 0.048 | 1.3e-104 | 4.37e-100 |
| Fibroblast | F4 ADAMDEC1+ Fib | CCL13 | 6.594 | 0.512 | 0.037 | 3.71e-40 | 1.25e-35 |
| Fibroblast | F4 ADAMDEC1+ Fib | VAT1L | 6.536 | 0.22 | 0.003 | 8.19e-37 | 2.76e-32 |
| Fibroblast | F4 ADAMDEC1+ Fib | HAPLN1 | 6.484 | 0.707 | 0.019 | 2.08e-95 | 7e-91 |
| Fibroblast | F4 ADAMDEC1+ Fib | EPHA7 | 6.026 | 0.171 | 0.004 | 2.08e-24 | 7.01e-20 |
| Fibroblast | F4 ADAMDEC1+ Fib | ADAM28 | 5.911 | 0.756 | 0.031 | 5.92e-85 | 2e-80 |

| compartment | subcluster | gene | avg_log2FC | pct.1 | pct.2 | p_val | p_val_adj |
|---|---|---|---|---|---|---|---|
| Fibroblast | F4 ADAMDEC1+ Fib | RBFOX1 | 5.773 | 0.122 | 0.003 | 1.81e-17 | 6.08e-13 |
| Fibroblast | F4 ADAMDEC1+ Fib | STMN2 | 4.998 | 0.439 | 0.03 | 2.28e-35 | 7.690000000000001e-31 |
| Fibroblast | F4 ADAMDEC1+ Fib | BMP5 | 4.956 | 0.39 | 0.014 | 8.279999999999999e-45 | 2.79e-40 |
| Fibroblast | F4 ADAMDEC1+ Fib | ENPP6 | 4.928 | 0.146 | 0.004 | 9.09e-20 | 3.06e-15 |
| Fibroblast | F4 ADAMDEC1+ Fib | CLEC14A | 4.873 | 0.122 | 0.008 | 1.39e-10 | 4.68e-06 |
| Fibroblast | F4 ADAMDEC1+ Fib | CXCL6 | 4.84 | 0.366 | 0.024 | 1.48e-29 | 4.98e-25 |
| Fibroblast | F4 ADAMDEC1+ Fib | P2RY14 | 4.832 | 0.366 | 0.022 | 9.430000000000001e-32 | 3.18e-27 |
| Fibroblast | F4 ADAMDEC1+ Fib | DACH1 | 4.803 | 0.146 | 0.01 | 3.66e-12 | 1.23e-07 |
| Fibroblast | F4 ADAMDEC1+ Fib | PCDH19 | 4.752 | 0.122 | 0.007 | 1.56e-11 | 5.26e-07 |
| Fibroblast | F4 ADAMDEC1+ Fib | TFPI2 | 4.723 | 0.61 | 0.05 | 1.77e-44 | 5.95e-40 |
| Fibroblast | F4 ADAMDEC1+ Fib | PAQR5 | 4.655 | 0.122 | 0.005 | 7.46e-14 | 2.51e-09 |
| Fibroblast | F4 ADAMDEC1+ Fib | CCL7 | 4.629 | 0.171 | 0.006 | 2.27e-20 | 7.64e-16 |
| Fibroblast | F4 ADAMDEC1+ Fib | PSD4 | 4.624 | 0.122 | 0.006 | 1.5e-12 | 5.05e-08 |
| Fibroblast | F4 ADAMDEC1+ Fib | SFTA1P | 4.59 | 0.317 | 0.015 | 1.66e-31 | 5.6e-27 |
| Fibroblast | F4 ADAMDEC1+ Fib | RAB38 | 4.518 | 0.122 | 0.007 | 1.66e-11 | 5.58e-07 |
| Fibroblast | F4 ADAMDEC1+ Fib | CCL8 | 4.447 | 0.195 | 0.029 | 1.82e-08 | 0.000613 |
| Fibroblast | F4 ADAMDEC1+ Fib | FAM105A | 4.417 | 0.366 | 0.031 | 1.32e-24 | 4.45e-20 |
| Fibroblast | F4 ADAMDEC1+ Fib | MCTP1 | 4.416 | 0.146 | 0.007 | 5.32e-15 | 1.79e-10 |
| Fibroblast | F4 ADAMDEC1+ Fib | LINC01082 | 4.229 | 0.756 | 0.092 | 9.9e-42 | 3.34e-37 |
| Fibroblast | F4 ADAMDEC1+ Fib | CCL11 | 4.188 | 0.854 | 0.137 | 2.17e-40 | 7.31e-36 |
| Fibroblast | F4 ADAMDEC1+ Fib | SCARA5 | 4.086 | 0.707 | 0.088 | 1.51e-38 | 5.1e-34 |
| Fibroblast | F4 ADAMDEC1+ Fib | SCT | 4.006 | 0.195 | 0.02 | 1.03e-11 | 3.48e-07 |
| Fibroblast | F4 ADAMDEC1+ Fib | FDCSP | 3.884 | 0.195 | 0.008 | 2.24e-21 | 7.54e-17 |
| Fibroblast | F4 ADAMDEC1+ Fib | ADGRL3 | 3.86 | 0.268 | 0.017 | 3.93e-22 | 1.32e-17 |
| Fibroblast | F4 ADAMDEC1+ Fib | HOXB-AS3 | 3.797 | 0.366 | 0.036 | 3.56e-22 | 1.2e-17 |
| Fibroblast | F4 ADAMDEC1+ Fib | PTGER2 | 3.779 | 0.537 | 0.068 | 4.43e-28 | 1.49e-23 |
| Fibroblast | F4 ADAMDEC1+ Fib | ZNF536 | 3.608 | 0.122 | 0.004 | 2.08e-15 | 6.99e-11 |
| Fibroblast | F4 ADAMDEC1+ Fib | FOXF2 | 3.535 | 0.293 | 0.026 | 9.27e-19 | 3.12e-14 |
| Fibroblast | F4 ADAMDEC1+ Fib | CXCL1 | 3.508 | 0.829 | 0.166 | 4.17e-31 | 1.41e-26 |
| Fibroblast | F4 ADAMDEC1+ Fib | ISL1 | 3.481 | 0.122 | 0.014 | 4.16e-07 | 0.014 |
| Fibroblast | F4 ADAMDEC1+ Fib | FABP5 | 3.48 | 0.659 | 0.242 | 1.58e-12 | 5.31e-08 |
| Fibroblast | F4 ADAMDEC1+ Fib | NCAM1 | 3.424 | 0.171 | 0.016 | 4.86e-11 | 1.64e-06 |
| Fibroblast | F4 ADAMDEC1+ Fib | RPS6KA2 | 3.362 | 0.244 | 0.029 | 1.11e-12 | 3.74e-08 |
| Fibroblast | F4 ADAMDEC1+ Fib | PITX1 | 3.3 | 0.585 | 0.079 | 2.76e-28 | 9.289999999999999e-24 |
| Fibroblast | F4 ADAMDEC1+ Fib | WNT2B | 3.275 | 0.415 | 0.067 | 8.14e-17 | 2.74e-12 |
| Fibroblast | F4 ADAMDEC1+ Fib | ATOH8 | 3.272 | 0.146 | 0.018 | 8.89e-08 | 0.003 |
| Fibroblast | F4 ADAMDEC1+ Fib | FAM150B | 3.262 | 0.146 | 0.015 | 5.18e-09 | 0.000175 |
| Fibroblast | F4 ADAMDEC1+ Fib | C5orf66-AS1 | 3.26 | 0.171 | 0.023 | 2.94e-08 | 0.00099 |
| Fibroblast | F4 ADAMDEC1+ Fib | TOX | 3.249 | 0.122 | 0.012 | 5.53e-08 | 0.00186 |
| Fibroblast | F4 ADAMDEC1+ Fib | FGFR4 | 3.206 | 0.244 | 0.019 | 1.64e-17 | 5.54e-13 |
| Fibroblast | F4 ADAMDEC1+ Fib | FENDRR | 3.196 | 0.268 | 0.049 | 8.96e-10 | 3.02e-05 |
| Fibroblast | F4 ADAMDEC1+ Fib | EDIL3 | 3.178 | 0.659 | 0.146 | 1.3e-20 | 4.37e-16 |
| Fibroblast | F4 ADAMDEC1+ Fib | EMID1 | 3.157 | 0.268 | 0.05 | 2.09e-09 | 7.04e-05 |
| Fibroblast | F4 ADAMDEC1+ Fib | HOXD11 | 3.135 | 0.146 | 0.019 | 1.94e-07 | 0.00652 |
| Fibroblast | F4 ADAMDEC1+ Fib | PDE7B | 3.127 | 0.244 | 0.03 | 2.5e-12 | 8.41e-08 |
| Fibroblast | F4 ADAMDEC1+ Fib | EFCC1 | 3.103 | 0.244 | 0.038 | 4.73e-10 | 1.59e-05 |
| Fibroblast | F4 ADAMDEC1+ Fib | IRF8 | 3.083 | 0.366 | 0.066 | 5.72e-13 | 1.93e-08 |
| Fibroblast | F4 ADAMDEC1+ Fib | HS3ST3B1 | 3.052 | 0.146 | 0.021 | 7.5e-07 | 0.0253 |
| Fibroblast | F4 ADAMDEC1+ Fib | HHIP | 3.04 | 0.171 | 0.03 | 1.39e-06 | 0.047 |
| Fibroblast | F4 ADAMDEC1+ Fib | TCF21 | 3.036 | 0.829 | 0.274 | 1.35e-20 | 4.54e-16 |
| Fibroblast | F4 ADAMDEC1+ Fib | FLJ37035 | 3.033 | 0.146 | 0.02 | 4.01e-07 | 0.0135 |
| Fibroblast | F4 ADAMDEC1+ Fib | FMO4 | 3.03 | 0.146 | 0.019 | 2.09e-07 | 0.00704 |
| Fibroblast | F4 ADAMDEC1+ Fib | CTSC | 2.985 | 0.902 | 0.428 | 1.12e-18 | 3.77e-14 |
| Fibroblast | F4 ADAMDEC1+ Fib | PRKAR2B | 2.958 | 0.293 | 0.054 | 2.88e-10 | 9.7e-06 |
| Fibroblast | F4 ADAMDEC1+ Fib | DLL1 | 2.949 | 0.341 | 0.08 | 1.79e-09 | 6.04e-05 |
| Fibroblast | F4 ADAMDEC1+ Fib | APOE | 2.938 | 0.951 | 0.53 | 9.04e-18 | 3.04e-13 |
| Fibroblast | F4 ADAMDEC1+ Fib | ST8SIA1 | 2.916 | 0.195 | 0.029 | 1.85e-08 | 0.000625 |
| Fibroblast | F4 ADAMDEC1+ Fib | HAAO | 2.902 | 0.585 | 0.108 | 5.230000000000001e-21 | 1.76e-16 |
| Fibroblast | F4 ADAMDEC1+ Fib | TSLP | 2.803 | 0.171 | 0.028 | 7.1e-07 | 0.0239 |
| Fibroblast | F4 ADAMDEC1+ Fib | CFD | 2.745 | 1.0 | 0.361 | 7.59e-26 | 2.56e-21 |
| Fibroblast | F4 ADAMDEC1+ Fib | CCL2 | 2.736 | 0.951 | 0.341 | 4.5e-20 | 1.52e-15 |
| Fibroblast | F4 ADAMDEC1+ Fib | PAPPA | 2.718 | 0.366 | 0.073 | 1.09e-11 | 3.67e-07 |
| Fibroblast | F4 ADAMDEC1+ Fib | POLA2 | 2.617 | 0.171 | 0.022 | 1.93e-08 | 0.000649 |
| Fibroblast | F4 ADAMDEC1+ Fib | SMPDL3A | 2.494 | 0.415 | 0.104 | 2.14e-10 | 7.21e-06 |

| compartment | subcluster | gene | avg_log2FC | pct.1 | pct.2 | p_val | p_val_adj |
|---|---|---|---|---|---|---|---|
| Fibroblast | F4 ADAMDEC1+ Fib | NEGR1 | 2.491 | 0.488 | 0.119 | 5.73e-13 | 1.93e-08 |
| Fibroblast | F4 ADAMDEC1+ Fib | TMEM150C | 2.405 | 0.244 | 0.049 | 7.57e-08 | 0.00255 |
| Fibroblast | F4 ADAMDEC1+ Fib | MATN2 | 2.389 | 0.537 | 0.118 | 3.66e-15 | 1.23e-10 |
| Fibroblast | F4 ADAMDEC1+ Fib | PLPP1 | 2.354 | 0.878 | 0.362 | 1.13e-17 | 3.81e-13 |
| Fibroblast | F4 ADAMDEC1+ Fib | RAB8B | 2.348 | 0.415 | 0.109 | 7.96e-10 | 2.68e-05 |
| Fibroblast | F4 ADAMDEC1+ Fib | HAS2 | 2.33 | 0.317 | 0.067 | 1.87e-09 | 6.29e-05 |
| Fibroblast | F4 ADAMDEC1+ Fib | ALDH16A1 | 2.269 | 0.317 | 0.09 | 6.15e-07 | 0.0207 |
| Fibroblast | F4 ADAMDEC1+ Fib | TM4SF1 | 2.266 | 0.585 | 0.186 | 8.49e-12 | 2.86e-07 |
| Fibroblast | F4 ADAMDEC1+ Fib | TMEM176A | 2.247 | 0.951 | 0.446 | 3.83e-21 | 1.29e-16 |
| Fibroblast | F4 ADAMDEC1+ Fib | NCOA7 | 2.243 | 0.61 | 0.238 | 7.02e-10 | 2.37e-05 |
| Fibroblast | F4 ADAMDEC1+ Fib | ITIH5 | 2.231 | 0.463 | 0.121 | 9.72e-11 | 3.28e-06 |
| Fibroblast | F4 ADAMDEC1+ Fib | EDNRB | 2.192 | 0.463 | 0.156 | 2.46e-08 | 0.000829 |
| Fibroblast | F4 ADAMDEC1+ Fib | IQGAP2 | 2.184 | 0.244 | 0.052 | 2.88e-07 | 0.0097 |
| Fibroblast | F4 ADAMDEC1+ Fib | MUC12 | 2.184 | 0.341 | 0.102 | 5.34e-07 | 0.018 |
| Fibroblast | F4 ADAMDEC1+ Fib | MAPK10 | 2.178 | 0.244 | 0.052 | 3.29e-07 | 0.0111 |
| Fibroblast | F4 ADAMDEC1+ Fib | TMEM176B | 2.142 | 1.0 | 0.511 | 3.47e-21 | 1.17e-16 |
| Fibroblast | F4 ADAMDEC1+ Fib | ECM1 | 2.138 | 0.732 | 0.313 | 5.04e-12 | 1.7e-07 |
| Fibroblast | F4 ADAMDEC1+ Fib | ETS2 | 2.132 | 0.634 | 0.176 | 8.35e-15 | 2.81e-10 |
| Fibroblast | F4 ADAMDEC1+ Fib | A2M | 2.099 | 0.951 | 0.492 | 3.52e-18 | 1.19e-13 |
| Fibroblast | F4 ADAMDEC1+ Fib | RCSD1 | 2.06 | 0.317 | 0.079 | 9.66e-08 | 0.00325 |
| Fibroblast | F4 ADAMDEC1+ Fib | BMP4 | 2.043 | 0.561 | 0.182 | 7.57e-11 | 2.55e-06 |
| Fibroblast | F4 ADAMDEC1+ Fib | PTGDS | 2.038 | 0.927 | 0.346 | 6.79e-17 | 2.29e-12 |
| Fibroblast | F4 ADAMDEC1+ Fib | CASP1 | 2.024 | 0.317 | 0.08 | 1.17e-07 | 0.00395 |
| Fibroblast | F4 ADAMDEC1+ Fib | CXCL14 | 1.998 | 0.902 | 0.324 | 2.33e-18 | 7.85e-14 |
| Fibroblast | F4 ADAMDEC1+ Fib | IL34 | 1.991 | 0.366 | 0.114 | 6.17e-07 | 0.0208 |
| Fibroblast | F4 ADAMDEC1+ Fib | RBP1 | 1.943 | 0.854 | 0.301 | 2.03e-16 | 6.83e-12 |
| Fibroblast | F4 ADAMDEC1+ Fib | GRK5 | 1.924 | 0.488 | 0.142 | 3.59e-10 | 1.21e-05 |
| Fibroblast | F4 ADAMDEC1+ Fib | ADH1B | 1.914 | 0.854 | 0.238 | 7.08e-19 | 2.38e-14 |
| Fibroblast | F4 ADAMDEC1+ Fib | ABCA8 | 1.894 | 0.634 | 0.223 | 3.06e-11 | 1.03e-06 |
| Fibroblast | F4 ADAMDEC1+ Fib | FBLN5 | 1.869 | 0.732 | 0.333 | 9.72e-11 | 3.28e-06 |
| Fibroblast | F4 ADAMDEC1+ Fib | WARS | 1.867 | 0.439 | 0.148 | 1.29e-07 | 0.00435 |
| Fibroblast | F4 ADAMDEC1+ Fib | PLPP3 | 1.846 | 0.634 | 0.294 | 1.39e-08 | 0.000468 |
| Fibroblast | F4 ADAMDEC1+ Fib | CYGB | 1.816 | 0.878 | 0.334 | 1.26e-16 | 4.23e-12 |
| Fibroblast | F4 ADAMDEC1+ Fib | QSOX1 | 1.812 | 0.732 | 0.291 | 2.71e-12 | 9.15e-08 |
| Fibroblast | F4 ADAMDEC1+ Fib | TGFBI | 1.788 | 0.756 | 0.289 | 3.75e-12 | 1.26e-07 |
| Fibroblast | F4 ADAMDEC1+ Fib | BCL3 | 1.782 | 0.439 | 0.16 | 1.08e-06 | 0.0364 |
| Fibroblast | F4 ADAMDEC1+ Fib | FKBP9 | 1.756 | 0.634 | 0.277 | 1.75e-08 | 0.00059 |
| Fibroblast | F4 ADAMDEC1+ Fib | LTBP4 | 1.732 | 0.854 | 0.453 | 7.29e-13 | 2.46e-08 |
| Fibroblast | F4 ADAMDEC1+ Fib | PTN | 1.682 | 0.634 | 0.262 | 7.57e-09 | 0.000255 |
| Fibroblast | F4 ADAMDEC1+ Fib | FABP4 | 1.668 | 0.293 | 0.075 | 6.85e-07 | 0.0231 |
| Fibroblast | F4 ADAMDEC1+ Fib | PGRMC1 | 1.664 | 0.829 | 0.419 | 9.09e-11 | 3.06e-06 |
| Fibroblast | F4 ADAMDEC1+ Fib | PLTP | 1.652 | 0.829 | 0.407 | 6.28e-11 | 2.12e-06 |
| Fibroblast | F4 ADAMDEC1+ Fib | PROCR | 1.605 | 0.585 | 0.257 | 8.6e-08 | 0.0029 |
| Fibroblast | F4 ADAMDEC1+ Fib | PDLIM1 | 1.603 | 0.805 | 0.474 | 3.66e-10 | 1.23e-05 |
| Fibroblast | F4 ADAMDEC1+ Fib | TAC3 | 1.59 | 0.122 | 0.014 | 6.17e-07 | 0.0208 |
| Fibroblast | F4 ADAMDEC1+ Fib | ADD3 | 1.586 | 0.561 | 0.269 | 1.46e-06 | 0.0491 |
| Fibroblast | F4 ADAMDEC1+ Fib | DKK3 | 1.584 | 0.78 | 0.397 | 2.46e-09 | 8.3e-05 |
| Fibroblast | F4 ADAMDEC1+ Fib | CD302 | 1.556 | 0.561 | 0.256 | 1.25e-06 | 0.042 |
| Fibroblast | F4 ADAMDEC1+ Fib | UBE2E3 | 1.543 | 0.683 | 0.395 | 6.36e-07 | 0.0214 |
| Fibroblast | F4 ADAMDEC1+ Fib | LSP1 | 1.53 | 0.683 | 0.267 | 1.67e-09 | 5.62e-05 |
| Fibroblast | F4 ADAMDEC1+ Fib | MFAP4 | 1.471 | 0.976 | 0.623 | 1.03e-13 | 3.48e-09 |
| Fibroblast | F4 ADAMDEC1+ Fib | FHL2 | 1.349 | 0.756 | 0.361 | 5.45e-09 | 0.000184 |
| Fibroblast | F4 ADAMDEC1+ Fib | IGFBP5 | 1.347 | 0.854 | 0.59 | 7.93e-08 | 0.00267 |
| Fibroblast | F4 ADAMDEC1+ Fib | EMILIN1 | 1.342 | 0.902 | 0.558 | 5.5e-10 | 1.85e-05 |
| Fibroblast | F4 ADAMDEC1+ Fib | IRF1 | 1.335 | 0.927 | 0.658 | 4.67e-10 | 1.57e-05 |
| Fibroblast | F4 ADAMDEC1+ Fib | XIST | 1.308 | 0.585 | 0.23 | 8.09e-08 | 0.00273 |
| Fibroblast | F4 ADAMDEC1+ Fib | NFKBIA | 1.184 | 1.0 | 0.87 | 6.76e-11 | 2.28e-06 |
| Fibroblast | F4 ADAMDEC1+ Fib | SEPP1 | 1.171 | 0.951 | 0.555 | 1.02e-07 | 0.00343 |
| Fibroblast | F4 ADAMDEC1+ Fib | SOCS3 | 1.166 | 0.927 | 0.72 | 4.61e-07 | 0.0155 |
| Fibroblast | F4 ADAMDEC1+ Fib | FHL1 | 1.099 | 0.805 | 0.433 | 6.63e-08 | 0.00223 |
| Fibroblast | F4 ADAMDEC1+ Fib | MEG3 | 1.095 | 0.878 | 0.492 | 1.19e-07 | 0.00402 |
| Fibroblast | F4 ADAMDEC1+ Fib | POSTN | 1.068 | 0.61 | 0.218 | 1.93e-08 | 0.000649 |
| Fibroblast | F4 ADAMDEC1+ Fib | FBLN1 | 1.056 | 0.976 | 0.605 | 3.44e-09 | 0.000116 |
| Fibroblast | F4 ADAMDEC1+ Fib | CEBPD | 1.025 | 0.951 | 0.764 | 1.85e-08 | 0.000623 |

| compartment | subcluster | gene | avg_log2FC | pct.1 | pct.2 | p_val | p_val_adj |
|---|---|---|---|---|---|---|---|
| Fibroblast | F4 ADAMDEC1+ Fib | CXCL12 | 0.991 | 0.902 | 0.451 | 2.27e-08 | 0.000766 |
| Fibroblast | F4 ADAMDEC1+ Fib | HLA-C | 0.755 | 1.0 | 0.897 | 5.83e-08 | 0.00196 |
| Fibroblast | F4 ADAMDEC1+ Fib | IGHA1 | 0.728 | 0.829 | 0.533 | 1.29e-06 | 0.0434 |
| Fibroblast | F4 ADAMDEC1+ Fib | JCHAIN | 0.661 | 0.732 | 0.338 | 7.68e-09 | 0.000259 |
| Fibroblast | F4 ADAMDEC1+ Fib | IGHA2 | 0.646 | 0.439 | 0.167 | 8.15e-07 | 0.0275 |
| Fibroblast | F4 ADAMDEC1+ Fib | EEF1A1 | 0.518 | 1.0 | 0.995 | 1.45e-08 | 0.000488 |
| Fibroblast | F4 ADAMDEC1+ Fib | RPS12 | 0.501 | 1.0 | 0.98 | 1.23e-06 | 0.0415 |

### Table S2

Table S2. Gene Ontology biological-process enrichment of subcluster marker genes.

| compartment | subcluster | GO_ID | Description | GeneRatio | BgRatio | FoldEnrichment | pvalue | p_adjust | Count | geneID |
|---|---|---|---|---|---|---|---|---|---|---|
| Malignant epithelial | C0 Is ECs | GO:0050792 | regulation of viral process | 11/175 | 141/18860 | 8.41 | 8.3e-08 | 0.000121 | 11 | LGALS1/CXCL8/IFITM3/ZFP36/IFITM1/HMGA2/BST2/SLPI/CD74/IFITM2/ISG15 |
| Malignant epithelial | C0 Is ECs | GO:0097193 | intrinsic apoptotic signaling pathway | 16/175 | 335/18860 | 5.15 | 9.77e-08 | 0.000121 | 16 | CAV1/DDIT4/PLAUR/MIF/GPX1/PHLDA3/CEBPB/CRIP1/IER3/PMAIP1/BBC3/TNFRSF1A/PPP1R15A/MUC1/CD74… |
| Malignant epithelial | C0 Is ECs | GO:1903900 | regulation of viral life cycle | 10/175 | 115/18860 | 9.37 | 1.18e-07 | 0.000121 | 10 | LGALS1/CXCL8/IFITM3/IFITM1/HMGA2/BST2/SLPI/CD74/IFITM2/ISG15 |
| Malignant epithelial | C0 Is ECs | GO:0050727 | regulation of inflammatory response | 18/175 | 460/18860 | 4.22 | 2.97e-07 | 0.0002 | 18 | LGALS1/NFKBIA/CTSC/SAA1/ANXA1/GPX1/MAP3K8/SOCS3/C3/KLF4/CEBPB/PTGES/BIRC3/IER3/IDO1/TNFRS… |
| Malignant epithelial | C0 Is ECs | GO:0045071 | negative regulation of viral genome replication | 7/175 | 50/18860 | 15.09 | 3.75e-07 | 0.0002 | 7 | IFITM3/IFITM1/HMGA2/BST2/SLPI/IFITM2/ISG15 |
| Malignant epithelial | C0 Is ECs | GO:0002237 | response to molecule of bacterial origin | 16/175 | 375/18860 | 4.6 | 4.48e-07 | 0.0002 | 16 | NFKBIA/CXCL2/CXCL8/MIF/GPX1/ZFP36/CEBPB/LCN2/JUND/ARF6/IDO1/SLPI/LITAF/RIPK2/NFKBIZ/TNFAI… |
| Malignant epithelial | C0 Is ECs | GO:0043122 | regulation of canonical NF-kappaB signal transduction | 15/175 | 332/18860 | 4.87 | 5.07e-07 | 0.0002 | 15 | CAV1/S100A4/LGALS1/NFKBIA/KLF4/SLC20A1/BIRC3/BST2/LITAF/RIPK2/TNFRSF1A/LTBR/SECTM1/CD74/T… |
| Malignant epithelial | C0 Is ECs | GO:0048525 | negative regulation of viral process | 8/175 | 76/18860 | 11.34 | 5.16e-07 | 0.0002 | 8 | IFITM3/ZFP36/IFITM1/HMGA2/BST2/SLPI/IFITM2/ISG15 |
| Malignant epithelial | C0 Is ECs | GO:0045069 | regulation of viral genome replication | 8/175 | 80/18860 | 10.78 | 7.68e-07 | 0.000262 | 8 | CXCL8/IFITM3/IFITM1/HMGA2/BST2/SLPI/IFITM2/ISG15 |
| Malignant epithelial | C0 Is ECs | GO:0008630 | intrinsic apoptotic signaling pathway in response to DNA damage | 9/175 | 111/18860 | 8.74 | 9.32e-07 | 0.000262 | 9 | DDIT4/MIF/PHLDA3/CRIP1/IER3/BBC3/TNFRSF1A/MUC1/CD74 |
| Malignant epithelial | C0 Is ECs | GO:0045444 | fat cell differentiation | 13/175 | 259/18860 | 5.41 | 9.33e-07 | 0.000262 | 13 | CEBPD/NR4A2/GPX1/KLF4/ZFP36/CEBPB/ZFP36L1/HMGA2/ADGRF1/TRIB2/ARL4A/LAMB3/ERO1A |
| Malignant epithelial | C0 Is ECs | GO:0007249 | canonical NF-kappaB signal transduction | 15/175 | 359/18860 | 4.5 | 1.35e-06 | 0.000349 | 15 | CAV1/S100A4/LGALS1/NFKBIA/KLF4/SLC20A1/BIRC3/BST2/LITAF/RIPK2/TNFRSF1A/LTBR/SECTM1/CD74/T… |
| Malignant epithelial | C0 Is ECs | GO:0001666 | response to hypoxia | 14/175 | 317/18860 | 4.76 | 1.61e-06 | 0.000384 | 14 | CAV1/DDIT4/NR4A2/PLAU/LCN2/LIF/ZFP36L1/ANGPTL4/PMAIP1/JUND/DIO3/BBC3/COMT/ERO1A |
| Malignant epithelial | C0 Is ECs | GO:0043123 | positive regulation of canonical NF-kappaB signal transduction | 12/175 | 232/18860 | 5.57 | 1.81e-06 | 0.000401 | 12 | CAV1/S100A4/LGALS1/SLC20A1/BIRC3/BST2/LITAF/RIPK2/TNFRSF1A/LTBR/SECTM1/CD74 |
| Malignant epithelial | C0 Is ECs | GO:0006935 | chemotaxis | 17/175 | 474/18860 | 3.87 | 2.12e-06 | 0.000409 | 17 | TYMP/S100A4/CXCL2/PLAUR/SAA1/ANXA1/CXCL8/MIF/PLAU/CXCL1/DUSP1/CXCL3/C3/CMTM7/CD74/CMTM3/C… |
| Malignant epithelial | C0 Is ECs | GO:0007162 | negative regulation of cell adhesion | 14/175 | 326/18860 | 4.63 | 2.24e-06 | 0.000409 | 14 | LGALS1/MYADM/ANXA1/DUSP1/KLF4/CEBPB/IRF1/IDO1/TACSTD2/MAP4K4/SOCS6/MUC1/CD74/RASA1 |
| Malignant epithelial | C0 Is ECs | GO:0042330 | taxis | 17/175 | 476/18860 | 3.85 | 2.25e-06 | 0.000409 | 17 | TYMP/S100A4/CXCL2/PLAUR/SAA1/ANXA1/CXCL8/MIF/PLAU/CXCL1/DUSP1/CXCL3/C3/CMTM7/CD74/CMTM3/C… |
| Malignant epithelial | C0 Is ECs | GO:0046597 | host-mediated suppression of symbiont invasion | 5/175 | 24/18860 | 22.45 | 2.39e-06 | 0.000411 | 5 | LY6E/IFITM3/IFITM1/CD74/IFITM2 |
| Malignant epithelial | C0 Is ECs | GO:0036293 | response to decreased oxygen levels | 14/175 | 330/18860 | 4.57 | 2.58e-06 | 0.000421 | 14 | CAV1/DDIT4/NR4A2/PLAU/LCN2/LIF/ZFP36L1/ANGPTL4/PMAIP1/JUND/DIO3/BBC3/COMT/ERO1A |

| compartment | subcluster | GO_ID | Description | GeneRatio | BgRatio | FoldEnrichment | pvalue | p_adjust | Count | geneID |
|---|---|---|---|---|---|---|---|---|---|---|
| Malignant epithelial | C0 Is ECs | GO:0019058 | viral life cycle | 13/175 | 298/18860 | 4.7 | 4.4e-06 | 0.000681 | 13 | CAV1/LGALS1/CXCL8/IFITM3/CAV2/IFITM1/HMGA2/BST2/SLPI/ANPEP/CD74/IFITM2/ISG15 |
| Malignant epithelial | C0 Is ECs | GO:0032496 | response to lipopolysaccharide | 14/175 | 350/18860 | 4.31 | 5.1e-06 | 0.000732 | 14 | NFKBIA/CXCL8/MIF/GPX1/ZFP36/CEBPB/LCN2/JUND/IDO1/SLPI/LITAF/RIPK2/NFKBIZ/TNFAIP3 |
| Malignant epithelial | C0 Is ECs | GO:0042771 | intrinsic apoptotic signaling pathway in response to DNA damage by p53 class mediator | 6/175 | 48/18860 | 13.47 | 5.2e-06 | 0.000732 | 6 | DDIT4/MIF/PHLDA3/BBC3/MUC1/CD74 |
| Malignant epithelial | C0 Is ECs | GO:1905954 | positive regulation of lipid localization | 8/175 | 104/18860 | 8.29 | 5.65e-06 | 0.000761 | 8 | CAV1/MIF/C3/PTGES/PLTP/ABCA7/PLIN2/NMB |
| Malignant epithelial | C0 Is ECs | GO:0070482 | response to oxygen levels | 14/175 | 357/18860 | 4.23 | 6.39e-06 | 0.000824 | 14 | CAV1/DDIT4/NR4A2/PLAU/LCN2/LIF/ZFP36L1/ANGPTL4/PMAIP1/JUND/DIO3/BBC3/COMT/ERO1A |
| Malignant epithelial | C0 Is ECs | GO:0051851 | host-mediated perturbation of symbiont process | 8/175 | 110/18860 | 7.84 | 8.58e-06 | 0.00106 | 8 | LY6E/IFITM3/CAV2/IFITM1/HMGA2/SLPI/CD74/IFITM2 |
| Malignant epithelial | C0 Is ECs | GO:0071216 | cellular response to biotic stimulus | 12/175 | 274/18860 | 4.72 | 9.98e-06 | 0.00119 | 12 | NFKBIA/CXCL8/MIF/ZFP36/CEBPB/CDA/LCN2/ARF6/LITAF/RIPK2/NFKBIZ/TNFAIP3 |
| Malignant epithelial | C0 Is ECs | GO:0048144 | fibroblast proliferation | 8/175 | 114/18860 | 7.56 | 1.12e-05 | 0.00128 | 8 | CAV1/MIF/GPX1/EREG/LIF/HMGA2/CD74/TGIF1 |
| Malignant epithelial | C0 Is ECs | GO:0019079 | viral genome replication | 8/175 | 117/18860 | 7.37 | 1.35e-05 | 0.00149 | 8 | CXCL8/IFITM3/IFITM1/HMGA2/BST2/SLPI/IFITM2/ISG15 |
| Malignant epithelial | C0 Is ECs | GO:0048146 | positive regulation of fibroblast proliferation | 6/175 | 57/18860 | 11.34 | 1.44e-05 | 0.00153 | 6 | MIF/EREG/LIF/HMGA2/CD74/TGIF1 |
| Malignant epithelial | C0 Is ECs | GO:0072332 | intrinsic apoptotic signaling pathway by p53 class mediator | 7/175 | 88/18860 | 8.57 | 1.77e-05 | 0.00183 | 7 | DDIT4/MIF/PHLDA3/PMAIP1/BBC3/MUC1/CD74 |
| Malignant epithelial | C0 Is ECs | GO:0048545 | response to steroid hormone | 13/175 | 343/18860 | 4.08 | 1.98e-05 | 0.00183 | 13 | CAV1/DDIT4/ANXA1/DUSP1/ZFP36/JUNB/ZFP36L1/PMEPA1/HMGA2/JUND/FOSB/TFPI/COMT |
| Malignant epithelial | C0 Is ECs | GO:0071219 | cellular response to molecule of bacterial origin | 11/175 | 247/18860 | 4.8 | 2.02e-05 | 0.00183 | 11 | NFKBIA/CXCL8/MIF/ZFP36/CEBPB/LCN2/ARF6/LITAF/RIPK2/NFKBIZ/TNFAIP3 |
| Malignant epithelial | C0 Is ECs | GO:0016032 | viral process | 14/175 | 396/18860 | 3.81 | 2.05e-05 | 0.00183 | 14 | CAV1/LGALS1/CXCL8/IFITM3/ZFP36/CAV2/IFITM1/HMGA2/BST2/SLPI/ANPEP/CD74/IFITM2/ISG15 |
| Malignant epithelial | C0 Is ECs | GO:0007492 | endoderm development | 7/175 | 90/18860 | 8.38 | 2.05e-05 | 0.00183 | 7 | DUSP1/ZFP36L1/HMGA2/LAMA3/DUSP5/MESP1/LAMB3 |
| Malignant epithelial | C0 Is ECs | GO:0035821 | modulation of process of another organism | 8/175 | 124/18860 | 6.95 | 2.07e-05 | 0.00183 | 8 | LY6E/IFITM3/CAV2/IFITM1/HMGA2/SLPI/CD74/IFITM2 |
| Malignant epithelial | C0 Is ECs | GO:0035456 | response to interferon-beta | 5/175 | 37/18860 | 14.56 | 2.23e-05 | 0.00187 | 5 | IFITM3/IFITM1/IRF1/BST2/IFITM2 |
| Malignant epithelial | C0 Is ECs | GO:0046631 | alpha-beta T cell activation | 10/175 | 206/18860 | 5.23 | 2.32e-05 | 0.00187 | 10 | LGALS1/ANXA1/SOCS3/CEBPB/JUNB/CDH26/IRF1/PRDM1/RIPK2/NFKBIZ |
| Malignant epithelial | C0 Is ECs | GO:0001706 | endoderm formation | 6/175 | 62/18860 | 10.43 | 2.34e-05 | 0.00187 | 6 | DUSP1/HMGA2/LAMA3/DUSP5/MESP1/LAMB3 |
| Malignant epithelial | C0 Is ECs | GO:1903037 | regulation of leukocyte cell-cell adhesion | 14/175 | 401/18860 | 3.76 | 2.35e-05 | 0.00187 | 14 | CAV1/LGALS1/ANXA1/MAP3K8/KLF4/CEBPB/IRF1/IDO1/RIPK2/SOCS6/ABL2/NFKBIZ/CD74/HAS2 |
| Malignant epithelial | C0 Is ECs | GO:0050863 | regulation of T cell activation | 14/175 | 404/18860 | 3.73 | 2.55e-05 | 0.00194 | 14 | CAV1/LGALS1/ANXA1/MAP3K8/CEBPB/JUNB/IRF1/IDO1/PRDM1/RIPK2/SOCS6/ABL2/NFKBIZ/CD74 |
| Malignant epithelial | C0 Is ECs | GO:0043616 | keratinocyte proliferation | 6/175 | 63/18860 | 10.26 | 2.57e-05 | 0.00194 | 6 | ZFP36/EREG/ZFP36L1/FAM3C/NFKBIZ/HAS2 |
| Malignant epithelial | C0 Is ECs | GO:2001233 | regulation of apoptotic signaling pathway | 14/175 | 406/18860 | 3.72 | 2.7e-05 | 0.00196 | 14 | CAV1/CTSC/NR4A2/PLAUR/MIF/GPX1/KLF4/IER3/PMAIP1/BBC3/MUC1/LTBR/CD74/TNFAIP3 |
| Malignant epithelial | C0 Is ECs | GO:0048145 | regulation of fibroblast proliferation | 7/175 | 94/18860 | 8.03 | 2.73e-05 | 0.00196 | 7 | CAV1/MIF/EREG/LIF/HMGA2/CD74/TGIF1 |
| Malignant epithelial | C0 Is ECs | GO:0070661 | leukocyte proliferation | 13/175 | 356/18860 | 3.94 | 2.92e-05 | 0.00205 | 13 | CLCF1/ANXA1/MIF/CEBPB/JUNB/IRF1/IDO1/BST2/RIPK2/NFKBIZ/CD74/NMB/TNFAIP3 |
| Malignant epithelial | C0 Is ECs | GO:0002544 | chronic inflammatory response | 4/175 | 21/18860 | 20.53 | 3.79e-05 | 0.00255 | 4 | CEBPB/IDO1/NFKBIZ/TNFAIP3 |

| compartment | subcluster | GO_ID | Description | GeneRatio | BgRatio | FoldEnrichment | pvalue | p_adjust | Count | geneID |
|---|---|---|---|---|---|---|---|---|---|---|
| Malignant epithelial | C0 Is ECs | GO:0035455 | response to interferon-alpha | 4/175 | 21/18860 | 20.53 | 3.79e-05 | 0.00255 | 4 | IFITM3/IFITM1/BST2/IFITM2 |
| Malignant epithelial | C0 Is ECs | GO:0030856 | regulation of epithelial cell differentiation | 9/175 | 177/18860 | 5.48 | 4.18e-05 | 0.0027 | 9 | CAV1/ZFP36/CEBPB/LIF/ZFP36L1/BTG1/TNFRSF1A/MESP1/XDH |
| Malignant epithelial | C0 Is ECs | GO:0043903 | regulation of biological process involved in symbiotic interaction | 5/175 | 42/18860 | 12.83 | 4.18e-05 | 0.0027 | 5 | CAV1/LGALS1/CXCL8/BST2/CD74 |
| Malignant epithelial | C0 Is ECs | GO:0035710 | CD4-positive, alpha-beta T cell activation | 8/175 | 138/18860 | 6.25 | 4.46e-05 | 0.0028 | 8 | LGALS1/ANXA1/SOCS3/CEBPB/JUNB/CDH26/RIPK2/NFKBIZ |
| Malignant epithelial | C0 Is ECs | GO:0050729 | positive regulation of inflammatory response | 9/175 | 179/18860 | 5.42 | 4.56e-05 | 0.0028 | 9 | LGALS1/NFKBIA/CTSC/MAP3K8/C3/CEBPB/IDO1/TNFRSF1A/NFKBIZ |
| Malignant epithelial | C0 Is ECs | GO:0010876 | lipid localization | 15/175 | 483/18860 | 3.35 | 4.62e-05 | 0.0028 | 15 | CAV1/NFKBIA/ANXA1/MIF/VMP1/C3/PTGES/PLTP/NME4/ABCA7/APOL1/PLIN2/NMB/COMT/FABP6 |
| Malignant epithelial | C0 Is ECs | GO:0002819 | regulation of adaptive immune response | 10/175 | 224/18860 | 4.81 | 4.74e-05 | 0.00282 | 10 | LGALS1/CLCF1/ANXA1/C3/JUNB/IRF1/RIPK2/NFKBIZ/SECTM1/TNFAIP3 |
| Malignant epithelial | C0 Is ECs | GO:0071356 | cellular response to tumor necrosis factor | 10/175 | 226/18860 | 4.77 | 5.11e-05 | 0.00298 | 10 | NFKBIA/CXCL8/ZFP36/LCN2/BIRC3/ZFP36L1/KLF2/TNFRSF1A/HAS2/TNFAIP3 |
| Malignant epithelial | C0 Is ECs | GO:1905952 | regulation of lipid localization | 9/175 | 183/18860 | 5.3 | 5.42e-05 | 0.0031 | 9 | CAV1/NFKBIA/MIF/C3/PTGES/PLTP/ABCA7/PLIN2/NMB |
| Malignant epithelial | C0 Is ECs | GO:0071222 | cellular response to lipopolysaccharide | 10/175 | 229/18860 | 4.71 | 5.71e-05 | 0.00317 | 10 | NFKBIA/CXCL8/MIF/ZFP36/CEBPB/LCN2/LITAF/RIPK2/NFKBIZ/TNFAIP3 |
| Malignant epithelial | C0 Is ECs | GO:0046632 | alpha-beta T cell differentiation | 8/175 | 143/18860 | 6.03 | 5.74e-05 | 0.00317 | 8 | LGALS1/ANXA1/SOCS3/JUNB/IRF1/PRDM1/RIPK2/NFKBIZ |
| Malignant epithelial | C0 Is ECs | GO:0050673 | epithelial cell proliferation | 15/175 | 496/18860 | 3.26 | 6.24e-05 | 0.00335 | 15 | CAV1/GPX1/ZFP36/CAV2/SULF2/CEBPB/EREG/ZFP36L1/HMGA2/TACSTD2/FAM3C/PRDM1/NFKBIZ/HAS2/TNFAI… |
| Malignant epithelial | C0 Is ECs | GO:0007159 | leukocyte cell-cell adhesion | 14/175 | 439/18860 | 3.44 | 6.28e-05 | 0.00335 | 14 | CAV1/LGALS1/ANXA1/MAP3K8/KLF4/CEBPB/IRF1/IDO1/RIPK2/SOCS6/ABL2/NFKBIZ/CD74/HAS2 |
| Malignant epithelial | C0 Is ECs | GO:1903706 | regulation of hemopoiesis | 14/175 | 443/18860 | 3.41 | 6.92e-05 | 0.00363 | 14 | LGALS1/NFKBIA/ANXA1/ZFP36/CEBPB/JUNB/LIF/ZFP36L1/IRF1/PRDM1/RIPK2/NFKBIZ/CD74/ISG15 |
| Malignant epithelial | C0 Is ECs | GO:0070663 | regulation of leukocyte proliferation | 11/175 | 284/18860 | 4.17 | 7.2e-05 | 0.00369 | 11 | CLCF1/ANXA1/MIF/CEBPB/IRF1/IDO1/BST2/RIPK2/CD74/NMB/TNFAIP3 |
| Malignant epithelial | C0 Is ECs | GO:0006953 | acute-phase response | 5/175 | 47/18860 | 11.47 | 7.27e-05 | 0.00369 | 5 | SAA1/CEBPB/PTGES/LCN2/SAA2 |
| Malignant epithelial | C0 Is ECs | GO:0042113 | B cell activation | 11/175 | 291/18860 | 4.07 | 8.94e-05 | 0.00434 | 11 | LGALS1/CLCF1/MIF/C3/KLF6/ZFP36L1/CMTM7/BST2/NFKBIZ/CD74/TNFAIP3 |
| Malignant epithelial | C0 Is ECs | GO:0048732 | gland development | 14/175 | 454/18860 | 3.32 | 8.98e-05 | 0.00434 | 14 | CAV1/GPX1/TGM2/SULF2/CEBPB/CRIP1/WNT10A/HMGA2/ARF6/ALDH1A3/ELF3/MESP1/TNFAIP3/XDH |
| Malignant epithelial | C0 Is ECs | GO:2000425 | regulation of apoptotic cell clearance | 3/175 | 10/18860 | 32.33 | 8.98e-05 | 0.00434 | 3 | C3/TGM2/ABCA7 |
| Malignant epithelial | C0 Is ECs | GO:0034612 | response to tumor necrosis factor | 10/175 | 244/18860 | 4.42 | 9.68e-05 | 0.00453 | 10 | NFKBIA/CXCL8/ZFP36/LCN2/BIRC3/ZFP36L1/KLF2/TNFRSF1A/HAS2/TNFAIP3 |
| Malignant epithelial | C0 Is ECs | GO:0006693 | prostaglandin metabolic process | 5/175 | 50/18860 | 10.78 | 9.82e-05 | 0.00453 | 5 | MIF/PTGES/TNFRSF1A/CD74/COMT |
| Malignant epithelial | C0 Is ECs | GO:0045622 | regulation of T-helper cell differentiation | 5/175 | 50/18860 | 10.78 | 9.82e-05 | 0.00453 | 5 | LGALS1/ANXA1/JUNB/RIPK2/NFKBIZ |
| Malignant epithelial | C0 Is ECs | GO:0006869 | lipid transport | 14/175 | 461/18860 | 3.27 | 0.000106 | 0.0048 | 14 | CAV1/NFKBIA/ANXA1/MIF/VMP1/PTGES/PLTP/NME4/ABCA7/APOL1/PLIN2/NMB/COMT/FABP6 |
| Malignant epithelial | C0 Is ECs | GO:0001570 | vasculogenesis | 6/175 | 82/18860 | 7.89 | 0.000114 | 0.00506 | 6 | CAV1/JUNB/ZFP36L1/HAS2/RRAS/RASA1 |
| Malignant epithelial | C0 Is ECs | GO:0046637 | regulation of alpha-beta T cell differentiation | 6/175 | 82/18860 | 7.89 | 0.000114 | 0.00506 | 6 | LGALS1/ANXA1/JUNB/PRDM1/RIPK2/NFKBIZ |
| Malignant epithelial | C0 Is ECs | GO:0006692 | prostanoid metabolic process | 5/175 | 52/18860 | 10.36 | 0.000119 | 0.00516 | 5 | MIF/PTGES/TNFRSF1A/CD74/COMT |

| compartment | subcluster | GO_ID | Description | GeneRatio | BgRatio | FoldEnrichment | pvalue | p_adjust | Count | geneID |
|---|---|---|---|---|---|---|---|---|---|---|
| Malignant epithelial | C0 Is ECs | GO:0007369 | gastrulation | 9/175 | 203/18860 | 4.78 | 0.00012 | 0.00516 | 9 | MYADM/DUSP1/KLF4/HMGA2/LAMA3/DUSP5/MESP1/LAMB3/TGIF1 |
| Malignant epithelial | C0 Is ECs | GO:0052372 | modulation by symbiont of entry into host | 4/175 | 28/18860 | 15.4 | 0.000123 | 0.00522 | 4 | CAV1/LGALS1/CXCL8/CD74 |
| Malignant epithelial | C0 Is ECs | GO:0140467 | integrated stress response signaling | 5/175 | 53/18860 | 10.17 | 0.00013 | 0.00544 | 5 | CEBPD/CEBPB/JUNB/ABCA7/PPP1R15A |
| Malignant epithelial | C0 Is ECs | GO:0032370 | positive regulation of lipid transport | 6/175 | 85/18860 | 7.61 | 0.00014 | 0.00576 | 6 | CAV1/MIF/PTGES/PLTP/ABCA7/NMB |
| Malignant epithelial | C0 Is ECs | GO:0070555 | response to interleukin-1 | 7/175 | 123/18860 | 6.13 | 0.000151 | 0.00606 | 7 | NFKBIA/CXCL8/CEBPB/LCN2/KLF2/RIPK2/HAS2 |
| Malignant epithelial | C0 Is ECs | GO:0002822 | regulation of adaptive immune response based on somatic recombination of immune receptors… | 9/175 | 210/18860 | 4.62 | 0.000155 | 0.00606 | 9 | LGALS1/CLCF1/ANXA1/C3/JUNB/RIPK2/NFKBIZ/SECTM1/TNFAIP3 |
| Malignant epithelial | C0 Is ECs | GO:1903038 | negative regulation of leukocyte cell-cell adhesion | 8/175 | 165/18860 | 5.23 | 0.000156 | 0.00606 | 8 | LGALS1/ANXA1/KLF4/CEBPB/IRF1/IDO1/SOCS6/CD74 |
| Malignant epithelial | C0 Is ECs | GO:0046634 | regulation of alpha-beta T cell activation | 7/175 | 124/18860 | 6.08 | 0.000159 | 0.00606 | 7 | LGALS1/ANXA1/JUNB/IRF1/PRDM1/RIPK2/NFKBIZ |
| Malignant epithelial | C0 Is ECs | GO:0002158 | osteoclast proliferation | 3/175 | 12/18860 | 26.94 | 0.000162 | 0.00606 | 3 | JUNB/NMB/TNFAIP3 |
| Malignant epithelial | C0 Is ECs | GO:0051549 | positive regulation of keratinocyte migration | 3/175 | 12/18860 | 26.94 | 0.000162 | 0.00606 | 3 | ARF6/MAP4K4/HAS2 |
| Malignant epithelial | C0 Is ECs | GO:1900153 | positive regulation of nuclear-transcribed mRNA catabolic process, deadenylation-dependen… | 3/175 | 12/18860 | 26.94 | 0.000162 | 0.00606 | 3 | ZFP36/ZFP36L1/TOB1 |
| Malignant epithelial | C0 Is ECs | GO:0031579 | membrane raft organization | 4/175 | 30/18860 | 14.37 | 0.000163 | 0.00606 | 4 | CAV1/MYADM/CAV2/ABCA7 |
| Malignant epithelial | C0 Is ECs | GO:0031589 | cell-substrate adhesion | 12/175 | 369/18860 | 3.5 | 0.000177 | 0.00652 | 12 | MYADM/PLAU/HACD1/TACSTD2/MAP4K4/RHOD/ITGA10/COL17A1/HAS2/RRAS/RASA1/BCAM |
| Malignant epithelial | C0 Is ECs | GO:0035872 | nucleotide-binding domain, leucine rich repeat containing receptor signaling pathway | 4/175 | 31/18860 | 13.91 | 0.000185 | 0.00674 | 4 | NFKBIA/BIRC3/RIPK2/TNFAIP3 |
| Malignant epithelial | C0 Is ECs | GO:0001701 | in utero embryonic development | 13/175 | 429/18860 | 3.27 | 0.000191 | 0.0068 | 13 | SOCS3/KLF4/CEBPB/JUNB/TM4SF1/FKBP10/LIF/ZFP36L1/KLF2/PRDM1/ELF3/CMTM3/SCO2 |
| Malignant epithelial | C0 Is ECs | GO:0042093 | T-helper cell differentiation | 6/175 | 90/18860 | 7.18 | 0.000191 | 0.0068 | 6 | LGALS1/ANXA1/SOCS3/JUNB/RIPK2/NFKBIZ |
| Malignant epithelial | C0 Is ECs | GO:0006690 | icosanoid metabolic process | 7/175 | 128/18860 | 5.89 | 0.000193 | 0.0068 | 7 | MIF/GPX1/PTGES/GGT5/TNFRSF1A/CD74/COMT |
| Malignant epithelial | C0 Is ECs | GO:0022612 | gland morphogenesis | 7/175 | 129/18860 | 5.85 | 0.000203 | 0.00699 | 7 | CAV1/TGM2/SULF2/CEBPB/CRIP1/ELF3/TNFAIP3 |
| Malignant epithelial | C0 Is ECs | GO:0001892 | embryonic placenta development | 6/175 | 91/18860 | 7.11 | 0.000203 | 0.00699 | 6 | SOCS3/CEBPB/JUNB/LIF/ZFP36L1/PRDM1 |
| Malignant epithelial | C0 Is ECs | GO:0002294 | CD4-positive, alpha-beta T cell differentiation involved in immune response | 6/175 | 92/18860 | 7.03 | 0.000216 | 0.00734 | 6 | LGALS1/ANXA1/SOCS3/JUNB/RIPK2/NFKBIZ |
| Malignant epithelial | C0 Is ECs | GO:0022408 | negative regulation of cell-cell adhesion | 9/175 | 220/18860 | 4.41 | 0.00022 | 0.00738 | 9 | LGALS1/MYADM/ANXA1/KLF4/CEBPB/IRF1/IDO1/SOCS6/CD74 |
| Malignant epithelial | C0 Is ECs | GO:0002287 | alpha-beta T cell activation involved in immune response | 6/175 | 93/18860 | 6.95 | 0.000229 | 0.00754 | 6 | LGALS1/ANXA1/SOCS3/JUNB/RIPK2/NFKBIZ |
| Malignant epithelial | C0 Is ECs | GO:0002293 | alpha-beta T cell differentiation involved in immune response | 6/175 | 93/18860 | 6.95 | 0.000229 | 0.00754 | 6 | LGALS1/ANXA1/SOCS3/JUNB/RIPK2/NFKBIZ |
| Malignant epithelial | C0 Is ECs | GO:0002696 | positive regulation of leukocyte activation | 12/175 | 381/18860 | 3.39 | 0.000238 | 0.00774 | 12 | CAV1/LGALS1/CTSC/CLCF1/ANXA1/MIF/MAP3K8/JUND/RIPK2/ABL2/NFKBIZ/CD74 |
| Malignant epithelial | C0 Is ECs | GO:0051384 | response to glucocorticoid | 7/175 | 133/18860 | 5.67 | 0.000245 | 0.00789 | 7 | DDIT4/ANXA1/DUSP1/ZFP36/ZFP36L1/FOSB/COMT |
| Malignant epithelial | C0 Is ECs | GO:0030098 | lymphocyte differentiation | 13/175 | 441/18860 | 3.18 | 0.000249 | 0.00796 | 13 | LGALS1/CLCF1/ANXA1/SOCS3/KLF6/JUNB/ZFP36L1/IRF1/CMTM7/PRDM1/RIPK2/NFKBIZ/CD74 |
| Malignant epithelial | C0 Is ECs | GO:0071347 | cellular response to interleukin-1 | 6/175 | 95/18860 | 6.81 | 0.000257 | 0.00812 | 6 | NFKBIA/CXCL8/CEBPB/LCN2/KLF2/HAS2 |
| Malignant epithelial | C0 Is ECs | GO:0044857 | plasma membrane raft organization | 3/175 | 14/18860 | 23.09 | 0.000265 | 0.00814 | 3 | CAV1/CAV2/ABCA7 |
| Malignant epithelial | C0 Is ECs | GO:0051547 | regulation of keratinocyte migration | 3/175 | 14/18860 | 23.09 | 0.000265 | 0.00814 | 3 | ARF6/MAP4K4/HAS2 |
| Malignant epithelial | C0 Is ECs | GO:0060326 | cell chemotaxis | 11/175 | 330/18860 | 3.59 | 0.000267 | 0.00814 | 11 | CXCL2/SAA1/ANXA1/CXCL8/MIF/CXCL1/DUSP1/CXCL3/C3/CD74/CKLF |

| compartment | subcluster | GO_ID | Description | GeneRatio | BgRatio | FoldEnrichment | pvalue | p_adjust | Count | geneID |
|---|---|---|---|---|---|---|---|---|---|---|
| Malignant epithelial | C0 Is ECs | GO:0001704 | formation of primary germ layer | 7/175 | 135/18860 | 5.59 | 0.000268 | 0.00814 | 7 | DUSP1/KLF4/HMGA2/LAMA3/DUSP5/MESP1/LAMB3 |
| Malignant epithelial | C0 Is ECs | GO:0070371 | ERK1 and ERK2 cascade | 11/175 | 331/18860 | 3.58 | 0.000274 | 0.00824 | 11 | MIF/DUSP1/KLF4/LIF/ZFP36L1/ABCA7/FAM3C/RIPK2/CD74/DUSP5/RRAS |
| Malignant epithelial | C0 Is ECs | GO:0045619 | regulation of lymphocyte differentiation | 9/175 | 227/18860 | 4.27 | 0.000277 | 0.00824 | 9 | LGALS1/ANXA1/JUNB/ZFP36L1/IRF1/PRDM1/RIPK2/NFKBIZ/CD74 |
| Malignant epithelial | C0 Is ECs | GO:0010810 | regulation of cell-substrate adhesion | 9/175 | 230/18860 | 4.22 | 0.000305 | 0.00899 | 9 | MYADM/PLAU/HACD1/TACSTD2/MAP4K4/RHOD/HAS2/RRAS/RASA1 |
| Malignant epithelial | C0 Is ECs | GO:0044403 | biological process involved in symbiotic interaction | 9/175 | 231/18860 | 4.2 | 0.000315 | 0.00903 | 9 | CAV1/LGALS1/CXCL8/GPX1/CAV2/BST2/ANPEP/APOL1/CD74 |
| Malignant epithelial | C0 Is ECs | GO:0051250 | negative regulation of lymphocyte activation | 8/175 | 183/18860 | 4.71 | 0.000316 | 0.00903 | 8 | LGALS1/ANXA1/CEBPB/IRF1/IDO1/SOCS6/CD74/TNFAIP3 |
| Malignant epithelial | C0 Is ECs | GO:0032102 | negative regulation of response to external stimulus | 13/175 | 452/18860 | 3.1 | 0.000316 | 0.00903 | 13 | PLAUR/SAA1/MIF/PLAU/GPX1/DUSP1/SOCS3/KLF4/IER3/TFPI/TNFRSF1A/TNFAIP3/ISG15 |
| Malignant epithelial | C0 Is ECs | GO:0043370 | regulation of CD4-positive, alpha-beta T cell differentiation | 5/175 | 64/18860 | 8.42 | 0.000318 | 0.00903 | 5 | LGALS1/ANXA1/JUNB/RIPK2/NFKBIZ |
| Malignant epithelial | C0 Is ECs | GO:0042060 | wound healing | 13/175 | 453/18860 | 3.09 | 0.000323 | 0.00908 | 13 | CAV1/PLAUR/SAA1/ANXA1/PLAU/GPX1/EREG/FKBP10/ANGPTL4/TFPI/RAP2B/TNFAIP3/JMJD1C |
| Malignant epithelial | C0 Is ECs | GO:0002292 | T cell differentiation involved in immune response | 6/175 | 101/18860 | 6.4 | 0.000358 | 0.00999 | 6 | LGALS1/ANXA1/SOCS3/JUNB/RIPK2/NFKBIZ |
| Malignant epithelial | C0 Is ECs | GO:0002286 | T cell activation involved in immune response | 7/175 | 142/18860 | 5.31 | 0.000365 | 0.0101 | 7 | LGALS1/ANXA1/SOCS3/JUNB/RIPK2/NFKBIZ/CD74 |
| Malignant epithelial | C0 Is ECs | GO:1902105 | regulation of leukocyte differentiation | 11/175 | 343/18860 | 3.46 | 0.000371 | 0.0102 | 11 | LGALS1/ANXA1/CEBPB/JUNB/LIF/ZFP36L1/IRF1/PRDM1/RIPK2/NFKBIZ/CD74 |
| Malignant epithelial | C0 Is ECs | GO:0050867 | positive regulation of cell activation | 12/175 | 401/18860 | 3.23 | 0.000378 | 0.0102 | 12 | CAV1/LGALS1/CTSC/CLCF1/ANXA1/MIF/MAP3K8/JUND/RIPK2/ABL2/NFKBIZ/CD74 |
| Malignant epithelial | C0 Is ECs | GO:0030595 | leukocyte chemotaxis | 9/175 | 237/18860 | 4.09 | 0.00038 | 0.0102 | 9 | SAA1/ANXA1/CXCL8/MIF/DUSP1/CXCL3/C3/CD74/CKLF |
| Malignant epithelial | C0 Is ECs | GO:0002460 | adaptive immune response based on somatic recombination of immune receptors built from im… | 12/175 | 402/18860 | 3.22 | 0.000386 | 0.0102 | 12 | LGALS1/CTSC/CLCF1/ANXA1/SOCS3/C3/JUNB/RIPK2/NFKBIZ/SECTM1/CD74/TNFAIP3 |
| Malignant epithelial | C0 Is ECs | GO:0008544 | epidermis development | 12/175 | 402/18860 | 3.22 | 0.000386 | 0.0102 | 12 | ANXA1/KLF4/ZFP36/EREG/ZFP36L1/WNT10A/EMP1/LAMA3/FAM3C/NFKBIZ/COL17A1/LAMB3 |
| Malignant epithelial | C0 Is ECs | GO:0070059 | intrinsic apoptotic signaling pathway in response to endoplasmic reticulum stress | 5/175 | 67/18860 | 8.04 | 0.000394 | 0.0103 | 5 | CEBPB/PMAIP1/BBC3/PPP1R15A/ERO1A |
| Malignant epithelial | C0 Is ECs | GO:0042542 | response to hydrogen peroxide | 6/175 | 103/18860 | 6.28 | 0.000398 | 0.0103 | 6 | GPX1/DUSP1/KLF4/LCN2/KLF2/TNFAIP3 |
| Malignant epithelial | C0 Is ECs | GO:1902166 | negative regulation of intrinsic apoptotic signaling pathway in response to DNA damage by… | 3/175 | 16/18860 | 20.21 | 0.000402 | 0.0103 | 3 | MIF/MUC1/CD74 |
| Malignant epithelial | C0 Is ECs | GO:0097191 | extrinsic apoptotic signaling pathway | 9/175 | 239/18860 | 4.06 | 0.000404 | 0.0103 | 9 | CAV1/GPX1/KLF4/LCN2/PMAIP1/TNFRSF1A/NFKBIZ/LTBR/TNFAIP3 |
| Malignant epithelial | C0 Is ECs | GO:0045766 | positive regulation of angiogenesis | 8/175 | 191/18860 | 4.51 | 0.00042 | 0.0106 | 8 | CXCL8/C3/TGM2/KLF4/HMGA2/ANGPTL4/BTG1/RRAS |
| Malignant epithelial | C0 Is ECs | GO:0038066 | p38MAPK cascade | 5/175 | 68/18860 | 7.92 | 0.000422 | 0.0106 | 5 | DUSP1/ZFP36/ZFP36L1/RIPK2/GADD45B |
| Malignant epithelial | C0 Is ECs | GO:0050868 | negative regulation of T cell activation | 7/175 | 146/18860 | 5.17 | 0.000432 | 0.0108 | 7 | LGALS1/ANXA1/CEBPB/IRF1/IDO1/SOCS6/CD74 |
| Malignant epithelial | C0 Is ECs | GO:0045580 | regulation of T cell differentiation | 8/175 | 194/18860 | 4.44 | 0.000466 | 0.0115 | 8 | LGALS1/ANXA1/JUNB/IRF1/PRDM1/RIPK2/NFKBIZ/CD74 |
| Malignant epithelial | C0 Is ECs | GO:1904018 | positive regulation of vasculature development | 8/175 | 195/18860 | 4.42 | 0.000482 | 0.0116 | 8 | CXCL8/C3/TGM2/KLF4/HMGA2/ANGPTL4/BTG1/RRAS |
| Malignant epithelial | C0 Is ECs | GO:2001242 | regulation of intrinsic apoptotic signaling pathway | 8/175 | 195/18860 | 4.42 | 0.000482 | 0.0116 | 8 | CAV1/PLAUR/MIF/GPX1/PMAIP1/BBC3/MUC1/CD74 |

| compartment | subcluster | GO_ID | Description | GeneRatio | BgRatio | FoldEnrichment | pvalue | p_adjust | Count | geneID |
|---|---|---|---|---|---|---|---|---|---|---|
| Malignant epithelial | C0 Is ECs | GO:2001234 | negative regulation of apoptotic signaling pathway | 9/175 | 245/18860 | 3.96 | 0.000483 | 0.0116 | 9 | NR4A2/PLAUR/MIF/GPX1/KLF4/IER3/MUC1/CD74/TNFAIP3 |
| Malignant epithelial | C0 Is ECs | GO:0006878 | intracellular copper ion homeostasis | 3/175 | 17/18860 | 19.02 | 0.000485 | 0.0116 | 3 | CP/MT2A/SCO2 |
| Malignant epithelial | C0 Is ECs | GO:0003158 | endothelium development | 7/175 | 149/18860 | 5.06 | 0.000488 | 0.0116 | 7 | MYADM/ID1/GPX1/LCN2/BTG1/TNFRSF1A/MESP1 |
| Malignant epithelial | C0 Is ECs | GO:0072331 | signal transduction by p53 class mediator | 8/175 | 196/18860 | 4.4 | 0.000499 | 0.0118 | 8 | DDIT4/MIF/PHLDA3/PMAIP1/BBC3/PPP1R15A/MUC1/CD74 |
| Malignant epithelial | C0 Is ECs | GO:0072539 | T-helper 17 cell differentiation | 4/175 | 41/18860 | 10.51 | 0.000555 | 0.013 | 4 | LGALS1/SOCS3/JUNB/NFKBIZ |
| Malignant epithelial | C0 Is ECs | GO:0002697 | regulation of immune effector process | 12/175 | 421/18860 | 3.07 | 0.000582 | 0.0134 | 12 | LGALS1/CLCF1/ANXA1/MIF/C3/JUNB/BST2/LITAF/RIPK2/NFKBIZ/SECTM1/CD74 |
| Malignant epithelial | C0 Is ECs | GO:0006631 | fatty acid metabolic process | 12/175 | 421/18860 | 3.07 | 0.000582 | 0.0134 | 12 | CAV1/MIF/GPX1/C3/PTGES/HACD1/ANGPTL4/GGT5/TNFRSF1A/CD74/COMT/PDP1 |
| Malignant epithelial | C0 Is ECs | GO:0043367 | CD4-positive, alpha-beta T cell differentiation | 6/175 | 111/18860 | 5.83 | 0.000593 | 0.0135 | 6 | LGALS1/ANXA1/SOCS3/JUNB/RIPK2/NFKBIZ |
| Malignant epithelial | C0 Is ECs | GO:0031960 | response to corticosteroid | 7/175 | 154/18860 | 4.9 | 0.000594 | 0.0135 | 7 | DDIT4/ANXA1/DUSP1/ZFP36/ZFP36L1/FOSB/COMT |
| Malignant epithelial | C0 Is ECs | GO:0032368 | regulation of lipid transport | 7/175 | 155/18860 | 4.87 | 0.000617 | 0.0139 | 7 | CAV1/NFKBIA/MIF/PTGES/PLTP/ABCA7/NMB |
| Malignant epithelial | C0 Is ECs | GO:0002526 | acute inflammatory response | 6/175 | 112/18860 | 5.77 | 0.000622 | 0.0139 | 6 | SAA1/C3/CEBPB/PTGES/LCN2/SAA2 |
| Malignant epithelial | C0 Is ECs | GO:0045765 | regulation of angiogenesis | 11/175 | 365/18860 | 3.25 | 0.000624 | 0.0139 | 11 | CXCL8/ID1/C3/TGM2/KLF4/HMGA2/ANGPTL4/BTG1/KLF2/RRAS/TNFAIP3 |
| Malignant epithelial | C0 Is ECs | GO:1902165 | regulation of intrinsic apoptotic signaling pathway in response to DNA damage by p53 clas… | 3/175 | 19/18860 | 17.02 | 0.000682 | 0.0151 | 3 | MIF/MUC1/CD74 |
| Malignant epithelial | C0 Is ECs | GO:0042325 | regulation of phosphorylation | 12/175 | 431/18860 | 3.0 | 0.000715 | 0.0157 | 12 | PLAUR/MIF/DUSP1/C3/EREG/LIF/HMGA2/GPRC5A/RIPK2/CD74/TRIB2/RAP2B |
| Malignant epithelial | C0 Is ECs | GO:0032309 | icosanoid secretion | 4/175 | 44/18860 | 9.8 | 0.000728 | 0.0157 | 4 | ANXA1/MIF/PTGES/NMB |
| Malignant epithelial | C0 Is ECs | GO:0043277 | apoptotic cell clearance | 4/175 | 44/18860 | 9.8 | 0.000728 | 0.0157 | 4 | ANXA1/C3/TGM2/ABCA7 |
| Malignant epithelial | C0 Is ECs | GO:1901342 | regulation of vasculature development | 11/175 | 375/18860 | 3.16 | 0.000779 | 0.0167 | 11 | CXCL8/ID1/C3/TGM2/KLF4/HMGA2/ANGPTL4/BTG1/KLF2/RRAS/TNFAIP3 |
| Malignant epithelial | C0 Is ECs | GO:0090199 | regulation of release of cytochrome c from mitochondria | 4/175 | 45/18860 | 9.58 | 0.000793 | 0.0169 | 4 | PLAUR/GPX1/PMAIP1/BBC3 |
| Malignant epithelial | C0 Is ECs | GO:0051546 | keratinocyte migration | 3/175 | 20/18860 | 16.17 | 0.000797 | 0.0169 | 3 | ARF6/MAP4K4/HAS2 |
| Malignant epithelial | C0 Is ECs | GO:0002695 | negative regulation of leukocyte activation | 8/175 | 212/18860 | 4.07 | 0.000834 | 0.0176 | 8 | LGALS1/ANXA1/CEBPB/IRF1/IDO1/SOCS6/CD74/TNFAIP3 |
| Malignant epithelial | C0 Is ECs | GO:0015908 | fatty acid transport | 6/175 | 119/18860 | 5.43 | 0.000855 | 0.0179 | 6 | ANXA1/MIF/PTGES/PLIN2/NMB/FABP6 |
| Malignant epithelial | C0 Is ECs | GO:0097696 | cell surface receptor signaling pathway via STAT | 7/175 | 165/18860 | 4.57 | 0.000893 | 0.0185 | 7 | CAV1/CLCF1/SOCS3/LIF/HMGA2/SOCS6/TNFRSF1A |
| Malignant epithelial | C0 Is ECs | GO:0055070 | copper ion homeostasis | 3/175 | 21/18860 | 15.4 | 0.000924 | 0.0186 | 3 | CP/MT2A/SCO2 |
| Malignant epithelial | C0 Is ECs | GO:0070431 | nucleotide-binding oligomerization domain containing 2 signaling pathway | 3/175 | 21/18860 | 15.4 | 0.000924 | 0.0186 | 3 | NFKBIA/RIPK2/TNFAIP3 |
| Malignant epithelial | C0 Is ECs | GO:0072567 | chemokine (C-X-C motif) ligand 2 production | 3/175 | 21/18860 | 15.4 | 0.000924 | 0.0186 | 3 | MIF/KLF4/CD74 |
| Malignant epithelial | C0 Is ECs | GO:2000341 | regulation of chemokine (C-X-C motif) ligand 2 production | 3/175 | 21/18860 | 15.4 | 0.000924 | 0.0186 | 3 | MIF/KLF4/CD74 |
| Malignant epithelial | C0 Is ECs | GO:0033559 | unsaturated fatty acid metabolic process | 6/175 | 121/18860 | 5.34 | 0.000933 | 0.0186 | 6 | MIF/GPX1/PTGES/TNFRSF1A/CD74/COMT |
| Malignant epithelial | C0 Is ECs | GO:0071621 | granulocyte chemotaxis | 6/175 | 121/18860 | 5.34 | 0.000933 | 0.0186 | 6 | SAA1/ANXA1/CXCL8/CXCL3/CD74/CKLF |
| Malignant epithelial | C0 Is ECs | GO:0009615 | response to virus | 12/175 | 445/18860 | 2.91 | 0.000943 | 0.0187 | 12 | DDIT4/IFITM3/LCN2/IFITM1/HMGA2/IRF1/PMAIP1/BST2/NMB/IFITM2/TNFAIP3/ISG15 |
| Malignant epithelial | C0 Is ECs | GO:0001932 | regulation of protein phosphorylation | 11/175 | 387/18860 | 3.06 | 0.00101 | 0.0198 | 11 | PLAUR/DUSP1/C3/EREG/LIF/HMGA2/GPRC5A/RIPK2/CD74/TRIB2/RAP2B |
| Malignant epithelial | C0 Is ECs | GO:0030593 | neutrophil chemotaxis | 5/175 | 83/18860 | 6.49 | 0.00105 | 0.0203 | 5 | SAA1/CXCL8/CXCL3/CD74/CKLF |

| compartment | subcluster | GO_ID | Description | GeneRatio | BgRatio | FoldEnrichment | pvalue | p_adjust | Count | geneID |
|---|---|---|---|---|---|---|---|---|---|---|
| Malignant epithelial | C0 Is ECs | GO:0030217 | T cell differentiation | 10/175 | 330/18860 | 3.27 | 0.00105 | 0.0203 | 10 | LGALS1/ANXA1/SOCS3/JUNB/ZFP36L1/IRF1/PRDM1/RIPK2/NFKBIZ/CD74 |
| Malignant epithelial | C0 Is ECs | GO:0032495 | response to muramyl dipeptide | 3/175 | 22/18860 | 14.7 | 0.00106 | 0.0203 | 3 | NFKBIA/RIPK2/TNFAIP3 |
| Malignant epithelial | C0 Is ECs | GO:0072574 | hepatocyte proliferation | 3/175 | 22/18860 | 14.7 | 0.00106 | 0.0203 | 3 | SULF2/CEBPB/TNFAIP3 |
| Malignant epithelial | C0 Is ECs | GO:0072575 | epithelial cell proliferation involved in liver morphogenesis | 3/175 | 22/18860 | 14.7 | 0.00106 | 0.0203 | 3 | SULF2/CEBPB/TNFAIP3 |
| Malignant epithelial | C0 Is ECs | GO:0071887 | leukocyte apoptotic process | 6/175 | 125/18860 | 5.17 | 0.0011 | 0.021 | 6 | ANXA1/MIF/IDO1/BBC3/NFKBIZ/CD74 |
| Malignant epithelial | C0 Is ECs | GO:0045446 | endothelial cell differentiation | 6/175 | 126/18860 | 5.13 | 0.00115 | 0.0216 | 6 | MYADM/ID1/GPX1/BTG1/TNFRSF1A/MESP1 |
| Malignant epithelial | C0 Is ECs | GO:0051607 | defense response to virus | 10/175 | 334/18860 | 3.23 | 0.00115 | 0.0216 | 10 | DDIT4/IFITM3/IFITM1/IRF1/PMAIP1/BST2/NMB/IFITM2/TNFAIP3/ISG15 |
| Malignant epithelial | C0 Is ECs | GO:0070227 | lymphocyte apoptotic process | 5/175 | 85/18860 | 6.34 | 0.00117 | 0.0218 | 5 | MIF/IDO1/BBC3/NFKBIZ/CD74 |
| Malignant epithelial | C0 Is ECs | GO:0070665 | positive regulation of leukocyte proliferation | 7/175 | 173/18860 | 4.36 | 0.00118 | 0.0218 | 7 | CLCF1/ANXA1/MIF/BST2/RIPK2/CD74/NMB |
| Malignant epithelial | C0 Is ECs | GO:0046426 | negative regulation of receptor signaling pathway via JAK-STAT | 3/175 | 23/18860 | 14.06 | 0.00121 | 0.0222 | 3 | CAV1/SOCS3/HMGA2 |
| Malignant epithelial | C0 Is ECs | GO:1900151 | regulation of nuclear-transcribed mRNA catabolic process, deadenylation-dependent decay | 3/175 | 23/18860 | 14.06 | 0.00121 | 0.0222 | 3 | ZFP36/ZFP36L1/TOB1 |
| Malignant epithelial | C0 Is ECs | GO:2000514 | regulation of CD4-positive, alpha-beta T cell activation | 5/175 | 86/18860 | 6.27 | 0.00123 | 0.0225 | 5 | LGALS1/ANXA1/JUNB/RIPK2/NFKBIZ |
| Malignant epithelial | C0 Is ECs | GO:0061082 | myeloid leukocyte cytokine production | 4/175 | 51/18860 | 8.45 | 0.00127 | 0.0229 | 4 | MIF/LITAF/RIPK2/CD74 |
| Malignant epithelial | C0 Is ECs | GO:0043434 | response to peptide hormone | 12/175 | 461/18860 | 2.81 | 0.00127 | 0.0229 | 12 | CAV1/NR4A2/SOCS3/KLF4/CAV2/EREG/ZFP36L1/JUND/BTG1/KLF2/AP3S1/COMT |
| Malignant epithelial | C0 Is ECs | GO:0034341 | response to type II interferon | 6/175 | 129/18860 | 5.01 | 0.0013 | 0.0232 | 6 | IFITM3/IFITM1/IRF1/BST2/CD74/IFITM2 |
| Malignant epithelial | C0 Is ECs | GO:0032635 | interleukin-6 production | 7/175 | 177/18860 | 4.26 | 0.00134 | 0.0236 | 7 | CEBPB/EREG/KLF2/RIPK2/CD74/NMB/TNFAIP3 |
| Malignant epithelial | C0 Is ECs | GO:0032675 | regulation of interleukin-6 production | 7/175 | 177/18860 | 4.26 | 0.00134 | 0.0236 | 7 | CEBPB/EREG/KLF2/RIPK2/CD74/NMB/TNFAIP3 |
| Malignant epithelial | C0 Is ECs | GO:0002367 | cytokine production involved in immune response | 6/175 | 130/18860 | 4.97 | 0.00135 | 0.0236 | 6 | MIF/BST2/LITAF/RIPK2/SECTM1/CD74 |
| Malignant epithelial | C0 Is ECs | GO:0002718 | regulation of cytokine production involved in immune response | 6/175 | 130/18860 | 4.97 | 0.00135 | 0.0236 | 6 | MIF/BST2/LITAF/RIPK2/SECTM1/CD74 |
| Malignant epithelial | C0 Is ECs | GO:0072576 | liver morphogenesis | 3/175 | 24/18860 | 13.47 | 0.00138 | 0.0238 | 3 | SULF2/CEBPB/TNFAIP3 |
| Malignant epithelial | C0 Is ECs | GO:1902254 | negative regulation of intrinsic apoptotic signaling pathway by p53 class mediator | 3/175 | 24/18860 | 13.47 | 0.00138 | 0.0238 | 3 | MIF/MUC1/CD74 |
| Malignant epithelial | C0 Is ECs | GO:0022409 | positive regulation of cell-cell adhesion | 10/175 | 343/18860 | 3.14 | 0.00141 | 0.024 | 10 | CAV1/LGALS1/PLAUR/ANXA1/MAP3K8/RIPK2/ABL2/NFKBIZ/CD74/HAS2 |
| Malignant epithelial | C0 Is ECs | GO:0007613 | memory | 6/175 | 131/18860 | 4.94 | 0.00141 | 0.024 | 6 | CALB1/CEBPB/LCN2/ABCA7/ADGRF1/COMT |
| Malignant epithelial | C0 Is ECs | GO:0051048 | negative regulation of secretion | 7/175 | 179/18860 | 4.21 | 0.00143 | 0.0242 | 7 | ANXA1/PIM3/LIF/TNFRSF1A/TFF2/CD74/NMB |
| Malignant epithelial | C0 Is ECs | GO:1903034 | regulation of response to wounding | 7/175 | 179/18860 | 4.21 | 0.00143 | 0.0242 | 7 | CAV1/PLAUR/ANXA1/PLAU/KLF4/TFPI/TNFAIP3 |
| Malignant epithelial | C0 Is ECs | GO:0035987 | endodermal cell differentiation | 4/175 | 53/18860 | 8.13 | 0.00147 | 0.0246 | 4 | HMGA2/LAMA3/MESP1/LAMB3 |
| Malignant epithelial | C0 Is ECs | GO:0050873 | brown fat cell differentiation | 4/175 | 53/18860 | 8.13 | 0.00147 | 0.0246 | 4 | CEBPB/ARL4A/LAMB3/ERO1A |
| Malignant epithelial | C0 Is ECs | GO:1901654 | response to ketone | 8/175 | 232/18860 | 3.72 | 0.00149 | 0.0247 | 8 | CAV1/DDIT4/DUSP1/PIM3/CDA/JUND/FOSB/COMT |
| Malignant epithelial | C0 Is ECs | GO:0051251 | positive regulation of lymphocyte activation | 10/175 | 346/18860 | 3.11 | 0.0015 | 0.0248 | 10 | CAV1/LGALS1/CLCF1/ANXA1/MIF/MAP3K8/RIPK2/ABL2/NFKBIZ/CD74 |
| Malignant epithelial | C0 Is ECs | GO:1903039 | positive regulation of leukocyte cell-cell adhesion | 9/175 | 289/18860 | 3.36 | 0.00155 | 0.0254 | 9 | CAV1/LGALS1/ANXA1/MAP3K8/RIPK2/ABL2/NFKBIZ/CD74/HAS2 |
| Malignant epithelial | C0 Is ECs | GO:0090200 | positive regulation of release of cytochrome c from mitochondria | 3/175 | 25/18860 | 12.93 | 0.00155 | 0.0254 | 3 | PLAUR/PMAIP1/BBC3 |

| compartment | subcluster | GO_ID | Description | GeneRatio | BgRatio | FoldEnrichment | pvalue | p_adjust | Count | geneID |
|---|---|---|---|---|---|---|---|---|---|---|
| Malignant epithelial | C0 Is ECs | GO:0045581 | negative regulation of T cell differentiation | 4/175 | 54/18860 | 7.98 | 0.00158 | 0.0257 | 4 | LGALS1/ANXA1/IRF1/CD74 |
| Malignant epithelial | C0 Is ECs | GO:0002285 | lymphocyte activation involved in immune response | 8/175 | 235/18860 | 3.67 | 0.00161 | 0.0259 | 8 | LGALS1/CLCF1/ANXA1/SOCS3/JUNB/RIPK2/NFKBIZ/CD74 |
| Malignant epithelial | C0 Is ECs | GO:0050866 | negative regulation of cell activation | 8/175 | 235/18860 | 3.67 | 0.00161 | 0.0259 | 8 | LGALS1/ANXA1/CEBPB/IRF1/IDO1/SOCS6/CD74/TNFAIP3 |
| Malignant epithelial | C0 Is ECs | GO:0071453 | cellular response to oxygen levels | 7/175 | 183/18860 | 4.12 | 0.00163 | 0.0259 | 7 | CAV1/LCN2/ZFP36L1/PMAIP1/JUND/BBC3/ERO1A |
| Malignant epithelial | C0 Is ECs | GO:0007259 | cell surface receptor signaling pathway via JAK-STAT | 6/175 | 135/18860 | 4.79 | 0.00164 | 0.0259 | 6 | CAV1/CLCF1/SOCS3/HMGA2/SOCS6/TNFRSF1A |
| Malignant epithelial | C0 Is ECs | GO:0031640 | killing of cells of another organism | 6/175 | 135/18860 | 4.79 | 0.00164 | 0.0259 | 6 | CXCL2/CXCL8/CXCL1/CXCL3/C3/APOL1 |
| Malignant epithelial | C0 Is ECs | GO:0141061 | disruption of cell in another organism | 6/175 | 135/18860 | 4.79 | 0.00164 | 0.0259 | 6 | CXCL2/CXCL8/CXCL1/CXCL3/C3/APOL1 |
| Malignant epithelial | C0 Is ECs | GO:0002440 | production of molecular mediator of immune response | 8/175 | 236/18860 | 3.65 | 0.00165 | 0.026 | 8 | CLCF1/MIF/BST2/LITAF/RIPK2/NFKBIZ/SECTM1/CD74 |
| Malignant epithelial | C0 Is ECs | GO:0010038 | response to metal ion | 10/175 | 351/18860 | 3.07 | 0.00167 | 0.0261 | 10 | CAV1/DUSP1/MT1E/JUNB/LCN2/CRIP1/JUND/MT2A/FOSB/ITPKC |
| Malignant epithelial | C0 Is ECs | GO:0072538 | T-helper 17 type immune response | 4/175 | 55/18860 | 7.84 | 0.00169 | 0.0263 | 4 | LGALS1/SOCS3/JUNB/NFKBIZ |
| Malignant epithelial | C0 Is ECs | GO:0097529 | myeloid leukocyte migration | 8/175 | 237/18860 | 3.64 | 0.0017 | 0.0263 | 8 | SAA1/ANXA1/CXCL8/MIF/DUSP1/CXCL3/CD74/CKLF |
| Malignant epithelial | C0 Is ECs | GO:0050678 | regulation of epithelial cell proliferation | 11/175 | 416/18860 | 2.85 | 0.00179 | 0.0275 | 11 | CAV1/GPX1/ZFP36/CAV2/SULF2/EREG/ZFP36L1/HMGA2/TACSTD2/HAS2/TNFAIP3 |
| Malignant epithelial | C0 Is ECs | GO:0071385 | cellular response to glucocorticoid stimulus | 4/175 | 56/18860 | 7.7 | 0.00181 | 0.0277 | 4 | DDIT4/ANXA1/ZFP36/ZFP36L1 |
| Malignant epithelial | C0 Is ECs | GO:0006979 | response to oxidative stress | 11/175 | 418/18860 | 2.84 | 0.00186 | 0.0283 | 11 | NR4A2/GPX1/DUSP1/KLF4/LCN2/BTG1/KLF2/ABL2/COMT/TNFAIP3/ERO1A |
| Malignant epithelial | C0 Is ECs | GO:0061041 | regulation of wound healing | 6/175 | 139/18860 | 4.65 | 0.0019 | 0.0287 | 6 | CAV1/PLAUR/ANXA1/PLAU/TFPI/TNFAIP3 |
| Malignant epithelial | C0 Is ECs | GO:0141060 | disruption of anatomical structure in another organism | 6/175 | 139/18860 | 4.65 | 0.0019 | 0.0287 | 6 | CXCL2/CXCL8/CXCL1/CXCL3/C3/APOL1 |
| Malignant epithelial | C0 Is ECs | GO:1905897 | regulation of response to endoplasmic reticulum stress | 5/175 | 95/18860 | 5.67 | 0.00192 | 0.0287 | 5 | CAV1/PMAIP1/BBC3/ABCA7/PPP1R15A |
| Malignant epithelial | C0 Is ECs | GO:0051480 | regulation of cytosolic calcium ion concentration | 4/175 | 57/18860 | 7.56 | 0.00193 | 0.0287 | 4 | CALB1/CAV1/CALB2/CAV2 |
| Malignant epithelial | C0 Is ECs | GO:0001783 | B cell apoptotic process | 3/175 | 27/18860 | 11.97 | 0.00195 | 0.0287 | 3 | MIF/NFKBIZ/CD74 |
| Malignant epithelial | C0 Is ECs | GO:0010875 | positive regulation of cholesterol efflux | 3/175 | 27/18860 | 11.97 | 0.00195 | 0.0287 | 3 | CAV1/PLTP/ABCA7 |
| Malignant epithelial | C0 Is ECs | GO:0045624 | positive regulation of T-helper cell differentiation | 3/175 | 27/18860 | 11.97 | 0.00195 | 0.0287 | 3 | ANXA1/RIPK2/NFKBIZ |
| Malignant epithelial | C0 Is ECs | GO:0007596 | blood coagulation | 8/175 | 243/18860 | 3.55 | 0.00199 | 0.0291 | 8 | CAV1/PLAUR/SAA1/PLAU/ANGPTL4/TFPI/RAP2B/JMJD1C |
| Malignant epithelial | C0 Is ECs | GO:0044409 | symbiont entry into host | 6/175 | 141/18860 | 4.59 | 0.00204 | 0.0298 | 6 | CAV1/LGALS1/CXCL8/CAV2/ANPEP/CD74 |
| Malignant epithelial | C0 Is ECs | GO:2000106 | regulation of leukocyte apoptotic process | 5/175 | 97/18860 | 5.56 | 0.0021 | 0.0306 | 5 | ANXA1/MIF/IDO1/BBC3/CD74 |
| Malignant epithelial | C0 Is ECs | GO:0007160 | cell-matrix adhesion | 8/175 | 246/18860 | 3.5 | 0.00214 | 0.031 | 8 | PLAU/MAP4K4/RHOD/ITGA10/COL17A1/RRAS/RASA1/BCAM |
| Malignant epithelial | C0 Is ECs | GO:0001836 | release of cytochrome c from mitochondria | 4/175 | 59/18860 | 7.31 | 0.00219 | 0.0314 | 4 | PLAUR/GPX1/PMAIP1/BBC3 |
| Malignant epithelial | C0 Is ECs | GO:0051592 | response to calcium ion | 6/175 | 143/18860 | 4.52 | 0.00219 | 0.0314 | 6 | CAV1/DUSP1/JUNB/JUND/FOSB/ITPKC |
| Malignant epithelial | C0 Is ECs | GO:0050817 | coagulation | 8/175 | 248/18860 | 3.48 | 0.00225 | 0.0321 | 8 | CAV1/PLAUR/SAA1/PLAU/ANGPTL4/TFPI/RAP2B/JMJD1C |
| Malignant epithelial | C0 Is ECs | GO:0007599 | hemostasis | 8/175 | 250/18860 | 3.45 | 0.00237 | 0.0333 | 8 | CAV1/PLAUR/SAA1/PLAU/ANGPTL4/TFPI/RAP2B/JMJD1C |
| Malignant epithelial | C0 Is ECs | GO:0048872 | homeostasis of number of cells | 10/175 | 369/18860 | 2.92 | 0.0024 | 0.0333 | 10 | ANXA1/MIF/ZFP36/ZFP36L1/PMAIP1/KLF2/NFKBIZ/CD74/TNFAIP3/ISG15 |
| Malignant epithelial | C0 Is ECs | GO:0045063 | T-helper 1 cell differentiation | 3/175 | 29/18860 | 11.15 | 0.0024 | 0.0333 | 3 | ANXA1/RIPK2/NFKBIZ |
| Malignant epithelial | C0 Is ECs | GO:0050482 | arachidonate secretion | 3/175 | 29/18860 | 11.15 | 0.0024 | 0.0333 | 3 | ANXA1/MIF/NMB |

| compartment | subcluster | GO_ID | Description | GeneRatio | BgRatio | FoldEnrichment | pvalue | p_adjust | Count | geneID |
|---|---|---|---|---|---|---|---|---|---|---|
| Malignant epithelial | C0 Is ECs | GO:1903963 | arachidonate transport | 3/175 | 29/18860 | 11.15 | 0.0024 | 0.0333 | 3 | ANXA1/MIF/NMB |
| Malignant epithelial | C0 Is ECs | GO:2000319 | regulation of T-helper 17 cell differentiation | 3/175 | 29/18860 | 11.15 | 0.0024 | 0.0333 | 3 | LGALS1/JUNB/NFKBIZ |
| Malignant epithelial | C0 Is ECs | GO:0043030 | regulation of macrophage activation | 4/175 | 61/18860 | 7.07 | 0.00248 | 0.0341 | 4 | CTSC/MIF/JUND/CD74 |
| Malignant epithelial | C0 Is ECs | GO:0071715 | icosanoid transport | 4/175 | 61/18860 | 7.07 | 0.00248 | 0.0341 | 4 | ANXA1/MIF/PTGES/NMB |
| Malignant epithelial | C0 Is ECs | GO:0030183 | B cell differentiation | 6/175 | 147/18860 | 4.4 | 0.00252 | 0.0345 | 6 | LGALS1/CLCF1/KLF6/ZFP36L1/CMTM7/NFKBIZ |
| Malignant epithelial | C0 Is ECs | GO:0046651 | lymphocyte proliferation | 9/175 | 312/18860 | 3.11 | 0.0026 | 0.0351 | 9 | CLCF1/ANXA1/MIF/CEBPB/IRF1/IDO1/RIPK2/NFKBIZ/CD74 |
| Malignant epithelial | C0 Is ECs | GO:0097530 | granulocyte migration | 6/175 | 148/18860 | 4.37 | 0.00261 | 0.0351 | 6 | SAA1/ANXA1/CXCL8/CXCL3/CD74/CKLF |
| Malignant epithelial | C0 Is ECs | GO:0042327 | positive regulation of phosphorylation | 8/175 | 254/18860 | 3.39 | 0.00261 | 0.0351 | 8 | PLAUR/MIF/C3/EREG/LIF/HMGA2/RIPK2/CD74 |
| Malignant epithelial | C0 Is ECs | GO:0050670 | regulation of lymphocyte proliferation | 8/175 | 254/18860 | 3.39 | 0.00261 | 0.0351 | 8 | CLCF1/ANXA1/MIF/CEBPB/IRF1/IDO1/RIPK2/CD74 |
| Malignant epithelial | C0 Is ECs | GO:0046456 | icosanoid biosynthetic process | 4/175 | 63/18860 | 6.84 | 0.00279 | 0.0372 | 4 | MIF/PTGES/GGT5/CD74 |
| Malignant epithelial | C0 Is ECs | GO:0070228 | regulation of lymphocyte apoptotic process | 4/175 | 63/18860 | 6.84 | 0.00279 | 0.0372 | 4 | MIF/IDO1/BBC3/CD74 |
| Malignant epithelial | C0 Is ECs | GO:0002700 | regulation of production of molecular mediator of immune response | 7/175 | 202/18860 | 3.73 | 0.00283 | 0.0376 | 7 | CLCF1/MIF/BST2/LITAF/RIPK2/SECTM1/CD74 |
| Malignant epithelial | C0 Is ECs | GO:0032944 | regulation of mononuclear cell proliferation | 8/175 | 258/18860 | 3.34 | 0.00287 | 0.038 | 8 | CLCF1/ANXA1/MIF/CEBPB/IRF1/IDO1/RIPK2/CD74 |
| Malignant epithelial | C0 Is ECs | GO:0001516 | prostaglandin biosynthetic process | 3/175 | 31/18860 | 10.43 | 0.00292 | 0.0381 | 3 | MIF/PTGES/CD74 |
| Malignant epithelial | C0 Is ECs | GO:0001975 | response to amphetamine | 3/175 | 31/18860 | 10.43 | 0.00292 | 0.0381 | 3 | NR4A2/FOSB/COMT |
| Malignant epithelial | C0 Is ECs | GO:1902230 | negative regulation of intrinsic apoptotic signaling pathway in response to DNA damage | 3/175 | 31/18860 | 10.43 | 0.00292 | 0.0381 | 3 | MIF/MUC1/CD74 |
| Malignant epithelial | C0 Is ECs | GO:0031663 | lipopolysaccharide-mediated signaling pathway | 4/175 | 64/18860 | 6.74 | 0.00295 | 0.0382 | 4 | NFKBIA/MIF/RIPK2/TNFAIP3 |
| Malignant epithelial | C0 Is ECs | GO:0045620 | negative regulation of lymphocyte differentiation | 4/175 | 64/18860 | 6.74 | 0.00295 | 0.0382 | 4 | LGALS1/ANXA1/IRF1/CD74 |
| Malignant epithelial | C0 Is ECs | GO:0032943 | mononuclear cell proliferation | 9/175 | 319/18860 | 3.04 | 0.00301 | 0.0388 | 9 | CLCF1/ANXA1/MIF/CEBPB/IRF1/IDO1/RIPK2/NFKBIZ/CD74 |
| Malignant epithelial | C0 Is ECs | GO:1990266 | neutrophil migration | 5/175 | 106/18860 | 5.08 | 0.0031 | 0.0397 | 5 | SAA1/CXCL8/CXCL3/CD74/CKLF |
| Malignant epithelial | C0 Is ECs | GO:0015909 | long-chain fatty acid transport | 4/175 | 65/18860 | 6.63 | 0.00312 | 0.0399 | 4 | ANXA1/MIF/PLIN2/NMB |
| Malignant epithelial | C0 Is ECs | GO:0032623 | interleukin-2 production | 4/175 | 66/18860 | 6.53 | 0.0033 | 0.0417 | 4 | ANXA1/ZFP36/RIPK2/TNFAIP3 |
| Malignant epithelial | C0 Is ECs | GO:0032663 | regulation of interleukin-2 production | 4/175 | 66/18860 | 6.53 | 0.0033 | 0.0417 | 4 | ANXA1/ZFP36/RIPK2/TNFAIP3 |
| Malignant epithelial | C0 Is ECs | GO:0071384 | cellular response to corticosteroid stimulus | 4/175 | 66/18860 | 6.53 | 0.0033 | 0.0417 | 4 | DDIT4/ANXA1/ZFP36/ZFP36L1 |
| Malignant epithelial | C0 Is ECs | GO:0150115 | cell-substrate junction organization | 5/175 | 109/18860 | 4.94 | 0.00349 | 0.0431 | 5 | ARF6/MAP4K4/LAMA3/RHOD/COL17A1 |
| Malignant epithelial | C0 Is ECs | GO:0046457 | prostanoid biosynthetic process | 3/175 | 33/18860 | 9.8 | 0.00349 | 0.0431 | 3 | MIF/PTGES/CD74 |
| Malignant epithelial | C0 Is ECs | GO:0048710 | regulation of astrocyte differentiation | 3/175 | 33/18860 | 9.8 | 0.00349 | 0.0431 | 3 | CLCF1/LIF/HMGA2 |
| Malignant epithelial | C0 Is ECs | GO:2000108 | positive regulation of leukocyte apoptotic process | 3/175 | 33/18860 | 9.8 | 0.00349 | 0.0431 | 3 | ANXA1/IDO1/BBC3 |
| Malignant epithelial | C0 Is ECs | GO:0050878 | regulation of body fluid levels | 10/175 | 389/18860 | 2.77 | 0.0035 | 0.0431 | 10 | CAV1/PLAUR/SAA1/PLAU/ANGPTL4/TFPI/RAP2B/HAS2/XDH/JMJD1C |
| Malignant epithelial | C0 Is ECs | GO:0050728 | negative regulation of inflammatory response | 7/175 | 210/18860 | 3.59 | 0.00351 | 0.0431 | 7 | SAA1/GPX1/SOCS3/KLF4/IER3/TNFRSF1A/TNFAIP3 |
| Malignant epithelial | C0 Is ECs | GO:0060759 | regulation of response to cytokine stimulus | 7/175 | 210/18860 | 3.59 | 0.00351 | 0.0431 | 7 | CAV1/KLF4/RIPK2/TNFRSF1A/CD74/TNFAIP3/ISG15 |
| Malignant epithelial | C0 Is ECs | GO:0050870 | positive regulation of T cell activation | 8/175 | 267/18860 | 3.23 | 0.00354 | 0.0433 | 8 | CAV1/LGALS1/ANXA1/MAP3K8/RIPK2/ABL2/NFKBIZ/CD74 |
| Malignant epithelial | C0 Is ECs | GO:0044854 | plasma membrane raft assembly | 2/175 | 10/18860 | 21.55 | 0.00367 | 0.0436 | 2 | CAV1/CAV2 |
| Malignant epithelial | C0 Is ECs | GO:0045647 | negative regulation of erythrocyte differentiation | 2/175 | 10/18860 | 21.55 | 0.00367 | 0.0436 | 2 | ZFP36/ZFP36L1 |
| Malignant epithelial | C0 Is ECs | GO:0045657 | positive regulation of monocyte differentiation | 2/175 | 10/18860 | 21.55 | 0.00367 | 0.0436 | 2 | ZFP36L1/CD74 |

| compartment | subcluster | GO_ID | Description | GeneRatio | BgRatio | FoldEnrichment | pvalue | p_adjust | Count | geneID |
|---|---|---|---|---|---|---|---|---|---|---|
| Malignant epithelial | C0 Is ECs | GO:0051593 | response to folic acid | 2/175 | 10/18860 | 21.55 | 0.00367 | 0.0436 | 2 | GPX1/NFKBIZ |
| Malignant epithelial | C0 Is ECs | GO:0060056 | mammary gland involution | 2/175 | 10/18860 | 21.55 | 0.00367 | 0.0436 | 2 | CAV1/ELF3 |
| Malignant epithelial | C0 Is ECs | GO:1903044 | protein localization to membrane raft | 2/175 | 10/18860 | 21.55 | 0.00367 | 0.0436 | 2 | CAV1/MYADM |
| Malignant epithelial | C0 Is ECs | GO:1990859 | cellular response to endothelin | 2/175 | 10/18860 | 21.55 | 0.00367 | 0.0436 | 2 | KLF4/KLF2 |
| Malignant epithelial | C0 Is ECs | GO:0001885 | endothelial cell development | 4/175 | 68/18860 | 6.34 | 0.00368 | 0.0436 | 4 | MYADM/ID1/GPX1/TNFRSF1A |
| Malignant epithelial | C0 Is ECs | GO:0071456 | cellular response to hypoxia | 6/175 | 159/18860 | 4.07 | 0.00371 | 0.0439 | 6 | LCN2/ZFP36L1/PMAIP1/JUND/BBC3/ERO1A |
| Malignant epithelial | C0 Is ECs | GO:0002828 | regulation of type 2 immune response | 3/175 | 34/18860 | 9.51 | 0.0038 | 0.0444 | 3 | ANXA1/IDO1/CD74 |
| Malignant epithelial | C0 Is ECs | GO:0009112 | nucleobase metabolic process | 3/175 | 34/18860 | 9.51 | 0.0038 | 0.0444 | 3 | TYMP/CDA/XDH |
| Malignant epithelial | C0 Is ECs | GO:0071498 | cellular response to fluid shear stress | 3/175 | 34/18860 | 9.51 | 0.0038 | 0.0444 | 3 | KLF4/KLF2/HAS2 |
| Malignant epithelial | C0 Is ECs | GO:0001890 | placenta development | 6/175 | 160/18860 | 4.04 | 0.00383 | 0.0445 | 6 | SOCS3/CEBPB/JUNB/LIF/ZFP36L1/PRDM1 |
| Malignant epithelial | C0 Is ECs | GO:0030198 | extracellular matrix organization | 9/175 | 331/18860 | 2.93 | 0.00384 | 0.0445 | 9 | CAV1/MIA/CAV2/SULF2/FKBP10/TNFRSF1A/ELF3/HAS2/ERO1A |
| Malignant epithelial | C0 Is ECs | GO:0043062 | extracellular structure organization | 9/175 | 332/18860 | 2.92 | 0.00391 | 0.0452 | 9 | CAV1/MIA/CAV2/SULF2/FKBP10/TNFRSF1A/ELF3/HAS2/ERO1A |
| Malignant epithelial | C0 Is ECs | GO:0045667 | regulation of osteoblast differentiation | 6/175 | 161/18860 | 4.02 | 0.00395 | 0.0454 | 6 | CEBPD/ID1/CEBPB/IFITM1/TOB1/JUND |
| Malignant epithelial | C0 Is ECs | GO:0045229 | external encapsulating structure organization | 9/175 | 333/18860 | 2.91 | 0.00399 | 0.0457 | 9 | CAV1/MIA/CAV2/SULF2/FKBP10/TNFRSF1A/ELF3/HAS2/ERO1A |
| Malignant epithelial | C0 Is ECs | GO:0050900 | leukocyte migration | 10/175 | 397/18860 | 2.71 | 0.00404 | 0.0461 | 10 | SAA1/ANXA1/CXCL8/MIF/DUSP1/CXCL3/C3/ABL2/CD74/CKLF |
| Malignant epithelial | C0 Is ECs | GO:0030193 | regulation of blood coagulation | 4/175 | 70/18860 | 6.16 | 0.00408 | 0.0464 | 4 | CAV1/PLAUR/PLAU/TFPI |
| Malignant epithelial | C0 Is ECs | GO:0048333 | mesodermal cell differentiation | 3/175 | 35/18860 | 9.24 | 0.00413 | 0.0467 | 3 | KLF4/HMGA2/MESP1 |
| Malignant epithelial | C0 Is ECs | GO:0061081 | positive regulation of myeloid leukocyte cytokine production involved in immune response | 3/175 | 35/18860 | 9.24 | 0.00413 | 0.0467 | 3 | MIF/RIPK2/CD74 |
| Malignant epithelial | C0 Is ECs | GO:0062197 | cellular response to chemical stress | 9/175 | 335/18860 | 2.9 | 0.00415 | 0.0467 | 9 | CAV1/NR4A2/GPX1/KLF4/LCN2/ZFP36L1/KLF2/ABL2/TNFAIP3 |
| Malignant epithelial | C0 Is ECs | GO:0070301 | cellular response to hydrogen peroxide | 4/175 | 71/18860 | 6.07 | 0.00429 | 0.0468 | 4 | KLF4/LCN2/KLF2/TNFAIP3 |
| Malignant epithelial | C0 Is ECs | GO:0002699 | positive regulation of immune effector process | 8/175 | 276/18860 | 3.12 | 0.00432 | 0.0468 | 8 | CLCF1/ANXA1/MIF/C3/RIPK2/NFKBIZ/SECTM1/CD74 |
| Malignant epithelial | C0 Is ECs | GO:2001243 | negative regulation of intrinsic apoptotic signaling pathway | 5/175 | 115/18860 | 4.69 | 0.00439 | 0.0468 | 5 | PLAUR/MIF/GPX1/MUC1/CD74 |
| Malignant epithelial | C0 Is ECs | GO:0051051 | negative regulation of transport | 11/175 | 468/18860 | 2.53 | 0.00439 | 0.0468 | 11 | CAV1/NFKBIA/ANXA1/PIM3/LIF/ARF6/ABCA7/TNFRSF1A/TFF2/CD74/NMB |
| Malignant epithelial | C0 Is ECs | GO:0002317 | plasma cell differentiation | 2/175 | 11/18860 | 19.59 | 0.00446 | 0.0468 | 2 | LGALS1/NFKBIZ |
| Malignant epithelial | C0 Is ECs | GO:0032060 | bleb assembly | 2/175 | 11/18860 | 19.59 | 0.00446 | 0.0468 | 2 | EMP3/EMP1 |
| Malignant epithelial | C0 Is ECs | GO:0034115 | negative regulation of heterotypic cell-cell adhesion | 2/175 | 11/18860 | 19.59 | 0.00446 | 0.0468 | 2 | MYADM/KLF4 |
| Malignant epithelial | C0 Is ECs | GO:0045625 | regulation of T-helper 1 cell differentiation | 2/175 | 11/18860 | 19.59 | 0.00446 | 0.0468 | 2 | ANXA1/RIPK2 |
| Malignant epithelial | C0 Is ECs | GO:0060426 | lung vasculature development | 2/175 | 11/18860 | 19.59 | 0.00446 | 0.0468 | 2 | ID1/LIF |
| Malignant epithelial | C0 Is ECs | GO:0070587 | regulation of cell-cell adhesion involved in gastrulation | 2/175 | 11/18860 | 19.59 | 0.00446 | 0.0468 | 2 | MYADM/KLF4 |
| Malignant epithelial | C0 Is ECs | GO:0071314 | cellular response to cocaine | 2/175 | 11/18860 | 19.59 | 0.00446 | 0.0468 | 2 | TGM2/COMT |
| Malignant epithelial | C0 Is ECs | GO:0120182 | regulation of focal adhesion disassembly | 2/175 | 11/18860 | 19.59 | 0.00446 | 0.0468 | 2 | ARF6/MAP4K4 |
| Malignant epithelial | C0 Is ECs | GO:1902237 | positive regulation of endoplasmic reticulum stress-induced intrinsic apoptotic signaling… | 2/175 | 11/18860 | 19.59 | 0.00446 | 0.0468 | 2 | PMAIP1/BBC3 |
| Malignant epithelial | C0 Is ECs | GO:1990839 | response to endothelin | 2/175 | 11/18860 | 19.59 | 0.00446 | 0.0468 | 2 | KLF4/KLF2 |
| Malignant epithelial | C0 Is ECs | GO:2000271 | positive regulation of fibroblast apoptotic process | 2/175 | 11/18860 | 19.59 | 0.00446 | 0.0468 | 2 | BTG1/BBC3 |
| Malignant epithelial | C0 Is ECs | GO:0001782 | B cell homeostasis | 3/175 | 36/18860 | 8.98 | 0.00448 | 0.0468 | 3 | MIF/CD74/TNFAIP3 |

| compartment | subcluster | GO_ID | Description | GeneRatio | BgRatio | FoldEnrichment | pvalue | p_adjust | Count | geneID |
|---|---|---|---|---|---|---|---|---|---|---|
| Malignant epithelial | C0 Is ECs | GO:0001893 | maternal placenta development | 3/175 | 36/18860 | 8.98 | 0.00448 | 0.0468 | 3 | JUNB/LIF/PRDM1 |
| Malignant epithelial | C0 Is ECs | GO:0032373 | positive regulation of sterol transport | 3/175 | 36/18860 | 8.98 | 0.00448 | 0.0468 | 3 | CAV1/PLTP/ABCA7 |
| Malignant epithelial | C0 Is ECs | GO:0032376 | positive regulation of cholesterol transport | 3/175 | 36/18860 | 8.98 | 0.00448 | 0.0468 | 3 | CAV1/PLTP/ABCA7 |
| Malignant epithelial | C0 Is ECs | GO:1904893 | negative regulation of receptor signaling pathway via STAT | 3/175 | 36/18860 | 8.98 | 0.00448 | 0.0468 | 3 | CAV1/SOCS3/HMGA2 |
| Malignant epithelial | C0 Is ECs | GO:0010632 | regulation of epithelial cell migration | 4/175 | 72/18860 | 5.99 | 0.00451 | 0.0468 | 4 | ARF6/TACSTD2/MAP4K4/HAS2 |
| Malignant epithelial | C0 Is ECs | GO:0034142 | toll-like receptor 4 signaling pathway | 4/175 | 72/18860 | 5.99 | 0.00451 | 0.0468 | 4 | CAV1/NFKBIA/RIPK2/TNFAIP3 |
| Malignant epithelial | C0 Is ECs | GO:1900046 | regulation of hemostasis | 4/175 | 72/18860 | 5.99 | 0.00451 | 0.0468 | 4 | CAV1/PLAUR/PLAU/TFPI |
| Malignant epithelial | C0 Is ECs | GO:0008637 | apoptotic mitochondrial changes | 5/175 | 116/18860 | 4.65 | 0.00455 | 0.0471 | 5 | PLAUR/GPX1/IER3/PMAIP1/BBC3 |
| Malignant epithelial | C0 Is ECs | GO:0001818 | negative regulation of cytokine production | 10/175 | 405/18860 | 2.66 | 0.00464 | 0.0478 | 10 | NFKBIA/ANXA1/KLF4/ZFP36/IDO1/BST2/KLF2/TRIB2/NMB/TNFAIP3 |
| Malignant epithelial | C0 Is ECs | GO:0036294 | cellular response to decreased oxygen levels | 6/175 | 167/18860 | 3.87 | 0.00472 | 0.0483 | 6 | LCN2/ZFP36L1/PMAIP1/JUND/BBC3/ERO1A |
| Malignant epithelial | C0 Is ECs | GO:2001236 | regulation of extrinsic apoptotic signaling pathway | 6/175 | 167/18860 | 3.87 | 0.00472 | 0.0483 | 6 | CAV1/GPX1/KLF4/PMAIP1/LTBR/TNFAIP3 |
| Malignant epithelial | C0 Is ECs | GO:0071383 | cellular response to steroid hormone stimulus | 7/175 | 222/18860 | 3.4 | 0.00475 | 0.0486 | 7 | DDIT4/ANXA1/ZFP36/ZFP36L1/PMEPA1/HMGA2/TFPI |
| Malignant epithelial | C0 Is ECs | GO:1900101 | regulation of endoplasmic reticulum unfolded protein response | 3/175 | 37/18860 | 8.74 | 0.00484 | 0.0489 | 3 | BBC3/ABCA7/PPP1R15A |
| Malignant epithelial | C0 Is ECs | GO:1902253 | regulation of intrinsic apoptotic signaling pathway by p53 class mediator | 3/175 | 37/18860 | 8.74 | 0.00484 | 0.0489 | 3 | MIF/MUC1/CD74 |
| Malignant epithelial | C0 Is ECs | GO:2000352 | negative regulation of endothelial cell apoptotic process | 3/175 | 37/18860 | 8.74 | 0.00484 | 0.0489 | 3 | ID1/ANGPTL4/TNFAIP3 |
| Malignant epithelial | C0 Is ECs | GO:0001776 | leukocyte homeostasis | 5/175 | 118/18860 | 4.57 | 0.00489 | 0.0493 | 5 | ANXA1/MIF/PMAIP1/CD74/TNFAIP3 |
| Malignant epithelial | C1 Das ECs | GO:0022600 | digestive system process | 10/166 | 116/18860 | 9.79 | 7.78e-08 | 0.000105 | 10 | TFF3/AQP1/MUC13/CLDN2/HEPH/LIMA1/SLC5A1/MDK/PLS1/KCNQ1 |
| Malignant epithelial | C1 Das ECs | GO:0007586 | digestion | 11/166 | 148/18860 | 8.44 | 7.96e-08 | 0.000105 | 11 | TFF3/AQP1/PRSS3/MUC13/CLDN2/HEPH/LIMA1/SLC5A1/MDK/PLS1/KCNQ1 |
| Malignant epithelial | C1 Das ECs | GO:0070633 | transepithelial transport | 6/166 | 40/18860 | 17.04 | 1.27e-06 | 0.00112 | 6 | AQP1/CLDN3/SLC12A2/CFTR/CXADR/SLC5A1 |
| Malignant epithelial | C1 Das ECs | GO:0050892 | intestinal absorption | 6/166 | 45/18860 | 15.15 | 2.6e-06 | 0.00171 | 6 | CLDN2/HEPH/LIMA1/SLC5A1/PLS1/KCNQ1 |
| Malignant epithelial | C1 Das ECs | GO:0048565 | digestive tract development | 8/166 | 141/18860 | 6.45 | 3.57e-05 | 0.0164 | 8 | AGR2/CDX1/ASS1/PKDCC/MYB/IFT172/CDX2/KCNQ1 |
| Malignant epithelial | C1 Das ECs | GO:0019730 | antimicrobial humoral response | 9/166 | 184/18860 | 5.56 | 3.74e-05 | 0.0164 | 9 | LGALS4/PRSS3/ANG/LYZ/CCL15/IGHG3/IGHM/IGHA1/RNASE4 |
| Malignant epithelial | C1 Das ECs | GO:0019731 | antibacterial humoral response | 6/166 | 78/18860 | 8.74 | 6.46e-05 | 0.0202 | 6 | LGALS4/ANG/IGHG3/IGHM/IGHA1/RNASE4 |
| Malignant epithelial | C1 Das ECs | GO:0055123 | digestive system development | 8/166 | 155/18860 | 5.86 | 6.99e-05 | 0.0202 | 8 | AGR2/CDX1/ASS1/PKDCC/MYB/IFT172/CDX2/KCNQ1 |
| Malignant epithelial | C1 Das ECs | GO:0030002 | intracellular monoatomic anion homeostasis | 3/166 | 10/18860 | 34.08 | 7.68e-05 | 0.0202 | 3 | SLC12A2/TBXAS1/KCNQ1 |
| Malignant epithelial | C1 Das ECs | GO:0030644 | intracellular chloride ion homeostasis | 3/166 | 10/18860 | 34.08 | 7.68e-05 | 0.0202 | 3 | SLC12A2/TBXAS1/KCNQ1 |
| Malignant epithelial | C1 Das ECs | GO:0071320 | cellular response to cAMP | 5/166 | 58/18860 | 9.79 | 0.000156 | 0.0354 | 5 | AQP1/ASS1/CFTR/FBP1/KCNQ1 |
| Malignant epithelial | C1 Das ECs | GO:0034767 | positive regulation of monoatomic ion transmembrane transport | 7/166 | 131/18860 | 6.07 | 0.000161 | 0.0354 | 7 | FXYD3/CFTR/CTSS/SRI/WNK2/HEPH/KCNQ1 |
| Malignant epithelial | C1 Das ECs | GO:0042045 | epithelial fluid transport | 3/166 | 13/18860 | 26.22 | 0.000179 | 0.0364 | 3 | AQP1/CFTR/SLC5A1 |
| Malignant epithelial | C2 WNT ECs | GO:0003401 | axis elongation | 5/176 | 31/18860 | 17.28 | 9.33e-06 | 0.0206 | 5 | NKD1/PTK7/WNT11/SPRY2/AREG |
| Malignant epithelial | C2 WNT ECs | GO:0016055 | Wnt signaling pathway | 16/176 | 487/18860 | 3.52 | 1.38e-05 | 0.0206 | 16 | NKD1/NOTUM/BAMBI/AXIN2/PTK7/SOX4/CD24/APCDD1/WNT11/COL6A1/RNF43/WIF1/WLS/IFRD1/LGR5/SNAI1 |
| Malignant epithelial | C2 WNT ECs | GO:0060485 | mesenchyme development | 13/176 | 347/18860 | 4.01 | 2.37e-05 | 0.0229 | 13 | BAMBI/AXIN2/PTEN/PTK7/SOX4/FGF19/WNT11/SPRY2/SEMA3F/RGCC/FN1/CITED2/SNAI1 |

| compartment | subcluster | GO_ID | Description | GeneRatio | BgRatio | FoldEnrichment | pvalue | p_adjust | Count | geneID |
|---|---|---|---|---|---|---|---|---|---|---|
| Malignant epithelial | C2 WNT ECs | GO:0001942 | hair follicle development | 7/176 | 99/18860 | 7.58 | 3.96e-05 | 0.0229 | 7 | APCDD1/GAL/COL6A1/KRT17/EDAR/LGR5/SNAI1 |
| Malignant epithelial | C2 WNT ECs | GO:0022404 | molting cycle process | 7/176 | 102/18860 | 7.35 | 4.79e-05 | 0.0229 | 7 | APCDD1/GAL/COL6A1/KRT17/EDAR/LGR5/SNAI1 |
| Malignant epithelial | C2 WNT ECs | GO:0022405 | hair cycle process | 7/176 | 102/18860 | 7.35 | 4.79e-05 | 0.0229 | 7 | APCDD1/GAL/COL6A1/KRT17/EDAR/LGR5/SNAI1 |
| Malignant epithelial | C2 WNT ECs | GO:0031214 | biomineral tissue development | 9/176 | 183/18860 | 5.27 | 5.66e-05 | 0.0229 | 9 | NOTUM/WDR72/AXIN2/ANKH/WNT11/COL6A1/ECM1/WLS/ANO6 |
| Malignant epithelial | C2 WNT ECs | GO:0048762 | mesenchymal cell differentiation | 11/176 | 278/18860 | 4.24 | 6.26e-05 | 0.0229 | 11 | BAMBI/AXIN2/PTEN/FGF19/WNT11/SPRY2/SEMA3F/RGCC/FN1/CITED2/SNAI1 |
| Malignant epithelial | C2 WNT ECs | GO:0002685 | regulation of leukocyte migration | 10/176 | 233/18860 | 4.6 | 6.92e-05 | 0.0229 | 10 | PERP/S100A14/ASCL2/RARRES2/LRCH1/ECM1/CD99/IL23A/PLCB1/ANO6 |
| Malignant epithelial | C2 WNT ECs | GO:0050900 | leukocyte migration | 13/176 | 397/18860 | 3.51 | 9.4e-05 | 0.0279 | 13 | DPEP1/PERP/S100A14/ASCL2/RARRES2/LRCH1/ECM1/VAV3/CD99/IL23A/PLCB1/ANO6/ITGB7 |
| Malignant epithelial | C2 WNT ECs | GO:0030111 | regulation of Wnt signaling pathway | 12/176 | 356/18860 | 3.61 | 0.000134 | 0.0359 | 12 | NKD1/NOTUM/BAMBI/AXIN2/PTK7/SOX4/APCDD1/WNT11/RNF43/WIF1/WLS/LGR5 |
| Malignant epithelial | C2 WNT ECs | GO:0042026 | protein refolding | 4/176 | 29/18860 | 14.78 | 0.000145 | 0.0359 | 4 | HSPA1A/CRYAB/FKBP1B/HSPA8 |
| Malignant epithelial | C2 WNT ECs | GO:0070857 | regulation of bile acid biosynthetic process | 3/176 | 12/18860 | 26.79 | 0.000165 | 0.0372 | 3 | CES1/PROX1/FGF19 |
| Malignant epithelial | C2 WNT ECs | GO:0042303 | molting cycle | 7/176 | 127/18860 | 5.91 | 0.000191 | 0.0372 | 7 | APCDD1/GAL/COL6A1/KRT17/EDAR/LGR5/SNAI1 |
| Malignant epithelial | C2 WNT ECs | GO:0042633 | hair cycle | 7/176 | 127/18860 | 5.91 | 0.000191 | 0.0372 | 7 | APCDD1/GAL/COL6A1/KRT17/EDAR/LGR5/SNAI1 |
| Malignant epithelial | C2 WNT ECs | GO:0030282 | bone mineralization | 7/176 | 128/18860 | 5.86 | 0.0002 | 0.0372 | 7 | NOTUM/AXIN2/ANKH/WNT11/COL6A1/ECM1/ANO6 |
| Malignant epithelial | C2 WNT ECs | GO:0042476 | odontogenesis | 7/176 | 133/18860 | 5.64 | 0.000254 | 0.0443 | 7 | ID3/AXIN2/ANKH/PERP/APCDD1/WLS/EDAR |
| Malignant epithelial | C2 WNT ECs | GO:0060070 | canonical Wnt signaling pathway | 11/176 | 333/18860 | 3.54 | 0.000303 | 0.045 | 11 | NKD1/NOTUM/BAMBI/AXIN2/PTK7/SOX4/WNT11/COL6A1/WLS/LGR5/SNAI1 |
| Malignant epithelial | C2 WNT ECs | GO:0098773 | skin epidermis development | 7/176 | 137/18860 | 5.48 | 0.000304 | 0.045 | 7 | APCDD1/GAL/COL6A1/KRT17/EDAR/LGR5/SNAI1 |
| Malignant epithelial | C2 WNT ECs | GO:1904251 | regulation of bile acid metabolic process | 3/176 | 15/18860 | 21.43 | 0.000335 | 0.045 | 3 | CES1/PROX1/FGF19 |
| Malignant epithelial | C2 WNT ECs | GO:0043627 | response to estrogen | 5/176 | 65/18860 | 8.24 | 0.000351 | 0.045 | 5 | MME/CD24/GAL/GSTM3/CITED2 |
| Malignant epithelial | C2 WNT ECs | GO:0007009 | plasma membrane organization | 8/176 | 185/18860 | 4.63 | 0.000353 | 0.045 | 8 | PTEN/COL6A1/MTSS1/A4GALT/CHMP4B/CLU/SERINC2/ANO6 |
| Malignant epithelial | C2 WNT ECs | GO:0001503 | ossification | 13/176 | 455/18860 | 3.06 | 0.000356 | 0.045 | 13 | NOTUM/BAMBI/ID3/ID4/AXIN2/PTEN/ANKH/WNT11/COL6A1/ECM1/AREG/ANO6/SNAI1 |
| Malignant epithelial | C2 WNT ECs | GO:0071674 | mononuclear cell migration | 9/176 | 235/18860 | 4.1 | 0.000372 | 0.045 | 9 | S100A14/ASCL2/RARRES2/LRCH1/ECM1/CD99/PLCB1/ANO6/ITGB7 |
| Malignant epithelial | C2 WNT ECs | GO:1900746 | regulation of vascular endothelial growth factor signaling pathway | 4/176 | 37/18860 | 11.58 | 0.000381 | 0.045 | 4 | PROX1/SPRY2/TSPAN32/SMOC2 |
| Malignant epithelial | C2 WNT ECs | GO:0097529 | myeloid leukocyte migration | 9/176 | 237/18860 | 4.07 | 0.000396 | 0.045 | 9 | DPEP1/PERP/S100A14/RARRES2/VAV3/CD99/IL23A/PLCB1/ANO6 |
| Malignant epithelial | C2 WNT ECs | GO:0060026 | convergent extension | 3/176 | 16/18860 | 20.09 | 0.000409 | 0.045 | 3 | NKD1/PTK7/WNT11 |
| Malignant epithelial | C2 WNT ECs | GO:0050921 | positive regulation of chemotaxis | 7/176 | 145/18860 | 5.17 | 0.000429 | 0.0455 | 7 | PERP/S100A14/RARRES2/IL23A/STX3/ANO6/SMOC2 |
| Malignant epithelial | C2 WNT ECs | GO:1902547 | regulation of cellular response to vascular endothelial growth factor stimulus | 4/176 | 39/18860 | 10.99 | 0.000467 | 0.0479 | 4 | PROX1/SPRY2/TSPAN32/SMOC2 |
| Malignant epithelial | C2 WNT ECs | GO:0097530 | granulocyte migration | 7/176 | 148/18860 | 5.07 | 0.000485 | 0.048 | 7 | DPEP1/PERP/S100A14/RARRES2/VAV3/CD99/IL23A |
| Malignant epithelial | C2 WNT ECs | GO:0071675 | regulation of mononuclear cell migration | 7/176 | 150/18860 | 5.0 | 0.000525 | 0.0492 | 7 | S100A14/ASCL2/RARRES2/LRCH1/ECM1/PLCB1/ANO6 |
| Malignant epithelial | C2 WNT ECs | GO:0045668 | negative regulation of osteoblast differentiation | 5/176 | 71/18860 | 7.55 | 0.000529 | 0.0492 | 5 | BAMBI/ID3/AXIN2/PTEN/AREG |
| Malignant epithelial | C2 WNT ECs | GO:0045600 | positive regulation of fat cell differentiation | 5/176 | 72/18860 | 7.44 | 0.000564 | 0.0497 | 5 | AXIN2/RARRES2/WIF1/ZBTB7C/KLF5 |

| compartment | subcluster | GO_ID | Description | GeneRatio | BgRatio | FoldEnrichment | pvalue | p_adjust | Count | geneID |
|---|---|---|---|---|---|---|---|---|---|---|
| Malignant epithelial | C2 WNT ECs | GO:0002687 | positive regulation of leukocyte migration | 7/176 | 152/18860 | 4.93 | 0.000568 | 0.0497 | 7 | PERP/S100A14/ASCL2/RARRES2/CD99/IL23A/ANO6 |
| Malignant epithelial | C3 Mp ECs | GO:0007059 | chromosome segregation | 75/178 | 434/18860 | 18.31 | 6.45e-76 | 1.29e-72 | 75 | CCNB1/CENPF/TPX2/BIRC5/NUSAP1/PTTG1/UBE2C/MKI67/AURKB/TOP2A/MAD2L1/PRC1/NUF2/CCNB2/ZWINT/… |
| Malignant epithelial | C3 Mp ECs | GO:0140014 | mitotic nuclear division | 64/178 | 281/18860 | 24.13 | 2.93e-72 | 2.93e-69 | 64 | CCNB1/CENPF/TPX2/BIRC5/NUSAP1/UBE2C/MKI67/AURKB/MAD2L1/PRC1/NUF2/UBE2S/ZWINT/PLK1/CDC20/C… |
| Malignant epithelial | C3 Mp ECs | GO:0000070 | mitotic sister chromatid segregation | 57/178 | 192/18860 | 31.46 | 2.28e-71 | 1.48e-68 | 57 | CCNB1/CENPF/TPX2/BIRC5/NUSAP1/UBE2C/AURKB/MAD2L1/PRC1/NUF2/ZWINT/PLK1/CDC20/CDK1/SPC25/DL… |
| Malignant epithelial | C3 Mp ECs | GO:0098813 | nuclear chromosome segregation | 66/178 | 323/18860 | 21.65 | 2.96e-71 | 1.48e-68 | 66 | CCNB1/CENPF/TPX2/BIRC5/NUSAP1/PTTG1/UBE2C/AURKB/TOP2A/MAD2L1/PRC1/NUF2/CCNB2/ZWINT/PLK1/C… |
| Malignant epithelial | C3 Mp ECs | GO:0000280 | nuclear division | 72/178 | 453/18860 | 16.84 | 7.66e-70 | 3.07e-67 | 72 | CCNB1/CENPF/TPX2/BIRC5/NUSAP1/PTTG1/UBE2C/MKI67/AURKB/TOP2A/MAD2L1/PRC1/NUF2/CCNB2/UBE2S/… |
| Malignant epithelial | C3 Mp ECs | GO:0000819 | sister chromatid segregation | 59/178 | 233/18860 | 26.83 | 2.47e-69 | 8.25e-67 | 59 | CCNB1/CENPF/TPX2/BIRC5/NUSAP1/UBE2C/AURKB/TOP2A/MAD2L1/PRC1/NUF2/ZWINT/PLK1/CDC20/CDK1/SP… |
| Malignant epithelial | C3 Mp ECs | GO:1902850 | microtubule cytoskeleton organization involved in mitosis | 38/178 | 170/18860 | 23.68 | 7.709999999999999e-42 | 2.2e-39 | 38 | CCNB1/TPX2/BIRC5/NUSAP1/CENPA/AURKB/MAD2L1/PRC1/NUF2/PLK1/CDC20/CDK1/SPC25/DLGAP5/AURKA/S… |
| Malignant epithelial | C3 Mp ECs | GO:1905818 | regulation of chromosome separation | 30/178 | 76/18860 | 41.82 | 1.23e-41 | 3.07e-39 | 30 | CCNB1/CENPF/BIRC5/UBE2C/AURKB/MAD2L1/NUF2/ZWINT/PLK1/CDC20/SPC25/DLGAP5/SMC4/BUB1/CDCA8/N… |
| Malignant epithelial | C3 Mp ECs | GO:0007051 | spindle organization | 40/178 | 209/18860 | 20.28 | 3.82e-41 | 8.499999999999999e-39 | 40 | CCNB1/TPX2/BIRC5/AURKB/PRC1/NUF2/CCNB2/PLK1/CDC20/ASPM/SPC25/DLGAP5/AURKA/STMN1/KIF23/KNS… |
| Malignant epithelial | C3 Mp ECs | GO:0051304 | chromosome separation | 30/178 | 83/18860 | 38.3 | 3.14e-40 | 6.27e-38 | 30 | CCNB1/CENPF/BIRC5/UBE2C/AURKB/MAD2L1/NUF2/ZWINT/PLK1/CDC20/SPC25/DLGAP5/SMC4/BUB1/CDCA8/N… |
| Malignant epithelial | C3 Mp ECs | GO:0051983 | regulation of chromosome segregation | 34/178 | 134/18860 | 26.88 | 1.7e-39 | 3.09e-37 | 34 | CCNB1/CENPF/BIRC5/UBE2C/AURKB/MAD2L1/NUF2/ZWINT/PLK1/CDC20/CDK1/SPC25/DLGAP5/CENPE/SMC4/B… |
| Malignant epithelial | C3 Mp ECs | GO:0051310 | metaphase chromosome alignment | 30/178 | 104/18860 | 30.56 | 9.01e-37 | 1.5e-34 | 30 | CCNB1/CENPF/BIRC5/AURKB/NUF2/ZWINT/CDK1/SPC25/KNSTRN/KIFC1/CENPE/NEK2/RACGAP1/KIF14/CDCA8… |
| Malignant epithelial | C3 Mp ECs | GO:0007088 | regulation of mitotic nuclear division | 31/178 | 119/18860 | 27.6 | 1.99e-36 | 3.06e-34 | 31 | CCNB1/CENPF/BIRC5/NUSAP1/UBE2C/MKI67/AURKB/MAD2L1/NUF2/ZWINT/PLK1/CDC20/SPC25/DLGAP5/AURK… |
| Malignant epithelial | C3 Mp ECs | GO:0007052 | mitotic spindle organization | 32/178 | 137/18860 | 24.75 | 6.6e-36 | 9.44e-34 | 32 | CCNB1/TPX2/BIRC5/AURKB/PRC1/NUF2/PLK1/CDC20/SPC25/DLGAP5/AURKA/STMN1/KIF23/KIFC1/CENPE/NE… |
| Malignant epithelial | C3 Mp ECs | GO:0050000 | chromosome localization | 31/178 | 124/18860 | 26.49 | 8.29e-36 | 1.11e-33 | 31 | CCNB1/CENPF/BIRC5/AURKB/NUF2/ZWINT/CDK1/SPC25/DLGAP5/KNSTRN/KIFC1/CENPE/NEK2/RACGAP1/KIF1… |

| compartment | subcluster | GO_ID | Description | GeneRatio | BgRatio | FoldEnrichment | pvalue | p_adjust | Count | geneID |
|---|---|---|---|---|---|---|---|---|---|---|
| Malignant epithelial | C3 Mp ECs | GO:0010965 | regulation of mitotic sister chromatid separation | 25/178 | 61/18860 | 43.42 | 2.71e-35 | 3.38e-33 | 25 | CCNB1/CENPF/BIRC5/UBE2C/AURKB/MAD2L1/NUF2/ZWINT/PLK1/CDC20/SPC25/DLGAP5/BUB1/CDCA8/ANAPC1… |
| Malignant epithelial | C3 Mp ECs | GO:0051783 | regulation of nuclear division | 32/178 | 147/18860 | 23.07 | 7.8e-35 | 9.17e-33 | 32 | CCNB1/CENPF/BIRC5/NUSAP1/UBE2C/MKI67/AURKB/MAD2L1/NUF2/ZWINT/PLK1/CDC20/SPC25/DLGAP5/AURK… |
| Malignant epithelial | C3 Mp ECs | GO:0051306 | mitotic sister chromatid separation | 25/178 | 64/18860 | 41.39 | 1.21e-34 | 1.34e-32 | 25 | CCNB1/CENPF/BIRC5/UBE2C/AURKB/MAD2L1/NUF2/ZWINT/PLK1/CDC20/SPC25/DLGAP5/BUB1/CDCA8/ANAPC1… |
| Malignant epithelial | C3 Mp ECs | GO:0033044 | regulation of chromosome organization | 37/178 | 244/18860 | 16.07 | 4.18e-34 | 4.4e-32 | 37 | CCNB1/CENPF/BIRC5/UBE2C/AURKB/MAD2L1/NUF2/ZWINT/PLK1/CDC20/CDK1/SPC25/DLGAP5/CENPE/NEK2/S… |
| Malignant epithelial | C3 Mp ECs | GO:0033045 | regulation of sister chromatid segregation | 28/178 | 107/18860 | 27.73 | 5.24e-33 | 5.24e-31 | 28 | CCNB1/CENPF/BIRC5/UBE2C/AURKB/MAD2L1/NUF2/ZWINT/PLK1/CDC20/CDK1/SPC25/DLGAP5/CENPE/BUB1/C… |
| Malignant epithelial | C3 Mp ECs | GO:0007091 | metaphase/anaphase transition of mitotic cell cycle | 27/178 | 96/18860 | 29.8 | 8.33e-33 | 7.93e-31 | 27 | CCNB1/CENPF/BIRC5/UBE2C/AURKB/MAD2L1/NUF2/ZWINT/PLK1/CDC20/SPC25/DLGAP5/CENPE/BUB1/CDCA8/… |
| Malignant epithelial | C3 Mp ECs | GO:0044784 | metaphase/anaphase transition of cell cycle | 27/178 | 99/18860 | 28.9 | 2.14e-32 | 1.95e-30 | 27 | CCNB1/CENPF/BIRC5/UBE2C/AURKB/MAD2L1/NUF2/ZWINT/PLK1/CDC20/SPC25/DLGAP5/CENPE/BUB1/CDCA8/… |
| Malignant epithelial | C3 Mp ECs | GO:0033047 | regulation of mitotic sister chromatid segregation | 23/178 | 57/18860 | 42.75 | 2.46e-32 | 2.14e-30 | 23 | CCNB1/CENPF/BIRC5/AURKB/MAD2L1/NUF2/ZWINT/PLK1/CDC20/CDK1/SPC25/BUB1/CDCA8/TRIP13/NDC80/S… |
| Malignant epithelial | C3 Mp ECs | GO:0033046 | negative regulation of sister chromatid segregation | 22/178 | 50/18860 | 46.62 | 5.200000000000001e-32 | 3.85e-30 | 22 | CCNB1/CENPF/BIRC5/AURKB/MAD2L1/NUF2/ZWINT/PLK1/CDC20/SPC25/BUB1/CDCA8/TRIP13/NDC80/SPC24/… |
| Malignant epithelial | C3 Mp ECs | GO:0033048 | negative regulation of mitotic sister chromatid segregation | 22/178 | 50/18860 | 46.62 | 5.200000000000001e-32 | 3.85e-30 | 22 | CCNB1/CENPF/BIRC5/AURKB/MAD2L1/NUF2/ZWINT/PLK1/CDC20/SPC25/BUB1/CDCA8/TRIP13/NDC80/SPC24/… |
| Malignant epithelial | C3 Mp ECs | GO:0045841 | negative regulation of mitotic metaphase/anaphase transition | 22/178 | 50/18860 | 46.62 | 5.200000000000001e-32 | 3.85e-30 | 22 | CCNB1/CENPF/BIRC5/AURKB/MAD2L1/NUF2/ZWINT/PLK1/CDC20/SPC25/BUB1/CDCA8/TRIP13/NDC80/SPC24/… |
| Malignant epithelial | C3 Mp ECs | GO:2000816 | negative regulation of mitotic sister chromatid separation | 22/178 | 50/18860 | 46.62 | 5.200000000000001e-32 | 3.85e-30 | 22 | CCNB1/CENPF/BIRC5/AURKB/MAD2L1/NUF2/ZWINT/PLK1/CDC20/SPC25/BUB1/CDCA8/TRIP13/NDC80/SPC24/… |
| Malignant epithelial | C3 Mp ECs | GO:0044772 | mitotic cell cycle phase transition | 44/178 | 461/18860 | 10.11 | 9.39e-32 | 6.7e-30 | 44 | CCNB1/CENPF/BIRC5/CDKN3/UBE2C/AURKB/MAD2L1/NUF2/CCNA2/CCNB2/UBE2S/ZWINT/FOXM1/PLK1/CDC20/… |
| Malignant epithelial | C3 Mp ECs | GO:0030071 | regulation of mitotic metaphase/anaphase transition | 26/178 | 92/18860 | 29.94 | 1.12e-31 | 7.739999999999999e-30 | 26 | CCNB1/CENPF/BIRC5/UBE2C/AURKB/MAD2L1/NUF2/ZWINT/PLK1/CDC20/SPC25/DLGAP5/CENPE/BUB1/CDCA8/… |
| Malignant epithelial | C3 Mp ECs | GO:1902100 | negative regulation of metaphase/anaphase transition of cell cycle | 22/178 | 52/18860 | 44.83 | 1.56e-31 | 1.01e-29 | 22 | CCNB1/CENPF/BIRC5/AURKB/MAD2L1/NUF2/ZWINT/PLK1/CDC20/SPC25/BUB1/CDCA8/TRIP13/NDC80/SPC24/… |
| Malignant epithelial | C3 Mp ECs | GO:1905819 | negative regulation of chromosome separation | 22/178 | 52/18860 | 44.83 | 1.56e-31 | 1.01e-29 | 22 | CCNB1/CENPF/BIRC5/AURKB/MAD2L1/NUF2/ZWINT/PLK1/CDC20/SPC25/BUB1/CDCA8/TRIP13/NDC80/SPC24/… |

| compartment | subcluster | GO_ID | Description | GeneRatio | BgRatio | FoldEnrichment | pvalue | p_adjust | Count | geneID |
|---|---|---|---|---|---|---|---|---|---|---|
| Malignant epithelial | C3 Mp ECs | GO:0051985 | negative regulation of chromosome segregation | 22/178 | 53/18860 | 43.98 | 2.65e-31 | 1.65e-29 | 22 | CCNB1/CENPF/BIRC5/AURKB/MAD2L1/NUF2/ZWINT/PLK1/CDC20/SPC25/BUB1/CDCA8/TRIP13/NDC80/SPC24/… |
| Malignant epithelial | C3 Mp ECs | GO:1902099 | regulation of metaphase/anaphase transition of cell cycle | 26/178 | 95/18860 | 29.0 | 2.89e-31 | 1.75e-29 | 26 | CCNB1/CENPF/BIRC5/UBE2C/AURKB/MAD2L1/NUF2/ZWINT/PLK1/CDC20/SPC25/DLGAP5/CENPE/BUB1/CDCA8/… |
| Malignant epithelial | C3 Mp ECs | GO:0090068 | positive regulation of cell cycle process | 36/178 | 278/18860 | 13.72 | 1.17e-30 | 6.86e-29 | 36 | CCNB1/BIRC5/NUSAP1/UBE2C/AURKB/MAD2L1/CDC20/CDK1/DLGAP5/AURKA/KIF23/RRM2/SMC4/RAD51AP1/BU… |
| Malignant epithelial | C3 Mp ECs | GO:0007094 | mitotic spindle assembly checkpoint signaling | 21/178 | 48/18860 | 46.36 | 1.58e-30 | 8.55e-29 | 21 | CCNB1/CENPF/BIRC5/AURKB/MAD2L1/NUF2/ZWINT/PLK1/CDC20/SPC25/BUB1/CDCA8/TRIP13/NDC80/SPC24/… |
| Malignant epithelial | C3 Mp ECs | GO:0071173 | spindle assembly checkpoint signaling | 21/178 | 48/18860 | 46.36 | 1.58e-30 | 8.55e-29 | 21 | CCNB1/CENPF/BIRC5/AURKB/MAD2L1/NUF2/ZWINT/PLK1/CDC20/SPC25/BUB1/CDCA8/TRIP13/NDC80/SPC24/… |
| Malignant epithelial | C3 Mp ECs | GO:0071174 | mitotic spindle checkpoint signaling | 21/178 | 48/18860 | 46.36 | 1.58e-30 | 8.55e-29 | 21 | CCNB1/CENPF/BIRC5/AURKB/MAD2L1/NUF2/ZWINT/PLK1/CDC20/SPC25/BUB1/CDCA8/TRIP13/NDC80/SPC24/… |
| Malignant epithelial | C3 Mp ECs | GO:0045839 | negative regulation of mitotic nuclear division | 22/178 | 57/18860 | 40.89 | 1.93e-30 | 1.02e-28 | 22 | CCNB1/CENPF/BIRC5/AURKB/MAD2L1/NUF2/ZWINT/PLK1/CDC20/SPC25/BUB1/CDCA8/TRIP13/NDC80/SPC24/… |
| Malignant epithelial | C3 Mp ECs | GO:0031577 | spindle checkpoint signaling | 21/178 | 49/18860 | 45.41 | 2.74e-30 | 1.41e-28 | 21 | CCNB1/CENPF/BIRC5/AURKB/MAD2L1/NUF2/ZWINT/PLK1/CDC20/SPC25/BUB1/CDCA8/TRIP13/NDC80/SPC24/… |
| Malignant epithelial | C3 Mp ECs | GO:0051784 | negative regulation of nuclear division | 22/178 | 64/18860 | 36.42 | 4.21e-29 | 2.11e-27 | 22 | CCNB1/CENPF/BIRC5/AURKB/MAD2L1/NUF2/ZWINT/PLK1/CDC20/SPC25/BUB1/CDCA8/TRIP13/NDC80/SPC24/… |
| Malignant epithelial | C3 Mp ECs | GO:0008608 | attachment of spindle microtubules to kinetochore | 20/178 | 55/18860 | 38.53 | 4e-27 | 1.94e-25 | 20 | CCNB1/BIRC5/AURKB/NUF2/CDK1/SPC25/KNSTRN/CENPE/NEK2/RACGAP1/CDCA8/SKA2/SPAG5/ECT2/NDC80/K… |
| Malignant epithelial | C3 Mp ECs | GO:0045787 | positive regulation of cell cycle | 36/178 | 349/18860 | 10.93 | 4.070000000000001e-27 | 1.94e-25 | 36 | CCNB1/BIRC5/NUSAP1/UBE2C/AURKB/MAD2L1/CDC20/CDK1/DLGAP5/AURKA/KIF23/RRM2/SMC4/RAD51AP1/BU… |
| Malignant epithelial | C3 Mp ECs | GO:0051225 | spindle assembly | 26/178 | 137/18860 | 20.11 | 9.810000000000001e-27 | 4.56e-25 | 26 | TPX2/BIRC5/AURKB/PRC1/CCNB2/PLK1/CDC20/ASPM/AURKA/KIF23/KIFC1/NEK2/KIF11/RACGAP1/KIF4A/TU… |
| Malignant epithelial | C3 Mp ECs | GO:2001251 | negative regulation of chromosome organization | 23/178 | 97/18860 | 25.12 | 3.8e-26 | 1.73e-24 | 23 | CCNB1/CENPF/BIRC5/AURKB/MAD2L1/NUF2/ZWINT/PLK1/CDC20/SPC25/BUB1/CDCA8/PIF1/TRIP13/NDC80/S… |
| Malignant epithelial | C3 Mp ECs | GO:1901990 | regulation of mitotic cell cycle phase transition | 35/178 | 357/18860 | 10.39 | 1.3e-25 | 5.789999999999999e-24 | 35 | CCNB1/CENPF/BIRC5/UBE2C/AURKB/MAD2L1/NUF2/ZWINT/PLK1/CDC20/CDK1/SPC25/DLGAP5/AURKA/ANLN/C… |
| Malignant epithelial | C3 Mp ECs | GO:1901987 | regulation of cell cycle phase transition | 38/178 | 468/18860 | 8.6 | 8.31e-25 | 3.61e-23 | 38 | CCNB1/CENPF/BIRC5/UBE2C/AURKB/MAD2L1/NUF2/ZWINT/PLK1/CDC20/CDK1/SPC25/DLGAP5/AURKA/ANLN/C… |
| Malignant epithelial | C3 Mp ECs | GO:0007080 | mitotic metaphase chromosome alignment | 19/178 | 63/18860 | 31.95 | 5.34e-24 | 2.27e-22 | 19 | CCNB1/BIRC5/AURKB/NUF2/CDK1/KIFC1/CENPE/KIF14/CDCA8/SKA2/KIF22/NDC80/KIF2C/PSRC1/TTK/SKA1… |

| compartment | subcluster | GO_ID | Description | GeneRatio | BgRatio | FoldEnrichment | pvalue | p_adjust | Count | geneID |
|---|---|---|---|---|---|---|---|---|---|---|
| Malignant epithelial | C3 Mp ECs | GO:0051321 | meiotic cell cycle | 31/178 | 299/18860 | 10.99 | 1.86e-23 | 7.74e-22 | 31 | PTTG1/TOP2A/NUF2/CCNB2/ZWINT/PLK1/CDC20/ASPM/AURKA/NEK2/SMC4/RAD51AP1/MND1/CKS2/SKA2/NCAP… |
| Malignant epithelial | C3 Mp ECs | GO:0140694 | membraneless organelle assembly | 36/178 | 446/18860 | 8.55 | 2.04e-23 | 8.34e-22 | 36 | CENPF/TPX2/BIRC5/CENPA/AURKB/PRC1/CCNB2/PLK1/CDC20/ASPM/DLGAP5/AURKA/KIF23/ANLN/KIFC1/CEN… |
| Malignant epithelial | C3 Mp ECs | GO:1903046 | meiotic cell cycle process | 28/178 | 227/18860 | 13.07 | 2.46e-23 | 9.86e-22 | 28 | PTTG1/TOP2A/NUF2/CCNB2/ZWINT/PLK1/CDC20/ASPM/AURKA/SMC4/RAD51AP1/MND1/CKS2/SKA2/NCAPD2/CD… |
| Malignant epithelial | C3 Mp ECs | GO:1901991 | negative regulation of mitotic cell cycle phase transition | 26/178 | 193/18860 | 14.27 | 1.01e-22 | 3.97e-21 | 26 | CCNB1/CENPF/BIRC5/AURKB/MAD2L1/NUF2/ZWINT/PLK1/CDC20/CDK1/SPC25/BUB1/CDCA8/PKMYT1/TRIP13/… |
| Malignant epithelial | C3 Mp ECs | GO:0007093 | mitotic cell cycle checkpoint signaling | 23/178 | 136/18860 | 17.92 | 1.57e-22 | 6.040000000000001e-21 | 23 | CCNB1/CENPF/BIRC5/AURKB/MAD2L1/NUF2/ZWINT/PLK1/CDC20/CDK1/SPC25/BUB1/CDCA8/TRIP13/NDC80/S… |
| Malignant epithelial | C3 Mp ECs | GO:0000075 | cell cycle checkpoint signaling | 25/178 | 187/18860 | 14.17 | 8.58e-22 | 3.24e-20 | 25 | CCNB1/CENPF/BIRC5/AURKB/MAD2L1/NUF2/ZWINT/PLK1/CDC20/CDK1/SPC25/BUB1/CDCA8/RHNO1/TRIP13/N… |
| Malignant epithelial | C3 Mp ECs | GO:1901988 | negative regulation of cell cycle phase transition | 28/178 | 274/18860 | 10.83 | 4.52e-21 | 1.67e-19 | 28 | CCNB1/CENPF/BIRC5/AURKB/MAD2L1/NUF2/ZWINT/PLK1/CDC20/CDK1/SPC25/BUB1/CDCA8/PKMYT1/RHNO1/T… |
| Malignant epithelial | C3 Mp ECs | GO:1905820 | positive regulation of chromosome separation | 14/178 | 33/18860 | 44.95 | 1.86e-20 | 6.75e-19 | 14 | BIRC5/UBE2C/AURKB/CDC20/DLGAP5/SMC4/CDCA8/NCAPD2/ANAPC11/NCAPG/SMC2/SKA1/NCAPH/SKA3 |
| Malignant epithelial | C3 Mp ECs | GO:0010948 | negative regulation of cell cycle process | 29/178 | 318/18860 | 9.66 | 2.07e-20 | 7.38e-19 | 29 | CCNB1/CENPF/BIRC5/AURKB/MAD2L1/NUF2/ZWINT/PLK1/CDC20/CDK1/SPC25/NEK2/BUB1/CDCA8/PKMYT1/RH… |
| Malignant epithelial | C3 Mp ECs | GO:0045930 | negative regulation of mitotic cell cycle | 26/178 | 244/18860 | 11.29 | 4.42e-20 | 1.55e-18 | 26 | CCNB1/CENPF/BIRC5/AURKB/MAD2L1/NUF2/ZWINT/PLK1/CDC20/CDK1/SPC25/BUB1/CDCA8/PKMYT1/TRIP13/… |
| Malignant epithelial | C3 Mp ECs | GO:0140013 | meiotic nuclear division | 24/178 | 203/18860 | 12.53 | 1.14e-19 | 3.92e-18 | 24 | PTTG1/TOP2A/NUF2/CCNB2/ZWINT/PLK1/CDC20/ASPM/AURKA/SMC4/RAD51AP1/MND1/CKS2/SKA2/TRIP13/ND… |
| Malignant epithelial | C3 Mp ECs | GO:0090307 | mitotic spindle assembly | 17/178 | 76/18860 | 23.7 | 4.06e-19 | 1.37e-17 | 17 | TPX2/BIRC5/AURKB/PRC1/PLK1/CDC20/KIF23/KIFC1/NEK2/KIF11/RACGAP1/KIF4A/CDCA8/MYBL2/KIF15/U… |
| Malignant epithelial | C3 Mp ECs | GO:0045786 | negative regulation of cell cycle | 29/178 | 399/18860 | 7.7 | 1.04e-17 | 3.47e-16 | 29 | CCNB1/CENPF/BIRC5/AURKB/MAD2L1/NUF2/ZWINT/PLK1/CDC20/CDK1/SPC25/NEK2/BUB1/CDCA8/PKMYT1/RH… |
| Malignant epithelial | C3 Mp ECs | GO:0051383 | kinetochore organization | 11/178 | 21/18860 | 55.5 | 1.26e-17 | 4.13e-16 | 11 | CENPF/CENPA/NUF2/DLGAP5/CENPW/CENPE/SMC4/CENPK/NDC80/SMC2/CENPH |
| Malignant epithelial | C3 Mp ECs | GO:0051984 | positive regulation of chromosome segregation | 12/178 | 30/18860 | 42.38 | 2.56e-17 | 8.27e-16 | 12 | CCNB1/BIRC5/AURKB/CDK1/SMC4/CDCA8/NCAPD2/NCAPG/SMC2/SKA1/NCAPH/SKA3 |
| Malignant epithelial | C3 Mp ECs | GO:0045132 | meiotic chromosome segregation | 17/178 | 97/18860 | 18.57 | 3.38e-17 | 1.07e-15 | 17 | PTTG1/NUF2/CCNB2/ZWINT/PLK1/ASPM/AURKA/SMC4/MND1/SKA2/TRIP13/NDC80/SMC2/TTK/SKA1/NCAPH/SK… |
| Malignant epithelial | C3 Mp ECs | GO:0051315 | attachment of mitotic spindle microtubules to kinetochore | 11/178 | 26/18860 | 44.83 | 2.65e-16 | 8.27e-15 | 11 | BIRC5/AURKB/NUF2/CDK1/CENPE/CDCA8/SKA2/NDC80/KIF2C/SKA1/SKA3 |

| compartment | subcluster | GO_ID | Description | GeneRatio | BgRatio | FoldEnrichment | pvalue | p_adjust | Count | geneID |
|---|---|---|---|---|---|---|---|---|---|---|
| Malignant epithelial | C3 Mp ECs | GO:0000281 | mitotic cytokinesis | 16/178 | 98/18860 | 17.3 | 9.69e-16 | 2.98e-14 | 16 | CEP55/BIRC5/NUSAP1/CENPA/AURKB/PLK1/STMN1/KIF23/ANLN/RACGAP1/KIF4A/CDCA8/CIT/IQGAP3/KIF20… |
| Malignant epithelial | C3 Mp ECs | GO:0010639 | negative regulation of organelle organization | 26/178 | 367/18860 | 7.51 | 1.09e-15 | 3.32e-14 | 26 | CCNB1/CENPF/TPX2/BIRC5/AURKB/MAD2L1/NUF2/ZWINT/PLK1/CDC20/SPC25/STMN1/BUB1/CDCA8/PIF1/TRI… |
| Malignant epithelial | C3 Mp ECs | GO:0051988 | regulation of attachment of spindle microtubules to kinetochore | 10/178 | 23/18860 | 46.07 | 4.47e-15 | 1.34e-13 | 10 | CCNB1/BIRC5/AURKB/CDK1/KNSTRN/NEK2/RACGAP1/CDCA8/SPAG5/ECT2 |
| Malignant epithelial | C3 Mp ECs | GO:0044839 | cell cycle G2/M phase transition | 18/178 | 162/18860 | 11.77 | 1.58e-14 | 4.64e-13 | 18 | CCNB1/CENPF/BIRC5/AURKB/CCNA2/CCNB2/FOXM1/PLK1/CDC20/CDK1/AURKA/KIF14/CDC25C/MELK/PKMYT1/… |
| Malignant epithelial | C3 Mp ECs | GO:0000910 | cytokinesis | 19/178 | 195/18860 | 10.32 | 3.23e-14 | 9.37e-13 | 19 | CEP55/BIRC5/NUSAP1/CENPA/AURKB/PRC1/PLK1/AURKA/STMN1/KIF23/ANLN/RACGAP1/KIF4A/KIF14/CDCA8… |
| Malignant epithelial | C3 Mp ECs | GO:0061640 | cytoskeleton-dependent cytokinesis | 16/178 | 126/18860 | 13.45 | 5.81e-14 | 1.66e-12 | 16 | CEP55/BIRC5/NUSAP1/CENPA/AURKB/PLK1/STMN1/KIF23/ANLN/RACGAP1/KIF4A/CDCA8/CIT/IQGAP3/KIF20… |
| Malignant epithelial | C3 Mp ECs | GO:0051231 | spindle elongation | 8/178 | 14/18860 | 60.55 | 1.54e-13 | 4.33e-12 | 8 | BIRC5/AURKB/PRC1/KIF23/KIF11/RACGAP1/KIF4A/CDCA8 |
| Malignant epithelial | C3 Mp ECs | GO:0034508 | centromere complex assembly | 10/178 | 31/18860 | 34.18 | 1.62e-13 | 4.51e-12 | 10 | CENPF/CENPA/DLGAP5/HJURP/CENPW/CENPE/CENPK/CENPN/OIP5/CENPH |
| Malignant epithelial | C3 Mp ECs | GO:0007143 | female meiotic nuclear division | 11/178 | 43/18860 | 27.1 | 1.72e-13 | 4.7e-12 | 11 | TOP2A/CCNB2/PLK1/AURKA/SKA2/TRIP13/NDC80/TTK/SKA1/NCAPH/SKA3 |
| Malignant epithelial | C3 Mp ECs | GO:1901970 | positive regulation of mitotic sister chromatid separation | 9/178 | 22/18860 | 43.35 | 2.17e-13 | 5.86e-12 | 9 | BIRC5/UBE2C/AURKB/CDC20/DLGAP5/CDCA8/ANAPC11/SKA1/SKA3 |
| Malignant epithelial | C3 Mp ECs | GO:0061982 | meiosis I cell cycle process | 16/178 | 142/18860 | 11.94 | 3.88e-13 | 1.03e-11 | 16 | PTTG1/TOP2A/CCNB2/ZWINT/PLK1/CDC20/AURKA/RAD51AP1/MND1/CKS2/CDC25C/PKMYT1/TRIP13/NDC80/RA… |
| Malignant epithelial | C3 Mp ECs | GO:0000086 | G2/M transition of mitotic cell cycle | 16/178 | 145/18860 | 11.69 | 5.39e-13 | 1.42e-11 | 16 | CCNB1/CENPF/BIRC5/AURKB/CCNA2/FOXM1/PLK1/CDC20/CDK1/AURKA/KIF14/CDC25C/MELK/PKMYT1/RAD21/… |
| Malignant epithelial | C3 Mp ECs | GO:0090231 | regulation of spindle checkpoint | 9/178 | 24/18860 | 39.73 | 5.61e-13 | 1.46e-11 | 9 | CCNB1/BIRC5/AURKB/MAD2L1/ZWINT/CDCA8/NDC80/SKA1/SKA3 |
| Malignant epithelial | C3 Mp ECs | GO:1901976 | regulation of cell cycle checkpoint | 11/178 | 48/18860 | 24.28 | 6.47e-13 | 1.66e-11 | 11 | CCNB1/BIRC5/AURKB/MAD2L1/ZWINT/CDCA8/NDC80/SKA1/RAD51/SKA3/BRCA1 |
| Malignant epithelial | C3 Mp ECs | GO:0051653 | spindle localization | 12/178 | 66/18860 | 19.26 | 1.09e-12 | 2.75e-11 | 12 | NUSAP1/CENPA/MAD2L1/PLK1/ASPM/SKA2/SPAG5/SAPCD2/NDC80/SKA1/SPDL1/SKA3 |
| Malignant epithelial | C3 Mp ECs | GO:2001252 | positive regulation of chromosome organization | 14/178 | 105/18860 | 14.13 | 1.16e-12 | 2.89e-11 | 14 | AURKB/CDK1/NEK2/SMC4/BUB1/NCAPD2/HNRNPA2B1/NCAPG/SMC2/SKA1/FEN1/NCAPH/SKA3/RAD21 |
| Malignant epithelial | C3 Mp ECs | GO:0051256 | mitotic spindle midzone assembly | 7/178 | 11/18860 | 67.43 | 1.89e-12 | 4.68e-11 | 7 | BIRC5/AURKB/PRC1/KIF23/RACGAP1/KIF4A/CDCA8 |
| Malignant epithelial | C3 Mp ECs | GO:0000022 | mitotic spindle elongation | 7/178 | 12/18860 | 61.81 | 4.51e-12 | 1.1e-10 | 7 | BIRC5/AURKB/PRC1/KIF23/RACGAP1/KIF4A/CDCA8 |
| Malignant epithelial | C3 Mp ECs | GO:0051293 | establishment of spindle localization | 11/178 | 61/18860 | 19.11 | 1.08e-11 | 2.6e-10 | 11 | NUSAP1/CENPA/MAD2L1/PLK1/SKA2/SPAG5/SAPCD2/NDC80/SKA1/SPDL1/SKA3 |
| Malignant epithelial | C3 Mp ECs | GO:0030261 | chromosome condensation | 10/178 | 46/18860 | 23.03 | 1.32e-11 | 3.15e-10 | 10 | CCNB1/NUSAP1/TOP2A/PLK1/CDK1/SMC4/NCAPD2/NCAPG/SMC2/NCAPH |
| Malignant epithelial | C3 Mp ECs | GO:0051294 | establishment of spindle orientation | 10/178 | 47/18860 | 22.54 | 1.67e-11 | 3.92e-10 | 10 | CENPA/MAD2L1/PLK1/SKA2/SPAG5/SAPCD2/NDC80/SKA1/SPDL1/SKA3 |

| compartment | subcluster | GO_ID | Description | GeneRatio | BgRatio | FoldEnrichment | pvalue | p_adjust | Count | geneID |
|---|---|---|---|---|---|---|---|---|---|---|
| Malignant epithelial | C3 Mp ECs | GO:0051255 | spindle midzone assembly | 7/178 | 14/18860 | 52.98 | 1.92e-11 | 4.47e-10 | 7 | BIRC5/AURKB/PRC1/KIF23/RACGAP1/KIF4A/CDCA8 |
| Malignant epithelial | C3 Mp ECs | GO:0000212 | meiotic spindle organization | 8/178 | 23/18860 | 36.85 | 2.34e-11 | 5.25e-10 | 8 | CCNB2/ASPM/AURKA/SKA2/NDC80/SKA1/SKA3/TUBG1 |
| Malignant epithelial | C3 Mp ECs | GO:0090266 | regulation of mitotic cell cycle spindle assembly checkpoint | 8/178 | 23/18860 | 36.85 | 2.34e-11 | 5.25e-10 | 8 | CCNB1/BIRC5/AURKB/MAD2L1/CDCA8/NDC80/SKA1/SKA3 |
| Malignant epithelial | C3 Mp ECs | GO:1903504 | regulation of mitotic spindle checkpoint | 8/178 | 23/18860 | 36.85 | 2.34e-11 | 5.25e-10 | 8 | CCNB1/BIRC5/AURKB/MAD2L1/CDCA8/NDC80/SKA1/SKA3 |
| Malignant epithelial | C3 Mp ECs | GO:0006310 | DNA recombination | 21/178 | 358/18860 | 6.22 | 2.7e-11 | 6.01e-10 | 21 | TOP2A/HMGB2/PLK1/RAD51AP1/NUCKS1/MND1/KPNA2/HMGB1/HMGB3/RPA3/RHNO1/PIF1/TRIP13/UHRF1/FEN1… |
| Malignant epithelial | C3 Mp ECs | GO:0090306 | meiotic spindle assembly | 7/178 | 15/18860 | 49.45 | 3.58e-11 | 7.86e-10 | 7 | CCNB2/ASPM/AURKA/SKA2/NDC80/SKA1/SKA3 |
| Malignant epithelial | C3 Mp ECs | GO:0044771 | meiotic cell cycle phase transition | 7/178 | 17/18860 | 43.63 | 1.06e-10 | 2.29e-09 | 7 | CCNB2/ZWINT/CDC20/CDC25C/PKMYT1/NDC80/TTK |
| Malignant epithelial | C3 Mp ECs | GO:0051382 | kinetochore assembly | 7/178 | 17/18860 | 43.63 | 1.06e-10 | 2.29e-09 | 7 | CENPF/CENPA/DLGAP5/CENPW/CENPE/CENPK/CENPH |
| Malignant epithelial | C3 Mp ECs | GO:0032465 | regulation of cytokinesis | 12/178 | 96/18860 | 13.24 | 1.08e-10 | 2.3e-09 | 12 | BIRC5/AURKB/PRC1/PLK1/AURKA/KIF23/RACGAP1/KIF14/CDCA8/CIT/KIF20A/ECT2 |
| Malignant epithelial | C3 Mp ECs | GO:0007076 | mitotic chromosome condensation | 7/178 | 18/18860 | 41.2 | 1.73e-10 | 3.64e-09 | 7 | NUSAP1/PLK1/SMC4/NCAPD2/NCAPG/SMC2/NCAPH |
| Malignant epithelial | C3 Mp ECs | GO:0065004 | protein-DNA complex assembly | 17/178 | 245/18860 | 7.35 | 1.77e-10 | 3.68e-09 | 17 | CENPF/CENPA/HMGB2/DLGAP5/HJURP/CENPW/CENPE/CENPK/CENPN/ASF1B/OIP5/HMGB1/ATAD2/RAD51/DNAJC… |
| Malignant epithelial | C3 Mp ECs | GO:0007127 | meiosis I | 13/178 | 132/18860 | 10.43 | 3.68e-10 | 7.6e-09 | 13 | PTTG1/TOP2A/CCNB2/ZWINT/PLK1/AURKA/RAD51AP1/MND1/CKS2/TRIP13/NDC80/RAD51/RAD21 |
| Malignant epithelial | C3 Mp ECs | GO:0007056 | spindle assembly involved in female meiosis | 6/178 | 12/18860 | 52.98 | 5.73e-10 | 1.17e-08 | 6 | CCNB2/AURKA/SKA2/NDC80/SKA1/SKA3 |
| Malignant epithelial | C3 Mp ECs | GO:0071824 | protein-DNA complex organization | 17/178 | 265/18860 | 6.8 | 5.95e-10 | 1.2e-08 | 17 | CENPF/CENPA/HMGB2/DLGAP5/HJURP/CENPW/CENPE/CENPK/CENPN/ASF1B/OIP5/HMGB1/ATAD2/RAD51/DNAJC… |
| Malignant epithelial | C3 Mp ECs | GO:0007292 | female gamete generation | 14/178 | 176/18860 | 8.43 | 1.3e-09 | 2.6e-08 | 14 | CCNB1/TOP2A/CCNB2/PLK1/ASPM/AURKA/SKA2/TRIP13/NDC80/TTK/SKA1/NCAPH/SKA3/FOSL1 |
| Malignant epithelial | C3 Mp ECs | GO:0060623 | regulation of chromosome condensation | 6/178 | 14/18860 | 45.41 | 1.83e-09 | 3.63e-08 | 6 | CCNB1/SMC4/NCAPD2/NCAPG/SMC2/NCAPH |
| Malignant epithelial | C3 Mp ECs | GO:0031109 | microtubule polymerization or depolymerization | 13/178 | 153/18860 | 9.0 | 2.31e-09 | 4.52e-08 | 13 | TPX2/AURKB/CDK1/STMN1/SKA2/KIF18B/KIF2C/PSRC1/SKA1/KIF18A/SKA3/TUBG1/MZT1 |
| Malignant epithelial | C3 Mp ECs | GO:1901989 | positive regulation of cell cycle phase transition | 12/178 | 126/18860 | 10.09 | 2.63e-09 | 5.12e-08 | 12 | CCNB1/BIRC5/UBE2C/CDC20/CDK1/DLGAP5/RRM2/CDC25C/FAM83D/ANAPC11/SKA1/SKA3 |
| Malignant epithelial | C3 Mp ECs | GO:0006302 | double-strand break repair | 18/178 | 334/18860 | 5.71 | 2.96e-09 | 5.68e-08 | 18 | HMGB2/FOXM1/PLK1/RAD51AP1/NUCKS1/CDCA5/HMGB1/RPA3/RHNO1/TRIP13/UHRF1/FEN1/RAD51/HELLS/RAD… |
| Malignant epithelial | C3 Mp ECs | GO:0051302 | regulation of cell division | 14/178 | 190/18860 | 7.81 | 3.53e-09 | 6.72e-08 | 14 | BIRC5/AURKB/PRC1/PLK1/ASPM/AURKA/KIF23/RACGAP1/KIF14/CDCA8/CIT/KIF20A/KIF18B/ECT2 |
| Malignant epithelial | C3 Mp ECs | GO:1901992 | positive regulation of mitotic cell cycle phase transition | 11/178 | 105/18860 | 11.1 | 4.48e-09 | 8.46e-08 | 11 | CCNB1/BIRC5/UBE2C/CDC20/CDK1/DLGAP5/RRM2/CDC25C/ANAPC11/SKA1/SKA3 |
| Malignant epithelial | C3 Mp ECs | GO:0010389 | regulation of G2/M transition of mitotic cell cycle | 11/178 | 109/18860 | 10.69 | 6.69e-09 | 1.25e-07 | 11 | CCNB1/CENPF/AURKB/PLK1/CDK1/AURKA/KIF14/CDC25C/PKMYT1/RAD21/BRCA1 |
| Malignant epithelial | C3 Mp ECs | GO:0051785 | positive regulation of nuclear division | 9/178 | 63/18860 | 15.14 | 7.41e-09 | 1.37e-07 | 9 | NUSAP1/UBE2C/CDC20/DLGAP5/AURKA/RAD51AP1/ANAPC11/SKA1/SKA3 |

| compartment | subcluster | GO_ID | Description | GeneRatio | BgRatio | FoldEnrichment | pvalue | p_adjust | Count | geneID |
|---|---|---|---|---|---|---|---|---|---|---|
| Malignant epithelial | C3 Mp ECs | GO:0045931 | positive regulation of mitotic cell cycle | 12/178 | 138/18860 | 9.21 | 7.47e-09 | 1.37e-07 | 12 | CCNB1/BIRC5/UBE2C/CDC20/CDK1/DLGAP5/AURKA/RRM2/CDC25C/ANAPC11/SKA1/SKA3 |
| Malignant epithelial | C3 Mp ECs | GO:0045842 | positive regulation of mitotic metaphase/anaphase transition | 6/178 | 18/18860 | 35.32 | 1.1e-08 | 2e-07 | 6 | UBE2C/CDC20/DLGAP5/ANAPC11/SKA1/SKA3 |
| Malignant epithelial | C3 Mp ECs | GO:0045840 | positive regulation of mitotic nuclear division | 8/178 | 47/18860 | 18.03 | 1.24e-08 | 2.23e-07 | 8 | NUSAP1/UBE2C/CDC20/DLGAP5/AURKA/ANAPC11/SKA1/SKA3 |
| Malignant epithelial | C3 Mp ECs | GO:0035825 | homologous recombination | 9/178 | 68/18860 | 14.02 | 1.48e-08 | 2.65e-07 | 9 | TOP2A/RAD51AP1/NUCKS1/MND1/TRIP13/UHRF1/RAD51/RAD21/BRCA1 |
| Malignant epithelial | C3 Mp ECs | GO:0051299 | centrosome separation | 6/178 | 19/18860 | 33.46 | 1.59e-08 | 2.79e-07 | 6 | PLK1/AURKA/NEK2/KIF11/RANBP1/KIF15 |
| Malignant epithelial | C3 Mp ECs | GO:1902101 | positive regulation of metaphase/anaphase transition of cell cycle | 6/178 | 19/18860 | 33.46 | 1.59e-08 | 2.79e-07 | 6 | UBE2C/CDC20/DLGAP5/ANAPC11/SKA1/SKA3 |
| Malignant epithelial | C3 Mp ECs | GO:0051445 | regulation of meiotic cell cycle | 9/178 | 70/18860 | 13.62 | 1.92e-08 | 3.35e-07 | 9 | ZWINT/CDC20/ASPM/RAD51AP1/CDC25C/PKMYT1/ANAPC11/TRIP13/TTK |
| Malignant epithelial | C3 Mp ECs | GO:1902749 | regulation of cell cycle G2/M phase transition | 11/178 | 122/18860 | 9.55 | 2.2e-08 | 3.79e-07 | 11 | CCNB1/CENPF/AURKB/PLK1/CDK1/AURKA/KIF14/CDC25C/PKMYT1/RAD21/BRCA1 |
| Malignant epithelial | C3 Mp ECs | GO:0034502 | protein localization to chromosome | 11/178 | 123/18860 | 9.48 | 2.4e-08 | 4.1e-07 | 11 | CENPA/AURKB/PLK1/CDK1/RHNO1/VRK1/TTK/SPDL1/BUB1B/BUB3/RAD21 |
| Malignant epithelial | C3 Mp ECs | GO:0032467 | positive regulation of cytokinesis | 8/178 | 51/18860 | 16.62 | 2.42e-08 | 4.11e-07 | 8 | BIRC5/AURKB/KIF23/RACGAP1/KIF14/CDCA8/CIT/ECT2 |
| Malignant epithelial | C3 Mp ECs | GO:0031055 | chromatin remodeling at centromere | 5/178 | 11/18860 | 48.16 | 3.12e-08 | 5.16e-07 | 5 | CENPA/HJURP/CENPW/CENPN/OIP5 |
| Malignant epithelial | C3 Mp ECs | GO:0034080 | CENP-A containing chromatin assembly | 5/178 | 11/18860 | 48.16 | 3.12e-08 | 5.16e-07 | 5 | CENPA/HJURP/CENPW/CENPN/OIP5 |
| Malignant epithelial | C3 Mp ECs | GO:0090267 | positive regulation of mitotic cell cycle spindle assembly checkpoint | 5/178 | 11/18860 | 48.16 | 3.12e-08 | 5.16e-07 | 5 | BIRC5/AURKB/MAD2L1/CDCA8/NDC80 |
| Malignant epithelial | C3 Mp ECs | GO:0090232 | positive regulation of spindle checkpoint | 5/178 | 12/18860 | 44.15 | 5.31e-08 | 8.71e-07 | 5 | BIRC5/AURKB/MAD2L1/CDCA8/NDC80 |
| Malignant epithelial | C3 Mp ECs | GO:0000725 | recombinational repair | 13/178 | 201/18860 | 6.85 | 6.15e-08 | 1e-06 | 13 | PLK1/RAD51AP1/NUCKS1/RPA3/RHNO1/UHRF1/FEN1/RAD51/HELLS/RAD21/BRCA1/RECQL4/PARPBP |
| Malignant epithelial | C3 Mp ECs | GO:0071459 | protein localization to chromosome, centromeric region | 7/178 | 41/18860 | 18.09 | 1.02e-07 | 1.64e-06 | 7 | CENPA/AURKB/CDK1/TTK/SPDL1/BUB1B/BUB3 |
| Malignant epithelial | C3 Mp ECs | GO:0040001 | establishment of mitotic spindle localization | 7/178 | 42/18860 | 17.66 | 1.21e-07 | 1.94e-06 | 7 | NUSAP1/CENPA/MAD2L1/PLK1/SAPCD2/NDC80/SPDL1 |
| Malignant epithelial | C3 Mp ECs | GO:0007062 | sister chromatid cohesion | 8/178 | 63/18860 | 13.45 | 1.34e-07 | 2.12e-06 | 8 | PLK1/CDC20/BUB1/CDCA5/KIF22/FEN1/BUB1B/RAD21 |
| Malignant epithelial | C3 Mp ECs | GO:0034501 | protein localization to kinetochore | 5/178 | 15/18860 | 35.32 | 1.97e-07 | 3.03e-06 | 5 | AURKB/CDK1/TTK/SPDL1/BUB3 |
| Malignant epithelial | C3 Mp ECs | GO:0090235 | regulation of metaphase plate congression | 5/178 | 15/18860 | 35.32 | 1.97e-07 | 3.03e-06 | 5 | BIRC5/AURKB/CDK1/CDCA8/SPAG5 |
| Malignant epithelial | C3 Mp ECs | GO:1903083 | protein localization to condensed chromosome | 5/178 | 15/18860 | 35.32 | 1.97e-07 | 3.03e-06 | 5 | AURKB/CDK1/TTK/SPDL1/BUB3 |
| Malignant epithelial | C3 Mp ECs | GO:1904666 | regulation of ubiquitin protein ligase activity | 5/178 | 15/18860 | 35.32 | 1.97e-07 | 3.03e-06 | 5 | UBE2C/MAD2L1/UBE2S/PLK1/CDC20 |
| Malignant epithelial | C3 Mp ECs | GO:0000724 | double-strand break repair via homologous recombination | 12/178 | 195/18860 | 6.52 | 3.44e-07 | 5.26e-06 | 12 | PLK1/RAD51AP1/NUCKS1/RPA3/UHRF1/FEN1/RAD51/HELLS/RAD21/BRCA1/RECQL4/PARPBP |
| Malignant epithelial | C3 Mp ECs | GO:0032886 | regulation of microtubule-based process | 14/178 | 276/18860 | 5.37 | 3.8e-07 | 5.76e-06 | 14 | TPX2/AURKB/PLK1/AURKA/STMN1/SKA2/SPAG5/TACC3/POC1A/PSRC1/SKA1/KIF18A/SKA3/BRCA1 |
| Malignant epithelial | C3 Mp ECs | GO:0071103 | DNA conformation change | 5/178 | 18/18860 | 29.43 | 5.49e-07 | 8.19e-06 | 5 | TOP2A/HMGB2/HNRNPA2B1/HMGB1/HMGB3 |
| Malignant epithelial | C3 Mp ECs | GO:1901978 | positive regulation of cell cycle checkpoint | 5/178 | 18/18860 | 29.43 | 5.49e-07 | 8.19e-06 | 5 | BIRC5/AURKB/MAD2L1/CDCA8/NDC80 |
| Malignant epithelial | C3 Mp ECs | GO:0032506 | cytokinetic process | 7/178 | 52/18860 | 14.26 | 5.54e-07 | 8.21e-06 | 7 | CEP55/AURKB/ANLN/RACGAP1/IQGAP3/KIF20A/ECT2 |
| Malignant epithelial | C3 Mp ECs | GO:0070507 | regulation of microtubule cytoskeleton organization | 11/178 | 168/18860 | 6.94 | 5.81e-07 | 8.55e-06 | 11 | TPX2/AURKB/PLK1/STMN1/SKA2/SPAG5/TACC3/PSRC1/SKA1/KIF18A/SKA3 |

| compartment | subcluster | GO_ID | Description | GeneRatio | BgRatio | FoldEnrichment | pvalue | p_adjust | Count | geneID |
|---|---|---|---|---|---|---|---|---|---|---|
| Malignant epithelial | C3 Mp ECs | GO:0000132 | establishment of mitotic spindle orientation | 6/178 | 37/18860 | 17.18 | 1.18e-06 | 1.73e-05 | 6 | CENPA/MAD2L1/PLK1/SAPCD2/NDC80/SPDL1 |
| Malignant epithelial | C3 Mp ECs | GO:0051656 | establishment of organelle localization | 16/178 | 400/18860 | 4.24 | 1.32e-06 | 1.91e-05 | 16 | NUSAP1/CENPA/MAD2L1/PLK1/CENPE/LMNB1/RAN/SKA2/SPAG5/TACC3/SAPCD2/LMNB2/NDC80/SKA1/SPDL1/S… |
| Malignant epithelial | C3 Mp ECs | GO:0044837 | actomyosin contractile ring organization | 4/178 | 10/18860 | 42.38 | 1.54e-06 | 2.2e-05 | 4 | ANLN/RACGAP1/IQGAP3/ECT2 |
| Malignant epithelial | C3 Mp ECs | GO:1902423 | regulation of attachment of mitotic spindle microtubules to kinetochore | 4/178 | 10/18860 | 42.38 | 1.54e-06 | 2.2e-05 | 4 | BIRC5/AURKB/CDK1/CDCA8 |
| Malignant epithelial | C3 Mp ECs | GO:0007098 | centrosome cycle | 10/178 | 150/18860 | 7.06 | 1.63e-06 | 2.31e-05 | 10 | PLK1/CDK1/AURKA/NEK2/KIF11/RANBP1/POC1A/NDC80/KIF15/BRCA1 |
| Malignant epithelial | C3 Mp ECs | GO:0051987 | positive regulation of attachment of spindle microtubules to kinetochore | 4/178 | 11/18860 | 38.53 | 2.4e-06 | 3.39e-05 | 4 | CCNB1/BIRC5/AURKB/CDCA8 |
| Malignant epithelial | C3 Mp ECs | GO:0070192 | chromosome organization involved in meiotic cell cycle | 7/178 | 66/18860 | 11.24 | 2.89e-06 | 4.04e-05 | 7 | SMC4/MND1/NCAPD2/TRIP13/SMC2/RAD51/NCAPH |
| Malignant epithelial | C3 Mp ECs | GO:0031145 | anaphase-promoting complex-dependent catabolic process | 5/178 | 25/18860 | 21.19 | 3.23e-06 | 4.48e-05 | 5 | UBE2C/UBE2S/PLK1/CDC20/ANAPC11 |
| Malignant epithelial | C3 Mp ECs | GO:0031023 | microtubule organizing center organization | 10/178 | 163/18860 | 6.5 | 3.45e-06 | 4.76e-05 | 10 | PLK1/CDK1/AURKA/NEK2/KIF11/RANBP1/POC1A/NDC80/KIF15/BRCA1 |
| Malignant epithelial | C3 Mp ECs | GO:0030010 | establishment of cell polarity | 10/178 | 164/18860 | 6.46 | 3.65e-06 | 5e-05 | 10 | CENPA/MAD2L1/PLK1/SKA2/SPAG5/SAPCD2/NDC80/SKA1/SPDL1/SKA3 |
| Malignant epithelial | C3 Mp ECs | GO:0006260 | DNA replication | 13/178 | 290/18860 | 4.75 | 3.95e-06 | 5.36e-05 | 13 | CCNA2/TK1/CDK1/NUCKS1/PTMS/ORC6/RPA3/RNASEH2A/FEN1/RAD51/PCNA/BRCA1/RECQL4 |
| Malignant epithelial | C3 Mp ECs | GO:0051438 | regulation of ubiquitin-protein transferase activity | 5/178 | 26/18860 | 20.38 | 3.96e-06 | 5.36e-05 | 5 | UBE2C/MAD2L1/UBE2S/PLK1/CDC20 |
| Malignant epithelial | C3 Mp ECs | GO:0051781 | positive regulation of cell division | 8/178 | 99/18860 | 8.56 | 4.43e-06 | 5.95e-05 | 8 | BIRC5/AURKB/KIF23/RACGAP1/KIF14/CDCA8/CIT/ECT2 |
| Malignant epithelial | C3 Mp ECs | GO:0007144 | female meiosis I | 4/178 | 13/18860 | 32.6 | 5.13e-06 | 6.75e-05 | 4 | CCNB2/AURKA/TRIP13/NDC80 |
| Malignant epithelial | C3 Mp ECs | GO:0051443 | positive regulation of ubiquitin-protein transferase activity | 4/178 | 13/18860 | 32.6 | 5.13e-06 | 6.75e-05 | 4 | UBE2C/UBE2S/PLK1/CDC20 |
| Malignant epithelial | C3 Mp ECs | GO:1903490 | positive regulation of mitotic cytokinesis | 4/178 | 13/18860 | 32.6 | 5.13e-06 | 6.75e-05 | 4 | BIRC5/AURKB/CDCA8/ECT2 |
| Malignant epithelial | C3 Mp ECs | GO:0051338 | regulation of transferase activity | 13/178 | 304/18860 | 4.53 | 6.58e-06 | 8.61e-05 | 13 | TPX2/CDKN3/UBE2C/MAD2L1/UBE2S/PLK1/CDC20/CENPE/KIF14/CDC25C/PKMYT1/ECT2/PCNA |
| Malignant epithelial | C3 Mp ECs | GO:0006213 | pyrimidine nucleoside metabolic process | 4/178 | 14/18860 | 30.27 | 7.13e-06 | 9.14e-05 | 4 | TK1/CDK1/DTYMK/ERH |
| Malignant epithelial | C3 Mp ECs | GO:0007100 | mitotic centrosome separation | 4/178 | 14/18860 | 30.27 | 7.13e-06 | 9.14e-05 | 4 | AURKA/NEK2/KIF11/RANBP1 |
| Malignant epithelial | C3 Mp ECs | GO:1901993 | regulation of meiotic cell cycle phase transition | 4/178 | 14/18860 | 30.27 | 7.13e-06 | 9.14e-05 | 4 | ZWINT/CDC25C/PKMYT1/TTK |
| Malignant epithelial | C3 Mp ECs | GO:0007019 | microtubule depolymerization | 6/178 | 52/18860 | 12.23 | 9.23e-06 | 0.000118 | 6 | TPX2/AURKB/STMN1/KIF18B/KIF2C/KIF18A |
| Malignant epithelial | C3 Mp ECs | GO:0044027 | negative regulation of gene expression via chromosomal CpG island methylation | 4/178 | 16/18860 | 26.49 | 1.28e-05 | 0.000161 | 4 | DNMT1/UHRF1/HELLS/BRCA1 |
| Malignant epithelial | C3 Mp ECs | GO:1902412 | regulation of mitotic cytokinesis | 4/178 | 16/18860 | 26.49 | 1.28e-05 | 0.000161 | 4 | BIRC5/AURKB/CDCA8/ECT2 |
| Malignant epithelial | C3 Mp ECs | GO:0007163 | establishment or maintenance of cell polarity | 11/178 | 236/18860 | 4.94 | 1.55e-05 | 0.000194 | 11 | CENPA/MAD2L1/PLK1/SKA2/SPAG5/SAPCD2/NDC80/KIF2C/SKA1/SPDL1/SKA3 |
| Malignant epithelial | C3 Mp ECs | GO:0051052 | regulation of DNA metabolic process | 16/178 | 497/18860 | 3.41 | 2.04e-05 | 0.000254 | 16 | AURKB/CCNA2/FOXM1/PLK1/CDK1/NEK2/RAD51AP1/NUCKS1/KPNA2/HNRNPA2B1/HMGB1/PIF1/RAD51/PCNA/BR… |
| Malignant epithelial | C3 Mp ECs | GO:0006298 | mismatch repair | 5/178 | 36/18860 | 14.72 | 2.11e-05 | 0.00026 | 5 | HMGB1/RPA3/RNASEH2A/PCNA/RNASEH2C |
| Malignant epithelial | C3 Mp ECs | GO:0007131 | reciprocal meiotic recombination | 6/178 | 60/18860 | 10.6 | 2.13e-05 | 0.00026 | 6 | TOP2A/RAD51AP1/MND1/TRIP13/RAD51/RAD21 |
| Malignant epithelial | C3 Mp ECs | GO:0140527 | reciprocal homologous recombination | 6/178 | 60/18860 | 10.6 | 2.13e-05 | 0.00026 | 6 | TOP2A/RAD51AP1/MND1/TRIP13/RAD51/RAD21 |

| compartment | subcluster | GO_ID | Description | GeneRatio | BgRatio | FoldEnrichment | pvalue | p_adjust | Count | geneID |
|---|---|---|---|---|---|---|---|---|---|---|
| Malignant epithelial | C3 Mp ECs | GO:0051347 | positive regulation of transferase activity | 9/178 | 161/18860 | 5.92 | 2.27e-05 | 0.000275 | 9 | TPX2/UBE2C/UBE2S/PLK1/CDC20/CENPE/KIF14/ECT2/PCNA |
| Malignant epithelial | C3 Mp ECs | GO:2000241 | regulation of reproductive process | 10/178 | 206/18860 | 5.14 | 2.69e-05 | 0.000324 | 10 | ZWINT/CDC20/ASPM/AURKA/RAD51AP1/CDC25C/PKMYT1/ANAPC11/TRIP13/TTK |
| Malignant epithelial | C3 Mp ECs | GO:1902410 | mitotic cytokinetic process | 5/178 | 38/18860 | 13.94 | 2.76e-05 | 0.000331 | 5 | CEP55/AURKB/IQGAP3/KIF20A/ECT2 |
| Malignant epithelial | C3 Mp ECs | GO:0000723 | telomere maintenance | 9/178 | 166/18860 | 5.74 | 2.89e-05 | 0.000344 | 9 | AURKB/NEK2/HNRNPA2B1/RPA3/PIF1/FEN1/RAD51/PCNA/RECQL4 |
| Malignant epithelial | C3 Mp ECs | GO:0033260 | nuclear DNA replication | 5/178 | 41/18860 | 12.92 | 4.03e-05 | 0.000476 | 5 | TK1/CDK1/FEN1/RAD51/PCNA |
| Malignant epithelial | C3 Mp ECs | GO:0051447 | negative regulation of meiotic cell cycle | 4/178 | 21/18860 | 20.18 | 4.05e-05 | 0.000476 | 4 | ZWINT/PKMYT1/TRIP13/TTK |
| Malignant epithelial | C3 Mp ECs | GO:0031110 | regulation of microtubule polymerization or depolymerization | 7/178 | 104/18860 | 7.13 | 5.83e-05 | 0.000682 | 7 | TPX2/AURKB/STMN1/SKA2/PSRC1/SKA1/SKA3 |
| Malignant epithelial | C3 Mp ECs | GO:0010972 | negative regulation of G2/M transition of mitotic cell cycle | 6/178 | 73/18860 | 8.71 | 6.55e-05 | 0.000762 | 6 | AURKB/PLK1/CDK1/PKMYT1/RAD21/BRCA1 |
| Malignant epithelial | C3 Mp ECs | GO:0044786 | cell cycle DNA replication | 5/178 | 46/18860 | 11.52 | 7.09e-05 | 0.00082 | 5 | TK1/CDK1/FEN1/RAD51/PCNA |
| Malignant epithelial | C3 Mp ECs | GO:0071478 | cellular response to radiation | 9/178 | 188/18860 | 5.07 | 7.61e-05 | 0.000867 | 9 | AURKB/RAD51AP1/NUCKS1/PBK/ECT2/RHNO1/RAD51/PCNA/BRCA1 |
| Malignant epithelial | C3 Mp ECs | GO:0071479 | cellular response to ionizing radiation | 6/178 | 75/18860 | 8.48 | 7.63e-05 | 0.000867 | 6 | RAD51AP1/NUCKS1/ECT2/RHNO1/RAD51/BRCA1 |
| Malignant epithelial | C3 Mp ECs | GO:1902750 | negative regulation of cell cycle G2/M phase transition | 6/178 | 75/18860 | 8.48 | 7.63e-05 | 0.000867 | 6 | AURKB/PLK1/CDK1/PKMYT1/RAD21/BRCA1 |
| Malignant epithelial | C3 Mp ECs | GO:0009116 | nucleoside metabolic process | 5/178 | 47/18860 | 11.27 | 7.88e-05 | 0.00089 | 5 | TK1/CDK1/DTYMK/NUDT1/ERH |
| Malignant epithelial | C3 Mp ECs | GO:0046785 | microtubule polymerization | 7/178 | 110/18860 | 6.74 | 8.33e-05 | 0.000936 | 7 | TPX2/CDK1/STMN1/PSRC1/SKA1/TUBG1/MZT1 |
| Malignant epithelial | C3 Mp ECs | GO:0090224 | regulation of spindle organization | 5/178 | 48/18860 | 11.04 | 8.73e-05 | 0.000975 | 5 | TPX2/PLK1/SPAG5/TACC3/PSRC1 |
| Malignant epithelial | C3 Mp ECs | GO:0032200 | telomere organization | 9/178 | 193/18860 | 4.94 | 9.3e-05 | 0.00103 | 9 | AURKB/NEK2/HNRNPA2B1/RPA3/PIF1/FEN1/RAD51/PCNA/RECQL4 |
| Malignant epithelial | C3 Mp ECs | GO:0006265 | DNA topological change | 3/178 | 10/18860 | 31.79 | 9.45e-05 | 0.00103 | 3 | TOP2A/HMGB2/HMGB1 |
| Malignant epithelial | C3 Mp ECs | GO:0033313 | meiotic cell cycle checkpoint signaling | 3/178 | 10/18860 | 31.79 | 9.45e-05 | 0.00103 | 3 | ZWINT/TRIP13/TTK |
| Malignant epithelial | C3 Mp ECs | GO:0045876 | positive regulation of sister chromatid cohesion | 3/178 | 10/18860 | 31.79 | 9.45e-05 | 0.00103 | 3 | BUB1/FEN1/RAD21 |
| Malignant epithelial | C3 Mp ECs | GO:0051307 | meiotic chromosome separation | 3/178 | 10/18860 | 31.79 | 9.45e-05 | 0.00103 | 3 | ZWINT/TTK/NCAPH |
| Malignant epithelial | C3 Mp ECs | GO:0045814 | negative regulation of gene expression, epigenetic | 9/178 | 197/18860 | 4.84 | 0.000109 | 0.00118 | 9 | LMNB1/OIP5/PHF19/HMGB1/LMNB2/DNMT1/UHRF1/HELLS/BRCA1 |
| Malignant epithelial | C3 Mp ECs | GO:0007077 | mitotic nuclear membrane disassembly | 3/178 | 11/18860 | 28.9 | 0.000129 | 0.00139 | 3 | PLK1/CDK1/VRK1 |
| Malignant epithelial | C3 Mp ECs | GO:0031507 | heterochromatin formation | 8/178 | 163/18860 | 5.2 | 0.000161 | 0.00172 | 8 | LMNB1/OIP5/HMGB1/LMNB2/DNMT1/UHRF1/HELLS/BRCA1 |
| Malignant epithelial | C3 Mp ECs | GO:0007064 | mitotic sister chromatid cohesion | 4/178 | 31/18860 | 13.67 | 0.000198 | 0.00209 | 4 | CDC20/BUB1/CDCA5/RAD21 |
| Malignant epithelial | C3 Mp ECs | GO:0010458 | exit from mitosis | 4/178 | 31/18860 | 13.67 | 0.000198 | 0.00209 | 4 | BIRC5/UBE2C/UBE2S/ANLN |
| Malignant epithelial | C3 Mp ECs | GO:0071897 | DNA biosynthetic process | 8/178 | 168/18860 | 5.05 | 0.000198 | 0.00209 | 8 | CENPF/CCNA2/TK1/CDK1/TYMS/PIF1/PCNA/CDKN2D |
| Malignant epithelial | C3 Mp ECs | GO:0009120 | deoxyribonucleoside metabolic process | 3/178 | 13/18860 | 24.45 | 0.000221 | 0.00231 | 3 | TK1/CDK1/DTYMK |
| Malignant epithelial | C3 Mp ECs | GO:0006261 | DNA-templated DNA replication | 8/178 | 171/18860 | 4.96 | 0.000224 | 0.00233 | 8 | TK1/CDK1/NUCKS1/ORC6/FEN1/RAD51/PCNA/BRCA1 |
| Malignant epithelial | C3 Mp ECs | GO:0031297 | replication fork processing | 5/178 | 59/18860 | 8.98 | 0.000235 | 0.00242 | 5 | CDK1/NUCKS1/RAD51/PCNA/BRCA1 |
| Malignant epithelial | C3 Mp ECs | GO:0072698 | protein localization to microtubule cytoskeleton | 5/178 | 59/18860 | 8.98 | 0.000235 | 0.00242 | 5 | PLK1/AURKA/SPAG5/FAM83D/TTK |
| Malignant epithelial | C3 Mp ECs | GO:0051642 | centrosome localization | 4/178 | 33/18860 | 12.84 | 0.000253 | 0.00259 | 4 | MAD2L1/ASPM/DLGAP5/AURKA |
| Malignant epithelial | C3 Mp ECs | GO:0061842 | microtubule organizing center localization | 4/178 | 33/18860 | 12.84 | 0.000253 | 0.00259 | 4 | MAD2L1/ASPM/DLGAP5/AURKA |
| Malignant epithelial | C3 Mp ECs | GO:0007018 | microtubule-based movement | 14/178 | 496/18860 | 2.99 | 0.000269 | 0.00273 | 14 | DLGAP5/KIF23/KIFC1/CENPE/KIF11/KIF4A/KIF14/KIF20A/TUBB4B/KIF18B/KIF22/KIF2C/KIF15/KIF18A |

| compartment | subcluster | GO_ID | Description | GeneRatio | BgRatio | FoldEnrichment | pvalue | p_adjust | Count | geneID |
|---|---|---|---|---|---|---|---|---|---|---|
| Malignant epithelial | C3 Mp ECs | GO:0090435 | protein localization to nuclear envelope | 3/178 | 14/18860 | 22.7 | 0.000279 | 0.00282 | 3 | PLK1/LMNB1/LMNB2 |
| Malignant epithelial | C3 Mp ECs | GO:0044843 | cell cycle G1/S phase transition | 10/178 | 275/18860 | 3.85 | 0.000293 | 0.00295 | 10 | CCNB1/CDKN3/CCNA2/CCNB2/CDK1/RRM2/KIF14/IQGAP3/FAM83D/CDKN2D |
| Malignant epithelial | C3 Mp ECs | GO:0040020 | regulation of meiotic nuclear division | 4/178 | 35/18860 | 12.11 | 0.00032 | 0.00316 | 4 | ZWINT/CDC20/RAD51AP1/TRIP13 |
| Malignant epithelial | C3 Mp ECs | GO:0006998 | nuclear envelope organization | 5/178 | 63/18860 | 8.41 | 0.00032 | 0.00316 | 5 | PLK1/CDK1/LMNB1/LMNB2/VRK1 |
| Malignant epithelial | C3 Mp ECs | GO:0031146 | SCF-dependent proteasomal ubiquitin-dependent protein catabolic process | 5/178 | 63/18860 | 8.41 | 0.00032 | 0.00316 | 5 | UBE2C/UBE2S/CDC20/KIF14/ANAPC11 |
| Malignant epithelial | C3 Mp ECs | GO:0031503 | protein-containing complex localization | 6/178 | 98/18860 | 6.49 | 0.000333 | 0.00329 | 6 | BIRC5/LMNB1/RAN/LMNB2/KIF2C/MZT1 |
| Malignant epithelial | C3 Mp ECs | GO:0051081 | nuclear membrane disassembly | 3/178 | 15/18860 | 21.19 | 0.000346 | 0.00339 | 3 | PLK1/CDK1/VRK1 |
| Malignant epithelial | C3 Mp ECs | GO:0051054 | positive regulation of DNA metabolic process | 10/178 | 283/18860 | 3.74 | 0.000368 | 0.00359 | 10 | AURKB/CCNA2/FOXM1/CDK1/NEK2/RAD51AP1/HNRNPA2B1/HMGB1/PCNA/BRCA1 |
| Malignant epithelial | C3 Mp ECs | GO:0045005 | DNA-templated DNA replication maintenance of fidelity | 5/178 | 66/18860 | 8.03 | 0.000397 | 0.00386 | 5 | CDK1/NUCKS1/RAD51/PCNA/BRCA1 |
| Malignant epithelial | C3 Mp ECs | GO:0009263 | deoxyribonucleotide biosynthetic process | 3/178 | 16/18860 | 19.87 | 0.000423 | 0.00397 | 3 | RRM2/DTYMK/TYMS |
| Malignant epithelial | C3 Mp ECs | GO:0009265 | 2'-deoxyribonucleotide biosynthetic process | 3/178 | 16/18860 | 19.87 | 0.000423 | 0.00397 | 3 | RRM2/DTYMK/TYMS |
| Malignant epithelial | C3 Mp ECs | GO:0030397 | membrane disassembly | 3/178 | 16/18860 | 19.87 | 0.000423 | 0.00397 | 3 | PLK1/CDK1/VRK1 |
| Malignant epithelial | C3 Mp ECs | GO:0036260 | RNA capping | 3/178 | 16/18860 | 19.87 | 0.000423 | 0.00397 | 3 | SNRPB/SNRPG/SNRPD1 |
| Malignant epithelial | C3 Mp ECs | GO:0046385 | deoxyribose phosphate biosynthetic process | 3/178 | 16/18860 | 19.87 | 0.000423 | 0.00397 | 3 | RRM2/DTYMK/TYMS |
| Malignant epithelial | C3 Mp ECs | GO:0048308 | organelle inheritance | 3/178 | 16/18860 | 19.87 | 0.000423 | 0.00397 | 3 | PLK1/CDK1/VRK1 |
| Malignant epithelial | C3 Mp ECs | GO:0048313 | Golgi inheritance | 3/178 | 16/18860 | 19.87 | 0.000423 | 0.00397 | 3 | PLK1/CDK1/VRK1 |
| Malignant epithelial | C3 Mp ECs | GO:0044380 | protein localization to cytoskeleton | 5/178 | 67/18860 | 7.91 | 0.000426 | 0.00398 | 5 | PLK1/AURKA/SPAG5/FAM83D/TTK |
| Malignant epithelial | C3 Mp ECs | GO:0045143 | homologous chromosome segregation | 5/178 | 68/18860 | 7.79 | 0.000456 | 0.00424 | 5 | PTTG1/ZWINT/PLK1/MND1/TRIP13 |
| Malignant epithelial | C3 Mp ECs | GO:0030866 | cortical actin cytoskeleton organization | 4/178 | 39/18860 | 10.87 | 0.000487 | 0.00451 | 4 | ANLN/RACGAP1/IQGAP3/ECT2 |
| Malignant epithelial | C3 Mp ECs | GO:0051177 | meiotic sister chromatid cohesion | 3/178 | 17/18860 | 18.7 | 0.00051 | 0.00468 | 3 | BUB1/BUB1B/RAD21 |
| Malignant epithelial | C3 Mp ECs | GO:1902969 | mitotic DNA replication | 3/178 | 17/18860 | 18.7 | 0.00051 | 0.00468 | 3 | TK1/CDK1/RAD51 |
| Malignant epithelial | C3 Mp ECs | GO:1901657 | glycosyl compound metabolic process | 5/178 | 71/18860 | 7.46 | 0.000557 | 0.00509 | 5 | TK1/CDK1/DTYMK/NUDT1/ERH |
| Malignant epithelial | C3 Mp ECs | GO:0000082 | G1/S transition of mitotic cell cycle | 9/178 | 246/18860 | 3.88 | 0.000563 | 0.00512 | 9 | CCNB1/CDKN3/CCNA2/CCNB2/CDK1/RRM2/KIF14/IQGAP3/CDKN2D |
| Malignant epithelial | C3 Mp ECs | GO:0031398 | positive regulation of protein ubiquitination | 6/178 | 109/18860 | 5.83 | 0.000589 | 0.00533 | 6 | UBE2C/UBE2S/PLK1/CDC20/FANCI/CAV1 |
| Malignant epithelial | C3 Mp ECs | GO:0070979 | protein K11-linked ubiquitination | 4/178 | 41/18860 | 10.34 | 0.000591 | 0.00533 | 4 | UBE2C/UBE2T/UBE2S/ANAPC11 |
| Malignant epithelial | C3 Mp ECs | GO:0007063 | regulation of sister chromatid cohesion | 3/178 | 18/18860 | 17.66 | 0.000608 | 0.00543 | 3 | BUB1/FEN1/RAD21 |
| Malignant epithelial | C3 Mp ECs | GO:0007096 | regulation of exit from mitosis | 3/178 | 18/18860 | 17.66 | 0.000608 | 0.00543 | 3 | BIRC5/UBE2C/ANLN |
| Malignant epithelial | C3 Mp ECs | GO:0040029 | epigenetic regulation of gene expression | 10/178 | 305/18860 | 3.47 | 0.000659 | 0.00586 | 10 | LMNB1/OIP5/PHF19/HMGB1/LMNB2/ATAD2/DNMT1/UHRF1/HELLS/BRCA1 |
| Malignant epithelial | C3 Mp ECs | GO:0072527 | pyrimidine-containing compound metabolic process | 5/178 | 74/18860 | 7.16 | 0.000674 | 0.00596 | 5 | TK1/CDK1/DTYMK/TYMS/ERH |
| Malignant epithelial | C3 Mp ECs | GO:0022411 | cellular component disassembly | 13/178 | 483/18860 | 2.85 | 0.000693 | 0.00611 | 13 | TPX2/AURKB/TOP2A/PLK1/CDK1/AURKA/STMN1/KIF18B/PIF1/ATAD2/VRK1/KIF2C/KIF18A |
| Malignant epithelial | C3 Mp ECs | GO:0036297 | interstrand cross-link repair | 4/178 | 43/18860 | 9.86 | 0.00071 | 0.00623 | 4 | RAD51AP1/NUCKS1/FANCI/RAD51 |
| Malignant epithelial | C3 Mp ECs | GO:0060236 | regulation of mitotic spindle organization | 4/178 | 44/18860 | 9.63 | 0.000776 | 0.00677 | 4 | TPX2/PLK1/TACC3/PSRC1 |
| Malignant epithelial | C3 Mp ECs | GO:0043085 | positive regulation of catalytic activity | 10/178 | 318/18860 | 3.33 | 0.000906 | 0.00788 | 10 | TPX2/UBE2C/UBE2S/PLK1/CDC20/ARHGAP11A/CENPE/KIF14/ECT2/PCNA |
| Malignant epithelial | C3 Mp ECs | GO:0072528 | pyrimidine-containing compound biosynthetic process | 4/178 | 46/18860 | 9.21 | 0.000919 | 0.00795 | 4 | TK1/CDK1/DTYMK/TYMS |

| compartment | subcluster | GO_ID | Description | GeneRatio | BgRatio | FoldEnrichment | pvalue | p_adjust | Count | geneID |
|---|---|---|---|---|---|---|---|---|---|---|
| Malignant epithelial | C3 Mp ECs | GO:0009163 | nucleoside biosynthetic process | 3/178 | 21/18860 | 15.14 | 0.00097 | 0.00833 | 3 | TK1/CDK1/DTYMK |
| Malignant epithelial | C3 Mp ECs | GO:0034404 | nucleobase-containing small molecule biosynthetic process | 3/178 | 21/18860 | 15.14 | 0.00097 | 0.00833 | 3 | TK1/CDK1/DTYMK |
| Malignant epithelial | C3 Mp ECs | GO:0051261 | protein depolymerization | 6/178 | 121/18860 | 5.25 | 0.00102 | 0.00871 | 6 | TPX2/AURKB/STMN1/KIF18B/KIF2C/KIF18A |
| Malignant epithelial | C3 Mp ECs | GO:0000387 | spliceosomal snRNP assembly | 4/178 | 51/18860 | 8.31 | 0.00136 | 0.0115 | 4 | SNRPB/LSM4/SNRPG/SNRPD1 |
| Malignant epithelial | C3 Mp ECs | GO:0061952 | midbody abscission | 3/178 | 24/18860 | 13.24 | 0.00145 | 0.0123 | 3 | CEP55/AURKB/KIF20A |
| Malignant epithelial | C3 Mp ECs | GO:0006284 | base-excision repair | 4/178 | 52/18860 | 8.15 | 0.00146 | 0.0123 | 4 | HMGB1/RPA3/FEN1/PCNA |
| Malignant epithelial | C3 Mp ECs | GO:1903322 | positive regulation of protein modification by small protein conjugation or removal | 6/178 | 132/18860 | 4.82 | 0.00159 | 0.0134 | 6 | UBE2C/UBE2S/PLK1/CDC20/FANCI/CAV1 |
| Malignant epithelial | C3 Mp ECs | GO:0000377 | RNA splicing, via transesterification reactions with bulged adenosine as nucleophile | 10/178 | 344/18860 | 3.08 | 0.00163 | 0.0135 | 10 | LSM5/SNRPB/HNRNPA2B1/LSM4/SNRPG/DDX39A/HNRNPA3/PSIP1/RBM8A/SNRPD1 |
| Malignant epithelial | C3 Mp ECs | GO:0000398 | mRNA splicing, via spliceosome | 10/178 | 344/18860 | 3.08 | 0.00163 | 0.0135 | 10 | LSM5/SNRPB/HNRNPA2B1/LSM4/SNRPG/DDX39A/HNRNPA3/PSIP1/RBM8A/SNRPD1 |
| Malignant epithelial | C3 Mp ECs | GO:1903241 | U2-type prespliceosome assembly | 3/178 | 25/18860 | 12.71 | 0.00163 | 0.0135 | 3 | SNRPB/SNRPG/SNRPD1 |
| Malignant epithelial | C3 Mp ECs | GO:1903902 | positive regulation of viral life cycle | 3/178 | 25/18860 | 12.71 | 0.00163 | 0.0135 | 3 | KPNA2/HMGB1/LGALS1 |
| Malignant epithelial | C3 Mp ECs | GO:1901875 | positive regulation of post-translational protein modification | 6/178 | 133/18860 | 4.78 | 0.00166 | 0.0136 | 6 | UBE2C/UBE2S/PLK1/CDC20/FANCI/CAV1 |
| Malignant epithelial | C3 Mp ECs | GO:0000375 | RNA splicing, via transesterification reactions | 10/178 | 348/18860 | 3.04 | 0.00178 | 0.0146 | 10 | LSM5/SNRPB/HNRNPA2B1/LSM4/SNRPG/DDX39A/HNRNPA3/PSIP1/RBM8A/SNRPD1 |
| Malignant epithelial | C3 Mp ECs | GO:0030865 | cortical cytoskeleton organization | 4/178 | 55/18860 | 7.71 | 0.0018 | 0.0147 | 4 | ANLN/RACGAP1/IQGAP3/ECT2 |
| Malignant epithelial | C3 Mp ECs | GO:0051258 | protein polymerization | 9/178 | 291/18860 | 3.28 | 0.00182 | 0.0148 | 9 | TPX2/UBE2C/UBE2S/CDK1/STMN1/PSRC1/SKA1/TUBG1/MZT1 |
| Malignant epithelial | C3 Mp ECs | GO:0085020 | protein K6-linked ubiquitination | 3/178 | 26/18860 | 12.23 | 0.00183 | 0.0148 | 3 | UBE2T/UBE2S/BRCA1 |
| Malignant epithelial | C3 Mp ECs | GO:1990166 | protein localization to site of double-strand break | 3/178 | 26/18860 | 12.23 | 0.00183 | 0.0148 | 3 | PLK1/CDK1/RHNO1 |
| Malignant epithelial | C3 Mp ECs | GO:0006282 | regulation of DNA repair | 8/178 | 236/18860 | 3.59 | 0.00184 | 0.0148 | 8 | FOXM1/PLK1/RAD51AP1/HMGB1/RAD51/PCNA/BRCA1/PARPBP |
| Malignant epithelial | C3 Mp ECs | GO:1901659 | glycosyl compound biosynthetic process | 3/178 | 27/18860 | 11.77 | 0.00205 | 0.0164 | 3 | TK1/CDK1/DTYMK |
| Malignant epithelial | C3 Mp ECs | GO:0007097 | nuclear migration | 3/178 | 28/18860 | 11.35 | 0.00228 | 0.0181 | 3 | LMNB1/TACC3/LMNB2 |
| Malignant epithelial | C3 Mp ECs | GO:0052372 | modulation by symbiont of entry into host | 3/178 | 28/18860 | 11.35 | 0.00228 | 0.0181 | 3 | HMGB1/CAV1/LGALS1 |
| Malignant epithelial | C3 Mp ECs | GO:0010971 | positive regulation of G2/M transition of mitotic cell cycle | 3/178 | 29/18860 | 10.96 | 0.00252 | 0.0199 | 3 | CCNB1/CDK1/CDC25C |
| Malignant epithelial | C3 Mp ECs | GO:0010212 | response to ionizing radiation | 6/178 | 145/18860 | 4.38 | 0.00256 | 0.0202 | 6 | RAD51AP1/NUCKS1/ECT2/RHNO1/RAD51/BRCA1 |
| Malignant epithelial | C3 Mp ECs | GO:0071168 | protein localization to chromatin | 4/178 | 62/18860 | 6.84 | 0.00279 | 0.0219 | 4 | CENPA/PLK1/VRK1/RAD21 |
| Malignant epithelial | C3 Mp ECs | GO:0031396 | regulation of protein ubiquitination | 7/178 | 200/18860 | 3.71 | 0.00295 | 0.0229 | 7 | UBE2C/MAD2L1/UBE2S/PLK1/CDC20/FANCI/CAV1 |
| Malignant epithelial | C3 Mp ECs | GO:0042770 | signal transduction in response to DNA damage | 7/178 | 200/18860 | 3.71 | 0.00295 | 0.0229 | 7 | FOXM1/GTSE1/PLK1/CDK1/RHNO1/RAD51/BRCA1 |
| Malignant epithelial | C3 Mp ECs | GO:0031401 | positive regulation of protein modification process | 10/178 | 375/18860 | 2.83 | 0.00305 | 0.0236 | 10 | TPX2/UBE2C/UBE2S/PLK1/CDC20/CENPE/KIF14/ECT2/FANCI/CAV1 |
| Malignant epithelial | C3 Mp ECs | GO:0048524 | positive regulation of viral process | 4/178 | 64/18860 | 6.62 | 0.00314 | 0.0242 | 4 | TOP2A/KPNA2/HMGB1/LGALS1 |
| Malignant epithelial | C3 Mp ECs | GO:1902751 | positive regulation of cell cycle G2/M phase transition | 3/178 | 32/18860 | 9.93 | 0.00335 | 0.0258 | 3 | CCNB1/CDK1/CDC25C |
| Malignant epithelial | C3 Mp ECs | GO:0000079 | regulation of cyclin-dependent protein serine/threonine kinase activity | 3/178 | 33/18860 | 9.63 | 0.00366 | 0.0279 | 3 | CDKN3/CDC25C/PKMYT1 |
| Malignant epithelial | C3 Mp ECs | GO:0051647 | nucleus localization | 3/178 | 33/18860 | 9.63 | 0.00366 | 0.0279 | 3 | LMNB1/TACC3/LMNB2 |
| Malignant epithelial | C3 Mp ECs | GO:0032984 | protein-containing complex disassembly | 8/178 | 264/18860 | 3.21 | 0.00367 | 0.0279 | 8 | TPX2/AURKB/STMN1/KIF18B/PIF1/ATAD2/KIF2C/KIF18A |

| compartment | subcluster | GO_ID | Description | GeneRatio | BgRatio | FoldEnrichment | pvalue | p_adjust | Count | geneID |
|---|---|---|---|---|---|---|---|---|---|---|
| Malignant epithelial | C3 Mp ECs | GO:0070303 | negative regulation of stress-activated protein kinase signaling cascade | 2/178 | 10/18860 | 21.19 | 0.00379 | 0.0286 | 2 | FOXM1/PBK |
| Malignant epithelial | C3 Mp ECs | GO:0097681 | double-strand break repair via alternative nonhomologous end joining | 2/178 | 10/18860 | 21.19 | 0.00379 | 0.0286 | 2 | PLK1/RHNO1 |
| Malignant epithelial | C3 Mp ECs | GO:0009411 | response to UV | 6/178 | 159/18860 | 4.0 | 0.00404 | 0.0303 | 6 | AURKB/PBK/RHNO1/FEN1/PCNA/CDKN2D |
| Malignant epithelial | C3 Mp ECs | GO:0006997 | nucleus organization | 6/178 | 160/18860 | 3.97 | 0.00416 | 0.0312 | 6 | HMGB2/PLK1/CDK1/LMNB1/LMNB2/VRK1 |
| Malignant epithelial | C3 Mp ECs | GO:0009314 | response to radiation | 11/178 | 462/18860 | 2.52 | 0.00453 | 0.0338 | 11 | AURKB/RAD51AP1/NUCKS1/PBK/ECT2/RHNO1/FEN1/RAD51/PCNA/CDKN2D/BRCA1 |
| Malignant epithelial | C3 Mp ECs | GO:0048144 | fibroblast proliferation | 5/178 | 114/18860 | 4.65 | 0.00454 | 0.0338 | 5 | CCNB1/CCNA2/CDK1/CKS2/CAV1 |
| Malignant epithelial | C3 Mp ECs | GO:0045835 | negative regulation of meiotic nuclear division | 2/178 | 11/18860 | 19.26 | 0.00461 | 0.0341 | 2 | ZWINT/TRIP13 |
| Malignant epithelial | C3 Mp ECs | GO:0007095 | mitotic G2 DNA damage checkpoint signaling | 3/178 | 36/18860 | 8.83 | 0.0047 | 0.0347 | 3 | PLK1/CDK1/BRCA1 |
| Malignant epithelial | C3 Mp ECs | GO:0000077 | DNA damage checkpoint signaling | 5/178 | 117/18860 | 4.53 | 0.00507 | 0.0373 | 5 | PLK1/CDK1/RHNO1/RAD51/BRCA1 |
| Malignant epithelial | C3 Mp ECs | GO:0000966 | RNA 5'-end processing | 3/178 | 38/18860 | 8.36 | 0.00547 | 0.0394 | 3 | SNRPB/SNRPG/SNRPD1 |
| Malignant epithelial | C3 Mp ECs | GO:0001556 | oocyte maturation | 3/178 | 38/18860 | 8.36 | 0.00547 | 0.0394 | 3 | CCNB1/AURKA/TRIP13 |
| Malignant epithelial | C3 Mp ECs | GO:0031112 | positive regulation of microtubule polymerization or depolymerization | 3/178 | 38/18860 | 8.36 | 0.00547 | 0.0394 | 3 | AURKB/PSRC1/SKA1 |
| Malignant epithelial | C3 Mp ECs | GO:0071539 | protein localization to centrosome | 3/178 | 38/18860 | 8.36 | 0.00547 | 0.0394 | 3 | PLK1/AURKA/SPAG5 |
| Malignant epithelial | C3 Mp ECs | GO:0035519 | protein K29-linked ubiquitination | 2/178 | 12/18860 | 17.66 | 0.00549 | 0.0394 | 2 | UBE2T/UBE2S |
| Malignant epithelial | C3 Mp ECs | GO:0070601 | centromeric sister chromatid cohesion | 2/178 | 12/18860 | 17.66 | 0.00549 | 0.0394 | 2 | BUB1/BUB1B |
| Malignant epithelial | C3 Mp ECs | GO:0072697 | protein localization to cell cortex | 2/178 | 12/18860 | 17.66 | 0.00549 | 0.0394 | 2 | PLK1/SAPCD2 |
| Malignant epithelial | C3 Mp ECs | GO:0032147 | activation of protein kinase activity | 3/178 | 39/18860 | 8.15 | 0.00589 | 0.0419 | 3 | TPX2/KIF14/ECT2 |
| Malignant epithelial | C3 Mp ECs | GO:1905508 | protein localization to microtubule organizing center | 3/178 | 39/18860 | 8.15 | 0.00589 | 0.0419 | 3 | PLK1/AURKA/SPAG5 |
| Malignant epithelial | C3 Mp ECs | GO:0042307 | positive regulation of protein import into nucleus | 3/178 | 40/18860 | 7.95 | 0.00632 | 0.0448 | 3 | CDK1/RAN/ECT2 |
| Malignant epithelial | C3 Mp ECs | GO:2000242 | negative regulation of reproductive process | 4/178 | 78/18860 | 5.43 | 0.00636 | 0.0449 | 4 | ZWINT/PKMYT1/TRIP13/TTK |
| Malignant epithelial | C3 Mp ECs | GO:0055015 | ventricular cardiac muscle cell development | 2/178 | 13/18860 | 16.3 | 0.00645 | 0.0453 | 2 | CCNB1/CDK1 |
| Malignant epithelial | C3 Mp ECs | GO:0061523 | cilium disassembly | 2/178 | 13/18860 | 16.3 | 0.00645 | 0.0453 | 2 | CDK1/AURKA |
| Malignant epithelial | C3 Mp ECs | GO:0071214 | cellular response to abiotic stimulus | 9/178 | 355/18860 | 2.69 | 0.00671 | 0.0467 | 9 | AURKB/RAD51AP1/NUCKS1/PBK/ECT2/RHNO1/RAD51/PCNA/BRCA1 |
| Malignant epithelial | C3 Mp ECs | GO:0104004 | cellular response to environmental stimulus | 9/178 | 355/18860 | 2.69 | 0.00671 | 0.0467 | 9 | AURKB/RAD51AP1/NUCKS1/PBK/ECT2/RHNO1/RAD51/PCNA/BRCA1 |
| Malignant epithelial | C3 Mp ECs | GO:0045859 | regulation of protein kinase activity | 7/178 | 234/18860 | 3.17 | 0.0069 | 0.0478 | 7 | TPX2/CDKN3/CENPE/KIF14/CDC25C/PKMYT1/ECT2 |
| Malignant epithelial | C3 Mp ECs | GO:0031570 | DNA integrity checkpoint signaling | 5/178 | 126/18860 | 4.2 | 0.0069 | 0.0478 | 5 | PLK1/CDK1/RHNO1/RAD51/BRCA1 |
| Malignant epithelial | C3 Mp ECs | GO:0006303 | double-strand break repair via nonhomologous end joining | 4/178 | 80/18860 | 5.3 | 0.00695 | 0.0479 | 4 | HMGB2/PLK1/HMGB1/RHNO1 |
| Malignant epithelial | C3 Mp ECs | GO:0043903 | regulation of biological process involved in symbiotic interaction | 3/178 | 42/18860 | 7.57 | 0.00724 | 0.0498 | 3 | HMGB1/CAV1/LGALS1 |
| Malignant epithelial | C4 T/r-h ECs | GO:0002181 | cytoplasmic translation | 71/181 | 208/18860 | 35.57 | 1.82e-94 | 3.96e-91 | 71 | RPL31/RPL37A/RPL36/RPS17/RPL35A/RPL41/RPL26/RPS25/RPS15/RPS28/RPLP2/RPS15A/RPL34/RPS29/RP… |
| Malignant epithelial | C4 T/r-h ECs | GO:0022613 | ribonucleoprotein complex biogenesis | 42/181 | 496/18860 | 8.82 | 7.38e-28 | 8.02e-25 | 42 | RPS17/RPL35A/RPL26/RPS25/RPS15/RPS28/RPS15A/RPS19/RPL35/RPS13/RPS24/RPL38/RPL24/RPL14/RPL… |
| Malignant epithelial | C4 T/r-h ECs | GO:0042254 | ribosome biogenesis | 35/181 | 318/18860 | 11.47 | 4.48e-27 | 3.24e-24 | 35 | RPS17/RPL35A/RPL26/RPS25/RPS15/RPS28/RPS15A/RPS19/RPL35/RPS13/RPS24/RPL38/RPL24/RPL14/RPL… |

| compartment | subcluster | GO_ID | Description | GeneRatio | BgRatio | FoldEnrichment | pvalue | p_adjust | Count | geneID |
|---|---|---|---|---|---|---|---|---|---|---|
| Malignant epithelial | C4 T/r-h ECs | GO:0042274 | ribosomal small subunit biogenesis | 23/181 | 107/18860 | 22.4 | 6.64e-25 | 3.61e-22 | 23 | RPS17/RPS25/RPS15/RPS28/RPS15A/RPS19/RPS13/RPS24/RPS23/RPS7/FAU/RPS27A/RPS27/RPS21/RPS8/R… |
| Malignant epithelial | C4 T/r-h ECs | GO:0006364 | rRNA processing | 21/181 | 223/18860 | 9.81 | 3.74e-15 | 1.63e-12 | 21 | RPS17/RPL35A/RPL26/RPS25/RPS15/RPS28/RPS19/RPL35/RPS24/RPL14/RPL11/RPS7/RPL7/RPS27/RPS21/… |
| Malignant epithelial | C4 T/r-h ECs | GO:0016072 | rRNA metabolic process | 21/181 | 270/18860 | 8.1 | 1.68e-13 | 6.09e-11 | 21 | RPS17/RPL35A/RPL26/RPS25/RPS15/RPS28/RPS19/RPL35/RPS24/RPL14/RPL11/RPS7/RPL7/RPS27/RPS21/… |
| Malignant epithelial | C4 T/r-h ECs | GO:0022618 | protein-RNA complex assembly | 18/181 | 231/18860 | 8.12 | 9.61e-12 | 2.9e-09 | 18 | RPS15/RPS28/RPS19/RPL38/RPL24/RPL11/FAU/RPS27/RPL13A/RPS6/RPS14/SNRPD2/RPL5/LUC7L3/DHX9/S… |
| Malignant epithelial | C4 T/r-h ECs | GO:0042255 | ribosome assembly | 11/181 | 60/18860 | 19.1 | 1.07e-11 | 2.9e-09 | 11 | RPS15/RPS28/RPS19/RPL38/RPL24/RPL11/FAU/RPS27/RPS6/RPS14/RPL5 |
| Malignant epithelial | C4 T/r-h ECs | GO:0071826 | protein-RNA complex organization | 18/181 | 240/18860 | 7.81 | 1.82e-11 | 4.4e-09 | 18 | RPS15/RPS28/RPS19/RPL38/RPL24/RPL11/FAU/RPS27/RPL13A/RPS6/RPS14/SNRPD2/RPL5/LUC7L3/DHX9/S… |
| Malignant epithelial | C4 T/r-h ECs | GO:0000028 | ribosomal small subunit assembly | 7/181 | 20/18860 | 36.47 | 4.66e-10 | 1.01e-07 | 7 | RPS15/RPS28/RPS19/FAU/RPS27/RPS6/RPS14 |
| Malignant epithelial | C4 T/r-h ECs | GO:1901798 | positive regulation of signal transduction by p53 class mediator | 7/181 | 36/18860 | 20.26 | 4.41e-08 | 8.7e-06 | 7 | RPL26/RPS15/RPS20/RPL37/RPL11/RPS7/RPL23 |
| Malignant epithelial | C4 T/r-h ECs | GO:0042273 | ribosomal large subunit biogenesis | 8/181 | 69/18860 | 12.08 | 3.13e-07 | 5.66e-05 | 8 | RPL35A/RPL26/RPL35/RPL24/RPL14/RPL11/RPL7/RPL5 |
| Malignant epithelial | C4 T/r-h ECs | GO:0051444 | negative regulation of ubiquitin-protein transferase activity | 4/181 | 11/18860 | 37.89 | 2.57e-06 | 0.000429 | 4 | RPL11/RPS7/RPL23/RPL5 |
| Malignant epithelial | C4 T/r-h ECs | GO:1901796 | regulation of signal transduction by p53 class mediator | 9/181 | 123/18860 | 7.62 | 2.91e-06 | 0.000452 | 9 | RPL26/RPS15/RPS20/RPL37/RPL11/RPS7/RPL23/MIF/RPL5 |
| Malignant epithelial | C4 T/r-h ECs | GO:0051438 | regulation of ubiquitin-protein transferase activity | 5/181 | 26/18860 | 20.04 | 4.3e-06 | 0.000623 | 5 | RPL11/RPS7/RPL23/RPL5/ARRDC3 |
| Malignant epithelial | C4 T/r-h ECs | GO:0140694 | membraneless organelle assembly | 16/181 | 446/18860 | 3.74 | 6.61e-06 | 0.000897 | 16 | RPS15/RPS28/RPS19/RPL38/RPL24/RPL11/RPS23/FAU/RPS27/RPS6/RPS14/MZT1/RPL5/DYNC1H1/EIF2S1/F… |
| Malignant epithelial | C4 T/r-h ECs | GO:1904666 | regulation of ubiquitin protein ligase activity | 4/181 | 15/18860 | 27.79 | 1.03e-05 | 0.00132 | 4 | RPL11/RPS7/RPL23/RPL5 |
| Malignant epithelial | C4 T/r-h ECs | GO:0030490 | maturation of SSU-rRNA | 6/181 | 54/18860 | 11.58 | 1.27e-05 | 0.00153 | 6 | RPS28/RPS19/RPS21/RPS8/RPS16/RPS14 |
| Malignant epithelial | C4 T/r-h ECs | GO:0072331 | signal transduction by p53 class mediator | 10/181 | 196/18860 | 5.32 | 2.03e-05 | 0.00232 | 10 | RPL26/RPS15/RPS20/RPL37/RPL11/RPS7/RPL23/MIF/RPL5/HINT1 |
| Malignant epithelial | C4 T/r-h ECs | GO:2000434 | regulation of protein neddylation | 4/181 | 20/18860 | 20.84 | 3.53e-05 | 0.00383 | 4 | RPL11/COPS7A/RPL5/COPS2 |
| Malignant epithelial | C4 T/r-h ECs | GO:0045116 | protein neddylation | 4/181 | 28/18860 | 14.89 | 0.00014 | 0.0145 | 4 | RPL11/COPS7A/RPL5/COPS2 |
| Malignant epithelial | C4 T/r-h ECs | GO:0000462 | maturation of SSU-rRNA from tricistronic rRNA transcript (SSU-rRNA, 5.8S rRNA, LSU-rRNA) | 4/181 | 35/18860 | 11.91 | 0.000341 | 0.0336 | 4 | RPS19/RPS21/RPS8/RPS16 |
| Malignant epithelial | C5 Am/CEA-like ECs | GO:0010669 | epithelial structure maintenance | 6/171 | 41/18860 | 16.14 | 1.76e-06 | 0.00386 | 6 | MUC13/CXADR/TFF1/TFF3/RBP4/LSR |
| Malignant epithelial | C5 Am/CEA-like ECs | GO:0032536 | regulation of cell projection size | 4/171 | 13/18860 | 33.94 | 4.38e-06 | 0.00386 | 4 | CDHR5/EZR/PLS1/CDHR2 |
| Malignant epithelial | C5 Am/CEA-like ECs | GO:0046598 | positive regulation of viral entry into host cell | 4/171 | 15/18860 | 29.41 | 8.24e-06 | 0.00386 | 4 | TMPRSS2/SMPD1/TMPRSS4/P4HB |
| Malignant epithelial | C5 Am/CEA-like ECs | GO:0075294 | positive regulation by symbiont of entry into host | 4/171 | 15/18860 | 29.41 | 8.24e-06 | 0.00386 | 4 | TMPRSS2/SMPD1/TMPRSS4/P4HB |
| Malignant epithelial | C5 Am/CEA-like ECs | GO:0007586 | digestion | 9/171 | 148/18860 | 6.71 | 8.36e-06 | 0.00386 | 9 | MUC13/PRSS3/EZR/HEPH/PLS1/TFF1/TFF3/RBP4/FABP2 |
| Malignant epithelial | C5 Am/CEA-like ECs | GO:0098742 | cell-cell adhesion via plasma-membrane adhesion molecules | 12/171 | 279/18860 | 4.74 | 9.46e-06 | 0.00386 | 12 | CEACAM5/CEACAM6/CDH17/CDHR5/BMP2/CXADR/CEACAM1/CD164/CLDN7/CLDN3/CDH1/CDHR2 |

| compartment | subcluster | GO_ID | Description | GeneRatio | BgRatio | FoldEnrichment | pvalue | p_adjust | Count | geneID |
|---|---|---|---|---|---|---|---|---|---|---|
| Malignant epithelial | C5 Am/CEA-like ECs | GO:0022600 | digestive system process | 8/171 | 116/18860 | 7.61 | 1.07e-05 | 0.00386 | 8 | MUC13/EZR/HEPH/PLS1/TFF1/TFF3/RBP4/FABP2 |
| Malignant epithelial | C5 Am/CEA-like ECs | GO:0032530 | regulation of microvillus organization | 4/171 | 18/18860 | 24.51 | 1.81e-05 | 0.0057 | 4 | CDHR5/EZR/PLS1/CDHR2 |
| Malignant epithelial | C5 Am/CEA-like ECs | GO:0046596 | regulation of viral entry into host cell | 4/171 | 21/18860 | 21.01 | 3.46e-05 | 0.00911 | 4 | TMPRSS2/SMPD1/TMPRSS4/P4HB |
| Malignant epithelial | C5 Am/CEA-like ECs | GO:1903828 | negative regulation of protein localization | 10/171 | 222/18860 | 4.97 | 3.61e-05 | 0.00911 | 10 | TMBIM1/FAM3D/TP53INP2/RSAD2/TMEM59/NDFIP2/TAX1BP3/RAB11FIP1/UFM1/DPH3 |
| Malignant epithelial | C5 Am/CEA-like ECs | GO:0050792 | regulation of viral process | 8/171 | 141/18860 | 6.26 | 4.41e-05 | 0.0101 | 8 | TMPRSS2/SMPD1/RSAD2/TRIM31/ISG20/DYNLT1/TMPRSS4/P4HB |
| Malignant epithelial | C5 Am/CEA-like ECs | GO:0032528 | microvillus organization | 4/171 | 25/18860 | 17.65 | 7.11e-05 | 0.0138 | 4 | CDHR5/EZR/PLS1/CDHR2 |
| Malignant epithelial | C5 Am/CEA-like ECs | GO:1903902 | positive regulation of viral life cycle | 4/171 | 25/18860 | 17.65 | 7.11e-05 | 0.0138 | 4 | TMPRSS2/SMPD1/TMPRSS4/P4HB |
| Malignant epithelial | C5 Am/CEA-like ECs | GO:0016236 | macroautophagy | 13/171 | 403/18860 | 3.56 | 8.15e-05 | 0.0147 | 13 | MAP1LC3B/HK2/TP53INP2/QSOX1/RNF186/ATP6V1G1/SQSTM1/RHEB/MAP1LC3A/ATP6V0D1/EFNB1/UFM1/ATP6… |
| Malignant epithelial | C5 Am/CEA-like ECs | GO:0030277 | maintenance of gastrointestinal epithelium | 4/171 | 27/18860 | 16.34 | 9.73e-05 | 0.0164 | 4 | MUC13/TFF1/TFF3/RBP4 |
| Malignant epithelial | C5 Am/CEA-like ECs | GO:0042572 | retinol metabolic process | 5/171 | 52/18860 | 10.61 | 0.000106 | 0.0166 | 5 | CYP3A5/DHRS9/RBP4/SDR16C5/PNPLA2 |
| Malignant epithelial | C5 Am/CEA-like ECs | GO:0052372 | modulation by symbiont of entry into host | 4/171 | 28/18860 | 15.76 | 0.000113 | 0.0166 | 4 | TMPRSS2/SMPD1/TMPRSS4/P4HB |
| Malignant epithelial | C5 Am/CEA-like ECs | GO:1903076 | regulation of protein localization to plasma membrane | 7/171 | 121/18860 | 6.38 | 0.000118 | 0.0166 | 7 | AGR2/EZR/TMBIM1/PLS1/SQSTM1/TMEM59/SPTBN1 |
| Malignant epithelial | C5 Am/CEA-like ECs | GO:0120254 | olefinic compound metabolic process | 8/171 | 169/18860 | 5.22 | 0.000157 | 0.0208 | 8 | CYP3A5/BMP2/DHRS9/CYP4F3/MGLL/RBP4/SDR16C5/PNPLA2 |
| Malignant epithelial | C5 Am/CEA-like ECs | GO:1905475 | regulation of protein localization to membrane | 8/171 | 177/18860 | 4.98 | 0.000215 | 0.0272 | 8 | AGR2/EZR/TMBIM1/PLS1/SQSTM1/TMEM59/SPTBN1/GSN |
| Malignant epithelial | C5 Am/CEA-like ECs | GO:0046718 | symbiont entry into host cell | 7/171 | 136/18860 | 5.68 | 0.000244 | 0.0284 | 7 | CD55/TMPRSS2/CXADR/SMPD1/DYNLT1/TMPRSS4/P4HB |
| Malignant epithelial | C5 Am/CEA-like ECs | GO:0009133 | nucleoside diphosphate biosynthetic process | 3/171 | 14/18860 | 23.63 | 0.000248 | 0.0284 | 3 | GUK1/ENTPD8/CMPK1 |
| Malignant epithelial | C5 Am/CEA-like ECs | GO:0051701 | biological process involved in interaction with host | 8/171 | 185/18860 | 4.77 | 0.000291 | 0.0319 | 8 | CD55/TMPRSS2/CXADR/SMPD1/TRIM31/DYNLT1/TMPRSS4/P4HB |
| Malignant epithelial | C5 Am/CEA-like ECs | GO:0044409 | symbiont entry into host | 7/171 | 141/18860 | 5.48 | 0.000304 | 0.032 | 7 | CD55/TMPRSS2/CXADR/SMPD1/DYNLT1/TMPRSS4/P4HB |
| Malignant epithelial | C5 Am/CEA-like ECs | GO:1903078 | positive regulation of protein localization to plasma membrane | 5/171 | 68/18860 | 8.11 | 0.00038 | 0.0375 | 5 | AGR2/EZR/PLS1/SQSTM1/SPTBN1 |
| Malignant epithelial | C5 Am/CEA-like ECs | GO:0034308 | primary alcohol metabolic process | 6/171 | 105/18860 | 6.3 | 0.00039 | 0.0375 | 6 | CYP3A5/BMP2/DHRS9/RBP4/SDR16C5/PNPLA2 |
| Malignant epithelial | C5 Am/CEA-like ECs | GO:0019058 | viral life cycle | 10/171 | 298/18860 | 3.7 | 0.000402 | 0.0375 | 10 | CD55/TMPRSS2/CXADR/SMPD1/RSAD2/TRIM31/ISG20/DYNLT1/TMPRSS4/P4HB |
| Malignant epithelial | C5 Am/CEA-like ECs | GO:0070482 | response to oxygen levels | 11/171 | 357/18860 | 3.4 | 0.000427 | 0.0385 | 11 | BMP2/CD24/CLDN3/HK2/FABP1/ATP6V1G1/NDRG1/MAP1LC3A/ATP6V0D1/P4HB/PDLIM1 |
| Malignant epithelial | C5 Am/CEA-like ECs | GO:0051014 | actin filament severing | 3/171 | 17/18860 | 19.46 | 0.000454 | 0.0395 | 3 | SCIN/VILL/GSN |
| Malignant epithelial | C5 Am/CEA-like ECs | GO:1901142 | insulin metabolic process | 3/171 | 18/18860 | 18.38 | 0.000541 | 0.044 | 3 | CEACAM1/CPE/P4HB |
| Malignant epithelial | C5 Am/CEA-like ECs | GO:0043903 | regulation of biological process involved in symbiotic interaction | 4/171 | 42/18860 | 10.5 | 0.000558 | 0.044 | 4 | TMPRSS2/SMPD1/TMPRSS4/P4HB |
| Malignant epithelial | C5 Am/CEA-like ECs | GO:0051693 | actin filament capping | 4/171 | 42/18860 | 10.5 | 0.000558 | 0.044 | 4 | SPTBN1/SCIN/VILL/GSN |
| Malignant epithelial | C5 Am/CEA-like ECs | GO:1904375 | regulation of protein localization to cell periphery | 7/171 | 158/18860 | 4.89 | 0.000603 | 0.0453 | 7 | AGR2/EZR/TMBIM1/PLS1/SQSTM1/TMEM59/SPTBN1 |
| Malignant epithelial | C5 Am/CEA-like ECs | GO:0009124 | nucleoside monophosphate biosynthetic process | 4/171 | 43/18860 | 10.26 | 0.000611 | 0.0453 | 4 | ENTPD8/UPP1/RFK/CMPK1 |
| Malignant epithelial | C5 Am/CEA-like ECs | GO:1903900 | regulation of viral life cycle | 6/171 | 115/18860 | 5.75 | 0.000633 | 0.0457 | 6 | TMPRSS2/SMPD1/RSAD2/ISG20/TMPRSS4/P4HB |
| Malignant epithelial | C5 Am/CEA-like ECs | GO:0007043 | cell-cell junction assembly | 7/171 | 162/18860 | 4.77 | 0.000699 | 0.0483 | 7 | CDH17/RHOC/CLDN7/CLDN3/OCLN/CDH1/LSR |
| Malignant epithelial | C5 Am/CEA-like ECs | GO:0009123 | nucleoside monophosphate metabolic process | 5/171 | 78/18860 | 7.07 | 0.000715 | 0.0483 | 5 | GUK1/ENTPD8/UPP1/RFK/CMPK1 |

| compartment | subcluster | GO_ID | Description | GeneRatio | BgRatio | FoldEnrichment | pvalue | p_adjust | Count | geneID |
|---|---|---|---|---|---|---|---|---|---|---|
| Malignant epithelial | C5 Am/CEA-like ECs | GO:0050892 | intestinal absorption | 4/171 | 45/18860 | 9.8 | 0.000728 | 0.0483 | 4 | EZR/HEPH/PLS1/FABP2 |
| Malignant epithelial | C5 Am/CEA-like ECs | GO:0002221 | pattern recognition receptor signaling pathway | 10/171 | 325/18860 | 3.39 | 0.000786 | 0.0494 | 10 | CTSS/SQSTM1/RSAD2/TRIM31/RIOK3/RNF39/LGR4/S100A14/TAX1BP1/LRRC19 |
| Malignant epithelial | C5 Am/CEA-like ECs | GO:0007157 | heterophilic cell-cell adhesion via plasma membrane cell adhesion molecules | 4/171 | 46/18860 | 9.59 | 0.000791 | 0.0494 | 4 | CEACAM5/CEACAM6/CXADR/CD164 |
| Malignant epithelial | C5 Am/CEA-like ECs | GO:1904377 | positive regulation of protein localization to cell periphery | 5/171 | 80/18860 | 6.89 | 0.000803 | 0.0494 | 5 | AGR2/EZR/PLS1/SQSTM1/SPTBN1 |
| Malignant epithelial | C6 DNA-r p ECs | GO:0006260 | DNA replication | 33/179 | 290/18860 | 11.99 | 3.53e-26 | 7.669999999999999e-23 | 33 | FAM111B/CDK1/GINS2/TK1/DSCC1/CDT1/CDC6/CCNE2/CDC45/RFC3/RAD51/PCNA/FEN1/PRIM1/RFC2/RPA3/E… |
| Malignant epithelial | C6 DNA-r p ECs | GO:0007059 | chromosome segregation | 38/179 | 434/18860 | 9.23 | 6.680000000000001e-26 | 7.669999999999999e-23 | 38 | CENPU/CDK1/SPC25/CDT1/CENPK/NCAPH/MND1/CDC6/ZWINT/NUSAP1/CCNE2/SKA3/DIAPH3/MYBL2/CENPH/CE… |
| Malignant epithelial | C6 DNA-r p ECs | GO:0006261 | DNA-templated DNA replication | 23/179 | 171/18860 | 14.17 | 4.01e-20 | 3.07e-17 | 23 | CDK1/TK1/CDT1/CDC6/CCNE2/CDC45/RFC3/RAD51/PCNA/FEN1/PRIM1/RFC2/ORC6/BRCA2/MCM7/CDK2/MCM4/… |
| Malignant epithelial | C6 DNA-r p ECs | GO:0000819 | sister chromatid segregation | 25/179 | 233/18860 | 11.31 | 2.36e-19 | 1.36e-16 | 25 | CDK1/SPC25/CDT1/CENPK/NCAPH/CDC6/ZWINT/NUSAP1/SKA3/MYBL2/MAD2L1/TRIP13/SMC4/PRC1/BIRC5/UB… |
| Malignant epithelial | C6 DNA-r p ECs | GO:0098813 | nuclear chromosome segregation | 28/179 | 323/18860 | 9.13 | 4.51e-19 | 2.07e-16 | 28 | CDK1/SPC25/CDT1/CENPK/NCAPH/MND1/CDC6/ZWINT/NUSAP1/CCNE2/SKA3/MYBL2/MAD2L1/TRIP13/SMC4/PR… |
| Malignant epithelial | C6 DNA-r p ECs | GO:0000070 | mitotic sister chromatid segregation | 21/179 | 192/18860 | 11.52 | 1.41e-16 | 5.38e-14 | 21 | CDK1/SPC25/CDT1/CENPK/NCAPH/ZWINT/NUSAP1/SKA3/MYBL2/MAD2L1/TRIP13/SMC4/PRC1/BIRC5/UBE2C/N… |
| Malignant epithelial | C6 DNA-r p ECs | GO:0000280 | nuclear division | 29/179 | 453/18860 | 6.75 | 3.6e-16 | 1.18e-13 | 29 | CDK1/SPC25/CDT1/PKMYT1/CENPK/NCAPH/MND1/ZWINT/NUSAP1/CCNE2/SKA3/RAD51/MYBL2/BRCA2/MAD2L1/… |
| Malignant epithelial | C6 DNA-r p ECs | GO:0051983 | regulation of chromosome segregation | 18/179 | 134/18860 | 14.15 | 5.8e-16 | 1.67e-13 | 18 | CDK1/SPC25/NCAPH/CDC6/ZWINT/SKA3/MAD2L1/TRIP13/SMC4/BIRC5/UBE2C/NCAPG2/AURKB/NDC80/RMI2/S… |
| Malignant epithelial | C6 DNA-r p ECs | GO:0000075 | cell cycle checkpoint signaling | 20/179 | 187/18860 | 11.27 | 1.2e-15 | 3.06e-13 | 20 | CDK1/SPC25/CDT1/CDC6/ZWINT/CDC45/SKA3/CLSPN/RAD51/E2F1/BRCA2/MAD2L1/TRIP13/WDR76/BIRC5/CD… |
| Malignant epithelial | C6 DNA-r p ECs | GO:1905818 | regulation of chromosome separation | 14/179 | 76/18860 | 19.41 | 1.19e-14 | 2.72e-12 | 14 | SPC25/NCAPH/ZWINT/SKA3/MAD2L1/TRIP13/SMC4/BIRC5/UBE2C/NCAPG2/AURKB/NDC80/SMC2/CDCA8 |
| Malignant epithelial | C6 DNA-r p ECs | GO:0006270 | DNA replication initiation | 11/179 | 37/18860 | 31.32 | 2.85e-14 | 5.67e-12 | 11 | CDT1/CDC6/CCNE2/CDC45/PRIM1/ORC6/MCM7/CDK2/MCM4/POLA2/MCM10 |
| Malignant epithelial | C6 DNA-r p ECs | GO:0140014 | mitotic nuclear division | 22/179 | 281/18860 | 8.25 | 2.96e-14 | 5.67e-12 | 22 | CDK1/SPC25/CDT1/PKMYT1/CENPK/NCAPH/ZWINT/NUSAP1/SKA3/MYBL2/MAD2L1/TRIP13/SMC4/PRC1/BIRC5/… |
| Malignant epithelial | C6 DNA-r p ECs | GO:0044772 | mitotic cell cycle phase transition | 27/179 | 461/18860 | 6.17 | 3.62e-14 | 6.4e-12 | 27 | RRM2/MELK/CDK1/SPC25/PKMYT1/CDC6/ZWINT/CCNE2/SKA3/CLSPN/E2F1/EIF4EBP1/MAD2L1/TRIP13/BIRC5… |
| Malignant epithelial | C6 DNA-r p ECs | GO:0051304 | chromosome separation | 14/179 | 83/18860 | 17.77 | 4.31e-14 | 7.07e-12 | 14 | SPC25/NCAPH/ZWINT/SKA3/MAD2L1/TRIP13/SMC4/BIRC5/UBE2C/NCAPG2/AURKB/NDC80/SMC2/CDCA8 |

| compartment | subcluster | GO_ID | Description | GeneRatio | BgRatio | FoldEnrichment | pvalue | p_adjust | Count | geneID |
|---|---|---|---|---|---|---|---|---|---|---|
| Malignant epithelial | C6 DNA-r p ECs | GO:1901987 | regulation of cell cycle phase transition | 27/179 | 468/18860 | 6.08 | 5.2e-14 | 7.97e-12 | 27 | RRM2/CDK1/SPC25/CDT1/PKMYT1/CDC6/ZWINT/CCNE2/CDC45/SKA3/CLSPN/RAD51/E2F1/BRCA2/MAD2L1/TRI… |
| Malignant epithelial | C6 DNA-r p ECs | GO:0033260 | nuclear DNA replication | 11/179 | 41/18860 | 28.27 | 1.02e-13 | 1.46e-11 | 11 | CDK1/TK1/CDT1/CDC45/RAD51/PCNA/FEN1/BRCA2/MCM4/GINS1/DONSON |
| Malignant epithelial | C6 DNA-r p ECs | GO:0051984 | positive regulation of chromosome segregation | 10/179 | 30/18860 | 35.12 | 1.17e-13 | 1.59e-11 | 10 | CDK1/NCAPH/CDC6/SKA3/SMC4/BIRC5/NCAPG2/AURKB/SMC2/CDCA8 |
| Malignant epithelial | C6 DNA-r p ECs | GO:1901988 | negative regulation of cell cycle phase transition | 21/179 | 274/18860 | 8.08 | 1.8e-13 | 2.29e-11 | 21 | CDK1/SPC25/CDT1/PKMYT1/CDC6/ZWINT/CDC45/SKA3/CLSPN/RAD51/E2F1/BRCA2/MAD2L1/TRIP13/WDR76/B… |
| Malignant epithelial | C6 DNA-r p ECs | GO:0033044 | regulation of chromosome organization | 20/179 | 244/18860 | 8.64 | 1.99e-13 | 2.4e-11 | 20 | CDK1/SPC25/NCAPH/CDC6/ZWINT/SKA3/FEN1/MAD2L1/TRIP13/SMC4/BIRC5/UBE2C/CDK2/NCAPG2/AURKB/ND… |
| Malignant epithelial | C6 DNA-r p ECs | GO:0044786 | cell cycle DNA replication | 11/179 | 46/18860 | 25.2 | 4.13e-13 | 4.75e-11 | 11 | CDK1/TK1/CDT1/CDC45/RAD51/PCNA/FEN1/BRCA2/MCM4/GINS1/DONSON |
| Malignant epithelial | C6 DNA-r p ECs | GO:0033045 | regulation of sister chromatid segregation | 14/179 | 107/18860 | 13.79 | 1.63e-12 | 1.78e-10 | 14 | CDK1/SPC25/CDC6/ZWINT/SKA3/MAD2L1/TRIP13/BIRC5/UBE2C/AURKB/NDC80/RMI2/CDCA8/TACC3 |
| Malignant epithelial | C6 DNA-r p ECs | GO:0010948 | negative regulation of cell cycle process | 21/179 | 318/18860 | 6.96 | 3.21e-12 | 3.35e-10 | 21 | CDK1/SPC25/CDT1/PKMYT1/CDC6/ZWINT/CDC45/SKA3/CLSPN/RAD51/E2F1/BRCA2/MAD2L1/TRIP13/WDR76/B… |
| Malignant epithelial | C6 DNA-r p ECs | GO:1901990 | regulation of mitotic cell cycle phase transition | 22/179 | 357/18860 | 6.49 | 3.73e-12 | 3.73e-10 | 22 | RRM2/CDK1/SPC25/PKMYT1/CDC6/ZWINT/CCNE2/SKA3/CLSPN/E2F1/MAD2L1/TRIP13/BIRC5/UBE2C/CDK2/AU… |
| Malignant epithelial | C6 DNA-r p ECs | GO:1905820 | positive regulation of chromosome separation | 9/179 | 33/18860 | 28.74 | 1.62e-11 | 1.55e-09 | 9 | NCAPH/SKA3/SMC4/BIRC5/UBE2C/NCAPG2/AURKB/SMC2/CDCA8 |
| Malignant epithelial | C6 DNA-r p ECs | GO:0045787 | positive regulation of cell cycle | 21/179 | 349/18860 | 6.34 | 1.87e-11 | 1.72e-09 | 21 | RRM2/CDK1/NCAPH/CDC6/NUSAP1/CCNE2/SKA3/FEN1/EIF4EBP1/BRCA2/MAD2L1/SMC4/BIRC5/UBE2C/NCAPG2… |
| Malignant epithelial | C6 DNA-r p ECs | GO:0090068 | positive regulation of cell cycle process | 19/179 | 278/18860 | 7.2 | 2.03e-11 | 1.79e-09 | 19 | RRM2/CDK1/NCAPH/CDC6/NUSAP1/CCNE2/SKA3/FEN1/MAD2L1/SMC4/BIRC5/UBE2C/NCAPG2/AURKB/NDC80/SM… |
| Malignant epithelial | C6 DNA-r p ECs | GO:1901976 | regulation of cell cycle checkpoint | 10/179 | 48/18860 | 21.95 | 2.21e-11 | 1.88e-09 | 10 | ZWINT/SKA3/RAD51/BRCA2/MAD2L1/WDR76/BIRC5/AURKB/NDC80/CDCA8 |
| Malignant epithelial | C6 DNA-r p ECs | GO:0045786 | negative regulation of cell cycle | 22/179 | 399/18860 | 5.81 | 3.29e-11 | 2.7e-09 | 22 | CDK1/SPC25/CDT1/PKMYT1/CDC6/ZWINT/CDC45/SKA3/CLSPN/RAD51/E2F1/BRCA2/MAD2L1/TRIP13/WDR76/B… |
| Malignant epithelial | C6 DNA-r p ECs | GO:0007093 | mitotic cell cycle checkpoint signaling | 14/179 | 136/18860 | 10.85 | 4.48e-11 | 3.55e-09 | 14 | CDK1/SPC25/CDC6/ZWINT/SKA3/CLSPN/MAD2L1/TRIP13/BIRC5/CDK2/AURKB/NDC80/CDCA8/DONSON |
| Malignant epithelial | C6 DNA-r p ECs | GO:0006302 | double-strand break repair | 20/179 | 334/18860 | 6.31 | 6.35e-11 | 4.87e-09 | 20 | GINS2/CDCA5/CDC45/RAD51/HMGB2/FEN1/RPA3/ESCO2/BRCA2/FANCB/MCM7/TRIP13/HELLS/MCM4/RECQL4/R… |
| Malignant epithelial | C6 DNA-r p ECs | GO:1902850 | microtubule cytoskeleton organization involved in mitosis | 15/179 | 170/18860 | 9.3 | 8.17e-11 | 6.05e-09 | 15 | CDK1/SPC25/NUSAP1/MYBL2/CENPH/MAD2L1/PRC1/DYNLT1/BIRC5/AURKB/NDC80/CDCA8/TACC3/UHRF1/WDR62 |
| Malignant epithelial | C6 DNA-r p ECs | GO:1902969 | mitotic DNA replication | 7/179 | 17/18860 | 43.38 | 1.11e-10 | 7.95e-09 | 7 | CDK1/TK1/CDC45/RAD51/BRCA2/MCM4/GINS1 |
| Malignant epithelial | C6 DNA-r p ECs | GO:0007091 | metaphase/anaphase transition of mitotic cell cycle | 12/179 | 96/18860 | 13.17 | 1.15e-10 | 8.03e-09 | 12 | SPC25/CDC6/ZWINT/SKA3/MAD2L1/TRIP13/BIRC5/UBE2C/AURKB/NDC80/CDCA8/TACC3 |

| compartment | subcluster | GO_ID | Description | GeneRatio | BgRatio | FoldEnrichment | pvalue | p_adjust | Count | geneID |
|---|---|---|---|---|---|---|---|---|---|---|
| Malignant epithelial | C6 DNA-r p ECs | GO:0033047 | regulation of mitotic sister chromatid segregation | 10/179 | 57/18860 | 18.48 | 1.35e-10 | 9.15e-09 | 10 | CDK1/SPC25/ZWINT/SKA3/MAD2L1/TRIP13/BIRC5/AURKB/NDC80/CDCA8 |
| Malignant epithelial | C6 DNA-r p ECs | GO:0044784 | metaphase/anaphase transition of cell cycle | 12/179 | 99/18860 | 12.77 | 1.66e-10 | 1.09e-08 | 12 | SPC25/CDC6/ZWINT/SKA3/MAD2L1/TRIP13/BIRC5/UBE2C/AURKB/NDC80/CDCA8/TACC3 |
| Malignant epithelial | C6 DNA-r p ECs | GO:0006310 | DNA recombination | 20/179 | 358/18860 | 5.89 | 2.18e-10 | 1.39e-08 | 20 | GINS2/MND1/CDC45/RAD51/HMGB2/FEN1/RPA3/BRCA2/FANCB/MCM7/TRIP13/HELLS/MCM4/RECQL4/RMI2/TOP… |
| Malignant epithelial | C6 DNA-r p ECs | GO:0010965 | regulation of mitotic sister chromatid separation | 10/179 | 61/18860 | 17.27 | 2.73e-10 | 1.7e-08 | 10 | SPC25/ZWINT/SKA3/MAD2L1/TRIP13/BIRC5/UBE2C/AURKB/NDC80/CDCA8 |
| Malignant epithelial | C6 DNA-r p ECs | GO:0051306 | mitotic sister chromatid separation | 10/179 | 64/18860 | 16.46 | 4.48e-10 | 2.71e-08 | 10 | SPC25/ZWINT/SKA3/MAD2L1/TRIP13/BIRC5/UBE2C/AURKB/NDC80/CDCA8 |
| Malignant epithelial | C6 DNA-r p ECs | GO:1901991 | negative regulation of mitotic cell cycle phase transition | 15/179 | 193/18860 | 8.19 | 4.9e-10 | 2.89e-08 | 15 | CDK1/SPC25/PKMYT1/CDC6/ZWINT/SKA3/CLSPN/MAD2L1/TRIP13/BIRC5/CDK2/AURKB/NDC80/CDCA8/DONSON |
| Malignant epithelial | C6 DNA-r p ECs | GO:0000724 | double-strand break repair via homologous recombination | 15/179 | 195/18860 | 8.1 | 5.66e-10 | 3.25e-08 | 15 | GINS2/CDC45/RAD51/FEN1/RPA3/BRCA2/FANCB/MCM7/HELLS/MCM4/RECQL4/RMI2/UHRF1/RAD51AP1/RAD51C |
| Malignant epithelial | C6 DNA-r p ECs | GO:0007094 | mitotic spindle assembly checkpoint signaling | 9/179 | 48/18860 | 19.76 | 6.23e-10 | 3.33e-08 | 9 | SPC25/ZWINT/SKA3/MAD2L1/TRIP13/BIRC5/AURKB/NDC80/CDCA8 |
| Malignant epithelial | C6 DNA-r p ECs | GO:0071173 | spindle assembly checkpoint signaling | 9/179 | 48/18860 | 19.76 | 6.23e-10 | 3.33e-08 | 9 | SPC25/ZWINT/SKA3/MAD2L1/TRIP13/BIRC5/AURKB/NDC80/CDCA8 |
| Malignant epithelial | C6 DNA-r p ECs | GO:0071174 | mitotic spindle checkpoint signaling | 9/179 | 48/18860 | 19.76 | 6.23e-10 | 3.33e-08 | 9 | SPC25/ZWINT/SKA3/MAD2L1/TRIP13/BIRC5/AURKB/NDC80/CDCA8 |
| Malignant epithelial | C6 DNA-r p ECs | GO:0031577 | spindle checkpoint signaling | 9/179 | 49/18860 | 19.35 | 7.57e-10 | 3.95e-08 | 9 | SPC25/ZWINT/SKA3/MAD2L1/TRIP13/BIRC5/AURKB/NDC80/CDCA8 |
| Malignant epithelial | C6 DNA-r p ECs | GO:0000725 | recombinational repair | 15/179 | 201/18860 | 7.86 | 8.64e-10 | 4.29e-08 | 15 | GINS2/CDC45/RAD51/FEN1/RPA3/BRCA2/FANCB/MCM7/HELLS/MCM4/RECQL4/RMI2/UHRF1/RAD51AP1/RAD51C |
| Malignant epithelial | C6 DNA-r p ECs | GO:0033046 | negative regulation of sister chromatid segregation | 9/179 | 50/18860 | 18.97 | 9.16e-10 | 4.29e-08 | 9 | SPC25/ZWINT/SKA3/MAD2L1/TRIP13/BIRC5/AURKB/NDC80/CDCA8 |
| Malignant epithelial | C6 DNA-r p ECs | GO:0033048 | negative regulation of mitotic sister chromatid segregation | 9/179 | 50/18860 | 18.97 | 9.16e-10 | 4.29e-08 | 9 | SPC25/ZWINT/SKA3/MAD2L1/TRIP13/BIRC5/AURKB/NDC80/CDCA8 |
| Malignant epithelial | C6 DNA-r p ECs | GO:0045841 | negative regulation of mitotic metaphase/anaphase transition | 9/179 | 50/18860 | 18.97 | 9.16e-10 | 4.29e-08 | 9 | SPC25/ZWINT/SKA3/MAD2L1/TRIP13/BIRC5/AURKB/NDC80/CDCA8 |
| Malignant epithelial | C6 DNA-r p ECs | GO:2000816 | negative regulation of mitotic sister chromatid separation | 9/179 | 50/18860 | 18.97 | 9.16e-10 | 4.29e-08 | 9 | SPC25/ZWINT/SKA3/MAD2L1/TRIP13/BIRC5/AURKB/NDC80/CDCA8 |
| Malignant epithelial | C6 DNA-r p ECs | GO:0030071 | regulation of mitotic metaphase/anaphase transition | 11/179 | 92/18860 | 12.6 | 1.14e-09 | 5.25e-08 | 11 | SPC25/CDC6/ZWINT/SKA3/MAD2L1/TRIP13/BIRC5/UBE2C/AURKB/NDC80/CDCA8 |
| Malignant epithelial | C6 DNA-r p ECs | GO:1902100 | negative regulation of metaphase/anaphase transition of cell cycle | 9/179 | 52/18860 | 18.24 | 1.32e-09 | 5.85e-08 | 9 | SPC25/ZWINT/SKA3/MAD2L1/TRIP13/BIRC5/AURKB/NDC80/CDCA8 |
| Malignant epithelial | C6 DNA-r p ECs | GO:1905819 | negative regulation of chromosome separation | 9/179 | 52/18860 | 18.24 | 1.32e-09 | 5.85e-08 | 9 | SPC25/ZWINT/SKA3/MAD2L1/TRIP13/BIRC5/AURKB/NDC80/CDCA8 |
| Malignant epithelial | C6 DNA-r p ECs | GO:0007088 | regulation of mitotic nuclear division | 12/179 | 119/18860 | 10.62 | 1.45e-09 | 6.28e-08 | 12 | SPC25/PKMYT1/ZWINT/NUSAP1/SKA3/MAD2L1/TRIP13/BIRC5/UBE2C/AURKB/NDC80/CDCA8 |
| Malignant epithelial | C6 DNA-r p ECs | GO:0051783 | regulation of nuclear division | 13/179 | 147/18860 | 9.32 | 1.51e-09 | 6.42e-08 | 13 | SPC25/PKMYT1/ZWINT/NUSAP1/SKA3/MAD2L1/TRIP13/BIRC5/UBE2C/AURKB/NDC80/CDCA8/RAD51AP1 |
| Malignant epithelial | C6 DNA-r p ECs | GO:0051985 | negative regulation of chromosome segregation | 9/179 | 53/18860 | 17.89 | 1.58e-09 | 6.6e-08 | 9 | SPC25/ZWINT/SKA3/MAD2L1/TRIP13/BIRC5/AURKB/NDC80/CDCA8 |

| compartment | subcluster | GO_ID | Description | GeneRatio | BgRatio | FoldEnrichment | pvalue | p_adjust | Count | geneID |
|---|---|---|---|---|---|---|---|---|---|---|
| Malignant epithelial | C6 DNA-r p ECs | GO:1902099 | regulation of metaphase/anaphase transition of cell cycle | 11/179 | 95/18860 | 12.2 | 1.62e-09 | 6.64e-08 | 11 | SPC25/CDC6/ZWINT/SKA3/MAD2L1/TRIP13/BIRC5/UBE2C/AURKB/NDC80/CDCA8 |
| Malignant epithelial | C6 DNA-r p ECs | GO:0090231 | regulation of spindle checkpoint | 7/179 | 24/18860 | 30.73 | 1.86e-09 | 7.51e-08 | 7 | ZWINT/SKA3/MAD2L1/BIRC5/AURKB/NDC80/CDCA8 |
| Malignant epithelial | C6 DNA-r p ECs | GO:0008608 | attachment of spindle microtubules to kinetochore | 9/179 | 55/18860 | 17.24 | 2.23e-09 | 8.84e-08 | 9 | CDK1/SPC25/CDT1/SKA3/BIRC5/AURKB/NDC80/DSN1/CDCA8 |
| Malignant epithelial | C6 DNA-r p ECs | GO:0045839 | negative regulation of mitotic nuclear division | 9/179 | 57/18860 | 16.64 | 3.11e-09 | 1.2e-07 | 9 | SPC25/ZWINT/SKA3/MAD2L1/TRIP13/BIRC5/AURKB/NDC80/CDCA8 |
| Malignant epithelial | C6 DNA-r p ECs | GO:0006271 | DNA strand elongation involved in DNA replication | 6/179 | 15/18860 | 42.15 | 3.13e-09 | 1.2e-07 | 6 | RFC3/PCNA/MCM7/MCM4/POLD2/GINS1 |
| Malignant epithelial | C6 DNA-r p ECs | GO:0051315 | attachment of mitotic spindle microtubules to kinetochore | 7/179 | 26/18860 | 28.37 | 3.48e-09 | 1.31e-07 | 7 | CDK1/CDT1/SKA3/BIRC5/AURKB/NDC80/CDCA8 |
| Malignant epithelial | C6 DNA-r p ECs | GO:0051321 | meiotic cell cycle | 17/179 | 299/18860 | 5.99 | 4.07e-09 | 1.51e-07 | 17 | PKMYT1/NCAPH/MND1/ZWINT/CCNE2/SKA3/RAD51/MNS1/BRCA2/TRIP13/SMC4/CDK2/NDC80/SMC2/TOP2A/RAD… |
| Malignant epithelial | C6 DNA-r p ECs | GO:1903046 | meiotic cell cycle process | 15/179 | 227/18860 | 6.96 | 4.61e-09 | 1.68e-07 | 15 | PKMYT1/NCAPH/MND1/ZWINT/CCNE2/SKA3/RAD51/BRCA2/TRIP13/SMC4/NDC80/SMC2/TOP2A/RAD51AP1/RAD5… |
| Malignant epithelial | C6 DNA-r p ECs | GO:0044839 | cell cycle G2/M phase transition | 13/179 | 162/18860 | 8.46 | 4.97e-09 | 1.78e-07 | 13 | MELK/CDK1/PKMYT1/CDC6/CLSPN/BIRC5/CALM1/CDK2/CCNA2/AURKB/NDC80/DONSON/RAD51C |
| Malignant epithelial | C6 DNA-r p ECs | GO:0140013 | meiotic nuclear division | 14/179 | 203/18860 | 7.27 | 8.89e-09 | 3.13e-07 | 14 | NCAPH/MND1/ZWINT/CCNE2/SKA3/RAD51/BRCA2/TRIP13/SMC4/NDC80/SMC2/TOP2A/RAD51AP1/RAD51C |
| Malignant epithelial | C6 DNA-r p ECs | GO:0051784 | negative regulation of nuclear division | 9/179 | 64/18860 | 14.82 | 8.98e-09 | 3.13e-07 | 9 | SPC25/ZWINT/SKA3/MAD2L1/TRIP13/BIRC5/AURKB/NDC80/CDCA8 |
| Malignant epithelial | C6 DNA-r p ECs | GO:0045930 | negative regulation of mitotic cell cycle | 15/179 | 244/18860 | 6.48 | 1.22e-08 | 4.2e-07 | 15 | CDK1/SPC25/PKMYT1/CDC6/ZWINT/SKA3/CLSPN/MAD2L1/TRIP13/BIRC5/CDK2/AURKB/NDC80/CDCA8/DONSON |
| Malignant epithelial | C6 DNA-r p ECs | GO:0000086 | G2/M transition of mitotic cell cycle | 12/179 | 145/18860 | 8.72 | 1.39e-08 | 4.71e-07 | 12 | MELK/CDK1/PKMYT1/CDC6/CLSPN/BIRC5/CALM1/CDK2/CCNA2/AURKB/DONSON/RAD51C |
| Malignant epithelial | C6 DNA-r p ECs | GO:0090267 | positive regulation of mitotic cell cycle spindle assembly checkpoint | 5/179 | 11/18860 | 47.89 | 3.21e-08 | 1.06e-06 | 5 | MAD2L1/BIRC5/AURKB/NDC80/CDCA8 |
| Malignant epithelial | C6 DNA-r p ECs | GO:0051383 | kinetochore organization | 6/179 | 21/18860 | 30.1 | 3.24e-08 | 1.06e-06 | 6 | CENPK/CENPH/SMC4/NDC80/SMC2/CENPW |
| Malignant epithelial | C6 DNA-r p ECs | GO:0031570 | DNA integrity checkpoint signaling | 11/179 | 126/18860 | 9.2 | 3.27e-08 | 1.06e-06 | 11 | CDK1/CDT1/CDC6/CDC45/CLSPN/RAD51/E2F1/BRCA2/WDR76/CDK2/DONSON |
| Malignant epithelial | C6 DNA-r p ECs | GO:0090232 | positive regulation of spindle checkpoint | 5/179 | 12/18860 | 43.9 | 5.46e-08 | 1.74e-06 | 5 | MAD2L1/BIRC5/AURKB/NDC80/CDCA8 |
| Malignant epithelial | C6 DNA-r p ECs | GO:0051310 | metaphase chromosome alignment | 10/179 | 104/18860 | 10.13 | 5.57e-08 | 1.75e-06 | 10 | CDK1/SPC25/CDT1/ZWINT/SKA3/BIRC5/AURKB/NDC80/DSN1/CDCA8 |
| Malignant epithelial | C6 DNA-r p ECs | GO:0090266 | regulation of mitotic cell cycle spindle assembly checkpoint | 6/179 | 23/18860 | 27.49 | 5.93e-08 | 1.82e-06 | 6 | SKA3/MAD2L1/BIRC5/AURKB/NDC80/CDCA8 |
| Malignant epithelial | C6 DNA-r p ECs | GO:1903504 | regulation of mitotic spindle checkpoint | 6/179 | 23/18860 | 27.49 | 5.93e-08 | 1.82e-06 | 6 | SKA3/MAD2L1/BIRC5/AURKB/NDC80/CDCA8 |
| Malignant epithelial | C6 DNA-r p ECs | GO:0007052 | mitotic spindle organization | 11/179 | 137/18860 | 8.46 | 7.78e-08 | 2.35e-06 | 11 | SPC25/MYBL2/CENPH/PRC1/BIRC5/AURKB/NDC80/CDCA8/TACC3/UHRF1/WDR62 |
| Malignant epithelial | C6 DNA-r p ECs | GO:0065004 | protein-DNA complex assembly | 14/179 | 245/18860 | 6.02 | 9.49e-08 | 2.83e-06 | 14 | ASF1B/CDT1/CENPK/CDC45/RAD51/ATAD2/HMGB2/CENPH/CENPP/CENPN/CENPW/OIP5/HMGB1/RAD51C |
| Malignant epithelial | C6 DNA-r p ECs | GO:0007051 | spindle organization | 13/179 | 209/18860 | 6.55 | 1.04e-07 | 3.06e-06 | 13 | SPC25/SKA3/MYBL2/CENPH/PRC1/BIRC5/AURKB/NDC80/CDCA8/TACC3/UHRF1/TUBB/WDR62 |
| Malignant epithelial | C6 DNA-r p ECs | GO:0061982 | meiosis I cell cycle process | 11/179 | 142/18860 | 8.16 | 1.12e-07 | 3.27e-06 | 11 | PKMYT1/MND1/ZWINT/CCNE2/RAD51/BRCA2/TRIP13/NDC80/TOP2A/RAD51AP1/RAD51C |

| compartment | subcluster | GO_ID | Description | GeneRatio | BgRatio | FoldEnrichment | pvalue | p_adjust | Count | geneID |
|---|---|---|---|---|---|---|---|---|---|---|
| Malignant epithelial | C6 DNA-r p ECs | GO:0006275 | regulation of DNA replication | 10/179 | 120/18860 | 8.78 | 2.17e-07 | 6.24e-06 | 10 | CDK1/DSCC1/CDT1/CDC6/PCNA/ESCO2/MCM7/CDK2/MCM4/CCNA2 |
| Malignant epithelial | C6 DNA-r p ECs | GO:0030261 | chromosome condensation | 7/179 | 46/18860 | 16.03 | 2.42e-07 | 6.85e-06 | 7 | CDK1/NCAPH/NUSAP1/SMC4/NCAPG2/SMC2/TOP2A |
| Malignant epithelial | C6 DNA-r p ECs | GO:0071824 | protein-DNA complex organization | 14/179 | 265/18860 | 5.57 | 2.49e-07 | 6.97e-06 | 14 | ASF1B/CDT1/CENPK/CDC45/RAD51/ATAD2/HMGB2/CENPH/CENPP/CENPN/CENPW/OIP5/HMGB1/RAD51C |
| Malignant epithelial | C6 DNA-r p ECs | GO:0009263 | deoxyribonucleotide biosynthetic process | 5/179 | 16/18860 | 32.93 | 2.92e-07 | 7.9e-06 | 5 | RRM2/TYMS/SHMT1/DTYMK/DUT |
| Malignant epithelial | C6 DNA-r p ECs | GO:0009265 | 2'-deoxyribonucleotide biosynthetic process | 5/179 | 16/18860 | 32.93 | 2.92e-07 | 7.9e-06 | 5 | RRM2/TYMS/SHMT1/DTYMK/DUT |
| Malignant epithelial | C6 DNA-r p ECs | GO:0046385 | deoxyribose phosphate biosynthetic process | 5/179 | 16/18860 | 32.93 | 2.92e-07 | 7.9e-06 | 5 | RRM2/TYMS/SHMT1/DTYMK/DUT |
| Malignant epithelial | C6 DNA-r p ECs | GO:0050000 | chromosome localization | 10/179 | 124/18860 | 8.5 | 2.96e-07 | 7.9e-06 | 10 | CDK1/SPC25/CDT1/ZWINT/SKA3/BIRC5/AURKB/NDC80/DSN1/CDCA8 |
| Malignant epithelial | C6 DNA-r p ECs | GO:0045132 | meiotic chromosome segregation | 9/179 | 97/18860 | 9.78 | 3.56e-07 | 9.31e-06 | 9 | NCAPH/MND1/ZWINT/CCNE2/SKA3/TRIP13/SMC4/NDC80/SMC2 |
| Malignant epithelial | C6 DNA-r p ECs | GO:2001251 | negative regulation of chromosome organization | 9/179 | 97/18860 | 9.78 | 3.56e-07 | 9.31e-06 | 9 | SPC25/ZWINT/SKA3/MAD2L1/TRIP13/BIRC5/AURKB/NDC80/CDCA8 |
| Malignant epithelial | C6 DNA-r p ECs | GO:0034508 | centromere complex assembly | 6/179 | 31/18860 | 20.39 | 4.06e-07 | 1.05e-05 | 6 | CENPK/CENPH/CENPP/CENPN/CENPW/OIP5 |
| Malignant epithelial | C6 DNA-r p ECs | GO:0007127 | meiosis I | 10/179 | 132/18860 | 7.98 | 5.3e-07 | 1.35e-05 | 10 | MND1/ZWINT/CCNE2/RAD51/BRCA2/TRIP13/NDC80/TOP2A/RAD51AP1/RAD51C |
| Malignant epithelial | C6 DNA-r p ECs | GO:1901978 | positive regulation of cell cycle checkpoint | 5/179 | 18/18860 | 29.27 | 5.64e-07 | 1.43e-05 | 5 | MAD2L1/BIRC5/AURKB/NDC80/CDCA8 |
| Malignant epithelial | C6 DNA-r p ECs | GO:2001252 | positive regulation of chromosome organization | 9/179 | 105/18860 | 9.03 | 7.02e-07 | 1.75e-05 | 9 | CDK1/NCAPH/SKA3/FEN1/SMC4/CDK2/NCAPG2/AURKB/SMC2 |
| Malignant epithelial | C6 DNA-r p ECs | GO:0045931 | positive regulation of mitotic cell cycle | 10/179 | 138/18860 | 7.64 | 7.99e-07 | 1.98e-05 | 10 | RRM2/CDK1/CDC6/CCNE2/SKA3/EIF4EBP1/BRCA2/BIRC5/UBE2C/RAD51C |
| Malignant epithelial | C6 DNA-r p ECs | GO:0000076 | DNA replication checkpoint signaling | 5/179 | 20/18860 | 26.34 | 1.01e-06 | 2.46e-05 | 5 | CDT1/CDC6/CDC45/CLSPN/DONSON |
| Malignant epithelial | C6 DNA-r p ECs | GO:0022616 | DNA strand elongation | 6/179 | 37/18860 | 17.09 | 1.22e-06 | 2.96e-05 | 6 | RFC3/PCNA/MCM7/MCM4/POLD2/GINS1 |
| Malignant epithelial | C6 DNA-r p ECs | GO:0051052 | regulation of DNA metabolic process | 18/179 | 497/18860 | 3.82 | 1.27e-06 | 3.01e-05 | 18 | CDK1/DSCC1/CDT1/CDC6/RFC3/RAD51/PCNA/RFC2/ESCO2/FANCB/MCM7/CDK2/MCM4/CCNA2/AURKB/RMI2/HMG… |
| Malignant epithelial | C6 DNA-r p ECs | GO:0140694 | membraneless organelle assembly | 17/179 | 446/18860 | 4.02 | 1.27e-06 | 3.01e-05 | 17 | CENPK/SKA3/DIAPH3/MYBL2/CENPH/LSM4/PRC1/BIRC5/CDK2/AURKB/NDC80/EIF6/CENPW/CDCA8/UHRF1/TUB… |
| Malignant epithelial | C6 DNA-r p ECs | GO:1902423 | regulation of attachment of mitotic spindle microtubules to kinetochore | 4/179 | 10/18860 | 42.15 | 1.58e-06 | 3.7e-05 | 4 | CDK1/BIRC5/AURKB/CDCA8 |
| Malignant epithelial | C6 DNA-r p ECs | GO:1901970 | positive regulation of mitotic sister chromatid separation | 5/179 | 22/18860 | 23.95 | 1.68e-06 | 3.91e-05 | 5 | SKA3/BIRC5/UBE2C/AURKB/CDCA8 |
| Malignant epithelial | C6 DNA-r p ECs | GO:0007080 | mitotic metaphase chromosome alignment | 7/179 | 63/18860 | 11.71 | 2.18e-06 | 5.01e-05 | 7 | CDK1/CDT1/SKA3/BIRC5/AURKB/NDC80/CDCA8 |
| Malignant epithelial | C6 DNA-r p ECs | GO:0031055 | chromatin remodeling at centromere | 4/179 | 11/18860 | 38.31 | 2.46e-06 | 5.48e-05 | 4 | CENPP/CENPN/CENPW/OIP5 |
| Malignant epithelial | C6 DNA-r p ECs | GO:0034080 | CENP-A containing chromatin assembly | 4/179 | 11/18860 | 38.31 | 2.46e-06 | 5.48e-05 | 4 | CENPP/CENPN/CENPW/OIP5 |
| Malignant epithelial | C6 DNA-r p ECs | GO:0051256 | mitotic spindle midzone assembly | 4/179 | 11/18860 | 38.31 | 2.46e-06 | 5.48e-05 | 4 | PRC1/BIRC5/AURKB/CDCA8 |
| Malignant epithelial | C6 DNA-r p ECs | GO:0070192 | chromosome organization involved in meiotic cell cycle | 7/179 | 66/18860 | 11.17 | 3e-06 | 6.62e-05 | 7 | NCAPH/MND1/CCNE2/RAD51/TRIP13/SMC4/SMC2 |
| Malignant epithelial | C6 DNA-r p ECs | GO:0000022 | mitotic spindle elongation | 4/179 | 12/18860 | 35.12 | 3.66e-06 | 7.88e-05 | 4 | PRC1/BIRC5/AURKB/CDCA8 |
| Malignant epithelial | C6 DNA-r p ECs | GO:0000727 | double-strand break repair via break-induced replication | 4/179 | 12/18860 | 35.12 | 3.66e-06 | 7.88e-05 | 4 | GINS2/CDC45/MCM7/MCM4 |
| Malignant epithelial | C6 DNA-r p ECs | GO:0035825 | homologous recombination | 7/179 | 68/18860 | 10.85 | 3.67e-06 | 7.88e-05 | 7 | MND1/RAD51/TRIP13/TOP2A/UHRF1/RAD51AP1/RAD51C |
| Malignant epithelial | C6 DNA-r p ECs | GO:0000723 | telomere maintenance | 10/179 | 166/18860 | 6.35 | 4.27e-06 | 9.1e-05 | 10 | CCNE2/RAD51/PCNA/FEN1/RPA3/BRCA2/CDK2/AURKB/RECQL4/RAD51C |

| compartment | subcluster | GO_ID | Description | GeneRatio | BgRatio | FoldEnrichment | pvalue | p_adjust | Count | geneID |
|---|---|---|---|---|---|---|---|---|---|---|
| Malignant epithelial | C6 DNA-r p ECs | GO:0072528 | pyrimidine-containing compound biosynthetic process | 6/179 | 46/18860 | 13.74 | 4.59e-06 | 9.69e-05 | 6 | CDK1/TK1/TYMS/SHMT1/DTYMK/DUT |
| Malignant epithelial | C6 DNA-r p ECs | GO:0071897 | DNA biosynthetic process | 10/179 | 168/18860 | 6.27 | 4.76e-06 | 9.94e-05 | 10 | CDK1/TK1/DSCC1/RFC3/PCNA/RFC2/TYMS/CCNA2/POLD1/POLD2 |
| Malignant epithelial | C6 DNA-r p ECs | GO:0051225 | spindle assembly | 9/179 | 137/18860 | 6.92 | 6.46e-06 | 0.000134 | 9 | SKA3/MYBL2/PRC1/BIRC5/AURKB/NDC80/CDCA8/UHRF1/TUBB |
| Malignant epithelial | C6 DNA-r p ECs | GO:1901992 | positive regulation of mitotic cell cycle phase transition | 8/179 | 105/18860 | 8.03 | 7.17e-06 | 0.000146 | 8 | RRM2/CDK1/CDC6/CCNE2/SKA3/BIRC5/UBE2C/RAD51C |
| Malignant epithelial | C6 DNA-r p ECs | GO:0051231 | spindle elongation | 4/179 | 14/18860 | 30.1 | 7.29e-06 | 0.000146 | 4 | PRC1/BIRC5/AURKB/CDCA8 |
| Malignant epithelial | C6 DNA-r p ECs | GO:0051255 | spindle midzone assembly | 4/179 | 14/18860 | 30.1 | 7.29e-06 | 0.000146 | 4 | PRC1/BIRC5/AURKB/CDCA8 |
| Malignant epithelial | C6 DNA-r p ECs | GO:0060623 | regulation of chromosome condensation | 4/179 | 14/18860 | 30.1 | 7.29e-06 | 0.000146 | 4 | NCAPH/SMC4/NCAPG2/SMC2 |
| Malignant epithelial | C6 DNA-r p ECs | GO:0010389 | regulation of G2/M transition of mitotic cell cycle | 8/179 | 109/18860 | 7.73 | 9.46e-06 | 0.000187 | 8 | CDK1/PKMYT1/CDC6/CLSPN/CDK2/AURKB/DONSON/RAD51C |
| Malignant epithelial | C6 DNA-r p ECs | GO:0090235 | regulation of metaphase plate congression | 4/179 | 15/18860 | 28.1 | 9.87e-06 | 0.000194 | 4 | CDK1/BIRC5/AURKB/CDCA8 |
| Malignant epithelial | C6 DNA-r p ECs | GO:0030174 | regulation of DNA-templated DNA replication initiation | 4/179 | 16/18860 | 26.34 | 1.31e-05 | 0.000254 | 4 | CDT1/MCM7/CDK2/MCM4 |
| Malignant epithelial | C6 DNA-r p ECs | GO:0000077 | DNA damage checkpoint signaling | 8/179 | 117/18860 | 7.2 | 1.59e-05 | 0.000307 | 8 | CDK1/CLSPN/RAD51/E2F1/BRCA2/WDR76/CDK2/DONSON |
| Malignant epithelial | C6 DNA-r p ECs | GO:0032200 | telomere organization | 10/179 | 193/18860 | 5.46 | 1.61e-05 | 0.000308 | 10 | CCNE2/RAD51/PCNA/FEN1/RPA3/BRCA2/CDK2/AURKB/RECQL4/RAD51C |
| Malignant epithelial | C6 DNA-r p ECs | GO:0051054 | positive regulation of DNA metabolic process | 12/179 | 283/18860 | 4.47 | 1.73e-05 | 0.000329 | 12 | CDK1/DSCC1/CDT1/RFC3/PCNA/RFC2/FANCB/CDK2/CCNA2/AURKB/HMGB1/RAD51AP1 |
| Malignant epithelial | C6 DNA-r p ECs | GO:1902749 | regulation of cell cycle G2/M phase transition | 8/179 | 122/18860 | 6.91 | 2.16e-05 | 0.000404 | 8 | CDK1/PKMYT1/CDC6/CLSPN/CDK2/AURKB/DONSON/RAD51C |
| Malignant epithelial | C6 DNA-r p ECs | GO:0007076 | mitotic chromosome condensation | 4/179 | 18/18860 | 23.41 | 2.16e-05 | 0.000404 | 4 | NCAPH/NUSAP1/SMC4/SMC2 |
| Malignant epithelial | C6 DNA-r p ECs | GO:0007131 | reciprocal meiotic recombination | 6/179 | 60/18860 | 10.54 | 2.2e-05 | 0.000404 | 6 | MND1/RAD51/TRIP13/TOP2A/RAD51AP1/RAD51C |
| Malignant epithelial | C6 DNA-r p ECs | GO:0140527 | reciprocal homologous recombination | 6/179 | 60/18860 | 10.54 | 2.2e-05 | 0.000404 | 6 | MND1/RAD51/TRIP13/TOP2A/RAD51AP1/RAD51C |
| Malignant epithelial | C6 DNA-r p ECs | GO:1901989 | positive regulation of cell cycle phase transition | 8/179 | 126/18860 | 6.69 | 2.73e-05 | 0.000497 | 8 | RRM2/CDK1/CDC6/CCNE2/SKA3/BIRC5/UBE2C/RAD51C |
| Malignant epithelial | C6 DNA-r p ECs | GO:0032465 | regulation of cytokinesis | 7/179 | 96/18860 | 7.68 | 3.61e-05 | 0.000654 | 7 | CDC6/BRCA2/PRC1/BIRC5/CALM1/AURKB/CDCA8 |
| Malignant epithelial | C6 DNA-r p ECs | GO:0045005 | DNA-templated DNA replication maintenance of fidelity | 6/179 | 66/18860 | 9.58 | 3.81e-05 | 0.000684 | 6 | CDK1/RAD51/PCNA/BRCA2/POLD1/DONSON |
| Malignant epithelial | C6 DNA-r p ECs | GO:0007143 | female meiotic nuclear division | 5/179 | 43/18860 | 12.25 | 5.23e-05 | 0.000932 | 5 | NCAPH/SKA3/TRIP13/NDC80/TOP2A |
| Malignant epithelial | C6 DNA-r p ECs | GO:0051988 | regulation of attachment of spindle microtubules to kinetochore | 4/179 | 23/18860 | 18.32 | 6.03e-05 | 0.00107 | 4 | CDK1/BIRC5/AURKB/CDCA8 |
| Malignant epithelial | C6 DNA-r p ECs | GO:0009394 | 2'-deoxyribonucleotide metabolic process | 5/179 | 45/18860 | 11.71 | 6.54e-05 | 0.00114 | 5 | RRM2/TYMS/SHMT1/DTYMK/DUT |
| Malignant epithelial | C6 DNA-r p ECs | GO:2000573 | positive regulation of DNA biosynthetic process | 5/179 | 45/18860 | 11.71 | 6.54e-05 | 0.00114 | 5 | DSCC1/RFC3/PCNA/RFC2/CCNA2 |
| Malignant epithelial | C6 DNA-r p ECs | GO:0010972 | negative regulation of G2/M transition of mitotic cell cycle | 6/179 | 73/18860 | 8.66 | 6.76e-05 | 0.00117 | 6 | CDK1/PKMYT1/CDC6/CLSPN/AURKB/DONSON |
| Malignant epithelial | C6 DNA-r p ECs | GO:0009262 | deoxyribonucleotide metabolic process | 5/179 | 46/18860 | 11.45 | 7.28e-05 | 0.00124 | 5 | RRM2/TYMS/SHMT1/DTYMK/DUT |
| Malignant epithelial | C6 DNA-r p ECs | GO:0072527 | pyrimidine-containing compound metabolic process | 6/179 | 74/18860 | 8.54 | 7.29e-05 | 0.00124 | 6 | CDK1/TK1/TYMS/SHMT1/DTYMK/DUT |
| Malignant epithelial | C6 DNA-r p ECs | GO:1902750 | negative regulation of cell cycle G2/M phase transition | 6/179 | 75/18860 | 8.43 | 7.87e-05 | 0.00133 | 6 | CDK1/PKMYT1/CDC6/CLSPN/AURKB/DONSON |
| Malignant epithelial | C6 DNA-r p ECs | GO:0090307 | mitotic spindle assembly | 6/179 | 76/18860 | 8.32 | 8.47e-05 | 0.00141 | 6 | MYBL2/PRC1/BIRC5/AURKB/CDCA8/UHRF1 |
| Malignant epithelial | C6 DNA-r p ECs | GO:0009219 | pyrimidine deoxyribonucleotide metabolic process | 4/179 | 25/18860 | 16.86 | 8.49e-05 | 0.00141 | 4 | TYMS/SHMT1/DTYMK/DUT |
| Malignant epithelial | C6 DNA-r p ECs | GO:0006265 | DNA topological change | 3/179 | 10/18860 | 31.61 | 9.61e-05 | 0.00159 | 3 | HMGB2/TOP2A/HMGB1 |

| compartment | subcluster | GO_ID | Description | GeneRatio | BgRatio | FoldEnrichment | pvalue | p_adjust | Count | geneID |
|---|---|---|---|---|---|---|---|---|---|---|
| Malignant epithelial | C6 DNA-r p ECs | GO:0000910 | cytokinesis | 9/179 | 195/18860 | 4.86 | 0.000105 | 0.00172 | 9 | CDC6/NUSAP1/BRCA2/PRC1/BIRC5/CALM1/AURKB/CDCA8/CKAP2 |
| Malignant epithelial | C6 DNA-r p ECs | GO:0000082 | G1/S transition of mitotic cell cycle | 10/179 | 246/18860 | 4.28 | 0.000125 | 0.00203 | 10 | RRM2/CDK1/CDC6/CCNE2/E2F1/EIF4EBP1/CDK2/CCNA2/CDKN2A/PSME2 |
| Malignant epithelial | C6 DNA-r p ECs | GO:0042770 | signal transduction in response to DNA damage | 9/179 | 200/18860 | 4.74 | 0.000127 | 0.00206 | 9 | CDK1/CLSPN/RAD51/E2F1/BRCA2/WDR76/CDK2/CDKN2A/DONSON |
| Malignant epithelial | C6 DNA-r p ECs | GO:0033314 | mitotic DNA replication checkpoint signaling | 3/179 | 11/18860 | 28.74 | 0.000131 | 0.00209 | 3 | CDC6/CLSPN/DONSON |
| Malignant epithelial | C6 DNA-r p ECs | GO:0051987 | positive regulation of attachment of spindle microtubules to kinetochore | 3/179 | 11/18860 | 28.74 | 0.000131 | 0.00209 | 3 | BIRC5/AURKB/CDCA8 |
| Malignant epithelial | C6 DNA-r p ECs | GO:0006284 | base-excision repair | 5/179 | 52/18860 | 10.13 | 0.000132 | 0.00209 | 5 | PCNA/FEN1/RPA3/POLD1/HMGB1 |
| Malignant epithelial | C6 DNA-r p ECs | GO:0051338 | regulation of transferase activity | 11/179 | 304/18860 | 3.81 | 0.00016 | 0.00252 | 11 | DSCC1/PKMYT1/CDC6/CCNE2/RFC3/CLSPN/PCNA/RFC2/MAD2L1/UBE2C/CDKN2A |
| Malignant epithelial | C6 DNA-r p ECs | GO:0009176 | pyrimidine deoxyribonucleoside monophosphate metabolic process | 3/179 | 12/18860 | 26.34 | 0.000174 | 0.00271 | 3 | TYMS/SHMT1/DUT |
| Malignant epithelial | C6 DNA-r p ECs | GO:0009120 | deoxyribonucleoside metabolic process | 3/179 | 13/18860 | 24.31 | 0.000224 | 0.00346 | 3 | CDK1/TK1/DTYMK |
| Malignant epithelial | C6 DNA-r p ECs | GO:1903490 | positive regulation of mitotic cytokinesis | 3/179 | 13/18860 | 24.31 | 0.000224 | 0.00346 | 3 | BIRC5/AURKB/CDCA8 |
| Malignant epithelial | C6 DNA-r p ECs | GO:0031297 | replication fork processing | 5/179 | 59/18860 | 8.93 | 0.000241 | 0.00369 | 5 | CDK1/RAD51/PCNA/BRCA2/DONSON |
| Malignant epithelial | C6 DNA-r p ECs | GO:0000079 | regulation of cyclin-dependent protein serine/threonine kinase activity | 4/179 | 33/18860 | 12.77 | 0.000259 | 0.00394 | 4 | PKMYT1/CDC6/CCNE2/CDKN2A |
| Malignant epithelial | C6 DNA-r p ECs | GO:0051293 | establishment of spindle localization | 5/179 | 61/18860 | 8.64 | 0.000282 | 0.00423 | 5 | NUSAP1/SKA3/MAD2L1/DYNLT1/NDC80 |
| Malignant epithelial | C6 DNA-r p ECs | GO:0006213 | pyrimidine nucleoside metabolic process | 3/179 | 14/18860 | 22.58 | 0.000283 | 0.00423 | 3 | CDK1/TK1/DTYMK |
| Malignant epithelial | C6 DNA-r p ECs | GO:0006287 | base-excision repair, gap-filling | 3/179 | 14/18860 | 22.58 | 0.000283 | 0.00423 | 3 | PCNA/FEN1/POLD1 |
| Malignant epithelial | C6 DNA-r p ECs | GO:0006221 | pyrimidine nucleotide biosynthetic process | 4/179 | 34/18860 | 12.4 | 0.000291 | 0.00432 | 4 | TYMS/SHMT1/DTYMK/DUT |
| Malignant epithelial | C6 DNA-r p ECs | GO:0044843 | cell cycle G1/S phase transition | 10/179 | 275/18860 | 3.83 | 0.000306 | 0.00451 | 10 | RRM2/CDK1/CDC6/CCNE2/E2F1/EIF4EBP1/CDK2/CCNA2/CDKN2A/PSME2 |
| Malignant epithelial | C6 DNA-r p ECs | GO:0007062 | sister chromatid cohesion | 5/179 | 63/18860 | 8.36 | 0.000328 | 0.0048 | 5 | DSCC1/CDCA5/FEN1/ESCO2/RAD51C |
| Malignant epithelial | C6 DNA-r p ECs | GO:0000722 | telomere maintenance via recombination | 3/179 | 15/18860 | 21.07 | 0.000352 | 0.00505 | 3 | RAD51/BRCA2/RAD51C |
| Malignant epithelial | C6 DNA-r p ECs | GO:0001833 | inner cell mass cell proliferation | 3/179 | 15/18860 | 21.07 | 0.000352 | 0.00505 | 3 | BRCA2/NCAPG2/GINS1 |
| Malignant epithelial | C6 DNA-r p ECs | GO:1904666 | regulation of ubiquitin protein ligase activity | 3/179 | 15/18860 | 21.07 | 0.000352 | 0.00505 | 3 | MAD2L1/UBE2C/CDKN2A |
| Malignant epithelial | C6 DNA-r p ECs | GO:0006298 | mismatch repair | 4/179 | 36/18860 | 11.71 | 0.000365 | 0.0052 | 4 | PCNA/RPA3/RNASEH2A/HMGB1 |
| Malignant epithelial | C6 DNA-r p ECs | GO:0051653 | spindle localization | 5/179 | 66/18860 | 7.98 | 0.000407 | 0.00578 | 5 | NUSAP1/SKA3/MAD2L1/DYNLT1/NDC80 |
| Malignant epithelial | C6 DNA-r p ECs | GO:0009130 | pyrimidine nucleoside monophosphate biosynthetic process | 3/179 | 16/18860 | 19.76 | 0.00043 | 0.00602 | 3 | TYMS/SHMT1/DUT |
| Malignant epithelial | C6 DNA-r p ECs | GO:1902412 | regulation of mitotic cytokinesis | 3/179 | 16/18860 | 19.76 | 0.00043 | 0.00602 | 3 | BIRC5/AURKB/CDCA8 |
| Malignant epithelial | C6 DNA-r p ECs | GO:0051302 | regulation of cell division | 8/179 | 190/18860 | 4.44 | 0.000472 | 0.00657 | 8 | CDC6/MDK/BRCA2/PRC1/BIRC5/CALM1/AURKB/CDCA8 |
| Malignant epithelial | C6 DNA-r p ECs | GO:0044771 | meiotic cell cycle phase transition | 3/179 | 17/18860 | 18.59 | 0.000518 | 0.00713 | 3 | PKMYT1/ZWINT/NDC80 |
| Malignant epithelial | C6 DNA-r p ECs | GO:0051382 | kinetochore assembly | 3/179 | 17/18860 | 18.59 | 0.000518 | 0.00713 | 3 | CENPK/CENPH/CENPW |
| Malignant epithelial | C6 DNA-r p ECs | GO:0071103 | DNA conformation change | 3/179 | 18/18860 | 17.56 | 0.000618 | 0.00845 | 3 | HMGB2/TOP2A/HMGB1 |
| Malignant epithelial | C6 DNA-r p ECs | GO:0040001 | establishment of mitotic spindle localization | 4/179 | 42/18860 | 10.03 | 0.000663 | 0.00896 | 4 | NUSAP1/MAD2L1/DYNLT1/NDC80 |
| Malignant epithelial | C6 DNA-r p ECs | GO:0045740 | positive regulation of DNA replication | 4/179 | 42/18860 | 10.03 | 0.000663 | 0.00896 | 4 | CDK1/CDT1/PCNA/CDK2 |
| Malignant epithelial | C6 DNA-r p ECs | GO:0009124 | nucleoside monophosphate biosynthetic process | 4/179 | 43/18860 | 9.8 | 0.000725 | 0.00963 | 4 | TK1/TYMS/SHMT1/DUT |

| compartment | subcluster | GO_ID | Description | GeneRatio | BgRatio | FoldEnrichment | pvalue | p_adjust | Count | geneID |
|---|---|---|---|---|---|---|---|---|---|---|
| Malignant epithelial | C6 DNA-r p ECs | GO:0036297 | interstrand cross-link repair | 4/179 | 43/18860 | 9.8 | 0.000725 | 0.00963 | 4 | FANCG/RAD51/FANCB/RAD51AP1 |
| Malignant epithelial | C6 DNA-r p ECs | GO:0090329 | regulation of DNA-templated DNA replication | 4/179 | 43/18860 | 9.8 | 0.000725 | 0.00963 | 4 | CDT1/MCM7/CDK2/MCM4 |
| Malignant epithelial | C6 DNA-r p ECs | GO:0044774 | mitotic DNA integrity checkpoint signaling | 5/179 | 75/18860 | 7.02 | 0.000735 | 0.0097 | 5 | CDK1/CDC6/CLSPN/CDK2/DONSON |
| Malignant epithelial | C6 DNA-r p ECs | GO:0010639 | negative regulation of organelle organization | 11/179 | 367/18860 | 3.16 | 0.000787 | 0.0103 | 11 | SPC25/ZWINT/SKA3/DIAPH3/MAD2L1/TRIP13/BIRC5/AURKB/NDC80/CDCA8/CKAP2 |
| Malignant epithelial | C6 DNA-r p ECs | GO:0046653 | tetrahydrofolate metabolic process | 3/179 | 20/18860 | 15.8 | 0.000851 | 0.0111 | 3 | TYMS/DHFR/SHMT1 |
| Malignant epithelial | C6 DNA-r p ECs | GO:0009163 | nucleoside biosynthetic process | 3/179 | 21/18860 | 15.05 | 0.000986 | 0.0127 | 3 | CDK1/TK1/DTYMK |
| Malignant epithelial | C6 DNA-r p ECs | GO:0034404 | nucleobase-containing small molecule biosynthetic process | 3/179 | 21/18860 | 15.05 | 0.000986 | 0.0127 | 3 | CDK1/TK1/DTYMK |
| Malignant epithelial | C6 DNA-r p ECs | GO:0051447 | negative regulation of meiotic cell cycle | 3/179 | 21/18860 | 15.05 | 0.000986 | 0.0127 | 3 | PKMYT1/ZWINT/TRIP13 |
| Malignant epithelial | C6 DNA-r p ECs | GO:0000731 | DNA synthesis involved in DNA repair | 4/179 | 47/18860 | 8.97 | 0.00102 | 0.0129 | 4 | RFC3/PCNA/POLD1/POLD2 |
| Malignant epithelial | C6 DNA-r p ECs | GO:0009116 | nucleoside metabolic process | 4/179 | 47/18860 | 8.97 | 0.00102 | 0.0129 | 4 | CDK1/TK1/ICMT/DTYMK |
| Malignant epithelial | C6 DNA-r p ECs | GO:0051294 | establishment of spindle orientation | 4/179 | 47/18860 | 8.97 | 0.00102 | 0.0129 | 4 | SKA3/MAD2L1/DYNLT1/NDC80 |
| Malignant epithelial | C6 DNA-r p ECs | GO:0009129 | pyrimidine nucleoside monophosphate metabolic process | 3/179 | 22/18860 | 14.37 | 0.00113 | 0.0142 | 3 | TYMS/SHMT1/DUT |
| Malignant epithelial | C6 DNA-r p ECs | GO:2000278 | regulation of DNA biosynthetic process | 5/179 | 85/18860 | 6.2 | 0.00129 | 0.0161 | 5 | DSCC1/RFC3/PCNA/RFC2/CCNA2 |
| Malignant epithelial | C6 DNA-r p ECs | GO:2000001 | regulation of DNA damage checkpoint | 3/179 | 23/18860 | 13.74 | 0.00129 | 0.0161 | 3 | RAD51/BRCA2/WDR76 |
| Malignant epithelial | C6 DNA-r p ECs | GO:0032467 | positive regulation of cytokinesis | 4/179 | 51/18860 | 8.26 | 0.00138 | 0.0171 | 4 | CDC6/BIRC5/AURKB/CDCA8 |
| Malignant epithelial | C6 DNA-r p ECs | GO:0001832 | blastocyst growth | 3/179 | 24/18860 | 13.17 | 0.00147 | 0.0179 | 3 | BRCA2/NCAPG2/GINS1 |
| Malignant epithelial | C6 DNA-r p ECs | GO:0006730 | one-carbon metabolic process | 3/179 | 24/18860 | 13.17 | 0.00147 | 0.0179 | 3 | TYMS/DHFR/SHMT1 |
| Malignant epithelial | C6 DNA-r p ECs | GO:0009147 | pyrimidine nucleoside triphosphate metabolic process | 3/179 | 24/18860 | 13.17 | 0.00147 | 0.0179 | 3 | TYMS/DTYMK/DUT |
| Malignant epithelial | C6 DNA-r p ECs | GO:0007292 | female gamete generation | 7/179 | 176/18860 | 4.19 | 0.00148 | 0.0179 | 7 | NCAPH/SKA3/MDK/BRCA2/TRIP13/NDC80/TOP2A |
| Malignant epithelial | C6 DNA-r p ECs | GO:2000045 | regulation of G1/S transition of mitotic cell cycle | 7/179 | 178/18860 | 4.14 | 0.00158 | 0.019 | 7 | RRM2/CDC6/CCNE2/E2F1/CDK2/CDKN2A/PSME2 |
| Malignant epithelial | C6 DNA-r p ECs | GO:0006220 | pyrimidine nucleotide metabolic process | 4/179 | 54/18860 | 7.8 | 0.00171 | 0.0204 | 4 | TYMS/SHMT1/DTYMK/DUT |
| Malignant epithelial | C6 DNA-r p ECs | GO:0044818 | mitotic G2/M transition checkpoint | 4/179 | 54/18860 | 7.8 | 0.00171 | 0.0204 | 4 | CDK1/CDC6/CLSPN/DONSON |
| Malignant epithelial | C6 DNA-r p ECs | GO:0019985 | translesion synthesis | 3/179 | 26/18860 | 12.16 | 0.00186 | 0.0219 | 3 | PCNA/POLD1/POLD2 |
| Malignant epithelial | C6 DNA-r p ECs | GO:0051438 | regulation of ubiquitin-protein transferase activity | 3/179 | 26/18860 | 12.16 | 0.00186 | 0.0219 | 3 | MAD2L1/UBE2C/CDKN2A |
| Malignant epithelial | C6 DNA-r p ECs | GO:0006760 | folic acid-containing compound metabolic process | 3/179 | 27/18860 | 11.71 | 0.00208 | 0.0243 | 3 | TYMS/DHFR/SHMT1 |
| Malignant epithelial | C6 DNA-r p ECs | GO:1901659 | glycosyl compound biosynthetic process | 3/179 | 27/18860 | 11.71 | 0.00208 | 0.0243 | 3 | CDK1/TK1/DTYMK |
| Malignant epithelial | C6 DNA-r p ECs | GO:0006312 | mitotic recombination | 3/179 | 28/18860 | 11.29 | 0.00231 | 0.0268 | 3 | RAD51/BRCA2/RAD51C |
| Malignant epithelial | C6 DNA-r p ECs | GO:0000281 | mitotic cytokinesis | 5/179 | 98/18860 | 5.38 | 0.00243 | 0.028 | 5 | NUSAP1/BIRC5/AURKB/CDCA8/CKAP2 |
| Malignant epithelial | C6 DNA-r p ECs | GO:0051781 | positive regulation of cell division | 5/179 | 99/18860 | 5.32 | 0.00254 | 0.0292 | 5 | CDC6/MDK/BIRC5/AURKB/CDCA8 |
| Malignant epithelial | C6 DNA-r p ECs | GO:0007141 | male meiosis I | 3/179 | 29/18860 | 10.9 | 0.00256 | 0.0293 | 3 | BRCA2/TRIP13/RAD51C |
| Malignant epithelial | C6 DNA-r p ECs | GO:0045814 | negative regulation of gene expression, epigenetic | 7/179 | 197/18860 | 3.74 | 0.0028 | 0.0318 | 7 | LMNB1/HELLS/CDK2/OIP5/HMGB1/UHRF1/PHF19 |
| Malignant epithelial | C6 DNA-r p ECs | GO:0006998 | nuclear envelope organization | 4/179 | 63/18860 | 6.69 | 0.00302 | 0.034 | 4 | CDK1/LMNB1/ICMT/VRK1 |
| Malignant epithelial | C6 DNA-r p ECs | GO:0051785 | positive regulation of nuclear division | 4/179 | 63/18860 | 6.69 | 0.00302 | 0.034 | 4 | NUSAP1/SKA3/UBE2C/RAD51AP1 |
| Malignant epithelial | C6 DNA-r p ECs | GO:0007064 | mitotic sister chromatid cohesion | 3/179 | 31/18860 | 10.2 | 0.00311 | 0.0348 | 3 | DSCC1/CDCA5/ESCO2 |
| Malignant epithelial | C6 DNA-r p ECs | GO:1902806 | regulation of cell cycle G1/S phase transition | 7/179 | 206/18860 | 3.58 | 0.00358 | 0.0399 | 7 | RRM2/CDC6/CCNE2/E2F1/CDK2/CDKN2A/PSME2 |
| Malignant epithelial | C6 DNA-r p ECs | GO:0033313 | meiotic cell cycle checkpoint signaling | 2/179 | 10/18860 | 21.07 | 0.00383 | 0.0422 | 2 | ZWINT/TRIP13 |

| compartment | subcluster | GO_ID | Description | GeneRatio | BgRatio | FoldEnrichment | pvalue | p_adjust | Count | geneID |
|---|---|---|---|---|---|---|---|---|---|---|
| Malignant epithelial | C6 DNA-r p ECs | GO:0035999 | tetrahydrofolate interconversion | 2/179 | 10/18860 | 21.07 | 0.00383 | 0.0422 | 2 | TYMS/SHMT1 |
| Malignant epithelial | C6 DNA-r p ECs | GO:0051307 | meiotic chromosome separation | 2/179 | 10/18860 | 21.07 | 0.00383 | 0.0422 | 2 | NCAPH/ZWINT |
| Malignant epithelial | C6 DNA-r p ECs | GO:0045143 | homologous chromosome segregation | 4/179 | 68/18860 | 6.2 | 0.00398 | 0.0436 | 4 | MND1/ZWINT/CCNE2/TRIP13 |
| Malignant epithelial | C6 DNA-r p ECs | GO:0042558 | pteridine-containing compound metabolic process | 3/179 | 34/18860 | 9.3 | 0.00405 | 0.0441 | 3 | TYMS/DHFR/SHMT1 |
| Malignant epithelial | C6 DNA-r p ECs | GO:0006997 | nucleus organization | 6/179 | 160/18860 | 3.95 | 0.00428 | 0.0463 | 6 | CDK1/HMGB2/LMNB1/ICMT/CDKN2A/VRK1 |
| Malignant epithelial | C6 DNA-r p ECs | GO:0040020 | regulation of meiotic nuclear division | 3/179 | 35/18860 | 9.03 | 0.0044 | 0.0472 | 3 | ZWINT/TRIP13/RAD51AP1 |
| Malignant epithelial | C6 DNA-r p ECs | GO:0051347 | positive regulation of transferase activity | 6/179 | 161/18860 | 3.93 | 0.00441 | 0.0472 | 6 | DSCC1/RFC3/CLSPN/PCNA/RFC2/UBE2C |
| Malignant epithelial | C6 DNA-r p ECs | GO:0051445 | regulation of meiotic cell cycle | 4/179 | 70/18860 | 6.02 | 0.00442 | 0.0472 | 4 | PKMYT1/ZWINT/TRIP13/RAD51AP1 |
| Malignant epithelial | C6 DNA-r p ECs | GO:1901293 | nucleoside phosphate biosynthetic process | 9/179 | 332/18860 | 2.86 | 0.00454 | 0.048 | 9 | RRM2/TK1/ACOT7/TYMS/ENO1/SHMT1/DTYMK/DUT/NAPRT |
| Malignant epithelial | C6 DNA-r p ECs | GO:0044773 | mitotic DNA damage checkpoint signaling | 4/179 | 71/18860 | 5.94 | 0.00465 | 0.048 | 4 | CDK1/CLSPN/CDK2/DONSON |
| Malignant epithelial | C6 DNA-r p ECs | GO:0070301 | cellular response to hydrogen peroxide | 4/179 | 71/18860 | 5.94 | 0.00465 | 0.048 | 4 | CDK1/PCNA/CDKN2A/PARK7 |
| Malignant epithelial | C6 DNA-r p ECs | GO:1901657 | glycosyl compound metabolic process | 4/179 | 71/18860 | 5.94 | 0.00465 | 0.048 | 4 | CDK1/TK1/ICMT/DTYMK |
| Malignant epithelial | C6 DNA-r p ECs | GO:0007077 | mitotic nuclear membrane disassembly | 2/179 | 11/18860 | 19.16 | 0.00466 | 0.048 | 2 | CDK1/VRK1 |
| Malignant epithelial | C6 DNA-r p ECs | GO:0045654 | positive regulation of megakaryocyte differentiation | 2/179 | 11/18860 | 19.16 | 0.00466 | 0.048 | 2 | HMGB2/TESC |
| Malignant epithelial | C6 DNA-r p ECs | GO:0045835 | negative regulation of meiotic nuclear division | 2/179 | 11/18860 | 19.16 | 0.00466 | 0.048 | 2 | ZWINT/TRIP13 |
| Malignant epithelial | C6 DNA-r p ECs | GO:0051444 | negative regulation of ubiquitin-protein transferase activity | 2/179 | 11/18860 | 19.16 | 0.00466 | 0.048 | 2 | MAD2L1/CDKN2A |
| Malignant epithelial | C6 DNA-r p ECs | GO:0031507 | heterochromatin formation | 6/179 | 163/18860 | 3.88 | 0.00468 | 0.048 | 6 | LMNB1/HELLS/CDK2/OIP5/HMGB1/UHRF1 |
| Malignant epithelial | C6 DNA-r p ECs | GO:0007095 | mitotic G2 DNA damage checkpoint signaling | 3/179 | 36/18860 | 8.78 | 0.00477 | 0.0487 | 3 | CDK1/CLSPN/DONSON |
| Malignant epithelial | C6 DNA-r p ECs | GO:0097193 | intrinsic apoptotic signaling pathway | 9/179 | 335/18860 | 2.83 | 0.00481 | 0.0489 | 9 | MELK/E2F1/ENO1/BRCA2/IFI27L1/HELLS/RTKN2/SIVA1/PARK7 |
| Fibroblast | F0 SFRP2+ Fib | GO:0030198 | extracellular matrix organization | 21/187 | 310/16509 | 5.98 | 5.33e-11 | 5.61e-08 | 21 | SFRP2/FBLN1/CCDC80/CST3/MMP2/DPT/LUM/FBLN5/TNXB/CTSK/ABI3BP/PDGFRA/MFAP4/LRP1/MMP23B/COL1… |
| Fibroblast | F0 SFRP2+ Fib | GO:0043062 | extracellular structure organization | 21/187 | 311/16509 | 5.96 | 5.66e-11 | 5.61e-08 | 21 | SFRP2/FBLN1/CCDC80/CST3/MMP2/DPT/LUM/FBLN5/TNXB/CTSK/ABI3BP/PDGFRA/MFAP4/LRP1/MMP23B/COL1… |
| Fibroblast | F0 SFRP2+ Fib | GO:0045229 | external encapsulating structure organization | 21/187 | 312/16509 | 5.94 | 6.01e-11 | 5.61e-08 | 21 | SFRP2/FBLN1/CCDC80/CST3/MMP2/DPT/LUM/FBLN5/TNXB/CTSK/ABI3BP/PDGFRA/MFAP4/LRP1/MMP23B/COL1… |
| Fibroblast | F0 SFRP2+ Fib | GO:0006956 | complement activation | 9/187 | 62/16509 | 12.82 | 3.06e-08 | 2.14e-05 | 9 | C3/C7/C1S/C1R/CFD/CLU/SERPING1/CFH/CFI |
| Fibroblast | F0 SFRP2+ Fib | GO:0031589 | cell-substrate adhesion | 18/187 | 354/16509 | 4.49 | 1.16e-07 | 6.52e-05 | 18 | FBLN1/CCDC80/SFRP1/APOD/FBLN5/TNXB/ABI3BP/FBLN2/CD34/MDK/DAB2/S100A10/CD63/THY1/PLPP3/SGC… |
| Fibroblast | F0 SFRP2+ Fib | GO:0032963 | collagen metabolic process | 10/187 | 105/16509 | 8.41 | 3.11e-07 | 0.000145 | 10 | CST3/MMP2/TNXB/CTSK/MFAP4/PCOLCE/MMP23B/COL1A2/CYGB/FAP |
| Fibroblast | F0 SFRP2+ Fib | GO:2000146 | negative regulation of cell motility | 16/187 | 306/16509 | 4.62 | 4.18e-07 | 0.000167 | 16 | SFRP2/FBLN1/SERPINF1/DCN/SFRP1/APOD/CXCL12/PODN/LRP1/CD63/THY1/SLIT2/NGFR/CYP1B1/MITF/CYGB |
| Fibroblast | F0 SFRP2+ Fib | GO:0001523 | retinoid metabolic process | 9/187 | 86/16509 | 9.24 | 5.5e-07 | 0.000192 | 9 | ADH1B/ALDH1A1/LRP1/SCPEP1/DHRS3/RARRES2/CYP1B1/AKR1C1/AKR1C3 |

| compartment | subcluster | GO_ID | Description | GeneRatio | BgRatio | FoldEnrichment | pvalue | p_adjust | Count | geneID |
|---|---|---|---|---|---|---|---|---|---|---|
| Fibroblast | F0 SFRP2+ Fib | GO:0016101 | diterpenoid metabolic process | 9/187 | 90/16509 | 8.83 | 8.12e-07 | 0.000251 | 9 | ADH1B/ALDH1A1/LRP1/SCPEP1/DHRS3/RARRES2/CYP1B1/AKR1C1/AKR1C3 |
| Fibroblast | F0 SFRP2+ Fib | GO:0040013 | negative regulation of locomotion | 16/187 | 324/16509 | 4.36 | 8.94e-07 | 0.000251 | 16 | SFRP2/FBLN1/SERPINF1/DCN/SFRP1/APOD/CXCL12/PODN/LRP1/CD63/THY1/SLIT2/NGFR/CYP1B1/MITF/CYGB |
| Fibroblast | F0 SFRP2+ Fib | GO:0006720 | isoprenoid metabolic process | 10/187 | 119/16509 | 7.42 | 1e-06 | 0.000255 | 10 | ADH1B/ALDH1A1/LRP1/NPC2/SCPEP1/DHRS3/RARRES2/CYP1B1/AKR1C1/AKR1C3 |
| Fibroblast | F0 SFRP2+ Fib | GO:0030336 | negative regulation of cell migration | 15/187 | 291/16509 | 4.55 | 1.17e-06 | 0.000272 | 15 | SFRP2/SERPINF1/DCN/SFRP1/APOD/CXCL12/PODN/LRP1/CD63/THY1/SLIT2/NGFR/CYP1B1/MITF/CYGB |
| Fibroblast | F0 SFRP2+ Fib | GO:0010811 | positive regulation of cell-substrate adhesion | 10/187 | 130/16509 | 6.79 | 2.25e-06 | 0.000469 | 10 | FBLN1/CCDC80/SFRP1/ABI3BP/FBLN2/MDK/DAB2/S100A10/THY1/PLPP3 |
| Fibroblast | F0 SFRP2+ Fib | GO:0030510 | regulation of BMP signaling pathway | 9/187 | 102/16509 | 7.79 | 2.34e-06 | 0.000469 | 9 | SFRP2/SFRP1/SFRP4/CHRDL1/GPC3/HTRA1/FSTL1/FBN1/CHRDL2 |
| Fibroblast | F0 SFRP2+ Fib | GO:0006721 | terpenoid metabolic process | 9/187 | 105/16509 | 7.57 | 2.99e-06 | 0.000524 | 9 | ADH1B/ALDH1A1/LRP1/SCPEP1/DHRS3/RARRES2/CYP1B1/AKR1C1/AKR1C3 |
| Fibroblast | F0 SFRP2+ Fib | GO:0090287 | regulation of cellular response to growth factor stimulus | 15/187 | 314/16509 | 4.22 | 2.99e-06 | 0.000524 | 15 | SFRP2/DCN/SFRP1/TNXB/SFRP4/CILP/CHRDL1/CD63/GPC3/HTRA1/SLIT2/FSTL1/NGFR/FBN1/CHRDL2 |
| Fibroblast | F0 SFRP2+ Fib | GO:0090288 | negative regulation of cellular response to growth factor stimulus | 9/187 | 106/16509 | 7.5 | 3.23e-06 | 0.000533 | 9 | SFRP2/DCN/SFRP1/CHRDL1/HTRA1/SLIT2/NGFR/FBN1/CHRDL2 |
| Fibroblast | F0 SFRP2+ Fib | GO:0006958 | complement activation, classical pathway | 6/187 | 39/16509 | 13.58 | 4.66e-06 | 0.000725 | 6 | C3/C7/C1S/C1R/CLU/CFI |
| Fibroblast | F0 SFRP2+ Fib | GO:0006959 | humoral immune response | 14/187 | 289/16509 | 4.28 | 5.5e-06 | 0.000811 | 14 | C3/C7/C1S/C1R/CFD/CXCL12/CLU/LGALS3/SERPING1/CFH/SPON2/CXCL14/RARRES2/CFI |
| Fibroblast | F0 SFRP2+ Fib | GO:0031099 | regeneration | 11/187 | 178/16509 | 5.46 | 5.91e-06 | 0.000828 | 11 | IGF1/MMP2/APOD/CXCL12/MDK/NFIB/THY1/ANXA1/GAS6/OMG/CPQ |
| Fibroblast | F0 SFRP2+ Fib | GO:1901342 | regulation of vasculature development | 14/187 | 294/16509 | 4.2 | 6.7e-06 | 0.000894 | 14 | C3/SFRP2/SERPINF1/DCN/SFRP1/GPNMB/FBLN5/CXCL12/CD34/MDK/PTGIS/THBS4/CYP1B1/HSPB6 |
| Fibroblast | F0 SFRP2+ Fib | GO:0048771 | tissue remodeling | 11/187 | 182/16509 | 5.34 | 7.31e-06 | 0.00093 | 11 | IGF1/CST3/MMP2/SFRP1/GPNMB/CTSK/MDK/THBS4/LEPR/RSPO3/MITF |
| Fibroblast | F0 SFRP2+ Fib | GO:2000050 | regulation of non-canonical Wnt signaling pathway | 5/187 | 25/16509 | 17.66 | 7.81e-06 | 0.000951 | 5 | SFRP1/SFRP4/DAB2/GPC3/RSPO3 |
| Fibroblast | F0 SFRP2+ Fib | GO:0002685 | regulation of leukocyte migration | 12/187 | 222/16509 | 4.77 | 8.76e-06 | 0.00102 | 12 | APOD/CXCL12/LGALS3/MDK/THY1/SLIT2/ANXA1/THBS4/JAM2/RARRES2/ICAM1/GAS6 |
| Fibroblast | F0 SFRP2+ Fib | GO:0085029 | extracellular matrix assembly | 6/187 | 44/16509 | 12.04 | 9.62e-06 | 0.00104 | 6 | FBLN5/TNXB/MFAP4/COL1A2/ELN/GAS6 |
| Fibroblast | F0 SFRP2+ Fib | GO:0048145 | regulation of fibroblast proliferation | 8/187 | 92/16509 | 7.68 | 9.64e-06 | 0.00104 | 8 | FBLN1/IGF1/SFRP1/PDGFRA/SOD2/NGFR/NUPR1/GAS6 |
| Fibroblast | F0 SFRP2+ Fib | GO:0050673 | epithelial cell proliferation | 17/187 | 438/16509 | 3.43 | 1.04e-05 | 0.00108 | 17 | SFRP2/SERPINF1/IGF1/IGFBP4/SFRP1/FGF7/CXCL12/CD34/MDK/DAB2/NFIB/GPC3/THBS4/NGFR/NUPR1/OSR… |
| Fibroblast | F0 SFRP2+ Fib | GO:0030509 | BMP signaling pathway | 10/187 | 156/16509 | 5.66 | 1.15e-05 | 0.00115 | 10 | SFRP2/SFRP1/SFRP4/CHRDL1/NFIA/GPC3/HTRA1/FSTL1/FBN1/CHRDL2 |
| Fibroblast | F0 SFRP2+ Fib | GO:0030199 | collagen fibril organization | 7/187 | 69/16509 | 8.96 | 1.28e-05 | 0.00123 | 7 | SFRP2/DPT/LUM/TNXB/COL1A2/COL14A1/CYP1B1 |
| Fibroblast | F0 SFRP2+ Fib | GO:0006909 | phagocytosis | 12/187 | 239/16509 | 4.43 | 1.84e-05 | 0.0017 | 12 | C3/C1S/SRPX/C1R/GSN/LRP1/COLEC12/SPON2/ANXA1/LEPR/CD302/GAS6 |
| Fibroblast | F0 SFRP2+ Fib | GO:2000052 | positive regulation of non-canonical Wnt signaling pathway | 4/187 | 15/16509 | 23.54 | 1.97e-05 | 0.0017 | 4 | SFRP1/DAB2/GPC3/RSPO3 |

| compartment | subcluster | GO_ID | Description | GeneRatio | BgRatio | FoldEnrichment | pvalue | p_adjust | Count | geneID |
|---|---|---|---|---|---|---|---|---|---|---|
| Fibroblast | F0 SFRP2+ Fib | GO:0051216 | cartilage development | 11/187 | 203/16509 | 4.78 | 2.04e-05 | 0.0017 | 11 | SFRP2/MGP/OGN/EFEMP1/LUM/CTSK/MDK/NFIA/NFIB/OSR2/CHRDL2 |
| Fibroblast | F0 SFRP2+ Fib | GO:0030178 | negative regulation of Wnt signaling pathway | 10/187 | 168/16509 | 5.25 | 2.19e-05 | 0.0017 | 10 | SFRP2/IGFBP4/SFRP1/SFRP4/LRP1/MDK/IGFBP6/DAB2/GPC3/BICC1 |
| Fibroblast | F0 SFRP2+ Fib | GO:0071772 | response to BMP | 10/187 | 168/16509 | 5.25 | 2.19e-05 | 0.0017 | 10 | SFRP2/SFRP1/SFRP4/CHRDL1/NFIA/GPC3/HTRA1/FSTL1/FBN1/CHRDL2 |
| Fibroblast | F0 SFRP2+ Fib | GO:0071773 | cellular response to BMP stimulus | 10/187 | 168/16509 | 5.25 | 2.19e-05 | 0.0017 | 10 | SFRP2/SFRP1/SFRP4/CHRDL1/NFIA/GPC3/HTRA1/FSTL1/FBN1/CHRDL2 |
| Fibroblast | F0 SFRP2+ Fib | GO:0090090 | negative regulation of canonical Wnt signaling pathway | 9/187 | 134/16509 | 5.93 | 2.19e-05 | 0.0017 | 9 | SFRP2/IGFBP4/SFRP1/SFRP4/MDK/IGFBP6/DAB2/GPC3/BICC1 |
| Fibroblast | F0 SFRP2+ Fib | GO:0042445 | hormone metabolic process | 12/187 | 244/16509 | 4.34 | 2.26e-05 | 0.00171 | 12 | ADH1B/ALDH1A1/CTSK/PDGFRA/DAB2/SCPEP1/DHRS3/HSD17B11/CYP1B1/AKR1C1/AKR1C3/CPQ |
| Fibroblast | F0 SFRP2+ Fib | GO:0034308 | primary alcohol metabolic process | 8/187 | 104/16509 | 6.79 | 2.38e-05 | 0.00174 | 8 | IGF1/ADH1B/ALDH1A1/DAB2/DHRS3/CYP1B1/AKR1C1/AKR1C3 |
| Fibroblast | F0 SFRP2+ Fib | GO:0045765 | regulation of angiogenesis | 13/187 | 287/16509 | 4.0 | 2.44e-05 | 0.00174 | 13 | C3/SFRP2/SERPINF1/DCN/SFRP1/GPNMB/FBLN5/CD34/MDK/PTGIS/THBS4/CYP1B1/HSPB6 |
| Fibroblast | F0 SFRP2+ Fib | GO:0061448 | connective tissue development | 13/187 | 288/16509 | 3.99 | 2.53e-05 | 0.00174 | 13 | SFRP2/MGP/OGN/IGF1/EFEMP1/LUM/CTSK/CD34/MDK/NFIA/NFIB/OSR2/CHRDL2 |
| Fibroblast | F0 SFRP2+ Fib | GO:0002455 | humoral immune response mediated by circulating immunoglobulin | 6/187 | 52/16509 | 10.19 | 2.57e-05 | 0.00174 | 6 | C3/C7/C1S/C1R/CLU/CFI |
| Fibroblast | F0 SFRP2+ Fib | GO:0001905 | activation of membrane attack complex | 4/187 | 16/16509 | 22.07 | 2.61e-05 | 0.00174 | 4 | C3/C1S/C1R/CLU |
| Fibroblast | F0 SFRP2+ Fib | GO:0048545 | response to steroid hormone | 14/187 | 334/16509 | 3.7 | 2.8e-05 | 0.00182 | 14 | SERPINF1/IGF1/SFRP1/PTGDS/MDK/DAB2/TXNIP/SLIT2/ANXA1/CDO1/TFPI/AKR1C3/LMO3/PLPP1 |
| Fibroblast | F0 SFRP2+ Fib | GO:0031638 | zymogen activation | 6/187 | 53/16509 | 9.99 | 2.88e-05 | 0.00183 | 6 | C1S/C1R/CFD/CTSH/SERPINE2/S100A10 |
| Fibroblast | F0 SFRP2+ Fib | GO:0050900 | leukocyte migration | 15/187 | 381/16509 | 3.48 | 2.99e-05 | 0.00186 | 15 | C3/MMP2/APOD/CXCL12/CD34/LGALS3/MDK/THY1/SLIT2/ANXA1/THBS4/JAM2/RARRES2/ICAM1/GAS6 |
| Fibroblast | F0 SFRP2+ Fib | GO:1904645 | response to amyloid-beta | 6/187 | 54/16509 | 9.81 | 3.2e-05 | 0.00195 | 6 | IGF1/MMP2/LRP1/NGFR/PRNP/ICAM1 |
| Fibroblast | F0 SFRP2+ Fib | GO:0006957 | complement activation, alternative pathway | 4/187 | 17/16509 | 20.77 | 3.38e-05 | 0.00201 | 4 | C3/C7/CFD/CFH |
| Fibroblast | F0 SFRP2+ Fib | GO:0007162 | negative regulation of cell adhesion | 13/187 | 297/16509 | 3.86 | 3.49e-05 | 0.00203 | 13 | FBLN1/PLA2G2A/MMP2/APOD/GPNMB/CXCL12/SERPINE2/LGALS3/MDK/FGL2/ANXA1/PRNP/CYP1B1 |
| Fibroblast | F0 SFRP2+ Fib | GO:0048144 | fibroblast proliferation | 8/187 | 111/16509 | 6.36 | 3.81e-05 | 0.00218 | 8 | FBLN1/IGF1/SFRP1/PDGFRA/SOD2/NGFR/NUPR1/GAS6 |
| Fibroblast | F0 SFRP2+ Fib | GO:0010810 | regulation of cell-substrate adhesion | 11/187 | 218/16509 | 4.45 | 3.94e-05 | 0.00221 | 11 | FBLN1/CCDC80/SFRP1/APOD/ABI3BP/FBLN2/MDK/DAB2/S100A10/THY1/PLPP3 |
| Fibroblast | F0 SFRP2+ Fib | GO:1902430 | negative regulation of amyloid-beta formation | 4/187 | 18/16509 | 19.62 | 4.31e-05 | 0.00237 | 4 | IGF1/CLU/SPON1/PRNP |
| Fibroblast | F0 SFRP2+ Fib | GO:0050866 | negative regulation of cell activation | 11/187 | 222/16509 | 4.37 | 4.65e-05 | 0.00247 | 11 | PLA2G2A/SFRP1/GPNMB/PDGFRA/SERPINE2/LGALS3/MDK/FGL2/ANXA1/PRNP/CYGB |
| Fibroblast | F0 SFRP2+ Fib | GO:0002687 | positive regulation of leukocyte migration | 9/187 | 148/16509 | 5.37 | 4.8e-05 | 0.00247 | 9 | CXCL12/LGALS3/MDK/THY1/THBS4/JAM2/RARRES2/ICAM1/GAS6 |
| Fibroblast | F0 SFRP2+ Fib | GO:0070613 | regulation of protein processing | 6/187 | 58/16509 | 9.13 | 4.84e-05 | 0.00247 | 6 | GSN/SERPINE2/SPON1/S100A10/TMEM98/PRNP |
| Fibroblast | F0 SFRP2+ Fib | GO:0071674 | mononuclear cell migration | 11/187 | 223/16509 | 4.35 | 4.84e-05 | 0.00247 | 11 | MMP2/APOD/CXCL12/LGALS3/MDK/SLIT2/ANXA1/JAM2/RARRES2/ICAM1/GAS6 |
| Fibroblast | F0 SFRP2+ Fib | GO:0090092 | regulation of transmembrane receptor protein serine/threonine kinase signaling pathway | 12/187 | 266/16509 | 3.98 | 5.24e-05 | 0.00261 | 12 | SFRP2/SFRP1/SFRP4/CILP/LRP1/CHRDL1/DAB2/GPC3/HTRA1/FSTL1/FBN1/CHRDL2 |

| compartment | subcluster | GO_ID | Description | GeneRatio | BgRatio | FoldEnrichment | pvalue | p_adjust | Count | geneID |
|---|---|---|---|---|---|---|---|---|---|---|
| Fibroblast | F0 SFRP2+ Fib | GO:0030514 | negative regulation of BMP signaling pathway | 6/187 | 59/16509 | 8.98 | 5.34e-05 | 0.00261 | 6 | SFRP2/SFRP1/CHRDL1/HTRA1/FBN1/CHRDL2 |
| Fibroblast | F0 SFRP2+ Fib | GO:1902992 | negative regulation of amyloid precursor protein catabolic process | 4/187 | 19/16509 | 18.59 | 5.41e-05 | 0.00261 | 4 | IGF1/CLU/SPON1/PRNP |
| Fibroblast | F0 SFRP2+ Fib | GO:0006910 | phagocytosis, recognition | 5/187 | 37/16509 | 11.93 | 5.74e-05 | 0.00273 | 5 | C3/C1S/C1R/COLEC12/SPON2 |
| Fibroblast | F0 SFRP2+ Fib | GO:0001906 | cell killing | 12/187 | 269/16509 | 3.94 | 5.84e-05 | 0.00273 | 12 | C3/C7/C1S/PLA2G2A/C1R/CXCL12/CTSH/CLU/LGALS3/CFH/CXCL14/ICAM1 |
| Fibroblast | F0 SFRP2+ Fib | GO:0048608 | reproductive structure development | 13/187 | 313/16509 | 3.67 | 5.98e-05 | 0.00274 | 13 | C3/SFRP2/SERPINF1/IGF1/MMP2/SFRP1/PDGFRA/SERPINE2/PSAP/SLIT2/BASP1/NUPR1/AKR1C3 |
| Fibroblast | F0 SFRP2+ Fib | GO:0050678 | regulation of epithelial cell proliferation | 14/187 | 359/16509 | 3.44 | 6.13e-05 | 0.00277 | 14 | SFRP2/SERPINF1/IGF1/SFRP1/FGF7/CXCL12/MDK/DAB2/NFIB/GPC3/THBS4/NGFR/NUPR1/OSR2 |
| Fibroblast | F0 SFRP2+ Fib | GO:0001503 | ossification | 15/187 | 407/16509 | 3.25 | 6.34e-05 | 0.00282 | 15 | SFRP2/MGP/IGF1/MMP2/SFRP1/MDK/CHRDL1/COL1A2/GPC3/VCAN/CLEC11A/DHRS3/TWIST2/OSR2/CHRDL2 |
| Fibroblast | F0 SFRP2+ Fib | GO:0061458 | reproductive system development | 13/187 | 318/16509 | 3.61 | 7.02e-05 | 0.00307 | 13 | C3/SFRP2/SERPINF1/IGF1/MMP2/SFRP1/PDGFRA/SERPINE2/PSAP/SLIT2/BASP1/NUPR1/AKR1C3 |
| Fibroblast | F0 SFRP2+ Fib | GO:1903317 | regulation of protein maturation | 6/187 | 63/16509 | 8.41 | 7.75e-05 | 0.00334 | 6 | GSN/SERPINE2/SPON1/S100A10/TMEM98/PRNP |
| Fibroblast | F0 SFRP2+ Fib | GO:0030449 | regulation of complement activation | 4/187 | 21/16509 | 16.82 | 8.2e-05 | 0.00348 | 4 | C3/CLU/SERPING1/CFH |
| Fibroblast | F0 SFRP2+ Fib | GO:0001656 | metanephros development | 7/187 | 92/16509 | 6.72 | 8.32e-05 | 0.00348 | 7 | CTSH/PDGFRA/CD34/GPC3/BASP1/OSR2/FBN1 |
| Fibroblast | F0 SFRP2+ Fib | GO:0007160 | cell-matrix adhesion | 11/187 | 238/16509 | 4.08 | 8.7e-05 | 0.00358 | 11 | SFRP1/APOD/FBLN5/TNXB/CD34/S100A10/CD63/THY1/PLPP3/SGCE/SVEP1 |
| Fibroblast | F0 SFRP2+ Fib | GO:0007178 | cell surface receptor protein serine/threonine kinase signaling pathway | 14/187 | 374/16509 | 3.3 | 9.48e-05 | 0.00381 | 14 | SFRP2/SFRP1/SFRP4/CILP/LRP1/CHRDL1/COL1A2/DAB2/NFIA/GPC3/HTRA1/FSTL1/FBN1/CHRDL2 |
| Fibroblast | F0 SFRP2+ Fib | GO:0030574 | collagen catabolic process | 5/187 | 41/16509 | 10.77 | 9.51e-05 | 0.00381 | 5 | CST3/MMP2/CTSK/MMP23B/FAP |
| Fibroblast | F0 SFRP2+ Fib | GO:2001233 | regulation of apoptotic signaling pathway | 14/187 | 376/16509 | 3.29 | 0.0001 | 0.00396 | 14 | SFRP2/IGF1/MMP2/SRPX/SFRP1/CXCL12/CTSH/CLU/LGALS3/SOD2/TNFSF10/NGFR/NUPR1/ICAM1 |
| Fibroblast | F0 SFRP2+ Fib | GO:0001655 | urogenital system development | 6/187 | 68/16509 | 7.79 | 0.000119 | 0.00454 | 6 | SERPINF1/IGF1/MMP2/SFRP1/PSAP/OSR2 |
| Fibroblast | F0 SFRP2+ Fib | GO:0043567 | regulation of insulin-like growth factor receptor signaling pathway | 4/187 | 23/16509 | 15.35 | 0.000119 | 0.00454 | 4 | IGF1/IGFBP4/CILP/IGFBP6 |
| Fibroblast | F0 SFRP2+ Fib | GO:0010656 | negative regulation of muscle cell apoptotic process | 5/187 | 43/16509 | 10.27 | 0.00012 | 0.00454 | 5 | SFRP2/IGF1/MDK/NUPR1/HSPB6 |
| Fibroblast | F0 SFRP2+ Fib | GO:0035265 | organ growth | 9/187 | 168/16509 | 4.73 | 0.000128 | 0.00466 | 9 | IGF1/FGF7/FBLN5/PSAP/COL14A1/PI16/LEPR/BASP1/BNC2 |
| Fibroblast | F0 SFRP2+ Fib | GO:0035567 | non-canonical Wnt signaling pathway | 6/187 | 69/16509 | 7.68 | 0.000129 | 0.00466 | 6 | SFRP2/SFRP1/SFRP4/DAB2/GPC3/RSPO3 |
| Fibroblast | F0 SFRP2+ Fib | GO:0031640 | killing of cells of another organism | 8/187 | 132/16509 | 5.35 | 0.00013 | 0.00466 | 8 | C3/C7/C1S/PLA2G2A/C1R/CXCL12/LGALS3/CXCL14 |
| Fibroblast | F0 SFRP2+ Fib | GO:0141061 | disruption of cell in another organism | 8/187 | 132/16509 | 5.35 | 0.00013 | 0.00466 | 8 | C3/C7/C1S/PLA2G2A/C1R/CXCL12/LGALS3/CXCL14 |
| Fibroblast | F0 SFRP2+ Fib | GO:1904646 | cellular response to amyloid-beta | 5/187 | 44/16509 | 10.03 | 0.000134 | 0.00476 | 5 | IGF1/LRP1/NGFR/PRNP/ICAM1 |
| Fibroblast | F0 SFRP2+ Fib | GO:0010660 | regulation of muscle cell apoptotic process | 6/187 | 70/16509 | 7.57 | 0.00014 | 0.0049 | 6 | SFRP2/IGF1/MDK/SOD2/NUPR1/HSPB6 |
| Fibroblast | F0 SFRP2+ Fib | GO:0050680 | negative regulation of epithelial cell proliferation | 8/187 | 134/16509 | 5.27 | 0.000144 | 0.00499 | 8 | SFRP2/SERPINF1/SFRP1/DAB2/NFIB/GPC3/NGFR/NUPR1 |
| Fibroblast | F0 SFRP2+ Fib | GO:0010876 | lipid localization | 15/187 | 441/16509 | 3.0 | 0.000154 | 0.00527 | 15 | C3/PLA2G2A/APOD/ABCA8/PLTP/CLU/LRP1/PSAP/NPC2/DAB2/ANXA1/AKR1C1/ABCA9/PLSCR4/CES1 |
| Fibroblast | F0 SFRP2+ Fib | GO:0141060 | disruption of anatomical structure in another organism | 8/187 | 136/16509 | 5.19 | 0.00016 | 0.00539 | 8 | C3/C7/C1S/PLA2G2A/C1R/CXCL12/LGALS3/CXCL14 |
| Fibroblast | F0 SFRP2+ Fib | GO:0034103 | regulation of tissue remodeling | 6/187 | 72/16509 | 7.36 | 0.000164 | 0.00539 | 6 | CST3/SFRP1/GPNMB/MDK/THBS4/LEPR |

| compartment | subcluster | GO_ID | Description | GeneRatio | BgRatio | FoldEnrichment | pvalue | p_adjust | Count | geneID |
|---|---|---|---|---|---|---|---|---|---|---|
| Fibroblast | F0 SFRP2+ Fib | GO:0008228 | opsonization | 4/187 | 25/16509 | 14.13 | 0.000167 | 0.00539 | 4 | C3/C1S/C1R/SPON2 |
| Fibroblast | F0 SFRP2+ Fib | GO:0010667 | negative regulation of cardiac muscle cell apoptotic process | 4/187 | 25/16509 | 14.13 | 0.000167 | 0.00539 | 4 | SFRP2/MDK/NUPR1/HSPB6 |
| Fibroblast | F0 SFRP2+ Fib | GO:0034249 | negative regulation of amide metabolic process | 4/187 | 25/16509 | 14.13 | 0.000167 | 0.00539 | 4 | IGF1/CLU/SPON1/PRNP |
| Fibroblast | F0 SFRP2+ Fib | GO:0141091 | transforming growth factor beta receptor superfamily signaling pathway | 13/187 | 348/16509 | 3.3 | 0.000172 | 0.00549 | 13 | SFRP2/SFRP1/SFRP4/CILP/CHRDL1/COL1A2/DAB2/NFIA/GPC3/HTRA1/FSTL1/FBN1/CHRDL2 |
| Fibroblast | F0 SFRP2+ Fib | GO:0006066 | alcohol metabolic process | 13/187 | 350/16509 | 3.28 | 0.000182 | 0.00574 | 13 | IGF1/ADH1B/ALDH1A1/NPC2/DAB2/DHRS3/PLPP3/LEPR/CYP1B1/AKR1C1/AKR1C3/PLPP1/CES1 |
| Fibroblast | F0 SFRP2+ Fib | GO:0010875 | positive regulation of cholesterol efflux | 4/187 | 26/16509 | 13.58 | 0.000196 | 0.0061 | 4 | ABCA8/PLTP/LRP1/CES1 |
| Fibroblast | F0 SFRP2+ Fib | GO:0006935 | chemotaxis | 15/187 | 452/16509 | 2.93 | 0.000201 | 0.0062 | 15 | C3/MMP2/GPNMB/FGF7/CXCL12/PDGFRA/LGALS3/MDK/CXCL14/SLIT2/ANXA1/THBS4/LSP1/RARRES2/GAS6 |
| Fibroblast | F0 SFRP2+ Fib | GO:0071675 | regulation of mononuclear cell migration | 8/187 | 141/16509 | 5.01 | 0.000205 | 0.00624 | 8 | APOD/CXCL12/LGALS3/MDK/SLIT2/JAM2/RARRES2/GAS6 |
| Fibroblast | F0 SFRP2+ Fib | GO:0042330 | taxis | 15/187 | 454/16509 | 2.92 | 0.000211 | 0.00636 | 15 | C3/MMP2/GPNMB/FGF7/CXCL12/PDGFRA/LGALS3/MDK/CXCL14/SLIT2/ANXA1/THBS4/LSP1/RARRES2/GAS6 |
| Fibroblast | F0 SFRP2+ Fib | GO:2001235 | positive regulation of apoptotic signaling pathway | 8/187 | 142/16509 | 4.97 | 0.000215 | 0.00641 | 8 | MMP2/SRPX/SFRP1/CTSH/CLU/TNFSF10/NGFR/NUPR1 |
| Fibroblast | F0 SFRP2+ Fib | GO:0048251 | elastic fiber assembly | 3/187 | 11/16509 | 24.08 | 0.000221 | 0.00644 | 3 | FBLN5/TNXB/MFAP4 |
| Fibroblast | F0 SFRP2+ Fib | GO:0042982 | amyloid precursor protein metabolic process | 6/187 | 76/16509 | 6.97 | 0.000221 | 0.00644 | 6 | IGF1/ITM2A/CLU/SPON1/ITM2B/PRNP |
| Fibroblast | F0 SFRP2+ Fib | GO:0060326 | cell chemotaxis | 12/187 | 311/16509 | 3.41 | 0.000228 | 0.00659 | 12 | C3/MMP2/CXCL12/PDGFRA/LGALS3/MDK/CXCL14/SLIT2/ANXA1/THBS4/RARRES2/GAS6 |
| Fibroblast | F0 SFRP2+ Fib | GO:0010657 | muscle cell apoptotic process | 6/187 | 77/16509 | 6.88 | 0.000237 | 0.00678 | 6 | SFRP2/IGF1/MDK/SOD2/NUPR1/HSPB6 |
| Fibroblast | F0 SFRP2+ Fib | GO:0010664 | negative regulation of striated muscle cell apoptotic process | 4/187 | 28/16509 | 12.61 | 0.000264 | 0.00746 | 4 | SFRP2/MDK/NUPR1/HSPB6 |
| Fibroblast | F0 SFRP2+ Fib | GO:0030595 | leukocyte chemotaxis | 10/187 | 227/16509 | 3.89 | 0.000269 | 0.00753 | 10 | C3/MMP2/CXCL12/LGALS3/MDK/SLIT2/ANXA1/THBS4/RARRES2/GAS6 |
| Fibroblast | F0 SFRP2+ Fib | GO:0030850 | prostate gland development | 5/187 | 51/16509 | 8.66 | 0.000272 | 0.00755 | 5 | SERPINF1/IGF1/MMP2/SFRP1/PSAP |
| Fibroblast | F0 SFRP2+ Fib | GO:0097278 | complement-dependent cytotoxicity | 3/187 | 12/16509 | 22.07 | 0.000292 | 0.00796 | 3 | C3/CLU/CFH |
| Fibroblast | F0 SFRP2+ Fib | GO:1903038 | negative regulation of leukocyte cell-cell adhesion | 8/187 | 149/16509 | 4.74 | 0.000298 | 0.00796 | 8 | PLA2G2A/GPNMB/CXCL12/LGALS3/MDK/FGL2/ANXA1/PRNP |
| Fibroblast | F0 SFRP2+ Fib | GO:0030195 | negative regulation of blood coagulation | 5/187 | 52/16509 | 8.49 | 0.000298 | 0.00796 | 5 | PDGFRA/SERPINE2/SERPING1/TFPI/FAP |
| Fibroblast | F0 SFRP2+ Fib | GO:0042572 | retinol metabolic process | 5/187 | 52/16509 | 8.49 | 0.000298 | 0.00796 | 5 | ADH1B/ALDH1A1/DHRS3/CYP1B1/AKR1C3 |
| Fibroblast | F0 SFRP2+ Fib | GO:0090101 | negative regulation of transmembrane receptor protein serine/threonine kinase signaling p… | 8/187 | 150/16509 | 4.71 | 0.000312 | 0.00819 | 8 | SFRP2/SFRP1/CILP/LRP1/CHRDL1/HTRA1/FBN1/CHRDL2 |
| Fibroblast | F0 SFRP2+ Fib | GO:0046209 | nitric oxide metabolic process | 6/187 | 81/16509 | 6.54 | 0.000313 | 0.00819 | 6 | IGF1/CLU/SOD2/PTGIS/CYP1B1/CYGB |
| Fibroblast | F0 SFRP2+ Fib | GO:1902106 | negative regulation of leukocyte differentiation | 7/187 | 114/16509 | 5.42 | 0.000317 | 0.00823 | 7 | SFRP1/MDK/FGL2/ANXA1/TMEM176B/TMEM176A/FBN1 |
| Fibroblast | F0 SFRP2+ Fib | GO:0001932 | regulation of protein phosphorylation | 13/187 | 371/16509 | 3.09 | 0.000321 | 0.00824 | 13 | C3/SFRP2/FBLN1/IGF1/SFRP1/FGF7/THY1/SLIT2/THBS4/COPS8/RARRES2/FBN1/GAS6 |
| Fibroblast | F0 SFRP2+ Fib | GO:0048260 | positive regulation of receptor-mediated endocytosis | 5/187 | 53/16509 | 8.33 | 0.000326 | 0.00824 | 5 | C3/SFRP4/CLU/DAB2/CD63 |
| Fibroblast | F0 SFRP2+ Fib | GO:1900047 | negative regulation of hemostasis | 5/187 | 53/16509 | 8.33 | 0.000326 | 0.00824 | 5 | PDGFRA/SERPINE2/SERPING1/TFPI/FAP |
| Fibroblast | F0 SFRP2+ Fib | GO:0014812 | muscle cell migration | 6/187 | 82/16509 | 6.46 | 0.000334 | 0.00836 | 6 | IGF1/LRP1/MDK/SLIT2/ANXA1/THBS4 |

| compartment | subcluster | GO_ID | Description | GeneRatio | BgRatio | FoldEnrichment | pvalue | p_adjust | Count | geneID |
|---|---|---|---|---|---|---|---|---|---|---|
| Fibroblast | F0 SFRP2+ Fib | GO:0006869 | lipid transport | 14/187 | 423/16509 | 2.92 | 0.000338 | 0.00838 | 14 | PLA2G2A/APOD/ABCA8/PLTP/CLU/LRP1/PSAP/NPC2/DAB2/ANXA1/AKR1C1/ABCA9/PLSCR4/CES1 |
| Fibroblast | F0 SFRP2+ Fib | GO:0030111 | regulation of Wnt signaling pathway | 12/187 | 325/16509 | 3.26 | 0.000341 | 0.00838 | 12 | SFRP2/IGFBP4/SFRP1/SFRP4/LRP1/MDK/IGFBP6/DAB2/GPC3/PLPP3/BICC1/RSPO3 |
| Fibroblast | F0 SFRP2+ Fib | GO:2001057 | reactive nitrogen species metabolic process | 6/187 | 83/16509 | 6.38 | 0.000357 | 0.0087 | 6 | IGF1/CLU/SOD2/PTGIS/CYP1B1/CYGB |
| Fibroblast | F0 SFRP2+ Fib | GO:0016485 | protein processing | 10/187 | 236/16509 | 3.74 | 0.000366 | 0.00884 | 10 | C1S/C1R/CFD/GSN/CTSH/SERPINE2/SPON1/S100A10/TMEM98/PRNP |
| Fibroblast | F0 SFRP2+ Fib | GO:0050819 | negative regulation of coagulation | 5/187 | 55/16509 | 8.03 | 0.000389 | 0.0093 | 5 | PDGFRA/SERPINE2/SERPING1/TFPI/FAP |
| Fibroblast | F0 SFRP2+ Fib | GO:0072001 | renal system development | 12/187 | 331/16509 | 3.2 | 0.000402 | 0.00954 | 12 | SERPINF1/SFRP1/CTSH/PDGFRA/CD34/NFIA/GPC3/SLIT2/BICC1/BASP1/OSR2/FBN1 |
| Fibroblast | F0 SFRP2+ Fib | GO:0006690 | icosanoid metabolic process | 7/187 | 119/16509 | 5.19 | 0.000412 | 0.00961 | 7 | PTGDS/GGT5/PTGIS/EPHX1/CYP1B1/AKR1C1/AKR1C3 |
| Fibroblast | F0 SFRP2+ Fib | GO:1903707 | negative regulation of hemopoiesis | 7/187 | 119/16509 | 5.19 | 0.000412 | 0.00961 | 7 | SFRP1/MDK/FGL2/ANXA1/TMEM176B/TMEM176A/FBN1 |
| Fibroblast | F0 SFRP2+ Fib | GO:0061138 | morphogenesis of a branching epithelium | 9/187 | 197/16509 | 4.03 | 0.000417 | 0.00966 | 9 | SFRP2/IGF1/SFRP1/FGF7/CTSH/MDK/GPC3/SLIT2/RSPO3 |
| Fibroblast | F0 SFRP2+ Fib | GO:0048146 | positive regulation of fibroblast proliferation | 5/187 | 56/16509 | 7.88 | 0.000423 | 0.00971 | 5 | FBLN1/IGF1/PDGFRA/NGFR/GAS6 |
| Fibroblast | F0 SFRP2+ Fib | GO:0009636 | response to toxic substance | 10/187 | 241/16509 | 3.66 | 0.000432 | 0.00984 | 10 | MGST1/FBLN5/ALDH1A1/SOD2/INMT/NUPR1/EPHX1/CYP1B1/CES1/CYGB |
| Fibroblast | F0 SFRP2+ Fib | GO:0045807 | positive regulation of endocytosis | 8/187 | 159/16509 | 4.44 | 0.000461 | 0.0104 | 8 | C3/SFRP4/CLU/LRP1/DAB2/CD63/GPC3/GAS6 |
| Fibroblast | F0 SFRP2+ Fib | GO:0002291 | T cell activation via T cell receptor contact with antigen bound to MHC molecule on antig… | 3/187 | 14/16509 | 18.92 | 0.000475 | 0.0106 | 3 | LGALS3/FGL2/ICAM1 |
| Fibroblast | F0 SFRP2+ Fib | GO:2000095 | regulation of Wnt signaling pathway, planar cell polarity pathway | 3/187 | 14/16509 | 18.92 | 0.000475 | 0.0106 | 3 | DAB2/GPC3/RSPO3 |
| Fibroblast | F0 SFRP2+ Fib | GO:0050435 | amyloid-beta metabolic process | 5/187 | 58/16509 | 7.61 | 0.000498 | 0.011 | 5 | IGF1/CLU/LRP1/SPON1/PRNP |
| Fibroblast | F0 SFRP2+ Fib | GO:0022408 | negative regulation of cell-cell adhesion | 9/187 | 203/16509 | 3.91 | 0.000519 | 0.0114 | 9 | PLA2G2A/GPNMB/CXCL12/SERPINE2/LGALS3/MDK/FGL2/ANXA1/PRNP |
| Fibroblast | F0 SFRP2+ Fib | GO:0033344 | cholesterol efflux | 5/187 | 59/16509 | 7.48 | 0.000539 | 0.0116 | 5 | ABCA8/PLTP/LRP1/NPC2/CES1 |
| Fibroblast | F0 SFRP2+ Fib | GO:0048008 | platelet-derived growth factor receptor signaling pathway | 5/187 | 59/16509 | 7.48 | 0.000539 | 0.0116 | 5 | APOD/PDGFRA/LRP1/PDGFRL/TXNIP |
| Fibroblast | F0 SFRP2+ Fib | GO:0045785 | positive regulation of cell adhesion | 15/187 | 498/16509 | 2.66 | 0.00056 | 0.012 | 15 | SFRP2/FBLN1/CCDC80/IGF1/SFRP1/CXCL12/ABI3BP/FBLN2/MDK/DAB2/S100A10/CD63/THY1/PLPP3/ANXA1 |
| Fibroblast | F0 SFRP2+ Fib | GO:0042063 | gliogenesis | 12/187 | 345/16509 | 3.07 | 0.000581 | 0.0123 | 12 | IGF1/SERPINE2/CLU/LRP1/MDK/NFIA/NFIB/VCAN/MXRA8/PLPP3/TMEM98/LEPR |
| Fibroblast | F0 SFRP2+ Fib | GO:1903019 | negative regulation of glycoprotein metabolic process | 3/187 | 15/16509 | 17.66 | 0.000589 | 0.0124 | 3 | CST3/ITM2A/ITM2B |
| Fibroblast | F0 SFRP2+ Fib | GO:0060828 | regulation of canonical Wnt signaling pathway | 10/187 | 252/16509 | 3.5 | 0.000612 | 0.0128 | 10 | SFRP2/IGFBP4/SFRP1/SFRP4/MDK/IGFBP6/DAB2/GPC3/BICC1/RSPO3 |
| Fibroblast | F0 SFRP2+ Fib | GO:0032373 | positive regulation of sterol transport | 4/187 | 35/16509 | 10.09 | 0.000634 | 0.0131 | 4 | ABCA8/PLTP/LRP1/CES1 |
| Fibroblast | F0 SFRP2+ Fib | GO:0032376 | positive regulation of cholesterol transport | 4/187 | 35/16509 | 10.09 | 0.000634 | 0.0131 | 4 | ABCA8/PLTP/LRP1/CES1 |
| Fibroblast | F0 SFRP2+ Fib | GO:0120254 | olefinic compound metabolic process | 8/187 | 167/16509 | 4.23 | 0.000639 | 0.0131 | 8 | ADH1B/ALDH1A1/DAB2/DHRS3/EPHX1/CYP1B1/AKR1C1/AKR1C3 |
| Fibroblast | F0 SFRP2+ Fib | GO:0001763 | morphogenesis of a branching structure | 9/187 | 210/16509 | 3.78 | 0.000662 | 0.0133 | 9 | SFRP2/IGF1/SFRP1/FGF7/CTSH/MDK/GPC3/SLIT2/RSPO3 |
| Fibroblast | F0 SFRP2+ Fib | GO:0060348 | bone development | 9/187 | 210/16509 | 3.78 | 0.000662 | 0.0133 | 9 | OGN/IGF1/FBLN5/SFRP4/DHRS3/LEPR/OSR2/FBN1/BNC2 |

| compartment | subcluster | GO_ID | Description | GeneRatio | BgRatio | FoldEnrichment | pvalue | p_adjust | Count | geneID |
|---|---|---|---|---|---|---|---|---|---|---|
| Fibroblast | F0 SFRP2+ Fib | GO:0022617 | extracellular matrix disassembly | 5/187 | 62/16509 | 7.12 | 0.000678 | 0.0136 | 5 | CST3/MMP2/CTSK/LRP1/FAP |
| Fibroblast | F0 SFRP2+ Fib | GO:1904018 | positive regulation of vasculature development | 8/187 | 169/16509 | 4.18 | 0.000691 | 0.0136 | 8 | C3/SFRP2/CXCL12/CD34/MDK/PTGIS/CYP1B1/HSPB6 |
| Fibroblast | F0 SFRP2+ Fib | GO:0010001 | glial cell differentiation | 10/187 | 256/16509 | 3.45 | 0.000691 | 0.0136 | 10 | IGF1/SERPINE2/CLU/LRP1/MDK/NFIA/NFIB/MXRA8/PLPP3/TMEM98 |
| Fibroblast | F0 SFRP2+ Fib | GO:0060766 | negative regulation of androgen receptor signaling pathway | 3/187 | 16/16509 | 16.55 | 0.000718 | 0.014 | 3 | IGF1/SFRP1/DAB2 |
| Fibroblast | F0 SFRP2+ Fib | GO:0061050 | regulation of cell growth involved in cardiac muscle cell development | 3/187 | 16/16509 | 16.55 | 0.000718 | 0.014 | 3 | IGF1/COL14A1/PI16 |
| Fibroblast | F0 SFRP2+ Fib | GO:1903035 | negative regulation of response to wounding | 6/187 | 95/16509 | 5.58 | 0.000735 | 0.0142 | 6 | PDGFRA/SERPINE2/MDK/SERPING1/TFPI/FAP |
| Fibroblast | F0 SFRP2+ Fib | GO:0071677 | positive regulation of mononuclear cell migration | 6/187 | 96/16509 | 5.52 | 0.000777 | 0.0147 | 6 | CXCL12/LGALS3/MDK/JAM2/RARRES2/GAS6 |
| Fibroblast | F0 SFRP2+ Fib | GO:0071383 | cellular response to steroid hormone stimulus | 9/187 | 215/16509 | 3.7 | 0.000783 | 0.0147 | 9 | SERPINF1/IGF1/SFRP1/DAB2/ANXA1/TFPI/AKR1C3/LMO3/PLPP1 |
| Fibroblast | F0 SFRP2+ Fib | GO:0010665 | regulation of cardiac muscle cell apoptotic process | 4/187 | 37/16509 | 9.54 | 0.000786 | 0.0147 | 4 | SFRP2/MDK/NUPR1/HSPB6 |
| Fibroblast | F0 SFRP2+ Fib | GO:0010874 | regulation of cholesterol efflux | 4/187 | 37/16509 | 9.54 | 0.000786 | 0.0147 | 4 | ABCA8/PLTP/LRP1/CES1 |
| Fibroblast | F0 SFRP2+ Fib | GO:1902003 | regulation of amyloid-beta formation | 4/187 | 37/16509 | 9.54 | 0.000786 | 0.0147 | 4 | IGF1/CLU/SPON1/PRNP |
| Fibroblast | F0 SFRP2+ Fib | GO:0051250 | negative regulation of lymphocyte activation | 8/187 | 174/16509 | 4.06 | 0.000835 | 0.0155 | 8 | PLA2G2A/SFRP1/GPNMB/LGALS3/MDK/FGL2/ANXA1/PRNP |
| Fibroblast | F0 SFRP2+ Fib | GO:0002693 | positive regulation of cellular extravasation | 3/187 | 17/16509 | 15.58 | 0.000865 | 0.0157 | 3 | MDK/THY1/ICAM1 |
| Fibroblast | F0 SFRP2+ Fib | GO:0010715 | regulation of extracellular matrix disassembly | 3/187 | 17/16509 | 15.58 | 0.000865 | 0.0157 | 3 | CST3/LRP1/FAP |
| Fibroblast | F0 SFRP2+ Fib | GO:1905954 | positive regulation of lipid localization | 6/187 | 98/16509 | 5.41 | 0.000866 | 0.0157 | 6 | C3/ABCA8/PLTP/LRP1/DAB2/CES1 |
| Fibroblast | F0 SFRP2+ Fib | GO:0002691 | regulation of cellular extravasation | 4/187 | 38/16509 | 9.29 | 0.000871 | 0.0157 | 4 | CXCL12/MDK/THY1/ICAM1 |
| Fibroblast | F0 SFRP2+ Fib | GO:0042325 | regulation of phosphorylation | 13/187 | 413/16509 | 2.78 | 0.000876 | 0.0157 | 13 | C3/SFRP2/FBLN1/IGF1/SFRP1/FGF7/THY1/SLIT2/THBS4/COPS8/RARRES2/FBN1/GAS6 |
| Fibroblast | F0 SFRP2+ Fib | GO:0032147 | activation of protein kinase activity | 4/187 | 39/16509 | 9.05 | 0.000962 | 0.0172 | 4 | IGF1/COPS8/FBN1/GAS6 |
| Fibroblast | F0 SFRP2+ Fib | GO:0050868 | negative regulation of T cell activation | 7/187 | 138/16509 | 4.48 | 0.000995 | 0.0176 | 7 | PLA2G2A/GPNMB/LGALS3/MDK/FGL2/ANXA1/PRNP |
| Fibroblast | F0 SFRP2+ Fib | GO:0009410 | response to xenobiotic stimulus | 13/187 | 420/16509 | 2.73 | 0.00102 | 0.0179 | 13 | SFRP2/MMP2/SFRP1/MDK/SOD2/TXNIP/AOX1/ADD3/PRNP/EPHX1/CYP1B1/AKR1C1/GAS6 |
| Fibroblast | F0 SFRP2+ Fib | GO:0042574 | retinal metabolic process | 3/187 | 18/16509 | 14.71 | 0.00103 | 0.0179 | 3 | CYP1B1/AKR1C1/AKR1C3 |
| Fibroblast | F0 SFRP2+ Fib | GO:0043691 | reverse cholesterol transport | 3/187 | 18/16509 | 14.71 | 0.00103 | 0.0179 | 3 | CLU/LRP1/CES1 |
| Fibroblast | F0 SFRP2+ Fib | GO:0042698 | ovulation cycle | 5/187 | 68/16509 | 6.49 | 0.00103 | 0.0179 | 5 | SERPINF1/MMP2/PDGFRA/MDK/SLIT2 |
| Fibroblast | F0 SFRP2+ Fib | GO:0010662 | regulation of striated muscle cell apoptotic process | 4/187 | 40/16509 | 8.83 | 0.00106 | 0.0182 | 4 | SFRP2/MDK/NUPR1/HSPB6 |
| Fibroblast | F0 SFRP2+ Fib | GO:0010721 | negative regulation of cell development | 10/187 | 271/16509 | 3.26 | 0.00107 | 0.0182 | 10 | IGF1/SFRP1/MDK/THY1/FGL2/ANXA1/TMEM176B/TMEM98/TMEM176A/FBN1 |
| Fibroblast | F0 SFRP2+ Fib | GO:0001822 | kidney development | 11/187 | 320/16509 | 3.03 | 0.00108 | 0.0182 | 11 | SERPINF1/SFRP1/CTSH/PDGFRA/CD34/GPC3/SLIT2/BICC1/BASP1/OSR2/FBN1 |
| Fibroblast | F0 SFRP2+ Fib | GO:0060541 | respiratory system development | 9/187 | 225/16509 | 3.53 | 0.00108 | 0.0182 | 9 | IGF1/FGF7/CTSH/PDGFRA/NFIB/GPC3/BASP1/HSD11B1/FBN1 |
| Fibroblast | F0 SFRP2+ Fib | GO:0009743 | response to carbohydrate | 9/187 | 226/16509 | 3.52 | 0.00111 | 0.0187 | 9 | SERPINF1/IGF1/COL6A3/CLEC12/COL6A2/SOD2/TXNIP/ICAM1/GAS6 |
| Fibroblast | F0 SFRP2+ Fib | GO:0010659 | cardiac muscle cell apoptotic process | 4/187 | 41/16509 | 8.61 | 0.00116 | 0.0192 | 4 | SFRP2/MDK/NUPR1/HSPB6 |
| Fibroblast | F0 SFRP2+ Fib | GO:0010712 | regulation of collagen metabolic process | 4/187 | 41/16509 | 8.61 | 0.00116 | 0.0192 | 4 | CST3/MFAP4/CYGB/FAP |
| Fibroblast | F0 SFRP2+ Fib | GO:0050921 | positive regulation of chemotaxis | 7/187 | 142/16509 | 4.35 | 0.00118 | 0.0192 | 7 | CXCL12/PDGFRA/MDK/SLIT2/THBS4/RARRES2/GAS6 |
| Fibroblast | F0 SFRP2+ Fib | GO:0030193 | regulation of blood coagulation | 5/187 | 70/16509 | 6.31 | 0.00118 | 0.0192 | 5 | PDGFRA/SERPINE2/SERPING1/TFPI/FAP |

| compartment | subcluster | GO_ID | Description | GeneRatio | BgRatio | FoldEnrichment | pvalue | p_adjust | Count | geneID |
|---|---|---|---|---|---|---|---|---|---|---|
| Fibroblast | F0 SFRP2+ Fib | GO:0035924 | cellular response to vascular endothelial growth factor stimulus | 5/187 | 70/16509 | 6.31 | 0.00118 | 0.0192 | 5 | DCN/TNXB/PDGFRA/CD63/ANXA1 |
| Fibroblast | F0 SFRP2+ Fib | GO:2000269 | regulation of fibroblast apoptotic process | 3/187 | 19/16509 | 13.94 | 0.00121 | 0.0196 | 3 | SFRP1/NUPR1/GAS6 |
| Fibroblast | F0 SFRP2+ Fib | GO:0150063 | visual system development | 13/187 | 429/16509 | 2.68 | 0.00124 | 0.0199 | 13 | C3/SERPINF1/EFEMP1/PDGFRA/CHRDL1/NFIA/NFIB/THY1/OSR2/FBN1/CYP1B1/MITF/SAMD11 |
| Fibroblast | F0 SFRP2+ Fib | GO:0030301 | cholesterol transport | 6/187 | 105/16509 | 5.04 | 0.00124 | 0.0199 | 6 | ABCA8/PLTP/CLU/LRP1/NPC2/CES1 |
| Fibroblast | F0 SFRP2+ Fib | GO:0061045 | negative regulation of wound healing | 5/187 | 71/16509 | 6.22 | 0.00126 | 0.02 | 5 | PDGFRA/SERPINE2/SERPING1/TFPI/FAP |
| Fibroblast | F0 SFRP2+ Fib | GO:0045123 | cellular extravasation | 5/187 | 72/16509 | 6.13 | 0.00134 | 0.021 | 5 | CXCL12/MDK/THY1/JAM2/ICAM1 |
| Fibroblast | F0 SFRP2+ Fib | GO:1900046 | regulation of hemostasis | 5/187 | 72/16509 | 6.13 | 0.00134 | 0.021 | 5 | PDGFRA/SERPINE2/SERPING1/TFPI/FAP |
| Fibroblast | F0 SFRP2+ Fib | GO:0030162 | regulation of proteolysis | 12/187 | 380/16509 | 2.79 | 0.00134 | 0.021 | 12 | CST3/GSN/SERPINE2/CLU/SPON1/DAB2/S100A10/TIMP2/TMEM98/NUPR1/PRNP/SH3D19 |
| Fibroblast | F0 SFRP2+ Fib | GO:0048246 | macrophage chemotaxis | 4/187 | 43/16509 | 8.21 | 0.00139 | 0.0216 | 4 | MMP2/LGALS3/MDK/RARRES2 |
| Fibroblast | F0 SFRP2+ Fib | GO:0006809 | nitric oxide biosynthetic process | 5/187 | 73/16509 | 6.05 | 0.00142 | 0.0216 | 5 | IGF1/CLU/SOD2/PTGIS/CYP1B1 |
| Fibroblast | F0 SFRP2+ Fib | GO:0033627 | cell adhesion mediated by integrin | 5/187 | 73/16509 | 6.05 | 0.00142 | 0.0216 | 5 | SFRP2/PLPP3/FBN1/ICAM1/CYP1B1 |
| Fibroblast | F0 SFRP2+ Fib | GO:0042246 | tissue regeneration | 5/187 | 73/16509 | 6.05 | 0.00142 | 0.0216 | 5 | IGF1/APOD/MDK/ANXA1/CPQ |
| Fibroblast | F0 SFRP2+ Fib | GO:0042060 | wound healing | 13/187 | 436/16509 | 2.63 | 0.00143 | 0.0216 | 13 | FBLN1/IGF1/PDGFRA/SERPINE2/CD34/SERPING1/S100A10/ANXA1/ANGPTL2/TFPI/GAS6/SVEP1/FAP |
| Fibroblast | F0 SFRP2+ Fib | GO:0048880 | sensory system development | 13/187 | 436/16509 | 2.63 | 0.00143 | 0.0216 | 13 | C3/SERPINF1/EFEMP1/PDGFRA/CHRDL1/NFIA/NFIB/THY1/OSR2/FBN1/CYP1B1/MITF/SAMD11 |
| Fibroblast | F0 SFRP2+ Fib | GO:0010658 | striated muscle cell apoptotic process | 4/187 | 44/16509 | 8.03 | 0.00152 | 0.0224 | 4 | SFRP2/MDK/NUPR1/HSPB6 |
| Fibroblast | F0 SFRP2+ Fib | GO:0043277 | apoptotic cell clearance | 4/187 | 44/16509 | 8.03 | 0.00152 | 0.0224 | 4 | C3/LRP1/ANXA1/GAS6 |
| Fibroblast | F0 SFRP2+ Fib | GO:1902991 | regulation of amyloid precursor protein catabolic process | 4/187 | 44/16509 | 8.03 | 0.00152 | 0.0224 | 4 | IGF1/CLU/SPON1/PRNP |
| Fibroblast | F0 SFRP2+ Fib | GO:1903018 | regulation of glycoprotein metabolic process | 4/187 | 44/16509 | 8.03 | 0.00152 | 0.0224 | 4 | IGF1/CST3/ITM2A/ITM2B |
| Fibroblast | F0 SFRP2+ Fib | GO:0000302 | response to reactive oxygen species | 8/187 | 191/16509 | 3.7 | 0.00152 | 0.0224 | 8 | MMP2/APOD/FBLN5/PDGFRA/SOD2/TXNIP/CYP1B1/CYGB |
| Fibroblast | F0 SFRP2+ Fib | GO:0006081 | aldehyde metabolic process | 5/187 | 75/16509 | 5.89 | 0.00161 | 0.0234 | 5 | ALDH1A1/DAB2/CYP1B1/AKR1C1/AKR1C3 |
| Fibroblast | F0 SFRP2+ Fib | GO:0050818 | regulation of coagulation | 5/187 | 75/16509 | 5.89 | 0.00161 | 0.0234 | 5 | PDGFRA/SERPINE2/SERPING1/TFPI/FAP |
| Fibroblast | F0 SFRP2+ Fib | GO:0010955 | negative regulation of protein processing | 3/187 | 21/16509 | 12.61 | 0.00164 | 0.0234 | 3 | SERPINE2/TMEM98/PRNP |
| Fibroblast | F0 SFRP2+ Fib | GO:0019430 | removal of superoxide radicals | 3/187 | 21/16509 | 12.61 | 0.00164 | 0.0234 | 3 | FBLN5/SOD2/CYGB |
| Fibroblast | F0 SFRP2+ Fib | GO:0071450 | cellular response to oxygen radical | 3/187 | 21/16509 | 12.61 | 0.00164 | 0.0234 | 3 | FBLN5/SOD2/CYGB |
| Fibroblast | F0 SFRP2+ Fib | GO:0071451 | cellular response to superoxide | 3/187 | 21/16509 | 12.61 | 0.00164 | 0.0234 | 3 | FBLN5/SOD2/CYGB |
| Fibroblast | F0 SFRP2+ Fib | GO:0006693 | prostaglandin metabolic process | 4/187 | 45/16509 | 7.85 | 0.00165 | 0.0234 | 4 | PTGDS/PTGIS/AKR1C1/AKR1C3 |
| Fibroblast | F0 SFRP2+ Fib | GO:0034205 | amyloid-beta formation | 4/187 | 45/16509 | 7.85 | 0.00165 | 0.0234 | 4 | IGF1/CLU/SPON1/PRNP |
| Fibroblast | F0 SFRP2+ Fib | GO:0031960 | response to corticosteroid | 7/187 | 151/16509 | 4.09 | 0.00168 | 0.0236 | 7 | SERPINF1/PTGDS/MDK/SLIT2/ANXA1/CDO1/AKR1C3 |
| Fibroblast | F0 SFRP2+ Fib | GO:0007596 | blood coagulation | 9/187 | 241/16509 | 3.3 | 0.00174 | 0.0243 | 9 | FBLN1/PDGFRA/SERPINE2/SERPING1/ANGPTL2/TFPI/GAS6/SVEP1/FAP |
| Fibroblast | F0 SFRP2+ Fib | GO:0002920 | regulation of humoral immune response | 4/187 | 46/16509 | 7.68 | 0.00179 | 0.025 | 4 | C3/CLU/SERPING1/CFH |
| Fibroblast | F0 SFRP2+ Fib | GO:0030324 | lung development | 8/187 | 197/16509 | 3.59 | 0.00185 | 0.0256 | 8 | IGF1/FGF7/CTSH/PDGFRA/NFIB/GPC3/HSD11B1/FBN1 |
| Fibroblast | F0 SFRP2+ Fib | GO:1905146 | lysosomal protein catabolic process | 3/187 | 22/16509 | 12.04 | 0.00188 | 0.0258 | 3 | CTSH/LRP1/PLBD1 |

| compartment | subcluster | GO_ID | Description | GeneRatio | BgRatio | FoldEnrichment | pvalue | p_adjust | Count | geneID |
|---|---|---|---|---|---|---|---|---|---|---|
| Fibroblast | F0 SFRP2+ Fib | GO:0015918 | sterol transport | 6/187 | 114/16509 | 4.65 | 0.00189 | 0.0258 | 6 | ABCA8/PLTP/CLU/LRP1/NPC2/CES1 |
| Fibroblast | F0 SFRP2+ Fib | GO:0033559 | unsaturated fatty acid metabolic process | 6/187 | 114/16509 | 4.65 | 0.00189 | 0.0258 | 6 | PTGDS/PTGIS/EPHX1/CYP1B1/AKR1C1/AKR1C3 |
| Fibroblast | F0 SFRP2+ Fib | GO:0050817 | coagulation | 9/187 | 245/16509 | 3.24 | 0.00194 | 0.0261 | 9 | FBLN1/PDGFRA/SERPINE2/SERPING1/ANGPTL2/TFPI/GAS6/SVEP1/FAP |
| Fibroblast | F0 SFRP2+ Fib | GO:0006692 | prostanoid metabolic process | 4/187 | 47/16509 | 7.51 | 0.00194 | 0.0261 | 4 | PTGDS/PTGIS/AKR1C1/AKR1C3 |
| Fibroblast | F0 SFRP2+ Fib | GO:0060071 | Wnt signaling pathway, planar cell polarity pathway | 4/187 | 47/16509 | 7.51 | 0.00194 | 0.0261 | 4 | SFRP1/DAB2/GPC3/RSPO3 |
| Fibroblast | F0 SFRP2+ Fib | GO:1902041 | regulation of extrinsic apoptotic signaling pathway via death domain receptors | 4/187 | 47/16509 | 7.51 | 0.00194 | 0.0261 | 4 | SFRP2/SFRP1/LGALS3/ICAM1 |
| Fibroblast | F0 SFRP2+ Fib | GO:0016055 | Wnt signaling pathway | 13/187 | 453/16509 | 2.53 | 0.002 | 0.0267 | 13 | SFRP2/IGFBP4/SFRP1/SFRP4/LRP1/MDK/IGFBP6/DAB2/GPC3/PLPP3/BICC1/RSPO3/MITF |
| Fibroblast | F0 SFRP2+ Fib | GO:0002695 | negative regulation of leukocyte activation | 8/187 | 200/16509 | 3.53 | 0.00203 | 0.0269 | 8 | PLA2G2A/SFRP1/GPNMB/LGALS3/MDK/FGL2/ANXA1/PRNP |
| Fibroblast | F0 SFRP2+ Fib | GO:0030323 | respiratory tube development | 8/187 | 201/16509 | 3.51 | 0.00209 | 0.0274 | 8 | IGF1/FGF7/CTSH/PDGFRA/NFIB/GPC3/HSD11B1/FBN1 |
| Fibroblast | F0 SFRP2+ Fib | GO:0038084 | vascular endothelial growth factor signaling pathway | 4/187 | 48/16509 | 7.36 | 0.0021 | 0.0274 | 4 | DCN/TNXB/PDGFRA/CD63 |
| Fibroblast | F0 SFRP2+ Fib | GO:0007599 | hemostasis | 9/187 | 248/16509 | 3.2 | 0.00211 | 0.0274 | 9 | FBLN1/PDGFRA/SERPINE2/SERPING1/ANGPTL2/TFPI/GAS6/SVEP1/FAP |
| Fibroblast | F0 SFRP2+ Fib | GO:0003298 | physiological muscle hypertrophy | 3/187 | 23/16509 | 11.52 | 0.00214 | 0.0274 | 3 | IGF1/COL14A1/PI16 |
| Fibroblast | F0 SFRP2+ Fib | GO:0003301 | physiological cardiac muscle hypertrophy | 3/187 | 23/16509 | 11.52 | 0.00214 | 0.0274 | 3 | IGF1/COL14A1/PI16 |
| Fibroblast | F0 SFRP2+ Fib | GO:0061049 | cell growth involved in cardiac muscle cell development | 3/187 | 23/16509 | 11.52 | 0.00214 | 0.0274 | 3 | IGF1/COL14A1/PI16 |
| Fibroblast | F0 SFRP2+ Fib | GO:1903318 | negative regulation of protein maturation | 3/187 | 23/16509 | 11.52 | 0.00214 | 0.0274 | 3 | SERPINE2/TMEM98/PRNP |
| Fibroblast | F0 SFRP2+ Fib | GO:0071621 | granulocyte chemotaxis | 6/187 | 117/16509 | 4.53 | 0.00216 | 0.0274 | 6 | LGALS3/MDK/SLIT2/ANXA1/THBS4/RARRES2 |
| Fibroblast | F0 SFRP2+ Fib | GO:0034284 | response to monosaccharide | 8/187 | 202/16509 | 3.5 | 0.00216 | 0.0274 | 8 | SERPINF1/IGF1/COL6A3/COL6A2/SOD2/TXNIP/ICAM1/GAS6 |
| Fibroblast | F0 SFRP2+ Fib | GO:0141193 | nuclear receptor-mediated signaling pathway | 8/187 | 202/16509 | 3.5 | 0.00216 | 0.0274 | 8 | IGF1/SFRP1/DAB2/PTGIS/DHRS3/AKR1C3/LMO3/PLPP1 |
| Fibroblast | F0 SFRP2+ Fib | GO:0001818 | negative regulation of cytokine production | 10/187 | 301/16509 | 2.93 | 0.00232 | 0.0291 | 10 | FBLN1/IGF1/APOD/GPNMB/CD34/ANXA1/C1QTNF3/PRNP/CHID1/GAS6 |
| Fibroblast | F0 SFRP2+ Fib | GO:0070371 | ERK1 and ERK2 cascade | 10/187 | 301/16509 | 2.93 | 0.00232 | 0.0291 | 10 | FBLN1/IGF1/PLA2G2A/GPNMB/CTSH/PDGFRA/DAB2/ICAM1/GAS6/LMO3 |
| Fibroblast | F0 SFRP2+ Fib | GO:0051402 | neuron apoptotic process | 10/187 | 302/16509 | 2.92 | 0.00237 | 0.0295 | 10 | MMP2/COL6A3/CLU/MDK/COL6A2/SOD2/NGFR/NUPR1/PRNP/TMBIM4 |
| Fibroblast | F0 SFRP2+ Fib | GO:0008625 | extrinsic apoptotic signaling pathway via death domain receptors | 5/187 | 82/16509 | 5.38 | 0.00238 | 0.0295 | 5 | SFRP2/SFRP1/LGALS3/TNFSF10/ICAM1 |
| Fibroblast | F0 SFRP2+ Fib | GO:0032370 | positive regulation of lipid transport | 5/187 | 82/16509 | 5.38 | 0.00238 | 0.0295 | 5 | ABCA8/PLTP/LRP1/DAB2/CES1 |
| Fibroblast | F0 SFRP2+ Fib | GO:0000303 | response to superoxide | 3/187 | 24/16509 | 11.04 | 0.00243 | 0.0298 | 3 | FBLN5/SOD2/CYGB |
| Fibroblast | F0 SFRP2+ Fib | GO:0010954 | positive regulation of protein processing | 3/187 | 24/16509 | 11.04 | 0.00243 | 0.0298 | 3 | GSN/SPON1/S100A10 |
| Fibroblast | F0 SFRP2+ Fib | GO:0002688 | regulation of leukocyte chemotaxis | 6/187 | 120/16509 | 4.41 | 0.00245 | 0.03 | 6 | CXCL12/MDK/SLIT2/THBS4/RARRES2/GAS6 |
| Fibroblast | F0 SFRP2+ Fib | GO:2001236 | regulation of extrinsic apoptotic signaling pathway | 7/187 | 162/16509 | 3.81 | 0.0025 | 0.0304 | 7 | SFRP2/IGF1/SRPX/SFRP1/LGALS3/TNFSF10/ICAM1 |
| Fibroblast | F0 SFRP2+ Fib | GO:2000027 | regulation of animal organ morphogenesis | 5/187 | 83/16509 | 5.32 | 0.00251 | 0.0305 | 5 | SFRP2/SFRP1/FGF7/CD34/NGFR |
| Fibroblast | F0 SFRP2+ Fib | GO:0018108 | peptidyl-tyrosine phosphorylation | 6/187 | 121/16509 | 4.38 | 0.00255 | 0.0308 | 6 | SFRP2/EFEMP1/SFRP1/FGF7/PDGFRA/THBS4 |
| Fibroblast | F0 SFRP2+ Fib | GO:0002548 | monocyte chemotaxis | 4/187 | 51/16509 | 6.92 | 0.00263 | 0.0312 | 4 | CXCL12/LGALS3/SLIT2/ANXA1 |
| Fibroblast | F0 SFRP2+ Fib | GO:0030521 | androgen receptor signaling pathway | 4/187 | 51/16509 | 6.92 | 0.00263 | 0.0312 | 4 | IGF1/SFRP1/DAB2/PLPP1 |

| compartment | subcluster | GO_ID | Description | GeneRatio | BgRatio | FoldEnrichment | pvalue | p_adjust | Count | geneID |
|---|---|---|---|---|---|---|---|---|---|---|
| Fibroblast | F0 SFRP2+ Fib | GO:0048009 | insulin-like growth factor receptor signaling pathway | 4/187 | 51/16509 | 6.92 | 0.00263 | 0.0312 | 4 | IGF1/IGFBP4/CILP/IGFBP6 |
| Fibroblast | F0 SFRP2+ Fib | GO:0071622 | regulation of granulocyte chemotaxis | 4/187 | 51/16509 | 6.92 | 0.00263 | 0.0312 | 4 | MDK/SLIT2/THBS4/RARRES2 |
| Fibroblast | F0 SFRP2+ Fib | GO:0018212 | peptidyl-tyrosine modification | 6/187 | 122/16509 | 4.34 | 0.00266 | 0.0312 | 6 | SFRP2/EFEMP1/SFRP1/FGF7/PDGFRA/THBS4 |
| Fibroblast | F0 SFRP2+ Fib | GO:0060070 | canonical Wnt signaling pathway | 10/187 | 307/16509 | 2.88 | 0.00267 | 0.0312 | 10 | SFRP2/IGFBP4/SFRP1/SFRP4/MDK/IGFBP6/DAB2/GPC3/BICC1/RSPO3 |
| Fibroblast | F0 SFRP2+ Fib | GO:0010563 | negative regulation of phosphorus metabolic process | 7/187 | 164/16509 | 3.77 | 0.00267 | 0.0312 | 7 | SFRP2/FBLN1/SFRP1/PID1/THY1/SLIT2/NUPR1 |
| Fibroblast | F0 SFRP2+ Fib | GO:0045936 | negative regulation of phosphate metabolic process | 7/187 | 164/16509 | 3.77 | 0.00267 | 0.0312 | 7 | SFRP2/FBLN1/SFRP1/PID1/THY1/SLIT2/NUPR1 |
| Fibroblast | F0 SFRP2+ Fib | GO:0043603 | amide metabolic process | 13/187 | 469/16509 | 2.45 | 0.00271 | 0.0314 | 13 | IGF1/CTSH/ABCA8/CLU/LRP1/GGT5/SPON1/SOD2/GSTM5/NNMT/PLPP3/PRNP/PLPP1 |
| Fibroblast | F0 SFRP2+ Fib | GO:0000305 | response to oxygen radical | 3/187 | 25/16509 | 10.59 | 0.00274 | 0.0314 | 3 | FBLN5/SOD2/CYGB |
| Fibroblast | F0 SFRP2+ Fib | GO:0044346 | fibroblast apoptotic process | 3/187 | 25/16509 | 10.59 | 0.00274 | 0.0314 | 3 | SFRP1/NUPR1/GAS6 |
| Fibroblast | F0 SFRP2+ Fib | GO:1903319 | positive regulation of protein maturation | 3/187 | 25/16509 | 10.59 | 0.00274 | 0.0314 | 3 | GSN/SPON1/S100A10 |
| Fibroblast | F0 SFRP2+ Fib | GO:0045766 | positive regulation of angiogenesis | 7/187 | 165/16509 | 3.75 | 0.00277 | 0.0316 | 7 | C3/SFRP2/CD34/MDK/PTGIS/CYP1B1/HSPB6 |
| Fibroblast | F0 SFRP2+ Fib | GO:0030308 | negative regulation of cell growth | 7/187 | 166/16509 | 3.72 | 0.00286 | 0.0324 | 7 | SFRP2/SFRP1/SERPINE2/DAB2/MEG3/SLIT2/PI16 |
| Fibroblast | F0 SFRP2+ Fib | GO:1903034 | regulation of response to wounding | 7/187 | 166/16509 | 3.72 | 0.00286 | 0.0324 | 7 | PDGFRA/SERPINE2/MDK/SERPING1/ANXA1/TFPI/FAP |
| Fibroblast | F0 SFRP2+ Fib | GO:0007159 | leukocyte cell-cell adhesion | 12/187 | 417/16509 | 2.54 | 0.00289 | 0.0327 | 12 | IGF1/PLA2G2A/GPNMB/CXCL12/LGALS3/MDK/THY1/FGL2/ANXA1/JAM2/PRNP/ICAM1 |
| Fibroblast | F0 SFRP2+ Fib | GO:0006706 | steroid catabolic process | 3/187 | 26/16509 | 10.19 | 0.00307 | 0.0343 | 3 | HSD11B1/HSD17B11/CYP1B1 |
| Fibroblast | F0 SFRP2+ Fib | GO:0007039 | protein catabolic process in the vacuole | 3/187 | 26/16509 | 10.19 | 0.00307 | 0.0343 | 3 | CTSH/LRP1/PLBD1 |
| Fibroblast | F0 SFRP2+ Fib | GO:0044342 | type B pancreatic cell proliferation | 3/187 | 26/16509 | 10.19 | 0.00307 | 0.0343 | 3 | IGF1/IGFBP4/NUPR1 |
| Fibroblast | F0 SFRP2+ Fib | GO:0046849 | bone remodeling | 5/187 | 87/16509 | 5.07 | 0.00309 | 0.0343 | 5 | SFRP1/CTSK/MDK/LEPR/MITF |
| Fibroblast | F0 SFRP2+ Fib | GO:0008037 | cell recognition | 7/187 | 169/16509 | 3.66 | 0.00316 | 0.035 | 7 | C3/C1S/C1R/LGALS3/COLEC12/SPON2/VCAN |
| Fibroblast | F0 SFRP2+ Fib | GO:0015850 | organic hydroxy compound transport | 9/187 | 264/16509 | 3.01 | 0.0032 | 0.0353 | 9 | CXCL12/ABCA8/PLTP/CLU/LRP1/NPC2/DAB2/AKR1C1/CES1 |
| Fibroblast | F0 SFRP2+ Fib | GO:0045926 | negative regulation of growth | 8/187 | 217/16509 | 3.25 | 0.00335 | 0.0368 | 8 | SFRP2/SFRP1/SERPINE2/DAB2/GPC3/MEG3/SLIT2/PI16 |
| Fibroblast | F0 SFRP2+ Fib | GO:0001654 | eye development | 12/187 | 425/16509 | 2.49 | 0.00337 | 0.0368 | 12 | SERPINF1/EFEMP1/PDGFRA/CHRDL1/NFIA/NFIB/THY1/OSR2/FBN1/CYP1B1/MITF/SAMD11 |
| Fibroblast | F0 SFRP2+ Fib | GO:0022612 | gland morphogenesis | 6/187 | 128/16509 | 4.14 | 0.00338 | 0.0368 | 6 | IGF1/MMP2/SFRP1/FGF7/MDK/NFIB |
| Fibroblast | F0 SFRP2+ Fib | GO:0016048 | detection of temperature stimulus | 3/187 | 27/16509 | 9.81 | 0.00342 | 0.0372 | 3 | CXCL12/LXN/NGFR |
| Fibroblast | F0 SFRP2+ Fib | GO:0031102 | neuron projection regeneration | 4/187 | 55/16509 | 6.42 | 0.00347 | 0.0373 | 4 | MMP2/APOD/THY1/OMG |
| Fibroblast | F0 SFRP2+ Fib | GO:1904035 | regulation of epithelial cell apoptotic process | 4/187 | 55/16509 | 6.42 | 0.00347 | 0.0373 | 4 | GSN/SFRP4/MDK/NUPR1 |
| Fibroblast | F0 SFRP2+ Fib | GO:0061041 | regulation of wound healing | 6/187 | 129/16509 | 4.11 | 0.00351 | 0.0376 | 6 | PDGFRA/SERPINE2/SERPING1/ANXA1/TFPI/FAP |
| Fibroblast | F0 SFRP2+ Fib | GO:0002263 | cell activation involved in immune response | 10/187 | 319/16509 | 2.77 | 0.00351 | 0.0376 | 10 | ITM2A/S100A13/PTGDS/LRP1/LGALS3/MDK/FGL2/ANXA1/TMEM98/ICAM1 |
| Fibroblast | F0 SFRP2+ Fib | GO:1905475 | regulation of protein localization to membrane | 7/187 | 173/16509 | 3.57 | 0.0036 | 0.0383 | 7 | GSN/LRP1/PID1/LGALS3/DAB2/GPC3/PRNP |
| Fibroblast | F0 SFRP2+ Fib | GO:0043010 | camera-type eye development | 11/187 | 374/16509 | 2.6 | 0.00364 | 0.0385 | 11 | SERPINF1/EFEMP1/PDGFRA/NFIA/NFIB/THY1/OSR2/FBN1/CYP1B1/MITF/SAMD11 |
| Fibroblast | F0 SFRP2+ Fib | GO:0051384 | response to glucocorticoid | 6/187 | 130/16509 | 4.07 | 0.00365 | 0.0385 | 6 | SERPINF1/PTGDS/MDK/SLIT2/ANXA1/CDO1 |
| Fibroblast | F0 SFRP2+ Fib | GO:0046456 | icosanoid biosynthetic process | 4/187 | 56/16509 | 6.31 | 0.0037 | 0.0388 | 4 | PTGDS/GGT5/PTGIS/AKR1C3 |

| compartment | subcluster | GO_ID | Description | GeneRatio | BgRatio | FoldEnrichment | pvalue | p_adjust | Count | geneID |
|---|---|---|---|---|---|---|---|---|---|---|
| Fibroblast | F0 SFRP2+ Fib | GO:2000351 | regulation of endothelial cell apoptotic process | 4/187 | 56/16509 | 6.31 | 0.0037 | 0.0388 | 4 | NGFR/ICAM1/AKR1C3/GAS6 |
| Fibroblast | F0 SFRP2+ Fib | GO:0031579 | membrane raft organization | 3/187 | 28/16509 | 9.46 | 0.0038 | 0.0396 | 3 | GSN/COLEC12/S100A10 |
| Fibroblast | F0 SFRP2+ Fib | GO:0060740 | prostate gland epithelium morphogenesis | 3/187 | 28/16509 | 9.46 | 0.0038 | 0.0396 | 3 | IGF1/MMP2/SFRP1 |
| Fibroblast | F0 SFRP2+ Fib | GO:0001836 | release of cytochrome c from mitochondria | 4/187 | 57/16509 | 6.2 | 0.00394 | 0.0409 | 4 | IGF1/CLU/SOD2/TNFSF10 |
| Fibroblast | F0 SFRP2+ Fib | GO:0001934 | positive regulation of protein phosphorylation | 8/187 | 223/16509 | 3.17 | 0.00395 | 0.0409 | 8 | C3/IGF1/FGF7/THBS4/COPS8/RARRES2/FBN1/GAS6 |
| Fibroblast | F0 SFRP2+ Fib | GO:0097529 | myeloid leukocyte migration | 8/187 | 224/16509 | 3.15 | 0.00406 | 0.0417 | 8 | MMP2/CXCL12/LGALS3/MDK/SLIT2/ANXA1/THBS4/RARRES2 |
| Fibroblast | F0 SFRP2+ Fib | GO:0048660 | regulation of smooth muscle cell proliferation | 6/187 | 133/16509 | 3.98 | 0.00408 | 0.0417 | 6 | OGN/IGF1/MMP2/APOD/SOD2/ELN |
| Fibroblast | F0 SFRP2+ Fib | GO:0070372 | regulation of ERK1 and ERK2 cascade | 9/187 | 274/16509 | 2.9 | 0.00409 | 0.0417 | 9 | FBLN1/IGF1/PLA2G2A/GPNMB/PDGFRA/DAB2/ICAM1/GAS6/LMO3 |
| Fibroblast | F0 SFRP2+ Fib | GO:0062012 | regulation of small molecule metabolic process | 10/187 | 326/16509 | 2.71 | 0.00409 | 0.0417 | 10 | IGF1/IGFBP4/PID1/DAB2/NNMT/C1QTNF3/LEPR/NUPR1/AKR1C3/CES1 |
| Fibroblast | F0 SFRP2+ Fib | GO:0014009 | glial cell proliferation | 4/187 | 58/16509 | 6.09 | 0.0042 | 0.0421 | 4 | CLU/NFIA/NFIB/LEPR |
| Fibroblast | F0 SFRP2+ Fib | GO:0032371 | regulation of sterol transport | 4/187 | 58/16509 | 6.09 | 0.0042 | 0.0421 | 4 | ABCA8/PLTP/LRP1/CES1 |
| Fibroblast | F0 SFRP2+ Fib | GO:0032374 | regulation of cholesterol transport | 4/187 | 58/16509 | 6.09 | 0.0042 | 0.0421 | 4 | ABCA8/PLTP/LRP1/CES1 |
| Fibroblast | F0 SFRP2+ Fib | GO:0042987 | amyloid precursor protein catabolic process | 4/187 | 58/16509 | 6.09 | 0.0042 | 0.0421 | 4 | IGF1/CLU/SPON1/PRNP |
| Fibroblast | F0 SFRP2+ Fib | GO:0046457 | prostanoid biosynthetic process | 3/187 | 29/16509 | 9.13 | 0.00421 | 0.0421 | 3 | PTGDS/PTGIS/AKR1C3 |
| Fibroblast | F0 SFRP2+ Fib | GO:0008202 | steroid metabolic process | 10/187 | 328/16509 | 2.69 | 0.00427 | 0.0426 | 10 | PDGFRA/NPC2/DAB2/LEPR/HSD11B1/HSD17B11/CYP1B1/AKR1C1/AKR1C3/CES1 |
| Fibroblast | F0 SFRP2+ Fib | GO:0002690 | positive regulation of leukocyte chemotaxis | 5/187 | 94/16509 | 4.7 | 0.0043 | 0.0427 | 5 | CXCL12/MDK/THBS4/RARRES2/GAS6 |
| Fibroblast | F0 SFRP2+ Fib | GO:0060420 | regulation of heart growth | 4/187 | 59/16509 | 5.99 | 0.00447 | 0.0437 | 4 | IGF1/COL14A1/PI16/BASP1 |
| Fibroblast | F0 SFRP2+ Fib | GO:1900024 | regulation of substrate adhesion-dependent cell spreading | 4/187 | 59/16509 | 5.99 | 0.00447 | 0.0437 | 4 | FBLN1/MDK/DAB2/S100A10 |
| Fibroblast | F0 SFRP2+ Fib | GO:1903053 | regulation of extracellular matrix organization | 4/187 | 59/16509 | 5.99 | 0.00447 | 0.0437 | 4 | CST3/TNXB/LRP1/FAP |
| Fibroblast | F0 SFRP2+ Fib | GO:0060041 | retina development in camera-type eye | 7/187 | 180/16509 | 3.43 | 0.00447 | 0.0437 | 7 | SERPINF1/PDGFRA/NFIA/NFIB/THY1/CYP1B1/SAMD11 |
| Fibroblast | F0 SFRP2+ Fib | GO:0043542 | endothelial cell migration | 8/187 | 228/16509 | 3.1 | 0.00452 | 0.0441 | 8 | SERPINF1/DCN/SLIT2/PLPP3/ANXA1/FSTL1/CYP1B1/FAP |
| Fibroblast | F0 SFRP2+ Fib | GO:0048659 | smooth muscle cell proliferation | 6/187 | 136/16509 | 3.89 | 0.00455 | 0.0442 | 6 | OGN/IGF1/MMP2/APOD/SOD2/ELN |
| Fibroblast | F0 SFRP2+ Fib | GO:0050860 | negative regulation of T cell receptor signaling pathway | 3/187 | 30/16509 | 8.83 | 0.00464 | 0.0443 | 3 | LGALS3/THY1/PRNP |
| Fibroblast | F0 SFRP2+ Fib | GO:0051894 | positive regulation of focal adhesion assembly | 3/187 | 30/16509 | 8.83 | 0.00464 | 0.0443 | 3 | SFRP1/S100A10/THY1 |
| Fibroblast | F0 SFRP2+ Fib | GO:0060512 | prostate gland morphogenesis | 3/187 | 30/16509 | 8.83 | 0.00464 | 0.0443 | 3 | IGF1/MMP2/SFRP1 |
| Fibroblast | F0 SFRP2+ Fib | GO:1900746 | regulation of vascular endothelial growth factor signaling pathway | 3/187 | 30/16509 | 8.83 | 0.00464 | 0.0443 | 3 | DCN/TNXB/CD63 |
| Fibroblast | F0 SFRP2+ Fib | GO:1902230 | negative regulation of intrinsic apoptotic signaling pathway in response to DNA damage | 3/187 | 30/16509 | 8.83 | 0.00464 | 0.0443 | 3 | SFRP2/CXCL12/CLU |
| Fibroblast | F0 SFRP2+ Fib | GO:0001954 | positive regulation of cell-matrix adhesion | 4/187 | 60/16509 | 5.89 | 0.00474 | 0.0449 | 4 | SFRP1/S100A10/THY1/PLPP3 |
| Fibroblast | F0 SFRP2+ Fib | GO:0045428 | regulation of nitric oxide biosynthetic process | 4/187 | 60/16509 | 5.89 | 0.00474 | 0.0449 | 4 | IGF1/CLU/SOD2/PTGIS |
| Fibroblast | F0 SFRP2+ Fib | GO:1905517 | macrophage migration | 4/187 | 60/16509 | 5.89 | 0.00474 | 0.0449 | 4 | MMP2/LGALS3/MDK/RARRES2 |
| Fibroblast | F0 SFRP2+ Fib | GO:0010906 | regulation of glucose metabolic process | 5/187 | 97/16509 | 4.55 | 0.00492 | 0.0464 | 5 | IGF1/IGFBP4/NNMT/C1QTNF3/LEPR |
| Fibroblast | F0 SFRP2+ Fib | GO:0097191 | extrinsic apoptotic signaling pathway | 8/187 | 232/16509 | 3.04 | 0.00501 | 0.047 | 8 | SFRP2/IGF1/SRPX/SFRP1/LGALS3/TNFSF10/NGFR/ICAM1 |
| Fibroblast | F0 SFRP2+ Fib | GO:0045620 | negative regulation of lymphocyte differentiation | 4/187 | 61/16509 | 5.79 | 0.00503 | 0.047 | 4 | SFRP1/MDK/FGL2/ANXA1 |
| Fibroblast | F0 SFRP2+ Fib | GO:0060425 | lung morphogenesis | 4/187 | 61/16509 | 5.79 | 0.00503 | 0.047 | 4 | IGF1/FGF7/CTSH/NFIB |

| compartment | subcluster | GO_ID | Description | GeneRatio | BgRatio | FoldEnrichment | pvalue | p_adjust | Count | geneID |
|---|---|---|---|---|---|---|---|---|---|---|
| Fibroblast | F0 SFRP2+ Fib | GO:0016042 | lipid catabolic process | 10/187 | 336/16509 | 2.63 | 0.00505 | 0.047 | 10 | PLA2G2A/PLBD1/PLPP3/HSD11B1/RARRES2/HSD17B11/CYP1B1/AKR1C3/PLPP1/CES1 |
| Fibroblast | F0 SFRP2+ Fib | GO:0090022 | regulation of neutrophil chemotaxis | 3/187 | 31/16509 | 8.54 | 0.00509 | 0.0472 | 3 | MDK/SLIT2/THBS4 |
| Fibroblast | F0 SFRP2+ Fib | GO:0001558 | regulation of cell growth | 11/187 | 392/16509 | 2.48 | 0.00516 | 0.0477 | 11 | SFRP2/IGF1/IGFBP4/SFRP1/CXCL12/SERPINE2/DAB2/COL14A1/MEG3/SLIT2/PI16 |
| Fibroblast | F0 SFRP2+ Fib | GO:2000377 | regulation of reactive oxygen species metabolic process | 6/187 | 140/16509 | 3.78 | 0.00523 | 0.0477 | 6 | FBLN5/PID1/SOD2/CYP1B1/AKR1C1/AKR1C3 |
| Fibroblast | F0 SFRP2+ Fib | GO:0014910 | regulation of smooth muscle cell migration | 4/187 | 62/16509 | 5.7 | 0.00533 | 0.0477 | 4 | IGF1/LRP1/MDK/SLIT2 |
| Fibroblast | F0 SFRP2+ Fib | GO:0072577 | endothelial cell apoptotic process | 4/187 | 62/16509 | 5.7 | 0.00533 | 0.0477 | 4 | NGFR/ICAM1/AKR1C3/GAS6 |
| Fibroblast | F0 SFRP2+ Fib | GO:0032102 | negative regulation of response to external stimulus | 11/187 | 394/16509 | 2.46 | 0.00536 | 0.0477 | 11 | IGF1/PDGFRA/SERPINE2/MDK/SERPING1/PTGIS/FGL2/SLIT2/C1QTNF3/TFPI/FAP |
| Fibroblast | F0 SFRP2+ Fib | GO:0048732 | gland development | 12/187 | 451/16509 | 2.35 | 0.00539 | 0.0477 | 12 | SERPINF1/IGF1/MMP2/SFRP1/FGF7/PDGFRA/SERPINE2/PSAP/MDK/SOD2/NFIB/CDO1 |
| Fibroblast | F0 SFRP2+ Fib | GO:0030638 | polyketide metabolic process | 2/187 | 10/16509 | 17.66 | 0.00541 | 0.0477 | 2 | AKR1C1/AKR1C3 |
| Fibroblast | F0 SFRP2+ Fib | GO:0033212 | iron import into cell | 2/187 | 10/16509 | 17.66 | 0.00541 | 0.0477 | 2 | CYBRD1/STEAP2 |
| Fibroblast | F0 SFRP2+ Fib | GO:0035360 | positive regulation of peroxisome proliferator activated receptor signaling pathway | 2/187 | 10/16509 | 17.66 | 0.00541 | 0.0477 | 2 | PTGIS/LMO3 |
| Fibroblast | F0 SFRP2+ Fib | GO:0035701 | hematopoietic stem cell migration | 2/187 | 10/16509 | 17.66 | 0.00541 | 0.0477 | 2 | JAM2/GAS6 |
| Fibroblast | F0 SFRP2+ Fib | GO:0044598 | doxorubicin metabolic process | 2/187 | 10/16509 | 17.66 | 0.00541 | 0.0477 | 2 | AKR1C1/AKR1C3 |
| Fibroblast | F0 SFRP2+ Fib | GO:0048865 | stem cell fate commitment | 2/187 | 10/16509 | 17.66 | 0.00541 | 0.0477 | 2 | SFRP2/SFRP1 |
| Fibroblast | F0 SFRP2+ Fib | GO:0060346 | bone trabecula formation | 2/187 | 10/16509 | 17.66 | 0.00541 | 0.0477 | 2 | MMP2/SFRP1 |
| Fibroblast | F0 SFRP2+ Fib | GO:2000271 | positive regulation of fibroblast apoptotic process | 2/187 | 10/16509 | 17.66 | 0.00541 | 0.0477 | 2 | SFRP1/NUPR1 |
| Fibroblast | F0 SFRP2+ Fib | GO:0002286 | T cell activation involved in immune response | 6/187 | 141/16509 | 3.76 | 0.00541 | 0.0477 | 6 | LGALS3/MDK/FGL2/ANXA1/TMEM98/ICAM1 |
| Fibroblast | F0 SFRP2+ Fib | GO:0034614 | cellular response to reactive oxygen species | 6/187 | 141/16509 | 3.76 | 0.00541 | 0.0477 | 6 | MMP2/FBLN5/PDGFRA/SOD2/CYP1B1/CYGB |
| Fibroblast | F0 SFRP2+ Fib | GO:0033687 | osteoblast proliferation | 3/187 | 32/16509 | 8.28 | 0.00557 | 0.0486 | 3 | SFRP1/FBLN5/OSR2 |
| Fibroblast | F0 SFRP2+ Fib | GO:0071404 | cellular response to low-density lipoprotein particle stimulus | 3/187 | 32/16509 | 8.28 | 0.00557 | 0.0486 | 3 | TXNIP/CES1/CD68 |
| Fibroblast | F0 SFRP2+ Fib | GO:1902547 | regulation of cellular response to vascular endothelial growth factor stimulus | 3/187 | 32/16509 | 8.28 | 0.00557 | 0.0486 | 3 | DCN/TNXB/CD63 |
| Fibroblast | F0 SFRP2+ Fib | GO:0080164 | regulation of nitric oxide metabolic process | 4/187 | 63/16509 | 5.61 | 0.00564 | 0.0491 | 4 | IGF1/CLU/SOD2/PTGIS |
| Fibroblast | F0 SFRP2+ Fib | GO:0006631 | fatty acid metabolic process | 11/187 | 397/16509 | 2.45 | 0.00567 | 0.0491 | 11 | C3/TNXB/PTGDS/GGT5/PTGIS/EPHX1/CYP1B1/AKR1C1/AKR1C3/CES1/CYGB |
| Fibroblast | F1 MYH11+ Fib | GO:0019730 | antimicrobial humoral response | 11/173 | 169/16509 | 6.21 | 1.68e-06 | 0.00305 | 11 | ADM/CXCL2/IGHA2/IGHG1/JCHAIN/IGHG4/CCL3L3/LYZ/SLPI/IGHA1/CCL3 |
| Fibroblast | F1 MYH11+ Fib | GO:0046688 | response to copper ion | 6/173 | 37/16509 | 15.47 | 2.16e-06 | 0.00305 | 6 | MT-CO1/MT-CYB/MT1F/MT1M/MT1X/MT1A |
| Fibroblast | F1 MYH11+ Fib | GO:0019646 | aerobic electron transport chain | 8/173 | 88/16509 | 8.68 | 3.9e-06 | 0.00305 | 8 | MT-ND2/MT-CO1/MT-CYB/MT-CO3/MT-ND4/MT-ND1/MT-ND3/MT-ND6 |
| Fibroblast | F1 MYH11+ Fib | GO:0006119 | oxidative phosphorylation | 9/173 | 124/16509 | 6.93 | 6.26e-06 | 0.00305 | 9 | MT-ND2/MT-ATP6/MT-CO1/MT-CYB/MT-CO3/MT-ND4/MT-ND1/MT-ND3/MT-ND6 |
| Fibroblast | F1 MYH11+ Fib | GO:0042773 | ATP synthesis coupled electron transport | 8/173 | 95/16509 | 8.04 | 6.92e-06 | 0.00305 | 8 | MT-ND2/MT-CO1/MT-CYB/MT-CO3/MT-ND4/MT-ND1/MT-ND3/MT-ND6 |
| Fibroblast | F1 MYH11+ Fib | GO:0042775 | mitochondrial ATP synthesis coupled electron transport | 8/173 | 95/16509 | 8.04 | 6.92e-06 | 0.00305 | 8 | MT-ND2/MT-CO1/MT-CYB/MT-CO3/MT-ND4/MT-ND1/MT-ND3/MT-ND6 |
| Fibroblast | F1 MYH11+ Fib | GO:0019731 | antibacterial humoral response | 7/173 | 69/16509 | 9.68 | 7.67e-06 | 0.00305 | 7 | ADM/IGHA2/IGHG1/JCHAIN/IGHG4/SLPI/IGHA1 |

| compartment | subcluster | GO_ID | Description | GeneRatio | BgRatio | FoldEnrichment | pvalue | p_adjust | Count | geneID |
|---|---|---|---|---|---|---|---|---|---|---|
| Fibroblast | F1 MYH11+ Fib | GO:1902600 | proton transmembrane transport | 10/173 | 167/16509 | 5.71 | 1.06e-05 | 0.00305 | 10 | MT-ND2/MT-ATP6/MT-CO1/MT-CYB/MT-CO3/MT-ND4/MT-ND1/MT-ND3/MT-ND6/RNASEK |
| Fibroblast | F1 MYH11+ Fib | GO:0010273 | detoxification of copper ion | 4/173 | 14/16509 | 27.27 | 1.07e-05 | 0.00305 | 4 | MT1F/MT1M/MT1X/MT1A |
| Fibroblast | F1 MYH11+ Fib | GO:1990169 | stress response to copper ion | 4/173 | 14/16509 | 27.27 | 1.07e-05 | 0.00305 | 4 | MT1F/MT1M/MT1X/MT1A |
| Fibroblast | F1 MYH11+ Fib | GO:0006959 | humoral immune response | 13/173 | 289/16509 | 4.29 | 1.15e-05 | 0.00305 | 13 | ADM/CXCL2/IGHA2/IGHG1/JCHAIN/IGHG4/CCL3L3/LYZ/SLPI/GPR183/IGHA1/CCL3/CLU |
| Fibroblast | F1 MYH11+ Fib | GO:0042776 | proton motive force-driven mitochondrial ATP synthesis | 6/173 | 52/16509 | 11.01 | 1.66e-05 | 0.00399 | 6 | MT-ND2/MT-ATP6/MT-ND4/MT-ND1/MT-ND3/MT-ND6 |
| Fibroblast | F1 MYH11+ Fib | GO:0045926 | negative regulation of growth | 11/173 | 217/16509 | 4.84 | 1.84e-05 | 0.00399 | 11 | MT1F/CRYAB/MT1M/TSPYL2/BTG1/CDKN1A/IFRD1/MT1X/SESN1/SPP1/MT1A |
| Fibroblast | F1 MYH11+ Fib | GO:0022904 | respiratory electron transport chain | 8/173 | 109/16509 | 7.0 | 1.91e-05 | 0.00399 | 8 | MT-ND2/MT-CO1/MT-CYB/MT-CO3/MT-ND4/MT-ND1/MT-ND3/MT-ND6 |
| Fibroblast | F1 MYH11+ Fib | GO:0015986 | proton motive force-driven ATP synthesis | 6/173 | 54/16509 | 10.6 | 2.07e-05 | 0.00403 | 6 | MT-ND2/MT-ATP6/MT-ND4/MT-ND1/MT-ND3/MT-ND6 |
| Fibroblast | F1 MYH11+ Fib | GO:0030316 | osteoclast differentiation | 8/173 | 112/16509 | 6.82 | 2.33e-05 | 0.00426 | 8 | FOS/JUNB/MAFB/GPR183/FCER1G/CORO1C/CCL3/TYROBP |
| Fibroblast | F1 MYH11+ Fib | GO:0009060 | aerobic respiration | 12/173 | 269/16509 | 4.26 | 2.72e-05 | 0.00469 | 12 | MT-ND2/MT-ATP6/MT-CO1/MT-CYB/MT-CO3/MT-ND4/MT-ND1/ZBTB20/MT-ND3/MT-ND6/FH/PDK4 |
| Fibroblast | F1 MYH11+ Fib | GO:0022900 | electron transport chain | 8/173 | 118/16509 | 6.47 | 3.4e-05 | 0.00552 | 8 | MT-ND2/MT-CO1/MT-CYB/MT-CO3/MT-ND4/MT-ND1/MT-ND3/MT-ND6 |
| Fibroblast | F1 MYH11+ Fib | GO:0009206 | purine ribonucleoside triphosphate biosynthetic process | 7/173 | 88/16509 | 7.59 | 3.82e-05 | 0.00573 | 7 | MT-ND2/MT-ATP6/MT-ND4/MT-ND1/MT-ND3/MT-ND6/NME7 |
| Fibroblast | F1 MYH11+ Fib | GO:0097501 | stress response to metal ion | 4/173 | 19/16509 | 20.09 | 3.99e-05 | 0.00573 | 4 | MT1F/MT1M/MT1X/MT1A |
| Fibroblast | F1 MYH11+ Fib | GO:0009145 | purine nucleoside triphosphate biosynthetic process | 7/173 | 89/16509 | 7.51 | 4.11e-05 | 0.00573 | 7 | MT-ND2/MT-ATP6/MT-ND4/MT-ND1/MT-ND3/MT-ND6/NME7 |
| Fibroblast | F1 MYH11+ Fib | GO:0045444 | fat cell differentiation | 11/173 | 240/16509 | 4.37 | 4.64e-05 | 0.00608 | 11 | ADIRF/MAFB/WDFY2/ID4/NR4A2/IFRD1/OPA3/NR4A1/HES1/ZFP36/ARL4A |
| Fibroblast | F1 MYH11+ Fib | GO:0045765 | regulation of angiogenesis | 12/173 | 287/16509 | 3.99 | 5.12e-05 | 0.00608 | 12 | HSPB1/ADM/RHOB/FOXC1/SEMA4A/FOXC2/ADAMTS1/ANGPT2/BTG1/ID1/KLF2/PLK2 |
| Fibroblast | F1 MYH11+ Fib | GO:0006958 | complement activation, classical pathway | 5/173 | 39/16509 | 12.23 | 5.15e-05 | 0.00608 | 5 | IGHA2/IGHG1/IGHG4/IGHA1/CLU |
| Fibroblast | F1 MYH11+ Fib | GO:0048545 | response to steroid hormone | 13/173 | 334/16509 | 3.71 | 5.19e-05 | 0.00608 | 13 | JUN/FOS/FOSB/JUNB/MT-ND3/EIF4E/SGK1/CLDN4/SPP1/DUSP1/LBH/BCL2L11/ZFP36 |
| Fibroblast | F1 MYH11+ Fib | GO:0009201 | ribonucleoside triphosphate biosynthetic process | 7/173 | 94/16509 | 7.11 | 5.85e-05 | 0.00657 | 7 | MT-ND2/MT-ATP6/MT-ND4/MT-ND1/MT-ND3/MT-ND6/NME7 |
| Fibroblast | F1 MYH11+ Fib | GO:0061687 | detoxification of inorganic compound | 4/173 | 21/16509 | 18.18 | 6.06e-05 | 0.00657 | 4 | MT1F/MT1M/MT1X/MT1A |
| Fibroblast | F1 MYH11+ Fib | GO:1901342 | regulation of vasculature development | 12/173 | 294/16509 | 3.9 | 6.45e-05 | 0.00664 | 12 | HSPB1/ADM/RHOB/FOXC1/SEMA4A/FOXC2/ADAMTS1/ANGPT2/BTG1/ID1/KLF2/PLK2 |
| Fibroblast | F1 MYH11+ Fib | GO:1902742 | apoptotic process involved in development | 5/173 | 41/16509 | 11.64 | 6.58e-05 | 0.00664 | 5 | FOXC1/FOXC2/CRYAB/KIF1B/BCL2L11 |
| Fibroblast | F1 MYH11+ Fib | GO:0010038 | response to metal ion | 13/173 | 345/16509 | 3.6 | 7.22e-05 | 0.00704 | 13 | MT-CO1/MT-CYB/JUN/FOS/FOSB/JUNB/MT1F/MT1M/MT1X/MICU3/PLN/DUSP1/MT1A |
| Fibroblast | F1 MYH11+ Fib | GO:0045333 | cellular respiration | 12/173 | 305/16509 | 3.75 | 9.16e-05 | 0.00864 | 12 | MT-ND2/MT-ATP6/MT-CO1/MT-CYB/MT-CO3/MT-ND4/MT-ND1/ZBTB20/MT-ND3/MT-ND6/FH/PDK4 |
| Fibroblast | F1 MYH11+ Fib | GO:0009142 | nucleoside triphosphate biosynthetic process | 7/173 | 102/16509 | 6.55 | 9.83e-05 | 0.00899 | 7 | MT-ND2/MT-ATP6/MT-ND4/MT-ND1/MT-ND3/MT-ND6/NME7 |
| Fibroblast | F1 MYH11+ Fib | GO:0071294 | cellular response to zinc ion | 4/173 | 24/16509 | 15.9 | 0.000105 | 0.00932 | 4 | MT1F/MT1M/MT1X/MT1A |
| Fibroblast | F1 MYH11+ Fib | GO:0016064 | immunoglobulin mediated immune response | 9/173 | 180/16509 | 4.77 | 0.000119 | 0.00989 | 9 | IGHA2/IGHG1/IGLC3/IGLC2/IGKC/IGHG4/FCER1G/IGHA1/CLU |

| compartment | subcluster | GO_ID | Description | GeneRatio | BgRatio | FoldEnrichment | pvalue | p_adjust | Count | geneID |
|---|---|---|---|---|---|---|---|---|---|---|
| Fibroblast | F1 MYH11+ Fib | GO:0071276 | cellular response to cadmium ion | 4/173 | 25/16509 | 15.27 | 0.000124 | 0.00989 | 4 | MT1F/MT1M/MT1X/MT1A |
| Fibroblast | F1 MYH11+ Fib | GO:0071280 | cellular response to copper ion | 4/173 | 25/16509 | 15.27 | 0.000124 | 0.00989 | 4 | MT1F/MT1M/MT1X/MT1A |
| Fibroblast | F1 MYH11+ Fib | GO:0046686 | response to cadmium ion | 5/173 | 47/16509 | 10.15 | 0.000128 | 0.00989 | 5 | MT-CYB/MT1F/MT1M/MT1X/MT1A |
| Fibroblast | F1 MYH11+ Fib | GO:0050853 | B cell receptor signaling pathway | 6/173 | 75/16509 | 7.63 | 0.000134 | 0.00989 | 6 | IGHA2/IGHG1/IGLC3/IGKC/IGHG4/IGHA1 |
| Fibroblast | F1 MYH11+ Fib | GO:0019724 | B cell mediated immunity | 9/173 | 183/16509 | 4.69 | 0.000135 | 0.00989 | 9 | IGHA2/IGHG1/IGLC3/IGLC2/IGKC/IGHG4/FCER1G/IGHA1/CLU |
| Fibroblast | F1 MYH11+ Fib | GO:0071248 | cellular response to metal ion | 9/173 | 183/16509 | 4.69 | 0.000135 | 0.00989 | 9 | JUN/FOS/FOSB/JUNB/MT1F/MT1M/MT1X/MICU3/MT1A |
| Fibroblast | F1 MYH11+ Fib | GO:0002460 | adaptive immune response based on somatic recombination of immune receptors built from im… | 13/173 | 370/16509 | 3.35 | 0.000145 | 0.0103 | 13 | JUNB/IGHA2/IGHG1/IGLC3/IGLC2/SEMA4A/IGKC/IGHG4/FCER1G/BRD2/IGHA1/HLA-DRA/CLU |
| Fibroblast | F1 MYH11+ Fib | GO:0043542 | endothelial cell migration | 10/173 | 228/16509 | 4.19 | 0.000148 | 0.0103 | 10 | HSPB1/RAB13/RHOB/SEMA4A/FOXC2/ANGPT2/ID1/PLK2/NR4A1/PTP4A3 |
| Fibroblast | F1 MYH11+ Fib | GO:0006754 | ATP biosynthetic process | 6/173 | 78/16509 | 7.34 | 0.000167 | 0.0112 | 6 | MT-ND2/MT-ATP6/MT-ND4/MT-ND1/MT-ND3/MT-ND6 |
| Fibroblast | F1 MYH11+ Fib | GO:0009260 | ribonucleotide biosynthetic process | 8/173 | 148/16509 | 5.16 | 0.000168 | 0.0112 | 8 | MT-ND2/MT-ATP6/MT-ND4/MT-ND1/MT-ND3/MT-ND6/UPP1/NME7 |
| Fibroblast | F1 MYH11+ Fib | GO:0006120 | mitochondrial electron transport, NADH to ubiquinone | 5/173 | 50/16509 | 9.54 | 0.000173 | 0.0112 | 5 | MT-ND2/MT-ND4/MT-ND1/MT-ND3/MT-ND6 |
| Fibroblast | F1 MYH11+ Fib | GO:0031960 | response to corticosteroid | 8/173 | 151/16509 | 5.06 | 0.000193 | 0.0122 | 8 | FOS/FOSB/MT-ND3/EIF4E/SGK1/DUSP1/BCL2L11/ZFP36 |
| Fibroblast | F1 MYH11+ Fib | GO:0060561 | apoptotic process involved in morphogenesis | 4/173 | 28/16509 | 13.63 | 0.000196 | 0.0122 | 4 | FOXC1/FOXC2/CRYAB/BCL2L11 |
| Fibroblast | F1 MYH11+ Fib | GO:0014032 | neural crest cell development | 6/173 | 81/16509 | 7.07 | 0.000206 | 0.0122 | 6 | FOXC1/SEMA4A/FOXC2/CITED2/CORO1C/HES1 |
| Fibroblast | F1 MYH11+ Fib | GO:0002455 | humoral immune response mediated by circulating immunoglobulin | 5/173 | 52/16509 | 9.18 | 0.000208 | 0.0122 | 5 | IGHA2/IGHG1/IGHG4/IGHA1/CLU |
| Fibroblast | F1 MYH11+ Fib | GO:0010043 | response to zinc ion | 5/173 | 52/16509 | 9.18 | 0.000208 | 0.0122 | 5 | MT1F/MT1M/MT1X/PLN/MT1A |
| Fibroblast | F1 MYH11+ Fib | GO:0070482 | response to oxygen levels | 12/173 | 335/16509 | 3.42 | 0.00022 | 0.0124 | 12 | MT-ND2/MT-ATP6/MT-CO1/MT-CYB/MT-ND4/FOS/MT-ND1/CRYAB/ANGPT2/CITED2/NR4A2/NOL3 |
| Fibroblast | F1 MYH11+ Fib | GO:0046390 | ribose phosphate biosynthetic process | 8/173 | 154/16509 | 4.96 | 0.00022 | 0.0124 | 8 | MT-ND2/MT-ATP6/MT-ND4/MT-ND1/MT-ND3/MT-ND6/UPP1/NME7 |
| Fibroblast | F1 MYH11+ Fib | GO:0015980 | energy derivation by oxidation of organic compounds | 13/173 | 393/16509 | 3.16 | 0.000262 | 0.0143 | 13 | MT-ND2/MT-ATP6/MT-CO1/MT-CYB/MT-CO3/MT-ND4/MT-ND1/ZBTB20/MT-ND3/MT-ND6/PHKG1/FH/PDK4 |
| Fibroblast | F1 MYH11+ Fib | GO:0030099 | myeloid cell differentiation | 14/173 | 448/16509 | 2.98 | 0.000272 | 0.0143 | 14 | JUN/FOS/JUNB/MAFB/GPR183/CITED2/KLF2/FCER1G/CORO1C/CCL3/TYROBP/ANKRD54/ZFP36/ARID4A |
| Fibroblast | F1 MYH11+ Fib | GO:0033002 | muscle cell proliferation | 9/173 | 201/16509 | 4.27 | 0.000273 | 0.0143 | 9 | JUN/FOS/FOXC1/FOXC2/ADAMTS1/PTGIR/CDKN1A/HBEGF/CNN1 |
| Fibroblast | F1 MYH11+ Fib | GO:0002573 | myeloid leukocyte differentiation | 10/173 | 246/16509 | 3.88 | 0.000274 | 0.0143 | 10 | JUN/FOS/JUNB/MAFB/GPR183/CITED2/FCER1G/CORO1C/CCL3/TYROBP |
| Fibroblast | F1 MYH11+ Fib | GO:0006935 | chemotaxis | 14/173 | 452/16509 | 2.96 | 0.000297 | 0.0148 | 14 | HSPB1/RAB13/CXCL2/SEMA4A/CCL3L3/ANGPT2/GPR183/HBEGF/PDGFA/FCER1G/CORO1C/CCL3/NR4A1/DUSP1 |
| Fibroblast | F1 MYH11+ Fib | GO:0060211 | regulation of nuclear-transcribed mRNA poly(A) tail shortening | 3/173 | 13/16509 | 22.02 | 0.000299 | 0.0148 | 3 | BTG2/TOB1/ZFP36 |
| Fibroblast | F1 MYH11+ Fib | GO:0060213 | positive regulation of nuclear-transcribed mRNA poly(A) tail shortening | 3/173 | 13/16509 | 22.02 | 0.000299 | 0.0148 | 3 | BTG2/TOB1/ZFP36 |
| Fibroblast | F1 MYH11+ Fib | GO:0048864 | stem cell development | 6/173 | 87/16509 | 6.58 | 0.000304 | 0.0148 | 6 | FOXC1/SEMA4A/FOXC2/CITED2/CORO1C/HES1 |

| compartment | subcluster | GO_ID | Description | GeneRatio | BgRatio | FoldEnrichment | pvalue | p_adjust | Count | geneID |
|---|---|---|---|---|---|---|---|---|---|---|
| Fibroblast | F1 MYH11+ Fib | GO:0042330 | taxis | 14/173 | 454/16509 | 2.94 | 0.000311 | 0.0149 | 14 | HSPB1/RAB13/CXCL2/SEMA4A/CCL3L3/ANGPT2/GPR183/HBEGF/PDGFA/FCER1G/CORO1C/CCL3/NR4A1/DUSP1 |
| Fibroblast | F1 MYH11+ Fib | GO:0001666 | response to hypoxia | 11/173 | 300/16509 | 3.5 | 0.00033 | 0.0156 | 11 | MT-ND2/MT-CO1/MT-CYB/MT-ND4/FOS/MT-ND1/CRYAB/ANGPT2/CITED2/NR4A2/NOL3 |
| Fibroblast | F1 MYH11+ Fib | GO:0007517 | muscle organ development | 12/173 | 355/16509 | 3.23 | 0.000372 | 0.0173 | 12 | SPEG/FOS/FOXC1/ATF3/FOXC2/CRYAB/BTG2/HBEGF/CITED2/RCAN1/NR4A1/SGCA |
| Fibroblast | F1 MYH11+ Fib | GO:0006882 | intracellular zinc ion homeostasis | 4/173 | 34/16509 | 11.23 | 0.000423 | 0.0193 | 4 | MT1F/MT1M/MT1X/MT1A |
| Fibroblast | F1 MYH11+ Fib | GO:0051384 | response to glucocorticoid | 7/173 | 130/16509 | 5.14 | 0.00044 | 0.0198 | 7 | FOS/FOSB/MT-ND3/EIF4E/DUSP1/BCL2L11/ZFP36 |
| Fibroblast | F1 MYH11+ Fib | GO:0060326 | cell chemotaxis | 11/173 | 311/16509 | 3.38 | 0.000447 | 0.0198 | 11 | HSPB1/RAB13/CXCL2/CCL3L3/GPR183/HBEGF/FCER1G/CORO1C/CCL3/NR4A1/DUSP1 |
| Fibroblast | F1 MYH11+ Fib | GO:0014033 | neural crest cell differentiation | 6/173 | 94/16509 | 6.09 | 0.000461 | 0.0202 | 6 | FOXC1/SEMA4A/FOXC2/CITED2/CORO1C/HES1 |
| Fibroblast | F1 MYH11+ Fib | GO:0036293 | response to decreased oxygen levels | 11/173 | 313/16509 | 3.35 | 0.000472 | 0.0202 | 11 | MT-ND2/MT-CO1/MT-CYB/MT-ND4/FOS/MT-ND1/CRYAB/ANGPT2/CITED2/NR4A2/NOL3 |
| Fibroblast | F1 MYH11+ Fib | GO:0006956 | complement activation | 5/173 | 62/16509 | 7.7 | 0.000476 | 0.0202 | 5 | IGHA2/IGHG1/IGHG4/IGHA1/CLU |
| Fibroblast | F1 MYH11+ Fib | GO:0009152 | purine ribonucleotide biosynthetic process | 7/173 | 132/16509 | 5.06 | 0.000483 | 0.0202 | 7 | MT-ND2/MT-ATP6/MT-ND4/MT-ND1/MT-ND3/MT-ND6/NME7 |
| Fibroblast | F1 MYH11+ Fib | GO:0042594 | response to starvation | 9/173 | 221/16509 | 3.89 | 0.000546 | 0.0225 | 9 | FOS/RNF152/ATF3/CDKN1A/SESN1/FADS1/UPP1/ZFP36/PDK4 |
| Fibroblast | F1 MYH11+ Fib | GO:0010257 | NADH dehydrogenase complex assembly | 4/173 | 37/16509 | 10.32 | 0.000587 | 0.0234 | 4 | MT-ND2/MT-ND4/MT-ND1/MT-ND6 |
| Fibroblast | F1 MYH11+ Fib | GO:0032981 | mitochondrial respiratory chain complex I assembly | 4/173 | 37/16509 | 10.32 | 0.000587 | 0.0234 | 4 | MT-ND2/MT-ND4/MT-ND1/MT-ND6 |
| Fibroblast | F1 MYH11+ Fib | GO:0071384 | cellular response to corticosteroid stimulus | 5/173 | 65/16509 | 7.34 | 0.000593 | 0.0234 | 5 | MT-ND3/EIF4E/SGK1/BCL2L11/ZFP36 |
| Fibroblast | F1 MYH11+ Fib | GO:0009612 | response to mechanical stimulus | 9/173 | 224/16509 | 3.83 | 0.000601 | 0.0235 | 9 | JUN/FOS/FOSB/BTG2/ANGPT2/CITED2/RCAN1/USP53/SUN1 |
| Fibroblast | F1 MYH11+ Fib | GO:1901654 | response to ketone | 9/173 | 226/16509 | 3.8 | 0.000641 | 0.0244 | 9 | FOS/FOSB/EIF4E/BTG2/SGK1/CLDN4/PLN/SPP1/DUSP1 |
| Fibroblast | F1 MYH11+ Fib | GO:0002292 | T cell differentiation involved in immune response | 6/173 | 100/16509 | 5.73 | 0.000642 | 0.0244 | 6 | JUNB/SEMA4A/GPR183/FCER1G/BRD2/HLA-DRA |
| Fibroblast | F1 MYH11+ Fib | GO:0002286 | T cell activation involved in immune response | 7/173 | 141/16509 | 4.74 | 0.000716 | 0.0268 | 7 | JUNB/SEMA4A/GPR183/FCER1G/BRD2/HLA-DRA/TYROBP |
| Fibroblast | F1 MYH11+ Fib | GO:0051592 | response to calcium ion | 7/173 | 142/16509 | 4.7 | 0.000746 | 0.0276 | 7 | MT-CYB/JUN/FOS/FOSB/JUNB/MICU3/DUSP1 |
| Fibroblast | F1 MYH11+ Fib | GO:0009267 | cellular response to starvation | 8/173 | 186/16509 | 4.1 | 0.000777 | 0.0284 | 8 | FOS/RNF152/ATF3/CDKN1A/SESN1/FADS1/UPP1/PDK4 |
| Fibroblast | F1 MYH11+ Fib | GO:0016525 | negative regulation of angiogenesis | 6/173 | 105/16509 | 5.45 | 0.00083 | 0.0296 | 6 | FOXC1/SEMA4A/ADAMTS1/ANGPT2/KLF2/PLK2 |
| Fibroblast | F1 MYH11+ Fib | GO:0032760 | positive regulation of tumor necrosis factor production | 6/173 | 105/16509 | 5.45 | 0.00083 | 0.0296 | 6 | HSPB1/ZBTB20/FCER1G/CCL3/CLU/TYROBP |
| Fibroblast | F1 MYH11+ Fib | GO:2000181 | negative regulation of blood vessel morphogenesis | 6/173 | 106/16509 | 5.4 | 0.000872 | 0.0307 | 6 | FOXC1/SEMA4A/ADAMTS1/ANGPT2/KLF2/PLK2 |
| Fibroblast | F1 MYH11+ Fib | GO:1901343 | negative regulation of vasculature development | 6/173 | 107/16509 | 5.35 | 0.000916 | 0.0319 | 6 | FOXC1/SEMA4A/ADAMTS1/ANGPT2/KLF2/PLK2 |
| Fibroblast | F1 MYH11+ Fib | GO:0006979 | response to oxidative stress | 12/173 | 394/16509 | 2.91 | 0.000932 | 0.0321 | 12 | MT-CO1/FOS/MT-ND1/MT-ND3/RHOB/CRYAB/BTG1/NR4A2/RCAN1/KLF2/SESN1/DUSP1 |
| Fibroblast | F1 MYH11+ Fib | GO:0045670 | regulation of osteoclast differentiation | 5/173 | 72/16509 | 6.63 | 0.000947 | 0.0322 | 5 | FOS/MAFB/CORO1C/CCL3/TYROBP |
| Fibroblast | F1 MYH11+ Fib | GO:0060263 | regulation of respiratory burst | 3/173 | 19/16509 | 15.07 | 0.000969 | 0.0323 | 3 | IGHA2/JCHAIN/IGHA1 |
| Fibroblast | F1 MYH11+ Fib | GO:0006091 | generation of precursor metabolites and energy | 13/173 | 452/16509 | 2.74 | 0.000972 | 0.0323 | 13 | MT-ND2/MT-ATP6/MT-CO1/MT-CYB/MT-CO3/MT-ND4/MT-ND1/ZBTB20/MT-ND3/MT-ND6/PHKG1/FH/PDK4 |

| compartment | subcluster | GO_ID | Description | GeneRatio | BgRatio | FoldEnrichment | pvalue | p_adjust | Count | geneID |
|---|---|---|---|---|---|---|---|---|---|---|
| Fibroblast | F1 MYH11+ Fib | GO:0002262 | myeloid cell homeostasis | 8/173 | 193/16509 | 3.96 | 0.000987 | 0.0325 | 8 | MAFB/CITED2/KLF2/FCER1G/ANKRD54/BCL2L11/ZFP36/ARID4A |
| Fibroblast | F1 MYH11+ Fib | GO:1903557 | positive regulation of tumor necrosis factor superfamily cytokine production | 6/173 | 109/16509 | 5.25 | 0.00101 | 0.0328 | 6 | HSPB1/ZBTB20/FCER1G/CCL3/CLU/TYROBP |
| Fibroblast | F1 MYH11+ Fib | GO:0014706 | striated muscle tissue development | 12/173 | 402/16509 | 2.85 | 0.00111 | 0.0356 | 12 | MYH11/FOS/FOXC1/SORBS2/ATF3/FOXC2/BTG2/CITED2/IFRD1/RCAN1/NR4A1/PLN |
| Fibroblast | F1 MYH11+ Fib | GO:0001774 | microglial cell activation | 4/173 | 45/16509 | 8.48 | 0.00124 | 0.0395 | 4 | JUN/CCL3/CLU/TYROBP |
| Fibroblast | F1 MYH11+ Fib | GO:0048872 | homeostasis of number of cells | 11/173 | 353/16509 | 2.97 | 0.00126 | 0.0398 | 11 | MAFB/GPR183/CITED2/KLF2/FCER1G/FH/ANKRD54/BCL2L11/ZFP36/UBAP2L/ARID4A |
| Fibroblast | F1 MYH11+ Fib | GO:0002449 | lymphocyte mediated immunity | 11/173 | 356/16509 | 2.95 | 0.00135 | 0.0418 | 11 | IGHA2/IGHG1/IGLC3/IGLC2/IGKC/IGHG4/FCER1G/IGHA1/HLA-DRA/CLU/TYROBP |
| Fibroblast | F1 MYH11+ Fib | GO:0035914 | skeletal muscle cell differentiation | 5/173 | 78/16509 | 6.12 | 0.00136 | 0.0418 | 5 | FOS/ATF3/BTG2/CITED2/NR4A1 |
| Fibroblast | F1 MYH11+ Fib | GO:0009165 | nucleotide biosynthetic process | 8/173 | 204/16509 | 3.74 | 0.00141 | 0.0429 | 8 | MT-ND2/MT-ATP6/MT-ND4/MT-ND1/MT-ND3/MT-ND6/UPP1/NME7 |
| Fibroblast | F1 MYH11+ Fib | GO:0002269 | leukocyte activation involved in inflammatory response | 4/173 | 48/16509 | 7.95 | 0.00158 | 0.0474 | 4 | JUN/CCL3/CLU/TYROBP |
| Fibroblast | F1 MYH11+ Fib | GO:0043534 | blood vessel endothelial cell migration | 6/173 | 119/16509 | 4.81 | 0.00159 | 0.0474 | 6 | HSPB1/FOXC2/ANGPT2/ID1/PLK2/NR4A1 |
| Fibroblast | F1 MYH11+ Fib | GO:0071277 | cellular response to calcium ion | 5/173 | 81/16509 | 5.89 | 0.00161 | 0.0475 | 5 | JUN/FOS/FOSB/JUNB/MICU3 |
| Fibroblast | F1 MYH11+ Fib | GO:0003151 | outflow tract morphogenesis | 5/173 | 82/16509 | 5.82 | 0.0017 | 0.0494 | 5 | JUN/FOXC1/FOXC2/CITED2/HES1 |
| Fibroblast | F1 MYH11+ Fib | GO:0060537 | muscle tissue development | 12/173 | 423/16509 | 2.71 | 0.0017 | 0.0494 | 12 | MYH11/FOS/FOXC1/SORBS2/ATF3/FOXC2/BTG2/CITED2/IFRD1/RCAN1/NR4A1/PLN |
| Fibroblast | F2 RGS5+ Fib | GO:0003012 | muscle system process | 31/176 | 447/16509 | 6.51 | 7.46e-17 | 1.05e-13 | 31 | MYH11/MYL9/ACTA2/TPM2/PLN/TBX2/CALD1/GJC1/CALM2/HEY2/CAV1/MYLK/CACNA1H/KCNA5/PDGFRB/DMPK/… |
| Fibroblast | F2 RGS5+ Fib | GO:0006936 | muscle contraction | 28/176 | 353/16509 | 7.44 | 8.71e-17 | 1.05e-13 | 28 | MYH11/MYL9/ACTA2/TPM2/PLN/TBX2/CALD1/GJC1/CALM2/CAV1/MYLK/CACNA1H/KCNA5/DMPK/SGCA/CNN1/HR… |
| Fibroblast | F2 RGS5+ Fib | GO:0007219 | Notch signaling pathway | 16/176 | 178/16509 | 8.43 | 7.59e-11 | 6.13e-08 | 16 | NOTCH3/PTP4A3/CDH6/RCAN2/PLN/TBX2/HEY2/HEYL/KCNA5/NRIP2/NEURL1B/JAG1/GRIP2/SORBS2/FOXC2/S… |
| Fibroblast | F2 RGS5+ Fib | GO:0042692 | muscle cell differentiation | 22/176 | 396/16509 | 5.21 | 2.48e-10 | 1.44e-07 | 22 | MYH11/MEF2C/CSRP2/MYL9/TBX2/HEY2/CAV2/CACNA1H/PDGFRB/DMPK/EHD2/TPM1/LDB3/LMOD1/ACTN4/SORT… |
| Fibroblast | F2 RGS5+ Fib | GO:0032835 | glomerulus development | 11/176 | 73/16509 | 14.13 | 2.96e-10 | 1.44e-07 | 11 | NOTCH3/MEF2C/GPR4/ACTA2/ANGPT2/HEYL/PLCE1/ENPEP/PDGFRB/JAG1/FOXC2 |
| Fibroblast | F2 RGS5+ Fib | GO:0072006 | nephron development | 14/176 | 159/16509 | 8.26 | 1.6e-09 | 6.45e-07 | 14 | NOTCH3/MEF2C/GPR4/ACTA2/ANGPT2/HEYL/PLCE1/LAMA5/ENPEP/PDGFRB/ITGA3/JAG1/FOXC2/NPNT |
| Fibroblast | F2 RGS5+ Fib | GO:0051146 | striated muscle cell differentiation | 18/176 | 294/16509 | 5.74 | 2.57e-09 | 8.88e-07 | 18 | MYH11/MEF2C/CSRP2/MYL9/HEY2/CAV2/CACNA1H/PDGFRB/DMPK/EHD2/TPM1/LDB3/LMOD1/SORT1/SORBS2/CS… |
| Fibroblast | F2 RGS5+ Fib | GO:0033002 | muscle cell proliferation | 15/176 | 201/16509 | 7.0 | 4.06e-09 | 1.23e-06 | 15 | RGS5/NOTCH3/MEF2C/TBX2/HEY2/CAV2/PDGFRB/CNN1/TPM1/NDRG2/ADAMTS1/RBPMS2/PTGIR/FOXC2/MYOCD |
| Fibroblast | F2 RGS5+ Fib | GO:0048844 | artery morphogenesis | 10/176 | 74/16509 | 12.68 | 5.85e-09 | 1.57e-06 | 10 | NOTCH3/TBX2/HEY2/MYLK/PDGFRB/ADGRF5/JAG1/FOXC2/ADAMTS9/MYOCD |
| Fibroblast | F2 RGS5+ Fib | GO:0008016 | regulation of heart contraction | 15/176 | 212/16509 | 6.64 | 8.39e-09 | 2.03e-06 | 15 | PLN/TBX2/GJC1/CALM2/HEY2/CAV1/CACNA1H/EPAS1/KCNA5/DMPK/HRC/TPM1/ATP1B2/CASQ2/SNTA1 |

| compartment | subcluster | GO_ID | Description | GeneRatio | BgRatio | FoldEnrichment | pvalue | p_adjust | Count | geneID |
|---|---|---|---|---|---|---|---|---|---|---|
| Fibroblast | F2 RGS5+ Fib | GO:0072001 | renal system development | 18/176 | 331/16509 | 5.1 | 1.64e-08 | 3.61e-06 | 18 | NOTCH3/MEF2C/GPR4/ACTA2/ANGPT2/HEYL/PDGFA/PLCE1/LAMA5/ENPEP/PDGFRB/ITGA3/ID3/JAG1/ADAMTS1… |
| Fibroblast | F2 RGS5+ Fib | GO:1903522 | regulation of blood circulation | 16/176 | 265/16509 | 5.66 | 2.5e-08 | 5.05e-06 | 16 | PLN/TBX2/GJC1/CALM2/HEY2/CAV1/CACNA1H/EPAS1/KCNA5/DMPK/HRC/TPM1/ATP1B2/CASQ2/ADRA2B/SNTA1 |
| Fibroblast | F2 RGS5+ Fib | GO:0006937 | regulation of muscle contraction | 13/176 | 168/16509 | 7.26 | 2.97e-08 | 5.54e-06 | 13 | MYL9/PLN/CALM2/CAV1/DMPK/CNN1/HRC/TPM1/CASQ2/PPP1R12B/ADRA2B/NPNT/MYOCD |
| Fibroblast | F2 RGS5+ Fib | GO:0001822 | kidney development | 17/176 | 320/16509 | 4.98 | 5.88e-08 | 1.02e-05 | 17 | NOTCH3/MEF2C/GPR4/ACTA2/ANGPT2/HEYL/PDGFA/PLCE1/LAMA5/ENPEP/PDGFRB/ITGA3/ID3/JAG1/ADAMTS1… |
| Fibroblast | F2 RGS5+ Fib | GO:0060047 | heart contraction | 15/176 | 252/16509 | 5.58 | 8.41e-08 | 1.36e-05 | 15 | PLN/TBX2/GJC1/CALM2/HEY2/CAV1/CACNA1H/EPAS1/KCNA5/DMPK/HRC/TPM1/ATP1B2/CASQ2/SNTA1 |
| Fibroblast | F2 RGS5+ Fib | GO:1904706 | negative regulation of vascular associated smooth muscle cell proliferation | 6/176 | 23/16509 | 24.47 | 1.17e-07 | 1.77e-05 | 6 | RGS5/MEF2C/CNN1/TPM1/NDRG2/MYOCD |
| Fibroblast | F2 RGS5+ Fib | GO:0003015 | heart process | 15/176 | 262/16509 | 5.37 | 1.4e-07 | 1.99e-05 | 15 | PLN/TBX2/GJC1/CALM2/HEY2/CAV1/CACNA1H/EPAS1/KCNA5/DMPK/HRC/TPM1/ATP1B2/CASQ2/SNTA1 |
| Fibroblast | F2 RGS5+ Fib | GO:0006939 | smooth muscle contraction | 10/176 | 104/16509 | 9.02 | 1.61e-07 | 2.17e-05 | 10 | MYH11/ACTA2/TBX2/CAV1/MYLK/CNN1/GRIP2/ADRA2B/NPNT/MYOCD |
| Fibroblast | F2 RGS5+ Fib | GO:0048660 | regulation of smooth muscle cell proliferation | 11/176 | 133/16509 | 7.76 | 1.81e-07 | 2.31e-05 | 11 | RGS5/NOTCH3/MEF2C/PDGFRB/CNN1/TPM1/NDRG2/ADAMTS1/RBPMS2/PTGIR/MYOCD |
| Fibroblast | F2 RGS5+ Fib | GO:0002027 | regulation of heart rate | 10/176 | 107/16509 | 8.77 | 2.11e-07 | 2.56e-05 | 10 | PLN/CALM2/HEY2/CAV1/EPAS1/KCNA5/HRC/TPM1/CASQ2/SNTA1 |
| Fibroblast | F2 RGS5+ Fib | GO:0048659 | smooth muscle cell proliferation | 11/176 | 136/16509 | 7.59 | 2.27e-07 | 2.62e-05 | 11 | RGS5/NOTCH3/MEF2C/PDGFRB/CNN1/TPM1/NDRG2/ADAMTS1/RBPMS2/PTGIR/MYOCD |
| Fibroblast | F2 RGS5+ Fib | GO:0060840 | artery development | 10/176 | 110/16509 | 8.53 | 2.75e-07 | 3.02e-05 | 10 | NOTCH3/TBX2/HEY2/MYLK/PDGFRB/ADGRF5/JAG1/FOXC2/ADAMTS9/MYOCD |
| Fibroblast | F2 RGS5+ Fib | GO:0072012 | glomerulus vasculature development | 6/176 | 29/16509 | 19.41 | 5.22e-07 | 5.5e-05 | 6 | NOTCH3/GPR4/ACTA2/ANGPT2/PDGFRB/FOXC2 |
| Fibroblast | F2 RGS5+ Fib | GO:0060537 | muscle tissue development | 18/176 | 423/16509 | 3.99 | 6.44e-07 | 6.5e-05 | 18 | MYH11/MEF2C/CSRP2/PLN/TBX2/GJC1/HEY2/CAV2/CAV1/HEYL/MYLK/PDGFRB/TPM1/SORBS2/CSRP1/FOXC2/A… |
| Fibroblast | F2 RGS5+ Fib | GO:0061437 | renal system vasculature development | 6/176 | 31/16509 | 18.16 | 7.96e-07 | 7.33e-05 | 6 | NOTCH3/GPR4/ACTA2/ANGPT2/PDGFRB/FOXC2 |
| Fibroblast | F2 RGS5+ Fib | GO:0061440 | kidney vasculature development | 6/176 | 31/16509 | 18.16 | 7.96e-07 | 7.33e-05 | 6 | NOTCH3/GPR4/ACTA2/ANGPT2/PDGFRB/FOXC2 |
| Fibroblast | F2 RGS5+ Fib | GO:0048662 | negative regulation of smooth muscle cell proliferation | 7/176 | 49/16509 | 13.4 | 8.18e-07 | 7.33e-05 | 7 | RGS5/MEF2C/CNN1/TPM1/NDRG2/PTGIR/MYOCD |
| Fibroblast | F2 RGS5+ Fib | GO:0055001 | muscle cell development | 12/176 | 192/16509 | 5.86 | 1.04e-06 | 8.95e-05 | 12 | MYH11/CSRP2/MYL9/HEY2/CAV2/PDGFRB/TPM1/LDB3/LMOD1/ACTN4/SORBS2/CSRP1 |
| Fibroblast | F2 RGS5+ Fib | GO:0010644 | cell communication by electrical coupling | 6/176 | 33/16509 | 17.05 | 1.18e-06 | 9.51e-05 | 6 | GJC1/CALM2/CAV1/HRC/ATP1B2/CASQ2 |
| Fibroblast | F2 RGS5+ Fib | GO:0030239 | myofibril assembly | 8/176 | 74/16509 | 10.14 | 1.18e-06 | 9.51e-05 | 8 | MYH11/CSRP2/MYL9/PDGFRB/TPM1/LDB3/LMOD1/CSRP1 |
| Fibroblast | F2 RGS5+ Fib | GO:0007517 | muscle organ development | 16/176 | 355/16509 | 4.23 | 1.32e-06 | 0.000103 | 16 | MEF2C/ITGA7/TAGLN/HEY2/CAV2/CAV1/HEYL/MYLK/CACNA1H/LAMA5/SGCA/TPM1/ID3/CRYAB/FOXC2/MYOCD |
| Fibroblast | F2 RGS5+ Fib | GO:0035909 | aorta morphogenesis | 6/176 | 34/16509 | 16.55 | 1.42e-06 | 0.000107 | 6 | TBX2/HEY2/MYLK/PDGFRB/JAG1/ADAMTS9 |

| compartment | subcluster | GO_ID | Description | GeneRatio | BgRatio | FoldEnrichment | pvalue | p_adjust | Count | geneID |
|---|---|---|---|---|---|---|---|---|---|---|
| Fibroblast | F2 RGS5+ Fib | GO:0055002 | striated muscle cell development | 11/176 | 164/16509 | 6.29 | 1.48e-06 | 0.000109 | 11 | MYH11/CSRP2/MYL9/HEY2/CAV2/PDGFRB/TPM1/LDB3/LMOD1/SORBS2/CSRP1 |
| Fibroblast | F2 RGS5+ Fib | GO:0090257 | regulation of muscle system process | 13/176 | 237/16509 | 5.15 | 1.59e-06 | 0.000113 | 13 | MYL9/PLN/CALM2/CAV1/DMPK/CNN1/HRC/TPM1/CASQ2/PPP1R12B/ADRA2B/NPNT/MYOCD |
| Fibroblast | F2 RGS5+ Fib | GO:0086065 | cell communication involved in cardiac conduction | 7/176 | 57/16509 | 11.52 | 2.34e-06 | 0.000162 | 7 | GJC1/CALM2/CAV1/KCNA5/HRC/ATP1B2/CASQ2 |
| Fibroblast | F2 RGS5+ Fib | GO:1904705 | regulation of vascular associated smooth muscle cell proliferation | 7/176 | 60/16509 | 10.94 | 3.33e-06 | 0.000224 | 7 | RGS5/MEF2C/CNN1/TPM1/NDRG2/ADAMTS1/MYOCD |
| Fibroblast | F2 RGS5+ Fib | GO:1990874 | vascular associated smooth muscle cell proliferation | 7/176 | 62/16509 | 10.59 | 4.17e-06 | 0.000273 | 7 | RGS5/MEF2C/CNN1/TPM1/NDRG2/ADAMTS1/MYOCD |
| Fibroblast | F2 RGS5+ Fib | GO:0086064 | cell communication by electrical coupling involved in cardiac conduction | 5/176 | 25/16509 | 18.76 | 5.81e-06 | 0.00037 | 5 | GJC1/CALM2/CAV1/HRC/ATP1B2 |
| Fibroblast | F2 RGS5+ Fib | GO:0048738 | cardiac muscle tissue development | 12/176 | 233/16509 | 4.83 | 7.71e-06 | 0.000475 | 12 | MYH11/MEF2C/PLN/TBX2/GJC1/HEY2/PDGFRB/TPM1/SORBS2/FOXC2/ADAMTS9/MYOCD |
| Fibroblast | F2 RGS5+ Fib | GO:0061337 | cardiac conduction | 8/176 | 95/16509 | 7.9 | 7.86e-06 | 0.000475 | 8 | PLN/GJC1/CALM2/CAV1/KCNA5/HRC/ATP1B2/CASQ2 |
| Fibroblast | F2 RGS5+ Fib | GO:0006941 | striated muscle contraction | 11/176 | 197/16509 | 5.24 | 8.71e-06 | 0.000515 | 11 | PLN/GJC1/CALM2/CAV1/KCNA5/DMPK/HRC/TPM1/RCSD1/CASQ2/SNTA1 |
| Fibroblast | F2 RGS5+ Fib | GO:0061041 | regulation of wound healing | 9/176 | 129/16509 | 6.54 | 9.92e-06 | 0.000572 | 9 | PTK2/WFDC1/ANGPT2/C1QTNF1/CAV1/MYLK/PDGFA/SMOC2/FOXC2 |
| Fibroblast | F2 RGS5+ Fib | GO:0010927 | cellular component assembly involved in morphogenesis | 9/176 | 130/16509 | 6.49 | 1.06e-05 | 0.000581 | 9 | MYH11/CSRP2/MYL9/PLN/PDGFRB/TPM1/LDB3/LMOD1/CSRP1 |
| Fibroblast | F2 RGS5+ Fib | GO:0032989 | cellular anatomical entity morphogenesis | 9/176 | 130/16509 | 6.49 | 1.06e-05 | 0.000581 | 9 | MYH11/CSRP2/MYL9/PLN/PDGFRB/TPM1/LDB3/LMOD1/CSRP1 |
| Fibroblast | F2 RGS5+ Fib | GO:0003206 | cardiac chamber morphogenesis | 9/176 | 131/16509 | 6.44 | 1.12e-05 | 0.000601 | 9 | MEF2C/TBX2/HEY2/HEYL/TPM1/JAG1/CPE/ADAMTS1/FOXC2 |
| Fibroblast | F2 RGS5+ Fib | GO:0003208 | cardiac ventricle morphogenesis | 7/176 | 72/16509 | 9.12 | 1.14e-05 | 0.000601 | 7 | MEF2C/HEY2/HEYL/TPM1/JAG1/CPE/FOXC2 |
| Fibroblast | F2 RGS5+ Fib | GO:0003205 | cardiac chamber development | 10/176 | 173/16509 | 5.42 | 1.67e-05 | 0.00086 | 10 | MEF2C/TBX2/HEY2/HEYL/TPM1/JAG1/CPE/ADAMTS1/FOXC2/MYOCD |
| Fibroblast | F2 RGS5+ Fib | GO:0042060 | wound healing | 16/176 | 436/16509 | 3.44 | 1.79e-05 | 0.000903 | 16 | MCAM/MYL9/PTK2/GPR4/WFDC1/ANGPT2/C1QTNF1/CAV1/MYLK/PDGFA/TPM1/CSRP1/SMOC2/ADRA2B/ENTPD1/F… |
| Fibroblast | F2 RGS5+ Fib | GO:0030048 | actin filament-based movement | 9/176 | 141/16509 | 5.99 | 2.03e-05 | 0.000957 | 9 | ACTA2/PLN/GJC1/CAV1/KCNA5/TPM1/ACTN4/CASQ2/SNTA1 |
| Fibroblast | F2 RGS5+ Fib | GO:0031529 | ruffle organization | 6/176 | 53/16509 | 10.62 | 2.04e-05 | 0.000957 | 6 | CAV1/CSPG4/TPM1/PFN1/EPS8/EPS8L1 |
| Fibroblast | F2 RGS5+ Fib | GO:0010649 | regulation of cell communication by electrical coupling | 4/176 | 16/16509 | 23.45 | 2.06e-05 | 0.000957 | 4 | CALM2/CAV1/HRC/CASQ2 |
| Fibroblast | F2 RGS5+ Fib | GO:0072109 | glomerular mesangium development | 4/176 | 16/16509 | 23.45 | 2.06e-05 | 0.000957 | 4 | GPR4/ACTA2/PDGFRB/FOXC2 |
| Fibroblast | F2 RGS5+ Fib | GO:0014812 | muscle cell migration | 7/176 | 82/16509 | 8.01 | 2.69e-05 | 0.0012 | 7 | MEF2C/PDGFA/PDGFRB/NET1/TPM1/ADAMTS1/MYOCD |
| Fibroblast | F2 RGS5+ Fib | GO:0031589 | cell-substrate adhesion | 14/176 | 354/16509 | 3.71 | 2.7e-05 | 0.0012 | 14 | BCAM/PTK2/ITGA7/ANGPT2/LAMA5/ITGA3/ATP1B2/JAG1/NTN4/ACTN4/EGFLAM/NPNT/ARHGEF7/ADAMTS9 |
| Fibroblast | F2 RGS5+ Fib | GO:0014706 | striated muscle tissue development | 15/176 | 402/16509 | 3.5 | 2.73e-05 | 0.0012 | 15 | MYH11/MEF2C/PLN/TBX2/GJC1/HEY2/CAV2/CAV1/HEYL/PDGFRB/TPM1/SORBS2/FOXC2/ADAMTS9/MYOCD |
| Fibroblast | F2 RGS5+ Fib | GO:0060048 | cardiac muscle contraction | 9/176 | 147/16509 | 5.74 | 2.83e-05 | 0.00122 | 9 | PLN/GJC1/CALM2/CAV1/KCNA5/HRC/TPM1/CASQ2/SNTA1 |
| Fibroblast | F2 RGS5+ Fib | GO:0072132 | mesenchyme morphogenesis | 6/176 | 58/16509 | 9.7 | 3.45e-05 | 0.00146 | 6 | ACTA2/TBX2/HEY2/HEYL/ACTG2/FOXC2 |
| Fibroblast | F2 RGS5+ Fib | GO:0070252 | actin-mediated cell contraction | 8/176 | 118/16509 | 6.36 | 3.84e-05 | 0.00158 | 8 | ACTA2/PLN/GJC1/CAV1/KCNA5/TPM1/CASQ2/SNTA1 |

| compartment | subcluster | GO_ID | Description | GeneRatio | BgRatio | FoldEnrichment | pvalue | p_adjust | Count | geneID |
|---|---|---|---|---|---|---|---|---|---|---|
| Fibroblast | F2 RGS5+ Fib | GO:0060485 | mesenchyme development | 13/176 | 319/16509 | 3.82 | 3.87e-05 | 0.00158 | 13 | MEF2C/PTK2/ACTA2/TBX2/FRZB/HEY2/HEYL/LAMA5/SEMA5B/JAG1/ACTG2/TGFB1I1/FOXC2 |
| Fibroblast | F2 RGS5+ Fib | GO:0032271 | regulation of protein polymerization | 10/176 | 191/16509 | 4.91 | 3.91e-05 | 0.00158 | 10 | ESAM/TPPP3/PFN1/MAP1B/LMOD1/DAAM2/VASP/EPS8/FKBP4/ARHGEF7 |
| Fibroblast | F2 RGS5+ Fib | GO:0014910 | regulation of smooth muscle cell migration | 6/176 | 62/16509 | 9.08 | 5.05e-05 | 0.00201 | 6 | MEF2C/PDGFA/PDGFRB/TPM1/ADAMTS1/MYOCD |
| Fibroblast | F2 RGS5+ Fib | GO:0051145 | smooth muscle cell differentiation | 6/176 | 64/16509 | 8.79 | 6.06e-05 | 0.00236 | 6 | MEF2C/TBX2/HEY2/RBPMS2/NPNT/MYOCD |
| Fibroblast | F2 RGS5+ Fib | GO:0035904 | aorta development | 6/176 | 65/16509 | 8.66 | 6.61e-05 | 0.00254 | 6 | TBX2/HEY2/MYLK/PDGFRB/JAG1/ADAMTS9 |
| Fibroblast | F2 RGS5+ Fib | GO:0097178 | ruffle assembly | 5/176 | 41/16509 | 11.44 | 7.14e-05 | 0.0027 | 5 | CAV1/CSPG4/PFN1/EPS8/EPS8L1 |
| Fibroblast | F2 RGS5+ Fib | GO:1903034 | regulation of response to wounding | 9/176 | 166/16509 | 5.09 | 7.32e-05 | 0.00273 | 9 | PTK2/WFDC1/ANGPT2/C1QTNF1/CAV1/MYLK/PDGFA/SMOC2/FOXC2 |
| Fibroblast | F2 RGS5+ Fib | GO:0003231 | cardiac ventricle development | 8/176 | 130/16509 | 5.77 | 7.65e-05 | 0.00278 | 8 | MEF2C/HEY2/HEYL/TPM1/JAG1/CPE/FOXC2/MYOCD |
| Fibroblast | F2 RGS5+ Fib | GO:0031032 | actomyosin structure organization | 10/176 | 207/16509 | 4.53 | 7.7e-05 | 0.00278 | 10 | MYH11/CSRP2/MYL9/PDGFRB/CNN1/TPM1/PFN1/LDB3/LMOD1/CSRP1 |
| Fibroblast | F2 RGS5+ Fib | GO:0014909 | smooth muscle cell migration | 6/176 | 68/16509 | 8.28 | 8.53e-05 | 0.00304 | 6 | MEF2C/PDGFA/PDGFRB/TPM1/ADAMTS1/MYOCD |
| Fibroblast | F2 RGS5+ Fib | GO:0010959 | regulation of metal ion transport | 13/176 | 351/16509 | 3.47 | 0.000102 | 0.00359 | 13 | PLN/CALM2/CAV1/MYLK/PDGFRB/DMPK/HRC/TESC/ATP1B2/ACTN4/NGF/CASQ2/SNTA1 |
| Fibroblast | F2 RGS5+ Fib | GO:0010881 | regulation of cardiac muscle contraction by regulation of the release of sequestered calc… | 4/176 | 24/16509 | 15.63 | 0.000112 | 0.00388 | 4 | PLN/CALM2/HRC/CASQ2 |
| Fibroblast | F2 RGS5+ Fib | GO:0071560 | cellular response to transforming growth factor beta stimulus | 11/176 | 263/16509 | 3.92 | 0.000123 | 0.00419 | 11 | MEF2C/PTK2/ACTA2/CAV2/CAV1/ITGA3/TGFB1I1/COL4A2/LRRC32/NPNT/MYOCD |
| Fibroblast | F2 RGS5+ Fib | GO:0030198 | extracellular matrix organization | 12/176 | 310/16509 | 3.63 | 0.000126 | 0.00423 | 12 | MYH11/CAV2/CAV1/OLFML2A/NTN4/ADAMTS1/SMOC2/COL4A2/FOXC2/EGFLAM/NPNT/ADAMTS9 |
| Fibroblast | F2 RGS5+ Fib | GO:0043062 | extracellular structure organization | 12/176 | 311/16509 | 3.62 | 0.00013 | 0.0043 | 12 | MYH11/CAV2/CAV1/OLFML2A/NTN4/ADAMTS1/SMOC2/COL4A2/FOXC2/EGFLAM/NPNT/ADAMTS9 |
| Fibroblast | F2 RGS5+ Fib | GO:0090130 | tissue migration | 7/176 | 105/16509 | 6.25 | 0.000131 | 0.0043 | 7 | PTK2/ACTA2/ITGA3/PFN1/DAAM2/ACTG2/MAPRE2 |
| Fibroblast | F2 RGS5+ Fib | GO:0045229 | external encapsulating structure organization | 12/176 | 312/16509 | 3.61 | 0.000133 | 0.00431 | 12 | MYH11/CAV2/CAV1/OLFML2A/NTN4/ADAMTS1/SMOC2/COL4A2/FOXC2/EGFLAM/NPNT/ADAMTS9 |
| Fibroblast | F2 RGS5+ Fib | GO:1904753 | negative regulation of vascular associated smooth muscle cell migration | 3/176 | 10/16509 | 28.14 | 0.000135 | 0.00431 | 3 | MEF2C/TPM1/MYOCD |
| Fibroblast | F2 RGS5+ Fib | GO:0007266 | Rho protein signal transduction | 8/176 | 142/16509 | 5.28 | 0.000142 | 0.00446 | 8 | GPR4/NET1/ITGA3/ARHGAP29/ARHGDIB/EPS8/ARHGEF7/EPS8L1 |
| Fibroblast | F2 RGS5+ Fib | GO:0071559 | response to transforming growth factor beta | 11/176 | 270/16509 | 3.82 | 0.000155 | 0.00476 | 11 | MEF2C/PTK2/ACTA2/CAV2/CAV1/ITGA3/TGFB1I1/COL4A2/LRRC32/NPNT/MYOCD |
| Fibroblast | F2 RGS5+ Fib | GO:1904752 | regulation of vascular associated smooth muscle cell migration | 4/176 | 26/16509 | 14.43 | 0.000155 | 0.00476 | 4 | MEF2C/TPM1/ADAMTS1/MYOCD |
| Fibroblast | F2 RGS5+ Fib | GO:0051258 | protein polymerization | 11/176 | 274/16509 | 3.77 | 0.000176 | 0.00529 | 11 | ESAM/TPPP3/PFN1/MAP1B/LMOD1/DAAM2/VASP/EPS8/CASQ2/FKBP4/ARHGEF7 |
| Fibroblast | F2 RGS5+ Fib | GO:0032970 | regulation of actin filament-based process | 13/176 | 371/16509 | 3.29 | 0.000177 | 0.00529 | 13 | ESAM/ACTA2/PLN/DSTN/CAV1/PDGFA/PDGFRB/TPM1/PFN1/LMOD1/DAAM2/VASP/EPS8 |
| Fibroblast | F2 RGS5+ Fib | GO:1901844 | regulation of cell communication by electrical coupling involved in cardiac conduction | 3/176 | 11/16509 | 25.58 | 0.000185 | 0.00545 | 3 | CALM2/CAV1/HRC |
| Fibroblast | F2 RGS5+ Fib | GO:0035850 | epithelial cell differentiation involved in kidney development | 5/176 | 50/16509 | 9.38 | 0.000187 | 0.00545 | 5 | MEF2C/GPR4/ACTA2/JAG1/FOXC2 |

| compartment | subcluster | GO_ID | Description | GeneRatio | BgRatio | FoldEnrichment | pvalue | p_adjust | Count | geneID |
|---|---|---|---|---|---|---|---|---|---|---|
| Fibroblast | F2 RGS5+ Fib | GO:0035051 | cardiocyte differentiation | 8/176 | 150/16509 | 5.0 | 0.000207 | 0.00589 | 8 | MYH11/MEF2C/TBX2/HEY2/PDGFRB/JAG1/SORBS2/MYOCD |
| Fibroblast | F2 RGS5+ Fib | GO:0060055 | angiogenesis involved in wound healing | 4/176 | 28/16509 | 13.4 | 0.000209 | 0.00589 | 4 | MCAM/GPR4/SMOC2/FOXC2 |
| Fibroblast | F2 RGS5+ Fib | GO:1904738 | vascular associated smooth muscle cell migration | 4/176 | 28/16509 | 13.4 | 0.000209 | 0.00589 | 4 | MEF2C/TPM1/ADAMTS1/MYOCD |
| Fibroblast | F2 RGS5+ Fib | GO:0045664 | regulation of neuron differentiation | 9/176 | 192/16509 | 4.4 | 0.000221 | 0.00614 | 9 | NOTCH3/MEF2C/HEY2/HEYL/MAP1B/JAG1/NGF/ADRA2B/ID4 |
| Fibroblast | F2 RGS5+ Fib | GO:1900027 | regulation of ruffle assembly | 4/176 | 29/16509 | 12.94 | 0.000241 | 0.00649 | 4 | CAV1/PFN1/EPS8/EPS8L1 |
| Fibroblast | F2 RGS5+ Fib | GO:0003151 | outflow tract morphogenesis | 6/176 | 82/16509 | 6.86 | 0.000241 | 0.00649 | 6 | MEF2C/TBX2/HEY2/HEYL/JAG1/FOXC2 |
| Fibroblast | F2 RGS5+ Fib | GO:1904238 | pericyte cell differentiation | 3/176 | 12/16509 | 23.45 | 0.000244 | 0.00649 | 3 | GPR4/ACTA2/FOXC2 |
| Fibroblast | F2 RGS5+ Fib | GO:0007015 | actin filament organization | 14/176 | 436/16509 | 3.01 | 0.000246 | 0.00649 | 14 | ESAM/MYL9/TPM2/CALD1/DSTN/CNN1/TPM1/PFN1/LMOD1/DAAM2/VASP/ARPC1A/CAP2/EPS8 |
| Fibroblast | F2 RGS5+ Fib | GO:0086009 | membrane repolarization | 5/176 | 53/16509 | 8.85 | 0.000247 | 0.00649 | 5 | CAV1/KCNA5/ATP1B2/CASQ2/SNTA1 |
| Fibroblast | F2 RGS5+ Fib | GO:0090066 | regulation of anatomical structure size | 15/176 | 495/16509 | 2.84 | 0.000276 | 0.00717 | 15 | ESAM/ACTA2/DSTN/CAV1/KCNA5/PFN1/MAP1B/LMOD1/DAAM2/GRIP2/NGF/VASP/EPS8/ADRA2B/FOXC2 |
| Fibroblast | F2 RGS5+ Fib | GO:0001667 | ameboidal-type cell migration | 10/176 | 244/16509 | 3.84 | 0.000294 | 0.00758 | 10 | PTK2/ACTA2/LAMA5/ITGA3/PFN1/ARHGDIB/SEMA5B/DAAM2/MAPRE2/ARHGEF7 |
| Fibroblast | F2 RGS5+ Fib | GO:0086003 | cardiac muscle cell contraction | 6/176 | 86/16509 | 6.54 | 0.000313 | 0.00786 | 6 | PLN/GJC1/CAV1/KCNA5/CASQ2/SNTA1 |
| Fibroblast | F2 RGS5+ Fib | GO:0001765 | membrane raft assembly | 3/176 | 13/16509 | 21.65 | 0.000315 | 0.00786 | 3 | CAV2/CAV1/PACSIN2 |
| Fibroblast | F2 RGS5+ Fib | GO:0044857 | plasma membrane raft organization | 3/176 | 13/16509 | 21.65 | 0.000315 | 0.00786 | 3 | CAV2/CAV1/PACSIN2 |
| Fibroblast | F2 RGS5+ Fib | GO:0006940 | regulation of smooth muscle contraction | 5/176 | 57/16509 | 8.23 | 0.000348 | 0.00859 | 5 | CAV1/CNN1/ADRA2B/NPNT/MYOCD |
| Fibroblast | F2 RGS5+ Fib | GO:0035023 | regulation of Rho protein signal transduction | 6/176 | 88/16509 | 6.4 | 0.000355 | 0.00867 | 6 | GPR4/NET1/ITGA3/ARHGDIB/EPS8/EPS8L1 |
| Fibroblast | F2 RGS5+ Fib | GO:0032387 | negative regulation of intracellular transport | 5/176 | 58/16509 | 8.09 | 0.000377 | 0.00911 | 5 | PLN/CABP1/MAP1B/PKIG/CRYAB |
| Fibroblast | F2 RGS5+ Fib | GO:0048762 | mesenchymal cell differentiation | 10/176 | 252/16509 | 3.72 | 0.00038 | 0.00911 | 10 | MEF2C/PTK2/FRZB/HEY2/HEYL/LAMA5/SEMA5B/JAG1/TGFB1I1/FOXC2 |
| Fibroblast | F2 RGS5+ Fib | GO:0014902 | myotube differentiation | 7/176 | 125/16509 | 5.25 | 0.000385 | 0.00914 | 7 | MEF2C/CAV2/CACNA1H/DMPK/EHD2/SORT1/MYOCD |
| Fibroblast | F2 RGS5+ Fib | GO:0038180 | nerve growth factor signaling pathway | 3/176 | 14/16509 | 20.1 | 0.000398 | 0.00925 | 3 | SORT1/NGF/NTF3 |
| Fibroblast | F2 RGS5+ Fib | GO:1900029 | positive regulation of ruffle assembly | 3/176 | 14/16509 | 20.1 | 0.000398 | 0.00925 | 3 | PFN1/EPS8/EPS8L1 |
| Fibroblast | F2 RGS5+ Fib | GO:0010882 | regulation of cardiac muscle contraction by calcium ion signaling | 4/176 | 33/16509 | 11.37 | 0.000401 | 0.00925 | 4 | PLN/CALM2/HRC/CASQ2 |
| Fibroblast | F2 RGS5+ Fib | GO:0048008 | platelet-derived growth factor receptor signaling pathway | 5/176 | 59/16509 | 7.95 | 0.000409 | 0.00925 | 5 | PDGFA/PDGFRB/CSPG4/PTGIR/MYOCD |
| Fibroblast | F2 RGS5+ Fib | GO:0061005 | cell differentiation involved in kidney development | 5/176 | 59/16509 | 7.95 | 0.000409 | 0.00925 | 5 | MEF2C/GPR4/ACTA2/JAG1/FOXC2 |
| Fibroblast | F2 RGS5+ Fib | GO:0086002 | cardiac muscle cell action potential involved in contraction | 5/176 | 60/16509 | 7.82 | 0.000442 | 0.00988 | 5 | GJC1/CAV1/KCNA5/CASQ2/SNTA1 |
| Fibroblast | F2 RGS5+ Fib | GO:0022612 | gland morphogenesis | 7/176 | 128/16509 | 5.13 | 0.000445 | 0.00988 | 7 | CRIP1/TBX2/CAV1/PDGFA/LAMA5/NTN4/ID4 |
| Fibroblast | F2 RGS5+ Fib | GO:0045598 | regulation of fat cell differentiation | 7/176 | 130/16509 | 5.05 | 0.000488 | 0.0107 | 7 | ADIRF/FRZB/CCDC3/JAG1/SORT1/ID4/TGFB1I1 |
| Fibroblast | F2 RGS5+ Fib | GO:0010763 | positive regulation of fibroblast migration | 3/176 | 15/16509 | 18.76 | 0.000493 | 0.0107 | 3 | PTK2/ACTA2/ARHGEF7 |
| Fibroblast | F2 RGS5+ Fib | GO:0035791 | platelet-derived growth factor receptor-beta signaling pathway | 3/176 | 15/16509 | 18.76 | 0.000493 | 0.0107 | 3 | PDGFA/PDGFRB/MYOCD |
| Fibroblast | F2 RGS5+ Fib | GO:0014033 | neural crest cell differentiation | 6/176 | 94/16509 | 5.99 | 0.000505 | 0.0108 | 6 | MEF2C/FRZB/LAMA5/SEMA5B/JAG1/FOXC2 |
| Fibroblast | F2 RGS5+ Fib | GO:0003179 | heart valve morphogenesis | 5/176 | 62/16509 | 7.56 | 0.000515 | 0.0109 | 5 | MEF2C/HEY2/HEYL/JAG1/ADAMTS9 |
| Fibroblast | F2 RGS5+ Fib | GO:1902903 | regulation of supramolecular fiber organization | 12/176 | 362/16509 | 3.11 | 0.000517 | 0.0109 | 12 | ESAM/DSTN/TPM1/PFN1/MAP1B/LMOD1/DAAM2/VASP/CRYAB/EPS8/FKBP4/ARHGEF7 |

| compartment | subcluster | GO_ID | Description | GeneRatio | BgRatio | FoldEnrichment | pvalue | p_adjust | Count | geneID |
|---|---|---|---|---|---|---|---|---|---|---|
| Fibroblast | F2 RGS5+ Fib | GO:1904062 | regulation of monoatomic cation transmembrane transport | 10/176 | 264/16509 | 3.55 | 0.000547 | 0.0114 | 10 | PLN/CALM2/CAV1/HRC/TESC/ATP1B2/ACTN4/NGF/CASQ2/SNTA1 |
| Fibroblast | F2 RGS5+ Fib | GO:0003007 | heart morphogenesis | 10/176 | 265/16509 | 3.54 | 0.000563 | 0.0115 | 10 | MEF2C/PTK2/TBX2/HEY2/HEYL/TPM1/JAG1/CPE/ADAMTS1/FOXC2 |
| Fibroblast | F2 RGS5+ Fib | GO:0010880 | regulation of release of sequestered calcium ion into cytosol by sarcoplasmic reticulum | 4/176 | 36/16509 | 10.42 | 0.000563 | 0.0115 | 4 | PLN/CALM2/HRC/CASQ2 |
| Fibroblast | F2 RGS5+ Fib | GO:0060306 | regulation of membrane repolarization | 4/176 | 36/16509 | 10.42 | 0.000563 | 0.0115 | 4 | CAV1/KCNA5/CASQ2/SNTA1 |
| Fibroblast | F2 RGS5+ Fib | GO:0043269 | regulation of monoatomic ion transport | 13/176 | 420/16509 | 2.9 | 0.00058 | 0.0117 | 13 | PLN/CALM2/CAV1/MYLK/PDGFRB/DMPK/HRC/TESC/ATP1B2/ACTN4/NGF/CASQ2/SNTA1 |
| Fibroblast | F2 RGS5+ Fib | GO:0006942 | regulation of striated muscle contraction | 6/176 | 100/16509 | 5.63 | 0.000702 | 0.014 | 6 | PLN/CALM2/CAV1/DMPK/HRC/CASQ2 |
| Fibroblast | F2 RGS5+ Fib | GO:0008064 | regulation of actin polymerization or depolymerization | 7/176 | 140/16509 | 4.69 | 0.000759 | 0.0151 | 7 | ESAM/DSTN/PFN1/LMOD1/DAAM2/VASP/EPS8 |
| Fibroblast | F2 RGS5+ Fib | GO:1902305 | regulation of sodium ion transmembrane transport | 4/176 | 39/16509 | 9.62 | 0.000767 | 0.0151 | 4 | TESC/ATP1B2/ACTN4/SNTA1 |
| Fibroblast | F2 RGS5+ Fib | GO:0030832 | regulation of actin filament length | 7/176 | 143/16509 | 4.59 | 0.000861 | 0.0168 | 7 | ESAM/DSTN/PFN1/LMOD1/DAAM2/VASP/EPS8 |
| Fibroblast | F2 RGS5+ Fib | GO:0097553 | calcium ion transmembrane import into cytosol | 8/176 | 186/16509 | 4.03 | 0.000869 | 0.0168 | 8 | PLN/CALM2/CAV1/CACNA1H/PLCE1/HRC/NGF/CASQ2 |
| Fibroblast | F2 RGS5+ Fib | GO:0014808 | release of sequestered calcium ion into cytosol by sarcoplasmic reticulum | 4/176 | 41/16509 | 9.15 | 0.000929 | 0.0179 | 4 | PLN/CALM2/HRC/CASQ2 |
| Fibroblast | F2 RGS5+ Fib | GO:0002028 | regulation of sodium ion transport | 5/176 | 71/16509 | 6.61 | 0.00096 | 0.0183 | 5 | DMPK/TESC/ATP1B2/ACTN4/SNTA1 |
| Fibroblast | F2 RGS5+ Fib | GO:0055012 | ventricular cardiac muscle cell differentiation | 3/176 | 19/16509 | 14.81 | 0.00102 | 0.0188 | 3 | MEF2C/HEY2/MYOCD |
| Fibroblast | F2 RGS5+ Fib | GO:0061042 | vascular wound healing | 3/176 | 19/16509 | 14.81 | 0.00102 | 0.0188 | 3 | MCAM/SMOC2/FOXC2 |
| Fibroblast | F2 RGS5+ Fib | GO:1902307 | positive regulation of sodium ion transmembrane transport | 3/176 | 19/16509 | 14.81 | 0.00102 | 0.0188 | 3 | TESC/ATP1B2/ACTN4 |
| Fibroblast | F2 RGS5+ Fib | GO:2000811 | negative regulation of anoikis | 3/176 | 19/16509 | 14.81 | 0.00102 | 0.0188 | 3 | PTK2/CAV1/TLE1 |
| Fibroblast | F2 RGS5+ Fib | GO:0019722 | calcium-mediated signaling | 9/176 | 238/16509 | 3.55 | 0.00104 | 0.0192 | 9 | RCAN2/PLN/CALM2/PLCE1/PDGFRB/DMPK/HRC/NUDT4/CASQ2 |
| Fibroblast | F2 RGS5+ Fib | GO:1903514 | release of sequestered calcium ion into cytosol by endoplasmic reticulum | 4/176 | 43/16509 | 8.73 | 0.00111 | 0.0203 | 4 | PLN/CALM2/HRC/CASQ2 |
| Fibroblast | F2 RGS5+ Fib | GO:0014855 | striated muscle cell proliferation | 5/176 | 74/16509 | 6.34 | 0.00116 | 0.0209 | 5 | MEF2C/TBX2/HEY2/CAV2/FOXC2 |
| Fibroblast | F2 RGS5+ Fib | GO:0003184 | pulmonary valve morphogenesis | 3/176 | 20/16509 | 14.07 | 0.00119 | 0.0212 | 3 | HEY2/HEYL/JAG1 |
| Fibroblast | F2 RGS5+ Fib | GO:0006123 | mitochondrial electron transport, cytochrome c to oxygen | 3/176 | 20/16509 | 14.07 | 0.00119 | 0.0212 | 3 | COX4I2/COX7A2/CYCS |
| Fibroblast | F2 RGS5+ Fib | GO:0045445 | myoblast differentiation | 6/176 | 111/16509 | 5.07 | 0.00121 | 0.0214 | 6 | MEF2C/TBX2/EPAS1/ID3/JAG1/MYOCD |
| Fibroblast | F2 RGS5+ Fib | GO:1902904 | negative regulation of supramolecular fiber organization | 7/176 | 152/16509 | 4.32 | 0.00123 | 0.0216 | 7 | PFN1/MAP1B/LMOD1/CRYAB/EPS8/FKBP4/ARHGEF7 |
| Fibroblast | F2 RGS5+ Fib | GO:0034765 | regulation of monoatomic ion transmembrane transport | 10/176 | 294/16509 | 3.19 | 0.00124 | 0.0217 | 10 | PLN/CALM2/CAV1/HRC/TESC/ATP1B2/ACTN4/NGF/CASQ2/SNTA1 |
| Fibroblast | F2 RGS5+ Fib | GO:0055017 | cardiac muscle tissue growth | 5/176 | 76/16509 | 6.17 | 0.0013 | 0.0226 | 5 | MEF2C/TBX2/HEY2/SORBS2/FOXC2 |
| Fibroblast | F2 RGS5+ Fib | GO:0003170 | heart valve development | 5/176 | 77/16509 | 6.09 | 0.00138 | 0.0238 | 5 | MEF2C/HEY2/HEYL/JAG1/ADAMTS9 |
| Fibroblast | F2 RGS5+ Fib | GO:0070296 | sarcoplasmic reticulum calcium ion transport | 4/176 | 47/16509 | 7.98 | 0.00156 | 0.0266 | 4 | PLN/CALM2/HRC/CASQ2 |
| Fibroblast | F2 RGS5+ Fib | GO:0060445 | branching involved in salivary gland morphogenesis | 3/176 | 22/16509 | 12.79 | 0.00158 | 0.0268 | 3 | PDGFA/LAMA5/NTN4 |
| Fibroblast | F2 RGS5+ Fib | GO:0051651 | maintenance of location in cell | 8/176 | 205/16509 | 3.66 | 0.00162 | 0.0272 | 8 | PLN/CHCHD10/CALM2/CAV1/PLCE1/HRC/NGF/CASQ2 |
| Fibroblast | F2 RGS5+ Fib | GO:0051056 | regulation of small GTPase mediated signal transduction | 10/176 | 305/16509 | 3.08 | 0.00163 | 0.0273 | 10 | GPR4/NET1/ITGA3/ARHGAP29/ARHGDIB/NGF/ARHGEF17/EPS8/MAPRE2/EPS8L1 |
| Fibroblast | F2 RGS5+ Fib | GO:0001974 | blood vessel remodeling | 4/176 | 48/16509 | 7.82 | 0.00169 | 0.0276 | 4 | MEF2C/EPAS1/JAG1/FOXC2 |

| compartment | subcluster | GO_ID | Description | GeneRatio | BgRatio | FoldEnrichment | pvalue | p_adjust | Count | geneID |
|---|---|---|---|---|---|---|---|---|---|---|
| Fibroblast | F2 RGS5+ Fib | GO:0010634 | positive regulation of epithelial cell migration | 4/176 | 48/16509 | 7.82 | 0.00169 | 0.0276 | 4 | PTK2/ITGA3/PFN1/MAPRE2 |
| Fibroblast | F2 RGS5+ Fib | GO:0038084 | vascular endothelial growth factor signaling pathway | 4/176 | 48/16509 | 7.82 | 0.00169 | 0.0276 | 4 | PTP4A3/PGF/PDGFRB/SMOC2 |
| Fibroblast | F2 RGS5+ Fib | GO:0048863 | stem cell differentiation | 9/176 | 256/16509 | 3.3 | 0.00173 | 0.0276 | 9 | MEF2C/TBX2/FRZB/LAMA5/SEMA5B/LBH/JAG1/FOXC2/MYOCD |
| Fibroblast | F2 RGS5+ Fib | GO:0001570 | vasculogenesis | 5/176 | 81/16509 | 5.79 | 0.00173 | 0.0276 | 5 | GJC1/HEY2/CAV1/PDGFRB/MYOCD |
| Fibroblast | F2 RGS5+ Fib | GO:0051279 | regulation of release of sequestered calcium ion into cytosol | 5/176 | 81/16509 | 5.79 | 0.00173 | 0.0276 | 5 | PLN/CALM2/HRC/NGF/CASQ2 |
| Fibroblast | F2 RGS5+ Fib | GO:0055117 | regulation of cardiac muscle contraction | 5/176 | 81/16509 | 5.79 | 0.00173 | 0.0276 | 5 | PLN/CALM2/CAV1/HRC/CASQ2 |
| Fibroblast | F2 RGS5+ Fib | GO:0055007 | cardiac muscle cell differentiation | 6/176 | 120/16509 | 4.69 | 0.00181 | 0.0284 | 6 | MYH11/MEF2C/HEY2/PDGFRB/SORBS2/MYOCD |
| Fibroblast | F2 RGS5+ Fib | GO:0045214 | sarcomere organization | 4/176 | 49/16509 | 7.66 | 0.00182 | 0.0284 | 4 | CSRP2/TPM1/LDB3/CSRP1 |
| Fibroblast | F2 RGS5+ Fib | GO:0045599 | negative regulation of fat cell differentiation | 4/176 | 49/16509 | 7.66 | 0.00182 | 0.0284 | 4 | JAG1/SORT1/ID4/TGFB1I1 |
| Fibroblast | F2 RGS5+ Fib | GO:0030833 | regulation of actin filament polymerization | 6/176 | 121/16509 | 4.65 | 0.00188 | 0.0292 | 6 | ESAM/PFN1/LMOD1/DAAM2/VASP/EPS8 |
| Fibroblast | F2 RGS5+ Fib | GO:0072009 | nephron epithelium development | 6/176 | 122/16509 | 4.61 | 0.00196 | 0.0303 | 6 | MEF2C/HEYL/LAMA5/JAG1/FOXC2/NPNT |
| Fibroblast | F2 RGS5+ Fib | GO:0090287 | regulation of cellular response to growth factor stimulus | 10/176 | 314/16509 | 2.99 | 0.00202 | 0.0306 | 10 | PTP4A3/CAV2/CAV1/ITGA3/SMOC2/RBPMS2/TGFB1I1/LRRC32/NPNT/MYOCD |
| Fibroblast | F2 RGS5+ Fib | GO:0060419 | heart growth | 5/176 | 84/16509 | 5.58 | 0.00204 | 0.0306 | 5 | MEF2C/TBX2/HEY2/SORBS2/FOXC2 |
| Fibroblast | F2 RGS5+ Fib | GO:0086001 | cardiac muscle cell action potential | 5/176 | 84/16509 | 5.58 | 0.00204 | 0.0306 | 5 | GJC1/CAV1/KCNA5/CASQ2/SNTA1 |
| Fibroblast | F2 RGS5+ Fib | GO:0003177 | pulmonary valve development | 3/176 | 24/16509 | 11.73 | 0.00204 | 0.0306 | 3 | HEY2/HEYL/JAG1 |
| Fibroblast | F2 RGS5+ Fib | GO:0014912 | negative regulation of smooth muscle cell migration | 3/176 | 24/16509 | 11.73 | 0.00204 | 0.0306 | 3 | MEF2C/TPM1/MYOCD |
| Fibroblast | F2 RGS5+ Fib | GO:0060038 | cardiac muscle cell proliferation | 4/176 | 51/16509 | 7.36 | 0.00211 | 0.0314 | 4 | MEF2C/TBX2/HEY2/FOXC2 |
| Fibroblast | F2 RGS5+ Fib | GO:0032273 | positive regulation of protein polymerization | 5/176 | 85/16509 | 5.52 | 0.00215 | 0.0317 | 5 | TPPP3/PFN1/MAP1B/LMOD1/VASP |
| Fibroblast | F2 RGS5+ Fib | GO:0051235 | maintenance of location | 8/176 | 215/16509 | 3.49 | 0.00218 | 0.032 | 8 | PLN/CHCHD10/CALM2/CAV1/PLCE1/HRC/NGF/CASQ2 |
| Fibroblast | F2 RGS5+ Fib | GO:0042310 | vasoconstriction | 5/176 | 86/16509 | 5.45 | 0.00226 | 0.0324 | 5 | ACTA2/CAV1/KCNA5/GRIP2/ADRA2B |
| Fibroblast | F2 RGS5+ Fib | GO:0055006 | cardiac cell development | 5/176 | 86/16509 | 5.45 | 0.00226 | 0.0324 | 5 | MYH11/HEY2/PDGFRB/JAG1/SORBS2 |
| Fibroblast | F2 RGS5+ Fib | GO:0003197 | endocardial cushion development | 4/176 | 52/16509 | 7.22 | 0.00227 | 0.0324 | 4 | TBX2/HEY2/HEYL/JAG1 |
| Fibroblast | F2 RGS5+ Fib | GO:0050926 | regulation of positive chemotaxis | 3/176 | 25/16509 | 11.26 | 0.00231 | 0.0324 | 3 | PGF/ANGPT2/NTF3 |
| Fibroblast | F2 RGS5+ Fib | GO:0060045 | positive regulation of cardiac muscle cell proliferation | 3/176 | 25/16509 | 11.26 | 0.00231 | 0.0324 | 3 | MEF2C/TBX2/HEY2 |
| Fibroblast | F2 RGS5+ Fib | GO:1905207 | regulation of cardiocyte differentiation | 3/176 | 25/16509 | 11.26 | 0.00231 | 0.0324 | 3 | MEF2C/HEY2/MYOCD |
| Fibroblast | F2 RGS5+ Fib | GO:2000209 | regulation of anoikis | 3/176 | 25/16509 | 11.26 | 0.00231 | 0.0324 | 3 | PTK2/CAV1/TLE1 |
| Fibroblast | F2 RGS5+ Fib | GO:0010810 | regulation of cell-substrate adhesion | 8/176 | 218/16509 | 3.44 | 0.00238 | 0.0333 | 8 | PTK2/ANGPT2/ITGA3/JAG1/ACTN4/EGFLAM/NPNT/ARHGEF7 |
| Fibroblast | F2 RGS5+ Fib | GO:0014829 | vascular associated smooth muscle contraction | 3/176 | 26/16509 | 10.82 | 0.00259 | 0.036 | 3 | ACTA2/GRIP2/ADRA2B |
| Fibroblast | F2 RGS5+ Fib | GO:0051209 | release of sequestered calcium ion into cytosol | 6/176 | 129/16509 | 4.36 | 0.0026 | 0.036 | 6 | PLN/CALM2/PLCE1/HRC/NGF/CASQ2 |
| Fibroblast | F2 RGS5+ Fib | GO:0007264 | small GTPase-mediated signal transduction | 13/176 | 498/16509 | 2.45 | 0.00267 | 0.0368 | 13 | GPR4/RASGRP2/PLCE1/NET1/ITGA3/ARHGAP29/ARHGDIB/NGF/ARHGEF17/EPS8/MAPRE2/ARHGEF7/EPS8L1 |
| Fibroblast | F2 RGS5+ Fib | GO:0051283 | negative regulation of sequestering of calcium ion | 6/176 | 130/16509 | 4.33 | 0.0027 | 0.037 | 6 | PLN/CALM2/PLCE1/HRC/NGF/CASQ2 |
| Fibroblast | F2 RGS5+ Fib | GO:0003018 | vascular process in circulatory system | 9/176 | 274/16509 | 3.08 | 0.00274 | 0.037 | 9 | PTP4A3/GPR4/ACTA2/CAV1/KCNA5/ATP1B2/GRIP2/ADRA2B/FOXC2 |
| Fibroblast | F2 RGS5+ Fib | GO:0008154 | actin polymerization or depolymerization | 7/176 | 175/16509 | 3.75 | 0.00274 | 0.037 | 7 | ESAM/DSTN/PFN1/LMOD1/DAAM2/VASP/EPS8 |
| Fibroblast | F2 RGS5+ Fib | GO:1901879 | regulation of protein depolymerization | 5/176 | 90/16509 | 5.21 | 0.00275 | 0.037 | 5 | DSTN/MAP1B/LMOD1/EPS8/NES |

| compartment | subcluster | GO_ID | Description | GeneRatio | BgRatio | FoldEnrichment | pvalue | p_adjust | Count | geneID |
|---|---|---|---|---|---|---|---|---|---|---|
| Fibroblast | F2 RGS5+ Fib | GO:0032956 | regulation of actin cytoskeleton organization | 10/176 | 328/16509 | 2.86 | 0.00277 | 0.0371 | 10 | ESAM/DSTN/PDGFA/PDGFRB/TPM1/PFN1/LMOD1/DAAM2/VASP/EPS8 |
| Fibroblast | F2 RGS5+ Fib | GO:0051282 | regulation of sequestering of calcium ion | 6/176 | 131/16509 | 4.3 | 0.00281 | 0.0374 | 6 | PLN/CALM2/PLCE1/HRC/NGF/CASQ2 |
| Fibroblast | F2 RGS5+ Fib | GO:0010460 | positive regulation of heart rate | 3/176 | 27/16509 | 10.42 | 0.00289 | 0.0378 | 3 | HEY2/HRC/TPM1 |
| Fibroblast | F2 RGS5+ Fib | GO:0099623 | regulation of cardiac muscle cell membrane repolarization | 3/176 | 27/16509 | 10.42 | 0.00289 | 0.0378 | 3 | KCNA5/CASQ2/SNTA1 |
| Fibroblast | F2 RGS5+ Fib | GO:0051924 | regulation of calcium ion transport | 8/176 | 225/16509 | 3.34 | 0.00289 | 0.0378 | 8 | PLN/CALM2/CAV1/MYLK/PDGFRB/HRC/NGF/CASQ2 |
| Fibroblast | F2 RGS5+ Fib | GO:0051480 | regulation of cytosolic calcium ion concentration | 4/176 | 56/16509 | 6.7 | 0.00298 | 0.0388 | 4 | PLN/CAV2/CAV1/HRC |
| Fibroblast | F2 RGS5+ Fib | GO:0043542 | endothelial cell migration | 8/176 | 228/16509 | 3.29 | 0.00313 | 0.0405 | 8 | PTP4A3/MEF2C/PTK2/ANGPT2/PACSIN2/SMOC2/FOXC2/ADAMTS9 |
| Fibroblast | F2 RGS5+ Fib | GO:0045666 | positive regulation of neuron differentiation | 5/176 | 93/16509 | 5.04 | 0.00318 | 0.0409 | 5 | MEF2C/HEYL/MAP1B/NGF/ADRA2B |
| Fibroblast | F2 RGS5+ Fib | GO:0003272 | endocardial cushion formation | 3/176 | 28/16509 | 10.05 | 0.00321 | 0.0409 | 3 | TBX2/HEY2/HEYL |
| Fibroblast | F2 RGS5+ Fib | GO:0031579 | membrane raft organization | 3/176 | 28/16509 | 10.05 | 0.00321 | 0.0409 | 3 | CAV2/CAV1/PACSIN2 |
| Fibroblast | F2 RGS5+ Fib | GO:0051208 | sequestering of calcium ion | 6/176 | 135/16509 | 4.17 | 0.00326 | 0.0413 | 6 | PLN/CALM2/PLCE1/HRC/NGF/CASQ2 |
| Fibroblast | F2 RGS5+ Fib | GO:0051495 | positive regulation of cytoskeleton organization | 7/176 | 182/16509 | 3.61 | 0.00341 | 0.0429 | 7 | DSTN/TPM1/PFN1/MAP1B/LMOD1/VASP/NES |
| Fibroblast | F2 RGS5+ Fib | GO:0045987 | positive regulation of smooth muscle contraction | 3/176 | 29/16509 | 9.7 | 0.00355 | 0.0443 | 3 | ADRA2B/NPNT/MYOCD |
| Fibroblast | F2 RGS5+ Fib | GO:0055023 | positive regulation of cardiac muscle tissue growth | 3/176 | 29/16509 | 9.7 | 0.00355 | 0.0443 | 3 | MEF2C/TBX2/HEY2 |
| Fibroblast | F2 RGS5+ Fib | GO:0061448 | connective tissue development | 9/176 | 288/16509 | 2.93 | 0.0038 | 0.0468 | 9 | CRIP1/MEF2C/GPR4/ACTA2/FRZB/PDGFRB/ID4/FOXC2/ARID5A |
| Fibroblast | F2 RGS5+ Fib | GO:0010631 | epithelial cell migration | 5/176 | 97/16509 | 4.84 | 0.00381 | 0.0468 | 5 | PTK2/ITGA3/PFN1/DAAM2/MAPRE2 |
| Fibroblast | F2 RGS5+ Fib | GO:0034446 | substrate adhesion-dependent cell spreading | 5/176 | 97/16509 | 4.84 | 0.00381 | 0.0468 | 5 | PTK2/LAMA5/NTN4/ACTN4/ARHGEF7 |
| Fibroblast | F2 RGS5+ Fib | GO:0017015 | regulation of transforming growth factor beta receptor signaling pathway | 6/176 | 140/16509 | 4.02 | 0.0039 | 0.0477 | 6 | CAV2/ITGA3/TGFB1I1/LRRC32/NPNT/MYOCD |
| Fibroblast | F2 RGS5+ Fib | GO:0030168 | platelet activation | 6/176 | 141/16509 | 3.99 | 0.00404 | 0.0487 | 6 | MYL9/C1QTNF1/PDGFA/CSRP1/ADRA2B/ENTPD1 |
| Fibroblast | F2 RGS5+ Fib | GO:0051147 | regulation of muscle cell differentiation | 6/176 | 141/16509 | 3.99 | 0.00404 | 0.0487 | 6 | MEF2C/HEY2/DMPK/RBPMS2/YBX1/MYOCD |
| Fibroblast | F2 RGS5+ Fib | GO:0007160 | cell-matrix adhesion | 8/176 | 238/16509 | 3.15 | 0.00406 | 0.0487 | 8 | BCAM/PTK2/ITGA7/ITGA3/JAG1/NPNT/ARHGEF7/ADAMTS9 |
| Fibroblast | F2 RGS5+ Fib | GO:0090303 | positive regulation of wound healing | 4/176 | 61/16509 | 6.15 | 0.00406 | 0.0487 | 4 | PTK2/MYLK/SMOC2/FOXC2 |
| Fibroblast | F2 RGS5+ Fib | GO:0031346 | positive regulation of cell projection organization | 10/176 | 347/16509 | 2.7 | 0.00413 | 0.0488 | 10 | SERPINI1/PLCE1/ITGA3/ATP1B2/PFN1/MAP1B/NGF/EPS8/ARHGEF7/EPS8L1 |
| Fibroblast | F2 RGS5+ Fib | GO:0072080 | nephron tubule development | 5/176 | 99/16509 | 4.74 | 0.00415 | 0.0488 | 5 | MEF2C/HEYL/LAMA5/JAG1/NPNT |
| Fibroblast | F2 RGS5+ Fib | GO:0034762 | regulation of transmembrane transport | 11/176 | 405/16509 | 2.55 | 0.00418 | 0.0488 | 11 | PLN/OAZ2/CALM2/CAV1/HRC/TESC/ATP1B2/ACTN4/NGF/CASQ2/SNTA1 |
| Fibroblast | F2 RGS5+ Fib | GO:0051051 | negative regulation of transport | 11/176 | 405/16509 | 2.55 | 0.00418 | 0.0488 | 11 | PLN/OAZ2/CALM2/CAV1/CABP1/PACSIN2/MAP1B/PKIG/CRYAB/CASQ2/ADRA2B |
| Fibroblast | F2 RGS5+ Fib | GO:0045444 | fat cell differentiation | 8/176 | 240/16509 | 3.13 | 0.00427 | 0.0488 | 8 | ADIRF/FRZB/CCDC3/ADGRF5/JAG1/SORT1/ID4/TGFB1I1 |
| Fibroblast | F2 RGS5+ Fib | GO:0030511 | positive regulation of transforming growth factor beta receptor signaling pathway | 3/176 | 31/16509 | 9.08 | 0.0043 | 0.0488 | 3 | TGFB1I1/NPNT/MYOCD |
| Fibroblast | F2 RGS5+ Fib | GO:0086011 | membrane repolarization during action potential | 3/176 | 31/16509 | 9.08 | 0.0043 | 0.0488 | 3 | CAV1/KCNA5/CASQ2 |
| Fibroblast | F2 RGS5+ Fib | GO:0086019 | cell-cell signaling involved in cardiac conduction | 3/176 | 31/16509 | 9.08 | 0.0043 | 0.0488 | 3 | GJC1/KCNA5/CASQ2 |
| Fibroblast | F2 RGS5+ Fib | GO:1903846 | positive regulation of cellular response to transforming growth factor beta stimulus | 3/176 | 31/16509 | 9.08 | 0.0043 | 0.0488 | 3 | TGFB1I1/NPNT/MYOCD |

| compartment | subcluster | GO_ID | Description | GeneRatio | BgRatio | FoldEnrichment | pvalue | p_adjust | Count | geneID |
|---|---|---|---|---|---|---|---|---|---|---|
| Fibroblast | F2 RGS5+ Fib | GO:0003229 | ventricular cardiac muscle tissue development | 4/176 | 62/16509 | 6.05 | 0.0043 | 0.0488 | 4 | HEY2/TPM1/FOXC2/ADAMTS9 |
| Fibroblast | F2 RGS5+ Fib | GO:0051148 | negative regulation of muscle cell differentiation | 4/176 | 62/16509 | 6.05 | 0.0043 | 0.0488 | 4 | HEY2/RBPMS2/YBX1/MYOCD |
| Fibroblast | F2 RGS5+ Fib | GO:1903844 | regulation of cellular response to transforming growth factor beta stimulus | 6/176 | 143/16509 | 3.94 | 0.00432 | 0.0488 | 6 | CAV2/ITGA3/TGFB1I1/LRRC32/NPNT/MYOCD |
| Fibroblast | F2 RGS5+ Fib | GO:0090132 | epithelium migration | 5/176 | 100/16509 | 4.69 | 0.00433 | 0.0488 | 5 | PTK2/ITGA3/PFN1/DAAM2/MAPRE2 |
| Fibroblast | F2 RGS5+ Fib | GO:0007596 | blood coagulation | 8/176 | 241/16509 | 3.11 | 0.00437 | 0.049 | 8 | MYL9/ANGPT2/C1QTNF1/CAV1/PDGFA/CSRP1/ADRA2B/ENTPD1 |
| Fibroblast | F2 RGS5+ Fib | GO:0006874 | intracellular calcium ion homeostasis | 9/176 | 295/16509 | 2.86 | 0.00445 | 0.0496 | 9 | PLN/CALM2/CAV2/CAV1/PLCE1/DMPK/HRC/NGF/CASQ2 |
| Fibroblast | F3 COL11A1+ mCAF | GO:0030199 | collagen fibril organization | 20/182 | 69/16509 | 26.29 | 1.76e-23 | 4.8e-20 | 20 | COL11A1/MMP11/COL5A1/COL1A1/COL3A1/LOXL2/COL5A2/COL1A2/SERPINH1/AEBP1/LOXL1/ADAMTS12/COL1… |
| Fibroblast | F3 COL11A1+ mCAF | GO:0030198 | extracellular matrix organization | 32/182 | 310/16509 | 9.36 | 4.13e-22 | 3.44e-19 | 32 | POSTN/COL11A1/SULF1/MMP11/COL5A1/COL1A1/COL3A1/LOXL2/COL5A2/COL1A2/PDPN/SERPINH1/ANTXR1/M… |
| Fibroblast | F3 COL11A1+ mCAF | GO:0043062 | extracellular structure organization | 32/182 | 311/16509 | 9.33 | 4.56e-22 | 3.44e-19 | 32 | POSTN/COL11A1/SULF1/MMP11/COL5A1/COL1A1/COL3A1/LOXL2/COL5A2/COL1A2/PDPN/SERPINH1/ANTXR1/M… |
| Fibroblast | F3 COL11A1+ mCAF | GO:0045229 | external encapsulating structure organization | 32/182 | 312/16509 | 9.3 | 5.03e-22 | 3.44e-19 | 32 | POSTN/COL11A1/SULF1/MMP11/COL5A1/COL1A1/COL3A1/LOXL2/COL5A2/COL1A2/PDPN/SERPINH1/ANTXR1/M… |
| Fibroblast | F3 COL11A1+ mCAF | GO:0032963 | collagen metabolic process | 17/182 | 105/16509 | 14.69 | 1.74e-15 | 9.51e-13 | 17 | INHBA/MMP11/COL5A1/COL1A1/COL1A2/CTSB/SERPINH1/MMP14/F2R/RCN3/FAP/CTSK/P3H4/COL6A1/MMP7/M… |
| Fibroblast | F3 COL11A1+ mCAF | GO:0035987 | endodermal cell differentiation | 12/182 | 51/16509 | 21.34 | 2.44e-13 | 1.11e-10 | 12 | COL11A1/INHBA/COL5A1/COL5A2/FN1/MMP14/COL12A1/MYH9/COL8A1/COL6A1/COL7A1/MMP2 |
| Fibroblast | F3 COL11A1+ mCAF | GO:0051216 | cartilage development | 19/182 | 203/16509 | 8.49 | 1.02e-12 | 4e-10 | 19 | COL11A1/SULF1/COL1A1/COL3A1/LOXL2/SERPINH1/BGN/ADAMTS12/MATN3/CTSK/LUM/COL6A1/RUNX2/TGFBI… |
| Fibroblast | F3 COL11A1+ mCAF | GO:0001706 | endoderm formation | 12/182 | 60/16509 | 18.14 | 1.98e-12 | 6.76e-10 | 12 | COL11A1/INHBA/COL5A1/COL5A2/FN1/MMP14/COL12A1/MYH9/COL8A1/COL6A1/COL7A1/MMP2 |
| Fibroblast | F3 COL11A1+ mCAF | GO:0001704 | formation of primary germ layer | 15/182 | 128/16509 | 10.63 | 1.08e-11 | 3.28e-09 | 15 | COL11A1/INHBA/COL5A1/COL5A2/FN1/MMP14/GJA1/COL12A1/MYH9/LEF1/COL8A1/COL6A1/COL7A1/MMP2/WN… |
| Fibroblast | F3 COL11A1+ mCAF | GO:0061448 | connective tissue development | 20/182 | 288/16509 | 6.3 | 6.18e-11 | 1.69e-08 | 20 | COL11A1/SULF1/COL5A1/COL1A1/COL3A1/LOXL2/SERPINH1/BGN/ADAMTS12/MATN3/CTSK/LUM/COL6A1/RUNX… |
| Fibroblast | F3 COL11A1+ mCAF | GO:0007492 | endoderm development | 12/182 | 88/16509 | 12.37 | 2.22e-10 | 5.52e-08 | 12 | COL11A1/INHBA/COL5A1/COL5A2/FN1/MMP14/COL12A1/MYH9/COL8A1/COL6A1/COL7A1/MMP2 |
| Fibroblast | F3 COL11A1+ mCAF | GO:0007369 | gastrulation | 15/182 | 192/16509 | 7.09 | 3.43e-09 | 7.83e-07 | 15 | COL11A1/INHBA/COL5A1/COL5A2/FN1/MMP14/GJA1/COL12A1/MYH9/LEF1/COL8A1/COL6A1/COL7A1/MMP2/WN… |
| Fibroblast | F3 COL11A1+ mCAF | GO:0060348 | bone development | 15/182 | 210/16509 | 6.48 | 1.16e-08 | 2.36e-06 | 15 | SULF1/COL1A1/COL3A1/PDGFC/SERPINH1/MMP14/GJA1/BGN/ADAMTS12/COL6A1/RUNX2/SULF2/PPIB/EPYC/T… |

| compartment | subcluster | GO_ID | Description | GeneRatio | BgRatio | FoldEnrichment | pvalue | p_adjust | Count | geneID |
|---|---|---|---|---|---|---|---|---|---|---|
| Fibroblast | F3 COL11A1+ mCAF | GO:0010755 | regulation of plasminogen activation | 6/182 | 16/16509 | 34.02 | 1.21e-08 | 2.36e-06 | 6 | PLAU/CTSZ/PLAT/S100A10/PLAUR/ENO1 |
| Fibroblast | F3 COL11A1+ mCAF | GO:0030574 | collagen catabolic process | 8/182 | 41/16509 | 17.7 | 1.31e-08 | 2.36e-06 | 8 | MMP11/CTSB/MMP14/FAP/CTSK/MMP7/MMP2/ADAMTS2 |
| Fibroblast | F3 COL11A1+ mCAF | GO:0031589 | cell-substrate adhesion | 19/182 | 354/16509 | 4.87 | 1.38e-08 | 2.36e-06 | 19 | POSTN/COL1A1/PLAU/COL3A1/FN1/PDPN/ANTXR1/MMP14/CDH11/P4HB/ADAMTS12/THY1/LEF1/COL8A1/SPOCK… |
| Fibroblast | F3 COL11A1+ mCAF | GO:0060541 | respiratory system development | 15/182 | 225/16509 | 6.05 | 2.94e-08 | 4.73e-06 | 15 | COL3A1/STRA6/PDPN/WNT2/MMP14/HOPX/RCN3/LEF1/COL6A1/PTK7/RBP4/BASP1/CTSZ/ADAMTS2/WNT5A |
| Fibroblast | F3 COL11A1+ mCAF | GO:0002062 | chondrocyte differentiation | 11/182 | 111/16509 | 8.99 | 3.93e-08 | 5.97e-06 | 11 | COL11A1/SULF1/COL3A1/LOXL2/SERPINH1/ADAMTS12/RUNX2/TGFBI/SULF2/MDK/CHST11 |
| Fibroblast | F3 COL11A1+ mCAF | GO:0010810 | regulation of cell-substrate adhesion | 14/182 | 218/16509 | 5.83 | 1.38e-07 | 1.98e-05 | 14 | POSTN/COL1A1/PLAU/FN1/PDPN/MMP14/P4HB/THY1/LEF1/COL8A1/SPOCK1/EFEMP2/S100A10/MDK |
| Fibroblast | F3 COL11A1+ mCAF | GO:0070613 | regulation of protein processing | 8/182 | 58/16509 | 12.51 | 2.24e-07 | 3.06e-05 | 8 | PLAU/MMP14/MYH9/CTSZ/PLAT/S100A10/PLAUR/ENO1 |
| Fibroblast | F3 COL11A1+ mCAF | GO:0031639 | plasminogen activation | 6/182 | 25/16509 | 21.77 | 2.46e-07 | 3.2e-05 | 6 | PLAU/CTSZ/PLAT/S100A10/PLAUR/ENO1 |
| Fibroblast | F3 COL11A1+ mCAF | GO:0048706 | embryonic skeletal system development | 11/182 | 134/16509 | 7.45 | 2.74e-07 | 3.4e-05 | 11 | COL11A1/SULF1/COL1A1/MMP14/KIAA1217/RUNX2/SULF2/RBP4/WNT5A/CHST11/TWIST1 |
| Fibroblast | F3 COL11A1+ mCAF | GO:0030324 | lung development | 13/182 | 197/16509 | 5.99 | 2.86e-07 | 3.4e-05 | 13 | COL3A1/STRA6/PDPN/WNT2/MMP14/HOPX/RCN3/COL6A1/PTK7/RBP4/CTSZ/ADAMTS2/WNT5A |
| Fibroblast | F3 COL11A1+ mCAF | GO:0030323 | respiratory tube development | 13/182 | 201/16509 | 5.87 | 3.6e-07 | 4.11e-05 | 13 | COL3A1/STRA6/PDPN/WNT2/MMP14/HOPX/RCN3/COL6A1/PTK7/RBP4/CTSZ/ADAMTS2/WNT5A |
| Fibroblast | F3 COL11A1+ mCAF | GO:1903317 | regulation of protein maturation | 8/182 | 63/16509 | 11.52 | 4.32e-07 | 4.73e-05 | 8 | PLAU/MMP14/MYH9/CTSZ/PLAT/S100A10/PLAUR/ENO1 |
| Fibroblast | F3 COL11A1+ mCAF | GO:0002063 | chondrocyte development | 6/182 | 31/16509 | 17.56 | 9.68e-07 | 0.000102 | 6 | COL11A1/SULF1/SERPINH1/RUNX2/SULF2/CHST11 |
| Fibroblast | F3 COL11A1+ mCAF | GO:0032964 | collagen biosynthetic process | 7/182 | 49/16509 | 12.96 | 1.02e-06 | 0.000104 | 7 | INHBA/COL5A1/COL1A1/SERPINH1/F2R/RCN3/P3H4 |
| Fibroblast | F3 COL11A1+ mCAF | GO:0031638 | zymogen activation | 7/182 | 53/16509 | 11.98 | 1.77e-06 | 0.000173 | 7 | PLAU/MMP14/CTSZ/PLAT/S100A10/PLAUR/ENO1 |
| Fibroblast | F3 COL11A1+ mCAF | GO:0001503 | ossification | 17/182 | 407/16509 | 3.79 | 2.74e-06 | 0.000259 | 17 | CTHRC1/COL1A1/COL1A2/IGFBP3/MMP14/CDH11/VCAN/ANKH/ADAMTS12/LEF1/COL6A1/RUNX2/ITGA11/MMP2/… |
| Fibroblast | F3 COL11A1+ mCAF | GO:1903053 | regulation of extracellular matrix organization | 7/182 | 59/16509 | 10.76 | 3.71e-06 | 0.000338 | 7 | PDPN/ANTXR1/AEBP1/FAP/COL6A1/EFEMP2/COLGALT1 |
| Fibroblast | F3 COL11A1+ mCAF | GO:0090596 | sensory organ morphogenesis | 14/182 | 295/16509 | 4.3 | 5.09e-06 | 0.000445 | 14 | CTHRC1/COL11A1/COL5A1/COL5A2/MFAP2/STRA6/SPARC/WNT2/THY1/COL8A1/PTK7/RBP4/WNT5A/TWIST1 |
| Fibroblast | F3 COL11A1+ mCAF | GO:0022617 | extracellular matrix disassembly | 7/182 | 62/16509 | 10.24 | 5.2e-06 | 0.000445 | 7 | MMP11/PDPN/MMP14/FAP/CTSK/MMP7/MMP2 |
| Fibroblast | F3 COL11A1+ mCAF | GO:0048771 | tissue remodeling | 11/182 | 182/16509 | 5.48 | 5.64e-06 | 0.000468 | 11 | CTHRC1/MMP14/GJA1/BGN/F2R/CTSK/P3H4/COL6A1/FAM114A1/MMP2/MDK |
| Fibroblast | F3 COL11A1+ mCAF | GO:0007160 | cell-matrix adhesion | 12/182 | 238/16509 | 4.57 | 1.34e-05 | 0.00108 | 12 | POSTN/PLAU/COL3A1/FN1/MMP14/CDH11/ADAMTS12/THY1/LEF1/EFEMP2/ITGA11/S100A10 |
| Fibroblast | F3 COL11A1+ mCAF | GO:0070482 | response to oxygen levels | 14/182 | 335/16509 | 3.79 | 2.14e-05 | 0.00167 | 14 | POSTN/INHBA/COL1A1/PLAU/LOXL2/MMP14/P4HB/COL6A1/PLOD2/MMP2/HSPG2/PLAT/ENO1/TWIST1 |
| Fibroblast | F3 COL11A1+ mCAF | GO:0048592 | eye morphogenesis | 10/182 | 174/16509 | 5.21 | 2.35e-05 | 0.00178 | 10 | COL5A1/COL5A2/MFAP2/STRA6/WNT2/THY1/COL8A1/RBP4/WNT5A/TWIST1 |

| compartment | subcluster | GO_ID | Description | GeneRatio | BgRatio | FoldEnrichment | pvalue | p_adjust | Count | geneID |
|---|---|---|---|---|---|---|---|---|---|---|
| Fibroblast | F3 COL11A1+ mCAF | GO:0071230 | cellular response to amino acid stimulus | 7/182 | 79/16509 | 8.04 | 2.62e-05 | 0.00194 | 7 | COL1A1/COL3A1/COL5A2/PDGFC/COL1A2/COL6A1/MMP2 |
| Fibroblast | F3 COL11A1+ mCAF | GO:0042060 | wound healing | 16/182 | 436/16509 | 3.33 | 2.71e-05 | 0.00195 | 16 | COL5A1/PLAU/COL3A1/FN1/PDPN/F2R/FAP/MYH9/CNN2/PTK7/PLAT/S100A10/PLAUR/PDIA3/WNT5A/MYL12A |
| Fibroblast | F3 COL11A1+ mCAF | GO:0051917 | regulation of fibrinolysis | 4/182 | 17/16509 | 21.34 | 3.04e-05 | 0.00213 | 4 | PLAU/FAP/PLAT/PLAUR |
| Fibroblast | F3 COL11A1+ mCAF | GO:0040036 | regulation of fibroblast growth factor receptor signaling pathway | 5/182 | 34/16509 | 13.34 | 3.31e-05 | 0.00226 | 5 | SULF1/GPC1/RUNX2/SULF2/WNT5A |
| Fibroblast | F3 COL11A1+ mCAF | GO:0009612 | response to mechanical stimulus | 11/182 | 224/16509 | 4.45 | 3.94e-05 | 0.00263 | 11 | POSTN/COL11A1/COL1A1/STRA6/MMP14/CNN2/COL6A1/MMP2/PIEZO2/ETV1/MDK |
| Fibroblast | F3 COL11A1+ mCAF | GO:0007517 | muscle organ development | 14/182 | 355/16509 | 3.58 | 4.04e-05 | 0.00263 | 14 | COL11A1/COL3A1/STRA6/WNT2/GPC1/LEF1/COL6A1/COL6A3/EFEMP2/BASP1/ITGA11/ETV1/FLNB/TWIST1 |
| Fibroblast | F3 COL11A1+ mCAF | GO:0036293 | response to decreased oxygen levels | 13/182 | 313/16509 | 3.77 | 4.52e-05 | 0.00285 | 13 | POSTN/INHBA/PLAU/LOXL2/MMP14/P4HB/COL6A1/PLOD2/MMP2/HSPG2/PLAT/ENO1/TWIST1 |
| Fibroblast | F3 COL11A1+ mCAF | GO:0010718 | positive regulation of epithelial to mesenchymal transition | 6/182 | 59/16509 | 9.22 | 4.59e-05 | 0.00285 | 6 | COL1A1/LOXL2/PDPN/LEF1/MDK/TWIST1 |
| Fibroblast | F3 COL11A1+ mCAF | GO:0040037 | negative regulation of fibroblast growth factor receptor signaling pathway | 4/182 | 19/16509 | 19.1 | 4.87e-05 | 0.00292 | 4 | SULF1/GPC1/SULF2/WNT5A |
| Fibroblast | F3 COL11A1+ mCAF | GO:0071229 | cellular response to acid chemical | 7/182 | 87/16509 | 7.3 | 4.9e-05 | 0.00292 | 7 | COL1A1/COL3A1/COL5A2/PDGFC/COL1A2/COL6A1/MMP2 |
| Fibroblast | F3 COL11A1+ mCAF | GO:0001935 | endothelial cell proliferation | 9/182 | 153/16509 | 5.34 | 5.04e-05 | 0.00293 | 9 | SULF1/LOXL2/SPARC/MMP14/MYDGF/COL8A1/HSPG2/WNT5A/MDK |
| Fibroblast | F3 COL11A1+ mCAF | GO:0016485 | protein processing | 11/182 | 236/16509 | 4.23 | 6.32e-05 | 0.0036 | 11 | PLAU/MMP14/P4HB/MYH9/ADAM19/CTSZ/ADAMTS2/PLAT/S100A10/PLAUR/ENO1 |
| Fibroblast | F3 COL11A1+ mCAF | GO:0060537 | muscle tissue development | 15/182 | 423/16509 | 3.22 | 7.18e-05 | 0.00401 | 15 | COL11A1/COL3A1/STRA6/WNT2/GJA1/GPC1/HOPX/LEF1/COL6A1/DSG2/EFEMP2/RBP4/VGLL4/FLNB/TWIST1 |
| Fibroblast | F3 COL11A1+ mCAF | GO:0007596 | blood coagulation | 11/182 | 241/16509 | 4.14 | 7.63e-05 | 0.00417 | 11 | PLAU/COL3A1/FN1/PDPN/F2R/FAP/MYH9/PLAT/PLAUR/PDIA3/MYL12A |
| Fibroblast | F3 COL11A1+ mCAF | GO:0045765 | regulation of angiogenesis | 12/182 | 287/16509 | 3.79 | 8.36e-05 | 0.00448 | 12 | SULF1/THBS2/SPARC/ADAM12/MYDGF/PKM/DSG2/HSPG2/SPHK1/WNT5A/MDK/TWIST1 |
| Fibroblast | F3 COL11A1+ mCAF | GO:0050817 | coagulation | 11/182 | 245/16509 | 4.07 | 8.84e-05 | 0.00465 | 11 | PLAU/COL3A1/FN1/PDPN/F2R/FAP/MYH9/PLAT/PLAUR/PDIA3/MYL12A |
| Fibroblast | F3 COL11A1+ mCAF | GO:0010811 | positive regulation of cell-substrate adhesion | 8/182 | 130/16509 | 5.58 | 9.68e-05 | 0.00484 | 8 | FN1/P4HB/THY1/LEF1/COL8A1/EFEMP2/S100A10/MDK |
| Fibroblast | F3 COL11A1+ mCAF | GO:0034446 | substrate adhesion-dependent cell spreading | 7/182 | 97/16509 | 6.55 | 9.83e-05 | 0.00484 | 7 | POSTN/FN1/PDPN/ANTXR1/P4HB/S100A10/MDK |
| Fibroblast | F3 COL11A1+ mCAF | GO:0007599 | hemostasis | 11/182 | 248/16509 | 4.02 | 9.85e-05 | 0.00484 | 11 | PLAU/COL3A1/FN1/PDPN/F2R/FAP/MYH9/PLAT/PLAUR/PDIA3/MYL12A |
| Fibroblast | F3 COL11A1+ mCAF | GO:0014909 | smooth muscle cell migration | 6/182 | 68/16509 | 8.0 | 0.000103 | 0.00484 | 6 | POSTN/PLAU/IGFBP3/S100A11/PLAT/MDK |
| Fibroblast | F3 COL11A1+ mCAF | GO:0061035 | regulation of cartilage development | 6/182 | 68/16509 | 8.0 | 0.000103 | 0.00484 | 6 | LOXL2/ADAMTS12/CTSK/RUNX2/WNT5A/MDK |
| Fibroblast | F3 COL11A1+ mCAF | GO:1905330 | regulation of morphogenesis of an epithelium | 6/182 | 68/16509 | 8.0 | 0.000103 | 0.00484 | 6 | SULF1/WNT2/GJA1/ADAMTS12/WNT5A/MDK |
| Fibroblast | F3 COL11A1+ mCAF | GO:1901342 | regulation of vasculature development | 12/182 | 294/16509 | 3.7 | 0.000105 | 0.00487 | 12 | SULF1/THBS2/SPARC/ADAM12/MYDGF/PKM/DSG2/HSPG2/SPHK1/WNT5A/MDK/TWIST1 |
| Fibroblast | F3 COL11A1+ mCAF | GO:0085029 | extracellular matrix assembly | 5/182 | 44/16509 | 10.31 | 0.000118 | 0.00533 | 5 | COL3A1/COL1A2/ANTXR1/COL6A1/EFEMP2 |
| Fibroblast | F3 COL11A1+ mCAF | GO:0001837 | epithelial to mesenchymal transition | 9/182 | 171/16509 | 4.77 | 0.000119 | 0.00533 | 9 | COL1A1/LOXL2/PDPN/WNT2/LEF1/DSG2/WNT5A/MDK/TWIST1 |

| compartment | subcluster | GO_ID | Description | GeneRatio | BgRatio | FoldEnrichment | pvalue | p_adjust | Count | geneID |
|---|---|---|---|---|---|---|---|---|---|---|
| Fibroblast | F3 COL11A1+ mCAF | GO:0001666 | response to hypoxia | 12/182 | 300/16509 | 3.63 | 0.000127 | 0.00555 | 12 | POSTN/INHBA/PLAU/LOXL2/MMP14/P4HB/PLOD2/MMP2/HSPG2/PLAT/ENO1/TWIST1 |
| Fibroblast | F3 COL11A1+ mCAF | GO:0010954 | positive regulation of protein processing | 4/182 | 24/16509 | 15.12 | 0.000128 | 0.00555 | 4 | MMP14/MYH9/S100A10/ENO1 |
| Fibroblast | F3 COL11A1+ mCAF | GO:0045785 | positive regulation of cell adhesion | 16/182 | 498/16509 | 2.91 | 0.00013 | 0.00557 | 16 | LGALS1/FN1/PDPN/P4HB/THY1/CD55/LEF1/COL8A1/ADAM19/DSG2/EFEMP2/S100A10/PLAUR/WNT5A/MDK/LGA… |
| Fibroblast | F3 COL11A1+ mCAF | GO:0048562 | embryonic organ morphogenesis | 12/182 | 302/16509 | 3.6 | 0.000135 | 0.00569 | 12 | CTHRC1/COL11A1/MFAP2/STRA6/SPARC/MMP14/RUNX2/PTK7/RBP4/WNT5A/CHST11/TWIST1 |
| Fibroblast | F3 COL11A1+ mCAF | GO:1903319 | positive regulation of protein maturation | 4/182 | 25/16509 | 14.51 | 0.000151 | 0.00625 | 4 | MMP14/MYH9/S100A10/ENO1 |
| Fibroblast | F3 COL11A1+ mCAF | GO:0031099 | regeneration | 9/182 | 178/16509 | 4.59 | 0.000161 | 0.0065 | 9 | POSTN/HOPX/THY1/NREP/COL6A1/SULF2/MMP2/HSPG2/MDK |
| Fibroblast | F3 COL11A1+ mCAF | GO:0044403 | biological process involved in symbiotic interaction | 10/182 | 219/16509 | 4.14 | 0.000162 | 0.0065 | 10 | LGALS1/FN1/CTSB/LRRC15/P4HB/CD55/MYH9/GAPDH/BST2/LGALS9 |
| Fibroblast | F3 COL11A1+ mCAF | GO:0071675 | regulation of mononuclear cell migration | 8/182 | 141/16509 | 5.15 | 0.00017 | 0.0067 | 8 | RARRES2/NBL1/MMP14/P4HB/CNN2/WNT5A/MDK/LGALS9 |
| Fibroblast | F3 COL11A1+ mCAF | GO:0090288 | negative regulation of cellular response to growth factor stimulus | 7/182 | 106/16509 | 5.99 | 0.000172 | 0.0067 | 7 | SULF1/NBL1/HTRA3/ADAMTS12/GPC1/SULF2/WNT5A |
| Fibroblast | F3 COL11A1+ mCAF | GO:2000242 | negative regulation of reproductive process | 6/182 | 75/16509 | 7.26 | 0.000177 | 0.00672 | 6 | SULF1/GJA1/MYH9/TPST2/PLAT/WNT5A |
| Fibroblast | F3 COL11A1+ mCAF | GO:0002685 | regulation of leukocyte migration | 10/182 | 222/16509 | 4.09 | 0.00018 | 0.00672 | 10 | RARRES2/NBL1/MMP14/P4HB/THY1/CNN2/PERP/WNT5A/MDK/LGALS9 |
| Fibroblast | F3 COL11A1+ mCAF | GO:0010717 | regulation of epithelial to mesenchymal transition | 7/182 | 107/16509 | 5.93 | 0.000182 | 0.00672 | 7 | COL1A1/LOXL2/PDPN/LEF1/DSG2/MDK/TWIST1 |
| Fibroblast | F3 COL11A1+ mCAF | GO:0032526 | response to retinoic acid | 7/182 | 107/16509 | 5.93 | 0.000182 | 0.00672 | 7 | COL1A1/WNT2/PTK7/RBP4/MMP2/MEST/WNT5A |
| Fibroblast | F3 COL11A1+ mCAF | GO:0048568 | embryonic organ development | 15/182 | 463/16509 | 2.94 | 0.000194 | 0.00708 | 15 | CTHRC1/COL11A1/MFAP2/STRA6/SPARC/WNT2/RARRES2/MMP14/LEF1/RUNX2/PTK7/RBP4/WNT5A/CHST11/TWI… |
| Fibroblast | F3 COL11A1+ mCAF | GO:0060900 | embryonic camera-type eye formation | 3/182 | 11/16509 | 24.74 | 0.000204 | 0.00721 | 3 | STRA6/WNT5A/TWIST1 |
| Fibroblast | F3 COL11A1+ mCAF | GO:0044344 | cellular response to fibroblast growth factor stimulus | 7/182 | 109/16509 | 5.83 | 0.000204 | 0.00721 | 7 | POSTN/SULF1/COL1A1/GPC1/RUNX2/SULF2/WNT5A |
| Fibroblast | F3 COL11A1+ mCAF | GO:0042730 | fibrinolysis | 4/182 | 27/16509 | 13.44 | 0.000206 | 0.00721 | 4 | PLAU/FAP/PLAT/PLAUR |
| Fibroblast | F3 COL11A1+ mCAF | GO:0060485 | mesenchyme development | 12/182 | 319/16509 | 3.41 | 0.000224 | 0.00775 | 12 | COL1A1/LOXL2/FN1/PDPN/WNT2/LEF1/DSG2/PTK7/BASP1/WNT5A/MDK/TWIST1 |
| Fibroblast | F3 COL11A1+ mCAF | GO:0048144 | fibroblast proliferation | 7/182 | 111/16509 | 5.72 | 0.000228 | 0.00775 | 7 | COL3A1/FN1/PDGFC/WNT2/FAM114A1/SPHK1/WNT5A |
| Fibroblast | F3 COL11A1+ mCAF | GO:0014823 | response to activity | 6/182 | 79/16509 | 6.89 | 0.000236 | 0.00775 | 6 | POSTN/FN1/RARRES2/COL6A1/RBP4/MMP2 |
| Fibroblast | F3 COL11A1+ mCAF | GO:0002687 | positive regulation of leukocyte migration | 8/182 | 148/16509 | 4.9 | 0.000237 | 0.00775 | 8 | RARRES2/MMP14/P4HB/THY1/PERP/WNT5A/MDK/LGALS9 |
| Fibroblast | F3 COL11A1+ mCAF | GO:0014850 | response to muscle activity | 4/182 | 28/16509 | 12.96 | 0.000238 | 0.00775 | 4 | POSTN/FN1/COL6A1/RBP4 |
| Fibroblast | F3 COL11A1+ mCAF | GO:0060065 | uterus development | 4/182 | 28/16509 | 12.96 | 0.000238 | 0.00775 | 4 | STRA6/COL6A1/RBP4/WNT5A |
| Fibroblast | F3 COL11A1+ mCAF | GO:0006029 | proteoglycan metabolic process | 7/182 | 112/16509 | 5.67 | 0.000241 | 0.00777 | 7 | COL11A1/SULF1/ADAMTS12/GPC1/CHPF/SULF2/CHST11 |
| Fibroblast | F3 COL11A1+ mCAF | GO:0043200 | response to amino acid | 7/182 | 114/16509 | 5.57 | 0.000269 | 0.00847 | 7 | COL1A1/COL3A1/COL5A2/PDGFC/COL1A2/COL6A1/MMP2 |
| Fibroblast | F3 COL11A1+ mCAF | GO:0007343 | egg activation | 3/182 | 12/16509 | 22.68 | 0.000269 | 0.00847 | 3 | MYH9/TPST2/PLAT |
| Fibroblast | F3 COL11A1+ mCAF | GO:0014812 | muscle cell migration | 6/182 | 82/16509 | 6.64 | 0.000289 | 0.00899 | 6 | POSTN/PLAU/IGFBP3/S100A11/PLAT/MDK |
| Fibroblast | F3 COL11A1+ mCAF | GO:0060324 | face development | 5/182 | 54/16509 | 8.4 | 0.000315 | 0.00957 | 5 | COL1A1/STRA6/LEF1/MMP2/WNT5A |
| Fibroblast | F3 COL11A1+ mCAF | GO:0071774 | response to fibroblast growth factor | 7/182 | 117/16509 | 5.43 | 0.000315 | 0.00957 | 7 | POSTN/SULF1/COL1A1/GPC1/RUNX2/SULF2/WNT5A |

| compartment | subcluster | GO_ID | Description | GeneRatio | BgRatio | FoldEnrichment | pvalue | p_adjust | Count | geneID |
|---|---|---|---|---|---|---|---|---|---|---|
| Fibroblast | F3 COL11A1+ mCAF | GO:0050900 | leukocyte migration | 13/182 | 381/16509 | 3.1 | 0.000318 | 0.00957 | 13 | RARRES2/NBL1/MMP14/P4HB/THY1/MYH9/CNN2/PPIB/PERP/MMP2/WNT5A/MDK/LGALS9 |
| Fibroblast | F3 COL11A1+ mCAF | GO:0050673 | epithelial cell proliferation | 14/182 | 438/16509 | 2.9 | 0.000364 | 0.0108 | 14 | SULF1/LOXL2/IGFBP3/SPARC/WNT2/MMP14/FAP/MYDGF/COL8A1/RUNX2/SULF2/HSPG2/WNT5A/MDK |
| Fibroblast | F3 COL11A1+ mCAF | GO:0048146 | positive regulation of fibroblast proliferation | 5/182 | 56/16509 | 8.1 | 0.000373 | 0.011 | 5 | FN1/PDGFC/WNT2/SPHK1/WNT5A |
| Fibroblast | F3 COL11A1+ mCAF | GO:0061036 | positive regulation of cartilage development | 4/182 | 32/16509 | 11.34 | 0.000404 | 0.0118 | 4 | LOXL2/RUNX2/WNT5A/MDK |
| Fibroblast | F3 COL11A1+ mCAF | GO:0061029 | eyelid development in camera-type eye | 3/182 | 14/16509 | 19.44 | 0.000439 | 0.0126 | 3 | INHBA/STRA6/TWIST1 |
| Fibroblast | F3 COL11A1+ mCAF | GO:0060325 | face morphogenesis | 4/182 | 33/16509 | 11.0 | 0.000455 | 0.013 | 4 | COL1A1/STRA6/LEF1/MMP2 |
| Fibroblast | F3 COL11A1+ mCAF | GO:0001952 | regulation of cell-matrix adhesion | 7/182 | 125/16509 | 5.08 | 0.000471 | 0.0132 | 7 | POSTN/PLAU/MMP14/THY1/LEF1/EFEMP2/S100A10 |
| Fibroblast | F3 COL11A1+ mCAF | GO:0060350 | endochondral bone morphogenesis | 5/182 | 59/16509 | 7.69 | 0.000477 | 0.0132 | 5 | COL1A1/COL3A1/SERPINH1/MMP14/RUNX2 |
| Fibroblast | F3 COL11A1+ mCAF | GO:1900024 | regulation of substrate adhesion-dependent cell spreading | 5/182 | 59/16509 | 7.69 | 0.000477 | 0.0132 | 5 | POSTN/PDPN/P4HB/S100A10/MDK |
| Fibroblast | F3 COL11A1+ mCAF | GO:0141091 | transforming growth factor beta receptor superfamily signaling pathway | 12/182 | 348/16509 | 3.13 | 0.000492 | 0.0133 | 12 | INHBA/SULF1/COL3A1/COL1A2/NBL1/HTRA3/NREP/LEF1/RUNX2/FSTL1/WNT5A/CHST11 |
| Fibroblast | F3 COL11A1+ mCAF | GO:0045766 | positive regulation of angiogenesis | 8/182 | 165/16509 | 4.4 | 0.000493 | 0.0133 | 8 | ADAM12/MYDGF/PKM/DSG2/SPHK1/WNT5A/MDK/TWIST1 |
| Fibroblast | F3 COL11A1+ mCAF | GO:0048762 | mesenchymal cell differentiation | 10/182 | 252/16509 | 3.6 | 0.000495 | 0.0133 | 10 | COL1A1/LOXL2/FN1/PDPN/WNT2/LEF1/DSG2/WNT5A/MDK/TWIST1 |
| Fibroblast | F3 COL11A1+ mCAF | GO:0016055 | Wnt signaling pathway | 14/182 | 453/16509 | 2.8 | 0.000509 | 0.0135 | 14 | CTHRC1/SULF1/COL1A1/SDC1/WNT2/LEF1/COL6A1/UBAC2/SULF2/PTK7/WNT5A/VGLL4/NXN/MDK |
| Fibroblast | F3 COL11A1+ mCAF | GO:1905517 | macrophage migration | 5/182 | 60/16509 | 7.56 | 0.000515 | 0.0135 | 5 | RARRES2/MMP14/CNN2/MMP2/MDK |
| Fibroblast | F3 COL11A1+ mCAF | GO:0001101 | response to acid chemical | 7/182 | 127/16509 | 5.0 | 0.000518 | 0.0135 | 7 | COL1A1/COL3A1/COL5A2/PDGFC/COL1A2/COL6A1/MMP2 |
| Fibroblast | F3 COL11A1+ mCAF | GO:0046598 | positive regulation of viral entry into host cell | 3/182 | 15/16509 | 18.14 | 0.000544 | 0.0138 | 3 | LGALS1/P4HB/LGALS9 |
| Fibroblast | F3 COL11A1+ mCAF | GO:0060572 | morphogenesis of an epithelial bud | 3/182 | 15/16509 | 18.14 | 0.000544 | 0.0138 | 3 | SULF1/WNT2/WNT5A |
| Fibroblast | F3 COL11A1+ mCAF | GO:0075294 | positive regulation by symbiont of entry into host | 3/182 | 15/16509 | 18.14 | 0.000544 | 0.0138 | 3 | LGALS1/P4HB/LGALS9 |
| Fibroblast | F3 COL11A1+ mCAF | GO:0071772 | response to BMP | 8/182 | 168/16509 | 4.32 | 0.000556 | 0.0138 | 8 | SULF1/NBL1/HTRA3/ADAMTS12/LEF1/RUNX2/FSTL1/WNT5A |
| Fibroblast | F3 COL11A1+ mCAF | GO:0071773 | cellular response to BMP stimulus | 8/182 | 168/16509 | 4.32 | 0.000556 | 0.0138 | 8 | SULF1/NBL1/HTRA3/ADAMTS12/LEF1/RUNX2/FSTL1/WNT5A |
| Fibroblast | F3 COL11A1+ mCAF | GO:0042476 | odontogenesis | 7/182 | 129/16509 | 4.92 | 0.000569 | 0.014 | 7 | INHBA/COL1A1/COL1A2/ANKH/LEF1/RUNX2/PERP |
| Fibroblast | F3 COL11A1+ mCAF | GO:1904018 | positive regulation of vasculature development | 8/182 | 169/16509 | 4.29 | 0.000578 | 0.0141 | 8 | ADAM12/MYDGF/PKM/DSG2/SPHK1/WNT5A/MDK/TWIST1 |
| Fibroblast | F3 COL11A1+ mCAF | GO:0014910 | regulation of smooth muscle cell migration | 5/182 | 62/16509 | 7.32 | 0.0006 | 0.0145 | 5 | POSTN/PLAU/IGFBP3/S100A11/MDK |
| Fibroblast | F3 COL11A1+ mCAF | GO:0060070 | canonical Wnt signaling pathway | 11/182 | 307/16509 | 3.25 | 0.000615 | 0.0148 | 11 | CTHRC1/COL1A1/SDC1/WNT2/LEF1/COL6A1/UBAC2/SULF2/PTK7/WNT5A/MDK |
| Fibroblast | F3 COL11A1+ mCAF | GO:1903035 | negative regulation of response to wounding | 6/182 | 95/16509 | 5.73 | 0.000638 | 0.0151 | 6 | PLAU/FAP/DSG2/PLAT/PLAUR/MDK |
| Fibroblast | F3 COL11A1+ mCAF | GO:0002474 | antigen processing and presentation of peptide antigen via MHC class I | 4/182 | 36/16509 | 10.08 | 0.000639 | 0.0151 | 4 | HLA-B/TAPBP/HLA-C/PDIA3 |
| Fibroblast | F3 COL11A1+ mCAF | GO:0018126 | protein hydroxylation | 3/182 | 16/16509 | 17.01 | 0.000664 | 0.0154 | 3 | P4HB/P3H4/PLOD2 |
| Fibroblast | F3 COL11A1+ mCAF | GO:0070234 | positive regulation of T cell apoptotic process | 3/182 | 16/16509 | 17.01 | 0.000664 | 0.0154 | 3 | PERP/WNT5A/LGALS9 |
| Fibroblast | F3 COL11A1+ mCAF | GO:0071677 | positive regulation of mononuclear cell migration | 6/182 | 96/16509 | 5.67 | 0.000674 | 0.0155 | 6 | RARRES2/MMP14/P4HB/WNT5A/MDK/LGALS9 |

| compartment | subcluster | GO_ID | Description | GeneRatio | BgRatio | FoldEnrichment | pvalue | p_adjust | Count | geneID |
|---|---|---|---|---|---|---|---|---|---|---|
| Fibroblast | F3 COL11A1+ mCAF | GO:0048048 | embryonic eye morphogenesis | 4/182 | 37/16509 | 9.81 | 0.00071 | 0.0162 | 4 | MFAP2/STRA6/WNT5A/TWIST1 |
| Fibroblast | F3 COL11A1+ mCAF | GO:0090287 | regulation of cellular response to growth factor stimulus | 11/182 | 314/16509 | 3.18 | 0.000741 | 0.0167 | 11 | SULF1/NBL1/HTRA3/ADAMTS12/GPC1/NREP/RUNX2/SULF2/FSTL1/WNT5A/CHST11 |
| Fibroblast | F3 COL11A1+ mCAF | GO:0060349 | bone morphogenesis | 6/182 | 98/16509 | 5.55 | 0.000752 | 0.0169 | 6 | COL1A1/COL3A1/SERPINH1/MMP14/RUNX2/TWIST1 |
| Fibroblast | F3 COL11A1+ mCAF | GO:0043903 | regulation of biological process involved in symbiotic interaction | 4/182 | 38/16509 | 9.55 | 0.000787 | 0.0174 | 4 | LGALS1/P4HB/BST2/LGALS9 |
| Fibroblast | F3 COL11A1+ mCAF | GO:0071604 | transforming growth factor beta production | 4/182 | 38/16509 | 9.55 | 0.000787 | 0.0174 | 4 | COL3A1/FN1/LUM/LGALS9 |
| Fibroblast | F3 COL11A1+ mCAF | GO:0035860 | glial cell-derived neurotrophic factor receptor signaling pathway | 3/182 | 17/16509 | 16.01 | 0.0008 | 0.0174 | 3 | SULF1/MMP14/SULF2 |
| Fibroblast | F3 COL11A1+ mCAF | GO:0080154 | regulation of fertilization | 3/182 | 17/16509 | 16.01 | 0.0008 | 0.0174 | 3 | MYH9/TPST2/PLAT |
| Fibroblast | F3 COL11A1+ mCAF | GO:0071674 | mononuclear cell migration | 9/182 | 223/16509 | 3.66 | 0.000837 | 0.018 | 9 | RARRES2/NBL1/MMP14/P4HB/CNN2/MMP2/WNT5A/MDK/LGALS9 |
| Fibroblast | F3 COL11A1+ mCAF | GO:0071559 | response to transforming growth factor beta | 10/182 | 270/16509 | 3.36 | 0.000843 | 0.018 | 10 | POSTN/COL1A1/COL3A1/MXRA5/COL1A2/WNT2/HTRA3/NREP/WNT5A/CHST11 |
| Fibroblast | F3 COL11A1+ mCAF | GO:0001649 | osteoblast differentiation | 9/182 | 224/16509 | 3.64 | 0.000864 | 0.0183 | 9 | CTHRC1/COL1A1/IGFBP3/VCAN/LEF1/COL6A1/RUNX2/ITGA11/TWIST1 |
| Fibroblast | F3 COL11A1+ mCAF | GO:0048705 | skeletal system morphogenesis | 9/182 | 225/16509 | 3.63 | 0.000891 | 0.0188 | 9 | COL11A1/COL1A1/COL3A1/SERPINH1/MMP14/RUNX2/MMP2/CHST11/TWIST1 |
| Fibroblast | F3 COL11A1+ mCAF | GO:0048593 | camera-type eye morphogenesis | 7/182 | 140/16509 | 4.54 | 0.000925 | 0.0192 | 7 | STRA6/WNT2/THY1/COL8A1/RBP4/WNT5A/TWIST1 |
| Fibroblast | F3 COL11A1+ mCAF | GO:0007178 | cell surface receptor protein serine/threonine kinase signaling pathway | 12/182 | 374/16509 | 2.91 | 0.000926 | 0.0192 | 12 | INHBA/SULF1/COL3A1/COL1A2/NBL1/HTRA3/NREP/LEF1/RUNX2/FSTL1/WNT5A/CHST11 |
| Fibroblast | F3 COL11A1+ mCAF | GO:0016264 | gap junction assembly | 3/182 | 18/16509 | 15.12 | 0.000952 | 0.0194 | 3 | GJB2/GJA1/HOPX |
| Fibroblast | F3 COL11A1+ mCAF | GO:0040013 | negative regulation of locomotion | 11/182 | 324/16509 | 3.08 | 0.000957 | 0.0194 | 11 | SULF1/COL3A1/IGFBP3/NBL1/CDH11/GJA1/DPYSL3/THY1/CNN2/WNT5A/BST2 |
| Fibroblast | F3 COL11A1+ mCAF | GO:0060323 | head morphogenesis | 4/182 | 40/16509 | 9.07 | 0.000958 | 0.0194 | 4 | COL1A1/STRA6/LEF1/MMP2 |
| Fibroblast | F3 COL11A1+ mCAF | GO:0034109 | homotypic cell-cell adhesion | 6/182 | 103/16509 | 5.28 | 0.000976 | 0.0196 | 6 | PDPN/MYH9/DSG2/PLAUR/PDIA3/MYL12A |
| Fibroblast | F3 COL11A1+ mCAF | GO:0030111 | regulation of Wnt signaling pathway | 11/182 | 325/16509 | 3.07 | 0.000981 | 0.0196 | 11 | CTHRC1/SULF1/COL1A1/LEF1/UBAC2/SULF2/PTK7/WNT5A/VGLL4/NXN/MDK |
| Fibroblast | F3 COL11A1+ mCAF | GO:0030193 | regulation of blood coagulation | 5/182 | 70/16509 | 6.48 | 0.00105 | 0.0207 | 5 | PLAU/F2R/FAP/PLAT/PLAUR |
| Fibroblast | F3 COL11A1+ mCAF | GO:0030162 | regulation of proteolysis | 12/182 | 380/16509 | 2.86 | 0.00106 | 0.0209 | 12 | PLAU/MMP14/RCN3/MYH9/GAPDH/PERP/CTSZ/PLAT/TIMP2/S100A10/PLAUR/ENO1 |
| Fibroblast | F3 COL11A1+ mCAF | GO:0051851 | host-mediated perturbation of symbiont process | 6/182 | 105/16509 | 5.18 | 0.00108 | 0.0211 | 6 | LRRC15/LEF1/LY6E/PPIB/ZBED1/IFI27 |
| Fibroblast | F3 COL11A1+ mCAF | GO:0030540 | female genitalia development | 3/182 | 19/16509 | 14.32 | 0.00112 | 0.0218 | 3 | STRA6/RBP4/WNT5A |
| Fibroblast | F3 COL11A1+ mCAF | GO:1900046 | regulation of hemostasis | 5/182 | 72/16509 | 6.3 | 0.00119 | 0.0229 | 5 | PLAU/F2R/FAP/PLAT/PLAUR |
| Fibroblast | F3 COL11A1+ mCAF | GO:0019058 | viral life cycle | 10/182 | 284/16509 | 3.19 | 0.00124 | 0.0236 | 10 | LGALS1/CTSB/IL32/LRRC15/P4HB/CD55/MYH9/ZBED1/BST2/LGALS9 |
| Fibroblast | F3 COL11A1+ mCAF | GO:0048701 | embryonic cranial skeleton morphogenesis | 4/182 | 43/16509 | 8.44 | 0.00126 | 0.024 | 4 | MMP14/RUNX2/CHST11/TWIST1 |
| Fibroblast | F3 COL11A1+ mCAF | GO:0046596 | regulation of viral entry into host cell | 3/182 | 20/16509 | 13.61 | 0.00131 | 0.0245 | 3 | LGALS1/P4HB/LGALS9 |
| Fibroblast | F3 COL11A1+ mCAF | GO:0070230 | positive regulation of lymphocyte apoptotic process | 3/182 | 20/16509 | 13.61 | 0.00131 | 0.0245 | 3 | PERP/WNT5A/LGALS9 |
| Fibroblast | F3 COL11A1+ mCAF | GO:0007229 | integrin-mediated signaling pathway | 6/182 | 110/16509 | 4.95 | 0.00137 | 0.0254 | 6 | PLAU/COL3A1/FN1/THY1/MYH9/ITGA11 |
| Fibroblast | F3 COL11A1+ mCAF | GO:0035107 | appendage morphogenesis | 7/182 | 150/16509 | 4.23 | 0.00138 | 0.0254 | 7 | COL3A1/LEF1/COL6A1/RUNX2/WNT5A/CHST11/TWIST1 |
| Fibroblast | F3 COL11A1+ mCAF | GO:0035108 | limb morphogenesis | 7/182 | 150/16509 | 4.23 | 0.00138 | 0.0254 | 7 | COL3A1/LEF1/COL6A1/RUNX2/WNT5A/CHST11/TWIST1 |
| Fibroblast | F3 COL11A1+ mCAF | GO:0060562 | epithelial tube morphogenesis | 11/182 | 340/16509 | 2.93 | 0.00141 | 0.0257 | 11 | CTHRC1/WNT2/MMP14/ADAMTS12/LEF1/PTK7/PERP/CTSZ/WNT5A/MDK/TWIST1 |

| compartment | subcluster | GO_ID | Description | GeneRatio | BgRatio | FoldEnrichment | pvalue | p_adjust | Count | geneID |
|---|---|---|---|---|---|---|---|---|---|---|
| Fibroblast | F3 COL11A1+ mCAF | GO:0050818 | regulation of coagulation | 5/182 | 75/16509 | 6.05 | 0.00143 | 0.0258 | 5 | PLAU/F2R/FAP/PLAT/PLAUR |
| Fibroblast | F3 COL11A1+ mCAF | GO:0030336 | negative regulation of cell migration | 10/182 | 291/16509 | 3.12 | 0.00148 | 0.0267 | 10 | SULF1/COL3A1/IGFBP3/NBL1/CDH11/GJA1/DPYSL3/THY1/CNN2/BST2 |
| Fibroblast | F3 COL11A1+ mCAF | GO:0010955 | negative regulation of protein processing | 3/182 | 21/16509 | 12.96 | 0.00151 | 0.0269 | 3 | PLAU/CTSZ/PLAT |
| Fibroblast | F3 COL11A1+ mCAF | GO:0019885 | antigen processing and presentation of endogenous peptide antigen via MHC class I | 3/182 | 21/16509 | 12.96 | 0.00151 | 0.0269 | 3 | HLA-B/TAPBP/HLA-C |
| Fibroblast | F3 COL11A1+ mCAF | GO:0006935 | chemotaxis | 13/182 | 452/16509 | 2.61 | 0.00154 | 0.0272 | 13 | PLAU/CMTM3/RARRES2/NBL1/LEF1/TYMP/PPIB/PERP/MMP2/PLAUR/WNT5A/MDK/LGALS9 |
| Fibroblast | F3 COL11A1+ mCAF | GO:0042330 | taxis | 13/182 | 454/16509 | 2.6 | 0.0016 | 0.0281 | 13 | PLAU/CMTM3/RARRES2/NBL1/LEF1/TYMP/PPIB/PERP/MMP2/PLAUR/WNT5A/MDK/LGALS9 |
| Fibroblast | F3 COL11A1+ mCAF | GO:1905521 | regulation of macrophage migration | 4/182 | 46/16509 | 7.89 | 0.00162 | 0.0283 | 4 | RARRES2/MMP14/CNN2/MDK |
| Fibroblast | F3 COL11A1+ mCAF | GO:0014706 | striated muscle tissue development | 12/182 | 402/16509 | 2.71 | 0.00171 | 0.0296 | 12 | COL11A1/WNT2/GJA1/GPC1/HOPX/LEF1/COL6A1/DSG2/RBP4/VGLL4/FLNB/TWIST1 |
| Fibroblast | F3 COL11A1+ mCAF | GO:0007162 | negative regulation of cell adhesion | 10/182 | 297/16509 | 3.05 | 0.00172 | 0.0296 | 10 | POSTN/LGALS1/COL1A1/MMP14/SPOCK1/TGFBI/MMP2/HSPG2/MDK/LGALS9 |
| Fibroblast | F3 COL11A1+ mCAF | GO:0030509 | BMP signaling pathway | 7/182 | 156/16509 | 4.07 | 0.00173 | 0.0296 | 7 | SULF1/NBL1/HTRA3/LEF1/RUNX2/FSTL1/WNT5A |
| Fibroblast | F3 COL11A1+ mCAF | GO:0006590 | thyroid hormone generation | 3/182 | 22/16509 | 12.37 | 0.00174 | 0.0296 | 3 | CTSB/CTSK/DIO2 |
| Fibroblast | F3 COL11A1+ mCAF | GO:0035821 | modulation of process of another organism | 6/182 | 116/16509 | 4.69 | 0.0018 | 0.0304 | 6 | LRRC15/LEF1/LY6E/PPIB/ZBED1/IFI27 |
| Fibroblast | F3 COL11A1+ mCAF | GO:0060428 | lung epithelium development | 4/182 | 48/16509 | 7.56 | 0.0019 | 0.032 | 4 | STRA6/WNT2/RCN3/COL6A1 |
| Fibroblast | F3 COL11A1+ mCAF | GO:0002483 | antigen processing and presentation of endogenous peptide antigen | 3/182 | 23/16509 | 11.83 | 0.00198 | 0.0329 | 3 | HLA-B/TAPBP/HLA-C |
| Fibroblast | F3 COL11A1+ mCAF | GO:1903318 | negative regulation of protein maturation | 3/182 | 23/16509 | 11.83 | 0.00198 | 0.0329 | 3 | PLAU/CTSZ/PLAT |
| Fibroblast | F3 COL11A1+ mCAF | GO:0006509 | membrane protein ectodomain proteolysis | 4/182 | 49/16509 | 7.4 | 0.00206 | 0.0337 | 4 | MYH9/MMP7/ADAM19/TIMP2 |
| Fibroblast | F3 COL11A1+ mCAF | GO:0032330 | regulation of chondrocyte differentiation | 4/182 | 49/16509 | 7.4 | 0.00206 | 0.0337 | 4 | LOXL2/ADAMTS12/RUNX2/MDK |
| Fibroblast | F3 COL11A1+ mCAF | GO:0002688 | regulation of leukocyte chemotaxis | 6/182 | 120/16509 | 4.54 | 0.00214 | 0.0347 | 6 | RARRES2/NBL1/PERP/WNT5A/MDK/LGALS9 |
| Fibroblast | F3 COL11A1+ mCAF | GO:2000146 | negative regulation of cell motility | 10/182 | 306/16509 | 2.96 | 0.00214 | 0.0347 | 10 | SULF1/COL3A1/IGFBP3/NBL1/CDH11/GJA1/DPYSL3/THY1/CNN2/BST2 |
| Fibroblast | F3 COL11A1+ mCAF | GO:0008543 | fibroblast growth factor receptor signaling pathway | 5/182 | 83/16509 | 5.46 | 0.00224 | 0.036 | 5 | SULF1/GPC1/RUNX2/SULF2/WNT5A |
| Fibroblast | F3 COL11A1+ mCAF | GO:0010171 | body morphogenesis | 4/182 | 51/16509 | 7.11 | 0.00238 | 0.0381 | 4 | COL1A1/STRA6/LEF1/MMP2 |
| Fibroblast | F3 COL11A1+ mCAF | GO:1903034 | regulation of response to wounding | 7/182 | 166/16509 | 3.83 | 0.00246 | 0.039 | 7 | PLAU/F2R/FAP/DSG2/PLAT/PLAUR/MDK |
| Fibroblast | F3 COL11A1+ mCAF | GO:0097193 | intrinsic apoptotic signaling pathway | 10/182 | 312/16509 | 2.91 | 0.00247 | 0.039 | 10 | P4HB/PYCR1/PERP/PTTG1IP/RPS27L/MMP2/SHISA5/PLAUR/ENO1/IFI27 |
| Fibroblast | F3 COL11A1+ mCAF | GO:0060571 | morphogenesis of an epithelial fold | 3/182 | 25/16509 | 10.89 | 0.00254 | 0.0394 | 3 | SULF1/WNT2/WNT5A |
| Fibroblast | F3 COL11A1+ mCAF | GO:1903055 | positive regulation of extracellular matrix organization | 3/182 | 25/16509 | 10.89 | 0.00254 | 0.0394 | 3 | PDPN/EFEMP2/COLGALT1 |
| Fibroblast | F3 COL11A1+ mCAF | GO:1903902 | positive regulation of viral life cycle | 3/182 | 25/16509 | 10.89 | 0.00254 | 0.0394 | 3 | LGALS1/P4HB/LGALS9 |
| Fibroblast | F3 COL11A1+ mCAF | GO:0030195 | negative regulation of blood coagulation | 4/182 | 52/16509 | 6.98 | 0.00256 | 0.0396 | 4 | PLAU/FAP/PLAT/PLAUR |
| Fibroblast | F3 COL11A1+ mCAF | GO:0071560 | cellular response to transforming growth factor beta stimulus | 9/182 | 263/16509 | 3.1 | 0.00261 | 0.0401 | 9 | POSTN/COL1A1/COL3A1/COL1A2/WNT2/HTRA3/NREP/WNT5A/CHST11 |
| Fibroblast | F3 COL11A1+ mCAF | GO:0001654 | eye development | 12/182 | 425/16509 | 2.56 | 0.0027 | 0.0412 | 12 | INHBA/COL5A1/COL5A2/MFAP2/STRA6/WNT2/THY1/COL8A1/RCN1/RBP4/WNT5A/TWIST1 |

| compartment | subcluster | GO_ID | Description | GeneRatio | BgRatio | FoldEnrichment | pvalue | p_adjust | Count | geneID |
|---|---|---|---|---|---|---|---|---|---|---|
| Fibroblast | F3 COL11A1+ mCAF | GO:0046660 | female sex differentiation | 6/182 | 126/16509 | 4.32 | 0.00273 | 0.0412 | 6 | INHBA/STRA6/MMP14/RBP4/MMP2/WNT5A |
| Fibroblast | F3 COL11A1+ mCAF | GO:0046849 | bone remodeling | 5/182 | 87/16509 | 5.21 | 0.00275 | 0.0412 | 5 | CTHRC1/GJA1/CTSK/P3H4/MDK |
| Fibroblast | F3 COL11A1+ mCAF | GO:1900047 | negative regulation of hemostasis | 4/182 | 53/16509 | 6.85 | 0.00275 | 0.0412 | 4 | PLAU/FAP/PLAT/PLAUR |
| Fibroblast | F3 COL11A1+ mCAF | GO:0007009 | plasma membrane organization | 7/182 | 170/16509 | 3.74 | 0.00281 | 0.0412 | 7 | COL5A1/MYH9/DMKN/TPST2/COL6A1/SERINC2/S100A10 |
| Fibroblast | F3 COL11A1+ mCAF | GO:0030194 | positive regulation of blood coagulation | 3/182 | 26/16509 | 10.47 | 0.00284 | 0.0412 | 3 | PLAU/F2R/PLAT |
| Fibroblast | F3 COL11A1+ mCAF | GO:0048745 | smooth muscle tissue development | 3/182 | 26/16509 | 10.47 | 0.00284 | 0.0412 | 3 | COL3A1/STRA6/EFEMP2 |
| Fibroblast | F3 COL11A1+ mCAF | GO:0052547 | regulation of peptidase activity | 3/182 | 26/16509 | 10.47 | 0.00284 | 0.0412 | 3 | RCN3/GAPDH/TIMP2 |
| Fibroblast | F3 COL11A1+ mCAF | GO:0060343 | trabecula formation | 3/182 | 26/16509 | 10.47 | 0.00284 | 0.0412 | 3 | COL1A1/RBP4/MMP2 |
| Fibroblast | F3 COL11A1+ mCAF | GO:0090103 | cochlea morphogenesis | 3/182 | 26/16509 | 10.47 | 0.00284 | 0.0412 | 3 | CTHRC1/PTK7/WNT5A |
| Fibroblast | F3 COL11A1+ mCAF | GO:1900048 | positive regulation of hemostasis | 3/182 | 26/16509 | 10.47 | 0.00284 | 0.0412 | 3 | PLAU/F2R/PLAT |
| Fibroblast | F3 COL11A1+ mCAF | GO:0150063 | visual system development | 12/182 | 429/16509 | 2.54 | 0.00292 | 0.042 | 12 | INHBA/COL5A1/COL5A2/MFAP2/STRA6/WNT2/THY1/COL8A1/RCN1/RBP4/WNT5A/TWIST1 |
| Fibroblast | F3 COL11A1+ mCAF | GO:0022612 | gland morphogenesis | 6/182 | 128/16509 | 4.25 | 0.00296 | 0.0423 | 6 | SULF1/SULF2/PERP/MMP2/WNT5A/MDK |
| Fibroblast | F3 COL11A1+ mCAF | GO:0043588 | skin development | 10/182 | 321/16509 | 2.83 | 0.00303 | 0.0431 | 10 | INHBA/COL5A1/COL1A1/COL3A1/COL5A2/COL1A2/COL6A1/ADAMTS2/WNT5A/FLNB |
| Fibroblast | F3 COL11A1+ mCAF | GO:0046718 | symbiont entry into host cell | 6/182 | 129/16509 | 4.22 | 0.00307 | 0.0436 | 6 | LGALS1/CTSB/P4HB/CD55/MYH9/LGALS9 |
| Fibroblast | F3 COL11A1+ mCAF | GO:0050819 | negative regulation of coagulation | 4/182 | 55/16509 | 6.6 | 0.00314 | 0.044 | 4 | PLAU/FAP/PLAT/PLAUR |
| Fibroblast | F3 COL11A1+ mCAF | GO:0060688 | regulation of morphogenesis of a branching structure | 4/182 | 55/16509 | 6.6 | 0.00314 | 0.044 | 4 | SULF1/WNT2/WNT5A/MDK |
| Fibroblast | F3 COL11A1+ mCAF | GO:0048596 | embryonic camera-type eye morphogenesis | 3/182 | 27/16509 | 10.08 | 0.00317 | 0.044 | 3 | STRA6/WNT5A/TWIST1 |
| Fibroblast | F3 COL11A1+ mCAF | GO:0052372 | modulation by symbiont of entry into host | 3/182 | 27/16509 | 10.08 | 0.00317 | 0.044 | 3 | LGALS1/P4HB/LGALS9 |
| Fibroblast | F3 COL11A1+ mCAF | GO:0001936 | regulation of endothelial cell proliferation | 6/182 | 130/16509 | 4.19 | 0.00319 | 0.044 | 6 | SULF1/SPARC/MYDGF/HSPG2/WNT5A/MDK |
| Fibroblast | F3 COL11A1+ mCAF | GO:0051701 | biological process involved in interaction with host | 7/182 | 174/16509 | 3.65 | 0.0032 | 0.044 | 7 | LGALS1/CTSB/LRRC15/P4HB/CD55/MYH9/LGALS9 |
| Fibroblast | F3 COL11A1+ mCAF | GO:0048880 | sensory system development | 12/182 | 436/16509 | 2.5 | 0.00332 | 0.0455 | 12 | INHBA/COL5A1/COL5A2/MFAP2/STRA6/WNT2/THY1/COL8A1/RCN1/RBP4/WNT5A/TWIST1 |
| Fibroblast | F3 COL11A1+ mCAF | GO:0042471 | ear morphogenesis | 6/182 | 132/16509 | 4.12 | 0.00344 | 0.0467 | 6 | CTHRC1/COL11A1/SPARC/PTK7/WNT5A/TWIST1 |
| Fibroblast | F3 COL11A1+ mCAF | GO:0097529 | myeloid leukocyte migration | 8/182 | 224/16509 | 3.24 | 0.00345 | 0.0467 | 8 | RARRES2/NBL1/MMP14/CNN2/PPIB/PERP/MMP2/MDK |
| Fibroblast | F3 COL11A1+ mCAF | GO:0048145 | regulation of fibroblast proliferation | 5/182 | 92/16509 | 4.93 | 0.0035 | 0.0468 | 5 | FN1/PDGFC/WNT2/SPHK1/WNT5A |
| Fibroblast | F3 COL11A1+ mCAF | GO:0060740 | prostate gland epithelium morphogenesis | 3/182 | 28/16509 | 9.72 | 0.00352 | 0.0468 | 3 | SULF1/MMP2/WNT5A |
| Fibroblast | F3 COL11A1+ mCAF | GO:0090594 | inflammatory response to wounding | 3/182 | 28/16509 | 9.72 | 0.00352 | 0.0468 | 3 | F2R/DSG2/MDK |
| Fibroblast | F3 COL11A1+ mCAF | GO:1905523 | positive regulation of macrophage migration | 3/182 | 28/16509 | 9.72 | 0.00352 | 0.0468 | 3 | RARRES2/MMP14/MDK |
| Fibroblast | F3 COL11A1+ mCAF | GO:0048704 | embryonic skeletal system morphogenesis | 5/182 | 93/16509 | 4.88 | 0.00366 | 0.0484 | 5 | COL11A1/MMP14/RUNX2/CHST11/TWIST1 |
| Fibroblast | F3 COL11A1+ mCAF | GO:0050878 | regulation of body fluid levels | 11/182 | 385/16509 | 2.59 | 0.00369 | 0.0485 | 11 | PLAU/COL3A1/FN1/PDPN/F2R/FAP/MYH9/PLAT/PLAUR/PDIA3/MYL12A |
| Fibroblast | F3 COL11A1+ mCAF | GO:0044409 | symbiont entry into host | 6/182 | 134/16509 | 4.06 | 0.00371 | 0.0485 | 6 | LGALS1/CTSB/P4HB/CD55/MYH9/LGALS9 |
| Fibroblast | F3 COL11A1+ mCAF | GO:0030595 | leukocyte chemotaxis | 8/182 | 227/16509 | 3.2 | 0.00374 | 0.0487 | 8 | RARRES2/NBL1/PPIB/PERP/MMP2/WNT5A/MDK/LGALS9 |
| Fibroblast | F3 COL11A1+ mCAF | GO:0002690 | positive regulation of leukocyte chemotaxis | 5/182 | 94/16509 | 4.82 | 0.00384 | 0.0497 | 5 | RARRES2/PERP/WNT5A/MDK/LGALS9 |
| Fibroblast | F3 COL11A1+ mCAF | GO:0001958 | endochondral ossification | 3/182 | 29/16509 | 9.38 | 0.0039 | 0.0499 | 3 | COL1A1/MMP14/RUNX2 |
| Fibroblast | F3 COL11A1+ mCAF | GO:0003156 | regulation of animal organ formation | 3/182 | 29/16509 | 9.38 | 0.0039 | 0.0499 | 3 | SULF1/WNT2/WNT5A |

| compartment | subcluster | GO_ID | Description | GeneRatio | BgRatio | FoldEnrichment | pvalue | p_adjust | Count | geneID |
|---|---|---|---|---|---|---|---|---|---|---|
| Fibroblast | F3 COL11A1+ mCAF | GO:0036075 | replacement ossification | 3/182 | 29/16509 | 9.38 | 0.0039 | 0.0499 | 3 | COL1A1/MMP14/RUNX2 |
| Fibroblast | F4 ADAMDEC1+ Fib | GO:0019730 | antimicrobial humoral response | 14/124 | 169/16509 | 11.03 | 3.31e-11 | 8.43e-08 | 14 | CCL11/CCL13/CXCL1/CXCL6/CCL7/CCL2/CXCL14/JCHAIN/CCL8/CXCL12/TSLP/IGHA2/BCL3/IGHA1 |
| Fibroblast | F4 ADAMDEC1+ Fib | GO:0006959 | humoral immune response | 16/124 | 289/16509 | 7.37 | 5.12e-10 | 6.52e-07 | 16 | CCL11/CCL13/CXCL1/CXCL6/CFD/CCL7/CCL2/CXCL14/A2M/JCHAIN/CCL8/CXCL12/TSLP/IGHA2/BCL3/IGHA1 |
| Fibroblast | F4 ADAMDEC1+ Fib | GO:0061844 | antimicrobial humoral immune response mediated by antimicrobial peptide | 10/124 | 122/16509 | 10.91 | 2.69e-08 | 2.28e-05 | 10 | CCL11/CCL13/CXCL1/CXCL6/CCL7/CCL2/CXCL14/CCL8/CXCL12/TSLP |
| Fibroblast | F4 ADAMDEC1+ Fib | GO:0048245 | eosinophil chemotaxis | 5/124 | 20/16509 | 33.28 | 3.12e-07 | 0.000199 | 5 | CCL11/CCL13/CCL7/CCL2/CCL8 |
| Fibroblast | F4 ADAMDEC1+ Fib | GO:0030198 | extracellular matrix organization | 13/124 | 310/16509 | 5.58 | 5.96e-07 | 0.000233 | 13 | FOXF2/FGFR4/MATN2/MFAP4/LTBP4/QSOX1/TGFBI/FBLN5/EMILIN1/HAS2/FBLN1/POSTN/BCL3 |
| Fibroblast | F4 ADAMDEC1+ Fib | GO:0043062 | extracellular structure organization | 13/124 | 311/16509 | 5.57 | 6.18e-07 | 0.000233 | 13 | FOXF2/FGFR4/MATN2/MFAP4/LTBP4/QSOX1/TGFBI/FBLN5/EMILIN1/HAS2/FBLN1/POSTN/BCL3 |
| Fibroblast | F4 ADAMDEC1+ Fib | GO:0045229 | external encapsulating structure organization | 13/124 | 312/16509 | 5.55 | 6.4e-07 | 0.000233 | 13 | FOXF2/FGFR4/MATN2/MFAP4/LTBP4/QSOX1/TGFBI/FBLN5/EMILIN1/HAS2/FBLN1/POSTN/BCL3 |
| Fibroblast | F4 ADAMDEC1+ Fib | GO:0072677 | eosinophil migration | 5/124 | 24/16509 | 27.74 | 8.36e-07 | 0.000235 | 5 | CCL11/CCL13/CCL7/CCL2/CCL8 |
| Fibroblast | F4 ADAMDEC1+ Fib | GO:0085029 | extracellular matrix assembly | 6/124 | 44/16509 | 18.16 | 8.89e-07 | 0.000235 | 6 | MFAP4/LTBP4/QSOX1/FBLN5/EMILIN1/HAS2 |
| Fibroblast | F4 ADAMDEC1+ Fib | GO:0048251 | elastic fiber assembly | 4/124 | 11/16509 | 48.41 | 9.61e-07 | 0.000235 | 4 | MFAP4/LTBP4/FBLN5/EMILIN1 |
| Fibroblast | F4 ADAMDEC1+ Fib | GO:0050673 | epithelial cell proliferation | 15/124 | 438/16509 | 4.56 | 1.02e-06 | 0.000235 | 15 | BMP5/CCL11/CCL2/APOE/DACH1/ECM1/BMP4/HAS2/PTN/ST8SIA1/CXCL12/EDNRB/IGFBP5/ATOH8/ISL1 |
| Fibroblast | F4 ADAMDEC1+ Fib | GO:0006935 | chemotaxis | 15/124 | 452/16509 | 4.42 | 1.5e-06 | 0.000311 | 15 | CCL11/CCL13/CXCL1/CXCL6/EPHA7/CCL7/CCL2/CXCL14/BMP4/LSP1/PTN/CCL8/CXCL12/EDNRB/IL34 |
| Fibroblast | F4 ADAMDEC1+ Fib | GO:0042330 | taxis | 15/124 | 454/16509 | 4.4 | 1.59e-06 | 0.000311 | 15 | CCL11/CCL13/CXCL1/CXCL6/EPHA7/CCL7/CCL2/CXCL14/BMP4/LSP1/PTN/CCL8/CXCL12/EDNRB/IL34 |
| Fibroblast | F4 ADAMDEC1+ Fib | GO:0050678 | regulation of epithelial cell proliferation | 13/124 | 359/16509 | 4.82 | 3.05e-06 | 0.000542 | 13 | BMP5/CCL11/CCL2/APOE/ECM1/BMP4/HAS2/PTN/ST8SIA1/CXCL12/EDNRB/ATOH8/ISL1 |
| Fibroblast | F4 ADAMDEC1+ Fib | GO:0070098 | chemokine-mediated signaling pathway | 7/124 | 84/16509 | 11.09 | 3.19e-06 | 0.000542 | 7 | CCL11/CCL13/CXCL6/CCL7/CCL2/CCL8/CXCL12 |
| Fibroblast | F4 ADAMDEC1+ Fib | GO:1905809 | negative regulation of synapse organization | 4/124 | 15/16509 | 35.5 | 3.88e-06 | 0.000587 | 4 | EPHA7/APOE/NEGR1/PGRMC1 |
| Fibroblast | F4 ADAMDEC1+ Fib | GO:0060326 | cell chemotaxis | 12/124 | 311/16509 | 5.14 | 3.92e-06 | 0.000587 | 12 | CCL11/CCL13/CXCL1/CXCL6/CCL7/CCL2/CXCL14/PTN/CCL8/CXCL12/EDNRB/IL34 |
| Fibroblast | F4 ADAMDEC1+ Fib | GO:1990868 | response to chemokine | 7/124 | 91/16509 | 10.24 | 5.46e-06 | 0.000732 | 7 | CCL11/CCL13/CXCL6/CCL7/CCL2/CCL8/CXCL12 |
| Fibroblast | F4 ADAMDEC1+ Fib | GO:1990869 | cellular response to chemokine | 7/124 | 91/16509 | 10.24 | 5.46e-06 | 0.000732 | 7 | CCL11/CCL13/CXCL6/CCL7/CCL2/CCL8/CXCL12 |
| Fibroblast | F4 ADAMDEC1+ Fib | GO:0050900 | leukocyte migration | 13/124 | 381/16509 | 4.54 | 5.83e-06 | 0.000742 | 13 | BMP5/CCL11/CCL13/CXCL6/CCL7/CCL2/ECM1/EMILIN1/PTN/CCL8/CXCL12/EDNRB/IL34 |
| Fibroblast | F4 ADAMDEC1+ Fib | GO:0097529 | myeloid leukocyte migration | 10/124 | 224/16509 | 5.94 | 7.33e-06 | 0.000889 | 10 | CCL11/CCL13/CXCL6/CCL7/CCL2/EMILIN1/CCL8/CXCL12/EDNRB/IL34 |
| Fibroblast | F4 ADAMDEC1+ Fib | GO:0030595 | leukocyte chemotaxis | 10/124 | 227/16509 | 5.87 | 8.24e-06 | 0.000954 | 10 | CCL11/CCL13/CXCL6/CCL7/CCL2/PTN/CCL8/CXCL12/EDNRB/IL34 |
| Fibroblast | F4 ADAMDEC1+ Fib | GO:0072073 | kidney epithelium development | 8/124 | 151/16509 | 7.05 | 1.83e-05 | 0.00203 | 8 | EPHA7/TCF21/WNT2B/BMP4/DLL1/EDNRB/HOXD11/HS3ST3B1 |
| Fibroblast | F4 ADAMDEC1+ Fib | GO:2000146 | negative regulation of cell motility | 11/124 | 306/16509 | 4.79 | 1.95e-05 | 0.00207 | 11 | BMP5/CCL2/APOE/CYGB/MCTP1/DACH1/BMP4/EMILIN1/FBLN1/CXCL12/IGFBP5 |
| Fibroblast | F4 ADAMDEC1+ Fib | GO:0071621 | granulocyte chemotaxis | 7/124 | 117/16509 | 7.97 | 2.85e-05 | 0.00289 | 7 | CCL11/CCL13/CXCL6/CCL7/CCL2/CCL8/IL34 |

| compartment | subcluster | GO_ID | Description | GeneRatio | BgRatio | FoldEnrichment | pvalue | p_adjust | Count | geneID |
|---|---|---|---|---|---|---|---|---|---|---|
| Fibroblast | F4 ADAMDEC1+ Fib | GO:0001763 | morphogenesis of a branching structure | 9/124 | 210/16509 | 5.71 | 2.95e-05 | 0.00289 | 9 | CCL11/EPHA7/TCF21/WNT2B/BMP4/HOXD11/SOCS3/HS3ST3B1/HHIP |
| Fibroblast | F4 ADAMDEC1+ Fib | GO:0040013 | negative regulation of locomotion | 11/124 | 324/16509 | 4.52 | 3.3e-05 | 0.00312 | 11 | BMP5/CCL2/APOE/CYGB/MCTP1/DACH1/BMP4/EMILIN1/FBLN1/CXCL12/IGFBP5 |
| Fibroblast | F4 ADAMDEC1+ Fib | GO:0072009 | nephron epithelium development | 7/124 | 122/16509 | 7.64 | 3.73e-05 | 0.00339 | 7 | TCF21/WNT2B/BMP4/DLL1/EDNRB/HOXD11/HS3ST3B1 |
| Fibroblast | F4 ADAMDEC1+ Fib | GO:0002685 | regulation of leukocyte migration | 9/124 | 222/16509 | 5.4 | 4.55e-05 | 0.004 | 9 | BMP5/CCL7/CCL2/ECM1/EMILIN1/PTN/CCL8/CXCL12/IL34 |
| Fibroblast | F4 ADAMDEC1+ Fib | GO:0001936 | regulation of endothelial cell proliferation | 7/124 | 130/16509 | 7.17 | 5.6e-05 | 0.00475 | 7 | CCL11/CCL2/APOE/ECM1/BMP4/CXCL12/ATOH8 |
| Fibroblast | F4 ADAMDEC1+ Fib | GO:0031640 | killing of cells of another organism | 7/124 | 132/16509 | 7.06 | 6.17e-05 | 0.00491 | 7 | CCL11/CCL13/CXCL1/CXCL6/CXCL14/CCL8/CXCL12 |
| Fibroblast | F4 ADAMDEC1+ Fib | GO:0141061 | disruption of cell in another organism | 7/124 | 132/16509 | 7.06 | 6.17e-05 | 0.00491 | 7 | CCL11/CCL13/CXCL1/CXCL6/CXCL14/CCL8/CXCL12 |
| Fibroblast | F4 ADAMDEC1+ Fib | GO:1902931 | negative regulation of alcohol biosynthetic process | 3/124 | 11/16509 | 36.31 | 6.53e-05 | 0.00504 | 3 | BMP5/APOE/DKK3 |
| Fibroblast | F4 ADAMDEC1+ Fib | GO:0030336 | negative regulation of cell migration | 10/124 | 291/16509 | 4.58 | 6.9e-05 | 0.00517 | 10 | BMP5/CCL2/APOE/CYGB/MCTP1/DACH1/BMP4/EMILIN1/CXCL12/IGFBP5 |
| Fibroblast | F4 ADAMDEC1+ Fib | GO:0141060 | disruption of anatomical structure in another organism | 7/124 | 136/16509 | 6.85 | 7.45e-05 | 0.00542 | 7 | CCL11/CCL13/CXCL1/CXCL6/CXCL14/CCL8/CXCL12 |
| Fibroblast | F4 ADAMDEC1+ Fib | GO:0071675 | regulation of mononuclear cell migration | 7/124 | 141/16509 | 6.61 | 9.36e-05 | 0.00645 | 7 | BMP5/CCL7/CCL2/ECM1/EMILIN1/CXCL12/IL34 |
| Fibroblast | F4 ADAMDEC1+ Fib | GO:0001658 | branching involved in ureteric bud morphogenesis | 5/124 | 61/16509 | 10.91 | 9.37e-05 | 0.00645 | 5 | TCF21/WNT2B/BMP4/HOXD11/HS3ST3B1 |
| Fibroblast | F4 ADAMDEC1+ Fib | GO:0072080 | nephron tubule development | 6/124 | 99/16509 | 8.07 | 0.000101 | 0.00667 | 6 | TCF21/WNT2B/BMP4/DLL1/HOXD11/HS3ST3B1 |
| Fibroblast | F4 ADAMDEC1+ Fib | GO:0097530 | granulocyte migration | 7/124 | 143/16509 | 6.52 | 0.000102 | 0.00667 | 7 | CCL11/CCL13/CXCL6/CCL7/CCL2/CCL8/IL34 |
| Fibroblast | F4 ADAMDEC1+ Fib | GO:0032341 | aldosterone metabolic process | 3/124 | 13/16509 | 30.72 | 0.000112 | 0.00713 | 3 | BMP5/DKK3/EDNRB |
| Fibroblast | F4 ADAMDEC1+ Fib | GO:0061138 | morphogenesis of a branching epithelium | 8/124 | 197/16509 | 5.41 | 0.00012 | 0.00748 | 8 | CCL11/TCF21/WNT2B/BMP4/HOXD11/SOCS3/HS3ST3B1/HHIP |
| Fibroblast | F4 ADAMDEC1+ Fib | GO:0061326 | renal tubule development | 6/124 | 104/16509 | 7.68 | 0.000132 | 0.00803 | 6 | TCF21/WNT2B/BMP4/DLL1/HOXD11/HS3ST3B1 |
| Fibroblast | F4 ADAMDEC1+ Fib | GO:0008212 | mineralocorticoid metabolic process | 3/124 | 14/16509 | 28.53 | 0.000142 | 0.00829 | 3 | BMP5/DKK3/EDNRB |
| Fibroblast | F4 ADAMDEC1+ Fib | GO:0050679 | positive regulation of epithelial cell proliferation | 8/124 | 202/16509 | 5.27 | 0.000143 | 0.00829 | 8 | BMP5/CCL11/ECM1/BMP4/HAS2/PTN/ST8SIA1/CXCL12 |
| Fibroblast | F4 ADAMDEC1+ Fib | GO:0060675 | ureteric bud morphogenesis | 5/124 | 67/16509 | 9.94 | 0.000147 | 0.00831 | 5 | TCF21/WNT2B/BMP4/HOXD11/HS3ST3B1 |
| Fibroblast | F4 ADAMDEC1+ Fib | GO:0001935 | endothelial cell proliferation | 7/124 | 153/16509 | 6.09 | 0.000156 | 0.00853 | 7 | CCL11/CCL2/APOE/ECM1/BMP4/CXCL12/ATOH8 |
| Fibroblast | F4 ADAMDEC1+ Fib | GO:0072171 | mesonephric tubule morphogenesis | 5/124 | 68/16509 | 9.79 | 0.000157 | 0.00853 | 5 | TCF21/WNT2B/BMP4/HOXD11/HS3ST3B1 |
| Fibroblast | F4 ADAMDEC1+ Fib | GO:0001906 | cell killing | 9/124 | 269/16509 | 4.45 | 0.000197 | 0.0103 | 9 | CCL11/CCL13/CXCL1/CXCL6/CTSC/CXCL14/CCL8/CXCL12/HLA-C |
| Fibroblast | F4 ADAMDEC1+ Fib | GO:0072006 | nephron development | 7/124 | 159/16509 | 5.86 | 0.000198 | 0.0103 | 7 | TCF21/WNT2B/BMP4/DLL1/EDNRB/HOXD11/HS3ST3B1 |
| Fibroblast | F4 ADAMDEC1+ Fib | GO:0010721 | negative regulation of cell development | 9/124 | 271/16509 | 4.42 | 0.000208 | 0.0106 | 9 | CCL11/EPHA7/TMEM176B/TMEM176A/NEGR1/BMP4/IRF1/PTN/EDNRB |
| Fibroblast | F4 ADAMDEC1+ Fib | GO:0048754 | branching morphogenesis of an epithelial tube | 7/124 | 164/16509 | 5.68 | 0.000239 | 0.0119 | 7 | CCL11/TCF21/WNT2B/BMP4/HOXD11/HS3ST3B1/HHIP |
| Fibroblast | F4 ADAMDEC1+ Fib | GO:0060562 | epithelial tube morphogenesis | 10/124 | 340/16509 | 3.92 | 0.000246 | 0.0119 | 10 | BMP5/CCL11/EPHA7/TCF21/WNT2B/BMP4/DLL1/HOXD11/HS3ST3B1/HHIP |
| Fibroblast | F4 ADAMDEC1+ Fib | GO:0010712 | regulation of collagen metabolic process | 4/124 | 41/16509 | 12.99 | 0.000248 | 0.0119 | 4 | CYGB/MFAP4/BMP4/EMILIN1 |
| Fibroblast | F4 ADAMDEC1+ Fib | GO:0072078 | nephron tubule morphogenesis | 5/124 | 76/16509 | 8.76 | 0.000266 | 0.0125 | 5 | TCF21/WNT2B/BMP4/HOXD11/HS3ST3B1 |
| Fibroblast | F4 ADAMDEC1+ Fib | GO:0071674 | mononuclear cell migration | 8/124 | 223/16509 | 4.78 | 0.000281 | 0.013 | 8 | BMP5/CCL7/CCL2/ECM1/EMILIN1/CXCL12/EDNRB/IL34 |
| Fibroblast | F4 ADAMDEC1+ Fib | GO:0072088 | nephron epithelium morphogenesis | 5/124 | 78/16509 | 8.53 | 0.0003 | 0.0137 | 5 | TCF21/WNT2B/BMP4/HOXD11/HS3ST3B1 |
| Fibroblast | F4 ADAMDEC1+ Fib | GO:0072028 | nephron morphogenesis | 5/124 | 80/16509 | 8.32 | 0.000338 | 0.0151 | 5 | TCF21/WNT2B/BMP4/HOXD11/HS3ST3B1 |

| compartment | subcluster | GO_ID | Description | GeneRatio | BgRatio | FoldEnrichment | pvalue | p_adjust | Count | geneID |
|---|---|---|---|---|---|---|---|---|---|---|
| Fibroblast | F4 ADAMDEC1+ Fib | GO:0061333 | renal tubule morphogenesis | 5/124 | 81/16509 | 8.22 | 0.000358 | 0.0157 | 5 | TCF21/WNT2B/BMP4/HOXD11/HS3ST3B1 |
| Fibroblast | F4 ADAMDEC1+ Fib | GO:0060263 | regulation of respiratory burst | 3/124 | 19/16509 | 21.02 | 0.000367 | 0.0158 | 3 | JCHAIN/IGHA2/IGHA1 |
| Fibroblast | F4 ADAMDEC1+ Fib | GO:0002686 | negative regulation of leukocyte migration | 4/124 | 50/16509 | 10.65 | 0.000535 | 0.0227 | 4 | BMP5/CCL2/EMILIN1/CXCL12 |
| Fibroblast | F4 ADAMDEC1+ Fib | GO:0035270 | endocrine system development | 6/124 | 136/16509 | 5.87 | 0.000564 | 0.0236 | 6 | BMP5/PITX1/BMP4/DLL1/DKK3/ISL1 |
| Fibroblast | F4 ADAMDEC1+ Fib | GO:0003323 | type B pancreatic cell development | 3/124 | 22/16509 | 18.16 | 0.000574 | 0.0236 | 3 | BMP5/BMP4/DLL1 |
| Fibroblast | F4 ADAMDEC1+ Fib | GO:0048762 | mesenchymal cell differentiation | 8/124 | 252/16509 | 4.23 | 0.000633 | 0.0256 | 8 | BMP5/TCF21/FOXF2/NCAM1/BMP4/HAS2/EDNRB/ISL1 |
| Fibroblast | F4 ADAMDEC1+ Fib | GO:0010894 | negative regulation of steroid biosynthetic process | 3/124 | 23/16509 | 17.37 | 0.000656 | 0.0261 | 3 | BMP5/APOE/DKK3 |
| Fibroblast | F4 ADAMDEC1+ Fib | GO:0003014 | renal system process | 6/124 | 141/16509 | 5.67 | 0.000682 | 0.0267 | 6 | BMP4/HAS2/JCHAIN/EDNRB/IGHA2/IGHA1 |
| Fibroblast | F4 ADAMDEC1+ Fib | GO:0001822 | kidney development | 9/124 | 320/16509 | 3.74 | 0.000696 | 0.0269 | 9 | EPHA7/TCF21/WNT2B/BMP4/DLL1/HAS2/EDNRB/HOXD11/HS3ST3B1 |
| Fibroblast | F4 ADAMDEC1+ Fib | GO:0002690 | positive regulation of leukocyte chemotaxis | 5/124 | 94/16509 | 7.08 | 0.00071 | 0.027 | 5 | CCL7/CCL2/PTN/CXCL12/IL34 |
| Fibroblast | F4 ADAMDEC1+ Fib | GO:0019216 | regulation of lipid metabolic process | 9/124 | 326/16509 | 3.68 | 0.000794 | 0.0298 | 9 | BMP5/APOE/PLPP1/FGFR4/FABP5/SCT/RAB38/DKK3/FMO4 |
| Fibroblast | F4 ADAMDEC1+ Fib | GO:0060571 | morphogenesis of an epithelial fold | 3/124 | 25/16509 | 15.98 | 0.000843 | 0.0298 | 3 | BMP5/WNT2B/BMP4 |
| Fibroblast | F4 ADAMDEC1+ Fib | GO:0001657 | ureteric bud development | 5/124 | 98/16509 | 6.79 | 0.000858 | 0.0298 | 5 | TCF21/WNT2B/BMP4/HOXD11/HS3ST3B1 |
| Fibroblast | F4 ADAMDEC1+ Fib | GO:0060993 | kidney morphogenesis | 5/124 | 98/16509 | 6.79 | 0.000858 | 0.0298 | 5 | TCF21/WNT2B/BMP4/HOXD11/HS3ST3B1 |
| Fibroblast | F4 ADAMDEC1+ Fib | GO:0002687 | positive regulation of leukocyte migration | 6/124 | 148/16509 | 5.4 | 0.000879 | 0.0298 | 6 | CCL7/CCL2/PTN/CCL8/CXCL12/IL34 |
| Fibroblast | F4 ADAMDEC1+ Fib | GO:0072001 | renal system development | 9/124 | 331/16509 | 3.62 | 0.000885 | 0.0298 | 9 | EPHA7/TCF21/WNT2B/BMP4/DLL1/HAS2/EDNRB/HOXD11/HS3ST3B1 |
| Fibroblast | F4 ADAMDEC1+ Fib | GO:0072163 | mesonephric epithelium development | 5/124 | 99/16509 | 6.72 | 0.000898 | 0.0298 | 5 | TCF21/WNT2B/BMP4/HOXD11/HS3ST3B1 |
| Fibroblast | F4 ADAMDEC1+ Fib | GO:0072164 | mesonephric tubule development | 5/124 | 99/16509 | 6.72 | 0.000898 | 0.0298 | 5 | TCF21/WNT2B/BMP4/HOXD11/HS3ST3B1 |
| Fibroblast | F4 ADAMDEC1+ Fib | GO:0050727 | regulation of inflammatory response | 10/124 | 402/16509 | 3.31 | 0.000909 | 0.0298 | 10 | CCL7/CTSC/APOE/NFKBIA/EDNRB/CASP1/ISL1/SOCS3/FABP4/TSLP |
| Fibroblast | F4 ADAMDEC1+ Fib | GO:0010976 | positive regulation of neuron projection development | 6/124 | 149/16509 | 5.36 | 0.000911 | 0.0298 | 6 | BMP5/STMN2/APOE/NEGR1/PTN/TOX |
| Fibroblast | F4 ADAMDEC1+ Fib | GO:0032371 | regulation of sterol transport | 4/124 | 58/16509 | 9.18 | 0.00094 | 0.0298 | 4 | APOE/ABCA8/PLTP/NFKBIA |
| Fibroblast | F4 ADAMDEC1+ Fib | GO:0032374 | regulation of cholesterol transport | 4/124 | 58/16509 | 9.18 | 0.00094 | 0.0298 | 4 | APOE/ABCA8/PLTP/NFKBIA |
| Fibroblast | F4 ADAMDEC1+ Fib | GO:0010875 | positive regulation of cholesterol efflux | 3/124 | 26/16509 | 15.36 | 0.000948 | 0.0298 | 3 | APOE/ABCA8/PLTP |
| Fibroblast | F4 ADAMDEC1+ Fib | GO:0090280 | positive regulation of calcium ion import | 3/124 | 26/16509 | 15.36 | 0.000948 | 0.0298 | 3 | CCL2/CXCL12/ISL1 |
| Fibroblast | F4 ADAMDEC1+ Fib | GO:0042742 | defense response to bacterium | 9/124 | 339/16509 | 3.53 | 0.00105 | 0.0322 | 9 | CXCL6/IRF8/EMILIN1/JCHAIN/CASP1/TSLP/IGHA2/BCL3/IGHA1 |
| Fibroblast | F4 ADAMDEC1+ Fib | GO:0045939 | negative regulation of steroid metabolic process | 3/124 | 27/16509 | 14.79 | 0.00106 | 0.0322 | 3 | BMP5/APOE/DKK3 |
| Fibroblast | F4 ADAMDEC1+ Fib | GO:1905517 | macrophage migration | 4/124 | 60/16509 | 8.88 | 0.00107 | 0.0322 | 4 | CCL2/EMILIN1/EDNRB/IL34 |
| Fibroblast | F4 ADAMDEC1+ Fib | GO:0001823 | mesonephros development | 5/124 | 103/16509 | 6.46 | 0.00107 | 0.0322 | 5 | TCF21/WNT2B/BMP4/HOXD11/HS3ST3B1 |
| Fibroblast | F4 ADAMDEC1+ Fib | GO:0060425 | lung morphogenesis | 4/124 | 61/16509 | 8.73 | 0.00114 | 0.0337 | 4 | TCF21/WNT2B/BMP4/HHIP |
| Fibroblast | F4 ADAMDEC1+ Fib | GO:0006956 | complement activation | 4/124 | 62/16509 | 8.59 | 0.00121 | 0.035 | 4 | CFD/A2M/IGHA2/IGHA1 |
| Fibroblast | F4 ADAMDEC1+ Fib | GO:0014910 | regulation of smooth muscle cell migration | 4/124 | 62/16509 | 8.59 | 0.00121 | 0.035 | 4 | BMP4/HAS2/POSTN/IGFBP5 |
| Fibroblast | F4 ADAMDEC1+ Fib | GO:0003309 | type B pancreatic cell differentiation | 3/124 | 29/16509 | 13.77 | 0.00131 | 0.0375 | 3 | BMP5/BMP4/DLL1 |
| Fibroblast | F4 ADAMDEC1+ Fib | GO:0003094 | glomerular filtration | 3/124 | 30/16509 | 13.31 | 0.00145 | 0.0405 | 3 | JCHAIN/IGHA2/IGHA1 |
| Fibroblast | F4 ADAMDEC1+ Fib | GO:1902882 | regulation of response to oxidative stress | 3/124 | 30/16509 | 13.31 | 0.00145 | 0.0405 | 3 | MCTP1/FBLN5/NCOA7 |
| Fibroblast | F4 ADAMDEC1+ Fib | GO:0045445 | myoblast differentiation | 5/124 | 111/16509 | 6.0 | 0.0015 | 0.0413 | 5 | PITX1/BMP4/DLL1/CCL8/ISL1 |
| Fibroblast | F4 ADAMDEC1+ Fib | GO:0009612 | response to mechanical stimulus | 7/124 | 224/16509 | 4.16 | 0.00152 | 0.0413 | 7 | NFKBIA/IRF1/PTN/POSTN/CXCL12/TMEM150C/CASP1 |
| Fibroblast | F4 ADAMDEC1+ Fib | GO:0060390 | regulation of SMAD protein signal transduction | 4/124 | 66/16509 | 8.07 | 0.00153 | 0.0413 | 4 | BMP5/BMP4/EMILIN1/ATOH8 |

| compartment | subcluster | GO_ID | Description | GeneRatio | BgRatio | FoldEnrichment | pvalue | p_adjust | Count | geneID |
|---|---|---|---|---|---|---|---|---|---|---|
| Fibroblast | F4 ADAMDEC1+ Fib | GO:0060037 | pharyngeal system development | 3/124 | 31/16509 | 12.88 | 0.00159 | 0.0423 | 3 | BMP5/BMP4/ISL1 |
| Fibroblast | F4 ADAMDEC1+ Fib | GO:0071676 | negative regulation of mononuclear cell migration | 3/124 | 31/16509 | 12.88 | 0.00159 | 0.0423 | 3 | BMP5/CCL2/EMILIN1 |
| Fibroblast | F4 ADAMDEC1+ Fib | GO:0050729 | positive regulation of inflammatory response | 6/124 | 167/16509 | 4.78 | 0.00164 | 0.043 | 6 | CCL7/CTSC/NFKBIA/CASP1/FABP4/TSLP |
| Fibroblast | F4 ADAMDEC1+ Fib | GO:0014909 | smooth muscle cell migration | 4/124 | 68/16509 | 7.83 | 0.0017 | 0.0443 | 4 | BMP4/HAS2/POSTN/IGFBP5 |
| Fibroblast | F4 ADAMDEC1+ Fib | GO:1904018 | positive regulation of vasculature development | 6/124 | 169/16509 | 4.73 | 0.00174 | 0.0446 | 6 | CCL11/ECM1/EMILIN1/DLL1/CXCL12/ISL1 |
| Fibroblast | F4 ADAMDEC1+ Fib | GO:0060441 | epithelial tube branching involved in lung morphogenesis | 3/124 | 32/16509 | 12.48 | 0.00175 | 0.0446 | 3 | WNT2B/BMP4/HHIP |
| Fibroblast | F4 ADAMDEC1+ Fib | GO:0045165 | cell fate commitment | 8/124 | 296/16509 | 3.6 | 0.00178 | 0.0448 | 8 | PITX1/WNT2B/ETS2/BMP4/DLL1/TOX/ISL1/SOCS3 |
| Fibroblast | F4 ADAMDEC1+ Fib | GO:2000401 | regulation of lymphocyte migration | 4/124 | 69/16509 | 7.72 | 0.0018 | 0.0449 | 4 | CCL7/CCL2/ECM1/CXCL12 |
| Fibroblast | F4 ADAMDEC1+ Fib | GO:0001837 | epithelial to mesenchymal transition | 6/124 | 171/16509 | 4.67 | 0.00184 | 0.0456 | 6 | BMP5/FOXF2/NCAM1/BMP4/HAS2/ISL1 |
| Fibroblast | F4 ADAMDEC1+ Fib | GO:0097205 | renal filtration | 3/124 | 33/16509 | 12.1 | 0.00191 | 0.0469 | 3 | JCHAIN/IGHA2/IGHA1 |

### Table S3

Table S3. Univariable and stage-adjusted multivariable Cox models for the four-gene score.

| cohort | model | term | n | HR | lower95 | upper95 | p |
|---|---|---|---|---|---|---|---|
| TCGA | Univariable | 4-gene score (High vs Low) | 593 | 1.48 | 1.01 | 2.18 | 0.0447 |
| TCGA | Multivariable (adj. stage) | 4-gene score (High vs Low) | 593 | 1.31 | 0.89 | 1.93 | 0.166 |
| GSE39582 | Univariable | 4-gene score (High vs Low) | 519 | 1.43 | 1.03 | 1.99 | 0.0332 |
| GSE39582 | Multivariable (adj. stage) | 4-gene score (High vs Low) | 519 | 1.54 | 1.1 | 2.15 | 0.0115 |

### Table S4

Table S4. Single-gene diagnostic performance in TCGA and GSE39582.

| cohort | gene | n | AUC | CI95 | sensitivity | specificity | accuracy |
|---|---|---|---|---|---|---|---|
| GSE39582 | SOX4 | 585 | 0.971 | 0.953-0.989 | 0.887 | 1.0 | 0.891 |
| GSE39582 | SLC1A5 | 585 | 0.926 | 0.895-0.958 | 0.836 | 0.947 | 0.839 |
| GSE39582 | TPT1 | 585 | 0.627 | 0.558-0.696 | 0.443 | 0.947 | 0.46 |
| GSE39582 | TSC22D1 | 585 | 0.544 | 0.459-0.628 | 0.466 | 0.789 | 0.477 |
| TCGA (Train) | SOX4 | 474 | 0.967 | 0.951-0.983 | 0.913 | 0.972 | 0.918 |
| TCGA (Train) | SLC1A5 | 474 | 0.958 | 0.936-0.98 | 0.893 | 0.972 | 0.899 |
| TCGA (Train) | TPT1 | 474 | 0.633 | 0.55-0.716 | 0.653 | 0.694 | 0.656 |
| TCGA (Train) | TSC22D1 | 474 | 0.593 | 0.523-0.663 | 0.381 | 0.917 | 0.422 |
| TCGA (Validation) | SOX4 | 202 | 0.966 | 0.94-0.993 | 0.904 | 0.933 | 0.906 |
| TCGA (Validation) | SLC1A5 | 202 | 0.924 | 0.871-0.977 | 0.866 | 0.8 | 0.861 |
| TCGA (Validation) | TSC22D1 | 202 | 0.681 | 0.594-0.767 | 0.385 | 1.0 | 0.431 |
| TCGA (Validation) | TPT1 | 202 | 0.657 | 0.557-0.757 | 0.642 | 0.667 | 0.644 |

### Table S5a

Table S5a. NicheNet ligand-activity ranking for the fibroblast-to-epithelial prediction.

| ligand | rank | auroc | aupr | aupr_corrected | pearson |
|---|---|---|---|---|---|
| TGFB1 | 1 | 0.5664 | 0.25 | 0.0492 | 0.0915 |
| ANG | 2 | 0.5583 | 0.2467 | 0.046 | 0.0686 |
| TSPAN3 | 3 | 0.5553 | 0.2454 | 0.0447 | 0.0817 |
| EFNA1 | 4 | 0.5565 | 0.2448 | 0.044 | 0.0768 |
| TIMP1 | 5 | 0.5514 | 0.2445 | 0.0438 | 0.0527 |
| PROS1 | 6 | 0.5492 | 0.244 | 0.0433 | 0.0659 |
| SERPING1 | 7 | 0.5571 | 0.2439 | 0.0432 | 0.0764 |
| CLDN3 | 8 | 0.5523 | 0.2436 | 0.0429 | 0.0755 |
| C1QB | 9 | 0.5547 | 0.2434 | 0.0426 | 0.073 |
| GDF11 | 10 | 0.5493 | 0.243 | 0.0423 | 0.0625 |
| SEMA5B | 11 | 0.5512 | 0.2428 | 0.042 | 0.0715 |
| ADAM9 | 12 | 0.5488 | 0.2425 | 0.0417 | 0.0661 |
| JAM3 | 13 | 0.5555 | 0.2422 | 0.0415 | 0.0744 |
| CSF1 | 14 | 0.5504 | 0.2418 | 0.0411 | 0.0702 |
| ANXA1 | 15 | 0.5475 | 0.2417 | 0.041 | 0.0744 |
| NRP1 | 16 | 0.5455 | 0.2416 | 0.0409 | 0.0625 |
| TGFB3 | 17 | 0.5521 | 0.2415 | 0.0407 | 0.0727 |
| THBS2 | 18 | 0.5397 | 0.2409 | 0.0402 | 0.0681 |
| MMP14 | 19 | 0.5502 | 0.2409 | 0.0402 | 0.0697 |
| EDIL3 | 20 | 0.5495 | 0.2407 | 0.0399 | 0.0731 |
| LAMC1 | 21 | 0.5459 | 0.2404 | 0.0397 | 0.0557 |
| MCAM | 22 | 0.5488 | 0.2403 | 0.0396 | 0.0608 |
| COL18A1 | 23 | 0.5479 | 0.2403 | 0.0395 | 0.0613 |
| LTBP1 | 24 | 0.5447 | 0.2401 | 0.0394 | 0.0675 |
| ENG | 25 | 0.5532 | 0.2401 | 0.0393 | 0.068 |
| ANXA2 | 26 | 0.5502 | 0.24 | 0.0393 | 0.0595 |
| COL14A1 | 27 | 0.5473 | 0.24 | 0.0393 | 0.0673 |
| VCAN | 28 | 0.5473 | 0.24 | 0.0393 | 0.0672 |
| NAMPT | 29 | 0.5461 | 0.24 | 0.0392 | 0.0634 |
| GPI | 30 | 0.548 | 0.2399 | 0.0392 | 0.0503 |
| PLAT | 31 | 0.5424 | 0.2398 | 0.039 | 0.0475 |
| CD44 | 32 | 0.5468 | 0.2396 | 0.0389 | 0.0581 |
| APOE | 33 | 0.5456 | 0.2396 | 0.0388 | 0.0617 |
| TFPI | 34 | 0.5449 | 0.2394 | 0.0387 | 0.0518 |
| COL5A1 | 35 | 0.5476 | 0.2394 | 0.0387 | 0.0588 |
| COL5A2 | 36 | 0.5473 | 0.2394 | 0.0386 | 0.0592 |
| GDF15 | 37 | 0.5458 | 0.2393 | 0.0385 | 0.0517 |
| COL6A1 | 38 | 0.5468 | 0.2393 | 0.0385 | 0.0648 |
| MDK | 39 | 0.5454 | 0.2392 | 0.0385 | 0.0585 |
| THBS1 | 40 | 0.5429 | 0.2392 | 0.0385 | 0.0556 |
| LUM | 41 | 0.5455 | 0.2391 | 0.0384 | 0.0639 |
| LGALS3BP | 42 | 0.5499 | 0.2391 | 0.0384 | 0.0624 |
| NID1 | 43 | 0.5339 | 0.239 | 0.0383 | 0.0605 |
| CTSD | 44 | 0.542 | 0.239 | 0.0383 | 0.0664 |
| S100A10 | 45 | 0.5468 | 0.2389 | 0.0382 | 0.0634 |
| GSTP1 | 46 | 0.544 | 0.2388 | 0.0381 | 0.0652 |
| FAM3C | 47 | 0.5475 | 0.2387 | 0.038 | 0.0542 |
| THY1 | 48 | 0.5445 | 0.2385 | 0.0378 | 0.0645 |
| LAMA5 | 49 | 0.5436 | 0.2385 | 0.0378 | 0.0646 |
| PTN | 50 | 0.55 | 0.2382 | 0.0374 | 0.0556 |
| HSP90B1 | 51 | 0.5427 | 0.2382 | 0.0374 | 0.0608 |
| COL4A2 | 52 | 0.5495 | 0.2381 | 0.0373 | 0.0572 |
| LGALS3 | 53 | 0.5472 | 0.238 | 0.0373 | 0.0524 |
| FN1 | 54 | 0.5471 | 0.2379 | 0.0372 | 0.0479 |
| COL4A1 | 55 | 0.5503 | 0.2379 | 0.0372 | 0.0542 |
| GRN | 56 | 0.548 | 0.2379 | 0.0372 | 0.0381 |
| UXS1 | 57 | 0.5474 | 0.2379 | 0.0372 | 0.0686 |
| LAMA4 | 58 | 0.5462 | 0.2378 | 0.037 | 0.0671 |
| PTPRK | 59 | 0.5465 | 0.2377 | 0.0369 | 0.0659 |

| ligand | rank | auroc | aupr | aupr_corrected | pearson |
|---|---|---|---|---|---|
| CXCL2 | 60 | 0.5459 | 0.2375 | 0.0368 | 0.0631 |
| SEMA4A | 61 | 0.542 | 0.2375 | 0.0367 | 0.0531 |
| LRPAP1 | 62 | 0.541 | 0.2373 | 0.0366 | 0.0608 |
| BMP1 | 63 | 0.5468 | 0.2373 | 0.0366 | 0.0704 |
| INHBA | 64 | 0.5459 | 0.2372 | 0.0365 | 0.0589 |
| CD99 | 65 | 0.5469 | 0.237 | 0.0362 | 0.0371 |
| RGMA | 66 | 0.5457 | 0.2369 | 0.0362 | 0.0678 |
| S100A4 | 67 | 0.5478 | 0.2369 | 0.0362 | 0.0641 |
| MFGE8 | 68 | 0.5425 | 0.2369 | 0.0361 | 0.0515 |
| A2M | 69 | 0.5476 | 0.2368 | 0.0361 | 0.0646 |
| DCN | 70 | 0.5445 | 0.2368 | 0.036 | 0.0411 |
| CFH | 71 | 0.5434 | 0.2365 | 0.0358 | 0.0648 |
| COL1A2 | 72 | 0.5376 | 0.2365 | 0.0357 | 0.0578 |
| TIMP3 | 73 | 0.5423 | 0.2364 | 0.0357 | 0.0596 |
| HLA-A | 74 | 0.5434 | 0.2364 | 0.0357 | 0.0633 |
| ADM | 75 | 0.5464 | 0.2364 | 0.0356 | 0.0362 |
| COL6A2 | 76 | 0.5413 | 0.2363 | 0.0355 | 0.0599 |
| BGN | 77 | 0.5454 | 0.2361 | 0.0354 | 0.0415 |
| CCL5 | 78 | 0.5442 | 0.236 | 0.0353 | 0.0346 |
| LGALS1 | 79 | 0.5454 | 0.236 | 0.0352 | 0.0319 |
| SDC2 | 80 | 0.5392 | 0.2358 | 0.035 | 0.0616 |
| ENTPD1 | 81 | 0.5471 | 0.2357 | 0.0349 | 0.0368 |
| BACE2 | 82 | 0.5389 | 0.2355 | 0.0348 | 0.0656 |
| COL3A1 | 83 | 0.542 | 0.2355 | 0.0347 | 0.0601 |
| ECM1 | 84 | 0.5414 | 0.2354 | 0.0347 | 0.0583 |
| CALR | 85 | 0.5475 | 0.2354 | 0.0347 | 0.0364 |
| PLTP | 86 | 0.5479 | 0.2352 | 0.0345 | 0.0416 |
| SEMA4C | 87 | 0.5481 | 0.2351 | 0.0343 | 0.0647 |
| JAG1 | 88 | 0.5383 | 0.2351 | 0.0343 | 0.043 |
| CD9 | 89 | 0.5463 | 0.2348 | 0.0341 | 0.0239 |
| CD320 | 90 | 0.5434 | 0.2348 | 0.034 | 0.0495 |
| COL1A1 | 91 | 0.5441 | 0.2347 | 0.0339 | 0.0502 |
| B2M | 92 | 0.542 | 0.2346 | 0.0339 | 0.0505 |
| COL6A3 | 93 | 0.5412 | 0.2346 | 0.0338 | 0.0585 |
| LAMB2 | 94 | 0.5386 | 0.2346 | 0.0338 | 0.0577 |
| GNAS | 95 | 0.5413 | 0.2345 | 0.0338 | 0.0261 |
| HBEGF | 96 | 0.5484 | 0.2342 | 0.0335 | 0.0176 |
| S100A9 | 97 | 0.5373 | 0.2341 | 0.0334 | 0.0578 |
| NUCB2 | 98 | 0.5329 | 0.2337 | 0.0329 | 0.0534 |
| HLA-DRB1 | 99 | 0.5392 | 0.2336 | 0.0328 | 0.0481 |
| CD47 | 100 | 0.5368 | 0.2332 | 0.0324 | 0.0456 |
| APP | 101 | 0.544 | 0.2332 | 0.0324 | 0.0319 |
| SPP1 | 102 | 0.5375 | 0.2331 | 0.0324 | 0.0428 |
| TGFBI | 103 | 0.5384 | 0.2331 | 0.0323 | 0.0557 |
| PSAP | 104 | 0.5399 | 0.2329 | 0.0322 | 0.0552 |
| EFEMP2 | 105 | 0.5382 | 0.2321 | 0.0314 | 0.0518 |
| ANGPTL4 | 106 | 0.5421 | 0.2321 | 0.0313 | 0.0365 |
| SERPINE2 | 107 | 0.5409 | 0.232 | 0.0312 | 0.0546 |
| HLA-DRA | 108 | 0.5344 | 0.2319 | 0.0311 | 0.0433 |
| ANGPT2 | 109 | 0.5433 | 0.2318 | 0.031 | 0.0315 |
| SARAF | 110 | 0.5403 | 0.2316 | 0.0308 | 0.0555 |
| IFITM1 | 111 | 0.5456 | 0.2311 | 0.0304 | 0.0544 |
| AREG | 112 | 0.5456 | 0.2308 | 0.03 | 0.029 |
| NPNT | 113 | 0.5424 | 0.23 | 0.0292 | 0.0543 |
| TIMP2 | 114 | 0.5349 | 0.2279 | 0.0272 | 0.0477 |
| ADGRE5 | 115 | 0.526 | 0.2269 | 0.0261 | 0.0374 |
| ANGPT1 | 116 | 0.5394 | 0.2265 | 0.0257 | 0.0226 |
| DKK3 | 117 | 0.5309 | 0.2251 | 0.0243 | 0.0438 |
| CXCL8 | 118 | 0.5366 | 0.2241 | 0.0234 | 0.0096 |
| HMGB1 | 119 | 0.5361 | 0.2241 | 0.0234 | 0.036 |
| MIF | 120 | 0.5349 | 0.2228 | 0.022 | 0.0276 |
| NGF | 121 | 0.5325 | 0.2223 | 0.0216 | 0.0218 |
| GAS6 | 122 | 0.5277 | 0.2216 | 0.0209 | 0.0394 |
| PAM | 123 | 0.5198 | 0.2213 | 0.0206 | 0.0302 |

| ligand | rank | auroc | aupr | aupr_corrected | pearson |
|---|---|---|---|---|---|
| CLEC11A | 124 | 0.5237 | 0.2206 | 0.0199 | 0.0354 |
| PDGFA | 125 | 0.5347 | 0.2176 | 0.0168 | -0.0014 |
| TNFSF12 | 126 | 0.5257 | 0.2166 | 0.0158 | 0.0033 |
| CTHRC1 | 127 | 0.5207 | 0.2164 | 0.0156 | 0.0306 |
| LTBP3 | 128 | 0.5091 | 0.2118 | 0.011 | 0.0146 |
| SPON2 | 129 | 0.5105 | 0.2094 | 0.0087 | 0.0162 |

### Table S5b

Table S5b. NicheNet ligand-target regulatory-potential links.

| ligand | target | regulatory_potential |
|---|---|---|
| TGFB1 | EDN1 | 0.3021 |
| TGFB1 | SGK1 | 0.2972 |
| TGFB1 | ID3 | 0.2959 |
| TGFB1 | GADD45B | 0.2937 |
| TGFB1 | KLF10 | 0.293 |
| TGFB1 | TGFBI | 0.282 |
| TGFB1 | THBS1 | 0.2811 |
| TGFB1 | FN1 | 0.281 |
| TGFB1 | ADORA2B | 0.2803 |
| TGFB1 | SOX4 | 0.276 |
| TGFB1 | SKIL | 0.2383 |
| TGFB1 | TNFRSF12A | 0.2369 |
| TGFB1 | CITED2 | 0.236 |
| TGFB1 | PDGFA | 0.2246 |
| TGFB1 | ID1 | 0.2243 |
| TGFB1 | JUN | 0.2204 |
| TGFB1 | LBH | 0.2197 |
| TGFB1 | LARP6 | 0.2165 |
| TGFB1 | HES1 | 0.2146 |
| TGFB3 | GADD45B | 0.1962 |
| TGFB3 | ABCA1 | 0.1909 |
| TGFB3 | CEBPB | 0.19 |
| TGFB3 | KLHL24 | 0.1894 |
| TGFB3 | GABARAPL1 | 0.1887 |
| TGFB1 | KLF6 | 0.1882 |
| TGFB1 | PLK2 | 0.1852 |
| TGFB1 | SMTN | 0.1845 |
| TGFB1 | ADM | 0.1801 |
| TGFB1 | GEM | 0.1797 |
| TGFB1 | TIMP3 | 0.1739 |
| TGFB1 | CYB5A | 0.1658 |
| TGFB1 | SNAI1 | 0.1657 |
| TGFB1 | PHLDB1 | 0.1632 |
| TGFB1 | ALDH3A2 | 0.1608 |
| GDF11 | ID1 | 0.1607 |
| TGFB1 | MME | 0.1562 |
| TGFB1 | CLDN1 | 0.1556 |
| TGFB1 | CCL3 | 0.1538 |
| TGFB3 | SOX9 | 0.1531 |
| TGFB3 | SKIL | 0.1499 |
| TGFB1 | UBL3 | 0.1495 |
| TGFB1 | ZFP36L2 | 0.1487 |
| TGFB1 | ALDH3A1 | 0.1485 |
| TGFB3 | PIM1 | 0.1481 |
| CSF1 | TGFBI | 0.1461 |
| CSF1 | NUPR1 | 0.1455 |
| GDF11 | PTK7 | 0.145 |
| GDF11 | EDN1 | 0.1447 |
| TGFB3 | ODC1 | 0.1445 |
| CSF1 | ABCA1 | 0.1443 |
| CSF1 | CTSZ | 0.1443 |
| CSF1 | CYB5A | 0.1441 |
| GDF11 | CLU | 0.1438 |
| CSF1 | DAB2 | 0.1438 |
| CSF1 | ODC1 | 0.1438 |
| GDF11 | FILIP1L | 0.1437 |
| GDF11 | PTPRR | 0.1436 |
| CSF1 | ACP2 | 0.1435 |
| TGFB3 | YPEL5 | 0.1434 |

| ligand | target | regulatory_potential |
|---|---|---|
| GDF11 | CHN2 | 0.1429 |
| CSF1 | G0S2 | 0.1428 |
| CSF1 | TUBA4A | 0.1427 |
| GDF11 | ADORA2B | 0.1424 |
| CSF1 | SLC39A8 | 0.1424 |
| GDF11 | ESF1 | 0.1421 |
| GDF11 | MMP11 | 0.142 |
| CSF1 | IDH1 | 0.1418 |
| GDF11 | ARL4D | 0.1416 |
| GDF11 | AKAP9 | 0.1415 |
| GDF11 | ABCA1 | 0.1413 |
| GDF11 | SGK1 | 0.1413 |
| CSF1 | PLA2G15 | 0.1411 |
| GDF11 | GADD45B | 0.141 |
| GDF11 | BCL2L11 | 0.1409 |
| CSF1 | RGS2 | 0.1408 |
| GDF11 | MFAP2 | 0.1406 |
| TGFB3 | DNASE2 | 0.1406 |
| GDF11 | HEY1 | 0.1403 |
| CSF1 | VAT1 | 0.1403 |
| GDF11 | SOX4 | 0.1398 |
| CSF1 | MARCKSL1 | 0.1397 |
| GDF11 | GDPD5 | 0.1393 |
| GDF11 | KLHL24 | 0.1391 |
| CSF1 | PLPP3 | 0.1385 |
| TGFB3 | BCAR3 | 0.1382 |
| CSF1 | FUCA1 | 0.1379 |
| CSF1 | DHRS3 | 0.1374 |
| GDF11 | PLPP3 | 0.1365 |
| GDF11 | MYCL | 0.1364 |
| CSF1 | IL1B | 0.1341 |
| TGFB3 | SMOX | 0.1331 |
| TGFB3 | SPP1 | 0.1325 |
| TGFB3 | PKIG | 0.1299 |
| TGFB3 | DUSP1 | 0.1291 |
| CSF1 | DUSP1 | 0.1279 |
| TGFB3 | MAFF | 0.1275 |
| TGFB3 | PNRC1 | 0.1267 |
| TGFB3 | THBS1 | 0.125 |
| TGFB3 | ID3 | 0.1249 |
| TGFB3 | TNFRSF11B | 0.1228 |
| TGFB3 | FILIP1L | 0.1208 |
| TGFB3 | AMOTL2 | 0.1202 |
| TGFB3 | CDC42EP3 | 0.1198 |
| TGFB3 | CCL3 | 0.1195 |
| TGFB3 | AREG | 0.1194 |
| TGFB3 | EVA1A | 0.1189 |
| TGFB3 | NET1 | 0.118 |
| TGFB3 | TUBA1A | 0.118 |
| TGFB3 | ADM | 0.1177 |
| TGFB3 | RGS2 | 0.1176 |
| TGFB3 | RGCC | 0.1175 |
| TGFB3 | TSPAN2 | 0.1175 |
| TGFB3 | GEM | 0.1172 |
| TGFB3 | BTG2 | 0.1168 |
| TGFB3 | DNAJB9 | 0.1168 |
| TGFB3 | CITED2 | 0.1159 |
| TGFB3 | TUBA4A | 0.1158 |
| TGFB3 | PLIN2 | 0.1156 |
| TGFB3 | G0S2 | 0.1154 |
| TGFB3 | RSBN1 | 0.1152 |
| TGFB3 | HILPDA | 0.1149 |
| TGFB3 | RASD1 | 0.1147 |
| TGFB3 | KDM7A | 0.114 |

| ligand | target | regulatory_potential |
| --- | --- | --- |
| TGFB3 | ALDH2 | 0.1138 |
| TGFB3 | MYLIP | 0.1138 |
| TGFB3 | YPEL3 | 0.1125 |
| TGFB3 | GLUL | 0.1123 |
| TGFB3 | ETS2 | 0.0828 |
| CSF1 | TCEAL4 | 0.0824 |
| CSF1 | HNMT | 0.082 |
| CSF1 | CCND1 | 0.0819 |
| TGFB3 | PAK1IP1 | 0.0818 |
| TGFB3 | BRIX1 | 0.0816 |
| TGFB3 | UBL3 | 0.0816 |
| TGFB3 | HSPA9 | 0.0815 |
| CSF1 | QDPR | 0.0812 |
| TGFB3 | ID2 | 0.0809 |
| CSF1 | LAMP1 | 0.0806 |
| CSF1 | NXT1 | 0.0805 |
| CSF1 | SLCO3A1 | 0.0805 |
| CSF1 | NXF1 | 0.0803 |
| CSF1 | RNF13 | 0.0803 |
| CSF1 | GRN | 0.0802 |
| CSF1 | GRPEL1 | 0.0802 |
| CSF1 | PEPD | 0.0801 |
| CSF1 | EHD1 | 0.08 |
| GDF11 | MTSS1 | 0.0799 |
| CSF1 | CTSA | 0.0798 |
| GDF11 | TUBB2B | 0.0797 |
| CSF1 | FKBP9 | 0.0797 |
| CSF1 | HEXB | 0.0794 |
| CSF1 | METTL9 | 0.0793 |
| CSF1 | NABP1 | 0.0791 |
| CSF1 | SLC11A2 | 0.079 |
| CSF1 | ALDH3A2 | 0.0789 |
| CSF1 | DNASE2 | 0.0788 |
| GDF11 | LGALSL | 0.0784 |
| GDF11 | EEA1 | 0.0779 |
| CSF1 | CRYZ | 0.0775 |
| CSF1 | PHYH | 0.0756 |
| CSF1 | FN1 | 0.0746 |
| CSF1 | THBS1 | 0.0743 |
| CSF1 | ID2 | 0.0741 |
| CSF1 | ID1 | 0.0739 |
| CSF1 | GDF15 | 0.0737 |
| CSF1 | PIM1 | 0.0737 |
| CSF1 | ATF3 | 0.0733 |
| CSF1 | JUN | 0.0732 |
| CSF1 | TIMP3 | 0.0728 |
| CSF1 | GADD45B | 0.0727 |
| CSF1 | CXCL2 | 0.0724 |
| CSF1 | SPP1 | 0.0719 |
| CSF1 | CCL3 | 0.0717 |
| TIMP1 | HMGCS2 | 0.0702 |
| GDF11 | CASC15 | 0.0696 |
| GDF11 | PIM1 | 0.0691 |
| GDF11 | RBP1 | 0.069 |
| NRP1 | SEMA3A | 0.066 |
| ANXA1 | IL1B | 0.0439 |
| ANXA1 | CXCL2 | 0.0401 |
| PROS1 | DDX18 | 0.0247 |
| PROS1 | DAPK2 | 0.0241 |
| TIMP1 | CCND1 | 0.0234 |
| PROS1 | CCND1 | 0.0232 |
| PROS1 | PHYH | 0.0231 |
| MMP14 | SPINT1 | 0.0224 |
| ADAM9 | CHIC2 | 0.0223 |

| ligand | target | regulatory_potential |
|---|---|---|
| ADAM9 | UQCRB | 0.0212 |
| ADAM9 | VPS4A | 0.0212 |
| ANG | CCND1 | 0.0205 |
| NRP1 | CCND1 | 0.0202 |
| TIMP1 | JUN | 0.0197 |
| CLDN3 | CCND1 | 0.0194 |
| THBS2 | CCND1 | 0.0186 |
| EDIL3 | CCND1 | 0.0186 |
| JAM3 | CCND1 | 0.0179 |
| ANXA1 | CCND1 | 0.0176 |
| PROS1 | JUN | 0.0171 |
| MMP14 | CCND1 | 0.0171 |
| TSPAN3 | CCND1 | 0.0169 |
| EDIL3 | JUN | 0.0167 |
| EFNA1 | CCND1 | 0.0166 |
| THBS2 | HES1 | 0.0166 |
| ANG | ID2 | 0.0158 |
| SERPING1 | CCND1 | 0.0157 |
| PROS1 | ID2 | 0.0154 |
| ANG | GDF15 | 0.0151 |
| TIMP1 | ID2 | 0.0151 |
| PROS1 | GADD45A | 0.015 |
| PROS1 | EDN1 | 0.0149 |
| PROS1 | ID1 | 0.0148 |
| ANG | ATF3 | 0.0145 |
| PROS1 | THBS1 | 0.0145 |
| C1QB | CCND1 | 0.0145 |
| PROS1 | ATF3 | 0.0143 |
| TIMP1 | EDN1 | 0.0142 |
| NRP1 | JUN | 0.0142 |
| ANG | JUN | 0.0141 |
| PROS1 | GDF15 | 0.014 |
| PROS1 | IL1B | 0.014 |
| TIMP1 | CEBPA | 0.0136 |
| ANG | EDN1 | 0.0134 |
| CLDN3 | JUN | 0.0134 |
| TIMP1 | ID1 | 0.0133 |
| NRP1 | ID2 | 0.0131 |
| TIMP1 | IL1B | 0.013 |
| PROS1 | CITED2 | 0.013 |
| SEMA5B | CCND1 | 0.013 |
| ANXA1 | JUN | 0.013 |
| ANG | GADD45A | 0.0129 |
| TIMP1 | GDF15 | 0.0129 |
| PROS1 | GADD45B | 0.0129 |
| PROS1 | ID3 | 0.0129 |
| ADAM9 | CCND1 | 0.0129 |
| CLDN3 | ID2 | 0.0128 |
| TIMP1 | ATF3 | 0.0127 |
| PROS1 | DDIT3 | 0.0126 |
| JAM3 | JUN | 0.0126 |
| ANG | DUSP1 | 0.0125 |
| TIMP1 | GADD45A | 0.0125 |
| NRP1 | ID1 | 0.0125 |
| THBS2 | ID1 | 0.0125 |
| PROS1 | APP | 0.0124 |
| PROS1 | FN1 | 0.0124 |
| ANG | ID3 | 0.0123 |
| PROS1 | HES1 | 0.0123 |
| PROS1 | SGK1 | 0.0123 |
| PROS1 | DUSP1 | 0.0122 |
| MMP14 | JUN | 0.0122 |
| TIMP1 | THBS1 | 0.0121 |
| TIMP1 | DDIT3 | 0.012 |

| ligand | target | regulatory_potential |
| --- | --- | --- |
| TIMP1 | DUSP1 | 0.012 |
| TIMP1 | ID3 | 0.012 |
| ANG | IL1B | 0.0119 |
| ANG | THBS1 | 0.0119 |
| CLDN3 | GDF15 | 0.0119 |
| ANG | PTEN | 0.0118 |
| TIMP1 | FN1 | 0.0118 |
| PROS1 | TGIF1 | 0.0118 |
| CLDN3 | ATF3 | 0.0118 |
| ANG | BTG2 | 0.0117 |
| ANG | ID1 | 0.0117 |
| PROS1 | CDK6 | 0.0117 |
| PROS1 | TRIB1 | 0.0117 |
| CLDN3 | EDN1 | 0.0117 |
| PROS1 | BTG2 | 0.0116 |
| PROS1 | PIM1 | 0.0116 |
| CLDN3 | THBS1 | 0.0116 |
| CLDN3 | SGK1 | 0.0115 |
| TIMP1 | PTEN | 0.0114 |
| CLDN3 | ID1 | 0.0114 |
| NRP1 | DUSP1 | 0.0114 |
| THBS2 | JUN | 0.0114 |
| MMP14 | ID2 | 0.0112 |
| ANG | SLC2A1 | 0.0111 |
| EFNA1 | JUN | 0.0111 |
| TIMP1 | HES1 | 0.0111 |
| ANXA1 | ID2 | 0.0111 |
| NRP1 | ATF3 | 0.0111 |
| NRP1 | GADD45A | 0.0111 |
| NRP1 | GDF15 | 0.0111 |
| THBS2 | HEY1 | 0.0111 |
| TIMP1 | APP | 0.011 |
| PROS1 | ODC1 | 0.011 |
| NRP1 | EDN1 | 0.011 |
| NRP1 | ID3 | 0.011 |
| EDIL3 | GADD45A | 0.011 |
| EDIL3 | THBS1 | 0.011 |
| PROS1 | CEBPB | 0.0109 |
| NRP1 | THBS1 | 0.0109 |
| ANG | GADD45B | 0.0108 |
| PROS1 | PTEN | 0.0108 |
| CLDN3 | GADD45A | 0.0108 |
| CLDN3 | IL1B | 0.0108 |
| JAM3 | ID2 | 0.0108 |
| NRP1 | IL1B | 0.0108 |
| MMP14 | ID1 | 0.0108 |
| ANG | DHRS3 | 0.0107 |
| ANG | TGFBI | 0.0107 |
| TIMP1 | CEBPB | 0.0107 |
| TIMP1 | TGIF1 | 0.0107 |
| PROS1 | PPP1R15A | 0.0107 |
| CLDN3 | CITED2 | 0.0107 |
| NRP1 | DDIT3 | 0.0107 |
| ANG | SGK1 | 0.0106 |
| TIMP1 | SLC2A1 | 0.0106 |
| PROS1 | ADM | 0.0106 |
| PROS1 | RBPJ | 0.0106 |
| PROS1 | ZMYND8 | 0.0106 |
| ANXA1 | ID1 | 0.0106 |
| NRP1 | HES1 | 0.0106 |
| THBS2 | EDN1 | 0.0106 |
| EDIL3 | ID2 | 0.0106 |
| ANG | CEBPA | 0.0105 |
| ANG | FN1 | 0.0105 |

| ligand | target | regulatory_potential |
|---|---|---|
| ANG | HES1 | 0.0105 |
| TIMP1 | NR1D1 | 0.0105 |
| TIMP1 | PIM1 | 0.0105 |
| TIMP1 | RARA | 0.0105 |
| TIMP1 | SPP1 | 0.0105 |
| PROS1 | CDKN1B | 0.0105 |
| CLDN3 | DDIT3 | 0.0105 |
| CLDN3 | DUSP1 | 0.0105 |
| ANXA1 | ATF3 | 0.0105 |
| NRP1 | PIM1 | 0.0105 |
| TSPAN3 | ID1 | 0.0104 |
| TIMP1 | TRIB1 | 0.0104 |
| PROS1 | ARID5B | 0.0104 |
| PROS1 | KLF5 | 0.0104 |
| NRP1 | TRIB1 | 0.0104 |
| ANG | TGIF1 | 0.0103 |
| EFNA1 | ID2 | 0.0103 |
| TIMP1 | SGK1 | 0.0103 |
| PROS1 | CXCL2 | 0.0103 |
| PROS1 | KLF6 | 0.0103 |
| PROS1 | TGFBI | 0.0103 |
| MMP14 | EDN1 | 0.0103 |
| EDIL3 | EDN1 | 0.0103 |
| ANG | ADM | 0.0102 |
| ANG | PLK2 | 0.0102 |
| TIMP1 | SOX9 | 0.0102 |
| PROS1 | EFNA1 | 0.0102 |
| PROS1 | PRSS23 | 0.0102 |
| PROS1 | RARA | 0.0102 |
| CLDN3 | FN1 | 0.0102 |
| ANXA1 | EDN1 | 0.0102 |
| TIMP1 | PDK4 | 0.0101 |
| PROS1 | DHRS3 | 0.0101 |
| PROS1 | HNRNPR | 0.0101 |
| PROS1 | RGS2 | 0.0101 |
| CLDN3 | ADM | 0.0101 |
| NRP1 | CITED2 | 0.0101 |
| NRP1 | TGIF1 | 0.0101 |
| ANG | CITED2 | 0.01 |
| ANG | DDIT3 | 0.01 |
| TIMP1 | BCL2L11 | 0.01 |
| TIMP1 | GEM | 0.01 |
| TIMP1 | NFE2L2 | 0.01 |
| PROS1 | SLC2A1 | 0.01 |
| JAM3 | HES1 | 0.01 |
| EDIL3 | CITED2 | 0.01 |
| TIMP1 | CITED2 | 0.0099 |
| PROS1 | SLC3A2 | 0.0099 |
| PROS1 | SOX9 | 0.0099 |
| CLDN3 | HES1 | 0.0099 |
| JAM3 | EDN1 | 0.0099 |
| JAM3 | IL1B | 0.0099 |
| EDIL3 | ATF3 | 0.0099 |
| ANG | CLDN1 | 0.0098 |
| TIMP1 | G0S2 | 0.0098 |
| PROS1 | MALAT1 | 0.0098 |
| CLDN3 | ID3 | 0.0098 |
| JAM3 | GADD45A | 0.0098 |
| ANXA1 | GDF15 | 0.0098 |
| NRP1 | GADD45B | 0.0098 |
| THBS2 | GDF15 | 0.0098 |
| EDIL3 | IL1B | 0.0098 |
| ANG | NFIA | 0.0097 |
| TSPAN3 | ID2 | 0.0097 |

| ligand | target | regulatory_potential |
| --- | --- | --- |
| TIMP1 | KLF6 | 0.0097 |
| PROS1 | GCLC | 0.0097 |
| PROS1 | HSP90AB1 | 0.0097 |
| PROS1 | IGFBP4 | 0.0097 |
| PROS1 | SLC1A5 | 0.0097 |
| SERPING1 | JUN | 0.0097 |
| JAM3 | ID1 | 0.0097 |
| NRP1 | CDK6 | 0.0097 |
| EDIL3 | ABCB1 | 0.0097 |
| EDIL3 | ID1 | 0.0097 |
| EDIL3 | ID3 | 0.0097 |
| ANG | CDKN1B | 0.0096 |
| TSPAN3 | CITED2 | 0.0096 |
| TSPAN3 | EDN1 | 0.0096 |
| TIMP1 | ODC1 | 0.0096 |
| PROS1 | ETFB | 0.0096 |
| PROS1 | NFE2L2 | 0.0096 |
| JAM3 | CITED2 | 0.0096 |
| ANXA1 | DUSP1 | 0.0096 |
| THBS2 | ID2 | 0.0096 |
| MMP14 | THBS1 | 0.0096 |
| EDIL3 | CDKN1B | 0.0096 |
| ANG | ODC1 | 0.0095 |
| TSPAN3 | HES1 | 0.0095 |
| TIMP1 | ABCA1 | 0.0095 |
| TIMP1 | CTNNB1 | 0.0095 |
| PROS1 | ABCB1 | 0.0095 |
| PROS1 | CEBPA | 0.0095 |
| PROS1 | FOXO3 | 0.0095 |
| PROS1 | RTN4 | 0.0095 |
| CLDN3 | BTG2 | 0.0095 |
| CLDN3 | PIM1 | 0.0095 |
| C1QB | JUN | 0.0095 |
| JAM3 | THBS1 | 0.0095 |
| THBS2 | DDIT3 | 0.0095 |
| THBS2 | GADD45A | 0.0095 |
| THBS2 | IL1B | 0.0095 |
| ANG | CEBPB | 0.0094 |
| ANG | EFNA1 | 0.0094 |
| ANG | HNRNPR | 0.0094 |
| ANG | RGS2 | 0.0094 |
| TSPAN3 | GDF15 | 0.0094 |
| TSPAN3 | JUN | 0.0094 |
| PROS1 | ETS2 | 0.0094 |
| ADAM9 | JUN | 0.0094 |
| JAM3 | DUSP1 | 0.0094 |
| JAM3 | ID3 | 0.0094 |
| ANXA1 | DDIT3 | 0.0094 |
| EDIL3 | ILK | 0.0094 |
| EDIL3 | MALAT1 | 0.0094 |
| ANG | FGFR3 | 0.0093 |
| ANG | MTF2 | 0.0093 |
| TSPAN3 | THBS1 | 0.0093 |
| TIMP1 | ADM | 0.0093 |
| TIMP1 | ETFB | 0.0093 |
| TIMP1 | TGFBI | 0.0093 |
| SERPING1 | ID2 | 0.0093 |
| CLDN3 | CTNNB1 | 0.0093 |
| CLDN3 | GADD45B | 0.0093 |
| CLDN3 | RARA | 0.0093 |
| SEMA5B | JUN | 0.0093 |
| JAM3 | CEBPB | 0.0093 |
| JAM3 | PIM1 | 0.0093 |
| ANXA1 | FN1 | 0.0093 |

| ligand | target | regulatory_potential |
|---|---|---|
| ANXA1 | GADD45A | 0.0093 |
| ANXA1 | GADD45B | 0.0093 |
| NRP1 | ADM | 0.0093 |
| EDIL3 | GDF15 | 0.0093 |
| TSPAN3 | SPP1 | 0.0092 |
| TIMP1 | CDK6 | 0.0092 |
| TIMP1 | DHRS3 | 0.0092 |
| TIMP1 | GADD45B | 0.0092 |
| TIMP1 | PLIN2 | 0.0092 |
| SERPING1 | GDF15 | 0.0092 |
| SERPING1 | IL1B | 0.0092 |
| CLDN3 | CDKN1B | 0.0092 |
| CLDN3 | IGFBP4 | 0.0092 |
| C1QB | IL1B | 0.0092 |
| JAM3 | ATF3 | 0.0092 |
| JAM3 | GDF15 | 0.0092 |
| ANXA1 | THBS1 | 0.0092 |
| NRP1 | FN1 | 0.0092 |
| THBS2 | THBS1 | 0.0092 |
| MMP14 | ATF3 | 0.0092 |
| EDIL3 | KLF6 | 0.0092 |
| ANG | NFE2L2 | 0.0091 |
| ANG | PPM1D | 0.0091 |
| TSPAN3 | IL1B | 0.0091 |
| TIMP1 | KLF10 | 0.0091 |
| TIMP1 | KLF5 | 0.0091 |
| CLDN3 | TGIF1 | 0.0091 |
| CLDN3 | UBC | 0.0091 |
| NRP1 | APP | 0.0091 |
| NRP1 | PTEN | 0.0091 |
| EDIL3 | APP | 0.0091 |
| ANG | IGFBP4 | 0.009 |
| ANG | PIM1 | 0.009 |
| ANG | RRM2B | 0.009 |
| TIMP1 | BTG2 | 0.009 |
| TIMP1 | TIMP3 | 0.009 |
| SERPING1 | GADD45A | 0.009 |
| CLDN3 | CEBPA | 0.009 |
| CLDN3 | CEBPB | 0.009 |
| CLDN3 | PTEN | 0.009 |
| JAM3 | ZMYND8 | 0.009 |
| ANXA1 | TRIB1 | 0.009 |
| NRP1 | BTG2 | 0.009 |
| NRP1 | SOX9 | 0.009 |
| MMP14 | FN1 | 0.009 |
| EDIL3 | HES1 | 0.009 |
| ANG | PROS1 | 0.0089 |
| TSPAN3 | GADD45A | 0.0089 |
| TIMP1 | CDKN1B | 0.0089 |
| TIMP1 | MALAT1 | 0.0089 |
| TIMP1 | RTN4 | 0.0089 |
| SERPING1 | EDN1 | 0.0089 |
| CLDN3 | EFNA1 | 0.0089 |
| JAM3 | GADD45B | 0.0089 |
| THBS2 | RBPJ | 0.0089 |
| EDIL3 | GADD45B | 0.0089 |
| EDIL3 | IGFBP4 | 0.0089 |
| EDIL3 | PRPF6 | 0.0089 |
| ANG | ANXA3 | 0.0088 |
| ANG | CTNNB1 | 0.0088 |
| ANG | ETS2 | 0.0088 |
| TSPAN3 | ETFB | 0.0088 |
| TIMP1 | CPT2 | 0.0088 |
| TIMP1 | IGFBP4 | 0.0088 |

| ligand | target | regulatory_potential |
|---|---|---|
| TIMP1 | NR0B2 | 0.0088 |
| TIMP1 | PPP1R15A | 0.0088 |
| TIMP1 | PRSS23 | 0.0088 |
| SERPING1 | ATF3 | 0.0088 |
| ANXA1 | HES1 | 0.0088 |
| NRP1 | HNRNPR | 0.0088 |
| NRP1 | KLF6 | 0.0088 |
| MMP14 | CITED2 | 0.0088 |
| MMP14 | DDIT3 | 0.0088 |
| MMP14 | DUSP1 | 0.0088 |
| EDIL3 | G0S2 | 0.0088 |
| EDIL3 | PTEN | 0.0088 |
| ANG | RND3 | 0.0087 |
| TIMP1 | MMD | 0.0087 |
| CLDN3 | APP | 0.0087 |
| C1QB | ID2 | 0.0087 |
| JAM3 | ADM | 0.0087 |
| JAM3 | DDIT3 | 0.0087 |
| JAM3 | FN1 | 0.0087 |
| ANXA1 | CITED2 | 0.0087 |
| NRP1 | SGK1 | 0.0087 |
| MMP14 | IL1B | 0.0087 |
| EDIL3 | CRYAB | 0.0087 |
| TSPAN3 | DHRS3 | 0.0086 |
| TSPAN3 | PIM1 | 0.0086 |
| TSPAN3 | TRIB1 | 0.0086 |
| EFNA1 | EDN1 | 0.0086 |
| SERPING1 | ID1 | 0.0086 |
| CLDN3 | CDK6 | 0.0086 |
| JAM3 | APP | 0.0086 |
| JAM3 | CEBPA | 0.0086 |
| JAM3 | GLB1 | 0.0086 |
| JAM3 | TRIB1 | 0.0086 |
| NRP1 | ODC1 | 0.0086 |
| THBS2 | ATF3 | 0.0086 |
| MMP14 | GADD45A | 0.0086 |
| MMP14 | GDF15 | 0.0086 |
| EDIL3 | ADM | 0.0086 |
| EDIL3 | ODC1 | 0.0086 |
| EFNA1 | ID1 | 0.0085 |
| CLDN3 | DHRS3 | 0.0085 |
| CLDN3 | FOXO3 | 0.0085 |
| CLDN3 | ODC1 | 0.0085 |
| CLDN3 | SLC2A1 | 0.0085 |
| CLDN3 | TRIB1 | 0.0085 |
| C1QB | GDF15 | 0.0085 |
| ADAM9 | ID2 | 0.0085 |
| ANXA1 | ID3 | 0.0085 |
| NRP1 | DHRS3 | 0.0085 |
| NRP1 | NFE2L2 | 0.0085 |
| MMP14 | ID3 | 0.0085 |
| MMP14 | SGK1 | 0.0085 |
| MMP14 | TRIB1 | 0.0085 |
| EDIL3 | BTG2 | 0.0085 |
| EDIL3 | FN1 | 0.0085 |
| EDIL3 | PIM1 | 0.0085 |
| EDIL3 | PITX2 | 0.0085 |
| SEMA5B | ID2 | 0.0084 |
| ANXA1 | ADM | 0.0084 |
| NRP1 | CEBPB | 0.0084 |
| NRP1 | RBPJ | 0.0084 |
| MMP14 | APP | 0.0084 |
| TSPAN3 | CDK6 | 0.0083 |
| EFNA1 | GADD45A | 0.0083 |

| ligand | target | regulatory_potential |
|---|---|---|
| SERPING1 | CITED2 | 0.0083 |
| SERPING1 | THBS1 | 0.0083 |
| CLDN3 | ABCB1 | 0.0083 |
| C1QB | ATF3 | 0.0083 |
| ANXA1 | SGK1 | 0.0083 |
| NRP1 | CDKN1B | 0.0083 |
| EDIL3 | DDIT3 | 0.0083 |
| EDIL3 | DUSP1 | 0.0083 |
| EDIL3 | GCLC | 0.0083 |
| TSPAN3 | RGS16 | 0.0082 |
| TSPAN3 | TGFBI | 0.0082 |
| SERPING1 | APP | 0.0082 |
| SERPING1 | PIM1 | 0.0082 |
| CLDN3 | HNRNPR | 0.0082 |
| CLDN3 | PRSS23 | 0.0082 |
| CLDN3 | SPP1 | 0.0082 |
| ANXA1 | PIM1 | 0.0082 |
| NRP1 | STMN1 | 0.0082 |
| NRP1 | ZMYND8 | 0.0082 |
| THBS2 | PTEN | 0.0082 |
| MMP14 | HES1 | 0.0082 |
| TSPAN3 | ADM | 0.0081 |
| TSPAN3 | ATF3 | 0.0081 |
| TSPAN3 | CEBPB | 0.0081 |
| TSPAN3 | CTNNB1 | 0.0081 |
| TSPAN3 | TPT1 | 0.0081 |
| CLDN3 | RTN4 | 0.0081 |
| SEMA5B | ID1 | 0.0081 |
| JAM3 | HNRNPR | 0.0081 |
| JAM3 | SGK1 | 0.0081 |
| NRP1 | KLF5 | 0.0081 |
| THBS2 | CITED2 | 0.0081 |
| THBS2 | FN1 | 0.0081 |
| EDIL3 | RBPJ | 0.0081 |
| EDIL3 | TSPAN12 | 0.0081 |
| EFNA1 | ATF3 | 0.008 |
| EFNA1 | GDF15 | 0.008 |
| EFNA1 | ID3 | 0.008 |
| EFNA1 | THBS1 | 0.008 |
| SERPING1 | DDIT3 | 0.008 |
| CLDN3 | ARID5B | 0.008 |
| CLDN3 | NFE2L2 | 0.008 |
| CLDN3 | PDK4 | 0.008 |
| ADAM9 | EDN1 | 0.008 |
| ANXA1 | CEBPB | 0.008 |
| NRP1 | ARID5B | 0.008 |
| THBS2 | DUSP1 | 0.008 |
| MMP14 | ADM | 0.008 |
| MMP14 | CDK6 | 0.008 |
| MMP14 | TGIF1 | 0.008 |
| EDIL3 | NR0B2 | 0.008 |
| EDIL3 | RTN4 | 0.008 |
| CLDN3 | IRS2 | 0.0079 |
| CLDN3 | KLF5 | 0.0079 |
| CLDN3 | RBPJ | 0.0079 |
| C1QB | APP | 0.0079 |
| C1QB | EDN1 | 0.0079 |
| ADAM9 | ID1 | 0.0079 |
| JAM3 | ODC1 | 0.0079 |
| ANXA1 | TGIF1 | 0.0079 |
| NRP1 | ABCB1 | 0.0079 |
| NRP1 | RTN4 | 0.0079 |
| NRP1 | SPP1 | 0.0079 |
| THBS2 | BTG2 | 0.0079 |

| ligand | target | regulatory_potential |
| --- | --- | --- |
| THBS2 | CDKN1B | 0.0079 |
| THBS2 | TPT1 | 0.0079 |
| MMP14 | GADD45B | 0.0079 |
| EDIL3 | SGK1 | 0.0079 |
| EDIL3 | TGIF1 | 0.0079 |
| EFNA1 | HES1 | 0.0078 |
| CLDN3 | ETFB | 0.0078 |
| CLDN3 | G0S2 | 0.0078 |
| CLDN3 | KLF6 | 0.0078 |
| CLDN3 | SOX9 | 0.0078 |
| CLDN3 | TGFBI | 0.0078 |
| C1QB | GADD45A | 0.0078 |
| JAM3 | ETFB | 0.0078 |
| JAM3 | NR1D1 | 0.0078 |
| JAM3 | PTEN | 0.0078 |
| JAM3 | SLC1A5 | 0.0078 |
| ANXA1 | APP | 0.0078 |
| ANXA1 | CDK6 | 0.0078 |
| NRP1 | CXCL2 | 0.0078 |
| THBS2 | DHRS3 | 0.0078 |
| THBS2 | SGK1 | 0.0078 |
| TSPAN3 | ID3 | 0.0077 |
| TSPAN3 | PITX2 | 0.0077 |
| EFNA1 | CITED2 | 0.0077 |
| CLDN3 | CD83 | 0.0077 |
| JAM3 | CDK6 | 0.0077 |
| JAM3 | NFE2L2 | 0.0077 |
| JAM3 | TGIF1 | 0.0077 |
| ANXA1 | BCL2L11 | 0.0077 |
| ANXA1 | BTG2 | 0.0077 |
| NRP1 | BCL2L11 | 0.0077 |
| NRP1 | CEBPA | 0.0077 |
| NRP1 | RGS2 | 0.0077 |
| NRP1 | SLC2A1 | 0.0077 |
| NRP1 | TGFBI | 0.0077 |
| NRP1 | TIMP3 | 0.0077 |
| THBS2 | AMOTL2 | 0.0077 |
| THBS2 | RARA | 0.0077 |
| THBS2 | TGIF1 | 0.0077 |
| MMP14 | PIM1 | 0.0077 |
| EDIL3 | CDK6 | 0.0077 |
| EDIL3 | FOXO3 | 0.0077 |
| EDIL3 | HES6 | 0.0077 |
| TSPAN3 | BRD2 | 0.0076 |
| TSPAN3 | GADD45B | 0.0076 |
| TSPAN3 | MAFF | 0.0076 |
| TSPAN3 | SOX9 | 0.0076 |
| EFNA1 | DDIT3 | 0.0076 |
| EFNA1 | DUSP1 | 0.0076 |
| EFNA1 | IL1B | 0.0076 |
| SERPING1 | HES1 | 0.0076 |
| ADAM9 | ATF3 | 0.0076 |
| JAM3 | IGFBP4 | 0.0076 |
| JAM3 | PRSS23 | 0.0076 |
| JAM3 | RBPJ | 0.0076 |
| ANXA1 | SLC2A1 | 0.0076 |
| ANXA1 | SOX9 | 0.0076 |
| NRP1 | ETFB | 0.0076 |
| NRP1 | FOXO3 | 0.0076 |
| NRP1 | MALAT1 | 0.0076 |
| NRP1 | SLC3A2 | 0.0076 |
| THBS2 | ARID5B | 0.0076 |
| THBS2 | CDK6 | 0.0076 |
| THBS2 | HSP90AB1 | 0.0076 |

| ligand | target | regulatory_potential |
| --- | --- | --- |
| THBS2 | ID3 | 0.0076 |
| THBS2 | SPP1 | 0.0076 |
| MMP14 | CTNNB1 | 0.0076 |
| EDIL3 | ATP1A1 | 0.0076 |
| EDIL3 | NEBL | 0.0076 |
| EDIL3 | SLC2A1 | 0.0076 |
| EDIL3 | TRIB1 | 0.0076 |
| TSPAN3 | DDX5 | 0.0075 |
| TSPAN3 | GEM | 0.0075 |
| TSPAN3 | HEY1 | 0.0075 |
| TSPAN3 | HNRNPR | 0.0075 |
| TSPAN3 | KLF5 | 0.0075 |
| TSPAN3 | NUPR1 | 0.0075 |
| TSPAN3 | ZMYND8 | 0.0075 |
| C1QB | FN1 | 0.0075 |
| C1QB | GADD45B | 0.0075 |
| C1QB | THBS1 | 0.0075 |
| SEMA5B | ATF3 | 0.0075 |
| SEMA5B | GADD45A | 0.0075 |
| SEMA5B | GDF15 | 0.0075 |
| JAM3 | BTG2 | 0.0075 |
| JAM3 | CTNNB1 | 0.0075 |
| JAM3 | RGS2 | 0.0075 |
| ANXA1 | ABCB1 | 0.0075 |
| ANXA1 | PTEN | 0.0075 |
| NRP1 | G0S2 | 0.0075 |
| NRP1 | PRSS23 | 0.0075 |
| NRP1 | RARA | 0.0075 |
| THBS2 | BCL2L11 | 0.0075 |
| THBS2 | FOXO3 | 0.0075 |
| THBS2 | PIM1 | 0.0075 |
| THBS2 | TRIB1 | 0.0075 |
| TSPAN3 | CRIP2 | 0.0074 |
| TSPAN3 | PTEN | 0.0074 |
| TSPAN3 | TGIF1 | 0.0074 |
| EFNA1 | FN1 | 0.0074 |
| SERPING1 | CEBPB | 0.0074 |
| SERPING1 | HNRNPR | 0.0074 |
| SERPING1 | ID3 | 0.0074 |
| SERPING1 | TGIF1 | 0.0074 |
| C1QB | ID1 | 0.0074 |
| SEMA5B | IL1B | 0.0074 |
| ADAM9 | GDF15 | 0.0074 |
| JAM3 | ABCB1 | 0.0074 |
| JAM3 | KLF6 | 0.0074 |
| JAM3 | PPP1R15A | 0.0074 |
| JAM3 | RND3 | 0.0074 |
| JAM3 | SOX9 | 0.0074 |
| ANXA1 | CDKN1B | 0.0074 |
| ANXA1 | NFE2L2 | 0.0074 |
| ANXA1 | RARA | 0.0074 |
| THBS2 | ORM1 | 0.0074 |
| TSPAN3 | CEBPA | 0.0073 |
| TSPAN3 | CLDN1 | 0.0073 |
| TSPAN3 | DDIT3 | 0.0073 |
| TSPAN3 | G0S2 | 0.0073 |
| TSPAN3 | TSC22D1 | 0.0073 |
| EFNA1 | SGK1 | 0.0073 |
| C1QB | CDK6 | 0.0073 |
| SEMA5B | EDN1 | 0.0073 |
| ADAM9 | GADD45A | 0.0073 |
| JAM3 | BCL2L11 | 0.0073 |
| JAM3 | CDKN1B | 0.0073 |
| JAM3 | G0S2 | 0.0073 |

| ligand | target | regulatory_potential |
|---|---|---|
| JAM3 | TGFBI | 0.0073 |
| ANXA1 | KLF6 | 0.0073 |
| THBS2 | GADD45B | 0.0073 |
| THBS2 | KLF5 | 0.0073 |
| THBS2 | TUBB2B | 0.0073 |
| TSPAN3 | BTG2 | 0.0072 |
| TSPAN3 | NFE2L2 | 0.0072 |
| EFNA1 | PIM1 | 0.0072 |
| SERPING1 | CDK6 | 0.0072 |
| SERPING1 | SGK1 | 0.0072 |
| SERPING1 | TRIB1 | 0.0072 |
| C1QB | ID3 | 0.0072 |
| ADAM9 | DDIT3 | 0.0072 |
| ADAM9 | THBS1 | 0.0072 |
| JAM3 | RARA | 0.0072 |
| ANXA1 | ODC1 | 0.0072 |
| ANXA1 | PPP1R15A | 0.0072 |
| ANXA1 | RBPJ | 0.0072 |
| THBS2 | KLF10 | 0.0072 |
| THBS2 | NFE2L2 | 0.0072 |
| MMP14 | DHRS3 | 0.0072 |
| TSPAN3 | CLU | 0.0071 |
| TSPAN3 | DUSP1 | 0.0071 |
| SERPING1 | CEBPA | 0.0071 |
| SERPING1 | DUSP1 | 0.0071 |
| SERPING1 | FN1 | 0.0071 |
| SERPING1 | GADD45B | 0.0071 |
| JAM3 | SLC2A1 | 0.0071 |
| ANXA1 | DHRS3 | 0.0071 |
| THBS2 | APP | 0.0071 |
| THBS2 | S100A8 | 0.0071 |
| MMP14 | FOXO3 | 0.0071 |
| MMP14 | ODC1 | 0.0071 |
| MMP14 | RBPJ | 0.0071 |
| TSPAN3 | APP | 0.007 |
| TSPAN3 | FN1 | 0.007 |
| TSPAN3 | NFIA | 0.007 |
| EFNA1 | CDK6 | 0.007 |
| EFNA1 | CDKN1B | 0.007 |
| EFNA1 | GADD45B | 0.007 |
| EFNA1 | PTEN | 0.007 |
| EFNA1 | TRIB1 | 0.007 |
| SERPING1 | CXCL2 | 0.007 |
| C1QB | BCL2L11 | 0.007 |
| C1QB | HES1 | 0.007 |
| SEMA5B | HES1 | 0.007 |
| SEMA5B | ID3 | 0.007 |
| SEMA5B | THBS1 | 0.007 |
| ADAM9 | ID3 | 0.007 |
| JAM3 | SLC38A2 | 0.007 |
| JAM3 | SLC3A2 | 0.007 |
| ANXA1 | KLF5 | 0.007 |
| ANXA1 | SPP1 | 0.007 |
| THBS2 | RBM39 | 0.007 |
| MMP14 | BCL2L11 | 0.007 |
| MMP14 | PTEN | 0.007 |
| TSPAN3 | CDKN1B | 0.0069 |
| TSPAN3 | RPL26 | 0.0069 |
| TSPAN3 | SGK1 | 0.0069 |
| SERPING1 | ODC1 | 0.0069 |
| C1QB | DDIT3 | 0.0069 |
| SEMA5B | DDIT3 | 0.0069 |
| SEMA5B | DUSP1 | 0.0069 |
| ADAM9 | IL1B | 0.0069 |

| ligand | target | regulatory_potential |
|---|---|---|
| ANXA1 | HNRNPR | 0.0069 |
| ANXA1 | HSP90AB1 | 0.0069 |
| THBS2 | GNAS | 0.0069 |
| THBS2 | RGS2 | 0.0069 |
| MMP14 | CDKN1B | 0.0069 |
| MMP14 | KLF5 | 0.0069 |
| MMP14 | NR1D1 | 0.0069 |
| TSPAN3 | HOXA3 | 0.0068 |
| EFNA1 | ADM | 0.0068 |
| C1QB | CITED2 | 0.0068 |
| C1QB | PTEN | 0.0068 |
| SEMA5B | CITED2 | 0.0068 |
| ADAM9 | DUSP1 | 0.0068 |
| ADAM9 | HES1 | 0.0068 |
| ANXA1 | MALAT1 | 0.0068 |
| ANXA1 | RTN4 | 0.0068 |
| ANXA1 | STMN1 | 0.0068 |
| THBS2 | ABCA1 | 0.0068 |
| THBS2 | SFN | 0.0068 |
| THBS2 | TGFBI | 0.0068 |
| THBS2 | ZMYND8 | 0.0068 |
| MMP14 | BTG2 | 0.0068 |
| MMP14 | CEBPA | 0.0068 |
| MMP14 | CEBPB | 0.0068 |
| MMP14 | KLF6 | 0.0068 |
| MMP14 | NFE2L2 | 0.0068 |
| MMP14 | RTN4 | 0.0068 |
| MMP14 | SLC2A1 | 0.0068 |
| MMP14 | ZMYND8 | 0.0068 |
| EFNA1 | BTG2 | 0.0067 |
| EFNA1 | DHRS3 | 0.0067 |
| SERPING1 | ADM | 0.0067 |
| SERPING1 | CDKN1B | 0.0067 |
| C1QB | PIM1 | 0.0067 |
| SEMA5B | TGIF1 | 0.0067 |
| ADAM9 | CITED2 | 0.0067 |
| ANXA1 | CEBPA | 0.0067 |
| ANXA1 | EFNA1 | 0.0067 |
| ANXA1 | SLC38A2 | 0.0067 |
| THBS2 | ODC1 | 0.0067 |
| MMP14 | HNRNPR | 0.0067 |
| MMP14 | SOX9 | 0.0067 |
| MMP14 | SPP1 | 0.0067 |
| MMP14 | STMN1 | 0.0067 |
| EFNA1 | ABCB1 | 0.0066 |
| EFNA1 | CTNNB1 | 0.0066 |
| EFNA1 | ODC1 | 0.0066 |
| EFNA1 | TGFBI | 0.0066 |
| EFNA1 | TGIF1 | 0.0066 |
| SERPING1 | DHRS3 | 0.0066 |
| C1QB | DUSP1 | 0.0066 |
| SEMA5B | GADD45B | 0.0066 |
| ANXA1 | GCLC | 0.0066 |
| ANXA1 | NR1D1 | 0.0066 |
| ANXA1 | RGS2 | 0.0066 |
| ANXA1 | UBC | 0.0066 |
| THBS2 | FCGRT | 0.0066 |
| THBS2 | MALAT1 | 0.0066 |
| THBS2 | RTN4 | 0.0066 |
| MMP14 | ABCA1 | 0.0066 |
| MMP14 | ANXA3 | 0.0066 |
| MMP14 | ETFB | 0.0066 |
| MMP14 | RARA | 0.0066 |
| SERPING1 | NFE2L2 | 0.0065 |

| ligand | target | regulatory_potential |
|---|---|---|
| SERPING1 | NUPR1 | 0.0065 |
| SERPING1 | SPP1 | 0.0065 |
| C1QB | ABCA1 | 0.0065 |
| C1QB | CXCL2 | 0.0065 |
| C1QB | TGIF1 | 0.0065 |
| ADAM9 | FN1 | 0.0065 |
| ADAM9 | PIM1 | 0.0065 |
| THBS2 | CTNNB1 | 0.0065 |
| THBS2 | ZC3H12A | 0.0065 |
| MMP14 | CLDN1 | 0.0065 |
| MMP14 | RGS2 | 0.0065 |
| EFNA1 | APP | 0.0064 |
| SERPING1 | PTEN | 0.0064 |
| SEMA5B | CDK6 | 0.0064 |
| SEMA5B | PIM1 | 0.0064 |
| ADAM9 | SGK1 | 0.0064 |
| MMP14 | AREG | 0.0064 |
| MMP14 | CLU | 0.0064 |
| MMP14 | MALAT1 | 0.0064 |
| MMP14 | SKIL | 0.0064 |
| MMP14 | TGFBI | 0.0064 |
| MMP14 | UBC | 0.0064 |
| EFNA1 | CEBPB | 0.0063 |
| EFNA1 | SLC2A1 | 0.0063 |
| SERPING1 | ARID5B | 0.0063 |
| SERPING1 | EFNA1 | 0.0063 |
| SERPING1 | RTN4 | 0.0063 |
| SERPING1 | SLC2A1 | 0.0063 |
| SERPING1 | ZMYND8 | 0.0063 |
| C1QB | ABCB1 | 0.0063 |
| SEMA5B | CEBPA | 0.0063 |
| SEMA5B | FN1 | 0.0063 |
| SEMA5B | SGK1 | 0.0063 |
| ADAM9 | APP | 0.0063 |
| ADAM9 | GADD45B | 0.0063 |
| MMP14 | PPP1R15A | 0.0063 |
| EFNA1 | BCL2L11 | 0.0062 |
| EFNA1 | HNRNPR | 0.0062 |
| EFNA1 | NFE2L2 | 0.0062 |
| EFNA1 | RBPJ | 0.0062 |
| SERPING1 | BCL2L11 | 0.0062 |
| SERPING1 | KLF5 | 0.0062 |
| C1QB | TRIB1 | 0.0062 |
| SEMA5B | RBPJ | 0.0062 |
| ADAM9 | SLC2A1 | 0.0062 |
| ADAM9 | TRIB1 | 0.0062 |
| EFNA1 | IGFBP4 | 0.0061 |
| SERPING1 | BAMBI | 0.0061 |
| SERPING1 | BTG2 | 0.0061 |
| SERPING1 | CLDN1 | 0.0061 |
| C1QB | BTG2 | 0.0061 |
| C1QB | CEBPB | 0.0061 |
| SEMA5B | ODC1 | 0.0061 |
| SEMA5B | TRIB1 | 0.0061 |
| ADAM9 | PTEN | 0.0061 |
| ADAM9 | TGIF1 | 0.0061 |
| EFNA1 | CEBPA | 0.006 |
| EFNA1 | FOXO3 | 0.006 |
| EFNA1 | KLF6 | 0.006 |
| SERPING1 | ETFB | 0.006 |
| SERPING1 | RARA | 0.006 |
| SERPING1 | SOX9 | 0.006 |
| C1QB | ADM | 0.006 |
| C1QB | HNRNPR | 0.006 |

| ligand | target | regulatory_potential |
|---|---|---|
| C1QB | SGK1 | 0.006 |
| SEMA5B | BTG2 | 0.006 |
| ADAM9 | ADM | 0.006 |
| ADAM9 | BTG2 | 0.006 |
| ADAM9 | CDK6 | 0.006 |
| ADAM9 | DHRS3 | 0.006 |
| ADAM9 | ODC1 | 0.006 |
| EFNA1 | RGS2 | 0.0059 |
| SEMA5B | CEBPB | 0.0059 |
| EFNA1 | ABCA1 | 0.0058 |
| EFNA1 | CXCL2 | 0.0058 |
| EFNA1 | RARA | 0.0058 |
| EFNA1 | SOX9 | 0.0058 |
| C1QB | CD83 | 0.0058 |
| C1QB | CDKN1B | 0.0058 |
| C1QB | DHRS3 | 0.0058 |
| C1QB | PRSS23 | 0.0058 |
| SEMA5B | ETFB | 0.0058 |
| SEMA5B | PTEN | 0.0058 |
| EFNA1 | ARID5B | 0.0057 |
| EFNA1 | CLDN1 | 0.0057 |
| EFNA1 | MALAT1 | 0.0057 |
| EFNA1 | NFIA | 0.0057 |
| EFNA1 | STMN1 | 0.0057 |
| EFNA1 | ZMYND8 | 0.0057 |
| C1QB | NFE2L2 | 0.0057 |
| C1QB | SOX9 | 0.0057 |
| SEMA5B | ADM | 0.0057 |
| SEMA5B | APP | 0.0057 |
| SEMA5B | DHRS3 | 0.0057 |
| SEMA5B | HNRNPR | 0.0057 |
| SEMA5B | SOX9 | 0.0057 |
| ADAM9 | BAMBI | 0.0057 |
| C1QB | GCLC | 0.0056 |
| C1QB | IGFBP4 | 0.0056 |
| C1QB | SLC2A1 | 0.0056 |
| C1QB | SPP1 | 0.0056 |
| SEMA5B | CDKN1B | 0.0056 |
| ADAM9 | BCL2L11 | 0.0056 |
| ADAM9 | CEBPA | 0.0056 |
| ADAM9 | RARA | 0.0056 |
| ADAM9 | RBPJ | 0.0056 |
| ADAM9 | RTN4 | 0.0056 |
| C1QB | CEBPA | 0.0055 |
| C1QB | ETFB | 0.0055 |
| C1QB | KLF10 | 0.0055 |
| C1QB | RBPJ | 0.0055 |
| C1QB | TGFBI | 0.0055 |
| SEMA5B | RARA | 0.0055 |
| SEMA5B | SLC2A1 | 0.0055 |
| SEMA5B | STMN1 | 0.0055 |
| SEMA5B | ZMYND8 | 0.0055 |
| ADAM9 | ABCA1 | 0.0055 |
| ADAM9 | CDKN1B | 0.0055 |
| ADAM9 | HNRNPR | 0.0055 |
| C1QB | EFNA1 | 0.0054 |
| C1QB | G0S2 | 0.0054 |
| C1QB | MAFF | 0.0054 |
| C1QB | ODC1 | 0.0054 |
| C1QB | STMN1 | 0.0054 |
| SEMA5B | EFNA1 | 0.0054 |
| SEMA5B | FOXO3 | 0.0054 |
| SEMA5B | KLF5 | 0.0054 |
| ADAM9 | FOXO3 | 0.0054 |

| ligand | target | regulatory_potential |
|---|---|---|
| ADAM9 | NFIA | 0.0054 |
| ADAM9 | SOX9 | 0.0054 |
| ADAM9 | STMN1 | 0.0054 |
| ADAM9 | ZMYND8 | 0.0054 |
| SEMA5B | BCL2L11 | 0.0053 |
| SEMA5B | CTNNB1 | 0.0053 |
| SEMA5B | KLF6 | 0.0053 |
| SEMA5B | MALAT1 | 0.0053 |
| SEMA5B | NFE2L2 | 0.0053 |
| SEMA5B | NR1D1 | 0.0053 |
| SEMA5B | PPP1R15A | 0.0053 |
| SEMA5B | RTN4 | 0.0053 |
| ADAM9 | CEBPB | 0.0053 |
| ADAM9 | IGFBP4 | 0.0053 |
| ADAM9 | KLF10 | 0.0053 |
| ADAM9 | SPP1 | 0.0053 |
| ADAM9 | TGFBI | 0.0053 |
| SEMA5B | NFIA | 0.0052 |
| SEMA5B | NR0B2 | 0.0052 |
| SEMA5B | TGFBI | 0.0052 |
| ADAM9 | EFNA1 | 0.0052 |
| ADAM9 | MALAT1 | 0.0052 |
| SEMA5B | CLU | 0.0051 |
| SEMA5B | CXCL2 | 0.0051 |
| SEMA5B | HNRNPF | 0.0051 |
| SEMA5B | SLC1A5 | 0.0051 |
| SEMA5B | SLC3A2 | 0.0051 |
| ADAM9 | ARID5B | 0.0051 |
| ADAM9 | CTNNB1 | 0.0051 |
| ADAM9 | KLF6 | 0.0051 |
| ADAM9 | NFE2L2 | 0.0051 |

### Table S6

Table S6. Gene sets used for the immune-context and therapeutic-association analyses.

| gene_set | category | n_genes | genes |
|---|---|---|---|
| Stromal | Immune / stromal activity | 15 | COL1A1; COL1A2; COL3A1; FN1; FAP; THY1; PDGFRB; ACTA2; DCN; LUM; POSTN; SPARC; TAGLN; MMP… |
| Immune | Immune / stromal activity | 15 | PTPRC; CD2; CD3D; CD3E; CD8A; GZMA; GZMB; PRF1; IL2RG; CXCL9; CXCL10; CCL5; HLA-DRA; LCK;… |
| CD8 T | Immune cell type | 6 | CD8A; CD8B; GZMK; GZMA; PRF1; EOMES |
| Cytotoxic | Immune cell type | 6 | GZMA; GZMB; GZMH; PRF1; NKG7; KLRD1 |
| Treg | Immune cell type | 5 | FOXP3; IL2RA; CTLA4; IKZF2; TNFRSF18 |
| Macrophage | Immune cell type | 6 | CD68; CD163; MSR1; MRC1; CSF1R; C1QA |
| NK | Immune cell type | 5 | NCR1; KLRF1; NCAM1; KIR2DL3; FCGR3A |
| B cell | Immune cell type | 5 | CD79A; CD79B; MS4A1; CD19; TNFRSF13C |
| DC | Immune cell type | 4 | CD1C; CLEC9A; LAMP3; FCER1A |
| Neutrophil | Immune cell type | 5 | FCGR3B; CSF3R; S100A8; S100A9; CXCR2 |
| CAF (exclusion) | Immune-exclusion / dysfunction axis | 8 | FAP; PDGFRB; ACTA2; COL1A1; COL11A1; THY1; POSTN; MMP2 |
| TGFB response | Immune-exclusion / dysfunction axis | 8 | TGFB1; TGFBR1; TGFBR2; SMAD2; SMAD3; SERPINE1; SKIL; JUNB |
| MDSC | Immune-exclusion / dysfunction axis | 6 | S100A8; S100A9; IL1B; CXCL8; ARG1; ITGAM |
| T-cell dysfunction | Immune-exclusion / dysfunction axis | 7 | PDCD1; CTLA4; LAG3; HAVCR2; TIGIT; TOX; ENTPD1 |
| IFNG response | Immune-exclusion / dysfunction axis | 7 | IFNG; STAT1; CXCL9; CXCL10; IDO1; GBP1; HLA-DRA |
| Cytolytic | Immune-exclusion / dysfunction axis | 2 | GZMA; PRF1 |
| 5-FU / TYMS | Drug-target / EMT pathway | 4 | TYMS; DPYD; UMPS; TK1 |
| Oxaliplatin / DDR | Drug-target / EMT pathway | 5 | ERCC1; XRCC1; BRCA1; ATM; ATR |
| Irinotecan / TOP1 | Drug-target / EMT pathway | 4 | TOP1; ABCB1; ABCG2; UGT1A1 |
| EGFR pathway | Drug-target / EMT pathway | 5 | EGFR; AREG; EREG; KRAS; BRAF |
| VEGF / angiogenesis | Drug-target / EMT pathway | 5 | VEGFA; KDR; FLT1; ANGPT2; PGF |
| EMT | Drug-target / EMT pathway | 6 | VIM; CDH2; SNAI1; SNAI2; ZEB1; TWIST1 |
| Mesenchymal (CMS4-like) | Molecular subtype axis | 12 | ZEB1; TWIST1; SNAI2; COL1A1; COL3A1; FN1; POSTN; THBS2; SPARC; TAGLN; FAP; INHBA |
