## Supplementary figures and images for "An RGS5-positive fibroblast to WNT-like epithelial TGFB axis in colorectal liver metastasis, replicated in an independent paired cohort"

### Additional_file_1.tiff

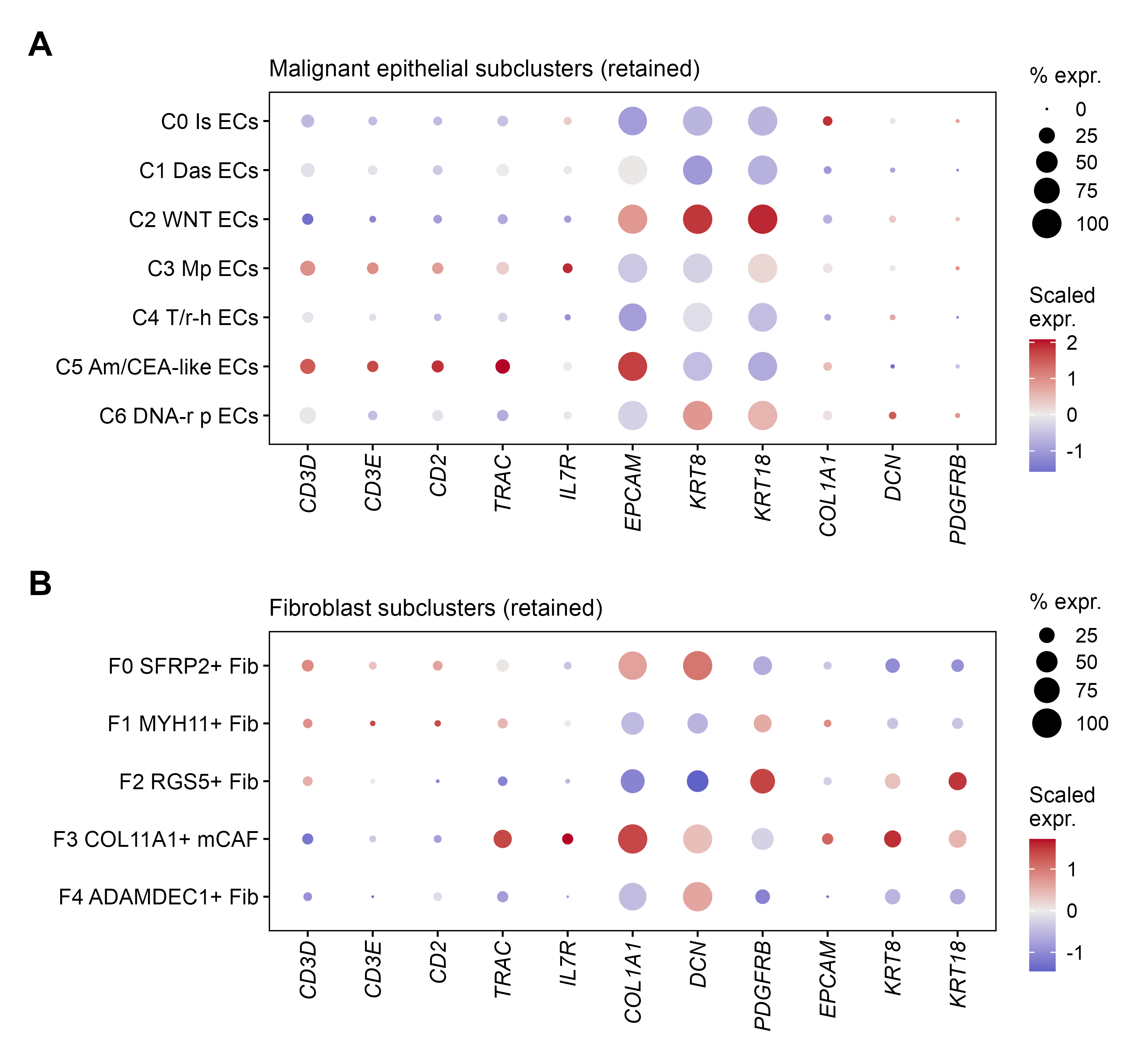

### Additional_file_4.tiff

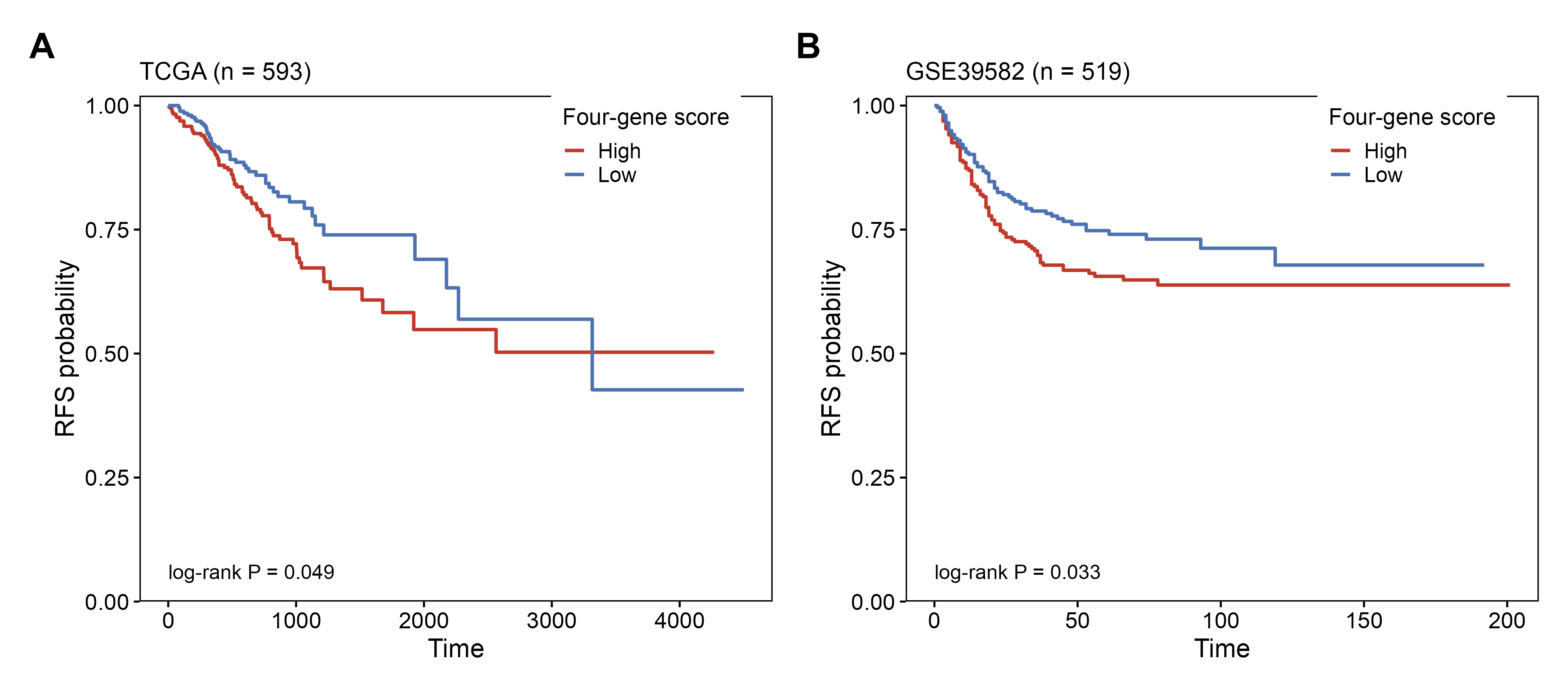
